# Core gene set of the species *Saccharomyces cerevisiae*

**DOI:** 10.1101/2023.09.07.545205

**Authors:** Fred S. Dietrich, Paul Magwene, John McCusker

**Affiliations:** Department of Molecular Genetics and Microbiology, Duke University, Durham, NC 27710 USA; Department of Biology, Duke University, Durham, NC 27710 USA

**Keywords:** Comparative Genomics, Fungal Genetics, Strain S288c, Introns, *Ashbya gossypii*, Transposable elements

## Abstract

Examination of the genome sequence of *Saccharomyces cerevisiae* strain S288c and 93 additional diverse strains allows identification of the 5885 genes that make up the core set of genes in this species and gives a better sense of the organization and plasticity of this genome. *S. cerevisiae* strains each contain dozens to hundreds of strain-specific genes. In addition to a variable content of retrotransposons Ty1-Ty6, some strains contain a novel transposable element, Ty7. Examination further shows that some annotated putative protein coding genes are likely artifacts. We propose altering approximately 5% of the current annotations in the widely used reference strain S288c. Potential null alleles are common and found in all 94 strains examined, with these potential null alleles typically containing a single stop codon or frameshift. There are also gene remnants, pseudogenes, and variable arrays of genes. Among the core genes there are now only 364 protein coding genes of unknown function, classified as uncharacterized in the Saccharomyces Genome Database. This work suggests that there is a role for carefully edited and annotated genome sequences in understanding the genome organization and content of a species. We propose that gene remnants be added to the repertoire of features found in the *S. cerevisiae* genome, and likely other fungal species.

## Introduction

It has now been more than 25 years since the public release and publication of the first complete annotated eukaryotic genome sequence, that of the *Saccharomyces cerevisiae* reference strain S288c, and 60 years since the publication of the first nucleotide sequence of a gene, an *S. cerevisiae* alanine tRNA (Holley et al. 1965). One of the questions answered when that first genome was completed was, “only 5885 protein encoding genes are believed to exist (Goffeau et al. 1996)”. Because the genome sequences of dozens of hemiascomycetes and more than 1000 genomes of strains of *S. cerevisiae* have been generated to varying degrees of completeness since then, it is now possible to ask a slightly different question, “what is the set of genes in the species *S. cerevisiae*?” To give a preliminary answer to this question we have re-examined 94 strains, the genome sequence of S288c as well as that of the 93 genomes published in Strope et al (Strope et al. 2015). In addition, this work depends on the tremendous amount of published data available on this species and closely related species. The work presented here would not be feasible without the compilation of much of this data into a single curated and searchable space, the *Saccharomyces* Genome Database (SGD) (Cherry et al. 2012). Additional data and analysis used here comes from work on *Ashbya gossypii* (Dietrich et al. 2004, Dietrich et al. 2013) the Yeast Gene Order Browser (YGOB) (Byrne and Wolfe 2005), and genomic and RNA-seq data from NCBI (Sayers et al. 2022) as well as proteomic data, and predicted protein structure (Jumper et al. 2021).

Much of what is reported here has long been known, but with the addition of whole genome sequences and additional data it is possible now to analyze the genome structure in greater depth with greater resolution of the features found in those genomes. For example, in the 1960’s it was known that there were at least five sets of genes for maltose utilization (then *MA*, now *MAL*), and that they are found at the ends of linkage groups (Halvorson et al. 1963, Mortimer and Hawthorne 1966) and these locations were variable between strains. Thus the idea that the distal ends of linkage groups, now telomeres, of *S. cerevisiae* have more extensive variation than exists in the central areas of the linkage groups, now chromosomes, can be traced back 60 years. Among the 94 strains, *MAL11* and paralogs can be found at eleven of the thirty-two chromosome ends (chromosome IL chromosome IIR chromosome IVL chromosome VIL chromosome VIIR chromosome IXL chromosome XL chromosome XR chromosome XIR chromosome XVL chromosome XVR). In this work we identify the location of the boundary between the less variable central region of each chromosome and the more variable subtelomeric region.

Another characteristic of *S. cerevisiae* strains is the presence of defective alleles. A pivotal event in the development of yeast genetics in the 1940’s was the discovery of an *S. cerevisiae* strain lacking homothallism (Winge 1949), making it possible to do crosses, study mating and genetics. It is now known that the *D* gene, now *HO*, is a null allele in strain S288c the result of missense mutations (Meiron et al. 1995). All 94 strains of *S. cerevisiae* that we are examining in this work contain multiple genes where the allele present is a potential null allele, containing a frameshift or a stop codon. Understanding that there are naturally occurring potential null alleles and that there are highly diverged genes often resulting from introgression are keys to establishing a list of the core genes for this species. It is important to note that these potential null alleles are here considered alleles, not pseudogenes or gene remnants, which are different phenomenon. Pseudogenes and gene remnants are two additional classes of gene related features that are not expected to encode a functional protein and cannot easily mutate back to a functional gene.

One aspect of moving from having a single genome sequence of *S. cerevisiae* to having thousands of genome sequences is the importance of having well defined terms to use in genome annotation. Fortunately for this species rules for chromosome numbering, gene names, and other needed terms have been adopted and are almost universally followed by the community, with a gene name registry formerly maintained by Robert Mortimer and now by SGD (von Borstel 1963, Mortimer and Hawthorne 1966, Mortimer et al. 1989, Cherry et al. 1997, SGD 1998, Engel et al. 2022). Some of the terms used in this work are defined in Table 1, following as closely as possible to the established terminology in the field. We propose the following terms for inclusion in the annotation of these genomes: core genes, non-core genes, variably located ubiquitous genes, central region of genome, subtelomeric region of genome, and gene remnants, as they provide additional insight into the history and genome plasticity of *S. cerevisiae* strains. Furthermore, it is important to have highly accurate annotations of the one or more reference genomes, so the reference genomes do not become a source of propagation of annotation errors.

**Table 1.** Nomenclature used in this paper.

|  |  |
| --- | --- |
| Gene | While sometimes “gene” is used to mean protein coding gene, in general and in this work gene refers to DNA elements encoding RNA products, such as tRNA molecules, as well as genes encoding proteins. Genomic structural features such as centromeres are not included here as genes. Dubious genes are not included here as genes. |
| Core Gene | A gene where an allele is at the same position and orientation relative to flanking genes in 95% or more of the strains examined. The alleles can be an intact copy of the gene, a truncated copy, a copy containing stop codons and/or frameshifts, a copy disrupted by a transposable element, or a copy of an ortholog introgressed from a sibling species. Other authors have used different definitions of a core gene, resulting in a different list of the core genes. A more precise list of the core genes by this definition will need to be based on a very large dataset. While not done in this work, a better definition of core gene would weigh the sequence conservation of the gene and adjacent genes, as highly similar sequences indicate recent identity by descent and thus bias the scoring. |
| Non-core genes | Genes found in less than 95% of strains at the same position and orientation in the genome. |
| Variably Located Ubiquitous Gene | A gene found in less than 95% of strains at the same position and orientation, thus a non-core gene, but found in more than 95% of strains at a combination of locations. |
| Central Region of Chromosome | The region of each chromosome where all core genes on that chromosome are found, ending with a core gene at each end. |
| Subtelomeric Region | The subtelomeric region at each chromosome end is between the final core gene and the terminal telomeric repeat. This definition of the subtelomeric region is based on genes differing from the definition based on *RAP1* silencing of chromosome ends (Wright et al. 1992, Kyrion et al. 1993, Martin et al. 1999). |
| Allele | This term is commonly used in reference both to naturally occurring alleles, and those engineered. In this work allele refers to natural gene variants found at the same location and in the same orientation. Alleles include those with, relative to other strains, synonymous and non-synonymous changes, as well as introgressed orthologs at the usual location for the gene and changes resulting in a premature stop codon, Ty insertion, and insertions and deletions, including those that are out of frame. While synthetic gene deletions are usually referred to as null alleles, naturally occurring alleles are sometimes mistakenly referred to as pseudogenes, and in this work we carefully distinguish between alleles and pseudogenes. |
| Potential null allele | An allele containing one or more stop codons, frameshift mutations, or a transposon insertion that disrupt the open reading frame or non-synonymous changes likely to result in the gene being non-functional. These should be considered potential null alleles as a single stop codon or frameshift can revert and frameshift and stop codon suppression occurs in *S. cerevisiae*, and in a heterozygous diploid a potential null allele can be gene converted back to a functional allele. |
| Truncated allele | An allele where a portion of the gene is missing due to deletion or Ty insertion. |
| Pseudogene | A non-functional copy of DNA at a location where that DNA is not usually found. This can be a fragment of mitochondrial DNA inserted in the nuclear genome, a partial tRNA gene, a piece of rDNA, or a piece of a protein coding gene or intergenic region. In *S. cerevisiae* many of these pseudogenes are found in the subtelomeric regions. |
| Gene Remnant | Not a gene, but a region of chromosomal DNA with remaining similarity to an ancestral gene that was at that location in the ancestor of *S. cerevisiae* but has accumulated multiple potentially null mutations. Recognizable fragments of the gene persist and can be recognized by similarity with genes in related species, or in some cases the paralog from the whole genome duplication (WGD). These gene fragments cannot easily be resurrected to a gene by mutation, or by gene conversion as the gene is generally not present in this species, and insufficient homology exists with other genes. The most likely chance of resurrection is introgression. In some cases these gene remnants are intermediates in the loss of genes that were duplicates from the WGD. In other cases they result from the process of gene duplication, where duplication of one gene can carry along a fragment of an adjoining gene. |
| Tandem | Cases of tandem gene duplication exist. In some cases, the gene copies are identical or nearly |
| Duplicate Gene | identical, and in other cases the copies are diverged. Some tandem duplicate genes are ubiquitous, while others are found in a subset of strains. Nomenclature for variable tandem arrays of genes is problematic as the number of genes can vary among strains, and in some cases the tandem duplicate genes can be identical in some strains, non-identical in other strains. When tandem duplicate genes are identical or near identical, then the tandem duplicate genes are here considered one core gene plus non-core genes. |
| Dispersed Duplicate Gene | Cases of non-adjacent duplicate genes exist, sometimes with a copy located in the subtelomeric region of a chromosomes. These are often intact genes, as opposed to pseudogenes that are gene fragments. In some cases it is unclear if a DNA segment encodes a pseudogene or a dispersed duplicate gene. |
| General *S. cerevisiae* nomenclature |  |
| Chromosome Names | These follow the chromosome numbering as described in the genetic maps of Robert Mortimer (Mortimer and Hawthorne 1966, Mortimer and Schild 1980, Mortimer and Schild 1985, Mortimer et al. 1989, Mortimer et al. 1992, Cherry et al. 1997), with the addition that in cases where there are translocations, the name of the chromosome is based on the origin of the centromere. In the 94 strains examined here, there has been no gain or loss of centromeres, other than in cases of aneuploidy. |
| Gene Names | Gene names follow the establish rules for *S. cerevisiae* as originally described and coordinated by Robert Mortimer, and more recently by SGD (SGD 1998). There are several difficulties with this naming convention that will need to be addressed in the future. These difficulties include names of genes not in the reference strain, names of tandemly duplicated genes, names of Ty/LTR elements, names of introgressed genes, and the names of subtelomeric genes that are located on different chromosomes in different strains. |
| Whole Genome Duplication (WGD) | A WGD occurred in the Hemiascomycete lineage leading to *S. cerevisiae*. In most cases one gene was lost from pairs of genes resulting from the whole genome duplication (Philippsen et al. 1997, Wolfe and Shields 1997). Some 600 pairs of duplicate genes remain in *S. cerevisiae* from that event in the ancestry of *S. cerevisiae*. In other post WGD species the set of remaining duplicate genes that have been retained is slightly different. Not all duplicate genes in *S. cerevisiae* are the result of the whole genome duplication; some predate the WGD, some post date the WGD. |
| Telomeric Sequence | The somewhat variable sequence found at telomeres of all *S. cerevisiae* chromosomes that is encoded by the *TLC1* RNA (Walmsley et al. 1984). |
| Verified Gene | A term pioneered by SGD to identify protein coding genes where something is known about the function of the gene. In this work we use the SGD assignment of which genes are verified to support that something is known about the function. In the future it may be desirable to further define what is meant by a gene of known function. |
| Uncharacterized Gene | A term pioneered by SGD to identify protein coding genes where evidence suggests they are real protein coding genes, but where the function is unknown. Also referred to as a gene of unknown function. |
| Dubious Gene | A term pioneered by SGD to identify regions previously annotated as a protein coding gene where there is little to no evidence that this open reading frame encodes a protein. This strictly speaking does not mean that the open reading frame is not a gene, only that there is insufficient data. If evidence is forthcoming, then it is possible for a dubious gene to be resurrected to the status of verified or uncharacterized gene. |

## Materials and Methods

Initial genome sequences of the 93 *S. cerevisiae* strains has been previously described (Strope et al. 2015). The sequences when originally submitted to GenBank in 2015 contained thousands of misassembles, gaps, and point errors. Sequences were reassembled with many gaps closed, many errors corrected, and the annotations updated in 2016. To further improve the assemblies by closing gaps and correcting errors, a more exhaustive assembly of the data was undertaken. Denovo sequence assembly was used for all assemblies as some sequences in this species are too diverged from reference genome assembly to work properly. All 1488 chromosomes contained one or more gaps or errors that have now been corrected. We are currently in the process of updating the sequences and the annotations in GenBank. Initial assemblies of the data using ABySS for the 93 genomes resulted, all together, in more than 300,000 contigs, with the exact number being difficult to determine as different assemblies using different parameters were used for the mitochondrial genome, the rDNA and for each strain. Alignment of these contigs with the S288c reference genomes identified that the majority of these *de novo* generated contigs are colinear with regions from the reference genome, though more than 12,000 contigs had unusual junctions. Localized assembly identified that some of the many unusual junctions were the result of deletion, inversion, translocation or other phenomena and were correct, while most of these contigs had incorrect junctions of fragments from different regions of the genome. These contigs were then broken. Localized assemblies were then carried out on all fragment ends, resulting in closing all gaps except for some subtelomeric gaps.

Gap closure required multiple rounds of sequence correction and examination of remaining gaps. The most useful tools were the use of BWA (Li and Durbin 2009), samtools and bcftools (Li et al. 2009) to align the raw reads for each genome back on the assembled genome. Pilon (Walker et al. 2014) was then used to identify errors in the sequence based on the sorted BAM files. Pilon is very good at finding errors in the assembly, though the suggested corrections are less accurate, so all corrections were individually checked. Correcting point errors adjacent to gaps often allowed assembly across those gaps. It appears that many of the gaps resulted from failure by both velvet (Zerbino and Birney 2008) and ABySS (Simpson et al. 2009) to extend a contig past a region where multiple point errors had been introduced. Furthermore, from each BWA alignment, reads that failed to align to the assembled genome were extracted and built into a new fastq file. These fastq files were then *de novo* assembled with either velvet or ABySS. Nearly all of these reads are low quality sequence reads that fail to assemble into contigs, though some small contigs are generated. Blast identified that many of these small contigs were missing from the genome being analyzed, but present in other *S. cerevisiae* genomes, suggesting that they were left out of the assembly. Addition of these fragments back into the genome often resulted in extension at the site of a gap and sometimes gap closure. Corrections at a site within the existing sequence also occur. Depth of coverage analysis was also used in each round of sequence editing. The depth of coverage of the Illumina sequence data used in this project is fairly uniform across the genome, and was previously used to identify aneuploidy, the number of rDNA repeats, and other aspects of the genome (Strope et al. 2015). Sequence regions as small as 200 bases that occur twice in the genome can be distinguished from sequences present only a single time. Regions present only a single time in the genome sequence that, based on depth of coverage analysis, should be present two or more times can thus be identified and the appropriate corrections made. Sequences included twice in the assembled genome that by depth of coverage should only have been present a single time were identified and similarly could be corrected. Illumina read pair pairing information was also examined. From the BWA alignments, scripts were used to extract the pairing information. Singlets, cases where only one sequence pair aligned in an unexpected way, were skipped. When there were multiple read pair anomalies in the same region of unique sequence where either the pairs were in the wrong orientation, or the pair length was two large or small, or they paired to different chromosomes, then these regions are examined. Similarly, all locations that were not spanned by any read pairs were examined.

Sites of errors were primarily repetitive sequence regions, regions of unusually low sequence coverage, and regions where there are a high number of low confidence base calls. The transposable elements Ty and the solo long terminal repeat (LTR) sequences from these elements were particularly problematic. Significant effort was made to verify the insertion location for each Ty and LTR element. Less effort was made to get the sequences of the Ty elements themselves correct, and there are still numerous sequence errors within the sequences of the Ty elements of these genomes where there are near identical copies. Furthermore, while all the gaps in the central regions of chromosomes and overall most gaps for each chromosome could be closed, the chromosome ends are problematic due to the high number of repeats, high sequence divergence from the reference genome, and that subtelomeric and telomeric regions are sometimes duplicated, sometimes lost, and sometimes translocated to other chromosomes. As described previously, (Strope et al. 2015), crosses were carried out so the number of translocations for each strain, usually zero, is known and so the assembled genomes should be colinear. This collinearity does not apply to the subtelomeric and telomeric sequences further complicating the editing of the sequences of these regions.

Additional tools used in this analysis include blast (Altschul et al. 1990), fasta (Pearson and Lipman 1988), Smith-Waterman alignment (Smith and Waterman 1981) clustal (Higgins and Sharp 1988), tRNA_scan_SE-1.3.1 (Lowe and Eddy 1997), PlasMapper (Wishart et al. 2023), EMBOSS (Rice et al. 2000), perl (Christiansen et al. 2012), C (Kernighan and Ritchie 1988), python, LAGAN (Brudno et al. 2003), MAFFT (Katoh et al. 2002), NCBI Entrez tools (Schuler et al. 1996), and the SRA toolkit (NCBI 2011).

The S288c sequence and annotation were taken from SGD (Cherry et al. 2012, Engel et al. 2022) in April, 2023, with updates checked in April 2025, where gene information was downloaded using the yeastmine tool (Balakrishnan et al. 2012) now Alliancemine (Alliance of Genome Resources 2024), with additional information from YGOB (Byrne and Wolfe 2005), and from NCBI, including from the Short Read Archive (Sayers et al. 2022), and the genome sequences of other hemiascomycetes, particularly *Saccharomyces paradoxus* and *A. gossypii*. The genomes of *A. gossypii, Ashbya aceri, Holleya sinecauda* (sometimes called *Eremothecium sinecaudum*)*, Zygosaccharomyces rouxii, Kluyveromyces lactis, Kluyveromyces marxianus, Torulaspora delbrueckii, Lachancea kluyveri* (formerly *Saccharomyces kluyveri*), *Lachancea waltii* (formerly *Kluyveromyces waltii*), *Lachancea thermotolerans* (formerly *Kluyveromyces thermotolerans*)*, Zygosaccharomyces bailii, Eremothecium coryli* (formerly *Nematospora coryli*) and *Eremothecium cymbalariae* were used to identify orthologs present before the whole genome duplication (WGD). Genome sequences of additional *Saccharomyces sensu stricto* species *Saccharomyces mikatae, Saccharomyces kudriavzevii, Saccharomyces uvarum, Saccharomyces jurei, Saccharomyces eubayanus, Saccharomyces arboricola,* and the hybrids *Saccharomyces bayanus* and *Saccharomyces pastorianus* were also used.

For identifying genes currently annotated as verified or uncharacterized that are likely to be dubious the following criteria were used. Lack of conservation of the open reading frame in other species. Lack of conservation of the open reading frame in *S. cerevisiae* strains. Overlap with adjacent genes. Lack of RNA-seq data supporting that the gene is transcribed with reads on the plus strand with reads starting in the 5’ region. Lack of evidence of a polyadenylation site. Unusual sequence, such as long homopolymer runs. Annotated to contain an intron where there is no evidence of intron splicing. Starting methionine missing in *S. cerevisiae* strains. Location unusually close to an adjacent gene. Lack of published data as to function. Another useful phenomenon is the ratio of non-synonymous to non-synonymous plus synonymous polymorphisms in a gene across many strains. For real genes, this ratio tends to be small. It should be noted that no combination of the above criteria proves that annotated protein coding gene does not encode a protein. It is the opinion of the authors that it is time to raise the bar and that published evidence that a gene is real should be necessary to keep questionable open reading frames from being relegated to dubious.

All sequence and annotation corrections are being submitted to GenBank with the same accession numbers as in Strope et al. Table S19 (Strope et al. 2015).

## Results

In this paper the primary data investigated is 93 strains (Strope et al. 2015) and S288c, referred to here as the 94 strains. Key terms used in section headers below are described in Table 1.

### Core Genes

The set of core genes in *S. cerevisiae* is here defined as those genes (1) present at the same position and orientation, relative to flanking genes, in more than 95% of strains, and (2) includes both protein coding genes and genes for which the final product is an RNA molecule. Non-transcribed features such as centromeres, terminal telomere repeats, and origins of replication are not included. The four genes found in the plasmid 2-micron are not core, because the plasmid is present in less than 95% of strain. As previously reported the gene order of the eight conserved protein coding genes within the 93 mitochondrial genomes and the tRNA, rRNA, and RNase P RNA are the same as in S288c, except for a region duplicated in YJM1242 (Vijayraghavan et al. 2019). Thus the conserved mitochondrial genes are core genes by the definition used here, as are the two L-BC genes (Vijayraghavan et al. 2023, Vijayraghavan et al. 2023).

In this analysis a gene is considered present even if in some strains it is an apparently defective allele, a truncated allele, or an introgressed copy of the gene from a *sensu stricto* species at the conserved location and orientation. While most genes in any strain are core genes, there are also non-core genes including ubiquitous genes of variable genomic location and strain-specific genes, those found in less than 95% of strains. The central region of the genome is defined as the region of each chromosome containing all of the core gene on that chromosome, ending with the final core gene at each end.

A total of 5885 core genes have been identified, the 5521 verified genes in Table S2, and the 364 uncharacterized genes in Table S3. The verified and uncharacterized definitions come from SGD (Hirschman et al. 2006), where “verified” means a real gene of known function, and “uncharacterized” means a real gene of unknown function. Of the core genes, 41 are in the mitochondrial genome, 2 are encoded on L-BC, and the remaining 5842 are encoded on the sixteen chromosomes. Of these core genes, 91.5% are conserved with *A. gossypii*, 98% are conserved with *A. gossypii* and/or at least one other pre-WGD hemiascomycete species. Only 1.5% of the core genes listed are found only in *Saccharomyces* species. Among those protein coding core genes found only in *S. cerevisiae*, most are uncharacterized, quite small, have no known function, meaning that while they are not obviously dubious genes, some of them may turn out not to be real protein coding genes. It should be noted that as more ascomycete genome sequences, and more accurate complete ascomycete genome sequences become available the number of genes in each category may change. Some sequences were not included in this analysis as they appear incorrect, such as *L. waltii* strain NCYC 2644 sequences AADM01000488.1 and AADM01000307.1 and AADM01000647.1 that appear to more likely be *Saccharomyces paradoxus* sequence (Kellis et al. 2004). The core set of genes reported here is somewhat larger than the core set of genes reported in Peter et al. (Peter et al. 2018) of 4990 and the 4900 gene core set reported by McCarthy and Fitzparick (McCarthy and Fitzpatrick 2019). This is due to differences in the definition of what is a core gene in these papers and how a core gene is defined here. The subtelomeric regions, as the term is used in this work, refers to the regions outside the central region, but not including the terminal telomeric repeat. These subtelomeric regions contain transporters, genes in synthetic and catabolic pathways, duplicate copies of genes, genes acquired by horizontal gene transfer, and other important genes that contribute to the diversity of this species, yet they are strain and positionally variable and as such are not here considered core genes.

Genes are here identified as being part of the core set even in cases where the sequence is quite diverged, as in the case of *FIT1*, where the S288c allele and the YJM248 allele share only 52.7% identity (Figure S2), or the case of *NTC20* where the S288c allele and the YJM1400 allele share 38% identity (Figure S3). The sequence divergence of *FIT1* and *NTC20* is primarily the result of introgression with the introgressed copy replacing the *S. cerevisiae* copy at the same location and orientation.

### Non-Core Genes

Genes that are not Core. Primarily genes that are found in less that 95% of strains.

### Variably Located Ubiquitous Genes

Nomenclature is complicated for genes that are variably located, such as many subtelomeric genes and transposable elements. In addition to non-core genes that are found in less than 95% of strains, there are genes found in more than 95% of strains, but not at the same location or in the same orientation. An example of this is the two protein coding genes, GAG and POL, found in the retrotransposon Ty2. 94/94 of the strains examined contain Ty2, but no single site of Ty2 is found in more than 95% of strains.

### Central Region of Chromosome

Examination of the genomes of the 94 strains confirms a well-known pattern where the gene content of the central region of each chromosome in the genome is highly conserved between strains, but the subtelomeric regions are significantly more variable in gene content and location (Teunissen and Steensma 1995, Li et al. 2014, Monerawela et al. 2015, McIlwain et al. 2016, Stefanini et al. 2022). To determine the core gene set, the central regions of all 93 strains were carefully edited with all gaps closed and numerous errors resolved. Significant effort was made to close the final gaps in the central regions of chromosomes. At the current time, there are no remaining gaps in what is described in this work as the central regions of the chromosomes, except for gaps in some *FLO1*, *FLO5*, *FLO9* genes, genes that are at the terminus of the central regions of three chromosome ends. For the 1488 chromosomes of the 93 strains from Strope et al, there are still 2200 gaps in the subtelomeric regions, telomeric regions, and at the edge of the central region in genes *FLO1*, *FLO5*, *FLO9*, and *FLO12*. Furthermore, 2172 chromosome ends out of 2976 are Ns, not the expected terminal repeat sequence. The location of remaining gaps in the telomeric and subtelomeric regions are shown in Table S14.

To identify the boundary of the central, conserved portion of the genome, the most distal gene on each end of each chromosome where 95% or more of the 94 strains have the same gene at that position and orientation was identified. The choice of 95% was arbitrary. Table 2 shows the most distal gene of the central region of the genomes for each chromosome end.

**Table 2.**
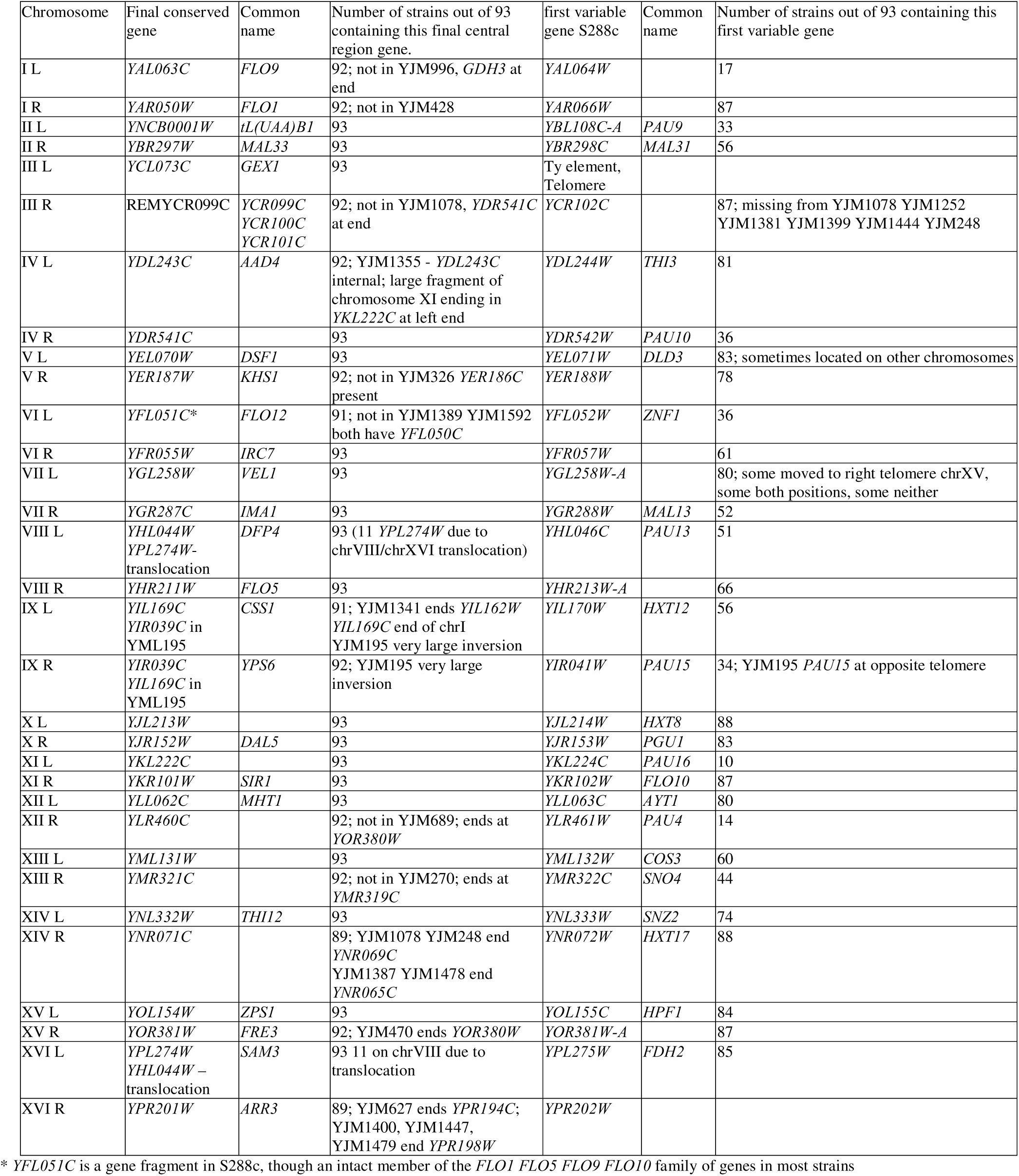
Final conserved gene for each chromosome arm.

In most cases an annotated gene marked the boundary of the central region of the genome. An exception is the left end of the central region of chromosome VI that is marked by *YFL051C*, a gene of unknown function. Examination of the 94 strains identifies that *YFL051C* is in S288c and some other strains a fragment of a gene of the *FLO1*, *FLO5*, *FLO9*, and *FLO10* family, while in other strains it is a full length flocculin gene. This gene is intact in more than half of the 93 strains. We propose that *YFL051C* be called *FLO12*, where the allele in S288c is a truncated allele (Figure S1). Another exception is the central region of the right end of chromosome III is marked by REMYCR099C, described below.

An analysis of the essential genes in *S. cerevisiae*, taken from SGD using the yeastmine tool (Skrzypek and Hirschman 2011), now part of the Alliancemine tool at the Alliance database (Alliance of Genome Resources 2024), shows that all essential genes are in the central region of the genome, as well as all tRNA genes found in S288c and all other known RNA coding genes. In addition, all 5’ UTR and coding region introns in S288c are found in the central region of the chromosomes, other than 14 introns currently annotated in putative subtelomeric genes.

Examination of RNA-seq data for these 14 putative subtelomeric introns using the RNA-Seq dataset SRR488142 (*Saccharomyces cerevisiae* S288c Project jgi.doe.gov) finds RNA spanning the splice sites but no evidence that these introns are spliced. These 14 introns shown in Table S1 are here considered dubious. Among the annotated introns eight have a 5’ GC splice site, rather than the 5’ GT splice site found for the other 277 introns. Six of these are among the dubious subtelomeric introns. A seventh one is in the gene *YJR079W*, which in this work is proposed to be a dubious gene. The eighth case is a real gene *YIL111W* (*COX5B*), and there is evidence in the SRR488142 RNA-seq dataset that this intron is spliced as annotated. The 5’ <u>GC</u>ATGT splice site is conserved in 94/94 of the genomes considered here and appears to be the only real case of an intron in S288c with a 5’ GC splice site. *YIL111W* (*COX5B*) in *S. paradoxus* also has a GC 5’ splice site, but *YIL111W* (*COX5B*) in the other *Saccharomyces sensu stricto* species have the more typical GT 5’ splice site.

The mitochondrial genome of *S. cerevisiae* is circular or circularly permuted and thus does not have variable sequence ends in the sense that chromosomes do. We consider the entire mitochondrial genome to be central. There is significant variation between *S. cerevisiae* mitochondrial genomes, but it is distributed across the genome in these 94 strains, as has been previously described (Vijayraghavan et al. 2019).

### Tandem Gene Duplication

In addition to single genes, tandem arrays of genes in the same orientation are also found in *S. cerevisiae.* For the analysis of core genes, tandemly duplicated genes where the copies are identical or nearly identical are counted only once in this analysis. For example, the 94 strains in this analysis contain from one to more than 10 copies of the *CUP1* locus (Zhao et al. 2014). *YHR053C* (*CUP1-1*) and *YHR055C* (*CUP1-2*) in strain S288c, are counted once *YHR053C*/*YHR055C* (*CUP1*) as a core gene. *ENA1*/*ENA2*/*ENA5*, *HXT6*/*HXT7*, and rDNA genes are similarly counted only once as a core gene for each array. For tandem genes where there are diverged copies such as *HXT4*/*HXT1*/*HXT5* and *PLB1*/*PLB2*, each is counted as a core gene as they are each found as diverged tandem genes in greater than 95% of strains. In three cases, *PHO3*/*PHO5*, *DOG1*/*DOG2*, and *ALD2*/*ALD3*, there are two diverged genes in S288c and many strains, but other strains have only one or the other or a hybrid of these genes. For these three genes, as 100% of strains have at least one gene, but less than 95% have both genes, these genes are here labeled as one core gene and one variable gene. The reason identical tandem duplicated genes and diverged tandem duplicated genes are treated differently is that identical tandem genes can potentially change copy number at higher frequency by unequal crossover.

Another example of diverged tandem genes is the DUP240 family of protein membrane genes. For *S. cerevisiae* there is a single copy of a DUP240 family gene *YCR007C* (*DFP3*), on chromosome III, found in all 94 strains, though in seven strains (YJM320, YJM248, YJM1304, YJM555, YJM689, YJM541, YJM554) the allele appears to be introgressed from another *sensu stricto* species. In S288c on chromosomes I there is a single DUP240 core gene *YAR023C* (*DFP1*) found in all 94 strains Adjacent to *YAR023C* (*DFP1*) is a tandem array in the opposite orientation of four DUP240 family genes *YAR027W* (*UIP3*), *YAR028W* (*KTD1*), *YAR031W* (*PRM9*), *YAR033W* (*MST28*). The 93 strains have zero to nine members of this array, with four of the strains (YJM270 YJM453 YJM1419 YJM1460) missing the array entirely, and strain YJM1434 containing a single gene in this array. Therefore, this array has one core gene. Thirty strains have the same four genes as found in S288c. In addition to the four genes found in S288c, a total of 23 different genes are found in the arrays of the other 59 strains, some of which may be introgressed from *S. paradoxus*. In some cases there are stop codons in the genes. A significant amount of genome variation can thus be attributed to the chromosome I DUP240 array. There is another DUP240 array on chromosome VII. In S288c this array contains two members, YGL051W (MST27) and YGL053W (PRM8). 67 of the 93 strains are missing this array, with three strains (YJM1208 YJM1386 YJM1419) containing the same two genes as in S288c, and the remaining 23 strains containing a varied combination of 16 different genes other than the S288c genes. Strain YJM1573 had six genes in the chromosome VII array, the most of any of these strains. These genes are not core genes, by the 95% rule. This array further contributes to genome diversity. Additional diversity for this DUP240 family is found in the 1011 *S. cerevisiae* strain sequences (Peter et al. 2018).

A summary of the tandem genes, including some of those not found in S288c is in Table S4. An example of a non-tandem gene duplication is shown in Figure S4 and includes *YJR108W* (*ABM1*), *YIL014C-A*, and *YIL102C* and possibly additional genes including some strain-specific genes and subtelomeric genes. These genes do not appear to be duplicate genes arising from the WGD event.

### Genes not in the Current Annotation

While some core genes are missing in up to 4 of the 93 strains from Strope et. al., we did not identify any core genes missing from S288c, and only one core gene is present in S288c but not in the current annotation. In Peter et al. it was reported that 3 additional genes are core genes that are not annotated in S288c. These they named *610-snap_masked-2999-BGP_1*, *584-snap_masked-1700-AIE_1*, and *611-snap_masked-3001-BGP_1* (Peter et al. 2018).

*Gene 610-snap_masked-2999-BGP_1* is a real protein coding gene that is a syntenic ortholog of *A. gossypii AER178W-A*. This gene is referred to in the YGOB browser in S288c as *YGOB_YBL026W-A* (Byrne and Wolfe 2005). The gene we are calling *YBL026W-A* is a core gene of unknown function in our analysis that was overlooked in the original annotation of *S. cerevisiae*.

In our analysis *584-snap_masked-1700-AIE_1*, located between *YPR148C* and *YPR149W* (*NCE102*), is only an intact reading frame in two of the 94 strains, YJM1078 and YJM248, where it appears to be an introgressed sequence from *S. paradoxus.* Though this sequence is found in two species, it appears that this is a dubious gene. The sequence is conserved, but the open reading frame is not conserved. *611-snap_masked-3001-BGP_1* is not a gene but a part of a gene remnant, as discussed below.

### Gene Remnants

In addition to genes, potential null alleles, pseudogenes, and chromosomal features such as centromeres and telomeres there are gene remnants in *S. cerevisiae* and other fungi, parts of which are often misidentified as small genes of unknown function. The term “gene remnant” has been used previously, initially to refer to gene fragments left over by V-J joining (Selsing et al. 1984). In this work we consider gene remnants to be a different category of feature from pseudogenes. Pseudogenes are pieces of DNA inserted at locations where they are not normally found. The definition of a pseudogene is “The term ‘pseudogene’ - defined as a region of DNA that displays significant homology to a functional gene but has mutations that prevent its expression”(Proudfoot 1980) making clear that this is not a mutant allele but is a distinct region of the genome from where the functional gene is found. Some people mistakenly use pseudogene to mean a null allele of a gene, at the genes normal location.

Gene remnants are former genes at the site where the gene was previously located, where there is no longer an intact gene, and they are truncated or have accumulated multiple stop codon and/or frameshift mutations. Comparative genomics can in many cases identify these gene remnants. These gene remnants are defined at the clade level; there is no intact gene in a clade, but intact genes can be found in strains or species outside the clade at the equivalent location. Understanding what gene remnants are present is important in understanding the gene set of the organism. In *S. cerevisiae* some of the genes that in this paper are identified as dubious are parts of gene remnants. They have homology to real protein coding genes, so they do appear to be genes. Genes recently acquired by horizontal gene transfer, introgression, gene duplication, or neofunctionalization are often cited as evidence of the recent evolution of a species, but gene loss is also important in understanding this evolution (Gladieux et al. 2014). Gene remnants are evidence of recent gene loss. For most duplicate genes from the WGD that have been lost since the WGD, there is no remaining sign of the lost gene, suggesting that they were not recently lost.

Because a region is identified as a gene remnant does not prove that none of the residual pieces are functional. For example, it has been known for more than 50 years that in *S. cerevisiae* you can have a gene with multiple point mutations, and with deletions and still have function or partial function, as has been shown for *HIS4*, a gene encoding three enzymes (Fink and Styles 1974), where gene fragments can still have one of the activities.

Below are eight examples of gene remnants in *S. cerevisiae*.

### REMYDR439C-A

One example of a gene remnant in *S. cerevisiae* is REMYDR439C-A. What we are calling “REMYDR439C-A” is not found as an intact gene in the 94 strains, or in over 1011 sequenced *S. cerevisiae* strains (Peter et al. 2018). REMYDR439C-A is found on the minus strand of S288c Chromosome IV at 1340798 to 1342041. It is a WGD paralog of uncharacterized gene *YML020W*. The localized gene order around *YML020W* and REMYDR439C-A is shown in Figure S5A.

*YDR439C-A* is still present as a gene in multiple strains of *S. paradoxus*. A partial alignment of REMYDR439C-A with *S. paradoxus YDR439C-A* and *S. cerevisiae YML020W* is shown in Figure S5B. *S. paradoxus* strain CBS432 *YDR439C-A* shares 42% protein identity with the *S. paradoxus* WGD paralog *YML020W*. The DNA sequences are 66% identical. The loss of *YDR439C-A* appears to have occurred in the ancestor of *S. cerevisiae* since the divergence of these species. This is a rare example of a duplicate gene pair from the WGD where one of the members was relatively recently lost, since the divergence of these closely related species. Note the stop codons in *S. cerevisiae*.

Additional evidence of this remnant in the *S. cerevisiae* genome is that in S288c the gap between adjacent genes *YDR439W* and *YDR440W* is 1720 nucleotides, larger than the typical distance between genes. The gene remnant is 1243 nucleotides, explaining 72% of this putative intergenic region.

There are 21 frameshifts in the alignments of *S. paradoxus YDR439C-A* and *S. cerevisiae* REMYDR439C-A. 14 of these are deletions, 202 bases in total, in *S. cerevisiae*, and 7 are insertions, 39 bases in total. Of the 21 frameshifts, 2 are in-frame, 19 are out of frame. There are also 5 in-frame stop codons in REMYDR439C-A.

### REMYFL041W-A

A second *S. cerevisiae* gene remnant REMYFL041W-A is located on S288c chromosome IV from 47982 to 48942 on the Watson strand and spans the Peter et al gene *611-snap_masked-3001-BGP_1*. Putative gene *611-snap_masked-3001-BGP_1* is only intact in about half of the 94 strains we examined. Furthermore, REMYFL041W-A overlaps uncharacterized gene *YFL041W-A*. Analysis of *YFL041W-A* shows that is has similarity to the C-terminal end of *YLR070C* (*XYL2*), a xylitol dehydrogenase (Richard et al. 1999). In *S. paradoxus* strains N44 and CBS432, *YFL041W-A* is a full-length WGD paralog of *YLR070C* (*XYL2*) (Philippsen et al. 1997, Wolfe and Shields 1997). Additional related non-core subtelomeric dehydrogenases are found in some strains, with two found in S288c, *SOR1* and *SOR2*. There are 78 copies of these genes in the 93 strains, with strains having from zero to two copies of these genes located near the telomeres on seven different chromosomes. Thus *S. cerevisiae* has lost one dehydrogenase, now REMYFL041W-A, since the divergence from *S. paradoxus* but some strains have gained additional dehydrogenases by acquiring duplicate sub-telomeric copies. Both dubious hypothetical gene *610-snap_masked-2999-BGP_1* and dubious gene *YFL041W-A* (Kessler et al. 2003) overlap the *S. paradoxus* gene but are in different reading frames. Part of the reason some people might be inclined to think that *YFL041W-A* is a real gene is that it has protein homology to real genes, though this homology is just fragments of a lost gene.

In addition, in *S. paradoxus* strains UWOPS91, UFRJ50816, and YPS138 (Bergstrom et al. 2014), *YFL041W-A* is a disrupted open reading frame, though much more intact than the *S. cerevisiae* copies. Thus *YFL041W-A* and *611-snap_masked-3001-BGP_1* are both dubious and are not uncharacterized genes nor pseudogenes but are parts of a gene remnant (Figure S6A). There is no intact *S. cerevisiae* copy of the *YFL041W-A* paralog of *YLR070C* in the 94 strains examined here or in the 1011 genome sequences from Peter et al. This remnant includes seven frameshifts and ten in-frame stop codons in S288c as well as numerous synonymous and non-synonymous polymorphisms (Figure S6B). A well conserved region of REMYFL041W-A and *YLR070C* is shown in Figure S6C. The remnant is somewhat shorter than the *S. paradoxus* gene and the orthologs found in other species, due to multiple deletions having occurred in the *S. cerevisiae* lineage.

Interestingly, *S. kudriavzevii* appears to have a gene remnant REMYLR070C, as well as a gene remnant REMYFL041W-A (Figure S6D). In this species both orthologs of *A. gossypii ABR229C* are lost, though there are two full length near-telomeric copies of these genes, where near-telomeric means that it is not known in this species if these are subtelomeric or are in the central region of those chromosomes. *S. mikatae* (Figure S6E) and *S. arboricola* (Figure S6F) also have gene remnants of *YLR070C* though intact *YFL041W-A* genes.

The most conserved portion of *S. cerevisiae* REMYFL041W-A with intact *YFL041W-A* genes is shown in figure S6G.

*YFL041W-A* and *YLR070C* and the gene remnants and the multiple non-WGD paralogs in these *Saccharomyces sensu stricto* species are highly diverged and suggest rapidly evolving dehydrogenases.

### REMYKL162C-A (REMPIR6)

A third *S. cerevisiae* gene remnant we are calling REMYKL162C-A (REMPIR6) is in S288c on the Crick strand of S288c chromosome XI from 145880 to 146662 adjacent to *YKL163W PIR3* (Figure S7A) and is not found as an intact gene in the 94 strains or the 1011 sequenced *S. cerevisiae* strains. This gene remnant overlaps the dubious hypothetical gene *YKL162C-A*. *A. gossypii* has three PIR genes and some *Saccharomyces sensu stricto* species have six, but *S. cerevisiae* has only five of these cell wall proteins (Toh-e et al. 1993, Mrsa et al. 1997) with a gene remnant of the ancestral location of the sixth gene. REMYKL162C-A (REMPIR6) is a WGD paralog of *YJL158C* (*CIS3, PIR4*). REMYKL162C-A contains four frameshifts and four in-frame stop codons when compared with the intact *PIR6* in *S. paradoxus* strain CBS432 and *S. mikatae* strain IFO1815. In *A. gossypii* and several other pre-WGD there are three intact genes, though there has been an inversion in the *A. gossypii* lineage so *AEL103W*, the ortholog of paralogs REMYKL162C-A REMPIR6 and *YJL158C CIS3*,*PIR4* is now separated from *AEL110W* and *AEL111C* by six genes and the gene orientation is reversed.

In REMYKL162C-A (REMPIR6) in S288c there are eight frameshifts, 6 of which are out of frame, 3 deletions, 4 bases total, and 5 insertions, 10 bases total (Figure S7B). There are also 5 in-frame stop codons. *S. paradoxus YKL162C-A PIR6* and *S. cerevisiae* REMYKL162C-A REMPIR6 have 75% nucleotide sequence identity. A conserved N-terminal end region of *PIR1* – *PIR5*, and REMPIR6 is shown in Figure S7C.

### REMYCR099C (REMVTH4)

A fourth *S. cerevisiae* gene remnant we are calling REMYCR099C (REMVTH4) is a paralog of *YBL017C* (*PEP1*) (Marcusson et al. 1994). *YBL017C* (*PEP1*) is the one syntenic copy of the gene *AFR018C* in *A. gossypii*. REMYCR099C spans *YCR099C* which we are proposing is dubious. In S288c there are two additional subtelomeric full-length copies, *YIL173W* (*VTH1*) and *YJL222W* (*VTH2*). There is an additional disrupted copy *YNR066C_YNR065C* (*VTH3*), disrupted by a single stop codon, with an intact open reading frame in 30 of the 93 strains. The gene remnant REMYCR099C (REMVTH4) is the most distal element in the central region of S288c chromosome III near the right telomere on the Crick strand from 300104 to 303030 and spans proposed dubious genes *YCR099C*, *YCR100C*, and *YCR101C*. It is disrupted by numerous frameshifts and stop codons in strain S288c. This region is a gene remnant, even though there is an intact copy in one of the 93 strains, strain YJM1252, and in 8 of the 1011 genome sequences (Peter et al. 2018). The YJM1252 *VTH4* gene and the 8 genes from the 1011 genome sequences are not an *S. cerevisiae* gene, but an introgressed *VTH4* gene from *S. paradoxus* that is 99% identical to that gene in *S. paradoxus* strain CBS432. No intact non-introgressed *S. cerevisiae VTH4* gene was identified. The multiple stop and frameshift mutations in *S. cerevisiae* REMYCR099C (REMVTH4) suggests that introgression would be the primary means of reviving *VTH4*.

While *A. gossypii* has one copy of this gene *AFR018C* and no paralogs, there are five features in *S. cerevisiae*; *PEP1* found in the central region of the genome, and non-tandem duplicate subtelomeric genes *VTH1* and *VTH2* (Westphal et al. 1996), the near telomeric gene *VTH3* that often has a stop codon in it, and near telomeric REMVTH4, which in at least one strain has been replaced by an introgressed *VTH4* gene. This suggests rapid evolution of this gene family in the *S. cerevisiae* lineage. In *S. uvarum* strain CBS7001 there is a tandem triplication of *YBL017C,* so this gene has been duplicated by both tandem and non-tandem duplication. The three tandem copies of *S. uvarum* strain CBS7001 *YBL017C* share approximately 92% protein identity with each other, so they have diverged significantly since the triplication. As with REMYFL041W-A, REMYCR099C (REMVTH4) seems to be part of a rapidly and recently evolving enzymatic activity.

REMYCR099C (REMVTH4) in S288c contains 16 frameshifts relative to the *S. paradoxus YCR099C* 9 of which are out of frame. 8 of these frameshifts are deletions, 1833 bases deleted, 8 of these frameshifts are insertions, 112 bases inserted. Five in-frame stop codons are found in REMYCR099C (REMVTH4). A partial alignment is shown in Figure S8.

### REMYCL001W-A

A fifth *S. cerevisiae* gene remnant is REMYCL001W-A, which includes genes here proposed to be dubious *YCL001W-A* and *YCL001W-B* and spans S288c Chromosome III from 113002 to 113982. This gene remnant is a WGD paralog of *YNL001W* (*DOM34*) (Figure S9A). In seven of the 93 strains (YJM320, YJM248, YJM1304, YJM555, YJM689, YJM541, YJM554) and 66 of the 1011 genome sequences (Peter et al. 2018) there is a full length *YCL001W-A* allele that has been acquired by introgression and is 100% identical to the intact allele found in *S. paradoxus* strain CBS432. This is similar to the case of *VTH4* described above where the gene was recovered by introgression. The allele found in S288c and the other 86 of the 93 strains contains multiple stop codons and a frameshift.

S288c REMYCL001W-A contains 12 frameshifts, 8 deletions, 94 bases, and 4 insertions, 12 bases. 9 of the frameshifts are out of frame in S288c. There are also three in-frame stop codons. Alignment of *S. paradoxus YCL001W-A* with S288c REMYCL001W-A is shown in Figure S9B.

### REMYHR054

A sixth *S. cerevisiae* gene remnant we are calling REMYHR054 arose differently, not through the accumulation of mutations in a WGD paralog. It is a duplicate 3’ end fragment of *YHR056C* (*RSC30*) or *YHR052W* (*CIC1*) or both that results from the tandem duplication of *CUP1* and is only found in strains with two or more tandem copies of *CUP1*(Figure S10A). It is here proposed that hypothetical gene *YHR054C* is dubious, and a gene remnant not a protein coding gene. As previously published, different *S. cerevisiae* strains contain at least five independent duplications of the *CUP1* locus (Figure S10B), so the length of REMYHR054 varies (Zhao et al. 2014). In this case the gene remnant name does not have a letter indicating the strand, as the formerly coding region can be on one or the other or both strands. Strains with three or more copies of *CUP1* have more than one copy of REMYHR054. In S288c REMYHR054 extends on Chromosome III from 213185 to 214249 on the Crick strand. This gene remnant is the one most similar to the initial use of the term, describing fragments left over by V-J joining (Selsing et al. 1984).

### REMYDR508W-A

A seventh *S. cerevisiae* gene remnant we are calling REMYDR508W-A is similar to REMYHR054 in that it arose from the tandem duplication of *YDR508C*, as shown in Figure S11A. REMYDR508W-A is a degraded fragment of *YDR510W*. REMYDR508W-A is found in 2/94 strains, YJM1400 and YJM1479, and 12 of the 1011 strains (Peter et al. 2018) but not in S288c. An alignment of *YDR510W* with REMYDR508W-A, both from strain YJM1400, is shown in Figure S11B, showing a frameshift and two stop codons as well as multiple non-synonymous polymorphisms. REMYDR508W-A is similar to REMYHR054 in that both originated from tandem duplication.

### REMYFR054C (REMFLO13)

An eighth *S. cerevisiae* gene remnant we are calling REMYFR054C (REMFLO13) is from 256524 to 260378 on Chromosome VI on the Crick strand and spans the open reading frame *YFR054C* that we are here calling dubious, but part of this gene remnant. This gene is a flocculation gene similar to *FLO1*/*FLO5*/*FLO9*/*FLO10*/*FLO12* and is found as an intact gene in some strains of *S. paradoxus, S. uvarum, S. kudriavzevii,* and *S. eubayanus*. Three strains, YJM248, YJM1252, and YJM1078 have intact versions of this gene that appear to be introgressed from *S. paradoxus*, and YJM1399 has a frameshifted copy of this gene, also apparently introgressed from *S. paradoxus*. The remaining 89 of the 93 strains plus S288c all have what appears to be the *S. cerevisiae* version with numerous stop codons and frameshift mutations. This gene remnant is in a gap of 9155 nucleotides between the flanking genes *YFR053C* (*HXK1*) and *YFR055W* (*IRC7*). This gene remnant is on the Crick strand of S288c chromosome VI between 256456 and 260378, in the central region of this chromosome, and is a WGD paralog of *YHR211W* (*FLO5*). This is a rather large gap between genes for *S. cerevisiae*, and this gene remnant explains forty percent of it.

### A gene remnant in *A. gossypii,* REMADL001C-A

Gene remnants seem to be found in most fungal species. While the above gene remnants were observed in *S. cerevisiae* in comparison with *A. gossypii* and species that diverged from the *Saccharomyces* lineage before the WGD, REMADL001C-A is an example of a gene remnant from a pre-duplication species. *A. gossypii* REMADL001C-A is an ortholog of *ADL001C-A* in *A. aceri* and *YIL001W* in *S. cerevisiae.* REMADL001C-A is found on the Watson strand of strain ATCC10895 of *A. gossypii* chromosome IV between 705800 and 706211. While this is an intact gene in *A, aceri*, in *A. gossypii* REMADL001C-A contains 22 frameshifts, 18 deletions of a total of 247 bases, and 4 insertions of a total of 5 bases. 18 of the insertions and deletions are out of frame. There are also seven in-frame stop codons in REMADL001C-A. The gene order around REMADL001C-A is shown in Figure S12.

### Variably located ubiquitous genes and strain-specific genes, plasmids, and transposable elements

In these 94 strains, there are also a few variably located ubiquitous genes and strain-specific genes. These include variably located ubiquitous genes in the central region of the chromosomes and those located in both the central region and the subtelomeric regions, as described in Table S5. These include sixteen non-core central region genes found in S288c. Strain-specific genes can be found both in the central region of the chromosomes, and more frequently in the subtelomeric regions.

A special case is the plasmid 2µ found only in 67 of the 93 strains(Strope et al. 2015). The four genes found on this plasmid are not part of the core set. Similarly, the single and double strand RNA virus genes though found in most strains are variable among strains and not core except for the two genes found in L-BC (Vijayraghavan et al. 2023, Vijayraghavan et al. 2023). Another special case is the transposable elements. While all 94 strains in this analysis contain the ubiquitous Ty element Ty2, a variable number of strains contain Ty1, Ty3, Ty4, and Ty5 (Table S6). As the location of those elements varies significantly, the Ty *GAG* and *POL* genes found in these transposable elements do not meet the definition of core genes used here, even in the case of Ty2. Only these five types of known Ty elements were found among the 94 strains, though a long terminal repeat (LTR) from an element we are calling Ty6 is located near the left telomere of chromosome II of strain YJM451, where Ty6 is a Ty element found in *S. cerevisiae* on contigs AFQ_1-6592, CDM_4-1998, and YDB-17588 from the 1011 genomes project (Peter et al. 2018), and found in many *S. paradoxus* strains including YPS138 and UFRJ50816 (Bergstrom et al. 2014). This element appears to be related to Ty4, and has been referred to as TSU4 (Bergman 2018).

### A Novel *S. cerevisiae* Transposon

A curious case of strain-specific genes is a circularly permuted 17kb set of five genes apparently derived by horizontal gene transfer from *Z. bailii* or a closely related species. Complete copies of this 17kb region are found in sixteen of the 93 strains, with partial portions of this 17kb unit found in an additional ten strains. Several strains contain more than one copy. The partial and full copies are at 22 genomic locations, a mixture of central and subtelomeric (Table S7). Among the 1011 strains from Peter et al. a blast search suggests at least 383 strains contain partial or complete copies of this 17kb element. This appears to be a novel transposon that we are calling Ty7, shown in Figure 1. Ty7 encodes 5 protein coding genes, including an ortholog of *OXP1* a 5-oxoprolinase (Kumar and Bachhawat 2010, Lu et al. 2010). In strain YJM1478, the circle is recombined into the *S. cerevisiae YKL215C* (*OXP1*) locus at a site of DNA homology within this protein coding gene, creating two hybrid *OXP1* genes flanking the 17kb insert. Some locations of the Ty7 insertion sites are shown in Table S7 and Figure 1. The remaining insertion sites do not have homology and do not have a duplication or deletion at the insertion site of more than 5 bases. The 17kb circularly permuted sequence does not contain direct or inverted repeat sequences. In one central region case, found in strains YJM271 and YJM193, two complete tandem copies of the 17kb unit were inserted into a site on chromosome XIII. The largest gene in this 17kb sequence, and the one most similar to an *S. cerevisiae* gene is an ortholog of *OXP1*. The other four genes are weakly similar to a flocculin gene, a transporter, and two are weakly similar to transcription factors. In *Zygosaccharomyces* species from which this transposon was acquired this gene cluster is located at a single location near a telomere; there is no evidence that this element is transposable in those species, though only a small number of genome sequences is available for these species, so this could change. Curiously, the *S. cerevisiae OXP1* gene is also located near a telomere, though the non-syntenic *A. gossypii* ortholog *ABR114C* is in the central region of the *A. gossypii* genome. *T. delbrueckii* also has a subtelomeric *OXP1* gene (*TDEL_0D00250*) that is not syntenic with the *A. gossypii, S. cerevisiae,* or *Zygosaccharomyces* genes, and in *L. kluyveri* the gene is at yet another non-syntenic location. *OXP1* thus seems to be unusual in the lack of conservation of location at which the gene is found. Although Ty1 through Ty6 elements are retrotransposons this was not known when Ty1 was first described (Cameron et al. 1979); Ty7 does not appear to be a retrotransposon. We are calling the encoded genes *TYG71* for the *OXP1* ortholog, and *TYG72*, *TYG73*, *TYG74*, and *TYG75* for the other four genes. No introns or ribosomal frameshifts are found in these genes. A single copy of Ty7 from strain YJM193 has been deposited in GenBank with accession number OR078387. An example of a mutation caused by this transformation is in strain YJM453 where the gene *YOL030W* (*GAS5*) is disrupted by a Ty7 element.

**Figure 1.**
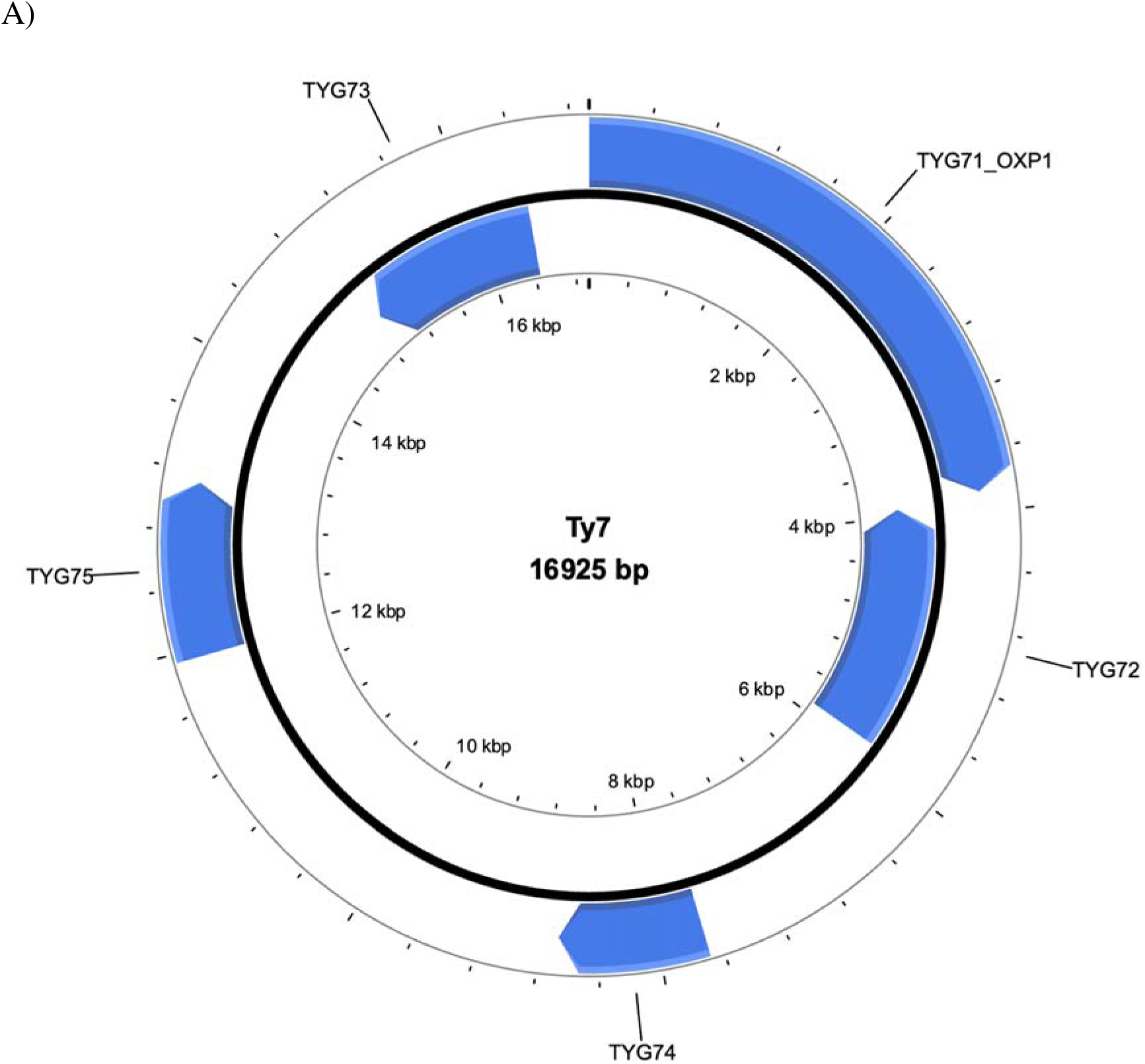
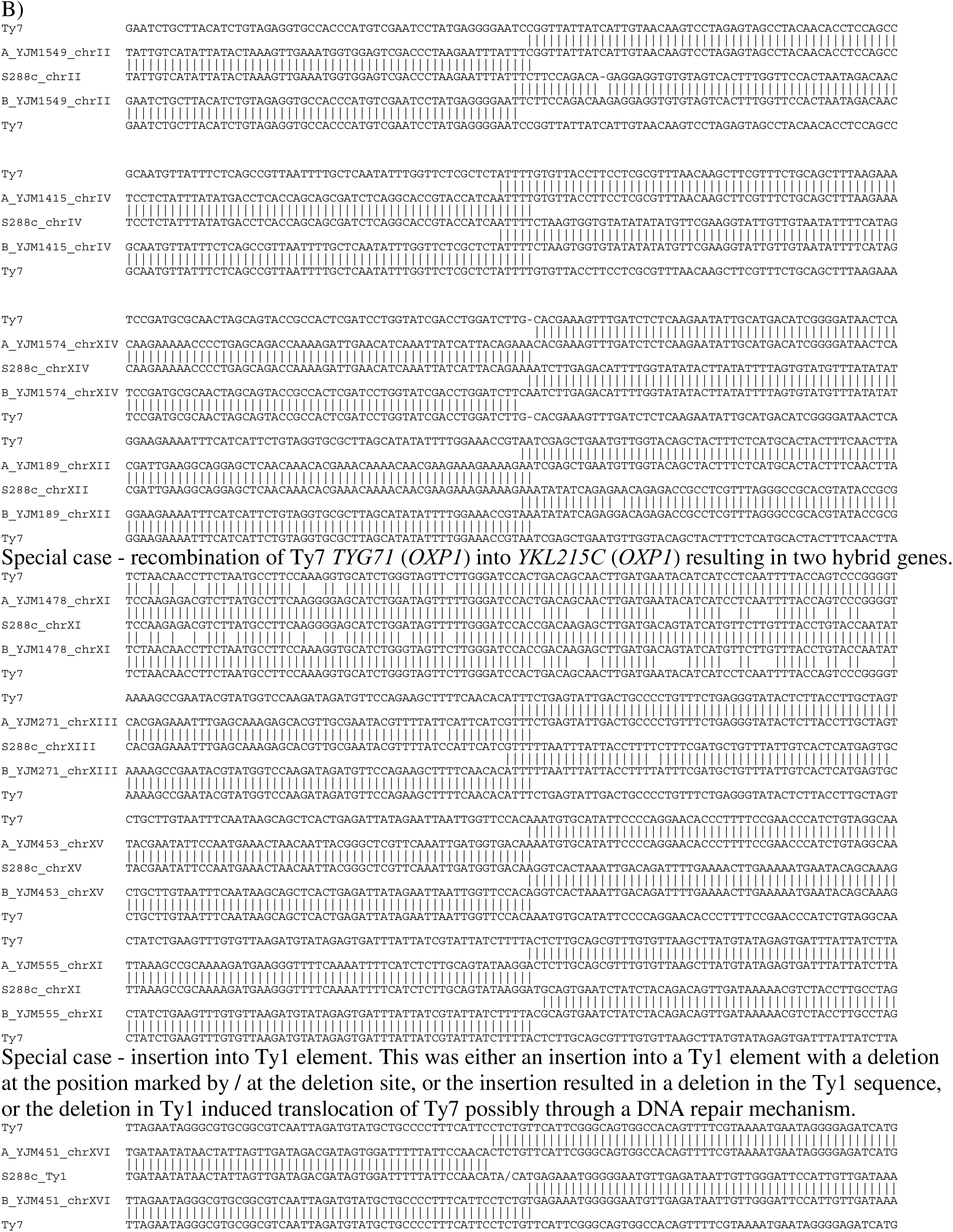
Ty7. A) Map of Ty7. Start codon of *TYG71* was arbitrarily selected as nucleotide one of this map. The five protein coding genes found in this transposon are shown. Image created using PlasMapper (Wishart et al. 2023). B) Insertion sites of Ty7. S288c is used for comparison as it contains no Ty7 elements. “Ty7” is the consensus Ty7 sequence. A_ and B_ are the two ends of the inserted Ty7. Note that Ty7 elements are circularly permuted and different sites in the element are found at the insertion junctions.

### Proposed Dubious Genes

In this analysis, 227 protein-coding genes in the current annotation of *S. cerevisiae* strain S288c are likely annotation artifacts, and thus these are not included in the lists of core genes and non-core genes for *S. cerevisiae*, and we consider them to be dubious. A list of these proposed dubious genes is in Table S8. This list does not include genes previously described as dubious in *S. cerevisiae* (Hirschman et al. 2006, Cherry et al. 2012). For most of these genes proposed as dubious, the open reading frame is quite short, they are not conserved in other species, and in many cases they are not even conserved among strains of *S. cerevisiae.* For all 237 proposed dubious hypothetical orfs, genes there is little to no experimental data supporting that they are actual genes. For the purposes of describing the set of genes in the species *S. cerevisiae*, they are dubious. There have been cases of genes listed as dubious that have later turned out to be genuine, such as *YIL156W-B*, a syntenic ortholog of *A. gossypii AFR298C* once listed in SGD as dubious. A gene more recently moved from dubious to verified is YIL059C, though uncharacterized might be a more accurate description. It overlaps with YIL060C, here proposed to be dubious. With sufficient evidence it is possible that one or more of these dubious genes may turn out to be genuine genes and not annotation errors. Without supporting evidence they should be considered dubious. One current dubious gene, *YBR012C*, appears not to be a dubious gene but instead *YBR012C* and *YBR013C* appear to be a single gene disrupted by a Ty1 element in S288c (Table S12), and thus we propose that *YBR012C* and *YBR013C* should be considered a single gene, a potential null allele in S288c. *YBR012C* is not a dubious gene as currently annotated, but an alias for *YBR013C*. Most of these dubious genes date back to the original annotation of the genome of *S. cerevisiae* strain S288c, when it was the first eukaryotic genome sequenced, and thus limited data was available for comparative analysis, with some additional dubious genes coming from later work (Kessler et al. 2003).

One case of a gene here proposed as dubious, *YJR079W*, contains a putative intron with a non-consensus 5’ splice site (GCATGT versus the consensus GTATGT). Out of 329 sequence reads at the putative splice site, there were no spliced transcripts found in SRR488142. The intron as well as the gene is dubious.

Another gene proposed here as dubious is *YER014C-A* (*BUD25*). This gene is not conserved in other species as a spliced open reading frame, likely due to the limited conservation of the overlapping, conserved *YER014W* (*HEM14*) gene. *HEM14* has only 44% protein identity between *S. cerevisiae* and *A. gossypii*. The putative *YER014C-A* open reading frame is disrupted in other species. While the open reading frame is present in 90/93 of the Strope et al strains, of 237 RNA-seq sequences spanning the putative *BUD25* splice site from the SRR488142 RNA-seq data set, all are unspliced.

Another case of a proposed dubious gene that contains an intron is more interesting. For proposed dubious gene *YBR090C*, the only conserved portion of this gene is the intron splice sites, branch site, and the intron’s location immediately upstream of *YBR089C-A* (*NHP6B*) (Figure S13). The putative coding sequence is not conserved. This is a real intron in a dubious gene, a 5’ UTR intron of *YBR089C-A* (*NHP6B*), not a separate protein-coding gene. Examination of RNA-Seq data SRR488142 shows evidence that this intron is spliced as expected. This 5’ UTR intron of *YBR089C-A* (*NHP6B*) is conserved in multiple species, and the gene is a syntenic ortholog of *ADL310W* in *A. gossypii*, which also has this 5’ UTR intron.

An interesting case of two proposed dubious genes are *YHR052C-B* and *YHR054C-B* recently published by Cuitong He (He et al. 2018). These putative genes were discovered using proteomics, showing quite convincingly that these proteins can be detected experimentally. During proteomic analysis the most common approach is to use the set of annotated proteins and identify those present or absent. There are generally multiple spectra that do not correspond to the annotated protein set. The work by He et al shows the value of examining these unexplained fragments. *YHR052C-B* and *YHR054C-B* appear to be real peptides, but here we are proposing that these are dubious genes (Figure S14). The reason for this is as follows.

These proteins are on the same strand and overlap with the identical tandem duplicate genes *YHR053C* (*CUP1-1*) and *YHR055C* (*CUP1-2*) over approximately the entire length of both proteins. *YHR052C-B* is in the +1-reading frame relative to *CUP1*. These are very short proteins, *YHR053C* 62 residues, *YHR052C-B* 58 residues. While the protein sequence of *YHR053C* is identical for all 94 strains, there are four polymorphisms in *YHR052C-B* among the sequences from the 94 strains changing the protein sequence, one is a conservative change, one loss of start codon, and two non-conservative changes. More strikingly, when comparing these open reading frames across the *Saccharomyces sensu stricto* species within *CUP1* there are forty-five positions where the protein sequences are identical, and eight positions with only conservative amino acid substitutions. In contrast for the *YHR052C-B* orthologs, there are only 14 conserved positions, and four positions with only conservative changes. Loss of start codon is found in several species. It thus appears that selection has been acting on *CUP1*, but not on *YHR052C-B*, with most of the polymorphism being in the 3^rd^ position, the wobble position, of residues of *CUP1*, but most of the polymorphism are in the 2^nd^ position of residues of *YHR052C-B*, a very unusual situation for a protein coding gene. We are proposing that *YHR052C-B* and *YHR054C-B* encode peptides, but these are not genes but are more analogous to the case of uORFs where translation occurs, but they seem to have a role in the regulation of translation of the associated gene and are not independent genetic elements. *YHR052C-B* and *YHR054C-B* thus appear to be features of *YHR053C* (*CUP1-1*) and *YHR055C* (*CUP1-2*), not distinct genes.

A similar case is *YKL104W-A* (He et al. 2018). *YKL104W-A* is on the opposite strand of *YKL103C* (*GFA1*), and is conserved in *S. cerevisiae*, and to a much lesser extent in *sensu stricto* species. As shown in Figure S15, when *YKL104W-A* is aligned to the S288c sequence, of the seventeen polymorphisms sixteen are in codon position one and one is in codon position two. None are in the third codon position, the wobble position, where polymorphisms are generally found in protein coding genes. Reading frame one of *YKL104W-A* is on the opposite strand from reading frame three of *YKL103C* (*GFA1*). Those sixteen polymorphisms are in the wobble position of *YKL103C* (*GFA1*). It appears that *YKL103C* is under selective pressure, not *YKL104W-A*. Thus, although *YKL104W-A* appears to encode a protein, it is not a real protein coding gene.

For gene *YLR379W-A*, this also appears to be dubious based on lack of conservation of the reading frame in other species, as well as limited conservation in *S. cerevisiae* (Figure S16). Multiple strains have two base frameshifts at the sequence AGAGAGAGAGAGA which occurs at fourteen locations in the S288c genome, though only three times in coding regions. The three coding region occurrences are dubious gene *YLR311C*, gene *YLR255C* proposed as dubious, and *YLR379W-A*, also proposed as dubious. In addition, single nucleotide polymorphisms in this gene are a mix of all three codon positions, not the usual pattern where most polymorphism are in the wobble position.

Yet another case of an erroneous intron annotation is *YJR112W-A*. In this case this is a real, core gene, but instead of a non-consensus sequence intron this gene contains a consensus plus one ribosomal frameshift site, CTTAGGC (Belcourt and Farabaugh 1990) (Figure S17). This gene has a conserved plus one ribosomal frameshift site also found at the same position in orthologs from several other species, including *A. gossypii, S. paradoxus,* and *S. bayanus.* Some other related species including *K. lactis* have neither an intron nor ribosomal frameshift in their ortholog. Supporting this being a case of ribosomal frameshifting in *S. cerevisiae* and not an intron is the non-consensus splice and branch sites, and the lack of evidence from RNA-seq data of splicing of the putative intron. For sequence reads from SRR488142 RNA-Seq data set, 0/171 reads matching the 5’ side of the putative splice site were spliced, and 0/492 reads matching the 3’ side of the putative splice site were spliced.

In *S. cerevisiae* there are only 285 annotated introns, not including the mitochondrial genome introns, tRNA introns, the dubious introns from *YER014C-A* (*BUD25*), *YJR079W*, *YJR112W-A*, and the dubious subtelomeric introns. All real introns in *S. cerevisiae* S288c are in core genes, and in no case among the 94 strains is an intron present in a core gene in some strains but missing from the gene in other strains. While the nuclear encoded protein and tRNA introns are highly conserved, the intron composition of the mitochondrial genomes is quite variable, as previously described (Vijayraghavan et al. 2019). Variation in the presence of mitochondrial introns, and the encoding of genes within mitochondrial introns is another phenomena that has long been known (Jacquier and Dujon 1985), but is much more clear with the availability of large numbers of complete mitochondrial genome sequences to compare.

The *NAG1* gene is on the minus strand and is completely contained within the *YGR031W* (*IMO32*) gene. While *NAG1* was reported to be highly conserved (Ma et al. 2008), it appears that the open reading frame is only conserved in *S. cerevisiae*, *S. uvarum*, and *S. arboricola*. In other *Saccharomyces sensu stricto* species, and other hemiascomycetes the open reading frame is not conserved (Figure S18), though segments of the sequence are conserved, corresponding to conserved regions of *IMO32*.

An interesting case of a dubious gene is *YNL269W* (*BSC4*). This gene has previously been described as a real gene, arising *de novo* by mutation (Cai et al. 2008). However, there is a lack of data as to any function of this gene The paper by Cai et al reports that by RT-PCR it is possible to detect transcription of this region, but this is a very low bar. Examination of publicly available NCBI short read archive illumina RNA-seq data, nanopore RNA-seq data, and Pacific Biosciences RNA-seq data for *S. cerevisiae* shows some transcripts spanning portions of this region, mostly on the wrong strand, and no evidence of transcription start sites or polyadenylation sites corresponding to the expected locations for this dubious gene. Furthermore, the Cai et all paper reports that “sequences from 29 strain samples that are from different localities or origins showed that it is fixed and the ORFs are conserved in all *S. cerevisiae* populations.” This is rather questionable, as among the 94 *S. cerevisiae* strains we have investigated, *YNL269W* (*BSC4*) is only intact in 49 strains, and in the 1011 strains it is only intact in 504 strains. In approximately half of all strains from each set, the open reading frame is disrupted by one or often more than one frameshift or stop codon (Figure S19). Frameshifts significantly outnumber stop codons, as this typical non-coding region contains multiple homopolymer runs that are prone to frameshift mutations.

Another aspect of *YNL269W* (*BSC4*) is that this dubious gene would be divergently transcribed from the adjacent real gene, *YNL270C* (*ALP1*), and that these coding regions are separated by only 38 bases. Examination of divergently transcribed genes in *S. cerevisiae,* not including dubious genes, and correcting several currently incorrectly annotated start codons (Figure S21), indicates that only 63 out of 1318 divergent pairs are less than 200 bases apart, and that the closest five divergent gene pairs are: YAL046C, YAL044W-A 133 bases apart, YLR118C, YLR119W 126 bases apart, YML049C, YML048W 121 bases apart, YGL221C, YGL220W 82 bases apart, and YDR512C, YDR513W 77 bases apart. This dubious gene is suspiciously close to the real adjacent gene. Overall it appears to be unlikely that *YNL269W* (*BSC4*) is a real protein coding gene, and thus here we are proposing that it is dubious. Cai et al propose this as an example of a protein coding gene that has arisen *de novo* by mutation. At the very best this is a terrible example of such a gene, as for this putative gene there is no evidence it is transcribed, no evidence of function, no evidence of localization or other information known about most verified genes. To show that genes can arise by mutations in non-coding regions, a better example is needed.

An example of the problem of open reading frames that are unlikely to be genes being annotated as genes is *YFR035W-A* (Wacholder and Carvunis 2023, Wacholder et al. 2023). In the work by Wacholder et al they examined all open reading frames, including those overlapping annotated genes and *YFR035W-A* was one of the two best candidates to be an overlooked gene, despite it almost entirely overlapping an annotated gene. *YFR035W-A* is buried under dubious gene *YFR034W-A* and a gene we are here proposing is dubious *YFR035C*. While it is reported that the deletion of *YFR035C* has a phenotype (Willingham et al. 2003), that deletion also deletes *YFR035W-A*. *YFR035W-A* is conserved in pre-WGD species and post-WGD species, whereas conservation of *YFR035C* is more limited. Also, as seen in Figure S28, the start codon from *S. paradoxus*, and the open reading frame conservation suggests that *YFR035W-A* is more like to be a real protein coding gene than *YFR035C*. We agree that *YFR035W-A* as published by Wacholder et al is a real gene of unknown function and that *YFR035C* should be relegated to dubious.

Another gene we are proposing is dubious is *YNL040C-A*, 26 amino acids in length. Out of the 93 strains, 91 full length, two 5’ ends missing, no in-frame start codons. Open reading frame not conserved in related species (Wacholder et al. 2023). Most transcripts spanning *YNL040C-A* are on the wrong strand in file SRR11550239 (Jenjaroenpun et al. 2021) and several other long read RNA-seq files investigated. The very short open reading frame has some similarity to a short open reading frame found in multiple bacterial species, including *Butyricimonas paravirosa* strain DSM 105722, but it is a small open reading frame on the minus strand of the coding region of a well conserved gene encoding NADH:ubiquinone reductase (Na(+)-transporting) subunit F. Some species have the conservation but are missing the start or stop codon.

Of the genes proposed here as dubious, 122 genes were previously proposed as dubious by a different method (Zhang and Wang 2000) and are marked in Table S8. The approach of Zhang and Wang is a version of codon usage, whereas the method used here to identify dubious genes is based on open reading frame conservation, protein sequence conservation, RNA-seq data, lack of known phenotype. These are two independent methods. Over 1000 additional small open reading frames can be found in the sequence of the 93 strains from Strope et al. that are at least equally likely to be species and strain-specific genes as the open reading frames that here have been relegated to dubious. Naming these 1000+ open reading frames as novel uncharacterized genes would serve no purpose. Among those 1000+ open reading frames, a small number of these species and strain-specific open reading frames may eventually be shown experimentally to be protein coding genes involved in specific processes, carrying out specific functions. It would only then be appropriate to welcome them into the pantheon of *S. cerevisiae* genes.

### Core genes found only in *S. cerevisiae*

Most of the protein coding genes previously described in *S. cerevisiae* S288c as dubious, and most of the genes here described as dubious are intact open reading frames only in *S. cerevisiae* and are generally only intact open reading frames in a subset of strains. Because *S. paradoxus* and *S. cerevisiae* share approximately 90% DNA sequence identity across the genome, generally these dubious genes show some protein sequence conservation with *S. paradoxus*, but the open reading frame is usually not conserved. Other than dubious genes and genes here proposed to be dubious there are thirty-nine core genes found only in *S. cerevisiae*, one of which looks possibly to be a real gene. That gene, *YMR008C-A,* is 155 amino acids and has the typical spacing for genes between the adjacent genes. Out of the 93 strains, 92 of these genes are full length, 1 frameshifted, no stop codons. Among the 1011 strains there are no stop codons and very few frameshift mutations. This open reading frame is not conserved in related species, though in *S. paradoxus* and *S. mikatae* is some sequence conservation but no open reading frame conservation. This gene does not appear to have a bacterial ortholog.

Additionally, *YGR227C-A* (*OTO1*) 57 amino acids Out of the 93 strains, 86 full length, 1 frameshifted, 6 frameshifted close to the 3 end, no stop codons. Among the 1011 strains, 28 had stop codon, and a similar ratio contain frameshifts. This open reading frame is not conserved in related species. A phenotype was found when overexpressing this putative small protein (Makanae et al. 2015) Another possible gene is *YDL204W-A* a putative gene of unknown function of 50 amino acids Out of the 93 strains, 37 are full length and 56 are disrupted by stop codon near 5’ end. This open reading frame not conserved in related species (Wacholder et al. 2023). Some transcripts span *YDL204W-A* and the downstream gene *YDL204W*. All of these 39 putative *S. cerevisiae* specific genes are small, and almost all are uncharacterized, and while we are not here proposing they are dubious, it is unclear if they are real protein coding genes.

An additional gene that on first inspection looks to be a gene found only in *S. cerevisiae* is *YMR106W-A* (Wacholder and Carvunis 2023, Wacholder et al. 2023). On closer inspection *YMR106W-A* appears to be a gene found in two species, *S. cerevisiae* and *S. paradoxus*, sometimes as a diverged tandem gene pair in both species, such as in *S. cerevisiae* strains YJM1400 and YJM1479. One form of the paralogs is 87 amino acids the other form 143 amino acids. Of the 93 strains, the shorter form is full length in 35 strains, disrupted by frameshifts in 58 strains, four different frameshifts, no in-frame stop codons. The longer form is only intact in the 2/93 strains mentioned, with partial copies, all of the amino terminal end, in the remaining 91 strains. In S288c the remnant of the amino terminal end of the longer gene runs from 482129 to 482209 on the Watson strand (Figure S30). Sixteen of the 1011 strains have full length, no internal stop codon copies of the longer form. As it appears that both the longer form and the shorter form are possibly real genes, and the shorter form is called *YMR106W-A*, we are calling the longer form *YMR106W-B*. The two genes are also intact only in a subset of *S. paradoxus* strains. While these both appear to be uncharacterized genes, it is also possible they should be considered dubious.

While there appear to be either very few or no *S. cerevisiae* core genes found only in *S. cerevisiae*, an interesting example of a real non-core gene found only in *S. cerevisiae* appears to be *YSC0033 STA1*/ *YSC0034 STA2*/*YSC0035 STA3*. This gene is found in only one of the 94 strains, YJM193, in the left subtelomeric region of chromosome IV. A full-length sequence is found in only one of the 1011 strains on contig AFP_1-7540. There may be additional copies of this gene in these 1011 strains, as the sequence contains repeats and assembly is problematic. This gene does not seem to be found in other species, and these two sequences and the additional protein sequences in GenBank are highly conserved. What is unusual about this gene is that it appears to be a chimeric gene, a fusion of *YIL099W SGA1* and *YIR019C MUC1*, as has been previously reported (Yamashita et al. 1987, Lo and Dranginis 1996, Krogerus et al. 2019). The high degree of sequence conservation suggests the chimera arose in this species.

### Overlapping protein coding regions

Among these proposed dubious genes, many overlap with other genes (Table S9). Seven overlap with Ty elements, seventeen overlap with RNA coding genes, mostly tRNA genes, and thirty-four overlap with protein coding genes. Among the genes here described as dubious is *YLR154W-C* (*TAR1*) (Coelho et al. 2002). While the published evidence suggests there is transcript and possibly even protein produced, not surprising considering there are as many as 100 copies of this region in the genome, the lack of conservation of the open reading frame (Figure S20 and Figure 1C of the Coelho et all paper), and the lack of a known function suggests that this open reading frame is unlikely to be a gene. A gene of unknown function overlapping with another gene is not on its own conclusive of an annotation error, but when combined with a general lack of evidence supporting these annotations, it contributes to the conclusion that these genes should be considered dubious. An additional two pairs of real genes currently annotated as overlapping (*YDR512C*, *YDR513W*; *YCR038C*, *YCR039C*) appear to overlap only due to errors in identification of the start codon. Until recently there was another pair of annotated overlapping genes, *YJR012C* and *YJR013W* but this has been resolved by mapping and correction of the start codon of *YJR012C* (Sadhu et al. 2018, Engel et al. 2022). For *YDR512C*, it appears that an internal start codon is the conserved start codon (Figure S21). The case of *YCR038C* is more complicated, and while it appears the current start codon is not correct, it will need to be investigated further as to which is the actual start codon.

Hidden among the numerous dubious cases there are real overlapping protein coding regions in *S. cerevisiae*, with seven overlapping gene pairs identified (Table S10). In these cases of well-characterized overlapping genes, as with the 26 pairs of overlapping genes in *A. gossypii* (Dietrich et al. 2013), they are all pairs of convergent protein coding genes. Two interesting cases are shown in Figure S22 where the overlapping region is conserved among related species. Interestingly, the relative frame of the overlap appears to be more conserved than the actual sequence of the overlap, particularly in the case of *VPS38* and *DCR2*.

While it appears that the only overlapping protein coding genes in *S. cerevisiae* are convergently transcribed, there is a particularly interesting case of *YNL155C-A* (Wacholder and Carvunis 2023, Wacholder et al. 2023) and *YNL156C* which both appear to be real, are highly conserved among the *Saccharomyces sensu stricto* species, and are on the same strand in two different reading frames and overlap. *YNL155C-A* is upstream of *YNL156C* and overlaps by 58 nucleotides, including the stop codon, in all *Saccharomyces sensu stricto* species, and in all cases the overlap is in the same relative reading frames. There are several interesting aspects of this case. There are no start codons in any of the three frames between the annotated start codons of *YNL155C-A* and *YNL156C*. As seen in Figure S29A, when examining long read mRNA and cDNA sequences generated using Nanopore and Pacific Biosciences sequencing the transcripts start both upstream and downstream of the *YNL155C-A* start codon and nearly all run past the stop codon of *YNL156C*. This is similar to but somewhat different than the case of *HTS1*, where there are two start codons both in-frame and where transcripts start either upstream or downstream of the first start codon (Natsoulis et al. 1986). Sequence alignment of the eight *Saccharomyces sensu stricto* species is shown if Figure S29B. Downstream of the putative −1 frameshift site between *YNL155C-A* and *YNL156C* there is possible RNA structure, similar to that in the putative −1 ribosomal frameshifting in *A. gossypii, A. aceri* and *H. sinecauda AFR597W* and in retroviruses(Farabaugh 1996, Farabaugh 1996, Brierley and Dos Ramos 2006, Brierley et al. 2010). While retrotransposons and some nuclear protein coding genes in *S. cerevisiae* have +1 ribosomal frameshifting, retroviruses generally have −1 ribosomal frameshifting.

While this will require experimental validation, it appears that *YNL155C-A* and *YNL156C* are a single gene and there is a longer and shorter form based on the transcription start site and −1 ribosomal frameshifting. The case of *A. gossypii AFR597W* is slightly different in that there is no in-frame start codon, the only start codon comes from the −1 frameshifting. For both *S. cerevisiae YNL156C* and *A. gossypii AFR597W* the −1 ribosomal frameshifting appears to be conserved in closely related species, but not in the orthologs in more distantly related species. −1 frameshifting is known to occur in *S. cerevisiae* in RNA encoded protein coding genes encoded on L-A and L-BC double stranded RNA (Dinman et al. 1991, Park et al. 1996) though has not previously been reported for DNA encoded protein coding genes.

Actually, there is one report but the paper was retracted as the results could not be replicated (Bekaert et al. 2005, Bekaert et al. 2006). The interesting thing about this retraction is one of the cases they proposed as a −1 ribosomal frameshift appears to be a −1 frameshift mutation, a null allele, in the non-core gene *YFL056C* (*AAD6*)/*YFL057C* (*AAD16*). There is an intact single non-core subtelomeric allele of this gene in 16 of the 93 genes with no frameshift mutations or internal stop codons, whereas in S288c and 15 of the 93 strains there is a single −1 frameshift mutation that appears to be a null mutation. In YJM1444 and YJM1419 there is a different frameshift mutation a +1 frameshift mutation at a different position in the gene. This gene, a putative aryl-alcohol dehydrogenase, has numerous paralogs. Full length intact copies of at least one of these paralogs is found in all 94 strains. Two paralogs are found at the end of the central region of the genome at two locations, Chromosome XIV left end *YNL331C* (*AAD14*) and Chromosome IV left end *YDL243C* (*AAD4*). There are also non-core copies in 15 sub-telomeric regions on Chromosomes IIL IIIR IVR VIL VIR VIIL VIIIL IXL XL XR IXL XIIR XVL XVR XVIL. Among all of these loci there are a mixture of intact copies and potentially null alleles, both stop codons and INDELs. While examination of any one strain could suggest the possibility of −1 or +1 frameshifting, examining of these paralogs from a large number of strains suggests numerous sub-telomeric defective copies of these paralogs is a more likely explanation. A case of ribosomal frameshifting would more likely be a conserved frameshift across many strains and multiple species.

In general, there are very few cases of eukaryotic protein coding genes, other that viruses and retrotransposons, annotated as using any type of ribosomal frameshifting. Whether this is due to most eukaryotes not using these translational mechanisms, or lack of careful annotation is not clear. While *YNL155C-A* and *YNL156C* appear to be a case of −1 frameshifting, it is also possible it is a case of a uORF overlapping with the main open reading frame, or a case of two independent genes that overlap.

### Transcripts spanning more than one gene

For the seven pairs of overlapping protein coding genes, an interesting phenomenon is observed. In some cases there are transcripts that span both genes in both directions. This is only clear from long read RNA-seq and cDNA-seq sequences generated using Nanopore and Pacific Biosciences methods. Nanopore sequences are direct mRNA sequencing, thus the sequences are strand specific. With short read RNA-seq reads using Illumina technology it is difficult to unambiguously identify these long transcripts.

For example, *YGR074W* (*SMD1*) and *YGR075C* (*PRP38*) are an overlapping convergent pair, and some nanopore sequenced RNA transcripts extend across both genes on the positive strand for *YGR075C* and the minus strand for *YGR084W*, for example from the NCBI SRA archive (Sayers et al. 2022): SRR32518276.739916, SRR32518275.971371, and SRR32518274.476520 (unpublished). Many sequences on the positive strand for *YGR074W* extend approximately across 300 nucleotides across *YGR075C* before terminating, examples: SRR32518276.867872, SRR32518277.1073419, SRR32518277.1552775 (unpublished).

Another example, *YNL246W* (*VPS75*) and *YNL245C* (*CWC25*) are an overlapping convergent pair, and some nanopore sequenced RNA transcripts extend across both genes on the positive strand for *YNL245C* and the minus strand for *YNL246W*, for example from the NCBI SRA archive (Sayers et al. 2022): SRR11550239.1065933 (Jenjaroenpun et al. 2021), SRR32518273.1620811, SRR32518274.202216, and SRR32518275.633887 (unpublished). Spanning both genes on the positive strand for *YNL246W* and the minus strand for *YNL245C*: SRR11550239.1065905, SRR11550239.1065906, SRR11550239.1065907, and SRR11550239.1065908 (Jenjaroenpun et al. 2021).

These multigene transcripts are also found for pairs of genes in the same orientation, such as transcripts that span *YHR189W* (*PTH1*) and *YHR190W* (*ERG11*) SRR32518277.1018672, SRR32518277.707160, and SRR32518284.3372292 (unpublished). Interestingly, *YHR189W* (*PTH1*) and *YHR190W* (*ERG11*) are adjacent genes in *A. gossypii*, *AFR445C* and *AFF444C*.

Another example is *YNL195C* and *YNL194C*, where transcripts spanning both genes include SRR11550239.1072387 (Jenjaroenpun et al. 2021) and SRR11589288.13457595, SRR11589288.15301624, and SRR11589288.15301627 (unpublished) sequenced using PacBio cDNA sequencing. *YNL195C* and *YNL194C* have adjacent orthologs in many pre-WGD species. A much larger number of transcripts span these two genes in dataset SRR11550239, sequencing RNA from heat shocked cells using nanopore RNA-seq (Jenjaroenpun et al. 2021). Longer transcripts from heat shocked cells has previously been reported (Yoon and Brem 2010).

What is unclear with these multigenic transcripts in *S. cerevisiae* is whether both proteins can be expressed off these transcripts, or just one or neither open reading frame. Thus it is unclear if these are operons.

### Potential null alleles

A phenomenon that has long been known and is quite widespread is that in *S. cerevisiae* there are genes with a stop codon or frameshift or disabling single nucleotide polymorphisms in the coding region. Well known cases of non-functional genes in S288c include *HO* and *HAP1* (Meiron et al. 1995, Gaisne et al. 1999). Another example is *YDL037C*/*YDL039C*, a single open reading frame in many strains and other *Saccharomyces* species, called *YSC0054 IMI1*, but is disrupted by a single stop codon in S288c and some other strains (Kowalec et al. 2015). While the *HO* gene is a null allele in strain S288c due to non-synonymous base substitutions (Meiron et al. 1995), many potential null alleles are the results of frameshifts or stop codons or transposon insertions. Some 444 core genes have a stop codon or frameshift (insertion or deletion) or transposon insertions, in one or more of the 94 strains, shown in Table S11. All 94 strains have some genes with potential null alleles, with the number of genes with potential null alleles per strain ranging from four to forty-four, with a median of 21. S288c is on the low end with only 16 genes with potential null alleles. In 95% percent of these potential null alleles there is only a single stop codon or frameshift. These genes with a single stop codon or frameshift are likely functional in the sense that a single point mutation could revert them. In the definition used here, a gene is still a core gene if it is present at the same location in 95% of the strains, even if in some cases it is a potential null allele. Furthermore, even when there is a frameshift or stop codon or transposon disruption in a protein-coding gene, this should be considered a single gene and given a single name. When the genome of S288c was first annotated, some genes were mistakenly given two names due to a frameshift or stop codon or transposon insertion because at that time in was not known that those fragments comprised a single gene. This is now misleading. These genes should be given a single name, with the second name relegated to an alias, and these genes identified as a potential null allele. Exactly how to define these genes with potential null alleles is complicated as some cases involve frameshifts or stop codons very close to the carboxy-terminal end of the protein, and thus may be fully functional, just with an alternative carboxy-terminal end.

An example of a gene with an elevated chance of a disabling mutation is *YLR046C* where a 9bp inverted micro-homology has led to a small inversion and introduction of a stop codon in two strains, YJM1592 and YJM1388, that theoretically can be reversed by a second inversion at the microhomology (Figure S23). An example of a gene with reduced likelihood of contracting a potential null allele is *HMO1*, where approximately half of the lysine residues are encoded by AAA, and half by AAG. The one run of seven consecutive lysines is encoded by (AAG)^7^, where a three base frameshift is possible, but a one base frameshift is not as likely as it would be if the seven consecutive lysines were encoded by (AAA)^7^. Overall, homopolymer runs are underrepresented in S288c protein coding regions in both essential and nonessential genes relative to non-coding regions (Figure S24).

Not all potential null alleles are stop codons/frameshifts/transposon insertions. An example of a potential null allele that is a non-synonymous substitution is the glycine to aspartic acid substitution at position 331 in uncharacterized gene *YDR111C* (*ALT2*) in strain S288c. In this WGD paralog *YLR089C* (*ALT1*) and in *ALT2* in all 93 strains there is a glycine at that position. In addition, the ortholog in *A. gossypii*, *Candida albicans, Schizosaccharomyces pombe*, *Cryptococcus neoformans*, *Homo sapiens*, *Arabidopsis thaliana, and even Escherichia coli* all have the glycine as well (Figure S25). It has been shown that the S288c *ALT2* is a non-functional paralog of *ALT1*, an alanine transaminase (Penalosa-Ruiz et al. 2012). Multiple sequence alignment suggests that residue 331 is the only highly conserved position between plants, animals, fungi, and bacteria that is mutated in S288c *ALT2*. The glycine to aspartic acid change at position 331 thus is a potential null allele, though experimental validation is needed to identify if this is the actual change that makes this a null allele, or whether this gene has a different function.

### Pseudogenes

Alleles with a stop codon or a frameshift where the gene is at the same location in other strains where the gene is intact are not pseudogenes. At times these have been mistakenly referred to as pseudogenes but they are instead alleles. For example the conserved gene *SDC25*, a WGD paralog of *CDC25* is a disrupted allele, a potential null allele, or a gene remnant, but it is not a pseudogene as reported (Sanchez et al. 2019) as it is at the same location where *CDC25*/*SDC25* is in the pre-WGD strains. There are also pseudogenes in *S. cerevisiae*. For example, strain YJM1208 contains a 376 base pair fragment of the 25S rDNA gene located in the subtelomeric region distal to *YPL275W* (*FDH2*). This is not a functional rDNA array, but is a small piece integrated at a location where rDNA is not usually found; a 25S pseudogene. In some strains small pieces of mitochondrial DNA are inserted into the nuclear genome, such as the mitochondrial DNA fragment found in strain S288c on in the subtelomeric region of chromosome X at bases 6998-7227 and the same piece in the subtelomeric region of chromosome IX at 7015-7244. This same fragment is also found near telomeres in strains YJM1381, YJM1083, YJM326, and YJM1460. Another pseudogene derived from mitochondrial DNA is a 95 base pair fragment from the middle of the *COX3* gene found near the left end of chromosome VII in several strains including YJM689, YJM683, YJM682. An example of a chromosomal protein coding gene derived pseudogene is the 830 bases derived from the 5’ end of *YDL015C* (*TSC3*) on chromosome IV that is inserted in the central region of chromosome VII between *YGR177C* (*ATF2*) and *YGR178C* (*PBP1*) in strains YJM1381 and YJM1199. Another example of a chromosomal protein coding gene derived pseudogene is a copy of *YMR096W* inserted into the chromosome XIII right end subtelomeric region containing frameshifts and a stop codon. These are pseudogenes, following the usage of the term in higher eukaryotes. The initial identification of eukaryotic pseudogenes came from identification of globin genes. In a review by Proudfoot(Proudfoot 1980) it is reported that “Studies from a variety of mammalian species reveal that in each case more gene sequences are present in the genome than necessary to encode the known globin polypeptides… the term ‘pseudogene’ – defined as a region of DNA that displays significant homology to a functional gene but has mutations that prevent its expression – has been coined for these aberrant genes”. To be a pseudogene a DNA segment is at a location distinct from the normal locus of the gene. In *S. cerevisiae* it is important to distinguish between a pseudogene and a gene that in some strains is a potential null allele. Many but not all pseudogenes are found in the subtelomeric regions.

### Start Codon Errors

While identifying the set of core genes in *S. cerevisiae*, it was clear that there are some errors in the current annotation, in addition to those listed above. There are 22 genes where it appears that the wrong start codon is annotated (Figure S21) as well as genes currently annotated as two genes (Table S12). This is based on sequence conservation among the 94 strains as well as sequence conservation with other species. These misannotated genes, as with the dubious genes, provide a somewhat inaccurate picture of the set of genes in *S. cerevisiae*. There are also genes where it’s unclear as to which start codon is the real start codon, or whether more than one start codon is used. It has long been known that there are real protein coding genes in *S. cerevisiae* where more than one start codon is used such as *HTS1* (Natsoulis et al. 1986).

### tRNA Genes

The set of tRNA genes is highly conserved among these strains. S288c has 299 nuclear encoded tRNA genes with 41 anticodons among the nuclear encoded tRNAs; the other 93 genes have very similar numbers. All tRNA genes are in the central region of the chromosomes, except for several non-core copies. This high level of conservation of the tRNA genes was not obvious in the original assembly of the 93 genomes, as Ty elements often insert adjacent to tRNA genes, so numerous assembly errors were associated with tRNA genes. The tRNA gene content is summarized in Table S13. Two serine tRNA’s, *YNCD0024C tX(XXX)D* and *YNCL0039W tX(XXX)L*, are currently misannotated in S288c due to a sequence error in each gene that likely arose from an issue with the ABI dye-primer sequencing originally used to sequence the genome that resulted in G/C compression errors (Yamakawa et al. 1996) (Figure S26).

### KHS1 and KHR1

An interesting example of how examining multiple genome sequences can resolve uncertainties in the annotation of the S288c genome is the case of *YSC0044* (*KHS1*) and *YSC0002* (*KHR1*) (Goto et al. 1990, Goto et al. 1991). In the current S288c annotation in SGD, these two genes are annotated as not in the reference strain. For *KHR1*, this gene is a non-core gene found in 66 of the 93 strains, and always at the same position. It is located on the complementary strand, between *YIL029C* and *tK(CUU)I* in the central region of chromosome IX. The sequence is somewhat variable, so the predicted protein in some strains is more similar to that in P22313.1 (Goto et al. 1990) and sometimes more similar to that in KZV10588.1 (McIlwain et al. 2016). *KHS1* is a more complicated issue. The gene is a core gene *YER187W* in the central region of chromosome V currently annotated as an uncharacterized gene, found in S288c and in 92/93 strains from Strope et. al. However the gene in S288c and many other strains contains a stop codon and is apparently non-functional. In addition, it appears that the original sequence of *KHS1* P39690.1 contains multiple sequence errors, mostly insertions, as well as an inverted DNA fragment connected at HindIII sites. While nearly all strains from the 1011 genome data set contain *KHS1*, in no case does the sequence closely match P39690.1, supporting the notion that the original sequence contained multiple errors, and that *YER187W* is *KHS1*as previously reported by Frank and Wolfe(Frank and Wolfe 2009).

An examination of the core genes in *S. cerevisiae* shows that out of 5885, 5375 have orthologs in *A. gossypii*, and an additional 392 genes have orthologs in *K. lactis*, or other pre-duplication species. More than 98% of core genes in *S. cerevisiae* are of vertical inheritance as they were in the ancestral hemiascomycete pre-WGD genome. For the 118 core genes in S288c that do not appear to have a pre-WGD ortholog, a few appear to have been acquired by horizontal gene transfer since the WGD (Hall et al. 2005, Hall and Dietrich 2007, Tapia et al. 2023), though most are of unclear origin.

### Genes of unknown function

Following the above description of the set of core genes in *S. cerevisiae*, the following question arises: how many core *S. cerevisiae* genes are of unknown function, those listed as “uncharacterized” in SGD? The current number is 364, of which 291 have orthologs in *A. gossypii* or other pre-WGD species suggesting that most of these 364 genes of unknown function are real genes. This number is decreasing as researchers identify function of previously uncharacterized genes. Some recent examples being the identification of the function of *YNR062C* and *YNR063W* as involved in resistance to pulcherrimin (Krause et al. 2018), *YBR063C* being identified as a nuclear membrane tethering protein (Eisenberg-Bord et al. 2021), *YDL180W* being identified as having a role in TORC1 signaling and localization (Wallace et al. 2022), *YGL036W* required for mRNA m6A methylation (Park et al. 2023), and YIL156W-B being identified as a mitochondrial fission factor (Fukuda et al. 2023). Nearly 94% of the core genes of *S. cerevisiae* are of known function. Among the genes of unknown function, 5 appear to have potential null alleles in the *S. cerevisiae* reference strain S288c. The community of *S. cerevisiae* researchers is thus not that far from the point of being able to say that “we know something about the function of every core gene in *S. cerevisiae*”. *S. cerevisiae* continues to be the standard for model organisms; In the future the standard for model organisms will be to know with some accuracy the set of core genes in the organism, and something about the function of every core gene. *S. cerevisiae* will likely be the first eukaryote to meet this threshold.

The total number of genes in the haploid genome of an *S. cerevisiae* strain is thus approximately 6000 genes, 5850 to 5855 core genes, 5 to 50 central, non-core genes including the 2µ and RNA virus encoded genes, Ty element *GAG* and *POL* genes, and 50-300 subtelomeric genes. These numbers are quite different than those reported in Peter et al (Peter et al. 2018), where they report the genome in this species being composed of some 4990 core genes, and a total of 7222 genes. Their numbers, while quite accurate, give the sense that a haploid yeast strain selected from a wild isolate will likely contain the 4990 core genes, and several hundred, perhaps a thousand or more variable genes. Our analysis suggests genomes in this species have a more conserved gene set, and that a strain will likely contain approximately 5885 core genes, plus 50-300 variable genes. This more constrained view of the gene set suggests that among strains much of the phenotypic variation is due to variation within the core set of genes with a contribution from the fewer variable genes, as opposed to a smaller set of core genes and a large set of variable genes, as seen in some bacteria such as *Escherichia coli* (Hayashi et al. 2001).

## Discussion

When the genome of *S. cerevisiae* was first sequenced in the 1990’s sequencing a eukaryotic genome was an expensive and labor-intensive process. Thus for *S. cerevisiae* as well as for *Caenorhabditis elegans*, *Drosophila melanogaster*, human, and other species there was the concept of a single reference genome sequence. That is, an *S. cerevisiae* gene came to mean the gene from the reference strain. With the technological advances in the nearly three decades since the first eukaryotic genome sequence, it is now possible to ask the question about a specific gene in a species, or about the set of genes for a species. This paper is in part an investigation into how the genome of *S. cerevisiae* is the same as and differs from the genome of *S. cerevisiae* strain S288c.

The set of genes for a species, the extent of potential null alleles, the extent of introgression from sibling species, the distinction between core genes and strain-specific genes, pseudogenes, transposons, gene remnants, and other aspects can best be identified from many genomic sequences and additional “omics” data as well as published data on specific genes.

While there is still a long way to go in understanding the genome structure and organization of *S. cerevisiae*, comparing multiple genomes of a single species can identify many aspects of the genome not clear from analysis of a single strain of this species. It should be noted that the term “core genes” as used in this paper refers to the core genes in this species. This is not the same as the core genes of the *Saccharomyces sensu stricto*, or the core genes of the hemiascomycetes. These are somewhat more complicated concepts.

It is clear from this analysis that 94 strains are far too few to get an accurate view of the genome of *S. cerevisiae.* Many features were found in only a single strain, suggesting that there are many additional features that will be identified by analysis of a larger number of strains. Some initial analysis suggests there are potential null alleles, pseudogenes, introgressed genes, and additional strain-specific genes in the 1011 *S. cerevisiae* genomes (Peter et al. 2018) that are not present in the 94 strains discussed here.

This work focusses on the core genes, not the strain-specific genes. The reason for this is that these genome sequences are still not complete in the subtelomeric regions where many strain-specific genes are found, so a list of the strain-specific genes for the 93 genomes is not yet possible. Improvements in sequencing technology since the data used here was generated, particularly recent advances in long sequence reads, means that complete, highly accurate genome sequences can now be more easily generated.

Another feature of this analysis is that the 94 strains examined here are haploids, or in the case of the 93 strains it is more accurate to say they are switched and mated homozygous diploids by the action of *HO*. Wild strains are typically diploid and often have extensive heterozygosity. In this analysis potential null alleles are found in over 400 protein coding genes; it is unclear if these potential null alleles are typically found in the population in a heterozygous or homozygous state. Phased diploid sequencing will potentially answer this question, and other questions about the genomes of wild strains of *S. cerevisiae*.

In this paper we propose that the *S. cerevisiae* S288c genome contains 5521 core genes of known function, more than 98% of which have orthologs in pre-WGD species. An additional 364 core genes of unknown function, labeled as uncharacterized in SGD, are present for a total of 5885 core genes, a number amazingly close to the 5885 genes reported by André Goffeau (Goffeau et al. 1996). Eleven of the core genes are potential null alleles in S288c, disrupted by a frameshift, a stop codon, or a transposon. There are also 22 non-core genes in the central regions of the genome, plus additional non-core genes in the subtelomeric regions. In the current annotation of 681 dubious genes, we conclude 680 are dubious, but *YBR012C* is not dubious but is a segment of a disrupted gene, the other half currently annotated as *YBR013C*. Here it is proposed that another 227 currently annotated genes should be relegated to dubious as they appear unlikely to be real genes but are instead annotation errors. In the absence of solid experimental evidence identifying a function for these open reading frames, their annotation as genes is misleading. Furthermore, there are eight gene remnants in S288c, fragments of former genes that have not completely disappeared. With more careful analysis, more gene remnants may be found. Two serine-tRNA genes appear to have sequence errors.

This work uncovers that there are seven real cases of convergent protein coding genes that overlap, and exactly one intron with a GC 5’ splice site, the remaining introns having the typical eukaryotic GT 5-splice site, and that some *S. cerevisiae* strains carry a novel transposon Ty7. More interestingly, reanalysis of the *S. cerevisiae* S288c genome suggests that, for example, the gene sets of *Candida albicans* and *S. cerevisiae* are more similar than has been previously thought due to the disrupted genes, the gene remnants, and the 237 proposed dubious genes, and other issues with the annotation.

No new data was presented in this paper. The data presented here has been publicly available in the fastq files deposited in in the NCBI short read archive for the initial publication of the 93 genomes (Strope et al. 2015), along with other publicly available data. There is now a massive amount of fungal data available, but the challenge is it is a labor-intensive process to extract information from data. That is what we have attempted to do here.

## Data Availability

The 93 *S. cerevisiae* strains are available through the Fungal Genetics Stock Center. Additional information on the strains is available (Strope et al. 2015).

The Illumina sequence data and annotated chromosomes are available through GenBank at the accession numbers in table S19 of Strope et al (Strope et al. 2015).

## Acknowledgements

The authors wish to thank Dr. Jennifer Tenor for careful reading of the manuscript.

