## Supplemental tables and figures for "Core gene set of the species *Saccharomyces cerevisiae*"

Table S1 Dubious subtelomeric introns based on examination of RNA-seq data SRR488142.

| Gene | Strand | Start | End | Chromosome | status | 5' Splice site |
| --- | --- | --- | --- | --- | --- | --- |
| YBL111C | minus | 4117 | 4215 | chrII | Intron dubious | GC |
| YIL177C | minus | 4599 | 4986 | chrIX | Intron dubious | GT |
| YEL076C-A | minus | 4323 | 4601 | chrV | Intron dubious | GT |
| YGR296W (YFR1-3) | plus | 1084883 | 1085030 | chrVII | Intron dubious | GT |
| YHL050C | minus | 1898 | 2670 | chrVIII | Intron dubious | GC |
| YHR218W | plus | 558616 | 558714 | chrVIII | Intron dubious | GC |
| YJL225C | minus | 4582 | 4969 | chrX | Intron dubious | GT |
| YLL067C | minus | 3916 | 4014 | chrXII | Intron dubious | GC |
| YLL066C | minus | 9451 | 9549 | chrXII | Intron dubious | GC |
| YLR464W | plus | 1067085 | 1067363 | chrXII | Intron dubious | GT |
| YML133C | minus | 3792 | 3890 | chrXIII | Intron dubious | GC |
| YNL339C (YFR1-6) | minus | 5932 | 6079 | chrXIV | Intron dubious | GT |
| YPL283C (YFR1-7) | minus | 5841 | 5988 | chrXVI | Intron dubious | GT |
| YPR202W | plus | 943051 | 943198 | chrXVI | Intron dubious | GT |

Table S2 Verified genes in the core set of genes in *S. cerevisiae.* The five pairs of columns are verified genes with orthologs in *A. gossypii*, verified genes with orthologs, or in some cases paralogs in pre-whole genome duplication species, genes that appear to be conserved only in post-WGD species both genus *Saccharomyces* and outside of this genus, genes conserved only among species of the genus *Saccharomyces,* and those present only in *S. cerevisiae.* * Verified genes more appropriately considered uncharacterized.

| preWGD, in  *A. gossypii* | Common name | preWGD, not in  *A. gossypii* | Common name | Broadly Post WGD | Common  name | *Saccharomyces*  *only* | Common  name | *S. cerevisiae*  only | Common  name |
| --- | --- | --- | --- | --- | --- | --- | --- | --- | --- |
| Number genes:5153 |  | Number genes: 323 |  | Number genes: 15 |  | Number genes: 26 |  | Number genes: 3 |  |
| YAL001C | TFC3 | YAL005C | SSA1 | YAR020C | PAU7 | YBR296C-A | TYC1 | YDL204W-A | * |
| YAL002W | VPS8 | YAL060W | BDH1 | YBR126W-A | MEO1 | YCR018C | SRD1 | YGR227C-A | OTO1 |
| YAL003W | EFB1 | YAL062W | GDH3 | YEL049W | PAU2 | YDR281C | PHM6 | YNL040C-A | * |
| YAL007C | ERP2 | YAL063C | FLO9 | YFL020C | PAU5 | YEL057C | SDD1 |  |  |
| YAL008W | FUN14 | YAR027W_28W_31W_33W | UIP3_KTD1_PRM9_MST2 | YHR053C | CUP1-1 | YEL066W | HPA3 |  |  |
| YAL009W | SPO7 | YAR050W | FLO1 | YHR055C | CUP1-2 | YFL050C | ALR2 |  |  |
| YAL010C | MDM10 | YBL005W | PDR3 | YJL037W | IRC18 | YGL015C | BIL2 |  |  |
| YAL011W | SWC3 | YBL010C | LAA2 | YJL038C | LOH1 | YGR066C | GID10 |  |  |
| YAL012W | CYS3 | YBR008C | FLR1 | YJR150C | DAN1 | YHL043W | ECM34 |  |  |
| YAL013W | DEP1 | YBR018C | GAL7 | YLL025W | PAU17 | YIL059C | * |  |  |
| YAL014C | SYN8 | YBR019C | GAL10 | YLL060C | GTT2 | YIR013C | GAT4 |  |  |
| YAL015C | NTG1 | YBR020W | GAL1 | YLR037C | PAU23 | YJL118W | * |  |  |
| YAL016W | TPD3 | YBR022W | POA1 | YML058W-A | HUG1 | YJR108W | ABM1 |  |  |
| YAL017W | PSK1 | YBR036C | CSG2 | YOR009W | TIR4 | YJR120W | DMO1 |  |  |
| YAL018C | LDS1 | YBR052C | RFS1 | YOR284W | HUA2 | YJR151C | DAN4 |  |  |
| YAL019W | FUN30 | YBR054W | YRO2 |  |  | YLR013W | GAT3 |  |  |
| YAL020C | ATS1 | YBR071W |  |  |  | YLR042C | NFG1 |  |  |
| YAL021C | CCR4 | YBR076W | ECM8 |  |  | YLR082C | SRL2 |  |  |
| YAL022C | FUN26 | YBR092C_093C | PHO3_PHO5 |  |  | YMR017W | SPO20 |  |  |
| YAL023C | PMT2 | YBR104W | YMC2 |  |  | YMR018W | PEX9 |  |  |
| YAL024C | LTE1 | YBR107C | IML3 |  |  | YMR106W-A | * |  |  |
| YAL025C | MAK16 | YBR137W |  |  |  | YOR031W | CRS5 |  |  |
| YAL026C | DRS2 | YBR149W | ARA1 |  |  | YPL021W | ECM23 |  |  |
| YAL027W | SAW1 | YBR150C | TBS1 |  |  | YPL027W | SMA1 |  |  |
| YAL028W | FRT2 | YBR161W | CSH1 |  |  | L-BC capsid |  |  |  |
| YAL029C | MYO4 | YBR194W | AIM4 |  |  | L-BC polymerase |  |  |  |
| YAL030W | SNC1 | YBR203W | COS111 |  |  |  |  |  |  |
| YAL031C | GIP4 | YBR227C | MCX1 |  |  |  |  |  |  |
| YAL032C | PRP45 | YBR293W | VBA2 |  |  |  |  |  |  |
| YAL033W | POP5 | YBR294W | SUL1 |  |  |  |  |  |  |
| YAL034C | FUN19 | YBR295W | PCA1 |  |  |  |  |  |  |
| YAL034W-A | MTW1 | YBR297W | MAL33 |  |  |  |  |  |  |
| YAL035W | FUN12 | YBR296C | PHO89 |  |  |  |  |  |  |
| YAL036C | RBG1 | YCL026C-A | FRM2 |  |  |  |  |  |  |
| YAL038W | CDC19 | YCL026C-B | HBN1 |  |  |  |  |  |  |
| YAL039C | CYC3 | YCL056C | PEX34 |  |  |  |  |  |  |
| YAL040C | CLN3 | YCL066W | HMLALPHA1 |  |  |  |  |  |  |
| YAL041W | CDC24 | YCL067C | HMLALPHA2 |  |  |  |  |  |  |
| YAL042W | ERV46 | YCL069W | VBA3 |  |  |  |  |  |  |
| YAL043C | PTA1 | YCL073C | GEX1 |  |  |  |  |  |  |
| YAL044C | GCV3 | YCR004C | YCP4 |  |  |  |  |  |  |
| YAL044W-A | BOL1 | YCR019W | MAK32 |  |  |  |  |  |  |
| YAL046C | BOL3 | YCR020C | PET18 |  |  |  |  |  |  |
| YAL047C | SPC72 | YCR021C | HSP30 |  |  |  |  |  |  |
| YAL048C | GEM1 | YCR027C | RHB1 |  |  |  |  |  |  |
| YAL049C | AIM2 | YCR039C | MATALPHA2 |  |  |  |  |  |  |
| YAL051W | OAF1 | YCR040W | MATALPHA1 |  |  |  |  |  |  |
| YAL053W | FLC2 | YCR089W | FIG2 |  |  |  |  |  |  |
| YAL054C | ACS1 | YCR096C | HMRA2 |  |  |  |  |  |  |
| YAL055W | PEX22 | YDL024C | DIA3 |  |  |  |  |  |  |
| YAL056W | GPB2 | YDL037C_039C | IMI1 |  |  |  |  |  |  |
| YAL058W | CNE1 | YDL124W |  |  |  |  |  |  |  |
| YAL059W | ECM1 | YDL154W | MSH5 |  |  |  |  |  |  |
| YAR002C-A | ERP1 | YDL170W | UGA3 |  |  |  |  |  |  |
| YAR002W | NUP60 | YDL200C | MGT1 |  |  |  |  |  |  |
| YAR003W | SWD1 | YDL206W | YCX1 |  |  |  |  |  |  |
| YAR007C | RFA1 | YDL223C | HBT1 |  |  |  |  |  |  |
| YAR008W | SEN34 | YDL227C | HO |  |  |  |  |  |  |
| YAR014C | BUD14 | YDL238C | GUD1 |  |  |  |  |  |  |
| YAR015W | ADE1 | YDL243C | AAD4 |  |  |  |  |  |  |
| YAR018C | KIN3 | YDR009W | GAL3 |  |  |  |  |  |  |
| YAR019C | CDC15 | YDR014W-A | HED1 |  |  |  |  |  |  |
| YAR035W | YAT1 | YDR019C | GCV1 |  |  |  |  |  |  |
| YAR042W | SWH1 | YDR032C | PST2 |  |  |  |  |  |  |
| YBL001C | ECM15 | YDR033W | MRH1 |  |  |  |  |  |  |
| YBL002W | HTB2 | YDR051C | DET1 |  |  |  |  |  |  |
| YBL003C | HTA2 | YDR070C | FMP16 |  |  |  |  |  |  |
| YBL004W | UTP20 | YDR072C | IPT1 |  |  |  |  |  |  |
| YBL006C | LDB7 | YDR090C | ILT1 |  |  |  |  |  |  |
| YBL007C | SLA1 | YDR119W-A | COX26 |  |  |  |  |  |  |
| YBL008W | HIR1 | YDR131C |  |  |  |  |  |  |  |
| YBL009W | ALK2 | YDR171W | HSP42 |  |  |  |  |  |  |
| YBL011W | SCT1 | YDR242W | AMD2 |  |  |  |  |  |  |
| YBL013W | FMT1 | YDR287W | INM2 |  |  |  |  |  |  |
| YBL014C | RRN6 | YDR294C | DPL1 |  |  |  |  |  |  |
| YBL015W | ACH1 | YDR314C | RAD34 |  |  |  |  |  |  |
| YBL016W | FUS3 | YDR383C | NKP1 |  |  |  |  |  |  |
| YBL017C | PEP1 | YDR399W | HPT1 |  |  |  |  |  |  |
| YBL018C | POP8 | YDR461W | MFA1 |  |  |  |  |  |  |
| YBL019W | APN2 | YDR506C | GMC1 |  |  |  |  |  |  |
| YBL020W | RFT1 | YDR519W | FPR2 |  |  |  |  |  |  |
| YBL021C | HAP3 | YDR533C | HSP31 |  |  |  |  |  |  |
| YBL022C | PIM1 | YDR534C | FIT1 |  |  |  |  |  |  |
| YBL023C | MCM2 | YDR538W | PAD1 |  |  |  |  |  |  |
| YBL024W | NCL1 | YDR539W | FDC1 |  |  |  |  |  |  |
| YBL025W | RRN10 | YDR541C |  |  |  |  |  |  |  |
| YBL026W | LSM2 | YEL004W | YEA4 |  |  |  |  |  |  |
| YBL027W | RPL19B | YEL040W | UTR2 |  |  |  |  |  |  |
| YBL028C |  | YEL065W | SIT1 |  |  |  |  |  |  |
| YBL029C-A |  | YEL069C | HXT13 |  |  |  |  |  |  |
| YBL030C | PET9 | YEL070W | DSF1 |  |  |  |  |  |  |
| YBL031W | SHE1 | YER028C | MIG3 |  |  |  |  |  |  |
| YBL032W | HEK2 | YER053C | PIC2 |  |  |  |  |  |  |
| YBL033C | RIB1 | YER053C-A |  |  |  |  |  |  |  |
| YBL034C | STU1 | YER184C | TOG1 |  |  |  |  |  |  |
| YBL035C | POL12 | YER185W | PUG1 |  |  |  |  |  |  |
| YBL036C |  | YFL003C | MSH4 |  |  |  |  |  |  |
| YBL037W | APL3 | YFL011W | HXT10 |  |  |  |  |  |  |
| YBL038W | MRPL16 | YFL030W | AGX1 |  |  |  |  |  |  |
| YBL039C | URA7 | YFR020W | CSS2 |  |  |  |  |  |  |
| YBL040C | ERD2 | YFR055W | IRC7 |  |  |  |  |  |  |
| YBL041W | PRE7 | YGL013C | PDR1 |  |  |  |  |  |  |
| YBL042C | FUI1 | YGL028C | SCW11 |  |  |  |  |  |  |
| YBL043W | ECM13 | YGL039W |  |  |  |  |  |  |  |
| YBL045C | COR1 | YGL067W | NPY1 |  |  |  |  |  |  |
| YBL046W | PSY4 | YGL090W | LIF1 |  |  |  |  |  |  |
| YBL047C | EDE1 | YGL117W | ARO5 |  |  |  |  |  |  |
| YBL049W | MOH1 | YGL121C | GPG1 |  |  |  |  |  |  |
| YBL050W | SEC17 | YGL157W | ARI1 |  |  |  |  |  |  |
| YBL051C | PIN4 | YGL186C | TPN1 |  |  |  |  |  |  |
| YBL052C | SAS3 | YGL196W | DSD1 |  |  |  |  |  |  |
| YBL054W | TOD6 | YGL209W | MIG2 |  |  |  |  |  |  |
| YBL055C |  | YGL249W | ZIP2 |  |  |  |  |  |  |
| YBL056W | PTC3 | YGL251C | HFM1 |  |  |  |  |  |  |
| YBL057C | PTH2 | YGL254W | FZF1 |  |  |  |  |  |  |
| YBL058W | SHP1 | YGL256W | ADH4 |  |  |  |  |  |  |
| YBL059C-A | CMC2 | YGL257C | MNT2 |  |  |  |  |  |  |
| YBL060W | YEL1 | YGL258W | VEL1 |  |  |  |  |  |  |
| YBL061C | SKT5 | YGR044C | RME1 |  |  |  |  |  |  |
| YBL063W | KIP1 | YGR154C | GTO1 |  |  |  |  |  |  |
| YBL064C | PRX1 | YGR177C | ATF2 |  |  |  |  |  |  |
| YBL066C | SEF1 | YGR192C | TDH3 |  |  |  |  |  |  |
| YBL067C | UBP13 | YGR199W | PMT6 |  |  |  |  |  |  |
| YBL068W | PRS4 | YGR213C | RTA1 |  |  |  |  |  |  |
| YBL069W | AST1 | YGR224W | AZR1 |  |  |  |  |  |  |
| YBL071W-A | KTI11 | YGR234W | YHB1 |  |  |  |  |  |  |
| YBL072C | RPS8A | YGR266W |  |  |  |  |  |  |  |
| YBL074C | AAR2 | YGR286C | BIO2 |  |  |  |  |  |  |
| YBL075C | SSA3 | YGR287C | IMA1 |  |  |  |  |  |  |
| YBL076C | ILS1 | YHL038C | CBP2 |  |  |  |  |  |  |
| YBL078C | ATG8 | YHL040C | ARN1 |  |  |  |  |  |  |
| YBL079W | NUP170 | YHR029C | YHI9 |  |  |  |  |  |  |
| YBL080C | PET112 | YHR046C | INM1 |  |  |  |  |  |  |
| YBL082C | ALG3 | YHR087W | RTC3 |  |  |  |  |  |  |
| YBL084C | CDC27 | YHR113W | APE4 |  |  |  |  |  |  |
| YBL085W | BOI1 | YHR136C | SPL2 |  |  |  |  |  |  |
| YBL087C | RPL23A | YHR137W | ARO9 |  |  |  |  |  |  |
| YBL088C | TEL1 | YHR139C | SPS100 |  |  |  |  |  |  |
| YBL089W | AVT5 | YHR153C | SPO16 |  |  |  |  |  |  |
| YBL090W | MRP21 | YHR157W | REC104 |  |  |  |  |  |  |
| YBL091C | MAP2 | YHR177W | ROF1 |  |  |  |  |  |  |
| YBL091C-A | SCS22 | YHR195W | NVJ1 |  |  |  |  |  |  |
| YBL092W | RPL32 | YHR198C | AIM18 |  |  |  |  |  |  |
| YBL093C | ROX3 | YHR199C-A | NBL1 |  |  |  |  |  |  |
| YBL097W | BRN1 | YHR209W | CRG1 |  |  |  |  |  |  |
| YBL098W | BNA4 | YIL009W | FAA3 |  |  |  |  |  |  |
| YBL099W | ATP1 | YIL073C | SPO22 |  |  |  |  |  |  |
| YBL101C | ECM21 | YIL085C | KTR7 |  |  |  |  |  |  |
| YBL102W | SFT2 | YIL164C_165C | NIT1 |  |  |  |  |  |  |
| YBL103C | RTG3 | YIL169C | CSS1 |  |  |  |  |  |  |
| YBL104C | SEA4 | YIR018W | YAP5 |  |  |  |  |  |  |
| YBL105C | PKC1 | YIR030C | DCG1 |  |  |  |  |  |  |
| YBL106C | SRO77 | YIR038C | GTT1 |  |  |  |  |  |  |
| YBL107C | MIX23 | YIR039C | YPS6 |  |  |  |  |  |  |
| YBR001C | NTH2 | YJL052W | TDH1 |  |  |  |  |  |  |
| YBR002C | RER2 | YJL082W | IML2 |  |  |  |  |  |  |
| YBR003W | COQ1 | YJL093C | TOK1 |  |  |  |  |  |  |
| YBR004C | GPI18 | YJL094C | KHA1 |  |  |  |  |  |  |
| YBR005W | RCR1 | YJL126W | NIT2 |  |  |  |  |  |  |
| YBR006W | UGA2 | YJL144W | ROQ1 |  |  |  |  |  |  |
| YBR007C | DSF2 | YJL151C | SNA3 |  |  |  |  |  |  |
| YBR009C | HHF1 | YJL153C | INO1 |  |  |  |  |  |  |
| YBR010W | HHT1 | YJL199C | MBB1 |  |  |  |  |  |  |
| YBR011C | IPP1 | YJL205C | NCE101 |  |  |  |  |  |  |
| YBR014C | GRX7 | YJL207C | LAA1 |  |  |  |  |  |  |
| YBR015C | MNN2 | YJL209W | CBP1 |  |  |  |  |  |  |
| YBR016W |  | YJL213W |  |  |  |  |  |  |  |
| YBR017C | KAP104 | YJR009C | TDH2 |  |  |  |  |  |  |
| YBR021W | FUR4 | YJR056C |  |  |  |  |  |  |  |
| YBR023C | CHS3 | YJR100C | AIM25 |  |  |  |  |  |  |
| YBR024W | SCO2 | YJR107C-A |  |  |  |  |  |  |  |
| YBR025C | OLA1 | YJR152W | DAL5 |  |  |  |  |  |  |
| YBR026C | ETR1 | YKL005C | BYE1 |  |  |  |  |  |  |
| YBR028C | YPK3 | YKL023C-A | MIN9 |  |  |  |  |  |  |
| YBR029C | CDS1 | YKL026C | GPX1 |  |  |  |  |  |  |
| YBR030W | RKM3 | YKL071W | OSI1 |  |  |  |  |  |  |
| YBR031W | RPL4A | YKL084W | HOT13 |  |  |  |  |  |  |
| YBR034C | HMT1 | YKL094W | YJU3 |  |  |  |  |  |  |
| YBR035C | PDX3 | YKL100C | YPF1 |  |  |  |  |  |  |
| YBR037C | SCO1 | YKL107W |  |  |  |  |  |  |  |
| YBR038W | CHS2 | YKL208W | CBT1 |  |  |  |  |  |  |
| YBR039W | ATP3 | YKL216W | URA1 |  |  |  |  |  |  |
| YBR040W | FIG1 | YKL218C | SRY1 |  |  |  |  |  |  |
| YBR041W | FAT1 | YKL219W | COS9 |  |  |  |  |  |  |
| YBR042C | CST26 | YKL220C | FRE2 |  |  |  |  |  |  |
| YBR043C | QDR3 | YKL221W | MCH2 |  |  |  |  |  |  |
| YBR044C | TCM62 | YKR018C |  |  |  |  |  |  |  |
| YBR045C | GIP1 | YKR066C | CCP1 |  |  |  |  |  |  |
| YBR046C | ZTA1 | YKR076W | ECM4 |  |  |  |  |  |  |
| YBR048W | RPS11B | YKR087C | OMA1 |  |  |  |  |  |  |
| YBR049C | REB1 | YKR101W | SIR1 |  |  |  |  |  |  |
| YBR050C | REG2 | YLL015W | BPT1 |  |  |  |  |  |  |
| YBR055C | PRP6 | YLL028W | TPO1 |  |  |  |  |  |  |
| YBR056W | MRX18 | YLL051C | FRE6 |  |  |  |  |  |  |
| YBR057C | MUM2 | YLL056C |  |  |  |  |  |  |  |
| YBR058C | UBP14 | YLL058W | HSU1 |  |  |  |  |  |  |
| YBR058C-A | TSC3 | YLR004C | THI73 |  |  |  |  |  |  |
| YBR059C | AKL1 | YLR011W | LOT6 |  |  |  |  |  |  |
| YBR060C | ORC2 | YLR014C | PPR1 |  |  |  |  |  |  |
| YBR061C | TRM7 | YLR040C | AFB1 |  |  |  |  |  |  |
| YBR063C | CNM1 | YLR053C | NRS1 |  |  |  |  |  |  |
| YBR065C | ECM2 | YLR081W | GAL2 |  |  |  |  |  |  |
| YBR066C | NRG2 | YLR097C | HRT3 |  |  |  |  |  |  |
| YBR067C | TIP1 | YLR109W | AHP1 |  |  |  |  |  |  |
| YBR068C | BAP2 | YLR126C |  |  |  |  |  |  |  |
| YBR069C | TAT1 | YLR146C | SPE4 |  |  |  |  |  |  |
| YBR070C | ALG14 | YLR266C | PDR8 |  |  |  |  |  |  |
| YBR072W | HSP26 | YLR286C | CTS1 |  |  |  |  |  |  |
| YBR073W | RDH54 | YLR297W |  |  |  |  |  |  |  |
| YBR074W | PFF1 | YLR301W | HRI1 |  |  |  |  |  |  |
| YBR077C | SLM4 | YLR315W | NKP2 |  |  |  |  |  |  |
| YBR078W | ECM33 | YLR346C | CIS1 |  |  |  |  |  |  |
| YBR079C | RPG1 | YLR348C | DIC1 |  |  |  |  |  |  |
| YBR080C | SEC18 | YLR363C | NMD4 |  |  |  |  |  |  |
| YBR081C | SPT7 | YLR394W | CST9 |  |  |  |  |  |  |
| YBR082C | UBC4 | YLR411W | CTR3 |  |  |  |  |  |  |
| YBR083W | TEC1 | YLR414C | PUN1 |  |  |  |  |  |  |
| YBR084C-A | RPL19A | YML037C | LFT1 |  |  |  |  |  |  |
| YBR084W | MIS1 | YML051W | GAL80 |  |  |  |  |  |  |
| YBR085C-A |  | YML054C | CYB2 |  |  |  |  |  |  |
| YBR085W | AAC3 | YML068W | ITT1 |  |  |  |  |  |  |
| YBR086C | IST2 | YML107C | PML39 |  |  |  |  |  |  |
| YBR087W | RFC5 | YML113W | DAT1 |  |  |  |  |  |  |
| YBR088C | POL30 | YML116W | ATR1 |  |  |  |  |  |  |
| YBR089C-A | NHP6B | YMR006C | PLB2 |  |  |  |  |  |  |
| YBR091C | TIM12 | YMR008C | PLB1 |  |  |  |  |  |  |
| YBR094W | PBY1 | YMR019W | STB4 |  |  |  |  |  |  |
| YBR095C | RXT2 | YMR030W | RSF1 |  |  |  |  |  |  |
| YBR097W | VPS15 | YMR056C | AAC1 |  |  |  |  |  |  |
| YBR098W | MMS4 | YMR110C | HFD1 |  |  |  |  |  |  |
| YBR101C | FES1 | YMR152W | YIM1 |  |  |  |  |  |  |
| YBR102C | EXO84 | YMR156C | TPP1 |  |  |  |  |  |  |
| YBR103W | SIF2 | YMR175W | SIP18 |  |  |  |  |  |  |
| YBR105C | VID24 | YMR189W | GCV2 |  |  |  |  |  |  |
| YBR106W | SND3 | YMR191W | SPG5 |  |  |  |  |  |  |
| YBR108W | AIM3 | YMR250W | GAD1 |  |  |  |  |  |  |
| YBR109C | CMD1 | YMR251W | GTO3 |  |  |  |  |  |  |
| YBR110W | ALG1 | YMR279C | ATR2 |  |  |  |  |  |  |
| YBR111C | YSA1 | YMR316W | DIA1 |  |  |  |  |  |  |
| YBR111W-A | SUS1 | YMR318C | ADH6 |  |  |  |  |  |  |
| YBR112C | CYC8 | YMR319C | FET4 |  |  |  |  |  |  |
| YBR114W | RAD16 | YNL018C |  |  |  |  |  |  |  |
| YBR115C | LYS2 | YNL029C | KTR5 |  |  |  |  |  |  |
| YBR117C | TKL2 | YNL034W |  |  |  |  |  |  |  |
| YBR118W | TEF2 | YNL040W |  |  |  |  |  |  |  |
| YBR119W | MUD1 | YNL125C | ESBP6 |  |  |  |  |  |  |
| YBR120C | CBP6 | YNL141W | AAH1 |  |  |  |  |  |  |
| YBR121C | GRS1 | YNL160W | YGP1 |  |  |  |  |  |  |
| YBR122C | MRPL36 | YNL190W |  |  |  |  |  |  |  |
| YBR123C | TFC1 | YNL196C | SLZ1 |  |  |  |  |  |  |
| YBR125C | PTC4 | YNL208W |  |  |  |  |  |  |  |
| YBR126C | TPS1 | YNL237W | YTP1 |  |  |  |  |  |  |
| YBR127C | VMA2 | YNL243W | SLA2 |  |  |  |  |  |  |
| YBR128C | ATG14 | YNL286W | CUS2 |  |  |  |  |  |  |
| YBR129C | OPY1 | YNL327W | EGT2 |  |  |  |  |  |  |
| YBR130C | SHE3 | YNL331C | AAD14 |  |  |  |  |  |  |
| YBR131W | CCZ1 | YNL332W | THI12 |  |  |  |  |  |  |
| YBR132C | AGP2 | YNR004W | SWM2 |  |  |  |  |  |  |
| YBR133C | HSL7 | YNR044W | AGA1 |  |  |  |  |  |  |
| YBR135W | CKS1 | YNR056C | BIO5 |  |  |  |  |  |  |
| YBR136W | MEC1 | YNR057C | BIO4 |  |  |  |  |  |  |
| YBR139W | ATG42 | YNR058W | BIO3 |  |  |  |  |  |  |
| YBR140C | IRA1 | YNR059W | MNT4 |  |  |  |  |  |  |
| YBR141C | BMT2 | YNR061C |  |  |  |  |  |  |  |
| YBR142W | MAK5 | YNR062C | PUL3 |  |  |  |  |  |  |
| YBR143C | SUP45 | YNR063W | PUL4 |  |  |  |  |  |  |
| YBR145W | ADH5 | YNR064C |  |  |  |  |  |  |  |
| YBR146W | MRPS9 | YNR069C | BSC5 |  |  |  |  |  |  |
| YBR147W | RTC2 | YOL011W | PLB3 |  |  |  |  |  |  |
| YBR148W | YSW1 | YOL015W | IRC10 |  |  |  |  |  |  |
| YBR151W | APD1 | YOL025W | LAG2 |  |  |  |  |  |  |
| YBR152W | SPP381 | YOL028C | YAP7 |  |  |  |  |  |  |
| YBR153W | RIB7 | YOL089C | HAL9 |  |  |  |  |  |  |
| YBR154C | RPB5 | YOL151W | GRE2 |  |  |  |  |  |  |
| YBR155W | CNS1 | YOL152W | FRE7 |  |  |  |  |  |  |
| YBR156C | SLI15 | YOL154W | ZPS1 |  |  |  |  |  |  |
| YBR157C | ICS2 | YOR010C | TIR2 |  |  |  |  |  |  |
| YBR158W | AMN1 | YOR049C | RSB1 |  |  |  |  |  |  |
| YBR159W | IFA38 | YOR071C | NRT1 |  |  |  |  |  |  |
| YBR160W | CDC28 | YOR084W | LPX1 |  |  |  |  |  |  |
| YBR162C | TOS1 | YOR087W | YVC1 |  |  |  |  |  |  |
| YBR162W-A | YSY6 | YOR114W | CIM1 |  |  |  |  |  |  |
| YBR163W | EXO5 | YOR131C |  |  |  |  |  |  |  |
| YBR164C | ARL1 | YOR137C | SIA1 |  |  |  |  |  |  |
| YBR165W | UBS1 | YOR138C | RUP1 |  |  |  |  |  |  |
| YBR166C | TYR1 | YOR162C | YRR1 |  |  |  |  |  |  |
| YBR167C | POP7 | YOR214C | SPR2 |  |  |  |  |  |  |
| YBR168W | PEX32 | YOR264W | DSE3 |  |  |  |  |  |  |
| YBR169C | SSE2 | YOR306C | MCH5 |  |  |  |  |  |  |
| YBR170C | NPL4 | YOR320C | GNT1 |  |  |  |  |  |  |
| YBR171W | SEC66 | YOR328W | PDR10 |  |  |  |  |  |  |
| YBR172C | SMY2 | YOR339C | UBC11 |  |  |  |  |  |  |
| YBR173C | UMP1 | YOR350C | MNE1 |  |  |  |  |  |  |
| YBR175W | SWD3 | YOR375C | GDH1 |  |  |  |  |  |  |
| YBR176W | ECM31 | YOR377W | ATF1 |  |  |  |  |  |  |
| YBR177C | EHT1 | YOR378W | AMF1 |  |  |  |  |  |  |
| YBR179C | FZO1 | YOR380W | RDR1 |  |  |  |  |  |  |
| YBR180W | DTR1 | YPL057C | SUR1 |  |  |  |  |  |  |
| YBR181C | RPS6B | YPL061W | ALD6 |  |  |  |  |  |  |
| YBR182C | SMP1 | YPL092W | SSU1 |  |  |  |  |  |  |
| YBR183W | YPC1 | YPL130W | SPO19 |  |  |  |  |  |  |
| YBR185C | MBA1 | YPL186C | UIP4 |  |  |  |  |  |  |
| YBR186W | PCH2 | YPL201C | YIG1 |  |  |  |  |  |  |
| YBR187W | GDT1 | YPL223C | GRE1 |  |  |  |  |  |  |
| YBR188C | NTC20 | YPR017C | DSS4 |  |  |  |  |  |  |
| YBR189W | RPS9B | YPR022C | SDD4 |  |  |  |  |  |  |
| YBR191W | RPL21A | YPR058W | YMC1 |  |  |  |  |  |  |
| YBR192W | RIM2 | YPR117W | HOB2 |  |  |  |  |  |  |
| YBR193C | MED8 | YPR121W | THI22 |  |  |  |  |  |  |
| YBR195C | MSI1 | YPR134W | MSS18 |  |  |  |  |  |  |
| YBR196C | PGI1 | YPR193C | HPA2 |  |  |  |  |  |  |
| YBR197C |  | YPR198W | SGE1 |  |  |  |  |  |  |
| YBR198C | TAF5 | YPR199C | ARR1 |  |  |  |  |  |  |
| YBR199W | KTR4 | YPR200C | ARR2 |  |  |  |  |  |  |
| YBR200W | BEM1 | YPR201W | ARR3 |  |  |  |  |  |  |
| YBR201W | DER1 | Q0060 | AI3 |  |  |  |  |  |  |
| YBR202W | MCM7 | Q0065 | AI4 |  |  |  |  |  |  |
| YBR204C | LDH1 | Q0070 | AI5_alpha |  |  |  |  |  |  |
| YBR205W | KTR3 | Q0110 | BI2 |  |  |  |  |  |  |
| YBR207W | FTH1 | Q0115 | BI3 |  |  |  |  |  |  |
| YBR208C | DUR12 | Q0120 | BI4 |  |  |  |  |  |  |
| YBR210W | ERV15 | YNCB0008W | GAL10-ncRNA |  |  |  |  |  |  |
| YBR211C | AME1 | YNCQ0017W | tR(ACG)Q2 |  |  |  |  |  |  |
| YBR212W | NGR1 |  |  |  |  |  |  |  |  |
| YBR213W | MET8 |  |  |  |  |  |  |  |  |
| YBR214W | SDS24 |  |  |  |  |  |  |  |  |
| YBR215W | HPC2 |  |  |  |  |  |  |  |  |
| YBR216C | YBP1 |  |  |  |  |  |  |  |  |
| YBR217W | ATG12 |  |  |  |  |  |  |  |  |
| YBR218C | PYC2 |  |  |  |  |  |  |  |  |
| YBR221C | PDB1 |  |  |  |  |  |  |  |  |
| YBR222C | PCS60 |  |  |  |  |  |  |  |  |
| YBR223C | TDP1 |  |  |  |  |  |  |  |  |
| YBR228W | SLX1 |  |  |  |  |  |  |  |  |
| YBR229C | ROT2 |  |  |  |  |  |  |  |  |
| YBR230C | OM14 |  |  |  |  |  |  |  |  |
| YBR230W-A | COQ21 |  |  |  |  |  |  |  |  |
| YBR231C | SWC5 |  |  |  |  |  |  |  |  |
| YBR233W | PBP2 |  |  |  |  |  |  |  |  |
| YBR233W-A | DAD3 |  |  |  |  |  |  |  |  |
| YBR234C | ARC40 |  |  |  |  |  |  |  |  |
| YBR235W | VHC1 |  |  |  |  |  |  |  |  |
| YBR236C | ABD1 |  |  |  |  |  |  |  |  |
| YBR237W | PRP5 |  |  |  |  |  |  |  |  |
| YBR238C |  |  |  |  |  |  |  |  |  |
| YBR239C | ERT1 |  |  |  |  |  |  |  |  |
| YBR240C | THI2 |  |  |  |  |  |  |  |  |
| YBR241C | VVS1 |  |  |  |  |  |  |  |  |
| YBR242W |  |  |  |  |  |  |  |  |  |
| YBR243C | ALG7 |  |  |  |  |  |  |  |  |
| YBR244W | GPX2 |  |  |  |  |  |  |  |  |
| YBR245C | ISW1 |  |  |  |  |  |  |  |  |
| YBR246W | RRT2 |  |  |  |  |  |  |  |  |
| YBR247C | ENP1 |  |  |  |  |  |  |  |  |
| YBR248C | HIS7 |  |  |  |  |  |  |  |  |
| YBR249C | ARO4 |  |  |  |  |  |  |  |  |
| YBR250W | SPO23 |  |  |  |  |  |  |  |  |
| YBR251W | MRPS5 |  |  |  |  |  |  |  |  |
| YBR252W | DUT1 |  |  |  |  |  |  |  |  |
| YBR253W | SRB6 |  |  |  |  |  |  |  |  |
| YBR254C | TRS20 |  |  |  |  |  |  |  |  |
| YBR255W | MTC4 |  |  |  |  |  |  |  |  |
| YBR256C | RIB5 |  |  |  |  |  |  |  |  |
| YBR257W | POP4 |  |  |  |  |  |  |  |  |
| YBR258C | SHG1 |  |  |  |  |  |  |  |  |
| YBR259W |  |  |  |  |  |  |  |  |  |
| YBR260C | RGD1 |  |  |  |  |  |  |  |  |
| YBR261C | TAE1 |  |  |  |  |  |  |  |  |
| YBR262C | MIC12 |  |  |  |  |  |  |  |  |
| YBR263W | SHM1 |  |  |  |  |  |  |  |  |
| YBR264C | YPT10 |  |  |  |  |  |  |  |  |
| YBR265W | TSC10 |  |  |  |  |  |  |  |  |
| YBR267W | REI1 |  |  |  |  |  |  |  |  |
| YBR268W | MRPL37 |  |  |  |  |  |  |  |  |
| YBR269C | SDH8 |  |  |  |  |  |  |  |  |
| YBR270C | BIT2 |  |  |  |  |  |  |  |  |
| YBR271W | EFM2 |  |  |  |  |  |  |  |  |
| YBR272C | HSM3 |  |  |  |  |  |  |  |  |
| YBR273C | UBX7 |  |  |  |  |  |  |  |  |
| YBR274W | CHK1 |  |  |  |  |  |  |  |  |
| YBR275C | RIF1 |  |  |  |  |  |  |  |  |
| YBR276C | PPS1 |  |  |  |  |  |  |  |  |
| YBR278W | DPB3 |  |  |  |  |  |  |  |  |
| YBR279W | PAF1 |  |  |  |  |  |  |  |  |
| YBR280C | SAF1 |  |  |  |  |  |  |  |  |
| YBR281C | DUG2 |  |  |  |  |  |  |  |  |
| YBR282W | MRPL27 |  |  |  |  |  |  |  |  |
| YBR283C | SSH1 |  |  |  |  |  |  |  |  |
| YBR285W | HAB1 |  |  |  |  |  |  |  |  |
| YBR286W | APE3 |  |  |  |  |  |  |  |  |
| YBR288C | APM3 |  |  |  |  |  |  |  |  |
| YBR289W | SNF5 |  |  |  |  |  |  |  |  |
| YBR290W | BSD2 |  |  |  |  |  |  |  |  |
| YBR291C | CTP1 |  |  |  |  |  |  |  |  |
| YCL001W | RER1 |  |  |  |  |  |  |  |  |
| YCL004W | PGS1 |  |  |  |  |  |  |  |  |
| YCL005W | LDB16 |  |  |  |  |  |  |  |  |
| YCL005W-A | VMA9 |  |  |  |  |  |  |  |  |
| YCL008C | STP22 |  |  |  |  |  |  |  |  |
| YCL009C | ILV6 |  |  |  |  |  |  |  |  |
| YCL010C | SGF29 |  |  |  |  |  |  |  |  |
| YCL011C | GBP2 |  |  |  |  |  |  |  |  |
| YCL012C |  |  |  |  |  |  |  |  |  |
| YCL014W | BUD3 |  |  |  |  |  |  |  |  |
| YCL016C | DCC1 |  |  |  |  |  |  |  |  |
| YCL017C | NFS1 |  |  |  |  |  |  |  |  |
| YCL018W | LEU2 |  |  |  |  |  |  |  |  |
| YCL024W | KCC4 |  |  |  |  |  |  |  |  |
| YCL025C | AGP1 |  |  |  |  |  |  |  |  |
| YCL027W | FUS1 |  |  |  |  |  |  |  |  |
| YCL028W | RNQ1 |  |  |  |  |  |  |  |  |
| YCL029C | BIK1 |  |  |  |  |  |  |  |  |
| YCL030C | HIS4 |  |  |  |  |  |  |  |  |
| YCL031C | RRP7 |  |  |  |  |  |  |  |  |
| YCL032W | STE50 |  |  |  |  |  |  |  |  |
| YCL033C | MXR2 |  |  |  |  |  |  |  |  |
| YCL034W | LSB5 |  |  |  |  |  |  |  |  |
| YCL035C | GRX1 |  |  |  |  |  |  |  |  |
| YCL036W | GFD2 |  |  |  |  |  |  |  |  |
| YCL037C | SRO9 |  |  |  |  |  |  |  |  |
| YCL038C | ATG22 |  |  |  |  |  |  |  |  |
| YCL039W | GID7 |  |  |  |  |  |  |  |  |
| YCL040W | GLK1 |  |  |  |  |  |  |  |  |
| YCL043C | PDI1 |  |  |  |  |  |  |  |  |
| YCL044C | MGR1 |  |  |  |  |  |  |  |  |
| YCL045C | EMC1 |  |  |  |  |  |  |  |  |
| YCL047C | POF1 |  |  |  |  |  |  |  |  |
| YCL048W | SPS22 |  |  |  |  |  |  |  |  |
| YCL050C | APA1 |  |  |  |  |  |  |  |  |
| YCL051W | LRE1 |  |  |  |  |  |  |  |  |
| YCL052C | PBN1 |  |  |  |  |  |  |  |  |
| YCL054W | SPB1 |  |  |  |  |  |  |  |  |
| YCL055W | KAR4 |  |  |  |  |  |  |  |  |
| YCL057C-A | MIC10 |  |  |  |  |  |  |  |  |
| YCL057W | PRD1 |  |  |  |  |  |  |  |  |
| YCL058W-A | ADF1 |  |  |  |  |  |  |  |  |
| YCL059C | KRR1 |  |  |  |  |  |  |  |  |
| YCL061C | MRC1 |  |  |  |  |  |  |  |  |
| YCL063W | VAC17 |  |  |  |  |  |  |  |  |
| YCL064C | CHA1 |  |  |  |  |  |  |  |  |
| YCR002C | CDC10 |  |  |  |  |  |  |  |  |
| YCR003W | MRPL32 |  |  |  |  |  |  |  |  |
| YCR005C | CIT2 |  |  |  |  |  |  |  |  |
| YCR008W | SAT4 |  |  |  |  |  |  |  |  |
| YCR009C | RVS161 |  |  |  |  |  |  |  |  |
| YCR010C | ADY2 |  |  |  |  |  |  |  |  |
| YCR011C | ADP1 |  |  |  |  |  |  |  |  |
| YCR012W | PGK1 |  |  |  |  |  |  |  |  |
| YCR014C | POL4 |  |  |  |  |  |  |  |  |
| YCR015C | CTO1 |  |  |  |  |  |  |  |  |
| YCR016W | RBP95 |  |  |  |  |  |  |  |  |
| YCR017C | CWH43 |  |  |  |  |  |  |  |  |
| YCR020C-A | MAK31 |  |  |  |  |  |  |  |  |
| YCR020W-B | HTL1 |  |  |  |  |  |  |  |  |
| YCR023C |  |  |  |  |  |  |  |  |  |
| YCR024C | SLM5 |  |  |  |  |  |  |  |  |
| YCR024C-A | PMP1 |  |  |  |  |  |  |  |  |
| YCR026C | NPP1 |  |  |  |  |  |  |  |  |
| YCR028C | FEN2 |  |  |  |  |  |  |  |  |
| YCR028C-A | RIM1 |  |  |  |  |  |  |  |  |
| YCR030C | SYP1 |  |  |  |  |  |  |  |  |
| YCR031C | RPS14A |  |  |  |  |  |  |  |  |
| YCR032W | BPH1 |  |  |  |  |  |  |  |  |
| YCR033W | SNT1 |  |  |  |  |  |  |  |  |
| YCR034W | ELO2 |  |  |  |  |  |  |  |  |
| YCR035C | RRP43 |  |  |  |  |  |  |  |  |
| YCR036W | RBK1 |  |  |  |  |  |  |  |  |
| YCR037C | PHO87 |  |  |  |  |  |  |  |  |
| YCR038C | BUD5 |  |  |  |  |  |  |  |  |
| YCR042C | TAF2 |  |  |  |  |  |  |  |  |
| YCR044C | PER1 |  |  |  |  |  |  |  |  |
| YCR045C | RRT12 |  |  |  |  |  |  |  |  |
| YCR046C | IMG1 |  |  |  |  |  |  |  |  |
| YCR047C | BUD23 |  |  |  |  |  |  |  |  |
| YCR048W | ARE1 |  |  |  |  |  |  |  |  |
| YCR052W | RSC6 |  |  |  |  |  |  |  |  |
| YCR053W | THR4 |  |  |  |  |  |  |  |  |
| YCR054C | CTR86 |  |  |  |  |  |  |  |  |
| YCR057C | PWP2 |  |  |  |  |  |  |  |  |
| YCR059C | YIH1 |  |  |  |  |  |  |  |  |
| YCR060W | TAH1 |  |  |  |  |  |  |  |  |
| YCR061W | TVS1 |  |  |  |  |  |  |  |  |
| YCR063W | BUD31 |  |  |  |  |  |  |  |  |
| YCR065W | HCM1 |  |  |  |  |  |  |  |  |
| YCR066W | RAD18 |  |  |  |  |  |  |  |  |
| YCR067C | SED4 |  |  |  |  |  |  |  |  |
| YCR068W | ATG15 |  |  |  |  |  |  |  |  |
| YCR069W | CPR4 |  |  |  |  |  |  |  |  |
| YCR071C | IMG2 |  |  |  |  |  |  |  |  |
| YCR072C | RSA4 |  |  |  |  |  |  |  |  |
| YCR073C | SSK22 |  |  |  |  |  |  |  |  |
| YCR073W-A | SOL2 |  |  |  |  |  |  |  |  |
| YCR075C | ERS1 |  |  |  |  |  |  |  |  |
| YCR075W-A | EGO2 |  |  |  |  |  |  |  |  |
| YCR076C | FUB1 |  |  |  |  |  |  |  |  |
| YCR077C | PAT1 |  |  |  |  |  |  |  |  |
| YCR079W | PTC6 |  |  |  |  |  |  |  |  |
| YCR081W | SRB8 |  |  |  |  |  |  |  |  |
| YCR082W | AHC2 |  |  |  |  |  |  |  |  |
| YCR083W | TRX3 |  |  |  |  |  |  |  |  |
| YCR084C | TUP1 |  |  |  |  |  |  |  |  |
| YCR086W | CSM1 |  |  |  |  |  |  |  |  |
| YCR087C-A |  |  |  |  |  |  |  |  |  |
| YCR088W | ABP1 |  |  |  |  |  |  |  |  |
| YCR091W | KIN82 |  |  |  |  |  |  |  |  |
| YCR092C | MSH3 |  |  |  |  |  |  |  |  |
| YCR093W | CDC39 |  |  |  |  |  |  |  |  |
| YCR094W | CDC50 |  |  |  |  |  |  |  |  |
| YCR097W | HMRA1 |  |  |  |  |  |  |  |  |
| YCR098C | GIT1 |  |  |  |  |  |  |  |  |
| YDL001W | RMD1 |  |  |  |  |  |  |  |  |
| YDL002C | NHP10 |  |  |  |  |  |  |  |  |
| YDL003W | MCD1 |  |  |  |  |  |  |  |  |
| YDL004W | ATP16 |  |  |  |  |  |  |  |  |
| YDL005C | MED2 |  |  |  |  |  |  |  |  |
| YDL006W | PTC1 |  |  |  |  |  |  |  |  |
| YDL007W | RPT2 |  |  |  |  |  |  |  |  |
| YDL008W | APC11 |  |  |  |  |  |  |  |  |
| YDL010W | GRX6 |  |  |  |  |  |  |  |  |
| YDL012C |  |  |  |  |  |  |  |  |  |
| YDL013W | SLX5 |  |  |  |  |  |  |  |  |
| YDL014W | NOP1 |  |  |  |  |  |  |  |  |
| YDL015C | TSC13 |  |  |  |  |  |  |  |  |
| YDL017W | CDC7 |  |  |  |  |  |  |  |  |
| YDL018C | ERP3 |  |  |  |  |  |  |  |  |
| YDL019C | OSH2 |  |  |  |  |  |  |  |  |
| YDL020C | RPN4 |  |  |  |  |  |  |  |  |
| YDL021W | GPM2 |  |  |  |  |  |  |  |  |
| YDL022W | GPD1 |  |  |  |  |  |  |  |  |
| YDL025C | RTK1 |  |  |  |  |  |  |  |  |
| YDL028C | MPS1 |  |  |  |  |  |  |  |  |
| YDL029W | ARP2 |  |  |  |  |  |  |  |  |
| YDL030W | PRP9 |  |  |  |  |  |  |  |  |
| YDL031W | DBP10 |  |  |  |  |  |  |  |  |
| YDL033C | SLM3 |  |  |  |  |  |  |  |  |
| YDL035C | GPR1 |  |  |  |  |  |  |  |  |
| YDL036C | PUS9 |  |  |  |  |  |  |  |  |
| YDL040C | NAT1 |  |  |  |  |  |  |  |  |
| YDL042C | SIR2 |  |  |  |  |  |  |  |  |
| YDL043C | PRP11 |  |  |  |  |  |  |  |  |
| YDL044C | MTF2 |  |  |  |  |  |  |  |  |
| YDL045C | FAD1 |  |  |  |  |  |  |  |  |
| YDL045W-A | MRP10 |  |  |  |  |  |  |  |  |
| YDL046W | NPC2 |  |  |  |  |  |  |  |  |
| YDL047W | SIT4 |  |  |  |  |  |  |  |  |
| YDL048C | STP4 |  |  |  |  |  |  |  |  |
| YDL049C | KNH1 |  |  |  |  |  |  |  |  |
| YDL051W | LHP1 |  |  |  |  |  |  |  |  |
| YDL052C | SLC1 |  |  |  |  |  |  |  |  |
| YDL053C | PBP4 |  |  |  |  |  |  |  |  |
| YDL054C | MCH1 |  |  |  |  |  |  |  |  |
| YDL055C | PSA1 |  |  |  |  |  |  |  |  |
| YDL056W | MBP1 |  |  |  |  |  |  |  |  |
| YDL058W | USO1 |  |  |  |  |  |  |  |  |
| YDL059C | RAD59 |  |  |  |  |  |  |  |  |
| YDL060W | TSR1 |  |  |  |  |  |  |  |  |
| YDL061C | RPS29B |  |  |  |  |  |  |  |  |
| YDL063C | SYO1 |  |  |  |  |  |  |  |  |
| YDL064W | UBC9 |  |  |  |  |  |  |  |  |
| YDL065C | PEX19 |  |  |  |  |  |  |  |  |
| YDL066W | IDP1 |  |  |  |  |  |  |  |  |
| YDL067C | COX9 |  |  |  |  |  |  |  |  |
| YDL069C | CBS1 |  |  |  |  |  |  |  |  |
| YDL070W | BDF2 |  |  |  |  |  |  |  |  |
| YDL072C | YET3 |  |  |  |  |  |  |  |  |
| YDL073W | AHK1 |  |  |  |  |  |  |  |  |
| YDL074C | BRE1 |  |  |  |  |  |  |  |  |
| YDL075W | RPL31A |  |  |  |  |  |  |  |  |
| YDL076C | RXT3 |  |  |  |  |  |  |  |  |
| YDL077C | VAM6 |  |  |  |  |  |  |  |  |
| YDL078C | MDH3 |  |  |  |  |  |  |  |  |
| YDL079C | MRK1 |  |  |  |  |  |  |  |  |
| YDL080C | THI3 |  |  |  |  |  |  |  |  |
| YDL081C | RPP1A |  |  |  |  |  |  |  |  |
| YDL082W | RPL13A |  |  |  |  |  |  |  |  |
| YDL083C | RPS16B |  |  |  |  |  |  |  |  |
| YDL084W | SUB2 |  |  |  |  |  |  |  |  |
| YDL085C-A |  |  |  |  |  |  |  |  |  |
| YDL085W | NDE2 |  |  |  |  |  |  |  |  |
| YDL087C | LUC7 |  |  |  |  |  |  |  |  |
| YDL088C | ASM4 |  |  |  |  |  |  |  |  |
| YDL089W | NUR1 |  |  |  |  |  |  |  |  |
| YDL090C | RAM1 |  |  |  |  |  |  |  |  |
| YDL091C | UBX3 |  |  |  |  |  |  |  |  |
| YDL092W | SRP14 |  |  |  |  |  |  |  |  |
| YDL093W | PMT5 |  |  |  |  |  |  |  |  |
| YDL095W | PMT1 |  |  |  |  |  |  |  |  |
| YDL097C | RPN6 |  |  |  |  |  |  |  |  |
| YDL098C | SNU23 |  |  |  |  |  |  |  |  |
| YDL099W | BUG1 |  |  |  |  |  |  |  |  |
| YDL100C | GET3 |  |  |  |  |  |  |  |  |
| YDL101C | DUN1 |  |  |  |  |  |  |  |  |
| YDL102W | POL3 |  |  |  |  |  |  |  |  |
| YDL103C | QRI1 |  |  |  |  |  |  |  |  |
| YDL104C | QRI7 |  |  |  |  |  |  |  |  |
| YDL105W | NSE4 |  |  |  |  |  |  |  |  |
| YDL106C | PHO2 |  |  |  |  |  |  |  |  |
| YDL107W | MSS2 |  |  |  |  |  |  |  |  |
| YDL108W | KIN28 |  |  |  |  |  |  |  |  |
| YDL110C | TMA17 |  |  |  |  |  |  |  |  |
| YDL111C | RRP42 |  |  |  |  |  |  |  |  |
| YDL112W | TRM3 |  |  |  |  |  |  |  |  |
| YDL113C | ATG20 |  |  |  |  |  |  |  |  |
| YDL115C | IWR1 |  |  |  |  |  |  |  |  |
| YDL116W | NUP84 |  |  |  |  |  |  |  |  |
| YDL117W | CYK3 |  |  |  |  |  |  |  |  |
| YDL119C | HEM25 |  |  |  |  |  |  |  |  |
| YDL120W | YFH1 |  |  |  |  |  |  |  |  |
| YDL121C | EXP1 |  |  |  |  |  |  |  |  |
| YDL122W | UBP1 |  |  |  |  |  |  |  |  |
| YDL123W | SNA4 |  |  |  |  |  |  |  |  |
| YDL125C | HNT1 |  |  |  |  |  |  |  |  |
| YDL126C | CDC48 |  |  |  |  |  |  |  |  |
| YDL127W | PCL2 |  |  |  |  |  |  |  |  |
| YDL128W | VCX1 |  |  |  |  |  |  |  |  |
| YDL129W |  |  |  |  |  |  |  |  |  |
| YDL130W | RPP1B |  |  |  |  |  |  |  |  |
| YDL130W-A | STF1 |  |  |  |  |  |  |  |  |
| YDL131W | LYS21 |  |  |  |  |  |  |  |  |
| YDL132W | CDC53 |  |  |  |  |  |  |  |  |
| YDL133C-A | RPL41B |  |  |  |  |  |  |  |  |
| YDL133W | SRF1 |  |  |  |  |  |  |  |  |
| YDL134C | PPH21 |  |  |  |  |  |  |  |  |
| YDL135C | RDI1 |  |  |  |  |  |  |  |  |
| YDL136W | RPL35B |  |  |  |  |  |  |  |  |
| YDL137W | ARF2 |  |  |  |  |  |  |  |  |
| YDL138W | RGT2 |  |  |  |  |  |  |  |  |
| YDL139C | SCM3 |  |  |  |  |  |  |  |  |
| YDL140C | RPO21 |  |  |  |  |  |  |  |  |
| YDL141W | BPL1 |  |  |  |  |  |  |  |  |
| YDL142C | CRD1 |  |  |  |  |  |  |  |  |
| YDL143W | CCT4 |  |  |  |  |  |  |  |  |
| YDL144C |  |  |  |  |  |  |  |  |  |
| YDL145C | COP1 |  |  |  |  |  |  |  |  |
| YDL146W | LDB17 |  |  |  |  |  |  |  |  |
| YDL147W | RPN5 |  |  |  |  |  |  |  |  |
| YDL148C | NOP14 |  |  |  |  |  |  |  |  |
| YDL149W | ATG9 |  |  |  |  |  |  |  |  |
| YDL150W | RPC53 |  |  |  |  |  |  |  |  |
| YDL153C | SAS10 |  |  |  |  |  |  |  |  |
| YDL155W | CLB3 |  |  |  |  |  |  |  |  |
| YDL156W | CMR1 |  |  |  |  |  |  |  |  |
| YDL159W | STE7 |  |  |  |  |  |  |  |  |
| YDL160C | DHH1 |  |  |  |  |  |  |  |  |
| YDL160C-A | MHF2 |  |  |  |  |  |  |  |  |
| YDL161W | ENT1 |  |  |  |  |  |  |  |  |
| YDL164C | CDC9 |  |  |  |  |  |  |  |  |
| YDL165W | CDC36 |  |  |  |  |  |  |  |  |
| YDL166C | FAP7 |  |  |  |  |  |  |  |  |
| YDL167C | NRP1 |  |  |  |  |  |  |  |  |
| YDL168W | SFA1 |  |  |  |  |  |  |  |  |
| YDL169C | UGX2 |  |  |  |  |  |  |  |  |
| YDL171C | GLT1 |  |  |  |  |  |  |  |  |
| YDL173W | PAR32 |  |  |  |  |  |  |  |  |
| YDL174C | DLD1 |  |  |  |  |  |  |  |  |
| YDL175C | AIR2 |  |  |  |  |  |  |  |  |
| YDL176W | GID12 |  |  |  |  |  |  |  |  |
| YDL178W | DLD2 |  |  |  |  |  |  |  |  |
| YDL179W | PCL9 |  |  |  |  |  |  |  |  |
| YDL180W | AIT1 |  |  |  |  |  |  |  |  |
| YDL181W | INH1 |  |  |  |  |  |  |  |  |
| YDL182W | LYS20 |  |  |  |  |  |  |  |  |
| YDL183C | MRX19 |  |  |  |  |  |  |  |  |
| YDL184C | RPL41A |  |  |  |  |  |  |  |  |
| YDL185W | VMA1 |  |  |  |  |  |  |  |  |
| YDL188C | PPH22 |  |  |  |  |  |  |  |  |
| YDL189W | RBS1 |  |  |  |  |  |  |  |  |
| YDL190C | UFD2 |  |  |  |  |  |  |  |  |
| YDL191W | RPL35A |  |  |  |  |  |  |  |  |
| YDL192W | ARF1 |  |  |  |  |  |  |  |  |
| YDL193W | NUS1 |  |  |  |  |  |  |  |  |
| YDL194W | SNF3 |  |  |  |  |  |  |  |  |
| YDL195W | SEC31 |  |  |  |  |  |  |  |  |
| YDL197C | ASF2 |  |  |  |  |  |  |  |  |
| YDL198C | GGC1 |  |  |  |  |  |  |  |  |
| YDL201W | TRM8 |  |  |  |  |  |  |  |  |
| YDL202W | MRPL11 |  |  |  |  |  |  |  |  |
| YDL203C | ACK1 |  |  |  |  |  |  |  |  |
| YDL204W | RTN2 |  |  |  |  |  |  |  |  |
| YDL205C | HEM3 |  |  |  |  |  |  |  |  |
| YDL207W | GLE1 |  |  |  |  |  |  |  |  |
| YDL208W | NHP2 |  |  |  |  |  |  |  |  |
| YDL209C | CWC2 |  |  |  |  |  |  |  |  |
| YDL210W | UGA4 |  |  |  |  |  |  |  |  |
| YDL212W | SHR3 |  |  |  |  |  |  |  |  |
| YDL213C | NOP6 |  |  |  |  |  |  |  |  |
| YDL214C | PRR2 |  |  |  |  |  |  |  |  |
| YDL215C | GDH2 |  |  |  |  |  |  |  |  |
| YDL216C | RRI1 |  |  |  |  |  |  |  |  |
| YDL217C | TIM22 |  |  |  |  |  |  |  |  |
| YDL219W | DTD1 |  |  |  |  |  |  |  |  |
| YDL220C | CDC13 |  |  |  |  |  |  |  |  |
| YDL222C | FMP45 |  |  |  |  |  |  |  |  |
| YDL224C | WHI4 |  |  |  |  |  |  |  |  |
| YDL225W | SHS1 |  |  |  |  |  |  |  |  |
| YDL226C | GCS1 |  |  |  |  |  |  |  |  |
| YDL229W | SSB1 |  |  |  |  |  |  |  |  |
| YDL230W | PTP1 |  |  |  |  |  |  |  |  |
| YDL231C | BRE4 |  |  |  |  |  |  |  |  |
| YDL232W | OST4 |  |  |  |  |  |  |  |  |
| YDL234C | GYP7 |  |  |  |  |  |  |  |  |
| YDL235C | YPD1 |  |  |  |  |  |  |  |  |
| YDL236W | PHO13 |  |  |  |  |  |  |  |  |
| YDL237W | AIM6 |  |  |  |  |  |  |  |  |
| YDL239C | ADY3 |  |  |  |  |  |  |  |  |
| YDL240W | LRG1 |  |  |  |  |  |  |  |  |
| YDR001C | NTH1 |  |  |  |  |  |  |  |  |
| YDR002W | YRB1 |  |  |  |  |  |  |  |  |
| YDR003W | RCR2 |  |  |  |  |  |  |  |  |
| YDR004W | RAD57 |  |  |  |  |  |  |  |  |
| YDR005C | MAF1 |  |  |  |  |  |  |  |  |
| YDR006C | SOK1 |  |  |  |  |  |  |  |  |
| YDR007W | TRP1 |  |  |  |  |  |  |  |  |
| YDR011W | SNQ2 |  |  |  |  |  |  |  |  |
| YDR012W | RPL4B |  |  |  |  |  |  |  |  |
| YDR013W | PSF1 |  |  |  |  |  |  |  |  |
| YDR014W | RAD61 |  |  |  |  |  |  |  |  |
| YDR016C | DAD1 |  |  |  |  |  |  |  |  |
| YDR017C | KCS1 |  |  |  |  |  |  |  |  |
| YDR018C | MLG1 |  |  |  |  |  |  |  |  |
| YDR021W | FAL1 |  |  |  |  |  |  |  |  |
| YDR022C | ATG31 |  |  |  |  |  |  |  |  |
| YDR023W | SES1 |  |  |  |  |  |  |  |  |
| YDR025W | RPS11A |  |  |  |  |  |  |  |  |
| YDR026C | NSI1 |  |  |  |  |  |  |  |  |
| YDR027C | VPS54 |  |  |  |  |  |  |  |  |
| YDR028C | REG1 |  |  |  |  |  |  |  |  |
| YDR030C | RAD28 |  |  |  |  |  |  |  |  |
| YDR031W | MIX14 |  |  |  |  |  |  |  |  |
| YDR034C | LYS14 |  |  |  |  |  |  |  |  |
| YDR035W | ARO3 |  |  |  |  |  |  |  |  |
| YDR036C | EHD3 |  |  |  |  |  |  |  |  |
| YDR037W | KRS1 |  |  |  |  |  |  |  |  |
| YDR038C | ENA5 |  |  |  |  |  |  |  |  |
| YDR039C | ENA2 |  |  |  |  |  |  |  |  |
| YDR040C | ENA1 |  |  |  |  |  |  |  |  |
| YDR041W | RSM10 |  |  |  |  |  |  |  |  |
| YDR043C | NRG1 |  |  |  |  |  |  |  |  |
| YDR044W | HEM13 |  |  |  |  |  |  |  |  |
| YDR045C | RPC11 |  |  |  |  |  |  |  |  |
| YDR046C | BAP3 |  |  |  |  |  |  |  |  |
| YDR047W | HEM12 |  |  |  |  |  |  |  |  |
| YDR049W | VMS1 |  |  |  |  |  |  |  |  |
| YDR050C | TPI1 |  |  |  |  |  |  |  |  |
| YDR052C | DBF4 |  |  |  |  |  |  |  |  |
| YDR054C | CDC34 |  |  |  |  |  |  |  |  |
| YDR055W | PST1 |  |  |  |  |  |  |  |  |
| YDR057W | YOS9 |  |  |  |  |  |  |  |  |
| YDR058C | TGL2 |  |  |  |  |  |  |  |  |
| YDR059C | UBC5 |  |  |  |  |  |  |  |  |
| YDR060W | MAK21 |  |  |  |  |  |  |  |  |
| YDR062W | LCB2 |  |  |  |  |  |  |  |  |
| YDR063W | AIM7 |  |  |  |  |  |  |  |  |
| YDR064W | RPS13 |  |  |  |  |  |  |  |  |
| YDR065W | RRG1 |  |  |  |  |  |  |  |  |
| YDR066C | RTR2 |  |  |  |  |  |  |  |  |
| YDR068W | DOS2 |  |  |  |  |  |  |  |  |
| YDR069C | DOA4 |  |  |  |  |  |  |  |  |
| YDR071C | PAA1 |  |  |  |  |  |  |  |  |
| YDR073W | SNF11 |  |  |  |  |  |  |  |  |
| YDR074W | TPS2 |  |  |  |  |  |  |  |  |
| YDR075W | PPH3 |  |  |  |  |  |  |  |  |
| YDR076W | RAD55 |  |  |  |  |  |  |  |  |
| YDR077W | SED1 |  |  |  |  |  |  |  |  |
| YDR078C | SHU2 |  |  |  |  |  |  |  |  |
| YDR079C-A | TFB5 |  |  |  |  |  |  |  |  |
| YDR079W | PET100 |  |  |  |  |  |  |  |  |
| YDR080W | VPS41 |  |  |  |  |  |  |  |  |
| YDR081C | PDC2 |  |  |  |  |  |  |  |  |
| YDR082W | STN1 |  |  |  |  |  |  |  |  |
| YDR083W | RRP8 |  |  |  |  |  |  |  |  |
| YDR084C | TVP23 |  |  |  |  |  |  |  |  |
| YDR085C | AFR1 |  |  |  |  |  |  |  |  |
| YDR086C | SSS1 |  |  |  |  |  |  |  |  |
| YDR087C | RRP1 |  |  |  |  |  |  |  |  |
| YDR088C | SLU7 |  |  |  |  |  |  |  |  |
| YDR089W | VTC5 |  |  |  |  |  |  |  |  |
| YDR091C | RLI1 |  |  |  |  |  |  |  |  |
| YDR092W | UBC13 |  |  |  |  |  |  |  |  |
| YDR093W | DNF2 |  |  |  |  |  |  |  |  |
| YDR096W | GIS1 |  |  |  |  |  |  |  |  |
| YDR097C | MSH6 |  |  |  |  |  |  |  |  |
| YDR098C | GRX3 |  |  |  |  |  |  |  |  |
| YDR099W | BMH2 |  |  |  |  |  |  |  |  |
| YDR100W | TVP15 |  |  |  |  |  |  |  |  |
| YDR101C | ARX1 |  |  |  |  |  |  |  |  |
| YDR103W | STE5 |  |  |  |  |  |  |  |  |
| YDR104C | SPO71 |  |  |  |  |  |  |  |  |
| YDR105C | TMS1 |  |  |  |  |  |  |  |  |
| YDR106W | ARP10 |  |  |  |  |  |  |  |  |
| YDR107C | TMN2 |  |  |  |  |  |  |  |  |
| YDR108W | TRS85 |  |  |  |  |  |  |  |  |
| YDR109C |  |  |  |  |  |  |  |  |  |
| YDR110W | FOB1 |  |  |  |  |  |  |  |  |
| YDR113C | PDS1 |  |  |  |  |  |  |  |  |
| YDR115W | MRX14 |  |  |  |  |  |  |  |  |
| YDR116C | MRPL1 |  |  |  |  |  |  |  |  |
| YDR117C | TMA64 |  |  |  |  |  |  |  |  |
| YDR118W | APC4 |  |  |  |  |  |  |  |  |
| YDR120C | TRM1 |  |  |  |  |  |  |  |  |
| YDR121W | DPB4 |  |  |  |  |  |  |  |  |
| YDR122W | KIN1 |  |  |  |  |  |  |  |  |
| YDR123C | INO2 |  |  |  |  |  |  |  |  |
| YDR125C | ECM18 |  |  |  |  |  |  |  |  |
| YDR126W | SWF1 |  |  |  |  |  |  |  |  |
| YDR127W | ARO1 |  |  |  |  |  |  |  |  |
| YDR128W | MTC5 |  |  |  |  |  |  |  |  |
| YDR129C | SAC6 |  |  |  |  |  |  |  |  |
| YDR130C | FIN1 |  |  |  |  |  |  |  |  |
| YDR132C | MRX16 |  |  |  |  |  |  |  |  |
| YDR135C | YCF1 |  |  |  |  |  |  |  |  |
| YDR137W | RGP1 |  |  |  |  |  |  |  |  |
| YDR138W | HPR1 |  |  |  |  |  |  |  |  |
| YDR139C | RUB1 |  |  |  |  |  |  |  |  |
| YDR140W | MTQ2 |  |  |  |  |  |  |  |  |
| YDR141C | DOP1 |  |  |  |  |  |  |  |  |
| YDR142C | PEX7 |  |  |  |  |  |  |  |  |
| YDR143C | SAN1 |  |  |  |  |  |  |  |  |
| YDR144C | MKC7 |  |  |  |  |  |  |  |  |
| YDR145W | TAF12 |  |  |  |  |  |  |  |  |
| YDR146C | SWI5 |  |  |  |  |  |  |  |  |
| YDR147W | EKI1 |  |  |  |  |  |  |  |  |
| YDR148C | KGD2 |  |  |  |  |  |  |  |  |
| YDR150W | NUM1 |  |  |  |  |  |  |  |  |
| YDR151C | CTH1 |  |  |  |  |  |  |  |  |
| YDR152W | GIR2 |  |  |  |  |  |  |  |  |
| YDR153C | ENT5 |  |  |  |  |  |  |  |  |
| YDR155C | CPR1 |  |  |  |  |  |  |  |  |
| YDR156W | RPA14 |  |  |  |  |  |  |  |  |
| YDR158W | HOM2 |  |  |  |  |  |  |  |  |
| YDR159W | SAC3 |  |  |  |  |  |  |  |  |
| YDR160W | SSY1 |  |  |  |  |  |  |  |  |
| YDR161W | ACL4 |  |  |  |  |  |  |  |  |
| YDR162C | NBP2 |  |  |  |  |  |  |  |  |
| YDR163W | CWC15 |  |  |  |  |  |  |  |  |
| YDR164C | SEC1 |  |  |  |  |  |  |  |  |
| YDR165W | TRM82 |  |  |  |  |  |  |  |  |
| YDR166C | SEC5 |  |  |  |  |  |  |  |  |
| YDR167W | TAF10 |  |  |  |  |  |  |  |  |
| YDR168W | CDC37 |  |  |  |  |  |  |  |  |
| YDR169C | STB3 |  |  |  |  |  |  |  |  |
| YDR170C | SEC7 |  |  |  |  |  |  |  |  |
| YDR172W | SUP35 |  |  |  |  |  |  |  |  |
| YDR173C | ARG82 |  |  |  |  |  |  |  |  |
| YDR174W | HMO1 |  |  |  |  |  |  |  |  |
| YDR175C | RSM24 |  |  |  |  |  |  |  |  |
| YDR176W | NGG1 |  |  |  |  |  |  |  |  |
| YDR177W | UBC1 |  |  |  |  |  |  |  |  |
| YDR178W | SDH4 |  |  |  |  |  |  |  |  |
| YDR179C | CSN9 |  |  |  |  |  |  |  |  |
| YDR179W-A | NVJ3 |  |  |  |  |  |  |  |  |
| YDR180W | SCC2 |  |  |  |  |  |  |  |  |
| YDR181C | SAS4 |  |  |  |  |  |  |  |  |
| YDR182W | CDC1 |  |  |  |  |  |  |  |  |
| YDR183W | PLP1 |  |  |  |  |  |  |  |  |
| YDR184C | ATC1 |  |  |  |  |  |  |  |  |
| YDR185C | UPS3 |  |  |  |  |  |  |  |  |
| YDR186C | SND1 |  |  |  |  |  |  |  |  |
| YDR188W | CCT6 |  |  |  |  |  |  |  |  |
| YDR189W | SLY1 |  |  |  |  |  |  |  |  |
| YDR190C | RVB1 |  |  |  |  |  |  |  |  |
| YDR191W | HST4 |  |  |  |  |  |  |  |  |
| YDR192C | NUP42 |  |  |  |  |  |  |  |  |
| YDR194C | MSS116 |  |  |  |  |  |  |  |  |
| YDR195W | REF2 |  |  |  |  |  |  |  |  |
| YDR196C | CAB5 |  |  |  |  |  |  |  |  |
| YDR197W | CBS2 |  |  |  |  |  |  |  |  |
| YDR198C | RKM2 |  |  |  |  |  |  |  |  |
| YDR200C | VPS64 |  |  |  |  |  |  |  |  |
| YDR201W | SPC19 |  |  |  |  |  |  |  |  |
| YDR202C | RAV2 |  |  |  |  |  |  |  |  |
| YDR204W | COQ4 |  |  |  |  |  |  |  |  |
| YDR205W | MSC2 |  |  |  |  |  |  |  |  |
| YDR206W | EBS1 |  |  |  |  |  |  |  |  |
| YDR207C | UME6 |  |  |  |  |  |  |  |  |
| YDR208W | MSS4 |  |  |  |  |  |  |  |  |
| YDR210W | CPP2 |  |  |  |  |  |  |  |  |
| YDR211W | GCD6 |  |  |  |  |  |  |  |  |
| YDR212W | TCP1 |  |  |  |  |  |  |  |  |
| YDR213W | UPC2 |  |  |  |  |  |  |  |  |
| YDR214W | AHA1 |  |  |  |  |  |  |  |  |
| YDR216W | ADR1 |  |  |  |  |  |  |  |  |
| YDR217C | RAD9 |  |  |  |  |  |  |  |  |
| YDR218C | SPR28 |  |  |  |  |  |  |  |  |
| YDR219C | MFB1 |  |  |  |  |  |  |  |  |
| YDR221W | GTB1 |  |  |  |  |  |  |  |  |
| YDR223W | CRF1 |  |  |  |  |  |  |  |  |
| YDR224C | HTB1 |  |  |  |  |  |  |  |  |
| YDR225W | HTA1 |  |  |  |  |  |  |  |  |
| YDR226W | ADK1 |  |  |  |  |  |  |  |  |
| YDR227W | SIR4 |  |  |  |  |  |  |  |  |
| YDR228C | PCF11 |  |  |  |  |  |  |  |  |
| YDR229W | IVY1 |  |  |  |  |  |  |  |  |
| YDR231C | COX20 |  |  |  |  |  |  |  |  |
| YDR232W | HEM1 |  |  |  |  |  |  |  |  |
| YDR233C | RTN1 |  |  |  |  |  |  |  |  |
| YDR234W | LYS4 |  |  |  |  |  |  |  |  |
| YDR235W | PRP42 |  |  |  |  |  |  |  |  |
| YDR236C | FMN1 |  |  |  |  |  |  |  |  |
| YDR237W | MRPL7 |  |  |  |  |  |  |  |  |
| YDR238C | SEC26 |  |  |  |  |  |  |  |  |
| YDR239C |  |  |  |  |  |  |  |  |  |
| YDR240C | SNU56 |  |  |  |  |  |  |  |  |
| YDR243C | PRP28 |  |  |  |  |  |  |  |  |
| YDR244W | PEX5 |  |  |  |  |  |  |  |  |
| YDR245W | MNN10 |  |  |  |  |  |  |  |  |
| YDR246W | TRS23 |  |  |  |  |  |  |  |  |
| YDR247W | VHS1 |  |  |  |  |  |  |  |  |
| YDR251W | PAM1 |  |  |  |  |  |  |  |  |
| YDR252W | BTT1 |  |  |  |  |  |  |  |  |
| YDR253C | MET32 |  |  |  |  |  |  |  |  |
| YDR254W | CHL4 |  |  |  |  |  |  |  |  |
| YDR255C | RMD5 |  |  |  |  |  |  |  |  |
| YDR256C | CTA1 |  |  |  |  |  |  |  |  |
| YDR257C | RKM4 |  |  |  |  |  |  |  |  |
| YDR258C | HSP78 |  |  |  |  |  |  |  |  |
| YDR259C | YAP6 |  |  |  |  |  |  |  |  |
| YDR260C | SWM1 |  |  |  |  |  |  |  |  |
| YDR261C | EXG2 |  |  |  |  |  |  |  |  |
| YDR263C | DIN7 |  |  |  |  |  |  |  |  |
| YDR264C | AKR1 |  |  |  |  |  |  |  |  |
| YDR265W | PEX10 |  |  |  |  |  |  |  |  |
| YDR266C | HEL2 |  |  |  |  |  |  |  |  |
| YDR267C | CIA1 |  |  |  |  |  |  |  |  |
| YDR268W | MSW1 |  |  |  |  |  |  |  |  |
| YDR270W | CCC2 |  |  |  |  |  |  |  |  |
| YDR272W | GLO2 |  |  |  |  |  |  |  |  |
| YDR273W | DON1 |  |  |  |  |  |  |  |  |
| YDR275W | TLD1 |  |  |  |  |  |  |  |  |
| YDR276C | PMP3 |  |  |  |  |  |  |  |  |
| YDR277C | MTH1 |  |  |  |  |  |  |  |  |
| YDR279W | RNH202 |  |  |  |  |  |  |  |  |
| YDR280W | RRP45 |  |  |  |  |  |  |  |  |
| YDR282C | MRX10 |  |  |  |  |  |  |  |  |
| YDR283C | GCN2 |  |  |  |  |  |  |  |  |
| YDR284C | DPP1 |  |  |  |  |  |  |  |  |
| YDR285W | ZIP1 |  |  |  |  |  |  |  |  |
| YDR288W | NSE3 |  |  |  |  |  |  |  |  |
| YDR289C | RTT103 |  |  |  |  |  |  |  |  |
| YDR291W | HRQ1 |  |  |  |  |  |  |  |  |
| YDR292C | SRP101 |  |  |  |  |  |  |  |  |
| YDR293C | SSD1 |  |  |  |  |  |  |  |  |
| YDR295C | HDA2 |  |  |  |  |  |  |  |  |
| YDR296W | MHR1 |  |  |  |  |  |  |  |  |
| YDR297W | SUR2 |  |  |  |  |  |  |  |  |
| YDR298C | ATP5 |  |  |  |  |  |  |  |  |
| YDR299W | BFR2 |  |  |  |  |  |  |  |  |
| YDR300C | PRO1 |  |  |  |  |  |  |  |  |
| YDR301W | CFT1 |  |  |  |  |  |  |  |  |
| YDR302W | GPI11 |  |  |  |  |  |  |  |  |
| YDR303C | RSC3 |  |  |  |  |  |  |  |  |
| YDR304C | CPR5 |  |  |  |  |  |  |  |  |
| YDR305C | HNT2 |  |  |  |  |  |  |  |  |
| YDR306C | PFU1 |  |  |  |  |  |  |  |  |
| YDR308C | SRB7 |  |  |  |  |  |  |  |  |
| YDR309C | GIC2 |  |  |  |  |  |  |  |  |
| YDR310C | SUM1 |  |  |  |  |  |  |  |  |
| YDR311W | TFB1 |  |  |  |  |  |  |  |  |
| YDR312W | SSF2 |  |  |  |  |  |  |  |  |
| YDR313C | PIB1 |  |  |  |  |  |  |  |  |
| YDR315C | IPK1 |  |  |  |  |  |  |  |  |
| YDR316W | OMS1 |  |  |  |  |  |  |  |  |
| YDR317W | HIM1 |  |  |  |  |  |  |  |  |
| YDR318W | MCM21 |  |  |  |  |  |  |  |  |
| YDR320C | SWA2 |  |  |  |  |  |  |  |  |
| YDR320C-A | DAD4 |  |  |  |  |  |  |  |  |
| YDR321W | ASP1 |  |  |  |  |  |  |  |  |
| YDR322C-A | TIM11 |  |  |  |  |  |  |  |  |
| YDR322W | MRPL35 |  |  |  |  |  |  |  |  |
| YDR323C | PEP7 |  |  |  |  |  |  |  |  |
| YDR324C | UTP4 |  |  |  |  |  |  |  |  |
| YDR325W | YCG1 |  |  |  |  |  |  |  |  |
| YDR326C | YSP2 |  |  |  |  |  |  |  |  |
| YDR328C | SKP1 |  |  |  |  |  |  |  |  |
| YDR329C | PEX3 |  |  |  |  |  |  |  |  |
| YDR330W | UBX5 |  |  |  |  |  |  |  |  |
| YDR331W | GPI8 |  |  |  |  |  |  |  |  |
| YDR332W | IRC3 |  |  |  |  |  |  |  |  |
| YDR333C | RQC1 |  |  |  |  |  |  |  |  |
| YDR334W | SWR1 |  |  |  |  |  |  |  |  |
| YDR335W | MSN5 |  |  |  |  |  |  |  |  |
| YDR336W | MRX8 |  |  |  |  |  |  |  |  |
| YDR337W | MRPS28 |  |  |  |  |  |  |  |  |
| YDR339C | FCF1 |  |  |  |  |  |  |  |  |
| YDR341C |  |  |  |  |  |  |  |  |  |
| YDR342C_343C | HXT7_HXT6 |  |  |  |  |  |  |  |  |
| YDR345C | HXT3 |  |  |  |  |  |  |  |  |
| YDR346C | SVF1 |  |  |  |  |  |  |  |  |
| YDR347W | MRP1 |  |  |  |  |  |  |  |  |
| YDR348C | PAL1 |  |  |  |  |  |  |  |  |
| YDR349C | YPS7 |  |  |  |  |  |  |  |  |
| YDR350C | ATP22 |  |  |  |  |  |  |  |  |
| YDR351W | SBE2 |  |  |  |  |  |  |  |  |
| YDR352W | YPQ2 |  |  |  |  |  |  |  |  |
| YDR353W | TRR1 |  |  |  |  |  |  |  |  |
| YDR354W | TRP4 |  |  |  |  |  |  |  |  |
| YDR356W | SPC110 |  |  |  |  |  |  |  |  |
| YDR357C | CNL1 |  |  |  |  |  |  |  |  |
| YDR358W | GGA1 |  |  |  |  |  |  |  |  |
| YDR359C | EAF1 |  |  |  |  |  |  |  |  |
| YDR361C | BCP1 |  |  |  |  |  |  |  |  |
| YDR362C | TFC6 |  |  |  |  |  |  |  |  |
| YDR363W | ESC2 |  |  |  |  |  |  |  |  |
| YDR363W-A | SEM1 |  |  |  |  |  |  |  |  |
| YDR364C | CDC40 |  |  |  |  |  |  |  |  |
| YDR365C | ESF1 |  |  |  |  |  |  |  |  |
| YDR367W | KEI1 |  |  |  |  |  |  |  |  |
| YDR368W | YPR1 |  |  |  |  |  |  |  |  |
| YDR369C | XRS2 |  |  |  |  |  |  |  |  |
| YDR370C | DXO1 |  |  |  |  |  |  |  |  |
| YDR371W | CTS2 |  |  |  |  |  |  |  |  |
| YDR372C | VPS74 |  |  |  |  |  |  |  |  |
| YDR373W | FRQ1 |  |  |  |  |  |  |  |  |
| YDR374C | PHO92 |  |  |  |  |  |  |  |  |
| YDR374W-A | WIP1 |  |  |  |  |  |  |  |  |
| YDR375C | BCS1 |  |  |  |  |  |  |  |  |
| YDR376W | ARH1 |  |  |  |  |  |  |  |  |
| YDR377W | ATP17 |  |  |  |  |  |  |  |  |
| YDR378C | LSM6 |  |  |  |  |  |  |  |  |
| YDR379C-A | SDH6 |  |  |  |  |  |  |  |  |
| YDR379W | RGA2 |  |  |  |  |  |  |  |  |
| YDR380W | ARO10 |  |  |  |  |  |  |  |  |
| YDR381C-A | COI1 |  |  |  |  |  |  |  |  |
| YDR381W | YRA1 |  |  |  |  |  |  |  |  |
| YDR382W | RPP2B |  |  |  |  |  |  |  |  |
| YDR384C | ATO3 |  |  |  |  |  |  |  |  |
| YDR385W | EFT2 |  |  |  |  |  |  |  |  |
| YDR386W | MUS81 |  |  |  |  |  |  |  |  |
| YDR388W | RVS167 |  |  |  |  |  |  |  |  |
| YDR389W | SAC7 |  |  |  |  |  |  |  |  |
| YDR390C | UBA2 |  |  |  |  |  |  |  |  |
| YDR392W | SPT3 |  |  |  |  |  |  |  |  |
| YDR393W | SHE9 |  |  |  |  |  |  |  |  |
| YDR394W | RPT3 |  |  |  |  |  |  |  |  |
| YDR395W | SXM1 |  |  |  |  |  |  |  |  |
| YDR397C | NCB2 |  |  |  |  |  |  |  |  |
| YDR398W | UTP5 |  |  |  |  |  |  |  |  |
| YDR400W | URH1 |  |  |  |  |  |  |  |  |
| YDR402C | DIT2 |  |  |  |  |  |  |  |  |
| YDR403W | DIT1 |  |  |  |  |  |  |  |  |
| YDR404C | RPB7 |  |  |  |  |  |  |  |  |
| YDR405W | MRP20 |  |  |  |  |  |  |  |  |
| YDR406W | PDR15 |  |  |  |  |  |  |  |  |
| YDR407C | TRS120 |  |  |  |  |  |  |  |  |
| YDR408C | ADE8 |  |  |  |  |  |  |  |  |
| YDR409W | SIZ1 |  |  |  |  |  |  |  |  |
| YDR410C | STE14 |  |  |  |  |  |  |  |  |
| YDR411C | DFM1 |  |  |  |  |  |  |  |  |
| YDR412W | RRP17 |  |  |  |  |  |  |  |  |
| YDR414C | ERD1 |  |  |  |  |  |  |  |  |
| YDR416W | SYF1 |  |  |  |  |  |  |  |  |
| YDR418W | RPL12B |  |  |  |  |  |  |  |  |
| YDR419W | RAD30 |  |  |  |  |  |  |  |  |
| YDR420W | HKR1 |  |  |  |  |  |  |  |  |
| YDR421W | ARO80 |  |  |  |  |  |  |  |  |
| YDR422C | SIP1 |  |  |  |  |  |  |  |  |
| YDR423C | CAD1 |  |  |  |  |  |  |  |  |
| YDR424C | DYN2 |  |  |  |  |  |  |  |  |
| YDR425W | SNX41 |  |  |  |  |  |  |  |  |
| YDR427W | RPN9 |  |  |  |  |  |  |  |  |
| YDR428C | BNA7 |  |  |  |  |  |  |  |  |
| YDR429C | TIF35 |  |  |  |  |  |  |  |  |
| YDR430C | CYM1 |  |  |  |  |  |  |  |  |
| YDR432W | NPL3 |  |  |  |  |  |  |  |  |
| YDR434W | GPI17 |  |  |  |  |  |  |  |  |
| YDR435C | PPM1 |  |  |  |  |  |  |  |  |
| YDR436W | PPZ2 |  |  |  |  |  |  |  |  |
| YDR437W | GPI19 |  |  |  |  |  |  |  |  |
| YDR438W | THI74 |  |  |  |  |  |  |  |  |
| YDR439W | LRS4 |  |  |  |  |  |  |  |  |
| YDR440W | DOT1 |  |  |  |  |  |  |  |  |
| YDR441C | APT2 |  |  |  |  |  |  |  |  |
| YDR443C | SSN2 |  |  |  |  |  |  |  |  |
| YDR446W | ECM11 |  |  |  |  |  |  |  |  |
| YDR447C | RPS17B |  |  |  |  |  |  |  |  |
| YDR448W | ADA2 |  |  |  |  |  |  |  |  |
| YDR449C | UTP6 |  |  |  |  |  |  |  |  |
| YDR450W | RPS18A |  |  |  |  |  |  |  |  |
| YDR451C | YHP1 |  |  |  |  |  |  |  |  |
| YDR452W | PPN1 |  |  |  |  |  |  |  |  |
| YDR453C | TSA2 |  |  |  |  |  |  |  |  |
| YDR454C | GUK1 |  |  |  |  |  |  |  |  |
| YDR456W | NHX1 |  |  |  |  |  |  |  |  |
| YDR457W | TOM1 |  |  |  |  |  |  |  |  |
| YDR458C | HEH2 |  |  |  |  |  |  |  |  |
| YDR459C | PFA5 |  |  |  |  |  |  |  |  |
| YDR460W | TFB3 |  |  |  |  |  |  |  |  |
| YDR462W | MRPL28 |  |  |  |  |  |  |  |  |
| YDR463W | STP1 |  |  |  |  |  |  |  |  |
| YDR464W | SPP41 |  |  |  |  |  |  |  |  |
| YDR465C | RMT2 |  |  |  |  |  |  |  |  |
| YDR466W | PKH3 |  |  |  |  |  |  |  |  |
| YDR468C | TLG1 |  |  |  |  |  |  |  |  |
| YDR469W | SDC1 |  |  |  |  |  |  |  |  |
| YDR470C | UGO1 |  |  |  |  |  |  |  |  |
| YDR471W | RPL27B |  |  |  |  |  |  |  |  |
| YDR472W | TRS31 |  |  |  |  |  |  |  |  |
| YDR473C | PRP3 |  |  |  |  |  |  |  |  |
| YDR475C | JIP4 |  |  |  |  |  |  |  |  |
| YDR477W | SNF1 |  |  |  |  |  |  |  |  |
| YDR478W | SNM1 |  |  |  |  |  |  |  |  |
| YDR479C | PEX29 |  |  |  |  |  |  |  |  |
| YDR480W | DIG2 |  |  |  |  |  |  |  |  |
| YDR481C | PHO8 |  |  |  |  |  |  |  |  |
| YDR482C | CWC21 |  |  |  |  |  |  |  |  |
| YDR483W | KRE2 |  |  |  |  |  |  |  |  |
| YDR484W | VPS52 |  |  |  |  |  |  |  |  |
| YDR485C | VPS72 |  |  |  |  |  |  |  |  |
| YDR486C | VPS60 |  |  |  |  |  |  |  |  |
| YDR487C | RIB3 |  |  |  |  |  |  |  |  |
| YDR488C | PAC11 |  |  |  |  |  |  |  |  |
| YDR489W | SLD5 |  |  |  |  |  |  |  |  |
| YDR490C | PKH1 |  |  |  |  |  |  |  |  |
| YDR492W | IZH1 |  |  |  |  |  |  |  |  |
| YDR493W | MZM1 |  |  |  |  |  |  |  |  |
| YDR494W | RSM28 |  |  |  |  |  |  |  |  |
| YDR495C | VPS3 |  |  |  |  |  |  |  |  |
| YDR496C | PUF6 |  |  |  |  |  |  |  |  |
| YDR497C | ITR1 |  |  |  |  |  |  |  |  |
| YDR498C | SEC20 |  |  |  |  |  |  |  |  |
| YDR499W | LCD1 |  |  |  |  |  |  |  |  |
| YDR500C | RPL37B |  |  |  |  |  |  |  |  |
| YDR501W | PLM2 |  |  |  |  |  |  |  |  |
| YDR502C | SAM2 |  |  |  |  |  |  |  |  |
| YDR503C | LPP1 |  |  |  |  |  |  |  |  |
| YDR505C | PSP1 |  |  |  |  |  |  |  |  |
| YDR507C | GIN4 |  |  |  |  |  |  |  |  |
| YDR508C | GNP1 |  |  |  |  |  |  |  |  |
| YDR510W | SMT3 |  |  |  |  |  |  |  |  |
| YDR511W | SDH7 |  |  |  |  |  |  |  |  |
| YDR512C | EMI1 |  |  |  |  |  |  |  |  |
| YDR513W | GRX2 |  |  |  |  |  |  |  |  |
| YDR514C |  |  |  |  |  |  |  |  |  |
| YDR515W | SLF1 |  |  |  |  |  |  |  |  |
| YDR516C | EMI2 |  |  |  |  |  |  |  |  |
| YDR517W | GRH1 |  |  |  |  |  |  |  |  |
| YDR518W | EUG1 |  |  |  |  |  |  |  |  |
| YDR522C | SPS2 |  |  |  |  |  |  |  |  |
| YDR523C | SPS1 |  |  |  |  |  |  |  |  |
| YDR524C | AGE1 |  |  |  |  |  |  |  |  |
| YDR525W-A | SNA2 |  |  |  |  |  |  |  |  |
| YDR527W | RBA50 |  |  |  |  |  |  |  |  |
| YDR528W | HLR1 |  |  |  |  |  |  |  |  |
| YDR529C | QCR7 |  |  |  |  |  |  |  |  |
| YDR530C | APA2 |  |  |  |  |  |  |  |  |
| YDR531W | CAB1 |  |  |  |  |  |  |  |  |
| YDR532C | KRE28 |  |  |  |  |  |  |  |  |
| YDR536W | STL1 |  |  |  |  |  |  |  |  |
| YDR540C | IRC4 |  |  |  |  |  |  |  |  |
| YEL001C | IRC22 |  |  |  |  |  |  |  |  |
| YEL002C | WBP1 |  |  |  |  |  |  |  |  |
| YEL003W | GIM4 |  |  |  |  |  |  |  |  |
| YEL005C | VAB2 |  |  |  |  |  |  |  |  |
| YEL006W | YEA6 |  |  |  |  |  |  |  |  |
| YEL007W | MIT1 |  |  |  |  |  |  |  |  |
| YEL009C | GCN4 |  |  |  |  |  |  |  |  |
| YEL011W | GLC3 |  |  |  |  |  |  |  |  |
| YEL012W | UBC8 |  |  |  |  |  |  |  |  |
| YEL013W | VAC8 |  |  |  |  |  |  |  |  |
| YEL015W | EDC3 |  |  |  |  |  |  |  |  |
| YEL016C | NPP2 |  |  |  |  |  |  |  |  |
| YEL017C-A | PMP2 |  |  |  |  |  |  |  |  |
| YEL017W | GTT3 |  |  |  |  |  |  |  |  |
| YEL018W | EAF5 |  |  |  |  |  |  |  |  |
| YEL019C | MMS21 |  |  |  |  |  |  |  |  |
| YEL020C | PXP1 |  |  |  |  |  |  |  |  |
| YEL020W-A | TIM9 |  |  |  |  |  |  |  |  |
| YEL021W | URA3 |  |  |  |  |  |  |  |  |
| YEL022W | GEA2 |  |  |  |  |  |  |  |  |
| YEL024W | RIP1 |  |  |  |  |  |  |  |  |
| YEL026W | SNU13 |  |  |  |  |  |  |  |  |
| YEL027W | VMA3 |  |  |  |  |  |  |  |  |
| YEL029C | BUD16 |  |  |  |  |  |  |  |  |
| YEL030W | ECM10 |  |  |  |  |  |  |  |  |
| YEL031W | SPF1 |  |  |  |  |  |  |  |  |
| YEL032W | MCM3 |  |  |  |  |  |  |  |  |
| YEL034W | HYP2 |  |  |  |  |  |  |  |  |
| YEL036C | ANP1 |  |  |  |  |  |  |  |  |
| YEL037C | RAD23 |  |  |  |  |  |  |  |  |
| YEL038W | UTR4 |  |  |  |  |  |  |  |  |
| YEL039C | CYC7 |  |  |  |  |  |  |  |  |
| YEL041W | YEF1 |  |  |  |  |  |  |  |  |
| YEL042W | GDA1 |  |  |  |  |  |  |  |  |
| YEL043W | GTA1 |  |  |  |  |  |  |  |  |
| YEL044W | IES6 |  |  |  |  |  |  |  |  |
| YEL046C | GLY1 |  |  |  |  |  |  |  |  |
| YEL047C | FRD1 |  |  |  |  |  |  |  |  |
| YEL048C | TCA17 |  |  |  |  |  |  |  |  |
| YEL050C | RML2 |  |  |  |  |  |  |  |  |
| YEL051W | VMA8 |  |  |  |  |  |  |  |  |
| YEL052W | AFG1 |  |  |  |  |  |  |  |  |
| YEL053C | MAK10 |  |  |  |  |  |  |  |  |
| YEL054C | RPL12A |  |  |  |  |  |  |  |  |
| YEL055C | POL5 |  |  |  |  |  |  |  |  |
| YEL056W | HAT2 |  |  |  |  |  |  |  |  |
| YEL058W | PCM1 |  |  |  |  |  |  |  |  |
| YEL059C-A | SOM1 |  |  |  |  |  |  |  |  |
| YEL060C | PRB1 |  |  |  |  |  |  |  |  |
| YEL061C | CIN8 |  |  |  |  |  |  |  |  |
| YEL062W | NPR2 |  |  |  |  |  |  |  |  |
| YEL063C | CAN1 |  |  |  |  |  |  |  |  |
| YEL064C | AVT2 |  |  |  |  |  |  |  |  |
| YER001W | MNN1 |  |  |  |  |  |  |  |  |
| YER002W | NOP16 |  |  |  |  |  |  |  |  |
| YER003C | PMI40 |  |  |  |  |  |  |  |  |
| YER004W | FMP52 |  |  |  |  |  |  |  |  |
| YER005W | YND1 |  |  |  |  |  |  |  |  |
| YER006W | NUG1 |  |  |  |  |  |  |  |  |
| YER007C-A | TMA20 |  |  |  |  |  |  |  |  |
| YER007W | PAC2 |  |  |  |  |  |  |  |  |
| YER008C | SEC3 |  |  |  |  |  |  |  |  |
| YER009W | NTF2 |  |  |  |  |  |  |  |  |
| YER010C |  |  |  |  |  |  |  |  |  |
| YER011W | TIR1 |  |  |  |  |  |  |  |  |
| YER012W | PRE1 |  |  |  |  |  |  |  |  |
| YER013W | PRP22 |  |  |  |  |  |  |  |  |
| YER014W | HEM14 |  |  |  |  |  |  |  |  |
| YER015W | FAA2 |  |  |  |  |  |  |  |  |
| YER016W | BIM1 |  |  |  |  |  |  |  |  |
| YER017C | AFG3 |  |  |  |  |  |  |  |  |
| YER018C | SPC25 |  |  |  |  |  |  |  |  |
| YER019C-A | SBH2 |  |  |  |  |  |  |  |  |
| YER019W | ISC1 |  |  |  |  |  |  |  |  |
| YER020W | GPA2 |  |  |  |  |  |  |  |  |
| YER021W | RPN3 |  |  |  |  |  |  |  |  |
| YER022W | SRB4 |  |  |  |  |  |  |  |  |
| YER023W | PRO3 |  |  |  |  |  |  |  |  |
| YER024W | YAT2 |  |  |  |  |  |  |  |  |
| YER025W | GCD11 |  |  |  |  |  |  |  |  |
| YER026C | CHO1 |  |  |  |  |  |  |  |  |
| YER027C | GAL83 |  |  |  |  |  |  |  |  |
| YER029C | SMB1 |  |  |  |  |  |  |  |  |
| YER030W | CHZ1 |  |  |  |  |  |  |  |  |
| YER031C | YPT31 |  |  |  |  |  |  |  |  |
| YER032W | FIR1 |  |  |  |  |  |  |  |  |
| YER033C | ZRG8 |  |  |  |  |  |  |  |  |
| YER034W |  |  |  |  |  |  |  |  |  |
| YER035W | EDC2 |  |  |  |  |  |  |  |  |
| YER036C | ARB1 |  |  |  |  |  |  |  |  |
| YER037W | PHM8 |  |  |  |  |  |  |  |  |
| YER038C | KRE29 |  |  |  |  |  |  |  |  |
| YER039C | HVG1 |  |  |  |  |  |  |  |  |
| YER040W | GLN3 |  |  |  |  |  |  |  |  |
| YER041W | YEN1 |  |  |  |  |  |  |  |  |
| YER042W | MXR1 |  |  |  |  |  |  |  |  |
| YER043C | SAH1 |  |  |  |  |  |  |  |  |
| YER044C | ERG28 |  |  |  |  |  |  |  |  |
| YER044C-A | MEI4 |  |  |  |  |  |  |  |  |
| YER045C | ACA1 |  |  |  |  |  |  |  |  |
| YER046W | SPO73 |  |  |  |  |  |  |  |  |
| YER047C | SAP1 |  |  |  |  |  |  |  |  |
| YER048C | CAJ1 |  |  |  |  |  |  |  |  |
| YER048W-A | ISD11 |  |  |  |  |  |  |  |  |
| YER049W | TPA1 |  |  |  |  |  |  |  |  |
| YER050C | RSM18 |  |  |  |  |  |  |  |  |
| YER051W | JHD1 |  |  |  |  |  |  |  |  |
| YER052C | HOM3 |  |  |  |  |  |  |  |  |
| YER054C | GIP2 |  |  |  |  |  |  |  |  |
| YER055C | HIS1 |  |  |  |  |  |  |  |  |
| YER056C | FCY2 |  |  |  |  |  |  |  |  |
| YER056C-A | RPL34A |  |  |  |  |  |  |  |  |
| YER057C | HMF1 |  |  |  |  |  |  |  |  |
| YER058W | PET117 |  |  |  |  |  |  |  |  |
| YER059W | PCL6 |  |  |  |  |  |  |  |  |
| YER060W | FCY21 |  |  |  |  |  |  |  |  |
| YER060W-A | FCY22 |  |  |  |  |  |  |  |  |
| YER061C | CEM1 |  |  |  |  |  |  |  |  |
| YER062C | GPP2 |  |  |  |  |  |  |  |  |
| YER063W | THO1 |  |  |  |  |  |  |  |  |
| YER064C | VHR2 |  |  |  |  |  |  |  |  |
| YER065C | ICL1 |  |  |  |  |  |  |  |  |
| YER067W | RGI1 |  |  |  |  |  |  |  |  |
| YER068W | MOT2 |  |  |  |  |  |  |  |  |
| YER069W | ARG56 |  |  |  |  |  |  |  |  |
| YER070W | RNR1 |  |  |  |  |  |  |  |  |
| YER071C | TDA2 |  |  |  |  |  |  |  |  |
| YER072W | VTC1 |  |  |  |  |  |  |  |  |
| YER073W | ALD5 |  |  |  |  |  |  |  |  |
| YER074W | RPS24A |  |  |  |  |  |  |  |  |
| YER074W-A | YOS1 |  |  |  |  |  |  |  |  |
| YER075C | PTP3 |  |  |  |  |  |  |  |  |
| YER078C | ICP55 |  |  |  |  |  |  |  |  |
| YER080W | AIM9 |  |  |  |  |  |  |  |  |
| YER081W | SER3 |  |  |  |  |  |  |  |  |
| YER082C | UTP7 |  |  |  |  |  |  |  |  |
| YER083C | GET2 |  |  |  |  |  |  |  |  |
| YER086W | ILV1 |  |  |  |  |  |  |  |  |
| YER087C-B | SBH1 |  |  |  |  |  |  |  |  |
| YER087W | AIM10 |  |  |  |  |  |  |  |  |
| YER088C | DOT6 |  |  |  |  |  |  |  |  |
| YER089C | PTC2 |  |  |  |  |  |  |  |  |
| YER090W | TRP2 |  |  |  |  |  |  |  |  |
| YER091C | MET6 |  |  |  |  |  |  |  |  |
| YER092W | IES5 |  |  |  |  |  |  |  |  |
| YER093C | TSC11 |  |  |  |  |  |  |  |  |
| YER093C-A | AIM11 |  |  |  |  |  |  |  |  |
| YER094C | PUP3 |  |  |  |  |  |  |  |  |
| YER095W | RAD51 |  |  |  |  |  |  |  |  |
| YER096W | SHC1 |  |  |  |  |  |  |  |  |
| YER098W | UBP9 |  |  |  |  |  |  |  |  |
| YER099C | PRS2 |  |  |  |  |  |  |  |  |
| YER100W | UBC6 |  |  |  |  |  |  |  |  |
| YER101C | AST2 |  |  |  |  |  |  |  |  |
| YER102W | RPS8B |  |  |  |  |  |  |  |  |
| YER103W | SSA4 |  |  |  |  |  |  |  |  |
| YER104W | RTT105 |  |  |  |  |  |  |  |  |
| YER105C | NUP157 |  |  |  |  |  |  |  |  |
| YER106W | MAM1 |  |  |  |  |  |  |  |  |
| YER107C | GLE2 |  |  |  |  |  |  |  |  |
| YER109C | FLO8 |  |  |  |  |  |  |  |  |
| YER110C | KAP123 |  |  |  |  |  |  |  |  |
| YER111C | SWI4 |  |  |  |  |  |  |  |  |
| YER112W | LSM4 |  |  |  |  |  |  |  |  |
| YER113C | TMN3 |  |  |  |  |  |  |  |  |
| YER114C | BOI2 |  |  |  |  |  |  |  |  |
| YER115C | SPR6 |  |  |  |  |  |  |  |  |
| YER116C | SLX8 |  |  |  |  |  |  |  |  |
| YER117W | RPL23B |  |  |  |  |  |  |  |  |
| YER118C | SHO1 |  |  |  |  |  |  |  |  |
| YER119C | AVT6 |  |  |  |  |  |  |  |  |
| YER120W | SCS2 |  |  |  |  |  |  |  |  |
| YER122C | GLO3 |  |  |  |  |  |  |  |  |
| YER123W | YCK3 |  |  |  |  |  |  |  |  |
| YER124C | DSE1 |  |  |  |  |  |  |  |  |
| YER125W | RSP5 |  |  |  |  |  |  |  |  |
| YER126C | NSA2 |  |  |  |  |  |  |  |  |
| YER127W | LCP5 |  |  |  |  |  |  |  |  |
| YER128W | VFA1 |  |  |  |  |  |  |  |  |
| YER129W | SAK1 |  |  |  |  |  |  |  |  |
| YER130C | COM2 |  |  |  |  |  |  |  |  |
| YER131W | RPS26B |  |  |  |  |  |  |  |  |
| YER132C | PMD1 |  |  |  |  |  |  |  |  |
| YER133W | GLC7 |  |  |  |  |  |  |  |  |
| YER134C |  |  |  |  |  |  |  |  |  |
| YER136W | GDI1 |  |  |  |  |  |  |  |  |
| YER137C |  |  |  |  |  |  |  |  |  |
| YER139C | RTR1 |  |  |  |  |  |  |  |  |
| YER140W | EMP65 |  |  |  |  |  |  |  |  |
| YER141W | COX15 |  |  |  |  |  |  |  |  |
| YER142C | MAG1 |  |  |  |  |  |  |  |  |
| YER143W | DDI1 |  |  |  |  |  |  |  |  |
| YER144C | UBP5 |  |  |  |  |  |  |  |  |
| YER145C | FTR1 |  |  |  |  |  |  |  |  |
| YER146W | LSM5 |  |  |  |  |  |  |  |  |
| YER147C | SCC4 |  |  |  |  |  |  |  |  |
| YER148W | SPT15 |  |  |  |  |  |  |  |  |
| YER149C | PEA2 |  |  |  |  |  |  |  |  |
| YER150W | SPI1 |  |  |  |  |  |  |  |  |
| YER151C | UBP3 |  |  |  |  |  |  |  |  |
| YER152C |  |  |  |  |  |  |  |  |  |
| YER153C | PET122 |  |  |  |  |  |  |  |  |
| YER154W | OXA1 |  |  |  |  |  |  |  |  |
| YER155C | BEM2 |  |  |  |  |  |  |  |  |
| YER156C | MYG1 |  |  |  |  |  |  |  |  |
| YER157W | COG3 |  |  |  |  |  |  |  |  |
| YER159C | BUR6 |  |  |  |  |  |  |  |  |
| YER161C | SPT2 |  |  |  |  |  |  |  |  |
| YER162C | RAD4 |  |  |  |  |  |  |  |  |
| YER163C | GCG1 |  |  |  |  |  |  |  |  |
| YER164W | CHD1 |  |  |  |  |  |  |  |  |
| YER165W | PAB1 |  |  |  |  |  |  |  |  |
| YER166W | DNF1 |  |  |  |  |  |  |  |  |
| YER167W | BCK2 |  |  |  |  |  |  |  |  |
| YER168C | CCA1 |  |  |  |  |  |  |  |  |
| YER169W | RPH1 |  |  |  |  |  |  |  |  |
| YER170W | ADK2 |  |  |  |  |  |  |  |  |
| YER171W | RAD3 |  |  |  |  |  |  |  |  |
| YER172C | BRR2 |  |  |  |  |  |  |  |  |
| YER173W | RAD24 |  |  |  |  |  |  |  |  |
| YER174C | GRX4 |  |  |  |  |  |  |  |  |
| YER175C | TMT1 |  |  |  |  |  |  |  |  |
| YER176W | ECM32 |  |  |  |  |  |  |  |  |
| YER177W | BMH1 |  |  |  |  |  |  |  |  |
| YER178W | PDA1 |  |  |  |  |  |  |  |  |
| YER179W | DMC1 |  |  |  |  |  |  |  |  |
| YER180C | ISC10 |  |  |  |  |  |  |  |  |
| YER180C-A | SLO1 |  |  |  |  |  |  |  |  |
| YER183C | FAU1 |  |  |  |  |  |  |  |  |
| YFL001W | DEG1 |  |  |  |  |  |  |  |  |
| YFL002C | SPB4 |  |  |  |  |  |  |  |  |
| YFL004W | VTC2 |  |  |  |  |  |  |  |  |
| YFL005W | SEC4 |  |  |  |  |  |  |  |  |
| YFL007W | BLM10 |  |  |  |  |  |  |  |  |
| YFL008W | SMC1 |  |  |  |  |  |  |  |  |
| YFL009W | CDC4 |  |  |  |  |  |  |  |  |
| YFL010C | WWM1 |  |  |  |  |  |  |  |  |
| YFL013C | IES1 |  |  |  |  |  |  |  |  |
| YFL014W | HSP12 |  |  |  |  |  |  |  |  |
| YFL016C | MDJ1 |  |  |  |  |  |  |  |  |
| YFL017C | GNA1 |  |  |  |  |  |  |  |  |
| YFL017W-A | SMX2 |  |  |  |  |  |  |  |  |
| YFL018C | LPD1 |  |  |  |  |  |  |  |  |
| YFL021W | GAT1 |  |  |  |  |  |  |  |  |
| YFL022C | FRS2 |  |  |  |  |  |  |  |  |
| YFL023W | BUD27 |  |  |  |  |  |  |  |  |
| YFL024C | EPL1 |  |  |  |  |  |  |  |  |
| YFL025C | BST1 |  |  |  |  |  |  |  |  |
| YFL026W | STE2 |  |  |  |  |  |  |  |  |
| YFL027C | GYP8 |  |  |  |  |  |  |  |  |
| YFL028C | CAF16 |  |  |  |  |  |  |  |  |
| YFL029C | CAK1 |  |  |  |  |  |  |  |  |
| YFL031W | HAC1 |  |  |  |  |  |  |  |  |
| YFL033C | RIM15 |  |  |  |  |  |  |  |  |
| YFL034C-A | RPL22B |  |  |  |  |  |  |  |  |
| YFL034C-B | MOB2 |  |  |  |  |  |  |  |  |
| YFL036W | RPO41 |  |  |  |  |  |  |  |  |
| YFL037W | TUB2 |  |  |  |  |  |  |  |  |
| YFL038C | YPT1 |  |  |  |  |  |  |  |  |
| YFL039C | ACT1 |  |  |  |  |  |  |  |  |
| YFL041W | FET5 |  |  |  |  |  |  |  |  |
| YFL042C | LAM5 |  |  |  |  |  |  |  |  |
| YFL044C | OTU1 |  |  |  |  |  |  |  |  |
| YFL045C | SEC53 |  |  |  |  |  |  |  |  |
| YFL046W | FMP32 |  |  |  |  |  |  |  |  |
| YFL047W | RGD2 |  |  |  |  |  |  |  |  |
| YFL048C | EMP47 |  |  |  |  |  |  |  |  |
| YFL049W | SWP82 |  |  |  |  |  |  |  |  |
| YFR001W | LOC1 |  |  |  |  |  |  |  |  |
| YFR002W | NIC96 |  |  |  |  |  |  |  |  |
| YFR003C | YPI1 |  |  |  |  |  |  |  |  |
| YFR004W | RPN11 |  |  |  |  |  |  |  |  |
| YFR005C | SAD1 |  |  |  |  |  |  |  |  |
| YFR007W | YFH7 |  |  |  |  |  |  |  |  |
| YFR008W | FAR7 |  |  |  |  |  |  |  |  |
| YFR009W | GCN20 |  |  |  |  |  |  |  |  |
| YFR010W | UBP6 |  |  |  |  |  |  |  |  |
| YFR011C | MIC19 |  |  |  |  |  |  |  |  |
| YFR012W | DCV1 |  |  |  |  |  |  |  |  |
| YFR013W | IOC3 |  |  |  |  |  |  |  |  |
| YFR014C | CMK1 |  |  |  |  |  |  |  |  |
| YFR015C | GSY1 |  |  |  |  |  |  |  |  |
| YFR016C | AIP5 |  |  |  |  |  |  |  |  |
| YFR017C | IGD1 |  |  |  |  |  |  |  |  |
| YFR019W | FAB1 |  |  |  |  |  |  |  |  |
| YFR021W | ATG18 |  |  |  |  |  |  |  |  |
| YFR022W | ROG3 |  |  |  |  |  |  |  |  |
| YFR023W | PES4 |  |  |  |  |  |  |  |  |
| YFR024C-A | LSB3 |  |  |  |  |  |  |  |  |
| YFR025C | HIS2 |  |  |  |  |  |  |  |  |
| YFR027W | ECO1 |  |  |  |  |  |  |  |  |
| YFR028C | CDC14 |  |  |  |  |  |  |  |  |
| YFR029W | PTR3 |  |  |  |  |  |  |  |  |
| YFR030W | MET10 |  |  |  |  |  |  |  |  |
| YFR031C | SMC2 |  |  |  |  |  |  |  |  |
| YFR031C-A | RPL2A |  |  |  |  |  |  |  |  |
| YFR032C-A | RPL29 |  |  |  |  |  |  |  |  |
| YFR033C | QCR6 |  |  |  |  |  |  |  |  |
| YFR034C | PHO4 |  |  |  |  |  |  |  |  |
| YFR035W-A |  |  |  |  |  |  |  |  |  |
| YFR036W | CDC26 |  |  |  |  |  |  |  |  |
| YFR037C | RSC8 |  |  |  |  |  |  |  |  |
| YFR038W | IRC5 |  |  |  |  |  |  |  |  |
| YFR039C | OSW7 |  |  |  |  |  |  |  |  |
| YFR040W | SAP155 |  |  |  |  |  |  |  |  |
| YFR041C | ERJ5 |  |  |  |  |  |  |  |  |
| YFR042W | KEG1 |  |  |  |  |  |  |  |  |
| YFR043C | IRC6 |  |  |  |  |  |  |  |  |
| YFR044C | DUG1 |  |  |  |  |  |  |  |  |
| YFR046C | CNN1 |  |  |  |  |  |  |  |  |
| YFR047C | BNA6 |  |  |  |  |  |  |  |  |
| YFR048W | RMD8 |  |  |  |  |  |  |  |  |
| YFR049W | KGD4 |  |  |  |  |  |  |  |  |
| YFR050C | PRE4 |  |  |  |  |  |  |  |  |
| YFR051C | RET2 |  |  |  |  |  |  |  |  |
| YFR052W | RPN12 |  |  |  |  |  |  |  |  |
| YFR053C | HXK1 |  |  |  |  |  |  |  |  |
| YGL001C | ERG26 |  |  |  |  |  |  |  |  |
| YGL002W | ERP6 |  |  |  |  |  |  |  |  |
| YGL003C | CDH1 |  |  |  |  |  |  |  |  |
| YGL004C | RPN14 |  |  |  |  |  |  |  |  |
| YGL005C | COG7 |  |  |  |  |  |  |  |  |
| YGL006W | PMC1 |  |  |  |  |  |  |  |  |
| YGL008C | PMA1 |  |  |  |  |  |  |  |  |
| YGL009C | LEU1 |  |  |  |  |  |  |  |  |
| YGL010W | MPO1 |  |  |  |  |  |  |  |  |
| YGL011C | SCL1 |  |  |  |  |  |  |  |  |
| YGL012W | ERG4 |  |  |  |  |  |  |  |  |
| YGL014W | PUF4 |  |  |  |  |  |  |  |  |
| YGL016W | KAP122 |  |  |  |  |  |  |  |  |
| YGL017W | ATE1 |  |  |  |  |  |  |  |  |
| YGL018C | JAC1 |  |  |  |  |  |  |  |  |
| YGL019W | CKB1 |  |  |  |  |  |  |  |  |
| YGL020C | GET1 |  |  |  |  |  |  |  |  |
| YGL021W | ALK1 |  |  |  |  |  |  |  |  |
| YGL022W | STT3 |  |  |  |  |  |  |  |  |
| YGL023C | PIB2 |  |  |  |  |  |  |  |  |
| YGL025C | PGD1 |  |  |  |  |  |  |  |  |
| YGL026C | TRP5 |  |  |  |  |  |  |  |  |
| YGL027C | CWH41 |  |  |  |  |  |  |  |  |
| YGL029W | CGR1 |  |  |  |  |  |  |  |  |
| YGL030W | RPL30 |  |  |  |  |  |  |  |  |
| YGL031C | RPL24A |  |  |  |  |  |  |  |  |
| YGL032C | AGA2 |  |  |  |  |  |  |  |  |
| YGL033W | HOP2 |  |  |  |  |  |  |  |  |
| YGL035C | MIG1 |  |  |  |  |  |  |  |  |
| YGL036W | VIR1 |  |  |  |  |  |  |  |  |
| YGL037C | PNC1 |  |  |  |  |  |  |  |  |
| YGL038C | OCH1 |  |  |  |  |  |  |  |  |
| YGL040C | HEM2 |  |  |  |  |  |  |  |  |
| YGL043W | DST1 |  |  |  |  |  |  |  |  |
| YGL044C | RNA15 |  |  |  |  |  |  |  |  |
| YGL045W | RIM8 |  |  |  |  |  |  |  |  |
| YGL047W | ALG13 |  |  |  |  |  |  |  |  |
| YGL048C | RPT6 |  |  |  |  |  |  |  |  |
| YGL049C | TIF4632 |  |  |  |  |  |  |  |  |
| YGL050W | TYW3 |  |  |  |  |  |  |  |  |
| YGL054C | ERV14 |  |  |  |  |  |  |  |  |
| YGL055W | OLE1 |  |  |  |  |  |  |  |  |
| YGL056C | SDS23 |  |  |  |  |  |  |  |  |
| YGL057C | GEP7 |  |  |  |  |  |  |  |  |
| YGL058W | RAD6 |  |  |  |  |  |  |  |  |
| YGL059W | PKP2 |  |  |  |  |  |  |  |  |
| YGL060W | YBP2 |  |  |  |  |  |  |  |  |
| YGL061C | DUO1 |  |  |  |  |  |  |  |  |
| YGL062W | PYC1 |  |  |  |  |  |  |  |  |
| YGL063W | PUS2 |  |  |  |  |  |  |  |  |
| YGL064C | MRH4 |  |  |  |  |  |  |  |  |
| YGL065C | ALG2 |  |  |  |  |  |  |  |  |
| YGL066W | SGF73 |  |  |  |  |  |  |  |  |
| YGL068W | MNP1 |  |  |  |  |  |  |  |  |
| YGL070C | RPB9 |  |  |  |  |  |  |  |  |
| YGL071W | AFT1 |  |  |  |  |  |  |  |  |
| YGL073W | HSF1 |  |  |  |  |  |  |  |  |
| YGL075C | MPS2 |  |  |  |  |  |  |  |  |
| YGL076C | RPL7A |  |  |  |  |  |  |  |  |
| YGL077C | HNM1 |  |  |  |  |  |  |  |  |
| YGL078C | DBP3 |  |  |  |  |  |  |  |  |
| YGL079W | KXD1 |  |  |  |  |  |  |  |  |
| YGL080W | MPC1 |  |  |  |  |  |  |  |  |
| YGL083W | SCY1 |  |  |  |  |  |  |  |  |
| YGL084C | GUP1 |  |  |  |  |  |  |  |  |
| YGL086W | MAD1 |  |  |  |  |  |  |  |  |
| YGL087C | MMS2 |  |  |  |  |  |  |  |  |
| YGL089C | MF(ALPHA)2 |  |  |  |  |  |  |  |  |
| YGL091C | NBP35 |  |  |  |  |  |  |  |  |
| YGL092W | NUP145 |  |  |  |  |  |  |  |  |
| YGL093W | SPC105 |  |  |  |  |  |  |  |  |
| YGL094C | PAN2 |  |  |  |  |  |  |  |  |
| YGL095C | VPS45 |  |  |  |  |  |  |  |  |
| YGL096W | TOS8 |  |  |  |  |  |  |  |  |
| YGL097W | SRM1 |  |  |  |  |  |  |  |  |
| YGL098W | USE1 |  |  |  |  |  |  |  |  |
| YGL099W | LSG1 |  |  |  |  |  |  |  |  |
| YGL100W | SEH1 |  |  |  |  |  |  |  |  |
| YGL101W | YGK1 |  |  |  |  |  |  |  |  |
| YGL103W | RPL28 |  |  |  |  |  |  |  |  |
| YGL104C | VPS73 |  |  |  |  |  |  |  |  |
| YGL105W | ARC1 |  |  |  |  |  |  |  |  |
| YGL106W | MLC1 |  |  |  |  |  |  |  |  |
| YGL107C | RMD9 |  |  |  |  |  |  |  |  |
| YGL108C |  |  |  |  |  |  |  |  |  |
| YGL110C | CUE3 |  |  |  |  |  |  |  |  |
| YGL111W | NSA1 |  |  |  |  |  |  |  |  |
| YGL112C | TAF6 |  |  |  |  |  |  |  |  |
| YGL113W | SLD3 |  |  |  |  |  |  |  |  |
| YGL115W | SNF4 |  |  |  |  |  |  |  |  |
| YGL116W | CDC20 |  |  |  |  |  |  |  |  |
| YGL119W | COQ8 |  |  |  |  |  |  |  |  |
| YGL120C | PRP43 |  |  |  |  |  |  |  |  |
| YGL122C | NAB2 |  |  |  |  |  |  |  |  |
| YGL123W | RPS2 |  |  |  |  |  |  |  |  |
| YGL124C | MON1 |  |  |  |  |  |  |  |  |
| YGL125W | MET13 |  |  |  |  |  |  |  |  |
| YGL126W | SCS3 |  |  |  |  |  |  |  |  |
| YGL127C | SOH1 |  |  |  |  |  |  |  |  |
| YGL128C | CWC23 |  |  |  |  |  |  |  |  |
| YGL129C | RSM23 |  |  |  |  |  |  |  |  |
| YGL130W | CEG1 |  |  |  |  |  |  |  |  |
| YGL131C | SNT2 |  |  |  |  |  |  |  |  |
| YGL133W | ITC1 |  |  |  |  |  |  |  |  |
| YGL134W | PCL10 |  |  |  |  |  |  |  |  |
| YGL135W | RPL1B |  |  |  |  |  |  |  |  |
| YGL136C | MRM2 |  |  |  |  |  |  |  |  |
| YGL137W | SEC27 |  |  |  |  |  |  |  |  |
| YGL139W | FLC3 |  |  |  |  |  |  |  |  |
| YGL141W | HUL5 |  |  |  |  |  |  |  |  |
| YGL142C | GPI10 |  |  |  |  |  |  |  |  |
| YGL143C | MRF1 |  |  |  |  |  |  |  |  |
| YGL144C | ROG1 |  |  |  |  |  |  |  |  |
| YGL145W | TIP20 |  |  |  |  |  |  |  |  |
| YGL146C | RRT6 |  |  |  |  |  |  |  |  |
| YGL147C | RPL9A |  |  |  |  |  |  |  |  |
| YGL148W | ARO2 |  |  |  |  |  |  |  |  |
| YGL150C | INO80 |  |  |  |  |  |  |  |  |
| YGL151W | NUT1 |  |  |  |  |  |  |  |  |
| YGL153W | PEX14 |  |  |  |  |  |  |  |  |
| YGL154C | LYS5 |  |  |  |  |  |  |  |  |
| YGL155W | CDC43 |  |  |  |  |  |  |  |  |
| YGL156W | AMS1 |  |  |  |  |  |  |  |  |
| YGL158W | RCK1 |  |  |  |  |  |  |  |  |
| YGL160W | AIM14 |  |  |  |  |  |  |  |  |
| YGL161C | YIP5 |  |  |  |  |  |  |  |  |
| YGL162W | SUT1 |  |  |  |  |  |  |  |  |
| YGL163C | RAD54 |  |  |  |  |  |  |  |  |
| YGL164C | YRB30 |  |  |  |  |  |  |  |  |
| YGL166W | CUP2 |  |  |  |  |  |  |  |  |
| YGL167C | PMR1 |  |  |  |  |  |  |  |  |
| YGL169W | SUA5 |  |  |  |  |  |  |  |  |
| YGL170C | SPO74 |  |  |  |  |  |  |  |  |
| YGL171W | ROK1 |  |  |  |  |  |  |  |  |
| YGL172W | NUP49 |  |  |  |  |  |  |  |  |
| YGL173C | XRN1 |  |  |  |  |  |  |  |  |
| YGL174W | BUD13 |  |  |  |  |  |  |  |  |
| YGL175C | SAE2 |  |  |  |  |  |  |  |  |
| YGL176C |  |  |  |  |  |  |  |  |  |
| YGL178W | MPT5 |  |  |  |  |  |  |  |  |
| YGL179C | TOS3 |  |  |  |  |  |  |  |  |
| YGL180W | ATG1 |  |  |  |  |  |  |  |  |
| YGL181W | GTS1 |  |  |  |  |  |  |  |  |
| YGL183C | MND1 |  |  |  |  |  |  |  |  |
| YGL184C | STR3 |  |  |  |  |  |  |  |  |
| YGL187C | COX4 |  |  |  |  |  |  |  |  |
| YGL189C | RPS26A |  |  |  |  |  |  |  |  |
| YGL190C | CDC55 |  |  |  |  |  |  |  |  |
| YGL191W | COX13 |  |  |  |  |  |  |  |  |
| YGL192W | IME4 |  |  |  |  |  |  |  |  |
| YGL194C | HOS2 |  |  |  |  |  |  |  |  |
| YGL195W | GCN1 |  |  |  |  |  |  |  |  |
| YGL197W | MDS3 |  |  |  |  |  |  |  |  |
| YGL198W | YIP4 |  |  |  |  |  |  |  |  |
| YGL200C | EMP24 |  |  |  |  |  |  |  |  |
| YGL201C | MCM6 |  |  |  |  |  |  |  |  |
| YGL202W | ARO8 |  |  |  |  |  |  |  |  |
| YGL203C | KEX1 |  |  |  |  |  |  |  |  |
| YGL205W | POX1 |  |  |  |  |  |  |  |  |
| YGL206C | CHC1 |  |  |  |  |  |  |  |  |
| YGL207W | SPT16 |  |  |  |  |  |  |  |  |
| YGL208W | SIP2 |  |  |  |  |  |  |  |  |
| YGL210W | YPT32 |  |  |  |  |  |  |  |  |
| YGL211W | NCS6 |  |  |  |  |  |  |  |  |
| YGL212W | VAM7 |  |  |  |  |  |  |  |  |
| YGL213C | SKI8 |  |  |  |  |  |  |  |  |
| YGL215W | CLG1 |  |  |  |  |  |  |  |  |
| YGL216W | KIP3 |  |  |  |  |  |  |  |  |
| YGL219C | MDM34 |  |  |  |  |  |  |  |  |
| YGL220W | BOL2 |  |  |  |  |  |  |  |  |
| YGL221C | NIF3 |  |  |  |  |  |  |  |  |
| YGL222C | EDC1 |  |  |  |  |  |  |  |  |
| YGL223C | COG1 |  |  |  |  |  |  |  |  |
| YGL224C | SDT1 |  |  |  |  |  |  |  |  |
| YGL225W | VRG4 |  |  |  |  |  |  |  |  |
| YGL226C-A | OST5 |  |  |  |  |  |  |  |  |
| YGL226W | MTC3 |  |  |  |  |  |  |  |  |
| YGL227W | VID30 |  |  |  |  |  |  |  |  |
| YGL228W | SHE10 |  |  |  |  |  |  |  |  |
| YGL229C | SAP4 |  |  |  |  |  |  |  |  |
| YGL231C | EMC4 |  |  |  |  |  |  |  |  |
| YGL232W | TAN1 |  |  |  |  |  |  |  |  |
| YGL233W | SEC15 |  |  |  |  |  |  |  |  |
| YGL234W | ADE57 |  |  |  |  |  |  |  |  |
| YGL236C | MTO1 |  |  |  |  |  |  |  |  |
| YGL237C | HAP2 |  |  |  |  |  |  |  |  |
| YGL238W | CSE1 |  |  |  |  |  |  |  |  |
| YGL240W | DOC1 |  |  |  |  |  |  |  |  |
| YGL241W | KAP114 |  |  |  |  |  |  |  |  |
| YGL242C | ANK1 |  |  |  |  |  |  |  |  |
| YGL243W | TAD1 |  |  |  |  |  |  |  |  |
| YGL244W | RTF1 |  |  |  |  |  |  |  |  |
| YGL245W | GUS1 |  |  |  |  |  |  |  |  |
| YGL246C | RAI1 |  |  |  |  |  |  |  |  |
| YGL247W | BRR6 |  |  |  |  |  |  |  |  |
| YGL248W | PDE1 |  |  |  |  |  |  |  |  |
| YGL250W | RMR1 |  |  |  |  |  |  |  |  |
| YGL252C | RTG2 |  |  |  |  |  |  |  |  |
| YGL253W | HXK2 |  |  |  |  |  |  |  |  |
| YGL255W | ZRT1 |  |  |  |  |  |  |  |  |
| YGR001C | EFM5 |  |  |  |  |  |  |  |  |
| YGR002C | SWC4 |  |  |  |  |  |  |  |  |
| YGR003W | CUL3 |  |  |  |  |  |  |  |  |
| YGR004W | PEX31 |  |  |  |  |  |  |  |  |
| YGR005C | TFG2 |  |  |  |  |  |  |  |  |
| YGR006W | PRP18 |  |  |  |  |  |  |  |  |
| YGR007W | ECT1 |  |  |  |  |  |  |  |  |
| YGR008C | STF2 |  |  |  |  |  |  |  |  |
| YGR009C | SEC9 |  |  |  |  |  |  |  |  |
| YGR010W | NMA2 |  |  |  |  |  |  |  |  |
| YGR012W | MCY1 |  |  |  |  |  |  |  |  |
| YGR013W | SNU71 |  |  |  |  |  |  |  |  |
| YGR014W | MSB2 |  |  |  |  |  |  |  |  |
| YGR017W |  |  |  |  |  |  |  |  |  |
| YGR019W | UGA1 |  |  |  |  |  |  |  |  |
| YGR020C | VMA7 |  |  |  |  |  |  |  |  |
| YGR021W | DPC29 |  |  |  |  |  |  |  |  |
| YGR023W | MTL1 |  |  |  |  |  |  |  |  |
| YGR024C | THG1 |  |  |  |  |  |  |  |  |
| YGR027C | RPS25A |  |  |  |  |  |  |  |  |
| YGR028W | MSP1 |  |  |  |  |  |  |  |  |
| YGR029W | ERV1 |  |  |  |  |  |  |  |  |
| YGR030C | POP6 |  |  |  |  |  |  |  |  |
| YGR031W | IMO32 |  |  |  |  |  |  |  |  |
| YGR032W | GSC2 |  |  |  |  |  |  |  |  |
| YGR033C | TIM21 |  |  |  |  |  |  |  |  |
| YGR034W | RPL26B |  |  |  |  |  |  |  |  |
| YGR036C | CAX4 |  |  |  |  |  |  |  |  |
| YGR037C | ACB1 |  |  |  |  |  |  |  |  |
| YGR038W | ORM1 |  |  |  |  |  |  |  |  |
| YGR040W | KSS1 |  |  |  |  |  |  |  |  |
| YGR041W | BUD9 |  |  |  |  |  |  |  |  |
| YGR042W | MTE1 |  |  |  |  |  |  |  |  |
| YGR043C | NQM1 |  |  |  |  |  |  |  |  |
| YGR046W | TAM41 |  |  |  |  |  |  |  |  |
| YGR047C | TFC4 |  |  |  |  |  |  |  |  |
| YGR048W | UFD1 |  |  |  |  |  |  |  |  |
| YGR049W | SCM4 |  |  |  |  |  |  |  |  |
| YGR054W |  |  |  |  |  |  |  |  |  |
| YGR055W | MUP1 |  |  |  |  |  |  |  |  |
| YGR056W | RSC1 |  |  |  |  |  |  |  |  |
| YGR057C | LST7 |  |  |  |  |  |  |  |  |
| YGR058W | PEF1 |  |  |  |  |  |  |  |  |
| YGR059W | SPR3 |  |  |  |  |  |  |  |  |
| YGR060W | ERG25 |  |  |  |  |  |  |  |  |
| YGR061C | ADE6 |  |  |  |  |  |  |  |  |
| YGR062C | COX18 |  |  |  |  |  |  |  |  |
| YGR063C | SPT4 |  |  |  |  |  |  |  |  |
| YGR065C | VHT1 |  |  |  |  |  |  |  |  |
| YGR068C | ART5 |  |  |  |  |  |  |  |  |
| YGR070W | ROM1 |  |  |  |  |  |  |  |  |
| YGR071C | ENV11 |  |  |  |  |  |  |  |  |
| YGR072W | UPF3 |  |  |  |  |  |  |  |  |
| YGR074W | SMD1 |  |  |  |  |  |  |  |  |
| YGR075C | PRP38 |  |  |  |  |  |  |  |  |
| YGR076C | MRPL25 |  |  |  |  |  |  |  |  |
| YGR077C | PEX8 |  |  |  |  |  |  |  |  |
| YGR078C | PAC10 |  |  |  |  |  |  |  |  |
| YGR079W |  |  |  |  |  |  |  |  |  |
| YGR080W | TWF1 |  |  |  |  |  |  |  |  |
| YGR081C | SLX9 |  |  |  |  |  |  |  |  |
| YGR082W | TOM20 |  |  |  |  |  |  |  |  |
| YGR083C | GCD2 |  |  |  |  |  |  |  |  |
| YGR084C | MRP13 |  |  |  |  |  |  |  |  |
| YGR085C | RPL11B |  |  |  |  |  |  |  |  |
| YGR086C | PIL1 |  |  |  |  |  |  |  |  |
| YGR087C | PDC6 |  |  |  |  |  |  |  |  |
| YGR088W | CTT1 |  |  |  |  |  |  |  |  |
| YGR089W | NNF2 |  |  |  |  |  |  |  |  |
| YGR090W | UTP22 |  |  |  |  |  |  |  |  |
| YGR091W | PRP31 |  |  |  |  |  |  |  |  |
| YGR092W | DBF2 |  |  |  |  |  |  |  |  |
| YGR093W | DRN1 |  |  |  |  |  |  |  |  |
| YGR094W | VAS1 |  |  |  |  |  |  |  |  |
| YGR095C | RRP46 |  |  |  |  |  |  |  |  |
| YGR096W | TPC1 |  |  |  |  |  |  |  |  |
| YGR097W | ASK10 |  |  |  |  |  |  |  |  |
| YGR098C | ESP1 |  |  |  |  |  |  |  |  |
| YGR099W | TEL2 |  |  |  |  |  |  |  |  |
| YGR100W | MDR1 |  |  |  |  |  |  |  |  |
| YGR101W | PCP1 |  |  |  |  |  |  |  |  |
| YGR102C | GTF1 |  |  |  |  |  |  |  |  |
| YGR103W | NOP7 |  |  |  |  |  |  |  |  |
| YGR104C | SRB5 |  |  |  |  |  |  |  |  |
| YGR105W | VMA21 |  |  |  |  |  |  |  |  |
| YGR106C | VOA1 |  |  |  |  |  |  |  |  |
| YGR108W | CLB1 |  |  |  |  |  |  |  |  |
| YGR109C | CLB6 |  |  |  |  |  |  |  |  |
| YGR110W | CLD1 |  |  |  |  |  |  |  |  |
| YGR112W | SHY1 |  |  |  |  |  |  |  |  |
| YGR113W | DAM1 |  |  |  |  |  |  |  |  |
| YGR116W | SPT6 |  |  |  |  |  |  |  |  |
| YGR118W | RPS23A |  |  |  |  |  |  |  |  |
| YGR119C | NUP57 |  |  |  |  |  |  |  |  |
| YGR120C | COG2 |  |  |  |  |  |  |  |  |
| YGR121C | MEP1 |  |  |  |  |  |  |  |  |
| YGR122W |  |  |  |  |  |  |  |  |  |
| YGR123C | PPT1 |  |  |  |  |  |  |  |  |
| YGR124W | ASN2 |  |  |  |  |  |  |  |  |
| YGR125W | VSB1 |  |  |  |  |  |  |  |  |
| YGR127W |  |  |  |  |  |  |  |  |  |
| YGR128C | UTP8 |  |  |  |  |  |  |  |  |
| YGR129W | SYF2 |  |  |  |  |  |  |  |  |
| YGR130C |  |  |  |  |  |  |  |  |  |
| YGR131W | FHN1 |  |  |  |  |  |  |  |  |
| YGR132C | PHB1 |  |  |  |  |  |  |  |  |
| YGR133W | PEX4 |  |  |  |  |  |  |  |  |
| YGR134W | CAF130 |  |  |  |  |  |  |  |  |
| YGR135W | PRE9 |  |  |  |  |  |  |  |  |
| YGR136W | LSB1 |  |  |  |  |  |  |  |  |
| YGR138C | TPO2 |  |  |  |  |  |  |  |  |
| YGR140W | CBF2 |  |  |  |  |  |  |  |  |
| YGR141W | VPS62 |  |  |  |  |  |  |  |  |
| YGR142W | BTN2 |  |  |  |  |  |  |  |  |
| YGR143W | SKN1 |  |  |  |  |  |  |  |  |
| YGR144W | THI4 |  |  |  |  |  |  |  |  |
| YGR145W | ENP2 |  |  |  |  |  |  |  |  |
| YGR146C | ECL1 |  |  |  |  |  |  |  |  |
| YGR147C | NAT2 |  |  |  |  |  |  |  |  |
| YGR148C | RPL24B |  |  |  |  |  |  |  |  |
| YGR149W | GPC1 |  |  |  |  |  |  |  |  |
| YGR150C | CCM1 |  |  |  |  |  |  |  |  |
| YGR152C | RSR1 |  |  |  |  |  |  |  |  |
| YGR155W | CYS4 |  |  |  |  |  |  |  |  |
| YGR156W | PTI1 |  |  |  |  |  |  |  |  |
| YGR157W | CHO2 |  |  |  |  |  |  |  |  |
| YGR158C | MTR3 |  |  |  |  |  |  |  |  |
| YGR159C | NSR1 |  |  |  |  |  |  |  |  |
| YGR162W | TIF4631 |  |  |  |  |  |  |  |  |
| YGR163W | GTR2 |  |  |  |  |  |  |  |  |
| YGR165W | MRPS35 |  |  |  |  |  |  |  |  |
| YGR166W | TRS65 |  |  |  |  |  |  |  |  |
| YGR167W | CLC1 |  |  |  |  |  |  |  |  |
| YGR168C | PEX35 |  |  |  |  |  |  |  |  |
| YGR169C | PUS6 |  |  |  |  |  |  |  |  |
| YGR169C-A | LSO2 |  |  |  |  |  |  |  |  |
| YGR170W | PSD2 |  |  |  |  |  |  |  |  |
| YGR171C | MSM1 |  |  |  |  |  |  |  |  |
| YGR172C | YIP1 |  |  |  |  |  |  |  |  |
| YGR173W | RBG2 |  |  |  |  |  |  |  |  |
| YGR174C | CBP4 |  |  |  |  |  |  |  |  |
| YGR175C | ERG1 |  |  |  |  |  |  |  |  |
| YGR178C | PBP1 |  |  |  |  |  |  |  |  |
| YGR179C | OKP1 |  |  |  |  |  |  |  |  |
| YGR180C | RNR4 |  |  |  |  |  |  |  |  |
| YGR181W | TIM13 |  |  |  |  |  |  |  |  |
| YGR183C | QCR9 |  |  |  |  |  |  |  |  |
| YGR184C | UBR1 |  |  |  |  |  |  |  |  |
| YGR185C | TYS1 |  |  |  |  |  |  |  |  |
| YGR186W | TFG1 |  |  |  |  |  |  |  |  |
| YGR187C | HGH1 |  |  |  |  |  |  |  |  |
| YGR188C | BUB1 |  |  |  |  |  |  |  |  |
| YGR189C | CRH1 |  |  |  |  |  |  |  |  |
| YGR191W | HIP1 |  |  |  |  |  |  |  |  |
| YGR193C | PDX1 |  |  |  |  |  |  |  |  |
| YGR194C | XKS1 |  |  |  |  |  |  |  |  |
| YGR195W | SKI6 |  |  |  |  |  |  |  |  |
| YGR196C | FYV8 |  |  |  |  |  |  |  |  |
| YGR197C | SNG1 |  |  |  |  |  |  |  |  |
| YGR198W | YPP1 |  |  |  |  |  |  |  |  |
| YGR200C | ELP2 |  |  |  |  |  |  |  |  |
| YGR202C | PCT1 |  |  |  |  |  |  |  |  |
| YGR203W | YCH1 |  |  |  |  |  |  |  |  |
| YGR204W | ADE3 |  |  |  |  |  |  |  |  |
| YGR205W | TDA10 |  |  |  |  |  |  |  |  |
| YGR206W | MVB12 |  |  |  |  |  |  |  |  |
| YGR207C | CIR1 |  |  |  |  |  |  |  |  |
| YGR208W | SER2 |  |  |  |  |  |  |  |  |
| YGR209C | TRX2 |  |  |  |  |  |  |  |  |
| YGR211W | ZPR1 |  |  |  |  |  |  |  |  |
| YGR212W | SLI1 |  |  |  |  |  |  |  |  |
| YGR214W | RPS0A |  |  |  |  |  |  |  |  |
| YGR215W | RSM27 |  |  |  |  |  |  |  |  |
| YGR216C | GPI1 |  |  |  |  |  |  |  |  |
| YGR217W | CCH1 |  |  |  |  |  |  |  |  |
| YGR218W | CRM1 |  |  |  |  |  |  |  |  |
| YGR220C | MRPL9 |  |  |  |  |  |  |  |  |
| YGR221C | TOS2 |  |  |  |  |  |  |  |  |
| YGR222W | PET54 |  |  |  |  |  |  |  |  |
| YGR223C | HSV2 |  |  |  |  |  |  |  |  |
| YGR225W | AMA1 |  |  |  |  |  |  |  |  |
| YGR227W | DIE2 |  |  |  |  |  |  |  |  |
| YGR229C | SMI1 |  |  |  |  |  |  |  |  |
| YGR230W | BNS1 |  |  |  |  |  |  |  |  |
| YGR231C | PHB2 |  |  |  |  |  |  |  |  |
| YGR232W | NAS6 |  |  |  |  |  |  |  |  |
| YGR233C | PHO81 |  |  |  |  |  |  |  |  |
| YGR235C | MIC26 |  |  |  |  |  |  |  |  |
| YGR236C | SPG1 |  |  |  |  |  |  |  |  |
| YGR238C | KEL2 |  |  |  |  |  |  |  |  |
| YGR239C | PEX21 |  |  |  |  |  |  |  |  |
| YGR240C | PFK1 |  |  |  |  |  |  |  |  |
| YGR241C | YAP1802 |  |  |  |  |  |  |  |  |
| YGR243W | MPC3 |  |  |  |  |  |  |  |  |
| YGR244C | LSC2 |  |  |  |  |  |  |  |  |
| YGR245C | SDA1 |  |  |  |  |  |  |  |  |
| YGR246C | BRF1 |  |  |  |  |  |  |  |  |
| YGR247W | CPD1 |  |  |  |  |  |  |  |  |
| YGR248W | SOL4 |  |  |  |  |  |  |  |  |
| YGR249W | MGA1 |  |  |  |  |  |  |  |  |
| YGR250C | RIE1 |  |  |  |  |  |  |  |  |
| YGR251W | NOP19 |  |  |  |  |  |  |  |  |
| YGR252W | GCN5 |  |  |  |  |  |  |  |  |
| YGR253C | PUP2 |  |  |  |  |  |  |  |  |
| YGR254W | ENO1 |  |  |  |  |  |  |  |  |
| YGR255C | COQ6 |  |  |  |  |  |  |  |  |
| YGR256W | GND2 |  |  |  |  |  |  |  |  |
| YGR257C | MTM1 |  |  |  |  |  |  |  |  |
| YGR258C | RAD2 |  |  |  |  |  |  |  |  |
| YGR260W | TNA1 |  |  |  |  |  |  |  |  |
| YGR261C | APL6 |  |  |  |  |  |  |  |  |
| YGR262C | BUD32 |  |  |  |  |  |  |  |  |
| YGR263C | SAY1 |  |  |  |  |  |  |  |  |
| YGR264C | MES1 |  |  |  |  |  |  |  |  |
| YGR267C | FOL2 |  |  |  |  |  |  |  |  |
| YGR268C | HUA1 |  |  |  |  |  |  |  |  |
| YGR270W | YTA7 |  |  |  |  |  |  |  |  |
| YGR271C-A | EFG1 |  |  |  |  |  |  |  |  |
| YGR271W | SLH1 |  |  |  |  |  |  |  |  |
| YGR274C | TAF1 |  |  |  |  |  |  |  |  |
| YGR275W | RTT102 |  |  |  |  |  |  |  |  |
| YGR276C | RNH70 |  |  |  |  |  |  |  |  |
| YGR277C | CAB4 |  |  |  |  |  |  |  |  |
| YGR278W | CWC22 |  |  |  |  |  |  |  |  |
| YGR279C | SCW4 |  |  |  |  |  |  |  |  |
| YGR280C | PXR1 |  |  |  |  |  |  |  |  |
| YGR281W | YOR1 |  |  |  |  |  |  |  |  |
| YGR282C | BGL2 |  |  |  |  |  |  |  |  |
| YGR283C | UPA1 |  |  |  |  |  |  |  |  |
| YGR284C | ERV29 |  |  |  |  |  |  |  |  |
| YGR285C | ZUO1 |  |  |  |  |  |  |  |  |
| YHL001W | RPL14B |  |  |  |  |  |  |  |  |
| YHL002W | HSE1 |  |  |  |  |  |  |  |  |
| YHL003C | LAG1 |  |  |  |  |  |  |  |  |
| YHL004W | MRP4 |  |  |  |  |  |  |  |  |
| YHL006C | SHU1 |  |  |  |  |  |  |  |  |
| YHL007C | STE20 |  |  |  |  |  |  |  |  |
| YHL009C | YAP3 |  |  |  |  |  |  |  |  |
| YHL010C | ETP1 |  |  |  |  |  |  |  |  |
| YHL011C | PRS3 |  |  |  |  |  |  |  |  |
| YHL013C | OTU2 |  |  |  |  |  |  |  |  |
| YHL014C | YLF2 |  |  |  |  |  |  |  |  |
| YHL015W | RPS20 |  |  |  |  |  |  |  |  |
| YHL016C | DUR3 |  |  |  |  |  |  |  |  |
| YHL019C | APM2 |  |  |  |  |  |  |  |  |
| YHL020C | OPI1 |  |  |  |  |  |  |  |  |
| YHL021C | AIM17 |  |  |  |  |  |  |  |  |
| YHL022C | SPO11 |  |  |  |  |  |  |  |  |
| YHL023C | NPR3 |  |  |  |  |  |  |  |  |
| YHL024W | RIM4 |  |  |  |  |  |  |  |  |
| YHL025W | SNF6 |  |  |  |  |  |  |  |  |
| YHL027W | RIM101 |  |  |  |  |  |  |  |  |
| YHL028W | WSC4 |  |  |  |  |  |  |  |  |
| YHL030W | ECM29 |  |  |  |  |  |  |  |  |
| YHL031C | GOS1 |  |  |  |  |  |  |  |  |
| YHL032C | GUT1 |  |  |  |  |  |  |  |  |
| YHL033C | RPL8A |  |  |  |  |  |  |  |  |
| YHL034C | SBP1 |  |  |  |  |  |  |  |  |
| YHL035C | VMR1 |  |  |  |  |  |  |  |  |
| YHL036W | MUP3 |  |  |  |  |  |  |  |  |
| YHL039W | EFM1 |  |  |  |  |  |  |  |  |
| YHR001W | OSH7 |  |  |  |  |  |  |  |  |
| YHR001W-A | QCR10 |  |  |  |  |  |  |  |  |
| YHR002W | LEU5 |  |  |  |  |  |  |  |  |
| YHR003C | TCD1 |  |  |  |  |  |  |  |  |
| YHR004C | NEM1 |  |  |  |  |  |  |  |  |
| YHR005C | GPA1 |  |  |  |  |  |  |  |  |
| YHR005C-A | TIM10 |  |  |  |  |  |  |  |  |
| YHR006W | STP2 |  |  |  |  |  |  |  |  |
| YHR007C | ERG11 |  |  |  |  |  |  |  |  |
| YHR008C | SOD2 |  |  |  |  |  |  |  |  |
| YHR009C | TDA3 |  |  |  |  |  |  |  |  |
| YHR010W | RPL27A |  |  |  |  |  |  |  |  |
| YHR011W | DIA4 |  |  |  |  |  |  |  |  |
| YHR012W | VPS29 |  |  |  |  |  |  |  |  |
| YHR013C | ARD1 |  |  |  |  |  |  |  |  |
| YHR014W | SPO13 |  |  |  |  |  |  |  |  |
| YHR015W | MIP6 |  |  |  |  |  |  |  |  |
| YHR016C | YSC84 |  |  |  |  |  |  |  |  |
| YHR017W | YSC83 |  |  |  |  |  |  |  |  |
| YHR018C | ARG4 |  |  |  |  |  |  |  |  |
| YHR019C | DED81 |  |  |  |  |  |  |  |  |
| YHR020W |  |  |  |  |  |  |  |  |  |
| YHR021C | RPS27B |  |  |  |  |  |  |  |  |
| YHR023W | MYO1 |  |  |  |  |  |  |  |  |
| YHR024C | MAS2 |  |  |  |  |  |  |  |  |
| YHR025W | THR1 |  |  |  |  |  |  |  |  |
| YHR026W | VMA16 |  |  |  |  |  |  |  |  |
| YHR027C | RPN1 |  |  |  |  |  |  |  |  |
| YHR028C | DAP2 |  |  |  |  |  |  |  |  |
| YHR030C | SLT2 |  |  |  |  |  |  |  |  |
| YHR031C | RRM3 |  |  |  |  |  |  |  |  |
| YHR032W | ERC1 |  |  |  |  |  |  |  |  |
| YHR034C | PIH1 |  |  |  |  |  |  |  |  |
| YHR035W | NEL1 |  |  |  |  |  |  |  |  |
| YHR036W | BRL1 |  |  |  |  |  |  |  |  |
| YHR037W | PUT2 |  |  |  |  |  |  |  |  |
| YHR038W | RRF1 |  |  |  |  |  |  |  |  |
| YHR039C | MSC7 |  |  |  |  |  |  |  |  |
| YHR039C-A | VMA10 |  |  |  |  |  |  |  |  |
| YHR040W | BCD1 |  |  |  |  |  |  |  |  |
| YHR041C | SRB2 |  |  |  |  |  |  |  |  |
| YHR042W | NCP1 |  |  |  |  |  |  |  |  |
| YHR043C_044C | DOG2_DOG1 |  |  |  |  |  |  |  |  |
| YHR045W | DDE1 |  |  |  |  |  |  |  |  |
| YHR047C | AAP1 |  |  |  |  |  |  |  |  |
| YHR049W | FSH1 |  |  |  |  |  |  |  |  |
| YHR050W | SMF2 |  |  |  |  |  |  |  |  |
| YHR051W | COX6 |  |  |  |  |  |  |  |  |
| YHR052W | CIC1 |  |  |  |  |  |  |  |  |
| YHR056C | RSC30 |  |  |  |  |  |  |  |  |
| YHR057C | CPR2 |  |  |  |  |  |  |  |  |
| YHR058C | MED6 |  |  |  |  |  |  |  |  |
| YHR059W | FYV4 |  |  |  |  |  |  |  |  |
| YHR060W | VMA22 |  |  |  |  |  |  |  |  |
| YHR061C | GIC1 |  |  |  |  |  |  |  |  |
| YHR062C | RPP1 |  |  |  |  |  |  |  |  |
| YHR063C | PAN5 |  |  |  |  |  |  |  |  |
| YHR064C | SSZ1 |  |  |  |  |  |  |  |  |
| YHR065C | RRP3 |  |  |  |  |  |  |  |  |
| YHR066W | SSF1 |  |  |  |  |  |  |  |  |
| YHR067W | HTD2 |  |  |  |  |  |  |  |  |
| YHR068W | DYS1 |  |  |  |  |  |  |  |  |
| YHR069C | RRP4 |  |  |  |  |  |  |  |  |
| YHR070W | TRM5 |  |  |  |  |  |  |  |  |
| YHR071W | PCL5 |  |  |  |  |  |  |  |  |
| YHR072W | ERG7 |  |  |  |  |  |  |  |  |
| YHR072W-A | NOP10 |  |  |  |  |  |  |  |  |
| YHR073W | OSH3 |  |  |  |  |  |  |  |  |
| YHR074W | QNS1 |  |  |  |  |  |  |  |  |
| YHR075C | PPE1 |  |  |  |  |  |  |  |  |
| YHR076W | PTC7 |  |  |  |  |  |  |  |  |
| YHR077C | NMD2 |  |  |  |  |  |  |  |  |
| YHR079C | IRE1 |  |  |  |  |  |  |  |  |
| YHR079C-A | SAE3 |  |  |  |  |  |  |  |  |
| YHR080C | LAM4 |  |  |  |  |  |  |  |  |
| YHR081W | LRP1 |  |  |  |  |  |  |  |  |
| YHR082C | KSP1 |  |  |  |  |  |  |  |  |
| YHR083W | SAM35 |  |  |  |  |  |  |  |  |
| YHR084W | STE12 |  |  |  |  |  |  |  |  |
| YHR085W | IPI1 |  |  |  |  |  |  |  |  |
| YHR086W | NAM8 |  |  |  |  |  |  |  |  |
| YHR088W | RPF1 |  |  |  |  |  |  |  |  |
| YHR089C | GAR1 |  |  |  |  |  |  |  |  |
| YHR090C | YNG2 |  |  |  |  |  |  |  |  |
| YHR091C | MSR1 |  |  |  |  |  |  |  |  |
| YHR092C | HXT4 |  |  |  |  |  |  |  |  |
| YHR094C | HXT1 |  |  |  |  |  |  |  |  |
| YHR096C | HXT5 |  |  |  |  |  |  |  |  |
| YHR097C | PAL2 |  |  |  |  |  |  |  |  |
| YHR098C | SFB3 |  |  |  |  |  |  |  |  |
| YHR099W | TRA1 |  |  |  |  |  |  |  |  |
| YHR100C | GEP4 |  |  |  |  |  |  |  |  |
| YHR101C | BIG1 |  |  |  |  |  |  |  |  |
| YHR102W | KIC1 |  |  |  |  |  |  |  |  |
| YHR103W | SBE22 |  |  |  |  |  |  |  |  |
| YHR104W | GRE3 |  |  |  |  |  |  |  |  |
| YHR105W | YPT35 |  |  |  |  |  |  |  |  |
| YHR106W | TRR2 |  |  |  |  |  |  |  |  |
| YHR107C | CDC12 |  |  |  |  |  |  |  |  |
| YHR108W | GGA2 |  |  |  |  |  |  |  |  |
| YHR109W | CTM1 |  |  |  |  |  |  |  |  |
| YHR110W | ERP5 |  |  |  |  |  |  |  |  |
| YHR111W | UBA4 |  |  |  |  |  |  |  |  |
| YHR112C |  |  |  |  |  |  |  |  |  |
| YHR114W | BZZ1 |  |  |  |  |  |  |  |  |
| YHR115C | DMA1 |  |  |  |  |  |  |  |  |
| YHR116W | COX23 |  |  |  |  |  |  |  |  |
| YHR117W | TOM71 |  |  |  |  |  |  |  |  |
| YHR118C | ORC6 |  |  |  |  |  |  |  |  |
| YHR119W | SET1 |  |  |  |  |  |  |  |  |
| YHR120W | MSH1 |  |  |  |  |  |  |  |  |
| YHR121W | LSM12 |  |  |  |  |  |  |  |  |
| YHR122W | CIA2 |  |  |  |  |  |  |  |  |
| YHR123W | EPT1 |  |  |  |  |  |  |  |  |
| YHR124W | NDT80 |  |  |  |  |  |  |  |  |
| YHR127W |  |  |  |  |  |  |  |  |  |
| YHR128W | FUR1 |  |  |  |  |  |  |  |  |
| YHR129C | ARP1 |  |  |  |  |  |  |  |  |
| YHR132C | ECM14 |  |  |  |  |  |  |  |  |
| YHR132W-A | IGO2 |  |  |  |  |  |  |  |  |
| YHR133C | NSG1 |  |  |  |  |  |  |  |  |
| YHR134W | WSS1 |  |  |  |  |  |  |  |  |
| YHR135C | YCK1 |  |  |  |  |  |  |  |  |
| YHR138C |  |  |  |  |  |  |  |  |  |
| YHR141C | RPL42B |  |  |  |  |  |  |  |  |
| YHR142W | CHS7 |  |  |  |  |  |  |  |  |
| YHR143W | DSE2 |  |  |  |  |  |  |  |  |
| YHR143W-A | RPC10 |  |  |  |  |  |  |  |  |
| YHR144C | DCD1 |  |  |  |  |  |  |  |  |
| YHR146W | CRP1 |  |  |  |  |  |  |  |  |
| YHR147C | MRPL6 |  |  |  |  |  |  |  |  |
| YHR148W | IMP3 |  |  |  |  |  |  |  |  |
| YHR149C | SKG6 |  |  |  |  |  |  |  |  |
| YHR150W | PEX28 |  |  |  |  |  |  |  |  |
| YHR151C | MTC6 |  |  |  |  |  |  |  |  |
| YHR152W | SPO12 |  |  |  |  |  |  |  |  |
| YHR154W | RTT107 |  |  |  |  |  |  |  |  |
| YHR155W | LAM1 |  |  |  |  |  |  |  |  |
| YHR156C | LIN1 |  |  |  |  |  |  |  |  |
| YHR158C | KEL1 |  |  |  |  |  |  |  |  |
| YHR160C | PEX18 |  |  |  |  |  |  |  |  |
| YHR161C | YAP1801 |  |  |  |  |  |  |  |  |
| YHR162W | MPC2 |  |  |  |  |  |  |  |  |
| YHR163W | SOL3 |  |  |  |  |  |  |  |  |
| YHR164C | DNA2 |  |  |  |  |  |  |  |  |
| YHR165C | PRP8 |  |  |  |  |  |  |  |  |
| YHR166C | CDC23 |  |  |  |  |  |  |  |  |
| YHR167W | THP2 |  |  |  |  |  |  |  |  |
| YHR168W | MTG2 |  |  |  |  |  |  |  |  |
| YHR169W | DBP8 |  |  |  |  |  |  |  |  |
| YHR170W | NMD3 |  |  |  |  |  |  |  |  |
| YHR171W | ATG7 |  |  |  |  |  |  |  |  |
| YHR172W | SPC97 |  |  |  |  |  |  |  |  |
| YHR174W | ENO2 |  |  |  |  |  |  |  |  |
| YHR175W | CTR2 |  |  |  |  |  |  |  |  |
| YHR176W | FMO1 |  |  |  |  |  |  |  |  |
| YHR178W | STB5 |  |  |  |  |  |  |  |  |
| YHR179W | OYE2 |  |  |  |  |  |  |  |  |
| YHR181W | SVP26 |  |  |  |  |  |  |  |  |
| YHR182W | RGD3 |  |  |  |  |  |  |  |  |
| YHR183W | GND1 |  |  |  |  |  |  |  |  |
| YHR184W | SSP1 |  |  |  |  |  |  |  |  |
| YHR185C | PFS1 |  |  |  |  |  |  |  |  |
| YHR186C | KOG1 |  |  |  |  |  |  |  |  |
| YHR187W | IKI1 |  |  |  |  |  |  |  |  |
| YHR188C | GPI16 |  |  |  |  |  |  |  |  |
| YHR189W | PTH1 |  |  |  |  |  |  |  |  |
| YHR190W | ERG9 |  |  |  |  |  |  |  |  |
| YHR191C | CTF8 |  |  |  |  |  |  |  |  |
| YHR192W | LNP1 |  |  |  |  |  |  |  |  |
| YHR193C | EGD2 |  |  |  |  |  |  |  |  |
| YHR194W | MDM31 |  |  |  |  |  |  |  |  |
| YHR196W | UTP9 |  |  |  |  |  |  |  |  |
| YHR197W | RIX1 |  |  |  |  |  |  |  |  |
| YHR199C | AIM46 |  |  |  |  |  |  |  |  |
| YHR200W | RPN10 |  |  |  |  |  |  |  |  |
| YHR201C | PPX1 |  |  |  |  |  |  |  |  |
| YHR202W | SMN1 |  |  |  |  |  |  |  |  |
| YHR203C | RPS4B |  |  |  |  |  |  |  |  |
| YHR204W | MNL1 |  |  |  |  |  |  |  |  |
| YHR205W | SCH9 |  |  |  |  |  |  |  |  |
| YHR206W | SKN7 |  |  |  |  |  |  |  |  |
| YHR207C | SET5 |  |  |  |  |  |  |  |  |
| YHR208W | BAT1 |  |  |  |  |  |  |  |  |
| YHR211W | FLO5 |  |  |  |  |  |  |  |  |
| YIL002C | INP51 |  |  |  |  |  |  |  |  |
| YIL003W | CFD1 |  |  |  |  |  |  |  |  |
| YIL004C | BET1 |  |  |  |  |  |  |  |  |
| YIL005W | EPS1 |  |  |  |  |  |  |  |  |
| YIL006W | YIA6 |  |  |  |  |  |  |  |  |
| YIL007C | NAS2 |  |  |  |  |  |  |  |  |
| YIL008W | URM1 |  |  |  |  |  |  |  |  |
| YIL009C-A | EST3 |  |  |  |  |  |  |  |  |
| YIL010W | DOT5 |  |  |  |  |  |  |  |  |
| YIL011W | TIR3 |  |  |  |  |  |  |  |  |
| YIL013C | PDR11 |  |  |  |  |  |  |  |  |
| YIL014W | MNT3 |  |  |  |  |  |  |  |  |
| YIL015W | BAR1 |  |  |  |  |  |  |  |  |
| YIL016W | SNL1 |  |  |  |  |  |  |  |  |
| YIL017C | VID28 |  |  |  |  |  |  |  |  |
| YIL018W | RPL2B |  |  |  |  |  |  |  |  |
| YIL019W | FAF1 |  |  |  |  |  |  |  |  |
| YIL020C | HIS6 |  |  |  |  |  |  |  |  |
| YIL021W | RPB3 |  |  |  |  |  |  |  |  |
| YIL022W | TIM44 |  |  |  |  |  |  |  |  |
| YIL023C | YKE4 |  |  |  |  |  |  |  |  |
| YIL024C | ATG45 |  |  |  |  |  |  |  |  |
| YIL026C | IRR1 |  |  |  |  |  |  |  |  |
| YIL027C | EMC5 |  |  |  |  |  |  |  |  |
| YIL030C | SSM4 |  |  |  |  |  |  |  |  |
| YIL031W | ULP2 |  |  |  |  |  |  |  |  |
| YIL033C | BCY1 |  |  |  |  |  |  |  |  |
| YIL034C | CAP2 |  |  |  |  |  |  |  |  |
| YIL035C | CKA1 |  |  |  |  |  |  |  |  |
| YIL036W | CST6 |  |  |  |  |  |  |  |  |
| YIL037C | PRM2 |  |  |  |  |  |  |  |  |
| YIL038C | NOT3 |  |  |  |  |  |  |  |  |
| YIL039W | TED1 |  |  |  |  |  |  |  |  |
| YIL040W | APQ12 |  |  |  |  |  |  |  |  |
| YIL041W | GVP36 |  |  |  |  |  |  |  |  |
| YIL042C | PKP1 |  |  |  |  |  |  |  |  |
| YIL043C | CBR1 |  |  |  |  |  |  |  |  |
| YIL044C | AGE2 |  |  |  |  |  |  |  |  |
| YIL045W | PIG2 |  |  |  |  |  |  |  |  |
| YIL046W | MET30 |  |  |  |  |  |  |  |  |
| YIL047C | SYG1 |  |  |  |  |  |  |  |  |
| YIL048W | NEO1 |  |  |  |  |  |  |  |  |
| YIL049W | DFG10 |  |  |  |  |  |  |  |  |
| YIL050W | PCL7 |  |  |  |  |  |  |  |  |
| YIL051C | MMF1 |  |  |  |  |  |  |  |  |
| YIL052C | RPL34B |  |  |  |  |  |  |  |  |
| YIL053W | GPP1 |  |  |  |  |  |  |  |  |
| YIL056W | VHR1 |  |  |  |  |  |  |  |  |
| YIL057C | RGI2 |  |  |  |  |  |  |  |  |
| YIL061C | SNP1 |  |  |  |  |  |  |  |  |
| YIL062C | ARC15 |  |  |  |  |  |  |  |  |
| YIL063C | YRB2 |  |  |  |  |  |  |  |  |
| YIL064W | EFM4 |  |  |  |  |  |  |  |  |
| YIL065C | FIS1 |  |  |  |  |  |  |  |  |
| YIL066C | RNR3 |  |  |  |  |  |  |  |  |
| YIL068C | SEC6 |  |  |  |  |  |  |  |  |
| YIL069C | RPS24B |  |  |  |  |  |  |  |  |
| YIL070C | MAM33 |  |  |  |  |  |  |  |  |
| YIL071C | PCI8 |  |  |  |  |  |  |  |  |
| YIL072W | HOP1 |  |  |  |  |  |  |  |  |
| YIL074C | SER33 |  |  |  |  |  |  |  |  |
| YIL075C | RPN2 |  |  |  |  |  |  |  |  |
| YIL076W | SEC28 |  |  |  |  |  |  |  |  |
| YIL078W | THS1 |  |  |  |  |  |  |  |  |
| YIL079C | AIR1 |  |  |  |  |  |  |  |  |
| YIL083C | CAB2 |  |  |  |  |  |  |  |  |
| YIL084C | SDS3 |  |  |  |  |  |  |  |  |
| YIL087C | AIM19 |  |  |  |  |  |  |  |  |
| YIL088C | AVT7 |  |  |  |  |  |  |  |  |
| YIL090W | ICE2 |  |  |  |  |  |  |  |  |
| YIL091C | UTP25 |  |  |  |  |  |  |  |  |
| YIL093C | RSM25 |  |  |  |  |  |  |  |  |
| YIL094C | LYS12 |  |  |  |  |  |  |  |  |
| YIL095W | PRK1 |  |  |  |  |  |  |  |  |
| YIL096C | BMT5 |  |  |  |  |  |  |  |  |
| YIL097W | FYV10 |  |  |  |  |  |  |  |  |
| YIL098C | FMC1 |  |  |  |  |  |  |  |  |
| YIL099W | SGA1 |  |  |  |  |  |  |  |  |
| YIL101C | XBP1 |  |  |  |  |  |  |  |  |
| YIL103W | DPH1 |  |  |  |  |  |  |  |  |
| YIL104C | SHQ1 |  |  |  |  |  |  |  |  |
| YIL105C | SLM1 |  |  |  |  |  |  |  |  |
| YIL106W | MOB1 |  |  |  |  |  |  |  |  |
| YIL107C | PFK26 |  |  |  |  |  |  |  |  |
| YIL108W |  |  |  |  |  |  |  |  |  |
| YIL109C | SEC24 |  |  |  |  |  |  |  |  |
| YIL110W | HPM1 |  |  |  |  |  |  |  |  |
| YIL111W | COX5B |  |  |  |  |  |  |  |  |
| YIL112W | HOS4 |  |  |  |  |  |  |  |  |
| YIL113W | SDP1 |  |  |  |  |  |  |  |  |
| YIL114C | POR2 |  |  |  |  |  |  |  |  |
| YIL115C | NUP159 |  |  |  |  |  |  |  |  |
| YIL116W | HIS5 |  |  |  |  |  |  |  |  |
| YIL117C | PRM5 |  |  |  |  |  |  |  |  |
| YIL118W | RHO3 |  |  |  |  |  |  |  |  |
| YIL119C | RPI1 |  |  |  |  |  |  |  |  |
| YIL120W | QDR1 |  |  |  |  |  |  |  |  |
| YIL121W | QDR2 |  |  |  |  |  |  |  |  |
| YIL122W | POG1 |  |  |  |  |  |  |  |  |
| YIL123W | SIM1 |  |  |  |  |  |  |  |  |
| YIL124W | AYR1 |  |  |  |  |  |  |  |  |
| YIL125W | KGD1 |  |  |  |  |  |  |  |  |
| YIL126W | STH1 |  |  |  |  |  |  |  |  |
| YIL128W | MET18 |  |  |  |  |  |  |  |  |
| YIL129C | TAO3 |  |  |  |  |  |  |  |  |
| YIL130W | ASG1 |  |  |  |  |  |  |  |  |
| YIL131C | FKH1 |  |  |  |  |  |  |  |  |
| YIL132C | CSM2 |  |  |  |  |  |  |  |  |
| YIL133C | RPL16A |  |  |  |  |  |  |  |  |
| YIL134W | FLX1 |  |  |  |  |  |  |  |  |
| YIL135C | VHS2 |  |  |  |  |  |  |  |  |
| YIL136W | OM45 |  |  |  |  |  |  |  |  |
| YIL137C | TMA108 |  |  |  |  |  |  |  |  |
| YIL138C | TPM2 |  |  |  |  |  |  |  |  |
| YIL139C | REV7 |  |  |  |  |  |  |  |  |
| YIL140W | AXL2 |  |  |  |  |  |  |  |  |
| YIL142W | CCT2 |  |  |  |  |  |  |  |  |
| YIL143C | SSL2 |  |  |  |  |  |  |  |  |
| YIL144W | NDC80 |  |  |  |  |  |  |  |  |
| YIL145C | PAN6 |  |  |  |  |  |  |  |  |
| YIL146C | ATG32 |  |  |  |  |  |  |  |  |
| YIL147C | SLN1 |  |  |  |  |  |  |  |  |
| YIL148W | RPL40A |  |  |  |  |  |  |  |  |
| YIL149C | MLP2 |  |  |  |  |  |  |  |  |
| YIL150C | MCM10 |  |  |  |  |  |  |  |  |
| YIL151C | ESL1 |  |  |  |  |  |  |  |  |
| YIL153W | RRD1 |  |  |  |  |  |  |  |  |
| YIL154C | IMP21 |  |  |  |  |  |  |  |  |
| YIL155C | GUT2 |  |  |  |  |  |  |  |  |
| YIL156W | UBP7 |  |  |  |  |  |  |  |  |
| YIL156W-B | ATG44 |  |  |  |  |  |  |  |  |
| YIL157C | COA1 |  |  |  |  |  |  |  |  |
| YIL158W | AIM20 |  |  |  |  |  |  |  |  |
| YIL159W | BNR1 |  |  |  |  |  |  |  |  |
| YIL160C | POT1 |  |  |  |  |  |  |  |  |
| YIL161W | SMU2 |  |  |  |  |  |  |  |  |
| YIL162W | SUC2 |  |  |  |  |  |  |  |  |
| YIL166C | SOA1 |  |  |  |  |  |  |  |  |
| YIR001C | SGN1 |  |  |  |  |  |  |  |  |
| YIR002C | MPH1 |  |  |  |  |  |  |  |  |
| YIR003W | AIM21 |  |  |  |  |  |  |  |  |
| YIR004W | DJP1 |  |  |  |  |  |  |  |  |
| YIR005W | IST3 |  |  |  |  |  |  |  |  |
| YIR006C | PAN1 |  |  |  |  |  |  |  |  |
| YIR007W | EGH1 |  |  |  |  |  |  |  |  |
| YIR008C | PRI1 |  |  |  |  |  |  |  |  |
| YIR009W | MSL1 |  |  |  |  |  |  |  |  |
| YIR010W | DSN1 |  |  |  |  |  |  |  |  |
| YIR011C | STS1 |  |  |  |  |  |  |  |  |
| YIR012W | SQT1 |  |  |  |  |  |  |  |  |
| YIR014W | VLD1 |  |  |  |  |  |  |  |  |
| YIR015W | RPR2 |  |  |  |  |  |  |  |  |
| YIR017C | MET28 |  |  |  |  |  |  |  |  |
| YIR019C | FLO11 |  |  |  |  |  |  |  |  |
| YIR021W | MRS1 |  |  |  |  |  |  |  |  |
| YIR022W | SEC11 |  |  |  |  |  |  |  |  |
| YIR023W | DAL81 |  |  |  |  |  |  |  |  |
| YIR024C | INA22 |  |  |  |  |  |  |  |  |
| YIR025W | MND2 |  |  |  |  |  |  |  |  |
| YIR026C | YVH1 |  |  |  |  |  |  |  |  |
| YIR027C | DAL1 |  |  |  |  |  |  |  |  |
| YIR028W | DAL4 |  |  |  |  |  |  |  |  |
| YIR029W | DAL2 |  |  |  |  |  |  |  |  |
| YIR031C | DAL7 |  |  |  |  |  |  |  |  |
| YIR032C | DAL3 |  |  |  |  |  |  |  |  |
| YIR033W | MGA2 |  |  |  |  |  |  |  |  |
| YIR034C | LYS1 |  |  |  |  |  |  |  |  |
| YIR035C | NRE1 |  |  |  |  |  |  |  |  |
| YIR036C | IRC24 |  |  |  |  |  |  |  |  |
| YIR037W | HYR1 |  |  |  |  |  |  |  |  |
| YJL001W | PRE3 |  |  |  |  |  |  |  |  |
| YJL002C | OST1 |  |  |  |  |  |  |  |  |
| YJL003W | COX16 |  |  |  |  |  |  |  |  |
| YJL004C | SYS1 |  |  |  |  |  |  |  |  |
| YJL005W | CYR1 |  |  |  |  |  |  |  |  |
| YJL006C | CTK2 |  |  |  |  |  |  |  |  |
| YJL008C | CCT8 |  |  |  |  |  |  |  |  |
| YJL010C | NOP9 |  |  |  |  |  |  |  |  |
| YJL011C | RPC17 |  |  |  |  |  |  |  |  |
| YJL012C | VTC4 |  |  |  |  |  |  |  |  |
| YJL013C | MAD3 |  |  |  |  |  |  |  |  |
| YJL014W | CCT3 |  |  |  |  |  |  |  |  |
| YJL019W | MPS3 |  |  |  |  |  |  |  |  |
| YJL020C | BBC1 |  |  |  |  |  |  |  |  |
| YJL023C | PET130 |  |  |  |  |  |  |  |  |
| YJL024C | APS3 |  |  |  |  |  |  |  |  |
| YJL025W | RRN7 |  |  |  |  |  |  |  |  |
| YJL026W | RNR2 |  |  |  |  |  |  |  |  |
| YJL029C | VPS53 |  |  |  |  |  |  |  |  |
| YJL030W | MAD2 |  |  |  |  |  |  |  |  |
| YJL031C | BET4 |  |  |  |  |  |  |  |  |
| YJL033W | HCA4 |  |  |  |  |  |  |  |  |
| YJL034W | KAR2 |  |  |  |  |  |  |  |  |
| YJL035C | TAD2 |  |  |  |  |  |  |  |  |
| YJL036W | SNX4 |  |  |  |  |  |  |  |  |
| YJL039C | NUP192 |  |  |  |  |  |  |  |  |
| YJL041W | NSP1 |  |  |  |  |  |  |  |  |
| YJL042W | MHP1 |  |  |  |  |  |  |  |  |
| YJL044C | GYP6 |  |  |  |  |  |  |  |  |
| YJL045W | SDH9 |  |  |  |  |  |  |  |  |
| YJL046W | AIM22 |  |  |  |  |  |  |  |  |
| YJL047C | RTT101 |  |  |  |  |  |  |  |  |
| YJL048C | UBX6 |  |  |  |  |  |  |  |  |
| YJL049W | CHM7 |  |  |  |  |  |  |  |  |
| YJL050W | MTR4 |  |  |  |  |  |  |  |  |
| YJL051W | IRC8 |  |  |  |  |  |  |  |  |
| YJL053W | PEP8 |  |  |  |  |  |  |  |  |
| YJL054W | TIM54 |  |  |  |  |  |  |  |  |
| YJL055W | LOG1 |  |  |  |  |  |  |  |  |
| YJL056C | ZAP1 |  |  |  |  |  |  |  |  |
| YJL057C | IKS1 |  |  |  |  |  |  |  |  |
| YJL058C | BIT61 |  |  |  |  |  |  |  |  |
| YJL059W | YHC3 |  |  |  |  |  |  |  |  |
| YJL060W | BNA3 |  |  |  |  |  |  |  |  |
| YJL061W | NUP82 |  |  |  |  |  |  |  |  |
| YJL062W | LAS21 |  |  |  |  |  |  |  |  |
| YJL062W-A | COA3 |  |  |  |  |  |  |  |  |
| YJL063C | MRPL8 |  |  |  |  |  |  |  |  |
| YJL065C | DLS1 |  |  |  |  |  |  |  |  |
| YJL066C | MPM1 |  |  |  |  |  |  |  |  |
| YJL068C |  |  |  |  |  |  |  |  |  |
| YJL069C | UTP18 |  |  |  |  |  |  |  |  |
| YJL071W | ARG2 |  |  |  |  |  |  |  |  |
| YJL072C | PSF2 |  |  |  |  |  |  |  |  |
| YJL073W | JEM1 |  |  |  |  |  |  |  |  |
| YJL074C | SMC3 |  |  |  |  |  |  |  |  |
| YJL076W | NET1 |  |  |  |  |  |  |  |  |
| YJL078C | PRY3 |  |  |  |  |  |  |  |  |
| YJL079C | PRY1 |  |  |  |  |  |  |  |  |
| YJL080C | SCP160 |  |  |  |  |  |  |  |  |
| YJL081C | ARP4 |  |  |  |  |  |  |  |  |
| YJL083W | TAX4 |  |  |  |  |  |  |  |  |
| YJL084C | ALY2 |  |  |  |  |  |  |  |  |
| YJL085W | EXO70 |  |  |  |  |  |  |  |  |
| YJL087C | TRL1 |  |  |  |  |  |  |  |  |
| YJL088W | ARG3 |  |  |  |  |  |  |  |  |
| YJL089W | SIP4 |  |  |  |  |  |  |  |  |
| YJL090C | DPB11 |  |  |  |  |  |  |  |  |
| YJL091C | GWT1 |  |  |  |  |  |  |  |  |
| YJL092W | SRS2 |  |  |  |  |  |  |  |  |
| YJL095W | BCK1 |  |  |  |  |  |  |  |  |
| YJL096W | MRPL49 |  |  |  |  |  |  |  |  |
| YJL097W | PHS1 |  |  |  |  |  |  |  |  |
| YJL098W | SAP185 |  |  |  |  |  |  |  |  |
| YJL099W | CHS6 |  |  |  |  |  |  |  |  |
| YJL100W | LSB6 |  |  |  |  |  |  |  |  |
| YJL101C | GSH1 |  |  |  |  |  |  |  |  |
| YJL102W | MEF2 |  |  |  |  |  |  |  |  |
| YJL103C | GSM1 |  |  |  |  |  |  |  |  |
| YJL104W | PAM16 |  |  |  |  |  |  |  |  |
| YJL105W | SET4 |  |  |  |  |  |  |  |  |
| YJL106W | IME2 |  |  |  |  |  |  |  |  |
| YJL107C_108C | PRM10 |  |  |  |  |  |  |  |  |
| YJL109C | UTP10 |  |  |  |  |  |  |  |  |
| YJL110C | GZF3 |  |  |  |  |  |  |  |  |
| YJL111W | CCT7 |  |  |  |  |  |  |  |  |
| YJL112W | MDV1 |  |  |  |  |  |  |  |  |
| YJL115W | ASF1 |  |  |  |  |  |  |  |  |
| YJL116C | NCA3 |  |  |  |  |  |  |  |  |
| YJL117W | PHO86 |  |  |  |  |  |  |  |  |
| YJL121C | RPE1 |  |  |  |  |  |  |  |  |
| YJL122W | ALB1 |  |  |  |  |  |  |  |  |
| YJL123C | MTC1 |  |  |  |  |  |  |  |  |
| YJL124C | LSM1 |  |  |  |  |  |  |  |  |
| YJL125C | GCD14 |  |  |  |  |  |  |  |  |
| YJL127C | SPT10 |  |  |  |  |  |  |  |  |
| YJL127C-B | MCO6 |  |  |  |  |  |  |  |  |
| YJL128C | PBS2 |  |  |  |  |  |  |  |  |
| YJL129C | TRK1 |  |  |  |  |  |  |  |  |
| YJL130C | URA2 |  |  |  |  |  |  |  |  |
| YJL131C | AIM23 |  |  |  |  |  |  |  |  |
| YJL133W | MRS3 |  |  |  |  |  |  |  |  |
| YJL134W | LCB3 |  |  |  |  |  |  |  |  |
| YJL136C | RPS21B |  |  |  |  |  |  |  |  |
| YJL137C | GLG2 |  |  |  |  |  |  |  |  |
| YJL138C | TIF2 |  |  |  |  |  |  |  |  |
| YJL139C | YUR1 |  |  |  |  |  |  |  |  |
| YJL140W | RPB4 |  |  |  |  |  |  |  |  |
| YJL141C | YAK1 |  |  |  |  |  |  |  |  |
| YJL143W | TIM17 |  |  |  |  |  |  |  |  |
| YJL145W | SFH5 |  |  |  |  |  |  |  |  |
| YJL146W | IDS2 |  |  |  |  |  |  |  |  |
| YJL148W | RPA34 |  |  |  |  |  |  |  |  |
| YJL149W | DAS1 |  |  |  |  |  |  |  |  |
| YJL154C | VPS35 |  |  |  |  |  |  |  |  |
| YJL155C | FBP26 |  |  |  |  |  |  |  |  |
| YJL156C | SSY5 |  |  |  |  |  |  |  |  |
| YJL157C | FAR1 |  |  |  |  |  |  |  |  |
| YJL158C | CIS3 |  |  |  |  |  |  |  |  |
| YJL159W | HSP150 |  |  |  |  |  |  |  |  |
| YJL162C | JJJ2 |  |  |  |  |  |  |  |  |
| YJL164C | TPK1 |  |  |  |  |  |  |  |  |
| YJL165C | HAL5 |  |  |  |  |  |  |  |  |
| YJL166W | QCR8 |  |  |  |  |  |  |  |  |
| YJL167W | ERG20 |  |  |  |  |  |  |  |  |
| YJL168C | SET2 |  |  |  |  |  |  |  |  |
| YJL170C | ASG7 |  |  |  |  |  |  |  |  |
| YJL171C | TOH1 |  |  |  |  |  |  |  |  |
| YJL172W | CPS1 |  |  |  |  |  |  |  |  |
| YJL173C | RFA3 |  |  |  |  |  |  |  |  |
| YJL174W | KRE9 |  |  |  |  |  |  |  |  |
| YJL176C | SWI3 |  |  |  |  |  |  |  |  |
| YJL177W | RPL17B |  |  |  |  |  |  |  |  |
| YJL178C | ATG27 |  |  |  |  |  |  |  |  |
| YJL179W | PFD1 |  |  |  |  |  |  |  |  |
| YJL180C | ATP12 |  |  |  |  |  |  |  |  |
| YJL183W | MNN11 |  |  |  |  |  |  |  |  |
| YJL184W | GON7 |  |  |  |  |  |  |  |  |
| YJL185C | ATG36 |  |  |  |  |  |  |  |  |
| YJL186W | MNN5 |  |  |  |  |  |  |  |  |
| YJL187C | SWE1 |  |  |  |  |  |  |  |  |
| YJL189W | RPL39 |  |  |  |  |  |  |  |  |
| YJL190C | RPS22A |  |  |  |  |  |  |  |  |
| YJL191W | RPS14B |  |  |  |  |  |  |  |  |
| YJL192C | SOP4 |  |  |  |  |  |  |  |  |
| YJL194W | CDC6 |  |  |  |  |  |  |  |  |
| YJL196C | ELO1 |  |  |  |  |  |  |  |  |
| YJL197W | UBP12 |  |  |  |  |  |  |  |  |
| YJL198W | PHO90 |  |  |  |  |  |  |  |  |
| YJL200C | ACO2 |  |  |  |  |  |  |  |  |
| YJL201W | ECM25 |  |  |  |  |  |  |  |  |
| YJL203W | PRP21 |  |  |  |  |  |  |  |  |
| YJL204C | RCY1 |  |  |  |  |  |  |  |  |
| YJL208C | NUC1 |  |  |  |  |  |  |  |  |
| YJL210W | PEX2 |  |  |  |  |  |  |  |  |
| YJL212C | OPT1 |  |  |  |  |  |  |  |  |
| YJR001W | AVT1 |  |  |  |  |  |  |  |  |
| YJR002W | MPP10 |  |  |  |  |  |  |  |  |
| YJR004C | SAG1 |  |  |  |  |  |  |  |  |
| YJR005C-A | LSO1 |  |  |  |  |  |  |  |  |
| YJR005W | APL1 |  |  |  |  |  |  |  |  |
| YJR006W | POL31 |  |  |  |  |  |  |  |  |
| YJR007W | SUI2 |  |  |  |  |  |  |  |  |
| YJR008W | MHO1 |  |  |  |  |  |  |  |  |
| YJR010C-A | SPC1 |  |  |  |  |  |  |  |  |
| YJR010W | MET3 |  |  |  |  |  |  |  |  |
| YJR011C | CAL4 |  |  |  |  |  |  |  |  |
| YJR012C |  |  |  |  |  |  |  |  |  |
| YJR013W | GPI14 |  |  |  |  |  |  |  |  |
| YJR014W | TMA22 |  |  |  |  |  |  |  |  |
| YJR016C | ILV3 |  |  |  |  |  |  |  |  |
| YJR017C | ESS1 |  |  |  |  |  |  |  |  |
| YJR019C | TES1 |  |  |  |  |  |  |  |  |
| YJR021C | REC107 |  |  |  |  |  |  |  |  |
| YJR022W | LSM8 |  |  |  |  |  |  |  |  |
| YJR024C | MDE1 |  |  |  |  |  |  |  |  |
| YJR025C | BNA1 |  |  |  |  |  |  |  |  |
| YJR031C | GEA1 |  |  |  |  |  |  |  |  |
| YJR032W | CPR7 |  |  |  |  |  |  |  |  |
| YJR033C | RAV1 |  |  |  |  |  |  |  |  |
| YJR034W | PET191 |  |  |  |  |  |  |  |  |
| YJR035W | RAD26 |  |  |  |  |  |  |  |  |
| YJR036C | HUL4 |  |  |  |  |  |  |  |  |
| YJR040W | GEF1 |  |  |  |  |  |  |  |  |
| YJR041C | URB2 |  |  |  |  |  |  |  |  |
| YJR042W | NUP85 |  |  |  |  |  |  |  |  |
| YJR043C | POL32 |  |  |  |  |  |  |  |  |
| YJR044C | VPS55 |  |  |  |  |  |  |  |  |
| YJR045C | SSC1 |  |  |  |  |  |  |  |  |
| YJR046W | TAH11 |  |  |  |  |  |  |  |  |
| YJR047C | ANB1 |  |  |  |  |  |  |  |  |
| YJR048W | CYC1 |  |  |  |  |  |  |  |  |
| YJR049C | UTR1 |  |  |  |  |  |  |  |  |
| YJR050W | ISY1 |  |  |  |  |  |  |  |  |
| YJR051W | OSM1 |  |  |  |  |  |  |  |  |
| YJR052W | RAD7 |  |  |  |  |  |  |  |  |
| YJR053W | BFA1 |  |  |  |  |  |  |  |  |
| YJR054W | KCH1 |  |  |  |  |  |  |  |  |
| YJR055W | HIT1 |  |  |  |  |  |  |  |  |
| YJR057W | CDC8 |  |  |  |  |  |  |  |  |
| YJR058C | APS2 |  |  |  |  |  |  |  |  |
| YJR059W | PTK2 |  |  |  |  |  |  |  |  |
| YJR060W | CBF1 |  |  |  |  |  |  |  |  |
| YJR062C | NTA1 |  |  |  |  |  |  |  |  |
| YJR063W | RPA12 |  |  |  |  |  |  |  |  |
| YJR064W | CCT5 |  |  |  |  |  |  |  |  |
| YJR065C | ARP3 |  |  |  |  |  |  |  |  |
| YJR066W | TOR1 |  |  |  |  |  |  |  |  |
| YJR067C | YAE1 |  |  |  |  |  |  |  |  |
| YJR068W | RFC2 |  |  |  |  |  |  |  |  |
| YJR069C | HAM1 |  |  |  |  |  |  |  |  |
| YJR070C | LIA1 |  |  |  |  |  |  |  |  |
| YJR072C | NPA3 |  |  |  |  |  |  |  |  |
| YJR073C | OPI3 |  |  |  |  |  |  |  |  |
| YJR074W | MOG1 |  |  |  |  |  |  |  |  |
| YJR075W | HOC1 |  |  |  |  |  |  |  |  |
| YJR076C | CDC11 |  |  |  |  |  |  |  |  |
| YJR077C | MIR1 |  |  |  |  |  |  |  |  |
| YJR078W | BNA2 |  |  |  |  |  |  |  |  |
| YJR080C | AIM24 |  |  |  |  |  |  |  |  |
| YJR082C | EAF6 |  |  |  |  |  |  |  |  |
| YJR083C | ACF4 |  |  |  |  |  |  |  |  |
| YJR084W |  |  |  |  |  |  |  |  |  |
| YJR085C | TMH11 |  |  |  |  |  |  |  |  |
| YJR086W | STE18 |  |  |  |  |  |  |  |  |
| YJR088C | EMC2 |  |  |  |  |  |  |  |  |
| YJR089W | BIR1 |  |  |  |  |  |  |  |  |
| YJR090C | GRR1 |  |  |  |  |  |  |  |  |
| YJR091C | JSN1 |  |  |  |  |  |  |  |  |
| YJR092W | BUD4 |  |  |  |  |  |  |  |  |
| YJR093C | FIP1 |  |  |  |  |  |  |  |  |
| YJR094C | IME1 |  |  |  |  |  |  |  |  |
| YJR094W-A | RPL43B |  |  |  |  |  |  |  |  |
| YJR095W | SFC1 |  |  |  |  |  |  |  |  |
| YJR096W |  |  |  |  |  |  |  |  |  |
| YJR097W | JJJ3 |  |  |  |  |  |  |  |  |
| YJR099W | YUH1 |  |  |  |  |  |  |  |  |
| YJR101W | RSM26 |  |  |  |  |  |  |  |  |
| YJR102C | VPS25 |  |  |  |  |  |  |  |  |
| YJR103W | URA8 |  |  |  |  |  |  |  |  |
| YJR104C | SOD1 |  |  |  |  |  |  |  |  |
| YJR105W | ADO1 |  |  |  |  |  |  |  |  |
| YJR106W | ECM27 |  |  |  |  |  |  |  |  |
| YJR109C | CPA2 |  |  |  |  |  |  |  |  |
| YJR110W | YMR1 |  |  |  |  |  |  |  |  |
| YJR111C | PXP2 |  |  |  |  |  |  |  |  |
| YJR112W | NNF1 |  |  |  |  |  |  |  |  |
| YJR113C | RSM7 |  |  |  |  |  |  |  |  |
| YJR117W | STE24 |  |  |  |  |  |  |  |  |
| YJR118C | ILM1 |  |  |  |  |  |  |  |  |
| YJR119C | JHD2 |  |  |  |  |  |  |  |  |
| YJR121W | ATP2 |  |  |  |  |  |  |  |  |
| YJR122W | IBA57 |  |  |  |  |  |  |  |  |
| YJR123W | RPS5 |  |  |  |  |  |  |  |  |
| YJR125C | ENT3 |  |  |  |  |  |  |  |  |
| YJR126C | VPS70 |  |  |  |  |  |  |  |  |
| YJR127C | RSF2 |  |  |  |  |  |  |  |  |
| YJR129C | EFM3 |  |  |  |  |  |  |  |  |
| YJR130C | STR2 |  |  |  |  |  |  |  |  |
| YJR131W | MNS1 |  |  |  |  |  |  |  |  |
| YJR132W | NMD5 |  |  |  |  |  |  |  |  |
| YJR133W | XPT1 |  |  |  |  |  |  |  |  |
| YJR134C | SGM1 |  |  |  |  |  |  |  |  |
| YJR135C | MCM22 |  |  |  |  |  |  |  |  |
| YJR135W-A | TIM8 |  |  |  |  |  |  |  |  |
| YJR136C | TTI2 |  |  |  |  |  |  |  |  |
| YJR137C | MET5 |  |  |  |  |  |  |  |  |
| YJR138W | IML1 |  |  |  |  |  |  |  |  |
| YJR139C | HOM6 |  |  |  |  |  |  |  |  |
| YJR140C | HIR3 |  |  |  |  |  |  |  |  |
| YJR141W | IPA1 |  |  |  |  |  |  |  |  |
| YJR142W |  |  |  |  |  |  |  |  |  |
| YJR143C | PMT4 |  |  |  |  |  |  |  |  |
| YJR144W | MGM101 |  |  |  |  |  |  |  |  |
| YJR145C | RPS4A |  |  |  |  |  |  |  |  |
| YJR147W | HMS2 |  |  |  |  |  |  |  |  |
| YJR148W | BAT2 |  |  |  |  |  |  |  |  |
| YJR155W | AAD10 |  |  |  |  |  |  |  |  |
| YJR156C | THI11 |  |  |  |  |  |  |  |  |
| YKL001C | MET14 |  |  |  |  |  |  |  |  |
| YKL002W | DID4 |  |  |  |  |  |  |  |  |
| YKL003C | MRP17 |  |  |  |  |  |  |  |  |
| YKL004W | AUR1 |  |  |  |  |  |  |  |  |
| YKL006C-A | SFT1 |  |  |  |  |  |  |  |  |
| YKL006W | RPL14A |  |  |  |  |  |  |  |  |
| YKL007W | CAP1 |  |  |  |  |  |  |  |  |
| YKL008C | LAC1 |  |  |  |  |  |  |  |  |
| YKL009W | MRT4 |  |  |  |  |  |  |  |  |
| YKL010C | UFD4 |  |  |  |  |  |  |  |  |
| YKL011C | CCE1 |  |  |  |  |  |  |  |  |
| YKL012W | PRP40 |  |  |  |  |  |  |  |  |
| YKL013C | ARC19 |  |  |  |  |  |  |  |  |
| YKL014C | URB1 |  |  |  |  |  |  |  |  |
| YKL015W | PUT3 |  |  |  |  |  |  |  |  |
| YKL016C | ATP7 |  |  |  |  |  |  |  |  |
| YKL017C | HCS1 |  |  |  |  |  |  |  |  |
| YKL018W | SWD2 |  |  |  |  |  |  |  |  |
| YKL019W | RAM2 |  |  |  |  |  |  |  |  |
| YKL020C | SPT23 |  |  |  |  |  |  |  |  |
| YKL021C | MAK11 |  |  |  |  |  |  |  |  |
| YKL022C | CDC16 |  |  |  |  |  |  |  |  |
| YKL023W | SKA1 |  |  |  |  |  |  |  |  |
| YKL024C | URA6 |  |  |  |  |  |  |  |  |
| YKL025C | PAN3 |  |  |  |  |  |  |  |  |
| YKL027W | TCD2 |  |  |  |  |  |  |  |  |
| YKL028W | TFA1 |  |  |  |  |  |  |  |  |
| YKL029C | MAE1 |  |  |  |  |  |  |  |  |
| YKL032C | IXR1 |  |  |  |  |  |  |  |  |
| YKL033W | TTI1 |  |  |  |  |  |  |  |  |
| YKL033W-A |  |  |  |  |  |  |  |  |  |
| YKL034W | TUL1 |  |  |  |  |  |  |  |  |
| YKL035W | UGP1 |  |  |  |  |  |  |  |  |
| YKL038W | RGT1 |  |  |  |  |  |  |  |  |
| YKL039W | PTM1 |  |  |  |  |  |  |  |  |
| YKL040C | NFU1 |  |  |  |  |  |  |  |  |
| YKL041W | VPS24 |  |  |  |  |  |  |  |  |
| YKL042W | SPC42 |  |  |  |  |  |  |  |  |
| YKL043W | PHD1 |  |  |  |  |  |  |  |  |
| YKL045W | PRI2 |  |  |  |  |  |  |  |  |
| YKL046C | DCW1 |  |  |  |  |  |  |  |  |
| YKL047W | ANR2 |  |  |  |  |  |  |  |  |
| YKL048C | ELM1 |  |  |  |  |  |  |  |  |
| YKL049C | CSE4 |  |  |  |  |  |  |  |  |
| YKL050C | LPX2 |  |  |  |  |  |  |  |  |
| YKL051W | SFK1 |  |  |  |  |  |  |  |  |
| YKL052C | ASK1 |  |  |  |  |  |  |  |  |
| YKL053C-A | MDM35 |  |  |  |  |  |  |  |  |
| YKL054C | DEF1 |  |  |  |  |  |  |  |  |
| YKL055C | OAR1 |  |  |  |  |  |  |  |  |
| YKL056C | TMA19 |  |  |  |  |  |  |  |  |
| YKL057C | NUP120 |  |  |  |  |  |  |  |  |
| YKL058W | TOA2 |  |  |  |  |  |  |  |  |
| YKL059C | MPE1 |  |  |  |  |  |  |  |  |
| YKL060C | FBA1 |  |  |  |  |  |  |  |  |
| YKL061W | BLI1 |  |  |  |  |  |  |  |  |
| YKL062W | MSN4 |  |  |  |  |  |  |  |  |
| YKL064W | MNR2 |  |  |  |  |  |  |  |  |
| YKL065C | YET1 |  |  |  |  |  |  |  |  |
| YKL065W-A | DPC7 |  |  |  |  |  |  |  |  |
| YKL067W | YNK1 |  |  |  |  |  |  |  |  |
| YKL068W | NUP100 |  |  |  |  |  |  |  |  |
| YKL069W |  |  |  |  |  |  |  |  |  |
| YKL072W | STB6 |  |  |  |  |  |  |  |  |
| YKL073W | LHS1 |  |  |  |  |  |  |  |  |
| YKL074C | MUD2 |  |  |  |  |  |  |  |  |
| YKL075C | AAN1 |  |  |  |  |  |  |  |  |
| YKL077W | PSG1 |  |  |  |  |  |  |  |  |
| YKL078W | DHR2 |  |  |  |  |  |  |  |  |
| YKL079W | SMY1 |  |  |  |  |  |  |  |  |
| YKL080W | VMA5 |  |  |  |  |  |  |  |  |
| YKL081W | TEF4 |  |  |  |  |  |  |  |  |
| YKL082C | RRP14 |  |  |  |  |  |  |  |  |
| YKL085W | MDH1 |  |  |  |  |  |  |  |  |
| YKL086W | SRX1 |  |  |  |  |  |  |  |  |
| YKL087C | CYT2 |  |  |  |  |  |  |  |  |
| YKL088W | CAB3 |  |  |  |  |  |  |  |  |
| YKL089W | MIF2 |  |  |  |  |  |  |  |  |
| YKL090W | CUE2 |  |  |  |  |  |  |  |  |
| YKL091C |  |  |  |  |  |  |  |  |  |
| YKL092C | BUD2 |  |  |  |  |  |  |  |  |
| YKL093W | MBR1 |  |  |  |  |  |  |  |  |
| YKL095W | YJU2 |  |  |  |  |  |  |  |  |
| YKL096W | CWP1 |  |  |  |  |  |  |  |  |
| YKL096W-A | CWP2 |  |  |  |  |  |  |  |  |
| YKL098W | MTC2 |  |  |  |  |  |  |  |  |
| YKL099C | UTP11 |  |  |  |  |  |  |  |  |
| YKL101W | HSL1 |  |  |  |  |  |  |  |  |
| YKL103C | APE1 |  |  |  |  |  |  |  |  |
| YKL104C | GFA1 |  |  |  |  |  |  |  |  |
| YKL105C | SEG2 |  |  |  |  |  |  |  |  |
| YKL106W | AAT1 |  |  |  |  |  |  |  |  |
| YKL108W | SLD2 |  |  |  |  |  |  |  |  |
| YKL109W | HAP4 |  |  |  |  |  |  |  |  |
| YKL110C | KTI12 |  |  |  |  |  |  |  |  |
| YKL112W | ABF1 |  |  |  |  |  |  |  |  |
| YKL113C | RAD27 |  |  |  |  |  |  |  |  |
| YKL114C | APN1 |  |  |  |  |  |  |  |  |
| YKL116C | PRR1 |  |  |  |  |  |  |  |  |
| YKL117W | SBA1 |  |  |  |  |  |  |  |  |
| YKL119C | VPH2 |  |  |  |  |  |  |  |  |
| YKL120W | OAC1 |  |  |  |  |  |  |  |  |
| YKL122C | SRP21 |  |  |  |  |  |  |  |  |
| YKL124W | SSH4 |  |  |  |  |  |  |  |  |
| YKL125W | RRN3 |  |  |  |  |  |  |  |  |
| YKL126W | YPK1 |  |  |  |  |  |  |  |  |
| YKL127W | PGM1 |  |  |  |  |  |  |  |  |
| YKL128C | PMU1 |  |  |  |  |  |  |  |  |
| YKL129C | MYO3 |  |  |  |  |  |  |  |  |
| YKL130C | SHE2 |  |  |  |  |  |  |  |  |
| YKL132C | RMA1 |  |  |  |  |  |  |  |  |
| YKL134C | OCT1 |  |  |  |  |  |  |  |  |
| YKL135C | APL2 |  |  |  |  |  |  |  |  |
| YKL137W | CMC1 |  |  |  |  |  |  |  |  |
| YKL138C | MRPL31 |  |  |  |  |  |  |  |  |
| YKL138C-A | HSK3 |  |  |  |  |  |  |  |  |
| YKL139W | CTK1 |  |  |  |  |  |  |  |  |
| YKL140W | TGL1 |  |  |  |  |  |  |  |  |
| YKL141W | SDH3 |  |  |  |  |  |  |  |  |
| YKL142W | MRP8 |  |  |  |  |  |  |  |  |
| YKL143W | LTV1 |  |  |  |  |  |  |  |  |
| YKL144C | RPC25 |  |  |  |  |  |  |  |  |
| YKL145W | RPT1 |  |  |  |  |  |  |  |  |
| YKL146W | AVT3 |  |  |  |  |  |  |  |  |
| YKL148C | SDH1 |  |  |  |  |  |  |  |  |
| YKL149C | DBR1 |  |  |  |  |  |  |  |  |
| YKL150W | MCR1 |  |  |  |  |  |  |  |  |
| YKL151C | NNR2 |  |  |  |  |  |  |  |  |
| YKL152C | GPM1 |  |  |  |  |  |  |  |  |
| YKL154W | SRP102 |  |  |  |  |  |  |  |  |
| YKL155C | RSM22 |  |  |  |  |  |  |  |  |
| YKL156W | RPS27A |  |  |  |  |  |  |  |  |
| YKL157W | APE2 |  |  |  |  |  |  |  |  |
| YKL159C | RCN1 |  |  |  |  |  |  |  |  |
| YKL160W | ELF1 |  |  |  |  |  |  |  |  |
| YKL161C | KDX1 |  |  |  |  |  |  |  |  |
| YKL163W | PIR3 |  |  |  |  |  |  |  |  |
| YKL164C | PIR1 |  |  |  |  |  |  |  |  |
| YKL165C | MCD4 |  |  |  |  |  |  |  |  |
| YKL166C | TPK3 |  |  |  |  |  |  |  |  |
| YKL167C | MRP49 |  |  |  |  |  |  |  |  |
| YKL168C | KKQ8 |  |  |  |  |  |  |  |  |
| YKL170W | MRPL38 |  |  |  |  |  |  |  |  |
| YKL171W | NNK1 |  |  |  |  |  |  |  |  |
| YKL172W | EBP2 |  |  |  |  |  |  |  |  |
| YKL173W | SNU114 |  |  |  |  |  |  |  |  |
| YKL174C | TPO5 |  |  |  |  |  |  |  |  |
| YKL175W | ZRT3 |  |  |  |  |  |  |  |  |
| YKL176C | LST4 |  |  |  |  |  |  |  |  |
| YKL178C | STE3 |  |  |  |  |  |  |  |  |
| YKL179C | COY1 |  |  |  |  |  |  |  |  |
| YKL180W | RPL17A |  |  |  |  |  |  |  |  |
| YKL181W | PRS1 |  |  |  |  |  |  |  |  |
| YKL182W | FAS1 |  |  |  |  |  |  |  |  |
| YKL183W | LOT5 |  |  |  |  |  |  |  |  |
| YKL184W | SPE1 |  |  |  |  |  |  |  |  |
| YKL185W | ASH1 |  |  |  |  |  |  |  |  |
| YKL186C | MTR2 |  |  |  |  |  |  |  |  |
| YKL187C | FAT3 |  |  |  |  |  |  |  |  |
| YKL188C | PXA2 |  |  |  |  |  |  |  |  |
| YKL189W | HYM1 |  |  |  |  |  |  |  |  |
| YKL190W | CNB1 |  |  |  |  |  |  |  |  |
| YKL191W | DPH2 |  |  |  |  |  |  |  |  |
| YKL192C | ACP1 |  |  |  |  |  |  |  |  |
| YKL193C | SDS22 |  |  |  |  |  |  |  |  |
| YKL194C | MST1 |  |  |  |  |  |  |  |  |
| YKL195W | MIA40 |  |  |  |  |  |  |  |  |
| YKL196C | YKT6 |  |  |  |  |  |  |  |  |
| YKL197C | PEX1 |  |  |  |  |  |  |  |  |
| YKL198C | PTK1 |  |  |  |  |  |  |  |  |
| YKL201C | MNN4 |  |  |  |  |  |  |  |  |
| YKL203C | TOR2 |  |  |  |  |  |  |  |  |
| YKL204W | EAP1 |  |  |  |  |  |  |  |  |
| YKL205W | LOS1 |  |  |  |  |  |  |  |  |
| YKL206C | ADD66 |  |  |  |  |  |  |  |  |
| YKL207W | EMC3 |  |  |  |  |  |  |  |  |
| YKL209C | STE6 |  |  |  |  |  |  |  |  |
| YKL210W | UBA1 |  |  |  |  |  |  |  |  |
| YKL211C | TRP3 |  |  |  |  |  |  |  |  |
| YKL212W | SAC1 |  |  |  |  |  |  |  |  |
| YKL213C | DOA1 |  |  |  |  |  |  |  |  |
| YKL214C | YRA2 |  |  |  |  |  |  |  |  |
| YKL215C | OXP1 |  |  |  |  |  |  |  |  |
| YKL217W | JEN1 |  |  |  |  |  |  |  |  |
| YKL222C |  |  |  |  |  |  |  |  |  |
| YKR001C | VPS1 |  |  |  |  |  |  |  |  |
| YKR002W | PAP1 |  |  |  |  |  |  |  |  |
| YKR003W | OSH6 |  |  |  |  |  |  |  |  |
| YKR004C | ECM9 |  |  |  |  |  |  |  |  |
| YKR006C | MRPL13 |  |  |  |  |  |  |  |  |
| YKR007W | MEH1 |  |  |  |  |  |  |  |  |
| YKR008W | RSC4 |  |  |  |  |  |  |  |  |
| YKR009C | FOX2 |  |  |  |  |  |  |  |  |
| YKR010C | TOF2 |  |  |  |  |  |  |  |  |
| YKR011C | FPT1 |  |  |  |  |  |  |  |  |
| YKR013W | PRY2 |  |  |  |  |  |  |  |  |
| YKR014C | YPT52 |  |  |  |  |  |  |  |  |
| YKR016W | MIC60 |  |  |  |  |  |  |  |  |
| YKR017C | HEL1 |  |  |  |  |  |  |  |  |
| YKR019C | IRS4 |  |  |  |  |  |  |  |  |
| YKR020W | VPS51 |  |  |  |  |  |  |  |  |
| YKR021W | ALY1 |  |  |  |  |  |  |  |  |
| YKR022C | NTR2 |  |  |  |  |  |  |  |  |
| YKR023W | RQT4 |  |  |  |  |  |  |  |  |
| YKR024C | DBP7 |  |  |  |  |  |  |  |  |
| YKR025W | RPC37 |  |  |  |  |  |  |  |  |
| YKR026C | GCN3 |  |  |  |  |  |  |  |  |
| YKR027W | BCH2 |  |  |  |  |  |  |  |  |
| YKR028W | SAP190 |  |  |  |  |  |  |  |  |
| YKR029C | SET3 |  |  |  |  |  |  |  |  |
| YKR030W | GMH1 |  |  |  |  |  |  |  |  |
| YKR031C | SPO14 |  |  |  |  |  |  |  |  |
| YKR034W | DAL80 |  |  |  |  |  |  |  |  |
| YKR035W-A | DID2 |  |  |  |  |  |  |  |  |
| YKR036C | CAF4 |  |  |  |  |  |  |  |  |
| YKR037C | SPC34 |  |  |  |  |  |  |  |  |
| YKR038C | KAE1 |  |  |  |  |  |  |  |  |
| YKR039W | GAP1 |  |  |  |  |  |  |  |  |
| YKR041W |  |  |  |  |  |  |  |  |  |
| YKR042W | UTH1 |  |  |  |  |  |  |  |  |
| YKR043C | SHB17 |  |  |  |  |  |  |  |  |
| YKR044W | UIP5 |  |  |  |  |  |  |  |  |
| YKR046C | PLN1 |  |  |  |  |  |  |  |  |
| YKR048C | NAP1 |  |  |  |  |  |  |  |  |
| YKR049C | FMP46 |  |  |  |  |  |  |  |  |
| YKR050W | TRK2 |  |  |  |  |  |  |  |  |
| YKR051W | HFL1 |  |  |  |  |  |  |  |  |
| YKR052C | MRS4 |  |  |  |  |  |  |  |  |
| YKR053C | YSR3 |  |  |  |  |  |  |  |  |
| YKR054C | DYN1 |  |  |  |  |  |  |  |  |
| YKR055W | RHO4 |  |  |  |  |  |  |  |  |
| YKR056W | TRM2 |  |  |  |  |  |  |  |  |
| YKR057W | RPS21A |  |  |  |  |  |  |  |  |
| YKR058W | GLG1 |  |  |  |  |  |  |  |  |
| YKR059W | TIF1 |  |  |  |  |  |  |  |  |
| YKR060W | UTP30 |  |  |  |  |  |  |  |  |
| YKR061W | KTR2 |  |  |  |  |  |  |  |  |
| YKR062W | TFA2 |  |  |  |  |  |  |  |  |
| YKR063C | LAS1 |  |  |  |  |  |  |  |  |
| YKR064W | OAF3 |  |  |  |  |  |  |  |  |
| YKR065C | PAM17 |  |  |  |  |  |  |  |  |
| YKR067W | GPT2 |  |  |  |  |  |  |  |  |
| YKR068C | BET3 |  |  |  |  |  |  |  |  |
| YKR069W | MET1 |  |  |  |  |  |  |  |  |
| YKR071C | DRE2 |  |  |  |  |  |  |  |  |
| YKR072C | SIS2 |  |  |  |  |  |  |  |  |
| YKR074W | AIM29 |  |  |  |  |  |  |  |  |
| YKR077W | MSA2 |  |  |  |  |  |  |  |  |
| YKR078W | VPS501 |  |  |  |  |  |  |  |  |
| YKR079C | TRZ1 |  |  |  |  |  |  |  |  |
| YKR080W | MTD1 |  |  |  |  |  |  |  |  |
| YKR081C | RPF2 |  |  |  |  |  |  |  |  |
| YKR082W | NUP133 |  |  |  |  |  |  |  |  |
| YKR083C | DAD2 |  |  |  |  |  |  |  |  |
| YKR084C | HBS1 |  |  |  |  |  |  |  |  |
| YKR085C | MRPL20 |  |  |  |  |  |  |  |  |
| YKR086W | PRP16 |  |  |  |  |  |  |  |  |
| YKR088C | TVP38 |  |  |  |  |  |  |  |  |
| YKR089C | TGL4 |  |  |  |  |  |  |  |  |
| YKR090W | PXL1 |  |  |  |  |  |  |  |  |
| YKR091W | SRL3 |  |  |  |  |  |  |  |  |
| YKR092C | SRP40 |  |  |  |  |  |  |  |  |
| YKR093W | PTR2 |  |  |  |  |  |  |  |  |
| YKR094C | RPL40B |  |  |  |  |  |  |  |  |
| YKR095W | MLP1 |  |  |  |  |  |  |  |  |
| YKR095W-A | PCC1 |  |  |  |  |  |  |  |  |
| YKR096W | ESL2 |  |  |  |  |  |  |  |  |
| YKR097W | PCK1 |  |  |  |  |  |  |  |  |
| YKR098C | UBP11 |  |  |  |  |  |  |  |  |
| YKR099W | BAS1 |  |  |  |  |  |  |  |  |
| YKR100C | SKG1 |  |  |  |  |  |  |  |  |
| YLL001W | DNM1 |  |  |  |  |  |  |  |  |
| YLL002W | RTT109 |  |  |  |  |  |  |  |  |
| YLL003W | SFI1 |  |  |  |  |  |  |  |  |
| YLL004W | ORC3 |  |  |  |  |  |  |  |  |
| YLL005C | SPO75 |  |  |  |  |  |  |  |  |
| YLL006W | MMM1 |  |  |  |  |  |  |  |  |
| YLL007C | LMO1 |  |  |  |  |  |  |  |  |
| YLL008W | DRS1 |  |  |  |  |  |  |  |  |
| YLL009C | COX17 |  |  |  |  |  |  |  |  |
| YLL010C | PSR1 |  |  |  |  |  |  |  |  |
| YLL011W | SOF1 |  |  |  |  |  |  |  |  |
| YLL012W | YEH1 |  |  |  |  |  |  |  |  |
| YLL013C | PUF3 |  |  |  |  |  |  |  |  |
| YLL014W | EMC6 |  |  |  |  |  |  |  |  |
| YLL018C | DPS1 |  |  |  |  |  |  |  |  |
| YLL018C-A | COX19 |  |  |  |  |  |  |  |  |
| YLL019C | KNS1 |  |  |  |  |  |  |  |  |
| YLL021W | SPA2 |  |  |  |  |  |  |  |  |
| YLL022C | HIF1 |  |  |  |  |  |  |  |  |
| YLL023C | POM33 |  |  |  |  |  |  |  |  |
| YLL024C | SSA2 |  |  |  |  |  |  |  |  |
| YLL026W | HSP104 |  |  |  |  |  |  |  |  |
| YLL027W | ISA1 |  |  |  |  |  |  |  |  |
| YLL029W | FRA1 |  |  |  |  |  |  |  |  |
| YLL031C | GPI13 |  |  |  |  |  |  |  |  |
| YLL032C |  |  |  |  |  |  |  |  |  |
| YLL033W | IRC19 |  |  |  |  |  |  |  |  |
| YLL034C | RIX7 |  |  |  |  |  |  |  |  |
| YLL035W | GRC3 |  |  |  |  |  |  |  |  |
| YLL036C | PRP19 |  |  |  |  |  |  |  |  |
| YLL038C | ENT4 |  |  |  |  |  |  |  |  |
| YLL039C | UBI4 |  |  |  |  |  |  |  |  |
| YLL040C | VPS13 |  |  |  |  |  |  |  |  |
| YLL041C | SDH2 |  |  |  |  |  |  |  |  |
| YLL042C | ATG10 |  |  |  |  |  |  |  |  |
| YLL043W | FPS1 |  |  |  |  |  |  |  |  |
| YLL045C | RPL8B |  |  |  |  |  |  |  |  |
| YLL046C | RNP1 |  |  |  |  |  |  |  |  |
| YLL048C | YBT1 |  |  |  |  |  |  |  |  |
| YLL049W | LDB18 |  |  |  |  |  |  |  |  |
| YLL050C | COF1 |  |  |  |  |  |  |  |  |
| YLL052C_053C | AQY2 |  |  |  |  |  |  |  |  |
| YLL055W | YCT1 |  |  |  |  |  |  |  |  |
| YLL057C | JLP1 |  |  |  |  |  |  |  |  |
| YLL061W | MMP1 |  |  |  |  |  |  |  |  |
| YLL062C | MHT1 |  |  |  |  |  |  |  |  |
| YLR002C | NOC3 |  |  |  |  |  |  |  |  |
| YLR003C | CMS1 |  |  |  |  |  |  |  |  |
| YLR005W | SSL1 |  |  |  |  |  |  |  |  |
| YLR006C | SSK1 |  |  |  |  |  |  |  |  |
| YLR007W | NSE1 |  |  |  |  |  |  |  |  |
| YLR008C | PAM18 |  |  |  |  |  |  |  |  |
| YLR009W | RLP24 |  |  |  |  |  |  |  |  |
| YLR010C | TEN1 |  |  |  |  |  |  |  |  |
| YLR015W | BRE2 |  |  |  |  |  |  |  |  |
| YLR016C | PML1 |  |  |  |  |  |  |  |  |
| YLR017W | MEU1 |  |  |  |  |  |  |  |  |
| YLR018C | POM34 |  |  |  |  |  |  |  |  |
| YLR019W | PSR2 |  |  |  |  |  |  |  |  |
| YLR020C | YEH2 |  |  |  |  |  |  |  |  |
| YLR021W | IRC25 |  |  |  |  |  |  |  |  |
| YLR022C | SDO1 |  |  |  |  |  |  |  |  |
| YLR023C | IZH3 |  |  |  |  |  |  |  |  |
| YLR024C | UBR2 |  |  |  |  |  |  |  |  |
| YLR025W | SNF7 |  |  |  |  |  |  |  |  |
| YLR026C | SED5 |  |  |  |  |  |  |  |  |
| YLR027C | AAT2 |  |  |  |  |  |  |  |  |
| YLR028C | ADE16 |  |  |  |  |  |  |  |  |
| YLR029C | RPL15A |  |  |  |  |  |  |  |  |
| YLR032W | RAD5 |  |  |  |  |  |  |  |  |
| YLR033W | RSC58 |  |  |  |  |  |  |  |  |
| YLR034C | SMF3 |  |  |  |  |  |  |  |  |
| YLR035C | MLH2 |  |  |  |  |  |  |  |  |
| YLR038C | COX12 |  |  |  |  |  |  |  |  |
| YLR039C | RIC1 |  |  |  |  |  |  |  |  |
| YLR043C | TRX1 |  |  |  |  |  |  |  |  |
| YLR044C | PDC1 |  |  |  |  |  |  |  |  |
| YLR045C | STU2 |  |  |  |  |  |  |  |  |
| YLR047C | FRE8 |  |  |  |  |  |  |  |  |
| YLR048W | RPS0B |  |  |  |  |  |  |  |  |
| YLR050C | EMA19 |  |  |  |  |  |  |  |  |
| YLR051C | FCF2 |  |  |  |  |  |  |  |  |
| YLR052W | IES3 |  |  |  |  |  |  |  |  |
| YLR054C | OSW2 |  |  |  |  |  |  |  |  |
| YLR055C | SPT8 |  |  |  |  |  |  |  |  |
| YLR056W | ERG3 |  |  |  |  |  |  |  |  |
| YLR057W | MNL2 |  |  |  |  |  |  |  |  |
| YLR058C | SHM2 |  |  |  |  |  |  |  |  |
| YLR059C | REX2 |  |  |  |  |  |  |  |  |
| YLR060W | FRS1 |  |  |  |  |  |  |  |  |
| YLR061W | RPL22A |  |  |  |  |  |  |  |  |
| YLR063W | BMT6 |  |  |  |  |  |  |  |  |
| YLR064W | PER33 |  |  |  |  |  |  |  |  |
| YLR065C | SND2 |  |  |  |  |  |  |  |  |
| YLR066W | SPC3 |  |  |  |  |  |  |  |  |
| YLR067C | PET309 |  |  |  |  |  |  |  |  |
| YLR068W | FYV7 |  |  |  |  |  |  |  |  |
| YLR069C | MEF1 |  |  |  |  |  |  |  |  |
| YLR070C | XYL2 |  |  |  |  |  |  |  |  |
| YLR071C | RGR1 |  |  |  |  |  |  |  |  |
| YLR072W | LAM6 |  |  |  |  |  |  |  |  |
| YLR073C | RFU1 |  |  |  |  |  |  |  |  |
| YLR074C | BUD20 |  |  |  |  |  |  |  |  |
| YLR075W | RPL10 |  |  |  |  |  |  |  |  |
| YLR077W | FMP25 |  |  |  |  |  |  |  |  |
| YLR078C | BOS1 |  |  |  |  |  |  |  |  |
| YLR079W | SIC1 |  |  |  |  |  |  |  |  |
| YLR080W | EMP46 |  |  |  |  |  |  |  |  |
| YLR083C | EMP70 |  |  |  |  |  |  |  |  |
| YLR084C | RAX2 |  |  |  |  |  |  |  |  |
| YLR085C | ARP6 |  |  |  |  |  |  |  |  |
| YLR086W | SMC4 |  |  |  |  |  |  |  |  |
| YLR087C | CSF1 |  |  |  |  |  |  |  |  |
| YLR088W | GAA1 |  |  |  |  |  |  |  |  |
| YLR089C | ALT1 |  |  |  |  |  |  |  |  |
| YLR090W | XDJ1 |  |  |  |  |  |  |  |  |
| YLR091W | GEP5 |  |  |  |  |  |  |  |  |
| YLR092W | SUL2 |  |  |  |  |  |  |  |  |
| YLR093C | NYV1 |  |  |  |  |  |  |  |  |
| YLR094C | GIS3 |  |  |  |  |  |  |  |  |
| YLR095C | IOC2 |  |  |  |  |  |  |  |  |
| YLR096W | KIN2 |  |  |  |  |  |  |  |  |
| YLR098C | CHA4 |  |  |  |  |  |  |  |  |
| YLR099C | ICT1 |  |  |  |  |  |  |  |  |
| YLR099W-A | MIM2 |  |  |  |  |  |  |  |  |
| YLR100W | ERG27 |  |  |  |  |  |  |  |  |
| YLR102C | APC9 |  |  |  |  |  |  |  |  |
| YLR103C | CDC45 |  |  |  |  |  |  |  |  |
| YLR105C | SEN2 |  |  |  |  |  |  |  |  |
| YLR106C | REA1 |  |  |  |  |  |  |  |  |
| YLR107W | REX3 |  |  |  |  |  |  |  |  |
| YLR108C |  |  |  |  |  |  |  |  |  |
| YLR110C | CCW12 |  |  |  |  |  |  |  |  |
| YLR113W | HOG1 |  |  |  |  |  |  |  |  |
| YLR114C | AVL9 |  |  |  |  |  |  |  |  |
| YLR115W | CFT2 |  |  |  |  |  |  |  |  |
| YLR116W | MSL5 |  |  |  |  |  |  |  |  |
| YLR117C | CLF1 |  |  |  |  |  |  |  |  |
| YLR118C | TML25 |  |  |  |  |  |  |  |  |
| YLR119W | SRN2 |  |  |  |  |  |  |  |  |
| YLR120C | YPS1 |  |  |  |  |  |  |  |  |
| YLR121C | YPS3 |  |  |  |  |  |  |  |  |
| YLR127C | APC2 |  |  |  |  |  |  |  |  |
| YLR128W | DCN1 |  |  |  |  |  |  |  |  |
| YLR129W | DIP2 |  |  |  |  |  |  |  |  |
| YLR130C | ZRT2 |  |  |  |  |  |  |  |  |
| YLR131C | ACE2 |  |  |  |  |  |  |  |  |
| YLR132C | USB1 |  |  |  |  |  |  |  |  |
| YLR133W | CKI1 |  |  |  |  |  |  |  |  |
| YLR134W | PDC5 |  |  |  |  |  |  |  |  |
| YLR135W | SLX4 |  |  |  |  |  |  |  |  |
| YLR136C | TIS11 |  |  |  |  |  |  |  |  |
| YLR137W | RKM5 |  |  |  |  |  |  |  |  |
| YLR138W | NHA1 |  |  |  |  |  |  |  |  |
| YLR139C | SLS1 |  |  |  |  |  |  |  |  |
| YLR141W | RRN5 |  |  |  |  |  |  |  |  |
| YLR142W | PUT1 |  |  |  |  |  |  |  |  |
| YLR143W | DPH6 |  |  |  |  |  |  |  |  |
| YLR144C | ACF2 |  |  |  |  |  |  |  |  |
| YLR145W | RMP1 |  |  |  |  |  |  |  |  |
| YLR146W-A |  |  |  |  |  |  |  |  |  |
| YLR147C | SMD3 |  |  |  |  |  |  |  |  |
| YLR148W | PEP3 |  |  |  |  |  |  |  |  |
| YLR149C | GID11 |  |  |  |  |  |  |  |  |
| YLR150W | STM1 |  |  |  |  |  |  |  |  |
| YLR151C | PCD1 |  |  |  |  |  |  |  |  |
| YLR153C | ACS2 |  |  |  |  |  |  |  |  |
| YLR154C | RNH203 |  |  |  |  |  |  |  |  |
| YLR163C | MAS1 |  |  |  |  |  |  |  |  |
| YLR164W | SHH4 |  |  |  |  |  |  |  |  |
| YLR165C | PUS5 |  |  |  |  |  |  |  |  |
| YLR166C | SEC10 |  |  |  |  |  |  |  |  |
| YLR167W | RPS31 |  |  |  |  |  |  |  |  |
| YLR168C | UPS2 |  |  |  |  |  |  |  |  |
| YLR170C | APS1 |  |  |  |  |  |  |  |  |
| YLR172C | DPH5 |  |  |  |  |  |  |  |  |
| YLR173W | TAG1 |  |  |  |  |  |  |  |  |
| YLR174W | IDP2 |  |  |  |  |  |  |  |  |
| YLR175W | CBF5 |  |  |  |  |  |  |  |  |
| YLR176C | RFX1 |  |  |  |  |  |  |  |  |
| YLR178C | TFS1 |  |  |  |  |  |  |  |  |
| YLR179C |  |  |  |  |  |  |  |  |  |
| YLR180W | SAM1 |  |  |  |  |  |  |  |  |
| YLR181C | VTA1 |  |  |  |  |  |  |  |  |
| YLR182W | SWI6 |  |  |  |  |  |  |  |  |
| YLR183C | TOS4 |  |  |  |  |  |  |  |  |
| YLR185W | RPL37A |  |  |  |  |  |  |  |  |
| YLR186W | EMG1 |  |  |  |  |  |  |  |  |
| YLR187W | SKG3 |  |  |  |  |  |  |  |  |
| YLR188W | MDL1 |  |  |  |  |  |  |  |  |
| YLR189C | ATG26 |  |  |  |  |  |  |  |  |
| YLR190W | MMR1 |  |  |  |  |  |  |  |  |
| YLR191W | PEX13 |  |  |  |  |  |  |  |  |
| YLR192C | HCR1 |  |  |  |  |  |  |  |  |
| YLR193C | UPS1 |  |  |  |  |  |  |  |  |
| YLR194C | NCW2 |  |  |  |  |  |  |  |  |
| YLR195C | NMT1 |  |  |  |  |  |  |  |  |
| YLR196W | PWP1 |  |  |  |  |  |  |  |  |
| YLR197W | NOP56 |  |  |  |  |  |  |  |  |
| YLR199C | PBA1 |  |  |  |  |  |  |  |  |
| YLR200W | YKE2 |  |  |  |  |  |  |  |  |
| YLR201C | COQ9 |  |  |  |  |  |  |  |  |
| YLR203C | MSS51 |  |  |  |  |  |  |  |  |
| YLR204W | QRI5 |  |  |  |  |  |  |  |  |
| YLR205C | HMX1 |  |  |  |  |  |  |  |  |
| YLR206W | ENT2 |  |  |  |  |  |  |  |  |
| YLR207W | HRD3 |  |  |  |  |  |  |  |  |
| YLR208W | SEC13 |  |  |  |  |  |  |  |  |
| YLR209C | PNP1 |  |  |  |  |  |  |  |  |
| YLR210W | CLB4 |  |  |  |  |  |  |  |  |
| YLR211C | ATG38 |  |  |  |  |  |  |  |  |
| YLR212C | TUB4 |  |  |  |  |  |  |  |  |
| YLR213C | CRR1 |  |  |  |  |  |  |  |  |
| YLR214W | FRE1 |  |  |  |  |  |  |  |  |
| YLR215C | CDC123 |  |  |  |  |  |  |  |  |
| YLR216C | CPR6 |  |  |  |  |  |  |  |  |
| YLR218C | COA4 |  |  |  |  |  |  |  |  |
| YLR219W | MSC3 |  |  |  |  |  |  |  |  |
| YLR220W | CCC1 |  |  |  |  |  |  |  |  |
| YLR221C | RSA3 |  |  |  |  |  |  |  |  |
| YLR222C | UTP13 |  |  |  |  |  |  |  |  |
| YLR223C | IFH1 |  |  |  |  |  |  |  |  |
| YLR224W | UCC1 |  |  |  |  |  |  |  |  |
| YLR226W | BUR2 |  |  |  |  |  |  |  |  |
| YLR227C | ADY4 |  |  |  |  |  |  |  |  |
| YLR228C | ECM22 |  |  |  |  |  |  |  |  |
| YLR229C | CDC42 |  |  |  |  |  |  |  |  |
| YLR231C | BNA5 |  |  |  |  |  |  |  |  |
| YLR233C | EST1 |  |  |  |  |  |  |  |  |
| YLR234W | TOP3 |  |  |  |  |  |  |  |  |
| YLR237W | THI7 |  |  |  |  |  |  |  |  |
| YLR238W | FAR10 |  |  |  |  |  |  |  |  |
| YLR239C | LIP2 |  |  |  |  |  |  |  |  |
| YLR240W | VPS34 |  |  |  |  |  |  |  |  |
| YLR241W | CSC1 |  |  |  |  |  |  |  |  |
| YLR242C | ARV1 |  |  |  |  |  |  |  |  |
| YLR243W | GPN3 |  |  |  |  |  |  |  |  |
| YLR244C | MAP1 |  |  |  |  |  |  |  |  |
| YLR245C | CDD1 |  |  |  |  |  |  |  |  |
| YLR246W | ERF2 |  |  |  |  |  |  |  |  |
| YLR247C | IRC20 |  |  |  |  |  |  |  |  |
| YLR248W | RCK2 |  |  |  |  |  |  |  |  |
| YLR249W | YEF3 |  |  |  |  |  |  |  |  |
| YLR250W | SSP120 |  |  |  |  |  |  |  |  |
| YLR251W | SYM1 |  |  |  |  |  |  |  |  |
| YLR253W | CQD2 |  |  |  |  |  |  |  |  |
| YLR254C | NDL1 |  |  |  |  |  |  |  |  |
| YLR256W | HAP1 |  |  |  |  |  |  |  |  |
| YLR257W |  |  |  |  |  |  |  |  |  |
| YLR258W | GSY2 |  |  |  |  |  |  |  |  |
| YLR259C | HSP60 |  |  |  |  |  |  |  |  |
| YLR260W | LCB5 |  |  |  |  |  |  |  |  |
| YLR262C | YPT6 |  |  |  |  |  |  |  |  |
| YLR262C-A | TMA7 |  |  |  |  |  |  |  |  |
| YLR263W | RED1 |  |  |  |  |  |  |  |  |
| YLR264W | RPS28B |  |  |  |  |  |  |  |  |
| YLR265C | NEJ1 |  |  |  |  |  |  |  |  |
| YLR268W | SEC22 |  |  |  |  |  |  |  |  |
| YLR270W | DCS1 |  |  |  |  |  |  |  |  |
| YLR272C | YCS4 |  |  |  |  |  |  |  |  |
| YLR273C | PIG1 |  |  |  |  |  |  |  |  |
| YLR274W | MCM5 |  |  |  |  |  |  |  |  |
| YLR275W | SMD2 |  |  |  |  |  |  |  |  |
| YLR276C | DBP9 |  |  |  |  |  |  |  |  |
| YLR277C | YSH1 |  |  |  |  |  |  |  |  |
| YLR281C | RSO55 |  |  |  |  |  |  |  |  |
| YLR284C | ECI1 |  |  |  |  |  |  |  |  |
| YLR285W | NNT1 |  |  |  |  |  |  |  |  |
| YLR287C-A | RPS30A |  |  |  |  |  |  |  |  |
| YLR288C | MEC3 |  |  |  |  |  |  |  |  |
| YLR289W | GUF1 |  |  |  |  |  |  |  |  |
| YLR290C | COQ11 |  |  |  |  |  |  |  |  |
| YLR291C | GCD7 |  |  |  |  |  |  |  |  |
| YLR292C | SEC72 |  |  |  |  |  |  |  |  |
| YLR293C | GSP1 |  |  |  |  |  |  |  |  |
| YLR295C | ATP14 |  |  |  |  |  |  |  |  |
| YLR298C | YHC1 |  |  |  |  |  |  |  |  |
| YLR299W | ECM38 |  |  |  |  |  |  |  |  |
| YLR300W | EXG1 |  |  |  |  |  |  |  |  |
| YLR303W | MET17 |  |  |  |  |  |  |  |  |
| YLR304C | ACO1 |  |  |  |  |  |  |  |  |
| YLR305C | STT4 |  |  |  |  |  |  |  |  |
| YLR306W | UBC12 |  |  |  |  |  |  |  |  |
| YLR307W | CDA1 |  |  |  |  |  |  |  |  |
| YLR308W | CDA2 |  |  |  |  |  |  |  |  |
| YLR309C | IMH1 |  |  |  |  |  |  |  |  |
| YLR310C | CDC25 |  |  |  |  |  |  |  |  |
| YLR312C | ATG39 |  |  |  |  |  |  |  |  |
| YLR312W-A | MRPL15 |  |  |  |  |  |  |  |  |
| YLR313C | SPH1 |  |  |  |  |  |  |  |  |
| YLR314C | CDC3 |  |  |  |  |  |  |  |  |
| YLR316C | TAD3 |  |  |  |  |  |  |  |  |
| YLR318W | EST2 |  |  |  |  |  |  |  |  |
| YLR319C | BUD6 |  |  |  |  |  |  |  |  |
| YLR320W | MMS22 |  |  |  |  |  |  |  |  |
| YLR321C | SFH1 |  |  |  |  |  |  |  |  |
| YLR323C | CWC24 |  |  |  |  |  |  |  |  |
| YLR324W | PEX30 |  |  |  |  |  |  |  |  |
| YLR325C | RPL38 |  |  |  |  |  |  |  |  |
| YLR327C | TMA10 |  |  |  |  |  |  |  |  |
| YLR328W | NMA1 |  |  |  |  |  |  |  |  |
| YLR329W | REC102 |  |  |  |  |  |  |  |  |
| YLR330W | CHS5 |  |  |  |  |  |  |  |  |
| YLR332W | MID2 |  |  |  |  |  |  |  |  |
| YLR333C | RPS25B |  |  |  |  |  |  |  |  |
| YLR335W | NUP2 |  |  |  |  |  |  |  |  |
| YLR336C | SGD1 |  |  |  |  |  |  |  |  |
| YLR337C | VRP1 |  |  |  |  |  |  |  |  |
| YLR340W | RPP0 |  |  |  |  |  |  |  |  |
| YLR341W | SPO77 |  |  |  |  |  |  |  |  |
| YLR342W | FKS1 |  |  |  |  |  |  |  |  |
| YLR343W | GAS2 |  |  |  |  |  |  |  |  |
| YLR344W | RPL26A |  |  |  |  |  |  |  |  |
| YLR347C | KAP95 |  |  |  |  |  |  |  |  |
| YLR350W | ORM2 |  |  |  |  |  |  |  |  |
| YLR351C | NIT3 |  |  |  |  |  |  |  |  |
| YLR352W | LUG1 |  |  |  |  |  |  |  |  |
| YLR353W | BUD8 |  |  |  |  |  |  |  |  |
| YLR354C | TAL1 |  |  |  |  |  |  |  |  |
| YLR355C | ILV5 |  |  |  |  |  |  |  |  |
| YLR356W | ATG33 |  |  |  |  |  |  |  |  |
| YLR357W | RSC2 |  |  |  |  |  |  |  |  |
| YLR359W | ADE13 |  |  |  |  |  |  |  |  |
| YLR360W | VPS38 |  |  |  |  |  |  |  |  |
| YLR361C | DCR2 |  |  |  |  |  |  |  |  |
| YLR362W | STE11 |  |  |  |  |  |  |  |  |
| YLR363W-A |  |  |  |  |  |  |  |  |  |
| YLR364W | GRX8 |  |  |  |  |  |  |  |  |
| YLR367W | RPS22B |  |  |  |  |  |  |  |  |
| YLR368W | MDM30 |  |  |  |  |  |  |  |  |
| YLR369W | SSQ1 |  |  |  |  |  |  |  |  |
| YLR370C | ARC18 |  |  |  |  |  |  |  |  |
| YLR371W | ROM2 |  |  |  |  |  |  |  |  |
| YLR372W | ELO3 |  |  |  |  |  |  |  |  |
| YLR373C | VID22 |  |  |  |  |  |  |  |  |
| YLR375W | STP3 |  |  |  |  |  |  |  |  |
| YLR376C | PSY3 |  |  |  |  |  |  |  |  |
| YLR377C | FBP1 |  |  |  |  |  |  |  |  |
| YLR378C | SEC61 |  |  |  |  |  |  |  |  |
| YLR380W | CSR1 |  |  |  |  |  |  |  |  |
| YLR381W | CTF3 |  |  |  |  |  |  |  |  |
| YLR382C | NAM2 |  |  |  |  |  |  |  |  |
| YLR383W | SMC6 |  |  |  |  |  |  |  |  |
| YLR384C | IKI3 |  |  |  |  |  |  |  |  |
| YLR385C | SWC7 |  |  |  |  |  |  |  |  |
| YLR386W | VAC14 |  |  |  |  |  |  |  |  |
| YLR387C | REH1 |  |  |  |  |  |  |  |  |
| YLR388W | RPS29A |  |  |  |  |  |  |  |  |
| YLR389C | STE23 |  |  |  |  |  |  |  |  |
| YLR390W | ECM19 |  |  |  |  |  |  |  |  |
| YLR390W-A | CCW14 |  |  |  |  |  |  |  |  |
| YLR392C | ART10 |  |  |  |  |  |  |  |  |
| YLR393W | ATP10 |  |  |  |  |  |  |  |  |
| YLR395C | COX8 |  |  |  |  |  |  |  |  |
| YLR396C | VPS33 |  |  |  |  |  |  |  |  |
| YLR397C | AFG2 |  |  |  |  |  |  |  |  |
| YLR398C | SKI2 |  |  |  |  |  |  |  |  |
| YLR399C | BDF1 |  |  |  |  |  |  |  |  |
| YLR401C | DUS3 |  |  |  |  |  |  |  |  |
| YLR403W | SFP1 |  |  |  |  |  |  |  |  |
| YLR404W | SEI1 |  |  |  |  |  |  |  |  |
| YLR405W | DUS4 |  |  |  |  |  |  |  |  |
| YLR406C | RPL31B |  |  |  |  |  |  |  |  |
| YLR408C | BLS1 |  |  |  |  |  |  |  |  |
| YLR409C | UTP21 |  |  |  |  |  |  |  |  |
| YLR410W | VIP1 |  |  |  |  |  |  |  |  |
| YLR412W | BER1 |  |  |  |  |  |  |  |  |
| YLR413W | INA1 |  |  |  |  |  |  |  |  |
| YLR417W | VPS36 |  |  |  |  |  |  |  |  |
| YLR418C | CDC73 |  |  |  |  |  |  |  |  |
| YLR420W | URA4 |  |  |  |  |  |  |  |  |
| YLR421C | RPN13 |  |  |  |  |  |  |  |  |
| YLR422W | DCK1 |  |  |  |  |  |  |  |  |
| YLR423C | ATG17 |  |  |  |  |  |  |  |  |
| YLR424W | SPP382 |  |  |  |  |  |  |  |  |
| YLR425W | TUS1 |  |  |  |  |  |  |  |  |
| YLR427W | MAG2 |  |  |  |  |  |  |  |  |
| YLR429W | CRN1 |  |  |  |  |  |  |  |  |
| YLR430W | SEN1 |  |  |  |  |  |  |  |  |
| YLR431C | ATG23 |  |  |  |  |  |  |  |  |
| YLR432W | IMD3 |  |  |  |  |  |  |  |  |
| YLR433C | CNA1 |  |  |  |  |  |  |  |  |
| YLR435W | TSR2 |  |  |  |  |  |  |  |  |
| YLR436C | ECM30 |  |  |  |  |  |  |  |  |
| YLR437C | DIF1 |  |  |  |  |  |  |  |  |
| YLR438C-A | LSM3 |  |  |  |  |  |  |  |  |
| YLR438W | CAR2 |  |  |  |  |  |  |  |  |
| YLR439W | MRPL4 |  |  |  |  |  |  |  |  |
| YLR440C | SEC39 |  |  |  |  |  |  |  |  |
| YLR441C | RPS1A |  |  |  |  |  |  |  |  |
| YLR442C | SIR3 |  |  |  |  |  |  |  |  |
| YLR443W | ECM7 |  |  |  |  |  |  |  |  |
| YLR445W | GMC2 |  |  |  |  |  |  |  |  |
| YLR446W | NGK1 |  |  |  |  |  |  |  |  |
| YLR447C | VMA6 |  |  |  |  |  |  |  |  |
| YLR448W | RPL6B |  |  |  |  |  |  |  |  |
| YLR449W | FPR4 |  |  |  |  |  |  |  |  |
| YLR450W | HMG2 |  |  |  |  |  |  |  |  |
| YLR451W | LEU3 |  |  |  |  |  |  |  |  |
| YLR452C | SST2 |  |  |  |  |  |  |  |  |
| YLR453C | RIF2 |  |  |  |  |  |  |  |  |
| YLR455W | PDP3 |  |  |  |  |  |  |  |  |
| YLR457C | NBP1 |  |  |  |  |  |  |  |  |
| YLR459W | GAB1 |  |  |  |  |  |  |  |  |
| YML001W | YPT7 |  |  |  |  |  |  |  |  |
| YML004C | GLO1 |  |  |  |  |  |  |  |  |
| YML005W | TRM12 |  |  |  |  |  |  |  |  |
| YML006C | GIS4 |  |  |  |  |  |  |  |  |
| YML007W | YAP1 |  |  |  |  |  |  |  |  |
| YML008C | ERG6 |  |  |  |  |  |  |  |  |
| YML009C | MRPL39 |  |  |  |  |  |  |  |  |
| YML010W | SPT5 |  |  |  |  |  |  |  |  |
| YML011C | RAD33 |  |  |  |  |  |  |  |  |
| YML012W | ERV25 |  |  |  |  |  |  |  |  |
| YML013W | UBX2 |  |  |  |  |  |  |  |  |
| YML014W | TRM9 |  |  |  |  |  |  |  |  |
| YML015C | TAF11 |  |  |  |  |  |  |  |  |
| YML016C | PPZ1 |  |  |  |  |  |  |  |  |
| YML017W | PSP2 |  |  |  |  |  |  |  |  |
| YML018C |  |  |  |  |  |  |  |  |  |
| YML019W | OST6 |  |  |  |  |  |  |  |  |
| YML021C | UNG1 |  |  |  |  |  |  |  |  |
| YML022W | APT1 |  |  |  |  |  |  |  |  |
| YML023C | NSE5 |  |  |  |  |  |  |  |  |
| YML024W | RPS17A |  |  |  |  |  |  |  |  |
| YML025C | YML6 |  |  |  |  |  |  |  |  |
| YML026C | RPS18B |  |  |  |  |  |  |  |  |
| YML027W | YOX1 |  |  |  |  |  |  |  |  |
| YML028W | TSA1 |  |  |  |  |  |  |  |  |
| YML029W | USA1 |  |  |  |  |  |  |  |  |
| YML030W | RCF1 |  |  |  |  |  |  |  |  |
| YML031W | NDC1 |  |  |  |  |  |  |  |  |
| YML032C | RAD52 |  |  |  |  |  |  |  |  |
| YML034W | SRC1 |  |  |  |  |  |  |  |  |
| YML035C | AMD1 |  |  |  |  |  |  |  |  |
| YML036W | CGI121 |  |  |  |  |  |  |  |  |
| YML038C | YMD8 |  |  |  |  |  |  |  |  |
| YML041C | VPS71 |  |  |  |  |  |  |  |  |
| YML042W | CAT2 |  |  |  |  |  |  |  |  |
| YML043C | RRN11 |  |  |  |  |  |  |  |  |
| YML046W | PRP39 |  |  |  |  |  |  |  |  |
| YML047C | PRM6 |  |  |  |  |  |  |  |  |
| YML048W | GSF2 |  |  |  |  |  |  |  |  |
| YML049C | RSE1 |  |  |  |  |  |  |  |  |
| YML050W | AIM32 |  |  |  |  |  |  |  |  |
| YML052W | SUR7 |  |  |  |  |  |  |  |  |
| YML055W | SPC2 |  |  |  |  |  |  |  |  |
| YML056C | IMD4 |  |  |  |  |  |  |  |  |
| YML057W | CMP2 |  |  |  |  |  |  |  |  |
| YML058W | SML1 |  |  |  |  |  |  |  |  |
| YML059C | NTE1 |  |  |  |  |  |  |  |  |
| YML060W | OGG1 |  |  |  |  |  |  |  |  |
| YML061C | PIF1 |  |  |  |  |  |  |  |  |
| YML062C | MFT1 |  |  |  |  |  |  |  |  |
| YML063W | RPS1B |  |  |  |  |  |  |  |  |
| YML064C | TEM1 |  |  |  |  |  |  |  |  |
| YML065W | ORC1 |  |  |  |  |  |  |  |  |
| YML066C | SMA2 |  |  |  |  |  |  |  |  |
| YML067C | ERV41 |  |  |  |  |  |  |  |  |
| YML069W | POB3 |  |  |  |  |  |  |  |  |
| YML070W | DAK1 |  |  |  |  |  |  |  |  |
| YML071C | COG8 |  |  |  |  |  |  |  |  |
| YML072C | TCB3 |  |  |  |  |  |  |  |  |
| YML073C | RPL6A |  |  |  |  |  |  |  |  |
| YML074C | FPR3 |  |  |  |  |  |  |  |  |
| YML075C | HMG1 |  |  |  |  |  |  |  |  |
| YML076C | WAR1 |  |  |  |  |  |  |  |  |
| YML077W | BET5 |  |  |  |  |  |  |  |  |
| YML078W | CPR3 |  |  |  |  |  |  |  |  |
| YML080W | DUS1 |  |  |  |  |  |  |  |  |
| YML081C-A | ATP18 |  |  |  |  |  |  |  |  |
| YML081W | TDA9 |  |  |  |  |  |  |  |  |
| YML083C |  |  |  |  |  |  |  |  |  |
| YML085C | TUB1 |  |  |  |  |  |  |  |  |
| YML086C | ALO1 |  |  |  |  |  |  |  |  |
| YML087C | AIM33 |  |  |  |  |  |  |  |  |
| YML088W | UFO1 |  |  |  |  |  |  |  |  |
| YML091C | RPM2 |  |  |  |  |  |  |  |  |
| YML092C | PRE8 |  |  |  |  |  |  |  |  |
| YML093W | UTP14 |  |  |  |  |  |  |  |  |
| YML094W | GIM5 |  |  |  |  |  |  |  |  |
| YML095C | RAD10 |  |  |  |  |  |  |  |  |
| YML097C | VPS9 |  |  |  |  |  |  |  |  |
| YML098W | TAF13 |  |  |  |  |  |  |  |  |
| YML099C | ARG81 |  |  |  |  |  |  |  |  |
| YML100W | TSL1 |  |  |  |  |  |  |  |  |
| YML101C | CUE4 |  |  |  |  |  |  |  |  |
| YML102W | CAC2 |  |  |  |  |  |  |  |  |
| YML103C | NUP188 |  |  |  |  |  |  |  |  |
| YML104C | MDM1 |  |  |  |  |  |  |  |  |
| YML105C | SEC65 |  |  |  |  |  |  |  |  |
| YML106W | URA5 |  |  |  |  |  |  |  |  |
| YML108W |  |  |  |  |  |  |  |  |  |
| YML109W | ZDS2 |  |  |  |  |  |  |  |  |
| YML110C | COQ5 |  |  |  |  |  |  |  |  |
| YML111W | BUL2 |  |  |  |  |  |  |  |  |
| YML112W | CTK3 |  |  |  |  |  |  |  |  |
| YML114C | TAF8 |  |  |  |  |  |  |  |  |
| YML115C | VAN1 |  |  |  |  |  |  |  |  |
| YML117W | NAB6 |  |  |  |  |  |  |  |  |
| YML118W | NGL3 |  |  |  |  |  |  |  |  |
| YML120C | NDI1 |  |  |  |  |  |  |  |  |
| YML121W | GTR1 |  |  |  |  |  |  |  |  |
| YML123C | PHO84 |  |  |  |  |  |  |  |  |
| YML124C | TUB3 |  |  |  |  |  |  |  |  |
| YML125C | PGA3 |  |  |  |  |  |  |  |  |
| YML126C | ERG13 |  |  |  |  |  |  |  |  |
| YML127W | RSC9 |  |  |  |  |  |  |  |  |
| YML128C | MSC1 |  |  |  |  |  |  |  |  |
| YML129C | COX14 |  |  |  |  |  |  |  |  |
| YML130C | ERO1 |  |  |  |  |  |  |  |  |
| YML131W |  |  |  |  |  |  |  |  |  |
| YMR001C | CDC5 |  |  |  |  |  |  |  |  |
| YMR002W | MIX17 |  |  |  |  |  |  |  |  |
| YMR003W | AIM34 |  |  |  |  |  |  |  |  |
| YMR004W | MVP1 |  |  |  |  |  |  |  |  |
| YMR005W | TAF4 |  |  |  |  |  |  |  |  |
| YMR009W | ADI1 |  |  |  |  |  |  |  |  |
| YMR010W | ANY1 |  |  |  |  |  |  |  |  |
| YMR011W | HXT2 |  |  |  |  |  |  |  |  |
| YMR012W | CLU1 |  |  |  |  |  |  |  |  |
| YMR013C | SEC59 |  |  |  |  |  |  |  |  |
| YMR014W | BUD22 |  |  |  |  |  |  |  |  |
| YMR015C | ERG5 |  |  |  |  |  |  |  |  |
| YMR016C | SOK2 |  |  |  |  |  |  |  |  |
| YMR020W | FMS1 |  |  |  |  |  |  |  |  |
| YMR021C | MAC1 |  |  |  |  |  |  |  |  |
| YMR022W | UBC7 |  |  |  |  |  |  |  |  |
| YMR023C | MSS1 |  |  |  |  |  |  |  |  |
| YMR024W | MRPL3 |  |  |  |  |  |  |  |  |
| YMR025W | CSI1 |  |  |  |  |  |  |  |  |
| YMR026C | PEX12 |  |  |  |  |  |  |  |  |
| YMR027W |  |  |  |  |  |  |  |  |  |
| YMR028W | TAP42 |  |  |  |  |  |  |  |  |
| YMR029C | FAR8 |  |  |  |  |  |  |  |  |
| YMR031C | EIS1 |  |  |  |  |  |  |  |  |
| YMR032W | HOF1 |  |  |  |  |  |  |  |  |
| YMR033W | ARP9 |  |  |  |  |  |  |  |  |
| YMR034C | RCH1 |  |  |  |  |  |  |  |  |
| YMR035W | IMP2 |  |  |  |  |  |  |  |  |
| YMR036C | MIH1 |  |  |  |  |  |  |  |  |
| YMR037C | MSN2 |  |  |  |  |  |  |  |  |
| YMR038C | CCS1 |  |  |  |  |  |  |  |  |
| YMR039C | SUB1 |  |  |  |  |  |  |  |  |
| YMR040W | YET2 |  |  |  |  |  |  |  |  |
| YMR041C | ARA2 |  |  |  |  |  |  |  |  |
| YMR042W | ARG80 |  |  |  |  |  |  |  |  |
| YMR043W | MCM1 |  |  |  |  |  |  |  |  |
| YMR044W | IOC4 |  |  |  |  |  |  |  |  |
| YMR047C | NUP116 |  |  |  |  |  |  |  |  |
| YMR048W | CSM3 |  |  |  |  |  |  |  |  |
| YMR049C | ERB1 |  |  |  |  |  |  |  |  |
| YMR052W | FAR3 |  |  |  |  |  |  |  |  |
| YMR053C | STB2 |  |  |  |  |  |  |  |  |
| YMR054W | STV1 |  |  |  |  |  |  |  |  |
| YMR055C | BUB2 |  |  |  |  |  |  |  |  |
| YMR058W | FET3 |  |  |  |  |  |  |  |  |
| YMR059W | SEN15 |  |  |  |  |  |  |  |  |
| YMR060C | SAM37 |  |  |  |  |  |  |  |  |
| YMR061W | RNA14 |  |  |  |  |  |  |  |  |
| YMR062C | ARG7 |  |  |  |  |  |  |  |  |
| YMR063W | RIM9 |  |  |  |  |  |  |  |  |
| YMR064W | AEP1 |  |  |  |  |  |  |  |  |
| YMR065W | KAR5 |  |  |  |  |  |  |  |  |
| YMR066W | SOV1 |  |  |  |  |  |  |  |  |
| YMR067C | UBX4 |  |  |  |  |  |  |  |  |
| YMR068W | AVO2 |  |  |  |  |  |  |  |  |
| YMR069W | NAT4 |  |  |  |  |  |  |  |  |
| YMR070W | MOT3 |  |  |  |  |  |  |  |  |
| YMR071C | TVP18 |  |  |  |  |  |  |  |  |
| YMR072W | ABF2 |  |  |  |  |  |  |  |  |
| YMR073C | IRC21 |  |  |  |  |  |  |  |  |
| YMR074C | SDD2 |  |  |  |  |  |  |  |  |
| YMR075W | RCO1 |  |  |  |  |  |  |  |  |
| YMR076C | PDS5 |  |  |  |  |  |  |  |  |
| YMR077C | VPS20 |  |  |  |  |  |  |  |  |
| YMR078C | CTF18 |  |  |  |  |  |  |  |  |
| YMR079W | SEC14 |  |  |  |  |  |  |  |  |
| YMR080C | NAM7 |  |  |  |  |  |  |  |  |
| YMR081C | ISF1 |  |  |  |  |  |  |  |  |
| YMR083W | ADH3 |  |  |  |  |  |  |  |  |
| YMR086W | SEG1 |  |  |  |  |  |  |  |  |
| YMR087W | PDL32 |  |  |  |  |  |  |  |  |
| YMR088C | VBA1 |  |  |  |  |  |  |  |  |
| YMR089C | YTA12 |  |  |  |  |  |  |  |  |
| YMR091C | NPL6 |  |  |  |  |  |  |  |  |
| YMR092C | AIP1 |  |  |  |  |  |  |  |  |
| YMR093W | UTP15 |  |  |  |  |  |  |  |  |
| YMR094W | CTF13 |  |  |  |  |  |  |  |  |
| YMR095C | SNO1 |  |  |  |  |  |  |  |  |
| YMR096W | SNZ1 |  |  |  |  |  |  |  |  |
| YMR097C | MTG1 |  |  |  |  |  |  |  |  |
| YMR098C | ATP25 |  |  |  |  |  |  |  |  |
| YMR099C |  |  |  |  |  |  |  |  |  |
| YMR100W | MUB1 |  |  |  |  |  |  |  |  |
| YMR101C | SRT1 |  |  |  |  |  |  |  |  |
| YMR102C | LAF1 |  |  |  |  |  |  |  |  |
| YMR104C | YPK2 |  |  |  |  |  |  |  |  |
| YMR105C | PGM2 |  |  |  |  |  |  |  |  |
| YMR106C | YKU80 |  |  |  |  |  |  |  |  |
| YMR107W | SPG4 |  |  |  |  |  |  |  |  |
| YMR108W | ILV2 |  |  |  |  |  |  |  |  |
| YMR109W | MYO5 |  |  |  |  |  |  |  |  |
| YMR111C | EUC1 |  |  |  |  |  |  |  |  |
| YMR112C | MED11 |  |  |  |  |  |  |  |  |
| YMR113W | FOL3 |  |  |  |  |  |  |  |  |
| YMR114C |  |  |  |  |  |  |  |  |  |
| YMR115W | MGR3 |  |  |  |  |  |  |  |  |
| YMR116C | ASC1 |  |  |  |  |  |  |  |  |
| YMR117C | SPC24 |  |  |  |  |  |  |  |  |
| YMR119W | ASI1 |  |  |  |  |  |  |  |  |
| YMR120C | ADE17 |  |  |  |  |  |  |  |  |
| YMR121C | RPL15B |  |  |  |  |  |  |  |  |
| YMR123W | PKR1 |  |  |  |  |  |  |  |  |
| YMR124W | EPO1 |  |  |  |  |  |  |  |  |
| YMR125W | STO1 |  |  |  |  |  |  |  |  |
| YMR127C | SAS2 |  |  |  |  |  |  |  |  |
| YMR128W | ECM16 |  |  |  |  |  |  |  |  |
| YMR129W | POM152 |  |  |  |  |  |  |  |  |
| YMR131C | RRB1 |  |  |  |  |  |  |  |  |
| YMR133W | REC114 |  |  |  |  |  |  |  |  |
| YMR134W | ERG29 |  |  |  |  |  |  |  |  |
| YMR135C | GID8 |  |  |  |  |  |  |  |  |
| YMR136W | GAT2 |  |  |  |  |  |  |  |  |
| YMR137C | PSO2 |  |  |  |  |  |  |  |  |
| YMR138W | CIN4 |  |  |  |  |  |  |  |  |
| YMR139W | RIM11 |  |  |  |  |  |  |  |  |
| YMR140W | SIP5 |  |  |  |  |  |  |  |  |
| YMR142C | RPL13B |  |  |  |  |  |  |  |  |
| YMR143W | RPS16A |  |  |  |  |  |  |  |  |
| YMR145C | NDE1 |  |  |  |  |  |  |  |  |
| YMR146C | TIF34 |  |  |  |  |  |  |  |  |
| YMR148W | LDO16 |  |  |  |  |  |  |  |  |
| YMR149W | SWP1 |  |  |  |  |  |  |  |  |
| YMR150C | IMP1 |  |  |  |  |  |  |  |  |
| YMR153W | NUP53 |  |  |  |  |  |  |  |  |
| YMR154C | RIM13 |  |  |  |  |  |  |  |  |
| YMR157C | AIM36 |  |  |  |  |  |  |  |  |
| YMR158W | MRPS8 |  |  |  |  |  |  |  |  |
| YMR159C | ATG16 |  |  |  |  |  |  |  |  |
| YMR160W | CVM1 |  |  |  |  |  |  |  |  |
| YMR161W | HLJ1 |  |  |  |  |  |  |  |  |
| YMR162C | DNF3 |  |  |  |  |  |  |  |  |
| YMR163C | INP2 |  |  |  |  |  |  |  |  |
| YMR164C | MSS11 |  |  |  |  |  |  |  |  |
| YMR165C | PAH1 |  |  |  |  |  |  |  |  |
| YMR166C | MME1 |  |  |  |  |  |  |  |  |
| YMR167W | MLH1 |  |  |  |  |  |  |  |  |
| YMR168C | CEP3 |  |  |  |  |  |  |  |  |
| YMR169C_170C | ALD3_ALD2 |  |  |  |  |  |  |  |  |
| YMR171C | EAR1 |  |  |  |  |  |  |  |  |
| YMR172W | HOT1 |  |  |  |  |  |  |  |  |
| YMR173W | DDR48 |  |  |  |  |  |  |  |  |
| YMR174C | PAI3 |  |  |  |  |  |  |  |  |
| YMR176W | ECM5 |  |  |  |  |  |  |  |  |
| YMR177W | MMT1 |  |  |  |  |  |  |  |  |
| YMR178W | FPY1 |  |  |  |  |  |  |  |  |
| YMR179W | SPT21 |  |  |  |  |  |  |  |  |
| YMR180C | CTL1 |  |  |  |  |  |  |  |  |
| YMR182C | RGM1 |  |  |  |  |  |  |  |  |
| YMR183C | SSO2 |  |  |  |  |  |  |  |  |
| YMR184W | ADD37 |  |  |  |  |  |  |  |  |
| YMR185W | RTP1 |  |  |  |  |  |  |  |  |
| YMR186W | HSC82 |  |  |  |  |  |  |  |  |
| YMR188C | MRPS17 |  |  |  |  |  |  |  |  |
| YMR190C | SGS1 |  |  |  |  |  |  |  |  |
| YMR192W | GYL1 |  |  |  |  |  |  |  |  |
| YMR193W | MRPL24 |  |  |  |  |  |  |  |  |
| YMR194C-B | CMC4 |  |  |  |  |  |  |  |  |
| YMR194W | RPL36A |  |  |  |  |  |  |  |  |
| YMR195W | ICY1 |  |  |  |  |  |  |  |  |
| YMR197C | VTI1 |  |  |  |  |  |  |  |  |
| YMR198W | CIK1 |  |  |  |  |  |  |  |  |
| YMR199W | CLN1 |  |  |  |  |  |  |  |  |
| YMR200W | ROT1 |  |  |  |  |  |  |  |  |
| YMR201C | RAD14 |  |  |  |  |  |  |  |  |
| YMR202W | ERG2 |  |  |  |  |  |  |  |  |
| YMR203W | TOM40 |  |  |  |  |  |  |  |  |
| YMR204C | INP1 |  |  |  |  |  |  |  |  |
| YMR205C | PFK2 |  |  |  |  |  |  |  |  |
| YMR207C | HFA1 |  |  |  |  |  |  |  |  |
| YMR208W | ERG12 |  |  |  |  |  |  |  |  |
| YMR210W | MGL2 |  |  |  |  |  |  |  |  |
| YMR211W | DML1 |  |  |  |  |  |  |  |  |
| YMR212C | EFR3 |  |  |  |  |  |  |  |  |
| YMR213W | CEF1 |  |  |  |  |  |  |  |  |
| YMR214W | SCJ1 |  |  |  |  |  |  |  |  |
| YMR215W | GAS3 |  |  |  |  |  |  |  |  |
| YMR216C | SKY1 |  |  |  |  |  |  |  |  |
| YMR217W | GUA1 |  |  |  |  |  |  |  |  |
| YMR218C | TRS130 |  |  |  |  |  |  |  |  |
| YMR219W | ESC1 |  |  |  |  |  |  |  |  |
| YMR220W | ERG8 |  |  |  |  |  |  |  |  |
| YMR222C | FSH2 |  |  |  |  |  |  |  |  |
| YMR223W | UBP8 |  |  |  |  |  |  |  |  |
| YMR224C | MRE11 |  |  |  |  |  |  |  |  |
| YMR225C | MRPL44 |  |  |  |  |  |  |  |  |
| YMR226C | ORA1 |  |  |  |  |  |  |  |  |
| YMR227C | TAF7 |  |  |  |  |  |  |  |  |
| YMR228W | MTF1 |  |  |  |  |  |  |  |  |
| YMR229C | RRP5 |  |  |  |  |  |  |  |  |
| YMR230W | RPS10B |  |  |  |  |  |  |  |  |
| YMR231W | PEP5 |  |  |  |  |  |  |  |  |
| YMR232W | FUS2 |  |  |  |  |  |  |  |  |
| YMR233W | TRI1 |  |  |  |  |  |  |  |  |
| YMR234W | RNH1 |  |  |  |  |  |  |  |  |
| YMR235C | RNA1 |  |  |  |  |  |  |  |  |
| YMR236W | TAF9 |  |  |  |  |  |  |  |  |
| YMR237W | BCH1 |  |  |  |  |  |  |  |  |
| YMR238W | DFG5 |  |  |  |  |  |  |  |  |
| YMR239C | RNT1 |  |  |  |  |  |  |  |  |
| YMR240C | CUS1 |  |  |  |  |  |  |  |  |
| YMR241W | YHM2 |  |  |  |  |  |  |  |  |
| YMR242C | RPL20A |  |  |  |  |  |  |  |  |
| YMR243C | ZRC1 |  |  |  |  |  |  |  |  |
| YMR244C-A | COA6 |  |  |  |  |  |  |  |  |
| YMR246W | FAA4 |  |  |  |  |  |  |  |  |
| YMR247C | RKR1 |  |  |  |  |  |  |  |  |
| YMR251W-A | HOR7 |  |  |  |  |  |  |  |  |
| YMR255W | GFD1 |  |  |  |  |  |  |  |  |
| YMR256C | COX7 |  |  |  |  |  |  |  |  |
| YMR257C | PET111 |  |  |  |  |  |  |  |  |
| YMR258C | ROY1 |  |  |  |  |  |  |  |  |
| YMR259C | TRM732 |  |  |  |  |  |  |  |  |
| YMR260C | TIF11 |  |  |  |  |  |  |  |  |
| YMR261C | TPS3 |  |  |  |  |  |  |  |  |
| YMR263W | SAP30 |  |  |  |  |  |  |  |  |
| YMR264W | CUE1 |  |  |  |  |  |  |  |  |
| YMR266W | RSN1 |  |  |  |  |  |  |  |  |
| YMR267W | PPA2 |  |  |  |  |  |  |  |  |
| YMR268C | PRP24 |  |  |  |  |  |  |  |  |
| YMR269W | TMA23 |  |  |  |  |  |  |  |  |
| YMR270C | RRN9 |  |  |  |  |  |  |  |  |
| YMR271C | URA10 |  |  |  |  |  |  |  |  |
| YMR272C | SCS7 |  |  |  |  |  |  |  |  |
| YMR273C | ZDS1 |  |  |  |  |  |  |  |  |
| YMR274C | RCE1 |  |  |  |  |  |  |  |  |
| YMR275C | BUL1 |  |  |  |  |  |  |  |  |
| YMR276W | DSK2 |  |  |  |  |  |  |  |  |
| YMR277W | FCP1 |  |  |  |  |  |  |  |  |
| YMR278W | PRM15 |  |  |  |  |  |  |  |  |
| YMR280C | CAT8 |  |  |  |  |  |  |  |  |
| YMR281W | GPI12 |  |  |  |  |  |  |  |  |
| YMR282C | AEP2 |  |  |  |  |  |  |  |  |
| YMR283C | RIT1 |  |  |  |  |  |  |  |  |
| YMR284W | YKU70 |  |  |  |  |  |  |  |  |
| YMR285C | NGL2 |  |  |  |  |  |  |  |  |
| YMR286W | MRPL33 |  |  |  |  |  |  |  |  |
| YMR287C | DSS1 |  |  |  |  |  |  |  |  |
| YMR288W | HSH155 |  |  |  |  |  |  |  |  |
| YMR289W | ABZ2 |  |  |  |  |  |  |  |  |
| YMR290C | HAS1 |  |  |  |  |  |  |  |  |
| YMR291W | TDA1 |  |  |  |  |  |  |  |  |
| YMR292W | GOT1 |  |  |  |  |  |  |  |  |
| YMR293C | HER2 |  |  |  |  |  |  |  |  |
| YMR294W | JNM1 |  |  |  |  |  |  |  |  |
| YMR295C | GSR1 |  |  |  |  |  |  |  |  |
| YMR296C | LCB1 |  |  |  |  |  |  |  |  |
| YMR297W | PRC1 |  |  |  |  |  |  |  |  |
| YMR298W | LIP1 |  |  |  |  |  |  |  |  |
| YMR299C | DYN3 |  |  |  |  |  |  |  |  |
| YMR300C | ADE4 |  |  |  |  |  |  |  |  |
| YMR301C | ATM1 |  |  |  |  |  |  |  |  |
| YMR302C | YME2 |  |  |  |  |  |  |  |  |
| YMR303C | ADH2 |  |  |  |  |  |  |  |  |
| YMR304W | UBP15 |  |  |  |  |  |  |  |  |
| YMR305C | SCW10 |  |  |  |  |  |  |  |  |
| YMR306W | FKS3 |  |  |  |  |  |  |  |  |
| YMR307W | GAS1 |  |  |  |  |  |  |  |  |
| YMR308C | PSE1 |  |  |  |  |  |  |  |  |
| YMR309C | NIP1 |  |  |  |  |  |  |  |  |
| YMR311C | GLC8 |  |  |  |  |  |  |  |  |
| YMR312W | ELP6 |  |  |  |  |  |  |  |  |
| YMR313C | TGL3 |  |  |  |  |  |  |  |  |
| YMR314W | PRE5 |  |  |  |  |  |  |  |  |
| YMR315W |  |  |  |  |  |  |  |  |  |
| YNL001W | DOM34 |  |  |  |  |  |  |  |  |
| YNL002C | RLP7 |  |  |  |  |  |  |  |  |
| YNL003C | PET8 |  |  |  |  |  |  |  |  |
| YNL004W | HRB1 |  |  |  |  |  |  |  |  |
| YNL005C | MRP7 |  |  |  |  |  |  |  |  |
| YNL006W | LST8 |  |  |  |  |  |  |  |  |
| YNL007C | SIS1 |  |  |  |  |  |  |  |  |
| YNL008C | ASI3 |  |  |  |  |  |  |  |  |
| YNL009W | IDP3 |  |  |  |  |  |  |  |  |
| YNL010W | PYP1 |  |  |  |  |  |  |  |  |
| YNL012W | SPO1 |  |  |  |  |  |  |  |  |
| YNL014W | HEF3 |  |  |  |  |  |  |  |  |
| YNL015W | PBI2 |  |  |  |  |  |  |  |  |
| YNL016W | PUB1 |  |  |  |  |  |  |  |  |
| YNL020C | ARK1 |  |  |  |  |  |  |  |  |
| YNL021W | HDA1 |  |  |  |  |  |  |  |  |
| YNL022C | RCM1 |  |  |  |  |  |  |  |  |
| YNL023C | FAP1 |  |  |  |  |  |  |  |  |
| YNL024C-A | KSH1 |  |  |  |  |  |  |  |  |
| YNL025C | SSN8 |  |  |  |  |  |  |  |  |
| YNL026W | SAM50 |  |  |  |  |  |  |  |  |
| YNL027W | CRZ1 |  |  |  |  |  |  |  |  |
| YNL030W | HHF2 |  |  |  |  |  |  |  |  |
| YNL031C | HHT2 |  |  |  |  |  |  |  |  |
| YNL032W | SIW14 |  |  |  |  |  |  |  |  |
| YNL035C |  |  |  |  |  |  |  |  |  |
| YNL036W | NCE103 |  |  |  |  |  |  |  |  |
| YNL037C | IDH1 |  |  |  |  |  |  |  |  |
| YNL038W | GPI15 |  |  |  |  |  |  |  |  |
| YNL039W | BDP1 |  |  |  |  |  |  |  |  |
| YNL041C | COG6 |  |  |  |  |  |  |  |  |
| YNL042W | BOP3 |  |  |  |  |  |  |  |  |
| YNL044W | YIP3 |  |  |  |  |  |  |  |  |
| YNL045W | LAP2 |  |  |  |  |  |  |  |  |
| YNL047C | SLM2 |  |  |  |  |  |  |  |  |
| YNL048W | ALG11 |  |  |  |  |  |  |  |  |
| YNL049C | SFB2 |  |  |  |  |  |  |  |  |
| YNL051W | COG5 |  |  |  |  |  |  |  |  |
| YNL052W | COX5A |  |  |  |  |  |  |  |  |
| YNL053W | MSG5 |  |  |  |  |  |  |  |  |
| YNL054W | VAC7 |  |  |  |  |  |  |  |  |
| YNL055C | POR1 |  |  |  |  |  |  |  |  |
| YNL056W | OCA2 |  |  |  |  |  |  |  |  |
| YNL059C | ARP5 |  |  |  |  |  |  |  |  |
| YNL061W | NOP2 |  |  |  |  |  |  |  |  |
| YNL062C | GCD10 |  |  |  |  |  |  |  |  |
| YNL063W | MTQ1 |  |  |  |  |  |  |  |  |
| YNL064C | YDJ1 |  |  |  |  |  |  |  |  |
| YNL065W | AQR1 |  |  |  |  |  |  |  |  |
| YNL066W | SUN4 |  |  |  |  |  |  |  |  |
| YNL067W | RPL9B |  |  |  |  |  |  |  |  |
| YNL068C | FKH2 |  |  |  |  |  |  |  |  |
| YNL069C | RPL16B |  |  |  |  |  |  |  |  |
| YNL070W | TOM7 |  |  |  |  |  |  |  |  |
| YNL071W | LAT1 |  |  |  |  |  |  |  |  |
| YNL072W | RNH201 |  |  |  |  |  |  |  |  |
| YNL073W | MSK1 |  |  |  |  |  |  |  |  |
| YNL074C | MLF3 |  |  |  |  |  |  |  |  |
| YNL075W | IMP4 |  |  |  |  |  |  |  |  |
| YNL076W | MKS1 |  |  |  |  |  |  |  |  |
| YNL077W | APJ1 |  |  |  |  |  |  |  |  |
| YNL078W | NIS1 |  |  |  |  |  |  |  |  |
| YNL079C | TPM1 |  |  |  |  |  |  |  |  |
| YNL080C | EOS1 |  |  |  |  |  |  |  |  |
| YNL081C | SWS2 |  |  |  |  |  |  |  |  |
| YNL082W | PMS1 |  |  |  |  |  |  |  |  |
| YNL083W | SAL1 |  |  |  |  |  |  |  |  |
| YNL084C | END3 |  |  |  |  |  |  |  |  |
| YNL085W | MKT1 |  |  |  |  |  |  |  |  |
| YNL086W | SNN1 |  |  |  |  |  |  |  |  |
| YNL087W | TCB2 |  |  |  |  |  |  |  |  |
| YNL088W | TOP2 |  |  |  |  |  |  |  |  |
| YNL090W | RHO2 |  |  |  |  |  |  |  |  |
| YNL091W | NST1 |  |  |  |  |  |  |  |  |
| YNL092W |  |  |  |  |  |  |  |  |  |
| YNL093W | YPT53 |  |  |  |  |  |  |  |  |
| YNL094W | APP1 |  |  |  |  |  |  |  |  |
| YNL096C | RPS7B |  |  |  |  |  |  |  |  |
| YNL097C | PHO23 |  |  |  |  |  |  |  |  |
| YNL098C | RAS2 |  |  |  |  |  |  |  |  |
| YNL099C | OCA1 |  |  |  |  |  |  |  |  |
| YNL100W | MIC27 |  |  |  |  |  |  |  |  |
| YNL101W | AVT4 |  |  |  |  |  |  |  |  |
| YNL102W | POL1 |  |  |  |  |  |  |  |  |
| YNL103W | MET4 |  |  |  |  |  |  |  |  |
| YNL104C | LEU4 |  |  |  |  |  |  |  |  |
| YNL106C | INP52 |  |  |  |  |  |  |  |  |
| YNL107W | YAF9 |  |  |  |  |  |  |  |  |
| YNL108C |  |  |  |  |  |  |  |  |  |
| YNL110C | NOP15 |  |  |  |  |  |  |  |  |
| YNL111C | CYB5 |  |  |  |  |  |  |  |  |
| YNL112W | DBP2 |  |  |  |  |  |  |  |  |
| YNL113W | RPC19 |  |  |  |  |  |  |  |  |
| YNL116W | DMA2 |  |  |  |  |  |  |  |  |
| YNL117W | MLS1 |  |  |  |  |  |  |  |  |
| YNL118C | DCP2 |  |  |  |  |  |  |  |  |
| YNL119W | NCS2 |  |  |  |  |  |  |  |  |
| YNL121C | TOM70 |  |  |  |  |  |  |  |  |
| YNL122C | MRP35 |  |  |  |  |  |  |  |  |
| YNL123W | NMA111 |  |  |  |  |  |  |  |  |
| YNL124W | NAF1 |  |  |  |  |  |  |  |  |
| YNL126W | SPC98 |  |  |  |  |  |  |  |  |
| YNL127W | FAR11 |  |  |  |  |  |  |  |  |
| YNL128W | TEP1 |  |  |  |  |  |  |  |  |
| YNL129W | NRK1 |  |  |  |  |  |  |  |  |
| YNL130C | CPT1 |  |  |  |  |  |  |  |  |
| YNL131W | TOM22 |  |  |  |  |  |  |  |  |
| YNL132W | KRE33 |  |  |  |  |  |  |  |  |
| YNL133C | FYV6 |  |  |  |  |  |  |  |  |
| YNL134C |  |  |  |  |  |  |  |  |  |
| YNL135C | FPR1 |  |  |  |  |  |  |  |  |
| YNL136W | EAF7 |  |  |  |  |  |  |  |  |
| YNL137C | NAM9 |  |  |  |  |  |  |  |  |
| YNL138W | SRV2 |  |  |  |  |  |  |  |  |
| YNL138W-A | YSF3 |  |  |  |  |  |  |  |  |
| YNL139C | THO2 |  |  |  |  |  |  |  |  |
| YNL142W | MEP2 |  |  |  |  |  |  |  |  |
| YNL145W | MFA2 |  |  |  |  |  |  |  |  |
| YNL147W | LSM7 |  |  |  |  |  |  |  |  |
| YNL148C | ALF1 |  |  |  |  |  |  |  |  |
| YNL149C | PGA2 |  |  |  |  |  |  |  |  |
| YNL151C | RPC31 |  |  |  |  |  |  |  |  |
| YNL152W | INN1 |  |  |  |  |  |  |  |  |
| YNL153C | GIM3 |  |  |  |  |  |  |  |  |
| YNL154C | YCK2 |  |  |  |  |  |  |  |  |
| YNL155W | CUZ1 |  |  |  |  |  |  |  |  |
| YNL156C | NSG2 |  |  |  |  |  |  |  |  |
| YNL157W | IGO1 |  |  |  |  |  |  |  |  |
| YNL158W | PGA1 |  |  |  |  |  |  |  |  |
| YNL159C | ASI2 |  |  |  |  |  |  |  |  |
| YNL161W | CBK1 |  |  |  |  |  |  |  |  |
| YNL162W | RPL42A |  |  |  |  |  |  |  |  |
| YNL163C | RIA1 |  |  |  |  |  |  |  |  |
| YNL164C | IBD2 |  |  |  |  |  |  |  |  |
| YNL166C | BNI5 |  |  |  |  |  |  |  |  |
| YNL167C | SKO1 |  |  |  |  |  |  |  |  |
| YNL169C | PSD1 |  |  |  |  |  |  |  |  |
| YNL172W | APC1 |  |  |  |  |  |  |  |  |
| YNL173C | MDG1 |  |  |  |  |  |  |  |  |
| YNL175C | NOP13 |  |  |  |  |  |  |  |  |
| YNL176C | TDA7 |  |  |  |  |  |  |  |  |
| YNL177C | MRPL22 |  |  |  |  |  |  |  |  |
| YNL178W | RPS3 |  |  |  |  |  |  |  |  |
| YNL180C | RHO5 |  |  |  |  |  |  |  |  |
| YNL182C | IPI3 |  |  |  |  |  |  |  |  |
| YNL183C | NPR1 |  |  |  |  |  |  |  |  |
| YNL185C | MRPL19 |  |  |  |  |  |  |  |  |
| YNL186W | UBP10 |  |  |  |  |  |  |  |  |
| YNL187W | SWT21 |  |  |  |  |  |  |  |  |
| YNL188W | KAR1 |  |  |  |  |  |  |  |  |
| YNL189W | SRP1 |  |  |  |  |  |  |  |  |
| YNL191W | DUG3 |  |  |  |  |  |  |  |  |
| YNL192W | CHS1 |  |  |  |  |  |  |  |  |
| YNL194C |  |  |  |  |  |  |  |  |  |
| YNL197C | WHI3 |  |  |  |  |  |  |  |  |
| YNL199C | GCR2 |  |  |  |  |  |  |  |  |
| YNL200C | NNR1 |  |  |  |  |  |  |  |  |
| YNL201C | PSY2 |  |  |  |  |  |  |  |  |
| YNL202W | SPS19 |  |  |  |  |  |  |  |  |
| YNL204C | SPS18 |  |  |  |  |  |  |  |  |
| YNL206C | RTT106 |  |  |  |  |  |  |  |  |
| YNL207W | RIO2 |  |  |  |  |  |  |  |  |
| YNL209W | SSB2 |  |  |  |  |  |  |  |  |
| YNL210W | MER1 |  |  |  |  |  |  |  |  |
| YNL212W | VID27 |  |  |  |  |  |  |  |  |
| YNL213C | RRG9 |  |  |  |  |  |  |  |  |
| YNL214W | PEX17 |  |  |  |  |  |  |  |  |
| YNL215W | IES2 |  |  |  |  |  |  |  |  |
| YNL216W | RAP1 |  |  |  |  |  |  |  |  |
| YNL217W | PPN2 |  |  |  |  |  |  |  |  |
| YNL218W | MGS1 |  |  |  |  |  |  |  |  |
| YNL219C | ALG9 |  |  |  |  |  |  |  |  |
| YNL220W | ADE12 |  |  |  |  |  |  |  |  |
| YNL221C | POP1 |  |  |  |  |  |  |  |  |
| YNL222W | SSU72 |  |  |  |  |  |  |  |  |
| YNL223W | ATG4 |  |  |  |  |  |  |  |  |
| YNL224C | SQS1 |  |  |  |  |  |  |  |  |
| YNL225C | CNM67 |  |  |  |  |  |  |  |  |
| YNL227C | JJJ1 |  |  |  |  |  |  |  |  |
| YNL229C | URE2 |  |  |  |  |  |  |  |  |
| YNL230C | ELA1 |  |  |  |  |  |  |  |  |
| YNL231C | PDR16 |  |  |  |  |  |  |  |  |
| YNL232W | CSL4 |  |  |  |  |  |  |  |  |
| YNL233W | BNI4 |  |  |  |  |  |  |  |  |
| YNL234W |  |  |  |  |  |  |  |  |  |
| YNL236W | SIN4 |  |  |  |  |  |  |  |  |
| YNL238W | KEX2 |  |  |  |  |  |  |  |  |
| YNL239W | LAP3 |  |  |  |  |  |  |  |  |
| YNL240C | NAR1 |  |  |  |  |  |  |  |  |
| YNL241C | ZWF1 |  |  |  |  |  |  |  |  |
| YNL242W | ATG2 |  |  |  |  |  |  |  |  |
| YNL244C | SUI1 |  |  |  |  |  |  |  |  |
| YNL245C | CWC25 |  |  |  |  |  |  |  |  |
| YNL246W | VPS75 |  |  |  |  |  |  |  |  |
| YNL247W | CRS1 |  |  |  |  |  |  |  |  |
| YNL248C | RPA49 |  |  |  |  |  |  |  |  |
| YNL249C | MPA43 |  |  |  |  |  |  |  |  |
| YNL250W | RAD50 |  |  |  |  |  |  |  |  |
| YNL251C | NRD1 |  |  |  |  |  |  |  |  |
| YNL252C | MRPL17 |  |  |  |  |  |  |  |  |
| YNL253W | TEX1 |  |  |  |  |  |  |  |  |
| YNL254C | RTC4 |  |  |  |  |  |  |  |  |
| YNL255C | GIS2 |  |  |  |  |  |  |  |  |
| YNL256W | FOL1 |  |  |  |  |  |  |  |  |
| YNL257C | SIP3 |  |  |  |  |  |  |  |  |
| YNL258C | DSL1 |  |  |  |  |  |  |  |  |
| YNL259C | ATX1 |  |  |  |  |  |  |  |  |
| YNL260C | LTO1 |  |  |  |  |  |  |  |  |
| YNL261W | ORC5 |  |  |  |  |  |  |  |  |
| YNL262W | POL2 |  |  |  |  |  |  |  |  |
| YNL263C | YIF1 |  |  |  |  |  |  |  |  |
| YNL264C | PDR17 |  |  |  |  |  |  |  |  |
| YNL265C | IST1 |  |  |  |  |  |  |  |  |
| YNL267W | PIK1 |  |  |  |  |  |  |  |  |
| YNL268W | LYP1 |  |  |  |  |  |  |  |  |
| YNL270C | ALP1 |  |  |  |  |  |  |  |  |
| YNL271C | BNI1 |  |  |  |  |  |  |  |  |
| YNL272C | SEC2 |  |  |  |  |  |  |  |  |
| YNL273W | TOF1 |  |  |  |  |  |  |  |  |
| YNL274C | GOR1 |  |  |  |  |  |  |  |  |
| YNL275W | BOR1 |  |  |  |  |  |  |  |  |
| YNL277W | MET2 |  |  |  |  |  |  |  |  |
| YNL278W | CAF120 |  |  |  |  |  |  |  |  |
| YNL279W | PRM1 |  |  |  |  |  |  |  |  |
| YNL280C | ERG24 |  |  |  |  |  |  |  |  |
| YNL281W | HCH1 |  |  |  |  |  |  |  |  |
| YNL282W | POP3 |  |  |  |  |  |  |  |  |
| YNL283C | WSC2 |  |  |  |  |  |  |  |  |
| YNL284C | MRPL10 |  |  |  |  |  |  |  |  |
| YNL287W | SEC21 |  |  |  |  |  |  |  |  |
| YNL288W | CAF40 |  |  |  |  |  |  |  |  |
| YNL289W | PCL1 |  |  |  |  |  |  |  |  |
| YNL290W | RFC3 |  |  |  |  |  |  |  |  |
| YNL291C | MID1 |  |  |  |  |  |  |  |  |
| YNL292W | PUS4 |  |  |  |  |  |  |  |  |
| YNL293W | MSB3 |  |  |  |  |  |  |  |  |
| YNL294C | RIM21 |  |  |  |  |  |  |  |  |
| YNL295W | MRX6 |  |  |  |  |  |  |  |  |
| YNL297C | MON2 |  |  |  |  |  |  |  |  |
| YNL298W | CLA4 |  |  |  |  |  |  |  |  |
| YNL299W | TRF5 |  |  |  |  |  |  |  |  |
| YNL301C | RPL18B |  |  |  |  |  |  |  |  |
| YNL302C | RPS19B |  |  |  |  |  |  |  |  |
| YNL304W | YPT11 |  |  |  |  |  |  |  |  |
| YNL305C | BXI1 |  |  |  |  |  |  |  |  |
| YNL306W | MRPS18 |  |  |  |  |  |  |  |  |
| YNL307C | MCK1 |  |  |  |  |  |  |  |  |
| YNL308C | KRI1 |  |  |  |  |  |  |  |  |
| YNL309W | STB1 |  |  |  |  |  |  |  |  |
| YNL310C | ZIM17 |  |  |  |  |  |  |  |  |
| YNL311C | SKP2 |  |  |  |  |  |  |  |  |
| YNL312W | RFA2 |  |  |  |  |  |  |  |  |
| YNL313C | EMW1 |  |  |  |  |  |  |  |  |
| YNL314W | DAL82 |  |  |  |  |  |  |  |  |
| YNL315C | ATP11 |  |  |  |  |  |  |  |  |
| YNL316C | PHA2 |  |  |  |  |  |  |  |  |
| YNL317W | PFS2 |  |  |  |  |  |  |  |  |
| YNL318C | HXT14 |  |  |  |  |  |  |  |  |
| YNL321W | VNX1 |  |  |  |  |  |  |  |  |
| YNL322C | KRE1 |  |  |  |  |  |  |  |  |
| YNL323W | LEM3 |  |  |  |  |  |  |  |  |
| YNL325C | FIG4 |  |  |  |  |  |  |  |  |
| YNL326C | PFA3 |  |  |  |  |  |  |  |  |
| YNL328C | MDJ2 |  |  |  |  |  |  |  |  |
| YNL329C | PEX6 |  |  |  |  |  |  |  |  |
| YNL330C | RPD3 |  |  |  |  |  |  |  |  |
| YNR001C | CIT1 |  |  |  |  |  |  |  |  |
| YNR002C | ATO2 |  |  |  |  |  |  |  |  |
| YNR003C | RPC34 |  |  |  |  |  |  |  |  |
| YNR006W | VPS27 |  |  |  |  |  |  |  |  |
| YNR007C | ATG3 |  |  |  |  |  |  |  |  |
| YNR008W | LRO1 |  |  |  |  |  |  |  |  |
| YNR009W | NRM1 |  |  |  |  |  |  |  |  |
| YNR010W | CSE2 |  |  |  |  |  |  |  |  |
| YNR011C | PRP2 |  |  |  |  |  |  |  |  |
| YNR012W | URK1 |  |  |  |  |  |  |  |  |
| YNR013C | PHO91 |  |  |  |  |  |  |  |  |
| YNR015W | SMM1 |  |  |  |  |  |  |  |  |
| YNR016C | ACC1 |  |  |  |  |  |  |  |  |
| YNR017W | TIM23 |  |  |  |  |  |  |  |  |
| YNR018W | RCF2 |  |  |  |  |  |  |  |  |
| YNR019W | ARE2 |  |  |  |  |  |  |  |  |
| YNR020C | ATP23 |  |  |  |  |  |  |  |  |
| YNR022C | MRPL50 |  |  |  |  |  |  |  |  |
| YNR023W | SNF12 |  |  |  |  |  |  |  |  |
| YNR024W | MPP6 |  |  |  |  |  |  |  |  |
| YNR026C | SEC12 |  |  |  |  |  |  |  |  |
| YNR027W | BUD17 |  |  |  |  |  |  |  |  |
| YNR028W | CPR8 |  |  |  |  |  |  |  |  |
| YNR029C | ZNG1 |  |  |  |  |  |  |  |  |
| YNR030W | ALG12 |  |  |  |  |  |  |  |  |
| YNR031C | SSK2 |  |  |  |  |  |  |  |  |
| YNR032C-A | HUB1 |  |  |  |  |  |  |  |  |
| YNR032W | PPG1 |  |  |  |  |  |  |  |  |
| YNR033W | ABZ1 |  |  |  |  |  |  |  |  |
| YNR034W | SOL1 |  |  |  |  |  |  |  |  |
| YNR034W-A | EGO4 |  |  |  |  |  |  |  |  |
| YNR035C | ARC35 |  |  |  |  |  |  |  |  |
| YNR036C | MRPS12 |  |  |  |  |  |  |  |  |
| YNR037C | RSM19 |  |  |  |  |  |  |  |  |
| YNR038W | DBP6 |  |  |  |  |  |  |  |  |
| YNR039C | ZRG17 |  |  |  |  |  |  |  |  |
| YNR040W | MRX15 |  |  |  |  |  |  |  |  |
| YNR041C | COQ2 |  |  |  |  |  |  |  |  |
| YNR043W | MVD1 |  |  |  |  |  |  |  |  |
| YNR045W | PET494 |  |  |  |  |  |  |  |  |
| YNR046W | TRM112 |  |  |  |  |  |  |  |  |
| YNR047W | FPK1 |  |  |  |  |  |  |  |  |
| YNR048W |  |  |  |  |  |  |  |  |  |
| YNR049C | MSO1 |  |  |  |  |  |  |  |  |
| YNR050C | LYS9 |  |  |  |  |  |  |  |  |
| YNR051C | BRE5 |  |  |  |  |  |  |  |  |
| YNR052C | POP2 |  |  |  |  |  |  |  |  |
| YNR053C | NOG2 |  |  |  |  |  |  |  |  |
| YNR054C | ESF2 |  |  |  |  |  |  |  |  |
| YNR055C | HOL1 |  |  |  |  |  |  |  |  |
| YNR060W | FRE4 |  |  |  |  |  |  |  |  |
| YNR067C | DSE4 |  |  |  |  |  |  |  |  |
| YNR070W | PDR18 |  |  |  |  |  |  |  |  |
| YOL001W | PHO80 |  |  |  |  |  |  |  |  |
| YOL002C | IZH2 |  |  |  |  |  |  |  |  |
| YOL003C | PFA4 |  |  |  |  |  |  |  |  |
| YOL004W | SIN3 |  |  |  |  |  |  |  |  |
| YOL005C | RPB11 |  |  |  |  |  |  |  |  |
| YOL006C | TOP1 |  |  |  |  |  |  |  |  |
| YOL007C | CSI2 |  |  |  |  |  |  |  |  |
| YOL008W | COQ10 |  |  |  |  |  |  |  |  |
| YOL009C | MDM12 |  |  |  |  |  |  |  |  |
| YOL010W | RCL1 |  |  |  |  |  |  |  |  |
| YOL012C | HTZ1 |  |  |  |  |  |  |  |  |
| YOL013C | HRD1 |  |  |  |  |  |  |  |  |
| YOL016C | CMK2 |  |  |  |  |  |  |  |  |
| YOL017W | ESC8 |  |  |  |  |  |  |  |  |
| YOL018C | TLG2 |  |  |  |  |  |  |  |  |
| YOL019W | TOS7 |  |  |  |  |  |  |  |  |
| YOL020W | TAT2 |  |  |  |  |  |  |  |  |
| YOL021C | DIS3 |  |  |  |  |  |  |  |  |
| YOL022C | TSR4 |  |  |  |  |  |  |  |  |
| YOL023W | IFM1 |  |  |  |  |  |  |  |  |
| YOL026C | MIM1 |  |  |  |  |  |  |  |  |
| YOL027C | MDM38 |  |  |  |  |  |  |  |  |
| YOL030W | GAS5 |  |  |  |  |  |  |  |  |
| YOL031C | SIL1 |  |  |  |  |  |  |  |  |
| YOL032W | OPI10 |  |  |  |  |  |  |  |  |
| YOL033W | MSE1 |  |  |  |  |  |  |  |  |
| YOL034W | SMC5 |  |  |  |  |  |  |  |  |
| YOL038W | PRE6 |  |  |  |  |  |  |  |  |
| YOL039W | RPP2A |  |  |  |  |  |  |  |  |
| YOL040C | RPS15 |  |  |  |  |  |  |  |  |
| YOL041C | NOP12 |  |  |  |  |  |  |  |  |
| YOL042W | NGL1 |  |  |  |  |  |  |  |  |
| YOL043C | NTG2 |  |  |  |  |  |  |  |  |
| YOL044W | PEX15 |  |  |  |  |  |  |  |  |
| YOL045W | PSK2 |  |  |  |  |  |  |  |  |
| YOL047C | LDS2 |  |  |  |  |  |  |  |  |
| YOL048C | RRT8 |  |  |  |  |  |  |  |  |
| YOL049W | GSH2 |  |  |  |  |  |  |  |  |
| YOL051W | GAL11 |  |  |  |  |  |  |  |  |
| YOL052C | SPE2 |  |  |  |  |  |  |  |  |
| YOL052C-A | DDR2 |  |  |  |  |  |  |  |  |
| YOL053W | AIM39 |  |  |  |  |  |  |  |  |
| YOL054W | PSH1 |  |  |  |  |  |  |  |  |
| YOL055C | THI20 |  |  |  |  |  |  |  |  |
| YOL056W | GPM3 |  |  |  |  |  |  |  |  |
| YOL057W |  |  |  |  |  |  |  |  |  |
| YOL058W | ARG1 |  |  |  |  |  |  |  |  |
| YOL059W | GPD2 |  |  |  |  |  |  |  |  |
| YOL060C | MAM3 |  |  |  |  |  |  |  |  |
| YOL061W | PRS5 |  |  |  |  |  |  |  |  |
| YOL062C | APM4 |  |  |  |  |  |  |  |  |
| YOL063C | CRT10 |  |  |  |  |  |  |  |  |
| YOL064C | MET22 |  |  |  |  |  |  |  |  |
| YOL065C | INP54 |  |  |  |  |  |  |  |  |
| YOL066C | RIB2 |  |  |  |  |  |  |  |  |
| YOL067C | RTG1 |  |  |  |  |  |  |  |  |
| YOL068C | HST1 |  |  |  |  |  |  |  |  |
| YOL069W | NUF2 |  |  |  |  |  |  |  |  |
| YOL070C | NBA1 |  |  |  |  |  |  |  |  |
| YOL071W | SDH5 |  |  |  |  |  |  |  |  |
| YOL072W | THP1 |  |  |  |  |  |  |  |  |
| YOL073C | DSC2 |  |  |  |  |  |  |  |  |
| YOL075C |  |  |  |  |  |  |  |  |  |
| YOL076W | MDM20 |  |  |  |  |  |  |  |  |
| YOL077C | BRX1 |  |  |  |  |  |  |  |  |
| YOL077W-A | ATP19 |  |  |  |  |  |  |  |  |
| YOL078W | AVO1 |  |  |  |  |  |  |  |  |
| YOL080C | REX4 |  |  |  |  |  |  |  |  |
| YOL081W | IRA2 |  |  |  |  |  |  |  |  |
| YOL082W | ATG19 |  |  |  |  |  |  |  |  |
| YOL083W | ATG34 |  |  |  |  |  |  |  |  |
| YOL084W | PHM7 |  |  |  |  |  |  |  |  |
| YOL086C | ADH1 |  |  |  |  |  |  |  |  |
| YOL086W-A | MHF1 |  |  |  |  |  |  |  |  |
| YOL087C | DUF1 |  |  |  |  |  |  |  |  |
| YOL088C | MPD2 |  |  |  |  |  |  |  |  |
| YOL090W | MSH2 |  |  |  |  |  |  |  |  |
| YOL091W | SPO21 |  |  |  |  |  |  |  |  |
| YOL092W | YPQ1 |  |  |  |  |  |  |  |  |
| YOL093W | TRM10 |  |  |  |  |  |  |  |  |
| YOL094C | RFC4 |  |  |  |  |  |  |  |  |
| YOL095C | HMI1 |  |  |  |  |  |  |  |  |
| YOL096C | COQ3 |  |  |  |  |  |  |  |  |
| YOL097C | WRS1 |  |  |  |  |  |  |  |  |
| YOL098C | SDD3 |  |  |  |  |  |  |  |  |
| YOL100W | PKH2 |  |  |  |  |  |  |  |  |
| YOL101C | IZH4 |  |  |  |  |  |  |  |  |
| YOL102C | TPT1 |  |  |  |  |  |  |  |  |
| YOL103W | ITR2 |  |  |  |  |  |  |  |  |
| YOL104C | NDJ1 |  |  |  |  |  |  |  |  |
| YOL105C | WSC3 |  |  |  |  |  |  |  |  |
| YOL108C | INO4 |  |  |  |  |  |  |  |  |
| YOL109W | ZEO1 |  |  |  |  |  |  |  |  |
| YOL110W | SHR5 |  |  |  |  |  |  |  |  |
| YOL111C | MDY2 |  |  |  |  |  |  |  |  |
| YOL112W | MSB4 |  |  |  |  |  |  |  |  |
| YOL113W | SKM1 |  |  |  |  |  |  |  |  |
| YOL114C | PTH4 |  |  |  |  |  |  |  |  |
| YOL115W | PAP2 |  |  |  |  |  |  |  |  |
| YOL116W | MSN1 |  |  |  |  |  |  |  |  |
| YOL117W | RRI2 |  |  |  |  |  |  |  |  |
| YOL119C | MCH4 |  |  |  |  |  |  |  |  |
| YOL120C | RPL18A |  |  |  |  |  |  |  |  |
| YOL121C | RPS19A |  |  |  |  |  |  |  |  |
| YOL122C | SMF1 |  |  |  |  |  |  |  |  |
| YOL123W | HRP1 |  |  |  |  |  |  |  |  |
| YOL124C | TRM11 |  |  |  |  |  |  |  |  |
| YOL125W | TRM13 |  |  |  |  |  |  |  |  |
| YOL126C | MDH2 |  |  |  |  |  |  |  |  |
| YOL127W | RPL25 |  |  |  |  |  |  |  |  |
| YOL128C | YGK3 |  |  |  |  |  |  |  |  |
| YOL129W | VPS68 |  |  |  |  |  |  |  |  |
| YOL130W | ALR1 |  |  |  |  |  |  |  |  |
| YOL132W | GAS4 |  |  |  |  |  |  |  |  |
| YOL133W | HRT1 |  |  |  |  |  |  |  |  |
| YOL135C | MED7 |  |  |  |  |  |  |  |  |
| YOL136C | PFK27 |  |  |  |  |  |  |  |  |
| YOL137W | BSC6 |  |  |  |  |  |  |  |  |
| YOL138C | RTC1 |  |  |  |  |  |  |  |  |
| YOL139C | CDC33 |  |  |  |  |  |  |  |  |
| YOL140W | ARG8 |  |  |  |  |  |  |  |  |
| YOL141W | PPM2 |  |  |  |  |  |  |  |  |
| YOL142W | RRP40 |  |  |  |  |  |  |  |  |
| YOL143C | RIB4 |  |  |  |  |  |  |  |  |
| YOL144W | NOP8 |  |  |  |  |  |  |  |  |
| YOL145C | CTR9 |  |  |  |  |  |  |  |  |
| YOL146W | PSF3 |  |  |  |  |  |  |  |  |
| YOL147C | PEX11 |  |  |  |  |  |  |  |  |
| YOL148C | SPT20 |  |  |  |  |  |  |  |  |
| YOL149W | DCP1 |  |  |  |  |  |  |  |  |
| YOR001W | RRP6 |  |  |  |  |  |  |  |  |
| YOR002W | ALG6 |  |  |  |  |  |  |  |  |
| YOR003W | YSP3 |  |  |  |  |  |  |  |  |
| YOR004W | UTP23 |  |  |  |  |  |  |  |  |
| YOR005C | DNL4 |  |  |  |  |  |  |  |  |
| YOR006C | TSR3 |  |  |  |  |  |  |  |  |
| YOR007C | SGT2 |  |  |  |  |  |  |  |  |
| YOR008C | SLG1 |  |  |  |  |  |  |  |  |
| YOR011W | AUS1 |  |  |  |  |  |  |  |  |
| YOR014W | RTS1 |  |  |  |  |  |  |  |  |
| YOR016C | ERP4 |  |  |  |  |  |  |  |  |
| YOR017W | PET127 |  |  |  |  |  |  |  |  |
| YOR018W | ROD1 |  |  |  |  |  |  |  |  |
| YOR019W |  |  |  |  |  |  |  |  |  |
| YOR020C | HSP10 |  |  |  |  |  |  |  |  |
| YOR020W-A | MCO10 |  |  |  |  |  |  |  |  |
| YOR021C | SFM1 |  |  |  |  |  |  |  |  |
| YOR022C | DDL1 |  |  |  |  |  |  |  |  |
| YOR023C | AHC1 |  |  |  |  |  |  |  |  |
| YOR025W | HST3 |  |  |  |  |  |  |  |  |
| YOR026W | BUB3 |  |  |  |  |  |  |  |  |
| YOR027W | STI1 |  |  |  |  |  |  |  |  |
| YOR028C | CIN5 |  |  |  |  |  |  |  |  |
| YOR030W | DFG16 |  |  |  |  |  |  |  |  |
| YOR032C | HMS1 |  |  |  |  |  |  |  |  |
| YOR033C | EXO1 |  |  |  |  |  |  |  |  |
| YOR034C | AKR2 |  |  |  |  |  |  |  |  |
| YOR035C | SHE4 |  |  |  |  |  |  |  |  |
| YOR036W | PEP12 |  |  |  |  |  |  |  |  |
| YOR037W | CYC2 |  |  |  |  |  |  |  |  |
| YOR038C | HIR2 |  |  |  |  |  |  |  |  |
| YOR039W | CKB2 |  |  |  |  |  |  |  |  |
| YOR040W | GLO4 |  |  |  |  |  |  |  |  |
| YOR042W | CUE5 |  |  |  |  |  |  |  |  |
| YOR043W | WHI2 |  |  |  |  |  |  |  |  |
| YOR044W | IRC23 |  |  |  |  |  |  |  |  |
| YOR045W | TOM6 |  |  |  |  |  |  |  |  |
| YOR046C | DBP5 |  |  |  |  |  |  |  |  |
| YOR047C | STD1 |  |  |  |  |  |  |  |  |
| YOR048C | RAT1 |  |  |  |  |  |  |  |  |
| YOR051C | ETT1 |  |  |  |  |  |  |  |  |
| YOR052C | TMC1 |  |  |  |  |  |  |  |  |
| YOR054C | VHS3 |  |  |  |  |  |  |  |  |
| YOR056C | NOB1 |  |  |  |  |  |  |  |  |
| YOR057W | SGT1 |  |  |  |  |  |  |  |  |
| YOR058C | ASE1 |  |  |  |  |  |  |  |  |
| YOR059C | LPL1 |  |  |  |  |  |  |  |  |
| YOR060C | SLD7 |  |  |  |  |  |  |  |  |
| YOR061W | CKA2 |  |  |  |  |  |  |  |  |
| YOR063W | RPL3 |  |  |  |  |  |  |  |  |
| YOR064C | YNG1 |  |  |  |  |  |  |  |  |
| YOR065W | CYT1 |  |  |  |  |  |  |  |  |
| YOR066W | MSA1 |  |  |  |  |  |  |  |  |
| YOR067C | ALG8 |  |  |  |  |  |  |  |  |
| YOR069W | VPS5 |  |  |  |  |  |  |  |  |
| YOR070C | GYP1 |  |  |  |  |  |  |  |  |
| YOR073W | SGO1 |  |  |  |  |  |  |  |  |
| YOR074C | CDC21 |  |  |  |  |  |  |  |  |
| YOR075W | UFE1 |  |  |  |  |  |  |  |  |
| YOR076C | SKI7 |  |  |  |  |  |  |  |  |
| YOR077W | RTS2 |  |  |  |  |  |  |  |  |
| YOR078W | BUD21 |  |  |  |  |  |  |  |  |
| YOR079C | ATX2 |  |  |  |  |  |  |  |  |
| YOR080W | DIA2 |  |  |  |  |  |  |  |  |
| YOR081C | TGL5 |  |  |  |  |  |  |  |  |
| YOR083W | WHI5 |  |  |  |  |  |  |  |  |
| YOR085W | OST3 |  |  |  |  |  |  |  |  |
| YOR086C | TCB1 |  |  |  |  |  |  |  |  |
| YOR089C | VPS21 |  |  |  |  |  |  |  |  |
| YOR090C | PTC5 |  |  |  |  |  |  |  |  |
| YOR091W | TMA46 |  |  |  |  |  |  |  |  |
| YOR092W | ECM3 |  |  |  |  |  |  |  |  |
| YOR094W | ARF3 |  |  |  |  |  |  |  |  |
| YOR095C | RKI1 |  |  |  |  |  |  |  |  |
| YOR096W | RPS7A |  |  |  |  |  |  |  |  |
| YOR098C | NUP1 |  |  |  |  |  |  |  |  |
| YOR099W | KTR1 |  |  |  |  |  |  |  |  |
| YOR100C | CRC1 |  |  |  |  |  |  |  |  |
| YOR101W | RAS1 |  |  |  |  |  |  |  |  |
| YOR103C | OST2 |  |  |  |  |  |  |  |  |
| YOR104W | PIN2 |  |  |  |  |  |  |  |  |
| YOR106W | VAM3 |  |  |  |  |  |  |  |  |
| YOR107W | RGS2 |  |  |  |  |  |  |  |  |
| YOR108W | LEU9 |  |  |  |  |  |  |  |  |
| YOR109W | INP53 |  |  |  |  |  |  |  |  |
| YOR110W | TFC7 |  |  |  |  |  |  |  |  |
| YOR112W | CEX1 |  |  |  |  |  |  |  |  |
| YOR113W | AZF1 |  |  |  |  |  |  |  |  |
| YOR115C | TRS33 |  |  |  |  |  |  |  |  |
| YOR116C | RPO31 |  |  |  |  |  |  |  |  |
| YOR117W | RPT5 |  |  |  |  |  |  |  |  |
| YOR118W | RTC5 |  |  |  |  |  |  |  |  |
| YOR119C | RIO1 |  |  |  |  |  |  |  |  |
| YOR120W | GCY1 |  |  |  |  |  |  |  |  |
| YOR122C | PFY1 |  |  |  |  |  |  |  |  |
| YOR123C | LEO1 |  |  |  |  |  |  |  |  |
| YOR124C | UBP2 |  |  |  |  |  |  |  |  |
| YOR125C | CAT5 |  |  |  |  |  |  |  |  |
| YOR126C | IAH1 |  |  |  |  |  |  |  |  |
| YOR127W | RGA1 |  |  |  |  |  |  |  |  |
| YOR128C | ADE2 |  |  |  |  |  |  |  |  |
| YOR129C | AFI1 |  |  |  |  |  |  |  |  |
| YOR130C | ORT1 |  |  |  |  |  |  |  |  |
| YOR132W | VPS17 |  |  |  |  |  |  |  |  |
| YOR133W | EFT1 |  |  |  |  |  |  |  |  |
| YOR134W | BAG7 |  |  |  |  |  |  |  |  |
| YOR136W | IDH2 |  |  |  |  |  |  |  |  |
| YOR140W | SFL1 |  |  |  |  |  |  |  |  |
| YOR141C | ARP8 |  |  |  |  |  |  |  |  |
| YOR142W | LSC1 |  |  |  |  |  |  |  |  |
| YOR143C | THI80 |  |  |  |  |  |  |  |  |
| YOR144C | ELG1 |  |  |  |  |  |  |  |  |
| YOR145C | PNO1 |  |  |  |  |  |  |  |  |
| YOR147W | MDM32 |  |  |  |  |  |  |  |  |
| YOR148C | SPP2 |  |  |  |  |  |  |  |  |
| YOR149C | SMP3 |  |  |  |  |  |  |  |  |
| YOR150W | MRPL23 |  |  |  |  |  |  |  |  |
| YOR151C | RPB2 |  |  |  |  |  |  |  |  |
| YOR152C | ATG40 |  |  |  |  |  |  |  |  |
| YOR153W | PDR5 |  |  |  |  |  |  |  |  |
| YOR154W | SLP1 |  |  |  |  |  |  |  |  |
| YOR155C | ISN1 |  |  |  |  |  |  |  |  |
| YOR156C | NFI1 |  |  |  |  |  |  |  |  |
| YOR157C | PUP1 |  |  |  |  |  |  |  |  |
| YOR158W | PET123 |  |  |  |  |  |  |  |  |
| YOR159C | SME1 |  |  |  |  |  |  |  |  |
| YOR160W | MTR10 |  |  |  |  |  |  |  |  |
| YOR161C | PNS1 |  |  |  |  |  |  |  |  |
| YOR163W | DDP1 |  |  |  |  |  |  |  |  |
| YOR164C | GET4 |  |  |  |  |  |  |  |  |
| YOR165W | SEY1 |  |  |  |  |  |  |  |  |
| YOR166C | SWT1 |  |  |  |  |  |  |  |  |
| YOR167C | RPS28A |  |  |  |  |  |  |  |  |
| YOR168W | GLN4 |  |  |  |  |  |  |  |  |
| YOR171C | LCB4 |  |  |  |  |  |  |  |  |
| YOR172W | YRM1 |  |  |  |  |  |  |  |  |
| YOR173W | DCS2 |  |  |  |  |  |  |  |  |
| YOR174W | MED4 |  |  |  |  |  |  |  |  |
| YOR175C | ALE1 |  |  |  |  |  |  |  |  |
| YOR176W | HEM15 |  |  |  |  |  |  |  |  |
| YOR177C | MPC54 |  |  |  |  |  |  |  |  |
| YOR178C | GAC1 |  |  |  |  |  |  |  |  |
| YOR179C | SYC1 |  |  |  |  |  |  |  |  |
| YOR180C | DCI1 |  |  |  |  |  |  |  |  |
| YOR181W | LAS17 |  |  |  |  |  |  |  |  |
| YOR182C | RPS30B |  |  |  |  |  |  |  |  |
| YOR184W | SER1 |  |  |  |  |  |  |  |  |
| YOR185C | GSP2 |  |  |  |  |  |  |  |  |
| YOR187W | TUF1 |  |  |  |  |  |  |  |  |
| YOR188W | MSB1 |  |  |  |  |  |  |  |  |
| YOR189W | IES4 |  |  |  |  |  |  |  |  |
| YOR190W | SPR1 |  |  |  |  |  |  |  |  |
| YOR191W | ULS1 |  |  |  |  |  |  |  |  |
| YOR192C | THI72 |  |  |  |  |  |  |  |  |
| YOR193W | PEX27 |  |  |  |  |  |  |  |  |
| YOR194C | TOA1 |  |  |  |  |  |  |  |  |
| YOR195W | SLK19 |  |  |  |  |  |  |  |  |
| YOR196C | LIP5 |  |  |  |  |  |  |  |  |
| YOR197W | MCA1 |  |  |  |  |  |  |  |  |
| YOR198C | BFR1 |  |  |  |  |  |  |  |  |
| YOR201C | MRM1 |  |  |  |  |  |  |  |  |
| YOR202W | HIS3 |  |  |  |  |  |  |  |  |
| YOR204W | DED1 |  |  |  |  |  |  |  |  |
| YOR205C | GEP3 |  |  |  |  |  |  |  |  |
| YOR206W | NOC2 |  |  |  |  |  |  |  |  |
| YOR207C | RET1 |  |  |  |  |  |  |  |  |
| YOR208W | PTP2 |  |  |  |  |  |  |  |  |
| YOR209C | NPT1 |  |  |  |  |  |  |  |  |
| YOR210W | RPB10 |  |  |  |  |  |  |  |  |
| YOR211C | MGM1 |  |  |  |  |  |  |  |  |
| YOR212W | STE4 |  |  |  |  |  |  |  |  |
| YOR213C | SAS5 |  |  |  |  |  |  |  |  |
| YOR215C | AIM41 |  |  |  |  |  |  |  |  |
| YOR216C | RUD3 |  |  |  |  |  |  |  |  |
| YOR217W | RFC1 |  |  |  |  |  |  |  |  |
| YOR219C | STE13 |  |  |  |  |  |  |  |  |
| YOR220W | RCN2 |  |  |  |  |  |  |  |  |
| YOR221C | MCT1 |  |  |  |  |  |  |  |  |
| YOR222W | ODC2 |  |  |  |  |  |  |  |  |
| YOR223W | DSC3 |  |  |  |  |  |  |  |  |
| YOR224C | RPB8 |  |  |  |  |  |  |  |  |
| YOR226C | ISU2 |  |  |  |  |  |  |  |  |
| YOR227W | HER1 |  |  |  |  |  |  |  |  |
| YOR228C | MCP1 |  |  |  |  |  |  |  |  |
| YOR229W | WTM2 |  |  |  |  |  |  |  |  |
| YOR230W | WTM1 |  |  |  |  |  |  |  |  |
| YOR231W | MKK1 |  |  |  |  |  |  |  |  |
| YOR232W | MGE1 |  |  |  |  |  |  |  |  |
| YOR233W | KIN4 |  |  |  |  |  |  |  |  |
| YOR234C | RPL33B |  |  |  |  |  |  |  |  |
| YOR236W | DFR1 |  |  |  |  |  |  |  |  |
| YOR237W | HES1 |  |  |  |  |  |  |  |  |
| YOR239W | ABP140 |  |  |  |  |  |  |  |  |
| YOR241W | MET7 |  |  |  |  |  |  |  |  |
| YOR242C | SSP2 |  |  |  |  |  |  |  |  |
| YOR243C | PUS7 |  |  |  |  |  |  |  |  |
| YOR244W | ESA1 |  |  |  |  |  |  |  |  |
| YOR245C | DGA1 |  |  |  |  |  |  |  |  |
| YOR246C | ENV9 |  |  |  |  |  |  |  |  |
| YOR247W | SRL1 |  |  |  |  |  |  |  |  |
| YOR249C | APC5 |  |  |  |  |  |  |  |  |
| YOR250C | CLP1 |  |  |  |  |  |  |  |  |
| YOR251C | TUM1 |  |  |  |  |  |  |  |  |
| YOR252W | TMA16 |  |  |  |  |  |  |  |  |
| YOR253W | NAT5 |  |  |  |  |  |  |  |  |
| YOR254C | SEC63 |  |  |  |  |  |  |  |  |
| YOR255W | OSW1 |  |  |  |  |  |  |  |  |
| YOR256C | TRE2 |  |  |  |  |  |  |  |  |
| YOR257W | CDC31 |  |  |  |  |  |  |  |  |
| YOR258W | HNT3 |  |  |  |  |  |  |  |  |
| YOR259C | RPT4 |  |  |  |  |  |  |  |  |
| YOR260W | GCD1 |  |  |  |  |  |  |  |  |
| YOR261C | RPN8 |  |  |  |  |  |  |  |  |
| YOR262W | GPN2 |  |  |  |  |  |  |  |  |
| YOR265W | RBL2 |  |  |  |  |  |  |  |  |
| YOR266W | PNT1 |  |  |  |  |  |  |  |  |
| YOR267C | HRK1 |  |  |  |  |  |  |  |  |
| YOR269W | PAC1 |  |  |  |  |  |  |  |  |
| YOR270C | VPH1 |  |  |  |  |  |  |  |  |
| YOR272W | YTM1 |  |  |  |  |  |  |  |  |
| YOR273C | TPO4 |  |  |  |  |  |  |  |  |
| YOR274W | MOD5 |  |  |  |  |  |  |  |  |
| YOR275C | RIM20 |  |  |  |  |  |  |  |  |
| YOR276W | CAF20 |  |  |  |  |  |  |  |  |
| YOR278W | HEM4 |  |  |  |  |  |  |  |  |
| YOR279C | RFM1 |  |  |  |  |  |  |  |  |
| YOR280C | FSH3 |  |  |  |  |  |  |  |  |
| YOR281C | PLP2 |  |  |  |  |  |  |  |  |
| YOR283W |  |  |  |  |  |  |  |  |  |
| YOR285W | RDL1 |  |  |  |  |  |  |  |  |
| YOR286W | RDL2 |  |  |  |  |  |  |  |  |
| YOR287C | RRP36 |  |  |  |  |  |  |  |  |
| YOR288C | MPD1 |  |  |  |  |  |  |  |  |
| YOR290C | SNF2 |  |  |  |  |  |  |  |  |
| YOR291W | YPK9 |  |  |  |  |  |  |  |  |
| YOR293W | RPS10A |  |  |  |  |  |  |  |  |
| YOR294W | RRS1 |  |  |  |  |  |  |  |  |
| YOR295W | UAF30 |  |  |  |  |  |  |  |  |
| YOR297C | TIM18 |  |  |  |  |  |  |  |  |
| YOR298C-A | MBF1 |  |  |  |  |  |  |  |  |
| YOR298W | MUM3 |  |  |  |  |  |  |  |  |
| YOR299W | BUD7 |  |  |  |  |  |  |  |  |
| YOR301W | RAX1 |  |  |  |  |  |  |  |  |
| YOR302W |  |  |  |  |  |  |  |  |  |
| YOR303W | CPA1 |  |  |  |  |  |  |  |  |
| YOR304C-A | BIL1 |  |  |  |  |  |  |  |  |
| YOR304W | ISW2 |  |  |  |  |  |  |  |  |
| YOR305W | RRG7 |  |  |  |  |  |  |  |  |
| YOR307C | SLY41 |  |  |  |  |  |  |  |  |
| YOR308C | SNU66 |  |  |  |  |  |  |  |  |
| YOR310C | NOP58 |  |  |  |  |  |  |  |  |
| YOR311C | DGK1 |  |  |  |  |  |  |  |  |
| YOR312C | RPL20B |  |  |  |  |  |  |  |  |
| YOR313C | SPS4 |  |  |  |  |  |  |  |  |
| YOR315W | SFG1 |  |  |  |  |  |  |  |  |
| YOR316C | COT1 |  |  |  |  |  |  |  |  |
| YOR317W | FAA1 |  |  |  |  |  |  |  |  |
| YOR319W | HSH49 |  |  |  |  |  |  |  |  |
| YOR321W | PMT3 |  |  |  |  |  |  |  |  |
| YOR322C | LDB19 |  |  |  |  |  |  |  |  |
| YOR323C | PRO2 |  |  |  |  |  |  |  |  |
| YOR324C | FRT1 |  |  |  |  |  |  |  |  |
| YOR326W | MYO2 |  |  |  |  |  |  |  |  |
| YOR327C | SNC2 |  |  |  |  |  |  |  |  |
| YOR329C | SCD5 |  |  |  |  |  |  |  |  |
| YOR330C | MIP1 |  |  |  |  |  |  |  |  |
| YOR332W | VMA4 |  |  |  |  |  |  |  |  |
| YOR334W | MRS2 |  |  |  |  |  |  |  |  |
| YOR335C | ALA1 |  |  |  |  |  |  |  |  |
| YOR336W | KRE5 |  |  |  |  |  |  |  |  |
| YOR337W | TEA1 |  |  |  |  |  |  |  |  |
| YOR340C | RPA43 |  |  |  |  |  |  |  |  |
| YOR341W | RPA190 |  |  |  |  |  |  |  |  |
| YOR342C |  |  |  |  |  |  |  |  |  |
| YOR344C | TYE7 |  |  |  |  |  |  |  |  |
| YOR346W | REV1 |  |  |  |  |  |  |  |  |
| YOR347C | PYK2 |  |  |  |  |  |  |  |  |
| YOR348C | PUT4 |  |  |  |  |  |  |  |  |
| YOR349W | CIN1 |  |  |  |  |  |  |  |  |
| YOR351C | MEK1 |  |  |  |  |  |  |  |  |
| YOR352W | TFB6 |  |  |  |  |  |  |  |  |
| YOR353C | SOG2 |  |  |  |  |  |  |  |  |
| YOR354C | MSC6 |  |  |  |  |  |  |  |  |
| YOR355W | GDS1 |  |  |  |  |  |  |  |  |
| YOR356W | CIR2 |  |  |  |  |  |  |  |  |
| YOR357C | SNX3 |  |  |  |  |  |  |  |  |
| YOR358W | HAP5 |  |  |  |  |  |  |  |  |
| YOR359W | VTS1 |  |  |  |  |  |  |  |  |
| YOR360C | PDE2 |  |  |  |  |  |  |  |  |
| YOR361C | PRT1 |  |  |  |  |  |  |  |  |
| YOR362C | PRE10 |  |  |  |  |  |  |  |  |
| YOR363C | PIP2 |  |  |  |  |  |  |  |  |
| YOR367W | SCP1 |  |  |  |  |  |  |  |  |
| YOR368W | RAD17 |  |  |  |  |  |  |  |  |
| YOR369C | RPS12 |  |  |  |  |  |  |  |  |
| YOR370C | MRS6 |  |  |  |  |  |  |  |  |
| YOR371C | GPB1 |  |  |  |  |  |  |  |  |
| YOR372C | NDD1 |  |  |  |  |  |  |  |  |
| YOR373W | NUD1 |  |  |  |  |  |  |  |  |
| YOR374W | ALD4 |  |  |  |  |  |  |  |  |
| YOR381W | FRE3 |  |  |  |  |  |  |  |  |
| YPL001W | HAT1 |  |  |  |  |  |  |  |  |
| YPL002C | SNF8 |  |  |  |  |  |  |  |  |
| YPL003W | ULA1 |  |  |  |  |  |  |  |  |
| YPL004C | LSP1 |  |  |  |  |  |  |  |  |
| YPL005W | AEP3 |  |  |  |  |  |  |  |  |
| YPL006W | NCR1 |  |  |  |  |  |  |  |  |
| YPL007C | TFC8 |  |  |  |  |  |  |  |  |
| YPL008W | CHL1 |  |  |  |  |  |  |  |  |
| YPL009C | RQC2 |  |  |  |  |  |  |  |  |
| YPL010W | RET3 |  |  |  |  |  |  |  |  |
| YPL011C | TAF3 |  |  |  |  |  |  |  |  |
| YPL012W | RRP12 |  |  |  |  |  |  |  |  |
| YPL013C | MRPS16 |  |  |  |  |  |  |  |  |
| YPL014W | CIP1 |  |  |  |  |  |  |  |  |
| YPL015C | HST2 |  |  |  |  |  |  |  |  |
| YPL016W | SWI1 |  |  |  |  |  |  |  |  |
| YPL017C | IRC15 |  |  |  |  |  |  |  |  |
| YPL018W | CTF19 |  |  |  |  |  |  |  |  |
| YPL019C | VTC3 |  |  |  |  |  |  |  |  |
| YPL020C | ULP1 |  |  |  |  |  |  |  |  |
| YPL022W | RAD1 |  |  |  |  |  |  |  |  |
| YPL023C | MET12 |  |  |  |  |  |  |  |  |
| YPL024W | RMI1 |  |  |  |  |  |  |  |  |
| YPL026C | SKS1 |  |  |  |  |  |  |  |  |
| YPL028W | ERG10 |  |  |  |  |  |  |  |  |
| YPL029W | SUV3 |  |  |  |  |  |  |  |  |
| YPL030W | TRM44 |  |  |  |  |  |  |  |  |
| YPL031C | PHO85 |  |  |  |  |  |  |  |  |
| YPL032C | SVL3 |  |  |  |  |  |  |  |  |
| YPL033C | SRL4 |  |  |  |  |  |  |  |  |
| YPL036W | PMA2 |  |  |  |  |  |  |  |  |
| YPL037C | EGD1 |  |  |  |  |  |  |  |  |
| YPL038W | MET31 |  |  |  |  |  |  |  |  |
| YPL040C | ISM1 |  |  |  |  |  |  |  |  |
| YPL042C | SSN3 |  |  |  |  |  |  |  |  |
| YPL043W | NOP4 |  |  |  |  |  |  |  |  |
| YPL045W | VPS16 |  |  |  |  |  |  |  |  |
| YPL046C | ELC1 |  |  |  |  |  |  |  |  |
| YPL047W | SGF11 |  |  |  |  |  |  |  |  |
| YPL048W | CAM1 |  |  |  |  |  |  |  |  |
| YPL049C | DIG1 |  |  |  |  |  |  |  |  |
| YPL050C | MNN9 |  |  |  |  |  |  |  |  |
| YPL051W | ARL3 |  |  |  |  |  |  |  |  |
| YPL052W | OAZ1 |  |  |  |  |  |  |  |  |
| YPL053C | KTR6 |  |  |  |  |  |  |  |  |
| YPL054W | LEE1 |  |  |  |  |  |  |  |  |
| YPL055C | LGE1 |  |  |  |  |  |  |  |  |
| YPL058C | PDR12 |  |  |  |  |  |  |  |  |
| YPL059W | GRX5 |  |  |  |  |  |  |  |  |
| YPL060W | MFM1 |  |  |  |  |  |  |  |  |
| YPL063W | TIM50 |  |  |  |  |  |  |  |  |
| YPL064C | CWC27 |  |  |  |  |  |  |  |  |
| YPL065W | VPS28 |  |  |  |  |  |  |  |  |
| YPL066W | RGL1 |  |  |  |  |  |  |  |  |
| YPL069C | BTS1 |  |  |  |  |  |  |  |  |
| YPL070W | MUK1 |  |  |  |  |  |  |  |  |
| YPL072W | UBP16 |  |  |  |  |  |  |  |  |
| YPL074W | YTA6 |  |  |  |  |  |  |  |  |
| YPL075W | GCR1 |  |  |  |  |  |  |  |  |
| YPL076W | GPI2 |  |  |  |  |  |  |  |  |
| YPL078C | ATP4 |  |  |  |  |  |  |  |  |
| YPL079W | RPL21B |  |  |  |  |  |  |  |  |
| YPL081W | RPS9A |  |  |  |  |  |  |  |  |
| YPL082C | MOT1 |  |  |  |  |  |  |  |  |
| YPL083C | SEN54 |  |  |  |  |  |  |  |  |
| YPL084W | BRO1 |  |  |  |  |  |  |  |  |
| YPL085W | SEC16 |  |  |  |  |  |  |  |  |
| YPL086C | ELP3 |  |  |  |  |  |  |  |  |
| YPL087W | YDC1 |  |  |  |  |  |  |  |  |
| YPL089C | RLM1 |  |  |  |  |  |  |  |  |
| YPL090C | RPS6A |  |  |  |  |  |  |  |  |
| YPL091W | GLR1 |  |  |  |  |  |  |  |  |
| YPL093W | NOG1 |  |  |  |  |  |  |  |  |
| YPL094C | SEC62 |  |  |  |  |  |  |  |  |
| YPL095C | EEB1 |  |  |  |  |  |  |  |  |
| YPL096C-A | ERI1 |  |  |  |  |  |  |  |  |
| YPL096W | PNG1 |  |  |  |  |  |  |  |  |
| YPL097W | MSY1 |  |  |  |  |  |  |  |  |
| YPL098C | MGR2 |  |  |  |  |  |  |  |  |
| YPL099C | INA17 |  |  |  |  |  |  |  |  |
| YPL100W | ATG21 |  |  |  |  |  |  |  |  |
| YPL101W | ELP4 |  |  |  |  |  |  |  |  |
| YPL103C | FMP30 |  |  |  |  |  |  |  |  |
| YPL104W | MSD1 |  |  |  |  |  |  |  |  |
| YPL105C | SYH1 |  |  |  |  |  |  |  |  |
| YPL106C | SSE1 |  |  |  |  |  |  |  |  |
| YPL109C | CQD1 |  |  |  |  |  |  |  |  |
| YPL110C | GDE1 |  |  |  |  |  |  |  |  |
| YPL111W | CAR1 |  |  |  |  |  |  |  |  |
| YPL112C | PEX25 |  |  |  |  |  |  |  |  |
| YPL113C |  |  |  |  |  |  |  |  |  |
| YPL115C | BEM3 |  |  |  |  |  |  |  |  |
| YPL116W | HOS3 |  |  |  |  |  |  |  |  |
| YPL117C | IDI1 |  |  |  |  |  |  |  |  |
| YPL118W | MRP51 |  |  |  |  |  |  |  |  |
| YPL119C | DBP1 |  |  |  |  |  |  |  |  |
| YPL120W | VPS30 |  |  |  |  |  |  |  |  |
| YPL121C | MEI5 |  |  |  |  |  |  |  |  |
| YPL122C | TFB2 |  |  |  |  |  |  |  |  |
| YPL123C | RNY1 |  |  |  |  |  |  |  |  |
| YPL124W | SPC29 |  |  |  |  |  |  |  |  |
| YPL125W | KAP120 |  |  |  |  |  |  |  |  |
| YPL126W | NAN1 |  |  |  |  |  |  |  |  |
| YPL127C | HHO1 |  |  |  |  |  |  |  |  |
| YPL128C | TBF1 |  |  |  |  |  |  |  |  |
| YPL129W | TAF14 |  |  |  |  |  |  |  |  |
| YPL131W | RPL5 |  |  |  |  |  |  |  |  |
| YPL132W | COX11 |  |  |  |  |  |  |  |  |
| YPL133C | RDS2 |  |  |  |  |  |  |  |  |
| YPL134C | ODC1 |  |  |  |  |  |  |  |  |
| YPL135W | ISU1 |  |  |  |  |  |  |  |  |
| YPL137C | GIP3 |  |  |  |  |  |  |  |  |
| YPL138C | SPP1 |  |  |  |  |  |  |  |  |
| YPL139C | UME1 |  |  |  |  |  |  |  |  |
| YPL140C | MKK2 |  |  |  |  |  |  |  |  |
| YPL141C | FRK1 |  |  |  |  |  |  |  |  |
| YPL143W | RPL33A |  |  |  |  |  |  |  |  |
| YPL144W | POC4 |  |  |  |  |  |  |  |  |
| YPL145C | KES1 |  |  |  |  |  |  |  |  |
| YPL146C | NOP53 |  |  |  |  |  |  |  |  |
| YPL147W | PXA1 |  |  |  |  |  |  |  |  |
| YPL148C | PPT2 |  |  |  |  |  |  |  |  |
| YPL149W | ATG5 |  |  |  |  |  |  |  |  |
| YPL151C | PRP46 |  |  |  |  |  |  |  |  |
| YPL152W | RRD2 |  |  |  |  |  |  |  |  |
| YPL153C | RAD53 |  |  |  |  |  |  |  |  |
| YPL154C | PEP4 |  |  |  |  |  |  |  |  |
| YPL155C | KIP2 |  |  |  |  |  |  |  |  |
| YPL156C | PRM4 |  |  |  |  |  |  |  |  |
| YPL157W | TGS1 |  |  |  |  |  |  |  |  |
| YPL158C | AIM44 |  |  |  |  |  |  |  |  |
| YPL159C | PET20 |  |  |  |  |  |  |  |  |
| YPL160W | CDC60 |  |  |  |  |  |  |  |  |
| YPL161C | BEM4 |  |  |  |  |  |  |  |  |
| YPL163C | SVS1 |  |  |  |  |  |  |  |  |
| YPL164C | MLH3 |  |  |  |  |  |  |  |  |
| YPL165C | SET6 |  |  |  |  |  |  |  |  |
| YPL166W | ATG29 |  |  |  |  |  |  |  |  |
| YPL167C | REV3 |  |  |  |  |  |  |  |  |
| YPL169C | MEX67 |  |  |  |  |  |  |  |  |
| YPL170W | DAP1 |  |  |  |  |  |  |  |  |
| YPL171C | OYE3 |  |  |  |  |  |  |  |  |
| YPL172C | COX10 |  |  |  |  |  |  |  |  |
| YPL173W | MRPL40 |  |  |  |  |  |  |  |  |
| YPL174C | NIP100 |  |  |  |  |  |  |  |  |
| YPL175W | SPT14 |  |  |  |  |  |  |  |  |
| YPL176C | TRE1 |  |  |  |  |  |  |  |  |
| YPL177C | CUP9 |  |  |  |  |  |  |  |  |
| YPL178W | CBC2 |  |  |  |  |  |  |  |  |
| YPL179W | PPQ1 |  |  |  |  |  |  |  |  |
| YPL180W | TCO89 |  |  |  |  |  |  |  |  |
| YPL181W | CTI6 |  |  |  |  |  |  |  |  |
| YPL183C | RTT10 |  |  |  |  |  |  |  |  |
| YPL183W-A | RTC6 |  |  |  |  |  |  |  |  |
| YPL184C | MRN1 |  |  |  |  |  |  |  |  |
| YPL187W | MF(ALPHA)1 |  |  |  |  |  |  |  |  |
| YPL188W | POS5 |  |  |  |  |  |  |  |  |
| YPL189C-A | COA2 |  |  |  |  |  |  |  |  |
| YPL189W | GUP2 |  |  |  |  |  |  |  |  |
| YPL190C | NAB3 |  |  |  |  |  |  |  |  |
| YPL191C | MIY2 |  |  |  |  |  |  |  |  |
| YPL192C | PRM3 |  |  |  |  |  |  |  |  |
| YPL193W | RSA1 |  |  |  |  |  |  |  |  |
| YPL194W | DDC1 |  |  |  |  |  |  |  |  |
| YPL195W | APL5 |  |  |  |  |  |  |  |  |
| YPL196W | OXR1 |  |  |  |  |  |  |  |  |
| YPL198W | RPL7B |  |  |  |  |  |  |  |  |
| YPL199C |  |  |  |  |  |  |  |  |  |
| YPL200W | CSM4 |  |  |  |  |  |  |  |  |
| YPL202C | AFT2 |  |  |  |  |  |  |  |  |
| YPL203W | TPK2 |  |  |  |  |  |  |  |  |
| YPL204W | HRR25 |  |  |  |  |  |  |  |  |
| YPL206C | PGC1 |  |  |  |  |  |  |  |  |
| YPL207W | TYW1 |  |  |  |  |  |  |  |  |
| YPL208W | RKM1 |  |  |  |  |  |  |  |  |
| YPL209C | IPL1 |  |  |  |  |  |  |  |  |
| YPL210C | SRP72 |  |  |  |  |  |  |  |  |
| YPL211W | NIP7 |  |  |  |  |  |  |  |  |
| YPL212C | PUS1 |  |  |  |  |  |  |  |  |
| YPL213W | LEA1 |  |  |  |  |  |  |  |  |
| YPL214C | THI6 |  |  |  |  |  |  |  |  |
| YPL215W | CBP3 |  |  |  |  |  |  |  |  |
| YPL217C | BMS1 |  |  |  |  |  |  |  |  |
| YPL218W | SAR1 |  |  |  |  |  |  |  |  |
| YPL219W | PCL8 |  |  |  |  |  |  |  |  |
| YPL220W | RPL1A |  |  |  |  |  |  |  |  |
| YPL221W | FLC1 |  |  |  |  |  |  |  |  |
| YPL222W | FMP40 |  |  |  |  |  |  |  |  |
| YPL224C | MMT2 |  |  |  |  |  |  |  |  |
| YPL225W | CHP1 |  |  |  |  |  |  |  |  |
| YPL226W | NEW1 |  |  |  |  |  |  |  |  |
| YPL227C | ALG5 |  |  |  |  |  |  |  |  |
| YPL228W | CET1 |  |  |  |  |  |  |  |  |
| YPL230W | USV1 |  |  |  |  |  |  |  |  |
| YPL231W | FAS2 |  |  |  |  |  |  |  |  |
| YPL232W | SSO1 |  |  |  |  |  |  |  |  |
| YPL233W | NSL1 |  |  |  |  |  |  |  |  |
| YPL234C | VMA11 |  |  |  |  |  |  |  |  |
| YPL235W | RVB2 |  |  |  |  |  |  |  |  |
| YPL236C | ENV7 |  |  |  |  |  |  |  |  |
| YPL237W | SUI3 |  |  |  |  |  |  |  |  |
| YPL239W | YAR1 |  |  |  |  |  |  |  |  |
| YPL240C | HSP82 |  |  |  |  |  |  |  |  |
| YPL241C | CIN2 |  |  |  |  |  |  |  |  |
| YPL242C | IQG1 |  |  |  |  |  |  |  |  |
| YPL243W | SRP68 |  |  |  |  |  |  |  |  |
| YPL244C | HUT1 |  |  |  |  |  |  |  |  |
| YPL246C | RBD2 |  |  |  |  |  |  |  |  |
| YPL248C | GAL4 |  |  |  |  |  |  |  |  |
| YPL249C | GYP5 |  |  |  |  |  |  |  |  |
| YPL249C-A | RPL36B |  |  |  |  |  |  |  |  |
| YPL250C | ATG41 |  |  |  |  |  |  |  |  |
| YPL252C | YAH1 |  |  |  |  |  |  |  |  |
| YPL253C | VIK1 |  |  |  |  |  |  |  |  |
| YPL254W | HFI1 |  |  |  |  |  |  |  |  |
| YPL255W | BBP1 |  |  |  |  |  |  |  |  |
| YPL256C | CLN2 |  |  |  |  |  |  |  |  |
| YPL258C | THI21 |  |  |  |  |  |  |  |  |
| YPL259C | APM1 |  |  |  |  |  |  |  |  |
| YPL260W | CUB1 |  |  |  |  |  |  |  |  |
| YPL262W | FUM1 |  |  |  |  |  |  |  |  |
| YPL263C | KEL3 |  |  |  |  |  |  |  |  |
| YPL265W | DIP5 |  |  |  |  |  |  |  |  |
| YPL266W | DIM1 |  |  |  |  |  |  |  |  |
| YPL267W | ACM1 |  |  |  |  |  |  |  |  |
| YPL268W | PLC1 |  |  |  |  |  |  |  |  |
| YPL269W | KAR9 |  |  |  |  |  |  |  |  |
| YPL270W | MDL2 |  |  |  |  |  |  |  |  |
| YPL271W | ATP15 |  |  |  |  |  |  |  |  |
| YPL273W | SAM4 |  |  |  |  |  |  |  |  |
| YPL274W | SAM3 |  |  |  |  |  |  |  |  |
| YPR001W | CIT3 |  |  |  |  |  |  |  |  |
| YPR002W | PDH1 |  |  |  |  |  |  |  |  |
| YPR004C | AIM45 |  |  |  |  |  |  |  |  |
| YPR005C | HAL1 |  |  |  |  |  |  |  |  |
| YPR006C | ICL2 |  |  |  |  |  |  |  |  |
| YPR007C | REC8 |  |  |  |  |  |  |  |  |
| YPR008W | HAA1 |  |  |  |  |  |  |  |  |
| YPR009W | SUT2 |  |  |  |  |  |  |  |  |
| YPR010C | RPA135 |  |  |  |  |  |  |  |  |
| YPR010C-A | MRA1 |  |  |  |  |  |  |  |  |
| YPR011C | MRX21 |  |  |  |  |  |  |  |  |
| YPR013C | CMR3 |  |  |  |  |  |  |  |  |
| YPR015C |  |  |  |  |  |  |  |  |  |
| YPR016C | TIF6 |  |  |  |  |  |  |  |  |
| YPR018W | RLF2 |  |  |  |  |  |  |  |  |
| YPR019W | MCM4 |  |  |  |  |  |  |  |  |
| YPR020W | ATP20 |  |  |  |  |  |  |  |  |
| YPR021C | AGC1 |  |  |  |  |  |  |  |  |
| YPR023C | EAF3 |  |  |  |  |  |  |  |  |
| YPR024W | YME1 |  |  |  |  |  |  |  |  |
| YPR025C | CCL1 |  |  |  |  |  |  |  |  |
| YPR026W | ATH1 |  |  |  |  |  |  |  |  |
| YPR028W | YOP1 |  |  |  |  |  |  |  |  |
| YPR029C | APL4 |  |  |  |  |  |  |  |  |
| YPR030W | CSR2 |  |  |  |  |  |  |  |  |
| YPR031W | NTO1 |  |  |  |  |  |  |  |  |
| YPR032W | SRO7 |  |  |  |  |  |  |  |  |
| YPR033C | HTS1 |  |  |  |  |  |  |  |  |
| YPR034W | ARP7 |  |  |  |  |  |  |  |  |
| YPR035W | GLN1 |  |  |  |  |  |  |  |  |
| YPR036W | VMA13 |  |  |  |  |  |  |  |  |
| YPR036W-A | SPO24 |  |  |  |  |  |  |  |  |
| YPR037C | ERV2 |  |  |  |  |  |  |  |  |
| YPR040W | TIP41 |  |  |  |  |  |  |  |  |
| YPR041W | TIF5 |  |  |  |  |  |  |  |  |
| YPR042C | PUF2 |  |  |  |  |  |  |  |  |
| YPR043W | RPL43A |  |  |  |  |  |  |  |  |
| YPR045C | THP3 |  |  |  |  |  |  |  |  |
| YPR046W | MCM16 |  |  |  |  |  |  |  |  |
| YPR047W | MSF1 |  |  |  |  |  |  |  |  |
| YPR048W | TAH18 |  |  |  |  |  |  |  |  |
| YPR049C | ATG11 |  |  |  |  |  |  |  |  |
| YPR051W | MAK3 |  |  |  |  |  |  |  |  |
| YPR052C | NHP6A |  |  |  |  |  |  |  |  |
| YPR054W | SMK1 |  |  |  |  |  |  |  |  |
| YPR055W | SEC8 |  |  |  |  |  |  |  |  |
| YPR056W | TFB4 |  |  |  |  |  |  |  |  |
| YPR057W | BRR1 |  |  |  |  |  |  |  |  |
| YPR060C | ARO7 |  |  |  |  |  |  |  |  |
| YPR061C | JID1 |  |  |  |  |  |  |  |  |
| YPR062W | FCY1 |  |  |  |  |  |  |  |  |
| YPR065W | ROX1 |  |  |  |  |  |  |  |  |
| YPR066W | UBA3 |  |  |  |  |  |  |  |  |
| YPR067W | ISA2 |  |  |  |  |  |  |  |  |
| YPR068C | HOS1 |  |  |  |  |  |  |  |  |
| YPR069C | SPE3 |  |  |  |  |  |  |  |  |
| YPR070W | MED1 |  |  |  |  |  |  |  |  |
| YPR072W | NOT5 |  |  |  |  |  |  |  |  |
| YPR073C | LTP1 |  |  |  |  |  |  |  |  |
| YPR074C | TKL1 |  |  |  |  |  |  |  |  |
| YPR075C | OPY2 |  |  |  |  |  |  |  |  |
| YPR079W | MRL1 |  |  |  |  |  |  |  |  |
| YPR080W | TEF1 |  |  |  |  |  |  |  |  |
| YPR081C | GRS2 |  |  |  |  |  |  |  |  |
| YPR082C | DIB1 |  |  |  |  |  |  |  |  |
| YPR083W | MDM36 |  |  |  |  |  |  |  |  |
| YPR085C | ASA1 |  |  |  |  |  |  |  |  |
| YPR086W | SUA7 |  |  |  |  |  |  |  |  |
| YPR088C | SRP54 |  |  |  |  |  |  |  |  |
| YPR091C | NVJ2 |  |  |  |  |  |  |  |  |
| YPR093C | ASR1 |  |  |  |  |  |  |  |  |
| YPR094W | RDS3 |  |  |  |  |  |  |  |  |
| YPR095C | SYT1 |  |  |  |  |  |  |  |  |
| YPR097W | LEC1 |  |  |  |  |  |  |  |  |
| YPR098C | TMH18 |  |  |  |  |  |  |  |  |
| YPR100W | MRPL51 |  |  |  |  |  |  |  |  |
| YPR101W | SNT309 |  |  |  |  |  |  |  |  |
| YPR102C | RPL11A |  |  |  |  |  |  |  |  |
| YPR103W | PRE2 |  |  |  |  |  |  |  |  |
| YPR104C | FHL1 |  |  |  |  |  |  |  |  |
| YPR105C | COG4 |  |  |  |  |  |  |  |  |
| YPR106W | ISR1 |  |  |  |  |  |  |  |  |
| YPR107C | YTH1 |  |  |  |  |  |  |  |  |
| YPR108W | RPN7 |  |  |  |  |  |  |  |  |
| YPR109W | GLD1 |  |  |  |  |  |  |  |  |
| YPR110C | RPC40 |  |  |  |  |  |  |  |  |
| YPR111W | DBF20 |  |  |  |  |  |  |  |  |
| YPR112C | MRD1 |  |  |  |  |  |  |  |  |
| YPR113W | PIS1 |  |  |  |  |  |  |  |  |
| YPR115W | RGC1 |  |  |  |  |  |  |  |  |
| YPR116W | RRG8 |  |  |  |  |  |  |  |  |
| YPR118W | MRI1 |  |  |  |  |  |  |  |  |
| YPR119W | CLB2 |  |  |  |  |  |  |  |  |
| YPR120C | CLB5 |  |  |  |  |  |  |  |  |
| YPR122W | AXL1 |  |  |  |  |  |  |  |  |
| YPR124W | CTR1 |  |  |  |  |  |  |  |  |
| YPR125W | YLH47 |  |  |  |  |  |  |  |  |
| YPR127W |  |  |  |  |  |  |  |  |  |
| YPR128C | ANT1 |  |  |  |  |  |  |  |  |
| YPR129W | SCD6 |  |  |  |  |  |  |  |  |
| YPR131C | NAT3 |  |  |  |  |  |  |  |  |
| YPR132W | RPS23B |  |  |  |  |  |  |  |  |
| YPR133C | SPN1 |  |  |  |  |  |  |  |  |
| YPR133W-A | TOM5 |  |  |  |  |  |  |  |  |
| YPR135W | CTF4 |  |  |  |  |  |  |  |  |
| YPR137W | RRP9 |  |  |  |  |  |  |  |  |
| YPR138C | MEP3 |  |  |  |  |  |  |  |  |
| YPR139C | LOA1 |  |  |  |  |  |  |  |  |
| YPR140W | TAZ1 |  |  |  |  |  |  |  |  |
| YPR141C | KAR3 |  |  |  |  |  |  |  |  |
| YPR143W | RRP15 |  |  |  |  |  |  |  |  |
| YPR144C | NOC4 |  |  |  |  |  |  |  |  |
| YPR145W | ASN1 |  |  |  |  |  |  |  |  |
| YPR147C |  |  |  |  |  |  |  |  |  |
| YPR148C |  |  |  |  |  |  |  |  |  |
| YPR149W | NCE102 |  |  |  |  |  |  |  |  |
| YPR151C | SUE1 |  |  |  |  |  |  |  |  |
| YPR152C | URN1 |  |  |  |  |  |  |  |  |
| YPR153W | MAY24 |  |  |  |  |  |  |  |  |
| YPR154W | PIN3 |  |  |  |  |  |  |  |  |
| YPR155C | NCA2 |  |  |  |  |  |  |  |  |
| YPR156C | TPO3 |  |  |  |  |  |  |  |  |
| YPR158W | CUR1 |  |  |  |  |  |  |  |  |
| YPR159W | KRE6 |  |  |  |  |  |  |  |  |
| YPR160W | GPH1 |  |  |  |  |  |  |  |  |
| YPR161C | SGV1 |  |  |  |  |  |  |  |  |
| YPR162C | ORC4 |  |  |  |  |  |  |  |  |
| YPR163C | TIF3 |  |  |  |  |  |  |  |  |
| YPR164W | MMS1 |  |  |  |  |  |  |  |  |
| YPR165W | RHO1 |  |  |  |  |  |  |  |  |
| YPR166C | MRP2 |  |  |  |  |  |  |  |  |
| YPR167C | MET16 |  |  |  |  |  |  |  |  |
| YPR168W | NUT2 |  |  |  |  |  |  |  |  |
| YPR169W | JIP5 |  |  |  |  |  |  |  |  |
| YPR170W-B |  |  |  |  |  |  |  |  |  |
| YPR171W | BSP1 |  |  |  |  |  |  |  |  |
| YPR173C | VPS4 |  |  |  |  |  |  |  |  |
| YPR174C | CSA1 |  |  |  |  |  |  |  |  |
| YPR175W | DPB2 |  |  |  |  |  |  |  |  |
| YPR176C | BET2 |  |  |  |  |  |  |  |  |
| YPR178W | PRP4 |  |  |  |  |  |  |  |  |
| YPR179C | HDA3 |  |  |  |  |  |  |  |  |
| YPR180W | AOS1 |  |  |  |  |  |  |  |  |
| YPR181C | SEC23 |  |  |  |  |  |  |  |  |
| YPR182W | SMX3 |  |  |  |  |  |  |  |  |
| YPR183W | DPM1 |  |  |  |  |  |  |  |  |
| YPR184W | GDB1 |  |  |  |  |  |  |  |  |
| YPR185W | ATG13 |  |  |  |  |  |  |  |  |
| YPR186C | PZF1 |  |  |  |  |  |  |  |  |
| YPR187W | RPO26 |  |  |  |  |  |  |  |  |
| YPR188C | MLC2 |  |  |  |  |  |  |  |  |
| YPR189W | SKI3 |  |  |  |  |  |  |  |  |
| YPR190C | RPC82 |  |  |  |  |  |  |  |  |
| YPR191W | QCR2 |  |  |  |  |  |  |  |  |
| YPR192W | AQY1 |  |  |  |  |  |  |  |  |
| YPR194C | OPT2 |  |  |  |  |  |  |  |  |
| Q0045 | COX1 |  |  |  |  |  |  |  |  |
| Q0080 | ATP8 |  |  |  |  |  |  |  |  |
| Q0085 | ATP6 |  |  |  |  |  |  |  |  |
| Q0105 | COB |  |  |  |  |  |  |  |  |
| Q0130 | OLI1 |  |  |  |  |  |  |  |  |
| Q0140 | VAR1 |  |  |  |  |  |  |  |  |
| Q0250 | COX2 |  |  |  |  |  |  |  |  |
| Q0275 | COX3 |  |  |  |  |  |  |  |  |
| YNCA0001W | HRA1 |  |  |  |  |  |  |  |  |
| YNCA0002W | TRN1 |  |  |  |  |  |  |  |  |
| YNCA0003W | SNR18 |  |  |  |  |  |  |  |  |
| YNCA0004W | TGA1 |  |  |  |  |  |  |  |  |
| YNCA0005W | SUP56 |  |  |  |  |  |  |  |  |
| YNCA0006C | tS(AGA)A |  |  |  |  |  |  |  |  |
| YNCB0001W | tL(UAA)B1 |  |  |  |  |  |  |  |  |
| YNCB0002W | tF(GAA)B |  |  |  |  |  |  |  |  |
| YNCB0003W | SNR56 |  |  |  |  |  |  |  |  |
| YNCB0004W | tI(AAU)B |  |  |  |  |  |  |  |  |
| YNCB0005C | tG(GCC)B |  |  |  |  |  |  |  |  |
| YNCB0006W | tS(AGA)B |  |  |  |  |  |  |  |  |
| YNCB0007W | tT(AGU)B |  |  |  |  |  |  |  |  |
| YNCB0009C | SNR161 |  |  |  |  |  |  |  |  |
| YNCB0010W | TLC1 |  |  |  |  |  |  |  |  |
| YNCB0011C | tV(UAC)B |  |  |  |  |  |  |  |  |
| YNCB0012W | tL(UAA)B2 |  |  |  |  |  |  |  |  |
| YNCB0013W | tQ(UUG)B |  |  |  |  |  |  |  |  |
| YNCB0014W | TBRT |  |  |  |  |  |  |  |  |
| YNCB0015W | tR(UCU)B |  |  |  |  |  |  |  |  |
| YNCB0016W | tD(GUC)B |  |  |  |  |  |  |  |  |
| YNCB0017C | tC(GCA)B |  |  |  |  |  |  |  |  |
| YNCB0018W | tE(UUC)B |  |  |  |  |  |  |  |  |
| YNCB0019C | LSR1 |  |  |  |  |  |  |  |  |
| YNCC0001C | tE(UUC)C |  |  |  |  |  |  |  |  |
| YNCC0002W | SUP53 |  |  |  |  |  |  |  |  |
| YNCC0003C | SNR43 |  |  |  |  |  |  |  |  |
| YNCC0004C | SUF2 |  |  |  |  |  |  |  |  |
| YNCC0005W | tN(GUU)C |  |  |  |  |  |  |  |  |
| YNCC0006C | SNR33 |  |  |  |  |  |  |  |  |
| YNCC0007C | SUF16 |  |  |  |  |  |  |  |  |
| YNCC0008W | tM(CAU)C |  |  |  |  |  |  |  |  |
| YNCC0009C | tK(CUU)C |  |  |  |  |  |  |  |  |
| YNCC0010C | tQ(UUG)C |  |  |  |  |  |  |  |  |
| YNCC0011W | SNR65 |  |  |  |  |  |  |  |  |
| YNCC0012C | SNR189 |  |  |  |  |  |  |  |  |
| YNCC0013W | SUP61 |  |  |  |  |  |  |  |  |
| YNCC0014W | tT(AGU)C |  |  |  |  |  |  |  |  |
| YNCD0001W | tG(GCC)D1 |  |  |  |  |  |  |  |  |
| YNCD0002C | SNR63 |  |  |  |  |  |  |  |  |
| YNCD0003W | tK(UUU)D |  |  |  |  |  |  |  |  |
| YNCD0004W | tA(AGC)D |  |  |  |  |  |  |  |  |
| YNCD0005C | tT(AGU)D |  |  |  |  |  |  |  |  |
| YNCD0006W | tS(AGA)D1 |  |  |  |  |  |  |  |  |
| YNCD0007C | tV(UAC)D |  |  |  |  |  |  |  |  |
| YNCD0008W | tL(UAA)D |  |  |  |  |  |  |  |  |
| YNCD0009W | tQ(UUG)D1 |  |  |  |  |  |  |  |  |
| YNCD0010C | SNR47 |  |  |  |  |  |  |  |  |
| YNCD0011W | tR(UCU)D |  |  |  |  |  |  |  |  |
| YNCD0012W | tD(GUC)D |  |  |  |  |  |  |  |  |
| YNCD0013C | tR(ACG)D |  |  |  |  |  |  |  |  |
| YNCD0014C | tQ(UUG)D2 |  |  |  |  |  |  |  |  |
| YNCD0015C | tI(AAU)D |  |  |  |  |  |  |  |  |
| YNCD0016C | tQ(UUG)D3 |  |  |  |  |  |  |  |  |
| YNCD0017W | tI(UAU)D |  |  |  |  |  |  |  |  |
| YNCD0018W | SUP2 |  |  |  |  |  |  |  |  |
| YNCD0019C | tS(AGA)D2 |  |  |  |  |  |  |  |  |
| YNCD0020W | tG(GCC)D2 |  |  |  |  |  |  |  |  |
| YNCD0021C | tE(CUC)D |  |  |  |  |  |  |  |  |
| YNCD0022C | tV(CAC)D |  |  |  |  |  |  |  |  |
| YNCD0023C | tF(GAA)D |  |  |  |  |  |  |  |  |
| YNCD0024C | tX(XXX)D |  |  |  |  |  |  |  |  |
| YNCD0025W | EMT1 |  |  |  |  |  |  |  |  |
| YNCD0026W | tK(CUU)D1 |  |  |  |  |  |  |  |  |
| YNCD0027C | SUF3 |  |  |  |  |  |  |  |  |
| YNCD0028W | tS(AGA)D3 |  |  |  |  |  |  |  |  |
| YNCD0029C | tK(CUU)D2 |  |  |  |  |  |  |  |  |
| YNCD0030W | SNR13 |  |  |  |  |  |  |  |  |
| YNCD0031C | tL(CAA)D |  |  |  |  |  |  |  |  |
| YNCD0032C | SNR84 |  |  |  |  |  |  |  |  |
| YNCE0001C | SNR80 |  |  |  |  |  |  |  |  |
| YNCE0002W | SNR67 |  |  |  |  |  |  |  |  |
| YNCE0003W | SNR53 |  |  |  |  |  |  |  |  |
| YNCE0004C | tG(GCC)E |  |  |  |  |  |  |  |  |
| YNCE0005W | tS(AGA)E |  |  |  |  |  |  |  |  |
| YNCE0006W | IMT4 |  |  |  |  |  |  |  |  |
| YNCE0007C | RPR1 |  |  |  |  |  |  |  |  |
| YNCE0008C | tQ(UUG)E2 |  |  |  |  |  |  |  |  |
| YNCE0009C | tK(CUU)E1 |  |  |  |  |  |  |  |  |
| YNCE0010W | tR(UCU)E |  |  |  |  |  |  |  |  |
| YNCE0011C | SNR14 |  |  |  |  |  |  |  |  |
| YNCE0012W | tE(UUC)E1 |  |  |  |  |  |  |  |  |
| YNCE0013W | tH(GUG)E1 |  |  |  |  |  |  |  |  |
| YNCE0014W | tQ(UUG)E1 |  |  |  |  |  |  |  |  |
| YNCE0015C | SUP19 |  |  |  |  |  |  |  |  |
| YNCE0016C | tA(UGC)E |  |  |  |  |  |  |  |  |
| YNCE0017W | SRG1 |  |  |  |  |  |  |  |  |
| YNCE0018W | tE(UUC)E2 |  |  |  |  |  |  |  |  |
| YNCE0019W | SNR4 |  |  |  |  |  |  |  |  |
| YNCE0020C | SNR52 |  |  |  |  |  |  |  |  |
| YNCE0021C | tH(GUG)E2 |  |  |  |  |  |  |  |  |
| YNCE0023W | tV(AAC)E1 |  |  |  |  |  |  |  |  |
| YNCE0024W | SCR1 |  |  |  |  |  |  |  |  |
| YNCE0025C | tI(AAU)E1 |  |  |  |  |  |  |  |  |
| YNCE0026W | tV(AAC)E2 |  |  |  |  |  |  |  |  |
| YNCE0027C | tE(UUC)E3 |  |  |  |  |  |  |  |  |
| YNCE0028C | tR(ACG)E |  |  |  |  |  |  |  |  |
| YNCE0029C | tI(AAU)E2 |  |  |  |  |  |  |  |  |
| YNCF0002W | SUF9 |  |  |  |  |  |  |  |  |
| YNCF0004C | tN(GUU)F |  |  |  |  |  |  |  |  |
| YNCF0005C | tF(GAA)F |  |  |  |  |  |  |  |  |
| YNCF0006W | SUF20 |  |  |  |  |  |  |  |  |
| YNCF0007W | SUP11 |  |  |  |  |  |  |  |  |
| YNCF0008C | tG(GCC)F2 |  |  |  |  |  |  |  |  |
| YNCF0009C | tS(GCU)F |  |  |  |  |  |  |  |  |
| YNCF0011C | tA(AGC)F |  |  |  |  |  |  |  |  |
| YNCF0012C | SUP6 |  |  |  |  |  |  |  |  |
| YNCF0014C | tK(CUU)F |  |  |  |  |  |  |  |  |
| YNCG0001C | RME3 |  |  |  |  |  |  |  |  |
| YNCG0002C | tV(AAC)G3 |  |  |  |  |  |  |  |  |
| YNCG0003C | tH(GUG)G1 |  |  |  |  |  |  |  |  |
| YNCG0004W | tK(UUU)G1 |  |  |  |  |  |  |  |  |
| YNCG0005W | tK(CUU)G1 |  |  |  |  |  |  |  |  |
| YNCG0006C | RME2 |  |  |  |  |  |  |  |  |
| YNCG0007C | tK(CUU)G2 |  |  |  |  |  |  |  |  |
| YNCG0008W | tL(CAA)G1 |  |  |  |  |  |  |  |  |
| YNCG0009C | tW(CCA)G1 |  |  |  |  |  |  |  |  |
| YNCG0010W | SNR82 |  |  |  |  |  |  |  |  |
| YNCG0011W | tH(GUG)G2 |  |  |  |  |  |  |  |  |
| YNCG0012W | SOE1 |  |  |  |  |  |  |  |  |
| YNCG0013W | SNR10 |  |  |  |  |  |  |  |  |
| YNCG0014C | SNR39 |  |  |  |  |  |  |  |  |
| YNCG0015C | SNR39B |  |  |  |  |  |  |  |  |
| YNCG0016C | tE(UUC)G2 |  |  |  |  |  |  |  |  |
| YNCG0017W | tR(UCU)G1 |  |  |  |  |  |  |  |  |
| YNCG0018C | tV(AAC)G1 |  |  |  |  |  |  |  |  |
| YNCG0019W | SUP54 |  |  |  |  |  |  |  |  |
| YNCG0020C | tF(GAA)G |  |  |  |  |  |  |  |  |
| YNCG0021W | tD(GUC)G1 |  |  |  |  |  |  |  |  |
| YNCG0022W | tE(UUC)G3 |  |  |  |  |  |  |  |  |
| YNCG0023C | tD(GUC)G2 |  |  |  |  |  |  |  |  |
| YNCG0024W | SNR46 |  |  |  |  |  |  |  |  |
| YNCG0025C | tS(AGA)G |  |  |  |  |  |  |  |  |
| YNCG0026W | SNR48 |  |  |  |  |  |  |  |  |
| YNCG0027W | tT(UGU)G1 |  |  |  |  |  |  |  |  |
| YNCG0028W | tL(GAG)G |  |  |  |  |  |  |  |  |
| YNCG0029C | tK(UUU)G2 |  |  |  |  |  |  |  |  |
| YNCG0030C | tC(GCA)G |  |  |  |  |  |  |  |  |
| YNCG0031W | tN(GUU)G |  |  |  |  |  |  |  |  |
| YNCG0032W | tR(UCU)G3 |  |  |  |  |  |  |  |  |
| YNCG0033W | tI(AAU)G |  |  |  |  |  |  |  |  |
| YNCG0034W | tA(AGC)G |  |  |  |  |  |  |  |  |
| YNCG0035W | SUF4 |  |  |  |  |  |  |  |  |
| YNCG0036W | tA(UGC)G |  |  |  |  |  |  |  |  |
| YNCG0037W | tV(AAC)G2 |  |  |  |  |  |  |  |  |
| YNCG0038C | tR(UCU)G2 |  |  |  |  |  |  |  |  |
| YNCG0039W | tG(GCC)G1 |  |  |  |  |  |  |  |  |
| YNCG0040C | tL(CAA)G3 |  |  |  |  |  |  |  |  |
| YNCG0041W | tK(CUU)G3 |  |  |  |  |  |  |  |  |
| YNCG0042C | tW(CCA)G2 |  |  |  |  |  |  |  |  |
| YNCG0043C | tG(GCC)G2 |  |  |  |  |  |  |  |  |
| YNCG0044C | SNR7-L |  |  |  |  |  |  |  |  |
| YNCG0045C | SNR7-S |  |  |  |  |  |  |  |  |
| YNCG0046W | tT(UGU)G2 |  |  |  |  |  |  |  |  |
| YNCH0001W | tH(GUG)H |  |  |  |  |  |  |  |  |
| YNCH0002C | tV(AAC)H |  |  |  |  |  |  |  |  |
| YNCH0003C | tT(AGU)H |  |  |  |  |  |  |  |  |
| YNCH0004C | tS(AGA)H |  |  |  |  |  |  |  |  |
| YNCH0005W | tQ(UUG)H |  |  |  |  |  |  |  |  |
| YNCH0006C | tA(AGC)H |  |  |  |  |  |  |  |  |
| YNCH0009C | tF(GAA)H1 |  |  |  |  |  |  |  |  |
| YNCH0010C | tF(GAA)H2 |  |  |  |  |  |  |  |  |
| YNCH0011W | SUT169 |  |  |  |  |  |  |  |  |
| YNCH0012W | SNR32 |  |  |  |  |  |  |  |  |
| YNCH0013C | SUF8 |  |  |  |  |  |  |  |  |
| YNCH0014W | SNR71 |  |  |  |  |  |  |  |  |
| YNCH0015W | tT(UGU)H |  |  |  |  |  |  |  |  |
| YNCH0016C | tV(CAC)H |  |  |  |  |  |  |  |  |
| YNCI0001W | SNR68 |  |  |  |  |  |  |  |  |
| YNCI0002W | tT(AGU)I1 |  |  |  |  |  |  |  |  |
| YNCI0003C | tI(AAU)I1 |  |  |  |  |  |  |  |  |
| YNCI0004W | tE(CUC)I |  |  |  |  |  |  |  |  |
| YNCI0005W | tI(AAU)I2 |  |  |  |  |  |  |  |  |
| YNCI0006W | SUP17 |  |  |  |  |  |  |  |  |
| YNCI0007C | tK(CUU)I |  |  |  |  |  |  |  |  |
| YNCI0008C | tD(GUC)I1 |  |  |  |  |  |  |  |  |
| YNCI0009W | tT(AGU)I2 |  |  |  |  |  |  |  |  |
| YNCI0010C | tD(GUC)I2 |  |  |  |  |  |  |  |  |
| YNCI0011W | tE(UUC)I |  |  |  |  |  |  |  |  |
| YNCI0012C | ICR1 |  |  |  |  |  |  |  |  |
| YNCI0013W | PWR1 |  |  |  |  |  |  |  |  |
| YNCJ0001C | tT(AGU)J |  |  |  |  |  |  |  |  |
| YNCJ0002C | tE(UUC)J |  |  |  |  |  |  |  |  |
| YNCJ0003C | SNR128 |  |  |  |  |  |  |  |  |
| YNCJ0004C | SNR190 |  |  |  |  |  |  |  |  |
| YNCJ0005C | tA(AGC)J |  |  |  |  |  |  |  |  |
| YNCJ0006W | tD(GUC)J1 |  |  |  |  |  |  |  |  |
| YNCJ0007C | SNR37 |  |  |  |  |  |  |  |  |
| YNCJ0008W | tR(ACG)J |  |  |  |  |  |  |  |  |
| YNCJ0009C | SNR60 |  |  |  |  |  |  |  |  |
| YNCJ0010W | SUP7 |  |  |  |  |  |  |  |  |
| YNCJ0011W | tR(UCU)J1 |  |  |  |  |  |  |  |  |
| YNCJ0012W | tD(GUC)J2 |  |  |  |  |  |  |  |  |
| YNCJ0013C | tD(GUC)J3 |  |  |  |  |  |  |  |  |
| YNCJ0014C | tR(UCU)J2 |  |  |  |  |  |  |  |  |
| YNCJ0015W | tV(AAC)J |  |  |  |  |  |  |  |  |
| YNCJ0016C | EMT5 |  |  |  |  |  |  |  |  |
| YNCJ0017C | tG(GCC)J1 |  |  |  |  |  |  |  |  |
| YNCJ0018W | tK(CUU)J |  |  |  |  |  |  |  |  |
| YNCJ0019C | tW(CCA)J |  |  |  |  |  |  |  |  |
| YNCJ0020W | EMT3 |  |  |  |  |  |  |  |  |
| YNCJ0021C | SUP51 |  |  |  |  |  |  |  |  |
| YNCJ0022C | IMT3 |  |  |  |  |  |  |  |  |
| YNCJ0023C | tS(AGA)J |  |  |  |  |  |  |  |  |
| YNCJ0024W | SUF23 |  |  |  |  |  |  |  |  |
| YNCJ0025W | HSX1 |  |  |  |  |  |  |  |  |
| YNCJ0026W | tD(GUC)J4 |  |  |  |  |  |  |  |  |
| YNCJ0027C | SUP4 |  |  |  |  |  |  |  |  |
| YNCJ0028C | IRT1 |  |  |  |  |  |  |  |  |
| YNCJ0029W | tL(UAG)J |  |  |  |  |  |  |  |  |
| YNCJ0030W | SNR3 |  |  |  |  |  |  |  |  |
| YNCK0001W | SNR64 |  |  |  |  |  |  |  |  |
| YNCK0002C | TRT2 |  |  |  |  |  |  |  |  |
| YNCK0003W | tN(GUU)K |  |  |  |  |  |  |  |  |
| YNCK0004W | tL(UAA)K |  |  |  |  |  |  |  |  |
| YNCK0005C | tE(UUC)K |  |  |  |  |  |  |  |  |
| YNCK0006C | tR(UCU)K |  |  |  |  |  |  |  |  |
| YNCK0007W | tK(CUU)K |  |  |  |  |  |  |  |  |
| YNCK0008W | tA(AGC)K1 |  |  |  |  |  |  |  |  |
| YNCK0009W | SNR38 |  |  |  |  |  |  |  |  |
| YNCK0010W | tW(CCA)K |  |  |  |  |  |  |  |  |
| YNCK0011C | tV(AAC)K1 |  |  |  |  |  |  |  |  |
| YNCK0012C | tH(GUG)K |  |  |  |  |  |  |  |  |
| YNCK0013W | SNR69 |  |  |  |  |  |  |  |  |
| YNCK0014W | tV(AAC)K2 |  |  |  |  |  |  |  |  |
| YNCK0015C | SNR87 |  |  |  |  |  |  |  |  |
| YNCK0016W | tL(CAA)K |  |  |  |  |  |  |  |  |
| YNCK0017W | tR(ACG)K |  |  |  |  |  |  |  |  |
| YNCK0018C | tD(GUC)K |  |  |  |  |  |  |  |  |
| YNCK0019W | tA(AGC)K2 |  |  |  |  |  |  |  |  |
| YNCK0020C | SNR42 |  |  |  |  |  |  |  |  |
| YNCK0021W | tK(UUU)K |  |  |  |  |  |  |  |  |
| YNCL0001W | tP(UGG)L |  |  |  |  |  |  |  |  |
| YNCL0002C | tS(AGA)L |  |  |  |  |  |  |  |  |
| YNCL0003W | SNR30 |  |  |  |  |  |  |  |  |
| YNCL0004C | tA(UGC)L |  |  |  |  |  |  |  |  |
| YNCL0005C | SNR79 |  |  |  |  |  |  |  |  |
| YNCL0006W | SNR6 |  |  |  |  |  |  |  |  |
| YNCL0007W | tR(ACG)L |  |  |  |  |  |  |  |  |
| YNCL0008W | tD(GUC)L1 |  |  |  |  |  |  |  |  |
| YNCL0009C | tQ(UUG)L |  |  |  |  |  |  |  |  |
| YNCL0010C | RDN37 |  |  |  |  |  |  |  |  |
| YNCL0018W | RDN5 |  |  |  |  |  |  |  |  |
| YNCL0032C | tL(UAG)L1 |  |  |  |  |  |  |  |  |
| YNCL0033C | tI(UAU)L |  |  |  |  |  |  |  |  |
| YNCL0034W | tL(CAA)L |  |  |  |  |  |  |  |  |
| YNCL0035C | tA(AGC)L |  |  |  |  |  |  |  |  |
| YNCL0036C | tV(AAC)L |  |  |  |  |  |  |  |  |
| YNCL0037W | tL(UAG)L2 |  |  |  |  |  |  |  |  |
| YNCL0038W | tI(AAU)L1 |  |  |  |  |  |  |  |  |
| YNCL0039W | tX(XXX)L |  |  |  |  |  |  |  |  |
| YNCL0040W | tD(GUC)L2 |  |  |  |  |  |  |  |  |
| YNCL0041C | SNR61 |  |  |  |  |  |  |  |  |
| YNCL0042C | SNR55 |  |  |  |  |  |  |  |  |
| YNCL0043C | SNR57 |  |  |  |  |  |  |  |  |
| YNCL0044W | tE(UUC)L |  |  |  |  |  |  |  |  |
| YNCL0045W | TRR4 |  |  |  |  |  |  |  |  |
| YNCL0046W | SNR44 |  |  |  |  |  |  |  |  |
| YNCL0047W | tK(UUU)L |  |  |  |  |  |  |  |  |
| YNCL0048W | SNR34 |  |  |  |  |  |  |  |  |
| YNCL0049C | tL(UAA)L |  |  |  |  |  |  |  |  |
| YNCL0050C | tN(GUU)L |  |  |  |  |  |  |  |  |
| YNCL0051C | tI(AAU)L2 |  |  |  |  |  |  |  |  |
| YNCM0001W | PHO94IncRNA |  |  |  |  |  |  |  |  |
| YNCM0002C | SNR85 |  |  |  |  |  |  |  |  |
| YNCM0003W | ZOD1 |  |  |  |  |  |  |  |  |
| YNCM0004C | tR(UCU)M2 |  |  |  |  |  |  |  |  |
| YNCM0005C | SNR54 |  |  |  |  |  |  |  |  |
| YNCM0006W | SUP5 |  |  |  |  |  |  |  |  |
| YNCM0007C | tG(GCC)M |  |  |  |  |  |  |  |  |
| YNCM0008C | SUF7 |  |  |  |  |  |  |  |  |
| YNCM0009C | tS(AGA)M |  |  |  |  |  |  |  |  |
| YNCM0010W | tE(UUC)M |  |  |  |  |  |  |  |  |
| YNCM0011W | SNR78 |  |  |  |  |  |  |  |  |
| YNCM0012W | SNR77 |  |  |  |  |  |  |  |  |
| YNCM0013W | SNR76 |  |  |  |  |  |  |  |  |
| YNCM0014W | SNR75 |  |  |  |  |  |  |  |  |
| YNCM0015W | SNR74 |  |  |  |  |  |  |  |  |
| YNCM0016W | SNR73 |  |  |  |  |  |  |  |  |
| YNCM0017W | SNR72 |  |  |  |  |  |  |  |  |
| YNCM0018C | tA(AGC)M1 |  |  |  |  |  |  |  |  |
| YNCM0019W | tF(GAA)M |  |  |  |  |  |  |  |  |
| YNCM0020W | tH(GUG)M |  |  |  |  |  |  |  |  |
| YNCM0021C | tV(AAC)M1 |  |  |  |  |  |  |  |  |
| YNCM0022C | tW(CCA)M |  |  |  |  |  |  |  |  |
| YNCM0023C | tV(AAC)M2 |  |  |  |  |  |  |  |  |
| YNCM0024W | tD(GUC)M |  |  |  |  |  |  |  |  |
| YNCM0025C | tK(CUU)M |  |  |  |  |  |  |  |  |
| YNCM0026C | SNR24 |  |  |  |  |  |  |  |  |
| YNCM0027W | tL(CAA)M |  |  |  |  |  |  |  |  |
| YNCM0028C | EMT4 |  |  |  |  |  |  |  |  |
| YNCM0029C | tV(AAC)M3 |  |  |  |  |  |  |  |  |
| YNCM0030W | SNR83 |  |  |  |  |  |  |  |  |
| YNCM0031W | SNR11 |  |  |  |  |  |  |  |  |
| YNCM0032C | RNA170 |  |  |  |  |  |  |  |  |
| YNCM0033C | tR(UCU)M1 |  |  |  |  |  |  |  |  |
| YNCM0034C | SNR86 |  |  |  |  |  |  |  |  |
| YNCM0035C | tA(AGC)M2 |  |  |  |  |  |  |  |  |
| YNCM0036C | CDC65 |  |  |  |  |  |  |  |  |
| YNCM0037W | SUP8 |  |  |  |  |  |  |  |  |
| YNCN0001W | SNR40 |  |  |  |  |  |  |  |  |
| YNCN0002C | SUF6 |  |  |  |  |  |  |  |  |
| YNCN0003W | tN(GUU)N1 |  |  |  |  |  |  |  |  |
| YNCN0004W | tT(AGU)N1 |  |  |  |  |  |  |  |  |
| YNCN0005C | SNR19 |  |  |  |  |  |  |  |  |
| YNCN0006W | tF(GAA)N |  |  |  |  |  |  |  |  |
| YNCN0007W | tL(CAA)N |  |  |  |  |  |  |  |  |
| YNCN0008C | tD(GUC)N |  |  |  |  |  |  |  |  |
| YNCN0009W | tP(UGG)N1 |  |  |  |  |  |  |  |  |
| YNCN0010C | tT(AGU)N2 |  |  |  |  |  |  |  |  |
| YNCN0011W | tP(UGG)N2 |  |  |  |  |  |  |  |  |
| YNCN0012C | tI(AAU)N1 |  |  |  |  |  |  |  |  |
| YNCN0013W | NME1 |  |  |  |  |  |  |  |  |
| YNCN0014W | SNR66 |  |  |  |  |  |  |  |  |
| YNCN0015W | tI(AAU)N2 |  |  |  |  |  |  |  |  |
| YNCN0016C | SUF10 |  |  |  |  |  |  |  |  |
| YNCN0017W | tN(GUU)N2 |  |  |  |  |  |  |  |  |
| YNCN0018W | SNR49 |  |  |  |  |  |  |  |  |
| YNCN0019C | SNR191 |  |  |  |  |  |  |  |  |
| YNCN0020C | tL(UAA)N |  |  |  |  |  |  |  |  |
| YNCO0001C | SUF1 |  |  |  |  |  |  |  |  |
| YNCO0002W | tT(AGU)O1 |  |  |  |  |  |  |  |  |
| YNCO0003C | SNR58 |  |  |  |  |  |  |  |  |
| YNCO0004C | tG(GCC)O1 |  |  |  |  |  |  |  |  |
| YNCO0005W | tN(GUU)O1 |  |  |  |  |  |  |  |  |
| YNCO0006W | SNR81 |  |  |  |  |  |  |  |  |
| YNCO0007W | SNR50 |  |  |  |  |  |  |  |  |
| YNCO0008W | tS(GCU)O |  |  |  |  |  |  |  |  |
| YNCO0009W | SUF17 |  |  |  |  |  |  |  |  |
| YNCO0010W | SUP3 |  |  |  |  |  |  |  |  |
| YNCO0011W | tP(UGG)O1 |  |  |  |  |  |  |  |  |
| YNCO0012C | tR(ACG)O |  |  |  |  |  |  |  |  |
| YNCO0013C | tT(AGU)O2 |  |  |  |  |  |  |  |  |
| YNCO0014C | SNR9 |  |  |  |  |  |  |  |  |
| YNCO0015C | SNR62 |  |  |  |  |  |  |  |  |
| YNCO0016W | tK(UUU)O |  |  |  |  |  |  |  |  |
| YNCO0017W | SUF11 |  |  |  |  |  |  |  |  |
| YNCO0018W | tN(GUU)O2 |  |  |  |  |  |  |  |  |
| YNCO0019W | tD(GUC)O |  |  |  |  |  |  |  |  |
| YNCO0020C | SUF5 |  |  |  |  |  |  |  |  |
| YNCO0021C | tV(AAC)O |  |  |  |  |  |  |  |  |
| YNCO0022C | SNR36 |  |  |  |  |  |  |  |  |
| YNCO0023W | IMT1 |  |  |  |  |  |  |  |  |
| YNCO0024C | SNR35 |  |  |  |  |  |  |  |  |
| YNCO0025W | SNR17A |  |  |  |  |  |  |  |  |
| YNCO0026W | SNR8 |  |  |  |  |  |  |  |  |
| YNCO0027C | SNR31 |  |  |  |  |  |  |  |  |
| YNCO0028W | SNR5 |  |  |  |  |  |  |  |  |
| YNCO0029C | tA(UGC)O |  |  |  |  |  |  |  |  |
| YNCO0030W | EMT2 |  |  |  |  |  |  |  |  |
| YNCO0031W | tP(UGG)O3 |  |  |  |  |  |  |  |  |
| YNCP0001C | tW(CCA)P |  |  |  |  |  |  |  |  |
| YNCP0002W | GAL4IncRNA |  |  |  |  |  |  |  |  |
| YNCP0003W | SNR59 |  |  |  |  |  |  |  |  |
| YNCP0004C | tE(UUC)P |  |  |  |  |  |  |  |  |
| YNCP0006C | IMT2 |  |  |  |  |  |  |  |  |
| YNCP0007C | tC(GCA)P1 |  |  |  |  |  |  |  |  |
| YNCP0008C | tF(GAA)P1 |  |  |  |  |  |  |  |  |
| YNCP0009W | tG(GCC)P1 |  |  |  |  |  |  |  |  |
| YNCP0010W | tK(CUU)P |  |  |  |  |  |  |  |  |
| YNCP0011C | tF(GAA)P2 |  |  |  |  |  |  |  |  |
| YNCP0012W | SUP16 |  |  |  |  |  |  |  |  |
| YNCP0013C | SNR51 |  |  |  |  |  |  |  |  |
| YNCP0014C | SNR70 |  |  |  |  |  |  |  |  |
| YNCP0015C | SNR41 |  |  |  |  |  |  |  |  |
| YNCP0016W | tT(UGU)P |  |  |  |  |  |  |  |  |
| YNCP0017C | tK(UUU)P |  |  |  |  |  |  |  |  |
| YNCP0018C | tC(GCA)P2 |  |  |  |  |  |  |  |  |
| YNCP0019W | tN(GUU)P |  |  |  |  |  |  |  |  |
| YNCP0020W | tI(AAU)P1 |  |  |  |  |  |  |  |  |
| YNCP0021W | SNR45 |  |  |  |  |  |  |  |  |
| YNCP0022W | tA(AGC)P |  |  |  |  |  |  |  |  |
| YNCP0023W | tG(GCC)P2 |  |  |  |  |  |  |  |  |
| YNCP0024C | tI(AAU)P2 |  |  |  |  |  |  |  |  |
| YNCQ0001W | tP(UGG)Q |  |  |  |  |  |  |  |  |
| YNCQ0002W | 15S_RRNA |  |  |  |  |  |  |  |  |
| YNCQ0003W | tW(UCA)Q |  |  |  |  |  |  |  |  |
| YNCQ0004W | tE(UUC)Q |  |  |  |  |  |  |  |  |
| YNCQ0005W | tS(UGA)Q2 |  |  |  |  |  |  |  |  |
| YNCQ0006W | 21S_RRNA |  |  |  |  |  |  |  |  |
| YNCQ0007W | tT(UGU)Q1 |  |  |  |  |  |  |  |  |
| YNCQ0008W | tC(GCA)Q |  |  |  |  |  |  |  |  |
| YNCQ0009W | tH(GUG)Q |  |  |  |  |  |  |  |  |
| YNCQ0010W | tL(UAA)Q |  |  |  |  |  |  |  |  |
| YNCQ0011W | tQ(UUG)Q |  |  |  |  |  |  |  |  |
| YNCQ0012W | tK(UUU)Q |  |  |  |  |  |  |  |  |
| YNCQ0013W | tR(UCU)Q1 |  |  |  |  |  |  |  |  |
| YNCQ0014W | tG(UCC)Q |  |  |  |  |  |  |  |  |
| YNCQ0015W | tD(GUC)Q |  |  |  |  |  |  |  |  |
| YNCQ0016W | tS(GCU)Q1 |  |  |  |  |  |  |  |  |
| YNCQ0018W | tA(UGC)Q |  |  |  |  |  |  |  |  |
| YNCQ0019W | tI(GAU)Q |  |  |  |  |  |  |  |  |
| YNCQ0020W | tY(GUA)Q |  |  |  |  |  |  |  |  |
| YNCQ0021W | tN(GUU)Q |  |  |  |  |  |  |  |  |
| YNCQ0022W | tM(CAU)Q1 |  |  |  |  |  |  |  |  |
| YNCQ0023W | tF(GAA)Q |  |  |  |  |  |  |  |  |
| YNCQ0024C | tT(UAG)Q2 |  |  |  |  |  |  |  |  |
| YNCQ0025W | tV(UAC)Q |  |  |  |  |  |  |  |  |
| YNCQ0026W | tM(CAU)Q2 |  |  |  |  |  |  |  |  |
| YNCQ0027W | RPM1 |  |  |  |  |  |  |  |  |

Table S3 Uncharacterized genes in the core set of genes in *S. cerevisiae.* The four pairs of columns are uncharacterized genes with orthologs in *A. gossypii*, uncharacterized genes with orthologs in pre-whole genome duplication species, uncharacterized genes that appear to be conserved only in post-WGD species, and uncharacterized genes that appear to be found only in *S. cerevisiae.*

| preWGD, in  *A. gossypii* | Common name | preWGD, not in  *A. gossypii* | Common name | Broadly Post WGD | Common name | *Saccharomyces* only | Common name | S. cerevisiae only | Common name |
| --- | --- | --- | --- | --- | --- | --- | --- | --- | --- |
| Number genes: 222 |  | Number genes: 69 |  | Number genes: 8 |  | Number genes: 29 |  | Number genes: 36 |  |
| YAL037W |  | YAR023C | DFP1 | YER076C |  | YBR184W |  | YBR196C-A |  |
| YBL026W-A |  | YAL061W | BDH2 | YGR153W |  | YBR221W-A |  | YDL204W-A |  |
| YBL029W |  | YBR047W | FMP23 | YHL044W | DFP4 | YCL021W-A |  | YDR246W-A |  |
| YBL039W-B | MIN6 | YBR201C-A | MIN7 | YOL014W |  | YCL068C |  | YEL035C | UTR5 |
| YBL059W | IAI11 | YCL002C |  | YNCH0007W | RUF5 | YCR007C | DFP3 | YER084W |  |
| YBL081W |  | YCL049C |  | YNCF0001C | RUF21 | YDL007C-A |  | YFR032C-B | MIN10 |
| YBL086C |  | YDL027C | MRX9 | YNCF0010C | RUF22 | YDL241W |  | YGL007W | BRP1 |
| YBL095W | MRX3 | YDL199C |  | YNCF0013W | RUF23 | YDR042C |  | YGL188C-A |  |
| YBR033W | EDS1 | YDL218W |  |  |  | YEL028W |  | YGR035W-A |  |
| YBR053C |  | YDR061W |  |  |  | YEL068C |  | YGR204C-A |  |
| YBR056W-A | MNC1 | YDR222W |  |  |  | YER078W-A |  | YGR240C-A |  |
| YBR062C |  | YDR387C | CIN10 |  |  | YER085C |  | YHL042W |  |
| YBR096W |  | YDR391C |  |  |  | YER121W |  | YHR022C-A |  |
| YBR138C |  | YFL051C | FLO12 |  |  | YFL012W |  | YHR050W-A |  |
| YBR220C |  | YGL041W-A | DPC13 |  |  | YFR012W-A |  | YHR086W-A |  |
| YBR225W |  | YGL159W |  |  |  | YGR161W-C |  | YIL054W |  |
| YBR255C-A | RCF3 | YGL185C |  |  |  | YGR201C |  | YJL027C |  |
| YBR284W |  | YGL194C-A |  |  |  | YIL014C-A |  | YJL052C-A |  |
| YBR287W |  | YGL230C |  |  |  | YIL092W |  | YKL183C-A |  |
| YCL048W-A |  | YGR015C | EAT1 |  |  | YIL102C |  | YLR285C-A |  |
| YCR043C |  | YGR035C |  |  |  | YJL043W |  | YML054C-A |  |
| YCR051W |  | YGR126W |  |  |  | YKL106C-A |  | YML100W-A |  |
| YCR090C |  | YHL026C |  |  |  | YLR036C |  | YMR001C-A |  |
| YCR095C | OCA4 | YHR048W | YHK8 |  |  | YLR125W |  | YMR008C-A |  |
| YDL057W |  | YHR210C |  |  |  | YML007C-A | MIN4 | YMR105W-A |  |
| YDL086W |  | YIL001W |  |  |  | YMR317W |  | YMR182W-A | MIN3 |
| YDL109C |  | YJL132W |  |  |  | YNL046W |  | YMR242W-A |  |
| YDL114W |  | YJL133C-A | DPI8 |  |  | YOR268C |  | YMR247W-A |  |
| YDL157C | DMO2 | YJL193W |  |  |  | YPR071W |  | YMR272W-B |  |
| YDL177C |  | YJL206C |  |  |  |  |  | YMR315W-A |  |
| YDL186W |  | YJR003C | MRX12 |  |  |  |  | YNL130C-A | DGR1 |
| YDL211C |  | YJR015W |  |  |  |  |  | YOL019W-A |  |
| YDL233W | MFG1 | YJR098C |  |  |  |  |  | YOL097W-A |  |
| YDR020C | DAS2 | YJR149W |  |  |  |  |  | YOR008C-A |  |
| YDR056C | EMC10 | YKL070W |  |  |  |  |  | YOR032W-A |  |
| YDR067C | OCA6 | YKR015C |  |  |  |  |  | YPL152W-A |  |
| YDR111C | ALT2 | YLL054C |  |  |  |  |  |  |  |
| YDR119W | VBA4 | YLR046C |  |  |  |  |  |  |  |
| YDR124W |  | YLR225C |  |  |  |  |  |  |  |
| YDR248C |  | YLR267W | BOP2 |  |  |  |  |  |  |
| YDR249C |  | YLR361C-A |  |  |  |  |  |  |  |
| YDR262W |  | YLR460C |  |  |  |  |  |  |  |
| YDR286C | MGP12 | YML053C |  |  |  |  |  |  |  |
| YDR307W | PMT7 | YML079W | CFF1 |  |  |  |  |  |  |
| YDR319C | YFT2 | YMR090W |  |  |  |  |  |  |  |
| YDR338C |  | YMR147W | LDO45 |  |  |  |  |  |  |
| YDR415C |  | YMR221C | FMP42 |  |  |  |  |  |  |
| YDR444W |  | YMR253C |  |  |  |  |  |  |  |
| YDR461C-A | CMI8 | YMR321C |  |  |  |  |  |  |  |
| YDR476C |  | YNL019C |  |  |  |  |  |  |  |
| YDR520C | URC2 | YNL033W |  |  |  |  |  |  |  |
| YEL023C |  | YNL115C |  |  |  |  |  |  |  |
| YEL025C |  | YNL146W |  |  |  |  |  |  |  |
| YER077C | MRX1 | YNL195C |  |  |  |  |  |  |  |
| YER079W |  | YNL300W | TOS6 |  |  |  |  |  |  |
| YER158C |  | YNR065C_066C |  |  |  |  |  |  |  |
| YER182W | FMP10 | YNR071C |  |  |  |  |  |  |  |
| YER187W |  | YOL029C |  |  |  |  |  |  |  |
| YFL034W | MIL1 | YOR012W |  |  |  |  |  |  |  |
| YFL040W |  | YOR097C |  |  |  |  |  |  |  |
| YFR006W |  | YOR186W |  |  |  |  |  |  |  |
| YFR018C |  | YPL039W |  |  |  |  |  |  |  |
| YFR032C | RRT5 | YPL067C | HTC1 |  |  |  |  |  |  |
| YFR045W | MRX20 | YPL088W |  |  |  |  |  |  |  |
| YGL081W |  | YPL245W |  |  |  |  |  |  |  |
| YGL082W |  | YPL264C |  |  |  |  |  |  |  |
| YGL085W | LCL3 | YPR027C |  |  |  |  |  |  |  |
| YGL114W |  | YPR078C |  |  |  |  |  |  |  |
| YGL138C |  | YPR196W |  |  |  |  |  |  |  |
| YGL140C |  |  |  |  |  |  |  |  |  |
| YGR016W |  |  |  |  |  |  |  |  |  |
| YGR026W |  |  |  |  |  |  |  |  |  |
| YGR052W | FMP48 |  |  |  |  |  |  |  |  |
| YGR053C | MCO32 |  |  |  |  |  |  |  |  |
| YGR067C |  |  |  |  |  |  |  |  |  |
| YGR111W |  |  |  |  |  |  |  |  |  |
| YGR117C |  |  |  |  |  |  |  |  |  |
| YGR161C | RTS3 |  |  |  |  |  |  |  |  |
| YGR210C |  |  |  |  |  |  |  |  |  |
| YGR237C |  |  |  |  |  |  |  |  |  |
| YGR273C |  |  |  |  |  |  |  |  |  |
| YHL008C |  |  |  |  |  |  |  |  |  |
| YHL012W |  |  |  |  |  |  |  |  |  |
| YHL017W |  |  |  |  |  |  |  |  |  |
| YHL018W | MCO14 |  |  |  |  |  |  |  |  |
| YHL029C | OCA5 |  |  |  |  |  |  |  |  |
| YHR022C |  |  |  |  |  |  |  |  |  |
| YHR033W |  |  |  |  |  |  |  |  |  |
| YHR078W |  |  |  |  |  |  |  |  |  |
| YHR126C | ANS1 |  |  |  |  |  |  |  |  |
| YHR131C |  |  |  |  |  |  |  |  |  |
| YHR140W |  |  |  |  |  |  |  |  |  |
| YHR159W | TDA11 |  |  |  |  |  |  |  |  |
| YIL002W-A | CMI7 |  |  |  |  |  |  |  |  |
| YIL055C |  |  |  |  |  |  |  |  |  |
| YIL067C |  |  |  |  |  |  |  |  |  |
| YIL077C | RCI37 |  |  |  |  |  |  |  |  |
| YIL102C-A |  |  |  |  |  |  |  |  |  |
| YIL127C | RRT14 |  |  |  |  |  |  |  |  |
| YIL152W | VPR1 |  |  |  |  |  |  |  |  |
| YIR016W |  |  |  |  |  |  |  |  |  |
| YJL016W | TPH3 |  |  |  |  |  |  |  |  |
| YJL070C |  |  |  |  |  |  |  |  |  |
| YJL147C | SMT1 |  |  |  |  |  |  |  |  |
| YJL160C | PIR5 |  |  |  |  |  |  |  |  |
| YJL161W | FMP33 |  |  |  |  |  |  |  |  |
| YJL163C |  |  |  |  |  |  |  |  |  |
| YJL181W | RBH1 |  |  |  |  |  |  |  |  |
| YJR030C | RBH2 |  |  |  |  |  |  |  |  |
| YJR039W | MLO127 |  |  |  |  |  |  |  |  |
| YJR061W | MNN14 |  |  |  |  |  |  |  |  |
| YJR107W | LIH1 |  |  |  |  |  |  |  |  |
| YJR112W-A |  |  |  |  |  |  |  |  |  |
| YJR115W |  |  |  |  |  |  |  |  |  |
| YJR116W | TDA4 |  |  |  |  |  |  |  |  |
| YJR124C |  |  |  |  |  |  |  |  |  |
| YJR154W |  |  |  |  |  |  |  |  |  |
| YKL018C-A | MCO12 |  |  |  |  |  |  |  |  |
| YKL063C |  |  |  |  |  |  |  |  |  |
| YKL068W-A |  |  |  |  |  |  |  |  |  |
| YKL121W | DGR2 |  |  |  |  |  |  |  |  |
| YKL133C | RCI50 |  |  |  |  |  |  |  |  |
| YKL162C |  |  |  |  |  |  |  |  |  |
| YKR005C |  |  |  |  |  |  |  |  |  |
| YKR045C |  |  |  |  |  |  |  |  |  |
| YKR070W |  |  |  |  |  |  |  |  |  |
| YKR075C |  |  |  |  |  |  |  |  |  |
| YLR001C |  |  |  |  |  |  |  |  |  |
| YLR012C |  |  |  |  |  |  |  |  |  |
| YLR031W_030W |  |  |  |  |  |  |  |  |  |
| YLR049C | MLO50 |  |  |  |  |  |  |  |  |
| YLR104W | LCL2 |  |  |  |  |  |  |  |  |
| YLR152C |  |  |  |  |  |  |  |  |  |
| YLR177W |  |  |  |  |  |  |  |  |  |
| YLR271W | CMG1 |  |  |  |  |  |  |  |  |
| YLR278C |  |  |  |  |  |  |  |  |  |
| YLR283W | PUT7 |  |  |  |  |  |  |  |  |
| YLR287C |  |  |  |  |  |  |  |  |  |
| YLR307C-A | DPA10 |  |  |  |  |  |  |  |  |
| YLR326W |  |  |  |  |  |  |  |  |  |
| YLR345W |  |  |  |  |  |  |  |  |  |
| YLR407W | GAG1 |  |  |  |  |  |  |  |  |
| YLR419W |  |  |  |  |  |  |  |  |  |
| YLR426W | TDA5 |  |  |  |  |  |  |  |  |
| YLR454W | FMP27 |  |  |  |  |  |  |  |  |
| YLR456W |  |  |  |  |  |  |  |  |  |
| YML002W_003W | VRL1 |  |  |  |  |  |  |  |  |
| YML020W |  |  |  |  |  |  |  |  |  |
| YML082W |  |  |  |  |  |  |  |  |  |
| YML096W |  |  |  |  |  |  |  |  |  |
| YML119W |  |  |  |  |  |  |  |  |  |
| YMR084W_085W |  |  |  |  |  |  |  |  |  |
| YMR118C | SHH3 |  |  |  |  |  |  |  |  |
| YMR122W-A | NCW1 |  |  |  |  |  |  |  |  |
| YMR126C | DLT1 |  |  |  |  |  |  |  |  |
| YMR130W | DPI35 |  |  |  |  |  |  |  |  |
| YMR132C | JLP2 |  |  |  |  |  |  |  |  |
| YMR144W | FDO1 |  |  |  |  |  |  |  |  |
| YMR155W |  |  |  |  |  |  |  |  |  |
| YMR181C |  |  |  |  |  |  |  |  |  |
| YMR187C |  |  |  |  |  |  |  |  |  |
| YMR196W |  |  |  |  |  |  |  |  |  |
| YMR206W |  |  |  |  |  |  |  |  |  |
| YMR209C |  |  |  |  |  |  |  |  |  |
| YMR244W |  |  |  |  |  |  |  |  |  |
| YMR252C | MLO1 |  |  |  |  |  |  |  |  |
| YMR262W |  |  |  |  |  |  |  |  |  |
| YMR265C |  |  |  |  |  |  |  |  |  |
| YMR310C | UPA2 |  |  |  |  |  |  |  |  |
| YNL011C |  |  |  |  |  |  |  |  |  |
| YNL024C | EFM6 |  |  |  |  |  |  |  |  |
| YNL050C |  |  |  |  |  |  |  |  |  |
| YNL058C |  |  |  |  |  |  |  |  |  |
| YNL095C |  |  |  |  |  |  |  |  |  |
| YNL144C |  |  |  |  |  |  |  |  |  |
| YNL162W-A |  |  |  |  |  |  |  |  |  |
| YNL165W |  |  |  |  |  |  |  |  |  |
| YNL168C | FMP41 |  |  |  |  |  |  |  |  |
| YNL181W | PBR1 |  |  |  |  |  |  |  |  |
| YNL193W |  |  |  |  |  |  |  |  |  |
| YNL211C | MRX7 |  |  |  |  |  |  |  |  |
| YNL320W |  |  |  |  |  |  |  |  |  |
| YNR014W |  |  |  |  |  |  |  |  |  |
| YNR021W |  |  |  |  |  |  |  |  |  |
| YNR068C |  |  |  |  |  |  |  |  |  |
| YOL024W |  |  |  |  |  |  |  |  |  |
| YOL036W |  |  |  |  |  |  |  |  |  |
| YOL107W |  |  |  |  |  |  |  |  |  |
| YOL131W |  |  |  |  |  |  |  |  |  |
| YOR062C |  |  |  |  |  |  |  |  |  |
| YOR093C | CMR2 |  |  |  |  |  |  |  |  |
| YOR111W |  |  |  |  |  |  |  |  |  |
| YOR238W |  |  |  |  |  |  |  |  |  |
| YOR271C | FSF1 |  |  |  |  |  |  |  |  |
| YOR289W |  |  |  |  |  |  |  |  |  |
| YOR292C |  |  |  |  |  |  |  |  |  |
| YOR296W |  |  |  |  |  |  |  |  |  |
| YOR338W |  |  |  |  |  |  |  |  |  |
| YOR365C |  |  |  |  |  |  |  |  |  |
| YPL034W |  |  |  |  |  |  |  |  |  |
| YPL041C | MRX11 |  |  |  |  |  |  |  |  |
| YPL068C |  |  |  |  |  |  |  |  |  |
| YPL071C |  |  |  |  |  |  |  |  |  |
| YPL077C |  |  |  |  |  |  |  |  |  |
| YPL107W | DPC25 |  |  |  |  |  |  |  |  |
| YPL108W |  |  |  |  |  |  |  |  |  |
| YPL150W |  |  |  |  |  |  |  |  |  |
| YPL162C |  |  |  |  |  |  |  |  |  |
| YPL168W | MRX4 |  |  |  |  |  |  |  |  |
| YPL216W |  |  |  |  |  |  |  |  |  |
| YPL229W |  |  |  |  |  |  |  |  |  |
| YPL247C |  |  |  |  |  |  |  |  |  |
| YPL272C | PBI1 |  |  |  |  |  |  |  |  |
| YPR003C |  |  |  |  |  |  |  |  |  |
| YPR063C |  |  |  |  |  |  |  |  |  |
| YPR084W |  |  |  |  |  |  |  |  |  |
| YPR089W |  |  |  |  |  |  |  |  |  |
| YPR114W |  |  |  |  |  |  |  |  |  |
| YPR145C-A |  |  |  |  |  |  |  |  |  |
| YPR157W | TDA6 |  |  |  |  |  |  |  |  |
| YPR172W |  |  |  |  |  |  |  |  |  |
| YNCF0003C | RUF20 |  |  |  |  |  |  |  |  |

Table S4 Tandem duplicate genes

| Name | Common  Name | In S288c? | Conserved/  Diverged | Core  Status | Number of copies |
| --- | --- | --- | --- | --- | --- |
| YHR053C  YHR055C | CUP1-1  CUP1-2 | Duplicated in S288c | Not diverged | One Core gene | Two+ copies 66/94 strains |
| YDR038C  YDR039C  YDR040C | ENA5  ENA1  ENA2 | Duplicated in S288c | Not diverged | One Core gene | Two+ copies 49/94 strains |
| YDR342C  YDR343C | HXT7  HXT6 | Duplicated in S288c | Not diverged  /diverged | One Core gene | one copy 6/94 strains  three copies 3/94 strains |
| YDR345C | HXT3 | Duplicated in S288c | Diverged | One Core gene | Present in 91/94 |
| YBR092C  YBR093C | PHO3  PHO5 | Duplicated in S288c | Diverged | One core/one not core | Two copies 88/94  At least one in all 94 strains |
| YHR043C  YHR044C | DOG2  DOG1 | Duplicated in S288c | Diverged | One core/one not core | Two copies 78/94  At least one in all 94 strains |
| YMR169C  YMR170C | ALD3  ALD2 | Duplicated in S288c | Diverged | One core/one not core | Two copies 87/94  At least one in all 94 strains |
| YER056C | FCY2 | Duplicated in S288c | Diverged | Core | Present in 94/94 |
| YER060W-A | FCY21 | Duplicated in S288c | Diverged | Not core | Present in 86/94 |
| YER060W | FCY22 | Duplicated in S288c | Diverged | Core | Present in 94/94 |
| YAR027W  YAR028W  YAR031W  YAR033W | UIP3  KTD1  PRM9  MST28 | Duplicated in S288c | Diverged | One core | Present zero copies 4/94  Present one to nine copies 90/94 |
| YGL051W  YGL053W | MST27  PRM8 | Duplicated in S288c | Diverged | Not core | One or more genes 26/94 |
| YHR092C  YHR094C  YHR096C | HXT4  HXT1  HXT5 | Duplicated in S288c | Diverged | Three genes Core | Present in 94/94 |
| YMR006C  YMR008C | PLB2  PLB1 | Duplicated in S288c | Diverged | Two genes Core | Present in 94/94 |
| YAR027W and YAR028W | UIP3  KTD1 |  |  | One gene Core for both | Duplicated in 2/94 YJM450, YJM193 |
| YDR508C | GNP1 |  |  | One gene Core | Duplicated in 2/94 YJM1400, YJM1479 +12 of 1011 |
| YIL162W | SUC2 |  |  | One gene Core | Duplicated in 1/94 YJM1439 subtelomeric tandem duplication |
| YLL039C | UBI4 |  |  | One gene Core | Duplicated in 19/94  NO – gene expansion – multiple lengths reference---  Yeast polyubiquitin unit regulates synaptonemal complex formation  and recombination during meiosis |
| YLR081W | GAL2 |  |  | One gene Core | Duplicated in 1/94 YJM1342 |
| YNL284C | MRPL10 and YNCN0003W |  |  | One gene Core | Duplicated in 1/94 YJM1083 |
| YOR285W | RDL1 |  |  | One gene Core | Duplicated in 1/94 YJM627 |
| YOR391C | HSP33 |  |  | One gene Core | Duplicated in 1/94 YJM1355  subtelomeric |
| YPR200C  YPR201W | ARR2  ARR3 |  |  | Two genes Core | Two genes duplicated in 1/94 in YJM271 partial duplication in S288c |

Table S5 Non-core genes in S288c located within the central regions.

| Gene | Common | Status |
| --- | --- | --- |
| YAR029W | DFP2 | Uncharacterized |
| YAR027W YAR028W YAR031W YAR033W | UIP3  KTD1  PRM9  MST2 | One core  Three non-core genes |
| YBL108C-A | PAU9 | Verified |
| YBR092C  YBR093C | PHO3  PHO5 | Verified  One non-core gene |
| YDR342C  YDR343C | HXT7  HXT6 | Verified  One non-core gene |
| YGL051W | MST27 | Verified |
| YGL053W | PRM8 | Verified |
| YHR043C  YHR044C | DOG2  DOG1 | Verified  One non-core gene |
| YIL029C | EMA17 | Uncharacterized |
| YLR155C | ASP3-1 | Verified |
| YLR157C | ASP3-2 | Verified |
| YLR158C | ASP3-3 | Verified |
| YLR160C | ASP3-4 | Verified |
| YMR169C  YMR170C | ALD3  ALD2 | Verified  One non-core gene |
| YPL257W |  | Uncharacterized |
| Q0255 | RF1 | Uncharacterized |
| Q0160 | SCEI | Verified |
| Q0050 | AI1 | Verified |
| Q0055 | AI2 | Verified |
| Q0075 | AI5_Beta | Uncharacterized |
| YNCP0005C | SNR17B |  |
| YNCE0022C | tK(CUU)E2 |  |
|  | Ty1-GAG | Verified |
|  | Ty1-POL | Verified |
|  | Ty2-GAG | Verified |
|  | Ty2-POL | Verified |
|  | Ty3-GAG | Verified |
|  | Ty3-POL | Verified |
|  | Ty4-GAG | Verified |
|  | Ty4-POL | Verified |
|  | Ty5-GAG | Verified |
|  | Ty5-POL | Verified |
| R0010W | FLP1 | Verified |
| R0020C | REP1 | Verified |
| R0040C | REP2 | Verified |
| R0030W | RAF1 | Verified |

Table S6 Transposon elements found in the 94 strains. Strains missing specific Ty elements listed as Ty count of 0. Number of Ty elements in each strain estimated by depth of sequence coverage. Depth of coverage of the LTR elements is not included. For Ty7 the chromosome is listed. “p” indicates a partial Ty7 element. These partial elements generally include the region around *OXP1*. “Telo” indicates unclear which subtelomeric region this element is in.

|  | Ty1 | Ty2 | Ty3 | Ty4 | Ty5 | Ty6 | Ty7 | Ty7  chr. | Sum |
| --- | --- | --- | --- | --- | --- | --- | --- | --- | --- |
| S288c | ~46 | ~22 | 3 | 3 | 2 | 0 |  |  | 76 |
| YJM189 | 1 | ~6 | 0 | 0 | 2 | 0 | 1 | XII | 10 |
| YJM193 | ~7 | ~12 | 4 | 2 | 1 | 0 | 1 | XIII | 27 |
| YJM195 | ~15 | ~8 | 2 | 0 | 0 | 0 |  |  | 25 |
| YJM244 | 0 | ~12 | 0 | 0 | 1 | 0 |  |  | 13 |
| YJM248 | 0 | ~30 | 0 | 0 | 2 | 0 |  |  | 32 |
| YJM270 | 0 | ~28 | 0 | 0 | 4 | 0 |  |  | 32 |
| YJM271 | ~9 | ~16 | 1 | 1 | 1 | 0 | 1 | XIII | 29 |
| YJM320 | ~19 | ~11 | 5 | 1 | 3 | 0 |  |  | 39 |
| YJM326 | ~21 | ~11 | ~7 | 3 | 1 | 0 |  |  | 43 |
| YJM428 | ~22 | ~21 | 4 | 5 | 3 | 0 |  |  | 55 |
| YJM450 | ~27 | ~13 | 4 | 2 | 1 | 0 |  |  | 47 |
| YJM451 | ~18 | ~10 | 1 | 3 | 3 | 0 | 2 | XVI  IV p | 37 |
| YJM453 | 1 | ~10 | 0 | 0 | 1 | 0 | 1 | XV | 13 |
| YJM456 | ~6 | ~9 | 5 | 5 | 2 | 0 |  |  | 27 |
| YJM470 | ~20 | ~13 | 4 | ~7 | 3 | 0 |  |  | 47 |
| YJM541 | ~22 | ~10 | ~6 | 0 | 3 | 0 | 1 | XI | 42 |
| YJM554 | ~18 | ~10 | 4 | 1 | 3 | 0 |  |  | 36 |
| YJM555 | ~17 | ~15 | ~7 | 1 | 2 | 0 | 1 | XI | 43 |
| YJM627 | ~9 | ~6 | 1 | 0 | 1 | 0 |  |  | 17 |
| YJM681 | ~12 | ~15 | ~8 | 1 | 2 | 0 |  |  | 38 |
| YJM682 | ~18 | ~12 | 5 | 1 | 1 | 0 |  |  | 37 |
| YJM683 | ~16 | ~14 | ~6 | 4 | 1 | 0 |  |  | 41 |
| YJM689 | ~14 | ~10 | 3 | 1 | 1 | 0 |  |  | 29 |
| YJM693 | ~18 | ~16 | 1 | 2 | 1 | 0 | 1 | X p | 39 |
| YJM969 | 0 | ~11 | 0 | 0 | 1 | 0 |  |  | 12 |
| YJM972 | 0 | ~8 | 0 | 0 | 2 | 0 |  |  | 10 |
| YJM975 | 0 | ~8 | 0 | 0 | 2 | 0 |  |  | 10 |
| YJM978 | 0 | ~6 | 0 | 0 | 2 | 0 |  |  | 8 |
| YJM981 | 0 | ~11 | 0 | 0 | 2 | 0 |  |  | 13 |
| YJM984 | 0 | ~8 | 0 | 0 | 2 | 0 |  |  | 10 |
| YJM987 | 0 | ~7 | 0 | 0 | 2 | 0 |  |  | 9 |
| YJM990 | 0 | ~8 | 0 | 0 | 2 | 0 |  |  | 10 |
| YJM993 | 0 | ~6 | 0 | 0 | 2 | 0 |  |  | 8 |
| YJM996 | 0 | ~13 | 0 | 0 | 2 | 0 |  |  | 15 |
| YJM1078 | 2 | ~35 | 0 | 0 | 2 | 0 |  |  | 39 |
| YJM1083 | ~19 | ~9 | 4 | 0 | 2 | 0 | 1 | IV p  XVI p  XV p | 37 |
| YJM1129 | ~9 | ~16 | 0 | 0 | 2 | 0 | 1 | XIII p | 28 |
| YJM1133 | ~10 | ~12 | 4 | 1 | 1 | 0 |  |  | 28 |
| YJM1190 | ~23 | ~9 | 2 | 0 | 2 | 0 | 1 | XI | 37 |
| YJM1199 | ~18 | ~9 | 5 | 1 | 2 | 0 |  |  | 35 |
| YJM1202 | ~22 | ~12 | 1 | 2 | 2 | 0 |  |  | 39 |
| YJM1208 | ~22 | ~8 | 1 | 3 | 2 | 0 | 1 | IV p | 37 |
| YJM1242 | 0 | ~11 | 0 | 0 | 2 | 0 |  |  | 13 |
| YJM1244 | 0 | ~15 | 0 | 0 | 2 | 0 |  |  | 17 |
| YJM1248 | ~20 | ~7 | 1 | 0 | 0 | 0 |  |  | 28 |
| YJM1250 | ~21 | ~15 | 1 | 0 | 3 | 0 | 1 | XIII | 41 |
| YJM1252 | 1 | ~31 | 0 | 0 | 2 | 0 |  |  | 34 |
| YJM1273 | 2 | ~9 | ~7 | 3 | 0 | 0 |  |  | 21 |
| YJM1304 | ~19 | ~13 | ~8 | 3 | 3 | 0 |  |  | 46 |
| YJM1307 | ~11 | ~18 | 3 | 1 | 1 | 0 |  |  | 34 |
| YJM1311 | ~21 | ~13 | 4 | 0 | 2 | 0 | 2 | VIII p  IV p | 42 |
| YJM1326 | ~18 | ~13 | 1 | 1 | 2 | 0 |  |  | 35 |
| YJM1332 | 5 | ~10 | 1 | 0 | 2 | 0 |  |  | 18 |
| YJM1336 | 1 | ~11 | 0 | 0 | 2 | 0 |  |  | 14 |
| YJM1338 | 3 | ~9 | 4 | 1 | 3 | 0 |  |  | 20 |
| YJM1341 | 1 | ~14 | 0 | 0 | 2 | 0 |  |  | 17 |
| YJM1342 | ~14 | ~18 | 0 | ~6 | 3 | 0 |  |  | 41 |
| YJM1355 | ~29 | ~18 | 1 | 1 | 4 | 0 | 3 | Telo  IV p  XI p | 56 |
| YJM1356 | 1 | ~14 | 0 | 0 | 2 | 0 |  |  | 17 |
| YJM1381 | ~17 | ~11 | 2 | 1 | 2 | 0 | 1 | IV p | 34 |
| YJM1383 | ~11 | ~15 | 4 | 0 | 1 | 0 | 1 | XI | 33 |
| YJM1385 | ~28 | ~15 | 4 | ~6 | 3 | 0 | 2 | VII  IV p | 58 |
| YJM1386 | ~15 | ~10 | 0 | 2 | 2 | 0 |  |  | 29 |
| YJM1387 | 0 | ~11 | 0 | 0 | 2 | 0 |  |  | 13 |
| YJM1388 | 4 | 2 | 1 | 3 | 0 | 0 |  |  | 10 |
| YJM1389 | ~10 | ~10 | 2 | 2 | 3 | 0 |  |  | 27 |
| YJM1399 | 0 | 1 | 1 | 0 | 0 | 0 |  |  | 2 |
| YJM1400 | ~6 | ~24 | ~6 | 3 | 2 | 0 |  |  | 41 |
| YJM1401 | ~7 | ~11 | ~7 | 4 | 1 | 0 | 1 | IV p | 31 |
| YJM1402 | 5 | ~16 | 1 | 1 | 2 | 0 |  |  | 25 |
| YJM1415 | 0 | ~6 | 0 | 0 | 2 | 0 | 1 | IV | 9 |
| YJM1417 | 0 | ~7 | 0 | 0 | 2 | 0 |  |  | 9 |
| YJM1418 | 2 | 1 | 5 | 1 | 2 | 0 |  |  | 11 |
| YJM1419 | ~20 | ~26 | 5 | ~6 | 2 | 0 |  |  | 59 |
| YJM1433 | ~6 | ~16 | 1 | 5 | 2 | 0 |  |  | 30 |
| YJM1434 | 2 | ~10 | ~7 | 2 | 0 | 0 |  |  | 21 |
| YJM1439 | ~17 | 5 | 2 | 0 | 0 | 0 |  |  | 24 |
| YJM1443 | ~6 | ~11 | ~7 | 4 | 0 | 0 |  |  | 28 |
| YJM1444 | ~19 | ~10 | ~6 | 0 | 3 | 0 |  |  | 38 |
| YJM1447 | 1 | 2 | 0 | 0 | 0 | 0 |  |  | 3 |
| YJM1450 | ~17 | ~12 | 5 | 0 | 1 | 0 |  |  | 35 |
| YJM1460 | ~16 | 3 | 2 | 5 | 2 | 0 |  |  | 28 |
| YJM1463 | ~9 | ~20 | 0 | 0 | 1 | 0 | 1 | IV p | 31 |
| YJM1477 | 0 | ~12 | 0 | 0 | 2 | 0 |  |  | 14 |
| YJM1478 | ~9 | ~22 | 3 | 0 | 0 | 0 | 1 | XI | 35 |
| YJM1479 | ~6 | ~23 | ~6 | 3 | 2 | 0 |  |  | 40 |
| YJM1526 | ~6 | ~19 | 1 | 0 | 3 | 0 | 1 | V p | 30 |
| YJM1527 | 1 | ~15 | 2 | 1 | 3 | 0 |  |  | 22 |
| YJM1549 | ~7 | ~9 | 3 | 1 | 2 | 0 | 2 | II  IV p | 24 |
| YJM1573 | 5 | ~18 | 2 | 3 | 3 | 0 |  |  | 31 |
| YJM1574 | ~6 | ~13 | 0 | 0 | 3 | 0 | 2 | XIV  XII | 24 |
| YJM1592 | ~10 | ~9 | 2 | 2 | 3 | 0 |  |  | 26 |
| YJM1615 | ~14 | ~7 | 4 | 3 | 2 | 0 | 1 | IV p | 31 |
| Totals | ~930 | ~1177 | ~220 | ~124 | ~172 | 0 | 33 |  | ~2656 |

Table S7 Locations of Ty7 elements in the 93 strains. Note that some strains contain more than one Ty7 element.

| Central | YJM189 | YJM1574 |  |  |  | chrXII | YLR378C | YLR380W |
| --- | --- | --- | --- | --- | --- | --- | --- | --- |
| Central (1) | YJM271 | YJM193 |  |  |  | chrXIII | YMR259C | YMR260C |
| Central | YJM451 |  |  |  |  | chrXVI | YNCP0024C | YPR171W |
| Central | YJM555 | YJM541 | YJM1190 | YJM1383 |  | chrXI | YKL063C | YKL062W |
| Central | YJM1478 |  |  |  |  | chrXI | YKL215C | YKL214C |
| Central (2) | YJM453 |  |  |  |  | chrXV | YOL030W |  |
| Central | YJM1574 |  |  |  |  | chrXIV | YNL038W | YNL037C |
| Central (2) | YJM1415 |  |  |  |  | chrIV | YDR263C |  |
| Subtelomeric | YJM1355 |  |  |  |  | chrIVR |  |  |
| Subtelomeric | YJM1463 | YJM1083 | YJM1311 | YJM1385 | YJM1615 | chrIVL |  |  |
| Subtelomeric | YJM1401 | YJM1208 | YJM451 | YJM1549 | YJM1381 | chrIVL |  |  |
| Subtelomeric | YJM1355 |  |  |  |  | chrXIL |  |  |
| Subtelomeric | YJM1250 | YJM1129 |  |  |  | chrXIIIL |  |  |
| Subtelomeric | YJM1385 |  |  |  |  | chrVIIL |  |  |
| Subtelomeric | YJM1083 |  |  |  |  | chrXVR |  |  |
| Subtelomeric | YJM1083 |  |  |  |  | chrXVIL |  |  |
| Subtelomeric | YJM1526 |  |  |  |  | chrVL |  |  |
| Subtelomeric | YJM1311 |  |  |  |  | chrVIIIR |  |  |

(1)Two tandem copies

(2) Ty7 inserted into gene

Table S8 Genes proposed as dubious, 227 in total. *YCL001W-A, YCL001W-B, YCR099C*, *YCR100C*, *YCR101C*, *YFL041W-A*, *YCR099C*, and *YHR054C* are proposed as dubious, but also parts of gene remnants. Those genes previously proposed as not protein coding genes(Zhang and Wang 2000) are marked with a “Z”. Genes from (Kessler et al. 2003) marked with a “K”.

|  | Name | Common |  | Name | Common |  | Name | Common |  | Name | Common |
| --- | --- | --- | --- | --- | --- | --- | --- | --- | --- | --- | --- |
|  | YAL037C-A |  |  | YDR524W-C |  | Z | YIL012W |  |  | YLR379W-A |  |
|  | YAR035C-A |  | Z | YDR525W | API2 |  | YIL014C-A |  |  | YLR406C-A |  |
| K | YBL008W-A |  | Z | YEL008W |  |  | YIL046W-A |  | K | YLR412C-A |  |
| Z | YBL044W |  |  | YEL009C-A |  | Z | YIL060W |  |  | YLR415C |  |
| Z | YBL048W | RRT1 | Z | YEL014C |  |  | YIL086C |  | Z | YML122C |  |
| Z | YBL071C |  | Z | YEL033W | MTC7 | Z | YIL089W |  | Z | YMR007W |  |
|  | YBL071C-B |  | Z | YEL067C |  |  | YIL102C |  | K | YMR030W-A |  |
|  | YBL100W-C |  | Z | YER014C-A | BUD25 |  | YIL105W-A |  | Z | YMR082C |  |
| Z | YBR027C |  |  | YER038W-A | FMP49 | K | YIL134C-A |  | Z | YMR103C |  |
|  | YBR032W |  |  | YER066W | RRT13 | Z | YIL163C |  | Z | YMR122C |  |
| K | YBR072C-A |  | Z | YER135C |  |  | YIR018C-A |  | Z | YMR141C |  |
| Z | YBR090C |  |  | YER138W-A |  | Z | YIR020C |  |  | YMR158C-A |  |
| Z | YBR144C |  | Z | YER145C-A |  |  | YIR021W-A |  |  | YMR175W-A |  |
|  | YBR182C-A |  |  | YER175W-A |  |  | YJL007C |  |  | YMR230W-A |  |
|  | YBR196C-B |  | Z | YER181C |  | Z | YJL028W |  | Z | YMR254C |  |
| K | YBR200W-A |  |  | YER186C |  |  | YJL047C-A |  | Z | YMR316C-A |  |
| Z | YBR209W |  |  | YFL010W-A | AUA1 | Z | YJL077C | ICS3 | Z | YMR320W |  |
|  | YBR219C |  | Z | YFL015C |  |  | YJL077W-A |  |  | YNL042W-B |  |
| Z | YBR292C |  | Z | YFL019C |  |  | YJL077W-B |  |  | YNL067W-B |  |
| Z | YCL001W-A |  | K | YFL041W-A |  | Z | YJL136W-A |  |  | YNL097C-B | PLS1 |
|  | YCL001W-B |  |  | YFR026C | ULI1 |  | YJR079W |  |  | YNL140C |  |
| Z | YCL042W |  |  | YFR035C |  | Z | YJR128W |  | Z | YNL143C |  |
|  | YCL054W-A | RDT1 | Z | YFR054C |  | Z | YJR146W |  |  | YNL146C-A |  |
| Z | YCL058C | FYV5 | Z | YGL006W-A |  | K | YJR151W-A |  | Z | YNL184C |  |
| Z | YCR001W |  |  | YGL007C-A |  | Z | YKL037W | AIM26 | Z | YNL269W | BSC4 |
| Z | YCR006C |  |  | YGL034C |  | Z | YKL044W | MMO1 |  | YNL277W-A |  |
| Z | YCR022C |  | K | YGL041C-B |  |  | YKL096C-B |  | Z | YOL013W-A |  |
|  | YCR024C-B |  | Z | YGL088W |  | Z | YKL097C |  | Z | YOL038C-A |  |
| Z | YCR025C |  | Z | YGL118C |  | Z | YKL102C |  |  | YOL085C |  |
| Z | YCR041W |  | Z | YGL149W | HUR1 |  | YKL104W-A |  | Z | YOL118C |  |
|  | YCR050C |  | Z | YGL168W |  | Z | YKR032W |  |  | YOR011W-A |  |
| Z | YCR085W |  | Z | YGL193C |  | Z | YKR073C |  | Z | YOR015W |  |
|  | YCR095W-A |  | Z | YGL204C |  | K | YLL006W-A |  | Z | YOR029W |  |
|  | YCR099C |  |  | YGL235W |  | Z | YLL030C | RRT7 |  | YOR034C-A |  |
|  | YCR100C | EMA35 | Z | YGR018C |  | Z | YLR111W |  | Z | YOR068C | VAM10 |
|  | YCR101C |  | Z | YGR025W | NAG1 | Z | YLR112W |  | Z | YOR072W |  |
| Z | YDL009C |  |  | YGR031C-A |  | Z | YLR122C |  |  | YOR072W-B |  |
| K | YDL022C-A |  | Z | YGR039W |  | K | YLR154C-G |  |  | YOR105W |  |
|  | YDL159W-A |  | Z | YGR045C |  | K | YLR154C-H |  |  | YOR161C-C |  |
|  | YDL242W |  |  | YGR050C |  |  | YLR154W-C | TAR1 |  | YOR192C-C |  |
|  | YDR003W-A |  | K | YGR121W-A |  | K | YLR156C-A |  | Z | YOR235W | IRC13 |
| Z | YDR010C |  |  | YGR146C-A |  | Z | YLR156W |  | Z | YOR314W |  |
| Z | YDR029W |  | Z | YGR164W |  | K | YLR157C-C |  | K | YOR316C-A |  |
| Z | YDR034C-A |  |  | YGR174W-A |  |  | YLR157W-D |  | Z | YOR343C |  |
|  | YDR034W-B |  | Z | YHL005C |  |  | YLR157W-E |  | Z | YOR376W |  |
| Z | YDR102C |  |  | YHL015W-A |  | K | YLR159C-A |  | K | YOR376W-A |  |
| Z | YDR114C |  | Z | YHL037C |  | Z | YLR159W |  | K | YPL038W-A |  |
| Z | YDR157W |  | Z | YHL041W |  | Z | YLR161W |  | Z | YPL056C | LCL1 |
| K | YDR169C-A |  |  | YHR007C-A | ECM12 | Z | YLR162W |  |  | YPL062W |  |
| K | YDR182W-A |  |  | YHR021W-A |  | K | YLR162W-A | RRT15 |  | YPL080C |  |
| K | YDR194W-A |  |  | YHR052C-B | AHT1 |  | YLR236C |  |  | YPL119C-A |  |
|  | YDR209C |  |  | YHR054C |  | Z | YLR255C |  | Z | YPL261C |  |
| Z | YDR215C |  |  | YHR054C-B |  | Z | YLR261C | VPS63 | Z | YPR014C |  |
| Z | YDR274C |  |  | YHR093W |  | Z | YLR264C-A |  | Z | YPR053C |  |
| Z | YDR278C |  | Z | YHR125W |  | Z | YLR296W |  | Z | YPR064W |  |
| Z | YDR344C |  | Z | YHR130C |  | Z | YLR302C |  | Z | YPR096C |  |
| Z | YDR366C | MOR1 | Z | YHR139C-A |  |  | YLR331C | JIP3 |  | YPR099C |  |
| Z | YDR504C | SPG3 | Z | YHR173C |  | K | YLR342W-A |  | K | YPR108W-A |  |
|  | YDR524C-B |  | K | YHR175W-A |  | Z | YLR358C |  |  | YPR159C-A |  |
|  |  |  |  | YHR180W |  | Z | YLR365W |  | Z | YPR195C |  |

Table S9 Proposed dubious genes that overlap with other genes or elements.

| Dubious  Overlap Ty | Dubious  Overlaps RNA: | RNA: | Dubious  Overlaps Prot: | Common  name | Protein it overlaps with: | Common  name |
| --- | --- | --- | --- | --- | --- | --- |
| YDR034C-A | YDR278C | YNCD0021C | YBR090C |  | YBR089C-A | NHP6B |
| YDR366C | YGL088W | YNCG0013W  SNR10 | YBR219C |  | YBR220C |  |
| YER138W-A | YGL193C | YNCG0006C  RME2 | YCL042W |  | YCL040W | GLK1 |
| YER181C | YGR164W | YNCG0038C | YCL058C | FYV5 | YCL058W-A | ADF1 |
| YIL060W | YIR020C | YNCI0012  ICR1 | YCR041W |  | YCR040W | MATALPHA1 |
| YMR158C-A | YJL007C | YNCJ0020W  EMT3 | YCR050C |  | YCR048W | ARE1 |
| YOR192C-C | YLR154C-G | rDNA | YDL009C |  | YDL010W | GRX6 |
|  | YLR154C-H | rDNA | YDL022C-A |  | YDL024C | DIA3 |
|  | YLR154W-C | rDNA | YDR114C |  | YDR115W | MRX14 |
|  | YLR156C-A | rDNA | YDR157W |  | YDR156W | RPA14 |
|  | YLR157C-C | rDNA | YDR209C |  | YDR210W |  |
|  | YLR159C-A | rDNA | YEL067C |  | YEL066W | HPA3 |
|  | YLR162W | rDNA | YER014C-A | BUD25 | YER014W | HEM14 |
|  | YLR162W-A | rDNA | YER038W-A |  | YER039C | HVG1 |
|  | YNL042W-B | YYNCN0009W | YER145C-A |  | YER146W | LSM5 |
|  | YPR108W-A | YNCP0016W | YFL010W-A | AUA1 | YFL010C | WWM1 |
|  | YPR159C-A | YNCP0023W | YGL041C-B |  | YGL041W-A | DPC13 |
|  |  |  | YGL168W | HUR1 | YGL167C | PMR1 |
|  |  |  | YGL235W |  | YGL236C | MTO1 |
|  |  |  | YGR018C |  | YGR017W |  |
|  |  |  | YGR031C-A | NAG1 | YGR031W | IMO32 |
|  |  |  | YHR021W-A | ECM12 | YHR022C |  |
|  |  |  | YIL105W-A |  | YIL105C | SLM1 |
|  |  |  | YJL077W-A |  | YJL077C | ICS3 |
|  |  |  | YJR079W |  | YJR080C | AIM24 |
|  |  |  | YJR128W |  | YJR127C | RSF2 |
|  |  |  | YJR146W |  | YJR147W | HMS2 |
|  |  |  | YLR261C | VPS63 | YLR261C | VPS63 |
|  |  |  | YLR331C | JIP3 | YLR331C | JIP3 |
|  |  |  | YLR358C |  | YLR357W | RSC2 |
|  |  |  | YMR316C-A |  | YMR316W | DIA1 |
|  |  |  | YNL140C |  | YNL139C | THO2 |
|  |  |  | YNL184C |  | YNL185C | MRPL19 |
|  |  |  | YOR068C | VAM10 | YOR069W | VPS5 |
|  |  |  | YOR105W |  | YOR104W | PIN2 |
|  |  |  | YPR053C |  | YPR052C | NHP6A |

Table S10 Seven pairs of protein coding genes that overlap each other in S288c.

| Protein Coding Gene Overlap | | | | | | | | | | 3' end | | *A. gossypii* |
| --- | --- | --- | --- | --- | --- | --- | --- | --- | --- | --- | --- | --- |
| Gene 1 | | Chr. | Start | Stop | Gene 2 | | Stop | Start | Overlap | | overlap | |
| YGR074W | SMD1 | VII | 635712 | 636152 | YGR075C | PRP38 | 636147 | 636875 | 6 bp | | Not adjacent | |
| YLR315W | NKP2 | IX | 764808 | 765269 | YLR316C | TAD3 | 765266 | 766358 | 4 bp | | (1) | |
| YLR360W | VPS38 | XII | 846103 | 847422 | YLR361C | DCR2 | 847388 | 849124 | 35 bp | | 14 bp | |
| YML096W |  | XIII | 79909 | 81486 | YML095C | RAD10 | 81481 | 82113 | 6 bp | | Not adjacent | |
| YNL246W | VPS75 | XIV | 185460 | 186349 | YNL245C | CWC25 | 186346 | 186885 | 4 bp | | No | |
| YPL018W | CTF19 | XVI | 517651 | 518760 | YPL017C | IRC15 | 518734 | 520233 | 27 bp | | No | |
| YPL166W | ATG29 | XVI | 237338 | 237979 | YPL165C | SET6 | 237956 | 239077 | 24 bp | | 24 bp | |

(1) YLR315W NKP1 has no homolog in *A. gossypii*

Table S11 Genes containing potential null alleles in one or more of the 94 strains.

| count | gene | common | 2nd name | common | type of disruption | description | strains |
| --- | --- | --- | --- | --- | --- | --- | --- |
| 1 | YAL002W | VPS8 |  |  | stop |  | YJM1592 |
| 2 | YAL062W | GDH3 |  |  | stop |  | YJM1419 |
| 3 | YAR020C | PAU7 |  |  | stop |  | S288c + additional |
| 4 | YAR031W | PRM9 |  |  | stop |  | YJM1592 + additional |
| 5 | YBL018C | POP8 |  |  | frame | plus 1 | YJM1399 |
| 6 | YBL106C | SRO77 |  |  | stop |  | YJM1129 |
| 6 | YBL106C | SRO77 |  |  | frame | polyA plus 2 | YJM1307 + additional |
| 6 | YBL106C | SRO77 |  |  | two frameshifts | polyT plus 1, polyA plus 1 | YJM1463 |
| 7 | YBR001C | NTH2 |  |  | stop |  | YJM689 |
| 8 | YBR013C |  | YBR012C |  | frameshift |  | YJM1381 + additional |
| 8 | YBR013C |  | YBR012C |  | insertion | Ty1 insertion | S288c |
| 9 | YBR033W | EDS1 |  |  | frame | 45 base deletion | YJM1356 + additional |
| 9 | YBR033W | EDS1 |  |  | frame | minus 1 | YJM1386 |
| 9 | YBR033W | EDS1 |  |  | frame | polyA plus 1 | YJM248 + additional |
| 10 | YBR063C | CNM1 |  |  | frame | polyA plus 1 | YJM1326 + additional |
| 10 | YBR063C | CNM1 |  |  | frame | minus 1 | YJM1574 + additional |
| 10 | YBR063C | CNM1 |  |  | multiple frameshifts | multiple frameshifts | YJM248 + additional |
| 10 | YBR063C | CNM1 |  |  | frame | polyA plus 5 | YJM470 |
| 11 | YBR069C | TAT1 |  |  | frame | polyA minus 1 | YJM1526 |
| 12 | YBR096W |  |  |  | stop |  | YJM1460 |
| 13 | YBR117C | TKL2 |  |  | stop |  | YJM1615 |
| 14 | YBR138C |  |  |  | stop |  | YJM1460 |
| 15 | YBR147W | RTC2 |  |  | frame | polyA plus 1 | YJM1083 |
| 16 | YBR152W | SPP381 |  |  | stop |  | YJM1202 + additional |
| 17 | YBR161W | CSH1 |  |  | frame | polyC plus 1 | YJM1332 |
| 17 | YBR161W | CSH1 |  |  | stop |  | YJM428 + additional |
| 18 | YBR184W |  |  |  | frame | polyA minus 1 | YJM627 + additional |
| 19 | YBR187W | GDT1 |  |  | frame | highly diverged | YJM1342 + additional |
| 20 | YBR239C | ERT1 |  |  | frame | minus 1 | YJM681 |
| 21 | YBR244W | GPX2 |  |  | stop |  | YJM270 |
| 22 | YBR284W |  |  |  | stop |  | YJM1592 + additional |
| 23 | YBR293W | VBA2 |  |  | stop |  | YJM1433 |
| 23 | YBR293W | VBA2 |  |  | frame | polyA minus 1 | YJM451 |
| 24 | YBR294W | SUL1 |  |  | frame | polyA plus 1 | YJM451 |
| 24 | YBR294W | SUL1 |  |  | stop |  | YJM453 + additional |
| 25 | YBR295W | PCA1 |  |  | frame | 29 base deletion | YJM1450 + additional |
| 26 | YBR296C | PHO89 |  |  | frame | plus 1 | YJM1549 |
| 26 | YBR296C | PHO89 |  |  | stop |  | YJM1549 |
| 27 | YBR297W | MAL33 |  |  | stop |  | YJM271 + additional |
| 28 | YCL033C | MXR2 |  |  | stop |  | YJM1248 + additional |
| 29 | YCL049C |  |  |  | frame | polyA plus 1 | YJM1244 + additional |
| 29 | YCL049C |  |  |  | stop |  | YJM693 + additional |
| 30 | YCL073C | GEX1 |  |  | two stops |  | YJM1356 + additional |
| 30 | YCL073C | GEX1 |  |  | frame | polyT minus 1 | YJM627 + additional |
| 30 | YCL073C | GEX1 |  |  | insertion | Ty1 insertion | YJM993 + additional |
| 31 | YCR018C | SRD1 |  |  | frame | polyA plus 1 | YJM1399 + additional |
| 31 | YCR018C | SRD1 |  |  | two frameshifts | polyA plus 1, plus 1 | YJM1592 + additional |
| 32 | YCR059C | YIH1 |  |  | stop |  | YJM326 + additional |
| 33 | YCR075C | ERS1 |  |  | stop |  | YJM1592 + additional |
| 34 | YCR098C | GIT1 |  |  | frame | 25bp deletion | YJM270 + additional |
| 34 | YCR098C | GIT1 |  |  | insertion | Ty4 insertion | YJM993 |
| 35 | YCR099C |  | YCR100C | EMA35 | frame | polyT plus 1 | S288c + additional |
| 35 | YCR099C |  | YCR100C | EMA35 | frame | polyT plus 1 | YJM1307 + additional |
| 35 | YCR099C |  | YCR100C | EMA35 | stop |  | YJM1401 |
| 36 | YCR101C |  |  |  | stop |  | YJM1419 + additional |
|  | YDL037C | BSC1 | YDL039C | PRM7 | stop |  | S288c + additional |
| 37 | YDL109C |  |  |  | insertion | Ty1 insertion | YJM1381 |
| 37 | YDL109C |  |  |  | two frameshifts | polyT plus 1,  polyA plus 1 | YJM1444 |
| 38 | YDL112W | TRM3 |  |  | frame | minus 1 | YJM1307 |
| 39 | YDL122W | UBP1 |  |  | stop |  | YJM1447 |
| 40 | YDL124W |  |  |  | frame | polyC plus 1 | YJM1419 |
| 41 | YDL138W | RGT2 |  |  | stop |  | YJM1399 |
| 42 | YDL170W | UGA3 |  |  | frame | polyA plus 1 | YJM683 + additional |
| 43 | YDL197C | ASF2 |  |  | stop |  | YJM1083 + additional |
| 44 | YDL227C | HO |  |  | frame | polyT plus 1 | YJM1083 |
| 45 | YDL231C | BRE4 |  |  | stop |  | YJM248 + additional |
| 46 | YDL233W | MFG1 |  |  | stop |  | YJM1477 + additional |
| 47 | YDL240W | LRG1 |  |  | stop |  | YJM1549 |
| 48 | YDL241W |  |  |  | stop |  | YJM1248 + additional |
| 49 | YDL243C | AAD4 |  |  | stop |  | YJM1383 + additional |
| 50 | YDR009W | GAL3 |  |  | stop |  | YJM627 + additional |
| 51 | YDR030C | RAD28 |  |  | stop |  | YJM453 |
| 52 | YDR055W | PST1 |  |  | frame | minus 1 | YJM1342 |
| 52 | YDR055W | PST1 |  |  | frame | minus 1 | YJM248 + additional |
| 53 | YDR066C | RTR2 |  |  | stop |  | YJM193 + additional |
| 54 | YDR093W | DNF2 |  |  | two frameshifts | polyT plus 1, polyA plus 1 | YJM689 |
| 55 | YDR098C | GRX3 |  |  | stop |  | YJM1199 + additional |
| 55 | YDR098C | GRX3 |  |  | insertion | Ty2 insertion | YJM470 |
| 56 | YDR107C | TMN2 |  |  | stop |  | YJM1415 |
| 57 | YDR109C |  |  |  | frame | polyA plus 1 | YJM1252 + additional |
| 58 | YDR111C | ALT2 |  |  | G->D |  | S288c |
| 58 | YDR111C | ALT2 |  |  | stop |  | YJM451 |
| 59 | YDR121W | DPB4 |  |  | stop |  | YJM270 |
| 60 | YDR125C | ECM18 |  |  | frame | 88 base deletion | YJM1304 + additional |
| 60 | YDR125C | ECM18 |  |  | frame | plus 1 | YJM1342 + additional |
| 61 | YDR144C | MKC7 |  |  | stop |  | YJM1208 + additional |
| 62 | YDR150W | NUM1 |  |  | stop |  | YJM1478 |
| 63 | YDR168W | CDC37 |  |  | frame | minus 1 | YJM1244 |
| 64 | YDR181C | SAS4 |  |  | stop |  | YJM693 + additional |
| 65 | YDR198C | RKM2 |  |  | stop |  | YJM1307 |
| 66 | YDR218C | SPR28 |  |  | frame | 11 base deletion | YJM1307 |
| 66 | YDR218C | SPR28 |  |  | frame | polyA plus 1 | YJM1383 + additional |
| 67 | YDR223W | CRF1 |  |  | stop |  | YJM1190 + additional |
| 67 | YDR223W | CRF1 |  |  | frame and stop | polyA plus 1 | YJM1381 + additional |
| 67 | YDR223W | CRF1 |  |  | frame | polyA minus 1 | YJM451 + additional |
| 68 | YDR229W | IVY1 |  |  | stop |  | YJM627 |
| 69 | YDR247W | VHS1 |  |  | stop |  | YJM270 |
| 70 | YDR258C | HSP78 |  |  | stop |  | YJM1383 |
| 71 | YDR261C | EXG2 |  |  | frame | minus 2 | YJM248 + additional |
| 72 | YDR325W | YCG1 |  |  | frame | plus 5 tandem duplication | YJM693 |
| 73 | YDR330W | UBX5 |  |  | stop |  | YJM555 |
| 74 | YDR338C |  |  |  | frame | polyT plus 1 | YJM1133 |
| 75 | YDR387C | CIN10 |  |  | frame | polyA plus 1 | YJM1388 |
| 76 | YDR441C | APT2 |  |  | frame | polyT plus 1 | YJM1190 |
| 77 | YDR446W | ECM11 |  |  | stop |  | YJM1526 + additional |
| 77 | YDR446W | ECM11 |  |  | insertion | Ty2 insertion | YJM681 |
| 78 | YDR453C | TSA2 |  |  | frame | 5 base deletion | YJM1381 |
| 79 | YDR490C | PKH1 |  |  | frame | polyA plus 1 | YJM1083 + additional |
| 79 | YDR490C | PKH1 |  |  | stop |  | YJM270 |
| 80 | YDR501W | PLM2 |  |  | stop |  | YJM428 |
| 81 | YDR522C | SPS2 |  |  | frame | plus 1 | YJM1252 + additional |
| 81 | YDR522C | SPS2 |  |  | frame | polyA minus 1 | YJM470 + additional |
| 81 | YDR522C | SPS2 |  |  | frame | minus 1 | YJM681 + additional |
| 82 | YDR528W | HLR1 |  |  | stop |  | YJM1460 |
| 83 | YDR534C | FIT1 |  |  | stop |  | YJM1242 + additional |
| 84 | YDR538W | PAD1 |  |  | stop |  | YJM1248 + additional |
| 85 | YDR539W | FDC1 |  |  | frame | minus 1 | YJM1439 + additional |
| 85 | YDR539W | FDC1 |  |  | stop |  | YJM1460 |
| 86 | YDR540C | IRC4 |  |  | frame | polyA plus 1 | YJM1478 + additional |
| 86 | YDR540C | IRC4 |  |  | frame | polyA minus 1 | YJM1573 + additional |
| 87 | YDR541C |  |  |  | frame | polyA minus 1 | YJM248 + additional |
| 88 | YEL025C |  |  |  | stop |  | YJM1385 |
| 89 | YEL030W | ECM10 |  |  | stop |  | YJM1399 |
| 90 | YEL035C | UTR5 |  |  | stop |  | YJM1399 |
| 91 | YEL040W | UTR2 |  |  | stop |  | YJM984 + additional |
| 92 | YEL069C | HXT13 |  |  | frame | polyG minus 1 | YJM1388 + additional |
| 92 | YEL069C | HXT13 |  |  | stop |  | YJM1460 + additional |
| 92 | YEL069C | HXT13 |  |  | frame | 16 base insertion | YJM1478 |
| 92 | YEL069C | HXT13 |  |  | frame | plus 1 | YJM248 + additional |
| 92 | YEL069C | HXT13 |  |  | frame and stop | polyT minus 1 | YJM271 + additional |
| 92 | YEL069C | HXT13 |  |  | N terminal deletion |  | YJM326 |
| 93 | YEL070W | DSF1 |  |  | stop |  | YJM1242 + additional |
| 93 | YEL070W | DSF1 |  |  | frame and stop | polyT minus 1 | YJM1356 + additional |
| 93 | YEL070W | DSF1 |  |  | two stops |  | YJM1388 + additional |
| 93 | YEL070W | DSF1 |  |  | N terminal deletion |  | YJM326 |
| 93 | YEL070W | DSF1 |  |  | insertion | Ty1 insertion | YJM693 |
| 94 | YER001W | MNN1 |  |  | stop |  | YJM248 + additional |
| 95 | YER028C | MIG3 |  |  | stop |  | YJM1592 + additional |
| 96 | YER039C | HVG1 | YER039C-A |  | stop - at residue 73 |  | S288c + additional |
| 96 | YER039C | HVG1 | YER039C-A |  | stop - at residue 151 |  | YJM1202 + additional |
| 96 | YER039C | HVG1 | YER039C-A |  | stop - at residue 180 |  | YJM1248 + additional |
| 96 | YER039C | HVG1 | YER039C-A |  | two stops 73 and 151 |  | YJM555 + additional |
| 97 | YER075C | PTP3 |  |  | stop |  | YJM1433 |
| 98 | YER076C |  |  |  | frame | polyT minus 1 | YJM1332 + additional |
| 98 | YER076C |  |  |  | stop |  | YJM1401 |
| 98 | YER076C |  |  |  | frame | minus 2 | YJM428 |
| 99 | YER098W | UBP9 |  |  | stop |  | YJM453 |
| 100 | YER101C | AST2 |  |  | frame | minus 1 | YJM1304 + additional |
| 101 | YER109C | FLO8 |  |  | stop |  | S288c |
| 102 | YER129W | SAK1 |  |  | stop |  | YJM470 |
| 103 | YER132C | PMD1 |  |  | stop |  | YJM1388 + additional |
| 104 | YER141W | COX15 |  |  | stop |  | YJM1615 |
| 105 | YER184C | TOG1 |  |  | stop |  | YJM271 + additional |
| 106 | YFL027C | GYP8 |  |  | frame | polyA plus 1 | YJM1478 |
| 107 | YFL033C | RIM15 |  |  | frame | minus 1 | YJM1383 |
| 108 | YFL041W-A |  |  |  | stop |  | YJM1381 + additional |
| 109 | YFL050C | ALR2 |  |  | stop |  | YJM1463 |
| 110 | YFL051C |  |  |  | C terminal deletion |  | S288c |
| 111 | YFR018C |  |  |  | frame | polyC minus 1 | YJM1311 |
| 112 | YFR022W | ROG3 |  |  | stop |  | YJM1615 |
| 113 | YFR025C | HIS2 |  |  | stop |  | YJM1326 |
| 114 | YFR035C |  |  |  | stop |  | YJM1463 |
| 115 | YFR055W | IRC7 |  |  | stop |  | YJM996 + additional |
| 116 | YGL004C | RPN14 |  |  | two frameshifts | polyT plus 1, polyA plus 1 | YJM1386 |
| 117 | YGL021W | ALK1 |  |  | frame | polyG plus 1 | YJM1386 |
| 117 | YGL021W | ALK1 |  |  | frame | minus 1 | YJM1418 + additional |
| 118 | YGL071W | AFT1 |  |  | stop near C-terminal |  | YJM1386 |
| 118 | YGL071W | AFT1 |  |  | stop |  | YJM270 |
| 119 | YGL089C | MF(ALPHA)2 |  |  | stop |  | YJM689 |
| 120 | YGL096W | TOS8 |  |  | Two frameshifts | poly(TA)TA minus 2, poly A plus 1 | YJM1443 + additional |
| 120 | YGL096W | TOS8 |  |  | C terminal deletion |  | YJM451 |
| 120 | YGL096W | TOS8 |  |  | frame | minus 5 | YJM555 |
| 121 | YGL117W |  |  |  | insertion | Ty1 insertion | YJM1199 + additional |
| 122 | YGL134W | PCL10 |  |  | frame | polyA minus 1 | YJM1592 + additional |
| 122 | YGL134W | PCL10 |  |  | stop |  | YJM996 + additional |
| 123 | YGL157W | ARI1 |  |  | stop |  | YJM555 + additional |
| 124 | YGL158W | RCK1 |  |  | stop |  | YJM1463 |
| 124 | YGL158W | RCK1 |  |  | frame | polyA plus 1 | YJM470 |
| 124 | YGL158W | RCK1 |  |  | frame | minus 1 | YJM683 + additional |
| 125 | YGL179C | TOS3 |  |  | frame | minus 1 | YJM1418 |
| 126 | YGL196W | DSD1 |  |  | frame | polyT minus 1 | YJM1078 |
| 127 | YGL202W | ARO8 |  |  | stop |  | YJM270 |
| 128 | YGL208W | SIP2 |  |  | stop |  | YJM1250 |
| 128 | YGL208W | SIP2 |  |  | frame | polyA plus 1 | YJM248 + additional |
| 129 | YGL220W | BOL2 |  |  | stop |  | YJM555 + additional |
| 130 | YGR035W-A |  |  |  | frame | polyT minus 1 | YJM1419 |
| 130 | YGR035W-A |  |  |  | frame | polyT minus 2 | YJM1447 |
| 130 | YGR035W-A |  |  |  | frame | polyA minus 1 | YJM683 + additional |
| 131 | YGR044C | RME1 |  |  | frame | 22 base tandem duplication | YJM1252 |
| 131 | YGR044C | RME1 |  |  | stop |  | YJM1342 |
| 132 | YGR052W | FMP48 |  |  | stop |  | YJM1401 + additional |
| 133 | YGR066C | GID10 |  |  | frame | polyT plus 1, polyA plus 1 | YJM1574 + additional |
| 134 | YGR067C |  |  |  | frame | minus 1 | YJM1381 + additional |
| 135 | YGR087C | PDC6 |  |  | stop |  | YJM1463 |
| 135 | YGR087C | PDC6 |  |  | frame | polyC minus 1 | YJM195 |
| 136 | YGR100W | MDR1 |  |  | stop |  | YJM627 |
| 137 | YGR124W | ASN2 |  |  | frame | 10 base tandem duplication | YJM682 + additional |
| 138 | YGR136W | LSB1 |  |  | stop |  | YJM972 + additional |
| 139 | YGR138C | TPO2 |  |  | insertion | Ty2 insertion | YJM1574 |
| 140 | YGR154C | GTO1 |  |  | frame | minus 1 | YJM1338 + additional |
| 141 | YGR184C | UBR1 |  |  | two frameshifts | polyT plus 1, polyA plus 1 | YJM689 + additional |
| 142 | YGR197C | SNG1 |  |  | frame | polyT plus 1 | YJM1083 + additional |
| 142 | YGR197C | SNG1 |  |  | frame | minus 1 | YJM1341 |
| 142 | YGR197C | SNG1 |  |  | frame | polyT plus 1 | YJM451 |
| 142 | YGR197C | SNG1 |  |  | frame | polyG minus 1 | YJM453 |
| 143 | YGR212W | SLI1 |  |  | stop |  | YJM1447 |
| 144 | YGR213C | RTA1 |  |  | stop |  | YJM627 + additional |
| 145 | YGR224W | AZR1 |  |  | two frameshifts | polyT plus 1, polyA plus 1 | YJM1463 |
| 145 | YGR224W | AZR1 |  |  | frame | polyT minus 1 | YJM996 + additional |
| 146 | YGR235C | MIC26 |  |  | stop |  | YJM693 |
| 147 | YGR238C | KEL2 |  |  | frame | polyA minus 1 | YJM1385 |
| 147 | YGR238C | KEL2 |  |  | frame | polyT minus 1 | YJM270 + additional |
| 147 | YGR238C | KEL2 |  |  | frame | polyC minus 1 | YJM271 + additional |
| 147 | YGR238C | KEL2 |  |  | stop |  | YJM689 |
| 148 | YGR248W | SOL4 |  |  | frame | polyA plus 1 | YJM1419 |
| 149 | YGR249W | MGA1 |  |  | stop |  | YJM1388 + additional |
| 150 | YGR256W | GND2 |  |  | stop |  | YJM1460 |
| 151 | YGR271W | SLH1 |  |  | stop |  | YJM1083 |
| 152 | YGR287C | IMA1 |  |  | stop |  | YJM1439 + additional |
| 152 | YGR287C | IMA1 |  |  | N terminal deletion |  | YJM993 + additional |
| 153 | YHL012W |  |  |  | frame | polyA plus 1 | YJM1342 |
| 153 | YHL012W |  |  |  | stop |  | YJM541 + additional |
| 154 | YHL016C | DUR3 |  |  | stop |  | YJM1478 |
| 155 | YHL018W | MCO14 |  |  | frame | polyG plus 1 | YJM1336 |
| 156 | YHL026C |  |  |  | frame | polyC plus 1 | YJM270 |
| 157 | YHL035C | VMR1 |  |  | frame | polyC plus 1 | YJM453 |
| 157 | YHL035C | VMR1 |  |  | stop |  | YJM681 + additional |
| 157 | YHL035C | VMR1 |  |  | frame | polyT plus 1 | YJM693 |
| 158 | YHL039W | EFM1 |  |  | frame | polyA minus 1 | YJM682 |
| 159 | YHL043W | ECM34 |  |  | stop near C-terminal |  | most strains |
| 159 | YHL043W | ECM34 |  |  | frame | polyT plus 1 | YJM1399 |
| 159 | YHL043W | ECM34 |  |  | stop |  | YJM1401 |
| 160 | YHL044W | DFP4 |  |  | stop |  | YJM1399 + additional |
| 160 | YHL044W | DFP4 |  |  | frame | polyC plus 1 | YJM195 |
| 160 | YHL044W | DFP4 |  |  | N terminal deletion |  | YJM554 |
| 161 | YHR022C |  |  |  | frame and stop | polyT plus 1 | YJM1242 + additional |
| 161 | YHR022C |  |  |  | frame | polyT plus 1 | YJM1250 + additional |
| 161 | YHR022C |  |  |  | frame and insertion | polyT plus 1, Ty2 insertion | YJM1419 + additional |
| 162 | YHR022C-A |  |  |  | stop |  | YJM1418 |
| 163 | YHR029C | YHI9 |  |  | frame | polyT minus 1 | YJM1250 |
| 164 | YHR033W |  |  |  | frame | 271 base deletion | YJM1450 + additional |
| 164 | YHR033W |  |  |  | stop |  | YJM681 + additional |
| 165 | YHR035W | NEL1 |  |  | frame | polyT plus 1 | YJM270 + additional |
| 165 | YHR035W | NEL1 |  |  | frame | minus 2 | YJM453 |
| 166 | YHR048W | YHK8 |  |  | stop |  | YJM1477 + additional |
| 167 | YHR084W | STE12 |  |  | stop |  | YJM1479 + additional |
| 168 | YHR104W | GRE3 |  |  | stop |  | YJM195 |
| 169 | YHR132C | ECM14 |  |  | stop |  | YJM1478 |
| 170 | YHR133C | NSG1 |  |  | stop |  | YJM270 |
| 171 | YHR136C | SPL2 |  |  | frame | 2 base deletion | YJM1078 + additional |
| 172 | YHR177W | ROF1 |  |  | frame | minus 4 | YJM1202 + additional |
| 173 | YHR210C |  |  |  | frame | polyG plus 1 | YJM1311 + additional |
| 173 | YHR210C |  |  |  | frame | 2 base deletion | YJM1592 + additional |
| 173 | YHR210C |  |  |  | frame | plus 2 | YJM189 + additional |
| 173 | YHR210C |  |  |  | two frameshifts | polyA plus 1, poly G plus 1 | YJM470 + additional |
| 174 | YIL009W | FAA3 |  |  | frame | 35 base tandem duplication | YJM1199 |
| 174 | YIL009W | FAA3 |  |  | frame | minus 1 | YJM554 + additional |
| 175 | YIL013C | PDR11 |  |  | frame | polyA minus 1 | YJM1355 |
| 176 | YIL077C | RCI37 |  |  | stop |  | YJM1447 |
| 177 | YIL092W |  |  |  | frame | polyA plus 1 | YJM1463 |
| 177 | YIL092W |  |  |  | frame | 28 base deletion | YJM1526 |
| 178 | YIL095W | PRK1 |  |  | insertion | Ty1 insertion | YJM1199 + additional |
| 179 | YIL096C | BMT5 |  |  | frame | polyA plus 1 | YJM1336 |
| 180 | YIL099W | SGA1 |  |  | stop |  | YJM1478 |
| 181 | YIL102C |  |  |  | frame | polyA plus 1 | YJM1083 |
| 182 | YIL114C | POR2 |  |  | frame | polyG plus 1 | YJM270 |
| 183 | YIL120W | QDR1 |  |  | stop |  | YJM1083 |
| 183 | YIL120W | QDR1 |  |  | frame | polyT plus 1 | YJM1478 |
| 184 | YIL131C | FKH1 |  |  | stop |  | YJM1311 |
| 185 | YIL145C | PAN6 |  |  | frame | 11 base insertion | YJM1208 |
| 186 | YIL149C | MLP2 |  |  | stop |  | YJM1479 |
| 187 | YIL162W | SUC2 |  |  | frame | polyA plus 2 | YJM1078 + additional |
| 187 | YIL162W | SUC2 |  |  | frame | minus 1 | YJM981 + additional |
| 188 | YIL164C | NIT1 |  |  | stop |  | YJM1383 + additional |
| 188 | YIL164C | NIT1 |  |  | frame | polyA plus 1 | YJM189 + additional |
| 189 | YIL165C |  |  |  | stop |  | YJM1383 + additional |
| 190 | YIL166C | SOA1 |  |  | frame | polyT plus 1 | YJM683 + additional |
| 191 | YIL167W | SDL1 | YIL168W | SDL1 | stop |  | S288c + additional |
| 191 | YIL167W | SDL1 | YIL168W | SDL1 | frame | polyA minus 1 | YJM1248 + additional |
| 191 | YIL167W | SDL1 | YIL168W | SDL1 | frame | minus 1 | YJM1447 |
| 192 | YIL169C | CSS1 |  |  | stop |  | S288c |
| 193 | YIR003W | AIM21 |  |  | stop |  | YJM193 |
| 194 | YIR019C | FLO11 |  |  | frame | polyG minus 1 | YJM1250 |
| 195 | YIR024C | INA22 |  |  | frame | polyA minus 2 | YJM1573 + additional |
| 196 | YIR028W | DAL4 |  |  | stop |  | YJM1415 + additional |
| 196 | YIR028W | DAL4 |  |  | frame | minus 1 | YJM627 + additional |
| 197 | YIR039C | YPS6 |  |  | insertion | Ty1 insertion | YJM1273 + additional |
| 197 | YIR039C | YPS6 |  |  | stop |  | YJM1418 |
| 197 | YIR039C | YPS6 |  |  | frame | polyG plus 1 | YJM1444 |
| 198 | YJL027C |  |  |  | stop |  | YJM1399 |
| 199 | YJL045W | SDH9 |  |  | frame | polyG minus 2 | YJM689 + additional |
| 200 | YJL073W | JEM1 |  |  | stop |  | YJM271 |
| 201 | YJL084C | ALY2 |  |  | frame | 19 base deletion | YJM1450 |
| 202 | YJL089W | SIP4 |  |  | frame | polyA plus 1 | YJM271 |
| 203 | YJL100W | LSB6 |  |  | stop |  | YJM1133 |
| 204 | YJL103C | GSM1 |  |  | stop |  | YJM1478 |
| 205 | YJL107C |  | YJL108C | PRM10 | frame | polyT minus 1 | S288c |
| 206 | YJL129C | TRK1 |  |  | frame | polyA minus 1 | YJM1388 |
| 207 | YJL137C | GLG2 |  |  | frame | polyT minus 2 | YJM1250 |
| 207 | YJL137C | GLG2 |  |  | frame | polyT plus 1 | YJM1307 |
| 207 | YJL137C | GLG2 |  |  | stop |  | YJM1386 |
| 207 | YJL137C | GLG2 |  |  | frame | polyA plus 1 | YJM555 |
| 208 | YJL144W | ROQ1 |  |  | frame | polyA plus 1 | YJM555 + additional |
| 209 | YJL160C | PIR5 |  |  | stop |  | YJM1549 + additional |
| 210 | YJL165C | HAL5 |  |  | frame | polyA plus 1 | YJM1355 |
| 211 | YJL181W | RBH1 |  |  | frame | 11 base deletion | YJM470 |
| 212 | YJL207C | LAA1 |  |  | frame | polyA minus 1 | YJM1250 |
| 213 | YJL213W |  |  |  | C terminal deletion |  | YJM1250 |
| 213 | YJL213W |  |  |  | frame | polyT plus 1 | YJM1307 |
| 214 | YJR010C-A | SPC1 |  |  | stop |  | YJM681 |
| 215 | YJR010W | MET3 |  |  | stop |  | YJM1355 |
| 216 | YJR011C | CAL4 |  |  | frame | polyA minus 1 | YJM1307 |
| 217 | YJR012C |  |  |  | frame | poly(AT) minus 2 | YJM1342 |
| 217 | YJR012C |  |  |  | frame | polyT minus 1 | YJM450 + additional |
| 218 | YJR015W |  |  |  | two stops |  | YJM1311 |
| 218 | YJR015W |  |  |  | stop |  | YJM1415 + additional |
| 218 | YJR015W |  |  |  | frame | add in details |  |
| 219 | YJR030C | RBH2 |  |  | insertion | Ty1 insertion | S288c |
| 220 | YJR031C | GEA1 |  |  | frame and stop | polyC minus 1 | YJM1417 |
| 221 | YJR050W | ISY1 |  |  | frame | polyA minus 1 | YJM1419 |
| 222 | YJR052W | RAD7 |  |  | frame | polyA minus 1 | YJM1332 |
| 223 | YJR062C | NTA1 |  |  | frame | polyG plus 1 | YJM1307 |
| 224 | YJR096W |  |  |  | frame | minus 1 | YJM456 |
| 225 | YJR098C |  |  |  | stop |  | YJM1356 |
| 226 | YJR120W | DMO1 |  |  | frame | minus 1 | YJM1401 |
| 227 | YJR127C | RSF2 |  |  | frame | polyT minus 1 | YJM1434 + additional |
| 227 | YJR127C | RSF2 |  |  | frame | polyA plus 1 | YJM1463 |
| 227 | YJR127C | RSF2 |  |  | three frameshifts | minus 5, minus 1, minus 1 | YJM248 + additional |
| 227 | YJR127C | RSF2 |  |  | frame | polyT plus 1 | YJM693 + additional |
| 228 | YJR149W |  |  |  | stop |  | YJM1355 |
| 228 | YJR149W |  |  |  | frame | minus 1 | YJM1615 + additional |
| 228 | YJR149W |  |  |  | stop near C-terminal |  | YJM470 + additional |
| 229 | YJR152W | DAL5 |  |  | frame | polyA minus 1 | YJM244 + additional |
| 230 | YKL050C | LPX2 |  |  | frame | polyA plus 1 | YJM1574 |
| 231 | YKL068W-A |  |  |  | stop |  | YJM1443 |
| 232 | YKL070W |  |  |  | stop |  | YJM1385 + additional |
| 232 | YKL070W |  |  |  | two frameshifts two stop codons | PolyC minus 1, minus 1 | YJM1399 |
| 232 | YKL070W |  |  |  | frame | polyT plus 1 | YJM1418 |
| 232 | YKL070W |  |  |  | frame | polyA minus 1 | YJM1574 + additional |
| 233 | YKL071W | OSI1 |  |  | frame | minus 1 | YJM689 + additional |
| 234 | YKL093W | MBR1 |  |  | stop |  | YJM1338 + additional |
| 235 | YKL101W | HSL1 |  |  | frame | polyA plus 1 | YJM271 |
| 236 | YKL114C | APN1 |  |  | stop |  | YJM689 |
| 237 | YKL162C |  |  |  | stop |  | YJM1418 |
| 237 | YKL162C |  |  |  | frame | polyC minus 1 | YJM996 + additional |
| 238 | YKL209C | STE6 |  |  | stop near C-terminal |  | YJM1399 |
| 238 | YKL209C | STE6 |  |  | stop |  | YJM1549 |
| 239 | YKL218C | SRY1 |  |  | stop |  | YJM1386 |
| 240 | YKL219W | COS9 |  |  | frame | multiple frameshifts | YJM1252 |
| 240 | YKL219W | COS9 |  |  | frame | polyA minus 1 | YJM1417 |
| 240 | YKL219W | COS9 |  |  | frame | polyG plus 1 | YJM470 |
| 241 | YKL220C | FRE2 |  |  | stop |  | YJM1463 |
| 241 | YKL220C | FRE2 |  |  | frame | polyA plus 1 | YJM270 |
| 242 | YKL221W | MCH2 |  |  | stop |  | YJM1447 |
| 242 | YKL221W | MCH2 |  |  | frame | plus 1 | YJM682 + additional |
| 243 | YKL222C |  |  |  | insertion | mitochondrial genome DNA insertion | YJM1463 |
| 243 | YKL222C |  |  |  | frame | PolyA plus 1 | YJM555 + additional |
| 244 | YKR015C |  |  |  | stop |  | YJM1386 |
| 245 | YKR039W | GAP1 |  |  | stop |  | YJM271 |
| 246 | YKR053C | YSR3 |  |  | frame | polyA minus 1 | YJM683 + additional |
| 247 | YKR061W | KTR2 |  |  | stop |  | YJM1311 |
| 248 | YKR064W | OAF3 |  |  | frame | polyA plus 1 | YJM270 + additional |
| 249 | YKR076W | ECM4 |  |  | stop |  | YJM1615 |
| 250 | YKR089C | TGL4 |  |  | stop |  | YJM195 |
| 251 | YKR093W | PTR2 |  |  | frame | plus 5 tandem duplication | YJM1415 |
| 251 | YKR093W | PTR2 |  |  | frame | poly C plus 1 | YJM1433 |
| 251 | YKR093W | PTR2 |  |  | frame | plus 1 | YJM1477 + additional |
| 251 | YKR093W | PTR2 |  |  | frame | minus 1 | YJM1574 + additional |
| 251 | YKR093W | PTR2 |  |  | stop |  | YJM270 + additional |
| 251 | YKR093W | PTR2 |  |  | frame and stop | minus 1 | YJM683 + additional |
| 252 | YKR096W | ESL2 |  |  | C terminal deletion |  | YJM1129 + additional |
| 252 | YKR096W | ESL2 |  |  | frame | polyC plus 1 | YJM244 |
| 253 | YKR098C | UBP11 |  |  | frame | polyA plus 1 | YJM1252 |
| 254 | YLL014W | EMC6 |  |  | stop |  | YJM689 |
| 255 | YLL016W | SDC25 | YLL017W | SDC25 | frame | polyT plus 1 | S288c + additional |
| 255 | YLL016W | SDC25 | YLL017W | SDC25 | frame | 19 base deletion | YJM1083 |
| 255 | YLL016W | SDC25 | YLL017W | SDC25 | frame | polyA plus 1 | YJM1388 |
| 255 | YLL016W | SDC25 | YLL017W | SDC25 | frame | minus 1 | YJM1419 + additional |
| 255 | YLL016W | SDC25 | YLL017W | SDC25 | frame | polyA plus 4 | YJM1460 |
| 255 | YLL016W | SDC25 | YLL017W | SDC25 | frame | plus 1 | YJM451 + additional |
| 256 | YLL046C | RNP1 |  |  | frame | polyA plus 1 | YJM1388 |
| 256 | YLL046C | RNP1 |  |  | frame | polyA plus 1 | YJM270 |
| 257 | YLL051C | FRE6 |  |  | frame | minus 1 | YJM1083 |
| 257 | YLL051C | FRE6 |  |  | stop |  | YJM1385 |
| 258 | YLL052C | AQY2 | YLL053C |  | frame | 11 base deletion | S288c + additional |
| 258 | YLL052C | AQY2 | YLL053C |  | frame | minus 1 | YJM1355 |
| 258 | YLL052C | AQY2 | YLL053C |  | frame | polyG minus 1 | YJM1447 |
| 258 | YLL052C | AQY2 | YLL053C |  | two stops |  | YJM1478 + additional |
| 258 | YLL052C | AQY2 | YLL053C |  | frame | polyA minus 1 | YJM1479 + additional |
| 259 | YLL053C |  |  |  | frame | polyG minus 1 | YJM1477 |
| 259 | YLL053C |  |  |  | stop |  | YJM1478 |
| 260 | YLL055W | YCT1 |  |  | frame | polyA minus 1 | YJM1252 |
| 260 | YLL055W | YCT1 |  |  | frame | minus 1 | YJM270 |
| 261 | YLL057C | JLP1 |  |  | stop |  | YJM627 + additional |
| 262 | YLL061W | MMP1 |  |  | frame | plus 1 | YJM1355 |
| 262 | YLL061W | MMP1 |  |  | frame | polyG minus 1 | YJM1401 |
| 262 | YLL061W | MMP1 |  |  | stop |  | YJM1463 + additional |
| 262 | YLL061W | MMP1 |  |  | two stops |  | YJM1592 + additional |
| 263 | YLR001C |  |  |  | stop |  | YJM975 + additional |
| 264 | YLR011W | LOT6 |  |  | frame | polyA minus 1 | YJM1386 |
| 265 | YLR030W |  | YLR031W |  | frame | minus 1 | S288c + additional |
| 265 | YLR030W |  | YLR031W |  | stop |  | YJM1447 |
| 265 | YLR030W |  | YLR031W |  | two frameshifts | polyC plus 2, polyA plus 2 | YJM1478 |
| 265 | YLR030W |  | YLR031W |  | frame | polyA plus 1 | YJM270 |
| 266 | YLR036C |  |  |  | stop near C-terminal |  | S288c + additional |
| 266 | YLR036C |  |  |  | stop |  | S288c + additional |
| 266 | YLR036C |  |  |  | frame | polyT minus q | YJM1592 + additional |
| 266 | YLR036C |  |  |  | two frameshifts | polyA plus q, polyT plus q | YJM451 |
| 266 | YLR036C |  |  |  | frame | polyA plus q | YJM451 |
| 266 | YLR036C |  |  |  | frame | polyG minus q | YJM682 + additional |
| 267 | YLR039C | RIC1 |  |  | frame | polyA plus 1 | YJM1463 |
| 268 | YLR046C |  |  |  | inversion | 37 base inversion | YJM1388 + additional |
| 268 | YLR046C |  |  |  | frame | polyG plus 1 | YJM1460 + additional |
| 268 | YLR046C |  |  |  | frame | polyT plus 1 | YJM1463 |
| 268 | YLR046C |  |  |  | inversion |  | YJM1592 + additional |
| 268 | YLR046C |  |  |  | frame | plus 1 | YJM195 |
| 269 | YLR049C | MLO50 |  |  | stop |  | YJM1592 |
| 270 | YLR057W | MNL2 |  |  | frame | polyA minus 1 | YJM1439 |
| 271 | YLR080W | EMP46 |  |  | frame | polyA plus 1 | YJM1479 + additional |
| 272 | YLR099C | ICT1 |  |  | stop |  | YJM1549 + additional |
| 273 | YLR121C | YPS3 |  |  | frame | polyA minus 1 | YJM248 + additional |
| 274 | YLR125W |  |  |  | stop |  | YJM1389 + additional |
| 274 | YLR125W |  |  |  | frame | polyA plus 1 | YJM1415 + additional |
| 275 | YLR130C | ZRT2 |  |  | frame | polyA plus 1 | YJM1250 |
| 276 | YLR136C | TIS11 |  |  | stop |  | YJM248 + additional |
| 277 | YLR152C |  |  |  | frame | polyG plus 1 | YJM1083 |
| 277 | YLR152C |  |  |  | frame | polyA plus 1 | YJM1307 + additional |
| 277 | YLR152C |  |  |  | stop |  | YJM1478 |
| 278 | YLR165C | PUS5 |  |  | frame | minus 1 | YJM1615 + additional |
| 279 | YLR187W | SKG3 |  |  | stop |  | YJM1419 + additional |
| 280 | YLR194C | NCW2 |  |  | stop |  | YJM248 |
| 281 | YLR214W | FRE1 |  |  | insertion | Ty1 LTR insertion | YJM270 |
| 281 | YLR214W | FRE1 |  |  | frame | polyA minus 1 | YJM996 + additional |
| 282 | YLR256W | HAP1 |  |  | insertion | Ty1 insertion | S288c |
| 283 | YLR266C | PDR8 |  |  | stop |  | YJM1447 |
| 284 | YLR273C | PIG1 |  |  | frame | minus 1 | YJM1447 |
| 284 | YLR273C | PIG1 |  |  | stop |  | YJM1574 |
| 285 | YLR278C |  |  |  | frame | polyA plus 1 | YJM1304 |
| 286 | YLR286C | CTS1 |  |  | stop |  | YJM1439 |
| 286 | YLR286C | CTS1 |  |  | stop |  | YJM189 |
| 287 | YLR300W | EXG1 |  |  | frame | minus 1 | YJM1443 |
| 288 | YLR312C | ATG39 |  |  | stop |  | YJM1479 + additional |
| 289 | YLR313C | SPH1 |  |  | frame | 2 base insertion | S288c + additional |
| 290 | YLR324W | PEX30 |  |  | deletion | C terminal end | YJM451 + additional |
| 291 | YLR346C | CIS1 |  |  | stop |  | YJM1385 |
| 292 | YLR390W | ECM19 |  |  | stop near C-terminal |  | YJM1342 |
| 292 | YLR390W | ECM19 |  |  | stop |  | YJM1463 |
| 293 | YLR414C | PUN1 |  |  | frame | polyT plus 1 | YJM248 + additional |
| 294 | YLR424W | SPP382 |  |  | frame | polyA minus 1 | YJM450 |
| 295 | YLR436C | ECM30 |  |  | stop |  | YJM555 |
| 296 | YML006C | GIS4 |  |  | stop |  | YJM189 |
| 297 | YML018C |  |  |  | frame | minus 1 | YJM541 + additional |
| 298 | YML052W | SUR7 |  |  | stop |  | YJM1307 |
| 299 | YML074C | FPR3 |  |  | frame | plus 1 | YJM428 + additional |
| 300 | YML088W | UFO1 |  |  | stop |  | YJM271 + additional |
| 301 | YML100W-A |  |  |  | frame | polyC minus 1 | S288c |
| 301 | YML100W-A |  |  |  | stop |  | YJM1386 |
| 302 | YML104C | MDM1 |  |  | frame | plus 1 | YJM1129 + additional |
| 303 | YMR001C-A |  |  |  | two frameshifts | poly(AT) plus 2, polyT plus 1 | YJM1478 |
| 304 | YMR030W | RSF1 |  |  | frame | polyA plus 1 | YJM1450 + additional |
| 304 | YMR030W | RSF1 |  |  | stop |  | YJM1477 |
| 304 | YMR030W | RSF1 |  |  | frame | minus 1 | YJM1592 + additional |
| 304 | YMR030W | RSF1 |  |  | frame | minus 1 | YJM987 + additional |
| 305 | YMR070W | MOT3 |  |  | stop |  | YJM1381 |
| 306 | YMR084W |  | YMR085W |  | frame | polyC minus 1 | S288c + additional |
| 306 | YMR084W |  | YMR085W |  | two stops |  | YJM1477 + additional |
| 307 | YMR096W | SNZ1 |  |  | stop |  | YJM1463 |
| 308 | YMR107W | SPG4 |  |  | stop |  | YJM1463 |
| 308 | YMR107W | SPG4 |  |  | insertion | Ty1 insertion | YJM450 |
| 309 | YMR110C | HFD1 |  |  | stop |  | YJM681 + additional |
| 310 | YMR132C | JLP2 |  |  | stop near C-terminal | Ty2 insertion in stop codon - 6 amino acid extension | YJM1355 |
| 311 | YMR136W | GAT2 |  |  | frame | polyA plus 1 | YJM1355 |
| 312 | YMR140W | SIP5 |  |  | frame | plus 1 | YJM683 + additional |
| 313 | YMR152W | YIM1 |  |  | stop |  | YJM1252 + additional |
| 314 | YMR159C | ATG16 |  |  | frame | polyA plus 1 | YJM1419 |
| 315 | YMR169C | ALD3 |  |  | stop |  | YJM1443 |
| 316 | YMR170C | ALD2 |  |  | stop |  | YJM1443 |
| 317 | YMR176W | ECM5 |  |  | frame | polyG minus 1 | YJM1592 + additional |
| 317 | YMR176W | ECM5 |  |  | frame | polyA plus 1 | YJM541 + additional |
| 318 | YMR180C | CTL1 |  |  | frame and stop | 23 base tandem insertion | YJM1326 + additional |
| 318 | YMR180C | CTL1 |  |  | frame | minus 1 | YJM1460 |
| 318 | YMR180C | CTL1 |  |  | stop |  | YJM270 + additional |
| 319 | YMR192W | GYL1 |  |  | stop |  | YJM1202 + additional |
| 320 | YMR204C | INP1 |  |  | stop |  | YJM1336 |
| 321 | YMR215W | GAS3 |  |  | frame | polyT plus 1 | YJM248 + additional |
| 322 | YMR251W | GTO3 |  |  | stop |  | YJM1463 |
| 323 | YMR265C |  |  |  | frame | plus 8 tandem duplication insertion | YJM1526 |
| 324 | YMR271C | URA10 |  |  | frame | polyA plus 1 | YJM1463 |
| 325 | YMR279C | ATR2 |  |  | frame | minus 1 | YJM1447 |
| 325 | YMR279C | ATR2 |  |  | frame | minus 1 | YJM1574 |
| 326 | YMR317W |  |  |  | frame | polyA minus 2 | YJM1460 + additional |
| 326 | YMR317W |  |  |  | stop |  | YJM1479 + additional |
| 327 | YNL001W | DOM34 |  |  | insertion | Ty2 insertion | YJM470 |
| 328 | YNL019C |  |  |  | frame | polyA minus 1 (same mutation as in YNL033W) | YJM972 |
| 329 | YNL033W |  |  |  | frame | polyA minus 1 (same mutation as in YNL019C) | YJM975 + additional |
| 330 | YNL040W |  |  |  | stop |  | YJM1615 |
| 331 | YNL045W | LAP2 |  |  | frame | polyC plus 1 | YJM450 + additional |
| 332 | YNL074C | MLF3 |  |  | stop |  | YJM1129 |
| 333 | YNL087W | TCB2 |  |  | frame | polyA minus 1 | YJM1208 |
| 333 | YNL087W | TCB2 |  |  | frame | polyT plus 1 | YJM270 |
| 333 | YNL087W | TCB2 |  |  | frame | polyT minus 1 | YJM682 + additional |
| 333 | YNL087W | TCB2 |  |  | frame | minus 1 | YJM683 |
| 333 | YNL087W | TCB2 |  |  | frame | polyT plus 1 | YJM689 + additional |
| 334 | YNL095C |  |  |  | frame | polyA minus 1 | YJM1549 + additional |
| 334 | YNL095C |  |  |  | frame | polyG plus 1 | YJM270 + additional |
| 334 | YNL095C |  |  |  | frame | polyG minus 1 | YJM627 |
| 335 | YNL100W | MIC27 |  |  | stop |  | YJM470 |
| 336 | YNL125C | ESBP6 |  |  | frame | polyC plus 1 | YJM1386 + additional |
| 336 | YNL125C | ESBP6 |  |  | frame | polyC minus 1 | YJM1463 |
| 337 | YNL159C | ASI2 |  |  | stop |  | YJM1478 |
| 338 | YNL175C | NOP13 |  |  | frame | polyA plus 1 | YJM1463 |
| 339 | YNL193W |  |  |  | frame | polyT plus 1 | YJM1385 |
| 340 | YNL194C |  |  |  | stop |  | YJM1388 + additional |
| 340 | YNL194C |  |  |  | frame | polyC plus 1 | YJM248 + additional |
| 341 | YNL195C |  |  |  | frame | polyA minus 1 | YJM1401 |
| 341 | YNL195C |  |  |  | stop |  | YJM1463 |
| 342 | YNL196C | SLZ1 |  |  | frame | polyA plus 1 | YJM271 + additional |
| 343 | YNL237W | YTP1 |  |  | frame | 140base deletion | YJM1415 |
| 344 | YNL239W | LAP3 |  |  | frame | 4 base tandem duplication insertion | YJM682 + additional |
| 345 | YNL275W | BOR1 |  |  | frame | minus 1 | YJM1304 |
| 346 | YNL279W | PRM1 |  |  | frame | minus 1 | YJM693 |
| 347 | YNL283C | WSC2 |  |  | stop |  | YJM270 |
| 348 | YNL331C | AAD14 |  |  | frame | minus 1 | YJM1338 |
| 348 | YNL331C | AAD14 |  |  | frame | polyT plus 1 | YJM270 |
| 349 | YNR002C | ATO2 |  |  | stop |  | YJM189 |
| 350 | YNR056C | BIO5 |  |  | stop |  | YJM1463 |
| 351 | YNR057C | BIO4 |  |  | stop |  | YJM1307 + additional |
| 352 | YNR059W | MNT4 |  |  | stop |  | YJM248 + additional |
| 353 | YNR060W | FRE4 |  |  | frame | polyA plus 1 | YJM1304 |
| 354 | YNR061C |  |  |  | frame | polyT plus 1 | YJM1387 + additional |
| 355 | YNR062C | PUL3 |  |  | frame | polyT plus 1 | YJM195 |
| 356 | YNR064C |  |  |  | stop |  | YJM248 + additional |
| 357 | YNR065C |  | YNR066C |  | stop |  | s288c + additional |
| 357 | YNR065C |  |  |  | frame | polyA minus 1 | YJM1244 |
| 357 | YNR065C |  |  |  | frame | polyC minus 1 | YJM555 |
| 358 | YNR066C |  |  |  | frame | polyA minus 1 | YJM1355 + additional |
| 358 | YNR066C |  |  |  | frame | minus 1 | YJM1574 + additional |
| 358 | YNR066C |  |  |  | frame | polyA plus 1 | YJM326 + additional |
| 358 | YNR066C |  |  |  | stop |  | YJM627 + additional |
| 359 | YNR068C |  |  |  | frame | polyC minus 1 | YJM1389 + additional |
| 359 | YNR068C |  |  |  | frame | polyA plus 1 | YJM1419 |
| 360 | YNR069C | BSC5 |  |  | frame | minus 1 | YJM1381 |
| 360 | YNR069C | BSC5 |  |  | stop |  | YJM1401 + additional |
| 360 | YNR069C | BSC5 |  |  | frame | polyT plus 1 | YJM1447 |
| 360 | YNR069C | BSC5 |  |  | frame | polyC minus 1 | YJM693 + additional |
| 361 | YNR070W | PDR18 |  |  | two frameshifts | plus 1, poly T plus 1 | YJM1463 |
| 361 | YNR070W | PDR18 |  |  | frame | plus 1 | YJM1549 + additional |
| 361 | YNR070W | PDR18 |  |  | frame | polyA plus 1 | YJM453 + additional |
| 362 | YNR071C |  |  |  | frame | polyG plus 1 | YJM1592 + additional |
| 362 | YNR071C |  |  |  | stop |  | YJM270 |
| 363 | YOL014W |  |  |  | stop |  | YJM1463 + additional |
| 364 | YOL019W-A |  |  |  | stop |  | YJM1592 + additional |
| 365 | YOL024W |  |  |  | stop |  | YJM1338 |
| 366 | YOL028C | YAP7 | YOL029C |  | stop |  | YJM1401 + additional |
| 366 | YOL028C | YAP7 | YOL029C |  | frame | poly(AT) minus 2 | YJM1443 |
| 366 | YOL028C | YAP7 | YOL029C |  | frame | polyA minus 1 | S288c + additional |
| 367 | YOL029C |  |  |  | frame | minus 2 | YJM1443 |
| 367 | YOL029C |  |  |  | stop |  | YJM470 + additional |
| 368 | YOL030W | GAS5 |  |  | two frameshifts | polyT plus 1, polyC plus 1 | YJM1078 + additional |
| 368 | YOL030W | GAS5 |  |  | stop |  | YJM1252 |
| 368 | YOL030W | GAS5 |  |  | insertion | 17kb transposable element insertion | YJM453 |
| 369 | YOL047C | LDS2 |  |  | frame | polyT plus 1 | YJM1389 + additional |
| 370 | YOL068C | HST1 |  |  | frame | polyT plus 1 | YJM1190 |
| 371 | YOL083W | ATG34 |  |  | frame | polyA minus 1 | YJM1083 |
| 371 | YOL083W | ATG34 |  |  | frame | polyT plus 1 | YJM1342 |
| 372 | YOL105C | WSC3 |  |  | frame | minus 1 | YJM1252 |
| 373 | YOL109W | ZEO1 |  |  | N terminal deletion |  | YJM1307 |
| 374 | YOL112W | MSB4 |  |  | stop |  | YJM1549 |
| 375 | YOL145C | CTR9 |  |  | stop |  | YJM456 |
| 376 | YOL151W | GRE2 |  |  | frame | polyA minus 1 | YJM1401 |
| 377 | YOL152W | FRE7 |  |  | frame | polyC minus 1 | YJM1387 |
| 377 | YOL152W | FRE7 |  |  | frame | polyA minus 1 | YJM1463 |
| 377 | YOL152W | FRE7 |  |  | frame | 46 base deletion | YJM195 + additional |
| 378 | YOL154W | ZPS1 |  |  | stop |  | YJM1381 + additional |
| 379 | YOR010C | TIR2 |  |  | stop |  | YJM1252 |
| 380 | YOR028C | CIN5 |  |  | frame | polyA plus 1 | YJM1415 |
| 381 | YOR030W | DFG16 |  |  | stop |  | YJM1252 |
| 382 | YOR032C | HMS1 |  |  | stop |  | YJM1615 + additional |
| 383 | YOR032W-A |  |  |  | stop |  | YJM470 + additional |
| 384 | YOR035C | SHE4 |  |  | frame | polyA minus 1 | YJM1383 |
| 385 | YOR043W | WHI2 |  |  | frame | 7 base tandem duplication insertion | YJM1342 |
| 385 | YOR043W | WHI2 |  |  | frame | polyT plus 1 | YJM1479 |
| 385 | YOR043W | WHI2 |  |  | stop |  | YJM248 + additional |
| 386 | YOR051C | ETT1 |  |  | frame | 19 base insertion | YJM1332 |
| 387 | YOR071C | NRT1 |  |  | two frameshifts and stop | plus 1, minus 1 | YJM1383 |
| 387 | YOR071C | NRT1 |  |  | frame and stop | polyT plus 1 | YJM1478 + additional |
| 387 | YOR071C | NRT1 |  |  | frame | plus 2 | YJM1574 + additional |
| 388 | YOR081C | TGL5 |  |  | stop |  | YJM1549 |
| 389 | YOR083W | WHI5 |  |  | stop |  | YJM451 |
| 390 | YOR107W | RGS2 |  |  | stop |  | YJM1078 + additional |
| 391 | YOR118W | RTC5 |  |  | frame | polyT plus 1 | YJM1355 |
| 392 | YOR120W | GCY1 |  |  | stop |  | YJM470 |
| 393 | YOR126C | IAH1 |  |  | stop |  | YJM1304 + additional |
| 394 | YOR148C | SPP2 |  |  | stop near C-terminal | polyA plus 2 moves stop codon four amino acids | YJM554 + additional |
| 394 | YOR148C | SPP2 |  |  | stop near C-terminal | polyA plus 1 moves stop codon two amino acids | YJM693 + additional |
| 395 | YOR186W |  |  |  | stop |  | YJM1248 + additional |
| 396 | YOR190W | SPR1 |  |  | stop |  | YJM271 |
| 397 | YOR192C | THI72 |  |  | frame | polyT plus 1 | YJM1083 |
| 397 | YOR192C | THI72 |  |  | stop |  | YJM1304 + additional |
| 398 | YOR197W | MCA1 |  |  | frame | polyC plus 1 | YJM193 |
| 399 | YOR220W | RCN2 |  |  | frame | minus 1 | YJM1419 |
| 399 | YOR220W | RCN2 |  |  | stop |  | YJM693 |
| 400 | YOR233W | KIN4 |  |  | stop |  | YJM270 |
| 401 | YOR284W | HUA2 |  |  | frame | polyA minus 1 | YJM248 + additional |
| 401 | YOR284W | HUA2 |  |  | stop |  | YJM681 + additional |
| 401 | YOR284W | HUA2 |  |  | frame | minus 1 | YJM996 + additional |
| 402 | YOR301W | RAX1 |  |  | stop |  | YJM1190 |
| 403 | YOR307C | SLY41 |  |  | stop |  | YJM248 |
| 404 | YOR316C | COT1 |  |  | stop |  | YJM1447 + additional |
| 404 | YOR316C | COT1 |  |  | frame | polyC minus 1 | YJM1450 + additional |
| 405 | YOR337W | TEA1 |  |  | stop |  | YJM193 + additional |
| 406 | YOR360C | PDE2 |  |  | frame | minus 1 | YJM1304 |
| 406 | YOR360C | PDE2 |  |  | frame | polyA minus 1 | YJM1355 |
| 407 | YOR361C | PRT1 |  |  | stop |  | YJM1336 |
| 408 | YOR365C |  |  |  | stop |  | YJM1356 |
| 409 | YOR378W | AMF1 |  |  | frame | polyT minus 1 | YJM1479 + additional |
| 410 | YOR380W | RDR1 |  |  | stop |  | YJM1549 + additional |
| 411 | YOR381W | FRE3 |  |  | stop |  | YJM1307 |
| 411 | YOR381W | FRE3 |  |  | frame | 7 base deletion | YJM1387 |
| 411 | YOR381W | FRE3 |  |  | frame | 8 base deletion | YJM1477 + additional |
| 411 | YOR381W | FRE3 |  |  | frame | polyA plus 1 | YJM453 + additional |
| 412 | YPL014W | CIP1 |  |  | frame | polyA minus 1 | YJM1615 + additional |
| 413 | YPL017C | IRC15 |  |  | frame | minus 1 | YJM1443 |
| 414 | YPL041C | MRX11 |  |  | stop |  | YJM470 |
| 415 | YPL054W | LEE1 |  |  | frame | minus 1 | YJM1355 |
| 416 | YPL058C | PDR12 |  |  | stop |  | YJM1549 |
| 417 | YPL092W | SSU1 |  |  | stop |  | YJM270 |
| 418 | YPL098C | MGR2 |  |  | stop |  | YJM1307 |
| 419 | YPL100W | ATG21 |  |  | stop |  | YJM195 |
| 420 | YPL147W | PXA1 |  |  | stop |  | YJM689 |
| 421 | YPL156C | PRM4 |  |  | frame | polyA plus 1 | YJM193 |
| 422 | YPL164C | MLH3 |  |  | stop |  | YJM195 |
| 423 | YPL165C | SET6 |  |  | stop |  | YJM1307 |
| 424 | YPL171C | OYE3 |  |  | frame | polyC minus 1 | YJM1342 |
| 424 | YPL171C | OYE3 |  |  | frame | 41 base deletion | YJM1399 |
| 424 | YPL171C | OYE3 |  |  | frame | minus 1 | YJM1417 |
| 424 | YPL171C | OYE3 |  |  | stop |  | YJM1479 + additional |
| 425 | YPL177C | CUP9 |  |  | stop |  | YJM1383 |
| 425 | YPL177C | CUP9 |  |  | frame | polyG plus 1 | YJM248 |
| 426 | YPL179W | PPQ1 |  |  | stop |  | YJM693 |
| 427 | YPL191C | MIY2 |  |  | frame | 5 base deletion | YJM270 + additional |
| 428 | YPL216W |  |  |  | frame | polyA plus 1 | YJM1250 |
| 428 | YPL216W |  |  |  | frame | minus 1 | YJM1338 |
| 428 | YPL216W |  |  |  | stop |  | YJM1385 + additional |
| 428 | YPL216W |  |  |  | frame | minus 1 | YJM1592 |
| 429 | YPL223C | GRE1 |  |  | stop |  | YJM1083 |
| 430 | YPL247C |  |  |  | stop |  | YJM271 |
| 431 | YPL248C | GAL4 |  |  | frame | polyA plus 1 | YJM270 + additional |
| 432 | YPL273W | SAM4 |  |  | stop |  | YJM990 + additional |
| 433 | YPL274W | SAM3 |  |  | stop |  | YJM1307 + additional |
| 433 | YPL274W | SAM3 |  |  | frame | polyG minus 1 | YJM1355 |
| 434 | YPR009W | SUT2 |  |  | stop |  | YJM1078 + additional |
| 434 | YPR009W | SUT2 |  |  | frame | minus 1 | YJM1355 |
| 435 | YPR065W | ROX1 |  |  | stop |  | YJM1248 + additional |
| 436 | YPR071W |  |  |  | stop |  | YJM1447 |
| 436 | YPR071W |  |  |  | C terminal deletion |  | YJM195 |
| 436 | YPR071W |  |  |  | C-terminal deletion |  | YJM195 |
| 437 | YPR078C |  |  |  | stop |  | YJM1460 |
| 437 | YPR078C |  |  |  | stop near C-terminal |  | YJM1460 |
| 438 | YPR091C | NVJ2 |  |  | stop |  | YJM428 |
| 438 | YPR091C | NVJ2 |  |  | stop near C-terminal |  | YJM689 + additional |
| 439 | YPR114W |  |  |  | stop |  | YJM450 |
| 440 | YPR117W | HOB2 |  |  | stop |  | YJM271 |
| 441 | YPR121W | THI22 |  |  | stop |  | YJM1307 |
| 442 | YPR192W | AQY1 |  |  | frame | polyA plus 1 | YJM1389 |
| 442 | YPR192W | AQY1 |  |  | frame | polyA plus 2 | YJM1401 |
| 442 | YPR192W | AQY1 |  |  | frame | polyA minus 1 | YJM627 + additional |
| 443 | YPR193C | HPA2 |  |  | stop |  | YJM1381 + additional |
| 443 | YPR193C | HPA2 |  |  | frame | polyT plus 1 | YJM1592 + additional |
| 444 | YPR194C | OPT2 |  |  | frame | PolyC minus 1 | YJM1311 |
| 444 | YPR194C | OPT2 |  |  | frame | minus 1 | YJM1385 |
| 444 | YPR194C | OPT2 |  |  | frame | minus 1 | YJM1433 |
| 444 | YPR194C | OPT2 |  |  | stop |  | YJM1574 + additional |
| 444 | YPR194C | OPT2 |  |  | frame | polyC plus 1 | YJM193 |
| 444 | YPR194C | OPT2 |  |  | frame | polyA plus 2 | YJM271 |
| 444 | YPR194C | OPT2 |  |  | frame | 17 base tandem insertion | YJM555 |
| 445 | YPR196W |  |  |  | frame | minus 1 | YJM1383 + additional |
| 445 | YPR196W |  |  |  | frame | polyA minus 1 | YJM1439 + additional |
| 445 | YPR196W |  |  |  | stop |  | YJM1477 + additional |

Table S12 Single genes annotated as two genes in S288c due to allele containing frameshift, stop codon, or transposon insertion.

| Name | Old Name 1 | Old Name 2 (alias) | Name | Core? | Location | *A. gossypii* ortholog |
| --- | --- | --- | --- | --- | --- | --- |
| YBR012C | YBR012C | YBR013C |  | core | Central |  |
| YER039C | YER039C | YER039C-A | HVG1 | core | Central | AFR236C |
| YHR212W-A | YHR212W-A | YHR213W |  | not core | Subtelomeric | Unclear, flocculin gene |
| YIL164C | YIL164C | YIL165C | NIT1 | core | Central |  |
| YIL170W | YIL170W | YIL171W | HXT12 | not core | Subtelomeric | Unclear, hexose transporter |
| YJL108C | YJL108C | YJL107C | PRM10 | core | Central | AFR075C |
| YLL016W | YLL016W | YLL017W | SDC25 | core | Central | ADL038W |
| YLL052C | YLL052C | YLL053C | AQY2 | core | Central |  |
| YLR030W | YLR030W | YLR031W |  | core | Central |  |
| YML002W | YML002W | YML003W | VRL1 | core | Central | ACR006C |
| YMR084W | YMR084W | YMR085W |  | core | Central |  |
| YNR065C | YNR065C | YNR066C | [VTH3] | core | Central |  |
| YPL275W | YPL275W | YPL276W | FDH2 | not core | Subtelomeric |  |
| YPL277C | YPL277C | YPL278C |  | not core | Subtelomeric |  |
| Former separate open reading frames merged in SGD to a single annotation. | | | | | | |
| YIL167W | YIL167W | YIL168W | SDL1 | core | Central |  |

Table S13 tRNA copy number in S288c plus 93 strains

| residue | anti-codon | # tRNA copies in S288c | fraction of 93 strains | Copies in majority of strains | strains with alternate number | | Number tRNA copies | Number of strains | | Number tRNA copies | Explanation |
| --- | --- | --- | --- | --- | --- | --- | --- | --- | --- | --- | --- |
| ARG | CCG | 1 | 93/93 | 1 |  |  |  |  |  |  |  |
| ARG | CCT | 1 | 93/93 | 1 |  |  |  |  |  |  |  |
| THR | CGT | 1 | 93/93 | 1 |  |  |  |  |  |  |  |
| GLN | CTG | 1 | 93/93 | 1 |  |  |  |  |  |  |  |
| LEU | GAG | 1 | 93/93 | 1 |  |  |  |  |  |  |  |
| SER | CGA | 1 | 93/93 | 1 |  |  |  |  |  |  |  |
| GLY | CCC | 2 | 93/93 | 2 |  |  |  |  |  |  |  |
| GLU | CTC | 2 | 93/93 | 2 |  |  |  |  |  |  |  |
| ILE | TAT | 2 | 93/93 | 2 |  |  |  |  |  |  |  |
| VAL | TAC | 2 | 93/93 | 2 |  |  |  |  |  |  |  |
| VAL | CAC | 2 | 93/93 | 2 |  |  |  |  |  |  |  |
| PRO | AGG | 2 | 92/93 | 2 | 1 | strain | 1 |  |  |  | 1 |
| PRO | GGG | 0 | 1/93 | 1 | 1 | strain | 1 |  |  |  | 1 |
| GLY | TCC | 3 | 93/93 | 3 |  |  |  |  |  |  |  |
| LEU | TAG | 3 | 93/93 | 3 |  |  |  |  |  |  |  |
| THR | TGT | 3 | 93/93 | 3 |  |  |  |  |  |  |  |
| SER | GCT | 2* | 93/93 | 4 |  |  |  |  |  |  | 2 |
| SER | TGA | 4 | 93/93 | 4 |  |  |  |  |  |  |  |
| CYS | GCA | 4 | 93/93 | 4 |  |  |  |  |  |  |  |
| ALA | TGC | 5 | 93/93 | 5 |  |  |  |  |  |  |  |
| TRP | CCA | 6 | 93/93 | 6 |  |  |  |  |  |  |  |
| ARG | ACG | 6 | 93/93 | 6 |  |  |  |  |  |  |  |
| LEU | TAA | 7 | 93/93 | 7 |  |  |  |  |  |  |  |
| LYS | TTT | 7 | 93/93 | 7 |  |  |  |  |  |  |  |
| HIS | GTG | 7 | 93/93 | 7 |  |  |  |  |  |  |  |
| TYR | GTA | 8 | 92/93 | 8 | 1 | strain | 7 |  |  |  | 3 |
| GLN | TTG | 9 | 93/93 | 9 |  |  |  |  |  |  |  |
| LEU | CAA | 10 | 93/93 | 10 |  |  |  |  |  |  |  |
| MET | CAT | 10 | 92/93 | 10 | 1 | strain | 9 |  |  |  | 4 |
| PHE | GAA | 10 | 92/93 | 10 | 1 | strain | 9 |  |  |  | 5 |
| PRO | TGG | 10 | 92/93 | 10 | 1 | strain | 9 |  |  |  | 6 |
| ASN | GTT | 10 | 87/93 | 10 | 1 | strain | 9 | 5 | strains | 11 | 7 |
| ALA | AGC | 11 | 93/93 | 11 |  |  |  |  |  |  |  |
| ARG | TCT | 11 | 89/93 | 11 | 4 | strains | 12 |  |  |  | 8 |
| SER | AGA | 11 | 91/93 | 11 | 2 | strains | 10 |  |  |  | 9 |
| THR | AGT | 11 | 90/93 | 11 | 3 | strains | 10 |  |  |  | 10 |
| ILE | AAT | 13 | 89/93 | 13 | 4 | strains | 12 |  |  |  | 11 |
| GLU | TTC | 14 | 93/93 | 14 |  |  |  |  |  |  |  |
| VAL | AAC | 14 | 93/93 | 14 |  |  |  |  |  |  |  |
| LYS | CTT | 14 | 85/93 | 14 | 7 | strains | 13 |  |  |  | 12 |
| ASP | GTC | 16 | 93/93 | 16 |  |  |  |  |  |  |  |
| GLY | GCC | 16 | 90/93 | 16 | 3 | strains | 15 |  |  |  | 13 |

Note that number of tRNA genes is for haploid genome. Aneuploid strains have additional tRNA genes not shown in this table.

Specific explanations:

1) tRNA SUF10 tP(AGG)N chromosome XIV mutated AGG->GGG anti-codon in strain YJM1355; only example found of change in anti-codon

2) due to sequence and annotation errors in S288c, tX(XXX)D and tX(XXX)L should be Ser GCT tRNAs

3) strain YJM1447 missing SUP7 tY(GUA)J1 from chromosome X

4) strain YJM1385 missing tM(CAU)C from chromosome III

5) strain YJM1549 small deletion in tRNA gene tF(GAA)G on chromosome VII

6) strain YJM1447 missing tP(UGG)O3 on chromosome XV

7) strains YJM1478, YJM1450, YJM1202, YJM681 - chromosome X extra copy at right telomere outside central portion of genome, apparently a horizontally acquired mitochondrial tRNA from Torulaspora sp.; YJM1202 also has an extra copy of the same mitochondrial tRNA on the left telomere of chromosome X; strains YJM1202 and YJM1199 missing tN(GUU)N1 from chromosome XIV; YJM1083, duplicated tN(GUU)N1 on chromosome XIV

8) strains YJM1078, YJM1252, YJM248 extra copy on chromosome XI between YKR038C and YKR039W; YJM470 extra copy on chromosome X between YJR055W and YJR056C

9) strains YJM1385, YJM555 missing tS(AGA)A on chromosome I

10) strain YJM1342 missing tT(AGU)N2 on chromosome XIV; strains YJM1208, YJM1615 missing tT(AGU)C on chromosome III

11) strain YJM1447 missing tI(AAU)E1 on chromosome V; strains YJM1400 YJM1479 YJM1342, missing tI(AAU)B on chromosome II

12) strains YJM456, YJM451, YJM270, YJM248, YJM1443, YJM1338 tK(CUU)E2 missing from chromosome V - not a core gene; YJM1385 missing tK(CUU)C from chromosome III

13) strains YJM1342, YJM1400, YJM1479 missing tG(GCC)B from chromosome II; two additional strains YJM1477 and YJM1242 appear to have mutated versions of tG(GCC)B

Table S14 Location of gaps currently in the 93 genome sequences.

| Strain | Chr. | Start Ns | End Ns | Length | location of Ns | Gene | identity | length |  |  | start | end | start | end | E-value | Score |
| --- | --- | --- | --- | --- | --- | --- | --- | --- | --- | --- | --- | --- | --- | --- | --- | --- |
| YJM189 | I | 1 | 10 | 186714 | leftend | YAL063C | 84.926 | 1632 | 242 | 4 | 1 | 1630 | 16787 | 15158 | 0 | 1648 |
| YJM189 | I | 186705 | 186714 | 186714 | rightend | YAR050W | 79.974 | 1558 | 306 | 6 | 76 | 1630 | 16712 | 15158 | 0 | 1144 |
| YJM189 | II | 1 | 10 | 792368 | leftend | YNCB0001W | 100 | 84 | 0 | 0 | 1 | 84 | 10422 | 10505 | 4.16E-38 | 156 |
| YJM189 | II | 1076 | 1085 | 792368 | middle |  |  |  |  |  |  |  |  |  |  |  |
| YJM189 | II | 791160 | 791169 | 792368 | middle |  |  |  |  |  |  |  |  |  |  |  |
| YJM189 | II | 791353 | 791362 | 792368 | middle |  |  |  |  |  |  |  |  |  |  |  |
| YJM189 | II | 792359 | 792368 | 792368 | rightend | YBR297W | 92.031 | 1418 | 97 | 12 | 1 | 1407 | 782654 | 784066 | 0 | 1978 |
| YJM189 | III | 1 | 10 | 302832 | leftend | YCL073C | 99.784 | 1848 | 4 | 0 | 1 | 1848 | 7347 | 5500 | 0 | 3391 |
| YJM189 | III | 300153 | 300162 | 302832 | middle |  |  |  |  |  |  |  |  |  |  |  |
| YJM189 | III | 302832 | 302832 | 302832 | rightend_complete | YCR101C | 99.636 | 549 | 2 | 0 | 1 | 549 | 295763 | 295215 | 0 | 1003 |
| YJM189 | IV | 1 | 10 | 1457068 | leftend | YDL243C | 95.051 | 990 | 49 | 0 | 1 | 990 | 5095 | 4106 | 0 | 1557 |
| YJM189 | IV | 761 | 770 | 1457068 | middle |  |  |  |  |  |  |  |  |  |  |  |
| YJM189 | IV | 2478 | 2487 | 1457068 | middle |  |  |  |  |  |  |  |  |  |  |  |
| YJM189 | IV | 1457059 | 1457068 | 1457068 | rightend | YDR541C | 99.42 | 1035 | 6 | 0 | 1 | 1035 | 1454220 | 1453186 | 0 | 1879 |
| YJM189 | V | 1 | 10 | 552580 | leftend | YEL070W | 99.072 | 1509 | 14 | 0 | 1 | 1509 | 14732 | 16240 | 0 | 2710 |
| YJM189 | V | 552571 | 552580 | 552580 | rightend | YER187W | 91.981 | 424 | 32 | 2 | 4 | 426 | 548615 | 549037 | 3.44E-169 | 593 |
| YJM189 | VI | 1 | 10 | 237316 | leftend | YFL051C | 98.451 | 452 | 7 | 0 | 1 | 452 | 4368 | 3917 | 0 | 797 |
| YJM189 | VI | 674 | 683 | 237316 | middle |  |  |  |  |  |  |  |  |  |  |  |
| YJM189 | VI | 237307 | 237316 | 237316 | rightend | YFR055W | 100 | 1023 | 0 | 0 | 1 | 1023 | 231033 | 232055 | 0 | 1890 |
| YJM189 | VII | 1 | 10 | 1049145 | leftend | YGL258W | 97.585 | 621 | 15 | 0 | 1 | 621 | 11022 | 11642 | 0 | 1064 |
| YJM189 | VII | 1049136 | 1049145 | 1049145 | rightend | YGR287C | 98.927 | 1770 | 19 | 0 | 1 | 1770 | 1030824 | 1029055 | 0 | 3164 |
| YJM189 | VIII | 1 | 10 | 924982 | leftend | YPL274W | 99.943 | 1764 | 1 | 0 | 1 | 1764 | 13321 | 15084 | 0 | 3253 |
| YJM189 | VIII | 1095 | 1104 | 924982 | middle |  |  |  |  |  |  |  |  |  |  |  |
| YJM189 | VIII | 3442 | 3451 | 924982 | middle |  |  |  |  |  |  |  |  |  |  |  |
| YJM189 | VIII | 879107 | 879116 | 924982 | middle |  |  |  |  |  |  |  |  |  |  |  |
| YJM189 | VIII | 924982 | 924982 | 924982 | rightend_complete | YHR211W | 92.088 | 2136 | 145 | 2 | 1096 | 3228 | 870189 | 872303 | 0 | 2987 |
| YJM189 | IX | 1 |  | 407187 | leftend_complete | YIL169C | 93.798 | 1548 | 90 | 3 | 265 | 1812 | 6528 | 4987 | 0 | 2322 |
| YJM189 | IX | 1746 | 1755 | 407187 | middle |  |  |  |  |  |  |  |  |  |  |  |
| YJM189 | IX | 407178 | 407187 | 407187 | rightend | YIR039C | 99.009 | 1614 | 16 | 0 | 1 | 1614 | 405918 | 404305 | 0 | 2892 |
| YJM189 | X | 1 | 10 | 728293 | leftend | YJL213W | 99.699 | 996 | 3 | 0 | 1 | 996 | 33052 | 34047 | 0 | 1823 |
| YJM189 | X | 1413 | 1422 | 728293 | middle |  |  |  |  |  |  |  |  |  |  |  |
| YJM189 | X | 728293 | 728293 | 728293 | rightend_complete | YJR152W | 99.816 | 1632 | 3 | 0 | 1 | 1632 | 702248 | 703879 | 0 | 2998 |
| YJM189 | XI | 1 |  | 661068 | leftend_complete | YKL222C | 99.906 | 2118 | 2 | 0 | 1 | 2118 | 4111 | 1994 | 0 | 3901 |
| YJM189 | XI | 418 | 427 | 661068 | middle |  |  |  |  |  |  |  |  |  |  |  |
| YJM189 | XI | 660675 | 660684 | 661068 | middle |  |  |  |  |  |  |  |  |  |  |  |
| YJM189 | XI | 661068 | 661068 | 661068 | rightend_complete | YKR101W | 99.898 | 1965 | 2 | 0 | 1 | 1965 | 638014 | 639978 | 0 | 3618 |
| YJM189 | XII | 1 | 10 | 1597679 | leftend | YLL062C | 99.692 | 975 | 3 | 0 | 1 | 975 | 1936 | 962 | 0 | 1784 |
| YJM189 | XII | 1597670 | 1597679 | 1597679 | rightend | YLR460C | 94.336 | 1130 | 62 | 2 | 3 | 1131 | 1595738 | 1594610 | 0 | 1731 |
| YJM189 | XIII | 1 | 10 | 897693 | leftend | YML131W | 100 | 1098 | 0 | 0 | 1 | 1098 | 2982 | 4079 | 0 | 2028 |
| YJM189 | XIII | 615 | 624 | 897693 | middle |  |  |  |  |  |  |  |  |  |  |  |
| YJM189 | XIII | 896013 | 896022 | 897693 | middle |  |  |  |  |  |  |  |  |  |  |  |
| YJM189 | XIII | 897684 | 897693 | 897693 | rightend | YMR321C | 99.361 | 313 | 2 | 0 | 6 | 318 | 888068 | 887756 | 2.48E-161 | 568 |
| YJM189 | XIV | 1 | 10 | 763066 | leftend | YNL332W | 98.631 | 1023 | 14 | 0 | 1 | 1023 | 6419 | 7441 | 0 | 1812 |
| YJM189 | XIV | 763057 | 763066 | 763066 | rightend | YNR071C | 99.028 | 1029 | 10 | 0 | 1 | 1029 | 744839 | 743811 | 0 | 1845 |
| YJM189 | XV | 1 | 10 | 1034423 | leftend | YOL154W | 99.467 | 750 | 4 | 0 | 1 | 750 | 7132 | 7881 | 0 | 1363 |
| YJM189 | XV | 1032284 | 1032293 | 1034423 | middle |  |  |  |  |  |  |  |  |  |  |  |
| YJM189 | XV | 1032799 | 1032808 | 1034423 | middle |  |  |  |  |  |  |  |  |  |  |  |
| YJM189 | XV | 1033982 | 1033991 | 1034423 | middle |  |  |  |  |  |  |  |  |  |  |  |
| YJM189 | XV | 1034414 | 1034423 | 1034423 | rightend | YOR381W | 99.953 | 2136 | 1 | 0 | 1 | 2136 | 1004392 | 1006527 | 0 | 3940 |
| YJM189 | XVI | 1 | 10 | 558910 | leftend | YHL044W | 98.446 | 708 | 11 | 0 | 1 | 708 | 8541 | 9248 | 0 | 1247 |
| YJM189 | XVI | 555464 | 555473 | 558910 | middle |  |  |  |  |  |  |  |  |  |  |  |
| YJM189 | XVI | 555788 | 555797 | 558910 | middle |  |  |  |  |  |  |  |  |  |  |  |
| YJM189 | XVI | 557309 | 557318 | 558910 | middle |  |  |  |  |  |  |  |  |  |  |  |
| YJM189 | XVI | 558901 | 558910 | 558910 | rightend | YPR201W | 100 | 1215 | 0 | 0 | 1 | 1215 | 551856 | 553070 | 0 | 2244 |
| YJM193 | I | 1 | 10 | 191304 | leftend | YAL063C | 85.11 | 1632 | 239 | 4 | 1 | 1630 | 12359 | 10730 | 0 | 1664 |
| YJM193 | I | 191295 | 191304 | 191304 | rightend | YAR050W | 80.295 | 1558 | 301 | 6 | 76 | 1630 | 12284 | 10730 | 0 | 1171 |
| YJM193 | II | 1 | 10 | 790321 | leftend | YNCB0001W | 100 | 84 | 0 | 0 | 1 | 84 | 9790 | 9873 | 4.16E-38 | 156 |
| YJM193 | II | 1642 | 1651 | 790321 | middle |  |  |  |  |  |  |  |  |  |  |  |
| YJM193 | II | 788225 | 788234 | 790321 | middle |  |  |  |  |  |  |  |  |  |  |  |
| YJM193 | II | 789561 | 789570 | 790321 | middle |  |  |  |  |  |  |  |  |  |  |  |
| YJM193 | II | 790312 | 790321 | 790321 | rightend | YBR297W | 92.172 | 1418 | 95 | 12 | 1 | 1407 | 779703 | 781115 | 0 | 1989 |
| YJM193 | III | 1 | 10 | 301412 | leftend | YCL073C | 98.647 | 1848 | 25 | 0 | 1 | 1848 | 7307 | 5460 | 0 | 3275 |
| YJM193 | III | 300832 | 300841 | 301412 | middle |  |  |  |  |  |  |  |  |  |  |  |
| YJM193 | III | 301412 | 301412 | 301412 | rightend_complete | YCR101C | 99.818 | 549 | 1 | 0 | 1 | 549 | 296714 | 296166 | 0 | 1009 |
| YJM193 | IV | 1 | 10 | 1471877 | leftend | YDL243C | 94.848 | 990 | 49 | 2 | 1 | 990 | 17559 | 16572 | 0 | 1544 |
| YJM193 | IV | 1471868 | 1471877 | 1471877 | rightend | YDR541C | 99.324 | 1035 | 7 | 0 | 1 | 1035 | 1469241 | 1468207 | 0 | 1873 |
| YJM193 | V | 1 | 10 | 550723 | leftend | YEL070W | 99.337 | 1509 | 10 | 0 | 1 | 1509 | 13900 | 15408 | 0 | 2732 |
| YJM193 | V | 622 | 631 | 550723 | middle |  |  |  |  |  |  |  |  |  |  |  |
| YJM193 | V | 550714 | 550723 | 550723 | rightend | YER187W | 91.981 | 424 | 32 | 2 | 4 | 426 | 548204 | 548626 | 3.44E-169 | 593 |
| YJM193 | VI | 1 | 10 | 240516 | leftend | YFL051C | 98.23 | 452 | 8 | 0 | 1 | 452 | 7541 | 7090 | 0 | 791 |
| YJM193 | VI | 1173 | 1182 | 240516 | middle |  |  |  |  |  |  |  |  |  |  |  |
| YJM193 | VI | 1870 | 1879 | 240516 | middle |  |  |  |  |  |  |  |  |  |  |  |
| YJM193 | VI | 240507 | 240516 | 240516 | rightend | YFR055W | 100 | 1023 | 0 | 0 | 1 | 1023 | 234164 | 235186 | 0 | 1890 |
| YJM193 | VII | 1 | 10 | 1048145 | leftend | YGL258W | 97.585 | 621 | 15 | 0 | 1 | 621 | 5021 | 5641 | 0 | 1064 |
| YJM193 | VII | 599 | 608 | 1048145 | middle |  |  |  |  |  |  |  |  |  |  |  |
| YJM193 | VII | 1004 | 1013 | 1048145 | middle |  |  |  |  |  |  |  |  |  |  |  |
| YJM193 | VII | 1048136 | 1048145 | 1048145 | rightend | YGR287C | 98.814 | 1770 | 21 | 0 | 1 | 1770 | 1025159 | 1023390 | 0 | 3153 |
| YJM193 | VIII | 1 | 10 | 542301 | leftend | YHL044W | 98.446 | 708 | 11 | 0 | 1 | 708 | 13898 | 14605 | 0 | 1247 |
| YJM193 | VIII | 756 | 765 | 542301 | middle |  |  |  |  |  |  |  |  |  |  |  |
| YJM193 | VIII | 6697 | 6706 | 542301 | middle |  |  |  |  |  |  |  |  |  |  |  |
| YJM193 | VIII | 12565 | 12574 | 542301 | middle |  |  |  |  |  |  |  |  |  |  |  |
| YJM193 | VIII | 537852 | 537861 | 542301 | middle |  |  |  |  |  |  |  |  |  |  |  |
| YJM193 | VIII | 540579 | 540588 | 542301 | middle |  |  |  |  |  |  |  |  |  |  |  |
| YJM193 | VIII | 542292 | 542301 | 542301 | rightend | YHR211W | 82.854 | 939 | 150 | 7 | 1 | 932 | 530595 | 531529 | 0 | 832 |
| YJM193 | IX | 1 |  | 417389 | leftend_complete | YIL169C | 94.513 | 1294 | 69 | 2 | 520 | 1812 | 10322 | 9030 | 0 | 1995 |
| YJM193 | IX | 7507 | 7516 | 417389 | middle |  |  |  |  |  |  |  |  |  |  |  |
| YJM193 | IX | 417380 | 417389 | 417389 | rightend | YIR039C | 99.628 | 1614 | 6 | 0 | 1 | 1614 | 411570 | 409957 | 0 | 2948 |
| YJM193 | X | 1 | 10 | 701769 | leftend | YJL213W | 99.498 | 996 | 5 | 0 | 1 | 996 | 20279 | 21274 | 0 | 1812 |
| YJM193 | X | 840 | 849 | 701769 | middle |  |  |  |  |  |  |  |  |  |  |  |
| YJM193 | X | 1488 | 1497 | 701769 | middle |  |  |  |  |  |  |  |  |  |  |  |
| YJM193 | X | 1722 | 1731 | 701769 | middle |  |  |  |  |  |  |  |  |  |  |  |
| YJM193 | X | 699791 | 699800 | 701769 | middle |  |  |  |  |  |  |  |  |  |  |  |
| YJM193 | X | 700029 | 700038 | 701769 | middle |  |  |  |  |  |  |  |  |  |  |  |
| YJM193 | X | 701769 | 701769 | 701769 | rightend_complete | YJR152W | 99.51 | 1632 | 8 | 0 | 1 | 1632 | 691308 | 692939 | 0 | 2970 |
| YJM193 | XI | 1 | 10 | 653492 | leftend | YKL222C | 99.433 | 2118 | 12 | 0 | 1 | 2118 | 4007 | 1890 | 0 | 3845 |
| YJM193 | XI | 807 | 816 | 653492 | middle |  |  |  |  |  |  |  |  |  |  |  |
| YJM193 | XI | 653483 | 653492 | 653492 | rightend | YKR101W | 99.847 | 1965 | 3 | 0 | 1 | 1965 | 638557 | 640521 | 0 | 3613 |
| YJM193 | XII | 1 |  | 2381521 | leftend_complete | YLL062C | 99.897 | 975 | 1 | 0 | 1 | 975 | 7355 | 6381 | 0 | 1796 |
| YJM193 | XII | 1786 | 1795 | 2381521 | middle |  |  |  |  |  |  |  |  |  |  |  |
| YJM193 | XII | 2015 | 2024 | 2381521 | middle |  |  |  |  |  |  |  |  |  |  |  |
| YJM193 | XII | 2284 | 2293 | 2381521 | middle |  |  |  |  |  |  |  |  |  |  |  |
| YJM193 | XII | 2380190 | 2380199 | 2381521 | middle |  |  |  |  |  |  |  |  |  |  |  |
| YJM193 | XII | 2380731 | 2380740 | 2381521 | middle |  |  |  |  |  |  |  |  |  |  |  |
| YJM193 | XII | 2381512 | 2381521 | 2381521 | rightend | YLR460C | 94.071 | 1130 | 65 | 2 | 3 | 1131 | 2378530 | 2377402 | 0 | 1714 |
| YJM193 | XIII | 1 | 10 | 938063 | leftend | YML131W | 99.909 | 1098 | 1 | 0 | 1 | 1098 | 4559 | 5656 | 0 | 2023 |
| YJM193 | XIII | 926149 | 926158 | 938063 | middle |  |  |  |  |  |  |  |  |  |  |  |
| YJM193 | XIII | 936112 | 936121 | 938063 | middle |  |  |  |  |  |  |  |  |  |  |  |
| YJM193 | XIII | 938063 | 938063 | 938063 | rightend_complete | YMR321C | 99.042 | 313 | 3 | 0 | 6 | 318 | 921317 | 921005 | 1.15E-159 | 562 |
| YJM193 | XIV | 1 | 10 | 757397 | leftend | YNL332W | 99.413 | 1023 | 4 | 1 | 1 | 1023 | 6851 | 7871 | 0 | 1855 |
| YJM193 | XIV | 1745 | 1754 | 757397 | middle |  |  |  |  |  |  |  |  |  |  |  |
| YJM193 | XIV | 756379 | 756388 | 757397 | middle |  |  |  |  |  |  |  |  |  |  |  |
| YJM193 | XIV | 757034 | 757043 | 757397 | middle |  |  |  |  |  |  |  |  |  |  |  |
| YJM193 | XIV | 757397 | 757397 | 757397 | rightend_complete | YNR071C | 99.903 | 1029 | 1 | 0 | 1 | 1029 | 746092 | 745064 | 0 | 1895 |
| YJM193 | XV | 1 | 10 | 1038625 | leftend | YOL154W | 99.333 | 750 | 5 | 0 | 1 | 750 | 21865 | 22614 | 0 | 1358 |
| YJM193 | XV | 1037522 | 1037531 | 1038625 | middle |  |  |  |  |  |  |  |  |  |  |  |
| YJM193 | XV | 1038246 | 1038255 | 1038625 | middle |  |  |  |  |  |  |  |  |  |  |  |
| YJM193 | XV | 1038616 | 1038625 | 1038625 | rightend | YOR381W | 99.953 | 2136 | 1 | 0 | 1 | 2136 | 1013612 | 1015747 | 0 | 3940 |
| YJM193 | XVI | 1 | 10 | 905173 | leftend | YPL274W | 99.717 | 1764 | 5 | 0 | 1 | 1764 | 12819 | 14582 | 0 | 3230 |
| YJM193 | XVI | 260 | 269 | 905173 | middle |  |  |  |  |  |  |  |  |  |  |  |
| YJM193 | XVI | 1151 | 1160 | 905173 | middle |  |  |  |  |  |  |  |  |  |  |  |
| YJM193 | XVI | 2670 | 2679 | 905173 | middle |  |  |  |  |  |  |  |  |  |  |  |
| YJM193 | XVI | 4191 | 4200 | 905173 | middle |  |  |  |  |  |  |  |  |  |  |  |
| YJM193 | XVI | 905173 | 905173 | 905173 | rightend_complete | YPR201W | 99.753 | 1213 | 3 | 0 | 1 | 1213 | 901924 | 903136 | 0 | 2224 |
| YJM195 | I | 1 | 10 | 194776 | leftend | YAL063C | 86.8 | 1447 | 187 | 4 | 46 | 1490 | 14508 | 13064 | 0 | 1611 |
| YJM195 | I | 752 | 761 | 194776 | middle |  |  |  |  |  |  |  |  |  |  |  |
| YJM195 | I | 1443 | 1452 | 194776 | middle |  |  |  |  |  |  |  |  |  |  |  |
| YJM195 | I | 11880 | 11889 | 194776 | middle |  |  |  |  |  |  |  |  |  |  |  |
| YJM195 | I | 14570 | 14579 | 194776 | middle |  |  |  |  |  |  |  |  |  |  |  |
| YJM195 | I | 186405 | 186414 | 194776 | middle |  |  |  |  |  |  |  |  |  |  |  |
| YJM195 | I | 187147 | 187156 | 194776 | middle |  |  |  |  |  |  |  |  |  |  |  |
| YJM195 | I | 194767 | 194776 | 194776 | rightend | YAR050W | 83.523 | 1493 | 240 | 6 | 1 | 1490 | 14553 | 13064 | 0 | 1389 |
| YJM195 | II | 1 | 10 | 779243 | leftend | YNCB0001W | 100 | 84 | 0 | 0 | 1 | 84 | 1636 | 1719 | 4.16E-38 | 156 |
| YJM195 | II | 779234 | 779243 | 779243 | rightend | YBR297W | 97.925 | 1205 | 25 | 0 | 1 | 1205 | 775059 | 776263 | 0 | 2087 |
| YJM195 | III | 1 | 10 | 304773 | leftend | YCL073C | 99.134 | 1848 | 16 | 0 | 1 | 1848 | 3787 | 1940 | 0 | 3325 |
| YJM195 | III | 295975 | 295984 | 304773 | middle |  |  |  |  |  |  |  |  |  |  |  |
| YJM195 | III | 304258 | 304267 | 304773 | middle |  |  |  |  |  |  |  |  |  |  |  |
| YJM195 | III | 304764 | 304773 | 304773 | rightend | YCR101C | 98.364 | 550 | 8 | 1 | 1 | 549 | 291662 | 291113 | 0 | 965 |
| YJM195 | IV | 1 | 10 | 1483958 | leftend | YDL243C | 91.313 | 990 | 85 | 1 | 1 | 990 | 29011 | 28023 | 0 | 1351 |
| YJM195 | IV | 1472992 | 1473001 | 1483958 | middle |  |  |  |  |  |  |  |  |  |  |  |
| YJM195 | IV | 1474399 | 1474408 | 1483958 | middle |  |  |  |  |  |  |  |  |  |  |  |
| YJM195 | IV | 1483949 | 1483958 | 1483958 | rightend | YDR541C | 99.71 | 1035 | 3 | 0 | 1 | 1035 | 1469804 | 1468770 | 0 | 1895 |
| YJM195 | V | 1 | 10 | 562401 | leftend | YEL070W | 99.271 | 1509 | 11 | 0 | 1 | 1509 | 14036 | 15544 | 0 | 2726 |
| YJM195 | V | 506 | 515 | 562401 | middle |  |  |  |  |  |  |  |  |  |  |  |
| YJM195 | V | 558723 | 558732 | 562401 | middle |  |  |  |  |  |  |  |  |  |  |  |
| YJM195 | V | 562392 | 562401 | 562401 | rightend | YER187W | 97.418 | 426 | 11 | 0 | 1 | 426 | 551441 | 551866 | 0 | 726 |
| YJM195 | VI | 1 | 10 | 272013 | leftend | YFL051C | 96.044 | 455 | 15 | 2 | 1 | 452 | 37014 | 36560 | 0 | 737 |
| YJM195 | VI | 2456 | 2465 | 272013 | middle |  |  |  |  |  |  |  |  |  |  |  |
| YJM195 | VI | 272004 | 272013 | 272013 | rightend | YFR055W | 99.119 | 1022 | 9 | 0 | 1 | 1022 | 265735 | 266756 | 0 | 1838 |
| YJM195 | VII | 1 | 10 | 1052409 | leftend | YGL258W | 99.195 | 621 | 5 | 0 | 1 | 621 | 11329 | 11949 | 0 | 1120 |
| YJM195 | VII | 604 | 613 | 1052409 | middle |  |  |  |  |  |  |  |  |  |  |  |
| YJM195 | VII | 7134 | 7143 | 1052409 | middle |  |  |  |  |  |  |  |  |  |  |  |
| YJM195 | VII | 1049779 | 1049788 | 1052409 | middle |  |  |  |  |  |  |  |  |  |  |  |
| YJM195 | VII | 1051377 | 1051386 | 1052409 | middle |  |  |  |  |  |  |  |  |  |  |  |
| YJM195 | VII | 1052400 | 1052409 | 1052409 | rightend | YGR287C | 97.119 | 1770 | 51 | 0 | 1 | 1770 | 1039640 | 1037871 | 0 | 2987 |
| YJM195 | VIII | 1 | 10 | 537071 | leftend | YHL044W | 97.884 | 709 | 14 | 1 | 1 | 708 | 8509 | 9217 | 0 | 1225 |
| YJM195 | VIII | 7270 | 7279 | 537071 | middle |  |  |  |  |  |  |  |  |  |  |  |
| YJM195 | VIII | 513128 | 513137 | 537071 | middle |  |  |  |  |  |  |  |  |  |  |  |
| YJM195 | VIII | 525162 | 525171 | 537071 | middle |  |  |  |  |  |  |  |  |  |  |  |
| YJM195 | VIII | 532558 | 532567 | 537071 | middle |  |  |  |  |  |  |  |  |  |  |  |
| YJM195 | VIII | 533477 | 533486 | 537071 | middle |  |  |  |  |  |  |  |  |  |  |  |
| YJM195 | VIII | 535971 | 535980 | 537071 | middle |  |  |  |  |  |  |  |  |  |  |  |
| YJM195 | VIII | 537071 | 537071 | 537071 | rightend_complete | YHR211W | 84.455 | 1100 | 158 | 9 | 1 | 1092 | 512033 | 513127 | 0 | 1072 |
| YJM195 | IX | 1 |  | 416269 | leftend_complete | YIR039C | 99.69 | 1614 | 5 | 0 | 1 | 1614 | 7759 | 9372 | 0 | 2953 |
| YJM195 | IX | 414808 | 414817 | 416269 | middle |  |  |  |  |  |  |  |  |  |  |  |
| YJM195 | IX | 415435 | 415444 | 416269 | middle |  |  |  |  |  |  |  |  |  |  |  |
| YJM195 | IX | 416260 | 416269 | 416269 | rightend | YIL169C | 95.405 | 1284 | 53 | 4 | 508 | 1791 | 408365 | 409642 | 0 | 2039 |
| YJM195 | X | 1 | 10 | 706565 | leftend | YJL213W | 99.398 | 996 | 6 | 0 | 1 | 996 | 25087 | 26082 | 0 | 1807 |
| YJM195 | X | 823 | 832 | 706565 | middle |  |  |  |  |  |  |  |  |  |  |  |
| YJM195 | X | 1567 | 1576 | 706565 | middle |  |  |  |  |  |  |  |  |  |  |  |
| YJM195 | X | 2925 | 2934 | 706565 | middle |  |  |  |  |  |  |  |  |  |  |  |
| YJM195 | X | 3134 | 3143 | 706565 | middle |  |  |  |  |  |  |  |  |  |  |  |
| YJM195 | X | 706556 | 706565 | 706565 | rightend | YJR152W | 99.632 | 1632 | 6 | 0 | 1 | 1632 | 696079 | 697710 | 0 | 2981 |
| YJM195 | XI | 1 | 10 | 662573 | leftend | YKL222C | 99.622 | 2118 | 8 | 0 | 1 | 2118 | 3231 | 1114 | 0 | 3868 |
| YJM195 | XI | 662564 | 662573 | 662573 | rightend | YKR101W | 99.695 | 1965 | 6 | 0 | 1 | 1965 | 638124 | 640088 | 0 | 3596 |
| YJM195 | XII | 1 | 10 | 2068728 | leftend | YLL062C | 99.179 | 975 | 8 | 0 | 1 | 975 | 4089 | 3115 | 0 | 1757 |
| YJM195 | XII | 2067334 | 2067343 | 2068728 | middle |  |  |  |  |  |  |  |  |  |  |  |
| YJM195 | XII | 2068003 | 2068012 | 2068728 | middle |  |  |  |  |  |  |  |  |  |  |  |
| YJM195 | XII | 2068719 | 2068728 | 2068728 | rightend | YLR460C | 96.817 | 1131 | 36 | 0 | 1 | 1131 | 2065659 | 2064529 | 0 | 1890 |
| YJM195 | XIII | 1 | 10 | 898382 | leftend | YML131W | 99.362 | 1098 | 7 | 0 | 1 | 1098 | 3204 | 4301 | 0 | 1989 |
| YJM195 | XIII | 898373 | 898382 | 898382 | rightend | YMR321C | 99.361 | 313 | 2 | 0 | 6 | 318 | 890411 | 890099 | 2.48E-161 | 568 |
| YJM195 | XIV | 1 | 10 | 757944 | leftend | YNL332W | 99.022 | 1023 | 10 | 0 | 1 | 1023 | 5883 | 6905 | 0 | 1834 |
| YJM195 | XIV | 531 | 540 | 757944 | middle |  |  |  |  |  |  |  |  |  |  |  |
| YJM195 | XIV | 757935 | 757944 | 757944 | rightend | YNR071C | 99.514 | 1029 | 5 | 0 | 1 | 1029 | 747601 | 746573 | 0 | 1873 |
| YJM195 | XV | 1 | 10 | 1031706 | leftend | YOL154W | 99.467 | 750 | 4 | 0 | 1 | 750 | 5464 | 6213 | 0 | 1363 |
| YJM195 | XV | 1031697 | 1031706 | 1031706 | rightend | YOR381W | 98.642 | 2136 | 29 | 0 | 1 | 2136 | 1004925 | 1007060 | 0 | 3784 |
| YJM195 | XVI | 1 | 10 | 911915 | leftend | YPL274W | 98.413 | 1764 | 28 | 0 | 1 | 1764 | 17809 | 19572 | 0 | 3103 |
| YJM195 | XVI | 450 | 459 | 911915 | middle |  |  |  |  |  |  |  |  |  |  |  |
| YJM195 | XVI | 695 | 704 | 911915 | middle |  |  |  |  |  |  |  |  |  |  |  |
| YJM195 | XVI | 911906 | 911915 | 911915 | rightend | YPR201W | 99.588 | 1215 | 5 | 0 | 1 | 1215 | 908830 | 910044 | 0 | 2217 |
| YJM244 | I | 1 | 10 | 244174 | leftend | YAL063C | 94.132 | 1261 | 40 | 14 | 2715 | 3969 | 179556 | 180788 | 0 | 1888 |
| YJM244 | I | 244174 | 244174 | 244174 | rightend_complete | YAR050W | 91.947 | 1428 | 94 | 5 | 2844 | 4250 | 179051 | 180478 | 0 | 1980 |
| YJM244 | II | 1 | 10 | 786202 | leftend | YNCB0001W | 100 | 84 | 0 | 0 | 1 | 84 | 4460 | 4543 | 4.16E-38 | 156 |
| YJM244 | II | 784 | 793 | 786202 | middle |  |  |  |  |  |  |  |  |  |  |  |
| YJM244 | II | 2158 | 2167 | 786202 | middle |  |  |  |  |  |  |  |  |  |  |  |
| YJM244 | II | 3164 | 3173 | 786202 | middle |  |  |  |  |  |  |  |  |  |  |  |
| YJM244 | II | 786193 | 786202 | 786202 | rightend | YBR297W | 92.243 | 1418 | 94 | 12 | 1 | 1407 | 776631 | 778043 | 0 | 1995 |
| YJM244 | III | 1 | 10 | 309673 | leftend | YCL073C | 99.513 | 1848 | 9 | 0 | 1 | 1848 | 7686 | 5839 | 0 | 3363 |
| YJM244 | III | 309673 | 309673 | 309673 | rightend_complete | YCR101C | 99.818 | 549 | 1 | 0 | 1 | 549 | 296113 | 295565 | 0 | 1009 |
| YJM244 | IV | 1 | 10 | 1479069 | leftend | YDL243C | 95.051 | 990 | 49 | 0 | 1 | 990 | 16829 | 15840 | 0 | 1557 |
| YJM244 | IV | 1479060 | 1479069 | 1479069 | rightend | YDR541C | 99.42 | 1035 | 6 | 0 | 1 | 1035 | 1476222 | 1475188 | 0 | 1879 |
| YJM244 | V | 1 | 10 | 557001 | leftend | YEL070W | 99.271 | 1509 | 11 | 0 | 1 | 1509 | 13885 | 15393 | 0 | 2726 |
| YJM244 | V | 554621 | 554630 | 557001 | middle |  |  |  |  |  |  |  |  |  |  |  |
| YJM244 | V | 557001 | 557001 | 557001 | rightend_complete | YER187W | 91.981 | 424 | 32 | 2 | 4 | 426 | 548314 | 548736 | 3.44E-169 | 593 |
| YJM244 | VI | 1 | 10 | 234326 | leftend | YFL051C | 98.451 | 452 | 7 | 0 | 1 | 452 | 1711 | 1260 | 0 | 797 |
| YJM244 | VI | 234317 | 234326 | 234326 | rightend | YFR055W | 100 | 1023 | 0 | 0 | 1 | 1023 | 227992 | 229014 | 0 | 1890 |
| YJM244 | VII | 1 |  | 1048623 | leftend_complete | YGL258W | 97.585 | 621 | 15 | 0 | 1 | 621 | 11243 | 11863 | 0 | 1064 |
| YJM244 | VII | 1048614 | 1048623 | 1048623 | rightend | YGR287C | 98.927 | 1770 | 19 | 0 | 1 | 1770 | 1030551 | 1028782 | 0 | 3164 |
| YJM244 | VIII | 1 | 10 | 519491 | leftend | YHL044W | 98.446 | 708 | 11 | 0 | 1 | 708 | 1305 | 2012 | 0 | 1247 |
| YJM244 | VIII | 519482 | 519491 | 519491 | rightend | YHR211W | 92.228 | 2136 | 142 | 2 | 1096 | 3228 | 509120 | 511234 | 0 | 3003 |
| YJM244 | IX | 1 | 10 | 420786 | leftend | YIL169C | 84.476 | 2158 | 265 | 38 | 520 | 2618 | 19337 | 17191 | 0 | 2065 |
| YJM244 | IX | 420777 | 420786 | 420786 | rightend | YIR039C | 99.009 | 1614 | 16 | 0 | 1 | 1614 | 419122 | 417509 | 0 | 2892 |
| YJM244 | X | 1 | 10 | 723573 | leftend | YJL213W | 99.699 | 996 | 3 | 0 | 1 | 996 | 33400 | 34395 | 0 | 1823 |
| YJM244 | X | 501 | 510 | 723573 | middle |  |  |  |  |  |  |  |  |  |  |  |
| YJM244 | X | 1412 | 1421 | 723573 | middle |  |  |  |  |  |  |  |  |  |  |  |
| YJM244 | X | 719485 | 719494 | 723573 | middle |  |  |  |  |  |  |  |  |  |  |  |
| YJM244 | X | 719740 | 719749 | 723573 | middle |  |  |  |  |  |  |  |  |  |  |  |
| YJM244 | X | 721189 | 721198 | 723573 | middle |  |  |  |  |  |  |  |  |  |  |  |
| YJM244 | X | 723573 | 723573 | 723573 | rightend_complete | YJR152W | 99.816 | 1632 | 2 | 1 | 1 | 1632 | 703109 | 704739 | 0 | 2996 |
| YJM244 | XI | 1 | 10 | 644857 | leftend | YKL222C | 99.953 | 2118 | 1 | 0 | 1 | 2118 | 3694 | 1577 | 0 | 3906 |
| YJM244 | XI | 644424 | 644433 | 644857 | middle |  |  |  |  |  |  |  |  |  |  |  |
| YJM244 | XI | 644857 | 644857 | 644857 | rightend_complete | YKR101W | 99.796 | 1965 | 4 | 0 | 1 | 1965 | 637474 | 639438 | 0 | 3607 |
| YJM244 | XII | 1 | 10 | 2351735 | leftend | YLL062C | 100 | 975 | 0 | 0 | 1 | 975 | 9392 | 8418 | 0 | 1801 |
| YJM244 | XII | 1432 | 1441 | 2351735 | middle |  |  |  |  |  |  |  |  |  |  |  |
| YJM244 | XII | 4291 | 4300 | 2351735 | middle |  |  |  |  |  |  |  |  |  |  |  |
| YJM244 | XII | 4904 | 4913 | 2351735 | middle |  |  |  |  |  |  |  |  |  |  |  |
| YJM244 | XII | 5131 | 5140 | 2351735 | middle |  |  |  |  |  |  |  |  |  |  |  |
| YJM244 | XII | 2349880 | 2349889 | 2351735 | middle |  |  |  |  |  |  |  |  |  |  |  |
| YJM244 | XII | 2351726 | 2351735 | 2351735 | rightend | YLR460C | 94.248 | 1130 | 63 | 2 | 3 | 1131 | 2348128 | 2347000 | 0 | 1725 |
| YJM244 | XIII | 1 | 10 | 895854 | leftend | YML131W | 99.909 | 1098 | 1 | 0 | 1 | 1098 | 2308 | 3405 | 0 | 2023 |
| YJM244 | XIII | 894561 | 894570 | 895854 | middle |  |  |  |  |  |  |  |  |  |  |  |
| YJM244 | XIII | 895845 | 895854 | 895854 | rightend | YMR321C | 99.361 | 313 | 2 | 0 | 6 | 318 | 887009 | 886697 | 2.48E-161 | 568 |
| YJM244 | XIV | 1 | 10 | 749738 | leftend | YNL332W | 98.586 | 495 | 7 | 0 | 529 | 1023 | 11 | 505 | 0 | 876 |
| YJM244 | XIV | 747974 | 747983 | 749738 | middle |  |  |  |  |  |  |  |  |  |  |  |
| YJM244 | XIV | 748221 | 748230 | 749738 | middle |  |  |  |  |  |  |  |  |  |  |  |
| YJM244 | XIV | 749738 | 749738 | 749738 | rightend_complete | YNR071C | 100 | 1029 | 0 | 0 | 1 | 1029 | 737877 | 736849 | 0 | 1901 |
| YJM244 | XV | 1 | 10 | 1028613 | leftend | YOL154W | 99.333 | 750 | 5 | 0 | 1 | 750 | 4010 | 4759 | 0 | 1358 |
| YJM244 | XV | 1028604 | 1028613 | 1028613 | rightend | YOR381W | 99.906 | 2137 | 1 | 1 | 1 | 2136 | 1001273 | 1003409 | 0 | 3934 |
| YJM244 | XVI | 1 |  | 959488 | leftend_complete | YPL274W | 99.887 | 1764 | 2 | 0 | 1 | 1764 | 70270 | 72033 | 0 | 3247 |
| YJM244 | XVI | 959479 | 959488 | 959488 | rightend | YPR201W | 100 | 1215 | 0 | 0 | 1 | 1215 | 956003 | 957217 | 0 | 2244 |
| YJM248 | I | 1 | 10 | 190892 | leftend | YAL063C | 85.539 | 1632 | 232 | 4 | 1 | 1630 | 19755 | 18126 | 0 | 1703 |
| YJM248 | I | 486 | 495 | 190892 | middle |  |  |  |  |  |  |  |  |  |  |  |
| YJM248 | I | 190883 | 190892 | 190892 | rightend | YAR050W | 80.552 | 1558 | 297 | 6 | 76 | 1630 | 19680 | 18126 | 0 | 1194 |
| YJM248 | II | 1 | 10 | 781350 | leftend | YNCB0001W | 100 | 84 | 0 | 0 | 1 | 84 | 1471 | 1554 | 4.16E-38 | 156 |
| YJM248 | II | 1110 | 1119 | 781350 | middle |  |  |  |  |  |  |  |  |  |  |  |
| YJM248 | II | 780091 | 780100 | 781350 | middle |  |  |  |  |  |  |  |  |  |  |  |
| YJM248 | II | 781341 | 781350 | 781350 | rightend | YBR297W | 99.005 | 1407 | 14 | 0 | 1 | 1407 | 771293 | 772699 | 0 | 2521 |
| YJM248 | III | 1 | 10 | 308186 | leftend | YCL073C | 99.729 | 1848 | 5 | 0 | 1 | 1848 | 7305 | 5458 | 0 | 3386 |
| YJM248 | III | 3735 | 3744 | 308186 | middle |  |  |  |  |  |  |  |  |  |  |  |
| YJM248 | III | 307803 | 307812 | 308186 | middle |  |  |  |  |  |  |  |  |  |  |  |
| YJM248 | III | 308186 | 308186 | 308186 | rightend_complete | YCR100C | 24.828 | 297 | 194 | 10 | 7 | 291 | 295492 | 294656 | 1.55E-13 | 70.1 |
| YJM248 | IV | 1 | 10 | 1519390 | leftend | YDL243C | 87.26 | 989 | 122 | 4 | 1 | 987 | 34055 | 33069 | 0 | 1125 |
| YJM248 | IV | 1519381 | 1519390 | 1519390 | rightend | YDR541C | 86.293 | 1036 | 137 | 5 | 1 | 1033 | 1516948 | 1515915 | 0 | 1122 |
| YJM248 | V | 1 | 10 | 552499 | leftend | YEL070W | 93.174 | 1509 | 103 | 0 | 1 | 1509 | 16091 | 17599 | 0 | 2217 |
| YJM248 | V | 552490 | 552499 | 552499 | rightend | YER187W | 91.981 | 424 | 32 | 2 | 4 | 426 | 549927 | 550349 | 3.44E-169 | 593 |
| YJM248 | VI | 1 | 10 | 233154 | leftend | YFL051C | 98.451 | 452 | 7 | 0 | 1 | 452 | 1092 | 641 | 0 | 797 |
| YJM248 | VI | 233145 | 233154 | 233154 | rightend | YFR055W | 90.873 | 1019 | 93 | 0 | 1 | 1019 | 229001 | 230019 | 0 | 1367 |
| YJM248 | VII | 1 |  | 1035068 | leftend_complete | YGL258W | 97.424 | 621 | 16 | 0 | 1 | 621 | 11148 | 11768 | 0 | 1059 |
| YJM248 | VII | 1035059 | 1035068 | 1035068 | rightend | YGR287C | 99.2 | 1626 | 13 | 0 | 145 | 1770 | 1031579 | 1029954 | 0 | 2931 |
| YJM248 | VIII | 1 | 10 | 529918 | leftend | YHL044W | 98.305 | 708 | 12 | 0 | 1 | 708 | 1769 | 2476 | 0 | 1242 |
| YJM248 | VIII | 529526 | 529535 | 529918 | middle |  |  |  |  |  |  |  |  |  |  |  |
| YJM248 | VIII | 529918 | 529918 | 529918 | rightend_complete | YHR211W | 82.428 | 939 | 154 | 7 | 1 | 932 | 518065 | 518999 | 0 | 809 |
| YJM248 | IX | 1 | 10 | 412987 | leftend | YIL169C | 96.068 | 1526 | 53 | 5 | 394 | 1919 | 7513 | 5995 | 0 | 2479 |
| YJM248 | IX | 412978 | 412987 | 412987 | rightend | YIR039C | 99.009 | 1614 | 16 | 0 | 1 | 1614 | 406721 | 405108 | 0 | 2892 |
| YJM248 | X | 1 | 10 | 737477 | leftend | YJL213W | 98.795 | 996 | 12 | 0 | 1 | 996 | 36448 | 37443 | 0 | 1773 |
| YJM248 | X | 736761 | 736770 | 737477 | middle |  |  |  |  |  |  |  |  |  |  |  |
| YJM248 | X | 737477 | 737477 | 737477 | rightend_complete | YJR152W | 92.463 | 1632 | 123 | 0 | 1 | 1632 | 705385 | 707016 | 0 | 2333 |
| YJM248 | XI | 1 | 10 | 656935 | leftend | YKL222C | 99.433 | 2118 | 12 | 0 | 1 | 2118 | 3337 | 1220 | 0 | 3845 |
| YJM248 | XI | 656926 | 656935 | 656935 | rightend | YKR101W | 99.847 | 1965 | 3 | 0 | 1 | 1965 | 637335 | 639299 | 0 | 3613 |
| YJM248 | XII | 1 | 10 | 1460600 | leftend | YLL062C | 100 | 975 | 0 | 0 | 1 | 975 | 3932 | 2958 | 0 | 1801 |
| YJM248 | XII | 1459754 | 1459763 | 1460600 | middle |  |  |  |  |  |  |  |  |  |  |  |
| YJM248 | XII | 1460591 | 1460600 | 1460600 | rightend | YLR460C | 88.729 | 417 | 47 | 0 | 715 | 1131 | 1452907 | 1452491 | 3.46E-143 | 510 |
| YJM248 | XIII | 1 | 10 | 914445 | leftend | YML131W | 100 | 1098 | 0 | 0 | 1 | 1098 | 4019 | 5116 | 0 | 2028 |
| YJM248 | XIII | 404 | 413 | 914445 | middle |  |  |  |  |  |  |  |  |  |  |  |
| YJM248 | XIII | 1630 | 1639 | 914445 | middle |  |  |  |  |  |  |  |  |  |  |  |
| YJM248 | XIII | 914436 | 914445 | 914445 | rightend | YMR321C | 99.042 | 313 | 3 | 0 | 6 | 318 | 889834 | 889522 | 1.15E-159 | 562 |
| YJM248 | XIV | 1 | 10 | 738869 | leftend | YNL332W | 97.361 | 1023 | 27 | 0 | 1 | 1023 | 6485 | 7507 | 0 | 1740 |
| YJM248 | XIV | 630 | 639 | 738869 | middle |  |  |  |  |  |  |  |  |  |  |  |
| YJM248 | XIV | 1449 | 1458 | 738869 | middle |  |  |  |  |  |  |  |  |  |  |  |
| YJM248 | XIV | 738869 | 738869 | 738869 | rightend_complete | YNR069C | 100 | 1470 | 0 | 0 | 1 | 1470 | 736574 | 735105 | 0 | 2715 |
| YJM248 | XV | 1 | 10 | 1040479 | leftend | YOL154W | 99.867 | 750 | 1 | 0 | 1 | 750 | 34188 | 34937 | 0 | 1380 |
| YJM248 | XV | 11264 | 11273 | 1040479 | middle |  |  |  |  |  |  |  |  |  |  |  |
| YJM248 | XV | 1039898 | 1039907 | 1040479 | middle |  |  |  |  |  |  |  |  |  |  |  |
| YJM248 | XV | 1040470 | 1040479 | 1040479 | rightend | YOR381W | 94.239 | 2135 | 121 | 2 | 1 | 2134 | 1032580 | 1034713 | 0 | 3260 |
| YJM248 | XVI | 1 | 10 | 910967 | leftend | YPL274W | 99.83 | 1764 | 3 | 0 | 1 | 1764 | 20565 | 22328 | 0 | 3241 |
| YJM248 | XVI | 260 | 269 | 910967 | middle |  |  |  |  |  |  |  |  |  |  |  |
| YJM248 | XVI | 4665 | 4674 | 910967 | middle |  |  |  |  |  |  |  |  |  |  |  |
| YJM248 | XVI | 5411 | 5420 | 910967 | middle |  |  |  |  |  |  |  |  |  |  |  |
| YJM248 | XVI | 5624 | 5633 | 910967 | middle |  |  |  |  |  |  |  |  |  |  |  |
| YJM248 | XVI | 5839 | 5848 | 910967 | middle |  |  |  |  |  |  |  |  |  |  |  |
| YJM248 | XVI | 910372 | 910381 | 910967 | middle |  |  |  |  |  |  |  |  |  |  |  |
| YJM248 | XVI | 910958 | 910967 | 910967 | rightend | YPR201W | 100 | 1215 | 0 | 0 | 1 | 1215 | 907636 | 908850 | 0 | 2244 |
| YJM270 | I | 1 | 10 | 182207 | leftend | YAL063C | 91.156 | 1289 | 57 | 12 | 2715 | 3969 | 175782 | 177047 | 0 | 1696 |
| YJM270 | I | 175697 | 175706 | 182207 | middle |  |  |  |  |  |  |  |  |  |  |  |
| YJM270 | I | 182198 | 182207 | 182207 | rightend | YAR050W | 96.709 | 1489 | 43 | 3 | 3126 | 4614 | 175565 | 177047 | 0 | 2492 |
| YJM270 | II | 1 | 10 | 786155 | leftend | YNCB0001W | 100 | 84 | 0 | 0 | 1 | 84 | 2401 | 2484 | 4.16E-38 | 156 |
| YJM270 | II | 1488 | 1497 | 786155 | middle |  |  |  |  |  |  |  |  |  |  |  |
| YJM270 | II | 784352 | 784361 | 786155 | middle |  |  |  |  |  |  |  |  |  |  |  |
| YJM270 | II | 784904 | 784913 | 786155 | middle |  |  |  |  |  |  |  |  |  |  |  |
| YJM270 | II | 786146 | 786155 | 786155 | rightend | YBR297W | 99.076 | 1407 | 13 | 0 | 1 | 1407 | 774731 | 776137 | 0 | 2527 |
| YJM270 | III | 1 | 10 | 302739 | leftend | YCL073C | 99.729 | 1848 | 5 | 0 | 1 | 1848 | 9783 | 7936 | 0 | 3386 |
| YJM270 | III | 302730 | 302739 | 302739 | rightend | YCR101C | 99.818 | 549 | 1 | 0 | 1 | 549 | 298612 | 298064 | 0 | 1009 |
| YJM270 | IV | 1 | 10 | 1461502 | leftend | YDL243C | 95.152 | 990 | 48 | 0 | 1 | 990 | 8101 | 7112 | 0 | 1563 |
| YJM270 | IV | 1461493 | 1461502 | 1461502 | rightend | YDR541C | 99.517 | 1035 | 5 | 0 | 1 | 1035 | 1458689 | 1457655 | 0 | 1884 |
| YJM270 | V | 1 | 10 | 547750 | leftend | YEL070W | 98.94 | 1509 | 16 | 0 | 1 | 1509 | 13940 | 15448 | 0 | 2699 |
| YJM270 | V | 547741 | 547750 | 547750 | rightend | YER187W | 99.296 | 426 | 3 | 0 | 1 | 426 | 544473 | 544898 | 0 | 771 |
| YJM270 | VI | 1 | 10 | 238367 | leftend | YFL051C | 98.23 | 452 | 8 | 0 | 1 | 452 | 5739 | 5288 | 0 | 791 |
| YJM270 | VI | 238358 | 238367 | 238367 | rightend | YFR055W | 99.902 | 1022 | 1 | 0 | 1 | 1022 | 232088 | 233109 | 0 | 1882 |
| YJM270 | VII | 1 |  | 1051614 | leftend_complete | YGL258W | 97.585 | 621 | 15 | 0 | 1 | 621 | 11850 | 12470 | 0 | 1064 |
| YJM270 | VII | 1051605 | 1051614 | 1051614 | rightend | YGR287C | 98.927 | 1770 | 19 | 0 | 1 | 1770 | 1032037 | 1030268 | 0 | 3164 |
| YJM270 | VIII | 1 | 10 | 522716 | leftend | YHL044W | 98.446 | 708 | 11 | 0 | 1 | 708 | 8653 | 9360 | 0 | 1247 |
| YJM270 | VIII | 516892 | 516901 | 522716 | middle |  |  |  |  |  |  |  |  |  |  |  |
| YJM270 | VIII | 522707 | 522716 | 522716 | rightend | YHR211W | 91.69 | 2130 | 172 | 3 | 1096 | 3221 | 513727 | 515855 | 0 | 2948 |
| YJM270 | IX | 1 | 10 | 407357 | leftend | YIL169C | 93.638 | 1336 | 77 | 4 | 478 | 1812 | 2918 | 1590 | 0 | 1989 |
| YJM270 | IX | 407348 | 407357 | 407357 | rightend | YIR039C | 99.071 | 1614 | 15 | 0 | 1 | 1614 | 401305 | 399692 | 0 | 2898 |
| YJM270 | X | 1 | 10 | 716369 | leftend | YJL213W | 99.598 | 996 | 4 | 0 | 1 | 996 | 25682 | 26677 | 0 | 1818 |
| YJM270 | X | 716360 | 716369 | 716369 | rightend | YJR152W | 99.939 | 1632 | 1 | 0 | 1 | 1632 | 696394 | 698025 | 0 | 3009 |
| YJM270 | XI | 1 | 10 | 660021 | leftend | YKL222C | 99.433 | 2118 | 12 | 0 | 1 | 2118 | 3105 | 988 | 0 | 3845 |
| YJM270 | XI | 659244 | 659253 | 660021 | middle |  |  |  |  |  |  |  |  |  |  |  |
| YJM270 | XI | 660021 | 660021 | 660021 | rightend_complete | YKR101W | 99.847 | 1965 | 3 | 0 | 1 | 1965 | 638122 | 640086 | 0 | 3613 |
| YJM270 | XII | 1 | 10 | 1821718 | leftend | YLL062C | 100 | 975 | 0 | 0 | 1 | 975 | 3381 | 2407 | 0 | 1801 |
| YJM270 | XII | 247 | 256 | 1821718 | middle |  |  |  |  |  |  |  |  |  |  |  |
| YJM270 | XII | 780 | 789 | 1821718 | middle |  |  |  |  |  |  |  |  |  |  |  |
| YJM270 | XII | 1821709 | 1821718 | 1821718 | rightend | YLR460C | 94.425 | 1130 | 61 | 2 | 3 | 1131 | 1820049 | 1818921 | 0 | 1736 |
| YJM270 | XIII | 1 | 10 | 886254 | leftend | YML131W | 99.818 | 1098 | 2 | 0 | 1 | 1098 | 3704 | 4801 | 0 | 2017 |
| YJM270 | XIII | 886245 | 886254 | 886254 | rightend | YMR319C | 99.458 | 1659 | 9 | 0 | 1 | 1659 | 885603 | 883945 | 0 | 3014 |
| YJM270 | XIV | 1 | 10 | 740315 | leftend | YNL332W | 98.799 | 333 | 4 | 0 | 691 | 1023 | 11 | 343 | 1.17E-168 | 593 |
| YJM270 | XIV | 739414 | 739423 | 740315 | middle |  |  |  |  |  |  |  |  |  |  |  |
| YJM270 | XIV | 740107 | 740116 | 740315 | middle |  |  |  |  |  |  |  |  |  |  |  |
| YJM270 | XIV | 740315 | 740315 | 740315 | rightend_complete | YNR071C | 99.806 | 1029 | 2 | 0 | 1 | 1029 | 737873 | 736845 | 0 | 1890 |
| YJM270 | XV | 1 | 10 | 1095805 | leftend | YOL154W | 99.467 | 750 | 4 | 0 | 1 | 750 | 7209 | 7958 | 0 | 1363 |
| YJM270 | XV | 1095805 | 1095805 | 1095805 | rightend_complete | YOR381W | 99.906 | 2137 | 1 | 1 | 1 | 2136 | 1004695 | 1006831 | 0 | 3934 |
| YJM270 | XVI | 1 | 10 | 920514 | leftend | YPL274W | 99.943 | 1764 | 1 | 0 | 1 | 1764 | 24889 | 26652 | 0 | 3253 |
| YJM270 | XVI | 337 | 346 | 920514 | middle |  |  |  |  |  |  |  |  |  |  |  |
| YJM270 | XVI | 656 | 665 | 920514 | middle |  |  |  |  |  |  |  |  |  |  |  |
| YJM270 | XVI | 2144 | 2153 | 920514 | middle |  |  |  |  |  |  |  |  |  |  |  |
| YJM270 | XVI | 916609 | 916618 | 920514 | middle |  |  |  |  |  |  |  |  |  |  |  |
| YJM270 | XVI | 920514 | 920514 | 920514 | rightend_complete | YPR201W | 100 | 1215 | 0 | 0 | 1 | 1215 | 913884 | 915098 | 0 | 2244 |
| YJM271 | I | 1 | 10 | 190144 | leftend | YAL063C | 90.088 | 1019 | 51 | 14 | 2966 | 3962 | 10450 | 9460 | 0 | 1277 |
| YJM271 | I | 12317 | 12326 | 190144 | middle |  |  |  |  |  |  |  |  |  |  |  |
| YJM271 | I | 189649 | 189658 | 190144 | middle |  |  |  |  |  |  |  |  |  |  |  |
| YJM271 | I | 190135 | 190144 | 190144 | rightend | YAR050W | 81.166 | 1492 | 250 | 24 | 1960 | 3437 | 12314 | 10840 | 0 | 1170 |
| YJM271 | II | 1 | 10 | 786401 | leftend | YNCB0001W | 100 | 84 | 0 | 0 | 1 | 84 | 5971 | 6054 | 4.16E-38 | 156 |
| YJM271 | II | 917 | 926 | 786401 | middle |  |  |  |  |  |  |  |  |  |  |  |
| YJM271 | II | 1853 | 1862 | 786401 | middle |  |  |  |  |  |  |  |  |  |  |  |
| YJM271 | II | 2360 | 2369 | 786401 | middle |  |  |  |  |  |  |  |  |  |  |  |
| YJM271 | II | 2767 | 2776 | 786401 | middle |  |  |  |  |  |  |  |  |  |  |  |
| YJM271 | II | 3495 | 3504 | 786401 | middle |  |  |  |  |  |  |  |  |  |  |  |
| YJM271 | II | 4985 | 4994 | 786401 | middle |  |  |  |  |  |  |  |  |  |  |  |
| YJM271 | II | 786392 | 786401 | 786401 | rightend | YBR297W | 92.172 | 1418 | 95 | 12 | 1 | 1407 | 777802 | 779214 | 0 | 1989 |
| YJM271 | III | 1 | 10 | 304974 | leftend | YCL073C | 99.621 | 1848 | 7 | 0 | 1 | 1848 | 8166 | 6319 | 0 | 3374 |
| YJM271 | III | 302734 | 302743 | 304974 | middle |  |  |  |  |  |  |  |  |  |  |  |
| YJM271 | III | 303219 | 303228 | 304974 | middle |  |  |  |  |  |  |  |  |  |  |  |
| YJM271 | III | 304974 | 304974 | 304974 | rightend_complete | YCR101C | 99.455 | 550 | 2 | 1 | 1 | 549 | 298614 | 298065 | 0 | 998 |
| YJM271 | IV | 1 | 10 | 1468828 | leftend | YDL243C | 94.848 | 990 | 49 | 2 | 1 | 990 | 10514 | 9527 | 0 | 1544 |
| YJM271 | IV | 3051 | 3060 | 1468828 | middle |  |  |  |  |  |  |  |  |  |  |  |
| YJM271 | IV | 1468819 | 1468828 | 1468828 | rightend | YDR541C | 99.324 | 1035 | 7 | 0 | 1 | 1035 | 1465979 | 1464945 | 0 | 1873 |
| YJM271 | V | 1 | 10 | 553297 | leftend | YEL070W | 99.139 | 1509 | 13 | 0 | 1 | 1509 | 14718 | 16226 | 0 | 2715 |
| YJM271 | V | 905 | 914 | 553297 | middle |  |  |  |  |  |  |  |  |  |  |  |
| YJM271 | V | 1440 | 1449 | 553297 | middle |  |  |  |  |  |  |  |  |  |  |  |
| YJM271 | V | 553288 | 553297 | 553297 | rightend | YER187W | 91.981 | 424 | 32 | 2 | 4 | 426 | 550588 | 551010 | 3.44E-169 | 593 |
| YJM271 | VI | 1 | 10 | 236899 | leftend | YFL051C | 98.451 | 452 | 7 | 0 | 1 | 452 | 3346 | 2895 | 0 | 797 |
| YJM271 | VI | 937 | 946 | 236899 | middle |  |  |  |  |  |  |  |  |  |  |  |
| YJM271 | VI | 1449 | 1458 | 236899 | middle |  |  |  |  |  |  |  |  |  |  |  |
| YJM271 | VI | 2255 | 2264 | 236899 | middle |  |  |  |  |  |  |  |  |  |  |  |
| YJM271 | VI | 236890 | 236899 | 236899 | rightend | YFR055W | 100 | 1023 | 0 | 0 | 1 | 1023 | 230547 | 231569 | 0 | 1890 |
| YJM271 | VII | 1 | 10 | 1058504 | leftend | YGL258W | 99.517 | 621 | 3 | 0 | 1 | 621 | 10041 | 10661 | 0 | 1131 |
| YJM271 | VII | 5981 | 5990 | 1058504 | middle |  |  |  |  |  |  |  |  |  |  |  |
| YJM271 | VII | 1058495 | 1058504 | 1058504 | rightend | YGR287C | 98.927 | 1770 | 19 | 0 | 1 | 1770 | 1032156 | 1030387 | 0 | 3166 |
| YJM271 | VIII | 1 | 10 | 538911 | leftend | YHL044W | 98.446 | 708 | 11 | 0 | 1 | 708 | 9796 | 10503 | 0 | 1247 |
| YJM271 | VIII | 908 | 917 | 538911 | middle |  |  |  |  |  |  |  |  |  |  |  |
| YJM271 | VIII | 1551 | 1560 | 538911 | middle |  |  |  |  |  |  |  |  |  |  |  |
| YJM271 | VIII | 532229 | 532238 | 538911 | middle |  |  |  |  |  |  |  |  |  |  |  |
| YJM271 | VIII | 538242 | 538251 | 538911 | middle |  |  |  |  |  |  |  |  |  |  |  |
| YJM271 | VIII | 538911 | 538911 | 538911 | rightend_complete | YHR211W | 82.854 | 939 | 150 | 7 | 1 | 932 | 527262 | 528196 | 0 | 832 |
| YJM271 | IX | 1 | 10 | 404938 | leftend | YIL169C | 93.74 | 1214 | 74 | 2 | 600 | 1812 | 3484 | 2272 | 0 | 1820 |
| YJM271 | IX | 803 | 812 | 404938 | middle |  |  |  |  |  |  |  |  |  |  |  |
| YJM271 | IX | 1289 | 1298 | 404938 | middle |  |  |  |  |  |  |  |  |  |  |  |
| YJM271 | IX | 404929 | 404938 | 404938 | rightend | YIR039C | 99.009 | 1614 | 16 | 0 | 1 | 1614 | 402934 | 401321 | 0 | 2892 |
| YJM271 | X | 1 |  | 704063 | leftend_complete | YJL213W | 99.498 | 996 | 5 | 0 | 1 | 996 | 24387 | 25382 | 0 | 1812 |
| YJM271 | X | 1178 | 1187 | 704063 | middle |  |  |  |  |  |  |  |  |  |  |  |
| YJM271 | X | 701108 | 701117 | 704063 | middle |  |  |  |  |  |  |  |  |  |  |  |
| YJM271 | X | 701758 | 701767 | 704063 | middle |  |  |  |  |  |  |  |  |  |  |  |
| YJM271 | X | 704063 | 704063 | 704063 | rightend_complete | YJR152W | 99.939 | 1632 | 1 | 0 | 1 | 1632 | 693491 | 695122 | 0 | 3009 |
| YJM271 | XI | 1 | 10 | 657639 | leftend | YKL222C | 99.433 | 2118 | 12 | 0 | 1 | 2118 | 3976 | 1859 | 0 | 3845 |
| YJM271 | XI | 681 | 690 | 657639 | middle |  |  |  |  |  |  |  |  |  |  |  |
| YJM271 | XI | 657630 | 657639 | 657639 | rightend | YKR101W | 99.847 | 1965 | 3 | 0 | 1 | 1965 | 639015 | 640979 | 0 | 3613 |
| YJM271 | XII | 1 | 10 | 2880618 | leftend | YLL062C | 99.897 | 975 | 1 | 0 | 1 | 975 | 4973 | 3999 | 0 | 1796 |
| YJM271 | XII | 562 | 571 | 2880618 | middle |  |  |  |  |  |  |  |  |  |  |  |
| YJM271 | XII | 2878616 | 2878625 | 2880618 | middle |  |  |  |  |  |  |  |  |  |  |  |
| YJM271 | XII | 2880618 | 2880618 | 2880618 | rightend_complete | YLR460C | 94.336 | 1130 | 62 | 2 | 3 | 1131 | 2876956 | 2875828 | 0 | 1731 |
| YJM271 | XIII | 1 | 10 | 928275 | leftend | YML131W | 99.909 | 1098 | 1 | 0 | 1 | 1098 | 3422 | 4519 | 0 | 2023 |
| YJM271 | XIII | 928266 | 928275 | 928275 | rightend | YMR321C | 99.361 | 313 | 2 | 0 | 6 | 318 | 920187 | 919875 | 2.48E-161 | 568 |
| YJM271 | XIV | 1 | 10 | 755906 | leftend | YNL332W | 99.707 | 1023 | 3 | 0 | 1 | 1023 | 5237 | 6259 | 0 | 1873 |
| YJM271 | XIV | 755897 | 755906 | 755906 | rightend | YNR071C | 99.903 | 1029 | 1 | 0 | 1 | 1029 | 744360 | 743332 | 0 | 1895 |
| YJM271 | XV | 1 | 10 | 1127247 | leftend | YOL154W | 99.333 | 750 | 5 | 0 | 1 | 750 | 21853 | 22602 | 0 | 1358 |
| YJM271 | XV | 1127247 | 1127247 | 1127247 | rightend_complete | YOR381W | 99.953 | 2136 | 1 | 0 | 1 | 2136 | 1013273 | 1015408 | 0 | 3940 |
| YJM271 | XVI | 1 | 10 | 916088 | leftend | YPL274W | 99.773 | 1764 | 4 | 0 | 1 | 1764 | 23877 | 25640 | 0 | 3236 |
| YJM271 | XVI | 916079 | 916088 | 916088 | rightend | YPR201W | 99.753 | 1215 | 3 | 0 | 1 | 1215 | 913374 | 914588 | 0 | 2228 |
| YJM320 | I | 1 | 10 | 199443 | leftend | YAL063C | 93.471 | 1256 | 71 | 9 | 2715 | 3962 | 13655 | 12403 | 0 | 1855 |
| YJM320 | I | 3285 | 3294 | 199443 | middle |  |  |  |  |  |  |  |  |  |  |  |
| YJM320 | I | 199434 | 199443 | 199443 | rightend | YAR050W | 91.713 | 1436 | 96 | 7 | 2869 | 4282 | 14135 | 12701 | 0 | 1971 |
| YJM320 | II | 1 | 10 | 792155 | leftend | YNCB0001W | 100 | 84 | 0 | 0 | 1 | 84 | 10208 | 10291 | 4.16E-38 | 156 |
| YJM320 | II | 792146 | 792155 | 792155 | rightend | YBR297W | 84.483 | 1334 | 203 | 3 | 1 | 1332 | 783633 | 784964 | 0 | 1314 |
| YJM320 | III | 1 |  | 305540 | leftend_complete | YCL073C | 99.621 | 1848 | 7 | 0 | 1 | 1848 | 7693 | 5846 | 0 | 3374 |
| YJM320 | III | 567 | 576 | 305540 | middle |  |  |  |  |  |  |  |  |  |  |  |
| YJM320 | III | 305540 | 305540 | 305540 | rightend_complete | YCR101C | 97.636 | 550 | 12 | 1 | 1 | 549 | 298590 | 298041 | 0 | 942 |
| YJM320 | IV | 1 | 10 | 1470637 | leftend | YDL243C | 95.051 | 990 | 49 | 0 | 1 | 990 | 20007 | 19018 | 0 | 1557 |
| YJM320 | IV | 3196 | 3205 | 1470637 | middle |  |  |  |  |  |  |  |  |  |  |  |
| YJM320 | IV | 1470628 | 1470637 | 1470637 | rightend | YDR541C | 100 | 1035 | 0 | 0 | 1 | 1035 | 1464605 | 1463571 | 0 | 1912 |
| YJM320 | V | 1 | 10 | 549798 | leftend | YEL070W | 99.072 | 1509 | 14 | 0 | 1 | 1509 | 13797 | 15305 | 0 | 2710 |
| YJM320 | V | 745 | 754 | 549798 | middle |  |  |  |  |  |  |  |  |  |  |  |
| YJM320 | V | 549789 | 549798 | 549798 | rightend | YER187W | 99.296 | 426 | 3 | 0 | 1 | 426 | 547867 | 548292 | 0 | 771 |
| YJM320 | VI | 1 | 10 | 280381 | leftend | YFL051C | 98.451 | 452 | 7 | 0 | 1 | 452 | 45351 | 44900 | 0 | 797 |
| YJM320 | VI | 1612 | 1621 | 280381 | middle |  |  |  |  |  |  |  |  |  |  |  |
| YJM320 | VI | 2757 | 2766 | 280381 | middle |  |  |  |  |  |  |  |  |  |  |  |
| YJM320 | VI | 280372 | 280381 | 280381 | rightend | YFR055W | 99.902 | 1023 | 1 | 0 | 1 | 1023 | 274231 | 275253 | 0 | 1884 |
| YJM320 | VII | 1 | 10 | 1049840 | leftend | YGL258W | 99.356 | 621 | 4 | 0 | 1 | 621 | 2124 | 2744 | 0 | 1125 |
| YJM320 | VII | 1048269 | 1048278 | 1049840 | middle |  |  |  |  |  |  |  |  |  |  |  |
| YJM320 | VII | 1049280 | 1049289 | 1049840 | middle |  |  |  |  |  |  |  |  |  |  |  |
| YJM320 | VII | 1049840 | 1049840 | 1049840 | rightend_complete | YGR287C | 99.379 | 1770 | 11 | 0 | 1 | 1770 | 1024796 | 1023027 | 0 | 3208 |
| YJM320 | VIII | 1 | 10 | 511809 | leftend | YHL044W | 98.446 | 708 | 11 | 0 | 1 | 708 | 2061 | 2768 | 0 | 1247 |
| YJM320 | VIII | 926 | 935 | 511809 | middle |  |  |  |  |  |  |  |  |  |  |  |
| YJM320 | VIII | 509535 | 509544 | 511809 | middle |  |  |  |  |  |  |  |  |  |  |  |
| YJM320 | VIII | 510153 | 510162 | 511809 | middle |  |  |  |  |  |  |  |  |  |  |  |
| YJM320 | VIII | 510361 | 510370 | 511809 | middle |  |  |  |  |  |  |  |  |  |  |  |
| YJM320 | VIII | 510817 | 510826 | 511809 | middle |  |  |  |  |  |  |  |  |  |  |  |
| YJM320 | VIII | 511193 | 511202 | 511809 | middle |  |  |  |  |  |  |  |  |  |  |  |
| YJM320 | VIII | 511809 | 511809 | 511809 | rightend_complete | YHR211W | 89.379 | 2175 | 181 | 26 | 1096 | 3221 | 507174 | 509347 | 0 | 2691 |
| YJM320 | IX | 1 | 10 | 411256 | leftend | YIL169C | 96.539 | 1907 | 56 | 8 | 562 | 2466 | 8385 | 6487 | 0 | 3147 |
| YJM320 | IX | 1294 | 1303 | 411256 | middle |  |  |  |  |  |  |  |  |  |  |  |
| YJM320 | IX | 410076 | 410085 | 411256 | middle |  |  |  |  |  |  |  |  |  |  |  |
| YJM320 | IX | 410553 | 410562 | 411256 | middle |  |  |  |  |  |  |  |  |  |  |  |
| YJM320 | IX | 411256 | 411256 | 411256 | rightend_complete | YIR039C | 99.009 | 1614 | 16 | 0 | 1 | 1614 | 407986 | 406373 | 0 | 2892 |
| YJM320 | X | 1 |  | 713149 | leftend_complete | YJL213W | 99.699 | 996 | 3 | 0 | 1 | 996 | 24416 | 25411 | 0 | 1823 |
| YJM320 | X | 560 | 569 | 713149 | middle |  |  |  |  |  |  |  |  |  |  |  |
| YJM320 | X | 1723 | 1732 | 713149 | middle |  |  |  |  |  |  |  |  |  |  |  |
| YJM320 | X | 5757 | 5766 | 713149 | middle |  |  |  |  |  |  |  |  |  |  |  |
| YJM320 | X | 15068 | 15077 | 713149 | middle |  |  |  |  |  |  |  |  |  |  |  |
| YJM320 | X | 710141 | 710150 | 713149 | middle |  |  |  |  |  |  |  |  |  |  |  |
| YJM320 | X | 712085 | 712094 | 713149 | middle |  |  |  |  |  |  |  |  |  |  |  |
| YJM320 | X | 712581 | 712590 | 713149 | middle |  |  |  |  |  |  |  |  |  |  |  |
| YJM320 | X | 713149 | 713149 | 713149 | rightend_complete | YJR152W | 99.755 | 1632 | 3 | 1 | 1 | 1632 | 694493 | 696123 | 0 | 2990 |
| YJM320 | XI | 1 | 10 | 662564 | leftend | YKL222C | 99.387 | 2119 | 12 | 1 | 1 | 2118 | 4037 | 1919 | 0 | 3840 |
| YJM320 | XI | 819 | 828 | 662564 | middle |  |  |  |  |  |  |  |  |  |  |  |
| YJM320 | XI | 661903 | 661912 | 662564 | middle |  |  |  |  |  |  |  |  |  |  |  |
| YJM320 | XI | 662564 | 662564 | 662564 | rightend_complete | YKR101W | 99.796 | 1965 | 4 | 0 | 1 | 1965 | 638412 | 640376 | 0 | 3607 |
| YJM320 | XII | 1 | 10 | 2829437 | leftend | YLL062C | 100 | 975 | 0 | 0 | 1 | 975 | 4792 | 3818 | 0 | 1801 |
| YJM320 | XII | 613 | 622 | 2829437 | middle |  |  |  |  |  |  |  |  |  |  |  |
| YJM320 | XII | 853 | 862 | 2829437 | middle |  |  |  |  |  |  |  |  |  |  |  |
| YJM320 | XII | 2827674 | 2827683 | 2829437 | middle |  |  |  |  |  |  |  |  |  |  |  |
| YJM320 | XII | 2829437 | 2829437 | 2829437 | rightend_complete | YLR460C | 93.982 | 1130 | 66 | 2 | 3 | 1131 | 2827673 | 2826545 | 0 | 1709 |
| YJM320 | XIII | 1 | 10 | 912087 | leftend | YML131W | 100 | 1098 | 0 | 0 | 1 | 1098 | 9284 | 10381 | 0 | 2028 |
| YJM320 | XIII | 5293 | 5302 | 912087 | middle |  |  |  |  |  |  |  |  |  |  |  |
| YJM320 | XIII | 6110 | 6119 | 912087 | middle |  |  |  |  |  |  |  |  |  |  |  |
| YJM320 | XIII | 906261 | 906270 | 912087 | middle |  |  |  |  |  |  |  |  |  |  |  |
| YJM320 | XIII | 908220 | 908229 | 912087 | middle |  |  |  |  |  |  |  |  |  |  |  |
| YJM320 | XIII | 912078 | 912087 | 912087 | rightend | YMR321C | 99.361 | 313 | 2 | 0 | 6 | 318 | 895181 | 894869 | 2.48E-161 | 568 |
| YJM320 | XIV | 1 | 10 | 772041 | leftend | YNL332W | 97.849 | 1023 | 22 | 0 | 1 | 1023 | 17877 | 18899 | 0 | 1768 |
| YJM320 | XIV | 813 | 822 | 772041 | middle |  |  |  |  |  |  |  |  |  |  |  |
| YJM320 | XIV | 1823 | 1832 | 772041 | middle |  |  |  |  |  |  |  |  |  |  |  |
| YJM320 | XIV | 2890 | 2899 | 772041 | middle |  |  |  |  |  |  |  |  |  |  |  |
| YJM320 | XIV | 3108 | 3117 | 772041 | middle |  |  |  |  |  |  |  |  |  |  |  |
| YJM320 | XIV | 8948 | 8957 | 772041 | middle |  |  |  |  |  |  |  |  |  |  |  |
| YJM320 | XIV | 767653 | 767662 | 772041 | middle |  |  |  |  |  |  |  |  |  |  |  |
| YJM320 | XIV | 772041 | 772041 | 772041 | rightend_complete | YNR071C | 100 | 1029 | 0 | 0 | 1 | 1029 | 757656 | 756628 | 0 | 1901 |
| YJM320 | XV | 1 | 10 | 1035743 | leftend | YOL154W | 99.333 | 750 | 5 | 0 | 1 | 750 | 10870 | 11619 | 0 | 1358 |
| YJM320 | XV | 1035734 | 1035743 | 1035743 | rightend | YOR381W | 99.86 | 2136 | 3 | 0 | 1 | 2136 | 1009176 | 1011311 | 0 | 3928 |
| YJM320 | XVI | 1 | 10 | 906635 | leftend | YPL274W | 98.186 | 1764 | 32 | 0 | 1 | 1764 | 11556 | 13319 | 0 | 3081 |
| YJM320 | XVI | 264 | 273 | 906635 | middle |  |  |  |  |  |  |  |  |  |  |  |
| YJM320 | XVI | 497 | 506 | 906635 | middle |  |  |  |  |  |  |  |  |  |  |  |
| YJM320 | XVI | 739 | 748 | 906635 | middle |  |  |  |  |  |  |  |  |  |  |  |
| YJM320 | XVI | 1138 | 1147 | 906635 | middle |  |  |  |  |  |  |  |  |  |  |  |
| YJM320 | XVI | 2782 | 2791 | 906635 | middle |  |  |  |  |  |  |  |  |  |  |  |
| YJM320 | XVI | 3290 | 3299 | 906635 | middle |  |  |  |  |  |  |  |  |  |  |  |
| YJM320 | XVI | 906626 | 906635 | 906635 | rightend | YPR201W | 99.588 | 1215 | 5 | 0 | 1 | 1215 | 903071 | 904285 | 0 | 2217 |
| YJM326 | I | 1 | 10 | 201779 | leftend | YAL063C | 89.416 | 2173 | 224 | 5 | 1 | 2170 | 15065 | 12896 | 0 | 2734 |
| YJM326 | I | 10308 | 10317 | 201779 | middle |  |  |  |  |  |  |  |  |  |  |  |
| YJM326 | I | 11469 | 11478 | 201779 | middle |  |  |  |  |  |  |  |  |  |  |  |
| YJM326 | I | 12476 | 12485 | 201779 | middle |  |  |  |  |  |  |  |  |  |  |  |
| YJM326 | I | 190988 | 190997 | 201779 | middle |  |  |  |  |  |  |  |  |  |  |  |
| YJM326 | I | 191292 | 191301 | 201779 | middle |  |  |  |  |  |  |  |  |  |  |  |
| YJM326 | I | 194402 | 194411 | 201779 | middle |  |  |  |  |  |  |  |  |  |  |  |
| YJM326 | I | 195050 | 195059 | 201779 | middle |  |  |  |  |  |  |  |  |  |  |  |
| YJM326 | I | 201770 | 201779 | 201779 | rightend | YAR050W | 84.702 | 2131 | 314 | 11 | 46 | 2170 | 15020 | 12896 | 0 | 2119 |
| YJM326 | II | 1 | 10 | 1203367 | leftend | YNCB0001W | 100 | 84 | 0 | 0 | 1 | 84 | 3703 | 3786 | 4.16E-38 | 156 |
| YJM326 | II | 1754 | 1763 | 1203367 | middle |  |  |  |  |  |  |  |  |  |  |  |
| YJM326 | II | 1202668 | 1202677 | 1203367 | middle |  |  |  |  |  |  |  |  |  |  |  |
| YJM326 | II | 1203367 | 1203367 | 1203367 | rightend_complete | YBR297W | 98.294 | 1407 | 24 | 0 | 1 | 1407 | 776894 | 778300 | 0 | 2466 |
| YJM326 | III | 1 |  | 307967 | leftend_complete | YCL073C | 98.972 | 1848 | 19 | 0 | 1 | 1848 | 5262 | 3415 | 0 | 3308 |
| YJM326 | III | 307958 | 307967 | 307967 | rightend | YCR101C | 99.818 | 549 | 1 | 0 | 1 | 549 | 294692 | 294144 | 0 | 1009 |
| YJM326 | IV | 1 | 10 | 1464916 | leftend | YDL243C | 95.051 | 990 | 49 | 0 | 1 | 990 | 2618 | 1629 | 0 | 1557 |
| YJM326 | IV | 1464907 | 1464916 | 1464916 | rightend | YDR541C | 99.614 | 1035 | 4 | 0 | 1 | 1035 | 1462106 | 1461072 | 0 | 1890 |
| YJM326 | V | 1 | 10 | 544198 | leftend | YEL070W | 98.387 | 124 | 2 | 0 | 1 | 124 | 13583 | 13706 | 8.91E-56 | 219 |
| YJM326 | V | 543598 | 543607 | 544198 | middle |  |  |  |  |  |  |  |  |  |  |  |
| YJM326 | V | 544189 | 544198 | 544198 | rightend | YER186C | 98.588 | 921 | 12 | 1 | 1 | 921 | 540593 | 539674 | 0 | 1628 |
| YJM326 | VI | 1 | 10 | 238902 | leftend | YFL051C | 98.451 | 452 | 7 | 0 | 1 | 452 | 757 | 306 | 0 | 797 |
| YJM326 | VI | 238893 | 238902 | 238902 | rightend | YFR055W | 99.609 | 1022 | 4 | 0 | 1 | 1022 | 229178 | 230199 | 0 | 1866 |
| YJM326 | VII | 1 |  | 1045004 | leftend_complete | YGL258W | 99.356 | 621 | 4 | 0 | 1 | 621 | 3111 | 3731 | 0 | 1125 |
| YJM326 | VII | 691 | 700 | 1045004 | middle |  |  |  |  |  |  |  |  |  |  |  |
| YJM326 | VII | 1041885 | 1041894 | 1045004 | middle |  |  |  |  |  |  |  |  |  |  |  |
| YJM326 | VII | 1043175 | 1043184 | 1045004 | middle |  |  |  |  |  |  |  |  |  |  |  |
| YJM326 | VII | 1045004 | 1045004 | 1045004 | rightend_complete | YGR287C | 98.531 | 1770 | 26 | 0 | 1 | 1770 | 1024751 | 1022982 | 0 | 3125 |
| YJM326 | VIII | 1 |  | 543197 | leftend_complete | YHL044W | 98.446 | 708 | 11 | 0 | 1 | 708 | 10054 | 10761 | 0 | 1247 |
| YJM326 | VIII | 1153 | 1162 | 543197 | middle |  |  |  |  |  |  |  |  |  |  |  |
| YJM326 | VIII | 515847 | 515856 | 543197 | middle |  |  |  |  |  |  |  |  |  |  |  |
| YJM326 | VIII | 516535 | 516544 | 543197 | middle |  |  |  |  |  |  |  |  |  |  |  |
| YJM326 | VIII | 517493 | 517502 | 543197 | middle |  |  |  |  |  |  |  |  |  |  |  |
| YJM326 | VIII | 529140 | 529149 | 543197 | middle |  |  |  |  |  |  |  |  |  |  |  |
| YJM326 | VIII | 532721 | 532730 | 543197 | middle |  |  |  |  |  |  |  |  |  |  |  |
| YJM326 | VIII | 534278 | 534287 | 543197 | middle |  |  |  |  |  |  |  |  |  |  |  |
| YJM326 | VIII | 538280 | 538289 | 543197 | middle |  |  |  |  |  |  |  |  |  |  |  |
| YJM326 | VIII | 539024 | 539033 | 543197 | middle |  |  |  |  |  |  |  |  |  |  |  |
| YJM326 | VIII | 540008 | 540017 | 543197 | middle |  |  |  |  |  |  |  |  |  |  |  |
| YJM326 | VIII | 542095 | 542104 | 543197 | middle |  |  |  |  |  |  |  |  |  |  |  |
| YJM326 | VIII | 543197 | 543197 | 543197 | rightend_complete | YHR211W | 83.197 | 1226 | 188 | 14 | 1 | 1217 | 514622 | 515838 | 0 | 1107 |
| YJM326 | IX | 1 | 10 | 418497 | leftend | YIL169C | 98.18 | 2802 | 41 | 8 | 194 | 2988 | 9615 | 6817 | 0 | 4883 |
| YJM326 | IX | 418488 | 418497 | 418497 | rightend | YIR039C | 99.071 | 1614 | 15 | 0 | 1 | 1614 | 411888 | 410275 | 0 | 2898 |
| YJM326 | X | 1 |  | 728074 | leftend_complete | YJL213W | 100 | 996 | 0 | 0 | 1 | 996 | 33059 | 34054 | 0 | 1840 |
| YJM326 | X | 826 | 835 | 728074 | middle |  |  |  |  |  |  |  |  |  |  |  |
| YJM326 | X | 1078 | 1087 | 728074 | middle |  |  |  |  |  |  |  |  |  |  |  |
| YJM326 | X | 2919 | 2928 | 728074 | middle |  |  |  |  |  |  |  |  |  |  |  |
| YJM326 | X | 726838 | 726847 | 728074 | middle |  |  |  |  |  |  |  |  |  |  |  |
| YJM326 | X | 727279 | 727288 | 728074 | middle |  |  |  |  |  |  |  |  |  |  |  |
| YJM326 | X | 728074 | 728074 | 728074 | rightend_complete | YJR152W | 99.755 | 1632 | 3 | 1 | 1 | 1632 | 702880 | 704510 | 0 | 2990 |
| YJM326 | XI | 1 | 10 | 661084 | leftend | YKL222C | 99.481 | 2118 | 11 | 0 | 1 | 2118 | 3488 | 1371 | 0 | 3851 |
| YJM326 | XI | 658900 | 658909 | 661084 | middle |  |  |  |  |  |  |  |  |  |  |  |
| YJM326 | XI | 661075 | 661084 | 661084 | rightend | YKR101W | 99.796 | 1965 | 4 | 0 | 1 | 1965 | 637619 | 639583 | 0 | 3607 |
| YJM326 | XII | 1 |  | 1682946 | leftend_complete | YLL062C | 100 | 975 | 0 | 0 | 1 | 975 | 4865 | 3891 | 0 | 1801 |
| YJM326 | XII | 859 | 868 | 1682946 | middle |  |  |  |  |  |  |  |  |  |  |  |
| YJM326 | XII | 1682937 | 1682946 | 1682946 | rightend | YLR460C | 99.735 | 1131 | 3 | 0 | 1 | 1131 | 1680977 | 1679847 | 0 | 2073 |
| YJM326 | XIII | 1 | 10 | 910012 | leftend | YML131W | 100 | 1098 | 0 | 0 | 1 | 1098 | 2961 | 4058 | 0 | 2028 |
| YJM326 | XIII | 684 | 693 | 910012 | middle |  |  |  |  |  |  |  |  |  |  |  |
| YJM326 | XIII | 905705 | 905714 | 910012 | middle |  |  |  |  |  |  |  |  |  |  |  |
| YJM326 | XIII | 907070 | 907079 | 910012 | middle |  |  |  |  |  |  |  |  |  |  |  |
| YJM326 | XIII | 907912 | 907921 | 910012 | middle |  |  |  |  |  |  |  |  |  |  |  |
| YJM326 | XIII | 908885 | 908894 | 910012 | middle |  |  |  |  |  |  |  |  |  |  |  |
| YJM326 | XIII | 910003 | 910012 | 910012 | rightend | YMR321C | 99.361 | 313 | 2 | 0 | 6 | 318 | 888468 | 888156 | 2.48E-161 | 568 |
| YJM326 | XIV | 1 |  | 770304 | leftend_complete | YNL332W | 98.631 | 1023 | 14 | 0 | 1 | 1023 | 17559 | 18581 | 0 | 1812 |
| YJM326 | XIV | 651 | 660 | 770304 | middle |  |  |  |  |  |  |  |  |  |  |  |
| YJM326 | XIV | 1373 | 1382 | 770304 | middle |  |  |  |  |  |  |  |  |  |  |  |
| YJM326 | XIV | 768514 | 768523 | 770304 | middle |  |  |  |  |  |  |  |  |  |  |  |
| YJM326 | XIV | 769598 | 769607 | 770304 | middle |  |  |  |  |  |  |  |  |  |  |  |
| YJM326 | XIV | 770295 | 770304 | 770304 | rightend | YNR071C | 99.611 | 1029 | 4 | 0 | 1 | 1029 | 758432 | 757404 | 0 | 1879 |
| YJM326 | XV | 1 |  | 1036869 | leftend_complete | YOL154W | 99.6 | 750 | 3 | 0 | 1 | 750 | 12125 | 12874 | 0 | 1369 |
| YJM326 | XV | 1861 | 1870 | 1036869 | middle |  |  |  |  |  |  |  |  |  |  |  |
| YJM326 | XV | 1036860 | 1036869 | 1036869 | rightend | YOR381W | 98.642 | 2136 | 29 | 0 | 1 | 2136 | 1009640 | 1011775 | 0 | 3784 |
| YJM326 | XVI | 1 | 10 | 898582 | leftend | YPL274W | 98.98 | 1764 | 18 | 0 | 1 | 1764 | 5141 | 6904 | 0 | 3158 |
| YJM326 | XVI | 897663 | 897672 | 898582 | middle |  |  |  |  |  |  |  |  |  |  |  |
| YJM326 | XVI | 897890 | 897899 | 898582 | middle |  |  |  |  |  |  |  |  |  |  |  |
| YJM326 | XVI | 898573 | 898582 | 898582 | rightend | YPR201W | 99.588 | 1215 | 5 | 0 | 1 | 1215 | 895149 | 896363 | 0 | 2217 |
| YJM428 | I | 1 |  | 198698 | leftend_complete | YAL063C | 85.057 | 783 | 111 | 6 | 1 | 780 | 20810 | 20031 | 0 | 793 |
| YJM428 | I | 1642 | 1651 | 198698 | middle |  |  |  |  |  |  |  |  |  |  |  |
| YJM428 | I | 20019 | 20028 | 198698 | middle |  |  |  |  |  |  |  |  |  |  |  |
| YJM428 | I | 198344 | 198353 | 198698 | middle |  |  |  |  |  |  |  |  |  |  |  |
| YJM428 | I | 198689 | 198698 | 198698 | rightend | YAR042W | 99.552 | 3570 | 13 | 1 | 1 | 3567 | 180602 | 184171 | 0 | 6501 |
| YJM428 | II | 1 | 10 | 778388 | leftend | YNCB0001W | 100 | 84 | 0 | 0 | 1 | 84 | 2856 | 2939 | 4.16E-38 | 156 |
| YJM428 | II | 686 | 695 | 778388 | middle |  |  |  |  |  |  |  |  |  |  |  |
| YJM428 | II | 1315 | 1324 | 778388 | middle |  |  |  |  |  |  |  |  |  |  |  |
| YJM428 | II | 778379 | 778388 | 778388 | rightend | YBR297W | 97.759 | 1205 | 27 | 0 | 1 | 1205 | 776073 | 777277 | 0 | 2076 |
| YJM428 | III | 1 | 10 | 313701 | leftend | YCL073C | 99.405 | 1848 | 11 | 0 | 1 | 1848 | 10775 | 8928 | 0 | 3352 |
| YJM428 | III | 3477 | 3486 | 313701 | middle |  |  |  |  |  |  |  |  |  |  |  |
| YJM428 | III | 313251 | 313260 | 313701 | middle |  |  |  |  |  |  |  |  |  |  |  |
| YJM428 | III | 313701 | 313701 | 313701 | rightend_complete | YCR101C | 99.818 | 549 | 1 | 0 | 1 | 549 | 300090 | 299542 | 0 | 1009 |
| YJM428 | IV | 1 |  | 1482351 | leftend_complete | YDL243C | 95.152 | 990 | 48 | 0 | 1 | 990 | 20800 | 19811 | 0 | 1563 |
| YJM428 | IV | 2168 | 2177 | 1482351 | middle |  |  |  |  |  |  |  |  |  |  |  |
| YJM428 | IV | 2872 | 2881 | 1482351 | middle |  |  |  |  |  |  |  |  |  |  |  |
| YJM428 | IV | 4532 | 4541 | 1482351 | middle |  |  |  |  |  |  |  |  |  |  |  |
| YJM428 | IV | 1482351 | 1482351 | 1482351 | rightend_complete | YDR541C | 99.614 | 1035 | 4 | 0 | 1 | 1035 | 1477671 | 1476637 | 0 | 1890 |
| YJM428 | V | 1 | 10 | 550477 | leftend | YEL070W | 99.404 | 1509 | 9 | 0 | 1 | 1509 | 12832 | 14340 | 0 | 2737 |
| YJM428 | V | 550468 | 550477 | 550477 | rightend | YER187W | 100 | 426 | 0 | 0 | 1 | 426 | 546262 | 546687 | 0 | 787 |
| YJM428 | VI | 1 | 10 | 260539 | leftend | YFL051C | 100 | 483 | 0 | 0 | 1 | 483 | 22697 | 22215 | 0 | 893 |
| YJM428 | VI | 708 | 717 | 260539 | middle |  |  |  |  |  |  |  |  |  |  |  |
| YJM428 | VI | 1145 | 1154 | 260539 | middle |  |  |  |  |  |  |  |  |  |  |  |
| YJM428 | VI | 260530 | 260539 | 260539 | rightend | YFR055W | 99.609 | 1022 | 4 | 0 | 1 | 1022 | 251072 | 252093 | 0 | 1866 |
| YJM428 | VII | 1 | 10 | 1068650 | leftend | YGL258W | 97.585 | 621 | 15 | 0 | 1 | 621 | 11128 | 11748 | 0 | 1064 |
| YJM428 | VII | 688 | 697 | 1068650 | middle |  |  |  |  |  |  |  |  |  |  |  |
| YJM428 | VII | 1065491 | 1065500 | 1068650 | middle |  |  |  |  |  |  |  |  |  |  |  |
| YJM428 | VII | 1065710 | 1065719 | 1068650 | middle |  |  |  |  |  |  |  |  |  |  |  |
| YJM428 | VII | 1068650 | 1068650 | 1068650 | rightend_complete | YGR287C | 99.831 | 1770 | 3 | 0 | 1 | 1770 | 1033155 | 1031386 | 0 | 3253 |
| YJM428 | VIII | 1 | 10 | 533352 | leftend | YHL044W | 98.588 | 708 | 10 | 0 | 1 | 708 | 7615 | 8322 | 0 | 1253 |
| YJM428 | VIII | 6467 | 6476 | 533352 | middle |  |  |  |  |  |  |  |  |  |  |  |
| YJM428 | VIII | 517802 | 517811 | 533352 | middle |  |  |  |  |  |  |  |  |  |  |  |
| YJM428 | VIII | 529405 | 529414 | 533352 | middle |  |  |  |  |  |  |  |  |  |  |  |
| YJM428 | VIII | 532530 | 532539 | 533352 | middle |  |  |  |  |  |  |  |  |  |  |  |
| YJM428 | VIII | 533352 | 533352 | 533352 | rightend_complete | YHR211W | 90.971 | 1761 | 135 | 2 | 1096 | 2853 | 516062 | 517801 | 0 | 2350 |
| YJM428 | IX | 1 |  | 412430 | leftend_complete | YIL169C | 96.725 | 1374 | 39 | 2 | 478 | 1851 | 6527 | 5160 | 0 | 2283 |
| YJM428 | IX | 1298 | 1307 | 412430 | middle |  |  |  |  |  |  |  |  |  |  |  |
| YJM428 | IX | 2621 | 2630 | 412430 | middle |  |  |  |  |  |  |  |  |  |  |  |
| YJM428 | IX | 410098 | 410107 | 412430 | middle |  |  |  |  |  |  |  |  |  |  |  |
| YJM428 | IX | 412430 | 412430 | 412430 | rightend_complete | YIR039C | 99.628 | 1614 | 6 | 0 | 1 | 1614 | 408576 | 406963 | 0 | 2948 |
| YJM428 | X | 1 | 10 | 707221 | leftend | YJL213W | 99.598 | 996 | 4 | 0 | 1 | 996 | 25579 | 26574 | 0 | 1818 |
| YJM428 | X | 755 | 764 | 707185 | middle |  |  |  |  |  |  |  |  |  |  |  |
| YJM428 | X | 1542 | 1551 | 707185 | middle |  |  |  |  |  |  |  |  |  |  |  |
| YJM428 | X | 706270 | 706279 | 707185 | middle |  |  |  |  |  |  |  |  |  |  |  |
| YJM428 | X | 707212 | 707221 | 707221 | rightend | YJR152W | 99.816 | 1632 | 3 | 0 | 1 | 1632 | 696207 | 697838 | 0 | 2998 |
| YJM428 | XI | 1 | 10 | 658019 | leftend | YKL222C | 99.387 | 2119 | 12 | 1 | 1 | 2118 | 3374 | 1256 | 0 | 3840 |
| YJM428 | XI | 657547 | 657556 | 658019 | middle |  |  |  |  |  |  |  |  |  |  |  |
| YJM428 | XI | 658019 | 658019 | 658019 | rightend_complete | YKR101W | 99.644 | 1965 | 7 | 0 | 1 | 1965 | 637838 | 639802 | 0 | 3591 |
| YJM428 | XII | 1 | 10 | 1588125 | leftend | YLL062C | 100 | 975 | 0 | 0 | 1 | 975 | 4595 | 3621 | 0 | 1801 |
| YJM428 | XII | 575 | 584 | 1585907 | middle |  |  |  |  |  |  |  |  |  |  |  |
| YJM428 | XII | 1588116 | 1588125 | 1588125 | rightend | YLR460C | 93.894 | 1130 | 67 | 2 | 3 | 1131 | 1587208 | 1586080 | 0 | 1703 |
| YJM428 | XIII | 1 | 10 | 912560 | leftend | YML131W | 99.454 | 1098 | 6 | 0 | 1 | 1098 | 7143 | 8240 | 0 | 1995 |
| YJM428 | XIII | 592 | 601 | 912560 | middle |  |  |  |  |  |  |  |  |  |  |  |
| YJM428 | XIII | 3552 | 3561 | 912560 | middle |  |  |  |  |  |  |  |  |  |  |  |
| YJM428 | XIII | 3758 | 3767 | 912560 | middle |  |  |  |  |  |  |  |  |  |  |  |
| YJM428 | XIII | 4677 | 4686 | 912560 | middle |  |  |  |  |  |  |  |  |  |  |  |
| YJM428 | XIII | 898027 | 898036 | 912560 | middle |  |  |  |  |  |  |  |  |  |  |  |
| YJM428 | XIII | 908201 | 908210 | 912560 | middle |  |  |  |  |  |  |  |  |  |  |  |
| YJM428 | XIII | 912560 | 912560 | 912560 | rightend_complete | YMR321C | 99.361 | 313 | 2 | 0 | 6 | 318 | 892068 | 891756 | 2.48E-161 | 568 |
| YJM428 | XIV | 1 | 10 | 752722 | leftend | YNL332W | 99.316 | 1023 | 7 | 0 | 1 | 1023 | 1533 | 2555 | 0 | 1851 |
| YJM428 | XIV | 720 | 729 | 752722 | middle |  |  |  |  |  |  |  |  |  |  |  |
| YJM428 | XIV | 1182 | 1191 | 752722 | middle |  |  |  |  |  |  |  |  |  |  |  |
| YJM428 | XIV | 751739 | 751748 | 752722 | middle |  |  |  |  |  |  |  |  |  |  |  |
| YJM428 | XIV | 752722 | 752722 | 752722 | rightend_complete | YNR071C | 100 | 1029 | 0 | 0 | 1 | 1029 | 741268 | 740240 | 0 | 1901 |
| YJM428 | XV | 1 |  | 1059936 | leftend_complete | YOL154W | 99.867 | 750 | 1 | 0 | 1 | 750 | 39214 | 39963 | 0 | 1380 |
| YJM428 | XV | 5750 | 5759 | 1059936 | middle |  |  |  |  |  |  |  |  |  |  |  |
| YJM428 | XV | 1056458 | 1056467 | 1059936 | middle |  |  |  |  |  |  |  |  |  |  |  |
| YJM428 | XV | 1058830 | 1058839 | 1059936 | middle |  |  |  |  |  |  |  |  |  |  |  |
| YJM428 | XV | 1059285 | 1059294 | 1059936 | middle |  |  |  |  |  |  |  |  |  |  |  |
| YJM428 | XV | 1059927 | 1059936 | 1059936 | rightend | YOR381W | 99.813 | 2136 | 4 | 0 | 1 | 2136 | 1036284 | 1038419 | 0 | 3923 |
| YJM428 | XVI | 1 |  | 902641 | leftend_complete | YPL274W | 98.243 | 1764 | 31 | 0 | 1 | 1764 | 11464 | 13227 | 0 | 3086 |
| YJM428 | XVI | 1608 | 1617 | 902641 | middle |  |  |  |  |  |  |  |  |  |  |  |
| YJM428 | XVI | 2452 | 2461 | 902641 | middle |  |  |  |  |  |  |  |  |  |  |  |
| YJM428 | XVI | 2707 | 2716 | 902641 | middle |  |  |  |  |  |  |  |  |  |  |  |
| YJM428 | XVI | 3050 | 3059 | 902641 | middle |  |  |  |  |  |  |  |  |  |  |  |
| YJM428 | XVI | 4841 | 4850 | 902641 | middle |  |  |  |  |  |  |  |  |  |  |  |
| YJM428 | XVI | 902641 | 902641 | 902641 | rightend_complete | YPR201W | 99.588 | 1215 | 5 | 0 | 1 | 1215 | 899393 | 900607 | 0 | 2217 |
| YJM450 | I | 1 |  | 237361 | leftend_complete | YAL063C | 89.891 | 2572 | 260 | 0 | 1 | 2572 | 15539 | 12968 | 0 | 3310 |
| YJM450 | I | 1312 | 1321 | 237361 | middle |  |  |  |  |  |  |  |  |  |  |  |
| YJM450 | I | 212123 | 212132 | 237361 | middle |  |  |  |  |  |  |  |  |  |  |  |
| YJM450 | I | 220480 | 220489 | 237361 | middle |  |  |  |  |  |  |  |  |  |  |  |
| YJM450 | I | 221954 | 221963 | 237361 | middle |  |  |  |  |  |  |  |  |  |  |  |
| YJM450 | I | 237361 | 237361 | 237361 | rightend_complete | YAR050W | 85.551 | 2796 | 396 | 8 | 46 | 2837 | 15494 | 12703 | 0 | 2918 |
| YJM450 | II | 1 | 10 | 784721 | leftend | YNCB0001W | 100 | 84 | 0 | 0 | 1 | 84 | 2057 | 2140 | 4.16E-38 | 156 |
| YJM450 | II | 541 | 550 | 784721 | middle |  |  |  |  |  |  |  |  |  |  |  |
| YJM450 | II | 784712 | 784721 | 784721 | rightend | YBR297W | 84.483 | 1334 | 203 | 3 | 1 | 1332 | 776329 | 777660 | 0 | 1314 |
| YJM450 | III | 1 | 10 | 299729 | leftend | YCL073C | 98.972 | 1848 | 19 | 0 | 1 | 1848 | 7306 | 5459 | 0 | 3308 |
| YJM450 | III | 299720 | 299729 | 299729 | rightend | YCR101C | 98.182 | 550 | 9 | 1 | 1 | 549 | 295141 | 294592 | 0 | 959 |
| YJM450 | IV | 1 | 10 | 1472363 | leftend | YDL243C | 100 | 990 | 0 | 0 | 1 | 990 | 18267 | 17278 | 0 | 1829 |
| YJM450 | IV | 1471716 | 1471725 | 1472363 | middle |  |  |  |  |  |  |  |  |  |  |  |
| YJM450 | IV | 1472363 | 1472363 | 1472363 | rightend_complete | YDR541C | 99.42 | 1035 | 6 | 0 | 1 | 1035 | 1468893 | 1467859 | 0 | 1879 |
| YJM450 | V | 1 |  | 558721 | leftend_complete | YEL070W | 99.271 | 1509 | 11 | 0 | 1 | 1509 | 19150 | 20658 | 0 | 2726 |
| YJM450 | V | 4137 | 4146 | 558721 | middle |  |  |  |  |  |  |  |  |  |  |  |
| YJM450 | V | 556214 | 556223 | 558721 | middle |  |  |  |  |  |  |  |  |  |  |  |
| YJM450 | V | 558018 | 558027 | 558721 | middle |  |  |  |  |  |  |  |  |  |  |  |
| YJM450 | V | 558712 | 558721 | 558721 | rightend | YER187W | 91.981 | 424 | 32 | 2 | 4 | 426 | 552258 | 552680 | 3.44E-169 | 593 |
| YJM450 | VI | 1 | 10 | 242261 | leftend | YFL051C | 98.451 | 452 | 7 | 0 | 1 | 452 | 1527 | 1076 | 0 | 797 |
| YJM450 | VI | 242261 | 242261 | 242261 | rightend_complete | YFR055W | 88.496 | 1017 | 115 | 2 | 1 | 1016 | 229758 | 230773 | 0 | 1229 |
| YJM450 | VII | 1 | 10 | 1036509 | leftend | YGL258W | 99.356 | 621 | 4 | 0 | 1 | 621 | 2384 | 3004 | 0 | 1125 |
| YJM450 | VII | 1036500 | 1036509 | 1036509 | rightend | YGR287C | 98.588 | 1770 | 25 | 0 | 1 | 1770 | 1024812 | 1023043 | 0 | 3131 |
| YJM450 | VIII | 1 | 10 | 511515 | leftend | YHL044W | 98.446 | 708 | 11 | 0 | 1 | 708 | 1165 | 1872 | 0 | 1247 |
| YJM450 | VIII | 511506 | 511515 | 511515 | rightend | YHR211W | 91.582 | 1960 | 141 | 2 | 1096 | 3052 | 509567 | 511505 | 0 | 2684 |
| YJM450 | IX | 1 | 10 | 412858 | leftend | YIL169C | 99.465 | 2989 | 12 | 4 | 1 | 2988 | 13470 | 10485 | 0 | 5428 |
| YJM450 | IX | 1004 | 1013 | 412858 | middle |  |  |  |  |  |  |  |  |  |  |  |
| YJM450 | IX | 1167 | 1176 | 412858 | middle |  |  |  |  |  |  |  |  |  |  |  |
| YJM450 | IX | 1848 | 1857 | 412858 | middle |  |  |  |  |  |  |  |  |  |  |  |
| YJM450 | IX | 2829 | 2838 | 412858 | middle |  |  |  |  |  |  |  |  |  |  |  |
| YJM450 | IX | 412849 | 412858 | 412858 | rightend | YIR039C | 98.947 | 1614 | 17 | 0 | 1 | 1614 | 411374 | 409761 | 0 | 2887 |
| YJM450 | X | 1 |  | 725638 | leftend_complete | YJL213W | 99.699 | 996 | 3 | 0 | 1 | 996 | 36587 | 37582 | 0 | 1823 |
| YJM450 | X | 724788 | 724797 | 725638 | middle |  |  |  |  |  |  |  |  |  |  |  |
| YJM450 | X | 725629 | 725638 | 725638 | rightend | YJR152W | 99.816 | 1632 | 3 | 0 | 1 | 1632 | 706551 | 708182 | 0 | 2998 |
| YJM450 | XI | 1 | 10 | 666110 | leftend | YKL222C | 100 | 2118 | 0 | 0 | 1 | 2118 | 7706 | 5589 | 0 | 3912 |
| YJM450 | XI | 908 | 917 | 666110 | middle |  |  |  |  |  |  |  |  |  |  |  |
| YJM450 | XI | 1717 | 1726 | 666110 | middle |  |  |  |  |  |  |  |  |  |  |  |
| YJM450 | XI | 2392 | 2401 | 666110 | middle |  |  |  |  |  |  |  |  |  |  |  |
| YJM450 | XI | 666101 | 666110 | 666110 | rightend | YKR101W | 99.695 | 1965 | 6 | 0 | 1 | 1965 | 641735 | 643699 | 0 | 3596 |
| YJM450 | XII | 1 |  | 1717464 | leftend_complete | YLL062C | 100 | 975 | 0 | 0 | 1 | 975 | 10809 | 9835 | 0 | 1801 |
| YJM450 | XII | 1716688 | 1716697 | 1717464 | middle |  |  |  |  |  |  |  |  |  |  |  |
| YJM450 | XII | 1717464 | 1717464 | 1717464 | rightend_complete | YLR460C | 99.823 | 1131 | 2 | 0 | 1 | 1131 | 1715497 | 1714367 | 0 | 2078 |
| YJM450 | XIII | 1 | 10 | 902422 | leftend | YML131W | 100 | 1098 | 0 | 0 | 1 | 1098 | 2362 | 3459 | 0 | 2028 |
| YJM450 | XIII | 897123 | 897132 | 902422 | middle |  |  |  |  |  |  |  |  |  |  |  |
| YJM450 | XIII | 902413 | 902422 | 902422 | rightend | YMR321C | 99.361 | 313 | 2 | 0 | 6 | 318 | 887618 | 887306 | 2.48E-161 | 568 |
| YJM450 | XIV | 1 | 10 | 770761 | leftend | YNL332W | 92.958 | 71 | 5 | 0 | 1 | 71 | 18751 | 18821 | 2.4E-21 | 104 |
| YJM450 | XIV | 485 | 494 | 770761 | middle |  |  |  |  |  |  |  |  |  |  |  |
| YJM450 | XIV | 768619 | 768628 | 770761 | middle |  |  |  |  |  |  |  |  |  |  |  |
| YJM450 | XIV | 770752 | 770761 | 770761 | rightend | YNR071C | 100 | 1029 | 0 | 0 | 1 | 1029 | 758141 | 757113 | 0 | 1901 |
| YJM450 | XV | 1 | 10 | 1047806 | leftend | YOL154W | 99.333 | 750 | 5 | 0 | 1 | 750 | 23121 | 23870 | 0 | 1358 |
| YJM450 | XV | 1039819 | 1039828 | 1047806 | middle |  |  |  |  |  |  |  |  |  |  |  |
| YJM450 | XV | 1045864 | 1045873 | 1047806 | middle |  |  |  |  |  |  |  |  |  |  |  |
| YJM450 | XV | 1046139 | 1046148 | 1047806 | middle |  |  |  |  |  |  |  |  |  |  |  |
| YJM450 | XV | 1047100 | 1047109 | 1047806 | middle |  |  |  |  |  |  |  |  |  |  |  |
| YJM450 | XV | 1047797 | 1047806 | 1047806 | rightend | YOR381W | 99.064 | 2136 | 20 | 0 | 1 | 2136 | 1020870 | 1023005 | 0 | 3834 |
| YJM450 | XVI | 1 |  | 913650 | leftend_complete | YPL274W | 99.943 | 1764 | 1 | 0 | 1 | 1764 | 17824 | 19587 | 0 | 3253 |
| YJM450 | XVI | 1241 | 1250 | 913650 | middle |  |  |  |  |  |  |  |  |  |  |  |
| YJM450 | XVI | 5062 | 5071 | 913650 | middle |  |  |  |  |  |  |  |  |  |  |  |
| YJM450 | XVI | 11570 | 11579 | 913650 | middle |  |  |  |  |  |  |  |  |  |  |  |
| YJM450 | XVI | 11867 | 11876 | 913650 | middle |  |  |  |  |  |  |  |  |  |  |  |
| YJM450 | XVI | 913641 | 913650 | 913650 | rightend | YPR201W | 99.588 | 1215 | 5 | 0 | 1 | 1215 | 910176 | 911390 | 0 | 2217 |
| YJM451 | I | 1 | 10 | 216633 | leftend | YAL063C | 84.368 | 2175 | 319 | 16 | 1 | 2170 | 13043 | 10885 | 0 | 2132 |
| YJM451 | I | 11305 | 11314 | 216633 | middle |  |  |  |  |  |  |  |  |  |  |  |
| YJM451 | I | 194850 | 194859 | 216633 | middle |  |  |  |  |  |  |  |  |  |  |  |
| YJM451 | I | 212610 | 212619 | 216633 | middle |  |  |  |  |  |  |  |  |  |  |  |
| YJM451 | I | 216624 | 216633 | 216633 | rightend | YAR050W | 80.628 | 2101 | 384 | 18 | 76 | 2170 | 12968 | 10885 | 0 | 1622 |
| YJM451 | II | 1 |  | 782976 | leftend_complete | YNCB0001W | 100 | 84 | 0 | 0 | 1 | 84 | 5369 | 5452 | 4.16E-38 | 156 |
| YJM451 | II | 3777 | 3786 | 782976 | middle |  |  |  |  |  |  |  |  |  |  |  |
| YJM451 | II | 782967 | 782976 | 782976 | rightend | YBR297W | 98.008 | 1205 | 24 | 0 | 1 | 1205 | 778776 | 779980 | 0 | 2093 |
| YJM451 | III | 1 |  | 303587 | leftend_complete | YCL073C | 99.026 | 1848 | 18 | 0 | 1 | 1848 | 7984 | 6137 | 0 | 3314 |
| YJM451 | III | 303578 | 303587 | 303587 | rightend | YCR101C | 92.021 | 564 | 28 | 12 | 1 | 549 | 298891 | 298330 | 0 | 776 |
| YJM451 | IV | 1 | 10 | 1474961 | leftend | YDL243C | 93.769 | 979 | 61 | 0 | 1 | 979 | 22865 | 21887 | 0 | 1471 |
| YJM451 | IV | 1138 | 1147 | 1474961 | middle |  |  |  |  |  |  |  |  |  |  |  |
| YJM451 | IV | 1474952 | 1474961 | 1474961 | rightend | YDR541C | 99.324 | 1035 | 7 | 0 | 1 | 1035 | 1472193 | 1471159 | 0 | 1873 |
| YJM451 | V | 1 | 10 | 550548 | leftend | YEL070W | 99.072 | 1509 | 14 | 0 | 1 | 1509 | 13275 | 14783 | 0 | 2710 |
| YJM451 | V | 549164 | 549173 | 550548 | middle |  |  |  |  |  |  |  |  |  |  |  |
| YJM451 | V | 550539 | 550548 | 550548 | rightend | YER187W | 100 | 426 | 0 | 0 | 1 | 426 | 544738 | 545163 | 0 | 787 |
| YJM451 | VI | 1 | 10 | 258666 | leftend | YFL051C | 100 | 483 | 0 | 0 | 1 | 483 | 20676 | 20194 | 0 | 893 |
| YJM451 | VI | 258657 | 258666 | 258666 | rightend | YFR055W | 99.511 | 1022 | 5 | 0 | 1 | 1022 | 249082 | 250103 | 0 | 1860 |
| YJM451 | VII | 1 |  | 1048876 | leftend_complete | YGL258W | 97.585 | 621 | 15 | 0 | 1 | 621 | 10943 | 11563 | 0 | 1064 |
| YJM451 | VII | 1041592 | 1041601 | 1048876 | middle |  |  |  |  |  |  |  |  |  |  |  |
| YJM451 | VII | 1044134 | 1044143 | 1048876 | middle |  |  |  |  |  |  |  |  |  |  |  |
| YJM451 | VII | 1044823 | 1044832 | 1048876 | middle |  |  |  |  |  |  |  |  |  |  |  |
| YJM451 | VII | 1045260 | 1045269 | 1048876 | middle |  |  |  |  |  |  |  |  |  |  |  |
| YJM451 | VII | 1046844 | 1046853 | 1048876 | middle |  |  |  |  |  |  |  |  |  |  |  |
| YJM451 | VII | 1048867 | 1048876 | 1048876 | rightend | YGR287C | 99.209 | 1770 | 14 | 0 | 1 | 1770 | 1029743 | 1027974 | 0 | 3192 |
| YJM451 | VIII | 1 | 10 | 530431 | leftend | YHL044W | 100 | 708 | 0 | 0 | 1 | 708 | 10657 | 11364 | 0 | 1308 |
| YJM451 | VIII | 909 | 918 | 530431 | middle |  |  |  |  |  |  |  |  |  |  |  |
| YJM451 | VIII | 1535 | 1544 | 530431 | middle |  |  |  |  |  |  |  |  |  |  |  |
| YJM451 | VIII | 516711 | 516720 | 530431 | middle |  |  |  |  |  |  |  |  |  |  |  |
| YJM451 | VIII | 524524 | 524533 | 530431 | middle |  |  |  |  |  |  |  |  |  |  |  |
| YJM451 | VIII | 528487 | 528496 | 530431 | middle |  |  |  |  |  |  |  |  |  |  |  |
| YJM451 | VIII | 529982 | 529991 | 530431 | middle |  |  |  |  |  |  |  |  |  |  |  |
| YJM451 | VIII | 530431 | 530431 | 530431 | rightend_complete | YHR211W | 91.892 | 1332 | 84 | 2 | 1590 | 2918 | 516721 | 518031 | 0 | 1840 |
| YJM451 | IX | 1 |  | 417778 | leftend_complete | YIL169C | 98.18 | 2802 | 41 | 8 | 194 | 2988 | 11367 | 8569 | 0 | 4883 |
| YJM451 | IX | 996 | 1005 | 417778 | middle |  |  |  |  |  |  |  |  |  |  |  |
| YJM451 | IX | 2294 | 2303 | 417778 | middle |  |  |  |  |  |  |  |  |  |  |  |
| YJM451 | IX | 3526 | 3535 | 417778 | middle |  |  |  |  |  |  |  |  |  |  |  |
| YJM451 | IX | 416470 | 416479 | 417778 | middle |  |  |  |  |  |  |  |  |  |  |  |
| YJM451 | IX | 417778 | 417778 | 417778 | rightend_complete | YIR039C | 99.504 | 1614 | 5 | 1 | 1 | 1614 | 410509 | 408899 | 0 | 2933 |
| YJM451 | X | 1 |  | 739929 | leftend_complete | YJL213W | 99.699 | 996 | 3 | 0 | 1 | 996 | 48607 | 49602 | 0 | 1823 |
| YJM451 | X | 636 | 645 | 739929 | middle |  |  |  |  |  |  |  |  |  |  |  |
| YJM451 | X | 1385 | 1394 | 739929 | middle |  |  |  |  |  |  |  |  |  |  |  |
| YJM451 | X | 2487 | 2496 | 739929 | middle |  |  |  |  |  |  |  |  |  |  |  |
| YJM451 | X | 739225 | 739234 | 739929 | middle |  |  |  |  |  |  |  |  |  |  |  |
| YJM451 | X | 739920 | 739929 | 739929 | rightend | YJR152W | 99.571 | 1632 | 7 | 0 | 1 | 1632 | 726242 | 727873 | 0 | 2976 |
| YJM451 | XI | 1 |  | 668508 | leftend_complete | YKL222C | 100 | 2118 | 0 | 0 | 1 | 2118 | 5311 | 3194 | 0 | 3912 |
| YJM451 | XI | 819 | 828 | 668508 | middle |  |  |  |  |  |  |  |  |  |  |  |
| YJM451 | XI | 1725 | 1734 | 668508 | middle |  |  |  |  |  |  |  |  |  |  |  |
| YJM451 | XI | 668499 | 668508 | 668508 | rightend | YKR101W | 99.695 | 1965 | 6 | 0 | 1 | 1965 | 640352 | 642316 | 0 | 3596 |
| YJM451 | XII | 1 | 10 | 1673785 | leftend | YLL062C | 100 | 975 | 0 | 0 | 1 | 975 | 5571 | 4597 | 0 | 1801 |
| YJM451 | XII | 1618 | 1627 | 1673785 | middle |  |  |  |  |  |  |  |  |  |  |  |
| YJM451 | XII | 1672047 | 1672056 | 1673785 | middle |  |  |  |  |  |  |  |  |  |  |  |
| YJM451 | XII | 1673785 | 1673785 | 1673785 | rightend_complete | YLR460C | 94.69 | 1130 | 58 | 2 | 3 | 1131 | 1669786 | 1668658 | 0 | 1753 |
| YJM451 | XIII | 1 | 10 | 895236 | leftend | YML131W | 98.816 | 1098 | 13 | 0 | 1 | 1098 | 4465 | 5562 | 0 | 1956 |
| YJM451 | XIII | 1639 | 1648 | 895236 | middle |  |  |  |  |  |  |  |  |  |  |  |
| YJM451 | XIII | 895236 | 895236 | 895236 | rightend_complete | YMR321C | 100 | 318 | 0 | 0 | 1 | 318 | 888792 | 888475 | 1.9E-167 | 588 |
| YJM451 | XIV | 1 | 10 | 776549 | leftend | YNL332W | 99.12 | 1023 | 9 | 0 | 1 | 1023 | 17201 | 18223 | 0 | 1840 |
| YJM451 | XIV | 1651 | 1660 | 776549 | middle |  |  |  |  |  |  |  |  |  |  |  |
| YJM451 | XIV | 11382 | 11391 | 776549 | middle |  |  |  |  |  |  |  |  |  |  |  |
| YJM451 | XIV | 776540 | 776549 | 776549 | rightend | YNR071C | 99.611 | 1029 | 4 | 0 | 1 | 1029 | 757251 | 756223 | 0 | 1879 |
| YJM451 | XV | 1 | 10 | 1048013 | leftend | YOL154W | 90.945 | 751 | 66 | 2 | 1 | 750 | 23659 | 24408 | 0 | 1009 |
| YJM451 | XV | 1047552 | 1047561 | 1048013 | middle |  |  |  |  |  |  |  |  |  |  |  |
| YJM451 | XV | 1048004 | 1048013 | 1048013 | rightend | YOR381W | 98.642 | 2136 | 29 | 0 | 1 | 2136 | 1020647 | 1022782 | 0 | 3784 |
| YJM451 | XVI | 1 | 10 | 912950 | leftend | YPL274W | 98.583 | 1764 | 25 | 0 | 1 | 1764 | 6022 | 7785 | 0 | 3120 |
| YJM451 | XVI | 540 | 549 | 912950 | middle |  |  |  |  |  |  |  |  |  |  |  |
| YJM451 | XVI | 1029 | 1038 | 912950 | middle |  |  |  |  |  |  |  |  |  |  |  |
| YJM451 | XVI | 1692 | 1701 | 912950 | middle |  |  |  |  |  |  |  |  |  |  |  |
| YJM451 | XVI | 912941 | 912950 | 912950 | rightend | YPR201W | 99.588 | 1215 | 5 | 0 | 1 | 1215 | 909995 | 911209 | 0 | 2217 |
| YJM453 | I | 1 | 10 | 169227 | leftend | YAL063C | 84.229 | 577 | 87 | 4 | 1 | 575 | 585 | 11 | 4.41E-158 | 558 |
| YJM453 | I | 169218 | 169227 | 169227 | rightend | YAR050W | 84.584 | 493 | 74 | 2 | 1 | 492 | 168726 | 169217 | 6.83E-137 | 488 |
| YJM453 | II | 1 | 10 | 783617 | leftend | YNCB0001W | 100 | 84 | 0 | 0 | 1 | 84 | 1963 | 2046 | 4.16E-38 | 156 |
| YJM453 | II | 1601 | 1610 | 783617 | middle |  |  |  |  |  |  |  |  |  |  |  |
| YJM453 | II | 783608 | 783617 | 783617 | rightend | YBR297W | 92.031 | 1418 | 97 | 12 | 1 | 1407 | 774763 | 776175 | 0 | 1978 |
| YJM453 | III | 1 | 10 | 301730 | leftend | YCL073C | 99.675 | 1848 | 6 | 0 | 1 | 1848 | 7759 | 5912 | 0 | 3380 |
| YJM453 | III | 300944 | 300953 | 301730 | middle |  |  |  |  |  |  |  |  |  |  |  |
| YJM453 | III | 301209 | 301218 | 301730 | middle |  |  |  |  |  |  |  |  |  |  |  |
| YJM453 | III | 301721 | 301730 | 301730 | rightend | YCR101C | 99.818 | 549 | 1 | 0 | 1 | 549 | 296274 | 295726 | 0 | 1009 |
| YJM453 | IV | 1 |  | 1455630 | leftend_complete | YDL243C | 95.051 | 990 | 49 | 0 | 1 | 990 | 6743 | 5754 | 0 | 1557 |
| YJM453 | IV | 206 | 215 | 1455630 | middle |  |  |  |  |  |  |  |  |  |  |  |
| YJM453 | IV | 1455621 | 1455630 | 1455630 | rightend | YDR541C | 99.42 | 1035 | 6 | 0 | 1 | 1035 | 1452963 | 1451929 | 0 | 1879 |
| YJM453 | V | 1 | 10 | 550315 | leftend | YEL070W | 99.072 | 1509 | 14 | 0 | 1 | 1509 | 13267 | 14775 | 0 | 2710 |
| YJM453 | V | 550306 | 550315 | 550315 | rightend | YER187W | 91.981 | 424 | 32 | 2 | 4 | 426 | 547743 | 548165 | 3.44E-169 | 593 |
| YJM453 | VI | 1 | 10 | 234224 | leftend | YFL051C | 98.451 | 452 | 7 | 0 | 1 | 452 | 1092 | 641 | 0 | 797 |
| YJM453 | VI | 234215 | 234224 | 234224 | rightend | YFR055W | 100 | 1023 | 0 | 0 | 1 | 1023 | 227759 | 228781 | 0 | 1890 |
| YJM453 | VII | 1 |  | 1034903 | leftend_complete | YGL258W | 97.585 | 621 | 15 | 0 | 1 | 621 | 10947 | 11567 | 0 | 1064 |
| YJM453 | VII | 494 | 503 | 1034903 | middle |  |  |  |  |  |  |  |  |  |  |  |
| YJM453 | VII | 1034894 | 1034903 | 1034903 | rightend | YGR287C | 98.893 | 1626 | 18 | 0 | 145 | 1770 | 1030202 | 1028577 | 0 | 2904 |
| YJM453 | VIII | 1 | 10 | 540948 | leftend | YHL044W | 98.446 | 708 | 11 | 0 | 1 | 708 | 1562 | 2269 | 0 | 1247 |
| YJM453 | VIII | 512393 | 512402 | 540948 | middle |  |  |  |  |  |  |  |  |  |  |  |
| YJM453 | VIII | 528546 | 528555 | 540948 | middle |  |  |  |  |  |  |  |  |  |  |  |
| YJM453 | VIII | 532922 | 532931 | 540948 | middle |  |  |  |  |  |  |  |  |  |  |  |
| YJM453 | VIII | 540099 | 540108 | 540948 | middle |  |  |  |  |  |  |  |  |  |  |  |
| YJM453 | VIII | 540939 | 540948 | 540948 | rightend | YHR211W | 92.041 | 2136 | 146 | 2 | 1096 | 3228 | 510257 | 512371 | 0 | 2981 |
| YJM453 | IX | 1 | 10 | 403683 | leftend | YIL169C | 93.992 | 1548 | 87 | 3 | 265 | 1812 | 2180 | 639 | 0 | 2338 |
| YJM453 | IX | 403674 | 403683 | 403683 | rightend | YIR039C | 99.009 | 1614 | 16 | 0 | 1 | 1614 | 402408 | 400795 | 0 | 2892 |
| YJM453 | X | 1 |  | 711199 | leftend_complete | YJL213W | 99.699 | 996 | 3 | 0 | 1 | 996 | 9610 | 10605 | 0 | 1823 |
| YJM453 | X | 699055 | 699064 | 711199 | middle |  |  |  |  |  |  |  |  |  |  |  |
| YJM453 | X | 710372 | 710381 | 711199 | middle |  |  |  |  |  |  |  |  |  |  |  |
| YJM453 | X | 711190 | 711199 | 711199 | rightend | YJR152W | 99.755 | 1632 | 3 | 1 | 1 | 1632 | 678828 | 680458 | 0 | 2990 |
| YJM453 | XI | 1 | 10 | 645786 | leftend | YKL222C | 99.528 | 2118 | 10 | 0 | 1 | 2118 | 3478 | 1361 | 0 | 3856 |
| YJM453 | XI | 645110 | 645119 | 645786 | middle |  |  |  |  |  |  |  |  |  |  |  |
| YJM453 | XI | 645786 | 645786 | 645786 | rightend_complete | YKR101W | 99.847 | 1965 | 3 | 0 | 1 | 1965 | 637410 | 639374 | 0 | 3613 |
| YJM453 | XII | 1 |  | 2083044 | leftend_complete | YLL062C | 100 | 975 | 0 | 0 | 1 | 975 | 4830 | 3856 | 0 | 1801 |
| YJM453 | XII | 198 | 207 | 2083044 | middle |  |  |  |  |  |  |  |  |  |  |  |
| YJM453 | XII | 793 | 802 | 2083044 | middle |  |  |  |  |  |  |  |  |  |  |  |
| YJM453 | XII | 2082449 | 2082458 | 2083044 | middle |  |  |  |  |  |  |  |  |  |  |  |
| YJM453 | XII | 2083035 | 2083044 | 2083044 | rightend | YLR460C | 94.611 | 1132 | 59 | 2 | 1 | 1131 | 2081209 | 2080079 | 0 | 1751 |
| YJM453 | XIII | 1 | 10 | 906916 | leftend | YML131W | 100 | 1098 | 0 | 0 | 1 | 1098 | 4378 | 5475 | 0 | 2028 |
| YJM453 | XIII | 1786 | 1795 | 906916 | middle |  |  |  |  |  |  |  |  |  |  |  |
| YJM453 | XIII | 906907 | 906916 | 906916 | rightend | YMR321C | 99.361 | 313 | 2 | 0 | 6 | 318 | 889373 | 889061 | 2.48E-161 | 568 |
| YJM453 | XIV | 1 |  | 756332 | leftend_complete | YNL332W | 98.338 | 1023 | 17 | 0 | 1 | 1023 | 7214 | 8236 | 0 | 1803 |
| YJM453 | XIV | 451 | 460 | 756332 | middle |  |  |  |  |  |  |  |  |  |  |  |
| YJM453 | XIV | 1073 | 1082 | 756332 | middle |  |  |  |  |  |  |  |  |  |  |  |
| YJM453 | XIV | 756323 | 756332 | 756332 | rightend | YNR071C | 99.903 | 1029 | 1 | 0 | 1 | 1029 | 746028 | 745000 | 0 | 1895 |
| YJM453 | XV | 1 | 10 | 1066085 | leftend | YOL154W | 99.467 | 750 | 4 | 0 | 1 | 750 | 22860 | 23609 | 0 | 1363 |
| YJM453 | XV | 1064917 | 1064926 | 1066085 | middle |  |  |  |  |  |  |  |  |  |  |  |
| YJM453 | XV | 1066085 | 1066085 | 1066085 | rightend_complete | YOR381W | 99.906 | 2137 | 1 | 1 | 1 | 2136 | 1036444 | 1038580 | 0 | 3934 |
| YJM453 | XVI | 1 | 10 | 908151 | leftend | YPL274W | 99.943 | 1764 | 1 | 0 | 1 | 1764 | 19536 | 21299 | 0 | 3253 |
| YJM453 | XVI | 3624 | 3633 | 908151 | middle |  |  |  |  |  |  |  |  |  |  |  |
| YJM453 | XVI | 4807 | 4816 | 908151 | middle |  |  |  |  |  |  |  |  |  |  |  |
| YJM453 | XVI | 908142 | 908151 | 908151 | rightend | YPR201W | 100 | 1215 | 0 | 0 | 1 | 1215 | 905637 | 906851 | 0 | 2244 |
| YJM456 | I | 1 |  | 201093 | leftend_complete | YAL063C | 92.485 | 998 | 46 | 3 | 2966 | 3962 | 11858 | 10889 | 0 | 1400 |
| YJM456 | I | 1651 | 1660 | 201093 | middle |  |  |  |  |  |  |  |  |  |  |  |
| YJM456 | I | 11861 | 11870 | 201093 | middle |  |  |  |  |  |  |  |  |  |  |  |
| YJM456 | I | 192184 | 192193 | 201093 | middle |  |  |  |  |  |  |  |  |  |  |  |
| YJM456 | I | 201084 | 201093 | 201093 | rightend | YAR050W | 98.049 | 769 | 15 | 0 | 3846 | 4614 | 192194 | 192962 | 0 | 1338 |
| YJM456 | II | 1 | 10 | 787988 | leftend | YNCB0001W | 100 | 84 | 0 | 0 | 1 | 84 | 9208 | 9291 | 4.16E-38 | 156 |
| YJM456 | II | 928 | 937 | 787988 | middle |  |  |  |  |  |  |  |  |  |  |  |
| YJM456 | II | 787149 | 787158 | 787988 | middle |  |  |  |  |  |  |  |  |  |  |  |
| YJM456 | II | 787979 | 787988 | 787988 | rightend | YBR297W | 98.008 | 1205 | 24 | 0 | 1 | 1205 | 782960 | 784164 | 0 | 2093 |
| YJM456 | III | 1 | 10 | 306080 | leftend | YCL073C | 99.784 | 1848 | 4 | 0 | 1 | 1848 | 7716 | 5869 | 0 | 3391 |
| YJM456 | III | 304371 | 304380 | 306080 | middle |  |  |  |  |  |  |  |  |  |  |  |
| YJM456 | III | 306071 | 306080 | 306080 | rightend | YCR101C | 98.545 | 550 | 7 | 1 | 1 | 549 | 297654 | 297105 | 0 | 970 |
| YJM456 | IV | 1 | 10 | 1465633 | leftend | YDL243C | 93.939 | 990 | 60 | 0 | 1 | 990 | 17818 | 16829 | 0 | 1496 |
| YJM456 | IV | 2247 | 2256 | 1465633 | middle |  |  |  |  |  |  |  |  |  |  |  |
| YJM456 | IV | 2465 | 2474 | 1465633 | middle |  |  |  |  |  |  |  |  |  |  |  |
| YJM456 | IV | 1465624 | 1465633 | 1465633 | rightend | YDR541C | 99.517 | 1035 | 5 | 0 | 1 | 1035 | 1462996 | 1461962 | 0 | 1884 |
| YJM456 | V | 1 | 10 | 550804 | leftend | YEL070W | 99.205 | 1509 | 12 | 0 | 1 | 1509 | 13997 | 15505 | 0 | 2721 |
| YJM456 | V | 550804 | 550804 | 550804 | rightend_complete | YER187W | 99.296 | 426 | 3 | 0 | 1 | 426 | 546657 | 547082 | 0 | 771 |
| YJM456 | VI | 1 |  | 277623 | leftend_complete | YFL051C | 100 | 452 | 0 | 0 | 1 | 452 | 43401 | 42950 | 0 | 835 |
| YJM456 | VI | 756 | 765 | 277623 | middle |  |  |  |  |  |  |  |  |  |  |  |
| YJM456 | VI | 2275 | 2284 | 277623 | middle |  |  |  |  |  |  |  |  |  |  |  |
| YJM456 | VI | 277614 | 277623 | 277623 | rightend | YFR055W | 99.902 | 1023 | 1 | 0 | 1 | 1023 | 271277 | 272299 | 0 | 1884 |
| YJM456 | VII | 1 |  | 1057768 | leftend_complete | YGL258W | 99.517 | 621 | 3 | 0 | 1 | 621 | 20153 | 20773 | 0 | 1131 |
| YJM456 | VII | 693 | 702 | 1057768 | middle |  |  |  |  |  |  |  |  |  |  |  |
| YJM456 | VII | 1446 | 1455 | 1057768 | middle |  |  |  |  |  |  |  |  |  |  |  |
| YJM456 | VII | 15785 | 15794 | 1057768 | middle |  |  |  |  |  |  |  |  |  |  |  |
| YJM456 | VII | 1052679 | 1052688 | 1057768 | middle |  |  |  |  |  |  |  |  |  |  |  |
| YJM456 | VII | 1053109 | 1053118 | 1057768 | middle |  |  |  |  |  |  |  |  |  |  |  |
| YJM456 | VII | 1056306 | 1056315 | 1057768 | middle |  |  |  |  |  |  |  |  |  |  |  |
| YJM456 | VII | 1057310 | 1057319 | 1057768 | middle |  |  |  |  |  |  |  |  |  |  |  |
| YJM456 | VII | 1057768 | 1057768 | 1057768 | rightend_complete | YGR287C | 90.463 | 1772 | 165 | 4 | 1 | 1770 | 1042278 | 1040509 | 0 | 2333 |
| YJM456 | VIII | 1 | 10 | 525630 | leftend | YHL044W | 98.446 | 708 | 11 | 0 | 1 | 708 | 8313 | 9020 | 0 | 1247 |
| YJM456 | VIII | 520125 | 520134 | 525630 | middle |  |  |  |  |  |  |  |  |  |  |  |
| YJM456 | VIII | 520376 | 520385 | 525630 | middle |  |  |  |  |  |  |  |  |  |  |  |
| YJM456 | VIII | 523140 | 523149 | 525630 | middle |  |  |  |  |  |  |  |  |  |  |  |
| YJM456 | VIII | 524113 | 524122 | 525630 | middle |  |  |  |  |  |  |  |  |  |  |  |
| YJM456 | VIII | 524928 | 524937 | 525630 | middle |  |  |  |  |  |  |  |  |  |  |  |
| YJM456 | VIII | 525621 | 525630 | 525630 | rightend | YHR211W | 81.246 | 1557 | 255 | 15 | 1 | 1530 | 518829 | 520375 | 0 | 1223 |
| YJM456 | IX | 1 |  | 425441 | leftend_complete | YIL169C | 96.921 | 1949 | 50 | 8 | 520 | 2466 | 20679 | 18739 | 0 | 3258 |
| YJM456 | IX | 1807 | 1816 | 425441 | middle |  |  |  |  |  |  |  |  |  |  |  |
| YJM456 | IX | 424343 | 424352 | 425441 | middle |  |  |  |  |  |  |  |  |  |  |  |
| YJM456 | IX | 425441 | 425441 | 425441 | rightend_complete | YIR039C | 99.009 | 1614 | 16 | 0 | 1 | 1614 | 422357 | 420744 | 0 | 2892 |
| YJM456 | X | 1 | 10 | 726951 | leftend | YJL213W | 99.699 | 996 | 3 | 0 | 1 | 996 | 33750 | 34745 | 0 | 1823 |
| YJM456 | X | 2085 | 2094 | 726951 | middle |  |  |  |  |  |  |  |  |  |  |  |
| YJM456 | X | 3723 | 3732 | 726951 | middle |  |  |  |  |  |  |  |  |  |  |  |
| YJM456 | X | 4357 | 4366 | 726951 | middle |  |  |  |  |  |  |  |  |  |  |  |
| YJM456 | X | 724635 | 724644 | 726951 | middle |  |  |  |  |  |  |  |  |  |  |  |
| YJM456 | X | 725666 | 725675 | 726951 | middle |  |  |  |  |  |  |  |  |  |  |  |
| YJM456 | X | 726428 | 726437 | 726951 | middle |  |  |  |  |  |  |  |  |  |  |  |
| YJM456 | X | 726951 | 726951 | 726951 | rightend_complete | YJR152W | 99.51 | 1632 | 8 | 0 | 1 | 1632 | 704178 | 705809 | 0 | 2970 |
| YJM456 | XI | 1 |  | 664018 | leftend_complete | YKL222C | 99.433 | 2118 | 12 | 0 | 1 | 2118 | 5619 | 3502 | 0 | 3845 |
| YJM456 | XI | 663564 | 663573 | 664018 | middle |  |  |  |  |  |  |  |  |  |  |  |
| YJM456 | XI | 664018 | 664018 | 664018 | rightend_complete | YKR101W | 99.847 | 1965 | 3 | 0 | 1 | 1965 | 640896 | 642860 | 0 | 3613 |
| YJM456 | XII | 1 | 10 | 1566252 | leftend | YLL062C | 99.692 | 975 | 3 | 0 | 1 | 975 | 8631 | 7657 | 0 | 1784 |
| YJM456 | XII | 668 | 677 | 1566252 | middle |  |  |  |  |  |  |  |  |  |  |  |
| YJM456 | XII | 1564045 | 1564054 | 1566252 | middle |  |  |  |  |  |  |  |  |  |  |  |
| YJM456 | XII | 1565261 | 1565270 | 1566252 | middle |  |  |  |  |  |  |  |  |  |  |  |
| YJM456 | XII | 1565523 | 1565532 | 1566252 | middle |  |  |  |  |  |  |  |  |  |  |  |
| YJM456 | XII | 1566252 | 1566252 | 1566252 | rightend_complete | YLR460C | 99.823 | 1131 | 2 | 0 | 1 | 1131 | 1558915 | 1557785 | 0 | 2078 |
| YJM456 | XIII | 1 | 10 | 894558 | leftend | YML131W | 100 | 1098 | 0 | 0 | 1 | 1098 | 2977 | 4074 | 0 | 2028 |
| YJM456 | XIII | 706 | 715 | 894558 | middle |  |  |  |  |  |  |  |  |  |  |  |
| YJM456 | XIII | 891515 | 891524 | 894558 | middle |  |  |  |  |  |  |  |  |  |  |  |
| YJM456 | XIII | 892439 | 892448 | 894558 | middle |  |  |  |  |  |  |  |  |  |  |  |
| YJM456 | XIII | 894549 | 894558 | 894558 | rightend | YMR321C | 100 | 318 | 0 | 0 | 1 | 318 | 887574 | 887257 | 1.9E-167 | 588 |
| YJM456 | XIV | 1 |  | 755556 | leftend_complete | YNL332W | 98.436 | 1023 | 16 | 0 | 1 | 1023 | 6036 | 7058 | 0 | 1801 |
| YJM456 | XIV | 725 | 734 | 755556 | middle |  |  |  |  |  |  |  |  |  |  |  |
| YJM456 | XIV | 748135 | 748144 | 755556 | middle |  |  |  |  |  |  |  |  |  |  |  |
| YJM456 | XIV | 755547 | 755556 | 755556 | rightend | YNR071C | 99.903 | 1029 | 1 | 0 | 1 | 1029 | 745701 | 744673 | 0 | 1895 |
| YJM456 | XV | 1 | 10 | 1051043 | leftend | YOL154W | 99.467 | 750 | 4 | 0 | 1 | 750 | 25979 | 26728 | 0 | 1363 |
| YJM456 | XV | 1050323 | 1050332 | 1051043 | middle |  |  |  |  |  |  |  |  |  |  |  |
| YJM456 | XV | 1051034 | 1051043 | 1051043 | rightend | YOR381W | 98.83 | 2136 | 25 | 0 | 1 | 2136 | 1022842 | 1024977 | 0 | 3807 |
| YJM456 | XVI | 1 |  | 898380 | leftend_complete | YPL274W | 99.887 | 1764 | 2 | 0 | 1 | 1764 | 8713 | 10476 | 0 | 3247 |
| YJM456 | XVI | 664 | 673 | 898380 | middle |  |  |  |  |  |  |  |  |  |  |  |
| YJM456 | XVI | 1298 | 1307 | 898380 | middle |  |  |  |  |  |  |  |  |  |  |  |
| YJM456 | XVI | 898380 | 898380 | 898380 | rightend_complete | YPR201W | 99.506 | 1215 | 6 | 0 | 1 | 1215 | 895115 | 896329 | 0 | 2211 |
| YJM470 | I | 1 | 10 | 196934 | leftend | YAL063C | 84.392 | 2441 | 363 | 16 | 1 | 2432 | 19823 | 17392 | 0 | 2381 |
| YJM470 | I | 1609 | 1618 | 196934 | middle |  |  |  |  |  |  |  |  |  |  |  |
| YJM470 | I | 2204 | 2213 | 196934 | middle |  |  |  |  |  |  |  |  |  |  |  |
| YJM470 | I | 2843 | 2852 | 196934 | middle |  |  |  |  |  |  |  |  |  |  |  |
| YJM470 | I | 196925 | 196934 | 196934 | rightend | YAR050W | 86.832 | 2696 | 304 | 18 | 1632 | 4282 | 19002 | 16313 | 0 | 2964 |
| YJM470 | II | 1 | 10 | 787113 | leftend | YNCB0001W | 100 | 84 | 0 | 0 | 1 | 84 | 1186 | 1269 | 4.16E-38 | 156 |
| YJM470 | II | 782503 | 782512 | 787113 | middle |  |  |  |  |  |  |  |  |  |  |  |
| YJM470 | II | 783511 | 783520 | 787113 | middle |  |  |  |  |  |  |  |  |  |  |  |
| YJM470 | II | 787104 | 787113 | 787113 | rightend | YBR297W | 93.182 | 1408 | 94 | 2 | 1 | 1407 | 771859 | 773265 | 0 | 2067 |
| YJM470 | III | 1 | 10 | 310342 | leftend | YCL073C | 98.593 | 1848 | 26 | 0 | 1 | 1848 | 7537 | 5690 | 0 | 3269 |
| YJM470 | III | 300465 | 300474 | 310342 | middle |  |  |  |  |  |  |  |  |  |  |  |
| YJM470 | III | 308708 | 308717 | 310342 | middle |  |  |  |  |  |  |  |  |  |  |  |
| YJM470 | III | 309787 | 309796 | 310342 | middle |  |  |  |  |  |  |  |  |  |  |  |
| YJM470 | III | 310333 | 310342 | 310342 | rightend | YCR101C | 98.909 | 550 | 5 | 1 | 1 | 549 | 295277 | 294728 | 0 | 981 |
| YJM470 | IV | 1 | 10 | 1467031 | leftend | YDL243C | 95.253 | 990 | 47 | 0 | 1 | 990 | 20482 | 19493 | 0 | 1568 |
| YJM470 | IV | 3996 | 4005 | 1467031 | middle |  |  |  |  |  |  |  |  |  |  |  |
| YJM470 | IV | 1463426 | 1463435 | 1467031 | middle |  |  |  |  |  |  |  |  |  |  |  |
| YJM470 | IV | 1465404 | 1465413 | 1467031 | middle |  |  |  |  |  |  |  |  |  |  |  |
| YJM470 | IV | 1467031 | 1467031 | 1467031 | rightend_complete | YDR541C | 99.903 | 1035 | 1 | 0 | 1 | 1035 | 1460662 | 1459628 | 0 | 1906 |
| YJM470 | V | 1 | 10 | 553362 | leftend | YEL070W | 99.205 | 1509 | 12 | 0 | 1 | 1509 | 12606 | 14114 | 0 | 2721 |
| YJM470 | V | 552380 | 552389 | 553362 | middle |  |  |  |  |  |  |  |  |  |  |  |
| YJM470 | V | 553362 | 553362 | 553362 | rightend_complete | YER187W | 97.653 | 426 | 10 | 0 | 1 | 426 | 545248 | 545673 | 0 | 732 |
| YJM470 | VI | 1 | 10 | 280958 | leftend | YFL051C | 99.115 | 452 | 4 | 0 | 1 | 452 | 46358 | 45907 | 0 | 813 |
| YJM470 | VI | 4707 | 4716 | 280958 | middle |  |  |  |  |  |  |  |  |  |  |  |
| YJM470 | VI | 9467 | 9476 | 280958 | middle |  |  |  |  |  |  |  |  |  |  |  |
| YJM470 | VI | 280949 | 280958 | 280958 | rightend | YFR055W | 100 | 1023 | 0 | 0 | 1 | 1023 | 274799 | 275821 | 0 | 1890 |
| YJM470 | VII | 1 | 10 | 1049196 | leftend | YGL258W | 99.678 | 621 | 2 | 0 | 1 | 621 | 23617 | 24237 | 0 | 1136 |
| YJM470 | VII | 1048128 | 1048137 | 1049196 | middle |  |  |  |  |  |  |  |  |  |  |  |
| YJM470 | VII | 1048342 | 1048351 | 1049196 | middle |  |  |  |  |  |  |  |  |  |  |  |
| YJM470 | VII | 1048774 | 1048783 | 1049196 | middle |  |  |  |  |  |  |  |  |  |  |  |
| YJM470 | VII | 1049187 | 1049196 | 1049196 | rightend | YGR287C | 98.644 | 1770 | 24 | 0 | 1 | 1770 | 1046085 | 1044316 | 0 | 3138 |
| YJM470 | VIII | 1 | 10 | 520915 | leftend | YHL044W | 98.446 | 708 | 11 | 0 | 1 | 708 | 11786 | 12493 | 0 | 1247 |
| YJM470 | VIII | 1172 | 1181 | 520915 | middle |  |  |  |  |  |  |  |  |  |  |  |
| YJM470 | VIII | 1986 | 1995 | 520915 | middle |  |  |  |  |  |  |  |  |  |  |  |
| YJM470 | VIII | 3355 | 3364 | 520915 | middle |  |  |  |  |  |  |  |  |  |  |  |
| YJM470 | VIII | 519864 | 519873 | 520915 | middle |  |  |  |  |  |  |  |  |  |  |  |
| YJM470 | VIII | 520906 | 520915 | 520915 | rightend | YHR211W | 80.741 | 1916 | 337 | 19 | 1 | 1898 | 516290 | 518191 | 0 | 1465 |
| YJM470 | IX | 1 |  | 419600 | leftend_complete | YIL169C | 79.417 | 1064 | 215 | 3 | 753 | 1814 | 10211 | 9150 | 0 | 749 |
| YJM470 | IX | 2311 | 2320 | 419600 | middle |  |  |  |  |  |  |  |  |  |  |  |
| YJM470 | IX | 3510 | 3519 | 419600 | middle |  |  |  |  |  |  |  |  |  |  |  |
| YJM470 | IX | 6433 | 6442 | 419600 | middle |  |  |  |  |  |  |  |  |  |  |  |
| YJM470 | IX | 419600 | 419600 | 419600 | rightend_complete | YIR039C | 99.938 | 1614 | 1 | 0 | 1 | 1614 | 411491 | 409878 | 0 | 2976 |
| YJM470 | X | 1 |  | 705605 | leftend_complete | YJL213W | 99.498 | 996 | 5 | 0 | 1 | 996 | 14887 | 15882 | 0 | 1812 |
| YJM470 | X | 1168 | 1177 | 705605 | middle |  |  |  |  |  |  |  |  |  |  |  |
| YJM470 | X | 705596 | 705605 | 705605 | rightend | YJR152W | 99.51 | 1632 | 8 | 0 | 1 | 1632 | 685905 | 687536 | 0 | 2970 |
| YJM470 | XI | 1 |  | 666573 | leftend_complete | YKL222C | 99.669 | 2118 | 7 | 0 | 1 | 2118 | 4315 | 2198 | 0 | 3873 |
| YJM470 | XI | 1294 | 1303 | 666573 | middle |  |  |  |  |  |  |  |  |  |  |  |
| YJM470 | XI | 666564 | 666573 | 666573 | rightend | YKR101W | 99.695 | 1965 | 6 | 0 | 1 | 1965 | 638707 | 640671 | 0 | 3596 |
| YJM470 | XII | 1 | 10 | 1923186 | leftend | YLL062C | 99.795 | 975 | 2 | 0 | 1 | 975 | 5358 | 4384 | 0 | 1790 |
| YJM470 | XII | 1031 | 1040 | 1923186 | middle |  |  |  |  |  |  |  |  |  |  |  |
| YJM470 | XII | 1922650 | 1922659 | 1923186 | middle |  |  |  |  |  |  |  |  |  |  |  |
| YJM470 | XII | 1923177 | 1923186 | 1923186 | rightend | YLR460C | 99.293 | 1131 | 8 | 0 | 1 | 1131 | 1921868 | 1920738 | 0 | 2045 |
| YJM470 | XIII | 1 | 10 | 893579 | leftend | YML131W | 99.909 | 1098 | 1 | 0 | 1 | 1098 | 2152 | 3249 | 0 | 2023 |
| YJM470 | XIII | 893579 | 893579 | 893579 | rightend_complete | YMR321C | 100 | 318 | 0 | 0 | 1 | 318 | 887094 | 886777 | 1.9E-167 | 588 |
| YJM470 | XIV | 1 | 10 | 758584 | leftend | YNL332W | 100 | 261 | 0 | 0 | 763 | 1023 | 10546 | 10806 | 2.64E-135 | 483 |
| YJM470 | XIV | 10536 | 10545 | 758584 | middle |  |  |  |  |  |  |  |  |  |  |  |
| YJM470 | XIV | 757694 | 757703 | 758584 | middle |  |  |  |  |  |  |  |  |  |  |  |
| YJM470 | XIV | 758575 | 758584 | 758584 | rightend | YNR071C | 99.903 | 1029 | 1 | 0 | 1 | 1029 | 749830 | 748802 | 0 | 1895 |
| YJM470 | XV | 1 | 10 | 1027177 | leftend | YOL154W | 99.6 | 750 | 3 | 0 | 1 | 750 | 22855 | 23604 | 0 | 1369 |
| YJM470 | XV | 1027168 | 1027177 | 1027177 | rightend | YOR380W | 99.452 | 1641 | 9 | 0 | 1 | 1641 | 1014416 | 1016056 | 0 | 2981 |
| YJM470 | XVI | 1 | 10 | 901892 | leftend | YPL274W | 98.526 | 1764 | 26 | 0 | 1 | 1764 | 16609 | 18372 | 0 | 3114 |
| YJM470 | XVI | 529 | 538 | 901892 | middle |  |  |  |  |  |  |  |  |  |  |  |
| YJM470 | XVI | 947 | 956 | 901892 | middle |  |  |  |  |  |  |  |  |  |  |  |
| YJM470 | XVI | 1439 | 1448 | 901892 | middle |  |  |  |  |  |  |  |  |  |  |  |
| YJM470 | XVI | 2485 | 2494 | 901892 | middle |  |  |  |  |  |  |  |  |  |  |  |
| YJM470 | XVI | 901883 | 901892 | 901892 | rightend | YPR201W | 99.753 | 1215 | 3 | 0 | 1 | 1215 | 898869 | 900083 | 0 | 2228 |
| YJM541 | I | 1 | 10 | 204568 | leftend | YAL063C | 93.471 | 1256 | 71 | 9 | 2715 | 3962 | 10953 | 9701 | 0 | 1855 |
| YJM541 | I | 604 | 613 | 204568 | middle |  |  |  |  |  |  |  |  |  |  |  |
| YJM541 | I | 194156 | 194165 | 204568 | middle |  |  |  |  |  |  |  |  |  |  |  |
| YJM541 | I | 196586 | 196595 | 204568 | middle |  |  |  |  |  |  |  |  |  |  |  |
| YJM541 | I | 204559 | 204568 | 204568 | rightend | YAR050W | 91.783 | 1436 | 95 | 7 | 2869 | 4282 | 11433 | 9999 | 0 | 1977 |
| YJM541 | II | 1 | 10 | 794629 | leftend | YNCB0001W | 100 | 84 | 0 | 0 | 1 | 84 | 2673 | 2756 | 4.16E-38 | 156 |
| YJM541 | II | 1093 | 1102 | 794629 | middle |  |  |  |  |  |  |  |  |  |  |  |
| YJM541 | II | 794620 | 794629 | 794629 | rightend | YBR297W | 98.863 | 1407 | 14 | 1 | 1 | 1407 | 776048 | 777452 | 0 | 2508 |
| YJM541 | III | 1 |  | 313470 | leftend_complete | YCL073C | 99.351 | 1848 | 12 | 0 | 1 | 1848 | 7974 | 6127 | 0 | 3347 |
| YJM541 | III | 312361 | 312370 | 313470 | middle |  |  |  |  |  |  |  |  |  |  |  |
| YJM541 | III | 312938 | 312947 | 313470 | middle |  |  |  |  |  |  |  |  |  |  |  |
| YJM541 | III | 313470 | 313470 | 313470 | rightend_complete | YCR101C | 99.636 | 549 | 2 | 0 | 1 | 549 | 298590 | 298042 | 0 | 1003 |
| YJM541 | IV | 1 | 10 | 1458308 | leftend | YDL243C | 95.051 | 990 | 49 | 0 | 1 | 990 | 9550 | 8561 | 0 | 1557 |
| YJM541 | IV | 3013 | 3022 | 1458308 | middle |  |  |  |  |  |  |  |  |  |  |  |
| YJM541 | IV | 1458299 | 1458308 | 1458308 | rightend | YDR541C | 100 | 1035 | 0 | 0 | 1 | 1035 | 1453835 | 1452801 | 0 | 1912 |
| YJM541 | V | 1 | 10 | 556212 | leftend | YEL070W | 99.271 | 1509 | 11 | 0 | 1 | 1509 | 13419 | 14927 | 0 | 2726 |
| YJM541 | V | 598 | 607 | 556212 | middle |  |  |  |  |  |  |  |  |  |  |  |
| YJM541 | V | 555517 | 555526 | 556212 | middle |  |  |  |  |  |  |  |  |  |  |  |
| YJM541 | V | 556203 | 556212 | 556212 | rightend | YER187W | 91.981 | 424 | 32 | 2 | 4 | 426 | 546828 | 547250 | 3.44E-169 | 593 |
| YJM541 | VI | 1 | 10 | 240751 | leftend | YFL051C | 98.451 | 452 | 7 | 0 | 1 | 452 | 6190 | 5739 | 0 | 797 |
| YJM541 | VI | 4998 | 5007 | 240751 | middle |  |  |  |  |  |  |  |  |  |  |  |
| YJM541 | VI | 240751 | 240751 | 240751 | rightend_complete | YFR055W | 100 | 1023 | 0 | 0 | 1 | 1023 | 234798 | 235820 | 0 | 1890 |
| YJM541 | VII | 1 | 10 | 1044800 | leftend | YGL258W | 96.457 | 621 | 22 | 0 | 1 | 621 | 3885 | 4505 | 0 | 1026 |
| YJM541 | VII | 1042344 | 1042353 | 1044800 | middle |  |  |  |  |  |  |  |  |  |  |  |
| YJM541 | VII | 1044791 | 1044800 | 1044800 | rightend | YGR287C | 99.209 | 1770 | 14 | 0 | 1 | 1770 | 1024847 | 1023078 | 0 | 3192 |
| YJM541 | VIII | 1 |  | 537428 | leftend_complete | YHL044W | 98.446 | 708 | 11 | 0 | 1 | 708 | 9442 | 10149 | 0 | 1247 |
| YJM541 | VIII | 7404 | 7413 | 537428 | middle |  |  |  |  |  |  |  |  |  |  |  |
| YJM541 | VIII | 8050 | 8059 | 537428 | middle |  |  |  |  |  |  |  |  |  |  |  |
| YJM541 | VIII | 533257 | 533266 | 537428 | middle |  |  |  |  |  |  |  |  |  |  |  |
| YJM541 | VIII | 534683 | 534692 | 537428 | middle |  |  |  |  |  |  |  |  |  |  |  |
| YJM541 | VIII | 537419 | 537428 | 537428 | rightend | YHR211W | 85.172 | 1632 | 230 | 8 | 1 | 1624 | 520642 | 522269 | 0 | 1663 |
| YJM541 | IX | 1 |  | 421423 | leftend_complete | YIL169C | 96.119 | 1443 | 48 | 4 | 478 | 1919 | 20538 | 19103 | 0 | 2348 |
| YJM541 | IX | 1602 | 1611 | 421423 | middle |  |  |  |  |  |  |  |  |  |  |  |
| YJM541 | IX | 3191 | 3200 | 421423 | middle |  |  |  |  |  |  |  |  |  |  |  |
| YJM541 | IX | 421414 | 421423 | 421423 | rightend | YIR039C | 99.009 | 1614 | 16 | 0 | 1 | 1614 | 419585 | 417972 | 0 | 2892 |
| YJM541 | X | 1 |  | 721411 | leftend_complete | YJL213W | 99.699 | 996 | 3 | 0 | 1 | 996 | 27971 | 28966 | 0 | 1823 |
| YJM541 | X | 2725 | 2734 | 721411 | middle |  |  |  |  |  |  |  |  |  |  |  |
| YJM541 | X | 721042 | 721051 | 721411 | middle |  |  |  |  |  |  |  |  |  |  |  |
| YJM541 | X | 721411 | 721411 | 721411 | rightend_complete | YJR152W | 99.571 | 1632 | 7 | 0 | 1 | 1632 | 697711 | 699342 | 0 | 2976 |
| YJM541 | XI | 1 |  | 674647 | leftend_complete | YKL222C | 99.387 | 2119 | 12 | 1 | 1 | 2118 | 3745 | 1627 | 0 | 3840 |
| YJM541 | XI | 559 | 568 | 674647 | middle |  |  |  |  |  |  |  |  |  |  |  |
| YJM541 | XI | 674638 | 674647 | 674647 | rightend | YKR101W | 99.847 | 1965 | 3 | 0 | 1 | 1965 | 655021 | 656985 | 0 | 3613 |
| YJM541 | XII | 1 |  | 2303819 | leftend_complete | YLL062C | 100 | 975 | 0 | 0 | 1 | 975 | 5508 | 4534 | 0 | 1801 |
| YJM541 | XII | 1215 | 1224 | 2303819 | middle |  |  |  |  |  |  |  |  |  |  |  |
| YJM541 | XII | 1569 | 1578 | 2303819 | middle |  |  |  |  |  |  |  |  |  |  |  |
| YJM541 | XII | 2302044 | 2302053 | 2303819 | middle |  |  |  |  |  |  |  |  |  |  |  |
| YJM541 | XII | 2303819 | 2303819 | 2303819 | rightend_complete | YLR460C | 93.982 | 1130 | 66 | 2 | 3 | 1131 | 2301067 | 2299939 | 0 | 1709 |
| YJM541 | XIII | 1 | 10 | 900382 | leftend | YML131W | 100 | 1098 | 0 | 0 | 1 | 1098 | 3971 | 5068 | 0 | 2028 |
| YJM541 | XIII | 797 | 806 | 900382 | middle |  |  |  |  |  |  |  |  |  |  |  |
| YJM541 | XIII | 894535 | 894544 | 900382 | middle |  |  |  |  |  |  |  |  |  |  |  |
| YJM541 | XIII | 899555 | 899564 | 900382 | middle |  |  |  |  |  |  |  |  |  |  |  |
| YJM541 | XIII | 900373 | 900382 | 900382 | rightend | YMR321C | 99.361 | 313 | 2 | 0 | 6 | 318 | 889129 | 888817 | 2.48E-161 | 568 |
| YJM541 | XIV | 1 | 10 | 752126 | leftend | YNL332W | 98.631 | 1023 | 14 | 0 | 1 | 1023 | 948 | 1970 | 0 | 1812 |
| YJM541 | XIV | 366 | 375 | 752126 | middle |  |  |  |  |  |  |  |  |  |  |  |
| YJM541 | XIV | 724 | 733 | 752126 | middle |  |  |  |  |  |  |  |  |  |  |  |
| YJM541 | XIV | 751410 | 751419 | 752126 | middle |  |  |  |  |  |  |  |  |  |  |  |
| YJM541 | XIV | 751669 | 751678 | 752126 | middle |  |  |  |  |  |  |  |  |  |  |  |
| YJM541 | XIV | 752126 | 752126 | 752126 | rightend_complete | YNR071C | 100 | 1029 | 0 | 0 | 1 | 1029 | 741111 | 740083 | 0 | 1901 |
| YJM541 | XV | 1 | 10 | 1051167 | leftend | YOL154W | 99.333 | 750 | 5 | 0 | 1 | 750 | 24872 | 25621 | 0 | 1358 |
| YJM541 | XV | 1046238 | 1046247 | 1051167 | middle |  |  |  |  |  |  |  |  |  |  |  |
| YJM541 | XV | 1046640 | 1046649 | 1051167 | middle |  |  |  |  |  |  |  |  |  |  |  |
| YJM541 | XV | 1046851 | 1046860 | 1051167 | middle |  |  |  |  |  |  |  |  |  |  |  |
| YJM541 | XV | 1050713 | 1050722 | 1051167 | middle |  |  |  |  |  |  |  |  |  |  |  |
| YJM541 | XV | 1051158 | 1051167 | 1051167 | rightend | YOR381W | 99.86 | 2136 | 3 | 0 | 1 | 2136 | 1023991 | 1026126 | 0 | 3928 |
| YJM541 | XVI | 1 |  | 906541 | leftend_complete | YPL274W | 99.887 | 1764 | 2 | 0 | 1 | 1764 | 11423 | 13186 | 0 | 3247 |
| YJM541 | XVI | 1761 | 1770 | 906541 | middle |  |  |  |  |  |  |  |  |  |  |  |
| YJM541 | XVI | 4890 | 4899 | 906541 | middle |  |  |  |  |  |  |  |  |  |  |  |
| YJM541 | XVI | 906532 | 906541 | 906541 | rightend | YPR201W | 99.588 | 1215 | 5 | 0 | 1 | 1215 | 902990 | 904204 | 0 | 2217 |
| YJM554 | I | 1 | 10 | 199625 | leftend | YAL063C | 97.709 | 1004 | 23 | 0 | 2966 | 3969 | 185453 | 186456 | 0 | 1727 |
| YJM554 | I | 608 | 617 | 199625 | middle |  |  |  |  |  |  |  |  |  |  |  |
| YJM554 | I | 199616 | 199625 | 199625 | rightend | YAR050W | 90.526 | 1425 | 94 | 8 | 2869 | 4253 | 11796 | 10373 | 0 | 1845 |
| YJM554 | II | 1 | 10 | 792253 | leftend | YNCB0001W | 100 | 84 | 0 | 0 | 1 | 84 | 547 | 630 | 4.16E-38 | 156 |
| YJM554 | II | 792244 | 792253 | 792253 | rightend | YBR297W | 98.863 | 1407 | 14 | 1 | 1 | 1407 | 773670 | 775074 | 0 | 2508 |
| YJM554 | III | 1 |  | 316995 | leftend_complete | YCL073C | 99.351 | 1848 | 12 | 0 | 1 | 1848 | 8040 | 6193 | 0 | 3347 |
| YJM554 | III | 316403 | 316412 | 316995 | middle |  |  |  |  |  |  |  |  |  |  |  |
| YJM554 | III | 316986 | 316995 | 316995 | rightend | YCR101C | 97.636 | 550 | 12 | 1 | 1 | 549 | 299142 | 298593 | 0 | 942 |
| YJM554 | IV | 1 |  | 1474585 | leftend_complete | YDL243C | 95.051 | 990 | 49 | 0 | 1 | 990 | 20708 | 19719 | 0 | 1557 |
| YJM554 | IV | 3977 | 3986 | 1474585 | middle |  |  |  |  |  |  |  |  |  |  |  |
| YJM554 | IV | 1474576 | 1474585 | 1474585 | rightend | YDR541C | 100 | 1035 | 0 | 0 | 1 | 1035 | 1470003 | 1468969 | 0 | 1912 |
| YJM554 | V | 1 | 10 | 553050 | leftend | YEL070W | 99.006 | 1509 | 15 | 0 | 1 | 1509 | 13419 | 14927 | 0 | 2704 |
| YJM554 | V | 598 | 607 | 553050 | middle |  |  |  |  |  |  |  |  |  |  |  |
| YJM554 | V | 549202 | 549211 | 553050 | middle |  |  |  |  |  |  |  |  |  |  |  |
| YJM554 | V | 550005 | 550014 | 553050 | middle |  |  |  |  |  |  |  |  |  |  |  |
| YJM554 | V | 553041 | 553050 | 553050 | rightend | YER187W | 91.981 | 424 | 32 | 2 | 4 | 426 | 546696 | 547118 | 3.44E-169 | 593 |
| YJM554 | VI | 1 | 10 | 276596 | leftend | YFL051C | 98.451 | 452 | 7 | 0 | 1 | 452 | 42544 | 42093 | 0 | 797 |
| YJM554 | VI | 276587 | 276596 | 276596 | rightend | YFR055W | 99.902 | 1023 | 1 | 0 | 1 | 1023 | 270247 | 271269 | 0 | 1884 |
| YJM554 | VII | 1 |  | 1054886 | leftend_complete | YGL258W | 96.457 | 621 | 22 | 0 | 1 | 621 | 4763 | 5383 | 0 | 1026 |
| YJM554 | VII | 202 | 211 | 1054886 | middle |  |  |  |  |  |  |  |  |  |  |  |
| YJM554 | VII | 1051210 | 1051219 | 1054886 | middle |  |  |  |  |  |  |  |  |  |  |  |
| YJM554 | VII | 1052143 | 1052152 | 1054886 | middle |  |  |  |  |  |  |  |  |  |  |  |
| YJM554 | VII | 1054886 | 1054886 | 1054886 | rightend_complete | YGR287C | 99.209 | 1770 | 14 | 0 | 1 | 1770 | 1028917 | 1027148 | 0 | 3192 |
| YJM554 | VIII | 1 | 10 | 509488 | leftend | YHL044W | 90.221 | 317 | 29 | 2 | 180 | 496 | 3673 | 3987 | 1.82E-114 | 412 |
| YJM554 | VIII | 424 | 433 | 509488 | middle |  |  |  |  |  |  |  |  |  |  |  |
| YJM554 | VIII | 1458 | 1467 | 509488 | middle |  |  |  |  |  |  |  |  |  |  |  |
| YJM554 | VIII | 509479 | 509488 | 509488 | rightend | YHR211W | 94.867 | 974 | 48 | 2 | 2249 | 3221 | 508315 | 509287 | 0 | 1520 |
| YJM554 | IX | 1 |  | 426295 | leftend_complete | YIL169C | 96.119 | 1443 | 48 | 4 | 478 | 1919 | 24339 | 22904 | 0 | 2348 |
| YJM554 | IX | 1631 | 1640 | 426295 | middle |  |  |  |  |  |  |  |  |  |  |  |
| YJM554 | IX | 2328 | 2337 | 426295 | middle |  |  |  |  |  |  |  |  |  |  |  |
| YJM554 | IX | 4599 | 4608 | 426295 | middle |  |  |  |  |  |  |  |  |  |  |  |
| YJM554 | IX | 7327 | 7336 | 426295 | middle |  |  |  |  |  |  |  |  |  |  |  |
| YJM554 | IX | 7764 | 7773 | 426295 | middle |  |  |  |  |  |  |  |  |  |  |  |
| YJM554 | IX | 8462 | 8471 | 426295 | middle |  |  |  |  |  |  |  |  |  |  |  |
| YJM554 | IX | 424831 | 424840 | 426295 | middle |  |  |  |  |  |  |  |  |  |  |  |
| YJM554 | IX | 425582 | 425591 | 426295 | middle |  |  |  |  |  |  |  |  |  |  |  |
| YJM554 | IX | 426295 | 426295 | 426295 | rightend_complete | YIR039C | 99.009 | 1614 | 16 | 0 | 1 | 1614 | 423364 | 421751 | 0 | 2892 |
| YJM554 | X | 1 | 10 | 713604 | leftend | YJL213W | 99.699 | 996 | 3 | 0 | 1 | 996 | 10060 | 11055 | 0 | 1823 |
| YJM554 | X | 700061 | 700070 | 713604 | middle |  |  |  |  |  |  |  |  |  |  |  |
| YJM554 | X | 710973 | 710982 | 713604 | middle |  |  |  |  |  |  |  |  |  |  |  |
| YJM554 | X | 712404 | 712413 | 713604 | middle |  |  |  |  |  |  |  |  |  |  |  |
| YJM554 | X | 713604 | 713604 | 713604 | rightend_complete | YJR152W | 99.571 | 1632 | 7 | 0 | 1 | 1632 | 679915 | 681546 | 0 | 2976 |
| YJM554 | XI | 1 | 10 | 660331 | leftend | YKL222C | 99.387 | 2119 | 12 | 1 | 1 | 2118 | 3219 | 1101 | 0 | 3840 |
| YJM554 | XI | 659195 | 659204 | 660331 | middle |  |  |  |  |  |  |  |  |  |  |  |
| YJM554 | XI | 660331 | 660331 | 660331 | rightend_complete | YKR101W | 99.796 | 1965 | 4 | 0 | 1 | 1965 | 636669 | 638633 | 0 | 3607 |
| YJM554 | XII | 1 |  | 2351370 | leftend_complete | YLL062C | 100 | 975 | 0 | 0 | 1 | 975 | 4623 | 3649 | 0 | 1801 |
| YJM554 | XII | 468 | 477 | 2351370 | middle |  |  |  |  |  |  |  |  |  |  |  |
| YJM554 | XII | 2351361 | 2351370 | 2351370 | rightend | YLR460C | 93.993 | 1132 | 66 | 2 | 1 | 1131 | 2345653 | 2344523 | 0 | 1712 |
| YJM554 | XIII | 1 | 10 | 900581 | leftend | YML131W | 100 | 1098 | 0 | 0 | 1 | 1098 | 2353 | 3450 | 0 | 2028 |
| YJM554 | XIII | 895659 | 895668 | 900581 | middle |  |  |  |  |  |  |  |  |  |  |  |
| YJM554 | XIII | 900572 | 900581 | 900581 | rightend | YMR321C | 99.361 | 313 | 2 | 0 | 6 | 318 | 887690 | 887378 | 2.48E-161 | 568 |
| YJM554 | XIV | 1 | 10 | 769332 | leftend | YNL332W | 99.12 | 1023 | 9 | 0 | 1 | 1023 | 1701 | 2723 | 0 | 1840 |
| YJM554 | XIV | 715 | 724 | 769332 | middle |  |  |  |  |  |  |  |  |  |  |  |
| YJM554 | XIV | 1073 | 1082 | 769332 | middle |  |  |  |  |  |  |  |  |  |  |  |
| YJM554 | XIV | 760689 | 760698 | 769332 | middle |  |  |  |  |  |  |  |  |  |  |  |
| YJM554 | XIV | 768601 | 768610 | 769332 | middle |  |  |  |  |  |  |  |  |  |  |  |
| YJM554 | XIV | 768860 | 768869 | 769332 | middle |  |  |  |  |  |  |  |  |  |  |  |
| YJM554 | XIV | 769332 | 769332 | 769332 | rightend_complete | YNR071C | 100 | 1029 | 0 | 0 | 1 | 1029 | 741490 | 740462 | 0 | 1901 |
| YJM554 | XV | 1 | 10 | 1048973 | leftend | YOL154W | 99.333 | 750 | 5 | 0 | 1 | 750 | 23029 | 23778 | 0 | 1358 |
| YJM554 | XV | 1048964 | 1048973 | 1048973 | rightend | YOR381W | 99.86 | 2136 | 3 | 0 | 1 | 2136 | 1021865 | 1024000 | 0 | 3928 |
| YJM554 | XVI | 1 | 10 | 922022 | leftend | YPL274W | 98.073 | 1764 | 34 | 0 | 1 | 1764 | 26240 | 28003 | 0 | 3070 |
| YJM554 | XVI | 609 | 618 | 922022 | middle |  |  |  |  |  |  |  |  |  |  |  |
| YJM554 | XVI | 9904 | 9913 | 922022 | middle |  |  |  |  |  |  |  |  |  |  |  |
| YJM554 | XVI | 10396 | 10405 | 922022 | middle |  |  |  |  |  |  |  |  |  |  |  |
| YJM554 | XVI | 11758 | 11767 | 922022 | middle |  |  |  |  |  |  |  |  |  |  |  |
| YJM554 | XVI | 921431 | 921440 | 922022 | middle |  |  |  |  |  |  |  |  |  |  |  |
| YJM554 | XVI | 922013 | 922022 | 922022 | rightend | YPR201W | 99.588 | 1215 | 5 | 0 | 1 | 1215 | 917789 | 919003 | 0 | 2217 |
| YJM555 | I | 1 | 10 | 209374 | leftend | YAL063C | 88.702 | 2434 | 271 | 3 | 1 | 2432 | 13374 | 10943 | 0 | 2968 |
| YJM555 | I | 209365 | 209374 | 209374 | rightend | YAR050W | 84.49 | 2392 | 361 | 9 | 46 | 2432 | 13329 | 10943 | 0 | 2353 |
| YJM555 | II | 1 | 10 | 784063 | leftend | YNCB0001W | 100 | 84 | 0 | 0 | 1 | 84 | 2945 | 3028 | 4.16E-38 | 156 |
| YJM555 | II | 1289 | 1298 | 784063 | middle |  |  |  |  |  |  |  |  |  |  |  |
| YJM555 | II | 784063 | 784063 | 784063 | rightend_complete | YBR297W | 98.008 | 1205 | 24 | 0 | 1 | 1205 | 775864 | 777068 | 0 | 2093 |
| YJM555 | III | 1 |  | 311524 | leftend_complete | YCL073C | 99.513 | 1848 | 9 | 0 | 1 | 1848 | 7994 | 6147 | 0 | 3363 |
| YJM555 | III | 310932 | 310941 | 311524 | middle |  |  |  |  |  |  |  |  |  |  |  |
| YJM555 | III | 311524 | 311524 | 311524 | rightend_complete | YCR101C | 92.021 | 564 | 28 | 12 | 1 | 549 | 297652 | 297091 | 0 | 776 |
| YJM555 | IV | 1 | 10 | 1459441 | leftend | YDL243C | 93.535 | 990 | 64 | 0 | 1 | 990 | 9025 | 8036 | 0 | 1474 |
| YJM555 | IV | 1331 | 1340 | 1459441 | middle |  |  |  |  |  |  |  |  |  |  |  |
| YJM555 | IV | 4322 | 4331 | 1459441 | middle |  |  |  |  |  |  |  |  |  |  |  |
| YJM555 | IV | 1459441 | 1459441 | 1459441 | rightend_complete | YDR541C | 100 | 1035 | 0 | 0 | 1 | 1035 | 1454734 | 1453700 | 0 | 1912 |
| YJM555 | V | 1 | 10 | 555202 | leftend | YEL070W | 99.139 | 1509 | 13 | 0 | 1 | 1509 | 12908 | 14416 | 0 | 2715 |
| YJM555 | V | 555202 | 555202 | 555202 | rightend_complete | YER187W | 91.981 | 424 | 32 | 2 | 4 | 426 | 546563 | 546985 | 3.44E-169 | 593 |
| YJM555 | VI | 1 | 10 | 281062 | leftend | YFL051C | 100 | 452 | 0 | 0 | 1 | 452 | 45503 | 45052 | 0 | 835 |
| YJM555 | VI | 1073 | 1082 | 281062 | middle |  |  |  |  |  |  |  |  |  |  |  |
| YJM555 | VI | 281053 | 281062 | 281062 | rightend | YFR055W | 99.902 | 1023 | 1 | 0 | 1 | 1023 | 274713 | 275735 | 0 | 1884 |
| YJM555 | VII | 1 |  | 1053060 | leftend_complete | YGL258W | 96.457 | 621 | 22 | 0 | 1 | 621 | 6820 | 7440 | 0 | 1026 |
| YJM555 | VII | 1053051 | 1053060 | 1053060 | rightend | YGR287C | 99.379 | 1770 | 11 | 0 | 1 | 1770 | 1028653 | 1026884 | 0 | 3208 |
| YJM555 | VIII | 1 | 10 | 531677 | leftend | YHL044W | 98.446 | 708 | 11 | 0 | 1 | 708 | 1165 | 1872 | 0 | 1247 |
| YJM555 | VIII | 514082 | 514091 | 531677 | middle |  |  |  |  |  |  |  |  |  |  |  |
| YJM555 | VIII | 531677 | 531677 | 531677 | rightend_complete | YHR211W | 92.135 | 2136 | 144 | 2 | 1096 | 3228 | 510426 | 512540 | 0 | 2992 |
| YJM555 | IX | 1 | 10 | 408566 | leftend | YIL169C | 96.539 | 1907 | 56 | 8 | 562 | 2466 | 6456 | 4558 | 0 | 3147 |
| YJM555 | IX | 407444 | 407453 | 408566 | middle |  |  |  |  |  |  |  |  |  |  |  |
| YJM555 | IX | 408566 | 408566 | 408566 | rightend_complete | YIR039C | 98.947 | 1614 | 17 | 0 | 1 | 1614 | 405524 | 403911 | 0 | 2887 |
| YJM555 | X | 1 | 10 | 719054 | leftend | YJL213W | 99.9 | 996 | 1 | 0 | 1 | 996 | 23711 | 24706 | 0 | 1834 |
| YJM555 | X | 529 | 538 | 719054 | middle |  |  |  |  |  |  |  |  |  |  |  |
| YJM555 | X | 800 | 809 | 719054 | middle |  |  |  |  |  |  |  |  |  |  |  |
| YJM555 | X | 717596 | 717605 | 719054 | middle |  |  |  |  |  |  |  |  |  |  |  |
| YJM555 | X | 719054 | 719054 | 719054 | rightend_complete | YJR152W | 99.755 | 1632 | 3 | 1 | 1 | 1632 | 693871 | 695501 | 0 | 2990 |
| YJM555 | XI | 1 | 10 | 676581 | leftend | YKL222C | 99.434 | 2119 | 11 | 1 | 1 | 2118 | 3353 | 1235 | 0 | 3845 |
| YJM555 | XI | 674838 | 674847 | 676581 | middle |  |  |  |  |  |  |  |  |  |  |  |
| YJM555 | XI | 676572 | 676581 | 676581 | rightend | YKR101W | 99.644 | 1965 | 7 | 0 | 1 | 1965 | 654784 | 656748 | 0 | 3591 |
| YJM555 | XII | 1 | 10 | 2585987 | leftend | YLL062C | 99.59 | 975 | 4 | 0 | 1 | 975 | 5889 | 4915 | 0 | 1779 |
| YJM555 | XII | 1537 | 1546 | 2585987 | middle |  |  |  |  |  |  |  |  |  |  |  |
| YJM555 | XII | 2584195 | 2584204 | 2585987 | middle |  |  |  |  |  |  |  |  |  |  |  |
| YJM555 | XII | 2585978 | 2585987 | 2585987 | rightend | YLR460C | 98.322 | 1132 | 17 | 2 | 1 | 1131 | 2582117 | 2580987 | 0 | 1984 |
| YJM555 | XIII | 1 | 10 | 923525 | leftend | YML131W | 100 | 1098 | 0 | 0 | 1 | 1098 | 13050 | 14147 | 0 | 2028 |
| YJM555 | XIII | 10015 | 10024 | 923525 | middle |  |  |  |  |  |  |  |  |  |  |  |
| YJM555 | XIII | 923516 | 923525 | 923525 | rightend | YMR321C | 99.361 | 313 | 2 | 0 | 6 | 318 | 898004 | 897692 | 2.48E-161 | 568 |
| YJM555 | XIV | 1 | 10 | 751042 | leftend | YNL332W | 98.631 | 1023 | 14 | 0 | 1 | 1023 | 2023 | 3045 | 0 | 1812 |
| YJM555 | XIV | 890 | 899 | 751042 | middle |  |  |  |  |  |  |  |  |  |  |  |
| YJM555 | XIV | 1110 | 1119 | 751042 | middle |  |  |  |  |  |  |  |  |  |  |  |
| YJM555 | XIV | 751042 | 751042 | 751042 | rightend_complete | YNR071C | 99.611 | 1029 | 4 | 0 | 1 | 1029 | 741047 | 740019 | 0 | 1879 |
| YJM555 | XV | 1 | 10 | 1048146 | leftend | YOL154W | 99.6 | 750 | 3 | 0 | 1 | 750 | 23117 | 23866 | 0 | 1369 |
| YJM555 | XV | 1048137 | 1048146 | 1048146 | rightend | YOR381W | 99.251 | 2136 | 16 | 0 | 1 | 2136 | 1020707 | 1022842 | 0 | 3856 |
| YJM555 | XVI | 1 |  | 917392 | leftend_complete | YPL274W | 99.887 | 1764 | 2 | 0 | 1 | 1764 | 25825 | 27588 | 0 | 3247 |
| YJM555 | XVI | 243 | 252 | 917392 | middle |  |  |  |  |  |  |  |  |  |  |  |
| YJM555 | XVI | 917383 | 917392 | 917392 | rightend | YPR201W | 99.588 | 1215 | 5 | 0 | 1 | 1215 | 913973 | 915187 | 0 | 2217 |
| YJM627 | I | 1 |  | 212290 | leftend_complete | YAL063C | 94.99 | 998 | 48 | 2 | 2966 | 3962 | 11315 | 10319 | 0 | 1565 |
| YJM627 | I | 921 | 930 | 212290 | middle |  |  |  |  |  |  |  |  |  |  |  |
| YJM627 | I | 1129 | 1138 | 212290 | middle |  |  |  |  |  |  |  |  |  |  |  |
| YJM627 | I | 208873 | 208882 | 212290 | middle |  |  |  |  |  |  |  |  |  |  |  |
| YJM627 | I | 209284 | 209293 | 212290 | middle |  |  |  |  |  |  |  |  |  |  |  |
| YJM627 | I | 210005 | 210014 | 212290 | middle |  |  |  |  |  |  |  |  |  |  |  |
| YJM627 | I | 210231 | 210240 | 212290 | middle |  |  |  |  |  |  |  |  |  |  |  |
| YJM627 | I | 211690 | 211699 | 212290 | middle |  |  |  |  |  |  |  |  |  |  |  |
| YJM627 | I | 212281 | 212290 | 212290 | rightend | YAR050W | 81.985 | 1632 | 290 | 4 | 1 | 1630 | 183273 | 184902 | 0 | 1382 |
| YJM627 | II | 1 |  | 797166 | leftend_complete | YNCB0001W | 100 | 84 | 0 | 0 | 1 | 84 | 2844 | 2927 | 4.16E-38 | 156 |
| YJM627 | II | 1121 | 1130 | 797166 | middle |  |  |  |  |  |  |  |  |  |  |  |
| YJM627 | II | 795084 | 795093 | 797166 | middle |  |  |  |  |  |  |  |  |  |  |  |
| YJM627 | II | 797157 | 797166 | 797166 | rightend | YBR297W | 99.858 | 1407 | 2 | 0 | 1 | 1407 | 776617 | 778023 | 0 | 2588 |
| YJM627 | III | 1 | 10 | 301130 | leftend | YCL073C | 99.405 | 1848 | 10 | 1 | 1 | 1848 | 7932 | 6086 | 0 | 3350 |
| YJM627 | III | 301121 | 301130 | 301130 | rightend | YCR101C | 99.818 | 549 | 1 | 0 | 1 | 549 | 296327 | 295779 | 0 | 1009 |
| YJM627 | IV | 1 | 10 | 1452834 | leftend | YDL243C | 95.055 | 991 | 48 | 1 | 1 | 990 | 6156 | 5166 | 0 | 1557 |
| YJM627 | IV | 4731 | 4740 | 1452834 | middle |  |  |  |  |  |  |  |  |  |  |  |
| YJM627 | IV | 1452825 | 1452834 | 1452834 | rightend | YDR541C | 99.614 | 1035 | 4 | 0 | 1 | 1035 | 1449775 | 1448741 | 0 | 1890 |
| YJM627 | V | 1 | 10 | 555786 | leftend | YEL070W | 99.139 | 1509 | 13 | 0 | 1 | 1509 | 15281 | 16789 | 0 | 2715 |
| YJM627 | V | 555786 | 555786 | 555786 | rightend_complete | YER187W | 91.981 | 424 | 32 | 2 | 4 | 426 | 549676 | 550098 | 3.44E-169 | 593 |
| YJM627 | VI | 1 | 10 | 277726 | leftend | YFL051C | 98.451 | 452 | 7 | 0 | 1 | 452 | 42943 | 42492 | 0 | 797 |
| YJM627 | VI | 277717 | 277726 | 277726 | rightend | YFR055W | 100 | 1023 | 0 | 0 | 1 | 1023 | 271461 | 272483 | 0 | 1890 |
| YJM627 | VII | 1 | 10 | 1038420 | leftend | YGL258W | 99.034 | 621 | 6 | 0 | 1 | 621 | 5720 | 6340 | 0 | 1114 |
| YJM627 | VII | 1038411 | 1038420 | 1038420 | rightend | YGR287C | 99.2 | 1626 | 13 | 0 | 145 | 1770 | 1030058 | 1028433 | 0 | 2931 |
| YJM627 | VIII | 1 | 10 | 531847 | leftend | YHL044W | 98.446 | 708 | 11 | 0 | 1 | 708 | 8431 | 9138 | 0 | 1247 |
| YJM627 | VIII | 520968 | 520977 | 531847 | middle |  |  |  |  |  |  |  |  |  |  |  |
| YJM627 | VIII | 531838 | 531847 | 531847 | rightend | YHR211W | 96.927 | 716 | 22 | 0 | 2513 | 3228 | 520978 | 521693 | 0 | 1201 |
| YJM627 | IX | 1 |  | 425599 | leftend_complete | YIL169C | 86.537 | 1337 | 165 | 11 | 562 | 1889 | 19847 | 18517 | 0 | 1458 |
| YJM627 | IX | 2205 | 2214 | 425599 | middle |  |  |  |  |  |  |  |  |  |  |  |
| YJM627 | IX | 425590 | 425599 | 425599 | rightend | YIR039C | 99.69 | 1614 | 5 | 0 | 1 | 1614 | 423986 | 422373 | 0 | 2953 |
| YJM627 | X | 1 | 10 | 718110 | leftend | YJL213W | 99.699 | 996 | 3 | 0 | 1 | 996 | 25594 | 26589 | 0 | 1823 |
| YJM627 | X | 718110 | 718110 | 718110 | rightend_complete | YJR152W | 99.694 | 1632 | 4 | 1 | 1 | 1632 | 697888 | 699518 | 0 | 2985 |
| YJM627 | XI | 1 | 10 | 664255 | leftend | YKL222C | 99.622 | 2118 | 8 | 0 | 1 | 2118 | 3345 | 1228 | 0 | 3868 |
| YJM627 | XI | 659967 | 659976 | 664255 | middle |  |  |  |  |  |  |  |  |  |  |  |
| YJM627 | XI | 662083 | 662092 | 664255 | middle |  |  |  |  |  |  |  |  |  |  |  |
| YJM627 | XI | 664255 | 664255 | 664255 | rightend_complete | YKR101W | 99.644 | 1965 | 7 | 0 | 1 | 1965 | 639030 | 640994 | 0 | 3591 |
| YJM627 | XII | 1 | 10 | 1935054 | leftend | YLL062C | 99.282 | 975 | 7 | 0 | 1 | 975 | 8781 | 7807 | 0 | 1762 |
| YJM627 | XII | 738 | 747 | 1935054 | middle |  |  |  |  |  |  |  |  |  |  |  |
| YJM627 | XII | 1933966 | 1933975 | 1935054 | middle |  |  |  |  |  |  |  |  |  |  |  |
| YJM627 | XII | 1935045 | 1935054 | 1935054 | rightend | YLR460C | 94.071 | 1130 | 65 | 2 | 3 | 1131 | 1931631 | 1930503 | 0 | 1714 |
| YJM627 | XIII | 1 | 10 | 905909 | leftend | YML131W | 99.545 | 1098 | 5 | 0 | 1 | 1098 | 2832 | 3929 | 0 | 2001 |
| YJM627 | XIII | 905900 | 905909 | 905909 | rightend | YMR321C | 99.361 | 313 | 2 | 0 | 6 | 318 | 888372 | 888060 | 2.48E-161 | 568 |
| YJM627 | XIV | 1 | 10 | 765506 | leftend | YNL332W | 98.143 | 1023 | 19 | 0 | 1 | 1023 | 6472 | 7494 | 0 | 1784 |
| YJM627 | XIV | 758223 | 758232 | 765506 | middle |  |  |  |  |  |  |  |  |  |  |  |
| YJM627 | XIV | 765497 | 765506 | 765506 | rightend | YNR071C | 99.514 | 1029 | 5 | 0 | 1 | 1029 | 745059 | 744031 | 0 | 1873 |
| YJM627 | XV | 1 | 10 | 1057804 | leftend | YOL154W | 99.467 | 750 | 4 | 0 | 1 | 750 | 28316 | 29065 | 0 | 1363 |
| YJM627 | XV | 1054900 | 1054909 | 1057804 | middle |  |  |  |  |  |  |  |  |  |  |  |
| YJM627 | XV | 1055266 | 1055275 | 1057804 | middle |  |  |  |  |  |  |  |  |  |  |  |
| YJM627 | XV | 1055793 | 1055802 | 1057804 | middle |  |  |  |  |  |  |  |  |  |  |  |
| YJM627 | XV | 1057304 | 1057313 | 1057804 | middle |  |  |  |  |  |  |  |  |  |  |  |
| YJM627 | XV | 1057795 | 1057804 | 1057804 | rightend | YOR381W | 99.906 | 2136 | 2 | 0 | 1 | 2136 | 1027380 | 1029515 | 0 | 3934 |
| YJM627 | XVI | 1 | 10 | 892629 | leftend | YPL274W | 99.943 | 1764 | 1 | 0 | 1 | 1764 | 16645 | 18408 | 0 | 3253 |
| YJM627 | XVI | 227 | 236 | 892629 | middle |  |  |  |  |  |  |  |  |  |  |  |
| YJM627 | XVI | 892620 | 892629 | 892629 | rightend | YPR194C | 99.375 | 2560 | 16 | 0 | 75 | 2634 | 891822 | 889263 | 0 | 4639 |
| YJM681 | I | 1 | 10 | 196537 | leftend | YAL063C | 83.064 | 1730 | 285 | 8 | 40 | 1765 | 184332 | 186057 | 0 | 1565 |
| YJM681 | I | 196528 | 196537 | 196537 | rightend | YAR050W | 81.72 | 1767 | 319 | 4 | 1 | 1765 | 184293 | 186057 | 0 | 1471 |
| YJM681 | II | 1 | 10 | 787096 | leftend | YNCB0001W | 100 | 84 | 0 | 0 | 1 | 84 | 3272 | 3355 | 4.16E-38 | 156 |
| YJM681 | II | 1054 | 1063 | 787096 | middle |  |  |  |  |  |  |  |  |  |  |  |
| YJM681 | II | 1569 | 1578 | 787096 | middle |  |  |  |  |  |  |  |  |  |  |  |
| YJM681 | II | 787087 | 787096 | 787096 | rightend | YBR297W | 98.091 | 1205 | 23 | 0 | 1 | 1205 | 777269 | 778473 | 0 | 2098 |
| YJM681 | III | 1 |  | 309168 | leftend_complete | YCL073C | 99.621 | 1848 | 7 | 0 | 1 | 1848 | 7678 | 5831 | 0 | 3374 |
| YJM681 | III | 309168 | 309168 | 309168 | rightend_complete | YCR101C | 92.021 | 564 | 28 | 12 | 1 | 549 | 295654 | 295093 | 0 | 776 |
| YJM681 | IV | 1 | 10 | 1458137 | leftend | YDL243C | 93.535 | 990 | 64 | 0 | 1 | 990 | 1736 | 747 | 0 | 1474 |
| YJM681 | IV | 1456214 | 1456223 | 1458137 | middle |  |  |  |  |  |  |  |  |  |  |  |
| YJM681 | IV | 1457205 | 1457214 | 1458137 | middle |  |  |  |  |  |  |  |  |  |  |  |
| YJM681 | IV | 1458128 | 1458137 | 1458137 | rightend | YDR541C | 99.42 | 1035 | 6 | 0 | 1 | 1035 | 1453588 | 1452554 | 0 | 1879 |
| YJM681 | V | 1 | 10 | 552545 | leftend | YEL070W | 99.072 | 1509 | 14 | 0 | 1 | 1509 | 13975 | 15483 | 0 | 2710 |
| YJM681 | V | 483 | 492 | 552545 | middle |  |  |  |  |  |  |  |  |  |  |  |
| YJM681 | V | 552545 | 552545 | 552545 | rightend_complete | YER187W | 99.765 | 426 | 1 | 0 | 1 | 426 | 548661 | 549086 | 0 | 782 |
| YJM681 | VI | 1 | 10 | 245882 | leftend | YFL051C | 98.23 | 452 | 8 | 0 | 1 | 452 | 8832 | 8381 | 0 | 791 |
| YJM681 | VI | 245873 | 245882 | 245882 | rightend | YFR055W | 100 | 1022 | 0 | 0 | 1 | 1022 | 236395 | 237416 | 0 | 1888 |
| YJM681 | VII | 1 | 10 | 1066552 | leftend | YGL258W | 96.457 | 621 | 22 | 0 | 1 | 621 | 12496 | 13116 | 0 | 1026 |
| YJM681 | VII | 1343 | 1352 | 1066552 | middle |  |  |  |  |  |  |  |  |  |  |  |
| YJM681 | VII | 1064616 | 1064625 | 1066552 | middle |  |  |  |  |  |  |  |  |  |  |  |
| YJM681 | VII | 1066543 | 1066552 | 1066552 | rightend | YGR287C | 99.831 | 1770 | 3 | 0 | 1 | 1770 | 1034051 | 1032282 | 0 | 3253 |
| YJM681 | VIII | 1 |  | 532638 | leftend_complete | YHL044W | 98.588 | 708 | 10 | 0 | 1 | 708 | 8501 | 9208 | 0 | 1253 |
| YJM681 | VIII | 7150 | 7159 | 532638 | middle |  |  |  |  |  |  |  |  |  |  |  |
| YJM681 | VIII | 529788 | 529797 | 532638 | middle |  |  |  |  |  |  |  |  |  |  |  |
| YJM681 | VIII | 531903 | 531912 | 532638 | middle |  |  |  |  |  |  |  |  |  |  |  |
| YJM681 | VIII | 532638 | 532638 | 532638 | rightend_complete | YHR211W | 92.135 | 2136 | 144 | 2 | 1096 | 3228 | 519368 | 521482 | 0 | 2992 |
| YJM681 | IX | 1 | 10 | 418681 | leftend | YIL169C | 96.539 | 1907 | 56 | 8 | 562 | 2466 | 17152 | 15254 | 0 | 3147 |
| YJM681 | IX | 722 | 731 | 418681 | middle |  |  |  |  |  |  |  |  |  |  |  |
| YJM681 | IX | 418672 | 418681 | 418681 | rightend | YIR039C | 98.947 | 1614 | 17 | 0 | 1 | 1614 | 416821 | 415208 | 0 | 2887 |
| YJM681 | X | 1 |  | 813303 | leftend_complete | YJL213W | 89.859 | 996 | 101 | 0 | 1 | 996 | 35831 | 36826 | 0 | 1280 |
| YJM681 | X | 199 | 208 | 813303 | middle |  |  |  |  |  |  |  |  |  |  |  |
| YJM681 | X | 730 | 739 | 813303 | middle |  |  |  |  |  |  |  |  |  |  |  |
| YJM681 | X | 968 | 977 | 813303 | middle |  |  |  |  |  |  |  |  |  |  |  |
| YJM681 | X | 1585 | 1594 | 813303 | middle |  |  |  |  |  |  |  |  |  |  |  |
| YJM681 | X | 2071 | 2080 | 813303 | middle |  |  |  |  |  |  |  |  |  |  |  |
| YJM681 | X | 3187 | 3196 | 813303 | middle |  |  |  |  |  |  |  |  |  |  |  |
| YJM681 | X | 3787 | 3796 | 813303 | middle |  |  |  |  |  |  |  |  |  |  |  |
| YJM681 | X | 813294 | 813303 | 813303 | rightend | YJR152W | 99.755 | 1632 | 3 | 1 | 1 | 1632 | 707521 | 709151 | 0 | 2990 |
| YJM681 | XI | 1 | 10 | 663672 | leftend | YKL222C | 99.481 | 2118 | 11 | 0 | 1 | 2118 | 3135 | 1018 | 0 | 3851 |
| YJM681 | XI | 661832 | 661841 | 663672 | middle |  |  |  |  |  |  |  |  |  |  |  |
| YJM681 | XI | 663663 | 663672 | 663672 | rightend | YKR101W | 99.847 | 1965 | 3 | 0 | 1 | 1965 | 637092 | 639056 | 0 | 3613 |
| YJM681 | XII | 1 | 10 | 1636790 | leftend | YLL062C | 99.59 | 975 | 4 | 0 | 1 | 975 | 4221 | 3247 | 0 | 1779 |
| YJM681 | XII | 1636379 | 1636388 | 1636790 | middle |  |  |  |  |  |  |  |  |  |  |  |
| YJM681 | XII | 1636790 | 1636790 | 1636790 | rightend_complete | YLR460C | 98.587 | 1132 | 14 | 2 | 1 | 1131 | 1635745 | 1634615 | 0 | 2001 |
| YJM681 | XIII | 1 | 10 | 898300 | leftend | YML131W | 99.818 | 1098 | 2 | 0 | 1 | 1098 | 3543 | 4640 | 0 | 2017 |
| YJM681 | XIII | 893867 | 893876 | 898300 | middle |  |  |  |  |  |  |  |  |  |  |  |
| YJM681 | XIII | 898006 | 898015 | 898300 | middle |  |  |  |  |  |  |  |  |  |  |  |
| YJM681 | XIII | 898291 | 898300 | 898300 | rightend | YMR321C | 99.361 | 313 | 2 | 0 | 6 | 318 | 888460 | 888148 | 2.48E-161 | 568 |
| YJM681 | XIV | 1 |  | 761499 | leftend_complete | YNL332W | 99.316 | 1023 | 7 | 0 | 1 | 1023 | 11543 | 12565 | 0 | 1851 |
| YJM681 | XIV | 4170 | 4179 | 761499 | middle |  |  |  |  |  |  |  |  |  |  |  |
| YJM681 | XIV | 761490 | 761499 | 761499 | rightend | YNR071C | 100 | 1029 | 0 | 0 | 1 | 1029 | 750864 | 749836 | 0 | 1901 |
| YJM681 | XV | 1 | 10 | 1046500 | leftend | YOL154W | 99.6 | 750 | 3 | 0 | 1 | 750 | 22689 | 23438 | 0 | 1369 |
| YJM681 | XV | 1046491 | 1046500 | 1046500 | rightend | YOR381W | 99.251 | 2136 | 16 | 0 | 1 | 2136 | 1018957 | 1021092 | 0 | 3856 |
| YJM681 | XVI | 1 |  | 904574 | leftend_complete | YPL274W | 99.887 | 1764 | 2 | 0 | 1 | 1764 | 13857 | 15620 | 0 | 3247 |
| YJM681 | XVI | 2939 | 2948 | 904574 | middle |  |  |  |  |  |  |  |  |  |  |  |
| YJM681 | XVI | 3879 | 3888 | 904574 | middle |  |  |  |  |  |  |  |  |  |  |  |
| YJM681 | XVI | 904565 | 904574 | 904574 | rightend | YPR201W | 99.588 | 1215 | 5 | 0 | 1 | 1215 | 901148 | 902362 | 0 | 2217 |
| YJM682 | I | 1 | 10 | 187172 | leftend | YAL063C | 87.966 | 2576 | 304 | 4 | 1 | 2573 | 14061 | 11489 | 0 | 3035 |
| YJM682 | I | 187163 | 187172 | 187172 | rightend | YAR050W | 82.219 | 3217 | 544 | 24 | 46 | 3248 | 14016 | 10814 | 0 | 2747 |
| YJM682 | II | 1 | 10 | 785701 | leftend | YNCB0001W | 100 | 84 | 0 | 0 | 1 | 84 | 9123 | 9206 | 4.16E-38 | 156 |
| YJM682 | II | 785692 | 785701 | 785701 | rightend | YBR297W | 98.008 | 1205 | 24 | 0 | 1 | 1205 | 781494 | 782698 | 0 | 2093 |
| YJM682 | III | 1 |  | 315685 | leftend_complete | YCL073C | 99.839 | 1242 | 2 | 0 | 607 | 1848 | 7373 | 6132 | 0 | 2283 |
| YJM682 | III | 310898 | 310907 | 315685 | middle |  |  |  |  |  |  |  |  |  |  |  |
| YJM682 | III | 313602 | 313611 | 315685 | middle |  |  |  |  |  |  |  |  |  |  |  |
| YJM682 | III | 315125 | 315134 | 315685 | middle |  |  |  |  |  |  |  |  |  |  |  |
| YJM682 | III | 315685 | 315685 | 315685 | rightend_complete | YCR101C | 99.818 | 549 | 1 | 0 | 1 | 549 | 298924 | 298376 | 0 | 1009 |
| YJM682 | IV | 1 | 10 | 1457606 | leftend | YDL243C | 95.051 | 990 | 49 | 0 | 1 | 990 | 4758 | 3769 | 0 | 1557 |
| YJM682 | IV | 1457597 | 1457606 | 1457606 | rightend | YDR541C | 99.614 | 1035 | 4 | 0 | 1 | 1035 | 1452808 | 1451774 | 0 | 1890 |
| YJM682 | V | 1 | 10 | 553524 | leftend | YEL070W | 99.139 | 1509 | 13 | 0 | 1 | 1509 | 13617 | 15125 | 0 | 2715 |
| YJM682 | V | 552721 | 552730 | 553524 | middle |  |  |  |  |  |  |  |  |  |  |  |
| YJM682 | V | 553515 | 553524 | 553524 | rightend | YER187W | 99.296 | 426 | 3 | 0 | 1 | 426 | 548345 | 548770 | 0 | 771 |
| YJM682 | VI | 1 |  | 251028 | leftend_complete | YFL051C | 98.673 | 452 | 6 | 0 | 1 | 452 | 12224 | 11773 | 0 | 802 |
| YJM682 | VI | 251019 | 251028 | 251028 | rightend | YFR055W | 99.609 | 1022 | 4 | 0 | 1 | 1022 | 241134 | 242155 | 0 | 1866 |
| YJM682 | VII | 1 |  | 1039128 | leftend_complete | YGL258W | 99.356 | 621 | 4 | 0 | 1 | 621 | 5859 | 6479 | 0 | 1125 |
| YJM682 | VII | 642 | 651 | 1039128 | middle |  |  |  |  |  |  |  |  |  |  |  |
| YJM682 | VII | 3439 | 3448 | 1039128 | middle |  |  |  |  |  |  |  |  |  |  |  |
| YJM682 | VII | 1039119 | 1039128 | 1039128 | rightend | YGR287C | 98.192 | 1770 | 32 | 0 | 1 | 1770 | 1026448 | 1024679 | 0 | 3092 |
| YJM682 | VIII | 1 | 10 | 530432 | leftend | YHL044W | 98.446 | 708 | 11 | 0 | 1 | 708 | 15238 | 15945 | 0 | 1247 |
| YJM682 | VIII | 1375 | 1384 | 530432 | middle |  |  |  |  |  |  |  |  |  |  |  |
| YJM682 | VIII | 530423 | 530432 | 530432 | rightend | YHR211W | 90.914 | 1761 | 136 | 2 | 1096 | 2853 | 528683 | 530422 | 0 | 2344 |
| YJM682 | IX | 1 |  | 439220 | leftend_complete | YIL169C | 96.539 | 1907 | 56 | 8 | 562 | 2466 | 20760 | 18862 | 0 | 3147 |
| YJM682 | IX | 1945 | 1954 | 439220 | middle |  |  |  |  |  |  |  |  |  |  |  |
| YJM682 | IX | 439211 | 439220 | 439220 | rightend | YIR039C | 99.071 | 1614 | 15 | 0 | 1 | 1614 | 423276 | 421663 | 0 | 2898 |
| YJM682 | X | 1 | 10 | 722851 | leftend | YJL213W | 100 | 996 | 0 | 0 | 1 | 996 | 26817 | 27812 | 0 | 1840 |
| YJM682 | X | 516 | 525 | 722851 | middle |  |  |  |  |  |  |  |  |  |  |  |
| YJM682 | X | 1646 | 1655 | 722851 | middle |  |  |  |  |  |  |  |  |  |  |  |
| YJM682 | X | 722240 | 722249 | 722851 | middle |  |  |  |  |  |  |  |  |  |  |  |
| YJM682 | X | 722851 | 722851 | 722851 | rightend_complete | YJR152W | 99.755 | 1632 | 3 | 1 | 1 | 1632 | 697594 | 699224 | 0 | 2990 |
| YJM682 | XI | 1 | 10 | 660079 | leftend | YKL222C | 99.481 | 2118 | 11 | 0 | 1 | 2118 | 3109 | 992 | 0 | 3851 |
| YJM682 | XI | 659450 | 659459 | 660079 | middle |  |  |  |  |  |  |  |  |  |  |  |
| YJM682 | XI | 660079 | 660079 | 660079 | rightend_complete | YKR101W | 99.796 | 1965 | 4 | 0 | 1 | 1965 | 637301 | 639265 | 0 | 3607 |
| YJM682 | XII | 1 | 10 | 1633276 | leftend | YLL062C | 99.897 | 975 | 1 | 0 | 1 | 975 | 10067 | 9093 | 0 | 1796 |
| YJM682 | XII | 1633267 | 1633276 | 1633276 | rightend | YLR460C | 97.963 | 1129 | 23 | 0 | 3 | 1131 | 1632071 | 1630943 | 0 | 1958 |
| YJM682 | XIII | 1 |  | 897736 | leftend_complete | YML131W | 99.909 | 1098 | 1 | 0 | 1 | 1098 | 2869 | 3966 | 0 | 2023 |
| YJM682 | XIII | 481 | 490 | 897736 | middle |  |  |  |  |  |  |  |  |  |  |  |
| YJM682 | XIII | 896022 | 896031 | 897736 | middle |  |  |  |  |  |  |  |  |  |  |  |
| YJM682 | XIII | 897736 | 897736 | 897736 | rightend_complete | YMR321C | 99.361 | 313 | 2 | 0 | 6 | 318 | 888077 | 887765 | 2.48E-161 | 568 |
| YJM682 | XIV | 1 | 10 | 769393 | leftend | YNL332W | 98.631 | 1023 | 14 | 0 | 1 | 1023 | 7199 | 8221 | 0 | 1812 |
| YJM682 | XIV | 769393 | 769393 | 769393 | rightend_complete | YNR071C | 99.611 | 1029 | 4 | 0 | 1 | 1029 | 747795 | 746767 | 0 | 1879 |
| YJM682 | XV | 1 | 10 | 1058008 | leftend | YOL154W | 99.6 | 750 | 3 | 0 | 1 | 750 | 29460 | 30209 | 0 | 1369 |
| YJM682 | XV | 1054122 | 1054131 | 1058008 | middle |  |  |  |  |  |  |  |  |  |  |  |
| YJM682 | XV | 1057999 | 1058008 | 1058008 | rightend | YOR381W | 98.642 | 2136 | 29 | 0 | 1 | 2136 | 1026913 | 1029048 | 0 | 3784 |
| YJM682 | XVI | 1 | 10 | 911646 | leftend | YPL274W | 98.583 | 1764 | 25 | 0 | 1 | 1764 | 18374 | 20137 | 0 | 3120 |
| YJM682 | XVI | 2774 | 2783 | 911646 | middle |  |  |  |  |  |  |  |  |  |  |  |
| YJM682 | XVI | 4432 | 4441 | 911646 | middle |  |  |  |  |  |  |  |  |  |  |  |
| YJM682 | XVI | 4674 | 4683 | 911646 | middle |  |  |  |  |  |  |  |  |  |  |  |
| YJM682 | XVI | 5142 | 5151 | 911646 | middle |  |  |  |  |  |  |  |  |  |  |  |
| YJM682 | XVI | 910300 | 910309 | 911646 | middle |  |  |  |  |  |  |  |  |  |  |  |
| YJM682 | XVI | 910517 | 910526 | 911646 | middle |  |  |  |  |  |  |  |  |  |  |  |
| YJM682 | XVI | 911637 | 911646 | 911646 | rightend | YPR201W | 99.588 | 1215 | 5 | 0 | 1 | 1215 | 906617 | 907831 | 0 | 2217 |
| YJM683 | I | 1 | 10 | 197208 | leftend | YAL063C | 85.037 | 1497 | 219 | 5 | 1 | 1495 | 12471 | 10978 | 0 | 1519 |
| YJM683 | I | 188672 | 188681 | 197208 | middle |  |  |  |  |  |  |  |  |  |  |  |
| YJM683 | I | 192703 | 192712 | 197208 | middle |  |  |  |  |  |  |  |  |  |  |  |
| YJM683 | I | 197208 | 197208 | 197208 | rightend_complete | YAR050W | 83.333 | 1776 | 261 | 22 | 1 | 1757 | 186861 | 188620 | 0 | 1607 |
| YJM683 | II | 1 | 10 | 782839 | leftend | YNCB0001W | 100 | 84 | 0 | 0 | 1 | 84 | 3650 | 3733 | 4.16E-38 | 156 |
| YJM683 | II | 1406 | 1415 | 782839 | middle |  |  |  |  |  |  |  |  |  |  |  |
| YJM683 | II | 2336 | 2345 | 782839 | middle |  |  |  |  |  |  |  |  |  |  |  |
| YJM683 | II | 781099 | 781108 | 782839 | middle |  |  |  |  |  |  |  |  |  |  |  |
| YJM683 | II | 782830 | 782839 | 782839 | rightend | YBR297W | 98.294 | 1407 | 24 | 0 | 1 | 1407 | 776602 | 778008 | 0 | 2466 |
| YJM683 | III | 1 |  | 310505 | leftend_complete | YCL073C | 98.755 | 1848 | 23 | 0 | 1 | 1848 | 5308 | 3461 | 0 | 3286 |
| YJM683 | III | 308863 | 308872 | 310505 | middle |  |  |  |  |  |  |  |  |  |  |  |
| YJM683 | III | 310496 | 310505 | 310505 | rightend | YCR101C | 99.818 | 549 | 1 | 0 | 1 | 549 | 295713 | 295165 | 0 | 1009 |
| YJM683 | IV | 1 | 10 | 1455075 | leftend | YDL243C | 95.051 | 990 | 49 | 0 | 1 | 990 | 2618 | 1629 | 0 | 1557 |
| YJM683 | IV | 1455066 | 1455075 | 1455075 | rightend | YDR541C | 99.614 | 1035 | 4 | 0 | 1 | 1035 | 1452281 | 1451247 | 0 | 1890 |
| YJM683 | V | 1 | 10 | 557102 | leftend | YEL070W | 99.139 | 1509 | 13 | 0 | 1 | 1509 | 17460 | 18968 | 0 | 2715 |
| YJM683 | V | 511 | 520 | 557102 | middle |  |  |  |  |  |  |  |  |  |  |  |
| YJM683 | V | 556156 | 556165 | 557102 | middle |  |  |  |  |  |  |  |  |  |  |  |
| YJM683 | V | 557102 | 557102 | 557102 | rightend_complete | YER187W | 99.296 | 426 | 3 | 0 | 1 | 426 | 552265 | 552690 | 0 | 771 |
| YJM683 | VI | 1 |  | 237014 | leftend_complete | YFL051C | 98.451 | 452 | 7 | 0 | 1 | 452 | 2959 | 2508 | 0 | 797 |
| YJM683 | VI | 237014 | 237014 | 237014 | rightend_complete | YFR055W | 100 | 1023 | 0 | 0 | 1 | 1023 | 231056 | 232078 | 0 | 1890 |
| YJM683 | VII | 1 | 10 | 1060392 | leftend | YGL258W | 99.356 | 621 | 4 | 0 | 1 | 621 | 3123 | 3743 | 0 | 1125 |
| YJM683 | VII | 1057429 | 1057438 | 1060392 | middle |  |  |  |  |  |  |  |  |  |  |  |
| YJM683 | VII | 1060383 | 1060392 | 1060392 | rightend | YGR287C | 99.209 | 1770 | 14 | 0 | 1 | 1770 | 1024479 | 1022710 | 0 | 3192 |
| YJM683 | VIII | 1 |  | 545260 | leftend_complete | YHL044W | 98.446 | 708 | 11 | 0 | 1 | 708 | 8825 | 9532 | 0 | 1247 |
| YJM683 | VIII | 545251 | 545260 | 545260 | rightend | YHR211W | 80.47 | 1915 | 328 | 29 | 1 | 1892 | 521048 | 522939 | 0 | 1423 |
| YJM683 | IX | 1 | 10 | 414673 | leftend | YIL169C | 98.254 | 2806 | 39 | 8 | 190 | 2988 | 8904 | 6102 | 0 | 4902 |
| YJM683 | IX | 413717 | 413726 | 414673 | middle |  |  |  |  |  |  |  |  |  |  |  |
| YJM683 | IX | 414673 | 414673 | 414673 | rightend_complete | YIR039C | 99.009 | 1614 | 16 | 0 | 1 | 1614 | 411563 | 409950 | 0 | 2892 |
| YJM683 | X | 1 |  | 722206 | leftend_complete | YJL213W | 100 | 996 | 0 | 0 | 1 | 996 | 25309 | 26304 | 0 | 1840 |
| YJM683 | X | 1948 | 1957 | 722206 | middle |  |  |  |  |  |  |  |  |  |  |  |
| YJM683 | X | 720722 | 720731 | 722206 | middle |  |  |  |  |  |  |  |  |  |  |  |
| YJM683 | X | 721579 | 721588 | 722206 | middle |  |  |  |  |  |  |  |  |  |  |  |
| YJM683 | X | 722206 | 722206 | 722206 | rightend_complete | YJR152W | 99.755 | 1632 | 3 | 1 | 1 | 1632 | 695586 | 697216 | 0 | 2990 |
| YJM683 | XI | 1 |  | 662991 | leftend_complete | YKL222C | 100 | 2118 | 0 | 0 | 1 | 2118 | 5661 | 3544 | 0 | 3912 |
| YJM683 | XI | 662982 | 662991 | 662991 | rightend | YKR101W | 99.796 | 1965 | 4 | 0 | 1 | 1965 | 640457 | 642421 | 0 | 3607 |
| YJM683 | XII | 1 | 10 | 1767114 | leftend | YLL062C | 99.897 | 975 | 1 | 0 | 1 | 975 | 4449 | 3475 | 0 | 1796 |
| YJM683 | XII | 1767114 | 1767114 | 1767114 | rightend_complete | YLR460C | 99.735 | 1131 | 3 | 0 | 1 | 1131 | 1762582 | 1761452 | 0 | 2073 |
| YJM683 | XIII | 1 | 10 | 905653 | leftend | YML131W | 99.909 | 1098 | 1 | 0 | 1 | 1098 | 2353 | 3450 | 0 | 2023 |
| YJM683 | XIII | 904132 | 904141 | 905653 | middle |  |  |  |  |  |  |  |  |  |  |  |
| YJM683 | XIII | 905653 | 905653 | 905653 | rightend_complete | YMR321C | 99.361 | 313 | 2 | 0 | 6 | 318 | 886931 | 886619 | 2.48E-161 | 568 |
| YJM683 | XIV | 1 | 10 | 766962 | leftend | YNL332W | 98.631 | 1023 | 14 | 0 | 1 | 1023 | 16755 | 17777 | 0 | 1812 |
| YJM683 | XIV | 764872 | 764881 | 766962 | middle |  |  |  |  |  |  |  |  |  |  |  |
| YJM683 | XIV | 766962 | 766962 | 766962 | rightend_complete | YNR071C | 99.708 | 1029 | 3 | 0 | 1 | 1029 | 757028 | 756000 | 0 | 1884 |
| YJM683 | XV | 1 | 10 | 1038379 | leftend | YOL154W | 99.733 | 750 | 2 | 0 | 1 | 750 | 10265 | 11014 | 0 | 1375 |
| YJM683 | XV | 1038370 | 1038379 | 1038379 | rightend | YOR381W | 98.596 | 2136 | 30 | 0 | 1 | 2136 | 1011106 | 1013241 | 0 | 3779 |
| YJM683 | XVI | 1 | 10 | 899188 | leftend | YPL274W | 99.546 | 1764 | 8 | 0 | 1 | 1764 | 6078 | 7841 | 0 | 3214 |
| YJM683 | XVI | 944 | 953 | 899188 | middle |  |  |  |  |  |  |  |  |  |  |  |
| YJM683 | XVI | 899179 | 899188 | 899188 | rightend | YPR201W | 99.588 | 1215 | 5 | 0 | 1 | 1215 | 896665 | 897879 | 0 | 2217 |
| YJM689 | I | 1 | 10 | 206966 | leftend | YAL063C | 89.884 | 2748 | 264 | 9 | 1222 | 3962 | 12179 | 9439 | 0 | 3522 |
| YJM689 | I | 198879 | 198888 | 206966 | middle |  |  |  |  |  |  |  |  |  |  |  |
| YJM689 | I | 206957 | 206966 | 206966 | rightend | YAR050W | 83.368 | 1906 | 305 | 11 | 1 | 1900 | 182782 | 184681 | 0 | 1753 |
| YJM689 | II | 1 | 10 | 779592 | leftend | YNCB0001W | 100 | 84 | 0 | 0 | 1 | 84 | 1654 | 1737 | 4.16E-38 | 156 |
| YJM689 | II | 779583 | 779592 | 779592 | rightend | YBR297W | 98.257 | 1205 | 21 | 0 | 1 | 1205 | 774632 | 775836 | 0 | 2109 |
| YJM689 | III | 1 | 10 | 313222 | leftend | YCL073C | 98.972 | 1848 | 19 | 0 | 1 | 1848 | 7605 | 5758 | 0 | 3308 |
| YJM689 | III | 312738 | 312747 | 313222 | middle |  |  |  |  |  |  |  |  |  |  |  |
| YJM689 | III | 313213 | 313222 | 313222 | rightend | YCR101C | 99.818 | 549 | 1 | 0 | 1 | 549 | 299183 | 298635 | 0 | 1009 |
| YJM689 | IV | 1 | 10 | 1465273 | leftend | YDL243C | 93.939 | 990 | 60 | 0 | 1 | 990 | 19843 | 18854 | 0 | 1496 |
| YJM689 | IV | 1033 | 1042 | 1465273 | middle |  |  |  |  |  |  |  |  |  |  |  |
| YJM689 | IV | 2681 | 2690 | 1465273 | middle |  |  |  |  |  |  |  |  |  |  |  |
| YJM689 | IV | 1465264 | 1465273 | 1465273 | rightend | YDR541C | 99.614 | 1035 | 4 | 0 | 1 | 1035 | 1462680 | 1461646 | 0 | 1890 |
| YJM689 | V | 1 | 10 | 553822 | leftend | YEL070W | 100 | 1509 | 0 | 0 | 1 | 1509 | 13493 | 15001 | 0 | 2787 |
| YJM689 | V | 553813 | 553822 | 553822 | rightend | YER187W | 91.981 | 424 | 32 | 2 | 4 | 426 | 549319 | 549741 | 3.44E-169 | 593 |
| YJM689 | VI | 1 | 10 | 235718 | leftend | YFL051C | 98.451 | 452 | 7 | 0 | 1 | 452 | 2392 | 1941 | 0 | 797 |
| YJM689 | VI | 1335 | 1344 | 235718 | middle |  |  |  |  |  |  |  |  |  |  |  |
| YJM689 | VI | 235709 | 235718 | 235718 | rightend | YFR055W | 99.804 | 1022 | 2 | 0 | 1 | 1022 | 229621 | 230642 | 0 | 1877 |
| YJM689 | VII | 1 | 10 | 1032356 | leftend | YGL258W | 99.356 | 621 | 4 | 0 | 1 | 621 | 2124 | 2744 | 0 | 1125 |
| YJM689 | VII | 1029397 | 1029406 | 1032356 | middle |  |  |  |  |  |  |  |  |  |  |  |
| YJM689 | VII | 1030808 | 1030817 | 1032356 | middle |  |  |  |  |  |  |  |  |  |  |  |
| YJM689 | VII | 1031038 | 1031047 | 1032356 | middle |  |  |  |  |  |  |  |  |  |  |  |
| YJM689 | VII | 1031744 | 1031753 | 1032356 | middle |  |  |  |  |  |  |  |  |  |  |  |
| YJM689 | VII | 1032347 | 1032356 | 1032356 | rightend | YGR287C | 98.305 | 1770 | 30 | 0 | 1 | 1770 | 1020617 | 1018848 | 0 | 3103 |
| YJM689 | VIII | 1 | 10 | 525444 | leftend | YHL044W | 98.729 | 708 | 9 | 0 | 1 | 708 | 9300 | 10007 | 0 | 1258 |
| YJM689 | VIII | 580 | 589 | 525444 | middle |  |  |  |  |  |  |  |  |  |  |  |
| YJM689 | VIII | 7828 | 7837 | 525444 | middle |  |  |  |  |  |  |  |  |  |  |  |
| YJM689 | VIII | 525435 | 525444 | 525444 | rightend | YHR211W | 82.535 | 939 | 153 | 7 | 1 | 932 | 513784 | 514718 | 0 | 815 |
| YJM689 | IX | 1 |  | 430289 | leftend_complete | YIL169C | 96.725 | 1374 | 39 | 2 | 478 | 1851 | 24198 | 22831 | 0 | 2283 |
| YJM689 | IX | 1938 | 1947 | 430289 | middle |  |  |  |  |  |  |  |  |  |  |  |
| YJM689 | IX | 2500 | 2509 | 430289 | middle |  |  |  |  |  |  |  |  |  |  |  |
| YJM689 | IX | 3625 | 3634 | 430289 | middle |  |  |  |  |  |  |  |  |  |  |  |
| YJM689 | IX | 5861 | 5870 | 430289 | middle |  |  |  |  |  |  |  |  |  |  |  |
| YJM689 | IX | 6494 | 6503 | 430289 | middle |  |  |  |  |  |  |  |  |  |  |  |
| YJM689 | IX | 7601 | 7610 | 430289 | middle |  |  |  |  |  |  |  |  |  |  |  |
| YJM689 | IX | 8489 | 8498 | 430289 | middle |  |  |  |  |  |  |  |  |  |  |  |
| YJM689 | IX | 426498 | 426507 | 430289 | middle |  |  |  |  |  |  |  |  |  |  |  |
| YJM689 | IX | 427908 | 427917 | 430289 | middle |  |  |  |  |  |  |  |  |  |  |  |
| YJM689 | IX | 429130 | 429139 | 430289 | middle |  |  |  |  |  |  |  |  |  |  |  |
| YJM689 | IX | 430280 | 430289 | 430289 | rightend | YIR039C | 99.69 | 1614 | 5 | 0 | 1 | 1614 | 424558 | 422945 | 0 | 2953 |
| YJM689 | X | 1 | 10 | 722426 | leftend | YJL213W | 99.398 | 996 | 6 | 0 | 1 | 996 | 31732 | 32727 | 0 | 1807 |
| YJM689 | X | 722426 | 722426 | 722426 | rightend_complete | YJR152W | 99.694 | 1632 | 4 | 1 | 1 | 1632 | 702209 | 703839 | 0 | 2985 |
| YJM689 | XI | 1 | 10 | 660470 | leftend | YKL222C | 99.575 | 2118 | 9 | 0 | 1 | 2118 | 3184 | 1067 | 0 | 3862 |
| YJM689 | XI | 660195 | 660204 | 660470 | middle |  |  |  |  |  |  |  |  |  |  |  |
| YJM689 | XI | 660470 | 660470 | 660470 | rightend_complete | YKR101W | 99.847 | 1965 | 3 | 0 | 1 | 1965 | 636930 | 638894 | 0 | 3613 |
| YJM689 | XII | 1 |  | 1625657 | leftend_complete | YLL062C | 99.897 | 975 | 1 | 0 | 1 | 975 | 4203 | 3229 | 0 | 1796 |
| YJM689 | XII | 2180 | 2189 | 1625657 | middle |  |  |  |  |  |  |  |  |  |  |  |
| YJM689 | XII | 1625648 | 1625657 | 1625657 | rightend | YOR380W | 99.756 | 1641 | 4 | 0 | 1 | 1641 | 1014715 | 1016355 | 0 | 3009 |
| YJM689 | XIII | 1 | 10 | 895136 | leftend | YML131W | 100 | 1098 | 0 | 0 | 1 | 1098 | 3562 | 4659 | 0 | 2028 |
| YJM689 | XIII | 623 | 632 | 895136 | middle |  |  |  |  |  |  |  |  |  |  |  |
| YJM689 | XIII | 1210 | 1219 | 895136 | middle |  |  |  |  |  |  |  |  |  |  |  |
| YJM689 | XIII | 893412 | 893421 | 895136 | middle |  |  |  |  |  |  |  |  |  |  |  |
| YJM689 | XIII | 895136 | 895136 | 895136 | rightend_complete | YMR321C | 99.371 | 318 | 2 | 0 | 1 | 318 | 888652 | 888335 | 4.12E-164 | 577 |
| YJM689 | XIV | 1 | 10 | 766264 | leftend | YNL332W | 98.81 | 504 | 6 | 0 | 520 | 1023 | 1391 | 1894 | 0 | 900 |
| YJM689 | XIV | 1381 | 1390 | 766264 | middle |  |  |  |  |  |  |  |  |  |  |  |
| YJM689 | XIV | 764324 | 764333 | 766264 | middle |  |  |  |  |  |  |  |  |  |  |  |
| YJM689 | XIV | 766255 | 766264 | 766264 | rightend | YNR071C | 100 | 1029 | 0 | 0 | 1 | 1029 | 739471 | 738443 | 0 | 1901 |
| YJM689 | XV | 1 | 10 | 1042880 | leftend | YOL154W | 100 | 750 | 0 | 0 | 1 | 750 | 22854 | 23603 | 0 | 1386 |
| YJM689 | XV | 1042871 | 1042880 | 1042880 | rightend | YOR381W | 100 | 2136 | 0 | 0 | 1 | 2136 | 1018970 | 1021105 | 0 | 3945 |
| YJM689 | XVI | 1 |  | 906795 | leftend_complete | YPL274W | 98.356 | 1764 | 29 | 0 | 1 | 1764 | 14840 | 16603 | 0 | 3097 |
| YJM689 | XVI | 502 | 511 | 906795 | middle |  |  |  |  |  |  |  |  |  |  |  |
| YJM689 | XVI | 905257 | 905266 | 906795 | middle |  |  |  |  |  |  |  |  |  |  |  |
| YJM689 | XVI | 906786 | 906795 | 906795 | rightend | YPR201W | 99.588 | 1215 | 5 | 0 | 1 | 1215 | 902309 | 903523 | 0 | 2217 |
| YJM693 | I | 1 | 10 | 196844 | leftend | YAL063C | 92.485 | 998 | 46 | 13 | 2966 | 3962 | 10430 | 9461 | 0 | 1400 |
| YJM693 | I | 196835 | 196844 | 196844 | rightend | YAR050W | 92.727 | 935 | 62 | 3 | 1 | 932 | 186274 | 187205 | 0 | 1345 |
| YJM693 | II | 1 | 10 | 784119 | leftend | YNCB0001W | 100 | 84 | 0 | 0 | 1 | 84 | 1575 | 1658 | 4.16E-38 | 156 |
| YJM693 | II | 1168 | 1177 | 784119 | middle |  |  |  |  |  |  |  |  |  |  |  |
| YJM693 | II | 783143 | 783152 | 784119 | middle |  |  |  |  |  |  |  |  |  |  |  |
| YJM693 | II | 784110 | 784119 | 784119 | rightend | YBR297W | 100 | 1407 | 0 | 0 | 1 | 1407 | 774171 | 775577 | 0 | 2599 |
| YJM693 | III | 1 |  | 302912 | leftend_complete | YCL073C | 99.919 | 1242 | 1 | 0 | 607 | 1848 | 7416 | 6175 | 0 | 2289 |
| YJM693 | III | 302903 | 302912 | 302912 | rightend | YCR101C | 97.636 | 550 | 12 | 1 | 1 | 549 | 298074 | 297525 | 0 | 942 |
| YJM693 | IV | 1 | 10 | 1453315 | leftend | YDL243C | 95.051 | 990 | 49 | 0 | 1 | 990 | 2618 | 1629 | 0 | 1557 |
| YJM693 | IV | 1451803 | 1451812 | 1453315 | middle |  |  |  |  |  |  |  |  |  |  |  |
| YJM693 | IV | 1453306 | 1453315 | 1453315 | rightend | YDR541C | 99.42 | 1035 | 6 | 0 | 1 | 1035 | 1448946 | 1447912 | 0 | 1879 |
| YJM693 | V | 1 | 10 | 556085 | leftend | YEL070W | 99.282 | 1114 | 8 | 0 | 396 | 1509 | 14559 | 15672 | 0 | 2013 |
| YJM693 | V | 549 | 558 | 556085 | middle |  |  |  |  |  |  |  |  |  |  |  |
| YJM693 | V | 555594 | 555603 | 556085 | middle |  |  |  |  |  |  |  |  |  |  |  |
| YJM693 | V | 556076 | 556085 | 556085 | rightend | YER187W | 91.745 | 424 | 33 | 2 | 4 | 426 | 551638 | 552060 | 1.6E-167 | 588 |
| YJM693 | VI | 1 | 10 | 266553 | leftend | YFL051C | 98.23 | 452 | 8 | 0 | 1 | 452 | 1481 | 1030 | 0 | 791 |
| YJM693 | VI | 266553 | 266553 | 266553 | rightend_complete | YFR055W | 99.902 | 1022 | 1 | 0 | 1 | 1022 | 260558 | 261579 | 0 | 1882 |
| YJM693 | VII | 1 | 10 | 1036798 | leftend | YGL258W | 97.262 | 621 | 17 | 0 | 1 | 621 | 10676 | 11296 | 0 | 1053 |
| YJM693 | VII | 1036242 | 1036251 | 1036798 | middle |  |  |  |  |  |  |  |  |  |  |  |
| YJM693 | VII | 1036789 | 1036798 | 1036798 | rightend | YGR287C | 98.362 | 1770 | 29 | 0 | 1 | 1770 | 1029730 | 1027961 | 0 | 3109 |
| YJM693 | VIII | 1 |  | 541871 | leftend_complete | YHL044W | 98.588 | 708 | 10 | 0 | 1 | 708 | 8217 | 8924 | 0 | 1253 |
| YJM693 | VIII | 1189 | 1198 | 541871 | middle |  |  |  |  |  |  |  |  |  |  |  |
| YJM693 | VIII | 538668 | 538677 | 541871 | middle |  |  |  |  |  |  |  |  |  |  |  |
| YJM693 | VIII | 540670 | 540679 | 541871 | middle |  |  |  |  |  |  |  |  |  |  |  |
| YJM693 | VIII | 540874 | 540883 | 541871 | middle |  |  |  |  |  |  |  |  |  |  |  |
| YJM693 | VIII | 541871 | 541871 | 541871 | rightend_complete | YHR211W | 82.535 | 939 | 153 | 7 | 1 | 932 | 528641 | 529575 | 0 | 815 |
| YJM693 | IX | 1 |  | 427691 | leftend_complete | YIL169C | 84.633 | 2128 | 278 | 31 | 520 | 2618 | 18042 | 15935 | 0 | 2073 |
| YJM693 | IX | 427682 | 427691 | 427691 | rightend | YIR039C | 99.752 | 1614 | 4 | 0 | 1 | 1614 | 419821 | 418208 | 0 | 2959 |
| YJM693 | X | 1 |  | 722992 | leftend_complete | YJL213W | 100 | 996 | 0 | 0 | 1 | 996 | 27577 | 28572 | 0 | 1840 |
| YJM693 | X | 721391 | 721400 | 722992 | middle |  |  |  |  |  |  |  |  |  |  |  |
| YJM693 | X | 722992 | 722992 | 722992 | rightend_complete | YJR152W | 99.51 | 1632 | 8 | 0 | 1 | 1632 | 697634 | 699265 | 0 | 2970 |
| YJM693 | XI | 1 |  | 650446 | leftend_complete | YKL222C | 99.481 | 2118 | 11 | 0 | 1 | 2118 | 4913 | 2796 | 0 | 3851 |
| YJM693 | XI | 1394 | 1403 | 650446 | middle |  |  |  |  |  |  |  |  |  |  |  |
| YJM693 | XI | 650437 | 650446 | 650446 | rightend | YKR101W | 99.542 | 1965 | 9 | 0 | 1 | 1965 | 639647 | 641611 | 0 | 3579 |
| YJM693 | XII | 1 | 10 | 2456655 | leftend | YLL062C | 100 | 975 | 0 | 0 | 1 | 975 | 3816 | 2842 | 0 | 1801 |
| YJM693 | XII | 2455255 | 2455264 | 2456655 | middle |  |  |  |  |  |  |  |  |  |  |  |
| YJM693 | XII | 2456655 | 2456655 | 2456655 | rightend_complete | YLR460C | 99.204 | 1131 | 9 | 0 | 1 | 1131 | 2450921 | 2449791 | 0 | 2039 |
| YJM693 | XIII | 1 | 10 | 901842 | leftend | YML131W | 99.909 | 1098 | 1 | 0 | 1 | 1098 | 2612 | 3709 | 0 | 2023 |
| YJM693 | XIII | 525 | 534 | 901842 | middle |  |  |  |  |  |  |  |  |  |  |  |
| YJM693 | XIII | 893627 | 893636 | 901842 | middle |  |  |  |  |  |  |  |  |  |  |  |
| YJM693 | XIII | 899280 | 899289 | 901842 | middle |  |  |  |  |  |  |  |  |  |  |  |
| YJM693 | XIII | 900725 | 900734 | 901842 | middle |  |  |  |  |  |  |  |  |  |  |  |
| YJM693 | XIII | 901360 | 901369 | 901842 | middle |  |  |  |  |  |  |  |  |  |  |  |
| YJM693 | XIII | 901842 | 901842 | 901842 | rightend_complete | YMR321C | 99.361 | 313 | 2 | 0 | 6 | 318 | 887694 | 887382 | 2.48E-161 | 568 |
| YJM693 | XIV | 1 | 10 | 759835 | leftend | YNL332W | 98.631 | 1023 | 14 | 0 | 1 | 1023 | 7116 | 8138 | 0 | 1812 |
| YJM693 | XIV | 758459 | 758468 | 759835 | middle |  |  |  |  |  |  |  |  |  |  |  |
| YJM693 | XIV | 759835 | 759835 | 759835 | rightend_complete | YNR071C | 100 | 1029 | 0 | 0 | 1 | 1029 | 746722 | 745694 | 0 | 1901 |
| YJM693 | XV | 1 | 10 | 1046677 | leftend | YOL154W | 99.467 | 750 | 3 | 1 | 1 | 750 | 23595 | 24343 | 0 | 1362 |
| YJM693 | XV | 1042448 | 1042457 | 1046677 | middle |  |  |  |  |  |  |  |  |  |  |  |
| YJM693 | XV | 1043047 | 1043056 | 1046677 | middle |  |  |  |  |  |  |  |  |  |  |  |
| YJM693 | XV | 1045850 | 1045859 | 1046677 | middle |  |  |  |  |  |  |  |  |  |  |  |
| YJM693 | XV | 1046668 | 1046677 | 1046677 | rightend | YOR381W | 98.736 | 2136 | 27 | 0 | 1 | 2136 | 1020636 | 1022771 | 0 | 3795 |
| YJM693 | XVI | 1 | 10 | 910596 | leftend | YPL274W | 99.773 | 1764 | 4 | 0 | 1 | 1764 | 16548 | 18311 | 0 | 3236 |
| YJM693 | XVI | 568 | 577 | 910596 | middle |  |  |  |  |  |  |  |  |  |  |  |
| YJM693 | XVI | 790 | 799 | 910596 | middle |  |  |  |  |  |  |  |  |  |  |  |
| YJM693 | XVI | 2125 | 2134 | 910596 | middle |  |  |  |  |  |  |  |  |  |  |  |
| YJM693 | XVI | 2723 | 2732 | 910596 | middle |  |  |  |  |  |  |  |  |  |  |  |
| YJM693 | XVI | 3809 | 3818 | 910596 | middle |  |  |  |  |  |  |  |  |  |  |  |
| YJM693 | XVI | 8760 | 8769 | 910596 | middle |  |  |  |  |  |  |  |  |  |  |  |
| YJM693 | XVI | 910587 | 910596 | 910596 | rightend | YPR201W | 100 | 1215 | 0 | 0 | 1 | 1215 | 907301 | 908515 | 0 | 2244 |
| YJM969 | I | 1 | 10 | 204570 | leftend | YAL063C | 85.784 | 1632 | 228 | 4 | 1 | 1630 | 12237 | 10608 | 0 | 1725 |
| YJM969 | I | 204561 | 204570 | 204570 | rightend | YAR050W | 82.699 | 1497 | 255 | 4 | 1 | 1495 | 181806 | 183300 | 0 | 1327 |
| YJM969 | II | 1 |  | 785903 | leftend_complete | YNCB0001W | 100 | 84 | 0 | 0 | 1 | 84 | 3768 | 3851 | 4.16E-38 | 156 |
| YJM969 | II | 785894 | 785903 | 785903 | rightend | YBR297W | 99.076 | 1407 | 13 | 0 | 1 | 1407 | 776213 | 777619 | 0 | 2527 |
| YJM969 | III | 1 | 10 | 308226 | leftend | YCL073C | 99.919 | 1242 | 1 | 0 | 607 | 1848 | 6406 | 5165 | 0 | 2289 |
| YJM969 | III | 308217 | 308226 | 308226 | rightend | YCR101C | 99.818 | 549 | 1 | 0 | 1 | 549 | 295401 | 294853 | 0 | 1009 |
| YJM969 | IV | 1 | 10 | 1458398 | leftend | YDL243C | 95.051 | 990 | 49 | 0 | 1 | 990 | 7920 | 6931 | 0 | 1557 |
| YJM969 | IV | 320 | 329 | 1458398 | middle |  |  |  |  |  |  |  |  |  |  |  |
| YJM969 | IV | 1458389 | 1458398 | 1458398 | rightend | YDR541C | 99.42 | 1035 | 6 | 0 | 1 | 1035 | 1455380 | 1454346 | 0 | 1879 |
| YJM969 | V | 1 | 10 | 553371 | leftend | YEL070W | 99.006 | 1509 | 15 | 0 | 1 | 1509 | 15140 | 16648 | 0 | 2704 |
| YJM969 | V | 798 | 807 | 553371 | middle |  |  |  |  |  |  |  |  |  |  |  |
| YJM969 | V | 553362 | 553371 | 553371 | rightend | YER187W | 91.981 | 424 | 32 | 2 | 4 | 426 | 549644 | 550066 | 3.44E-169 | 593 |
| YJM969 | VI | 1 | 10 | 242185 | leftend | YFL051C | 98.23 | 452 | 8 | 0 | 1 | 452 | 9319 | 8868 | 0 | 791 |
| YJM969 | VI | 242176 | 242185 | 242185 | rightend | YFR055W | 99.902 | 1023 | 1 | 0 | 1 | 1023 | 235657 | 236679 | 0 | 1884 |
| YJM969 | VII | 1 | 10 | 1037008 | leftend | YGL258W | 97.585 | 621 | 15 | 0 | 1 | 621 | 10513 | 11133 | 0 | 1064 |
| YJM969 | VII | 1036999 | 1037008 | 1037008 | rightend | YGR287C | 99.016 | 1626 | 16 | 0 | 145 | 1770 | 1030528 | 1028903 | 0 | 2915 |
| YJM969 | VIII | 1 |  | 885636 | leftend_complete | YPL274W | 100 | 1764 | 0 | 0 | 1 | 1764 | 19806 | 21569 | 0 | 3258 |
| YJM969 | VIII | 2192 | 2201 | 885636 | middle |  |  |  |  |  |  |  |  |  |  |  |
| YJM969 | VIII | 5111 | 5120 | 885636 | middle |  |  |  |  |  |  |  |  |  |  |  |
| YJM969 | VIII | 885627 | 885636 | 885636 | rightend | YHR211W | 82.535 | 939 | 153 | 7 | 1 | 932 | 874097 | 875031 | 0 | 815 |
| YJM969 | IX | 1 | 10 | 406191 | leftend | YIL169C | 85.481 | 1901 | 227 | 28 | 520 | 2382 | 1903 | 14 | 0 | 1936 |
| YJM969 | IX | 406191 | 406191 | 406191 | rightend_complete | YIR039C | 98.947 | 1614 | 17 | 0 | 1 | 1614 | 402222 | 400609 | 0 | 2887 |
| YJM969 | X | 1 | 10 | 717168 | leftend | YJL213W | 99.699 | 996 | 3 | 0 | 1 | 996 | 27019 | 28014 | 0 | 1823 |
| YJM969 | X | 697 | 706 | 717168 | middle |  |  |  |  |  |  |  |  |  |  |  |
| YJM969 | X | 717168 | 717168 | 717168 | rightend_complete | YJR152W | 99.755 | 1632 | 3 | 1 | 1 | 1632 | 697020 | 698650 | 0 | 2990 |
| YJM969 | XI | 1 | 10 | 659218 | leftend | YKL222C | 99.339 | 2119 | 13 | 1 | 1 | 2118 | 3161 | 1043 | 0 | 3834 |
| YJM969 | XI | 658623 | 658632 | 659218 | middle |  |  |  |  |  |  |  |  |  |  |  |
| YJM969 | XI | 659209 | 659218 | 659218 | rightend | YKR101W | 99.847 | 1965 | 3 | 0 | 1 | 1965 | 637378 | 639342 | 0 | 3613 |
| YJM969 | XII | 1 |  | 1746932 | leftend_complete | YLL062C | 99.897 | 975 | 1 | 0 | 1 | 975 | 6319 | 5345 | 0 | 1796 |
| YJM969 | XII | 1746923 | 1746932 | 1746932 | rightend | YLR460C | 94.071 | 1130 | 65 | 2 | 3 | 1131 | 1745219 | 1744091 | 0 | 1714 |
| YJM969 | XIII | 1 | 10 | 910459 | leftend | YML131W | 100 | 1098 | 0 | 0 | 1 | 1098 | 2891 | 3988 | 0 | 2028 |
| YJM969 | XIII | 910242 | 910251 | 910459 | middle |  |  |  |  |  |  |  |  |  |  |  |
| YJM969 | XIII | 910450 | 910459 | 910459 | rightend | YMR321C | 99.361 | 313 | 2 | 0 | 6 | 318 | 887452 | 887140 | 2.48E-161 | 568 |
| YJM969 | XIV | 1 |  | 759378 | leftend_complete | YNL332W | 98.534 | 1023 | 15 | 0 | 1 | 1023 | 10219 | 11241 | 0 | 1807 |
| YJM969 | XIV | 758596 | 758605 | 759378 | middle |  |  |  |  |  |  |  |  |  |  |  |
| YJM969 | XIV | 759091 | 759100 | 759378 | middle |  |  |  |  |  |  |  |  |  |  |  |
| YJM969 | XIV | 759378 | 759378 | 759378 | rightend_complete | YNR071C | 100 | 1029 | 0 | 0 | 1 | 1029 | 748675 | 747647 | 0 | 1901 |
| YJM969 | XV | 1 | 10 | 1047922 | leftend | YOL154W | 99.467 | 750 | 4 | 0 | 1 | 750 | 22582 | 23331 | 0 | 1363 |
| YJM969 | XV | 1046635 | 1046644 | 1047922 | middle |  |  |  |  |  |  |  |  |  |  |  |
| YJM969 | XV | 1047007 | 1047016 | 1047922 | middle |  |  |  |  |  |  |  |  |  |  |  |
| YJM969 | XV | 1047913 | 1047922 | 1047922 | rightend | YOR381W | 100 | 2136 | 0 | 0 | 1 | 2136 | 1019557 | 1021692 | 0 | 3945 |
| YJM969 | XVI | 1 | 10 | 557050 | leftend | YHL044W | 98.446 | 708 | 11 | 0 | 1 | 708 | 8151 | 8858 | 0 | 1247 |
| YJM969 | XVI | 557041 | 557050 | 557050 | rightend | YPR201W | 100 | 1215 | 0 | 0 | 1 | 1215 | 551123 | 552337 | 0 | 2244 |
| YJM972 | I | 1 | 10 | 211191 | leftend | YAL063C | 94.132 | 1261 | 40 | 14 | 2715 | 3969 | 187735 | 188967 | 0 | 1888 |
| YJM972 | I | 211191 | 211191 | 211191 | rightend_complete | YAR050W | 91.106 | 1428 | 103 | 7 | 2844 | 4250 | 187233 | 188657 | 0 | 1912 |
| YJM972 | II | 1 | 10 | 784917 | leftend | YNCB0001W | 100 | 84 | 0 | 0 | 1 | 84 | 2231 | 2314 | 4.16E-38 | 156 |
| YJM972 | II | 487 | 496 | 784917 | middle |  |  |  |  |  |  |  |  |  |  |  |
| YJM972 | II | 784908 | 784917 | 784917 | rightend | YBR297W | 99.076 | 1407 | 13 | 0 | 1 | 1407 | 774722 | 776128 | 0 | 2527 |
| YJM972 | III | 1 |  | 309517 | leftend_complete | YCL073C | 99.729 | 1848 | 5 | 0 | 1 | 1848 | 8250 | 6403 | 0 | 3386 |
| YJM972 | III | 319 | 328 | 309517 | middle |  |  |  |  |  |  |  |  |  |  |  |
| YJM972 | III | 309508 | 309517 | 309517 | rightend | YCR101C | 99.818 | 549 | 1 | 0 | 1 | 549 | 296613 | 296065 | 0 | 1009 |
| YJM972 | IV | 1 | 10 | 1479996 | leftend | YDL243C | 95.051 | 990 | 49 | 0 | 1 | 990 | 18250 | 17261 | 0 | 1557 |
| YJM972 | IV | 1479987 | 1479996 | 1479996 | rightend | YDR541C | 99.42 | 1035 | 6 | 0 | 1 | 1035 | 1477249 | 1476215 | 0 | 1879 |
| YJM972 | V | 1 | 10 | 551568 | leftend | YEL070W | 99.006 | 1509 | 15 | 0 | 1 | 1509 | 12832 | 14340 | 0 | 2704 |
| YJM972 | V | 551559 | 551568 | 551568 | rightend | YER187W | 91.981 | 424 | 32 | 2 | 4 | 426 | 547638 | 548060 | 3.44E-169 | 593 |
| YJM972 | VI | 1 | 10 | 234509 | leftend | YFL051C | 98.451 | 452 | 7 | 0 | 1 | 452 | 1969 | 1518 | 0 | 797 |
| YJM972 | VI | 768 | 777 | 234509 | middle |  |  |  |  |  |  |  |  |  |  |  |
| YJM972 | VI | 234500 | 234509 | 234509 | rightend | YFR055W | 99.902 | 1023 | 1 | 0 | 1 | 1023 | 228315 | 229337 | 0 | 1884 |
| YJM972 | VII | 1 | 10 | 1037045 | leftend | YGL258W | 97.585 | 621 | 15 | 0 | 1 | 621 | 4778 | 5398 | 0 | 1064 |
| YJM972 | VII | 1037036 | 1037045 | 1037045 | rightend | YGR287C | 99.262 | 1626 | 12 | 0 | 145 | 1770 | 1024755 | 1023130 | 0 | 2937 |
| YJM972 | VIII | 1 | 10 | 893833 | leftend | YPL274W | 99.887 | 1764 | 2 | 0 | 1 | 1764 | 15825 | 17588 | 0 | 3247 |
| YJM972 | VIII | 1095 | 1104 | 893833 | middle |  |  |  |  |  |  |  |  |  |  |  |
| YJM972 | VIII | 893833 | 893833 | 893833 | rightend_complete | YHR211W | 82.535 | 939 | 153 | 7 | 1 | 932 | 881708 | 882642 | 0 | 815 |
| YJM972 | IX | 1 | 10 | 410479 | leftend | YIL169C | 93.928 | 1548 | 88 | 3 | 265 | 1812 | 7569 | 6028 | 0 | 2333 |
| YJM972 | IX | 409999 | 410008 | 410479 | middle |  |  |  |  |  |  |  |  |  |  |  |
| YJM972 | IX | 410479 | 410479 | 410479 | rightend_complete | YIR039C | 98.947 | 1614 | 17 | 0 | 1 | 1614 | 408148 | 406535 | 0 | 2887 |
| YJM972 | X | 1 | 10 | 717651 | leftend | YJL213W | 99.699 | 996 | 3 | 0 | 1 | 996 | 27701 | 28696 | 0 | 1823 |
| YJM972 | X | 1162 | 1171 | 717651 | middle |  |  |  |  |  |  |  |  |  |  |  |
| YJM972 | X | 1379 | 1388 | 717651 | middle |  |  |  |  |  |  |  |  |  |  |  |
| YJM972 | X | 717651 | 717651 | 717651 | rightend_complete | YJR152W | 99.755 | 1632 | 3 | 1 | 1 | 1632 | 697326 | 698956 | 0 | 2990 |
| YJM972 | XI | 1 | 10 | 658567 | leftend | YKL222C | 99.339 | 2119 | 13 | 1 | 1 | 2118 | 3201 | 1083 | 0 | 3834 |
| YJM972 | XI | 658558 | 658567 | 658567 | rightend | YKR101W | 99.796 | 1965 | 4 | 0 | 1 | 1965 | 637400 | 639364 | 0 | 3607 |
| YJM972 | XII | 1 | 10 | 2341640 | leftend | YLL062C | 99.897 | 975 | 1 | 0 | 1 | 975 | 4016 | 3042 | 0 | 1796 |
| YJM972 | XII | 2341631 | 2341640 | 2341640 | rightend | YLR460C | 93.982 | 1130 | 66 | 2 | 3 | 1131 | 2339879 | 2338751 | 0 | 1709 |
| YJM972 | XIII | 1 | 10 | 910034 | leftend | YML131W | 100 | 1098 | 0 | 0 | 1 | 1098 | 2329 | 3426 | 0 | 2028 |
| YJM972 | XIII | 910025 | 910034 | 910034 | rightend | YMR321C | 99.361 | 313 | 2 | 0 | 6 | 318 | 887263 | 886951 | 2.48E-161 | 568 |
| YJM972 | XIV | 1 | 10 | 756737 | leftend | YNL332W | 98.631 | 1023 | 14 | 0 | 1 | 1023 | 6015 | 7037 | 0 | 1812 |
| YJM972 | XIV | 775 | 784 | 756737 | middle |  |  |  |  |  |  |  |  |  |  |  |
| YJM972 | XIV | 756371 | 756380 | 756737 | middle |  |  |  |  |  |  |  |  |  |  |  |
| YJM972 | XIV | 756737 | 756737 | 756737 | rightend_complete | YNR071C | 100 | 1029 | 0 | 0 | 1 | 1029 | 744497 | 743469 | 0 | 1901 |
| YJM972 | XV | 1 | 10 | 1037074 | leftend | YOL154W | 99.467 | 750 | 4 | 0 | 1 | 750 | 23001 | 23750 | 0 | 1363 |
| YJM972 | XV | 1036187 | 1036196 | 1037074 | middle |  |  |  |  |  |  |  |  |  |  |  |
| YJM972 | XV | 1037065 | 1037074 | 1037074 | rightend | YOR381W | 100 | 2136 | 0 | 0 | 1 | 2136 | 1020095 | 1022230 | 0 | 3945 |
| YJM972 | XVI | 1 | 10 | 558198 | leftend | YHL044W | 98.446 | 708 | 11 | 0 | 1 | 708 | 8687 | 9394 | 0 | 1247 |
| YJM972 | XVI | 555689 | 555698 | 558198 | middle |  |  |  |  |  |  |  |  |  |  |  |
| YJM972 | XVI | 557539 | 557548 | 558198 | middle |  |  |  |  |  |  |  |  |  |  |  |
| YJM972 | XVI | 558189 | 558198 | 558198 | rightend | YPR201W | 100 | 1215 | 0 | 0 | 1 | 1215 | 552032 | 553246 | 0 | 2244 |
| YJM975 | I | 1 | 10 | 195504 | leftend | YAL063C | 85.571 | 1497 | 212 | 4 | 1 | 1495 | 12166 | 10672 | 0 | 1565 |
| YJM975 | I | 195495 | 195504 | 195504 | rightend | YAR050W | 82.164 | 1497 | 263 | 4 | 1 | 1495 | 181729 | 183223 | 0 | 1282 |
| YJM975 | II | 1 | 10 | 785640 | leftend | YNCB0001W | 100 | 84 | 0 | 0 | 1 | 84 | 1647 | 1730 | 4.16E-38 | 156 |
| YJM975 | II | 784051 | 784060 | 785640 | middle |  |  |  |  |  |  |  |  |  |  |  |
| YJM975 | II | 785631 | 785640 | 785640 | rightend | YBR297W | 99.076 | 1407 | 13 | 0 | 1 | 1407 | 774065 | 775471 | 0 | 2527 |
| YJM975 | III | 1 |  | 309531 | leftend_complete | YCL073C | 99.678 | 1242 | 4 | 0 | 607 | 1848 | 7374 | 6133 | 0 | 2272 |
| YJM975 | III | 309522 | 309531 | 309531 | rightend | YCR101C | 99.818 | 549 | 1 | 0 | 1 | 549 | 296705 | 296157 | 0 | 1009 |
| YJM975 | IV | 1 | 10 | 1469105 | leftend | YDL243C | 95.051 | 990 | 49 | 0 | 1 | 990 | 19312 | 18323 | 0 | 1557 |
| YJM975 | IV | 1687 | 1696 | 1469105 | middle |  |  |  |  |  |  |  |  |  |  |  |
| YJM975 | IV | 1469096 | 1469105 | 1469105 | rightend | YDR541C | 99.42 | 1035 | 6 | 0 | 1 | 1035 | 1466472 | 1465438 | 0 | 1879 |
| YJM975 | V | 1 | 10 | 550353 | leftend | YEL070W | 99.006 | 1509 | 15 | 0 | 1 | 1509 | 12840 | 14348 | 0 | 2704 |
| YJM975 | V | 550344 | 550353 | 550353 | rightend | YER187W | 91.981 | 424 | 32 | 2 | 4 | 426 | 547368 | 547790 | 3.44E-169 | 593 |
| YJM975 | VI | 1 | 10 | 233509 | leftend | YFL051C | 98.451 | 452 | 7 | 0 | 1 | 452 | 1092 | 641 | 0 | 797 |
| YJM975 | VI | 233509 | 233509 | 233509 | rightend_complete | YFR055W | 99.902 | 1023 | 1 | 0 | 1 | 1023 | 227440 | 228462 | 0 | 1884 |
| YJM975 | VII | 1 | 10 | 1041704 | leftend | YGL258W | 97.585 | 621 | 15 | 0 | 1 | 621 | 10623 | 11243 | 0 | 1064 |
| YJM975 | VII | 1041695 | 1041704 | 1041704 | rightend | YGR287C | 99.016 | 1626 | 16 | 0 | 145 | 1770 | 1030476 | 1028851 | 0 | 2915 |
| YJM975 | VIII | 1 | 10 | 533482 | leftend | YHL044W | 98.446 | 708 | 11 | 0 | 1 | 708 | 1304 | 2011 | 0 | 1247 |
| YJM975 | VIII | 533473 | 533482 | 533482 | rightend | YHR211W | 82.535 | 939 | 153 | 7 | 1 | 932 | 522030 | 522964 | 0 | 815 |
| YJM975 | IX | 1 | 10 | 409597 | leftend | YIL169C | 84.176 | 2155 | 271 | 38 | 520 | 2615 | 7216 | 5073 | 0 | 2026 |
| YJM975 | IX | 409301 | 409310 | 409597 | middle |  |  |  |  |  |  |  |  |  |  |  |
| YJM975 | IX | 409588 | 409597 | 409597 | rightend | YIR039C | 99.009 | 1614 | 16 | 0 | 1 | 1614 | 407646 | 406033 | 0 | 2892 |
| YJM975 | X | 1 |  | 720007 | leftend_complete | YJL213W | 99.699 | 996 | 3 | 0 | 1 | 996 | 29917 | 30912 | 0 | 1823 |
| YJM975 | X | 2250 | 2259 | 720007 | middle |  |  |  |  |  |  |  |  |  |  |  |
| YJM975 | X | 3611 | 3620 | 720007 | middle |  |  |  |  |  |  |  |  |  |  |  |
| YJM975 | X | 720007 | 720007 | 720007 | rightend_complete | YJR152W | 99.755 | 1632 | 3 | 1 | 1 | 1632 | 699924 | 701554 | 0 | 2990 |
| YJM975 | XI | 1 | 10 | 661607 | leftend | YKL222C | 99.481 | 2118 | 11 | 0 | 1 | 2118 | 3293 | 1176 | 0 | 3851 |
| YJM975 | XI | 661177 | 661186 | 661607 | middle |  |  |  |  |  |  |  |  |  |  |  |
| YJM975 | XI | 661607 | 661607 | 661607 | rightend_complete | YKR101W | 99.796 | 1965 | 4 | 0 | 1 | 1965 | 637177 | 639141 | 0 | 3607 |
| YJM975 | XII | 1 | 10 | 1618966 | leftend | YLL062C | 99.897 | 975 | 1 | 0 | 1 | 975 | 5887 | 4913 | 0 | 1796 |
| YJM975 | XII | 2073 | 2082 | 1618966 | middle |  |  |  |  |  |  |  |  |  |  |  |
| YJM975 | XII | 1618957 | 1618966 | 1618966 | rightend | YLR460C | 94.071 | 1130 | 65 | 2 | 3 | 1131 | 1617205 | 1616077 | 0 | 1714 |
| YJM975 | XIII | 1 | 10 | 914102 | leftend | YML131W | 100 | 1098 | 0 | 0 | 1 | 1098 | 6290 | 7387 | 0 | 2028 |
| YJM975 | XIII | 2019 | 2028 | 914102 | middle |  |  |  |  |  |  |  |  |  |  |  |
| YJM975 | XIII | 2713 | 2722 | 914102 | middle |  |  |  |  |  |  |  |  |  |  |  |
| YJM975 | XIII | 914093 | 914102 | 914102 | rightend | YMR321C | 99.361 | 313 | 2 | 0 | 6 | 318 | 891097 | 890785 | 2.48E-161 | 568 |
| YJM975 | XIV | 1 | 10 | 755304 | leftend | YNL332W | 98.631 | 1023 | 14 | 0 | 1 | 1023 | 5239 | 6261 | 0 | 1812 |
| YJM975 | XIV | 753842 | 753851 | 755304 | middle |  |  |  |  |  |  |  |  |  |  |  |
| YJM975 | XIV | 754131 | 754140 | 755304 | middle |  |  |  |  |  |  |  |  |  |  |  |
| YJM975 | XIV | 755295 | 755304 | 755304 | rightend | YNR071C | 100 | 1029 | 0 | 0 | 1 | 1029 | 743357 | 742329 | 0 | 1901 |
| YJM975 | XV | 1 | 10 | 1049782 | leftend | YOL154W | 99.467 | 750 | 4 | 0 | 1 | 750 | 23864 | 24613 | 0 | 1363 |
| YJM975 | XV | 962 | 971 | 1049782 | middle |  |  |  |  |  |  |  |  |  |  |  |
| YJM975 | XV | 1048240 | 1048249 | 1049782 | middle |  |  |  |  |  |  |  |  |  |  |  |
| YJM975 | XV | 1048774 | 1048783 | 1049782 | middle |  |  |  |  |  |  |  |  |  |  |  |
| YJM975 | XV | 1049773 | 1049782 | 1049782 | rightend | YOR381W | 100 | 2136 | 0 | 0 | 1 | 2136 | 1020969 | 1023104 | 0 | 3945 |
| YJM975 | XVI | 1 | 10 | 911644 | leftend | YPL274W | 100 | 1764 | 0 | 0 | 1 | 1764 | 17783 | 19546 | 0 | 3258 |
| YJM975 | XVI | 647 | 656 | 911644 | middle |  |  |  |  |  |  |  |  |  |  |  |
| YJM975 | XVI | 1587 | 1596 | 911644 | middle |  |  |  |  |  |  |  |  |  |  |  |
| YJM975 | XVI | 2905 | 2914 | 911644 | middle |  |  |  |  |  |  |  |  |  |  |  |
| YJM975 | XVI | 909438 | 909447 | 911644 | middle |  |  |  |  |  |  |  |  |  |  |  |
| YJM975 | XVI | 911635 | 911644 | 911644 | rightend | YPR201W | 100 | 1215 | 0 | 0 | 1 | 1215 | 905765 | 906979 | 0 | 2244 |
| YJM978 | I | 1 | 10 | 196060 | leftend | YAL063C | 94.132 | 1261 | 40 | 14 | 2715 | 3969 | 173095 | 174327 | 0 | 1888 |
| YJM978 | I | 196051 | 196060 | 196060 | rightend | YAR050W | 91.106 | 1428 | 103 | 7 | 2844 | 4250 | 172593 | 174017 | 0 | 1912 |
| YJM978 | II | 1 |  | 786576 | leftend_complete | YNCB0001W | 100 | 84 | 0 | 0 | 1 | 84 | 3778 | 3861 | 4.16E-38 | 156 |
| YJM978 | II | 786567 | 786576 | 786576 | rightend | YBR297W | 99.076 | 1407 | 13 | 0 | 1 | 1407 | 776248 | 777654 | 0 | 2527 |
| YJM978 | III | 1 |  | 310218 | leftend_complete | YCL073C | 99.675 | 1848 | 6 | 0 | 1 | 1848 | 7687 | 5840 | 0 | 3380 |
| YJM978 | III | 325 | 334 | 310218 | middle |  |  |  |  |  |  |  |  |  |  |  |
| YJM978 | III | 566 | 575 | 310218 | middle |  |  |  |  |  |  |  |  |  |  |  |
| YJM978 | III | 309329 | 309338 | 310218 | middle |  |  |  |  |  |  |  |  |  |  |  |
| YJM978 | III | 309549 | 309558 | 310218 | middle |  |  |  |  |  |  |  |  |  |  |  |
| YJM978 | III | 310218 | 310218 | 310218 | rightend_complete | YCR101C | 99.818 | 549 | 1 | 0 | 1 | 549 | 295735 | 295187 | 0 | 1009 |
| YJM978 | IV | 1 | 10 | 1453104 | leftend | YDL243C | 95.051 | 990 | 49 | 0 | 1 | 990 | 1845 | 856 | 0 | 1557 |
| YJM978 | IV | 1453095 | 1453104 | 1453104 | rightend | YDR541C | 99.42 | 1035 | 6 | 0 | 1 | 1035 | 1450507 | 1449473 | 0 | 1879 |
| YJM978 | V | 1 | 10 | 554610 | leftend | YEL070W | 99.006 | 1509 | 15 | 0 | 1 | 1509 | 16182 | 17690 | 0 | 2704 |
| YJM978 | V | 671 | 680 | 554610 | middle |  |  |  |  |  |  |  |  |  |  |  |
| YJM978 | V | 554601 | 554610 | 554610 | rightend | YER187W | 91.981 | 424 | 32 | 2 | 4 | 426 | 550645 | 551067 | 3.44E-169 | 593 |
| YJM978 | VI | 1 | 10 | 233198 | leftend | YFL051C | 98.451 | 452 | 7 | 0 | 1 | 452 | 863 | 412 | 0 | 797 |
| YJM978 | VI | 233198 | 233198 | 233198 | rightend_complete | YFR055W | 99.902 | 1023 | 1 | 0 | 1 | 1023 | 227201 | 228223 | 0 | 1884 |
| YJM978 | VII | 1 |  | 1038299 | leftend_complete | YGL258W | 97.585 | 621 | 15 | 0 | 1 | 621 | 11336 | 11956 | 0 | 1064 |
| YJM978 | VII | 394 | 403 | 1038299 | middle |  |  |  |  |  |  |  |  |  |  |  |
| YJM978 | VII | 1038290 | 1038299 | 1038299 | rightend | YGR287C | 99.262 | 1626 | 12 | 0 | 145 | 1770 | 1030950 | 1029325 | 0 | 2937 |
| YJM978 | VIII | 1 |  | 882461 | leftend_complete | YPL274W | 99.943 | 1764 | 1 | 0 | 1 | 1764 | 23188 | 24951 | 0 | 3253 |
| YJM978 | VIII | 882452 | 882461 | 882461 | rightend | YHR211W | 82.535 | 939 | 153 | 7 | 1 | 932 | 873902 | 874836 | 0 | 815 |
| YJM978 | IX | 1 | 10 | 407974 | leftend | YIL169C | 93.928 | 1548 | 88 | 3 | 265 | 1812 | 6108 | 4567 | 0 | 2333 |
| YJM978 | IX | 407965 | 407974 | 407974 | rightend | YIR039C | 98.947 | 1614 | 17 | 0 | 1 | 1614 | 406310 | 404697 | 0 | 2887 |
| YJM978 | X | 1 | 10 | 717211 | leftend | YJL213W | 99.699 | 996 | 3 | 0 | 1 | 996 | 27005 | 28000 | 0 | 1823 |
| YJM978 | X | 697 | 706 | 717211 | middle |  |  |  |  |  |  |  |  |  |  |  |
| YJM978 | X | 717211 | 717211 | 717211 | rightend_complete | YJR152W | 99.755 | 1632 | 3 | 1 | 1 | 1632 | 696832 | 698462 | 0 | 2990 |
| YJM978 | XI | 1 | 10 | 658200 | leftend | YKL222C | 99.339 | 2119 | 13 | 1 | 1 | 2118 | 3201 | 1083 | 0 | 3834 |
| YJM978 | XI | 658191 | 658200 | 658200 | rightend | YKR101W | 99.796 | 1965 | 4 | 0 | 1 | 1965 | 637060 | 639024 | 0 | 3607 |
| YJM978 | XII | 1 | 10 | 1911963 | leftend | YLL062C | 99.897 | 975 | 1 | 0 | 1 | 975 | 5165 | 4191 | 0 | 1796 |
| YJM978 | XII | 1150 | 1159 | 1911963 | middle |  |  |  |  |  |  |  |  |  |  |  |
| YJM978 | XII | 1911475 | 1911484 | 1911963 | middle |  |  |  |  |  |  |  |  |  |  |  |
| YJM978 | XII | 1911954 | 1911963 | 1911963 | rightend | YLR460C | 93.982 | 1130 | 66 | 2 | 3 | 1131 | 1909723 | 1908595 | 0 | 1709 |
| YJM978 | XIII | 1 | 10 | 911077 | leftend | YML131W | 100 | 1098 | 0 | 0 | 1 | 1098 | 2984 | 4081 | 0 | 2028 |
| YJM978 | XIII | 367 | 376 | 911077 | middle |  |  |  |  |  |  |  |  |  |  |  |
| YJM978 | XIII | 911068 | 911077 | 911077 | rightend | YMR321C | 99.361 | 313 | 2 | 0 | 6 | 318 | 888167 | 887855 | 2.48E-161 | 568 |
| YJM978 | XIV | 1 | 10 | 755784 | leftend | YNL332W | 98.631 | 1023 | 14 | 0 | 1 | 1023 | 5399 | 6421 | 0 | 1812 |
| YJM978 | XIV | 755775 | 755784 | 755784 | rightend | YNR071C | 100 | 1029 | 0 | 0 | 1 | 1029 | 744056 | 743028 | 0 | 1901 |
| YJM978 | XV | 1 | 10 | 1047584 | leftend | YOL154W | 99.467 | 750 | 4 | 0 | 1 | 750 | 22924 | 23673 | 0 | 1363 |
| YJM978 | XV | 1045722 | 1045731 | 1047584 | middle |  |  |  |  |  |  |  |  |  |  |  |
| YJM978 | XV | 1046296 | 1046305 | 1047584 | middle |  |  |  |  |  |  |  |  |  |  |  |
| YJM978 | XV | 1047584 | 1047584 | 1047584 | rightend_complete | YOR381W | 100 | 2136 | 0 | 0 | 1 | 2136 | 1020131 | 1022266 | 0 | 3945 |
| YJM978 | XVI | 1 |  | 556953 | leftend_complete | YHL044W | 98.446 | 708 | 11 | 0 | 1 | 708 | 8852 | 9559 | 0 | 1247 |
| YJM978 | XVI | 555094 | 555103 | 556953 | middle |  |  |  |  |  |  |  |  |  |  |  |
| YJM978 | XVI | 556944 | 556953 | 556953 | rightend | YPR201W | 100 | 1215 | 0 | 0 | 1 | 1215 | 551713 | 552927 | 0 | 2244 |
| YJM981 | I | 1 | 10 | 196939 | leftend | YAL063C | 94.132 | 1261 | 40 | 14 | 2715 | 3969 | 173477 | 174709 | 0 | 1888 |
| YJM981 | I | 196939 | 196939 | 196939 | rightend_complete | YAR050W | 91.106 | 1428 | 103 | 7 | 2844 | 4250 | 172975 | 174399 | 0 | 1912 |
| YJM981 | II | 1 | 10 | 785031 | leftend | YNCB0001W | 100 | 84 | 0 | 0 | 1 | 84 | 2675 | 2758 | 4.16E-38 | 156 |
| YJM981 | II | 785022 | 785031 | 785031 | rightend | YBR297W | 99.076 | 1407 | 13 | 0 | 1 | 1407 | 774839 | 776245 | 0 | 2527 |
| YJM981 | III | 1 | 10 | 309306 | leftend | YCL073C | 99.729 | 1848 | 5 | 0 | 1 | 1848 | 8340 | 6493 | 0 | 3386 |
| YJM981 | III | 596 | 605 | 309306 | middle |  |  |  |  |  |  |  |  |  |  |  |
| YJM981 | III | 309297 | 309306 | 309306 | rightend | YCR101C | 99.818 | 549 | 1 | 0 | 1 | 549 | 296402 | 295854 | 0 | 1009 |
| YJM981 | IV | 1 | 10 | 1482018 | leftend | YDL243C | 95.051 | 990 | 49 | 0 | 1 | 990 | 16910 | 15921 | 0 | 1557 |
| YJM981 | IV | 1482009 | 1482018 | 1482018 | rightend | YDR541C | 99.42 | 1035 | 6 | 0 | 1 | 1035 | 1479421 | 1478387 | 0 | 1879 |
| YJM981 | V | 1 | 10 | 550159 | leftend | YEL070W | 99.006 | 1509 | 15 | 0 | 1 | 1509 | 12828 | 14336 | 0 | 2704 |
| YJM981 | V | 550150 | 550159 | 550159 | rightend | YER187W | 91.981 | 424 | 32 | 2 | 4 | 426 | 547651 | 548073 | 3.44E-169 | 593 |
| YJM981 | VI | 1 | 10 | 234048 | leftend | YFL051C | 98.23 | 452 | 8 | 0 | 1 | 452 | 1630 | 1179 | 0 | 791 |
| YJM981 | VI | 768 | 777 | 234048 | middle |  |  |  |  |  |  |  |  |  |  |  |
| YJM981 | VI | 234039 | 234048 | 234048 | rightend | YFR055W | 99.902 | 1023 | 1 | 0 | 1 | 1023 | 227945 | 228967 | 0 | 1884 |
| YJM981 | VII | 1 | 10 | 1036860 | leftend | YGL258W | 97.585 | 621 | 15 | 0 | 1 | 621 | 4820 | 5440 | 0 | 1064 |
| YJM981 | VII | 1036851 | 1036860 | 1036860 | rightend | YGR287C | 99.262 | 1626 | 12 | 0 | 145 | 1770 | 1024573 | 1022948 | 0 | 2937 |
| YJM981 | VIII | 1 | 10 | 884210 | leftend | YPL274W | 99.943 | 1764 | 1 | 0 | 1 | 1764 | 18092 | 19855 | 0 | 3253 |
| YJM981 | VIII | 3277 | 3286 | 884210 | middle |  |  |  |  |  |  |  |  |  |  |  |
| YJM981 | VIII | 884210 | 884210 | 884210 | rightend_complete | YHR211W | 82.535 | 939 | 153 | 7 | 1 | 932 | 872092 | 873026 | 0 | 815 |
| YJM981 | IX | 1 | 10 | 403996 | leftend | YIL169C | 85.481 | 1901 | 227 | 26 | 520 | 2382 | 1905 | 16 | 0 | 1936 |
| YJM981 | IX | 403987 | 403996 | 403996 | rightend | YIR039C | 99.009 | 1614 | 16 | 0 | 1 | 1614 | 402136 | 400523 | 0 | 2892 |
| YJM981 | X | 1 | 10 | 715685 | leftend | YJL213W | 99.699 | 996 | 3 | 0 | 1 | 996 | 25793 | 26788 | 0 | 1823 |
| YJM981 | X | 715685 | 715685 | 715685 | rightend_complete | YJR152W | 99.755 | 1632 | 3 | 1 | 1 | 1632 | 695583 | 697213 | 0 | 2990 |
| YJM981 | XI | 1 | 10 | 658890 | leftend | YKL222C | 99.339 | 2119 | 13 | 1 | 1 | 2118 | 3293 | 1175 | 0 | 3834 |
| YJM981 | XI | 658881 | 658890 | 658890 | rightend | YKR101W | 99.796 | 1965 | 4 | 0 | 1 | 1965 | 637536 | 639500 | 0 | 3607 |
| YJM981 | XII | 1 | 10 | 2291245 | leftend | YLL062C | 100 | 975 | 0 | 0 | 1 | 975 | 5224 | 4250 | 0 | 1801 |
| YJM981 | XII | 1209 | 1218 | 2291245 | middle |  |  |  |  |  |  |  |  |  |  |  |
| YJM981 | XII | 2291236 | 2291245 | 2291245 | rightend | YLR460C | 93.982 | 1130 | 66 | 2 | 3 | 1131 | 2289086 | 2287958 | 0 | 1709 |
| YJM981 | XIII | 1 | 10 | 911845 | leftend | YML131W | 100 | 1098 | 0 | 0 | 1 | 1098 | 4027 | 5124 | 0 | 2028 |
| YJM981 | XIII | 396 | 405 | 911845 | middle |  |  |  |  |  |  |  |  |  |  |  |
| YJM981 | XIII | 1835 | 1844 | 911845 | middle |  |  |  |  |  |  |  |  |  |  |  |
| YJM981 | XIII | 911836 | 911845 | 911845 | rightend | YMR321C | 99.361 | 313 | 2 | 0 | 6 | 318 | 888836 | 888524 | 2.48E-161 | 568 |
| YJM981 | XIV | 1 | 10 | 755237 | leftend | YNL332W | 98.631 | 1023 | 14 | 0 | 1 | 1023 | 6012 | 7034 | 0 | 1812 |
| YJM981 | XIV | 775 | 784 | 755237 | middle |  |  |  |  |  |  |  |  |  |  |  |
| YJM981 | XIV | 754574 | 754583 | 755237 | middle |  |  |  |  |  |  |  |  |  |  |  |
| YJM981 | XIV | 755237 | 755237 | 755237 | rightend_complete | YNR071C | 100 | 1029 | 0 | 0 | 1 | 1029 | 744105 | 743077 | 0 | 1901 |
| YJM981 | XV | 1 | 10 | 1052679 | leftend | YOL154W | 99.467 | 750 | 4 | 0 | 1 | 750 | 22743 | 23492 | 0 | 1363 |
| YJM981 | XV | 1051991 | 1052000 | 1052679 | middle |  |  |  |  |  |  |  |  |  |  |  |
| YJM981 | XV | 1052670 | 1052679 | 1052679 | rightend | YOR381W | 100 | 2136 | 0 | 0 | 1 | 2136 | 1019674 | 1021809 | 0 | 3945 |
| YJM981 | XVI | 1 | 10 | 556055 | leftend | YHL044W | 98.446 | 708 | 11 | 0 | 1 | 708 | 9759 | 10466 | 0 | 1247 |
| YJM981 | XVI | 1053 | 1062 | 556055 | middle |  |  |  |  |  |  |  |  |  |  |  |
| YJM981 | XVI | 556046 | 556055 | 556055 | rightend | YPR201W | 100 | 1215 | 0 | 0 | 1 | 1215 | 553142 | 554356 | 0 | 2244 |
| YJM984 | I | 1 | 10 | 202872 | leftend | YAL063C | 94.132 | 1261 | 40 | 14 | 2715 | 3969 | 188066 | 189298 | 0 | 1888 |
| YJM984 | I | 464 | 473 | 202872 | middle |  |  |  |  |  |  |  |  |  |  |  |
| YJM984 | I | 202863 | 202872 | 202872 | rightend | YAR050W | 91.106 | 1428 | 103 | 7 | 2844 | 4250 | 187564 | 188988 | 0 | 1912 |
| YJM984 | II | 1 | 10 | 785604 | leftend | YNCB0001W | 100 | 84 | 0 | 0 | 1 | 84 | 3839 | 3922 | 4.16E-38 | 156 |
| YJM984 | II | 562 | 571 | 785604 | middle |  |  |  |  |  |  |  |  |  |  |  |
| YJM984 | II | 770 | 779 | 785604 | middle |  |  |  |  |  |  |  |  |  |  |  |
| YJM984 | II | 785595 | 785604 | 785604 | rightend | YBR297W | 99.076 | 1407 | 13 | 0 | 1 | 1407 | 776306 | 777712 | 0 | 2527 |
| YJM984 | III | 1 | 10 | 309333 | leftend | YCL073C | 99.567 | 1848 | 8 | 0 | 1 | 1848 | 7745 | 5898 | 0 | 3369 |
| YJM984 | III | 309333 | 309333 | 309333 | rightend_complete | YCR101C | 99.818 | 549 | 1 | 0 | 1 | 549 | 295815 | 295267 | 0 | 1009 |
| YJM984 | IV | 1 | 10 | 1471292 | leftend | YDL243C | 95.051 | 990 | 49 | 0 | 1 | 990 | 16910 | 15921 | 0 | 1557 |
| YJM984 | IV | 1471283 | 1471292 | 1471292 | rightend | YDR541C | 99.42 | 1035 | 6 | 0 | 1 | 1035 | 1468659 | 1467625 | 0 | 1879 |
| YJM984 | V | 1 | 10 | 542791 | leftend | YEL070W | 98.873 | 1509 | 17 | 0 | 1 | 1509 | 5605 | 7113 | 0 | 2693 |
| YJM984 | V | 542782 | 542791 | 542791 | rightend | YER187W | 91.981 | 424 | 32 | 2 | 4 | 426 | 540383 | 540805 | 3.44E-169 | 593 |
| YJM984 | VI | 1 | 10 | 233520 | leftend | YFL051C | 98.451 | 452 | 7 | 0 | 1 | 452 | 863 | 412 | 0 | 797 |
| YJM984 | VI | 233511 | 233520 | 233520 | rightend | YFR055W | 99.902 | 1023 | 1 | 0 | 1 | 1023 | 227191 | 228213 | 0 | 1884 |
| YJM984 | VII | 1 | 10 | 1033768 | leftend | YGL258W | 97.585 | 621 | 15 | 0 | 1 | 621 | 11007 | 11627 | 0 | 1064 |
| YJM984 | VII | 483 | 492 | 1033768 | middle |  |  |  |  |  |  |  |  |  |  |  |
| YJM984 | VII | 1033759 | 1033768 | 1033768 | rightend | YGR287C | 99.016 | 1626 | 16 | 0 | 145 | 1770 | 1031144 | 1029519 | 0 | 2915 |
| YJM984 | VIII | 1 | 10 | 521979 | leftend | YHL044W | 98.446 | 708 | 11 | 0 | 1 | 708 | 1418 | 2125 | 0 | 1247 |
| YJM984 | VIII | 521970 | 521979 | 521979 | rightend | YHR211W | 82.428 | 939 | 154 | 7 | 1 | 932 | 510611 | 511545 | 0 | 809 |
| YJM984 | IX | 1 | 10 | 410441 | leftend | YIL169C | 93.928 | 1548 | 88 | 3 | 265 | 1812 | 8917 | 7376 | 0 | 2333 |
| YJM984 | IX | 410432 | 410441 | 410441 | rightend | YIR039C | 99.009 | 1614 | 16 | 0 | 1 | 1614 | 408777 | 407164 | 0 | 2892 |
| YJM984 | X | 1 | 10 | 718533 | leftend | YJL213W | 99.699 | 996 | 3 | 0 | 1 | 996 | 27073 | 28068 | 0 | 1823 |
| YJM984 | X | 521 | 530 | 718533 | middle |  |  |  |  |  |  |  |  |  |  |  |
| YJM984 | X | 1288 | 1297 | 718533 | middle |  |  |  |  |  |  |  |  |  |  |  |
| YJM984 | X | 717008 | 717017 | 718533 | middle |  |  |  |  |  |  |  |  |  |  |  |
| YJM984 | X | 718533 | 718533 | 718533 | rightend_complete | YJR152W | 99.755 | 1632 | 3 | 1 | 1 | 1632 | 696749 | 698379 | 0 | 2990 |
| YJM984 | XI | 1 | 10 | 662218 | leftend | YKL222C | 99.339 | 2119 | 13 | 1 | 1 | 2118 | 3293 | 1175 | 0 | 3834 |
| YJM984 | XI | 661805 | 661814 | 662218 | middle |  |  |  |  |  |  |  |  |  |  |  |
| YJM984 | XI | 662218 | 662218 | 662218 | rightend_complete | YKR101W | 99.847 | 1965 | 3 | 0 | 1 | 1965 | 637208 | 639172 | 0 | 3613 |
| YJM984 | XII | 1 | 10 | 2453294 | leftend | YLL062C | 100 | 975 | 0 | 0 | 1 | 975 | 4633 | 3659 | 0 | 1801 |
| YJM984 | XII | 397 | 406 | 2453294 | middle |  |  |  |  |  |  |  |  |  |  |  |
| YJM984 | XII | 819 | 828 | 2453294 | middle |  |  |  |  |  |  |  |  |  |  |  |
| YJM984 | XII | 2453285 | 2453294 | 2453294 | rightend | YLR460C | 93.982 | 1130 | 66 | 2 | 3 | 1131 | 2451533 | 2450405 | 0 | 1709 |
| YJM984 | XIII | 1 | 10 | 910749 | leftend | YML131W | 100 | 1098 | 0 | 0 | 1 | 1098 | 3760 | 4857 | 0 | 2028 |
| YJM984 | XIII | 648 | 657 | 910749 | middle |  |  |  |  |  |  |  |  |  |  |  |
| YJM984 | XIII | 910740 | 910749 | 910749 | rightend | YMR321C | 99.361 | 313 | 2 | 0 | 6 | 318 | 888258 | 887946 | 2.48E-161 | 568 |
| YJM984 | XIV | 1 | 10 | 755178 | leftend | YNL332W | 98.631 | 1023 | 14 | 0 | 1 | 1023 | 6230 | 7252 | 0 | 1812 |
| YJM984 | XIV | 763 | 772 | 755178 | middle |  |  |  |  |  |  |  |  |  |  |  |
| YJM984 | XIV | 754658 | 754667 | 755178 | middle |  |  |  |  |  |  |  |  |  |  |  |
| YJM984 | XIV | 755169 | 755178 | 755178 | rightend | YNR071C | 100 | 1029 | 0 | 0 | 1 | 1029 | 744198 | 743170 | 0 | 1901 |
| YJM984 | XV | 1 | 10 | 1055074 | leftend | YOL154W | 99.467 | 750 | 4 | 0 | 1 | 750 | 22808 | 23557 | 0 | 1363 |
| YJM984 | XV | 1055065 | 1055074 | 1055074 | rightend | YOR381W | 100 | 2136 | 0 | 0 | 1 | 2136 | 1019998 | 1022133 | 0 | 3945 |
| YJM984 | XVI | 1 | 10 | 910079 | leftend | YPL274W | 99.83 | 1764 | 3 | 0 | 1 | 1764 | 17487 | 19250 | 0 | 3241 |
| YJM984 | XVI | 2432 | 2441 | 910079 | middle |  |  |  |  |  |  |  |  |  |  |  |
| YJM984 | XVI | 2754 | 2763 | 910079 | middle |  |  |  |  |  |  |  |  |  |  |  |
| YJM984 | XVI | 908243 | 908252 | 910079 | middle |  |  |  |  |  |  |  |  |  |  |  |
| YJM984 | XVI | 908490 | 908499 | 910079 | middle |  |  |  |  |  |  |  |  |  |  |  |
| YJM984 | XVI | 910070 | 910079 | 910079 | rightend | YPR201W | 100 | 1215 | 0 | 0 | 1 | 1215 | 905538 | 906752 | 0 | 2244 |
| YJM987 | I | 1 | 10 | 211794 | leftend | YAL063C | 94.132 | 1261 | 40 | 14 | 2715 | 3969 | 195650 | 196882 | 0 | 1888 |
| YJM987 | I | 211785 | 211794 | 211794 | rightend | YAR050W | 91.106 | 1428 | 103 | 7 | 2844 | 4250 | 195148 | 196572 | 0 | 1912 |
| YJM987 | II | 1 | 10 | 785315 | leftend | YNCB0001W | 100 | 84 | 0 | 0 | 1 | 84 | 2719 | 2802 | 4.16E-38 | 156 |
| YJM987 | II | 785306 | 785315 | 785315 | rightend | YBR297W | 99.076 | 1407 | 13 | 0 | 1 | 1407 | 775124 | 776530 | 0 | 2527 |
| YJM987 | III | 1 | 10 | 309400 | leftend | YCL073C | 100 | 1242 | 0 | 0 | 607 | 1848 | 7495 | 6254 | 0 | 2294 |
| YJM987 | III | 309391 | 309400 | 309400 | rightend | YCR101C | 99.818 | 549 | 1 | 0 | 1 | 549 | 296496 | 295948 | 0 | 1009 |
| YJM987 | IV | 1 | 10 | 1483626 | leftend | YDL243C | 95.051 | 990 | 49 | 0 | 1 | 990 | 16910 | 15921 | 0 | 1557 |
| YJM987 | IV | 1483617 | 1483626 | 1483626 | rightend | YDR541C | 99.42 | 1035 | 6 | 0 | 1 | 1035 | 1480888 | 1479854 | 0 | 1879 |
| YJM987 | V | 1 | 10 | 552848 | leftend | YEL070W | 98.608 | 1509 | 21 | 0 | 1 | 1509 | 12126 | 13634 | 0 | 2671 |
| YJM987 | V | 552839 | 552848 | 552848 | rightend | YER187W | 91.981 | 424 | 32 | 2 | 4 | 426 | 546568 | 546990 | 3.44E-169 | 593 |
| YJM987 | VI | 1 | 10 | 233479 | leftend | YFL051C | 98.23 | 452 | 8 | 0 | 1 | 452 | 863 | 412 | 0 | 791 |
| YJM987 | VI | 233470 | 233479 | 233479 | rightend | YFR055W | 99.902 | 1023 | 1 | 0 | 1 | 1023 | 227210 | 228232 | 0 | 1884 |
| YJM987 | VII | 1 | 10 | 1042635 | leftend | YGL258W | 97.585 | 621 | 15 | 0 | 1 | 621 | 4577 | 5197 | 0 | 1064 |
| YJM987 | VII | 1042626 | 1042635 | 1042635 | rightend | YGR287C | 99.016 | 1626 | 16 | 0 | 145 | 1770 | 1030346 | 1028721 | 0 | 2915 |
| YJM987 | VIII | 1 | 10 | 876714 | leftend | YPL274W | 99.887 | 1764 | 2 | 0 | 1 | 1764 | 10782 | 12545 | 0 | 3247 |
| YJM987 | VIII | 876705 | 876714 | 876714 | rightend | YHR211W | 82.535 | 939 | 153 | 7 | 1 | 932 | 865342 | 866276 | 0 | 815 |
| YJM987 | IX | 1 | 10 | 404193 | leftend | YIL169C | 93.928 | 1548 | 88 | 3 | 265 | 1812 | 2349 | 808 | 0 | 2333 |
| YJM987 | IX | 404184 | 404193 | 404193 | rightend | YIR039C | 99.009 | 1614 | 16 | 0 | 1 | 1614 | 402333 | 400720 | 0 | 2892 |
| YJM987 | X | 1 | 10 | 716810 | leftend | YJL213W | 99.699 | 996 | 3 | 0 | 1 | 996 | 26765 | 27760 | 0 | 1823 |
| YJM987 | X | 444 | 453 | 716810 | middle |  |  |  |  |  |  |  |  |  |  |  |
| YJM987 | X | 716810 | 716810 | 716810 | rightend_complete | YJR152W | 99.755 | 1632 | 3 | 1 | 1 | 1632 | 696401 | 698031 | 0 | 2990 |
| YJM987 | XI | 1 | 10 | 658828 | leftend | YKL222C | 99.339 | 2119 | 13 | 1 | 1 | 2118 | 3293 | 1175 | 0 | 3834 |
| YJM987 | XI | 658819 | 658828 | 658828 | rightend | YKR101W | 99.847 | 1965 | 3 | 0 | 1 | 1965 | 637235 | 639199 | 0 | 3613 |
| YJM987 | XII | 1 | 10 | 2344195 | leftend | YLL062C | 100 | 975 | 0 | 0 | 1 | 975 | 4099 | 3125 | 0 | 1801 |
| YJM987 | XII | 2343247 | 2343256 | 2344195 | middle |  |  |  |  |  |  |  |  |  |  |  |
| YJM987 | XII | 2344186 | 2344195 | 2344195 | rightend | YLR460C | 93.982 | 1130 | 66 | 2 | 3 | 1131 | 2340913 | 2339785 | 0 | 1709 |
| YJM987 | XIII | 1 | 10 | 901594 | leftend | YML131W | 100 | 1098 | 0 | 0 | 1 | 1098 | 3431 | 4528 | 0 | 2028 |
| YJM987 | XIII | 901585 | 901594 | 901594 | rightend | YMR321C | 99.361 | 313 | 2 | 0 | 6 | 318 | 887999 | 887687 | 2.48E-161 | 568 |
| YJM987 | XIV | 1 | 10 | 757482 | leftend | YNL332W | 98.631 | 1023 | 14 | 0 | 1 | 1023 | 7896 | 8918 | 0 | 1812 |
| YJM987 | XIV | 351 | 360 | 757482 | middle |  |  |  |  |  |  |  |  |  |  |  |
| YJM987 | XIV | 587 | 596 | 757482 | middle |  |  |  |  |  |  |  |  |  |  |  |
| YJM987 | XIV | 757473 | 757482 | 757482 | rightend | YNR071C | 100 | 1029 | 0 | 0 | 1 | 1029 | 746076 | 745048 | 0 | 1901 |
| YJM987 | XV | 1 | 10 | 1054158 | leftend | YOL154W | 99.467 | 750 | 4 | 0 | 1 | 750 | 23362 | 24111 | 0 | 1363 |
| YJM987 | XV | 1053655 | 1053664 | 1054158 | middle |  |  |  |  |  |  |  |  |  |  |  |
| YJM987 | XV | 1054149 | 1054158 | 1054158 | rightend | YOR381W | 100 | 2136 | 0 | 0 | 1 | 2136 | 1020633 | 1022768 | 0 | 3945 |
| YJM987 | XVI | 1 | 10 | 554614 | leftend | YHL044W | 98.446 | 708 | 11 | 0 | 1 | 708 | 8553 | 9260 | 0 | 1247 |
| YJM987 | XVI | 554605 | 554614 | 554614 | rightend | YPR201W | 100 | 1215 | 0 | 0 | 1 | 1215 | 551591 | 552805 | 0 | 2244 |
| YJM990 | I | 1 | 10 | 187666 | leftend | YAL063C | 85.784 | 1632 | 228 | 4 | 1 | 1630 | 16858 | 15229 | 0 | 1725 |
| YJM990 | I | 187657 | 187666 | 187666 | rightend | YAR050W | 87.925 | 1234 | 134 | 8 | 1 | 1225 | 186421 | 187648 | 0 | 1439 |
| YJM990 | II | 1 | 10 | 786922 | leftend | YNCB0001W | 100 | 84 | 0 | 0 | 1 | 84 | 4064 | 4147 | 4.16E-38 | 156 |
| YJM990 | II | 903 | 912 | 786922 | middle |  |  |  |  |  |  |  |  |  |  |  |
| YJM990 | II | 786913 | 786922 | 786922 | rightend | YBR297W | 99.076 | 1407 | 13 | 0 | 1 | 1407 | 776164 | 777570 | 0 | 2527 |
| YJM990 | III | 1 |  | 310325 | leftend_complete | YCL073C | 99.729 | 1848 | 5 | 0 | 1 | 1848 | 8368 | 6521 | 0 | 3386 |
| YJM990 | III | 608 | 617 | 310325 | middle |  |  |  |  |  |  |  |  |  |  |  |
| YJM990 | III | 309813 | 309822 | 310325 | middle |  |  |  |  |  |  |  |  |  |  |  |
| YJM990 | III | 310025 | 310034 | 310325 | middle |  |  |  |  |  |  |  |  |  |  |  |
| YJM990 | III | 310316 | 310325 | 310325 | rightend | YCR101C | 99.818 | 549 | 1 | 0 | 1 | 549 | 296437 | 295889 | 0 | 1009 |
| YJM990 | IV | 1 |  | 1469177 | leftend_complete | YDL243C | 95.051 | 990 | 49 | 0 | 1 | 990 | 19060 | 18071 | 0 | 1557 |
| YJM990 | IV | 3297 | 3306 | 1469177 | middle |  |  |  |  |  |  |  |  |  |  |  |
| YJM990 | IV | 1469168 | 1469177 | 1469177 | rightend | YDR541C | 99.42 | 1035 | 6 | 0 | 1 | 1035 | 1466544 | 1465510 | 0 | 1879 |
| YJM990 | V | 1 | 10 | 550381 | leftend | YEL070W | 98.873 | 1509 | 17 | 0 | 1 | 1509 | 13038 | 14546 | 0 | 2693 |
| YJM990 | V | 222 | 231 | 550381 | middle |  |  |  |  |  |  |  |  |  |  |  |
| YJM990 | V | 550372 | 550381 | 550381 | rightend | YER187W | 91.981 | 424 | 32 | 2 | 4 | 426 | 547809 | 548231 | 3.44E-169 | 593 |
| YJM990 | VI | 1 | 10 | 233135 | leftend | YFL051C | 98.451 | 452 | 7 | 0 | 1 | 452 | 863 | 412 | 0 | 797 |
| YJM990 | VI | 233135 | 233135 | 233135 | rightend_complete | YFR055W | 99.902 | 1023 | 1 | 0 | 1 | 1023 | 227197 | 228219 | 0 | 1884 |
| YJM990 | VII | 1 | 10 | 1036609 | leftend | YGL258W | 97.585 | 621 | 15 | 0 | 1 | 621 | 10757 | 11377 | 0 | 1064 |
| YJM990 | VII | 1036600 | 1036609 | 1036609 | rightend | YGR287C | 99.016 | 1626 | 16 | 0 | 145 | 1770 | 1030121 | 1028496 | 0 | 2915 |
| YJM990 | VIII | 1 | 10 | 521992 | leftend | YHL044W | 98.446 | 708 | 11 | 0 | 1 | 708 | 1304 | 2011 | 0 | 1247 |
| YJM990 | VIII | 521757 | 521766 | 521992 | middle |  |  |  |  |  |  |  |  |  |  |  |
| YJM990 | VIII | 521983 | 521992 | 521992 | rightend | YHR211W | 82.535 | 939 | 153 | 7 | 1 | 932 | 518304 | 519238 | 0 | 815 |
| YJM990 | IX | 1 | 10 | 410313 | leftend | YIL169C | 93.928 | 1548 | 88 | 3 | 265 | 1812 | 8386 | 6845 | 0 | 2333 |
| YJM990 | IX | 410304 | 410313 | 410313 | rightend | YIR039C | 98.947 | 1614 | 17 | 0 | 1 | 1614 | 408649 | 407036 | 0 | 2887 |
| YJM990 | X | 1 | 10 | 716746 | leftend | YJL213W | 99.699 | 996 | 3 | 0 | 1 | 996 | 26314 | 27309 | 0 | 1823 |
| YJM990 | X | 716746 | 716746 | 716746 | rightend_complete | YJR152W | 99.755 | 1632 | 3 | 1 | 1 | 1632 | 696478 | 698108 | 0 | 2990 |
| YJM990 | XI | 1 | 10 | 658750 | leftend | YKL222C | 99.339 | 2119 | 13 | 1 | 1 | 2118 | 3293 | 1175 | 0 | 3834 |
| YJM990 | XI | 658741 | 658750 | 658750 | rightend | YKR101W | 99.796 | 1965 | 4 | 0 | 1 | 1965 | 637423 | 639387 | 0 | 3607 |
| YJM990 | XII | 1 | 10 | 1827355 | leftend | YLL062C | 99.897 | 975 | 1 | 0 | 1 | 975 | 4706 | 3732 | 0 | 1796 |
| YJM990 | XII | 691 | 700 | 1827355 | middle |  |  |  |  |  |  |  |  |  |  |  |
| YJM990 | XII | 1827346 | 1827355 | 1827355 | rightend | YLR460C | 94.071 | 1130 | 65 | 2 | 3 | 1131 | 1816460 | 1815332 | 0 | 1714 |
| YJM990 | XIII | 1 | 10 | 912929 | leftend | YML131W | 100 | 1098 | 0 | 0 | 1 | 1098 | 4797 | 5894 | 0 | 2028 |
| YJM990 | XIII | 912920 | 912929 | 912929 | rightend | YMR321C | 99.361 | 313 | 2 | 0 | 6 | 318 | 889619 | 889307 | 2.48E-161 | 568 |
| YJM990 | XIV | 1 | 10 | 758021 | leftend | YNL332W | 98.631 | 1023 | 14 | 0 | 1 | 1023 | 5638 | 6660 | 0 | 1812 |
| YJM990 | XIV | 756490 | 756499 | 758021 | middle |  |  |  |  |  |  |  |  |  |  |  |
| YJM990 | XIV | 758012 | 758021 | 758021 | rightend | YNR071C | 100 | 1029 | 0 | 0 | 1 | 1029 | 743805 | 742777 | 0 | 1901 |
| YJM990 | XV | 1 | 10 | 1054403 | leftend | YOL154W | 99.467 | 750 | 4 | 0 | 1 | 750 | 22974 | 23723 | 0 | 1363 |
| YJM990 | XV | 1052899 | 1052908 | 1054403 | middle |  |  |  |  |  |  |  |  |  |  |  |
| YJM990 | XV | 1053611 | 1053620 | 1054403 | middle |  |  |  |  |  |  |  |  |  |  |  |
| YJM990 | XV | 1054058 | 1054067 | 1054403 | middle |  |  |  |  |  |  |  |  |  |  |  |
| YJM990 | XV | 1054394 | 1054403 | 1054403 | rightend | YOR381W | 100 | 2136 | 0 | 0 | 1 | 2136 | 1019794 | 1021929 | 0 | 3945 |
| YJM990 | XVI | 1 | 10 | 907818 | leftend | YPL274W | 99.83 | 1764 | 3 | 0 | 1 | 1764 | 18664 | 20427 | 0 | 3241 |
| YJM990 | XVI | 3670 | 3679 | 907818 | middle |  |  |  |  |  |  |  |  |  |  |  |
| YJM990 | XVI | 3920 | 3929 | 907818 | middle |  |  |  |  |  |  |  |  |  |  |  |
| YJM990 | XVI | 907809 | 907818 | 907818 | rightend | YPR201W | 100 | 1215 | 0 | 0 | 1 | 1215 | 904795 | 906009 | 0 | 2244 |
| YJM993 | I | 1 | 10 | 199866 | leftend | YAL063C | 85.784 | 1632 | 228 | 4 | 1 | 1630 | 18244 | 16615 | 0 | 1725 |
| YJM993 | I | 199866 | 199866 | 199866 | rightend_complete | YAR050W | 92.678 | 956 | 67 | 2 | 1 | 953 | 187810 | 188765 | 0 | 1375 |
| YJM993 | II | 1 | 10 | 788161 | leftend | YNCB0001W | 100 | 84 | 0 | 0 | 1 | 84 | 5869 | 5952 | 4.16E-38 | 156 |
| YJM993 | II | 788152 | 788161 | 788161 | rightend | YBR297W | 99.076 | 1407 | 13 | 0 | 1 | 1407 | 777967 | 779373 | 0 | 2527 |
| YJM993 | III | 1 | 10 | 309786 | leftend | YCL073C | 100 | 1242 | 0 | 0 | 607 | 1848 | 7277 | 6036 | 0 | 2294 |
| YJM993 | III | 309777 | 309786 | 309786 | rightend | YCR101C | 99.818 | 549 | 1 | 0 | 1 | 549 | 296578 | 296030 | 0 | 1009 |
| YJM993 | IV | 1 | 10 | 1459003 | leftend | YDL243C | 95.051 | 990 | 49 | 0 | 1 | 990 | 1918 | 929 | 0 | 1557 |
| YJM993 | IV | 1458994 | 1459003 | 1459003 | rightend | YDR541C | 99.42 | 1035 | 6 | 0 | 1 | 1035 | 1454969 | 1453935 | 0 | 1879 |
| YJM993 | V | 1 | 10 | 554810 | leftend | YEL070W | 98.873 | 1509 | 17 | 0 | 1 | 1509 | 14000 | 15508 | 0 | 2693 |
| YJM993 | V | 554801 | 554810 | 554810 | rightend | YER187W | 91.981 | 424 | 32 | 2 | 4 | 426 | 548494 | 548916 | 3.44E-169 | 593 |
| YJM993 | VI | 1 | 10 | 233873 | leftend | YFL051C | 98.23 | 452 | 8 | 0 | 1 | 452 | 1583 | 1132 | 0 | 791 |
| YJM993 | VI | 233873 | 233873 | 233873 | rightend_complete | YFR055W | 99.902 | 1023 | 1 | 0 | 1 | 1023 | 227923 | 228945 | 0 | 1884 |
| YJM993 | VII | 1 | 10 | 1043722 | leftend | YGL258W | 97.585 | 621 | 15 | 0 | 1 | 621 | 11236 | 11856 | 0 | 1064 |
| YJM993 | VII | 1043713 | 1043722 | 1043722 | rightend | YGR287C | 99.262 | 1626 | 12 | 0 | 145 | 1770 | 1031433 | 1029808 | 0 | 2937 |
| YJM993 | VIII | 1 | 10 | 519406 | leftend | YHL044W | 98.446 | 708 | 11 | 0 | 1 | 708 | 5526 | 6233 | 0 | 1247 |
| YJM993 | VIII | 519397 | 519406 | 519406 | rightend | YHR211W | 82.428 | 939 | 154 | 7 | 1 | 932 | 514367 | 515301 | 0 | 809 |
| YJM993 | IX | 1 | 10 | 404651 | leftend | YIL169C | 93.928 | 1548 | 88 | 3 | 265 | 1812 | 2349 | 808 | 0 | 2333 |
| YJM993 | IX | 404642 | 404651 | 404651 | rightend | YIR039C | 98.947 | 1614 | 17 | 0 | 1 | 1614 | 402581 | 400968 | 0 | 2887 |
| YJM993 | X | 1 | 10 | 722171 | leftend | YJL213W | 99.699 | 996 | 3 | 0 | 1 | 996 | 31786 | 32781 | 0 | 1823 |
| YJM993 | X | 722171 | 722171 | 722171 | rightend_complete | YJR152W | 99.755 | 1632 | 3 | 1 | 1 | 1632 | 701709 | 703339 | 0 | 2990 |
| YJM993 | XI | 1 | 10 | 658858 | leftend | YKL222C | 99.339 | 2119 | 13 | 1 | 1 | 2118 | 3293 | 1175 | 0 | 3834 |
| YJM993 | XI | 658849 | 658858 | 658858 | rightend | YKR101W | 99.796 | 1965 | 4 | 0 | 1 | 1965 | 637190 | 639154 | 0 | 3607 |
| YJM993 | XII | 1 | 10 | 1881170 | leftend | YLL062C | 100 | 975 | 0 | 0 | 1 | 975 | 4169 | 3195 | 0 | 1801 |
| YJM993 | XII | 1881161 | 1881170 | 1881170 | rightend | YLR460C | 93.982 | 1130 | 66 | 2 | 3 | 1131 | 1879011 | 1877883 | 0 | 1709 |
| YJM993 | XIII | 1 | 10 | 913885 | leftend | YML131W | 100 | 1098 | 0 | 0 | 1 | 1098 | 6276 | 7373 | 0 | 2028 |
| YJM993 | XIII | 913876 | 913885 | 913885 | rightend | YMR321C | 99.361 | 313 | 2 | 0 | 6 | 318 | 891118 | 890806 | 2.48E-161 | 568 |
| YJM993 | XIV | 1 | 10 | 757188 | leftend | YNL332W | 98.534 | 1023 | 15 | 0 | 1 | 1023 | 6866 | 7888 | 0 | 1807 |
| YJM993 | XIV | 757179 | 757188 | 757188 | rightend | YNR071C | 100 | 1029 | 0 | 0 | 1 | 1029 | 745331 | 744303 | 0 | 1901 |
| YJM993 | XV | 1 | 10 | 1052584 | leftend | YOL154W | 99.467 | 750 | 4 | 0 | 1 | 750 | 22975 | 23724 | 0 | 1363 |
| YJM993 | XV | 1052575 | 1052584 | 1052584 | rightend | YOR381W | 100 | 2136 | 0 | 0 | 1 | 2136 | 1020215 | 1022350 | 0 | 3945 |
| YJM993 | XVI | 1 | 10 | 905363 | leftend | YPL274W | 99.887 | 1764 | 2 | 0 | 1 | 1764 | 14741 | 16504 | 0 | 3247 |
| YJM993 | XVI | 905354 | 905363 | 905363 | rightend | YPR201W | 100 | 1215 | 0 | 0 | 1 | 1215 | 900494 | 901708 | 0 | 2244 |
| YJM996 | I | 1 |  | 182382 | leftend_complete | YAL062W | 99.054 | 1374 | 13 | 0 | 1 | 1374 | 179 | 1552 | 0 | 2466 |
| YJM996 | I | 182373 | 182382 | 182382 | rightend | YAR050W | 91.106 | 1428 | 103 | 7 | 2844 | 4250 | 166998 | 168422 | 0 | 1912 |
| YJM996 | II | 1 | 10 | 787769 | leftend | YNCB0001W | 100 | 84 | 0 | 0 | 1 | 84 | 5349 | 5432 | 4.16E-38 | 156 |
| YJM996 | II | 209 | 218 | 787769 | middle |  |  |  |  |  |  |  |  |  |  |  |
| YJM996 | II | 557 | 566 | 787769 | middle |  |  |  |  |  |  |  |  |  |  |  |
| YJM996 | II | 3149 | 3158 | 787769 | middle |  |  |  |  |  |  |  |  |  |  |  |
| YJM996 | II | 787760 | 787769 | 787769 | rightend | YBR297W | 99.076 | 1407 | 13 | 0 | 1 | 1407 | 777787 | 779193 | 0 | 2527 |
| YJM996 | III | 1 | 10 | 309807 | leftend | YCL073C | 99.784 | 1848 | 4 | 0 | 1 | 1848 | 8702 | 6855 | 0 | 3391 |
| YJM996 | III | 633 | 642 | 309807 | middle |  |  |  |  |  |  |  |  |  |  |  |
| YJM996 | III | 309798 | 309807 | 309807 | rightend | YCR101C | 99.818 | 549 | 1 | 0 | 1 | 549 | 296779 | 296231 | 0 | 1009 |
| YJM996 | IV | 1 | 10 | 1453867 | leftend | YDL243C | 95.051 | 990 | 49 | 0 | 1 | 990 | 1845 | 856 | 0 | 1557 |
| YJM996 | IV | 1453858 | 1453867 | 1453867 | rightend | YDR541C | 99.42 | 1035 | 6 | 0 | 1 | 1035 | 1451129 | 1450095 | 0 | 1879 |
| YJM996 | V | 1 | 10 | 554145 | leftend | YEL070W | 99.006 | 1509 | 15 | 0 | 1 | 1509 | 15408 | 16916 | 0 | 2704 |
| YJM996 | V | 554136 | 554145 | 554145 | rightend | YER187W | 91.981 | 424 | 32 | 2 | 4 | 426 | 550210 | 550632 | 3.44E-169 | 593 |
| YJM996 | VI | 1 | 10 | 235207 | leftend | YFL051C | 98.23 | 452 | 8 | 0 | 1 | 452 | 2348 | 1897 | 0 | 791 |
| YJM996 | VI | 897 | 906 | 235207 | middle |  |  |  |  |  |  |  |  |  |  |  |
| YJM996 | VI | 235198 | 235207 | 235207 | rightend | YFR055W | 99.902 | 1023 | 1 | 0 | 1 | 1023 | 228996 | 230018 | 0 | 1884 |
| YJM996 | VII | 1 |  | 1036847 | leftend_complete | YGL258W | 97.585 | 621 | 15 | 0 | 1 | 621 | 10525 | 11145 | 0 | 1064 |
| YJM996 | VII | 1036838 | 1036847 | 1036847 | rightend | YGR287C | 99.016 | 1626 | 16 | 0 | 145 | 1770 | 1030359 | 1028734 | 0 | 2915 |
| YJM996 | VIII | 1 | 10 | 894600 | leftend | YPL274W | 99.943 | 1764 | 1 | 0 | 1 | 1764 | 21584 | 23347 | 0 | 3253 |
| YJM996 | VIII | 1250 | 1259 | 894600 | middle |  |  |  |  |  |  |  |  |  |  |  |
| YJM996 | VIII | 894591 | 894600 | 894600 | rightend | YHR211W | 82.428 | 939 | 154 | 7 | 1 | 932 | 888815 | 889749 | 0 | 809 |
| YJM996 | IX | 1 | 10 | 411092 | leftend | YIL169C | 93.928 | 1548 | 88 | 3 | 265 | 1812 | 9116 | 7575 | 0 | 2333 |
| YJM996 | IX | 411083 | 411092 | 411092 | rightend | YIR039C | 99.009 | 1614 | 16 | 0 | 1 | 1614 | 409623 | 408010 | 0 | 2892 |
| YJM996 | X | 1 |  | 718334 | leftend_complete | YJL213W | 99.699 | 996 | 3 | 0 | 1 | 996 | 28166 | 29161 | 0 | 1823 |
| YJM996 | X | 1612 | 1621 | 718334 | middle |  |  |  |  |  |  |  |  |  |  |  |
| YJM996 | X | 1846 | 1855 | 718334 | middle |  |  |  |  |  |  |  |  |  |  |  |
| YJM996 | X | 718334 | 718334 | 718334 | rightend_complete | YJR152W | 99.755 | 1632 | 3 | 1 | 1 | 1632 | 697955 | 699585 | 0 | 2990 |
| YJM996 | XI | 1 | 10 | 660367 | leftend | YKL222C | 99.481 | 2118 | 11 | 0 | 1 | 2118 | 3686 | 1569 | 0 | 3851 |
| YJM996 | XI | 208 | 217 | 660367 | middle |  |  |  |  |  |  |  |  |  |  |  |
| YJM996 | XI | 659630 | 659639 | 660367 | middle |  |  |  |  |  |  |  |  |  |  |  |
| YJM996 | XI | 660367 | 660367 | 660367 | rightend_complete | YKR101W | 99.796 | 1965 | 4 | 0 | 1 | 1965 | 637629 | 639593 | 0 | 3607 |
| YJM996 | XII | 1 | 10 | 2116169 | leftend | YLL062C | 100 | 975 | 0 | 0 | 1 | 975 | 4778 | 3804 | 0 | 1801 |
| YJM996 | XII | 763 | 772 | 2116169 | middle |  |  |  |  |  |  |  |  |  |  |  |
| YJM996 | XII | 2115617 | 2115626 | 2116169 | middle |  |  |  |  |  |  |  |  |  |  |  |
| YJM996 | XII | 2115832 | 2115841 | 2116169 | middle |  |  |  |  |  |  |  |  |  |  |  |
| YJM996 | XII | 2116160 | 2116169 | 2116169 | rightend | YLR460C | 94.071 | 1130 | 65 | 2 | 3 | 1131 | 2113873 | 2112745 | 0 | 1714 |
| YJM996 | XIII | 1 |  | 899300 | leftend_complete | YML131W | 100 | 1098 | 0 | 0 | 1 | 1098 | 4739 | 5836 | 0 | 2028 |
| YJM996 | XIII | 640 | 649 | 899300 | middle |  |  |  |  |  |  |  |  |  |  |  |
| YJM996 | XIII | 1460 | 1469 | 899300 | middle |  |  |  |  |  |  |  |  |  |  |  |
| YJM996 | XIII | 899083 | 899092 | 899300 | middle |  |  |  |  |  |  |  |  |  |  |  |
| YJM996 | XIII | 899291 | 899300 | 899300 | rightend | YMR321C | 99.361 | 313 | 2 | 0 | 6 | 318 | 889657 | 889345 | 2.48E-161 | 568 |
| YJM996 | XIV | 1 | 10 | 755379 | leftend | YNL332W | 98.534 | 1023 | 15 | 0 | 1 | 1023 | 5586 | 6608 | 0 | 1807 |
| YJM996 | XIV | 755370 | 755379 | 755379 | rightend | YNR071C | 100 | 1029 | 0 | 0 | 1 | 1029 | 743993 | 742965 | 0 | 1901 |
| YJM996 | XV | 1 |  | 1045769 | leftend_complete | YOL154W | 99.467 | 750 | 4 | 0 | 1 | 750 | 29310 | 30059 | 0 | 1363 |
| YJM996 | XV | 1045406 | 1045415 | 1045769 | middle |  |  |  |  |  |  |  |  |  |  |  |
| YJM996 | XV | 1045769 | 1045769 | 1045769 | rightend_complete | YOR381W | 100 | 2136 | 0 | 0 | 1 | 2136 | 1026403 | 1028538 | 0 | 3945 |
| YJM996 | XVI | 1 |  | 558427 | leftend_complete | YHL044W | 98.446 | 708 | 11 | 0 | 1 | 708 | 8853 | 9560 | 0 | 1247 |
| YJM996 | XVI | 555442 | 555451 | 558427 | middle |  |  |  |  |  |  |  |  |  |  |  |
| YJM996 | XVI | 557435 | 557444 | 558427 | middle |  |  |  |  |  |  |  |  |  |  |  |
| YJM996 | XVI | 558418 | 558427 | 558427 | rightend | YPR201W | 100 | 1215 | 0 | 0 | 1 | 1215 | 552216 | 553430 | 0 | 2244 |
| YJM1078 | I | 1 | 10 | 183535 | leftend | YAL063C | 86.27 | 1362 | 183 | 4 | 1 | 1360 | 12825 | 11466 | 0 | 1476 |
| YJM1078 | I | 11042 | 11051 | 183535 | middle |  |  |  |  |  |  |  |  |  |  |  |
| YJM1078 | I | 183526 | 183535 | 183535 | rightend | YAR050W | 83.708 | 1246 | 199 | 4 | 1 | 1244 | 182282 | 183525 | 0 | 1173 |
| YJM1078 | II | 1 | 10 | 783609 | leftend | YNCB0001W | 100 | 84 | 0 | 0 | 1 | 84 | 1818 | 1901 | 4.16E-38 | 156 |
| YJM1078 | II | 783600 | 783609 | 783609 | rightend | YBR297W | 99.073 | 1403 | 13 | 0 | 1 | 1403 | 772973 | 774375 | 0 | 2519 |
| YJM1078 | III | 1 | 10 | 322339 | leftend | YCL073C | 99.621 | 1848 | 7 | 0 | 1 | 1848 | 10103 | 8256 | 0 | 3374 |
| YJM1078 | III | 322329 | 322339 | 322339 | rightend | YDR541C | 86.293 | 1036 | 137 | 5 | 1 | 1033 | 321529 | 320496 | 0 | 1122 |
| YJM1078 | IV | 1 | 10 | 1483189 | leftend | YDL243C | 95.051 | 990 | 49 | 0 | 1 | 990 | 2623 | 1634 | 0 | 1557 |
| YJM1078 | IV | 1483180 | 1483189 | 1483189 | rightend | YDR541C | 86.293 | 1036 | 137 | 5 | 1 | 1033 | 1479188 | 1478155 | 0 | 1122 |
| YJM1078 | V | 1 |  | 557633 | leftend_complete | YEL070W | 93.174 | 1509 | 103 | 0 | 1 | 1509 | 18186 | 19694 | 0 | 2217 |
| YJM1078 | V | 919 | 928 | 557633 | middle |  |  |  |  |  |  |  |  |  |  |  |
| YJM1078 | V | 557624 | 557633 | 557633 | rightend | YER187W | 99.296 | 426 | 3 | 0 | 1 | 426 | 554034 | 554459 | 0 | 771 |
| YJM1078 | VI | 1 | 10 | 252711 | leftend | YFL051C | 98.451 | 452 | 7 | 0 | 1 | 452 | 1698 | 1247 | 0 | 797 |
| YJM1078 | VI | 252702 | 252711 | 252711 | rightend | YFR055W | 90.873 | 1019 | 93 | 0 | 1 | 1019 | 229493 | 230511 | 0 | 1367 |
| YJM1078 | VII | 1 | 10 | 1037823 | leftend | YGL258W | 97.585 | 621 | 15 | 0 | 1 | 621 | 4018 | 4638 | 0 | 1064 |
| YJM1078 | VII | 1037823 | 1037823 | 1037823 | rightend_complete | YGR287C | 99.139 | 1626 | 14 | 0 | 145 | 1770 | 1025159 | 1023534 | 0 | 2926 |
| YJM1078 | VIII | 1 | 10 | 528800 | leftend | YHL044W | 98.446 | 708 | 11 | 0 | 1 | 708 | 3707 | 4414 | 0 | 1247 |
| YJM1078 | VIII | 981 | 990 | 528800 | middle |  |  |  |  |  |  |  |  |  |  |  |
| YJM1078 | VIII | 527293 | 527302 | 528800 | middle |  |  |  |  |  |  |  |  |  |  |  |
| YJM1078 | VIII | 527852 | 527861 | 528800 | middle |  |  |  |  |  |  |  |  |  |  |  |
| YJM1078 | VIII | 528791 | 528800 | 528800 | rightend | YHR211W | 82.428 | 939 | 154 | 7 | 1 | 932 | 515598 | 516532 | 0 | 809 |
| YJM1078 | IX | 1 | 10 | 425169 | leftend | YIL169C | 96.238 | 1648 | 55 | 5 | 272 | 1919 | 20118 | 18478 | 0 | 2693 |
| YJM1078 | IX | 425160 | 425169 | 425169 | rightend | YIR039C | 99.009 | 1614 | 16 | 0 | 1 | 1614 | 418838 | 417225 | 0 | 2892 |
| YJM1078 | X | 1 | 10 | 720411 | leftend | YJL213W | 99.699 | 996 | 3 | 0 | 1 | 996 | 31010 | 32005 | 0 | 1823 |
| YJM1078 | X | 719824 | 719833 | 720411 | middle |  |  |  |  |  |  |  |  |  |  |  |
| YJM1078 | X | 720411 | 720411 | 720411 | rightend_complete | YJR152W | 91.115 | 1632 | 145 | 0 | 1 | 1632 | 699976 | 701607 | 0 | 2211 |
| YJM1078 | XI | 1 | 10 | 665505 | leftend | YKL222C | 89.099 | 2119 | 229 | 2 | 1 | 2118 | 9679 | 7562 | 0 | 2632 |
| YJM1078 | XI | 665119 | 665128 | 665505 | middle |  |  |  |  |  |  |  |  |  |  |  |
| YJM1078 | XI | 665496 | 665505 | 665505 | rightend | YKR101W | 99.847 | 1965 | 3 | 0 | 1 | 1965 | 644331 | 646295 | 0 | 3613 |
| YJM1078 | XII | 1 | 10 | 1766831 | leftend | YLL062C | 100 | 975 | 0 | 0 | 1 | 975 | 3932 | 2958 | 0 | 1801 |
| YJM1078 | XII | 1766822 | 1766831 | 1766831 | rightend | YLR460C | 88.729 | 417 | 47 | 0 | 715 | 1131 | 1761252 | 1760836 | 3.46E-143 | 510 |
| YJM1078 | XIII | 1 | 10 | 908802 | leftend | YML131W | 100 | 1098 | 0 | 0 | 1 | 1098 | 5704 | 6801 | 0 | 2028 |
| YJM1078 | XIII | 2809 | 2818 | 908802 | middle |  |  |  |  |  |  |  |  |  |  |  |
| YJM1078 | XIII | 3068 | 3077 | 908802 | middle |  |  |  |  |  |  |  |  |  |  |  |
| YJM1078 | XIII | 908802 | 908802 | 908802 | rightend_complete | YMR321C | 99.361 | 313 | 2 | 0 | 6 | 318 | 890923 | 890611 | 2.48E-161 | 568 |
| YJM1078 | XIV | 1 | 10 | 738480 | leftend | YNL332W | 98.631 | 1023 | 14 | 0 | 1 | 1023 | 6187 | 7209 | 0 | 1812 |
| YJM1078 | XIV | 738480 | 738480 | 738480 | rightend_complete | YNR069C | 100 | 1470 | 0 | 0 | 1 | 1470 | 736249 | 734780 | 0 | 2715 |
| YJM1078 | XV | 1 | 10 | 1030520 | leftend | YOL154W | 99.867 | 750 | 1 | 0 | 1 | 750 | 23584 | 24333 | 0 | 1380 |
| YJM1078 | XV | 1030511 | 1030520 | 1030520 | rightend | YOR381W | 94.239 | 2135 | 121 | 2 | 1 | 2134 | 1021432 | 1023565 | 0 | 3260 |
| YJM1078 | XVI | 1 | 10 | 908543 | leftend | YPL274W | 99.83 | 1764 | 3 | 0 | 1 | 1764 | 18163 | 19926 | 0 | 3241 |
| YJM1078 | XVI | 3020 | 3029 | 908543 | middle |  |  |  |  |  |  |  |  |  |  |  |
| YJM1078 | XVI | 908543 | 908543 | 908543 | rightend_complete | YPR201W | 100 | 1215 | 0 | 0 | 1 | 1215 | 905253 | 906467 | 0 | 2244 |
| YJM1083 | I | 1 | 10 | 212046 | leftend | YAL063C | 89.745 | 2038 | 203 | 5 | 1 | 2035 | 13495 | 11461 | 0 | 2601 |
| YJM1083 | I | 212037 | 212046 | 212046 | rightend | YAR050W | 92.802 | 2209 | 112 | 12 | 2449 | 4614 | 189540 | 191744 | 0 | 3155 |
| YJM1083 | II | 1 |  | 787579 | leftend_complete | YNCB0001W | 100 | 84 | 0 | 0 | 1 | 84 | 10379 | 10462 | 4.16E-38 | 156 |
| YJM1083 | II | 787570 | 787579 | 787579 | rightend | YBR297W | 97.759 | 1205 | 27 | 0 | 1 | 1205 | 783377 | 784581 | 0 | 2076 |
| YJM1083 | III | 1 | 10 | 301172 | leftend | YCL073C | 98.593 | 1848 | 26 | 0 | 1 | 1848 | 7621 | 5774 | 0 | 3269 |
| YJM1083 | III | 301163 | 301172 | 301172 | rightend | YCR101C | 99.636 | 549 | 2 | 0 | 1 | 549 | 296488 | 295940 | 0 | 1003 |
| YJM1083 | IV | 1 | 10 | 1474624 | leftend | YDL243C | 93.769 | 979 | 61 | 0 | 1 | 979 | 15905 | 14927 | 0 | 1471 |
| YJM1083 | IV | 1474615 | 1474624 | 1474624 | rightend | YDR541C | 99.517 | 1035 | 5 | 0 | 1 | 1035 | 1469604 | 1468570 | 0 | 1884 |
| YJM1083 | V | 1 | 10 | 553383 | leftend | YEL070W | 99.735 | 1509 | 4 | 0 | 1 | 1509 | 16069 | 17577 | 0 | 2765 |
| YJM1083 | V | 553374 | 553383 | 553383 | rightend | YER187W | 99.765 | 425 | 1 | 0 | 2 | 426 | 549169 | 549593 | 0 | 780 |
| YJM1083 | VI | 1 | 10 | 288687 | leftend | YFL051C | 100 | 452 | 0 | 0 | 1 | 452 | 51056 | 50605 | 0 | 835 |
| YJM1083 | VI | 1965 | 1974 | 288687 | middle |  |  |  |  |  |  |  |  |  |  |  |
| YJM1083 | VI | 2503 | 2512 | 288687 | middle |  |  |  |  |  |  |  |  |  |  |  |
| YJM1083 | VI | 3489 | 3498 | 288687 | middle |  |  |  |  |  |  |  |  |  |  |  |
| YJM1083 | VI | 288678 | 288687 | 288687 | rightend | YFR055W | 99.511 | 1022 | 5 | 0 | 1 | 1022 | 278941 | 279962 | 0 | 1860 |
| YJM1083 | VII | 1 | 10 | 1048989 | leftend | YGL258W | 99.356 | 621 | 4 | 0 | 1 | 621 | 2391 | 3011 | 0 | 1125 |
| YJM1083 | VII | 1047604 | 1047613 | 1048989 | middle |  |  |  |  |  |  |  |  |  |  |  |
| YJM1083 | VII | 1048980 | 1048989 | 1048989 | rightend | YGR287C | 98.701 | 1770 | 23 | 0 | 1 | 1770 | 1022996 | 1021227 | 0 | 3142 |
| YJM1083 | VIII | 1 |  | 526378 | leftend_complete | YHL044W | 98.588 | 708 | 10 | 0 | 1 | 708 | 8839 | 9546 | 0 | 1253 |
| YJM1083 | VIII | 519403 | 519412 | 526378 | middle |  |  |  |  |  |  |  |  |  |  |  |
| YJM1083 | VIII | 520442 | 520451 | 526378 | middle |  |  |  |  |  |  |  |  |  |  |  |
| YJM1083 | VIII | 522389 | 522398 | 526378 | middle |  |  |  |  |  |  |  |  |  |  |  |
| YJM1083 | VIII | 525642 | 525651 | 526378 | middle |  |  |  |  |  |  |  |  |  |  |  |
| YJM1083 | VIII | 526369 | 526378 | 526378 | rightend | YHR211W | 85.3 | 1034 | 139 | 10 | 144 | 1169 | 519413 | 520441 | 0 | 1055 |
| YJM1083 | IX | 1 |  | 413951 | leftend_complete | YIL169C | 92.549 | 1369 | 96 | 4 | 444 | 1812 | 4884 | 3522 | 0 | 1958 |
| YJM1083 | IX | 776 | 785 | 413951 | middle |  |  |  |  |  |  |  |  |  |  |  |
| YJM1083 | IX | 413367 | 413376 | 413951 | middle |  |  |  |  |  |  |  |  |  |  |  |
| YJM1083 | IX | 413951 | 413951 | 413951 | rightend_complete | YIR039C | 99.009 | 1614 | 16 | 0 | 1 | 1614 | 411299 | 409686 | 0 | 2892 |
| YJM1083 | X | 1 |  | 700137 | leftend_complete | YJL213W | 99.699 | 996 | 3 | 0 | 1 | 996 | 10157 | 11152 | 0 | 1823 |
| YJM1083 | X | 378 | 387 | 700137 | middle |  |  |  |  |  |  |  |  |  |  |  |
| YJM1083 | X | 1120 | 1129 | 700137 | middle |  |  |  |  |  |  |  |  |  |  |  |
| YJM1083 | X | 697149 | 697158 | 700137 | middle |  |  |  |  |  |  |  |  |  |  |  |
| YJM1083 | X | 698006 | 698015 | 700137 | middle |  |  |  |  |  |  |  |  |  |  |  |
| YJM1083 | X | 699341 | 699350 | 700137 | middle |  |  |  |  |  |  |  |  |  |  |  |
| YJM1083 | X | 699806 | 699815 | 700137 | middle |  |  |  |  |  |  |  |  |  |  |  |
| YJM1083 | X | 700137 | 700137 | 700137 | rightend_complete | YJR152W | 100 | 1632 | 0 | 0 | 1 | 1632 | 680219 | 681850 | 0 | 3014 |
| YJM1083 | XI | 1 | 10 | 662304 | leftend | YKL222C | 99.906 | 2118 | 2 | 0 | 1 | 2118 | 3183 | 1066 | 0 | 3901 |
| YJM1083 | XI | 662295 | 662304 | 662304 | rightend | YKR101W | 99.695 | 1965 | 6 | 0 | 1 | 1965 | 637553 | 639517 | 0 | 3596 |
| YJM1083 | XII | 1 |  | 1781389 | leftend_complete | YLL062C | 99.59 | 975 | 4 | 0 | 1 | 975 | 11977 | 11003 | 0 | 1779 |
| YJM1083 | XII | 1096 | 1105 | 1781389 | middle |  |  |  |  |  |  |  |  |  |  |  |
| YJM1083 | XII | 1779019 | 1779028 | 1781389 | middle |  |  |  |  |  |  |  |  |  |  |  |
| YJM1083 | XII | 1781380 | 1781389 | 1781389 | rightend | YLR460C | 93.982 | 1130 | 66 | 2 | 3 | 1131 | 1777274 | 1776146 | 0 | 1709 |
| YJM1083 | XIII | 1 | 10 | 891852 | leftend | YML131W | 99.909 | 1098 | 1 | 0 | 1 | 1098 | 4815 | 5912 | 0 | 2023 |
| YJM1083 | XIII | 571 | 580 | 891852 | middle |  |  |  |  |  |  |  |  |  |  |  |
| YJM1083 | XIII | 1584 | 1593 | 891852 | middle |  |  |  |  |  |  |  |  |  |  |  |
| YJM1083 | XIII | 2457 | 2466 | 891852 | middle |  |  |  |  |  |  |  |  |  |  |  |
| YJM1083 | XIII | 890213 | 890222 | 891852 | middle |  |  |  |  |  |  |  |  |  |  |  |
| YJM1083 | XIII | 891852 | 891852 | 891852 | rightend_complete | YMR321C | 100 | 318 | 0 | 0 | 1 | 318 | 888034 | 887717 | 1.9E-167 | 588 |
| YJM1083 | XIV | 1 | 10 | 771803 | leftend | YNL332W | 99.902 | 1023 | 1 | 0 | 1 | 1023 | 10979 | 12001 | 0 | 1884 |
| YJM1083 | XIV | 803 | 812 | 771803 | middle |  |  |  |  |  |  |  |  |  |  |  |
| YJM1083 | XIV | 771794 | 771803 | 771803 | rightend | YNR071C | 99.514 | 1029 | 5 | 0 | 1 | 1029 | 753176 | 752148 | 0 | 1873 |
| YJM1083 | XV | 1 | 10 | 1058412 | leftend | YOL154W | 99.867 | 750 | 1 | 0 | 1 | 750 | 20201 | 20950 | 0 | 1380 |
| YJM1083 | XV | 1045022 | 1045031 | 1058412 | middle |  |  |  |  |  |  |  |  |  |  |  |
| YJM1083 | XV | 1058403 | 1058412 | 1058412 | rightend | YOR381W | 98.642 | 2136 | 29 | 0 | 1 | 2136 | 1016445 | 1018580 | 0 | 3784 |
| YJM1083 | XVI | 1 | 10 | 923698 | leftend | YPL274W | 99.83 | 1764 | 3 | 0 | 1 | 1764 | 31909 | 33672 | 0 | 3241 |
| YJM1083 | XVI | 15772 | 15781 | 923698 | middle |  |  |  |  |  |  |  |  |  |  |  |
| YJM1083 | XVI | 921559 | 921568 | 923698 | middle |  |  |  |  |  |  |  |  |  |  |  |
| YJM1083 | XVI | 922624 | 922633 | 923698 | middle |  |  |  |  |  |  |  |  |  |  |  |
| YJM1083 | XVI | 923689 | 923698 | 923698 | rightend | YPR201W | 99.588 | 1215 | 5 | 0 | 1 | 1215 | 918443 | 919657 | 0 | 2217 |
| YJM1129 | I | 1 | 10 | 180745 | leftend | YAL063C | 94.522 | 1004 | 28 | 16 | 2966 | 3969 | 1008 | 32 | 0 | 1524 |
| YJM1129 | I | 1970 | 1979 | 180745 | middle |  |  |  |  |  |  |  |  |  |  |  |
| YJM1129 | I | 180736 | 180745 | 180745 | rightend | YAR050W | 84.043 | 1222 | 191 | 4 | 1 | 1220 | 179516 | 180735 | 0 | 1173 |
| YJM1129 | II | 1 | 10 | 791955 | leftend | YNCB0001W | 100 | 84 | 0 | 0 | 1 | 84 | 7029 | 7112 | 4.16E-38 | 156 |
| YJM1129 | II | 2353 | 2362 | 791955 | middle |  |  |  |  |  |  |  |  |  |  |  |
| YJM1129 | II | 3367 | 3376 | 791955 | middle |  |  |  |  |  |  |  |  |  |  |  |
| YJM1129 | II | 4019 | 4028 | 791955 | middle |  |  |  |  |  |  |  |  |  |  |  |
| YJM1129 | II | 791946 | 791955 | 791955 | rightend | YBR297W | 99.716 | 1407 | 4 | 0 | 1 | 1407 | 780128 | 781534 | 0 | 2577 |
| YJM1129 | III | 1 | 10 | 309740 | leftend | YCL073C | 99.892 | 1848 | 2 | 0 | 1 | 1848 | 7456 | 5609 | 0 | 3402 |
| YJM1129 | III | 300534 | 300543 | 309740 | middle |  |  |  |  |  |  |  |  |  |  |  |
| YJM1129 | III | 309167 | 309176 | 309740 | middle |  |  |  |  |  |  |  |  |  |  |  |
| YJM1129 | III | 309740 | 309740 | 309740 | rightend_complete | YCR101C | 99.818 | 549 | 1 | 0 | 1 | 549 | 295600 | 295052 | 0 | 1009 |
| YJM1129 | IV | 1 |  | 1473665 | leftend_complete | YDL243C | 95.152 | 990 | 48 | 0 | 1 | 990 | 18558 | 17569 | 0 | 1563 |
| YJM1129 | IV | 1473656 | 1473665 | 1473665 | rightend | YDR541C | 99.324 | 1035 | 7 | 0 | 1 | 1035 | 1469321 | 1468287 | 0 | 1873 |
| YJM1129 | V | 1 |  | 557243 | leftend_complete | YEL070W | 98.873 | 1509 | 17 | 0 | 1 | 1509 | 17761 | 19269 | 0 | 2693 |
| YJM1129 | V | 4352 | 4361 | 557243 | middle |  |  |  |  |  |  |  |  |  |  |  |
| YJM1129 | V | 557234 | 557243 | 557243 | rightend | YER187W | 91.981 | 424 | 32 | 2 | 4 | 426 | 552551 | 552973 | 3.44E-169 | 593 |
| YJM1129 | VI | 1 |  | 234622 | leftend_complete | YFL051C | 98.23 | 452 | 8 | 0 | 1 | 452 | 1712 | 1261 | 0 | 791 |
| YJM1129 | VI | 192 | 201 | 234622 | middle |  |  |  |  |  |  |  |  |  |  |  |
| YJM1129 | VI | 234613 | 234622 | 234622 | rightend | YFR055W | 100 | 1023 | 0 | 0 | 1 | 1023 | 228359 | 229381 | 0 | 1890 |
| YJM1129 | VII | 1 |  | 1048825 | leftend_complete | YGL258W | 100 | 621 | 0 | 0 | 1 | 621 | 4657 | 5277 | 0 | 1147 |
| YJM1129 | VII | 517 | 526 | 1048825 | middle |  |  |  |  |  |  |  |  |  |  |  |
| YJM1129 | VII | 1048816 | 1048825 | 1048825 | rightend | YGR287C | 98.588 | 1770 | 25 | 0 | 1 | 1770 | 1025740 | 1023971 | 0 | 3131 |
| YJM1129 | VIII | 1 | 10 | 894204 | leftend | YPL274W | 99.887 | 1764 | 2 | 0 | 1 | 1764 | 14647 | 16410 | 0 | 3247 |
| YJM1129 | VIII | 883694 | 883703 | 894204 | middle |  |  |  |  |  |  |  |  |  |  |  |
| YJM1129 | VIII | 891667 | 891676 | 894204 | middle |  |  |  |  |  |  |  |  |  |  |  |
| YJM1129 | VIII | 892610 | 892619 | 894204 | middle |  |  |  |  |  |  |  |  |  |  |  |
| YJM1129 | VIII | 894195 | 894204 | 894204 | rightend | YHR211W | 92.416 | 2136 | 138 | 2 | 1096 | 3228 | 870211 | 872325 | 0 | 3025 |
| YJM1129 | IX | 1 | 10 | 412280 | leftend | YIL169C | 91.62 | 1420 | 117 | 2 | 394 | 1812 | 9156 | 7738 | 0 | 1962 |
| YJM1129 | IX | 412271 | 412280 | 412280 | rightend | YIR039C | 99.009 | 1614 | 16 | 0 | 1 | 1614 | 410419 | 408806 | 0 | 2892 |
| YJM1129 | X | 1 | 10 | 721636 | leftend | YJL213W | 99.699 | 996 | 3 | 0 | 1 | 996 | 30835 | 31830 | 0 | 1823 |
| YJM1129 | X | 4552 | 4561 | 721636 | middle |  |  |  |  |  |  |  |  |  |  |  |
| YJM1129 | X | 721636 | 721636 | 721636 | rightend_complete | YJR152W | 99.939 | 1632 | 1 | 0 | 1 | 1632 | 701104 | 702735 | 0 | 3009 |
| YJM1129 | XI | 1 | 10 | 665750 | leftend | YKL222C | 99.433 | 2118 | 12 | 0 | 1 | 2118 | 3203 | 1086 | 0 | 3845 |
| YJM1129 | XI | 665130 | 665139 | 665750 | middle |  |  |  |  |  |  |  |  |  |  |  |
| YJM1129 | XI | 665750 | 665750 | 665750 | rightend_complete | YKR101W | 99.847 | 1965 | 3 | 0 | 1 | 1965 | 637133 | 639097 | 0 | 3613 |
| YJM1129 | XII | 1 |  | 2440959 | leftend_complete | YLL062C | 99.795 | 975 | 2 | 0 | 1 | 975 | 6992 | 6018 | 0 | 1790 |
| YJM1129 | XII | 4487 | 4496 | 2440959 | middle |  |  |  |  |  |  |  |  |  |  |  |
| YJM1129 | XII | 2440950 | 2440959 | 2440959 | rightend | YLR460C | 94.248 | 1130 | 63 | 2 | 3 | 1131 | 2439209 | 2438081 | 0 | 1725 |
| YJM1129 | XIII | 1 | 10 | 918165 | leftend | YML131W | 100 | 1098 | 0 | 0 | 1 | 1098 | 22903 | 24000 | 0 | 2028 |
| YJM1129 | XIII | 1428 | 1437 | 918165 | middle |  |  |  |  |  |  |  |  |  |  |  |
| YJM1129 | XIII | 918156 | 918165 | 918165 | rightend | YMR321C | 99.361 | 313 | 2 | 0 | 6 | 318 | 907890 | 907578 | 2.48E-161 | 568 |
| YJM1129 | XIV | 1 | 10 | 758911 | leftend | YNL332W | 98.631 | 1023 | 14 | 0 | 1 | 1023 | 8549 | 9571 | 0 | 1812 |
| YJM1129 | XIV | 757430 | 757439 | 758911 | middle |  |  |  |  |  |  |  |  |  |  |  |
| YJM1129 | XIV | 758911 | 758911 | 758911 | rightend_complete | YNR071C | 99.903 | 1029 | 1 | 0 | 1 | 1029 | 746935 | 745907 | 0 | 1895 |
| YJM1129 | XV | 1 | 10 | 1049998 | leftend | YOL154W | 99.6 | 750 | 3 | 0 | 1 | 750 | 23036 | 23785 | 0 | 1369 |
| YJM1129 | XV | 1048027 | 1048036 | 1049998 | middle |  |  |  |  |  |  |  |  |  |  |  |
| YJM1129 | XV | 1049989 | 1049998 | 1049998 | rightend | YOR381W | 100 | 2136 | 0 | 0 | 1 | 2136 | 1020797 | 1022932 | 0 | 3945 |
| YJM1129 | XVI | 1 | 10 | 559831 | leftend | YHL044W | 98.446 | 708 | 11 | 0 | 1 | 708 | 8330 | 9037 | 0 | 1247 |
| YJM1129 | XVI | 559822 | 559831 | 559831 | rightend | YPR201W | 100 | 1215 | 0 | 0 | 1 | 1215 | 556975 | 558189 | 0 | 2244 |
| YJM1133 | I | 1 | 10 | 189480 | leftend | YAL063C | 92.229 | 1274 | 70 | 21 | 2715 | 3962 | 10805 | 9535 | 0 | 1777 |
| YJM1133 | I | 189471 | 189480 | 189480 | rightend | YAR050W | 90.646 | 1454 | 95 | 7 | 2869 | 4282 | 11285 | 9833 | 0 | 1893 |
| YJM1133 | II | 1 | 10 | 800176 | leftend | YNCB0001W | 100 | 84 | 0 | 0 | 1 | 84 | 5511 | 5594 | 4.16E-38 | 156 |
| YJM1133 | II | 649 | 658 | 800176 | middle |  |  |  |  |  |  |  |  |  |  |  |
| YJM1133 | II | 2000 | 2009 | 800176 | middle |  |  |  |  |  |  |  |  |  |  |  |
| YJM1133 | II | 800167 | 800176 | 800176 | rightend | YBR297W | 98.091 | 1205 | 23 | 0 | 1 | 1205 | 779987 | 781191 | 0 | 2098 |
| YJM1133 | III | 1 |  | 300819 | leftend_complete | YCL073C | 99.459 | 1848 | 10 | 0 | 1 | 1848 | 5259 | 3412 | 0 | 3358 |
| YJM1133 | III | 300810 | 300819 | 300819 | rightend | YCR101C | 99.636 | 549 | 2 | 0 | 1 | 549 | 295830 | 295282 | 0 | 1003 |
| YJM1133 | IV | 1 | 10 | 1472425 | leftend | YDL243C | 93.845 | 991 | 59 | 2 | 1 | 990 | 17622 | 16633 | 0 | 1491 |
| YJM1133 | IV | 598 | 607 | 1472425 | middle |  |  |  |  |  |  |  |  |  |  |  |
| YJM1133 | IV | 891 | 900 | 1472425 | middle |  |  |  |  |  |  |  |  |  |  |  |
| YJM1133 | IV | 1472011 | 1472020 | 1472425 | middle |  |  |  |  |  |  |  |  |  |  |  |
| YJM1133 | IV | 1472416 | 1472425 | 1472425 | rightend | YDR541C | 99.614 | 1035 | 4 | 0 | 1 | 1035 | 1460700 | 1459666 | 0 | 1890 |
| YJM1133 | V | 1 |  | 550488 | leftend_complete | YEL070W | 99.205 | 1509 | 12 | 0 | 1 | 1509 | 14671 | 16179 | 0 | 2721 |
| YJM1133 | V | 569 | 578 | 550488 | middle |  |  |  |  |  |  |  |  |  |  |  |
| YJM1133 | V | 789 | 798 | 550488 | middle |  |  |  |  |  |  |  |  |  |  |  |
| YJM1133 | V | 1386 | 1395 | 550488 | middle |  |  |  |  |  |  |  |  |  |  |  |
| YJM1133 | V | 550479 | 550488 | 550488 | rightend | YER187W | 91.981 | 424 | 32 | 2 | 4 | 426 | 547713 | 548135 | 3.44E-169 | 593 |
| YJM1133 | VI | 1 | 10 | 273929 | leftend | YFL051C | 100 | 452 | 0 | 0 | 1 | 452 | 37457 | 37006 | 0 | 835 |
| YJM1133 | VI | 273920 | 273929 | 273929 | rightend | YFR055W | 99.609 | 1022 | 4 | 0 | 1 | 1022 | 264442 | 265463 | 0 | 1866 |
| YJM1133 | VII | 1 | 10 | 1056905 | leftend | YGL258W | 97.585 | 621 | 15 | 0 | 1 | 621 | 10550 | 11170 | 0 | 1064 |
| YJM1133 | VII | 1056905 | 1056905 | 1056905 | rightend_complete | YGR287C | 99.209 | 1770 | 14 | 0 | 1 | 1770 | 1030442 | 1028673 | 0 | 3192 |
| YJM1133 | VIII | 1 |  | 540105 | leftend_complete | YHL044W | 98.446 | 708 | 11 | 0 | 1 | 708 | 8810 | 9517 | 0 | 1247 |
| YJM1133 | VIII | 540096 | 540105 | 540105 | rightend | YHR211W | 87.679 | 2443 | 227 | 31 | 826 | 3228 | 515964 | 518372 | 0 | 2776 |
| YJM1133 | IX | 1 |  | 424439 | leftend_complete | YIL169C | 95.158 | 1611 | 42 | 13 | 318 | 1919 | 20271 | 18688 | 0 | 2510 |
| YJM1133 | IX | 1465 | 1474 | 424439 | middle |  |  |  |  |  |  |  |  |  |  |  |
| YJM1133 | IX | 3064 | 3073 | 424439 | middle |  |  |  |  |  |  |  |  |  |  |  |
| YJM1133 | IX | 424059 | 424068 | 424439 | middle |  |  |  |  |  |  |  |  |  |  |  |
| YJM1133 | IX | 424439 | 424439 | 424439 | rightend_complete | YIR039C | 99.009 | 1614 | 16 | 0 | 1 | 1614 | 418118 | 416505 | 0 | 2892 |
| YJM1133 | X | 1 |  | 718481 | leftend_complete | YJL213W | 99.699 | 996 | 3 | 0 | 1 | 996 | 28083 | 29078 | 0 | 1823 |
| YJM1133 | X | 1693 | 1702 | 718481 | middle |  |  |  |  |  |  |  |  |  |  |  |
| YJM1133 | X | 2559 | 2568 | 718481 | middle |  |  |  |  |  |  |  |  |  |  |  |
| YJM1133 | X | 717942 | 717951 | 718481 | middle |  |  |  |  |  |  |  |  |  |  |  |
| YJM1133 | X | 718472 | 718481 | 718481 | rightend | YJR152W | 99.755 | 1632 | 3 | 1 | 1 | 1632 | 697812 | 699442 | 0 | 2990 |
| YJM1133 | XI | 1 | 10 | 666012 | leftend | YKL222C | 99.528 | 2118 | 10 | 0 | 1 | 2118 | 3371 | 1254 | 0 | 3856 |
| YJM1133 | XI | 665498 | 665507 | 666012 | middle |  |  |  |  |  |  |  |  |  |  |  |
| YJM1133 | XI | 666012 | 666012 | 666012 | rightend_complete | YKR101W | 99.847 | 1965 | 3 | 0 | 1 | 1965 | 637192 | 639156 | 0 | 3613 |
| YJM1133 | XII | 1 | 10 | 1700759 | leftend | YLL062C | 99.897 | 975 | 1 | 0 | 1 | 975 | 1875 | 901 | 0 | 1796 |
| YJM1133 | XII | 1698484 | 1698493 | 1700759 | middle |  |  |  |  |  |  |  |  |  |  |  |
| YJM1133 | XII | 1699088 | 1699097 | 1700759 | middle |  |  |  |  |  |  |  |  |  |  |  |
| YJM1133 | XII | 1700759 | 1700759 | 1700759 | rightend_complete | YLR460C | 98.762 | 1131 | 14 | 0 | 1 | 1131 | 1696886 | 1695756 | 0 | 2012 |
| YJM1133 | XIII | 1 | 10 | 898793 | leftend | YML131W | 100 | 1098 | 0 | 0 | 1 | 1098 | 3837 | 4934 | 0 | 2028 |
| YJM1133 | XIII | 601 | 610 | 898793 | middle |  |  |  |  |  |  |  |  |  |  |  |
| YJM1133 | XIII | 898189 | 898198 | 898793 | middle |  |  |  |  |  |  |  |  |  |  |  |
| YJM1133 | XIII | 898784 | 898793 | 898793 | rightend | YMR321C | 99.361 | 313 | 2 | 0 | 6 | 318 | 889940 | 889628 | 2.48E-161 | 568 |
| YJM1133 | XIV | 1 |  | 757662 | leftend_complete | YNL332W | 98.827 | 1023 | 12 | 0 | 1 | 1023 | 9154 | 10176 | 0 | 1823 |
| YJM1133 | XIV | 579 | 588 | 757662 | middle |  |  |  |  |  |  |  |  |  |  |  |
| YJM1133 | XIV | 6536 | 6545 | 757662 | middle |  |  |  |  |  |  |  |  |  |  |  |
| YJM1133 | XIV | 757653 | 757662 | 757662 | rightend | YNR071C | 99.611 | 1029 | 4 | 0 | 1 | 1029 | 748924 | 747896 | 0 | 1879 |
| YJM1133 | XV | 1 | 10 | 1042213 | leftend | YOL154W | 99.6 | 750 | 3 | 0 | 1 | 750 | 10829 | 11578 | 0 | 1369 |
| YJM1133 | XV | 1042213 | 1042213 | 1042213 | rightend_complete | YOR381W | 99.906 | 2136 | 2 | 0 | 1 | 2136 | 1008938 | 1011073 | 0 | 3934 |
| YJM1133 | XVI | 1 | 10 | 906596 | leftend | YPL274W | 98.129 | 1764 | 33 | 0 | 1 | 1764 | 17797 | 19560 | 0 | 3075 |
| YJM1133 | XVI | 1479 | 1488 | 906596 | middle |  |  |  |  |  |  |  |  |  |  |  |
| YJM1133 | XVI | 2050 | 2059 | 906596 | middle |  |  |  |  |  |  |  |  |  |  |  |
| YJM1133 | XVI | 2741 | 2750 | 906596 | middle |  |  |  |  |  |  |  |  |  |  |  |
| YJM1133 | XVI | 3257 | 3266 | 906596 | middle |  |  |  |  |  |  |  |  |  |  |  |
| YJM1133 | XVI | 906587 | 906596 | 906596 | rightend | YPR201W | 99.588 | 1215 | 5 | 0 | 1 | 1215 | 903203 | 904417 | 0 | 2217 |
| YJM1190 | I | 1 | 10 | 196015 | leftend | YAL063C | 91.72 | 1256 | 72 | 10 | 2715 | 3962 | 10443 | 9212 | 0 | 1714 |
| YJM1190 | I | 11194 | 11203 | 196015 | middle |  |  |  |  |  |  |  |  |  |  |  |
| YJM1190 | I | 188486 | 188495 | 196015 | middle |  |  |  |  |  |  |  |  |  |  |  |
| YJM1190 | I | 196006 | 196015 | 196015 | rightend | YAR050W | 92.895 | 1689 | 110 | 8 | 2599 | 4282 | 11193 | 9510 | 0 | 2447 |
| YJM1190 | II | 1 | 10 | 789596 | leftend | YNCB0001W | 100 | 84 | 0 | 0 | 1 | 84 | 3779 | 3862 | 4.16E-38 | 156 |
| YJM1190 | II | 1057 | 1066 | 789596 | middle |  |  |  |  |  |  |  |  |  |  |  |
| YJM1190 | II | 789596 | 789596 | 789596 | rightend_complete | YBR297W | 98.863 | 1407 | 14 | 1 | 1 | 1407 | 776751 | 778155 | 0 | 2508 |
| YJM1190 | III | 1 |  | 299381 | leftend_complete | YCL073C | 98.864 | 1848 | 21 | 0 | 1 | 1848 | 5257 | 3410 | 0 | 3297 |
| YJM1190 | III | 299104 | 299113 | 299381 | middle |  |  |  |  |  |  |  |  |  |  |  |
| YJM1190 | III | 299381 | 299381 | 299381 | rightend_complete | YCR101C | 98.364 | 550 | 8 | 1 | 1 | 549 | 294659 | 294110 | 0 | 965 |
| YJM1190 | IV | 1 |  | 1467476 | leftend_complete | YDL243C | 95.152 | 990 | 48 | 0 | 1 | 990 | 18489 | 17500 | 0 | 1563 |
| YJM1190 | IV | 1466606 | 1466615 | 1467476 | middle |  |  |  |  |  |  |  |  |  |  |  |
| YJM1190 | IV | 1467467 | 1467476 | 1467476 | rightend | YDR541C | 99.42 | 1035 | 6 | 0 | 1 | 1035 | 1463994 | 1462960 | 0 | 1879 |
| YJM1190 | V | 1 | 10 | 551345 | leftend | YEL070W | 99.139 | 1509 | 13 | 0 | 1 | 1509 | 13401 | 14909 | 0 | 2715 |
| YJM1190 | V | 551336 | 551345 | 551345 | rightend | YER187W | 91.981 | 424 | 32 | 2 | 4 | 426 | 548937 | 549359 | 3.44E-169 | 593 |
| YJM1190 | VI | 1 | 10 | 274246 | leftend | YFL051C | 98.451 | 452 | 7 | 0 | 1 | 452 | 36260 | 35809 | 0 | 797 |
| YJM1190 | VI | 274237 | 274246 | 274246 | rightend | YFR055W | 99.511 | 1022 | 5 | 0 | 1 | 1022 | 264779 | 265800 | 0 | 1860 |
| YJM1190 | VII | 1 | 10 | 1061252 | leftend | YGL258W | 99.678 | 621 | 2 | 0 | 1 | 621 | 10831 | 11451 | 0 | 1136 |
| YJM1190 | VII | 1061243 | 1061252 | 1061252 | rightend | YGR287C | 99.209 | 1770 | 14 | 0 | 1 | 1770 | 1032939 | 1031170 | 0 | 3192 |
| YJM1190 | VIII | 1 |  | 531761 | leftend_complete | YHL044W | 98.588 | 708 | 10 | 0 | 1 | 708 | 8867 | 9574 | 0 | 1253 |
| YJM1190 | VIII | 522787 | 522796 | 531761 | middle |  |  |  |  |  |  |  |  |  |  |  |
| YJM1190 | VIII | 529212 | 529221 | 531761 | middle |  |  |  |  |  |  |  |  |  |  |  |
| YJM1190 | VIII | 531761 | 531761 | 531761 | rightend_complete | YHR211W | 85.567 | 1552 | 212 | 8 | 1 | 1544 | 521239 | 522786 | 0 | 1615 |
| YJM1190 | IX | 1 |  | 406781 | leftend_complete | YIL169C | 94.092 | 1168 | 67 | 2 | 646 | 1812 | 5101 | 3935 | 0 | 1773 |
| YJM1190 | IX | 1363 | 1372 | 406781 | middle |  |  |  |  |  |  |  |  |  |  |  |
| YJM1190 | IX | 406158 | 406167 | 406781 | middle |  |  |  |  |  |  |  |  |  |  |  |
| YJM1190 | IX | 406772 | 406781 | 406781 | rightend | YIR039C | 99.69 | 1614 | 5 | 0 | 1 | 1614 | 404555 | 402942 | 0 | 2953 |
| YJM1190 | X | 1 |  | 726230 | leftend_complete | YJL213W | 99.799 | 996 | 2 | 0 | 1 | 996 | 36565 | 37560 | 0 | 1829 |
| YJM1190 | X | 726230 | 726230 | 726230 | rightend_complete | YJR152W | 99.755 | 1632 | 3 | 1 | 1 | 1632 | 705981 | 707611 | 0 | 2990 |
| YJM1190 | XI | 1 | 10 | 683070 | leftend | YKL222C | 99.387 | 2119 | 12 | 1 | 1 | 2118 | 4167 | 2049 | 0 | 3840 |
| YJM1190 | XI | 846 | 855 | 683070 | middle |  |  |  |  |  |  |  |  |  |  |  |
| YJM1190 | XI | 683070 | 683070 | 683070 | rightend_complete | YKR101W | 99.644 | 1965 | 7 | 0 | 1 | 1965 | 654758 | 656722 | 0 | 3591 |
| YJM1190 | XII | 1 | 10 | 2259125 | leftend | YLL062C | 100 | 975 | 0 | 0 | 1 | 975 | 6445 | 5471 | 0 | 1801 |
| YJM1190 | XII | 1532 | 1541 | 2259125 | middle |  |  |  |  |  |  |  |  |  |  |  |
| YJM1190 | XII | 2522 | 2531 | 2259125 | middle |  |  |  |  |  |  |  |  |  |  |  |
| YJM1190 | XII | 2258374 | 2258383 | 2259125 | middle |  |  |  |  |  |  |  |  |  |  |  |
| YJM1190 | XII | 2259125 | 2259125 | 2259125 | rightend_complete | YLR460C | 100 | 1131 | 0 | 0 | 1 | 1131 | 2255358 | 2254228 | 0 | 2089 |
| YJM1190 | XIII | 1 | 10 | 894228 | leftend | YML131W | 99.454 | 1098 | 6 | 0 | 1 | 1098 | 2087 | 3184 | 0 | 1995 |
| YJM1190 | XIII | 894228 | 894228 | 894228 | rightend_complete | YMR321C | 100 | 318 | 0 | 0 | 1 | 318 | 887798 | 887481 | 1.9E-167 | 588 |
| YJM1190 | XIV | 1 | 10 | 756042 | leftend | YNL332W | 92.958 | 71 | 5 | 0 | 1 | 71 | 6929 | 6999 | 2.4E-21 | 104 |
| YJM1190 | XIV | 1424 | 1433 | 756042 | middle |  |  |  |  |  |  |  |  |  |  |  |
| YJM1190 | XIV | 755553 | 755562 | 756042 | middle |  |  |  |  |  |  |  |  |  |  |  |
| YJM1190 | XIV | 756033 | 756042 | 756042 | rightend | YNR071C | 99.903 | 1029 | 1 | 0 | 1 | 1029 | 743948 | 742920 | 0 | 1895 |
| YJM1190 | XV | 1 | 10 | 1050602 | leftend | YOL154W | 99.067 | 750 | 7 | 0 | 1 | 750 | 23124 | 23873 | 0 | 1347 |
| YJM1190 | XV | 1044295 | 1044304 | 1050602 | middle |  |  |  |  |  |  |  |  |  |  |  |
| YJM1190 | XV | 1049692 | 1049701 | 1050602 | middle |  |  |  |  |  |  |  |  |  |  |  |
| YJM1190 | XV | 1049902 | 1049911 | 1050602 | middle |  |  |  |  |  |  |  |  |  |  |  |
| YJM1190 | XV | 1050593 | 1050602 | 1050602 | rightend | YOR381W | 98.689 | 2136 | 28 | 0 | 1 | 2136 | 1018541 | 1020676 | 0 | 3790 |
| YJM1190 | XVI | 1 | 10 | 906037 | leftend | YPL274W | 98.583 | 1764 | 25 | 0 | 1 | 1764 | 16571 | 18334 | 0 | 3120 |
| YJM1190 | XVI | 1282 | 1291 | 906037 | middle |  |  |  |  |  |  |  |  |  |  |  |
| YJM1190 | XVI | 3394 | 3403 | 906037 | middle |  |  |  |  |  |  |  |  |  |  |  |
| YJM1190 | XVI | 905643 | 905652 | 906037 | middle |  |  |  |  |  |  |  |  |  |  |  |
| YJM1190 | XVI | 906028 | 906037 | 906037 | rightend | YPR201W | 99.588 | 1215 | 5 | 0 | 1 | 1215 | 902838 | 904052 | 0 | 2217 |
| YJM1199 | I | 1 |  | 207131 | leftend_complete | YAL063C | 87.868 | 1665 | 192 | 8 | 1 | 1660 | 22641 | 20982 | 0 | 1947 |
| YJM1199 | I | 2216 | 2225 | 207131 | middle |  |  |  |  |  |  |  |  |  |  |  |
| YJM1199 | I | 2468 | 2477 | 207131 | middle |  |  |  |  |  |  |  |  |  |  |  |
| YJM1199 | I | 20312 | 20321 | 207131 | middle |  |  |  |  |  |  |  |  |  |  |  |
| YJM1199 | I | 20968 | 20977 | 207131 | middle |  |  |  |  |  |  |  |  |  |  |  |
| YJM1199 | I | 199004 | 199013 | 207131 | middle |  |  |  |  |  |  |  |  |  |  |  |
| YJM1199 | I | 199836 | 199845 | 207131 | middle |  |  |  |  |  |  |  |  |  |  |  |
| YJM1199 | I | 207122 | 207131 | 207131 | rightend | YAR050W | 99.027 | 822 | 8 | 0 | 3274 | 4095 | 199014 | 199835 | 0 | 1474 |
| YJM1199 | II | 1 | 10 | 782289 | leftend | YNCB0001W | 100 | 84 | 0 | 0 | 1 | 84 | 2422 | 2505 | 4.16E-38 | 156 |
| YJM1199 | II | 1048 | 1057 | 782289 | middle |  |  |  |  |  |  |  |  |  |  |  |
| YJM1199 | II | 780596 | 780605 | 782289 | middle |  |  |  |  |  |  |  |  |  |  |  |
| YJM1199 | II | 782280 | 782289 | 782289 | rightend | YBR297W | 98.257 | 1205 | 21 | 0 | 1 | 1205 | 776006 | 777210 | 0 | 2109 |
| YJM1199 | III | 1 | 10 | 309070 | leftend | YCL073C | 98.918 | 1848 | 20 | 0 | 1 | 1848 | 7825 | 5978 | 0 | 3302 |
| YJM1199 | III | 309061 | 309070 | 309070 | rightend | YCR101C | 99.818 | 549 | 1 | 0 | 1 | 549 | 296026 | 295478 | 0 | 1009 |
| YJM1199 | IV | 1 |  | 1465095 | leftend_complete | YDL243C | 93.845 | 991 | 59 | 2 | 1 | 990 | 18433 | 17444 | 0 | 1491 |
| YJM1199 | IV | 1463599 | 1463608 | 1465095 | middle |  |  |  |  |  |  |  |  |  |  |  |
| YJM1199 | IV | 1465095 | 1465095 | 1465095 | rightend_complete | YDR541C | 99.517 | 1035 | 5 | 0 | 1 | 1035 | 1460876 | 1459842 | 0 | 1884 |
| YJM1199 | V | 1 | 10 | 553511 | leftend | YEL070W | 99.072 | 1509 | 14 | 0 | 1 | 1509 | 13277 | 14785 | 0 | 2710 |
| YJM1199 | V | 551631 | 551640 | 553511 | middle |  |  |  |  |  |  |  |  |  |  |  |
| YJM1199 | V | 553511 | 553511 | 553511 | rightend_complete | YER187W | 91.981 | 424 | 32 | 2 | 4 | 426 | 548104 | 548526 | 3.44E-169 | 593 |
| YJM1199 | VI | 1 | 10 | 243087 | leftend | YFL051C | 98.451 | 452 | 7 | 0 | 1 | 452 | 5916 | 5465 | 0 | 797 |
| YJM1199 | VI | 607 | 616 | 243087 | middle |  |  |  |  |  |  |  |  |  |  |  |
| YJM1199 | VI | 243078 | 243087 | 243087 | rightend | YFR055W | 99.413 | 1022 | 6 | 0 | 1 | 1022 | 233529 | 234550 | 0 | 1855 |
| YJM1199 | VII | 1 | 10 | 1068067 | leftend | YGL258W | 97.424 | 621 | 16 | 0 | 1 | 621 | 19158 | 19778 | 0 | 1059 |
| YJM1199 | VII | 1064263 | 1064272 | 1068067 | middle |  |  |  |  |  |  |  |  |  |  |  |
| YJM1199 | VII | 1068067 | 1068067 | 1068067 | rightend_complete | YGR287C | 98.814 | 1770 | 21 | 0 | 1 | 1770 | 1041248 | 1039479 | 0 | 3153 |
| YJM1199 | VIII | 1 |  | 540055 | leftend_complete | YHL044W | 98.446 | 708 | 11 | 0 | 1 | 708 | 8796 | 9503 | 0 | 1247 |
| YJM1199 | VIII | 531906 | 531915 | 540055 | middle |  |  |  |  |  |  |  |  |  |  |  |
| YJM1199 | VIII | 536898 | 536907 | 540055 | middle |  |  |  |  |  |  |  |  |  |  |  |
| YJM1199 | VIII | 540055 | 540055 | 540055 | rightend_complete | YHR211W | 93.183 | 1731 | 94 | 2 | 1501 | 3228 | 518898 | 520607 | 0 | 2521 |
| YJM1199 | IX | 1 |  | 427528 | leftend_complete | YIL169C | 96.487 | 1907 | 57 | 8 | 562 | 2466 | 21861 | 19963 | 0 | 3142 |
| YJM1199 | IX | 2675 | 2684 | 427528 | middle |  |  |  |  |  |  |  |  |  |  |  |
| YJM1199 | IX | 426700 | 426709 | 427528 | middle |  |  |  |  |  |  |  |  |  |  |  |
| YJM1199 | IX | 427528 | 427528 | 427528 | rightend_complete | YIR039C | 99.009 | 1614 | 16 | 0 | 1 | 1614 | 420822 | 419209 | 0 | 2892 |
| YJM1199 | X | 1 | 10 | 723832 | leftend | YJL213W | 99.699 | 996 | 3 | 0 | 1 | 996 | 34356 | 35351 | 0 | 1823 |
| YJM1199 | X | 714 | 723 | 723832 | middle |  |  |  |  |  |  |  |  |  |  |  |
| YJM1199 | X | 723823 | 723832 | 723832 | rightend | YJR152W | 99.755 | 1632 | 3 | 1 | 1 | 1632 | 703703 | 705333 | 0 | 2990 |
| YJM1199 | XI | 1 | 10 | 666191 | leftend | YKL222C | 99.481 | 2118 | 11 | 0 | 1 | 2118 | 3974 | 1857 | 0 | 3851 |
| YJM1199 | XI | 775 | 784 | 666191 | middle |  |  |  |  |  |  |  |  |  |  |  |
| YJM1199 | XI | 665744 | 665753 | 666191 | middle |  |  |  |  |  |  |  |  |  |  |  |
| YJM1199 | XI | 666191 | 666191 | 666191 | rightend_complete | YKR101W | 99.847 | 1965 | 3 | 0 | 1 | 1965 | 637787 | 639751 | 0 | 3613 |
| YJM1199 | XII | 1 | 10 | 1630995 | leftend | YLL062C | 99.59 | 975 | 4 | 0 | 1 | 975 | 7838 | 6864 | 0 | 1779 |
| YJM1199 | XII | 691 | 700 | 1630995 | middle |  |  |  |  |  |  |  |  |  |  |  |
| YJM1199 | XII | 1630986 | 1630995 | 1630995 | rightend | YLR460C | 99.823 | 1131 | 2 | 0 | 1 | 1131 | 1628765 | 1627635 | 0 | 2078 |
| YJM1199 | XIII | 1 |  | 900978 | leftend_complete | YML131W | 100 | 1098 | 0 | 0 | 1 | 1098 | 2902 | 3999 | 0 | 2028 |
| YJM1199 | XIII | 289 | 298 | 900978 | middle |  |  |  |  |  |  |  |  |  |  |  |
| YJM1199 | XIII | 895508 | 895517 | 900978 | middle |  |  |  |  |  |  |  |  |  |  |  |
| YJM1199 | XIII | 900969 | 900978 | 900978 | rightend | YMR321C | 99.361 | 313 | 2 | 0 | 6 | 318 | 888472 | 888160 | 2.48E-161 | 568 |
| YJM1199 | XIV | 1 |  | 759900 | leftend_complete | YNL332W | 92.958 | 71 | 5 | 0 | 1 | 71 | 9677 | 9747 | 2.4E-21 | 104 |
| YJM1199 | XIV | 758032 | 758041 | 759900 | middle |  |  |  |  |  |  |  |  |  |  |  |
| YJM1199 | XIV | 758798 | 758807 | 759900 | middle |  |  |  |  |  |  |  |  |  |  |  |
| YJM1199 | XIV | 759900 | 759900 | 759900 | rightend_complete | YNR071C | 100 | 1029 | 0 | 0 | 1 | 1029 | 748297 | 747269 | 0 | 1901 |
| YJM1199 | XV | 1 | 10 | 1047277 | leftend | YOL154W | 99.867 | 750 | 1 | 0 | 1 | 750 | 20654 | 21403 | 0 | 1380 |
| YJM1199 | XV | 1045673 | 1045682 | 1047277 | middle |  |  |  |  |  |  |  |  |  |  |  |
| YJM1199 | XV | 1047268 | 1047277 | 1047277 | rightend | YOR381W | 100 | 2136 | 0 | 0 | 1 | 2136 | 1018368 | 1020503 | 0 | 3945 |
| YJM1199 | XVI | 1 | 10 | 912000 | leftend | YPL274W | 99.376 | 1764 | 11 | 0 | 1 | 1764 | 18239 | 20002 | 0 | 3197 |
| YJM1199 | XVI | 3586 | 3595 | 912000 | middle |  |  |  |  |  |  |  |  |  |  |  |
| YJM1199 | XVI | 911535 | 911544 | 912000 | middle |  |  |  |  |  |  |  |  |  |  |  |
| YJM1199 | XVI | 911991 | 912000 | 912000 | rightend | YPR201W | 99.588 | 1215 | 5 | 0 | 1 | 1215 | 908435 | 909649 | 0 | 2217 |
| YJM1202 | I | 1 |  | 212216 | leftend_complete | YAL063C | 88.471 | 2342 | 197 | 38 | 1642 | 3969 | 197014 | 199296 | 0 | 2761 |
| YJM1202 | I | 210648 | 210657 | 212216 | middle |  |  |  |  |  |  |  |  |  |  |  |
| YJM1202 | I | 212207 | 212216 | 212216 | rightend | YAR050W | 92.882 | 1995 | 122 | 7 | 2304 | 4282 | 197001 | 198991 | 0 | 2880 |
| YJM1202 | II | 1 |  | 780867 | leftend_complete | YNCB0001W | 100 | 84 | 0 | 0 | 1 | 84 | 3381 | 3464 | 4.16E-38 | 156 |
| YJM1202 | II | 780858 | 780867 | 780867 | rightend | YBR297W | 98.257 | 1205 | 21 | 0 | 1 | 1205 | 776635 | 777839 | 0 | 2109 |
| YJM1202 | III | 1 | 10 | 308414 | leftend | YCL073C | 99.838 | 1848 | 3 | 0 | 1 | 1848 | 7777 | 5930 | 0 | 3397 |
| YJM1202 | III | 308405 | 308414 | 308414 | rightend | YCR101C | 99.818 | 549 | 1 | 0 | 1 | 549 | 295387 | 294839 | 0 | 1009 |
| YJM1202 | IV | 1 | 10 | 1465717 | leftend | YDL243C | 95.253 | 990 | 47 | 0 | 1 | 990 | 17638 | 16649 | 0 | 1568 |
| YJM1202 | IV | 1465717 | 1465717 | 1465717 | rightend_complete | YDR541C | 99.517 | 1035 | 5 | 0 | 1 | 1035 | 1460983 | 1459949 | 0 | 1884 |
| YJM1202 | V | 1 | 10 | 556739 | leftend | YEL070W | 99.072 | 1509 | 14 | 0 | 1 | 1509 | 13391 | 14899 | 0 | 2710 |
| YJM1202 | V | 556739 | 556739 | 556739 | rightend_complete | YER187W | 91.981 | 424 | 32 | 2 | 4 | 426 | 548231 | 548653 | 3.44E-169 | 593 |
| YJM1202 | VI | 1 |  | 240195 | leftend_complete | YFL051C | 98.23 | 452 | 8 | 0 | 1 | 452 | 6712 | 6261 | 0 | 791 |
| YJM1202 | VI | 240195 | 240195 | 240195 | rightend_complete | YFR055W | 99.902 | 1023 | 1 | 0 | 1 | 1023 | 234216 | 235238 | 0 | 1884 |
| YJM1202 | VII | 1 |  | 1059632 | leftend_complete | YGL258W | 97.424 | 621 | 16 | 0 | 1 | 621 | 10956 | 11576 | 0 | 1059 |
| YJM1202 | VII | 6470 | 6479 | 1059632 | middle |  |  |  |  |  |  |  |  |  |  |  |
| YJM1202 | VII | 1059632 | 1059632 | 1059632 | rightend_complete | YGR287C | 98.87 | 1770 | 20 | 0 | 1 | 1770 | 1032577 | 1030808 | 0 | 3158 |
| YJM1202 | VIII | 1 | 10 | 529959 | leftend | YHL044W | 98.446 | 708 | 11 | 0 | 1 | 708 | 3070 | 3777 | 0 | 1247 |
| YJM1202 | VIII | 523074 | 523083 | 529959 | middle |  |  |  |  |  |  |  |  |  |  |  |
| YJM1202 | VIII | 529959 | 529959 | 529959 | rightend_complete | YHR211W | 87.273 | 2428 | 266 | 26 | 826 | 3221 | 511409 | 513825 | 0 | 2732 |
| YJM1202 | IX | 1 |  | 427811 | leftend_complete | YIL169C | 96.539 | 1907 | 56 | 8 | 562 | 2466 | 20068 | 18170 | 0 | 3147 |
| YJM1202 | IX | 622 | 631 | 427811 | middle |  |  |  |  |  |  |  |  |  |  |  |
| YJM1202 | IX | 427811 | 427811 | 427811 | rightend_complete | YIR039C | 99.938 | 1614 | 1 | 0 | 1 | 1614 | 419982 | 418369 | 0 | 2976 |
| YJM1202 | X | 1 |  | 898819 | leftend_complete | YJL213W | 99.699 | 996 | 3 | 0 | 1 | 996 | 123036 | 124031 | 0 | 1823 |
| YJM1202 | X | 898819 | 898819 | 898819 | rightend_complete | YJR152W | 99.755 | 1632 | 3 | 1 | 1 | 1632 | 792749 | 794379 | 0 | 2990 |
| YJM1202 | XI | 1 | 10 | 666631 | leftend | YKL222C | 99.528 | 2118 | 10 | 0 | 1 | 2118 | 6835 | 4718 | 0 | 3856 |
| YJM1202 | XI | 1316 | 1325 | 666631 | middle |  |  |  |  |  |  |  |  |  |  |  |
| YJM1202 | XI | 1903 | 1912 | 666631 | middle |  |  |  |  |  |  |  |  |  |  |  |
| YJM1202 | XI | 2435 | 2444 | 666631 | middle |  |  |  |  |  |  |  |  |  |  |  |
| YJM1202 | XI | 3463 | 3472 | 666631 | middle |  |  |  |  |  |  |  |  |  |  |  |
| YJM1202 | XI | 666130 | 666139 | 666631 | middle |  |  |  |  |  |  |  |  |  |  |  |
| YJM1202 | XI | 666631 | 666631 | 666631 | rightend_complete | YKR101W | 99.847 | 1965 | 3 | 0 | 1 | 1965 | 640748 | 642712 | 0 | 3613 |
| YJM1202 | XII | 1 |  | 1797287 | leftend_complete | YLL062C | 99.59 | 975 | 4 | 0 | 1 | 975 | 6357 | 5383 | 0 | 1779 |
| YJM1202 | XII | 1797287 | 1797287 | 1797287 | rightend_complete | YLR460C | 99.823 | 1131 | 2 | 0 | 1 | 1131 | 1792812 | 1791682 | 0 | 2078 |
| YJM1202 | XIII | 1 | 10 | 910036 | leftend | YML131W | 100 | 1098 | 0 | 0 | 1 | 1098 | 4700 | 5797 | 0 | 2028 |
| YJM1202 | XIII | 2425 | 2434 | 910036 | middle |  |  |  |  |  |  |  |  |  |  |  |
| YJM1202 | XIII | 910036 | 910036 | 910036 | rightend_complete | YMR321C | 99.361 | 313 | 2 | 0 | 6 | 318 | 890678 | 890366 | 2.48E-161 | 568 |
| YJM1202 | XIV | 1 | 10 | 757370 | leftend | YNL332W | 98.535 | 1024 | 14 | 1 | 1 | 1023 | 6487 | 7510 | 0 | 1807 |
| YJM1202 | XIV | 378 | 387 | 757370 | middle |  |  |  |  |  |  |  |  |  |  |  |
| YJM1202 | XIV | 630 | 639 | 757370 | middle |  |  |  |  |  |  |  |  |  |  |  |
| YJM1202 | XIV | 756839 | 756848 | 757370 | middle |  |  |  |  |  |  |  |  |  |  |  |
| YJM1202 | XIV | 757361 | 757370 | 757370 | rightend | YNR071C | 100 | 1029 | 0 | 0 | 1 | 1029 | 746607 | 745579 | 0 | 1901 |
| YJM1202 | XV | 1 | 10 | 1066582 | leftend | YOL154W | 99.867 | 750 | 1 | 0 | 1 | 750 | 35133 | 35882 | 0 | 1380 |
| YJM1202 | XV | 1064465 | 1064474 | 1066582 | middle |  |  |  |  |  |  |  |  |  |  |  |
| YJM1202 | XV | 1066573 | 1066582 | 1066582 | rightend | YOR381W | 98.83 | 2136 | 25 | 0 | 1 | 2136 | 1033720 | 1035855 | 0 | 3807 |
| YJM1202 | XVI | 1 |  | 906061 | leftend_complete | YPL274W | 99.887 | 1764 | 2 | 0 | 1 | 1764 | 16531 | 18294 | 0 | 3247 |
| YJM1202 | XVI | 906052 | 906061 | 906061 | rightend | YPR201W | 99.588 | 1215 | 5 | 0 | 1 | 1215 | 902369 | 903583 | 0 | 2217 |
| YJM1208 | I | 1 | 10 | 216487 | leftend | YAL063C | 86.111 | 1908 | 249 | 14 | 1 | 1900 | 12474 | 10575 | 0 | 2041 |
| YJM1208 | I | 9927 | 9936 | 216487 | middle |  |  |  |  |  |  |  |  |  |  |  |
| YJM1208 | I | 189401 | 189410 | 216487 | middle |  |  |  |  |  |  |  |  |  |  |  |
| YJM1208 | I | 210763 | 210772 | 216487 | middle |  |  |  |  |  |  |  |  |  |  |  |
| YJM1208 | I | 216478 | 216487 | 216487 | rightend | YAR050W | 93.586 | 1107 | 41 | 4 | 3535 | 4614 | 189411 | 190514 | 0 | 1624 |
| YJM1208 | II | 1 | 10 | 780639 | leftend | YNCB0001W | 100 | 84 | 0 | 0 | 1 | 84 | 1582 | 1665 | 4.16E-38 | 156 |
| YJM1208 | II | 780630 | 780639 | 780639 | rightend | YBR297W | 98.008 | 1205 | 24 | 0 | 1 | 1205 | 776403 | 777607 | 0 | 2093 |
| YJM1208 | III | 1 | 10 | 299923 | leftend | YCL073C | 99.892 | 1848 | 2 | 0 | 1 | 1848 | 8393 | 6546 | 0 | 3402 |
| YJM1208 | III | 299914 | 299923 | 299923 | rightend | YCR101C | 100 | 549 | 0 | 0 | 1 | 549 | 295397 | 294849 | 0 | 1014 |
| YJM1208 | IV | 1 | 10 | 1493437 | leftend | YDL243C | 93.769 | 979 | 61 | 0 | 1 | 979 | 22305 | 21327 | 0 | 1471 |
| YJM1208 | IV | 1491715 | 1491724 | 1493437 | middle |  |  |  |  |  |  |  |  |  |  |  |
| YJM1208 | IV | 1493437 | 1493437 | 1493437 | rightend_complete | YDR541C | 99.903 | 1035 | 1 | 0 | 1 | 1035 | 1486835 | 1485801 | 0 | 1906 |
| YJM1208 | V | 1 |  | 554157 | leftend_complete | YEL070W | 99.669 | 1509 | 5 | 0 | 1 | 1509 | 16767 | 18275 | 0 | 2760 |
| YJM1208 | V | 554148 | 554157 | 554157 | rightend | YER187W | 99.531 | 426 | 2 | 0 | 1 | 426 | 550507 | 550932 | 0 | 776 |
| YJM1208 | VI | 1 | 10 | 263455 | leftend | YFL051C | 100 | 483 | 0 | 0 | 1 | 483 | 28331 | 27849 | 0 | 893 |
| YJM1208 | VI | 263455 | 263455 | 263455 | rightend_complete | YFR055W | 99.902 | 1023 | 1 | 0 | 1 | 1023 | 257428 | 258450 | 0 | 1884 |
| YJM1208 | VII | 1 |  | 1043993 | leftend_complete | YGL258W | 99.356 | 621 | 4 | 0 | 1 | 621 | 8702 | 9322 | 0 | 1125 |
| YJM1208 | VII | 6541 | 6550 | 1043993 | middle |  |  |  |  |  |  |  |  |  |  |  |
| YJM1208 | VII | 1043984 | 1043993 | 1043993 | rightend | YGR287C | 99.209 | 1770 | 14 | 0 | 1 | 1770 | 1032111 | 1030342 | 0 | 3192 |
| YJM1208 | VIII | 1 |  | 530127 | leftend_complete | YHL044W | 100 | 708 | 0 | 0 | 1 | 708 | 9730 | 10437 | 0 | 1308 |
| YJM1208 | VIII | 1162 | 1171 | 530127 | middle |  |  |  |  |  |  |  |  |  |  |  |
| YJM1208 | VIII | 1913 | 1922 | 530127 | middle |  |  |  |  |  |  |  |  |  |  |  |
| YJM1208 | VIII | 519931 | 519940 | 530127 | middle |  |  |  |  |  |  |  |  |  |  |  |
| YJM1208 | VIII | 522700 | 522709 | 530127 | middle |  |  |  |  |  |  |  |  |  |  |  |
| YJM1208 | VIII | 525303 | 525312 | 530127 | middle |  |  |  |  |  |  |  |  |  |  |  |
| YJM1208 | VIII | 530127 | 530127 | 530127 | rightend_complete | YHR211W | 83.197 | 1226 | 188 | 14 | 1 | 1217 | 518706 | 519922 | 0 | 1107 |
| YJM1208 | IX | 1 | 10 | 443326 | leftend | YIL169C | 96.725 | 1374 | 39 | 2 | 478 | 1851 | 30847 | 29480 | 0 | 2283 |
| YJM1208 | IX | 11739 | 11748 | 443326 | middle |  |  |  |  |  |  |  |  |  |  |  |
| YJM1208 | IX | 443326 | 443326 | 443326 | rightend_complete | YIR039C | 99.814 | 1614 | 3 | 0 | 1 | 1614 | 435554 | 433941 | 0 | 2964 |
| YJM1208 | X | 1 |  | 730593 | leftend_complete | YJL213W | 99.699 | 996 | 3 | 0 | 1 | 996 | 38968 | 39963 | 0 | 1823 |
| YJM1208 | X | 1161 | 1170 | 730593 | middle |  |  |  |  |  |  |  |  |  |  |  |
| YJM1208 | X | 2159 | 2168 | 730593 | middle |  |  |  |  |  |  |  |  |  |  |  |
| YJM1208 | X | 727799 | 727808 | 730593 | middle |  |  |  |  |  |  |  |  |  |  |  |
| YJM1208 | X | 728528 | 728537 | 730593 | middle |  |  |  |  |  |  |  |  |  |  |  |
| YJM1208 | X | 729403 | 729412 | 730593 | middle |  |  |  |  |  |  |  |  |  |  |  |
| YJM1208 | X | 730593 | 730593 | 730593 | rightend_complete | YJR152W | 99.939 | 1632 | 1 | 0 | 1 | 1632 | 714203 | 715834 | 0 | 3009 |
| YJM1208 | XI | 1 | 10 | 660758 | leftend | YKL222C | 99.622 | 2118 | 8 | 0 | 1 | 2118 | 3412 | 1295 | 0 | 3868 |
| YJM1208 | XI | 660253 | 660262 | 660758 | middle |  |  |  |  |  |  |  |  |  |  |  |
| YJM1208 | XI | 660758 | 660758 | 660758 | rightend_complete | YKR101W | 99.796 | 1965 | 4 | 0 | 1 | 1965 | 637590 | 639554 | 0 | 3607 |
| YJM1208 | XII | 1 |  | 2358591 | leftend_complete | YLL062C | 99.282 | 975 | 7 | 0 | 1 | 975 | 5784 | 4810 | 0 | 1762 |
| YJM1208 | XII | 1563 | 1572 | 2358591 | middle |  |  |  |  |  |  |  |  |  |  |  |
| YJM1208 | XII | 2356879 | 2356888 | 2358591 | middle |  |  |  |  |  |  |  |  |  |  |  |
| YJM1208 | XII | 2358591 | 2358591 | 2358591 | rightend_complete | YLR460C | 94.159 | 1130 | 64 | 2 | 3 | 1131 | 2354379 | 2353251 | 0 | 1720 |
| YJM1208 | XIII | 1 |  | 905258 | leftend_complete | YML131W | 99.545 | 1098 | 5 | 0 | 1 | 1098 | 13313 | 14410 | 0 | 2001 |
| YJM1208 | XIII | 455 | 464 | 905258 | middle |  |  |  |  |  |  |  |  |  |  |  |
| YJM1208 | XIII | 901606 | 901615 | 905258 | middle |  |  |  |  |  |  |  |  |  |  |  |
| YJM1208 | XIII | 903617 | 903626 | 905258 | middle |  |  |  |  |  |  |  |  |  |  |  |
| YJM1208 | XIII | 904586 | 904595 | 905258 | middle |  |  |  |  |  |  |  |  |  |  |  |
| YJM1208 | XIII | 905258 | 905258 | 905258 | rightend_complete | YMR321C | 99.371 | 318 | 2 | 0 | 1 | 318 | 898388 | 898071 | 4.12E-164 | 577 |
| YJM1208 | XIV | 1 |  | 768218 | leftend_complete | YNL332W | 99.902 | 1023 | 1 | 0 | 1 | 1023 | 8186 | 9208 | 0 | 1884 |
| YJM1208 | XIV | 766636 | 766645 | 768218 | middle |  |  |  |  |  |  |  |  |  |  |  |
| YJM1208 | XIV | 768218 | 768218 | 768218 | rightend_complete | YNR071C | 99.611 | 1029 | 4 | 0 | 1 | 1029 | 748170 | 747142 | 0 | 1879 |
| YJM1208 | XV | 1 | 10 | 1052488 | leftend | YOL154W | 99.467 | 750 | 4 | 0 | 1 | 750 | 32939 | 33688 | 0 | 1363 |
| YJM1208 | XV | 1049257 | 1049266 | 1052488 | middle |  |  |  |  |  |  |  |  |  |  |  |
| YJM1208 | XV | 1049759 | 1049768 | 1052488 | middle |  |  |  |  |  |  |  |  |  |  |  |
| YJM1208 | XV | 1050196 | 1050205 | 1052488 | middle |  |  |  |  |  |  |  |  |  |  |  |
| YJM1208 | XV | 1051251 | 1051260 | 1052488 | middle |  |  |  |  |  |  |  |  |  |  |  |
| YJM1208 | XV | 1052488 | 1052488 | 1052488 | rightend_complete | YOR381W | 99.345 | 2136 | 14 | 0 | 1 | 2136 | 1032009 | 1034144 | 0 | 3868 |
| YJM1208 | XVI | 1 | 10 | 903515 | leftend | YPL274W | 98.129 | 1764 | 33 | 0 | 1 | 1764 | 10825 | 12588 | 0 | 3075 |
| YJM1208 | XVI | 3070 | 3079 | 903515 | middle |  |  |  |  |  |  |  |  |  |  |  |
| YJM1208 | XVI | 901325 | 901334 | 903515 | middle |  |  |  |  |  |  |  |  |  |  |  |
| YJM1208 | XVI | 903506 | 903515 | 903515 | rightend | YPR201W | 99.918 | 1215 | 1 | 0 | 1 | 1215 | 897560 | 898774 | 0 | 2239 |
| YJM1242 | I | 1 | 10 | 184441 | leftend | YAL063C | 91.363 | 1262 | 52 | 27 | 2715 | 3969 | 173957 | 175168 | 0 | 1674 |
| YJM1242 | I | 172473 | 172482 | 184441 | middle |  |  |  |  |  |  |  |  |  |  |  |
| YJM1242 | I | 184432 | 184441 | 184441 | rightend | YAR050W | 93.193 | 1807 | 113 | 8 | 2449 | 4250 | 173057 | 174858 | 0 | 2647 |
| YJM1242 | II | 1 | 10 | 783816 | leftend | YNCB0001W | 100 | 84 | 0 | 0 | 1 | 84 | 1725 | 1808 | 4.16E-38 | 156 |
| YJM1242 | II | 782667 | 782676 | 783816 | middle |  |  |  |  |  |  |  |  |  |  |  |
| YJM1242 | II | 783807 | 783816 | 783816 | rightend | YBR297W | 92.243 | 1418 | 94 | 12 | 1 | 1407 | 774158 | 775570 | 0 | 1995 |
| YJM1242 | III | 1 | 10 | 308503 | leftend | YCL073C | 99.784 | 1848 | 4 | 0 | 1 | 1848 | 7607 | 5760 | 0 | 3391 |
| YJM1242 | III | 308494 | 308503 | 308503 | rightend | YCR101C | 99.818 | 549 | 1 | 0 | 1 | 549 | 295676 | 295128 | 0 | 1009 |
| YJM1242 | IV | 1 | 10 | 1456121 | leftend | YDL243C | 95.051 | 990 | 49 | 0 | 1 | 990 | 6538 | 5549 | 0 | 1557 |
| YJM1242 | IV | 1456112 | 1456121 | 1456121 | rightend | YDR541C | 99.42 | 1035 | 6 | 0 | 1 | 1035 | 1453273 | 1452239 | 0 | 1879 |
| YJM1242 | V | 1 | 10 | 557091 | leftend | YEL070W | 99.271 | 1509 | 11 | 0 | 1 | 1509 | 13406 | 14914 | 0 | 2726 |
| YJM1242 | V | 557091 | 557091 | 557091 | rightend_complete | YER187W | 91.745 | 424 | 33 | 2 | 4 | 426 | 548242 | 548664 | 1.6E-167 | 588 |
| YJM1242 | VI | 1 | 10 | 233737 | leftend | YFL051C | 98.451 | 452 | 7 | 0 | 1 | 452 | 1092 | 641 | 0 | 797 |
| YJM1242 | VI | 233737 | 233737 | 233737 | rightend_complete | YFR055W | 100 | 1023 | 0 | 0 | 1 | 1023 | 227771 | 228793 | 0 | 1890 |
| YJM1242 | VII | 1 |  | 1038102 | leftend_complete | YGL258W | 97.585 | 621 | 15 | 0 | 1 | 621 | 11236 | 11856 | 0 | 1064 |
| YJM1242 | VII | 1038093 | 1038102 | 1038102 | rightend | YGR287C | 99.139 | 1626 | 14 | 0 | 145 | 1770 | 1031164 | 1029539 | 0 | 2926 |
| YJM1242 | VIII | 1 | 10 | 539885 | leftend | YHL044W | 98.305 | 708 | 12 | 0 | 1 | 708 | 9082 | 9789 | 0 | 1242 |
| YJM1242 | VIII | 6983 | 6992 | 539885 | middle |  |  |  |  |  |  |  |  |  |  |  |
| YJM1242 | VIII | 7638 | 7647 | 539885 | middle |  |  |  |  |  |  |  |  |  |  |  |
| YJM1242 | VIII | 538865 | 538874 | 539885 | middle |  |  |  |  |  |  |  |  |  |  |  |
| YJM1242 | VIII | 539678 | 539687 | 539885 | middle |  |  |  |  |  |  |  |  |  |  |  |
| YJM1242 | VIII | 539885 | 539885 | 539885 | rightend_complete | YHR211W | 98.465 | 977 | 15 | 0 | 2252 | 3228 | 516482 | 517458 | 0 | 1722 |
| YJM1242 | IX | 1 | 10 | 402940 | leftend | YIL169C | 93.53 | 1252 | 79 | 2 | 562 | 1812 | 2182 | 932 | 0 | 1862 |
| YJM1242 | IX | 402931 | 402940 | 402940 | rightend | YIR039C | 99.009 | 1614 | 16 | 0 | 1 | 1614 | 401276 | 399663 | 0 | 2892 |
| YJM1242 | X | 1 | 10 | 728113 | leftend | YJL213W | 99.699 | 996 | 3 | 0 | 1 | 996 | 26311 | 27306 | 0 | 1823 |
| YJM1242 | X | 715061 | 715070 | 728113 | middle |  |  |  |  |  |  |  |  |  |  |  |
| YJM1242 | X | 726449 | 726458 | 728113 | middle |  |  |  |  |  |  |  |  |  |  |  |
| YJM1242 | X | 728104 | 728113 | 728113 | rightend | YJR152W | 99.755 | 1632 | 3 | 1 | 1 | 1632 | 695335 | 696965 | 0 | 2990 |
| YJM1242 | XI | 1 | 10 | 660026 | leftend | YKL222C | 99.386 | 2118 | 13 | 0 | 1 | 2118 | 3266 | 1149 | 0 | 3840 |
| YJM1242 | XI | 659404 | 659413 | 660026 | middle |  |  |  |  |  |  |  |  |  |  |  |
| YJM1242 | XI | 660026 | 660026 | 660026 | rightend_complete | YKR101W | 99.847 | 1965 | 3 | 0 | 1 | 1965 | 637114 | 639078 | 0 | 3613 |
| YJM1242 | XII | 1 | 10 | 1602038 | leftend | YLL062C | 100 | 975 | 0 | 0 | 1 | 975 | 1968 | 994 | 0 | 1801 |
| YJM1242 | XII | 1602029 | 1602038 | 1602038 | rightend | YLR460C | 94.336 | 1130 | 62 | 2 | 3 | 1131 | 1600389 | 1599261 | 0 | 1731 |
| YJM1242 | XIII | 1 | 10 | 902131 | leftend | YML131W | 99.818 | 1098 | 2 | 0 | 1 | 1098 | 4978 | 6075 | 0 | 2017 |
| YJM1242 | XIII | 1938 | 1947 | 902131 | middle |  |  |  |  |  |  |  |  |  |  |  |
| YJM1242 | XIII | 2671 | 2680 | 902131 | middle |  |  |  |  |  |  |  |  |  |  |  |
| YJM1242 | XIII | 896935 | 896944 | 902131 | middle |  |  |  |  |  |  |  |  |  |  |  |
| YJM1242 | XIII | 902122 | 902131 | 902131 | rightend | YMR321C | 99.361 | 313 | 2 | 0 | 6 | 318 | 889914 | 889602 | 2.48E-161 | 568 |
| YJM1242 | XIV | 1 | 10 | 759231 | leftend | YNL332W | 98.631 | 1023 | 14 | 0 | 1 | 1023 | 8596 | 9618 | 0 | 1812 |
| YJM1242 | XIV | 294 | 303 | 759231 | middle |  |  |  |  |  |  |  |  |  |  |  |
| YJM1242 | XIV | 1396 | 1405 | 759231 | middle |  |  |  |  |  |  |  |  |  |  |  |
| YJM1242 | XIV | 1633 | 1642 | 759231 | middle |  |  |  |  |  |  |  |  |  |  |  |
| YJM1242 | XIV | 2498 | 2507 | 759231 | middle |  |  |  |  |  |  |  |  |  |  |  |
| YJM1242 | XIV | 3077 | 3086 | 759231 | middle |  |  |  |  |  |  |  |  |  |  |  |
| YJM1242 | XIV | 759222 | 759231 | 759231 | rightend | YNR071C | 100 | 1029 | 0 | 0 | 1 | 1029 | 747038 | 746010 | 0 | 1901 |
| YJM1242 | XV | 1 | 10 | 1034307 | leftend | YOL154W | 99.467 | 750 | 4 | 0 | 1 | 750 | 9037 | 9786 | 0 | 1363 |
| YJM1242 | XV | 1032999 | 1033008 | 1034307 | middle |  |  |  |  |  |  |  |  |  |  |  |
| YJM1242 | XV | 1034298 | 1034307 | 1034307 | rightend | YOR381W | 99.532 | 2136 | 2 | 1 | 1 | 2136 | 1005752 | 1007879 | 0 | 3882 |
| YJM1242 | XVI | 1 | 10 | 895313 | leftend | YPL274W | 99.943 | 1764 | 1 | 0 | 1 | 1764 | 5759 | 7522 | 0 | 3253 |
| YJM1242 | XVI | 1037 | 1046 | 895313 | middle |  |  |  |  |  |  |  |  |  |  |  |
| YJM1242 | XVI | 1544 | 1553 | 895313 | middle |  |  |  |  |  |  |  |  |  |  |  |
| YJM1242 | XVI | 894848 | 894857 | 895313 | middle |  |  |  |  |  |  |  |  |  |  |  |
| YJM1242 | XVI | 895304 | 895313 | 895313 | rightend | YPR201W | 100 | 1215 | 0 | 0 | 1 | 1215 | 891837 | 893051 | 0 | 2244 |
| YJM1244 | I | 1 | 10 | 194316 | leftend | YAL063C | 95.02 | 1004 | 23 | 11 | 2966 | 3969 | 172759 | 173735 | 0 | 1552 |
| YJM1244 | I | 194307 | 194316 | 194316 | rightend | YAR050W | 81.713 | 1635 | 292 | 5 | 1 | 1630 | 170580 | 172212 | 0 | 1356 |
| YJM1244 | II | 1 | 10 | 784789 | leftend | YNCB0001W | 100 | 84 | 0 | 0 | 1 | 84 | 1647 | 1730 | 4.16E-38 | 156 |
| YJM1244 | II | 784165 | 784174 | 784789 | middle |  |  |  |  |  |  |  |  |  |  |  |
| YJM1244 | II | 784544 | 784553 | 784789 | middle |  |  |  |  |  |  |  |  |  |  |  |
| YJM1244 | II | 784789 | 784789 | 784789 | rightend_complete | YBR297W | 99.574 | 1407 | 4 | 1 | 1 | 1407 | 773812 | 775216 | 0 | 2564 |
| YJM1244 | III | 1 |  | 309207 | leftend_complete | YCL073C | 99.729 | 1848 | 5 | 0 | 1 | 1848 | 7982 | 6135 | 0 | 3386 |
| YJM1244 | III | 309198 | 309207 | 309207 | rightend | YCR101C | 99.636 | 549 | 2 | 0 | 1 | 549 | 296376 | 295828 | 0 | 1003 |
| YJM1244 | IV | 1 | 10 | 1487556 | leftend | YDL243C | 95.152 | 990 | 48 | 0 | 1 | 990 | 16922 | 15933 | 0 | 1563 |
| YJM1244 | IV | 1487547 | 1487556 | 1487556 | rightend | YDR541C | 99.42 | 1035 | 6 | 0 | 1 | 1035 | 1484921 | 1483887 | 0 | 1879 |
| YJM1244 | V | 1 | 10 | 550102 | leftend | YEL070W | 99.139 | 1509 | 13 | 0 | 1 | 1509 | 12824 | 14332 | 0 | 2715 |
| YJM1244 | V | 550093 | 550102 | 550102 | rightend | YER187W | 91.981 | 424 | 32 | 2 | 4 | 426 | 547097 | 547519 | 3.44E-169 | 593 |
| YJM1244 | VI | 1 | 10 | 241094 | leftend | YFL051C | 98.451 | 452 | 7 | 0 | 1 | 452 | 7962 | 7511 | 0 | 797 |
| YJM1244 | VI | 6199 | 6208 | 241094 | middle |  |  |  |  |  |  |  |  |  |  |  |
| YJM1244 | VI | 241085 | 241094 | 241094 | rightend | YFR055W | 99.902 | 1023 | 1 | 0 | 1 | 1023 | 234639 | 235661 | 0 | 1884 |
| YJM1244 | VII | 1 | 10 | 1057043 | leftend | YGL258W | 97.585 | 621 | 15 | 0 | 1 | 621 | 10787 | 11407 | 0 | 1064 |
| YJM1244 | VII | 6226 | 6235 | 1057043 | middle |  |  |  |  |  |  |  |  |  |  |  |
| YJM1244 | VII | 1057034 | 1057043 | 1057043 | rightend | YGR287C | 98.701 | 1770 | 23 | 0 | 1 | 1770 | 1031472 | 1029703 | 0 | 3142 |
| YJM1244 | VIII | 1 | 10 | 524609 | leftend | YHL044W | 98.446 | 708 | 11 | 0 | 1 | 708 | 8609 | 9316 | 0 | 1247 |
| YJM1244 | VIII | 6775 | 6784 | 524609 | middle |  |  |  |  |  |  |  |  |  |  |  |
| YJM1244 | VIII | 7309 | 7318 | 524609 | middle |  |  |  |  |  |  |  |  |  |  |  |
| YJM1244 | VIII | 524600 | 524609 | 524609 | rightend | YHR211W | 82.535 | 939 | 153 | 7 | 1 | 932 | 514142 | 515076 | 0 | 815 |
| YJM1244 | IX | 1 | 10 | 403880 | leftend | YIL169C | 85.483 | 1977 | 238 | 26 | 444 | 2382 | 3557 | 1592 | 0 | 2015 |
| YJM1244 | IX | 1577 | 1586 | 403880 | middle |  |  |  |  |  |  |  |  |  |  |  |
| YJM1244 | IX | 403476 | 403485 | 403880 | middle |  |  |  |  |  |  |  |  |  |  |  |
| YJM1244 | IX | 403880 | 403880 | 403880 | rightend_complete | YIR039C | 99.009 | 1614 | 16 | 0 | 1 | 1614 | 402016 | 400403 | 0 | 2892 |
| YJM1244 | X | 1 | 10 | 728283 | leftend | YJL213W | 99.699 | 996 | 3 | 0 | 1 | 996 | 26694 | 27689 | 0 | 1823 |
| YJM1244 | X | 881 | 890 | 728283 | middle |  |  |  |  |  |  |  |  |  |  |  |
| YJM1244 | X | 716625 | 716634 | 728283 | middle |  |  |  |  |  |  |  |  |  |  |  |
| YJM1244 | X | 727518 | 727527 | 728283 | middle |  |  |  |  |  |  |  |  |  |  |  |
| YJM1244 | X | 727813 | 727822 | 728283 | middle |  |  |  |  |  |  |  |  |  |  |  |
| YJM1244 | X | 728274 | 728283 | 728283 | rightend | YJR152W | 99.755 | 1632 | 3 | 1 | 1 | 1632 | 696679 | 698309 | 0 | 2990 |
| YJM1244 | XI | 1 | 10 | 659862 | leftend | YKL222C | 99.433 | 2118 | 12 | 0 | 1 | 2118 | 3284 | 1167 | 0 | 3845 |
| YJM1244 | XI | 659853 | 659862 | 659862 | rightend | YKR101W | 99.796 | 1965 | 4 | 0 | 1 | 1965 | 637191 | 639155 | 0 | 3607 |
| YJM1244 | XII | 1 | 10 | 2121296 | leftend | YLL062C | 100 | 975 | 0 | 0 | 1 | 975 | 5550 | 4576 | 0 | 1801 |
| YJM1244 | XII | 1544 | 1553 | 2121296 | middle |  |  |  |  |  |  |  |  |  |  |  |
| YJM1244 | XII | 2121287 | 2121296 | 2121296 | rightend | YLR460C | 94.248 | 1130 | 63 | 2 | 3 | 1131 | 2119543 | 2118415 | 0 | 1725 |
| YJM1244 | XIII | 1 | 10 | 913653 | leftend | YML131W | 100 | 1098 | 0 | 0 | 1 | 1098 | 5046 | 6143 | 0 | 2028 |
| YJM1244 | XIII | 665 | 674 | 913653 | middle |  |  |  |  |  |  |  |  |  |  |  |
| YJM1244 | XIII | 2718 | 2727 | 913653 | middle |  |  |  |  |  |  |  |  |  |  |  |
| YJM1244 | XIII | 908033 | 908042 | 913653 | middle |  |  |  |  |  |  |  |  |  |  |  |
| YJM1244 | XIII | 913644 | 913653 | 913653 | rightend | YMR321C | 99.361 | 313 | 2 | 0 | 6 | 318 | 890508 | 890196 | 2.48E-161 | 568 |
| YJM1244 | XIV | 1 |  | 749299 | leftend_complete | YNL332W | 98.586 | 495 | 7 | 0 | 529 | 1023 | 439 | 933 | 0 | 876 |
| YJM1244 | XIV | 205 | 214 | 749299 | middle |  |  |  |  |  |  |  |  |  |  |  |
| YJM1244 | XIV | 429 | 438 | 749299 | middle |  |  |  |  |  |  |  |  |  |  |  |
| YJM1244 | XIV | 749290 | 749299 | 749299 | rightend | YNR071C | 100 | 1029 | 0 | 0 | 1 | 1029 | 738773 | 737745 | 0 | 1901 |
| YJM1244 | XV | 1 | 10 | 1049993 | leftend | YOL154W | 99.467 | 750 | 4 | 0 | 1 | 750 | 23162 | 23911 | 0 | 1363 |
| YJM1244 | XV | 1047306 | 1047315 | 1049993 | middle |  |  |  |  |  |  |  |  |  |  |  |
| YJM1244 | XV | 1048548 | 1048557 | 1049993 | middle |  |  |  |  |  |  |  |  |  |  |  |
| YJM1244 | XV | 1048934 | 1048943 | 1049993 | middle |  |  |  |  |  |  |  |  |  |  |  |
| YJM1244 | XV | 1049984 | 1049993 | 1049993 | rightend | YOR381W | 99.86 | 2136 | 3 | 0 | 1 | 2136 | 1020203 | 1022338 | 0 | 3928 |
| YJM1244 | XVI | 1 | 10 | 919857 | leftend | YPL274W | 99.943 | 1764 | 1 | 0 | 1 | 1764 | 22644 | 24407 | 0 | 3253 |
| YJM1244 | XVI | 1249 | 1258 | 919857 | middle |  |  |  |  |  |  |  |  |  |  |  |
| YJM1244 | XVI | 3231 | 3240 | 919857 | middle |  |  |  |  |  |  |  |  |  |  |  |
| YJM1244 | XVI | 3697 | 3706 | 919857 | middle |  |  |  |  |  |  |  |  |  |  |  |
| YJM1244 | XVI | 6220 | 6229 | 919857 | middle |  |  |  |  |  |  |  |  |  |  |  |
| YJM1244 | XVI | 6434 | 6443 | 919857 | middle |  |  |  |  |  |  |  |  |  |  |  |
| YJM1244 | XVI | 7914 | 7923 | 919857 | middle |  |  |  |  |  |  |  |  |  |  |  |
| YJM1244 | XVI | 914510 | 914519 | 919857 | middle |  |  |  |  |  |  |  |  |  |  |  |
| YJM1244 | XVI | 916559 | 916568 | 919857 | middle |  |  |  |  |  |  |  |  |  |  |  |
| YJM1244 | XVI | 917544 | 917553 | 919857 | middle |  |  |  |  |  |  |  |  |  |  |  |
| YJM1244 | XVI | 919857 | 919857 | 919857 | rightend_complete | YPR201W | 100 | 1215 | 0 | 0 | 1 | 1215 | 911776 | 912990 | 0 | 2244 |
| YJM1248 | I | 1 |  | 200251 | leftend_complete | YAL063C | 94.244 | 1859 | 103 | 4 | 2106 | 3962 | 15342 | 13486 | 0 | 2837 |
| YJM1248 | I | 701 | 710 | 200251 | middle |  |  |  |  |  |  |  |  |  |  |  |
| YJM1248 | I | 15343 | 15352 | 200251 | middle |  |  |  |  |  |  |  |  |  |  |  |
| YJM1248 | I | 15771 | 15780 | 200251 | middle |  |  |  |  |  |  |  |  |  |  |  |
| YJM1248 | I | 188105 | 188114 | 200251 | middle |  |  |  |  |  |  |  |  |  |  |  |
| YJM1248 | I | 200242 | 200251 | 200251 | rightend | YAR050W | 91.745 | 957 | 49 | 11 | 3349 | 4282 | 14733 | 13784 | 0 | 1303 |
| YJM1248 | II | 1 | 10 | 780938 | leftend | YNCB0001W | 100 | 84 | 0 | 0 | 1 | 84 | 2490 | 2573 | 4.16E-38 | 156 |
| YJM1248 | II | 315 | 324 | 780938 | middle |  |  |  |  |  |  |  |  |  |  |  |
| YJM1248 | II | 1046 | 1055 | 780938 | middle |  |  |  |  |  |  |  |  |  |  |  |
| YJM1248 | II | 2025 | 2034 | 780938 | middle |  |  |  |  |  |  |  |  |  |  |  |
| YJM1248 | II | 780929 | 780938 | 780938 | rightend | YBR297W | 97.925 | 1205 | 25 | 0 | 1 | 1205 | 776815 | 778019 | 0 | 2087 |
| YJM1248 | III | 1 | 10 | 307684 | leftend | YCL073C | 98.81 | 1848 | 21 | 1 | 1 | 1848 | 3927 | 2081 | 0 | 3290 |
| YJM1248 | III | 307675 | 307684 | 307684 | rightend | YCR101C | 98.364 | 550 | 8 | 1 | 1 | 549 | 294578 | 294029 | 0 | 965 |
| YJM1248 | IV | 1 | 10 | 1447577 | leftend | YDL243C | 91.111 | 990 | 87 | 1 | 1 | 990 | 2519 | 1531 | 0 | 1339 |
| YJM1248 | IV | 1447568 | 1447577 | 1447577 | rightend | YDR541C | 99.614 | 1035 | 4 | 0 | 1 | 1035 | 1444518 | 1443484 | 0 | 1890 |
| YJM1248 | V | 1 | 10 | 552061 | leftend | YEL070W | 99.205 | 1509 | 12 | 0 | 1 | 1509 | 13006 | 14514 | 0 | 2721 |
| YJM1248 | V | 552052 | 552061 | 552061 | rightend | YER187W | 97.183 | 426 | 12 | 0 | 1 | 426 | 548392 | 548817 | 0 | 721 |
| YJM1248 | VI | 1 | 10 | 274979 | leftend | YFL051C | 98.451 | 452 | 7 | 0 | 1 | 452 | 39397 | 38946 | 0 | 797 |
| YJM1248 | VI | 274970 | 274979 | 274979 | rightend | YFR055W | 99.217 | 1022 | 8 | 0 | 1 | 1022 | 268771 | 269792 | 0 | 1844 |
| YJM1248 | VII | 1 | 10 | 1042722 | leftend | YGL258W | 99.034 | 621 | 6 | 0 | 1 | 621 | 3925 | 4545 | 0 | 1114 |
| YJM1248 | VII | 1039672 | 1039681 | 1042722 | middle |  |  |  |  |  |  |  |  |  |  |  |
| YJM1248 | VII | 1041790 | 1041799 | 1042722 | middle |  |  |  |  |  |  |  |  |  |  |  |
| YJM1248 | VII | 1042713 | 1042722 | 1042722 | rightend | YGR287C | 96.16 | 1771 | 66 | 2 | 1 | 1770 | 1030050 | 1028281 | 0 | 2892 |
| YJM1248 | VIII | 1 | 10 | 515493 | leftend | YHL044W | 98.023 | 708 | 14 | 0 | 1 | 708 | 1463 | 2170 | 0 | 1230 |
| YJM1248 | VIII | 506467 | 506476 | 515493 | middle |  |  |  |  |  |  |  |  |  |  |  |
| YJM1248 | VIII | 515484 | 515493 | 515493 | rightend | YHR211W | 84.032 | 1002 | 149 | 7 | 1 | 995 | 504258 | 505255 | 0 | 953 |
| YJM1248 | IX | 1 | 10 | 423431 | leftend | YIL169C | 84.703 | 1752 | 224 | 21 | 753 | 2467 | 1755 | 11 | 0 | 1711 |
| YJM1248 | IX | 423422 | 423431 | 423431 | rightend | YIR039C | 99.69 | 1614 | 5 | 0 | 1 | 1614 | 415266 | 413653 | 0 | 2953 |
| YJM1248 | X | 1 | 10 | 727248 | leftend | YJL213W | 99.598 | 996 | 4 | 0 | 1 | 996 | 30096 | 31091 | 0 | 1818 |
| YJM1248 | X | 1885 | 1894 | 727248 | middle |  |  |  |  |  |  |  |  |  |  |  |
| YJM1248 | X | 3576 | 3585 | 727248 | middle |  |  |  |  |  |  |  |  |  |  |  |
| YJM1248 | X | 3828 | 3837 | 727248 | middle |  |  |  |  |  |  |  |  |  |  |  |
| YJM1248 | X | 4815 | 4824 | 727248 | middle |  |  |  |  |  |  |  |  |  |  |  |
| YJM1248 | X | 5708 | 5717 | 727248 | middle |  |  |  |  |  |  |  |  |  |  |  |
| YJM1248 | X | 726394 | 726403 | 727248 | middle |  |  |  |  |  |  |  |  |  |  |  |
| YJM1248 | X | 727239 | 727248 | 727248 | rightend | YJR152W | 99.694 | 1632 | 5 | 0 | 1 | 1632 | 702365 | 703996 | 0 | 2987 |
| YJM1248 | XI | 1 | 10 | 663981 | leftend | YKL222C | 99.622 | 2118 | 8 | 0 | 1 | 2118 | 3301 | 1184 | 0 | 3868 |
| YJM1248 | XI | 663972 | 663981 | 663981 | rightend | YKR101W | 99.644 | 1965 | 7 | 0 | 1 | 1965 | 639548 | 641512 | 0 | 3591 |
| YJM1248 | XII | 1 | 10 | 1939382 | leftend | YLL062C | 99.282 | 975 | 7 | 0 | 1 | 975 | 7906 | 6932 | 0 | 1762 |
| YJM1248 | XII | 1929059 | 1929068 | 1939382 | middle |  |  |  |  |  |  |  |  |  |  |  |
| YJM1248 | XII | 1939373 | 1939382 | 1939382 | rightend | YLR460C | 95.933 | 1131 | 46 | 0 | 1 | 1131 | 1926952 | 1925822 | 0 | 1834 |
| YJM1248 | XIII | 1 | 10 | 901407 | leftend | YML131W | 99.545 | 1098 | 5 | 0 | 1 | 1098 | 3401 | 4498 | 0 | 2001 |
| YJM1248 | XIII | 898251 | 898260 | 901407 | middle |  |  |  |  |  |  |  |  |  |  |  |
| YJM1248 | XIII | 901398 | 901407 | 901407 | rightend | YMR321C | 99.361 | 313 | 2 | 0 | 6 | 318 | 889760 | 889448 | 2.48E-161 | 568 |
| YJM1248 | XIV | 1 | 10 | 759732 | leftend | YNL332W | 98.534 | 1023 | 15 | 0 | 1 | 1023 | 3096 | 4118 | 0 | 1807 |
| YJM1248 | XIV | 1111 | 1120 | 759732 | middle |  |  |  |  |  |  |  |  |  |  |  |
| YJM1248 | XIV | 2939 | 2948 | 759732 | middle |  |  |  |  |  |  |  |  |  |  |  |
| YJM1248 | XIV | 759723 | 759732 | 759732 | rightend | YNR071C | 99.514 | 1029 | 5 | 0 | 1 | 1029 | 749253 | 748225 | 0 | 1873 |
| YJM1248 | XV | 1 | 10 | 1033043 | leftend | YOL154W | 99.467 | 750 | 4 | 0 | 1 | 750 | 5273 | 6022 | 0 | 1363 |
| YJM1248 | XV | 1031620 | 1031629 | 1033043 | middle |  |  |  |  |  |  |  |  |  |  |  |
| YJM1248 | XV | 1032339 | 1032348 | 1033043 | middle |  |  |  |  |  |  |  |  |  |  |  |
| YJM1248 | XV | 1033034 | 1033043 | 1033043 | rightend | YOR381W | 98.876 | 2136 | 24 | 0 | 1 | 2136 | 1004603 | 1006738 | 0 | 3812 |
| YJM1248 | XVI | 1 | 10 | 912323 | leftend | YPL274W | 98.356 | 1764 | 29 | 0 | 1 | 1764 | 16176 | 17939 | 0 | 3097 |
| YJM1248 | XVI | 1452 | 1461 | 912323 | middle |  |  |  |  |  |  |  |  |  |  |  |
| YJM1248 | XVI | 912323 | 912323 | 912323 | rightend_complete | YPR201W | 99.588 | 1215 | 5 | 0 | 1 | 1215 | 908756 | 909970 | 0 | 2217 |
| YJM1250 | I | 1 | 10 | 191544 | leftend | YAL063C | 84.075 | 1595 | 246 | 8 | 40 | 1630 | 189090 | 190680 | 0 | 1531 |
| YJM1250 | I | 10958 | 10967 | 191544 | middle |  |  |  |  |  |  |  |  |  |  |  |
| YJM1250 | I | 191535 | 191544 | 191544 | rightend | YAR050W | 82.659 | 1632 | 279 | 4 | 1 | 1630 | 189051 | 190680 | 0 | 1443 |
| YJM1250 | II | 1 | 10 | 784997 | leftend | YNCB0001W | 100 | 84 | 0 | 0 | 1 | 84 | 2319 | 2402 | 4.16E-38 | 156 |
| YJM1250 | II | 1966 | 1975 | 784997 | middle |  |  |  |  |  |  |  |  |  |  |  |
| YJM1250 | II | 784988 | 784997 | 784997 | rightend | YBR297W | 99.005 | 1407 | 14 | 0 | 1 | 1407 | 774911 | 776317 | 0 | 2521 |
| YJM1250 | III | 1 | 10 | 309489 | leftend | YCL073C | 99.838 | 1848 | 3 | 0 | 1 | 1848 | 7489 | 5642 | 0 | 3397 |
| YJM1250 | III | 300680 | 300689 | 309489 | middle |  |  |  |  |  |  |  |  |  |  |  |
| YJM1250 | III | 309489 | 309489 | 309489 | rightend_complete | YCR101C | 99.818 | 549 | 1 | 0 | 1 | 549 | 296118 | 295570 | 0 | 1009 |
| YJM1250 | IV | 1 |  | 1456085 | leftend_complete | YDL243C | 95.051 | 990 | 49 | 0 | 1 | 990 | 8969 | 7980 | 0 | 1557 |
| YJM1250 | IV | 420 | 429 | 1456085 | middle |  |  |  |  |  |  |  |  |  |  |  |
| YJM1250 | IV | 421 | 430 | 1456085 | middle |  |  |  |  |  |  |  |  |  |  |  |
| YJM1250 | IV | 1456076 | 1456085 | 1456085 | rightend | YDR541C | 99.324 | 1035 | 7 | 0 | 1 | 1035 | 1453427 | 1452393 | 0 | 1873 |
| YJM1250 | V | 1 | 10 | 550400 | leftend | YEL070W | 98.211 | 1509 | 27 | 0 | 1 | 1509 | 12814 | 14322 | 0 | 2638 |
| YJM1250 | V | 548793 | 548802 | 550400 | middle |  |  |  |  |  |  |  |  |  |  |  |
| YJM1250 | V | 550391 | 550400 | 550400 | rightend | YER187W | 91.981 | 424 | 32 | 2 | 4 | 426 | 546230 | 546652 | 3.44E-169 | 593 |
| YJM1250 | VI | 1 | 10 | 233940 | leftend | YFL051C | 98.451 | 452 | 7 | 0 | 1 | 452 | 1193 | 742 | 0 | 797 |
| YJM1250 | VI | 233931 | 233940 | 233940 | rightend | YFR055W | 99.902 | 1023 | 1 | 0 | 1 | 1023 | 227590 | 228612 | 0 | 1884 |
| YJM1250 | VII | 1 | 10 | 1035658 | leftend | YGL258W | 94.686 | 621 | 33 | 0 | 1 | 621 | 10615 | 11235 | 0 | 965 |
| YJM1250 | VII | 1035649 | 1035658 | 1035658 | rightend | YGR287C | 87.281 | 1769 | 223 | 2 | 1 | 1768 | 1030682 | 1028915 | 0 | 2019 |
| YJM1250 | VIII | 1 | 10 | 534326 | leftend | YHL044W | 98.446 | 708 | 11 | 0 | 1 | 708 | 2265 | 2972 | 0 | 1247 |
| YJM1250 | VIII | 910 | 919 | 534326 | middle |  |  |  |  |  |  |  |  |  |  |  |
| YJM1250 | VIII | 534317 | 534326 | 534326 | rightend | YHR211W | 89.547 | 2296 | 198 | 18 | 966 | 3228 | 510199 | 512485 | 0 | 2872 |
| YJM1250 | IX | 1 | 10 | 420984 | leftend | YIL169C | 97.92 | 1442 | 24 | 2 | 478 | 1919 | 20004 | 18569 | 0 | 2492 |
| YJM1250 | IX | 1015 | 1024 | 420984 | middle |  |  |  |  |  |  |  |  |  |  |  |
| YJM1250 | IX | 5132 | 5141 | 420984 | middle |  |  |  |  |  |  |  |  |  |  |  |
| YJM1250 | IX | 420975 | 420984 | 420984 | rightend | YIR039C | 99.628 | 1614 | 6 | 0 | 1 | 1614 | 419336 | 417723 | 0 | 2948 |
| YJM1250 | X | 1 |  | 736380 | leftend_complete | YJL213W | 88.797 | 482 | 54 | 0 | 515 | 996 | 37059 | 37540 | 3.89E-168 | 592 |
| YJM1250 | X | 2311 | 2320 | 736380 | middle |  |  |  |  |  |  |  |  |  |  |  |
| YJM1250 | X | 3730 | 3739 | 736380 | middle |  |  |  |  |  |  |  |  |  |  |  |
| YJM1250 | X | 733798 | 733807 | 736380 | middle |  |  |  |  |  |  |  |  |  |  |  |
| YJM1250 | X | 735211 | 735220 | 736380 | middle |  |  |  |  |  |  |  |  |  |  |  |
| YJM1250 | X | 735446 | 735455 | 736380 | middle |  |  |  |  |  |  |  |  |  |  |  |
| YJM1250 | X | 736380 | 736380 | 736380 | rightend_complete | YJR152W | 99.816 | 1632 | 3 | 0 | 1 | 1632 | 708973 | 710604 | 0 | 2998 |
| YJM1250 | XI | 1 | 10 | 673546 | leftend | YKL222C | 99.481 | 2118 | 11 | 0 | 1 | 2118 | 4229 | 2112 | 0 | 3851 |
| YJM1250 | XI | 936 | 945 | 673546 | middle |  |  |  |  |  |  |  |  |  |  |  |
| YJM1250 | XI | 673537 | 673546 | 673546 | rightend | YKR101W | 99.847 | 1965 | 3 | 0 | 1 | 1965 | 645397 | 647361 | 0 | 3613 |
| YJM1250 | XII | 1 | 10 | 1650453 | leftend | YLL062C | 100 | 975 | 0 | 0 | 1 | 975 | 8028 | 7054 | 0 | 1801 |
| YJM1250 | XII | 2883 | 2892 | 1650453 | middle |  |  |  |  |  |  |  |  |  |  |  |
| YJM1250 | XII | 3991 | 4000 | 1650453 | middle |  |  |  |  |  |  |  |  |  |  |  |
| YJM1250 | XII | 1650444 | 1650453 | 1650453 | rightend | YLR460C | 94.159 | 1130 | 64 | 2 | 3 | 1131 | 1648152 | 1647024 | 0 | 1720 |
| YJM1250 | XIII | 1 | 10 | 927585 | leftend | YML131W | 100 | 1098 | 0 | 0 | 1 | 1098 | 19562 | 20659 | 0 | 2028 |
| YJM1250 | XIII | 927585 | 927585 | 927585 | rightend_complete | YMR321C | 99.361 | 313 | 2 | 0 | 6 | 318 | 904231 | 903919 | 2.48E-161 | 568 |
| YJM1250 | XIV | 1 | 10 | 758913 | leftend | YNL332W | 98.436 | 1023 | 16 | 0 | 1 | 1023 | 6806 | 7828 | 0 | 1801 |
| YJM1250 | XIV | 1161 | 1170 | 758913 | middle |  |  |  |  |  |  |  |  |  |  |  |
| YJM1250 | XIV | 756609 | 756618 | 758913 | middle |  |  |  |  |  |  |  |  |  |  |  |
| YJM1250 | XIV | 758904 | 758913 | 758913 | rightend | YNR071C | 100 | 1029 | 0 | 0 | 1 | 1029 | 746256 | 745228 | 0 | 1901 |
| YJM1250 | XV | 1 | 10 | 1030411 | leftend | YOL154W | 99.467 | 750 | 4 | 0 | 1 | 750 | 7260 | 8009 | 0 | 1363 |
| YJM1250 | XV | 1024463 | 1024472 | 1030411 | middle |  |  |  |  |  |  |  |  |  |  |  |
| YJM1250 | XV | 1030402 | 1030411 | 1030411 | rightend | YOR381W | 99.906 | 2136 | 2 | 0 | 1 | 2136 | 1006106 | 1008241 | 0 | 3934 |
| YJM1250 | XVI | 1 | 10 | 917777 | leftend | YPL274W | 99.887 | 1764 | 2 | 0 | 1 | 1764 | 19956 | 21719 | 0 | 3247 |
| YJM1250 | XVI | 875 | 884 | 917777 | middle |  |  |  |  |  |  |  |  |  |  |  |
| YJM1250 | XVI | 4813 | 4822 | 917777 | middle |  |  |  |  |  |  |  |  |  |  |  |
| YJM1250 | XVI | 5028 | 5037 | 917777 | middle |  |  |  |  |  |  |  |  |  |  |  |
| YJM1250 | XVI | 915535 | 915544 | 917777 | middle |  |  |  |  |  |  |  |  |  |  |  |
| YJM1250 | XVI | 916582 | 916591 | 917777 | middle |  |  |  |  |  |  |  |  |  |  |  |
| YJM1250 | XVI | 917768 | 917777 | 917777 | rightend | YPR201W | 100 | 1215 | 0 | 0 | 1 | 1215 | 912679 | 913893 | 0 | 2244 |
| YJM1252 | I | 1 | 10 | 199003 | leftend | YAL063C | 84.075 | 1595 | 246 | 8 | 40 | 1630 | 181559 | 183149 | 0 | 1531 |
| YJM1252 | I | 198994 | 199003 | 199003 | rightend | YAR050W | 82.659 | 1632 | 279 | 4 | 1 | 1630 | 181520 | 183149 | 0 | 1443 |
| YJM1252 | II | 1 | 10 | 782129 | leftend | YNCB0001W | 100 | 84 | 0 | 0 | 1 | 84 | 2817 | 2900 | 4.16E-38 | 156 |
| YJM1252 | II | 1162 | 1171 | 782129 | middle |  |  |  |  |  |  |  |  |  |  |  |
| YJM1252 | II | 782120 | 782129 | 782129 | rightend | YBR297W | 98.934 | 1407 | 15 | 0 | 1 | 1407 | 771821 | 773227 | 0 | 2516 |
| YJM1252 | III | 1 |  | 320060 | leftend_complete | YCL073C | 99.729 | 1848 | 5 | 0 | 1 | 1848 | 7693 | 5846 | 0 | 3386 |
| YJM1252 | III | 235 | 244 | 320060 | middle |  |  |  |  |  |  |  |  |  |  |  |
| YJM1252 | III | 320051 | 320060 | 320060 | rightend | YCR101C | 78.136 | 558 | 105 | 14 | 1 | 549 | 298938 | 298389 | 1.29E-92 | 339 |
| YJM1252 | IV | 1 |  | 1457448 | leftend_complete | YDL243C | 83.282 | 981 | 158 | 6 | 1 | 978 | 12009 | 11032 | 0 | 898 |
| YJM1252 | IV | 493 | 502 | 1457448 | middle |  |  |  |  |  |  |  |  |  |  |  |
| YJM1252 | IV | 1457439 | 1457448 | 1457448 | rightend | YDR541C | 86.293 | 1036 | 137 | 5 | 1 | 1033 | 1455859 | 1454826 | 0 | 1122 |
| YJM1252 | V | 1 | 10 | 554646 | leftend | YEL070W | 90.722 | 1509 | 140 | 0 | 1 | 1509 | 15100 | 16608 | 0 | 2012 |
| YJM1252 | V | 554130 | 554139 | 554646 | middle |  |  |  |  |  |  |  |  |  |  |  |
| YJM1252 | V | 554646 | 554646 | 554646 | rightend_complete | YER187W | 99.296 | 426 | 3 | 0 | 1 | 426 | 550786 | 551211 | 0 | 771 |
| YJM1252 | VI | 1 | 10 | 237749 | leftend | YFL051C | 98.451 | 452 | 7 | 0 | 1 | 452 | 5227 | 4776 | 0 | 797 |
| YJM1252 | VI | 237740 | 237749 | 237749 | rightend | YFR055W | 90.775 | 1019 | 94 | 0 | 1 | 1019 | 233476 | 234494 | 0 | 1362 |
| YJM1252 | VII | 1 | 10 | 1031254 | leftend | YGL258W | 97.585 | 621 | 15 | 0 | 1 | 621 | 4115 | 4735 | 0 | 1064 |
| YJM1252 | VII | 1031245 | 1031254 | 1031254 | rightend | YGR287C | 99.2 | 1626 | 13 | 0 | 145 | 1770 | 1023901 | 1022276 | 0 | 2931 |
| YJM1252 | VIII | 1 | 10 | 520019 | leftend | YHL044W | 98.446 | 708 | 11 | 0 | 1 | 708 | 2281 | 2988 | 0 | 1247 |
| YJM1252 | VIII | 263 | 272 | 520019 | middle |  |  |  |  |  |  |  |  |  |  |  |
| YJM1252 | VIII | 719 | 728 | 520019 | middle |  |  |  |  |  |  |  |  |  |  |  |
| YJM1252 | VIII | 518568 | 518577 | 520019 | middle |  |  |  |  |  |  |  |  |  |  |  |
| YJM1252 | VIII | 519301 | 519310 | 520019 | middle |  |  |  |  |  |  |  |  |  |  |  |
| YJM1252 | VIII | 520019 | 520019 | 520019 | rightend_complete | YHR211W | 82.428 | 939 | 154 | 7 | 1 | 932 | 506519 | 507453 | 0 | 809 |
| YJM1252 | IX | 1 | 10 | 414951 | leftend | YIL169C | 96.879 | 1442 | 38 | 3 | 478 | 1919 | 7429 | 5995 | 0 | 2407 |
| YJM1252 | IX | 413894 | 413903 | 414951 | middle |  |  |  |  |  |  |  |  |  |  |  |
| YJM1252 | IX | 414942 | 414951 | 414951 | rightend | YIR039C | 98.885 | 1614 | 18 | 0 | 1 | 1614 | 407409 | 405796 | 0 | 2881 |
| YJM1252 | X | 1 | 10 | 730533 | leftend | YJL213W | 99.699 | 996 | 3 | 0 | 1 | 996 | 39361 | 40356 | 0 | 1823 |
| YJM1252 | X | 840 | 849 | 730533 | middle |  |  |  |  |  |  |  |  |  |  |  |
| YJM1252 | X | 2284 | 2293 | 730533 | middle |  |  |  |  |  |  |  |  |  |  |  |
| YJM1252 | X | 2875 | 2884 | 730533 | middle |  |  |  |  |  |  |  |  |  |  |  |
| YJM1252 | X | 730533 | 730533 | 730533 | rightend_complete | YJR152W | 91.115 | 1632 | 145 | 0 | 1 | 1632 | 710344 | 711975 | 0 | 2211 |
| YJM1252 | XI | 1 | 10 | 662669 | leftend | YKL222C | 89.099 | 2119 | 229 | 2 | 1 | 2118 | 10115 | 7998 | 0 | 2632 |
| YJM1252 | XI | 533 | 542 | 662669 | middle |  |  |  |  |  |  |  |  |  |  |  |
| YJM1252 | XI | 783 | 792 | 662669 | middle |  |  |  |  |  |  |  |  |  |  |  |
| YJM1252 | XI | 662660 | 662669 | 662669 | rightend | YKR101W | 99.847 | 1965 | 3 | 0 | 1 | 1965 | 644869 | 646833 | 0 | 3613 |
| YJM1252 | XII | 1 |  | 2004038 | leftend_complete | YLL062C | 100 | 975 | 0 | 0 | 1 | 975 | 4449 | 3475 | 0 | 1801 |
| YJM1252 | XII | 239 | 248 | 2004038 | middle |  |  |  |  |  |  |  |  |  |  |  |
| YJM1252 | XII | 2002970 | 2002979 | 2004038 | middle |  |  |  |  |  |  |  |  |  |  |  |
| YJM1252 | XII | 2003493 | 2003502 | 2004038 | middle |  |  |  |  |  |  |  |  |  |  |  |
| YJM1252 | XII | 2004029 | 2004038 | 2004038 | rightend | YLR460C | 94.248 | 1130 | 63 | 2 | 3 | 1131 | 2000994 | 1999866 | 0 | 1725 |
| YJM1252 | XIII | 1 | 10 | 914991 | leftend | YML131W | 99.818 | 1098 | 2 | 0 | 1 | 1098 | 3017 | 4114 | 0 | 2017 |
| YJM1252 | XIII | 914982 | 914991 | 914991 | rightend | YMR321C | 99.361 | 313 | 2 | 0 | 6 | 318 | 887416 | 887104 | 2.48E-161 | 568 |
| YJM1252 | XIV | 1 |  | 761430 | leftend_complete | YNL332W | 98.534 | 1023 | 15 | 0 | 1 | 1023 | 5841 | 6863 | 0 | 1807 |
| YJM1252 | XIV | 523 | 532 | 761430 | middle |  |  |  |  |  |  |  |  |  |  |  |
| YJM1252 | XIV | 760260 | 760269 | 761430 | middle |  |  |  |  |  |  |  |  |  |  |  |
| YJM1252 | XIV | 760884 | 760893 | 761430 | middle |  |  |  |  |  |  |  |  |  |  |  |
| YJM1252 | XIV | 761430 | 761430 | 761430 | rightend_complete | YNR071C | 98.542 | 1029 | 15 | 0 | 1 | 1029 | 744744 | 743716 | 0 | 1818 |
| YJM1252 | XV | 1 |  | 1041599 | leftend_complete | YOL154W | 99.6 | 750 | 3 | 0 | 1 | 750 | 34806 | 35555 | 0 | 1369 |
| YJM1252 | XV | 612 | 621 | 1041599 | middle |  |  |  |  |  |  |  |  |  |  |  |
| YJM1252 | XV | 11660 | 11669 | 1041599 | middle |  |  |  |  |  |  |  |  |  |  |  |
| YJM1252 | XV | 1040972 | 1040981 | 1041599 | middle |  |  |  |  |  |  |  |  |  |  |  |
| YJM1252 | XV | 1041590 | 1041599 | 1041599 | rightend | YOR381W | 94.192 | 2135 | 122 | 2 | 1 | 2134 | 1032260 | 1034393 | 0 | 3254 |
| YJM1252 | XVI | 1 | 10 | 915850 | leftend | YPL274W | 99.773 | 1764 | 4 | 0 | 1 | 1764 | 24761 | 26524 | 0 | 3236 |
| YJM1252 | XVI | 915841 | 915850 | 915850 | rightend | YPR201W | 99.918 | 1215 | 1 | 0 | 1 | 1215 | 912597 | 913811 | 0 | 2239 |
| YJM1273 | I | 1 | 10 | 213122 | leftend | YAL063C | 84.452 | 2547 | 376 | 13 | 1 | 2537 | 12241 | 9705 | 0 | 2492 |
| YJM1273 | I | 9695 | 9704 | 213122 | middle |  |  |  |  |  |  |  |  |  |  |  |
| YJM1273 | I | 189208 | 189217 | 213122 | middle |  |  |  |  |  |  |  |  |  |  |  |
| YJM1273 | I | 190624 | 190633 | 213122 | middle |  |  |  |  |  |  |  |  |  |  |  |
| YJM1273 | I | 213113 | 213122 | 213122 | rightend | YAR050W | 81.116 | 2473 | 445 | 15 | 76 | 2537 | 12166 | 9705 | 0 | 1960 |
| YJM1273 | II | 1 |  | 781279 | leftend_complete | YNCB0001W | 100 | 84 | 0 | 0 | 1 | 84 | 2421 | 2504 | 4.16E-38 | 156 |
| YJM1273 | II | 1125 | 1134 | 781279 | middle |  |  |  |  |  |  |  |  |  |  |  |
| YJM1273 | II | 781270 | 781279 | 781279 | rightend | YBR297W | 98.257 | 1205 | 21 | 0 | 1 | 1205 | 775846 | 777050 | 0 | 2109 |
| YJM1273 | III | 1 | 10 | 309827 | leftend | YCL073C | 99.08 | 1848 | 17 | 0 | 1 | 1848 | 7492 | 5645 | 0 | 3319 |
| YJM1273 | III | 2906 | 2915 | 309827 | middle |  |  |  |  |  |  |  |  |  |  |  |
| YJM1273 | III | 309827 | 309827 | 309827 | rightend_complete | YCR101C | 98 | 550 | 9 | 2 | 1 | 549 | 296178 | 295630 | 0 | 953 |
| YJM1273 | IV | 1 | 10 | 1467376 | leftend | YDL243C | 93.131 | 990 | 68 | 0 | 1 | 990 | 16180 | 15191 | 0 | 1452 |
| YJM1273 | IV | 1467367 | 1467376 | 1467376 | rightend | YDR541C | 99.517 | 1035 | 5 | 0 | 1 | 1035 | 1464532 | 1463498 | 0 | 1884 |
| YJM1273 | V | 1 | 10 | 554621 | leftend | YEL070W | 99.205 | 1509 | 12 | 0 | 1 | 1509 | 14702 | 16210 | 0 | 2721 |
| YJM1273 | V | 554612 | 554621 | 554621 | rightend | YER187W | 99.296 | 426 | 3 | 0 | 1 | 426 | 548107 | 548532 | 0 | 771 |
| YJM1273 | VI | 1 | 10 | 268577 | leftend | YFL051C | 100 | 452 | 0 | 0 | 1 | 452 | 30209 | 29758 | 0 | 835 |
| YJM1273 | VI | 268568 | 268577 | 268577 | rightend | YFR055W | 99.609 | 1022 | 4 | 0 | 1 | 1022 | 258881 | 259902 | 0 | 1866 |
| YJM1273 | VII | 1 |  | 1077943 | leftend_complete | YGL258W | 99.678 | 621 | 2 | 0 | 1 | 621 | 15558 | 16178 | 0 | 1136 |
| YJM1273 | VII | 6673 | 6682 | 1077943 | middle |  |  |  |  |  |  |  |  |  |  |  |
| YJM1273 | VII | 1077943 | 1077943 | 1077943 | rightend_complete | YGR287C | 99.605 | 1770 | 7 | 0 | 1 | 1770 | 1041288 | 1039519 | 0 | 3230 |
| YJM1273 | VIII | 1 |  | 517899 | leftend_complete | YHL044W | 98.729 | 708 | 9 | 0 | 1 | 708 | 9971 | 10678 | 0 | 1258 |
| YJM1273 | VIII | 1657 | 1666 | 517899 | middle |  |  |  |  |  |  |  |  |  |  |  |
| YJM1273 | VIII | 515554 | 515563 | 517899 | middle |  |  |  |  |  |  |  |  |  |  |  |
| YJM1273 | VIII | 516563 | 516572 | 517899 | middle |  |  |  |  |  |  |  |  |  |  |  |
| YJM1273 | VIII | 517899 | 517899 | 517899 | rightend_complete | YHR211W | 81.473 | 1317 | 221 | 13 | 1 | 1304 | 514247 | 515553 | 0 | 1059 |
| YJM1273 | IX | 1 |  | 416536 | leftend_complete | YIL169C | 96.159 | 2265 | 76 | 7 | 352 | 2612 | 8476 | 6219 | 0 | 3690 |
| YJM1273 | IX | 1472 | 1481 | 416536 | middle |  |  |  |  |  |  |  |  |  |  |  |
| YJM1273 | IX | 416527 | 416536 | 416536 | rightend | YIR039C | 99.62 | 1317 | 5 | 0 | 1 | 1317 | 409068 | 407752 | 0 | 2405 |
| YJM1273 | X | 1 | 10 | 717058 | leftend | YJL213W | 99.799 | 996 | 2 | 0 | 1 | 996 | 25941 | 26936 | 0 | 1829 |
| YJM1273 | X | 717058 | 717058 | 717058 | rightend_complete | YJR152W | 99.449 | 1632 | 9 | 0 | 1 | 1632 | 696055 | 697686 | 0 | 2964 |
| YJM1273 | XI | 1 | 10 | 663591 | leftend | YKL222C | 99.669 | 2118 | 7 | 0 | 1 | 2118 | 3476 | 1359 | 0 | 3873 |
| YJM1273 | XI | 663582 | 663591 | 663591 | rightend | YKR101W | 99.644 | 1965 | 7 | 0 | 1 | 1965 | 637711 | 639675 | 0 | 3591 |
| YJM1273 | XII | 1 | 10 | 1829475 | leftend | YLL062C | 99.59 | 975 | 4 | 0 | 1 | 975 | 4491 | 3517 | 0 | 1779 |
| YJM1273 | XII | 1826608 | 1826617 | 1829475 | middle |  |  |  |  |  |  |  |  |  |  |  |
| YJM1273 | XII | 1827700 | 1827709 | 1829475 | middle |  |  |  |  |  |  |  |  |  |  |  |
| YJM1273 | XII | 1828276 | 1828285 | 1829475 | middle |  |  |  |  |  |  |  |  |  |  |  |
| YJM1273 | XII | 1829475 | 1829475 | 1829475 | rightend_complete | YLR460C | 99.469 | 1131 | 6 | 0 | 1 | 1131 | 1825817 | 1824687 | 0 | 2056 |
| YJM1273 | XIII | 1 | 10 | 913945 | leftend | YML131W | 99.545 | 1098 | 5 | 0 | 1 | 1098 | 3708 | 4805 | 0 | 2001 |
| YJM1273 | XIII | 913936 | 913945 | 913945 | rightend | YMR321C | 99.361 | 313 | 2 | 0 | 6 | 318 | 888026 | 887714 | 2.48E-161 | 568 |
| YJM1273 | XIV | 1 | 10 | 763090 | leftend | YNL332W | 99.022 | 1023 | 10 | 0 | 1 | 1023 | 12188 | 13210 | 0 | 1834 |
| YJM1273 | XIV | 5814 | 5823 | 763090 | middle |  |  |  |  |  |  |  |  |  |  |  |
| YJM1273 | XIV | 763081 | 763090 | 763090 | rightend | YNR071C | 99.611 | 1029 | 4 | 0 | 1 | 1029 | 752626 | 751598 | 0 | 1879 |
| YJM1273 | XV | 1 |  | 1065730 | leftend_complete | YOL154W | 99.733 | 750 | 2 | 0 | 1 | 750 | 35495 | 36244 | 0 | 1375 |
| YJM1273 | XV | 1060966 | 1060975 | 1065730 | middle |  |  |  |  |  |  |  |  |  |  |  |
| YJM1273 | XV | 1065721 | 1065730 | 1065730 | rightend | YOR381W | 98.876 | 2136 | 24 | 0 | 1 | 2136 | 1032271 | 1034406 | 0 | 3812 |
| YJM1273 | XVI | 1 |  | 905630 | leftend_complete | YPL274W | 98.186 | 1764 | 32 | 0 | 1 | 1764 | 10045 | 11808 | 0 | 3081 |
| YJM1273 | XVI | 916 | 925 | 905630 | middle |  |  |  |  |  |  |  |  |  |  |  |
| YJM1273 | XVI | 905621 | 905630 | 905630 | rightend | YPR201W | 99.506 | 1215 | 6 | 0 | 1 | 1215 | 900017 | 901231 | 0 | 2211 |
| YJM1304 | I | 1 | 10 | 215530 | leftend | YAL063C | 91.242 | 1256 | 72 | 20 | 2715 | 3962 | 10514 | 9289 | 0 | 1676 |
| YJM1304 | I | 11343 | 11352 | 215530 | middle |  |  |  |  |  |  |  |  |  |  |  |
| YJM1304 | I | 184835 | 184844 | 215530 | middle |  |  |  |  |  |  |  |  |  |  |  |
| YJM1304 | I | 206572 | 206581 | 215530 | middle |  |  |  |  |  |  |  |  |  |  |  |
| YJM1304 | I | 215521 | 215530 | 215530 | rightend | YAR050W | 92.4 | 1750 | 102 | 13 | 2530 | 4253 | 11333 | 9589 | 0 | 2466 |
| YJM1304 | II | 1 | 10 | 779472 | leftend | YNCB0001W | 100 | 84 | 0 | 0 | 1 | 84 | 2171 | 2254 | 4.16E-38 | 156 |
| YJM1304 | II | 732 | 741 | 779472 | middle |  |  |  |  |  |  |  |  |  |  |  |
| YJM1304 | II | 1636 | 1645 | 779472 | middle |  |  |  |  |  |  |  |  |  |  |  |
| YJM1304 | II | 779463 | 779472 | 779472 | rightend | YBR297W | 98.008 | 1205 | 24 | 0 | 1 | 1205 | 775235 | 776439 | 0 | 2093 |
| YJM1304 | III | 1 | 10 | 311026 | leftend | YCL073C | 99.675 | 1848 | 6 | 0 | 1 | 1848 | 7091 | 5244 | 0 | 3380 |
| YJM1304 | III | 311017 | 311026 | 311026 | rightend | YCR101C | 99.818 | 549 | 1 | 0 | 1 | 549 | 297765 | 297217 | 0 | 1009 |
| YJM1304 | IV | 1 |  | 1471989 | leftend_complete | YDL243C | 100 | 990 | 0 | 0 | 1 | 990 | 18474 | 17485 | 0 | 1829 |
| YJM1304 | IV | 1469567 | 1469576 | 1471989 | middle |  |  |  |  |  |  |  |  |  |  |  |
| YJM1304 | IV | 1471335 | 1471344 | 1471989 | middle |  |  |  |  |  |  |  |  |  |  |  |
| YJM1304 | IV | 1471989 | 1471989 | 1471989 | rightend_complete | YDR541C | 99.614 | 1035 | 4 | 0 | 1 | 1035 | 1466701 | 1465667 | 0 | 1890 |
| YJM1304 | V | 1 | 10 | 551515 | leftend | YEL070W | 99.205 | 1509 | 12 | 0 | 1 | 1509 | 13324 | 14832 | 0 | 2721 |
| YJM1304 | V | 550306 | 550315 | 551515 | middle |  |  |  |  |  |  |  |  |  |  |  |
| YJM1304 | V | 551506 | 551515 | 551515 | rightend | YER187W | 91.981 | 424 | 32 | 2 | 4 | 426 | 547540 | 547962 | 3.44E-169 | 593 |
| YJM1304 | VI | 1 | 10 | 261614 | leftend | YFL051C | 100 | 483 | 0 | 0 | 1 | 483 | 17508 | 17026 | 0 | 893 |
| YJM1304 | VI | 808 | 817 | 261614 | middle |  |  |  |  |  |  |  |  |  |  |  |
| YJM1304 | VI | 261614 | 261614 | 261614 | rightend_complete | YFR055W | 99.511 | 1022 | 5 | 0 | 1 | 1022 | 244695 | 245716 | 0 | 1860 |
| YJM1304 | VII | 1 | 10 | 1067040 | leftend | YGL258W | 99.356 | 621 | 4 | 0 | 1 | 621 | 22355 | 22975 | 0 | 1125 |
| YJM1304 | VII | 1067031 | 1067040 | 1067040 | rightend | YGR287C | 98.814 | 1770 | 21 | 0 | 1 | 1770 | 1044032 | 1042263 | 0 | 3153 |
| YJM1304 | VIII | 1 | 10 | 517704 | leftend | YHL044W | 98.588 | 708 | 10 | 0 | 1 | 708 | 1397 | 2104 | 0 | 1253 |
| YJM1304 | VIII | 517704 | 517704 | 517704 | rightend_complete | YHR211W | 85.537 | 1210 | 163 | 8 | 1 | 1202 | 505273 | 506478 | 0 | 1254 |
| YJM1304 | IX | 1 |  | 423706 | leftend_complete | YIL169C | 96.539 | 1907 | 56 | 8 | 562 | 2466 | 19883 | 17985 | 0 | 3147 |
| YJM1304 | IX | 249 | 258 | 423706 | middle |  |  |  |  |  |  |  |  |  |  |  |
| YJM1304 | IX | 423697 | 423706 | 423706 | rightend | YIR039C | 99.938 | 1614 | 1 | 0 | 1 | 1614 | 417480 | 415867 | 0 | 2976 |
| YJM1304 | X | 1 |  | 697006 | leftend_complete | YJL213W | 99.699 | 996 | 3 | 0 | 1 | 996 | 14368 | 15363 | 0 | 1823 |
| YJM1304 | X | 2408 | 2417 | 697006 | middle |  |  |  |  |  |  |  |  |  |  |  |
| YJM1304 | X | 3056 | 3065 | 697006 | middle |  |  |  |  |  |  |  |  |  |  |  |
| YJM1304 | X | 3304 | 3313 | 697006 | middle |  |  |  |  |  |  |  |  |  |  |  |
| YJM1304 | X | 697006 | 697006 | 697006 | rightend_complete | YJR152W | 100 | 1632 | 0 | 0 | 1 | 1632 | 684819 | 686450 | 0 | 3014 |
| YJM1304 | XI | 1 | 10 | 664159 | leftend | YKL222C | 99.481 | 2118 | 11 | 0 | 1 | 2118 | 3200 | 1083 | 0 | 3851 |
| YJM1304 | XI | 664150 | 664159 | 664159 | rightend | YKR101W | 99.695 | 1965 | 6 | 0 | 1 | 1965 | 636679 | 638643 | 0 | 3596 |
| YJM1304 | XII | 1 | 10 | 1713679 | leftend | YLL062C | 99.692 | 975 | 3 | 0 | 1 | 975 | 3274 | 2300 | 0 | 1784 |
| YJM1304 | XII | 1308 | 1317 | 1713679 | middle |  |  |  |  |  |  |  |  |  |  |  |
| YJM1304 | XII | 1711971 | 1711980 | 1713679 | middle |  |  |  |  |  |  |  |  |  |  |  |
| YJM1304 | XII | 1713679 | 1713679 | 1713679 | rightend_complete | YLR460C | 98.762 | 1131 | 14 | 0 | 1 | 1131 | 1710760 | 1709630 | 0 | 2012 |
| YJM1304 | XIII | 1 | 10 | 911087 | leftend | YML131W | 99.454 | 1098 | 6 | 0 | 1 | 1098 | 2617 | 3714 | 0 | 1995 |
| YJM1304 | XIII | 904475 | 904484 | 911087 | middle |  |  |  |  |  |  |  |  |  |  |  |
| YJM1304 | XIII | 911078 | 911087 | 911087 | rightend | YMR321C | 99.361 | 313 | 2 | 0 | 6 | 318 | 887022 | 886710 | 2.48E-161 | 568 |
| YJM1304 | XIV | 1 | 10 | 750381 | leftend | YNL332W | 99.316 | 1023 | 7 | 0 | 1 | 1023 | 5888 | 6910 | 0 | 1851 |
| YJM1304 | XIV | 750372 | 750381 | 750381 | rightend | YNR071C | 100 | 1029 | 0 | 0 | 1 | 1029 | 745036 | 744008 | 0 | 1901 |
| YJM1304 | XV | 1 |  | 1022162 | leftend_complete | YOL154W | 99.467 | 750 | 4 | 0 | 1 | 750 | 8354 | 9103 | 0 | 1363 |
| YJM1304 | XV | 1022153 | 1022162 | 1022162 | rightend | YOR381W | 98.736 | 2136 | 27 | 0 | 1 | 2136 | 1005602 | 1007737 | 0 | 3795 |
| YJM1304 | XVI | 1 | 10 | 907568 | leftend | YPL274W | 98.129 | 1764 | 33 | 0 | 1 | 1764 | 14649 | 16412 | 0 | 3075 |
| YJM1304 | XVI | 907559 | 907568 | 907568 | rightend | YPR201W | 99.835 | 1215 | 2 | 0 | 1 | 1215 | 904792 | 906006 | 0 | 2233 |
| YJM1307 | I | 1 | 10 | 199293 | leftend | YAL063C | 85.416 | 1865 | 264 | 7 | 40 | 1900 | 185948 | 187808 | 0 | 1930 |
| YJM1307 | I | 10591 | 10600 | 199293 | middle |  |  |  |  |  |  |  |  |  |  |  |
| YJM1307 | I | 10992 | 11001 | 199293 | middle |  |  |  |  |  |  |  |  |  |  |  |
| YJM1307 | I | 199284 | 199293 | 199293 | rightend | YAR050W | 91.09 | 2458 | 205 | 12 | 2164 | 4614 | 186722 | 189172 | 0 | 3314 |
| YJM1307 | II | 1 | 10 | 786389 | leftend | YNCB0001W | 100 | 84 | 0 | 0 | 1 | 84 | 2846 | 2929 | 4.16E-38 | 156 |
| YJM1307 | II | 893 | 902 | 786389 | middle |  |  |  |  |  |  |  |  |  |  |  |
| YJM1307 | II | 1459 | 1468 | 786389 | middle |  |  |  |  |  |  |  |  |  |  |  |
| YJM1307 | II | 785744 | 785753 | 786389 | middle |  |  |  |  |  |  |  |  |  |  |  |
| YJM1307 | II | 786380 | 786389 | 786389 | rightend | YBR297W | 97.759 | 1205 | 27 | 0 | 1 | 1205 | 774920 | 776124 | 0 | 2076 |
| YJM1307 | III | 1 | 10 | 303769 | leftend | YCL073C | 99.675 | 1848 | 6 | 0 | 1 | 1848 | 7980 | 6133 | 0 | 3380 |
| YJM1307 | III | 301632 | 301641 | 303769 | middle |  |  |  |  |  |  |  |  |  |  |  |
| YJM1307 | III | 303231 | 303240 | 303769 | middle |  |  |  |  |  |  |  |  |  |  |  |
| YJM1307 | III | 303769 | 303769 | 303769 | rightend_complete | YCR101C | 99.455 | 550 | 2 | 1 | 1 | 549 | 297378 | 296829 | 0 | 998 |
| YJM1307 | IV | 1 | 10 | 1484074 | leftend | YDL243C | 93.037 | 991 | 66 | 3 | 1 | 990 | 15993 | 15005 | 0 | 1445 |
| YJM1307 | IV | 1484065 | 1484074 | 1484074 | rightend | YDR541C | 99.227 | 1035 | 8 | 0 | 1 | 1035 | 1478905 | 1477871 | 0 | 1868 |
| YJM1307 | V | 1 |  | 552948 | leftend_complete | YEL070W | 99.072 | 1509 | 14 | 0 | 1 | 1509 | 13730 | 15238 | 0 | 2710 |
| YJM1307 | V | 552939 | 552948 | 552948 | rightend | YER187W | 99.765 | 426 | 1 | 0 | 1 | 426 | 548482 | 548907 | 0 | 782 |
| YJM1307 | VI | 1 | 10 | 259965 | leftend | YFL051C | 98.451 | 452 | 7 | 0 | 1 | 452 | 25586 | 25135 | 0 | 797 |
| YJM1307 | VI | 1459 | 1468 | 259965 | middle |  |  |  |  |  |  |  |  |  |  |  |
| YJM1307 | VI | 259956 | 259965 | 259965 | rightend | YFR055W | 99.804 | 1022 | 2 | 0 | 1 | 1022 | 253904 | 254925 | 0 | 1877 |
| YJM1307 | VII | 1 |  | 1047096 | leftend_complete | YGL258W | 97.424 | 621 | 16 | 0 | 1 | 621 | 11281 | 11901 | 0 | 1059 |
| YJM1307 | VII | 1045328 | 1045337 | 1047096 | middle |  |  |  |  |  |  |  |  |  |  |  |
| YJM1307 | VII | 1047087 | 1047096 | 1047096 | rightend | YGR287C | 98.757 | 1770 | 22 | 0 | 1 | 1770 | 1030472 | 1028703 | 0 | 3147 |
| YJM1307 | VIII | 1 | 10 | 531481 | leftend | YHL044W | 98.588 | 708 | 10 | 0 | 1 | 708 | 7199 | 7906 | 0 | 1253 |
| YJM1307 | VIII | 528177 | 528186 | 531481 | middle |  |  |  |  |  |  |  |  |  |  |  |
| YJM1307 | VIII | 531481 | 531481 | 531481 | rightend_complete | YBR297W | 75.57 | 1404 | 313 | 23 | 22 | 1407 | 525228 | 526619 | 0 | 665 |
| YJM1307 | IX | 1 | 10 | 419401 | leftend | YIL169C | 97.946 | 1947 | 34 | 4 | 520 | 2466 | 17088 | 15148 | 0 | 3369 |
| YJM1307 | IX | 1867 | 1876 | 419401 | middle |  |  |  |  |  |  |  |  |  |  |  |
| YJM1307 | IX | 7465 | 7474 | 419401 | middle |  |  |  |  |  |  |  |  |  |  |  |
| YJM1307 | IX | 419392 | 419401 | 419401 | rightend | YIR039C | 99.566 | 1614 | 7 | 0 | 1 | 1614 | 417791 | 416178 | 0 | 2942 |
| YJM1307 | X | 1 |  | 707065 | leftend_complete | YJL213W | 99.498 | 997 | 4 | 1 | 1 | 996 | 26187 | 27183 | 0 | 1812 |
| YJM1307 | X | 3765 | 3774 | 707065 | middle |  |  |  |  |  |  |  |  |  |  |  |
| YJM1307 | X | 704269 | 704278 | 707065 | middle |  |  |  |  |  |  |  |  |  |  |  |
| YJM1307 | X | 704774 | 704783 | 707065 | middle |  |  |  |  |  |  |  |  |  |  |  |
| YJM1307 | X | 705258 | 705267 | 707065 | middle |  |  |  |  |  |  |  |  |  |  |  |
| YJM1307 | X | 707065 | 707065 | 707065 | rightend_complete | YJR152W | 99.387 | 1632 | 10 | 0 | 1 | 1632 | 696471 | 698102 | 0 | 2959 |
| YJM1307 | XI | 1 | 10 | 654436 | leftend | YKL222C | 99.481 | 2118 | 11 | 0 | 1 | 2118 | 11773 | 9656 | 0 | 3851 |
| YJM1307 | XI | 654427 | 654436 | 654436 | rightend | YKR101W | 99.796 | 1965 | 4 | 0 | 1 | 1965 | 646362 | 648326 | 0 | 3607 |
| YJM1307 | XII | 1 | 10 | 2391209 | leftend | YLL062C | 99.487 | 975 | 5 | 0 | 1 | 975 | 4812 | 3838 | 0 | 1773 |
| YJM1307 | XII | 1000 | 1009 | 2391209 | middle |  |  |  |  |  |  |  |  |  |  |  |
| YJM1307 | XII | 2389940 | 2389949 | 2391209 | middle |  |  |  |  |  |  |  |  |  |  |  |
| YJM1307 | XII | 2391209 | 2391209 | 2391209 | rightend_complete | YLR460C | 95.044 | 1130 | 54 | 2 | 3 | 1131 | 2388144 | 2387016 | 0 | 1775 |
| YJM1307 | XIII | 1 | 10 | 895222 | leftend | YML131W | 99.089 | 1098 | 10 | 0 | 1 | 1098 | 2991 | 4088 | 0 | 1973 |
| YJM1307 | XIII | 895213 | 895222 | 895222 | rightend | YMR321C | 99.361 | 313 | 2 | 0 | 6 | 318 | 887027 | 886715 | 2.48E-161 | 568 |
| YJM1307 | XIV | 1 | 10 | 768687 | leftend | YNL332W | 98.827 | 1023 | 12 | 0 | 1 | 1023 | 9822 | 10844 | 0 | 1823 |
| YJM1307 | XIV | 768678 | 768687 | 768687 | rightend | YNR071C | 99.514 | 1029 | 5 | 0 | 1 | 1029 | 749375 | 748347 | 0 | 1873 |
| YJM1307 | XV | 1 |  | 1036839 | leftend_complete | YOL154W | 99.733 | 750 | 2 | 0 | 1 | 750 | 20552 | 21301 | 0 | 1375 |
| YJM1307 | XV | 1027598 | 1027607 | 1036839 | middle |  |  |  |  |  |  |  |  |  |  |  |
| YJM1307 | XV | 1036830 | 1036839 | 1036839 | rightend | YOR381W | 94.939 | 2134 | 108 | 0 | 1 | 2134 | 1018925 | 1021058 | 0 | 3343 |
| YJM1307 | XVI | 1 | 10 | 914338 | leftend | YPL274W | 99.546 | 1764 | 8 | 0 | 1 | 1764 | 24535 | 26298 | 0 | 3214 |
| YJM1307 | XVI | 957 | 966 | 914338 | middle |  |  |  |  |  |  |  |  |  |  |  |
| YJM1307 | XVI | 914329 | 914338 | 914338 | rightend | YPR201W | 100 | 1215 | 0 | 0 | 1 | 1215 | 911201 | 912415 | 0 | 2244 |
| YJM1311 | I | 1 |  | 214632 | leftend_complete | YAL063C | 93.894 | 1261 | 43 | 14 | 2715 | 3969 | 194619 | 195851 | 0 | 1871 |
| YJM1311 | I | 6605 | 6614 | 214632 | middle |  |  |  |  |  |  |  |  |  |  |  |
| YJM1311 | I | 214623 | 214632 | 214632 | rightend | YAR050W | 90.021 | 1423 | 114 | 10 | 2851 | 4250 | 194124 | 195541 | 0 | 1816 |
| YJM1311 | II | 1 |  | 788187 | leftend_complete | YNCB0001W | 100 | 84 | 0 | 0 | 1 | 84 | 4489 | 4572 | 4.16E-38 | 156 |
| YJM1311 | II | 3260 | 3269 | 788187 | middle |  |  |  |  |  |  |  |  |  |  |  |
| YJM1311 | II | 788178 | 788187 | 788187 | rightend | YBR297W | 92.172 | 1418 | 95 | 12 | 1 | 1407 | 777878 | 779290 | 0 | 1989 |
| YJM1311 | III | 1 | 10 | 310473 | leftend | YCL073C | 99.026 | 1848 | 18 | 0 | 1 | 1848 | 7989 | 6142 | 0 | 3314 |
| YJM1311 | III | 310473 | 310473 | 310473 | rightend_complete | YCR101C | 99.818 | 549 | 1 | 0 | 1 | 549 | 296922 | 296374 | 0 | 1009 |
| YJM1311 | IV | 1 |  | 1476927 | leftend_complete | YDL243C | 93.769 | 979 | 61 | 0 | 1 | 979 | 17722 | 16744 | 0 | 1471 |
| YJM1311 | IV | 1476918 | 1476927 | 1476927 | rightend | YDR541C | 99.614 | 1035 | 4 | 0 | 1 | 1035 | 1473118 | 1472084 | 0 | 1890 |
| YJM1311 | V | 1 | 10 | 556975 | leftend | YEL070W | 99.404 | 1509 | 9 | 0 | 1 | 1509 | 13391 | 14899 | 0 | 2737 |
| YJM1311 | V | 556975 | 556975 | 556975 | rightend_complete | YER187W | 91.981 | 424 | 32 | 2 | 4 | 426 | 548271 | 548693 | 3.44E-169 | 593 |
| YJM1311 | VI | 1 | 10 | 238636 | leftend | YFL051C | 98.451 | 452 | 7 | 0 | 1 | 452 | 1164 | 713 | 0 | 797 |
| YJM1311 | VI | 238627 | 238636 | 238636 | rightend | YFR055W | 99.609 | 1022 | 4 | 0 | 1 | 1022 | 228972 | 229993 | 0 | 1866 |
| YJM1311 | VII | 1 |  | 1058158 | leftend_complete | YGL258W | 97.585 | 621 | 15 | 0 | 1 | 621 | 11150 | 11770 | 0 | 1064 |
| YJM1311 | VII | 1056697 | 1056706 | 1058158 | middle |  |  |  |  |  |  |  |  |  |  |  |
| YJM1311 | VII | 1056942 | 1056951 | 1058158 | middle |  |  |  |  |  |  |  |  |  |  |  |
| YJM1311 | VII | 1058149 | 1058158 | 1058158 | rightend | YGR287C | 99.887 | 1770 | 2 | 0 | 1 | 1770 | 1033075 | 1031306 | 0 | 3258 |
| YJM1311 | VIII | 1 | 10 | 557632 | leftend | YHL044W | 98.588 | 708 | 10 | 0 | 1 | 708 | 7697 | 8404 | 0 | 1253 |
| YJM1311 | VIII | 539080 | 539089 | 557632 | middle |  |  |  |  |  |  |  |  |  |  |  |
| YJM1311 | VIII | 557623 | 557632 | 557632 | rightend | YHR211W | 97.164 | 1728 | 49 | 0 | 1501 | 3228 | 515114 | 516841 | 0 | 2920 |
| YJM1311 | IX | 1 | 10 | 402403 | leftend | YIL169C | 92.475 | 1422 | 105 | 2 | 394 | 1814 | 2174 | 754 | 0 | 2032 |
| YJM1311 | IX | 402394 | 402403 | 402403 | rightend | YIR039C | 99.009 | 1614 | 16 | 0 | 1 | 1614 | 401128 | 399515 | 0 | 2892 |
| YJM1311 | X | 1 |  | 716653 | leftend_complete | YJL213W | 99.699 | 996 | 3 | 0 | 1 | 996 | 27829 | 28824 | 0 | 1823 |
| YJM1311 | X | 1512 | 1521 | 716653 | middle |  |  |  |  |  |  |  |  |  |  |  |
| YJM1311 | X | 716644 | 716653 | 716653 | rightend | YJR152W | 99.755 | 1632 | 3 | 1 | 1 | 1632 | 698665 | 700295 | 0 | 2990 |
| YJM1311 | XI | 1 | 10 | 657192 | leftend | YKL222C | 99.387 | 2119 | 12 | 1 | 1 | 2118 | 3353 | 1235 | 0 | 3840 |
| YJM1311 | XI | 656375 | 656384 | 657192 | middle |  |  |  |  |  |  |  |  |  |  |  |
| YJM1311 | XI | 657192 | 657192 | 657192 | rightend_complete | YKR101W | 99.796 | 1965 | 4 | 0 | 1 | 1965 | 638086 | 640050 | 0 | 3607 |
| YJM1311 | XII | 1 |  | 1858724 | leftend_complete | YLL062C | 100 | 975 | 0 | 0 | 1 | 975 | 5981 | 5007 | 0 | 1801 |
| YJM1311 | XII | 1967 | 1976 | 1858724 | middle |  |  |  |  |  |  |  |  |  |  |  |
| YJM1311 | XII | 1856873 | 1856882 | 1858724 | middle |  |  |  |  |  |  |  |  |  |  |  |
| YJM1311 | XII | 1858715 | 1858724 | 1858724 | rightend | YLR460C | 94.071 | 1130 | 65 | 2 | 3 | 1131 | 1856757 | 1855629 | 0 | 1714 |
| YJM1311 | XIII | 1 |  | 898272 | leftend_complete | YML131W | 99.454 | 1098 | 6 | 0 | 1 | 1098 | 4694 | 5791 | 0 | 1995 |
| YJM1311 | XIII | 896213 | 896222 | 898272 | middle |  |  |  |  |  |  |  |  |  |  |  |
| YJM1311 | XIII | 898263 | 898272 | 898272 | rightend | YMR321C | 99.361 | 313 | 2 | 0 | 6 | 318 | 889152 | 888840 | 2.48E-161 | 568 |
| YJM1311 | XIV | 1 | 10 | 756170 | leftend | YNL332W | 98.338 | 1023 | 17 | 0 | 1 | 1023 | 6822 | 7844 | 0 | 1796 |
| YJM1311 | XIV | 789 | 798 | 756170 | middle |  |  |  |  |  |  |  |  |  |  |  |
| YJM1311 | XIV | 1501 | 1510 | 756170 | middle |  |  |  |  |  |  |  |  |  |  |  |
| YJM1311 | XIV | 756161 | 756170 | 756170 | rightend | YNR071C | 100 | 1029 | 0 | 0 | 1 | 1029 | 746091 | 745063 | 0 | 1901 |
| YJM1311 | XV | 1 | 10 | 1043478 | leftend | YOL154W | 99.467 | 750 | 4 | 0 | 1 | 750 | 19752 | 20501 | 0 | 1363 |
| YJM1311 | XV | 1040754 | 1040763 | 1043478 | middle |  |  |  |  |  |  |  |  |  |  |  |
| YJM1311 | XV | 1043469 | 1043478 | 1043478 | rightend | YOR381W | 99.906 | 2136 | 2 | 0 | 1 | 2136 | 1016163 | 1018298 | 0 | 3934 |
| YJM1311 | XVI | 1 |  | 904934 | leftend_complete | YPL274W | 98.639 | 1764 | 24 | 0 | 1 | 1764 | 19990 | 21753 | 0 | 3125 |
| YJM1311 | XVI | 5700 | 5709 | 904934 | middle |  |  |  |  |  |  |  |  |  |  |  |
| YJM1311 | XVI | 8006 | 8015 | 904934 | middle |  |  |  |  |  |  |  |  |  |  |  |
| YJM1311 | XVI | 904925 | 904934 | 904934 | rightend | YPR201W | 99.671 | 1215 | 4 | 0 | 1 | 1215 | 901911 | 903125 | 0 | 2222 |
| YJM1326 | I | 1 | 10 | 200912 | leftend | YAL063C | 94.721 | 1004 | 32 | 6 | 2966 | 3969 | 19251 | 18269 | 0 | 1541 |
| YJM1326 | I | 7619 | 7628 | 200912 | middle |  |  |  |  |  |  |  |  |  |  |  |
| YJM1326 | I | 8169 | 8178 | 200912 | middle |  |  |  |  |  |  |  |  |  |  |  |
| YJM1326 | I | 18238 | 18247 | 200912 | middle |  |  |  |  |  |  |  |  |  |  |  |
| YJM1326 | I | 196247 | 196256 | 200912 | middle |  |  |  |  |  |  |  |  |  |  |  |
| YJM1326 | I | 200903 | 200912 | 200912 | rightend | YAR050W | 86.354 | 1363 | 167 | 11 | 1 | 1352 | 194889 | 196243 | 0 | 1469 |
| YJM1326 | II | 1 | 10 | 786341 | leftend | YNCB0001W | 100 | 84 | 0 | 0 | 1 | 84 | 3198 | 3281 | 4.16E-38 | 156 |
| YJM1326 | II | 503 | 512 | 786341 | middle |  |  |  |  |  |  |  |  |  |  |  |
| YJM1326 | II | 1616 | 1625 | 786341 | middle |  |  |  |  |  |  |  |  |  |  |  |
| YJM1326 | II | 786332 | 786341 | 786341 | rightend | YBR297W | 98.863 | 1407 | 14 | 1 | 1 | 1407 | 776359 | 777763 | 0 | 2508 |
| YJM1326 | III | 1 | 10 | 308698 | leftend | YCL073C | 99.729 | 1848 | 5 | 0 | 1 | 1848 | 7111 | 5264 | 0 | 3386 |
| YJM1326 | III | 300700 | 300709 | 308698 | middle |  |  |  |  |  |  |  |  |  |  |  |
| YJM1326 | III | 308128 | 308137 | 308698 | middle |  |  |  |  |  |  |  |  |  |  |  |
| YJM1326 | III | 308689 | 308698 | 308698 | rightend | YCR101C | 92.021 | 564 | 28 | 12 | 1 | 549 | 295821 | 295260 | 0 | 776 |
| YJM1326 | IV | 1 | 10 | 1474836 | leftend | YDL243C | 95.152 | 990 | 48 | 0 | 1 | 990 | 19926 | 18937 | 0 | 1563 |
| YJM1326 | IV | 3195 | 3204 | 1474836 | middle |  |  |  |  |  |  |  |  |  |  |  |
| YJM1326 | IV | 1474827 | 1474836 | 1474836 | rightend | YDR541C | 99.42 | 1035 | 6 | 0 | 1 | 1035 | 1469717 | 1468683 | 0 | 1879 |
| YJM1326 | V | 1 | 10 | 554200 | leftend | YEL070W | 98.675 | 1509 | 20 | 0 | 1 | 1509 | 12814 | 14322 | 0 | 2676 |
| YJM1326 | V | 554191 | 554200 | 554200 | rightend | YER187W | 91.981 | 424 | 32 | 2 | 4 | 426 | 547920 | 548342 | 3.44E-169 | 593 |
| YJM1326 | VI | 1 |  | 275380 | leftend_complete | YFL051C | 100 | 452 | 0 | 0 | 1 | 452 | 39714 | 39263 | 0 | 835 |
| YJM1326 | VI | 1175 | 1184 | 275380 | middle |  |  |  |  |  |  |  |  |  |  |  |
| YJM1326 | VI | 3101 | 3110 | 275380 | middle |  |  |  |  |  |  |  |  |  |  |  |
| YJM1326 | VI | 3309 | 3318 | 275380 | middle |  |  |  |  |  |  |  |  |  |  |  |
| YJM1326 | VI | 275371 | 275380 | 275380 | rightend | YFR055W | 99.902 | 1023 | 1 | 0 | 1 | 1023 | 269031 | 270053 | 0 | 1884 |
| YJM1326 | VII | 1 |  | 1050128 | leftend_complete | YGL258W | 99.678 | 621 | 2 | 0 | 1 | 621 | 11146 | 11766 | 0 | 1136 |
| YJM1326 | VII | 6852 | 6861 | 1050128 | middle |  |  |  |  |  |  |  |  |  |  |  |
| YJM1326 | VII | 1048769 | 1048778 | 1050128 | middle |  |  |  |  |  |  |  |  |  |  |  |
| YJM1326 | VII | 1050128 | 1050128 | 1050128 | rightend_complete | YGR287C | 98.814 | 1770 | 21 | 0 | 1 | 1770 | 1030093 | 1028324 | 0 | 3153 |
| YJM1326 | VIII | 1 |  | 559601 | leftend_complete | YHL044W | 98.588 | 708 | 10 | 0 | 1 | 708 | 9382 | 10089 | 0 | 1253 |
| YJM1326 | VIII | 1521 | 1530 | 559601 | middle |  |  |  |  |  |  |  |  |  |  |  |
| YJM1326 | VIII | 559592 | 559601 | 559601 | rightend | YHR211W | 92.135 | 2136 | 144 | 2 | 1096 | 3228 | 530797 | 532911 | 0 | 2992 |
| YJM1326 | IX | 1 |  | 420914 | leftend_complete | YIL169C | 94.513 | 1294 | 69 | 2 | 520 | 1812 | 7137 | 5845 | 0 | 1995 |
| YJM1326 | IX | 845 | 854 | 420914 | middle |  |  |  |  |  |  |  |  |  |  |  |
| YJM1326 | IX | 418348 | 418357 | 420914 | middle |  |  |  |  |  |  |  |  |  |  |  |
| YJM1326 | IX | 420701 | 420710 | 420914 | middle |  |  |  |  |  |  |  |  |  |  |  |
| YJM1326 | IX | 420914 | 420914 | 420914 | rightend_complete | YIR039C | 99.009 | 1614 | 16 | 0 | 1 | 1614 | 412306 | 410693 | 0 | 2892 |
| YJM1326 | X | 1 | 10 | 714396 | leftend | YJL213W | 99.598 | 996 | 4 | 0 | 1 | 996 | 26916 | 27911 | 0 | 1818 |
| YJM1326 | X | 647 | 656 | 714396 | middle |  |  |  |  |  |  |  |  |  |  |  |
| YJM1326 | X | 712580 | 712589 | 714396 | middle |  |  |  |  |  |  |  |  |  |  |  |
| YJM1326 | X | 713384 | 713393 | 714396 | middle |  |  |  |  |  |  |  |  |  |  |  |
| YJM1326 | X | 714396 | 714396 | 714396 | rightend_complete | YJR152W | 99.755 | 1632 | 3 | 1 | 1 | 1632 | 696429 | 698059 | 0 | 2990 |
| YJM1326 | XI | 1 | 10 | 660576 | leftend | YKL222C | 99.387 | 2119 | 12 | 1 | 1 | 2118 | 3121 | 1003 | 0 | 3840 |
| YJM1326 | XI | 660567 | 660576 | 660576 | rightend | YKR101W | 99.796 | 1965 | 4 | 0 | 1 | 1965 | 637384 | 639348 | 0 | 3607 |
| YJM1326 | XII | 1 | 10 | 1675522 | leftend | YLL062C | 99.59 | 975 | 4 | 0 | 1 | 975 | 4038 | 3064 | 0 | 1779 |
| YJM1326 | XII | 1674230 | 1674239 | 1675522 | middle |  |  |  |  |  |  |  |  |  |  |  |
| YJM1326 | XII | 1674778 | 1674787 | 1675522 | middle |  |  |  |  |  |  |  |  |  |  |  |
| YJM1326 | XII | 1675513 | 1675522 | 1675522 | rightend | YLR460C | 94.336 | 1130 | 62 | 2 | 3 | 1131 | 1672686 | 1671558 | 0 | 1731 |
| YJM1326 | XIII | 1 | 10 | 896154 | leftend | YML131W | 100 | 1098 | 0 | 0 | 1 | 1098 | 2308 | 3405 | 0 | 2028 |
| YJM1326 | XIII | 892601 | 892610 | 896154 | middle |  |  |  |  |  |  |  |  |  |  |  |
| YJM1326 | XIII | 893218 | 893227 | 896154 | middle |  |  |  |  |  |  |  |  |  |  |  |
| YJM1326 | XIII | 895095 | 895104 | 896154 | middle |  |  |  |  |  |  |  |  |  |  |  |
| YJM1326 | XIII | 896145 | 896154 | 896154 | rightend | YMR321C | 99.361 | 313 | 2 | 0 | 6 | 318 | 887195 | 886883 | 2.48E-161 | 568 |
| YJM1326 | XIV | 1 | 10 | 754932 | leftend | YNL332W | 98.436 | 1023 | 16 | 0 | 1 | 1023 | 5367 | 6389 | 0 | 1801 |
| YJM1326 | XIV | 754923 | 754932 | 754932 | rightend | YNR071C | 100 | 1029 | 0 | 0 | 1 | 1029 | 744228 | 743200 | 0 | 1901 |
| YJM1326 | XV | 1 | 10 | 1062841 | leftend | YOL154W | 99.6 | 750 | 3 | 0 | 1 | 750 | 35988 | 36737 | 0 | 1369 |
| YJM1326 | XV | 1059698 | 1059707 | 1062841 | middle |  |  |  |  |  |  |  |  |  |  |  |
| YJM1326 | XV | 1060219 | 1060228 | 1062841 | middle |  |  |  |  |  |  |  |  |  |  |  |
| YJM1326 | XV | 1062832 | 1062841 | 1062841 | rightend | YOR381W | 100 | 2136 | 0 | 0 | 1 | 2136 | 1036397 | 1038532 | 0 | 3945 |
| YJM1326 | XVI | 1 |  | 903962 | leftend_complete | YPL274W | 98.583 | 1764 | 25 | 0 | 1 | 1764 | 15265 | 17028 | 0 | 3120 |
| YJM1326 | XVI | 1777 | 1786 | 903962 | middle |  |  |  |  |  |  |  |  |  |  |  |
| YJM1326 | XVI | 4560 | 4569 | 903962 | middle |  |  |  |  |  |  |  |  |  |  |  |
| YJM1326 | XVI | 903953 | 903962 | 903962 | rightend | YPR201W | 99.671 | 1215 | 4 | 0 | 1 | 1215 | 901226 | 902440 | 0 | 2222 |
| YJM1332 | I | 1 | 10 | 196554 | leftend | YAL063C | 84.804 | 1632 | 243 | 5 | 1 | 1630 | 1737 | 109 | 0 | 1635 |
| YJM1332 | I | 196554 | 196554 | 196554 | rightend_complete | YAR050W | 82.659 | 1632 | 279 | 4 | 1 | 1630 | 171199 | 172828 | 0 | 1443 |
| YJM1332 | II | 1 | 10 | 783237 | leftend | YNCB0001W | 100 | 84 | 0 | 0 | 1 | 84 | 1903 | 1986 | 4.16E-38 | 156 |
| YJM1332 | II | 1165 | 1174 | 783237 | middle |  |  |  |  |  |  |  |  |  |  |  |
| YJM1332 | II | 782668 | 782677 | 783237 | middle |  |  |  |  |  |  |  |  |  |  |  |
| YJM1332 | II | 783228 | 783237 | 783237 | rightend | YBR297W | 99.502 | 1407 | 5 | 1 | 1 | 1407 | 774080 | 775484 | 0 | 2558 |
| YJM1332 | III | 1 |  | 309904 | leftend_complete | YCL073C | 99.758 | 1242 | 3 | 0 | 607 | 1848 | 7437 | 6196 | 0 | 2278 |
| YJM1332 | III | 301368 | 301377 | 309904 | middle |  |  |  |  |  |  |  |  |  |  |  |
| YJM1332 | III | 309443 | 309452 | 309904 | middle |  |  |  |  |  |  |  |  |  |  |  |
| YJM1332 | III | 309904 | 309904 | 309904 | rightend_complete | YCR101C | 99.818 | 549 | 1 | 0 | 1 | 549 | 296676 | 296128 | 0 | 1009 |
| YJM1332 | IV | 1 | 10 | 1474144 | leftend | YDL243C | 95.051 | 990 | 49 | 0 | 1 | 990 | 18325 | 17336 | 0 | 1557 |
| YJM1332 | IV | 1473669 | 1473678 | 1474144 | middle |  |  |  |  |  |  |  |  |  |  |  |
| YJM1332 | IV | 1474135 | 1474144 | 1474144 | rightend | YDR541C | 99.517 | 1035 | 5 | 0 | 1 | 1035 | 1471043 | 1470009 | 0 | 1884 |
| YJM1332 | V | 1 |  | 554144 | leftend_complete | YEL070W | 99.337 | 1509 | 10 | 0 | 1 | 1509 | 15510 | 17018 | 0 | 2732 |
| YJM1332 | V | 554135 | 554144 | 554144 | rightend | YER187W | 91.981 | 424 | 32 | 2 | 4 | 426 | 549690 | 550112 | 3.44E-169 | 593 |
| YJM1332 | VI | 1 | 10 | 233882 | leftend | YFL051C | 98.451 | 452 | 7 | 0 | 1 | 452 | 1227 | 776 | 0 | 797 |
| YJM1332 | VI | 233882 | 233882 | 233882 | rightend_complete | YFR055W | 99.902 | 1022 | 1 | 0 | 1 | 1022 | 227887 | 228908 | 0 | 1882 |
| YJM1332 | VII | 1 | 10 | 1053572 | leftend | YGL258W | 97.585 | 621 | 15 | 0 | 1 | 621 | 4488 | 5108 | 0 | 1064 |
| YJM1332 | VII | 1053563 | 1053572 | 1053572 | rightend | YGR287C | 98.701 | 1770 | 23 | 0 | 1 | 1770 | 1030555 | 1028786 | 0 | 3142 |
| YJM1332 | VIII | 1 |  | 531790 | leftend_complete | YHL044W | 98.446 | 708 | 11 | 0 | 1 | 708 | 9789 | 10496 | 0 | 1247 |
| YJM1332 | VIII | 530714 | 530723 | 531790 | middle |  |  |  |  |  |  |  |  |  |  |  |
| YJM1332 | VIII | 531255 | 531264 | 531790 | middle |  |  |  |  |  |  |  |  |  |  |  |
| YJM1332 | VIII | 531790 | 531790 | 531790 | rightend_complete | YHR211W | 92.041 | 2136 | 146 | 2 | 1096 | 3228 | 516913 | 519027 | 0 | 2981 |
| YJM1332 | IX | 1 |  | 410749 | leftend_complete | YIL169C | 93.344 | 1202 | 78 | 2 | 612 | 1812 | 9344 | 8144 | 0 | 1775 |
| YJM1332 | IX | 410740 | 410749 | 410749 | rightend | YIR039C | 98.947 | 1614 | 17 | 0 | 1 | 1614 | 409485 | 407872 | 0 | 2887 |
| YJM1332 | X | 1 |  | 716517 | leftend_complete | YJL213W | 99.699 | 996 | 3 | 0 | 1 | 996 | 36555 | 37550 | 0 | 1823 |
| YJM1332 | X | 716508 | 716517 | 716517 | rightend | YJR152W | 99.877 | 1632 | 2 | 0 | 1 | 1632 | 706110 | 707741 | 0 | 3003 |
| YJM1332 | XI | 1 | 10 | 667413 | leftend | YKL222C | 99.434 | 2119 | 11 | 1 | 1 | 2118 | 3474 | 1356 | 0 | 3845 |
| YJM1332 | XI | 667413 | 667413 | 667413 | rightend_complete | YKR101W | 99.847 | 1965 | 3 | 0 | 1 | 1965 | 638863 | 640827 | 0 | 3613 |
| YJM1332 | XII | 1 |  | 1683509 | leftend_complete | YLL062C | 99.897 | 975 | 1 | 0 | 1 | 975 | 5186 | 4212 | 0 | 1796 |
| YJM1332 | XII | 981 | 990 | 1683509 | middle |  |  |  |  |  |  |  |  |  |  |  |
| YJM1332 | XII | 1683500 | 1683509 | 1683509 | rightend | YLR460C | 94.071 | 1130 | 65 | 2 | 3 | 1131 | 1681759 | 1680631 | 0 | 1714 |
| YJM1332 | XIII | 1 |  | 910213 | leftend_complete | YML131W | 100 | 1098 | 0 | 0 | 1 | 1098 | 4662 | 5759 | 0 | 2028 |
| YJM1332 | XIII | 910204 | 910213 | 910213 | rightend | YMR321C | 99.361 | 313 | 2 | 0 | 6 | 318 | 887515 | 887203 | 2.48E-161 | 568 |
| YJM1332 | XIV | 1 | 10 | 755653 | leftend | YNL332W | 97.947 | 1023 | 21 | 0 | 1 | 1023 | 5859 | 6881 | 0 | 1773 |
| YJM1332 | XIV | 754803 | 754812 | 755653 | middle |  |  |  |  |  |  |  |  |  |  |  |
| YJM1332 | XIV | 755653 | 755653 | 755653 | rightend_complete | YNR071C | 100 | 1029 | 0 | 0 | 1 | 1029 | 744218 | 743190 | 0 | 1901 |
| YJM1332 | XV | 1 | 10 | 1047394 | leftend | YOL154W | 99.6 | 750 | 3 | 0 | 1 | 750 | 20638 | 21387 | 0 | 1369 |
| YJM1332 | XV | 1047385 | 1047394 | 1047394 | rightend | YOR381W | 99.906 | 2137 | 1 | 1 | 1 | 2136 | 1018748 | 1020884 | 0 | 3934 |
| YJM1332 | XVI | 1 | 10 | 909728 | leftend | YPL274W | 99.943 | 1764 | 1 | 0 | 1 | 1764 | 17884 | 19647 | 0 | 3253 |
| YJM1332 | XVI | 221 | 230 | 909728 | middle |  |  |  |  |  |  |  |  |  |  |  |
| YJM1332 | XVI | 1144 | 1153 | 909728 | middle |  |  |  |  |  |  |  |  |  |  |  |
| YJM1332 | XVI | 3001 | 3010 | 909728 | middle |  |  |  |  |  |  |  |  |  |  |  |
| YJM1332 | XVI | 909719 | 909728 | 909728 | rightend | YPR201W | 100 | 1215 | 0 | 0 | 1 | 1215 | 906676 | 907890 | 0 | 2244 |
| YJM1336 | I | 1 | 10 | 181729 | leftend | YAL063C | 88.34 | 1012 | 114 | 4 | 1 | 1010 | 11266 | 10257 | 0 | 1212 |
| YJM1336 | I | 10010 | 10019 | 181729 | middle |  |  |  |  |  |  |  |  |  |  |  |
| YJM1336 | I | 10247 | 10256 | 181729 | middle |  |  |  |  |  |  |  |  |  |  |  |
| YJM1336 | I | 181720 | 181729 | 181729 | rightend | YAR050W | 84.047 | 934 | 145 | 4 | 1 | 932 | 180781 | 181712 | 0 | 896 |
| YJM1336 | II | 1 | 10 | 783630 | leftend | YNCB0001W | 100 | 84 | 0 | 0 | 1 | 84 | 1278 | 1361 | 4.16E-38 | 156 |
| YJM1336 | II | 796 | 805 | 783630 | middle |  |  |  |  |  |  |  |  |  |  |  |
| YJM1336 | II | 783621 | 783630 | 783630 | rightend | YBR297W | 98.863 | 1407 | 14 | 1 | 1 | 1407 | 773657 | 775061 | 0 | 2508 |
| YJM1336 | III | 1 |  | 300210 | leftend_complete | YCL073C | 99.892 | 1848 | 2 | 0 | 1 | 1848 | 7620 | 5773 | 0 | 3402 |
| YJM1336 | III | 447 | 456 | 300210 | middle |  |  |  |  |  |  |  |  |  |  |  |
| YJM1336 | III | 299835 | 299844 | 300210 | middle |  |  |  |  |  |  |  |  |  |  |  |
| YJM1336 | III | 300210 | 300210 | 300210 | rightend_complete | YCR101C | 99.818 | 549 | 1 | 0 | 1 | 549 | 295717 | 295169 | 0 | 1009 |
| YJM1336 | IV | 1 | 10 | 1462581 | leftend | YDL243C | 95.051 | 990 | 49 | 0 | 1 | 990 | 16922 | 15933 | 0 | 1557 |
| YJM1336 | IV | 1461809 | 1461818 | 1462581 | middle |  |  |  |  |  |  |  |  |  |  |  |
| YJM1336 | IV | 1462572 | 1462581 | 1462581 | rightend | YDR541C | 99.517 | 1035 | 5 | 0 | 1 | 1035 | 1459024 | 1457990 | 0 | 1884 |
| YJM1336 | V | 1 | 10 | 553936 | leftend | YEL070W | 99.271 | 1509 | 11 | 0 | 1 | 1509 | 12823 | 14331 | 0 | 2726 |
| YJM1336 | V | 553927 | 553936 | 553936 | rightend | YER187W | 91.981 | 424 | 32 | 2 | 4 | 426 | 547656 | 548078 | 3.44E-169 | 593 |
| YJM1336 | VI | 1 | 10 | 241600 | leftend | YFL051C | 98.451 | 452 | 7 | 0 | 1 | 452 | 8165 | 7714 | 0 | 797 |
| YJM1336 | VI | 6619 | 6628 | 241600 | middle |  |  |  |  |  |  |  |  |  |  |  |
| YJM1336 | VI | 241591 | 241600 | 241600 | rightend | YFR055W | 99.902 | 1023 | 1 | 0 | 1 | 1023 | 235295 | 236317 | 0 | 1884 |
| YJM1336 | VII | 1 | 10 | 1049747 | leftend | YGL258W | 97.585 | 621 | 15 | 0 | 1 | 621 | 10579 | 11199 | 0 | 1064 |
| YJM1336 | VII | 481 | 490 | 1049747 | middle |  |  |  |  |  |  |  |  |  |  |  |
| YJM1336 | VII | 6089 | 6098 | 1049747 | middle |  |  |  |  |  |  |  |  |  |  |  |
| YJM1336 | VII | 1049738 | 1049747 | 1049747 | rightend | YGR287C | 98.701 | 1770 | 23 | 0 | 1 | 1770 | 1031934 | 1030165 | 0 | 3142 |
| YJM1336 | VIII | 1 | 10 | 537872 | leftend | YHL044W | 98.446 | 708 | 11 | 0 | 1 | 708 | 8199 | 8906 | 0 | 1247 |
| YJM1336 | VIII | 536119 | 536128 | 537872 | middle |  |  |  |  |  |  |  |  |  |  |  |
| YJM1336 | VIII | 536642 | 536651 | 537872 | middle |  |  |  |  |  |  |  |  |  |  |  |
| YJM1336 | VIII | 537359 | 537368 | 537872 | middle |  |  |  |  |  |  |  |  |  |  |  |
| YJM1336 | VIII | 537872 | 537872 | 537872 | rightend_complete | YHR211W | 92.135 | 2136 | 144 | 2 | 1096 | 3228 | 520289 | 522403 | 0 | 2992 |
| YJM1336 | IX | 1 | 10 | 410830 | leftend | YIL169C | 94.747 | 1542 | 79 | 2 | 272 | 1812 | 9395 | 7855 | 0 | 2398 |
| YJM1336 | IX | 1771 | 1780 | 410830 | middle |  |  |  |  |  |  |  |  |  |  |  |
| YJM1336 | IX | 4586 | 4595 | 410830 | middle |  |  |  |  |  |  |  |  |  |  |  |
| YJM1336 | IX | 410821 | 410830 | 410830 | rightend | YIR039C | 99.071 | 1614 | 15 | 0 | 1 | 1614 | 409654 | 408041 | 0 | 2898 |
| YJM1336 | X | 1 | 10 | 716545 | leftend | YJL213W | 99.699 | 996 | 3 | 0 | 1 | 996 | 26286 | 27281 | 0 | 1823 |
| YJM1336 | X | 716545 | 716545 | 716545 | rightend_complete | YJR152W | 99.755 | 1632 | 3 | 1 | 1 | 1632 | 696091 | 697721 | 0 | 2990 |
| YJM1336 | XI | 1 | 10 | 669328 | leftend | YKL222C | 99.386 | 2118 | 13 | 0 | 1 | 2118 | 12903 | 10786 | 0 | 3840 |
| YJM1336 | XI | 9399 | 9408 | 669328 | middle |  |  |  |  |  |  |  |  |  |  |  |
| YJM1336 | XI | 9768 | 9777 | 669328 | middle |  |  |  |  |  |  |  |  |  |  |  |
| YJM1336 | XI | 669319 | 669328 | 669328 | rightend | YKR101W | 99.796 | 1965 | 4 | 0 | 1 | 1965 | 646793 | 648757 | 0 | 3607 |
| YJM1336 | XII | 1 |  | 2229726 | leftend_complete | YLL062C | 100 | 975 | 0 | 0 | 1 | 975 | 5050 | 4076 | 0 | 1801 |
| YJM1336 | XII | 504 | 513 | 2229726 | middle |  |  |  |  |  |  |  |  |  |  |  |
| YJM1336 | XII | 981 | 990 | 2229726 | middle |  |  |  |  |  |  |  |  |  |  |  |
| YJM1336 | XII | 2229717 | 2229726 | 2229726 | rightend | YLR460C | 94.159 | 1130 | 64 | 2 | 3 | 1131 | 2228057 | 2226929 | 0 | 1720 |
| YJM1336 | XIII | 1 | 10 | 903749 | leftend | YML131W | 100 | 1098 | 0 | 0 | 1 | 1098 | 2770 | 3867 | 0 | 2028 |
| YJM1336 | XIII | 463 | 472 | 903749 | middle |  |  |  |  |  |  |  |  |  |  |  |
| YJM1336 | XIII | 903740 | 903749 | 903749 | rightend | YMR321C | 99.361 | 313 | 2 | 0 | 6 | 318 | 888066 | 887754 | 2.48E-161 | 568 |
| YJM1336 | XIV | 1 | 10 | 749436 | leftend | YNL332W | 99.197 | 249 | 2 | 0 | 775 | 1023 | 11 | 259 | 2.67E-125 | 449 |
| YJM1336 | XIV | 748828 | 748837 | 749436 | middle |  |  |  |  |  |  |  |  |  |  |  |
| YJM1336 | XIV | 749427 | 749436 | 749436 | rightend | YNR071C | 99.903 | 1029 | 1 | 0 | 1 | 1029 | 737700 | 736672 | 0 | 1895 |
| YJM1336 | XV | 1 |  | 1059602 | leftend_complete | YOL154W | 99.467 | 750 | 4 | 0 | 1 | 750 | 29057 | 29806 | 0 | 1363 |
| YJM1336 | XV | 1058504 | 1058513 | 1059602 | middle |  |  |  |  |  |  |  |  |  |  |  |
| YJM1336 | XV | 1059009 | 1059018 | 1059602 | middle |  |  |  |  |  |  |  |  |  |  |  |
| YJM1336 | XV | 1059602 | 1059602 | 1059602 | rightend_complete | YOR381W | 99.953 | 2136 | 1 | 0 | 1 | 2136 | 1025728 | 1027863 | 0 | 3940 |
| YJM1336 | XVI | 1 |  | 904506 | leftend_complete | YPL274W | 99.943 | 1764 | 1 | 0 | 1 | 1764 | 15332 | 17095 | 0 | 3253 |
| YJM1336 | XVI | 2467 | 2476 | 904506 | middle |  |  |  |  |  |  |  |  |  |  |  |
| YJM1336 | XVI | 904497 | 904506 | 904506 | rightend | YPR201W | 100 | 1215 | 0 | 0 | 1 | 1215 | 901784 | 902998 | 0 | 2244 |
| YJM1338 | I | 1 |  | 192904 | leftend_complete | YAL063C | 87.454 | 1092 | 133 | 4 | 1 | 1090 | 15425 | 14336 | 0 | 1254 |
| YJM1338 | I | 2720 | 2729 | 192904 | middle |  |  |  |  |  |  |  |  |  |  |  |
| YJM1338 | I | 4259 | 4268 | 192904 | middle |  |  |  |  |  |  |  |  |  |  |  |
| YJM1338 | I | 14165 | 14174 | 192904 | middle |  |  |  |  |  |  |  |  |  |  |  |
| YJM1338 | I | 186152 | 186161 | 192904 | middle |  |  |  |  |  |  |  |  |  |  |  |
| YJM1338 | I | 192895 | 192904 | 192904 | rightend | YAR050W | 82.937 | 1178 | 197 | 4 | 1 | 1176 | 184976 | 186151 | 0 | 1059 |
| YJM1338 | II | 1 |  | 789959 | leftend_complete | YNCB0001W | 100 | 84 | 0 | 0 | 1 | 84 | 10070 | 10153 | 4.16E-38 | 156 |
| YJM1338 | II | 789950 | 789959 | 789959 | rightend | YBR297W | 98.091 | 1205 | 23 | 0 | 1 | 1205 | 783277 | 784481 | 0 | 2098 |
| YJM1338 | III | 1 |  | 306516 | leftend_complete | YCL073C | 99.026 | 1848 | 18 | 0 | 1 | 1848 | 8013 | 6166 | 0 | 3314 |
| YJM1338 | III | 301653 | 301662 | 306516 | middle |  |  |  |  |  |  |  |  |  |  |  |
| YJM1338 | III | 305274 | 305283 | 306516 | middle |  |  |  |  |  |  |  |  |  |  |  |
| YJM1338 | III | 305780 | 305789 | 306516 | middle |  |  |  |  |  |  |  |  |  |  |  |
| YJM1338 | III | 306507 | 306516 | 306516 | rightend | YCR101C | 99.636 | 549 | 2 | 0 | 1 | 549 | 296705 | 296157 | 0 | 1003 |
| YJM1338 | IV | 1 | 10 | 1468499 | leftend | YDL243C | 95.051 | 990 | 49 | 0 | 1 | 990 | 18897 | 17908 | 0 | 1557 |
| YJM1338 | IV | 743 | 752 | 1468499 | middle |  |  |  |  |  |  |  |  |  |  |  |
| YJM1338 | IV | 1736 | 1745 | 1468499 | middle |  |  |  |  |  |  |  |  |  |  |  |
| YJM1338 | IV | 1465759 | 1465768 | 1468499 | middle |  |  |  |  |  |  |  |  |  |  |  |
| YJM1338 | IV | 1468490 | 1468499 | 1468499 | rightend | YDR541C | 99.903 | 1035 | 1 | 0 | 1 | 1035 | 1462029 | 1460995 | 0 | 1906 |
| YJM1338 | V | 1 | 10 | 550585 | leftend | YEL070W | 99.271 | 1509 | 11 | 0 | 1 | 1509 | 13402 | 14910 | 0 | 2726 |
| YJM1338 | V | 550585 | 550585 | 550585 | rightend_complete | YER187W | 99.296 | 426 | 3 | 0 | 1 | 426 | 546451 | 546876 | 0 | 771 |
| YJM1338 | VI | 1 | 10 | 238465 | leftend | YFL051C | 98.23 | 452 | 8 | 0 | 1 | 452 | 1176 | 725 | 0 | 791 |
| YJM1338 | VI | 238456 | 238465 | 238465 | rightend | YFR055W | 99.511 | 1022 | 5 | 0 | 1 | 1022 | 228766 | 229787 | 0 | 1860 |
| YJM1338 | VII | 1 | 10 | 1070648 | leftend | YGL258W | 99.517 | 621 | 3 | 0 | 1 | 621 | 9996 | 10616 | 0 | 1131 |
| YJM1338 | VII | 5945 | 5954 | 1070648 | middle |  |  |  |  |  |  |  |  |  |  |  |
| YJM1338 | VII | 1066798 | 1066807 | 1070648 | middle |  |  |  |  |  |  |  |  |  |  |  |
| YJM1338 | VII | 1069852 | 1069861 | 1070648 | middle |  |  |  |  |  |  |  |  |  |  |  |
| YJM1338 | VII | 1070639 | 1070648 | 1070648 | rightend | YGR287C | 99.831 | 1770 | 3 | 0 | 1 | 1770 | 1034187 | 1032418 | 0 | 3253 |
| YJM1338 | VIII | 1 | 10 | 541580 | leftend | YHL044W | 98.87 | 708 | 8 | 0 | 1 | 708 | 8096 | 8803 | 0 | 1264 |
| YJM1338 | VIII | 535984 | 535993 | 541580 | middle |  |  |  |  |  |  |  |  |  |  |  |
| YJM1338 | VIII | 540360 | 540369 | 541580 | middle |  |  |  |  |  |  |  |  |  |  |  |
| YJM1338 | VIII | 541254 | 541263 | 541580 | middle |  |  |  |  |  |  |  |  |  |  |  |
| YJM1338 | VIII | 541580 | 541580 | 541580 | rightend_complete | YHR211W | 92.416 | 2136 | 138 | 2 | 1096 | 3228 | 519697 | 521811 | 0 | 3025 |
| YJM1338 | IX | 1 |  | 425197 | leftend_complete | YIL169C | 84.633 | 2128 | 278 | 31 | 520 | 2618 | 18055 | 15948 | 0 | 2073 |
| YJM1338 | IX | 424124 | 424133 | 425197 | middle |  |  |  |  |  |  |  |  |  |  |  |
| YJM1338 | IX | 425197 | 425197 | 425197 | rightend_complete | YIR039C | 98.947 | 1614 | 17 | 0 | 1 | 1614 | 417892 | 416279 | 0 | 2887 |
| YJM1338 | X | 1 |  | 709676 | leftend_complete | YJL213W | 99.699 | 996 | 3 | 0 | 1 | 996 | 28873 | 29868 | 0 | 1823 |
| YJM1338 | X | 3296 | 3305 | 709676 | middle |  |  |  |  |  |  |  |  |  |  |  |
| YJM1338 | X | 708623 | 708632 | 709676 | middle |  |  |  |  |  |  |  |  |  |  |  |
| YJM1338 | X | 709676 | 709676 | 709676 | rightend_complete | YJR152W | 99.51 | 1632 | 8 | 0 | 1 | 1632 | 698562 | 700193 | 0 | 2970 |
| YJM1338 | XI | 1 | 10 | 656625 | leftend | YKL222C | 99.433 | 2118 | 12 | 0 | 1 | 2118 | 3165 | 1048 | 0 | 3845 |
| YJM1338 | XI | 656616 | 656625 | 656625 | rightend | YKR101W | 99.542 | 1965 | 9 | 0 | 1 | 1965 | 637750 | 639714 | 0 | 3579 |
| YJM1338 | XII | 1 | 10 | 1574353 | leftend | YLL062C | 99.692 | 975 | 3 | 0 | 1 | 975 | 4465 | 3491 | 0 | 1784 |
| YJM1338 | XII | 1574344 | 1574353 | 1574353 | rightend | YLR460C | 98.585 | 1131 | 16 | 0 | 1 | 1131 | 1573350 | 1572220 | 0 | 2001 |
| YJM1338 | XIII | 1 | 10 | 904981 | leftend | YML131W | 99.545 | 1098 | 5 | 0 | 1 | 1098 | 2806 | 3903 | 0 | 2001 |
| YJM1338 | XIII | 900373 | 900382 | 904981 | middle |  |  |  |  |  |  |  |  |  |  |  |
| YJM1338 | XIII | 901817 | 901826 | 904981 | middle |  |  |  |  |  |  |  |  |  |  |  |
| YJM1338 | XIII | 904981 | 904981 | 904981 | rightend_complete | YMR321C | 99.361 | 313 | 2 | 0 | 6 | 318 | 888840 | 888528 | 2.48E-161 | 568 |
| YJM1338 | XIV | 1 |  | 766542 | leftend_complete | YNL332W | 98.631 | 1023 | 14 | 0 | 1 | 1023 | 7781 | 8803 | 0 | 1812 |
| YJM1338 | XIV | 1062 | 1071 | 766542 | middle |  |  |  |  |  |  |  |  |  |  |  |
| YJM1338 | XIV | 2264 | 2273 | 766542 | middle |  |  |  |  |  |  |  |  |  |  |  |
| YJM1338 | XIV | 766533 | 766542 | 766542 | rightend | YNR071C | 99.611 | 1029 | 4 | 0 | 1 | 1029 | 746191 | 745163 | 0 | 1879 |
| YJM1338 | XV | 1 |  | 1036000 | leftend_complete | YOL154W | 99.467 | 750 | 4 | 0 | 1 | 750 | 9152 | 9901 | 0 | 1363 |
| YJM1338 | XV | 1034001 | 1034010 | 1036000 | middle |  |  |  |  |  |  |  |  |  |  |  |
| YJM1338 | XV | 1035231 | 1035240 | 1036000 | middle |  |  |  |  |  |  |  |  |  |  |  |
| YJM1338 | XV | 1035491 | 1035500 | 1036000 | middle |  |  |  |  |  |  |  |  |  |  |  |
| YJM1338 | XV | 1035991 | 1036000 | 1036000 | rightend | YOR381W | 100 | 2136 | 0 | 0 | 1 | 2136 | 1005440 | 1007575 | 0 | 3945 |
| YJM1338 | XVI | 1 | 10 | 902795 | leftend | YPL274W | 99.376 | 1764 | 11 | 0 | 1 | 1764 | 12958 | 14721 | 0 | 3197 |
| YJM1338 | XVI | 902795 | 902795 | 902795 | rightend_complete | YPR201W | 99.506 | 1215 | 6 | 0 | 1 | 1215 | 899517 | 900731 | 0 | 2211 |
| YJM1341 | I | 1 | 10 | 198769 | leftend | YAL063C | 95.02 | 1004 | 23 | 11 | 2966 | 3969 | 178743 | 179719 | 0 | 1552 |
| YJM1341 | I | 5894 | 5903 | 198769 | middle |  |  |  |  |  |  |  |  |  |  |  |
| YJM1341 | I | 198760 | 198769 | 198769 | rightend | YAR050W | 82.414 | 1632 | 283 | 4 | 1 | 1630 | 176387 | 178016 | 0 | 1421 |
| YJM1341 | II | 1 | 10 | 787593 | leftend | YNCB0001W | 100 | 84 | 0 | 0 | 1 | 84 | 6531 | 6614 | 4.16E-38 | 156 |
| YJM1341 | II | 1438 | 1447 | 787593 | middle |  |  |  |  |  |  |  |  |  |  |  |
| YJM1341 | II | 2925 | 2934 | 787593 | middle |  |  |  |  |  |  |  |  |  |  |  |
| YJM1341 | II | 4109 | 4118 | 787593 | middle |  |  |  |  |  |  |  |  |  |  |  |
| YJM1341 | II | 4874 | 4883 | 787593 | middle |  |  |  |  |  |  |  |  |  |  |  |
| YJM1341 | II | 787584 | 787593 | 787593 | rightend | YBR297W | 91.961 | 1418 | 98 | 12 | 1 | 1407 | 778989 | 780401 | 0 | 1973 |
| YJM1341 | III | 1 |  | 309163 | leftend_complete | YCL073C | 99.838 | 1848 | 3 | 0 | 1 | 1848 | 8049 | 6202 | 0 | 3397 |
| YJM1341 | III | 309154 | 309163 | 309163 | rightend | YCR101C | 99.818 | 549 | 1 | 0 | 1 | 549 | 296124 | 295576 | 0 | 1009 |
| YJM1341 | IV | 1 | 10 | 1496458 | leftend | YDL243C | 100 | 990 | 0 | 0 | 1 | 990 | 18966 | 17977 | 0 | 1829 |
| YJM1341 | IV | 642 | 651 | 1496458 | middle |  |  |  |  |  |  |  |  |  |  |  |
| YJM1341 | IV | 1496449 | 1496458 | 1496458 | rightend | YDR541C | 99.42 | 1035 | 6 | 0 | 1 | 1035 | 1493626 | 1492592 | 0 | 1879 |
| YJM1341 | V | 1 | 10 | 550745 | leftend | YEL070W | 99.205 | 1509 | 12 | 0 | 1 | 1509 | 13229 | 14737 | 0 | 2721 |
| YJM1341 | V | 549634 | 549643 | 550745 | middle |  |  |  |  |  |  |  |  |  |  |  |
| YJM1341 | V | 550745 | 550745 | 550745 | rightend_complete | YER187W | 91.745 | 424 | 33 | 2 | 4 | 426 | 547071 | 547493 | 1.6E-167 | 588 |
| YJM1341 | VI | 1 | 10 | 233851 | leftend | YFL051C | 98.451 | 452 | 7 | 0 | 1 | 452 | 1092 | 641 | 0 | 797 |
| YJM1341 | VI | 233842 | 233851 | 233851 | rightend | YFR055W | 100 | 1023 | 0 | 0 | 1 | 1023 | 227414 | 228436 | 0 | 1890 |
| YJM1341 | VII | 1 | 10 | 1039115 | leftend | YGL258W | 97.585 | 621 | 15 | 0 | 1 | 621 | 11974 | 12594 | 0 | 1064 |
| YJM1341 | VII | 1037632 | 1037641 | 1039115 | middle |  |  |  |  |  |  |  |  |  |  |  |
| YJM1341 | VII | 1039106 | 1039115 | 1039115 | rightend | YGR287C | 98.893 | 1626 | 18 | 0 | 145 | 1770 | 1032128 | 1030503 | 0 | 2904 |
| YJM1341 | VIII | 1 | 10 | 519862 | leftend | YHL044W | 98.446 | 708 | 11 | 0 | 1 | 708 | 1650 | 2357 | 0 | 1247 |
| YJM1341 | VIII | 512923 | 512932 | 519862 | middle |  |  |  |  |  |  |  |  |  |  |  |
| YJM1341 | VIII | 518816 | 518825 | 519862 | middle |  |  |  |  |  |  |  |  |  |  |  |
| YJM1341 | VIII | 519862 | 519862 | 519862 | rightend_complete | YHR211W | 82.535 | 939 | 153 | 7 | 1 | 932 | 509629 | 510563 | 0 | 815 |
| YJM1341 | IX | 1 | 10 | 392791 | leftend | YIL162W | 99.062 | 1599 | 15 | 0 | 1 | 1599 | 2451 | 4049 | 0 | 2870 |
| YJM1341 | IX | 392309 | 392318 | 392791 | middle |  |  |  |  |  |  |  |  |  |  |  |
| YJM1341 | IX | 392791 | 392791 | 392791 | rightend_complete | YIR039C | 99.071 | 1614 | 15 | 0 | 1 | 1614 | 390678 | 389065 | 0 | 2898 |
| YJM1341 | X | 1 | 10 | 716839 | leftend | YJL213W | 99.699 | 996 | 3 | 0 | 1 | 996 | 26312 | 27307 | 0 | 1823 |
| YJM1341 | X | 716839 | 716839 | 716839 | rightend_complete | YJR152W | 99.816 | 1632 | 3 | 0 | 1 | 1632 | 696581 | 698212 | 0 | 2998 |
| YJM1341 | XI | 1 | 10 | 658521 | leftend | YKL222C | 99.953 | 2118 | 1 | 0 | 1 | 2118 | 4076 | 1959 | 0 | 3906 |
| YJM1341 | XI | 569 | 578 | 658521 | middle |  |  |  |  |  |  |  |  |  |  |  |
| YJM1341 | XI | 657759 | 657768 | 658521 | middle |  |  |  |  |  |  |  |  |  |  |  |
| YJM1341 | XI | 657988 | 657997 | 658521 | middle |  |  |  |  |  |  |  |  |  |  |  |
| YJM1341 | XI | 658521 | 658521 | 658521 | rightend_complete | YKR101W | 99.847 | 1965 | 3 | 0 | 1 | 1965 | 637956 | 639920 | 0 | 3613 |
| YJM1341 | XII | 1 | 10 | 1884752 | leftend | YLL062C | 100 | 975 | 0 | 0 | 1 | 975 | 4208 | 3234 | 0 | 1801 |
| YJM1341 | XII | 1884087 | 1884096 | 1884752 | middle |  |  |  |  |  |  |  |  |  |  |  |
| YJM1341 | XII | 1884743 | 1884752 | 1884752 | rightend | YLR460C | 93.982 | 1130 | 66 | 2 | 3 | 1131 | 1881535 | 1880407 | 0 | 1709 |
| YJM1341 | XIII | 1 |  | 896699 | leftend_complete | YML131W | 99.909 | 1098 | 1 | 0 | 1 | 1098 | 4071 | 5168 | 0 | 2023 |
| YJM1341 | XIII | 252 | 261 | 896699 | middle |  |  |  |  |  |  |  |  |  |  |  |
| YJM1341 | XIII | 1047 | 1056 | 896699 | middle |  |  |  |  |  |  |  |  |  |  |  |
| YJM1341 | XIII | 1252 | 1261 | 896699 | middle |  |  |  |  |  |  |  |  |  |  |  |
| YJM1341 | XIII | 1560 | 1569 | 896699 | middle |  |  |  |  |  |  |  |  |  |  |  |
| YJM1341 | XIII | 896690 | 896699 | 896699 | rightend | YMR321C | 99.361 | 313 | 2 | 0 | 6 | 318 | 888901 | 888589 | 2.48E-161 | 568 |
| YJM1341 | XIV | 1 | 10 | 753941 | leftend | YNL332W | 98.631 | 1023 | 14 | 0 | 1 | 1023 | 5110 | 6132 | 0 | 1812 |
| YJM1341 | XIV | 753932 | 753941 | 753941 | rightend | YNR071C | 100 | 1029 | 0 | 0 | 1 | 1029 | 743854 | 742826 | 0 | 1901 |
| YJM1341 | XV | 1 |  | 1087200 | leftend_complete | YOL154W | 99.467 | 750 | 4 | 0 | 1 | 750 | 23089 | 23838 | 0 | 1363 |
| YJM1341 | XV | 1087191 | 1087200 | 1087200 | rightend | YOR381W | 100 | 2136 | 0 | 0 | 1 | 2136 | 1020725 | 1022860 | 0 | 3945 |
| YJM1341 | XVI | 1 |  | 917745 | leftend_complete | YPL274W | 99.83 | 1764 | 3 | 0 | 1 | 1764 | 24816 | 26579 | 0 | 3241 |
| YJM1341 | XVI | 2404 | 2413 | 917745 | middle |  |  |  |  |  |  |  |  |  |  |  |
| YJM1341 | XVI | 6367 | 6376 | 917745 | middle |  |  |  |  |  |  |  |  |  |  |  |
| YJM1341 | XVI | 6875 | 6884 | 917745 | middle |  |  |  |  |  |  |  |  |  |  |  |
| YJM1341 | XVI | 7766 | 7775 | 917745 | middle |  |  |  |  |  |  |  |  |  |  |  |
| YJM1341 | XVI | 916914 | 916923 | 917745 | middle |  |  |  |  |  |  |  |  |  |  |  |
| YJM1341 | XVI | 917736 | 917745 | 917745 | rightend | YPR201W | 100 | 1215 | 0 | 0 | 1 | 1215 | 913920 | 915134 | 0 | 2244 |
| YJM1342 | I | 1 | 10 | 202731 | leftend | YAL063C | 92.777 | 983 | 23 | 8 | 3014 | 3969 | 193557 | 194518 | 0 | 1378 |
| YJM1342 | I | 701 | 710 | 202731 | middle |  |  |  |  |  |  |  |  |  |  |  |
| YJM1342 | I | 1726 | 1735 | 202731 | middle |  |  |  |  |  |  |  |  |  |  |  |
| YJM1342 | I | 2474 | 2483 | 202731 | middle |  |  |  |  |  |  |  |  |  |  |  |
| YJM1342 | I | 3352 | 3361 | 202731 | middle |  |  |  |  |  |  |  |  |  |  |  |
| YJM1342 | I | 13733 | 13742 | 202731 | middle |  |  |  |  |  |  |  |  |  |  |  |
| YJM1342 | I | 192879 | 192888 | 202731 | middle |  |  |  |  |  |  |  |  |  |  |  |
| YJM1342 | I | 202722 | 202731 | 202731 | rightend | YAR050W | 97.089 | 962 | 28 | 0 | 3653 | 4614 | 193557 | 194518 | 0 | 1622 |
| YJM1342 | II | 1 | 10 | 780359 | leftend | YNCB0001W | 100 | 84 | 0 | 0 | 1 | 84 | 1797 | 1880 | 4.16E-38 | 156 |
| YJM1342 | II | 780350 | 780359 | 780359 | rightend | YBR297W | 98.174 | 1205 | 22 | 0 | 1 | 1205 | 776163 | 777367 | 0 | 2104 |
| YJM1342 | III | 1 |  | 311834 | leftend_complete | YCL073C | 98.972 | 1848 | 19 | 0 | 1 | 1848 | 8088 | 6241 | 0 | 3308 |
| YJM1342 | III | 311825 | 311834 | 311834 | rightend | YCR101C | 98.364 | 550 | 8 | 1 | 1 | 549 | 297869 | 297320 | 0 | 965 |
| YJM1342 | IV | 1 | 10 | 1452138 | leftend | YDL243C | 85.903 | 986 | 139 | 0 | 1 | 986 | 3511 | 2526 | 0 | 1051 |
| YJM1342 | IV | 1450285 | 1450294 | 1452138 | middle |  |  |  |  |  |  |  |  |  |  |  |
| YJM1342 | IV | 1450915 | 1450924 | 1452138 | middle |  |  |  |  |  |  |  |  |  |  |  |
| YJM1342 | IV | 1452129 | 1452138 | 1452138 | rightend | YDR541C | 99.903 | 1035 | 1 | 0 | 1 | 1035 | 1447463 | 1446429 | 0 | 1906 |
| YJM1342 | V | 1 | 10 | 562785 | leftend | YEL070W | 99.337 | 1509 | 10 | 0 | 1 | 1509 | 13212 | 14720 | 0 | 2732 |
| YJM1342 | V | 558507 | 558516 | 562785 | middle |  |  |  |  |  |  |  |  |  |  |  |
| YJM1342 | V | 562785 | 562785 | 562785 | rightend_complete | YER187W | 97.653 | 426 | 10 | 0 | 1 | 426 | 547293 | 547718 | 0 | 732 |
| YJM1342 | VI | 1 | 10 | 291442 | leftend | YFL051C | 99.779 | 452 | 1 | 0 | 1 | 452 | 52421 | 51970 | 0 | 830 |
| YJM1342 | VI | 291433 | 291442 | 291442 | rightend | YFR055W | 99.609 | 1022 | 4 | 0 | 1 | 1022 | 281316 | 282337 | 0 | 1866 |
| YJM1342 | VII | 1 | 10 | 1054471 | leftend | YGL258W | 97.746 | 621 | 14 | 0 | 1 | 621 | 17824 | 18444 | 0 | 1070 |
| YJM1342 | VII | 6487 | 6496 | 1054471 | middle |  |  |  |  |  |  |  |  |  |  |  |
| YJM1342 | VII | 1054462 | 1054471 | 1054471 | rightend | YGR287C | 91.479 | 1772 | 147 | 4 | 1 | 1770 | 1041564 | 1039795 | 0 | 2433 |
| YJM1342 | VIII | 1 | 10 | 536278 | leftend | YHL044W | 98.729 | 708 | 9 | 0 | 1 | 708 | 7569 | 8276 | 0 | 1258 |
| YJM1342 | VIII | 514513 | 514522 | 536278 | middle |  |  |  |  |  |  |  |  |  |  |  |
| YJM1342 | VIII | 536269 | 536278 | 536278 | rightend | YHR211W | 83.087 | 1218 | 195 | 7 | 1 | 1211 | 513276 | 514489 | 0 | 1098 |
| YJM1342 | IX | 1 |  | 413983 | leftend_complete | YIL169C | 96.417 | 1535 | 40 | 6 | 394 | 1919 | 7964 | 6436 | 0 | 2516 |
| YJM1342 | IX | 2126 | 2135 | 413983 | middle |  |  |  |  |  |  |  |  |  |  |  |
| YJM1342 | IX | 3068 | 3077 | 413983 | middle |  |  |  |  |  |  |  |  |  |  |  |
| YJM1342 | IX | 413974 | 413983 | 413983 | rightend | YIR039C | 99.814 | 1614 | 3 | 0 | 1 | 1614 | 407497 | 405884 | 0 | 2964 |
| YJM1342 | X | 1 |  | 712566 | leftend_complete | YJL213W | 99.398 | 996 | 6 | 0 | 1 | 996 | 16948 | 17943 | 0 | 1807 |
| YJM1342 | X | 1038 | 1047 | 712566 | middle |  |  |  |  |  |  |  |  |  |  |  |
| YJM1342 | X | 2033 | 2042 | 712566 | middle |  |  |  |  |  |  |  |  |  |  |  |
| YJM1342 | X | 2830 | 2839 | 712566 | middle |  |  |  |  |  |  |  |  |  |  |  |
| YJM1342 | X | 712566 | 712566 | 712566 | rightend_complete | YJR152W | 99.51 | 1632 | 8 | 0 | 1 | 1632 | 688556 | 690187 | 0 | 2970 |
| YJM1342 | XI | 1 | 10 | 670041 | leftend | YKL222C | 99.575 | 2118 | 9 | 0 | 1 | 2118 | 4370 | 2253 | 0 | 3862 |
| YJM1342 | XI | 670032 | 670041 | 670041 | rightend | YKR101W | 99.44 | 1965 | 11 | 0 | 1 | 1965 | 641345 | 643309 | 0 | 3568 |
| YJM1342 | XII | 1 | 10 | 1901633 | leftend | YLL062C | 99.692 | 975 | 3 | 0 | 1 | 975 | 4417 | 3443 | 0 | 1784 |
| YJM1342 | XII | 1899858 | 1899867 | 1901633 | middle |  |  |  |  |  |  |  |  |  |  |  |
| YJM1342 | XII | 1900091 | 1900100 | 1901633 | middle |  |  |  |  |  |  |  |  |  |  |  |
| YJM1342 | XII | 1901633 | 1901633 | 1901633 | rightend_complete | YLR460C | 93.993 | 1132 | 66 | 2 | 1 | 1131 | 1898104 | 1896974 | 0 | 1712 |
| YJM1342 | XIII | 1 | 10 | 918622 | leftend | YML131W | 99.362 | 1098 | 7 | 0 | 1 | 1098 | 4082 | 5179 | 0 | 1989 |
| YJM1342 | XIII | 916089 | 916098 | 918622 | middle |  |  |  |  |  |  |  |  |  |  |  |
| YJM1342 | XIII | 918613 | 918622 | 918622 | rightend | YMR321C | 98.403 | 313 | 5 | 0 | 6 | 318 | 889090 | 888778 | 2.5E-156 | 551 |
| YJM1342 | XIV | 1 | 10 | 750429 | leftend | YNL332W | 98.534 | 1023 | 15 | 0 | 1 | 1023 | 189 | 1211 | 0 | 1807 |
| YJM1342 | XIV | 750420 | 750429 | 750429 | rightend | YNR071C | 99.611 | 1029 | 4 | 0 | 1 | 1029 | 739738 | 738710 | 0 | 1879 |
| YJM1342 | XV | 1 | 10 | 1068470 | leftend | YOL154W | 99.6 | 750 | 3 | 0 | 1 | 750 | 39840 | 40589 | 0 | 1369 |
| YJM1342 | XV | 1066316 | 1066325 | 1068470 | middle |  |  |  |  |  |  |  |  |  |  |  |
| YJM1342 | XV | 1066750 | 1066759 | 1068470 | middle |  |  |  |  |  |  |  |  |  |  |  |
| YJM1342 | XV | 1068461 | 1068470 | 1068470 | rightend | YOR381W | 97.801 | 2137 | 45 | 2 | 1 | 2136 | 1039045 | 1041180 | 0 | 3685 |
| YJM1342 | XVI | 1 | 10 | 912017 | leftend | YPL274W | 98.129 | 1764 | 33 | 0 | 1 | 1764 | 16647 | 18410 | 0 | 3075 |
| YJM1342 | XVI | 1697 | 1706 | 912017 | middle |  |  |  |  |  |  |  |  |  |  |  |
| YJM1342 | XVI | 2058 | 2067 | 912017 | middle |  |  |  |  |  |  |  |  |  |  |  |
| YJM1342 | XVI | 912017 | 912017 | 912017 | rightend_complete | YPR201W | 99.424 | 1215 | 6 | 1 | 1 | 1215 | 908753 | 909966 | 0 | 2204 |
| YJM1355 | I | 1 | 10 | 200807 | leftend | YAL063C | 91.483 | 998 | 56 | 13 | 2966 | 3962 | 13725 | 12756 | 0 | 1345 |
| YJM1355 | I | 3244 | 3253 | 200807 | middle |  |  |  |  |  |  |  |  |  |  |  |
| YJM1355 | I | 14592 | 14601 | 200807 | middle |  |  |  |  |  |  |  |  |  |  |  |
| YJM1355 | I | 186810 | 186819 | 200807 | middle |  |  |  |  |  |  |  |  |  |  |  |
| YJM1355 | I | 200807 | 200807 | 200807 | rightend_complete | YAR050W | 92.687 | 670 | 28 | 1 | 3605 | 4253 | 13725 | 13056 | 0 | 946 |
| YJM1355 | II | 1 | 10 | 793333 | leftend | YNCB0001W | 100 | 84 | 0 | 0 | 1 | 84 | 10321 | 10404 | 4.16E-38 | 156 |
| YJM1355 | II | 793333 | 793333 | 793333 | rightend_complete | YBR297W | 84.408 | 1334 | 204 | 3 | 1 | 1332 | 784579 | 785910 | 0 | 1308 |
| YJM1355 | III | 1 | 10 | 306241 | leftend | YCL073C | 99.784 | 1848 | 4 | 0 | 1 | 1848 | 11598 | 9751 | 0 | 3391 |
| YJM1355 | III | 1153 | 1162 | 306241 | middle |  |  |  |  |  |  |  |  |  |  |  |
| YJM1355 | III | 305774 | 305783 | 306241 | middle |  |  |  |  |  |  |  |  |  |  |  |
| YJM1355 | III | 306241 | 306241 | 306241 | rightend_complete | YCR101C | 97.636 | 550 | 12 | 1 | 1 | 549 | 300730 | 300181 | 0 | 942 |
| YJM1355 | IV | 1 | 10 | 1807692 | leftend | YKL222C | 99.481 | 2118 | 11 | 0 | 1 | 2118 | 17368 | 15251 | 0 | 3851 |
| YJM1355 | IV | 819 | 828 | 1807692 | middle |  |  |  |  |  |  |  |  |  |  |  |
| YJM1355 | IV | 1806833 | 1806842 | 1807692 | middle |  |  |  |  |  |  |  |  |  |  |  |
| YJM1355 | IV | 1807683 | 1807692 | 1807692 | rightend | YDR541C | 99.517 | 1035 | 5 | 0 | 1 | 1035 | 1804111 | 1803077 | 0 | 1884 |
| YJM1355 | V | 1 | 10 | 564684 | leftend | YEL070W | 99.139 | 1509 | 13 | 0 | 1 | 1509 | 14421 | 15929 | 0 | 2715 |
| YJM1355 | V | 838 | 847 | 564684 | middle |  |  |  |  |  |  |  |  |  |  |  |
| YJM1355 | V | 563957 | 563966 | 564684 | middle |  |  |  |  |  |  |  |  |  |  |  |
| YJM1355 | V | 564675 | 564684 | 564684 | rightend | YER187W | 99.765 | 426 | 1 | 0 | 1 | 426 | 547882 | 548307 | 0 | 782 |
| YJM1355 | VI | 1 |  | 256359 | leftend_complete | YFL051C | 100 | 483 | 0 | 0 | 1 | 483 | 21980 | 21498 | 0 | 893 |
| YJM1355 | VI | 2784 | 2793 | 256359 | middle |  |  |  |  |  |  |  |  |  |  |  |
| YJM1355 | VI | 4242 | 4251 | 256359 | middle |  |  |  |  |  |  |  |  |  |  |  |
| YJM1355 | VI | 256350 | 256359 | 256359 | rightend | YFR055W | 100 | 1023 | 0 | 0 | 1 | 1023 | 249895 | 250917 | 0 | 1890 |
| YJM1355 | VII | 1 | 10 | 1036919 | leftend | YGL258W | 97.585 | 621 | 15 | 0 | 1 | 621 | 3922 | 4542 | 0 | 1064 |
| YJM1355 | VII | 1036910 | 1036919 | 1036919 | rightend | YGR287C | 98.588 | 1770 | 25 | 0 | 1 | 1770 | 1025521 | 1023752 | 0 | 3131 |
| YJM1355 | VIII | 1 | 10 | 532456 | leftend | YHL044W | 98.446 | 708 | 11 | 0 | 1 | 708 | 17415 | 18122 | 0 | 1247 |
| YJM1355 | VIII | 530732 | 530741 | 532456 | middle |  |  |  |  |  |  |  |  |  |  |  |
| YJM1355 | VIII | 531500 | 531509 | 532456 | middle |  |  |  |  |  |  |  |  |  |  |  |
| YJM1355 | VIII | 532447 | 532456 | 532456 | rightend | YHR211W | 87.845 | 1086 | 111 | 4 | 1231 | 2316 | 523970 | 525034 | 0 | 1254 |
| YJM1355 | IX | 1 | 10 | 437061 | leftend | YIL169C | 83.885 | 2116 | 271 | 38 | 562 | 2618 | 32039 | 29935 | 0 | 1954 |
| YJM1355 | IX | 437052 | 437061 | 437061 | rightend | YIR039C | 99.009 | 1614 | 16 | 0 | 1 | 1614 | 430826 | 429213 | 0 | 2892 |
| YJM1355 | X | 1 | 10 | 695767 | leftend | YJL213W | 99.699 | 996 | 3 | 0 | 1 | 996 | 11547 | 12542 | 0 | 1823 |
| YJM1355 | X | 486 | 495 | 695767 | middle |  |  |  |  |  |  |  |  |  |  |  |
| YJM1355 | X | 694631 | 694640 | 695767 | middle |  |  |  |  |  |  |  |  |  |  |  |
| YJM1355 | X | 695758 | 695767 | 695767 | rightend | YJR152W | 99.877 | 1632 | 2 | 0 | 1 | 1632 | 682185 | 683816 | 0 | 3003 |
| YJM1355 | XI | 1 | 10 | 408743 | leftend | YKL222C | 99.764 | 2118 | 5 | 0 | 1 | 2118 | 5196 | 3079 | 0 | 3884 |
| YJM1355 | XI | 408175 | 408184 | 408743 | middle |  |  |  |  |  |  |  |  |  |  |  |
| YJM1355 | XI | 408743 | 408743 | 408743 | rightend_complete | YKR101W | 99.847 | 1965 | 3 | 0 | 1 | 1965 | 386822 | 388786 | 0 | 3613 |
| YJM1355 | XII | 1 | 10 | 2159678 | leftend | YLL062C | 99.897 | 975 | 1 | 0 | 1 | 975 | 9387 | 8413 | 0 | 1796 |
| YJM1355 | XII | 1605 | 1614 | 2159678 | middle |  |  |  |  |  |  |  |  |  |  |  |
| YJM1355 | XII | 2761 | 2770 | 2159678 | middle |  |  |  |  |  |  |  |  |  |  |  |
| YJM1355 | XII | 3485 | 3494 | 2159678 | middle |  |  |  |  |  |  |  |  |  |  |  |
| YJM1355 | XII | 4238 | 4247 | 2159678 | middle |  |  |  |  |  |  |  |  |  |  |  |
| YJM1355 | XII | 4937 | 4946 | 2159678 | middle |  |  |  |  |  |  |  |  |  |  |  |
| YJM1355 | XII | 2156144 | 2156153 | 2159678 | middle |  |  |  |  |  |  |  |  |  |  |  |
| YJM1355 | XII | 2157168 | 2157177 | 2159678 | middle |  |  |  |  |  |  |  |  |  |  |  |
| YJM1355 | XII | 2157826 | 2157835 | 2159678 | middle |  |  |  |  |  |  |  |  |  |  |  |
| YJM1355 | XII | 2159678 | 2159678 | 2159678 | rightend_complete | YLR460C | 94.159 | 1130 | 64 | 2 | 3 | 1131 | 2155199 | 2154071 | 0 | 1720 |
| YJM1355 | XIII | 1 | 10 | 903180 | leftend | YML131W | 99.909 | 1098 | 1 | 0 | 1 | 1098 | 3333 | 4430 | 0 | 2023 |
| YJM1355 | XIII | 443 | 452 | 903180 | middle |  |  |  |  |  |  |  |  |  |  |  |
| YJM1355 | XIII | 903171 | 903180 | 903180 | rightend | YMR321C | 99.361 | 313 | 2 | 0 | 6 | 318 | 888727 | 888415 | 2.48E-161 | 568 |
| YJM1355 | XIV | 1 |  | 768602 | leftend_complete | YNL332W | 98.925 | 1023 | 11 | 0 | 1 | 1023 | 8714 | 9736 | 0 | 1829 |
| YJM1355 | XIV | 1752 | 1761 | 768602 | middle |  |  |  |  |  |  |  |  |  |  |  |
| YJM1355 | XIV | 768157 | 768166 | 768602 | middle |  |  |  |  |  |  |  |  |  |  |  |
| YJM1355 | XIV | 768593 | 768602 | 768602 | rightend | YNR071C | 99.514 | 1029 | 5 | 0 | 1 | 1029 | 748623 | 747595 | 0 | 1873 |
| YJM1355 | XV | 1 |  | 1061198 | leftend_complete | YOL154W | 99.467 | 750 | 4 | 0 | 1 | 750 | 26916 | 27665 | 0 | 1363 |
| YJM1355 | XV | 1061189 | 1061198 | 1061198 | rightend | YOR381W | 99.906 | 2137 | 1 | 1 | 1 | 2136 | 1025830 | 1027966 | 0 | 3934 |
| YJM1355 | XVI | 1 | 10 | 898738 | leftend | YPL274W | 99.433 | 1764 | 9 | 1 | 1 | 1764 | 7454 | 9216 | 0 | 3201 |
| YJM1355 | XVI | 897228 | 897237 | 898738 | middle |  |  |  |  |  |  |  |  |  |  |  |
| YJM1355 | XVI | 898729 | 898738 | 898738 | rightend | YPR201W | 100 | 1215 | 0 | 0 | 1 | 1215 | 894215 | 895429 | 0 | 2244 |
| YJM1356 | I | 1 | 10 | 186466 | leftend | YAL063C | 85.4 | 1500 | 212 | 5 | 1 | 1495 | 12627 | 11130 | 0 | 1550 |
| YJM1356 | I | 186457 | 186466 | 186466 | rightend | YAR050W | 79.944 | 1426 | 277 | 7 | 76 | 1495 | 12552 | 11130 | 0 | 1040 |
| YJM1356 | II | 1 | 10 | 786459 | leftend | YNCB0001W | 100 | 84 | 0 | 0 | 1 | 84 | 3528 | 3611 | 4.16E-38 | 156 |
| YJM1356 | II | 364 | 373 | 786459 | middle |  |  |  |  |  |  |  |  |  |  |  |
| YJM1356 | II | 1939 | 1948 | 786459 | middle |  |  |  |  |  |  |  |  |  |  |  |
| YJM1356 | II | 3036 | 3045 | 786459 | middle |  |  |  |  |  |  |  |  |  |  |  |
| YJM1356 | II | 785554 | 785563 | 786459 | middle |  |  |  |  |  |  |  |  |  |  |  |
| YJM1356 | II | 786450 | 786459 | 786459 | rightend | YBR297W | 98.792 | 1407 | 15 | 1 | 1 | 1407 | 775872 | 777276 | 0 | 2503 |
| YJM1356 | III | 1 |  | 301246 | leftend_complete | YCL073C | 99.838 | 1848 | 3 | 0 | 1 | 1848 | 8102 | 6255 | 0 | 3397 |
| YJM1356 | III | 301237 | 301246 | 301246 | rightend | YCR101C | 99.636 | 549 | 2 | 0 | 1 | 549 | 296721 | 296173 | 0 | 1003 |
| YJM1356 | IV | 1 | 10 | 1468104 | leftend | YDL243C | 95.051 | 990 | 49 | 0 | 1 | 990 | 16921 | 15932 | 0 | 1557 |
| YJM1356 | IV | 1468095 | 1468104 | 1468104 | rightend | YDR541C | 99.324 | 1035 | 7 | 0 | 1 | 1035 | 1465256 | 1464222 | 0 | 1873 |
| YJM1356 | V | 1 | 10 | 558974 | leftend | YEL070W | 98.807 | 1509 | 17 | 1 | 1 | 1509 | 13409 | 14916 | 0 | 2686 |
| YJM1356 | V | 557097 | 557106 | 558974 | middle |  |  |  |  |  |  |  |  |  |  |  |
| YJM1356 | V | 558643 | 558652 | 558974 | middle |  |  |  |  |  |  |  |  |  |  |  |
| YJM1356 | V | 558965 | 558974 | 558974 | rightend | YER187W | 91.981 | 424 | 32 | 2 | 4 | 426 | 548292 | 548714 | 3.44E-169 | 593 |
| YJM1356 | VI | 1 | 10 | 236131 | leftend | YFL051C | 98.451 | 452 | 7 | 0 | 1 | 452 | 2652 | 2201 | 0 | 797 |
| YJM1356 | VI | 1304 | 1313 | 236131 | middle |  |  |  |  |  |  |  |  |  |  |  |
| YJM1356 | VI | 1561 | 1570 | 236131 | middle |  |  |  |  |  |  |  |  |  |  |  |
| YJM1356 | VI | 236131 | 236131 | 236131 | rightend_complete | YFR055W | 100 | 1023 | 0 | 0 | 1 | 1023 | 230106 | 231128 | 0 | 1890 |
| YJM1356 | VII | 1 | 10 | 1050266 | leftend | YGL258W | 97.585 | 621 | 15 | 0 | 1 | 621 | 10261 | 10881 | 0 | 1064 |
| YJM1356 | VII | 5881 | 5890 | 1050266 | middle |  |  |  |  |  |  |  |  |  |  |  |
| YJM1356 | VII | 1050257 | 1050266 | 1050266 | rightend | YGR287C | 98.588 | 1770 | 25 | 0 | 1 | 1770 | 1030781 | 1029012 | 0 | 3131 |
| YJM1356 | VIII | 1 | 10 | 532860 | leftend | YHL044W | 98.446 | 708 | 11 | 0 | 1 | 708 | 9201 | 9908 | 0 | 1247 |
| YJM1356 | VIII | 6904 | 6913 | 532860 | middle |  |  |  |  |  |  |  |  |  |  |  |
| YJM1356 | VIII | 7823 | 7832 | 532860 | middle |  |  |  |  |  |  |  |  |  |  |  |
| YJM1356 | VIII | 531578 | 531587 | 532860 | middle |  |  |  |  |  |  |  |  |  |  |  |
| YJM1356 | VIII | 532192 | 532201 | 532860 | middle |  |  |  |  |  |  |  |  |  |  |  |
| YJM1356 | VIII | 532851 | 532860 | 532860 | rightend | YHR211W | 92.416 | 2136 | 138 | 2 | 1096 | 3228 | 514291 | 516405 | 0 | 3025 |
| YJM1356 | IX | 1 | 10 | 403975 | leftend | YIL169C | 90.958 | 1294 | 115 | 2 | 520 | 1812 | 2371 | 1079 | 0 | 1740 |
| YJM1356 | IX | 403966 | 403975 | 403975 | rightend | YIR039C | 99.009 | 1614 | 16 | 0 | 1 | 1614 | 402311 | 400698 | 0 | 2892 |
| YJM1356 | X | 1 | 10 | 726636 | leftend | YJL213W | 99.699 | 996 | 3 | 0 | 1 | 996 | 42373 | 43368 | 0 | 1823 |
| YJM1356 | X | 725592 | 725601 | 726636 | middle |  |  |  |  |  |  |  |  |  |  |  |
| YJM1356 | X | 726636 | 726636 | 726636 | rightend_complete | YJR152W | 99.755 | 1632 | 3 | 1 | 1 | 1632 | 711456 | 713086 | 0 | 2990 |
| YJM1356 | XI | 1 | 10 | 670775 | leftend | YKL222C | 99.386 | 2118 | 13 | 0 | 1 | 2118 | 11506 | 9389 | 0 | 3840 |
| YJM1356 | XI | 670280 | 670289 | 670775 | middle |  |  |  |  |  |  |  |  |  |  |  |
| YJM1356 | XI | 670775 | 670775 | 670775 | rightend_complete | YKR101W | 99.847 | 1965 | 3 | 0 | 1 | 1965 | 645338 | 647302 | 0 | 3613 |
| YJM1356 | XII | 1 |  | 1956208 | leftend_complete | YLL062C | 100 | 975 | 0 | 0 | 1 | 975 | 5382 | 4408 | 0 | 1801 |
| YJM1356 | XII | 1376 | 1385 | 1956208 | middle |  |  |  |  |  |  |  |  |  |  |  |
| YJM1356 | XII | 1956199 | 1956208 | 1956208 | rightend | YLR460C | 94.336 | 1130 | 62 | 2 | 3 | 1131 | 1954455 | 1953327 | 0 | 1731 |
| YJM1356 | XIII | 1 | 10 | 899430 | leftend | YML131W | 100 | 1098 | 0 | 0 | 1 | 1098 | 3311 | 4408 | 0 | 2028 |
| YJM1356 | XIII | 897978 | 897987 | 899430 | middle |  |  |  |  |  |  |  |  |  |  |  |
| YJM1356 | XIII | 899421 | 899430 | 899430 | rightend | YMR321C | 99.361 | 313 | 2 | 0 | 6 | 318 | 888637 | 888325 | 2.48E-161 | 568 |
| YJM1356 | XIV | 1 | 10 | 767709 | leftend | YNL332W | 98.631 | 1023 | 14 | 0 | 1 | 1023 | 16992 | 18014 | 0 | 1812 |
| YJM1356 | XIV | 740 | 749 | 767709 | middle |  |  |  |  |  |  |  |  |  |  |  |
| YJM1356 | XIV | 767700 | 767709 | 767709 | rightend | YNR071C | 100 | 1029 | 0 | 0 | 1 | 1029 | 755745 | 754717 | 0 | 1901 |
| YJM1356 | XV | 1 |  | 1057054 | leftend_complete | YOL154W | 99.467 | 750 | 4 | 0 | 1 | 750 | 7705 | 8454 | 0 | 1363 |
| YJM1356 | XV | 1057045 | 1057054 | 1057054 | rightend | YOR381W | 99.906 | 2137 | 1 | 1 | 1 | 2136 | 1004899 | 1007035 | 0 | 3934 |
| YJM1356 | XVI | 1 | 10 | 895498 | leftend | YPL274W | 99.943 | 1764 | 1 | 0 | 1 | 1764 | 6537 | 8300 | 0 | 3253 |
| YJM1356 | XVI | 895489 | 895498 | 895498 | rightend | YPR201W | 100 | 1215 | 0 | 0 | 1 | 1215 | 892651 | 893865 | 0 | 2244 |
| YJM1381 | I | 1 | 10 | 181440 | leftend | YAL063C | 92.241 | 1740 | 135 | 0 | 1759 | 3498 | 178536 | 180275 | 0 | 2466 |
| YJM1381 | I | 525 | 534 | 181440 | middle |  |  |  |  |  |  |  |  |  |  |  |
| YJM1381 | I | 6960 | 6969 | 181440 | middle |  |  |  |  |  |  |  |  |  |  |  |
| YJM1381 | I | 180276 | 180285 | 181440 | middle |  |  |  |  |  |  |  |  |  |  |  |
| YJM1381 | I | 181431 | 181440 | 181440 | rightend | YAR050W | 82.051 | 1638 | 284 | 5 | 1 | 1630 | 177717 | 179352 | 0 | 1387 |
| YJM1381 | II | 1 | 10 | 788418 | leftend | YNCB0001W | 100 | 84 | 0 | 0 | 1 | 84 | 4069 | 4152 | 4.16E-38 | 156 |
| YJM1381 | II | 1129 | 1138 | 788418 | middle |  |  |  |  |  |  |  |  |  |  |  |
| YJM1381 | II | 2549 | 2558 | 788418 | middle |  |  |  |  |  |  |  |  |  |  |  |
| YJM1381 | II | 787524 | 787533 | 788418 | middle |  |  |  |  |  |  |  |  |  |  |  |
| YJM1381 | II | 788418 | 788418 | 788418 | rightend_complete | YBR297W | 98.863 | 1407 | 14 | 1 | 1 | 1407 | 777562 | 778966 | 0 | 2508 |
| YJM1381 | III | 1 |  | 341639 | leftend_complete | YCL073C | 99.729 | 1848 | 5 | 0 | 1 | 1848 | 7946 | 6099 | 0 | 3386 |
| YJM1381 | III | 332539 | 332548 | 341639 | middle |  |  |  |  |  |  |  |  |  |  |  |
| YJM1381 | III | 341630 | 341639 | 341639 | rightend | YCR101C | 98.182 | 550 | 9 | 1 | 1 | 549 | 297704 | 297155 | 0 | 959 |
| YJM1381 | IV | 1 | 10 | 1460900 | leftend | YDL243C | 93.769 | 979 | 61 | 0 | 1 | 979 | 10239 | 9261 | 0 | 1471 |
| YJM1381 | IV | 1460900 | 1460900 | 1460900 | rightend_complete | YDR541C | 99.903 | 1035 | 1 | 0 | 1 | 1035 | 1456178 | 1455144 | 0 | 1906 |
| YJM1381 | V | 1 | 10 | 554916 | leftend | YEL070W | 99.271 | 1509 | 11 | 0 | 1 | 1509 | 15258 | 16766 | 0 | 2726 |
| YJM1381 | V | 734 | 743 | 554916 | middle |  |  |  |  |  |  |  |  |  |  |  |
| YJM1381 | V | 554524 | 554533 | 554916 | middle |  |  |  |  |  |  |  |  |  |  |  |
| YJM1381 | V | 554916 | 554916 | 554916 | rightend_complete | YER187W | 91.745 | 424 | 33 | 2 | 4 | 426 | 549786 | 550208 | 1.6E-167 | 588 |
| YJM1381 | VI | 1 | 10 | 250706 | leftend | YFL051C | 100 | 483 | 0 | 0 | 1 | 483 | 17328 | 16846 | 0 | 893 |
| YJM1381 | VI | 250697 | 250706 | 250706 | rightend | YFR055W | 100 | 1023 | 0 | 0 | 1 | 1023 | 244392 | 245414 | 0 | 1890 |
| YJM1381 | VII | 1 |  | 1041764 | leftend_complete | YGL258W | 97.585 | 621 | 15 | 0 | 1 | 621 | 11638 | 12258 | 0 | 1064 |
| YJM1381 | VII | 489 | 498 | 1041764 | middle |  |  |  |  |  |  |  |  |  |  |  |
| YJM1381 | VII | 696 | 705 | 1041764 | middle |  |  |  |  |  |  |  |  |  |  |  |
| YJM1381 | VII | 1040426 | 1040435 | 1041764 | middle |  |  |  |  |  |  |  |  |  |  |  |
| YJM1381 | VII | 1041755 | 1041764 | 1041764 | rightend | YGR287C | 98.192 | 1770 | 32 | 0 | 1 | 1770 | 1034032 | 1032263 | 0 | 3092 |
| YJM1381 | VIII | 1 | 10 | 549273 | leftend | YHL044W | 98.729 | 708 | 9 | 0 | 1 | 708 | 7936 | 8643 | 0 | 1258 |
| YJM1381 | VIII | 523059 | 523068 | 549273 | middle |  |  |  |  |  |  |  |  |  |  |  |
| YJM1381 | VIII | 524307 | 524316 | 549273 | middle |  |  |  |  |  |  |  |  |  |  |  |
| YJM1381 | VIII | 539976 | 539985 | 549273 | middle |  |  |  |  |  |  |  |  |  |  |  |
| YJM1381 | VIII | 542726 | 542735 | 549273 | middle |  |  |  |  |  |  |  |  |  |  |  |
| YJM1381 | VIII | 545461 | 545470 | 549273 | middle |  |  |  |  |  |  |  |  |  |  |  |
| YJM1381 | VIII | 545913 | 545922 | 549273 | middle |  |  |  |  |  |  |  |  |  |  |  |
| YJM1381 | VIII | 548312 | 548321 | 549273 | middle |  |  |  |  |  |  |  |  |  |  |  |
| YJM1381 | VIII | 549264 | 549273 | 549273 | rightend | YHR211W | 91.811 | 1209 | 45 | 12 | 2047 | 3228 | 524317 | 525498 | 0 | 1635 |
| YJM1381 | IX | 1 |  | 425605 | leftend_complete | YIL169C | 97.92 | 1442 | 24 | 2 | 478 | 1919 | 18520 | 17085 | 0 | 2492 |
| YJM1381 | IX | 411 | 420 | 425605 | middle |  |  |  |  |  |  |  |  |  |  |  |
| YJM1381 | IX | 2037 | 2046 | 425605 | middle |  |  |  |  |  |  |  |  |  |  |  |
| YJM1381 | IX | 425596 | 425605 | 425605 | rightend | YIR039C | 99.69 | 1614 | 5 | 0 | 1 | 1614 | 423475 | 421862 | 0 | 2953 |
| YJM1381 | X | 1 | 10 | 734239 | leftend | YJL213W | 99.699 | 996 | 3 | 0 | 1 | 996 | 30439 | 31434 | 0 | 1823 |
| YJM1381 | X | 1878 | 1887 | 734239 | middle |  |  |  |  |  |  |  |  |  |  |  |
| YJM1381 | X | 731933 | 731942 | 734239 | middle |  |  |  |  |  |  |  |  |  |  |  |
| YJM1381 | X | 733770 | 733779 | 734239 | middle |  |  |  |  |  |  |  |  |  |  |  |
| YJM1381 | X | 734239 | 734239 | 734239 | rightend_complete | YJR152W | 100 | 1632 | 0 | 0 | 1 | 1632 | 707153 | 708784 | 0 | 3014 |
| YJM1381 | XI | 1 | 10 | 658578 | leftend | YKL222C | 99.528 | 2118 | 10 | 0 | 1 | 2118 | 2612 | 495 | 0 | 3856 |
| YJM1381 | XI | 658568 | 658578 | 658578 | rightend | YKR101W | 99.847 | 1965 | 3 | 0 | 1 | 1965 | 630882 | 632846 | 0 | 3613 |
| YJM1381 | XII | 1 | 10 | 1760731 | leftend | YLL062C | 99.795 | 975 | 2 | 0 | 1 | 975 | 6548 | 5574 | 0 | 1790 |
| YJM1381 | XII | 661 | 670 | 1760731 | middle |  |  |  |  |  |  |  |  |  |  |  |
| YJM1381 | XII | 1498 | 1507 | 1760731 | middle |  |  |  |  |  |  |  |  |  |  |  |
| YJM1381 | XII | 1990 | 1999 | 1760731 | middle |  |  |  |  |  |  |  |  |  |  |  |
| YJM1381 | XII | 2200 | 2209 | 1760731 | middle |  |  |  |  |  |  |  |  |  |  |  |
| YJM1381 | XII | 1759964 | 1759973 | 1760731 | middle |  |  |  |  |  |  |  |  |  |  |  |
| YJM1381 | XII | 1760722 | 1760731 | 1760731 | rightend | YLR460C | 94.159 | 1130 | 64 | 2 | 3 | 1131 | 1757791 | 1756663 | 0 | 1720 |
| YJM1381 | XIII | 1 | 10 | 904894 | leftend | YML131W | 99.909 | 1098 | 1 | 0 | 1 | 1098 | 2308 | 3405 | 0 | 2023 |
| YJM1381 | XIII | 904885 | 904894 | 904894 | rightend | YMR321C | 99.361 | 313 | 2 | 0 | 6 | 318 | 885785 | 885473 | 2.48E-161 | 568 |
| YJM1381 | XIV | 1 |  | 761051 | leftend_complete | YNL332W | 98.631 | 1023 | 14 | 0 | 1 | 1023 | 7498 | 8520 | 0 | 1812 |
| YJM1381 | XIV | 1837 | 1846 | 761051 | middle |  |  |  |  |  |  |  |  |  |  |  |
| YJM1381 | XIV | 758188 | 758197 | 761051 | middle |  |  |  |  |  |  |  |  |  |  |  |
| YJM1381 | XIV | 760304 | 760313 | 761051 | middle |  |  |  |  |  |  |  |  |  |  |  |
| YJM1381 | XIV | 761042 | 761051 | 761051 | rightend | YNR071C | 99.417 | 1029 | 6 | 0 | 1 | 1029 | 747719 | 746691 | 0 | 1868 |
| YJM1381 | XV | 1 | 10 | 1032129 | leftend | YOL154W | 99.733 | 750 | 2 | 0 | 1 | 750 | 23037 | 23786 | 0 | 1375 |
| YJM1381 | XV | 1032120 | 1032129 | 1032129 | rightend | YOR381W | 94.239 | 2135 | 121 | 2 | 1 | 2134 | 1022897 | 1025030 | 0 | 3260 |
| YJM1381 | XVI | 1 | 10 | 909399 | leftend | YPL274W | 99.15 | 1764 | 15 | 0 | 1 | 1764 | 21986 | 23749 | 0 | 3175 |
| YJM1381 | XVI | 909399 | 909399 | 909399 | rightend_complete | YPR201W | 99.67 | 1213 | 4 | 0 | 1 | 1213 | 906132 | 907344 | 0 | 2218 |
| YJM1383 | I | 1 | 10 | 202489 | leftend | YAL063C | 84.972 | 1770 | 259 | 5 | 1 | 1765 | 13015 | 11248 | 0 | 1788 |
| YJM1383 | I | 461 | 470 | 202489 | middle |  |  |  |  |  |  |  |  |  |  |  |
| YJM1383 | I | 184587 | 184596 | 202489 | middle |  |  |  |  |  |  |  |  |  |  |  |
| YJM1383 | I | 193737 | 193746 | 202489 | middle |  |  |  |  |  |  |  |  |  |  |  |
| YJM1383 | I | 202480 | 202489 | 202489 | rightend | YAR050W | 88.533 | 1936 | 197 | 17 | 2329 | 4253 | 12172 | 10251 | 0 | 2322 |
| YJM1383 | II | 1 | 10 | 785698 | leftend | YNCB0001W | 100 | 84 | 0 | 0 | 1 | 84 | 3978 | 4061 | 4.16E-38 | 156 |
| YJM1383 | II | 1729 | 1738 | 785698 | middle |  |  |  |  |  |  |  |  |  |  |  |
| YJM1383 | II | 3275 | 3284 | 785698 | middle |  |  |  |  |  |  |  |  |  |  |  |
| YJM1383 | II | 3493 | 3502 | 785698 | middle |  |  |  |  |  |  |  |  |  |  |  |
| YJM1383 | II | 785689 | 785698 | 785698 | rightend | YBR297W | 99.076 | 1407 | 13 | 0 | 1 | 1407 | 776133 | 777539 | 0 | 2527 |
| YJM1383 | III | 1 |  | 301574 | leftend_complete | YCL073C | 99.675 | 1848 | 6 | 0 | 1 | 1848 | 7932 | 6085 | 0 | 3380 |
| YJM1383 | III | 317 | 326 | 301574 | middle |  |  |  |  |  |  |  |  |  |  |  |
| YJM1383 | III | 301565 | 301574 | 301574 | rightend | YCR101C | 100 | 549 | 0 | 0 | 1 | 549 | 296677 | 296129 | 0 | 1014 |
| YJM1383 | IV | 1 |  | 1463876 | leftend_complete | YDL243C | 93.838 | 990 | 61 | 0 | 1 | 990 | 12716 | 11727 | 0 | 1491 |
| YJM1383 | IV | 1462596 | 1462605 | 1463876 | middle |  |  |  |  |  |  |  |  |  |  |  |
| YJM1383 | IV | 1463867 | 1463876 | 1463876 | rightend | YDR541C | 99.324 | 1035 | 7 | 0 | 1 | 1035 | 1459756 | 1458722 | 0 | 1873 |
| YJM1383 | V | 1 |  | 552666 | leftend_complete | YEL070W | 99.139 | 1509 | 13 | 0 | 1 | 1509 | 15850 | 17358 | 0 | 2715 |
| YJM1383 | V | 2225 | 2234 | 552666 | middle |  |  |  |  |  |  |  |  |  |  |  |
| YJM1383 | V | 552657 | 552666 | 552666 | rightend | YER187W | 99.296 | 426 | 3 | 0 | 1 | 426 | 549160 | 549585 | 0 | 771 |
| YJM1383 | VI | 1 | 10 | 234049 | leftend | YFL051C | 98.451 | 452 | 7 | 0 | 1 | 452 | 1470 | 1019 | 0 | 797 |
| YJM1383 | VI | 234040 | 234049 | 234049 | rightend | YFR055W | 99.902 | 1022 | 1 | 0 | 1 | 1022 | 227832 | 228853 | 0 | 1882 |
| YJM1383 | VII | 1 | 10 | 1052398 | leftend | YGL258W | 97.585 | 621 | 15 | 0 | 1 | 621 | 11277 | 11897 | 0 | 1064 |
| YJM1383 | VII | 1052389 | 1052398 | 1052398 | rightend | YGR287C | 98.757 | 1770 | 22 | 0 | 1 | 1770 | 1032873 | 1031104 | 0 | 3147 |
| YJM1383 | VIII | 1 | 10 | 537866 | leftend | YHL044W | 98.446 | 708 | 11 | 0 | 1 | 708 | 8994 | 9701 | 0 | 1247 |
| YJM1383 | VIII | 518644 | 518653 | 537866 | middle |  |  |  |  |  |  |  |  |  |  |  |
| YJM1383 | VIII | 533181 | 533190 | 537866 | middle |  |  |  |  |  |  |  |  |  |  |  |
| YJM1383 | VIII | 537650 | 537659 | 537866 | middle |  |  |  |  |  |  |  |  |  |  |  |
| YJM1383 | VIII | 537866 | 537866 | 537866 | rightend_complete | YHR211W | 92.625 | 1261 | 59 | 14 | 1996 | 3228 | 519379 | 520633 | 0 | 1783 |
| YJM1383 | IX | 1 | 10 | 409658 | leftend | YIL169C | 96.118 | 2164 | 73 | 7 | 520 | 2679 | 8290 | 6134 | 0 | 3520 |
| YJM1383 | IX | 409649 | 409658 | 409658 | rightend | YIR039C | 98.947 | 1614 | 17 | 0 | 1 | 1614 | 408026 | 406413 | 0 | 2887 |
| YJM1383 | X | 1 | 10 | 727362 | leftend | YJL213W | 99.699 | 996 | 3 | 0 | 1 | 996 | 25572 | 26567 | 0 | 1823 |
| YJM1383 | X | 726435 | 726444 | 727362 | middle |  |  |  |  |  |  |  |  |  |  |  |
| YJM1383 | X | 727362 | 727362 | 727362 | rightend_complete | YJR152W | 100 | 1632 | 0 | 0 | 1 | 1632 | 695593 | 697224 | 0 | 3014 |
| YJM1383 | XI | 1 | 10 | 698638 | leftend | YKL222C | 99.386 | 2118 | 13 | 0 | 1 | 2118 | 3183 | 1066 | 0 | 3840 |
| YJM1383 | XI | 696930 | 696939 | 698638 | middle |  |  |  |  |  |  |  |  |  |  |  |
| YJM1383 | XI | 698638 | 698638 | 698638 | rightend_complete | YKR101W | 99.491 | 1965 | 10 | 0 | 1 | 1965 | 671852 | 673816 | 0 | 3574 |
| YJM1383 | XII | 1 |  | 1875050 | leftend_complete | YLL062C | 99.795 | 975 | 2 | 0 | 1 | 975 | 2221 | 1247 | 0 | 1790 |
| YJM1383 | XII | 255 | 264 | 1875050 | middle |  |  |  |  |  |  |  |  |  |  |  |
| YJM1383 | XII | 1874822 | 1874831 | 1875050 | middle |  |  |  |  |  |  |  |  |  |  |  |
| YJM1383 | XII | 1875050 | 1875050 | 1875050 | rightend_complete | YLR460C | 98.849 | 1129 | 13 | 0 | 3 | 1131 | 1864541 | 1863413 | 0 | 2013 |
| YJM1383 | XIII | 1 | 10 | 908617 | leftend | YML131W | 100 | 1098 | 0 | 0 | 1 | 1098 | 4778 | 5875 | 0 | 2028 |
| YJM1383 | XIII | 647 | 656 | 908617 | middle |  |  |  |  |  |  |  |  |  |  |  |
| YJM1383 | XIII | 2507 | 2516 | 908617 | middle |  |  |  |  |  |  |  |  |  |  |  |
| YJM1383 | XIII | 907985 | 907994 | 908617 | middle |  |  |  |  |  |  |  |  |  |  |  |
| YJM1383 | XIII | 908608 | 908617 | 908617 | rightend | YMR321C | 99.361 | 313 | 2 | 0 | 6 | 318 | 889375 | 889063 | 2.48E-161 | 568 |
| YJM1383 | XIV | 1 |  | 759072 | leftend_complete | YNL332W | 98.534 | 1023 | 15 | 0 | 1 | 1023 | 8030 | 9052 | 0 | 1807 |
| YJM1383 | XIV | 1219 | 1228 | 759072 | middle |  |  |  |  |  |  |  |  |  |  |  |
| YJM1383 | XIV | 757606 | 757615 | 759072 | middle |  |  |  |  |  |  |  |  |  |  |  |
| YJM1383 | XIV | 757815 | 757824 | 759072 | middle |  |  |  |  |  |  |  |  |  |  |  |
| YJM1383 | XIV | 759072 | 759072 | 759072 | rightend_complete | YNR071C | 100 | 1029 | 0 | 0 | 1 | 1029 | 747140 | 746112 | 0 | 1901 |
| YJM1383 | XV | 1 | 10 | 1037579 | leftend | YOL154W | 99.467 | 750 | 4 | 0 | 1 | 750 | 23867 | 24616 | 0 | 1363 |
| YJM1383 | XV | 482 | 491 | 1037579 | middle |  |  |  |  |  |  |  |  |  |  |  |
| YJM1383 | XV | 2061 | 2070 | 1037579 | middle |  |  |  |  |  |  |  |  |  |  |  |
| YJM1383 | XV | 1037570 | 1037579 | 1037579 | rightend | YOR381W | 98.689 | 2136 | 28 | 0 | 1 | 2136 | 1022232 | 1024367 | 0 | 3790 |
| YJM1383 | XVI | 1 | 10 | 915935 | leftend | YPL274W | 99.49 | 1764 | 9 | 0 | 1 | 1764 | 22751 | 24514 | 0 | 3208 |
| YJM1383 | XVI | 915146 | 915155 | 915935 | middle |  |  |  |  |  |  |  |  |  |  |  |
| YJM1383 | XVI | 915935 | 915935 | 915935 | rightend_complete | YPR201W | 100 | 1215 | 0 | 0 | 1 | 1215 | 912037 | 913251 | 0 | 2244 |
| YJM1385 | I | 1 |  | 218690 | leftend_complete | YAL063C | 96.567 | 1136 | 37 | 2 | 2828 | 3962 | 13483 | 12349 | 0 | 1881 |
| YJM1385 | I | 1220 | 1229 | 218690 | middle |  |  |  |  |  |  |  |  |  |  |  |
| YJM1385 | I | 3177 | 3186 | 218690 | middle |  |  |  |  |  |  |  |  |  |  |  |
| YJM1385 | I | 13484 | 13493 | 218690 | middle |  |  |  |  |  |  |  |  |  |  |  |
| YJM1385 | I | 190497 | 190506 | 218690 | middle |  |  |  |  |  |  |  |  |  |  |  |
| YJM1385 | I | 209214 | 209223 | 218690 | middle |  |  |  |  |  |  |  |  |  |  |  |
| YJM1385 | I | 218681 | 218690 | 218690 | rightend | YAR050W | 93.079 | 838 | 34 | 7 | 3466 | 4282 | 13481 | 12647 | 0 | 1205 |
| YJM1385 | II | 1 | 10 | 779078 | leftend | YNCB0001W | 100 | 84 | 0 | 0 | 1 | 84 | 3924 | 4007 | 4.16E-38 | 156 |
| YJM1385 | II | 670 | 679 | 779078 | middle |  |  |  |  |  |  |  |  |  |  |  |
| YJM1385 | II | 1474 | 1483 | 779078 | middle |  |  |  |  |  |  |  |  |  |  |  |
| YJM1385 | II | 2393 | 2402 | 779078 | middle |  |  |  |  |  |  |  |  |  |  |  |
| YJM1385 | II | 779069 | 779078 | 779078 | rightend | YBR297W | 100 | 1382 | 0 | 0 | 1 | 1382 | 777687 | 779068 | 0 | 2553 |
| YJM1385 | III | 1 |  | 310682 | leftend_complete | YCL073C | 99.784 | 1848 | 4 | 0 | 1 | 1848 | 8015 | 6168 | 0 | 3391 |
| YJM1385 | III | 310673 | 310682 | 310682 | rightend | YCR101C | 99.818 | 549 | 1 | 0 | 1 | 549 | 297653 | 297105 | 0 | 1009 |
| YJM1385 | IV | 1 | 10 | 1461841 | leftend | YDL243C | 93.737 | 990 | 62 | 0 | 1 | 990 | 3447 | 2458 | 0 | 1485 |
| YJM1385 | IV | 940 | 949 | 1461841 | middle |  |  |  |  |  |  |  |  |  |  |  |
| YJM1385 | IV | 1461221 | 1461230 | 1461841 | middle |  |  |  |  |  |  |  |  |  |  |  |
| YJM1385 | IV | 1461832 | 1461841 | 1461841 | rightend | YDR541C | 99.614 | 1035 | 4 | 0 | 1 | 1035 | 1458192 | 1457158 | 0 | 1890 |
| YJM1385 | V | 1 |  | 554664 | leftend_complete | YEL070W | 99.337 | 1509 | 10 | 0 | 1 | 1509 | 18447 | 19955 | 0 | 2732 |
| YJM1385 | V | 5038 | 5047 | 554664 | middle |  |  |  |  |  |  |  |  |  |  |  |
| YJM1385 | V | 554213 | 554222 | 554664 | middle |  |  |  |  |  |  |  |  |  |  |  |
| YJM1385 | V | 554664 | 554664 | 554664 | rightend_complete | YER187W | 91.981 | 424 | 32 | 2 | 4 | 426 | 551650 | 552072 | 3.44E-169 | 593 |
| YJM1385 | VI | 1 |  | 314219 | leftend_complete | YFL051C | 99.115 | 452 | 4 | 0 | 1 | 452 | 48014 | 47563 | 0 | 813 |
| YJM1385 | VI | 314219 | 314219 | 314219 | rightend_complete | YFR055W | 99.511 | 1022 | 5 | 0 | 1 | 1022 | 308237 | 309258 | 0 | 1860 |
| YJM1385 | VII | 1 | 10 | 1095605 | leftend | YGL258W | 99.517 | 621 | 3 | 0 | 1 | 621 | 53112 | 53732 | 0 | 1131 |
| YJM1385 | VII | 13749 | 13758 | 1095605 | middle |  |  |  |  |  |  |  |  |  |  |  |
| YJM1385 | VII | 1094645 | 1094654 | 1095605 | middle |  |  |  |  |  |  |  |  |  |  |  |
| YJM1385 | VII | 1095596 | 1095605 | 1095605 | rightend | YGR287C | 99.262 | 1626 | 12 | 0 | 145 | 1770 | 1082784 | 1081159 | 0 | 2937 |
| YJM1385 | VIII | 1 | 10 | 526315 | leftend | YHL044W | 98.446 | 708 | 11 | 0 | 1 | 708 | 8834 | 9541 | 0 | 1247 |
| YJM1385 | VIII | 1103 | 1112 | 526315 | middle |  |  |  |  |  |  |  |  |  |  |  |
| YJM1385 | VIII | 7438 | 7447 | 526315 | middle |  |  |  |  |  |  |  |  |  |  |  |
| YJM1385 | VIII | 522680 | 522689 | 526315 | middle |  |  |  |  |  |  |  |  |  |  |  |
| YJM1385 | VIII | 526306 | 526315 | 526315 | rightend | YHR211W | 81.86 | 2376 | 368 | 28 | 1 | 2337 | 515457 | 517808 | 0 | 1941 |
| YJM1385 | IX | 1 | 10 | 416768 | leftend | YIL169C | 94.057 | 1649 | 84 | 8 | 272 | 1919 | 8562 | 6927 | 0 | 2490 |
| YJM1385 | IX | 1098 | 1107 | 416768 | middle |  |  |  |  |  |  |  |  |  |  |  |
| YJM1385 | IX | 416768 | 416768 | 416768 | rightend_complete | YIR039C | 99.69 | 1614 | 5 | 0 | 1 | 1614 | 409010 | 407397 | 0 | 2953 |
| YJM1385 | X | 1 |  | 709051 | leftend_complete | YJL213W | 99.598 | 996 | 4 | 0 | 1 | 996 | 12613 | 13608 | 0 | 1818 |
| YJM1385 | X | 1143 | 1152 | 709051 | middle |  |  |  |  |  |  |  |  |  |  |  |
| YJM1385 | X | 3305 | 3314 | 709051 | middle |  |  |  |  |  |  |  |  |  |  |  |
| YJM1385 | X | 4030 | 4039 | 709051 | middle |  |  |  |  |  |  |  |  |  |  |  |
| YJM1385 | X | 709042 | 709051 | 709051 | rightend | YJR152W | 100 | 1632 | 0 | 0 | 1 | 1632 | 683848 | 685479 | 0 | 3014 |
| YJM1385 | XI | 1 | 10 | 660570 | leftend | YKL222C | 99.433 | 2118 | 12 | 0 | 1 | 2118 | 3289 | 1172 | 0 | 3845 |
| YJM1385 | XI | 660561 | 660570 | 660570 | rightend | YKR101W | 99.746 | 1965 | 5 | 0 | 1 | 1965 | 637842 | 639806 | 0 | 3602 |
| YJM1385 | XII | 1 | 10 | 2188600 | leftend | YLL062C | 100 | 975 | 0 | 0 | 1 | 975 | 3908 | 2934 | 0 | 1801 |
| YJM1385 | XII | 2184905 | 2184914 | 2188600 | middle |  |  |  |  |  |  |  |  |  |  |  |
| YJM1385 | XII | 2185926 | 2185935 | 2188600 | middle |  |  |  |  |  |  |  |  |  |  |  |
| YJM1385 | XII | 2188167 | 2188176 | 2188600 | middle |  |  |  |  |  |  |  |  |  |  |  |
| YJM1385 | XII | 2188591 | 2188600 | 2188600 | rightend | YLR460C | 100 | 1131 | 0 | 0 | 1 | 1131 | 2178287 | 2177157 | 0 | 2089 |
| YJM1385 | XIII | 1 |  | 895951 | leftend_complete | YML131W | 99.909 | 1098 | 1 | 0 | 1 | 1098 | 3035 | 4132 | 0 | 2023 |
| YJM1385 | XIII | 682 | 691 | 895951 | middle |  |  |  |  |  |  |  |  |  |  |  |
| YJM1385 | XIII | 892410 | 892419 | 895951 | middle |  |  |  |  |  |  |  |  |  |  |  |
| YJM1385 | XIII | 892627 | 892636 | 895951 | middle |  |  |  |  |  |  |  |  |  |  |  |
| YJM1385 | XIII | 895951 | 895951 | 895951 | rightend_complete | YMR321C | 99.042 | 313 | 3 | 0 | 6 | 318 | 888898 | 888586 | 1.15E-159 | 562 |
| YJM1385 | XIV | 1 | 10 | 758233 | leftend | YNL332W | 99.218 | 1023 | 8 | 0 | 1 | 1023 | 6666 | 7688 | 0 | 1845 |
| YJM1385 | XIV | 1341 | 1350 | 758233 | middle |  |  |  |  |  |  |  |  |  |  |  |
| YJM1385 | XIV | 757670 | 757679 | 758233 | middle |  |  |  |  |  |  |  |  |  |  |  |
| YJM1385 | XIV | 758224 | 758233 | 758233 | rightend | YNR071C | 100 | 1029 | 0 | 0 | 1 | 1029 | 746385 | 745357 | 0 | 1901 |
| YJM1385 | XV | 1 | 10 | 1049990 | leftend | YOL154W | 99.467 | 750 | 4 | 0 | 1 | 750 | 24278 | 25027 | 0 | 1363 |
| YJM1385 | XV | 1224 | 1233 | 1049990 | middle |  |  |  |  |  |  |  |  |  |  |  |
| YJM1385 | XV | 1047517 | 1047526 | 1049990 | middle |  |  |  |  |  |  |  |  |  |  |  |
| YJM1385 | XV | 1048806 | 1048815 | 1049990 | middle |  |  |  |  |  |  |  |  |  |  |  |
| YJM1385 | XV | 1049506 | 1049515 | 1049990 | middle |  |  |  |  |  |  |  |  |  |  |  |
| YJM1385 | XV | 1049981 | 1049990 | 1049990 | rightend | YOR381W | 98.736 | 2136 | 27 | 0 | 1 | 2136 | 1022236 | 1024371 | 0 | 3795 |
| YJM1385 | XVI | 1 | 10 | 906875 | leftend | YPL274W | 98.356 | 1764 | 28 | 1 | 1 | 1764 | 14379 | 16141 | 0 | 3096 |
| YJM1385 | XVI | 500 | 509 | 906875 | middle |  |  |  |  |  |  |  |  |  |  |  |
| YJM1385 | XVI | 893 | 902 | 906875 | middle |  |  |  |  |  |  |  |  |  |  |  |
| YJM1385 | XVI | 2039 | 2048 | 906875 | middle |  |  |  |  |  |  |  |  |  |  |  |
| YJM1385 | XVI | 3671 | 3680 | 906875 | middle |  |  |  |  |  |  |  |  |  |  |  |
| YJM1385 | XVI | 3874 | 3883 | 906875 | middle |  |  |  |  |  |  |  |  |  |  |  |
| YJM1385 | XVI | 905883 | 905892 | 906875 | middle |  |  |  |  |  |  |  |  |  |  |  |
| YJM1385 | XVI | 906866 | 906875 | 906875 | rightend | YPR201W | 99.588 | 1215 | 5 | 0 | 1 | 1215 | 902909 | 904123 | 0 | 2217 |
| YJM1386 | I | 1 | 10 | 199344 | leftend | YAL063C | 91.173 | 1892 | 167 | 0 | 1624 | 3515 | 187457 | 189348 | 0 | 2569 |
| YJM1386 | I | 752 | 761 | 199344 | middle |  |  |  |  |  |  |  |  |  |  |  |
| YJM1386 | I | 13533 | 13542 | 199344 | middle |  |  |  |  |  |  |  |  |  |  |  |
| YJM1386 | I | 189349 | 189358 | 199344 | middle |  |  |  |  |  |  |  |  |  |  |  |
| YJM1386 | I | 193400 | 193409 | 199344 | middle |  |  |  |  |  |  |  |  |  |  |  |
| YJM1386 | I | 195169 | 195178 | 199344 | middle |  |  |  |  |  |  |  |  |  |  |  |
| YJM1386 | I | 196486 | 196495 | 199344 | middle |  |  |  |  |  |  |  |  |  |  |  |
| YJM1386 | I | 198073 | 198082 | 199344 | middle |  |  |  |  |  |  |  |  |  |  |  |
| YJM1386 | I | 198486 | 198495 | 199344 | middle |  |  |  |  |  |  |  |  |  |  |  |
| YJM1386 | I | 199335 | 199344 | 199344 | rightend | YAR050W | 89.183 | 2718 | 278 | 13 | 1902 | 4614 | 10329 | 7623 | 0 | 3376 |
| YJM1386 | II | 1 |  | 785373 | leftend_complete | YNCB0001W | 100 | 84 | 0 | 0 | 1 | 84 | 3222 | 3305 | 4.16E-38 | 156 |
| YJM1386 | II | 636 | 645 | 785373 | middle |  |  |  |  |  |  |  |  |  |  |  |
| YJM1386 | II | 1900 | 1909 | 785373 | middle |  |  |  |  |  |  |  |  |  |  |  |
| YJM1386 | II | 785364 | 785373 | 785373 | rightend | YBR297W | 99.005 | 1407 | 14 | 0 | 1 | 1407 | 775603 | 777009 | 0 | 2521 |
| YJM1386 | III | 1 | 10 | 310305 | leftend | YCL073C | 99.729 | 1848 | 5 | 0 | 1 | 1848 | 7495 | 5648 | 0 | 3386 |
| YJM1386 | III | 310296 | 310305 | 310305 | rightend | YCR101C | 92.362 | 563 | 28 | 10 | 1 | 549 | 296640 | 296079 | 0 | 787 |
| YJM1386 | IV | 1 | 10 | 1458677 | leftend | YDL243C | 95.051 | 990 | 49 | 0 | 1 | 990 | 4872 | 3883 | 0 | 1557 |
| YJM1386 | IV | 1458003 | 1458012 | 1458677 | middle |  |  |  |  |  |  |  |  |  |  |  |
| YJM1386 | IV | 1458668 | 1458677 | 1458677 | rightend | YDR541C | 99.903 | 1035 | 1 | 0 | 1 | 1035 | 1452940 | 1451906 | 0 | 1906 |
| YJM1386 | V | 1 | 10 | 557389 | leftend | YEL070W | 99.271 | 1509 | 11 | 0 | 1 | 1509 | 16654 | 18162 | 0 | 2726 |
| YJM1386 | V | 1435 | 1444 | 557389 | middle |  |  |  |  |  |  |  |  |  |  |  |
| YJM1386 | V | 1754 | 1763 | 557389 | middle |  |  |  |  |  |  |  |  |  |  |  |
| YJM1386 | V | 3249 | 3258 | 557389 | middle |  |  |  |  |  |  |  |  |  |  |  |
| YJM1386 | V | 557380 | 557389 | 557389 | rightend | YER187W | 97.653 | 426 | 10 | 0 | 1 | 426 | 550244 | 550669 | 0 | 732 |
| YJM1386 | VI | 1 | 10 | 257857 | leftend | YFL051C | 100 | 483 | 0 | 0 | 1 | 483 | 23327 | 22845 | 0 | 893 |
| YJM1386 | VI | 6153 | 6162 | 257857 | middle |  |  |  |  |  |  |  |  |  |  |  |
| YJM1386 | VI | 257848 | 257857 | 257857 | rightend | YFR055W | 100 | 1023 | 0 | 0 | 1 | 1023 | 251505 | 252527 | 0 | 1890 |
| YJM1386 | VII | 1 |  | 1041763 | leftend_complete | YGL258W | 97.262 | 621 | 17 | 0 | 1 | 621 | 11577 | 12197 | 0 | 1053 |
| YJM1386 | VII | 1041754 | 1041763 | 1041763 | rightend | YGR287C | 99.016 | 1626 | 16 | 0 | 145 | 1770 | 1034458 | 1032833 | 0 | 2915 |
| YJM1386 | VIII | 1 | 10 | 534336 | leftend | YHL044W | 98.446 | 708 | 11 | 0 | 1 | 708 | 10306 | 11013 | 0 | 1247 |
| YJM1386 | VIII | 1824 | 1833 | 534336 | middle |  |  |  |  |  |  |  |  |  |  |  |
| YJM1386 | VIII | 2287 | 2296 | 534336 | middle |  |  |  |  |  |  |  |  |  |  |  |
| YJM1386 | VIII | 516095 | 516104 | 534336 | middle |  |  |  |  |  |  |  |  |  |  |  |
| YJM1386 | VIII | 526966 | 526975 | 534336 | middle |  |  |  |  |  |  |  |  |  |  |  |
| YJM1386 | VIII | 528090 | 528099 | 534336 | middle |  |  |  |  |  |  |  |  |  |  |  |
| YJM1386 | VIII | 528851 | 528860 | 534336 | middle |  |  |  |  |  |  |  |  |  |  |  |
| YJM1386 | VIII | 529145 | 529154 | 534336 | middle |  |  |  |  |  |  |  |  |  |  |  |
| YJM1386 | VIII | 529919 | 529928 | 534336 | middle |  |  |  |  |  |  |  |  |  |  |  |
| YJM1386 | VIII | 531067 | 531076 | 534336 | middle |  |  |  |  |  |  |  |  |  |  |  |
| YJM1386 | VIII | 534336 | 534336 | 534336 | rightend_complete | YHR211W | 83.406 | 916 | 141 | 7 | 1 | 909 | 515183 | 516094 | 0 | 839 |
| YJM1386 | IX | 1 |  | 442415 | leftend_complete | YIL169C | 94.545 | 1650 | 86 | 4 | 272 | 1919 | 39876 | 38229 | 0 | 2545 |
| YJM1386 | IX | 673 | 682 | 442415 | middle |  |  |  |  |  |  |  |  |  |  |  |
| YJM1386 | IX | 10158 | 10167 | 442415 | middle |  |  |  |  |  |  |  |  |  |  |  |
| YJM1386 | IX | 441592 | 441601 | 442415 | middle |  |  |  |  |  |  |  |  |  |  |  |
| YJM1386 | IX | 442415 | 442415 | 442415 | rightend_complete | YIR039C | 98.947 | 1614 | 17 | 0 | 1 | 1614 | 440253 | 438640 | 0 | 2887 |
| YJM1386 | X | 1 |  | 718739 | leftend_complete | YJL213W | 99.699 | 996 | 3 | 0 | 1 | 996 | 23295 | 24290 | 0 | 1823 |
| YJM1386 | X | 637 | 646 | 718739 | middle |  |  |  |  |  |  |  |  |  |  |  |
| YJM1386 | X | 1490 | 1499 | 718739 | middle |  |  |  |  |  |  |  |  |  |  |  |
| YJM1386 | X | 717808 | 717817 | 718739 | middle |  |  |  |  |  |  |  |  |  |  |  |
| YJM1386 | X | 718730 | 718739 | 718739 | rightend | YJR152W | 99.449 | 1632 | 9 | 0 | 1 | 1632 | 693622 | 695253 | 0 | 2964 |
| YJM1386 | XI | 1 | 10 | 667204 | leftend | YKL222C | 99.481 | 2118 | 11 | 0 | 1 | 2118 | 4305 | 2188 | 0 | 3851 |
| YJM1386 | XI | 663640 | 663649 | 667204 | middle |  |  |  |  |  |  |  |  |  |  |  |
| YJM1386 | XI | 667204 | 667204 | 667204 | rightend_complete | YKR101W | 99.695 | 1965 | 6 | 0 | 1 | 1965 | 638794 | 640758 | 0 | 3596 |
| YJM1386 | XII | 1 | 10 | 1901677 | leftend | YLL062C | 99.282 | 975 | 7 | 0 | 1 | 975 | 5573 | 4599 | 0 | 1762 |
| YJM1386 | XII | 1296 | 1305 | 1901677 | middle |  |  |  |  |  |  |  |  |  |  |  |
| YJM1386 | XII | 1897232 | 1897241 | 1901677 | middle |  |  |  |  |  |  |  |  |  |  |  |
| YJM1386 | XII | 1899279 | 1899288 | 1901677 | middle |  |  |  |  |  |  |  |  |  |  |  |
| YJM1386 | XII | 1901005 | 1901014 | 1901677 | middle |  |  |  |  |  |  |  |  |  |  |  |
| YJM1386 | XII | 1901668 | 1901677 | 1901677 | rightend | YLR460C | 100 | 1131 | 0 | 0 | 1 | 1131 | 1892013 | 1890883 | 0 | 2089 |
| YJM1386 | XIII | 1 | 10 | 898765 | leftend | YML131W | 100 | 1098 | 0 | 0 | 1 | 1098 | 2522 | 3619 | 0 | 2028 |
| YJM1386 | XIII | 436 | 445 | 898765 | middle |  |  |  |  |  |  |  |  |  |  |  |
| YJM1386 | XIII | 896715 | 896724 | 898765 | middle |  |  |  |  |  |  |  |  |  |  |  |
| YJM1386 | XIII | 898756 | 898765 | 898765 | rightend | YMR321C | 99.361 | 313 | 2 | 0 | 6 | 318 | 887797 | 887485 | 2.48E-161 | 568 |
| YJM1386 | XIV | 1 |  | 768815 | leftend_complete | YNL332W | 99.343 | 609 | 4 | 0 | 415 | 1023 | 13663 | 14271 | 0 | 1103 |
| YJM1386 | XIV | 871 | 880 | 768815 | middle |  |  |  |  |  |  |  |  |  |  |  |
| YJM1386 | XIV | 3682 | 3691 | 768815 | middle |  |  |  |  |  |  |  |  |  |  |  |
| YJM1386 | XIV | 13653 | 13662 | 768815 | middle |  |  |  |  |  |  |  |  |  |  |  |
| YJM1386 | XIV | 768806 | 768815 | 768815 | rightend | YNR071C | 100 | 1029 | 0 | 0 | 1 | 1029 | 752383 | 751355 | 0 | 1901 |
| YJM1386 | XV | 1 | 10 | 1050121 | leftend | YOL154W | 99.467 | 750 | 4 | 0 | 1 | 750 | 26356 | 27105 | 0 | 1363 |
| YJM1386 | XV | 3409 | 3418 | 1050121 | middle |  |  |  |  |  |  |  |  |  |  |  |
| YJM1386 | XV | 1049602 | 1049611 | 1050121 | middle |  |  |  |  |  |  |  |  |  |  |  |
| YJM1386 | XV | 1050112 | 1050121 | 1050121 | rightend | YOR381W | 99.906 | 2136 | 2 | 0 | 1 | 2136 | 1022211 | 1024346 | 0 | 3934 |
| YJM1386 | XVI | 1 | 10 | 902600 | leftend | YPL274W | 99.546 | 1764 | 8 | 0 | 1 | 1764 | 7684 | 9447 | 0 | 3214 |
| YJM1386 | XVI | 1088 | 1097 | 902600 | middle |  |  |  |  |  |  |  |  |  |  |  |
| YJM1386 | XVI | 1571 | 1580 | 902600 | middle |  |  |  |  |  |  |  |  |  |  |  |
| YJM1386 | XVI | 902591 | 902600 | 902600 | rightend | YPR201W | 99.588 | 1215 | 5 | 0 | 1 | 1215 | 898931 | 900145 | 0 | 2217 |
| YJM1387 | I | 1 | 10 | 187836 | leftend | YAL063C | 91.309 | 955 | 53 | 11 | 2715 | 3662 | 172801 | 173732 | 0 | 1277 |
| YJM1387 | I | 187415 | 187424 | 187836 | middle |  |  |  |  |  |  |  |  |  |  |  |
| YJM1387 | I | 187827 | 187836 | 187836 | rightend | YAR050W | 93.879 | 1650 | 71 | 9 | 2992 | 4614 | 172447 | 174093 | 0 | 2460 |
| YJM1387 | II | 1 | 10 | 779654 | leftend | YNCB0001W | 100 | 84 | 0 | 0 | 1 | 84 | 3089 | 3172 | 4.16E-38 | 156 |
| YJM1387 | II | 1696 | 1705 | 779654 | middle |  |  |  |  |  |  |  |  |  |  |  |
| YJM1387 | II | 779645 | 779654 | 779654 | rightend | YBR297W | 97.676 | 1205 | 28 | 0 | 1 | 1205 | 775453 | 776657 | 0 | 2071 |
| YJM1387 | III | 1 |  | 316209 | leftend_complete | YCL073C | 99.946 | 1848 | 1 | 0 | 1 | 1848 | 10281 | 8434 | 0 | 3408 |
| YJM1387 | III | 2371 | 2380 | 316209 | middle |  |  |  |  |  |  |  |  |  |  |  |
| YJM1387 | III | 311999 | 312008 | 316209 | middle |  |  |  |  |  |  |  |  |  |  |  |
| YJM1387 | III | 315551 | 315560 | 316209 | middle |  |  |  |  |  |  |  |  |  |  |  |
| YJM1387 | III | 316209 | 316209 | 316209 | rightend_complete | YCR101C | 99.636 | 549 | 2 | 0 | 1 | 549 | 298734 | 298186 | 0 | 1003 |
| YJM1387 | IV | 1 | 10 | 1476680 | leftend | YDL243C | 95.051 | 990 | 49 | 0 | 1 | 990 | 7949 | 6960 | 0 | 1557 |
| YJM1387 | IV | 1476680 | 1476680 | 1476680 | rightend_complete | YDR541C | 99.42 | 1035 | 6 | 0 | 1 | 1035 | 1472012 | 1470978 | 0 | 1879 |
| YJM1387 | V | 1 | 10 | 551496 | leftend | YEL070W | 98.476 | 1509 | 22 | 1 | 1 | 1509 | 13410 | 14917 | 0 | 2658 |
| YJM1387 | V | 551487 | 551496 | 551496 | rightend | YER187W | 91.981 | 424 | 32 | 2 | 4 | 426 | 547947 | 548369 | 3.44E-169 | 593 |
| YJM1387 | VI | 1 | 10 | 237035 | leftend | YFL051C | 98.23 | 452 | 8 | 0 | 1 | 452 | 4390 | 3939 | 0 | 791 |
| YJM1387 | VI | 1894 | 1903 | 237035 | middle |  |  |  |  |  |  |  |  |  |  |  |
| YJM1387 | VI | 2127 | 2136 | 237035 | middle |  |  |  |  |  |  |  |  |  |  |  |
| YJM1387 | VI | 237035 | 237035 | 237035 | rightend_complete | YFR055W | 100 | 1023 | 0 | 0 | 1 | 1023 | 230697 | 231719 | 0 | 1890 |
| YJM1387 | VII | 1 | 10 | 1048701 | leftend | YGL258W | 97.585 | 621 | 15 | 0 | 1 | 621 | 4790 | 5410 | 0 | 1064 |
| YJM1387 | VII | 1048692 | 1048701 | 1048701 | rightend | YGR287C | 98.757 | 1770 | 22 | 0 | 1 | 1770 | 1025635 | 1023866 | 0 | 3147 |
| YJM1387 | VIII | 1 | 10 | 543521 | leftend | YHL044W | 98.446 | 708 | 11 | 0 | 1 | 708 | 7515 | 8222 | 0 | 1247 |
| YJM1387 | VIII | 542868 | 542877 | 543521 | middle |  |  |  |  |  |  |  |  |  |  |  |
| YJM1387 | VIII | 543512 | 543521 | 543521 | rightend | YHR211W | 98.265 | 980 | 17 | 0 | 2249 | 3228 | 517356 | 518335 | 0 | 1716 |
| YJM1387 | IX | 1 | 10 | 443117 | leftend | YIL169C | 95.115 | 1699 | 66 | 11 | 198 | 1880 | 41735 | 40038 | 0 | 2662 |
| YJM1387 | IX | 443108 | 443117 | 443117 | rightend | YIR039C | 99.566 | 1614 | 7 | 0 | 1 | 1614 | 441941 | 440328 | 0 | 2942 |
| YJM1387 | X | 1 |  | 723543 | leftend_complete | YJL213W | 99.699 | 996 | 3 | 0 | 1 | 996 | 26930 | 27925 | 0 | 1823 |
| YJM1387 | X | 667 | 676 | 723543 | middle |  |  |  |  |  |  |  |  |  |  |  |
| YJM1387 | X | 721389 | 721398 | 723543 | middle |  |  |  |  |  |  |  |  |  |  |  |
| YJM1387 | X | 722179 | 722188 | 723543 | middle |  |  |  |  |  |  |  |  |  |  |  |
| YJM1387 | X | 723543 | 723543 | 723543 | rightend_complete | YJR152W | 99.755 | 1632 | 3 | 1 | 1 | 1632 | 697224 | 698854 | 0 | 2990 |
| YJM1387 | XI | 1 | 10 | 665838 | leftend | YKL222C | 99.481 | 2118 | 11 | 0 | 1 | 2118 | 4684 | 2567 | 0 | 3851 |
| YJM1387 | XI | 987 | 996 | 665838 | middle |  |  |  |  |  |  |  |  |  |  |  |
| YJM1387 | XI | 663050 | 663059 | 665838 | middle |  |  |  |  |  |  |  |  |  |  |  |
| YJM1387 | XI | 665303 | 665312 | 665838 | middle |  |  |  |  |  |  |  |  |  |  |  |
| YJM1387 | XI | 665838 | 665838 | 665838 | rightend_complete | YKR101W | 99.796 | 1965 | 4 | 0 | 1 | 1965 | 638927 | 640891 | 0 | 3607 |
| YJM1387 | XII | 1 |  | 2118914 | leftend_complete | YLL062C | 100 | 975 | 0 | 0 | 1 | 975 | 6137 | 5163 | 0 | 1801 |
| YJM1387 | XII | 1128 | 1137 | 2118914 | middle |  |  |  |  |  |  |  |  |  |  |  |
| YJM1387 | XII | 2121 | 2130 | 2118914 | middle |  |  |  |  |  |  |  |  |  |  |  |
| YJM1387 | XII | 2116674 | 2116683 | 2118914 | middle |  |  |  |  |  |  |  |  |  |  |  |
| YJM1387 | XII | 2117122 | 2117131 | 2118914 | middle |  |  |  |  |  |  |  |  |  |  |  |
| YJM1387 | XII | 2118914 | 2118914 | 2118914 | rightend_complete | YLR460C | 99.558 | 1131 | 5 | 0 | 1 | 1131 | 2114283 | 2113153 | 0 | 2061 |
| YJM1387 | XIII | 1 | 10 | 902057 | leftend | YML131W | 100 | 1098 | 0 | 0 | 1 | 1098 | 4094 | 5191 | 0 | 2028 |
| YJM1387 | XIII | 559 | 568 | 902057 | middle |  |  |  |  |  |  |  |  |  |  |  |
| YJM1387 | XIII | 1075 | 1084 | 902057 | middle |  |  |  |  |  |  |  |  |  |  |  |
| YJM1387 | XIII | 896335 | 896344 | 902057 | middle |  |  |  |  |  |  |  |  |  |  |  |
| YJM1387 | XIII | 902048 | 902057 | 902057 | rightend | YMR321C | 99.361 | 313 | 2 | 0 | 6 | 318 | 889876 | 889564 | 2.48E-161 | 568 |
| YJM1387 | XIV | 1 | 10 | 733697 | leftend | YNL332W | 98.534 | 1023 | 15 | 0 | 1 | 1023 | 7124 | 8146 | 0 | 1807 |
| YJM1387 | XIV | 485 | 494 | 733697 | middle |  |  |  |  |  |  |  |  |  |  |  |
| YJM1387 | XIV | 731093 | 731102 | 733697 | middle |  |  |  |  |  |  |  |  |  |  |  |
| YJM1387 | XIV | 733697 | 733697 | 733697 | rightend_complete | YNR065C | 98.48 | 1908 | 28 | 1 | 1445 | 3351 | 726328 | 724421 | 0 | 3362 |
| YJM1387 | XV | 1 | 10 | 1104070 | leftend | YOL154W | 99.467 | 750 | 4 | 0 | 1 | 750 | 9196 | 9945 | 0 | 1363 |
| YJM1387 | XV | 374 | 383 | 1104070 | middle |  |  |  |  |  |  |  |  |  |  |  |
| YJM1387 | XV | 1104070 | 1104070 | 1104070 | rightend_complete | YOR381W | 98.408 | 2136 | 27 | 1 | 1 | 2136 | 1008757 | 1010885 | 0 | 3749 |
| YJM1387 | XVI | 1 | 10 | 908756 | leftend | YPL274W | 98.583 | 1764 | 25 | 0 | 1 | 1764 | 12445 | 14208 | 0 | 3120 |
| YJM1387 | XVI | 1338 | 1347 | 908756 | middle |  |  |  |  |  |  |  |  |  |  |  |
| YJM1387 | XVI | 901452 | 901461 | 908756 | middle |  |  |  |  |  |  |  |  |  |  |  |
| YJM1387 | XVI | 902175 | 902184 | 908756 | middle |  |  |  |  |  |  |  |  |  |  |  |
| YJM1387 | XVI | 907666 | 907675 | 908756 | middle |  |  |  |  |  |  |  |  |  |  |  |
| YJM1387 | XVI | 908747 | 908756 | 908756 | rightend | YPR201W | 100 | 1215 | 0 | 0 | 1 | 1215 | 898342 | 899556 | 0 | 2244 |
| YJM1388 | I | 1 | 10 | 190219 | leftend | YAL063C | 84.672 | 1768 | 265 | 6 | 1 | 1765 | 14090 | 12326 | 0 | 1759 |
| YJM1388 | I | 502 | 511 | 190219 | middle |  |  |  |  |  |  |  |  |  |  |  |
| YJM1388 | I | 11250 | 11259 | 190219 | middle |  |  |  |  |  |  |  |  |  |  |  |
| YJM1388 | I | 190210 | 190219 | 190219 | rightend | YAR050W | 86.409 | 1994 | 221 | 16 | 2329 | 4278 | 13247 | 11260 | 0 | 2135 |
| YJM1388 | II | 1 | 10 | 784547 | leftend | YNCB0001W | 100 | 84 | 0 | 0 | 1 | 84 | 2263 | 2346 | 4.16E-38 | 156 |
| YJM1388 | II | 270 | 279 | 784547 | middle |  |  |  |  |  |  |  |  |  |  |  |
| YJM1388 | II | 961 | 970 | 784547 | middle |  |  |  |  |  |  |  |  |  |  |  |
| YJM1388 | II | 784538 | 784547 | 784547 | rightend | YBR297W | 97.925 | 1205 | 25 | 0 | 1 | 1205 | 774686 | 775890 | 0 | 2087 |
| YJM1388 | III | 1 | 10 | 297344 | leftend | YCL073C | 98.81 | 1848 | 22 | 0 | 1 | 1848 | 4109 | 2262 | 0 | 3291 |
| YJM1388 | III | 297335 | 297344 | 297344 | rightend | YCR101C | 98 | 550 | 10 | 1 | 1 | 549 | 292554 | 292005 | 0 | 953 |
| YJM1388 | IV | 1 | 10 | 1465402 | leftend | YDL243C | 93.38 | 997 | 59 | 4 | 1 | 990 | 16184 | 15188 | 0 | 1469 |
| YJM1388 | IV | 1465393 | 1465402 | 1465402 | rightend | YDR541C | 99.517 | 1035 | 5 | 0 | 1 | 1035 | 1462599 | 1461565 | 0 | 1884 |
| YJM1388 | V | 1 | 10 | 550985 | leftend | YEL070W | 99.47 | 1509 | 8 | 0 | 1 | 1509 | 14466 | 15974 | 0 | 2743 |
| YJM1388 | V | 550985 | 550985 | 550985 | rightend_complete | YER187W | 99.765 | 426 | 1 | 0 | 1 | 426 | 546823 | 547248 | 0 | 782 |
| YJM1388 | VI | 1 | 10 | 271644 | leftend | YFL051C | 99.793 | 483 | 1 | 0 | 1 | 483 | 33091 | 32609 | 0 | 887 |
| YJM1388 | VI | 1214 | 1223 | 271644 | middle |  |  |  |  |  |  |  |  |  |  |  |
| YJM1388 | VI | 2882 | 2891 | 271644 | middle |  |  |  |  |  |  |  |  |  |  |  |
| YJM1388 | VI | 4786 | 4795 | 271644 | middle |  |  |  |  |  |  |  |  |  |  |  |
| YJM1388 | VI | 271635 | 271644 | 271644 | rightend | YFR055W | 99.609 | 1022 | 4 | 0 | 1 | 1022 | 261264 | 262285 | 0 | 1866 |
| YJM1388 | VII | 1 |  | 1041329 | leftend_complete | YGL258W | 99.356 | 621 | 4 | 0 | 1 | 621 | 10677 | 11297 | 0 | 1125 |
| YJM1388 | VII | 6658 | 6667 | 1041329 | middle |  |  |  |  |  |  |  |  |  |  |  |
| YJM1388 | VII | 1038156 | 1038165 | 1041329 | middle |  |  |  |  |  |  |  |  |  |  |  |
| YJM1388 | VII | 1040252 | 1040261 | 1041329 | middle |  |  |  |  |  |  |  |  |  |  |  |
| YJM1388 | VII | 1041329 | 1041329 | 1041329 | rightend_complete | YGR287C | 99.04 | 1770 | 17 | 0 | 1 | 1770 | 1032478 | 1030709 | 0 | 3175 |
| YJM1388 | VIII | 1 | 10 | 523691 | leftend | YHL044W | 98.588 | 708 | 10 | 0 | 1 | 708 | 7862 | 8569 | 0 | 1253 |
| YJM1388 | VIII | 513111 | 513120 | 523691 | middle |  |  |  |  |  |  |  |  |  |  |  |
| YJM1388 | VIII | 523682 | 523691 | 523691 | rightend | YHR211W | 86.867 | 967 | 105 | 14 | 1 | 947 | 511451 | 512415 | 0 | 1062 |
| YJM1388 | IX | 1 |  | 411555 | leftend_complete | YIL169C | 98.197 | 1442 | 20 | 2 | 478 | 1919 | 7986 | 6551 | 0 | 2514 |
| YJM1388 | IX | 601 | 610 | 411555 | middle |  |  |  |  |  |  |  |  |  |  |  |
| YJM1388 | IX | 1356 | 1365 | 411555 | middle |  |  |  |  |  |  |  |  |  |  |  |
| YJM1388 | IX | 410339 | 410348 | 411555 | middle |  |  |  |  |  |  |  |  |  |  |  |
| YJM1388 | IX | 411310 | 411319 | 411555 | middle |  |  |  |  |  |  |  |  |  |  |  |
| YJM1388 | IX | 411546 | 411555 | 411555 | rightend | YIR039C | 99.69 | 1614 | 5 | 0 | 1 | 1614 | 408570 | 406957 | 0 | 2953 |
| YJM1388 | X | 1 | 10 | 717545 | leftend | YJL213W | 99.9 | 996 | 1 | 0 | 1 | 996 | 27775 | 28770 | 0 | 1834 |
| YJM1388 | X | 1780 | 1789 | 717545 | middle |  |  |  |  |  |  |  |  |  |  |  |
| YJM1388 | X | 3573 | 3582 | 717545 | middle |  |  |  |  |  |  |  |  |  |  |  |
| YJM1388 | X | 3970 | 3979 | 717545 | middle |  |  |  |  |  |  |  |  |  |  |  |
| YJM1388 | X | 4922 | 4931 | 717545 | middle |  |  |  |  |  |  |  |  |  |  |  |
| YJM1388 | X | 5172 | 5181 | 717545 | middle |  |  |  |  |  |  |  |  |  |  |  |
| YJM1388 | X | 5975 | 5984 | 717545 | middle |  |  |  |  |  |  |  |  |  |  |  |
| YJM1388 | X | 717536 | 717545 | 717545 | rightend | YJR152W | 99.51 | 1632 | 8 | 0 | 1 | 1632 | 703231 | 704862 | 0 | 2970 |
| YJM1388 | XI | 1 | 10 | 656049 | leftend | YKL222C | 99.858 | 2118 | 3 | 0 | 1 | 2118 | 7440 | 5323 | 0 | 3895 |
| YJM1388 | XI | 656040 | 656049 | 656049 | rightend | YKR101W | 99.593 | 1965 | 8 | 0 | 1 | 1965 | 640824 | 642788 | 0 | 3585 |
| YJM1388 | XII | 1 | 10 | 1922188 | leftend | YLL062C | 99.795 | 975 | 2 | 0 | 1 | 975 | 4103 | 3129 | 0 | 1790 |
| YJM1388 | XII | 1920291 | 1920300 | 1922188 | middle |  |  |  |  |  |  |  |  |  |  |  |
| YJM1388 | XII | 1920751 | 1920760 | 1922188 | middle |  |  |  |  |  |  |  |  |  |  |  |
| YJM1388 | XII | 1922188 | 1922188 | 1922188 | rightend_complete | YLR460C | 99.646 | 1131 | 4 | 0 | 1 | 1131 | 1918642 | 1917512 | 0 | 2067 |
| YJM1388 | XIII | 1 | 10 | 890133 | leftend | YML131W | 98.998 | 1098 | 11 | 0 | 1 | 1098 | 4452 | 5549 | 0 | 1967 |
| YJM1388 | XIII | 890124 | 890133 | 890133 | rightend | YMR321C | 99.371 | 318 | 2 | 0 | 1 | 318 | 889137 | 888820 | 4.12E-164 | 577 |
| YJM1388 | XIV | 1 | 10 | 763033 | leftend | YNL332W | 99.12 | 1023 | 9 | 0 | 1 | 1023 | 5356 | 6378 | 0 | 1840 |
| YJM1388 | XIV | 1133 | 1142 | 763033 | middle |  |  |  |  |  |  |  |  |  |  |  |
| YJM1388 | XIV | 763033 | 763033 | 763033 | rightend_complete | YNR071C | 99.515 | 1030 | 4 | 1 | 1 | 1029 | 744791 | 743762 | 0 | 1873 |
| YJM1388 | XV | 1 | 10 | 1050534 | leftend | YOL154W | 99.867 | 750 | 1 | 0 | 1 | 750 | 24732 | 25481 | 0 | 1380 |
| YJM1388 | XV | 1049986 | 1049995 | 1050534 | middle |  |  |  |  |  |  |  |  |  |  |  |
| YJM1388 | XV | 1050534 | 1050534 | 1050534 | rightend_complete | YOR381W | 98.689 | 2136 | 28 | 0 | 1 | 2136 | 1023297 | 1025432 | 0 | 3790 |
| YJM1388 | XVI | 1 |  | 910633 | leftend_complete | YPL274W | 98.526 | 1764 | 26 | 0 | 1 | 1764 | 15485 | 17248 | 0 | 3114 |
| YJM1388 | XVI | 789 | 798 | 910633 | middle |  |  |  |  |  |  |  |  |  |  |  |
| YJM1388 | XVI | 910633 | 910633 | 910633 | rightend_complete | YPR201W | 99.671 | 1215 | 4 | 0 | 1 | 1215 | 907321 | 908535 | 0 | 2222 |
| YJM1389 | I | 1 | 10 | 192335 | leftend | YAL063C | 87.69 | 1779 | 156 | 34 | 1912 | 3660 | 190580 | 192325 | 0 | 2013 |
| YJM1389 | I | 274 | 283 | 192335 | middle |  |  |  |  |  |  |  |  |  |  |  |
| YJM1389 | I | 3681 | 3690 | 192335 | middle |  |  |  |  |  |  |  |  |  |  |  |
| YJM1389 | I | 192326 | 192335 | 192335 | rightend | YAR050W | 91.477 | 1760 | 94 | 12 | 2574 | 4278 | 190567 | 192325 | 0 | 2368 |
| YJM1389 | II | 1 | 10 | 783602 | leftend | YNCB0001W | 100 | 84 | 0 | 0 | 1 | 84 | 2203 | 2286 | 4.16E-38 | 156 |
| YJM1389 | II | 619 | 628 | 783188 | middle |  |  |  |  |  |  |  |  |  |  |  |
| YJM1389 | II | 783593 | 783602 | 783602 | rightend | YBR297W | 97.925 | 1205 | 25 | 0 | 1 | 1205 | 773378 | 774582 | 0 | 2087 |
| YJM1389 | III | 1 | 10 | 300382 | leftend | YCL073C | 98.755 | 1848 | 23 | 0 | 1 | 1848 | 6946 | 5099 | 0 | 3286 |
| YJM1389 | III | 300373 | 300382 | 300382 | rightend | YCR101C | 98 | 550 | 10 | 1 | 1 | 549 | 295533 | 294984 | 0 | 953 |
| YJM1389 | IV | 1 | 10 | 1470174 | leftend | YDL243C | 93.838 | 990 | 61 | 0 | 1 | 990 | 16179 | 15190 | 0 | 1491 |
| YJM1389 | IV | 1470165 | 1470174 | 1470174 | rightend | YDR541C | 99.517 | 1035 | 5 | 0 | 1 | 1035 | 1458303 | 1457269 | 0 | 1884 |
| YJM1389 | V | 1 | 10 | 551257 | leftend | YEL070W | 99.47 | 1509 | 8 | 0 | 1 | 1509 | 14466 | 15974 | 0 | 2743 |
| YJM1389 | V | 551257 | 551257 | 551257 | rightend_complete | YER187W | 99.765 | 426 | 1 | 0 | 1 | 426 | 547096 | 547521 | 0 | 782 |
| YJM1389 | VI | 1 |  | 241334 | leftend_complete | YFL050C | 99.321 | 1472 | 10 | 0 | 1 | 1472 | 6179 | 4708 | 0 | 2663 |
| YJM1389 | VI | 1329 | 1338 | 240546 | middle |  |  |  |  |  |  |  |  |  |  |  |
| YJM1389 | VI | 1556 | 1565 | 240546 | middle |  |  |  |  |  |  |  |  |  |  |  |
| YJM1389 | VI | 240063 | 240072 | 240546 | middle |  |  |  |  |  |  |  |  |  |  |  |
| YJM1389 | VI | 241334 | 241334 | 241334 | rightend_complete | YFR055W | 99.609 | 1022 | 4 | 0 | 1 | 1022 | 228892 | 229913 | 0 | 1866 |
| YJM1389 | VII | 1 | 10 | 1053418 | leftend | YGL258W | 99.195 | 621 | 5 | 0 | 1 | 621 | 7726 | 8346 | 0 | 1120 |
| YJM1389 | VII | 899 | 908 | 1053418 | middle |  |  |  |  |  |  |  |  |  |  |  |
| YJM1389 | VII | 3343 | 3352 | 1053418 | middle |  |  |  |  |  |  |  |  |  |  |  |
| YJM1389 | VII | 1050374 | 1050383 | 1053418 | middle |  |  |  |  |  |  |  |  |  |  |  |
| YJM1389 | VII | 1053409 | 1053418 | 1053418 | rightend | YGR287C | 99.04 | 1770 | 17 | 0 | 1 | 1770 | 1034511 | 1032742 | 0 | 3175 |
| YJM1389 | VIII | 1 |  | 513935 | leftend_complete | YHL044W | 98.588 | 708 | 10 | 0 | 1 | 708 | 9422 | 10129 | 0 | 1253 |
| YJM1389 | VIII | 1292 | 1301 | 513935 | middle |  |  |  |  |  |  |  |  |  |  |  |
| YJM1389 | VIII | 513926 | 513935 | 513935 | rightend | YHR210C | 99.025 | 1026 | 8 | 1 | 1 | 1026 | 507803 | 506780 | 0 | 1838 |
| YJM1389 | IX | 1 | 10 | 422114 | leftend | YIL169C | 98.45 | 2322 | 30 | 2 | 478 | 2799 | 17634 | 15319 | 0 | 4084 |
| YJM1389 | IX | 420097 | 420106 | 422114 | middle |  |  |  |  |  |  |  |  |  |  |  |
| YJM1389 | IX | 420615 | 420624 | 422114 | middle |  |  |  |  |  |  |  |  |  |  |  |
| YJM1389 | IX | 422114 | 422114 | 422114 | rightend_complete | YIR039C | 99.69 | 1614 | 5 | 0 | 1 | 1614 | 418636 | 417023 | 0 | 2953 |
| YJM1389 | X | 1 | 10 | 721937 | leftend | YJL213W | 99.9 | 996 | 1 | 0 | 1 | 996 | 21817 | 22812 | 0 | 1834 |
| YJM1389 | X | 1256 | 1265 | 721937 | middle |  |  |  |  |  |  |  |  |  |  |  |
| YJM1389 | X | 2942 | 2951 | 721937 | middle |  |  |  |  |  |  |  |  |  |  |  |
| YJM1389 | X | 4402 | 4411 | 721937 | middle |  |  |  |  |  |  |  |  |  |  |  |
| YJM1389 | X | 721937 | 721937 | 721937 | rightend_complete | YJR152W | 99.449 | 1632 | 9 | 0 | 1 | 1632 | 692155 | 693786 | 0 | 2964 |
| YJM1389 | XI | 1 | 10 | 664729 | leftend | YKL222C | 99.858 | 2118 | 3 | 0 | 1 | 2118 | 3333 | 1216 | 0 | 3895 |
| YJM1389 | XI | 664720 | 664729 | 664729 | rightend | YKR101W | 99.644 | 1965 | 7 | 0 | 1 | 1965 | 637346 | 639310 | 0 | 3591 |
| YJM1389 | XII | 1 | 10 | 1650334 | leftend | YLL062C | 99.795 | 975 | 2 | 0 | 1 | 975 | 4157 | 3183 | 0 | 1790 |
| YJM1389 | XII | 1648908 | 1648917 | 1650334 | middle |  |  |  |  |  |  |  |  |  |  |  |
| YJM1389 | XII | 1650334 | 1650334 | 1650334 | rightend_complete | YLR460C | 100 | 1131 | 0 | 0 | 1 | 1131 | 1647125 | 1645995 | 0 | 2089 |
| YJM1389 | XIII | 1 | 10 | 890662 | leftend | YML131W | 98.998 | 1098 | 11 | 0 | 1 | 1098 | 4613 | 5710 | 0 | 1967 |
| YJM1389 | XIII | 890653 | 890662 | 890662 | rightend | YMR321C | 99.371 | 318 | 2 | 0 | 1 | 318 | 889574 | 889257 | 4.12E-164 | 577 |
| YJM1389 | XIV | 1 | 10 | 754162 | leftend | YNL332W | 99.413 | 1023 | 6 | 0 | 1 | 1023 | 11968 | 12990 | 0 | 1857 |
| YJM1389 | XIV | 4533 | 4542 | 754815 | middle |  |  |  |  |  |  |  |  |  |  |  |
| YJM1389 | XIV | 5417 | 5426 | 754815 | middle |  |  |  |  |  |  |  |  |  |  |  |
| YJM1389 | XIV | 6070 | 6079 | 754815 | middle |  |  |  |  |  |  |  |  |  |  |  |
| YJM1389 | XIV | 754153 | 754162 | 754162 | rightend | YNR071C | 99.515 | 1030 | 4 | 1 | 1 | 1029 | 752095 | 751066 | 0 | 1873 |
| YJM1389 | XV | 1 | 10 | 1035246 | leftend | YOL154W | 99.6 | 750 | 3 | 0 | 1 | 750 | 8233 | 8982 | 0 | 1369 |
| YJM1389 | XV | 1035237 | 1035246 | 1035246 | rightend | YOR381W | 98.642 | 2136 | 29 | 0 | 1 | 2136 | 1006372 | 1008507 | 0 | 3784 |
| YJM1389 | XVI | 1 | 10 | 896221 | leftend | YPL274W | 98.526 | 1764 | 26 | 0 | 1 | 1764 | 5382 | 7145 | 0 | 3114 |
| YJM1389 | XVI | 896221 | 896221 | 896221 | rightend_complete | YPR201W | 99.671 | 1215 | 4 | 0 | 1 | 1215 | 892922 | 894136 | 0 | 2222 |
| YJM1399 | I | 1 | 10 | 207219 | leftend | YAL063C | 88.143 | 1147 | 132 | 4 | 1 | 1145 | 14101 | 12957 | 0 | 1362 |
| YJM1399 | I | 12945 | 12954 | 207219 | middle |  |  |  |  |  |  |  |  |  |  |  |
| YJM1399 | I | 192542 | 192551 | 207219 | middle |  |  |  |  |  |  |  |  |  |  |  |
| YJM1399 | I | 193241 | 193250 | 207219 | middle |  |  |  |  |  |  |  |  |  |  |  |
| YJM1399 | I | 207210 | 207219 | 207219 | rightend | YAR050W | 83.834 | 1497 | 222 | 15 | 1 | 1487 | 190518 | 192004 | 0 | 1406 |
| YJM1399 | II | 1 | 10 | 797654 | leftend | YNCB0001W | 100 | 84 | 0 | 0 | 1 | 84 | 6534 | 6617 | 4.16E-38 | 156 |
| YJM1399 | II | 919 | 928 | 797654 | middle |  |  |  |  |  |  |  |  |  |  |  |
| YJM1399 | II | 1175 | 1184 | 797654 | middle |  |  |  |  |  |  |  |  |  |  |  |
| YJM1399 | II | 4952 | 4961 | 797654 | middle |  |  |  |  |  |  |  |  |  |  |  |
| YJM1399 | II | 797645 | 797654 | 797654 | rightend | YBR297W | 84.483 | 1334 | 203 | 3 | 1 | 1332 | 780028 | 781359 | 0 | 1314 |
| YJM1399 | III | 1 | 10 | 299305 | leftend | YCL073C | 99.188 | 1848 | 15 | 0 | 1 | 1848 | 4036 | 2189 | 0 | 3330 |
| YJM1399 | III | 290921 | 290930 | 299305 | middle |  |  |  |  |  |  |  |  |  |  |  |
| YJM1399 | III | 291155 | 291164 | 299305 | middle |  |  |  |  |  |  |  |  |  |  |  |
| YJM1399 | III | 299296 | 299305 | 299305 | rightend | YCR100C | 92.813 | 487 | 33 | 2 | 463 | 948 | 290919 | 290434 | 0 | 704 |
| YJM1399 | IV | 1 | 10 | 1466740 | leftend | YDL243C | 94.242 | 990 | 57 | 0 | 1 | 990 | 17864 | 16875 | 0 | 1513 |
| YJM1399 | IV | 1465969 | 1465978 | 1466740 | middle |  |  |  |  |  |  |  |  |  |  |  |
| YJM1399 | IV | 1466195 | 1466204 | 1466740 | middle |  |  |  |  |  |  |  |  |  |  |  |
| YJM1399 | IV | 1466468 | 1466477 | 1466740 | middle |  |  |  |  |  |  |  |  |  |  |  |
| YJM1399 | IV | 1466731 | 1466740 | 1466740 | rightend | YDR541C | 99.42 | 1035 | 6 | 0 | 1 | 1035 | 1457300 | 1456266 | 0 | 1879 |
| YJM1399 | V | 1 | 10 | 551869 | leftend | YEL070W | 98.608 | 1509 | 21 | 0 | 1 | 1509 | 13195 | 14703 | 0 | 2671 |
| YJM1399 | V | 551869 | 551869 | 551869 | rightend_complete | YER187W | 98.357 | 426 | 7 | 0 | 1 | 426 | 547880 | 548305 | 0 | 749 |
| YJM1399 | VI | 1 | 10 | 266830 | leftend | YFL051C | 98.009 | 452 | 9 | 0 | 1 | 452 | 36045 | 35594 | 0 | 785 |
| YJM1399 | VI | 266821 | 266830 | 266830 | rightend | YFR055W | 86.092 | 1021 | 135 | 5 | 1 | 1016 | 262094 | 263112 | 0 | 1092 |
| YJM1399 | VII | 1 | 10 | 1057805 | leftend | YGL258W | 97.585 | 621 | 15 | 0 | 1 | 621 | 3244 | 3864 | 0 | 1064 |
| YJM1399 | VII | 1057796 | 1057805 | 1057805 | rightend | YGR287C | 99.096 | 1770 | 16 | 0 | 1 | 1770 | 1023865 | 1022096 | 0 | 3181 |
| YJM1399 | VIII | 1 | 10 | 530699 | leftend | YHL044W | 98.305 | 708 | 12 | 0 | 1 | 708 | 6583 | 7290 | 0 | 1242 |
| YJM1399 | VIII | 527164 | 527173 | 530699 | middle |  |  |  |  |  |  |  |  |  |  |  |
| YJM1399 | VIII | 527454 | 527463 | 530699 | middle |  |  |  |  |  |  |  |  |  |  |  |
| YJM1399 | VIII | 528824 | 528833 | 530699 | middle |  |  |  |  |  |  |  |  |  |  |  |
| YJM1399 | VIII | 529120 | 529129 | 530699 | middle |  |  |  |  |  |  |  |  |  |  |  |
| YJM1399 | VIII | 530690 | 530699 | 530699 | rightend | YHR211W | 81.435 | 1756 | 311 | 12 | 13 | 1762 | 509705 | 511451 | 0 | 1423 |
| YJM1399 | IX | 1 | 10 | 415616 | leftend | YIL169C | 95.026 | 1568 | 42 | 6 | 352 | 1919 | 9565 | 8034 | 0 | 2431 |
| YJM1399 | IX | 449 | 458 | 415616 | middle |  |  |  |  |  |  |  |  |  |  |  |
| YJM1399 | IX | 703 | 712 | 415616 | middle |  |  |  |  |  |  |  |  |  |  |  |
| YJM1399 | IX | 931 | 940 | 415616 | middle |  |  |  |  |  |  |  |  |  |  |  |
| YJM1399 | IX | 413840 | 413849 | 415616 | middle |  |  |  |  |  |  |  |  |  |  |  |
| YJM1399 | IX | 415384 | 415393 | 415616 | middle |  |  |  |  |  |  |  |  |  |  |  |
| YJM1399 | IX | 415616 | 415616 | 415616 | rightend_complete | YIR039C | 98.885 | 1614 | 18 | 0 | 1 | 1614 | 408351 | 406738 | 0 | 2881 |
| YJM1399 | X | 1 | 10 | 715078 | leftend | YJL213W | 99.297 | 996 | 7 | 0 | 1 | 996 | 22357 | 23352 | 0 | 1801 |
| YJM1399 | X | 520 | 529 | 715078 | middle |  |  |  |  |  |  |  |  |  |  |  |
| YJM1399 | X | 877 | 886 | 715078 | middle |  |  |  |  |  |  |  |  |  |  |  |
| YJM1399 | X | 715078 | 715078 | 715078 | rightend_complete | YJR152W | 99.877 | 1632 | 2 | 0 | 1 | 1632 | 693207 | 694838 | 0 | 3003 |
| YJM1399 | XI | 1 | 10 | 664222 | leftend | YKL222C | 99.103 | 2118 | 19 | 0 | 1 | 2118 | 3090 | 973 | 0 | 3807 |
| YJM1399 | XI | 664213 | 664222 | 664222 | rightend | YKR101W | 98.984 | 1968 | 17 | 1 | 1 | 1965 | 638927 | 640894 | 0 | 3520 |
| YJM1399 | XII | 1 | 10 | 1697961 | leftend | YLL062C | 99.59 | 975 | 4 | 0 | 1 | 975 | 4385 | 3411 | 0 | 1779 |
| YJM1399 | XII | 1697539 | 1697548 | 1697961 | middle |  |  |  |  |  |  |  |  |  |  |  |
| YJM1399 | XII | 1697952 | 1697961 | 1697961 | rightend | YLR460C | 98.408 | 1131 | 18 | 0 | 1 | 1131 | 1696484 | 1695354 | 0 | 1989 |
| YJM1399 | XIII | 1 | 10 | 918295 | leftend | YML131W | 100 | 1098 | 0 | 0 | 1 | 1098 | 3213 | 4310 | 0 | 2028 |
| YJM1399 | XIII | 846 | 855 | 918295 | middle |  |  |  |  |  |  |  |  |  |  |  |
| YJM1399 | XIII | 918286 | 918295 | 918295 | rightend | YMR321C | 99.042 | 313 | 3 | 0 | 6 | 318 | 893273 | 892961 | 1.15E-159 | 562 |
| YJM1399 | XIV | 1 |  | 762042 | leftend_complete | YNL332W | 98.827 | 1023 | 12 | 0 | 1 | 1023 | 12308 | 13330 | 0 | 1823 |
| YJM1399 | XIV | 6807 | 6816 | 762042 | middle |  |  |  |  |  |  |  |  |  |  |  |
| YJM1399 | XIV | 762033 | 762042 | 762042 | rightend | YNR071C | 98.834 | 1029 | 12 | 0 | 1 | 1029 | 751745 | 750717 | 0 | 1834 |
| YJM1399 | XV | 1 | 10 | 1044812 | leftend | YOL154W | 95.2 | 750 | 36 | 0 | 1 | 750 | 18981 | 19730 | 0 | 1186 |
| YJM1399 | XV | 494 | 503 | 1044812 | middle |  |  |  |  |  |  |  |  |  |  |  |
| YJM1399 | XV | 1044309 | 1044318 | 1044812 | middle |  |  |  |  |  |  |  |  |  |  |  |
| YJM1399 | XV | 1044812 | 1044812 | 1044812 | rightend_complete | YOR381W | 99.251 | 2136 | 16 | 0 | 1 | 2136 | 1016069 | 1018204 | 0 | 3856 |
| YJM1399 | XVI | 1 |  | 903281 | leftend_complete | YPL274W | 99.546 | 1764 | 8 | 0 | 1 | 1764 | 10468 | 12231 | 0 | 3214 |
| YJM1399 | XVI | 438 | 447 | 903281 | middle |  |  |  |  |  |  |  |  |  |  |  |
| YJM1399 | XVI | 707 | 716 | 903281 | middle |  |  |  |  |  |  |  |  |  |  |  |
| YJM1399 | XVI | 903272 | 903281 | 903281 | rightend | YPR201W | 99.588 | 1215 | 5 | 0 | 1 | 1215 | 900461 | 901675 | 0 | 2217 |
| YJM1400 | I | 1 | 10 | 185163 | leftend | YAL063C | 84.264 | 1271 | 156 | 26 | 1777 | 3037 | 179753 | 180989 | 0 | 1199 |
| YJM1400 | I | 879 | 888 | 185163 | middle |  |  |  |  |  |  |  |  |  |  |  |
| YJM1400 | I | 1460 | 1469 | 185163 | middle |  |  |  |  |  |  |  |  |  |  |  |
| YJM1400 | I | 179699 | 179708 | 185163 | middle |  |  |  |  |  |  |  |  |  |  |  |
| YJM1400 | I | 181462 | 181471 | 185163 | middle |  |  |  |  |  |  |  |  |  |  |  |
| YJM1400 | I | 185154 | 185163 | 185163 | rightend | YAR050W | 86.413 | 1288 | 149 | 13 | 2408 | 3676 | 179709 | 180989 | 0 | 1386 |
| YJM1400 | II | 1 | 10 | 780974 | leftend | YNCB0001W | 100 | 84 | 0 | 0 | 1 | 84 | 2724 | 2807 | 4.16E-38 | 156 |
| YJM1400 | II | 1207 | 1216 | 780974 | middle |  |  |  |  |  |  |  |  |  |  |  |
| YJM1400 | II | 780965 | 780974 | 780974 | rightend | YBR297W | 97.842 | 1205 | 26 | 0 | 1 | 1205 | 776774 | 777978 | 0 | 2082 |
| YJM1400 | III | 1 |  | 313944 | leftend_complete | YCL073C | 98.755 | 1848 | 23 | 0 | 1 | 1848 | 10873 | 9026 | 0 | 3286 |
| YJM1400 | III | 3378 | 3387 | 313944 | middle |  |  |  |  |  |  |  |  |  |  |  |
| YJM1400 | III | 305095 | 305104 | 313944 | middle |  |  |  |  |  |  |  |  |  |  |  |
| YJM1400 | III | 313944 | 313944 | 313944 | rightend_complete | YCR101C | 98.727 | 550 | 6 | 1 | 1 | 549 | 300510 | 299961 | 0 | 976 |
| YJM1400 | IV | 1 | 10 | 1454635 | leftend | YDL243C | 92.072 | 883 | 68 | 2 | 1 | 882 | 892 | 11 | 0 | 1242 |
| YJM1400 | IV | 1453220 | 1453229 | 1454635 | middle |  |  |  |  |  |  |  |  |  |  |  |
| YJM1400 | IV | 1454626 | 1454635 | 1454635 | rightend | YDR541C | 99.807 | 1035 | 2 | 0 | 1 | 1035 | 1448236 | 1447202 | 0 | 1901 |
| YJM1400 | V | 1 | 10 | 551189 | leftend | YEL070W | 99.204 | 1507 | 12 | 0 | 3 | 1509 | 12755 | 14261 | 0 | 2717 |
| YJM1400 | V | 549729 | 549738 | 551189 | middle |  |  |  |  |  |  |  |  |  |  |  |
| YJM1400 | V | 551189 | 551189 | 551189 | rightend_complete | YER187W | 98.122 | 426 | 8 | 0 | 1 | 426 | 546195 | 546620 | 0 | 743 |
| YJM1400 | VI | 1 | 10 | 285869 | leftend | YFL051C | 98.23 | 452 | 8 | 0 | 1 | 452 | 47046 | 46595 | 0 | 791 |
| YJM1400 | VI | 285860 | 285869 | 285869 | rightend | YFR055W | 99.609 | 1022 | 4 | 0 | 1 | 1022 | 275627 | 276648 | 0 | 1866 |
| YJM1400 | VII | 1 | 10 | 1053087 | leftend | YGL258W | 93.72 | 621 | 39 | 0 | 1 | 621 | 9857 | 10477 | 0 | 931 |
| YJM1400 | VII | 6056 | 6065 | 1053087 | middle |  |  |  |  |  |  |  |  |  |  |  |
| YJM1400 | VII | 1042744 | 1042753 | 1053087 | middle |  |  |  |  |  |  |  |  |  |  |  |
| YJM1400 | VII | 1044386 | 1044395 | 1053087 | middle |  |  |  |  |  |  |  |  |  |  |  |
| YJM1400 | VII | 1051996 | 1052005 | 1053087 | middle |  |  |  |  |  |  |  |  |  |  |  |
| YJM1400 | VII | 1053078 | 1053087 | 1053087 | rightend | YGR287C | 99.887 | 1770 | 2 | 0 | 1 | 1770 | 1032838 | 1031069 | 0 | 3258 |
| YJM1400 | VIII | 1 | 10 | 549818 | leftend | YHL044W | 99.011 | 708 | 7 | 0 | 1 | 708 | 7983 | 8690 | 0 | 1269 |
| YJM1400 | VIII | 522906 | 522915 | 549818 | middle |  |  |  |  |  |  |  |  |  |  |  |
| YJM1400 | VIII | 538491 | 538500 | 549818 | middle |  |  |  |  |  |  |  |  |  |  |  |
| YJM1400 | VIII | 539494 | 539503 | 549818 | middle |  |  |  |  |  |  |  |  |  |  |  |
| YJM1400 | VIII | 549200 | 549209 | 549818 | middle |  |  |  |  |  |  |  |  |  |  |  |
| YJM1400 | VIII | 549809 | 549818 | 549818 | rightend | YHR211W | 84.644 | 1335 | 194 | 7 | 1 | 1328 | 521575 | 522905 | 0 | 1319 |
| YJM1400 | IX | 1 | 10 | 426254 | leftend | YIL169C | 90.732 | 1284 | 104 | 10 | 612 | 1880 | 17201 | 15918 | 0 | 1698 |
| YJM1400 | IX | 1265 | 1274 | 426254 | middle |  |  |  |  |  |  |  |  |  |  |  |
| YJM1400 | IX | 2360 | 2369 | 426254 | middle |  |  |  |  |  |  |  |  |  |  |  |
| YJM1400 | IX | 425969 | 425978 | 426254 | middle |  |  |  |  |  |  |  |  |  |  |  |
| YJM1400 | IX | 426245 | 426254 | 426254 | rightend | YIR039C | 99.69 | 1614 | 5 | 0 | 1 | 1614 | 419245 | 417632 | 0 | 2953 |
| YJM1400 | X | 1 |  | 693032 | leftend_complete | YJL213W | 98.896 | 996 | 11 | 0 | 1 | 996 | 9863 | 10858 | 0 | 1779 |
| YJM1400 | X | 1427 | 1436 | 693032 | middle |  |  |  |  |  |  |  |  |  |  |  |
| YJM1400 | X | 691543 | 691552 | 693032 | middle |  |  |  |  |  |  |  |  |  |  |  |
| YJM1400 | X | 691782 | 691791 | 693032 | middle |  |  |  |  |  |  |  |  |  |  |  |
| YJM1400 | X | 692781 | 692790 | 693032 | middle |  |  |  |  |  |  |  |  |  |  |  |
| YJM1400 | X | 693023 | 693032 | 693032 | rightend | YJR152W | 99.571 | 1632 | 7 | 0 | 1 | 1632 | 680350 | 681981 | 0 | 2976 |
| YJM1400 | XI | 1 | 10 | 663815 | leftend | YKL222C | 99.669 | 2118 | 7 | 0 | 1 | 2118 | 3529 | 1412 | 0 | 3873 |
| YJM1400 | XI | 663806 | 663815 | 663815 | rightend | YKR101W | 99.389 | 1965 | 12 | 0 | 1 | 1965 | 638528 | 640492 | 0 | 3563 |
| YJM1400 | XII | 1 | 10 | 1651531 | leftend | YLL062C | 98.872 | 975 | 11 | 0 | 1 | 975 | 4352 | 3378 | 0 | 1740 |
| YJM1400 | XII | 1650909 | 1650918 | 1651531 | middle |  |  |  |  |  |  |  |  |  |  |  |
| YJM1400 | XII | 1651522 | 1651531 | 1651531 | rightend | YLR460C | 98.587 | 1132 | 14 | 2 | 1 | 1131 | 1649867 | 1648737 | 0 | 2001 |
| YJM1400 | XIII | 1 |  | 918211 | leftend_complete | YML131W | 99.454 | 1098 | 6 | 0 | 1 | 1098 | 4777 | 5874 | 0 | 1995 |
| YJM1400 | XIII | 1155 | 1164 | 918211 | middle |  |  |  |  |  |  |  |  |  |  |  |
| YJM1400 | XIII | 917335 | 917344 | 918211 | middle |  |  |  |  |  |  |  |  |  |  |  |
| YJM1400 | XIII | 918202 | 918211 | 918211 | rightend | YMR321C | 99.361 | 313 | 2 | 0 | 6 | 318 | 892517 | 892205 | 2.48E-161 | 568 |
| YJM1400 | XIV | 1 | 10 | 762820 | leftend | YNL332W | 98.729 | 1023 | 13 | 0 | 1 | 1023 | 6663 | 7685 | 0 | 1818 |
| YJM1400 | XIV | 1154 | 1163 | 762820 | middle |  |  |  |  |  |  |  |  |  |  |  |
| YJM1400 | XIV | 4979 | 4988 | 762820 | middle |  |  |  |  |  |  |  |  |  |  |  |
| YJM1400 | XIV | 762811 | 762820 | 762820 | rightend | YNR071C | 99.514 | 1029 | 5 | 0 | 1 | 1029 | 746267 | 745239 | 0 | 1873 |
| YJM1400 | XV | 1 | 10 | 1057848 | leftend | YOL154W | 99.467 | 750 | 4 | 0 | 1 | 750 | 24072 | 24821 | 0 | 1363 |
| YJM1400 | XV | 1092 | 1101 | 1057848 | middle |  |  |  |  |  |  |  |  |  |  |  |
| YJM1400 | XV | 1057839 | 1057848 | 1057848 | rightend | YOR381W | 98.689 | 2136 | 28 | 0 | 1 | 2136 | 1025080 | 1027215 | 0 | 3790 |
| YJM1400 | XVI | 1 | 10 | 915319 | leftend | YPL274W | 98.243 | 1764 | 31 | 0 | 1 | 1764 | 20375 | 22138 | 0 | 3086 |
| YJM1400 | XVI | 915310 | 915319 | 915319 | rightend | YPR198W | 99.203 | 1632 | 13 | 0 | 1 | 1632 | 906804 | 908435 | 0 | 2942 |
| YJM1401 | I | 1 | 10 | 191492 | leftend | YAL063C | 90.501 | 1916 | 136 | 15 | 2059 | 3969 | 177214 | 179088 | 0 | 2488 |
| YJM1401 | I | 815 | 824 | 191492 | middle |  |  |  |  |  |  |  |  |  |  |  |
| YJM1401 | I | 1496 | 1505 | 191492 | middle |  |  |  |  |  |  |  |  |  |  |  |
| YJM1401 | I | 2380 | 2389 | 191492 | middle |  |  |  |  |  |  |  |  |  |  |  |
| YJM1401 | I | 191483 | 191492 | 191492 | rightend | YAR050W | 90.949 | 1602 | 110 | 19 | 2709 | 4282 | 177189 | 178783 | 0 | 2122 |
| YJM1401 | II | 1 | 10 | 778741 | leftend | YNCB0001W | 100 | 84 | 0 | 0 | 1 | 84 | 2061 | 2144 | 4.16E-38 | 156 |
| YJM1401 | II | 763 | 772 | 778741 | middle |  |  |  |  |  |  |  |  |  |  |  |
| YJM1401 | II | 778732 | 778741 | 778741 | rightend | YBR297W | 97.759 | 1205 | 27 | 0 | 1 | 1205 | 774540 | 775744 | 0 | 2076 |
| YJM1401 | III | 1 | 10 | 310898 | leftend | YCL073C | 98.918 | 1848 | 20 | 0 | 1 | 1848 | 7061 | 5214 | 0 | 3302 |
| YJM1401 | III | 310010 | 310019 | 310898 | middle |  |  |  |  |  |  |  |  |  |  |  |
| YJM1401 | III | 310898 | 310898 | 310898 | rightend_complete | YCR101C | 98.364 | 550 | 8 | 1 | 1 | 549 | 296249 | 295700 | 0 | 965 |
| YJM1401 | IV | 1 | 10 | 1475893 | leftend | YDL243C | 93.769 | 979 | 61 | 0 | 1 | 979 | 26678 | 25700 | 0 | 1471 |
| YJM1401 | IV | 4188 | 4197 | 1475893 | middle |  |  |  |  |  |  |  |  |  |  |  |
| YJM1401 | IV | 1475298 | 1475307 | 1475893 | middle |  |  |  |  |  |  |  |  |  |  |  |
| YJM1401 | IV | 1475884 | 1475893 | 1475893 | rightend | YDR541C | 99.903 | 1035 | 1 | 0 | 1 | 1035 | 1472451 | 1471417 | 0 | 1906 |
| YJM1401 | V | 1 |  | 554967 | leftend_complete | YEL070W | 99.271 | 1509 | 11 | 0 | 1 | 1509 | 15302 | 16810 | 0 | 2726 |
| YJM1401 | V | 1484 | 1493 | 554967 | middle |  |  |  |  |  |  |  |  |  |  |  |
| YJM1401 | V | 554077 | 554086 | 554967 | middle |  |  |  |  |  |  |  |  |  |  |  |
| YJM1401 | V | 554967 | 554967 | 554967 | rightend_complete | YER187W | 99.765 | 426 | 1 | 0 | 1 | 426 | 549916 | 550341 | 0 | 782 |
| YJM1401 | VI | 1 | 10 | 274564 | leftend | YFL051C | 99.115 | 452 | 4 | 0 | 1 | 452 | 36922 | 36471 | 0 | 813 |
| YJM1401 | VI | 274555 | 274564 | 274564 | rightend | YFR055W | 99.609 | 1022 | 4 | 0 | 1 | 1022 | 264900 | 265921 | 0 | 1866 |
| YJM1401 | VII | 1 | 10 | 1056077 | leftend | YGL258W | 99.356 | 621 | 4 | 0 | 1 | 621 | 7832 | 8452 | 0 | 1125 |
| YJM1401 | VII | 5678 | 5687 | 1056077 | middle |  |  |  |  |  |  |  |  |  |  |  |
| YJM1401 | VII | 1055324 | 1055333 | 1056077 | middle |  |  |  |  |  |  |  |  |  |  |  |
| YJM1401 | VII | 1056068 | 1056077 | 1056077 | rightend | YGR287C | 98.644 | 1770 | 24 | 0 | 1 | 1770 | 1031529 | 1029760 | 0 | 3136 |
| YJM1401 | VIII | 1 | 10 | 536281 | leftend | YHL044W | 98.729 | 708 | 9 | 0 | 1 | 708 | 9130 | 9837 | 0 | 1258 |
| YJM1401 | VIII | 1124 | 1133 | 536281 | middle |  |  |  |  |  |  |  |  |  |  |  |
| YJM1401 | VIII | 514496 | 514505 | 536281 | middle |  |  |  |  |  |  |  |  |  |  |  |
| YJM1401 | VIII | 516164 | 516173 | 536281 | middle |  |  |  |  |  |  |  |  |  |  |  |
| YJM1401 | VIII | 529244 | 529253 | 536281 | middle |  |  |  |  |  |  |  |  |  |  |  |
| YJM1401 | VIII | 535165 | 535174 | 536281 | middle |  |  |  |  |  |  |  |  |  |  |  |
| YJM1401 | VIII | 536272 | 536281 | 536281 | rightend | YHR211W | 81.405 | 2377 | 384 | 26 | 1 | 2337 | 513107 | 515465 | 0 | 1906 |
| YJM1401 | IX | 1 |  | 423746 | leftend_complete | YIL169C | 96.921 | 1949 | 50 | 8 | 520 | 2466 | 20825 | 18885 | 0 | 3258 |
| YJM1401 | IX | 1767 | 1776 | 423746 | middle |  |  |  |  |  |  |  |  |  |  |  |
| YJM1401 | IX | 2357 | 2366 | 423746 | middle |  |  |  |  |  |  |  |  |  |  |  |
| YJM1401 | IX | 423737 | 423746 | 423746 | rightend | YIR039C | 99.628 | 1614 | 6 | 0 | 1 | 1614 | 421602 | 419989 | 0 | 2948 |
| YJM1401 | X | 1 |  | 714430 | leftend_complete | YJL213W | 99.699 | 996 | 3 | 0 | 1 | 996 | 25709 | 26704 | 0 | 1823 |
| YJM1401 | X | 2145 | 2154 | 714430 | middle |  |  |  |  |  |  |  |  |  |  |  |
| YJM1401 | X | 2761 | 2770 | 714430 | middle |  |  |  |  |  |  |  |  |  |  |  |
| YJM1401 | X | 3247 | 3256 | 714430 | middle |  |  |  |  |  |  |  |  |  |  |  |
| YJM1401 | X | 3946 | 3955 | 714430 | middle |  |  |  |  |  |  |  |  |  |  |  |
| YJM1401 | X | 4548 | 4557 | 714430 | middle |  |  |  |  |  |  |  |  |  |  |  |
| YJM1401 | X | 711432 | 711441 | 714430 | middle |  |  |  |  |  |  |  |  |  |  |  |
| YJM1401 | X | 712719 | 712728 | 714430 | middle |  |  |  |  |  |  |  |  |  |  |  |
| YJM1401 | X | 714421 | 714430 | 714430 | rightend | YJR152W | 99.571 | 1632 | 7 | 0 | 1 | 1632 | 697257 | 698888 | 0 | 2976 |
| YJM1401 | XI | 1 | 10 | 658771 | leftend | YKL222C | 99.811 | 2118 | 4 | 0 | 1 | 2118 | 3348 | 1231 | 0 | 3890 |
| YJM1401 | XI | 658762 | 658771 | 658771 | rightend | YKR101W | 99.695 | 1965 | 6 | 0 | 1 | 1965 | 638522 | 640486 | 0 | 3596 |
| YJM1401 | XII | 1 | 10 | 1506199 | leftend | YLL062C | 99.795 | 975 | 2 | 0 | 1 | 975 | 5435 | 4461 | 0 | 1790 |
| YJM1401 | XII | 1298 | 1307 | 1506199 | middle |  |  |  |  |  |  |  |  |  |  |  |
| YJM1401 | XII | 1503723 | 1503732 | 1506199 | middle |  |  |  |  |  |  |  |  |  |  |  |
| YJM1401 | XII | 1505553 | 1505562 | 1506199 | middle |  |  |  |  |  |  |  |  |  |  |  |
| YJM1401 | XII | 1506199 | 1506199 | 1506199 | rightend_complete | YLR460C | 99.558 | 1131 | 5 | 0 | 1 | 1131 | 1502472 | 1501342 | 0 | 2061 |
| YJM1401 | XIII | 1 | 10 | 908733 | leftend | YML131W | 99.454 | 1098 | 6 | 0 | 1 | 1098 | 2887 | 3984 | 0 | 1995 |
| YJM1401 | XIII | 908724 | 908733 | 908733 | rightend | YMR321C | 99.361 | 313 | 2 | 0 | 6 | 318 | 886994 | 886682 | 2.48E-161 | 568 |
| YJM1401 | XIV | 1 |  | 752633 | leftend_complete | YNL332W | 99.511 | 1023 | 5 | 0 | 1 | 1023 | 2254 | 3276 | 0 | 1862 |
| YJM1401 | XIV | 752624 | 752633 | 752633 | rightend | YNR071C | 99.514 | 1029 | 5 | 0 | 1 | 1029 | 742570 | 741542 | 0 | 1873 |
| YJM1401 | XV | 1 | 10 | 1041536 | leftend | YOL154W | 99.467 | 750 | 4 | 0 | 1 | 750 | 9320 | 10069 | 0 | 1363 |
| YJM1401 | XV | 1041527 | 1041536 | 1041536 | rightend | YOR381W | 98.642 | 2136 | 29 | 0 | 1 | 2136 | 1009548 | 1011683 | 0 | 3784 |
| YJM1401 | XVI | 1 | 10 | 902229 | leftend | YPL274W | 98.243 | 1764 | 31 | 0 | 1 | 1764 | 11737 | 13500 | 0 | 3086 |
| YJM1401 | XVI | 258 | 267 | 902229 | middle |  |  |  |  |  |  |  |  |  |  |  |
| YJM1401 | XVI | 902220 | 902229 | 902229 | rightend | YPR201W | 99.588 | 1215 | 5 | 0 | 1 | 1215 | 898803 | 900017 | 0 | 2217 |
| YJM1402 | I | 1 | 10 | 202503 | leftend | YAL063C | 95.291 | 998 | 45 | 2 | 2966 | 3962 | 14110 | 13114 | 0 | 1581 |
| YJM1402 | I | 2261 | 2270 | 202503 | middle |  |  |  |  |  |  |  |  |  |  |  |
| YJM1402 | I | 3916 | 3925 | 202503 | middle |  |  |  |  |  |  |  |  |  |  |  |
| YJM1402 | I | 191076 | 191085 | 202503 | middle |  |  |  |  |  |  |  |  |  |  |  |
| YJM1402 | I | 191382 | 191391 | 202503 | middle |  |  |  |  |  |  |  |  |  |  |  |
| YJM1402 | I | 202494 | 202503 | 202503 | rightend | YAR050W | 81.768 | 2183 | 359 | 26 | 1 | 2162 | 188869 | 191033 | 0 | 1790 |
| YJM1402 | II | 1 | 10 | 778278 | leftend | YNCB0001W | 100 | 84 | 0 | 0 | 1 | 84 | 1164 | 1247 | 4.16E-38 | 156 |
| YJM1402 | II | 778269 | 778278 | 778278 | rightend | YBR297W | 98.257 | 1205 | 21 | 0 | 1 | 1205 | 774081 | 775285 | 0 | 2109 |
| YJM1402 | III | 1 |  | 301914 | leftend_complete | YCL073C | 99.188 | 1848 | 15 | 0 | 1 | 1848 | 8002 | 6155 | 0 | 3330 |
| YJM1402 | III | 301320 | 301329 | 301914 | middle |  |  |  |  |  |  |  |  |  |  |  |
| YJM1402 | III | 301905 | 301914 | 301914 | rightend | YCR101C | 97.636 | 550 | 12 | 1 | 1 | 549 | 296403 | 295854 | 0 | 942 |
| YJM1402 | IV | 1 | 10 | 1463982 | leftend | YDL243C | 93.131 | 990 | 68 | 0 | 1 | 990 | 16195 | 15206 | 0 | 1452 |
| YJM1402 | IV | 1462290 | 1462299 | 1463982 | middle |  |  |  |  |  |  |  |  |  |  |  |
| YJM1402 | IV | 1463982 | 1463982 | 1463982 | rightend_complete | YDR541C | 99.517 | 1035 | 5 | 0 | 1 | 1035 | 1459567 | 1458533 | 0 | 1884 |
| YJM1402 | V | 1 | 10 | 555577 | leftend | YEL070W | 99.536 | 1509 | 7 | 0 | 1 | 1509 | 18906 | 20414 | 0 | 2748 |
| YJM1402 | V | 555577 | 555577 | 555577 | rightend_complete | YER187W | 99.061 | 426 | 4 | 0 | 1 | 426 | 551452 | 551877 | 0 | 765 |
| YJM1402 | VI | 1 |  | 272573 | leftend_complete | YFL051C | 100 | 452 | 0 | 0 | 1 | 452 | 34348 | 33897 | 0 | 835 |
| YJM1402 | VI | 765 | 774 | 272573 | middle |  |  |  |  |  |  |  |  |  |  |  |
| YJM1402 | VI | 1609 | 1618 | 272573 | middle |  |  |  |  |  |  |  |  |  |  |  |
| YJM1402 | VI | 272564 | 272573 | 272573 | rightend | YFR055W | 99.609 | 1022 | 4 | 0 | 1 | 1022 | 262906 | 263927 | 0 | 1866 |
| YJM1402 | VII | 1 |  | 1079139 | leftend_complete | YGL258W | 99.678 | 621 | 2 | 0 | 1 | 621 | 11361 | 11981 | 0 | 1136 |
| YJM1402 | VII | 6588 | 6597 | 1079139 | middle |  |  |  |  |  |  |  |  |  |  |  |
| YJM1402 | VII | 1074635 | 1074644 | 1079139 | middle |  |  |  |  |  |  |  |  |  |  |  |
| YJM1402 | VII | 1077670 | 1077679 | 1079139 | middle |  |  |  |  |  |  |  |  |  |  |  |
| YJM1402 | VII | 1079130 | 1079139 | 1079139 | rightend | YGR287C | 99.548 | 1770 | 8 | 0 | 1 | 1770 | 1040253 | 1038484 | 0 | 3225 |
| YJM1402 | VIII | 1 |  | 532763 | leftend_complete | YHL044W | 98.729 | 708 | 9 | 0 | 1 | 708 | 8801 | 9508 | 0 | 1258 |
| YJM1402 | VIII | 529969 | 529978 | 532763 | middle |  |  |  |  |  |  |  |  |  |  |  |
| YJM1402 | VIII | 530654 | 530663 | 532763 | middle |  |  |  |  |  |  |  |  |  |  |  |
| YJM1402 | VIII | 531585 | 531594 | 532763 | middle |  |  |  |  |  |  |  |  |  |  |  |
| YJM1402 | VIII | 532763 | 532763 | 532763 | rightend_complete | YHR211W | 89.781 | 2104 | 191 | 2 | 818 | 2918 | 513732 | 515814 | 0 | 2673 |
| YJM1402 | IX | 1 |  | 417171 | leftend_complete | YIL169C | 96.131 | 2223 | 75 | 7 | 394 | 2612 | 9621 | 7406 | 0 | 3618 |
| YJM1402 | IX | 583 | 592 | 417171 | middle |  |  |  |  |  |  |  |  |  |  |  |
| YJM1402 | IX | 417171 | 417171 | 417171 | rightend_complete | YIR039C | 99.69 | 1614 | 5 | 0 | 1 | 1614 | 410000 | 408387 | 0 | 2953 |
| YJM1402 | X | 1 | 10 | 719591 | leftend | YJL213W | 99.799 | 996 | 2 | 0 | 1 | 996 | 28623 | 29618 | 0 | 1829 |
| YJM1402 | X | 958 | 967 | 719591 | middle |  |  |  |  |  |  |  |  |  |  |  |
| YJM1402 | X | 1483 | 1492 | 719591 | middle |  |  |  |  |  |  |  |  |  |  |  |
| YJM1402 | X | 719591 | 719591 | 719591 | rightend_complete | YJR152W | 99.571 | 1632 | 7 | 0 | 1 | 1632 | 698749 | 700380 | 0 | 2976 |
| YJM1402 | XI | 1 | 10 | 658674 | leftend | YKL222C | 99.622 | 2118 | 8 | 0 | 1 | 2118 | 3235 | 1118 | 0 | 3868 |
| YJM1402 | XI | 657446 | 657455 | 658674 | middle |  |  |  |  |  |  |  |  |  |  |  |
| YJM1402 | XI | 658665 | 658674 | 658674 | rightend | YKR101W | 99.542 | 1965 | 9 | 0 | 1 | 1965 | 637166 | 639130 | 0 | 3579 |
| YJM1402 | XII | 1 |  | 1866513 | leftend_complete | YLL062C | 99.59 | 975 | 4 | 0 | 1 | 975 | 5340 | 4366 | 0 | 1779 |
| YJM1402 | XII | 1385 | 1394 | 1866513 | middle |  |  |  |  |  |  |  |  |  |  |  |
| YJM1402 | XII | 1865655 | 1865664 | 1866513 | middle |  |  |  |  |  |  |  |  |  |  |  |
| YJM1402 | XII | 1866504 | 1866513 | 1866513 | rightend | YLR460C | 99.646 | 1131 | 4 | 0 | 1 | 1131 | 1863029 | 1861899 | 0 | 2067 |
| YJM1402 | XIII | 1 | 10 | 916503 | leftend | YML131W | 99.545 | 1098 | 5 | 0 | 1 | 1098 | 4287 | 5384 | 0 | 2001 |
| YJM1402 | XIII | 916494 | 916503 | 916503 | rightend | YMR321C | 99.042 | 313 | 3 | 0 | 6 | 318 | 891589 | 891277 | 1.15E-159 | 562 |
| YJM1402 | XIV | 1 | 10 | 753728 | leftend | YNL332W | 99.12 | 1023 | 9 | 0 | 1 | 1023 | 5472 | 6494 | 0 | 1840 |
| YJM1402 | XIV | 459 | 468 | 753728 | middle |  |  |  |  |  |  |  |  |  |  |  |
| YJM1402 | XIV | 753719 | 753728 | 753728 | rightend | YNR071C | 99.611 | 1029 | 4 | 0 | 1 | 1029 | 743545 | 742517 | 0 | 1879 |
| YJM1402 | XV | 1 |  | 1073508 | leftend_complete | YOL154W | 99.333 | 750 | 5 | 0 | 1 | 750 | 35470 | 36219 | 0 | 1358 |
| YJM1402 | XV | 1064697 | 1064706 | 1073508 | middle |  |  |  |  |  |  |  |  |  |  |  |
| YJM1402 | XV | 1072507 | 1072516 | 1073508 | middle |  |  |  |  |  |  |  |  |  |  |  |
| YJM1402 | XV | 1073034 | 1073043 | 1073508 | middle |  |  |  |  |  |  |  |  |  |  |  |
| YJM1402 | XV | 1073499 | 1073508 | 1073508 | rightend | YOR381W | 98.876 | 2136 | 24 | 0 | 1 | 2136 | 1032536 | 1034671 | 0 | 3812 |
| YJM1402 | XVI | 1 |  | 904619 | leftend_complete | YPL274W | 98.186 | 1764 | 32 | 0 | 1 | 1764 | 12996 | 14759 | 0 | 3081 |
| YJM1402 | XVI | 699 | 708 | 904619 | middle |  |  |  |  |  |  |  |  |  |  |  |
| YJM1402 | XVI | 904619 | 904619 | 904619 | rightend_complete | YPR201W | 99.588 | 1215 | 5 | 0 | 1 | 1215 | 901374 | 902588 | 0 | 2217 |
| YJM1415 | I | 1 | 10 | 204762 | leftend | YAL063C | 95.02 | 1004 | 23 | 11 | 2966 | 3969 | 182362 | 183338 | 0 | 1552 |
| YJM1415 | I | 10235 | 10244 | 204762 | middle |  |  |  |  |  |  |  |  |  |  |  |
| YJM1415 | I | 204753 | 204762 | 204762 | rightend | YAR050W | 83.242 | 1092 | 179 | 4 | 1 | 1090 | 180729 | 181818 | 0 | 1000 |
| YJM1415 | II | 1 | 10 | 786575 | leftend | YNCB0001W | 100 | 84 | 0 | 0 | 1 | 84 | 2823 | 2906 | 4.16E-38 | 156 |
| YJM1415 | II | 223 | 232 | 786575 | middle |  |  |  |  |  |  |  |  |  |  |  |
| YJM1415 | II | 868 | 877 | 786575 | middle |  |  |  |  |  |  |  |  |  |  |  |
| YJM1415 | II | 786566 | 786575 | 786575 | rightend | YBR297W | 98.863 | 1407 | 14 | 1 | 1 | 1407 | 775217 | 776621 | 0 | 2508 |
| YJM1415 | III | 1 |  | 300758 | leftend_complete | YCL073C | 99.892 | 1848 | 2 | 0 | 1 | 1848 | 8126 | 6279 | 0 | 3402 |
| YJM1415 | III | 721 | 730 | 300758 | middle |  |  |  |  |  |  |  |  |  |  |  |
| YJM1415 | III | 300749 | 300758 | 300758 | rightend | YCR101C | 99.636 | 549 | 2 | 0 | 1 | 549 | 296233 | 295685 | 0 | 1003 |
| YJM1415 | IV | 1 | 10 | 1466831 | leftend | YDL243C | 95.051 | 990 | 49 | 0 | 1 | 990 | 2618 | 1629 | 0 | 1557 |
| YJM1415 | IV | 1466822 | 1466831 | 1466831 | rightend | YDR541C | 99.42 | 1035 | 6 | 0 | 1 | 1035 | 1463981 | 1462947 | 0 | 1879 |
| YJM1415 | V | 1 | 10 | 556650 | leftend | YEL070W | 98.94 | 1509 | 16 | 0 | 1 | 1509 | 13450 | 14958 | 0 | 2699 |
| YJM1415 | V | 556641 | 556650 | 556650 | rightend | YER187W | 91.981 | 424 | 32 | 2 | 4 | 426 | 547633 | 548055 | 3.44E-169 | 593 |
| YJM1415 | VI | 1 | 10 | 235631 | leftend | YFL051C | 98.451 | 452 | 7 | 0 | 1 | 452 | 3696 | 3245 | 0 | 797 |
| YJM1415 | VI | 235631 | 235631 | 235631 | rightend_complete | YFR055W | 100 | 1023 | 0 | 0 | 1 | 1023 | 229681 | 230703 | 0 | 1890 |
| YJM1415 | VII | 1 |  | 1050290 | leftend_complete | YGL258W | 97.585 | 621 | 15 | 0 | 1 | 621 | 11211 | 11831 | 0 | 1064 |
| YJM1415 | VII | 1050281 | 1050290 | 1050290 | rightend | YGR287C | 98.87 | 1770 | 20 | 0 | 1 | 1770 | 1030659 | 1028890 | 0 | 3158 |
| YJM1415 | VIII | 1 |  | 518144 | leftend_complete | YHL044W | 98.305 | 708 | 12 | 0 | 1 | 708 | 10502 | 11209 | 0 | 1242 |
| YJM1415 | VIII | 1282 | 1291 | 518144 | middle |  |  |  |  |  |  |  |  |  |  |  |
| YJM1415 | VIII | 2323 | 2332 | 518144 | middle |  |  |  |  |  |  |  |  |  |  |  |
| YJM1415 | VIII | 518135 | 518144 | 518144 | rightend | YHR211W | 82.535 | 939 | 153 | 7 | 1 | 932 | 514406 | 515340 | 0 | 815 |
| YJM1415 | IX | 1 | 10 | 403684 | leftend | YIL169C | 94.436 | 1294 | 70 | 2 | 520 | 1812 | 1931 | 639 | 0 | 1989 |
| YJM1415 | IX | 403675 | 403684 | 403684 | rightend | YIR039C | 99.009 | 1614 | 16 | 0 | 1 | 1614 | 402215 | 400602 | 0 | 2892 |
| YJM1415 | X | 1 | 10 | 721505 | leftend | YJL213W | 99.699 | 996 | 3 | 0 | 1 | 996 | 26939 | 27934 | 0 | 1823 |
| YJM1415 | X | 556 | 565 | 721505 | middle |  |  |  |  |  |  |  |  |  |  |  |
| YJM1415 | X | 720987 | 720996 | 721505 | middle |  |  |  |  |  |  |  |  |  |  |  |
| YJM1415 | X | 721505 | 721505 | 721505 | rightend_complete | YJR152W | 99.571 | 1632 | 3 | 2 | 1 | 1632 | 696564 | 698191 | 0 | 2972 |
| YJM1415 | XI | 1 | 10 | 668938 | leftend | YKL222C | 99.386 | 2118 | 13 | 0 | 1 | 2118 | 11502 | 9385 | 0 | 3840 |
| YJM1415 | XI | 668677 | 668686 | 668938 | middle |  |  |  |  |  |  |  |  |  |  |  |
| YJM1415 | XI | 668938 | 668938 | 668938 | rightend_complete | YKR101W | 99.796 | 1965 | 4 | 0 | 1 | 1965 | 645734 | 647698 | 0 | 3607 |
| YJM1415 | XII | 1 | 10 | 2267457 | leftend | YLL062C | 100 | 975 | 0 | 0 | 1 | 975 | 4921 | 3947 | 0 | 1801 |
| YJM1415 | XII | 915 | 924 | 2267457 | middle |  |  |  |  |  |  |  |  |  |  |  |
| YJM1415 | XII | 2264149 | 2264158 | 2267457 | middle |  |  |  |  |  |  |  |  |  |  |  |
| YJM1415 | XII | 2267448 | 2267457 | 2267457 | rightend | YLR460C | 94.248 | 1130 | 63 | 2 | 3 | 1131 | 2262405 | 2261277 | 0 | 1725 |
| YJM1415 | XIII | 1 | 10 | 900418 | leftend | YML131W | 100 | 1098 | 0 | 0 | 1 | 1098 | 4422 | 5519 | 0 | 2028 |
| YJM1415 | XIII | 1403 | 1412 | 900418 | middle |  |  |  |  |  |  |  |  |  |  |  |
| YJM1415 | XIII | 895680 | 895689 | 900418 | middle |  |  |  |  |  |  |  |  |  |  |  |
| YJM1415 | XIII | 900409 | 900418 | 900418 | rightend | YMR321C | 99.361 | 313 | 2 | 0 | 6 | 318 | 889094 | 888782 | 2.48E-161 | 568 |
| YJM1415 | XIV | 1 | 10 | 755540 | leftend | YNL332W | 98.631 | 1023 | 14 | 0 | 1 | 1023 | 7109 | 8131 | 0 | 1812 |
| YJM1415 | XIV | 1052 | 1061 | 755540 | middle |  |  |  |  |  |  |  |  |  |  |  |
| YJM1415 | XIV | 1952 | 1961 | 755540 | middle |  |  |  |  |  |  |  |  |  |  |  |
| YJM1415 | XIV | 753921 | 753930 | 755540 | middle |  |  |  |  |  |  |  |  |  |  |  |
| YJM1415 | XIV | 754605 | 754614 | 755540 | middle |  |  |  |  |  |  |  |  |  |  |  |
| YJM1415 | XIV | 755531 | 755540 | 755540 | rightend | YNR071C | 100 | 1029 | 0 | 0 | 1 | 1029 | 743627 | 742599 | 0 | 1901 |
| YJM1415 | XV | 1 | 10 | 1097882 | leftend | YOL154W | 99.467 | 750 | 4 | 0 | 1 | 750 | 9141 | 9890 | 0 | 1363 |
| YJM1415 | XV | 1097882 | 1097882 | 1097882 | rightend_complete | YOR381W | 99.953 | 2136 | 1 | 0 | 1 | 2136 | 1006378 | 1008513 | 0 | 3940 |
| YJM1415 | XVI | 1 |  | 908017 | leftend_complete | YPL274W | 99.887 | 1764 | 2 | 0 | 1 | 1764 | 16539 | 18302 | 0 | 3247 |
| YJM1415 | XVI | 905811 | 905820 | 908017 | middle |  |  |  |  |  |  |  |  |  |  |  |
| YJM1415 | XVI | 906867 | 906876 | 908017 | middle |  |  |  |  |  |  |  |  |  |  |  |
| YJM1415 | XVI | 908008 | 908017 | 908017 | rightend | YPR201W | 100 | 1215 | 0 | 0 | 1 | 1215 | 902533 | 903747 | 0 | 2244 |
| YJM1417 | I | 1 | 10 | 184639 | leftend | YAL063C | 86.3 | 781 | 99 | 8 | 40 | 816 | 173741 | 174517 | 0 | 843 |
| YJM1417 | I | 174534 | 174543 | 184639 | middle |  |  |  |  |  |  |  |  |  |  |  |
| YJM1417 | I | 184630 | 184639 | 184639 | rightend | YAR050W | 94.957 | 813 | 41 | 0 | 1 | 813 | 173702 | 174514 | 0 | 1275 |
| YJM1417 | II | 1 | 10 | 785529 | leftend | YNCB0001W | 100 | 84 | 0 | 0 | 1 | 84 | 3255 | 3338 | 4.16E-38 | 156 |
| YJM1417 | II | 785520 | 785529 | 785529 | rightend | YBR297W | 98.792 | 1407 | 17 | 0 | 1 | 1407 | 775443 | 776849 | 0 | 2505 |
| YJM1417 | III | 1 | 10 | 309164 | leftend | YCL073C | 99.892 | 1848 | 2 | 0 | 1 | 1848 | 7375 | 5528 | 0 | 3402 |
| YJM1417 | III | 309155 | 309164 | 309164 | rightend | YCR101C | 99.818 | 549 | 1 | 0 | 1 | 549 | 296121 | 295573 | 0 | 1009 |
| YJM1417 | IV | 1 | 10 | 1458513 | leftend | YDL243C | 95.051 | 990 | 49 | 0 | 1 | 990 | 6538 | 5549 | 0 | 1557 |
| YJM1417 | IV | 1458504 | 1458513 | 1458513 | rightend | YDR541C | 99.42 | 1035 | 6 | 0 | 1 | 1035 | 1455561 | 1454527 | 0 | 1879 |
| YJM1417 | V | 1 | 10 | 557828 | leftend | YEL070W | 98.873 | 1509 | 17 | 0 | 1 | 1509 | 17402 | 18910 | 0 | 2693 |
| YJM1417 | V | 2811 | 2820 | 557828 | middle |  |  |  |  |  |  |  |  |  |  |  |
| YJM1417 | V | 3887 | 3896 | 557828 | middle |  |  |  |  |  |  |  |  |  |  |  |
| YJM1417 | V | 557819 | 557828 | 557828 | rightend | YER187W | 91.981 | 424 | 32 | 2 | 4 | 426 | 551512 | 551934 | 3.44E-169 | 593 |
| YJM1417 | VI | 1 | 10 | 234675 | leftend | YFL051C | 98.451 | 452 | 7 | 0 | 1 | 452 | 1771 | 1320 | 0 | 797 |
| YJM1417 | VI | 909 | 918 | 234675 | middle |  |  |  |  |  |  |  |  |  |  |  |
| YJM1417 | VI | 234666 | 234675 | 234675 | rightend | YFR055W | 100 | 1023 | 0 | 0 | 1 | 1023 | 228426 | 229448 | 0 | 1890 |
| YJM1417 | VII | 1 | 10 | 1046990 | leftend | YGL258W | 97.585 | 621 | 15 | 0 | 1 | 621 | 10686 | 11306 | 0 | 1064 |
| YJM1417 | VII | 1046981 | 1046990 | 1046990 | rightend | YGR287C | 98.701 | 1770 | 23 | 0 | 1 | 1770 | 1031496 | 1029727 | 0 | 3142 |
| YJM1417 | VIII | 1 | 10 | 525119 | leftend | YHL044W | 98.446 | 708 | 11 | 0 | 1 | 708 | 1304 | 2011 | 0 | 1247 |
| YJM1417 | VIII | 524444 | 524453 | 525119 | middle |  |  |  |  |  |  |  |  |  |  |  |
| YJM1417 | VIII | 525110 | 525119 | 525119 | rightend | YHR211W | 93.258 | 2136 | 120 | 2 | 1096 | 3228 | 508774 | 510888 | 0 | 3125 |
| YJM1417 | IX | 1 | 10 | 407909 | leftend | YIL169C | 93.028 | 1420 | 97 | 2 | 394 | 1812 | 6100 | 4682 | 0 | 2073 |
| YJM1417 | IX | 407900 | 407909 | 407909 | rightend | YIR039C | 99.009 | 1614 | 16 | 0 | 1 | 1614 | 406634 | 405021 | 0 | 2892 |
| YJM1417 | X | 1 | 10 | 726496 | leftend | YJL213W | 99.598 | 996 | 4 | 0 | 1 | 996 | 26267 | 27262 | 0 | 1818 |
| YJM1417 | X | 715686 | 715695 | 726496 | middle |  |  |  |  |  |  |  |  |  |  |  |
| YJM1417 | X | 726487 | 726496 | 726496 | rightend | YJR152W | 99.755 | 1632 | 3 | 1 | 1 | 1632 | 695729 | 697359 | 0 | 2990 |
| YJM1417 | XI | 1 | 10 | 663026 | leftend | YKL222C | 99.481 | 2118 | 11 | 0 | 1 | 2118 | 3473 | 1356 | 0 | 3851 |
| YJM1417 | XI | 662291 | 662300 | 663026 | middle |  |  |  |  |  |  |  |  |  |  |  |
| YJM1417 | XI | 663026 | 663026 | 663026 | rightend_complete | YKR101W | 99.847 | 1965 | 3 | 0 | 1 | 1965 | 637073 | 639037 | 0 | 3613 |
| YJM1417 | XII | 1 | 10 | 2125796 | leftend | YLL062C | 100 | 975 | 0 | 0 | 1 | 975 | 10386 | 9412 | 0 | 1801 |
| YJM1417 | XII | 931 | 940 | 2125796 | middle |  |  |  |  |  |  |  |  |  |  |  |
| YJM1417 | XII | 6343 | 6352 | 2125796 | middle |  |  |  |  |  |  |  |  |  |  |  |
| YJM1417 | XII | 2124600 | 2124609 | 2125796 | middle |  |  |  |  |  |  |  |  |  |  |  |
| YJM1417 | XII | 2125796 | 2125796 | 2125796 | rightend_complete | YLR460C | 94.071 | 1130 | 65 | 2 | 3 | 1131 | 2122940 | 2121812 | 0 | 1714 |
| YJM1417 | XIII | 1 | 10 | 895509 | leftend | YML131W | 100 | 1098 | 0 | 0 | 1 | 1098 | 4203 | 5300 | 0 | 2028 |
| YJM1417 | XIII | 511 | 520 | 895509 | middle |  |  |  |  |  |  |  |  |  |  |  |
| YJM1417 | XIII | 1343 | 1352 | 895509 | middle |  |  |  |  |  |  |  |  |  |  |  |
| YJM1417 | XIII | 1896 | 1905 | 895509 | middle |  |  |  |  |  |  |  |  |  |  |  |
| YJM1417 | XIII | 895500 | 895509 | 895509 | rightend | YMR321C | 99.361 | 313 | 2 | 0 | 6 | 318 | 888914 | 888602 | 2.48E-161 | 568 |
| YJM1417 | XIV | 1 | 10 | 755420 | leftend | YNL332W | 98.631 | 1023 | 14 | 0 | 1 | 1023 | 5635 | 6657 | 0 | 1812 |
| YJM1417 | XIV | 754968 | 754977 | 755420 | middle |  |  |  |  |  |  |  |  |  |  |  |
| YJM1417 | XIV | 755411 | 755420 | 755420 | rightend | YNR071C | 100 | 1029 | 0 | 0 | 1 | 1029 | 744370 | 743342 | 0 | 1901 |
| YJM1417 | XV | 1 | 10 | 1034506 | leftend | YOL154W | 99.467 | 750 | 4 | 0 | 1 | 750 | 7883 | 8632 | 0 | 1363 |
| YJM1417 | XV | 1032904 | 1032913 | 1034506 | middle |  |  |  |  |  |  |  |  |  |  |  |
| YJM1417 | XV | 1034146 | 1034155 | 1034506 | middle |  |  |  |  |  |  |  |  |  |  |  |
| YJM1417 | XV | 1034497 | 1034506 | 1034506 | rightend | YOR381W | 99.86 | 2136 | 3 | 0 | 1 | 2136 | 1004562 | 1006697 | 0 | 3928 |
| YJM1417 | XVI | 1 |  | 893696 | leftend_complete | YPL274W | 99.943 | 1764 | 1 | 0 | 1 | 1764 | 4407 | 6170 | 0 | 3253 |
| YJM1417 | XVI | 626 | 635 | 893696 | middle |  |  |  |  |  |  |  |  |  |  |  |
| YJM1417 | XVI | 893687 | 893696 | 893696 | rightend | YPR201W | 100 | 1215 | 0 | 0 | 1 | 1215 | 890849 | 892063 | 0 | 2244 |
| YJM1418 | I | 1 |  | 227395 | leftend_complete | YAL063C | 95.02 | 1004 | 29 | 6 | 2966 | 3969 | 201370 | 202352 | 0 | 1557 |
| YJM1418 | I | 18247 | 18256 | 227395 | middle |  |  |  |  |  |  |  |  |  |  |  |
| YJM1418 | I | 18468 | 18477 | 227395 | middle |  |  |  |  |  |  |  |  |  |  |  |
| YJM1418 | I | 18988 | 18997 | 227395 | middle |  |  |  |  |  |  |  |  |  |  |  |
| YJM1418 | I | 217715 | 217724 | 227395 | middle |  |  |  |  |  |  |  |  |  |  |  |
| YJM1418 | I | 227386 | 227395 | 227395 | rightend | YAR050W | 82.884 | 1782 | 274 | 21 | 1 | 1765 | 199143 | 200910 | 0 | 1572 |
| YJM1418 | II | 1 |  | 788972 | leftend_complete | YNCB0001W | 100 | 84 | 0 | 0 | 1 | 84 | 10095 | 10178 | 4.16E-38 | 156 |
| YJM1418 | II | 788963 | 788972 | 788972 | rightend | YBR297W | 98.091 | 1205 | 23 | 0 | 1 | 1205 | 784676 | 785880 | 0 | 2098 |
| YJM1418 | III | 1 |  | 301892 | leftend_complete | YCL073C | 99.08 | 1848 | 17 | 0 | 1 | 1848 | 8115 | 6268 | 0 | 3319 |
| YJM1418 | III | 301342 | 301351 | 301892 | middle |  |  |  |  |  |  |  |  |  |  |  |
| YJM1418 | III | 301892 | 301892 | 301892 | rightend_complete | YCR101C | 98.182 | 550 | 8 | 2 | 1 | 549 | 296550 | 296002 | 0 | 959 |
| YJM1418 | IV | 1 | 10 | 1461741 | leftend | YDL243C | 94.04 | 990 | 59 | 0 | 1 | 990 | 15603 | 14614 | 0 | 1502 |
| YJM1418 | IV | 1460791 | 1460800 | 1461741 | middle |  |  |  |  |  |  |  |  |  |  |  |
| YJM1418 | IV | 1461732 | 1461741 | 1461741 | rightend | YDR541C | 99.42 | 1035 | 6 | 0 | 1 | 1035 | 1457451 | 1456417 | 0 | 1879 |
| YJM1418 | V | 1 | 10 | 550020 | leftend | YEL070W | 99.205 | 1509 | 12 | 0 | 1 | 1509 | 14481 | 15989 | 0 | 2721 |
| YJM1418 | V | 550011 | 550020 | 550020 | rightend | YER187W | 98.122 | 426 | 8 | 0 | 1 | 426 | 548695 | 549120 | 0 | 743 |
| YJM1418 | VI | 1 |  | 282011 | leftend_complete | YFL051C | 96.681 | 452 | 15 | 0 | 1 | 452 | 48218 | 47767 | 0 | 752 |
| YJM1418 | VI | 282011 | 282011 | 282011 | rightend_complete | YFR055W | 99.804 | 1022 | 2 | 0 | 1 | 1022 | 275892 | 276913 | 0 | 1877 |
| YJM1418 | VII | 1 |  | 1077448 | leftend_complete | YGL258W | 97.746 | 621 | 14 | 0 | 1 | 621 | 10618 | 11238 | 0 | 1070 |
| YJM1418 | VII | 6361 | 6370 | 1077448 | middle |  |  |  |  |  |  |  |  |  |  |  |
| YJM1418 | VII | 1076334 | 1076343 | 1077448 | middle |  |  |  |  |  |  |  |  |  |  |  |
| YJM1418 | VII | 1077439 | 1077448 | 1077448 | rightend | YGR287C | 99.096 | 1770 | 16 | 0 | 1 | 1770 | 1040514 | 1038745 | 0 | 3181 |
| YJM1418 | VIII | 1 |  | 518659 | leftend_complete | YHL044W | 98.729 | 708 | 9 | 0 | 1 | 708 | 8902 | 9609 | 0 | 1258 |
| YJM1418 | VIII | 513203 | 513212 | 518659 | middle |  |  |  |  |  |  |  |  |  |  |  |
| YJM1418 | VIII | 517774 | 517783 | 518659 | middle |  |  |  |  |  |  |  |  |  |  |  |
| YJM1418 | VIII | 518650 | 518659 | 518659 | rightend | YHR211W | 81.287 | 1336 | 233 | 10 | 1 | 1326 | 511865 | 513193 | 0 | 1066 |
| YJM1418 | IX | 1 |  | 419275 | leftend_complete | YIL169C | 96.026 | 2265 | 79 | 7 | 352 | 2612 | 10744 | 8487 | 0 | 3674 |
| YJM1418 | IX | 648 | 657 | 419275 | middle |  |  |  |  |  |  |  |  |  |  |  |
| YJM1418 | IX | 1850 | 1859 | 419275 | middle |  |  |  |  |  |  |  |  |  |  |  |
| YJM1418 | IX | 2856 | 2865 | 419275 | middle |  |  |  |  |  |  |  |  |  |  |  |
| YJM1418 | IX | 419275 | 419275 | 419275 | rightend_complete | YIR039C | 98.885 | 1614 | 18 | 0 | 1 | 1614 | 411445 | 409832 | 0 | 2881 |
| YJM1418 | X | 1 |  | 705380 | leftend_complete | YJL213W | 99.9 | 996 | 1 | 0 | 1 | 996 | 24618 | 25613 | 0 | 1834 |
| YJM1418 | X | 2405 | 2414 | 705380 | middle |  |  |  |  |  |  |  |  |  |  |  |
| YJM1418 | X | 2801 | 2810 | 705380 | middle |  |  |  |  |  |  |  |  |  |  |  |
| YJM1418 | X | 705371 | 705380 | 705380 | rightend | YJR152W | 97.794 | 1632 | 36 | 0 | 1 | 1632 | 694970 | 696601 | 0 | 2815 |
| YJM1418 | XI | 1 | 10 | 661844 | leftend | YKL222C | 99.528 | 2118 | 10 | 0 | 1 | 2118 | 3557 | 1440 | 0 | 3856 |
| YJM1418 | XI | 661835 | 661844 | 661844 | rightend | YKR101W | 99.338 | 1965 | 13 | 0 | 1 | 1965 | 638465 | 640429 | 0 | 3557 |
| YJM1418 | XII | 1 | 10 | 1753119 | leftend | YLL062C | 99.282 | 975 | 7 | 0 | 1 | 975 | 4347 | 3373 | 0 | 1762 |
| YJM1418 | XII | 1753110 | 1753119 | 1753119 | rightend | YLR460C | 98.322 | 1132 | 17 | 2 | 1 | 1131 | 1752077 | 1750947 | 0 | 1984 |
| YJM1418 | XIII | 1 | 10 | 916587 | leftend | YML131W | 99.545 | 1098 | 5 | 0 | 1 | 1098 | 3627 | 4724 | 0 | 2001 |
| YJM1418 | XIII | 911786 | 911795 | 916587 | middle |  |  |  |  |  |  |  |  |  |  |  |
| YJM1418 | XIII | 915292 | 915301 | 916587 | middle |  |  |  |  |  |  |  |  |  |  |  |
| YJM1418 | XIII | 916578 | 916587 | 916587 | rightend | YMR321C | 99.042 | 313 | 3 | 0 | 6 | 318 | 888865 | 888553 | 1.15E-159 | 562 |
| YJM1418 | XIV | 1 | 10 | 755931 | leftend | YNL332W | 98.729 | 1023 | 13 | 0 | 1 | 1023 | 6590 | 7612 | 0 | 1818 |
| YJM1418 | XIV | 755922 | 755931 | 755931 | rightend | YNR071C | 99.125 | 1029 | 9 | 0 | 1 | 1029 | 745691 | 744663 | 0 | 1851 |
| YJM1418 | XV | 1 | 10 | 1068778 | leftend | YOL154W | 98.267 | 750 | 13 | 0 | 1 | 750 | 35094 | 35843 | 0 | 1314 |
| YJM1418 | XV | 1066210 | 1066219 | 1068778 | middle |  |  |  |  |  |  |  |  |  |  |  |
| YJM1418 | XV | 1068769 | 1068778 | 1068778 | rightend | YOR381W | 98.549 | 2136 | 31 | 0 | 1 | 2136 | 1039196 | 1041331 | 0 | 3773 |
| YJM1418 | XVI | 1 |  | 899938 | leftend_complete | YPL274W | 97.789 | 1764 | 39 | 0 | 1 | 1764 | 5933 | 7696 | 0 | 3042 |
| YJM1418 | XVI | 550 | 559 | 899938 | middle |  |  |  |  |  |  |  |  |  |  |  |
| YJM1418 | XVI | 899938 | 899938 | 899938 | rightend_complete | YPR201W | 99.753 | 1215 | 3 | 0 | 1 | 1215 | 896628 | 897842 | 0 | 2228 |
| YJM1419 | I | 1 | 10 | 192532 | leftend | YAL063C | 93.086 | 998 | 46 | 8 | 2966 | 3962 | 12212 | 11237 | 0 | 1439 |
| YJM1419 | I | 997 | 1006 | 192532 | middle |  |  |  |  |  |  |  |  |  |  |  |
| YJM1419 | I | 1692 | 1701 | 192532 | middle |  |  |  |  |  |  |  |  |  |  |  |
| YJM1419 | I | 12261 | 12270 | 192532 | middle |  |  |  |  |  |  |  |  |  |  |  |
| YJM1419 | I | 12867 | 12876 | 192532 | middle |  |  |  |  |  |  |  |  |  |  |  |
| YJM1419 | I | 13334 | 13343 | 192532 | middle |  |  |  |  |  |  |  |  |  |  |  |
| YJM1419 | I | 14149 | 14158 | 192532 | middle |  |  |  |  |  |  |  |  |  |  |  |
| YJM1419 | I | 179261 | 179270 | 192532 | middle |  |  |  |  |  |  |  |  |  |  |  |
| YJM1419 | I | 192523 | 192532 | 192532 | rightend | YAR050W | 84.711 | 1282 | 186 | 4 | 1 | 1274 | 177981 | 179260 | 0 | 1273 |
| YJM1419 | II | 1 | 10 | 837317 | leftend | YNCB0001W | 100 | 84 | 0 | 0 | 1 | 84 | 59848 | 59931 | 4.16E-38 | 156 |
| YJM1419 | II | 835277 | 835286 | 837317 | middle |  |  |  |  |  |  |  |  |  |  |  |
| YJM1419 | II | 837308 | 837317 | 837317 | rightend | YBR297W | 98.008 | 1205 | 24 | 0 | 1 | 1205 | 833028 | 834232 | 0 | 2093 |
| YJM1419 | III | 1 |  | 311826 | leftend_complete | YCL073C | 98.972 | 1848 | 19 | 0 | 1 | 1848 | 8069 | 6222 | 0 | 3308 |
| YJM1419 | III | 302550 | 302559 | 311826 | middle |  |  |  |  |  |  |  |  |  |  |  |
| YJM1419 | III | 311817 | 311826 | 311826 | rightend | YCR101C | 98.182 | 550 | 9 | 1 | 1 | 549 | 298185 | 297636 | 0 | 959 |
| YJM1419 | IV | 1 | 10 | 1451671 | leftend | YDL243C | 93.037 | 991 | 65 | 4 | 1 | 990 | 2868 | 1881 | 0 | 1445 |
| YJM1419 | IV | 1449821 | 1449830 | 1451671 | middle |  |  |  |  |  |  |  |  |  |  |  |
| YJM1419 | IV | 1451671 | 1451671 | 1451671 | rightend_complete | YDR541C | 99.903 | 1035 | 1 | 0 | 1 | 1035 | 1447019 | 1445985 | 0 | 1906 |
| YJM1419 | V | 1 | 10 | 552650 | leftend | YEL070W | 99.139 | 1509 | 13 | 0 | 1 | 1509 | 13494 | 15002 | 0 | 2715 |
| YJM1419 | V | 552650 | 552650 | 552650 | rightend_complete | YER187W | 99.531 | 426 | 2 | 0 | 1 | 426 | 548484 | 548909 | 0 | 776 |
| YJM1419 | VI | 1 | 10 | 254571 | leftend | YFL051C | 100 | 483 | 0 | 0 | 1 | 483 | 19579 | 19097 | 0 | 893 |
| YJM1419 | VI | 1441 | 1450 | 254571 | middle |  |  |  |  |  |  |  |  |  |  |  |
| YJM1419 | VI | 2218 | 2227 | 254571 | middle |  |  |  |  |  |  |  |  |  |  |  |
| YJM1419 | VI | 3287 | 3296 | 254571 | middle |  |  |  |  |  |  |  |  |  |  |  |
| YJM1419 | VI | 254562 | 254571 | 254571 | rightend | YFR055W | 99.511 | 1022 | 5 | 0 | 1 | 1022 | 248180 | 249201 | 0 | 1860 |
| YJM1419 | VII | 1 | 10 | 1068571 | leftend | YGL258W | 99.678 | 621 | 2 | 0 | 1 | 621 | 4277 | 4897 | 0 | 1136 |
| YJM1419 | VII | 1057671 | 1057680 | 1068571 | middle |  |  |  |  |  |  |  |  |  |  |  |
| YJM1419 | VII | 1068562 | 1068571 | 1068571 | rightend | YGR287C | 99.944 | 1770 | 1 | 0 | 1 | 1770 | 1028455 | 1026686 | 0 | 3264 |
| YJM1419 | VIII | 1 |  | 535520 | leftend_complete | YHL044W | 98.588 | 708 | 10 | 0 | 1 | 708 | 8918 | 9625 | 0 | 1253 |
| YJM1419 | VIII | 518848 | 518857 | 535520 | middle |  |  |  |  |  |  |  |  |  |  |  |
| YJM1419 | VIII | 530553 | 530562 | 535520 | middle |  |  |  |  |  |  |  |  |  |  |  |
| YJM1419 | VIII | 535511 | 535520 | 535520 | rightend | YHR211W | 81.12 | 1589 | 273 | 16 | 1 | 1574 | 517271 | 518847 | 0 | 1247 |
| YJM1419 | IX | 1 |  | 414898 | leftend_complete | YIL169C | 97.384 | 2676 | 61 | 6 | 1 | 2667 | 5777 | 3102 | 0 | 4545 |
| YJM1419 | IX | 1415 | 1424 | 414898 | middle |  |  |  |  |  |  |  |  |  |  |  |
| YJM1419 | IX | 413625 | 413634 | 414898 | middle |  |  |  |  |  |  |  |  |  |  |  |
| YJM1419 | IX | 414898 | 414898 | 414898 | rightend_complete | YIR039C | 99.009 | 1614 | 16 | 0 | 1 | 1614 | 411630 | 410017 | 0 | 2892 |
| YJM1419 | X | 1 |  | 718246 | leftend_complete | YJL213W | 99.699 | 996 | 3 | 0 | 1 | 996 | 24046 | 25041 | 0 | 1823 |
| YJM1419 | X | 841 | 850 | 718246 | middle |  |  |  |  |  |  |  |  |  |  |  |
| YJM1419 | X | 718237 | 718246 | 718246 | rightend | YJR152W | 99.571 | 1632 | 7 | 0 | 1 | 1632 | 694343 | 695974 | 0 | 2976 |
| YJM1419 | XI | 1 | 10 | 662410 | leftend | YKL222C | 99.858 | 2118 | 3 | 0 | 1 | 2118 | 3332 | 1215 | 0 | 3895 |
| YJM1419 | XI | 662401 | 662410 | 662410 | rightend | YKR101W | 99.695 | 1965 | 6 | 0 | 1 | 1965 | 637851 | 639815 | 0 | 3596 |
| YJM1419 | XII | 1 |  | 2880384 | leftend_complete | YLL062C | 99.282 | 975 | 7 | 0 | 1 | 975 | 6290 | 5316 | 0 | 1762 |
| YJM1419 | XII | 1777 | 1786 | 2880384 | middle |  |  |  |  |  |  |  |  |  |  |  |
| YJM1419 | XII | 2878568 | 2878577 | 2880384 | middle |  |  |  |  |  |  |  |  |  |  |  |
| YJM1419 | XII | 2880384 | 2880384 | 2880384 | rightend_complete | YLR460C | 99.204 | 1131 | 9 | 0 | 1 | 1131 | 2873720 | 2872590 | 0 | 2039 |
| YJM1419 | XIII | 1 | 10 | 894032 | leftend | YML131W | 99.545 | 1098 | 5 | 0 | 1 | 1098 | 3717 | 4814 | 0 | 2001 |
| YJM1419 | XIII | 894032 | 894032 | 894032 | rightend_complete | YMR321C | 100 | 318 | 0 | 0 | 1 | 318 | 887558 | 887241 | 1.9E-167 | 588 |
| YJM1419 | XIV | 1 | 10 | 756891 | leftend | YNL332W | 99.12 | 1023 | 9 | 0 | 1 | 1023 | 6853 | 7875 | 0 | 1840 |
| YJM1419 | XIV | 1113 | 1122 | 756891 | middle |  |  |  |  |  |  |  |  |  |  |  |
| YJM1419 | XIV | 1394 | 1403 | 756891 | middle |  |  |  |  |  |  |  |  |  |  |  |
| YJM1419 | XIV | 756882 | 756891 | 756891 | rightend | YNR071C | 99.32 | 1029 | 7 | 0 | 1 | 1029 | 746352 | 745324 | 0 | 1862 |
| YJM1419 | XV | 1 |  | 1053231 | leftend_complete | YOL154W | 99.733 | 750 | 2 | 0 | 1 | 750 | 34968 | 35717 | 0 | 1375 |
| YJM1419 | XV | 675 | 684 | 1053231 | middle |  |  |  |  |  |  |  |  |  |  |  |
| YJM1419 | XV | 1049084 | 1049093 | 1053231 | middle |  |  |  |  |  |  |  |  |  |  |  |
| YJM1419 | XV | 1053222 | 1053231 | 1053231 | rightend | YOR381W | 98.736 | 2136 | 27 | 0 | 1 | 2136 | 1032709 | 1034844 | 0 | 3795 |
| YJM1419 | XVI | 1 | 10 | 905215 | leftend | YPL274W | 98.129 | 1764 | 33 | 0 | 1 | 1764 | 16589 | 18352 | 0 | 3075 |
| YJM1419 | XVI | 351 | 360 | 905215 | middle |  |  |  |  |  |  |  |  |  |  |  |
| YJM1419 | XVI | 1494 | 1503 | 905215 | middle |  |  |  |  |  |  |  |  |  |  |  |
| YJM1419 | XVI | 1936 | 1945 | 905215 | middle |  |  |  |  |  |  |  |  |  |  |  |
| YJM1419 | XVI | 905206 | 905215 | 905215 | rightend | YPR201W | 99.588 | 1215 | 5 | 0 | 1 | 1215 | 901795 | 903009 | 0 | 2217 |
| YJM1433 | I | 1 | 10 | 198468 | leftend | YAL063C | 85.73 | 1773 | 237 | 14 | 1 | 1765 | 12359 | 10595 | 0 | 1858 |
| YJM1433 | I | 9902 | 9911 | 198468 | middle |  |  |  |  |  |  |  |  |  |  |  |
| YJM1433 | I | 187515 | 187524 | 198468 | middle |  |  |  |  |  |  |  |  |  |  |  |
| YJM1433 | I | 194242 | 194251 | 198468 | middle |  |  |  |  |  |  |  |  |  |  |  |
| YJM1433 | I | 197692 | 197701 | 198468 | middle |  |  |  |  |  |  |  |  |  |  |  |
| YJM1433 | I | 198459 | 198468 | 198468 | rightend | YAR050W | 90.165 | 1088 | 98 | 5 | 1 | 1082 | 186392 | 187476 | 0 | 1408 |
| YJM1433 | II | 1 |  | 781406 | leftend_complete | YNCB0001W | 100 | 84 | 0 | 0 | 1 | 84 | 4753 | 4836 | 4.16E-38 | 156 |
| YJM1433 | II | 2071 | 2080 | 781406 | middle |  |  |  |  |  |  |  |  |  |  |  |
| YJM1433 | II | 3450 | 3459 | 781406 | middle |  |  |  |  |  |  |  |  |  |  |  |
| YJM1433 | II | 781397 | 781406 | 781406 | rightend | YBR297W | 97.925 | 1205 | 25 | 0 | 1 | 1205 | 775581 | 776785 | 0 | 2087 |
| YJM1433 | III | 1 |  | 308678 | leftend_complete | YCL073C | 99.351 | 1848 | 12 | 0 | 1 | 1848 | 7999 | 6152 | 0 | 3347 |
| YJM1433 | III | 300378 | 300387 | 308678 | middle |  |  |  |  |  |  |  |  |  |  |  |
| YJM1433 | III | 308669 | 308678 | 308678 | rightend | YCR101C | 99.636 | 549 | 2 | 0 | 1 | 549 | 295676 | 295128 | 0 | 1003 |
| YJM1433 | IV | 1 | 10 | 1451585 | leftend | YDL243C | 95.152 | 990 | 48 | 0 | 1 | 990 | 2507 | 1518 | 0 | 1563 |
| YJM1433 | IV | 1451585 | 1451585 | 1451585 | rightend_complete | YDR541C | 99.903 | 1035 | 1 | 0 | 1 | 1035 | 1446851 | 1445817 | 0 | 1906 |
| YJM1433 | V | 1 |  | 558587 | leftend_complete | YEL070W | 99.139 | 1509 | 13 | 0 | 1 | 1509 | 16038 | 17546 | 0 | 2715 |
| YJM1433 | V | 2393 | 2402 | 558587 | middle |  |  |  |  |  |  |  |  |  |  |  |
| YJM1433 | V | 558578 | 558587 | 558587 | rightend | YER187W | 91.981 | 424 | 32 | 2 | 4 | 426 | 556015 | 556437 | 3.44E-169 | 593 |
| YJM1433 | VI | 1 | 10 | 233975 | leftend | YFL051C | 98.451 | 452 | 7 | 0 | 1 | 452 | 1092 | 641 | 0 | 797 |
| YJM1433 | VI | 233966 | 233975 | 233975 | rightend | YFR055W | 100 | 1023 | 0 | 0 | 1 | 1023 | 227799 | 228821 | 0 | 1890 |
| YJM1433 | VII | 1 | 10 | 1055327 | leftend | YGL258W | 97.424 | 621 | 16 | 0 | 1 | 621 | 10683 | 11303 | 0 | 1059 |
| YJM1433 | VII | 1055318 | 1055327 | 1055327 | rightend | YGR287C | 98.644 | 1770 | 24 | 0 | 1 | 1770 | 1032262 | 1030493 | 0 | 3136 |
| YJM1433 | VIII | 1 |  | 546302 | leftend_complete | YHL044W | 98.446 | 708 | 11 | 0 | 1 | 708 | 9524 | 10231 | 0 | 1247 |
| YJM1433 | VIII | 815 | 824 | 546302 | middle |  |  |  |  |  |  |  |  |  |  |  |
| YJM1433 | VIII | 533178 | 533187 | 546302 | middle |  |  |  |  |  |  |  |  |  |  |  |
| YJM1433 | VIII | 535478 | 535487 | 546302 | middle |  |  |  |  |  |  |  |  |  |  |  |
| YJM1433 | VIII | 537044 | 537053 | 546302 | middle |  |  |  |  |  |  |  |  |  |  |  |
| YJM1433 | VIII | 538774 | 538783 | 546302 | middle |  |  |  |  |  |  |  |  |  |  |  |
| YJM1433 | VIII | 544266 | 544275 | 546302 | middle |  |  |  |  |  |  |  |  |  |  |  |
| YJM1433 | VIII | 545345 | 545354 | 546302 | middle |  |  |  |  |  |  |  |  |  |  |  |
| YJM1433 | VIII | 546302 | 546302 | 546302 | rightend_complete | YHR211W | 95.458 | 2400 | 109 | 0 | 829 | 3228 | 520133 | 522532 | 0 | 3829 |
| YJM1433 | IX | 1 |  | 428883 | leftend_complete | YIL169C | 84.633 | 2128 | 278 | 31 | 520 | 2618 | 18025 | 15918 | 0 | 2073 |
| YJM1433 | IX | 428883 | 428883 | 428883 | rightend_complete | YIR039C | 99.938 | 1614 | 1 | 0 | 1 | 1614 | 420743 | 419130 | 0 | 2976 |
| YJM1433 | X | 1 |  | 732993 | leftend_complete | YJL213W | 99.699 | 996 | 3 | 0 | 1 | 996 | 36816 | 37811 | 0 | 1823 |
| YJM1433 | X | 731098 | 731107 | 732993 | middle |  |  |  |  |  |  |  |  |  |  |  |
| YJM1433 | X | 732993 | 732993 | 732993 | rightend_complete | YJR152W | 99.755 | 1632 | 3 | 1 | 1 | 1632 | 706170 | 707800 | 0 | 2990 |
| YJM1433 | XI | 1 | 10 | 661648 | leftend | YKL222C | 99.433 | 2118 | 12 | 0 | 1 | 2118 | 3487 | 1370 | 0 | 3845 |
| YJM1433 | XI | 659655 | 659664 | 661648 | middle |  |  |  |  |  |  |  |  |  |  |  |
| YJM1433 | XI | 661639 | 661648 | 661648 | rightend | YKR101W | 99.847 | 1965 | 3 | 0 | 1 | 1965 | 637107 | 639071 | 0 | 3613 |
| YJM1433 | XII | 1 | 10 | 1924591 | leftend | YLL062C | 99.795 | 975 | 2 | 0 | 1 | 975 | 4134 | 3160 | 0 | 1790 |
| YJM1433 | XII | 1922695 | 1922704 | 1924591 | middle |  |  |  |  |  |  |  |  |  |  |  |
| YJM1433 | XII | 1924591 | 1924591 | 1924591 | rightend_complete | YLR460C | 94.071 | 1130 | 65 | 2 | 3 | 1131 | 1920470 | 1919342 | 0 | 1714 |
| YJM1433 | XIII | 1 | 10 | 896560 | leftend | YML131W | 100 | 1098 | 0 | 0 | 1 | 1098 | 2272 | 3369 | 0 | 2028 |
| YJM1433 | XIII | 890991 | 891000 | 896560 | middle |  |  |  |  |  |  |  |  |  |  |  |
| YJM1433 | XIII | 896551 | 896560 | 896560 | rightend | YMR321C | 99.361 | 313 | 2 | 0 | 6 | 318 | 886006 | 885694 | 2.48E-161 | 568 |
| YJM1433 | XIV | 1 |  | 758157 | leftend_complete | YNL332W | 98.631 | 1023 | 14 | 0 | 1 | 1023 | 6616 | 7638 | 0 | 1812 |
| YJM1433 | XIV | 1249 | 1258 | 758157 | middle |  |  |  |  |  |  |  |  |  |  |  |
| YJM1433 | XIV | 757449 | 757458 | 758157 | middle |  |  |  |  |  |  |  |  |  |  |  |
| YJM1433 | XIV | 758148 | 758157 | 758157 | rightend | YNR071C | 99.903 | 1029 | 1 | 0 | 1 | 1029 | 746078 | 745050 | 0 | 1895 |
| YJM1433 | XV | 1 | 10 | 1090584 | leftend | YOL154W | 99.733 | 750 | 2 | 0 | 1 | 750 | 2042 | 2791 | 0 | 1375 |
| YJM1433 | XV | 1090584 | 1090584 | 1090584 | rightend_complete | YOR381W | 99.813 | 2136 | 4 | 0 | 1 | 2136 | 999300 | 1001435 | 0 | 3923 |
| YJM1433 | XVI | 1 | 10 | 912175 | leftend | YPL274W | 99.206 | 1764 | 14 | 0 | 1 | 1764 | 19341 | 21104 | 0 | 3181 |
| YJM1433 | XVI | 471 | 480 | 912175 | middle |  |  |  |  |  |  |  |  |  |  |  |
| YJM1433 | XVI | 4467 | 4476 | 912175 | middle |  |  |  |  |  |  |  |  |  |  |  |
| YJM1433 | XVI | 911096 | 911105 | 912175 | middle |  |  |  |  |  |  |  |  |  |  |  |
| YJM1433 | XVI | 911364 | 911373 | 912175 | middle |  |  |  |  |  |  |  |  |  |  |  |
| YJM1433 | XVI | 912166 | 912175 | 912175 | rightend | YPR201W | 100 | 1215 | 0 | 0 | 1 | 1215 | 908371 | 909585 | 0 | 2244 |
| YJM1434 | I | 1 | 10 | 209085 | leftend | YAL063C | 84.514 | 2583 | 380 | 13 | 1 | 2573 | 15007 | 12435 | 0 | 2536 |
| YJM1434 | I | 11179 | 11188 | 209085 | middle |  |  |  |  |  |  |  |  |  |  |  |
| YJM1434 | I | 12233 | 12242 | 209085 | middle |  |  |  |  |  |  |  |  |  |  |  |
| YJM1434 | I | 185446 | 185455 | 209085 | middle |  |  |  |  |  |  |  |  |  |  |  |
| YJM1434 | I | 207702 | 207711 | 209085 | middle |  |  |  |  |  |  |  |  |  |  |  |
| YJM1434 | I | 209085 | 209085 | 209085 | rightend_complete | YAR050W | 80.665 | 2705 | 493 | 21 | 76 | 2765 | 14932 | 12243 | 0 | 2071 |
| YJM1434 | II | 1 |  | 780397 | leftend_complete | YNCB0001W | 100 | 84 | 0 | 0 | 1 | 84 | 2739 | 2822 | 4.16E-38 | 156 |
| YJM1434 | II | 1438 | 1447 | 780397 | middle |  |  |  |  |  |  |  |  |  |  |  |
| YJM1434 | II | 780388 | 780397 | 780397 | rightend | YBR297W | 98.257 | 1205 | 21 | 0 | 1 | 1205 | 776161 | 777365 | 0 | 2109 |
| YJM1434 | III | 1 | 10 | 307611 | leftend | YCL073C | 98.972 | 1848 | 19 | 0 | 1 | 1848 | 4821 | 2974 | 0 | 3308 |
| YJM1434 | III | 298624 | 298633 | 307611 | middle |  |  |  |  |  |  |  |  |  |  |  |
| YJM1434 | III | 307611 | 307611 | 307611 | rightend_complete | YCR101C | 98 | 550 | 9 | 2 | 1 | 549 | 293512 | 292964 | 0 | 953 |
| YJM1434 | IV | 1 | 10 | 1468354 | leftend | YDL243C | 93.131 | 990 | 68 | 0 | 1 | 990 | 16180 | 15191 | 0 | 1452 |
| YJM1434 | IV | 1468345 | 1468354 | 1468354 | rightend | YDR541C | 99.517 | 1035 | 5 | 0 | 1 | 1035 | 1458866 | 1457832 | 0 | 1884 |
| YJM1434 | V | 1 | 10 | 555055 | leftend | YEL070W | 100 | 1509 | 0 | 0 | 1 | 1509 | 17865 | 19373 | 0 | 2787 |
| YJM1434 | V | 555055 | 555055 | 555055 | rightend_complete | YER187W | 99.296 | 426 | 3 | 0 | 1 | 426 | 551257 | 551682 | 0 | 771 |
| YJM1434 | VI | 1 | 10 | 275842 | leftend | YFL051C | 100 | 452 | 0 | 0 | 1 | 452 | 37334 | 36883 | 0 | 835 |
| YJM1434 | VI | 275833 | 275842 | 275842 | rightend | YFR055W | 99.609 | 1022 | 4 | 0 | 1 | 1022 | 266146 | 267167 | 0 | 1866 |
| YJM1434 | VII | 1 | 10 | 1055014 | leftend | YGL258W | 99.678 | 621 | 2 | 0 | 1 | 621 | 4880 | 5500 | 0 | 1136 |
| YJM1434 | VII | 1052400 | 1052409 | 1055014 | middle |  |  |  |  |  |  |  |  |  |  |  |
| YJM1434 | VII | 1052625 | 1052634 | 1055014 | middle |  |  |  |  |  |  |  |  |  |  |  |
| YJM1434 | VII | 1055014 | 1055014 | 1055014 | rightend_complete | YGR287C | 98.079 | 1770 | 34 | 0 | 1 | 1770 | 1031861 | 1030092 | 0 | 3081 |
| YJM1434 | VIII | 1 | 10 | 513771 | leftend | YHL044W | 98.729 | 708 | 9 | 0 | 1 | 708 | 8582 | 9289 | 0 | 1258 |
| YJM1434 | VIII | 513762 | 513771 | 513771 | rightend | YHR211W | 81.473 | 1317 | 221 | 13 | 1 | 1304 | 512455 | 513761 | 0 | 1059 |
| YJM1434 | IX | 1 | 10 | 416925 | leftend | YIL169C | 96.159 | 2265 | 76 | 7 | 352 | 2612 | 7743 | 5486 | 0 | 3690 |
| YJM1434 | IX | 416307 | 416316 | 416925 | middle |  |  |  |  |  |  |  |  |  |  |  |
| YJM1434 | IX | 416925 | 416925 | 416925 | rightend_complete | YIR039C | 99.62 | 1317 | 5 | 0 | 1 | 1317 | 408266 | 406950 | 0 | 2405 |
| YJM1434 | X | 1 | 10 | 726651 | leftend | YJL213W | 99.799 | 996 | 2 | 0 | 1 | 996 | 30631 | 31626 | 0 | 1829 |
| YJM1434 | X | 1673 | 1682 | 726651 | middle |  |  |  |  |  |  |  |  |  |  |  |
| YJM1434 | X | 7288 | 7297 | 726651 | middle |  |  |  |  |  |  |  |  |  |  |  |
| YJM1434 | X | 721869 | 721878 | 726651 | middle |  |  |  |  |  |  |  |  |  |  |  |
| YJM1434 | X | 723377 | 723386 | 726651 | middle |  |  |  |  |  |  |  |  |  |  |  |
| YJM1434 | X | 724819 | 724828 | 726651 | middle |  |  |  |  |  |  |  |  |  |  |  |
| YJM1434 | X | 726642 | 726651 | 726651 | rightend | YJR152W | 99.449 | 1632 | 9 | 0 | 1 | 1632 | 700374 | 702005 | 0 | 2964 |
| YJM1434 | XI | 1 | 10 | 662644 | leftend | YKL222C | 99.669 | 2118 | 7 | 0 | 1 | 2118 | 3476 | 1359 | 0 | 3873 |
| YJM1434 | XI | 662635 | 662644 | 662644 | rightend | YKR101W | 99.644 | 1965 | 7 | 0 | 1 | 1965 | 637717 | 639681 | 0 | 3591 |
| YJM1434 | XII | 1 | 10 | 1806645 | leftend | YLL062C | 99.59 | 975 | 4 | 0 | 1 | 975 | 10483 | 9509 | 0 | 1779 |
| YJM1434 | XII | 1805339 | 1805348 | 1806645 | middle |  |  |  |  |  |  |  |  |  |  |  |
| YJM1434 | XII | 1806645 | 1806645 | 1806645 | rightend_complete | YLR460C | 99.469 | 1131 | 6 | 0 | 1 | 1131 | 1804548 | 1803418 | 0 | 2056 |
| YJM1434 | XIII | 1 | 10 | 896154 | leftend | YML131W | 99.545 | 1098 | 5 | 0 | 1 | 1098 | 3708 | 4805 | 0 | 2001 |
| YJM1434 | XIII | 896145 | 896154 | 896154 | rightend | YMR321C | 99.361 | 313 | 2 | 0 | 6 | 318 | 888311 | 887999 | 2.48E-161 | 568 |
| YJM1434 | XIV | 1 |  | 759684 | leftend_complete | YNL332W | 99.022 | 1023 | 10 | 0 | 1 | 1023 | 8196 | 9218 | 0 | 1834 |
| YJM1434 | XIV | 759675 | 759684 | 759684 | rightend | YNR071C | 99.611 | 1029 | 4 | 0 | 1 | 1029 | 748636 | 747608 | 0 | 1879 |
| YJM1434 | XV | 1 |  | 1064549 | leftend_complete | YOL154W | 99.733 | 750 | 2 | 0 | 1 | 750 | 35870 | 36619 | 0 | 1375 |
| YJM1434 | XV | 984 | 993 | 1064549 | middle |  |  |  |  |  |  |  |  |  |  |  |
| YJM1434 | XV | 1059886 | 1059895 | 1064549 | middle |  |  |  |  |  |  |  |  |  |  |  |
| YJM1434 | XV | 1063925 | 1063934 | 1064549 | middle |  |  |  |  |  |  |  |  |  |  |  |
| YJM1434 | XV | 1064549 | 1064549 | 1064549 | rightend_complete | YOR381W | 98.876 | 2136 | 24 | 0 | 1 | 2136 | 1032642 | 1034777 | 0 | 3812 |
| YJM1434 | XVI | 1 |  | 903384 | leftend_complete | YPL274W | 98.696 | 1764 | 23 | 0 | 1 | 1764 | 10095 | 11858 | 0 | 3131 |
| YJM1434 | XVI | 922 | 931 | 903384 | middle |  |  |  |  |  |  |  |  |  |  |  |
| YJM1434 | XVI | 903384 | 903384 | 903384 | rightend_complete | YPR201W | 99.506 | 1215 | 6 | 0 | 1 | 1215 | 900068 | 901282 | 0 | 2211 |
| YJM1439 | I | 1 | 10 | 198472 | leftend | YAL063C | 87.557 | 1093 | 132 | 4 | 1 | 1091 | 14889 | 13799 | 0 | 1262 |
| YJM1439 | I | 632 | 641 | 198472 | middle |  |  |  |  |  |  |  |  |  |  |  |
| YJM1439 | I | 2024 | 2033 | 198472 | middle |  |  |  |  |  |  |  |  |  |  |  |
| YJM1439 | I | 13414 | 13423 | 198472 | middle |  |  |  |  |  |  |  |  |  |  |  |
| YJM1439 | I | 185419 | 185428 | 198472 | middle |  |  |  |  |  |  |  |  |  |  |  |
| YJM1439 | I | 198463 | 198472 | 198472 | rightend | YAR050W | 80.727 | 991 | 187 | 4 | 103 | 1091 | 14787 | 13799 | 0 | 769 |
| YJM1439 | II | 1 | 10 | 779897 | leftend | YNCB0001W | 100 | 84 | 0 | 0 | 1 | 84 | 1434 | 1517 | 4.16E-38 | 156 |
| YJM1439 | II | 980 | 989 | 779897 | middle |  |  |  |  |  |  |  |  |  |  |  |
| YJM1439 | II | 779888 | 779897 | 779897 | rightend | YBR297W | 97.925 | 1205 | 25 | 0 | 1 | 1205 | 775774 | 776978 | 0 | 2087 |
| YJM1439 | III | 1 | 10 | 299675 | leftend | YCL073C | 98.81 | 1848 | 21 | 1 | 1 | 1848 | 3927 | 2081 | 0 | 3290 |
| YJM1439 | III | 299666 | 299675 | 299675 | rightend | YCR101C | 98.364 | 550 | 8 | 1 | 1 | 549 | 294670 | 294121 | 0 | 965 |
| YJM1439 | IV | 1 | 10 | 1451607 | leftend | YDL243C | 91.111 | 990 | 87 | 1 | 1 | 990 | 6906 | 5918 | 0 | 1339 |
| YJM1439 | IV | 1451598 | 1451607 | 1451607 | rightend | YDR541C | 99.614 | 1035 | 4 | 0 | 1 | 1035 | 1448342 | 1447308 | 0 | 1890 |
| YJM1439 | V | 1 | 10 | 552152 | leftend | YEL070W | 99.205 | 1509 | 12 | 0 | 1 | 1509 | 14449 | 15957 | 0 | 2721 |
| YJM1439 | V | 552143 | 552152 | 552152 | rightend | YER187W | 97.183 | 426 | 12 | 0 | 1 | 426 | 550198 | 550623 | 0 | 721 |
| YJM1439 | VI | 1 | 10 | 272674 | leftend | YFL051C | 98.451 | 452 | 7 | 0 | 1 | 452 | 38204 | 37753 | 0 | 797 |
| YJM1439 | VI | 272665 | 272674 | 272674 | rightend | YFR055W | 99.217 | 1022 | 8 | 0 | 1 | 1022 | 266501 | 267522 | 0 | 1844 |
| YJM1439 | VII | 1 | 10 | 1042268 | leftend | YGL258W | 99.034 | 621 | 6 | 0 | 1 | 621 | 3553 | 4173 | 0 | 1114 |
| YJM1439 | VII | 1039801 | 1039810 | 1042268 | middle |  |  |  |  |  |  |  |  |  |  |  |
| YJM1439 | VII | 1041575 | 1041584 | 1042268 | middle |  |  |  |  |  |  |  |  |  |  |  |
| YJM1439 | VII | 1042259 | 1042268 | 1042268 | rightend | YGR287C | 96.16 | 1771 | 66 | 2 | 1 | 1770 | 1030181 | 1028412 | 0 | 2892 |
| YJM1439 | VIII | 1 | 10 | 517165 | leftend | YHL044W | 98.023 | 708 | 14 | 0 | 1 | 708 | 2424 | 3131 | 0 | 1230 |
| YJM1439 | VIII | 508037 | 508046 | 517165 | middle |  |  |  |  |  |  |  |  |  |  |  |
| YJM1439 | VIII | 517156 | 517165 | 517165 | rightend | YHR211W | 84.032 | 1002 | 149 | 7 | 1 | 995 | 506293 | 507290 | 0 | 953 |
| YJM1439 | IX | 1 | 10 | 417301 | leftend | YIL169C | 79.207 | 2145 | 374 | 42 | 753 | 2842 | 13288 | 11161 | 0 | 1424 |
| YJM1439 | IX | 10906 | 10915 | 417301 | middle |  |  |  |  |  |  |  |  |  |  |  |
| YJM1439 | IX | 416730 | 416739 | 417301 | middle |  |  |  |  |  |  |  |  |  |  |  |
| YJM1439 | IX | 417292 | 417301 | 417301 | rightend | YIR039C | 99.69 | 1614 | 5 | 0 | 1 | 1614 | 415126 | 413513 | 0 | 2953 |
| YJM1439 | X | 1 | 10 | 724203 | leftend | YJL213W | 99.598 | 996 | 4 | 0 | 1 | 996 | 29894 | 30889 | 0 | 1818 |
| YJM1439 | X | 1828 | 1837 | 724203 | middle |  |  |  |  |  |  |  |  |  |  |  |
| YJM1439 | X | 1939 | 1948 | 724203 | middle |  |  |  |  |  |  |  |  |  |  |  |
| YJM1439 | X | 4395 | 4404 | 724203 | middle |  |  |  |  |  |  |  |  |  |  |  |
| YJM1439 | X | 4647 | 4656 | 724203 | middle |  |  |  |  |  |  |  |  |  |  |  |
| YJM1439 | X | 5506 | 5515 | 724203 | middle |  |  |  |  |  |  |  |  |  |  |  |
| YJM1439 | X | 712911 | 712920 | 724203 | middle |  |  |  |  |  |  |  |  |  |  |  |
| YJM1439 | X | 723510 | 723519 | 724203 | middle |  |  |  |  |  |  |  |  |  |  |  |
| YJM1439 | X | 724194 | 724203 | 724203 | rightend | YJR152W | 99.694 | 1632 | 5 | 0 | 1 | 1632 | 702837 | 704468 | 0 | 2987 |
| YJM1439 | XI | 1 | 10 | 663928 | leftend | YKL222C | 99.622 | 2118 | 8 | 0 | 1 | 2118 | 3301 | 1184 | 0 | 3868 |
| YJM1439 | XI | 663919 | 663928 | 663928 | rightend | YKR101W | 99.644 | 1965 | 7 | 0 | 1 | 1965 | 639494 | 641458 | 0 | 3591 |
| YJM1439 | XII | 1 | 10 | 2303699 | leftend | YLL062C | 99.282 | 975 | 7 | 0 | 1 | 975 | 8423 | 7449 | 0 | 1762 |
| YJM1439 | XII | 2301860 | 2301869 | 2303699 | middle |  |  |  |  |  |  |  |  |  |  |  |
| YJM1439 | XII | 2303690 | 2303699 | 2303699 | rightend | YLR460C | 96.021 | 1131 | 45 | 0 | 1 | 1131 | 2299753 | 2298623 | 0 | 1840 |
| YJM1439 | XIII | 1 | 10 | 915393 | leftend | YML131W | 99.545 | 1098 | 5 | 0 | 1 | 1098 | 3401 | 4498 | 0 | 2001 |
| YJM1439 | XIII | 913213 | 913222 | 915393 | middle |  |  |  |  |  |  |  |  |  |  |  |
| YJM1439 | XIII | 915384 | 915393 | 915393 | rightend | YMR321C | 99.361 | 313 | 2 | 0 | 6 | 318 | 889783 | 889471 | 2.48E-161 | 568 |
| YJM1439 | XIV | 1 | 10 | 764046 | leftend | YNL332W | 98.436 | 1023 | 16 | 0 | 1 | 1023 | 12786 | 13808 | 0 | 1801 |
| YJM1439 | XIV | 1111 | 1120 | 764046 | middle |  |  |  |  |  |  |  |  |  |  |  |
| YJM1439 | XIV | 1853 | 1862 | 764046 | middle |  |  |  |  |  |  |  |  |  |  |  |
| YJM1439 | XIV | 2883 | 2892 | 764046 | middle |  |  |  |  |  |  |  |  |  |  |  |
| YJM1439 | XIV | 4919 | 4928 | 764046 | middle |  |  |  |  |  |  |  |  |  |  |  |
| YJM1439 | XIV | 12629 | 12638 | 764046 | middle |  |  |  |  |  |  |  |  |  |  |  |
| YJM1439 | XIV | 764037 | 764046 | 764046 | rightend | YNR071C | 99.514 | 1029 | 5 | 0 | 1 | 1029 | 753568 | 752540 | 0 | 1873 |
| YJM1439 | XV | 1 | 10 | 1050943 | leftend | YOL154W | 99.467 | 750 | 4 | 0 | 1 | 750 | 16092 | 16841 | 0 | 1363 |
| YJM1439 | XV | 1050568 | 1050577 | 1050943 | middle |  |  |  |  |  |  |  |  |  |  |  |
| YJM1439 | XV | 1050934 | 1050943 | 1050943 | rightend | YOR381W | 98.876 | 2136 | 24 | 0 | 1 | 2136 | 1015477 | 1017612 | 0 | 3812 |
| YJM1439 | XVI | 1 | 10 | 912769 | leftend | YPL274W | 98.469 | 1764 | 27 | 0 | 1 | 1764 | 16141 | 17904 | 0 | 3109 |
| YJM1439 | XVI | 1417 | 1426 | 912769 | middle |  |  |  |  |  |  |  |  |  |  |  |
| YJM1439 | XVI | 912760 | 912769 | 912769 | rightend | YPR201W | 99.588 | 1215 | 5 | 0 | 1 | 1215 | 908148 | 909362 | 0 | 2217 |
| YJM1443 | I | 1 | 10 | 198556 | leftend | YAL063C | 85.113 | 1498 | 217 | 6 | 1 | 1495 | 20722 | 19228 | 0 | 1526 |
| YJM1443 | I | 18142 | 18151 | 198556 | middle |  |  |  |  |  |  |  |  |  |  |  |
| YJM1443 | I | 18709 | 18718 | 198556 | middle |  |  |  |  |  |  |  |  |  |  |  |
| YJM1443 | I | 197845 | 197854 | 198556 | middle |  |  |  |  |  |  |  |  |  |  |  |
| YJM1443 | I | 198547 | 198556 | 198556 | rightend | YAR050W | 88.625 | 1222 | 132 | 5 | 1 | 1217 | 196620 | 197839 | 0 | 1480 |
| YJM1443 | II | 1 | 10 | 777748 | leftend | YNCB0001W | 100 | 84 | 0 | 0 | 1 | 84 | 1194 | 1277 | 4.16E-38 | 156 |
| YJM1443 | II | 777739 | 777748 | 777748 | rightend | YBR297W | 98.257 | 1205 | 21 | 0 | 1 | 1205 | 773616 | 774820 | 0 | 2109 |
| YJM1443 | III | 1 |  | 305211 | leftend_complete | YCL073C | 98.972 | 1848 | 19 | 0 | 1 | 1848 | 5267 | 3420 | 0 | 3308 |
| YJM1443 | III | 300444 | 300453 | 305211 | middle |  |  |  |  |  |  |  |  |  |  |  |
| YJM1443 | III | 305202 | 305211 | 305211 | rightend | YCR101C | 98.545 | 550 | 7 | 1 | 1 | 549 | 295756 | 295207 | 0 | 970 |
| YJM1443 | IV | 1 | 10 | 1465909 | leftend | YDL243C | 93.03 | 990 | 69 | 0 | 1 | 990 | 15721 | 14732 | 0 | 1447 |
| YJM1443 | IV | 1460669 | 1460678 | 1465909 | middle |  |  |  |  |  |  |  |  |  |  |  |
| YJM1443 | IV | 1461333 | 1461342 | 1465909 | middle |  |  |  |  |  |  |  |  |  |  |  |
| YJM1443 | IV | 1464253 | 1464262 | 1465909 | middle |  |  |  |  |  |  |  |  |  |  |  |
| YJM1443 | IV | 1465909 | 1465909 | 1465909 | rightend_complete | YDR541C | 99.903 | 1035 | 1 | 0 | 1 | 1035 | 1457905 | 1456871 | 0 | 1906 |
| YJM1443 | V | 1 | 10 | 548855 | leftend | YEL070W | 99.536 | 1509 | 7 | 0 | 1 | 1509 | 12763 | 14271 | 0 | 2748 |
| YJM1443 | V | 548846 | 548855 | 548855 | rightend | YER187W | 99.296 | 426 | 3 | 0 | 1 | 426 | 544752 | 545177 | 0 | 771 |
| YJM1443 | VI | 1 | 10 | 275927 | leftend | YFL051C | 100 | 452 | 0 | 0 | 1 | 452 | 35451 | 35000 | 0 | 835 |
| YJM1443 | VI | 275927 | 275927 | 275927 | rightend_complete | YFR055W | 88.496 | 1017 | 115 | 2 | 1 | 1016 | 264462 | 265477 | 0 | 1229 |
| YJM1443 | VII | 1 | 10 | 1064909 | leftend | YGL258W | 99.517 | 621 | 3 | 0 | 1 | 621 | 11278 | 11898 | 0 | 1131 |
| YJM1443 | VII | 5852 | 5861 | 1064909 | middle |  |  |  |  |  |  |  |  |  |  |  |
| YJM1443 | VII | 1063901 | 1063910 | 1064909 | middle |  |  |  |  |  |  |  |  |  |  |  |
| YJM1443 | VII | 1064900 | 1064909 | 1064909 | rightend | YGR287C | 99.831 | 1770 | 3 | 0 | 1 | 1770 | 1031818 | 1030049 | 0 | 3253 |
| YJM1443 | VIII | 1 |  | 537996 | leftend_complete | YHL044W | 98.446 | 708 | 11 | 0 | 1 | 708 | 8362 | 9069 | 0 | 1247 |
| YJM1443 | VIII | 7094 | 7103 | 537996 | middle |  |  |  |  |  |  |  |  |  |  |  |
| YJM1443 | VIII | 514516 | 514525 | 537996 | middle |  |  |  |  |  |  |  |  |  |  |  |
| YJM1443 | VIII | 526116 | 526125 | 537996 | middle |  |  |  |  |  |  |  |  |  |  |  |
| YJM1443 | VIII | 537996 | 537996 | 537996 | rightend_complete | YHR211W | 80.84 | 1571 | 268 | 19 | 1 | 1553 | 512942 | 514497 | 0 | 1203 |
| YJM1443 | IX | 1 |  | 417361 | leftend_complete | YIL169C | 96.176 | 2223 | 74 | 7 | 394 | 2612 | 7793 | 5578 | 0 | 3624 |
| YJM1443 | IX | 608 | 617 | 417361 | middle |  |  |  |  |  |  |  |  |  |  |  |
| YJM1443 | IX | 1660 | 1669 | 417361 | middle |  |  |  |  |  |  |  |  |  |  |  |
| YJM1443 | IX | 414237 | 414246 | 417361 | middle |  |  |  |  |  |  |  |  |  |  |  |
| YJM1443 | IX | 415139 | 415148 | 417361 | middle |  |  |  |  |  |  |  |  |  |  |  |
| YJM1443 | IX | 417361 | 417361 | 417361 | rightend_complete | YIR039C | 98.575 | 1614 | 22 | 1 | 1 | 1614 | 407226 | 405614 | 0 | 2852 |
| YJM1443 | X | 1 | 10 | 716892 | leftend | YJL213W | 99.799 | 996 | 2 | 0 | 1 | 996 | 26665 | 27660 | 0 | 1829 |
| YJM1443 | X | 712553 | 712562 | 716892 | middle |  |  |  |  |  |  |  |  |  |  |  |
| YJM1443 | X | 716892 | 716892 | 716892 | rightend_complete | YJR152W | 99.51 | 1632 | 8 | 0 | 1 | 1632 | 696794 | 698425 | 0 | 2970 |
| YJM1443 | XI | 1 | 10 | 657721 | leftend | YKL222C | 99.575 | 2118 | 9 | 0 | 1 | 2118 | 3176 | 1059 | 0 | 3862 |
| YJM1443 | XI | 657712 | 657721 | 657721 | rightend | YKR101W | 99.288 | 1965 | 14 | 0 | 1 | 1965 | 637598 | 639562 | 0 | 3552 |
| YJM1443 | XII | 1 | 10 | 1581839 | leftend | YLL062C | 99.795 | 975 | 2 | 0 | 1 | 975 | 3785 | 2811 | 0 | 1790 |
| YJM1443 | XII | 1580877 | 1580886 | 1581839 | middle |  |  |  |  |  |  |  |  |  |  |  |
| YJM1443 | XII | 1581830 | 1581839 | 1581839 | rightend | YLR460C | 98.851 | 1131 | 13 | 0 | 1 | 1131 | 1580330 | 1579200 | 0 | 2017 |
| YJM1443 | XIII | 1 | 10 | 902314 | leftend | YML131W | 99.545 | 1098 | 5 | 0 | 1 | 1098 | 4286 | 5383 | 0 | 2001 |
| YJM1443 | XIII | 1318 | 1327 | 902314 | middle |  |  |  |  |  |  |  |  |  |  |  |
| YJM1443 | XIII | 902305 | 902314 | 902314 | rightend | YMR321C | 99.361 | 313 | 2 | 0 | 6 | 318 | 887341 | 887029 | 2.48E-161 | 568 |
| YJM1443 | XIV | 1 |  | 768330 | leftend_complete | YNL332W | 98.631 | 1023 | 14 | 0 | 1 | 1023 | 14406 | 15428 | 0 | 1812 |
| YJM1443 | XIV | 7399 | 7408 | 768330 | middle |  |  |  |  |  |  |  |  |  |  |  |
| YJM1443 | XIV | 7832 | 7841 | 768330 | middle |  |  |  |  |  |  |  |  |  |  |  |
| YJM1443 | XIV | 8818 | 8827 | 768330 | middle |  |  |  |  |  |  |  |  |  |  |  |
| YJM1443 | XIV | 768321 | 768330 | 768330 | rightend | YNR071C | 99.611 | 1029 | 4 | 0 | 1 | 1029 | 754188 | 753160 | 0 | 1879 |
| YJM1443 | XV | 1 | 10 | 1054388 | leftend | YOL154W | 99.733 | 750 | 2 | 0 | 1 | 750 | 32273 | 33022 | 0 | 1375 |
| YJM1443 | XV | 9248 | 9257 | 1054388 | middle |  |  |  |  |  |  |  |  |  |  |  |
| YJM1443 | XV | 1054388 | 1054388 | 1054388 | rightend_complete | YOR381W | 98.83 | 2136 | 25 | 0 | 1 | 2136 | 1029196 | 1031331 | 0 | 3807 |
| YJM1443 | XVI | 1 | 10 | 916010 | leftend | YPL274W | 98.243 | 1764 | 31 | 0 | 1 | 1764 | 22732 | 24495 | 0 | 3086 |
| YJM1443 | XVI | 916001 | 916010 | 916010 | rightend | YPR201W | 99.506 | 1215 | 6 | 0 | 1 | 1215 | 912912 | 914126 | 0 | 2211 |
| YJM1444 | I | 1 | 10 | 209940 | leftend | YAL063C | 79.842 | 2153 | 390 | 29 | 40 | 2170 | 188383 | 190513 | 0 | 1531 |
| YJM1444 | I | 895 | 904 | 209940 | middle |  |  |  |  |  |  |  |  |  |  |  |
| YJM1444 | I | 196174 | 196183 | 209940 | middle |  |  |  |  |  |  |  |  |  |  |  |
| YJM1444 | I | 209940 | 209940 | 209940 | rightend_complete | YAR050W | 84.84 | 2190 | 292 | 25 | 1 | 2170 | 188344 | 190513 | 0 | 2169 |
| YJM1444 | II | 1 | 10 | 788486 | leftend | YNCB0001W | 100 | 84 | 0 | 0 | 1 | 84 | 9077 | 9160 | 4.16E-38 | 156 |
| YJM1444 | II | 699 | 708 | 788486 | middle |  |  |  |  |  |  |  |  |  |  |  |
| YJM1444 | II | 788477 | 788486 | 788486 | rightend | YBR297W | 98.257 | 1205 | 21 | 0 | 1 | 1205 | 784169 | 785373 | 0 | 2109 |
| YJM1444 | III | 1 | 10 | 299650 | leftend | YCL073C | 98.864 | 1848 | 21 | 0 | 1 | 1848 | 7913 | 6066 | 0 | 3297 |
| YJM1444 | III | 572 | 581 | 299650 | middle |  |  |  |  |  |  |  |  |  |  |  |
| YJM1444 | III | 297588 | 297597 | 299650 | middle |  |  |  |  |  |  |  |  |  |  |  |
| YJM1444 | III | 298441 | 298450 | 299650 | middle |  |  |  |  |  |  |  |  |  |  |  |
| YJM1444 | III | 299641 | 299650 | 299650 | rightend | YCR101C | 98.364 | 550 | 8 | 1 | 1 | 549 | 296420 | 295871 | 0 | 965 |
| YJM1444 | IV | 1 | 10 | 1449719 | leftend | YDL243C | 93.34 | 991 | 64 | 2 | 1 | 990 | 3327 | 2338 | 0 | 1463 |
| YJM1444 | IV | 1370 | 1379 | 1449719 | middle |  |  |  |  |  |  |  |  |  |  |  |
| YJM1444 | IV | 1449710 | 1449719 | 1449719 | rightend | YDR541C | 99.71 | 1035 | 3 | 0 | 1 | 1035 | 1447063 | 1446029 | 0 | 1895 |
| YJM1444 | V | 1 | 10 | 551705 | leftend | YEL070W | 99.337 | 1509 | 10 | 0 | 1 | 1509 | 13817 | 15325 | 0 | 2732 |
| YJM1444 | V | 551251 | 551260 | 551705 | middle |  |  |  |  |  |  |  |  |  |  |  |
| YJM1444 | V | 551696 | 551705 | 551705 | rightend | YER187W | 91.981 | 424 | 32 | 2 | 4 | 426 | 548747 | 549169 | 3.44E-169 | 593 |
| YJM1444 | VI | 1 | 10 | 264667 | leftend | YFL051C | 100 | 483 | 0 | 0 | 1 | 483 | 23335 | 22853 | 0 | 893 |
| YJM1444 | VI | 264658 | 264667 | 264667 | rightend | YFR055W | 88.496 | 1017 | 115 | 2 | 1 | 1016 | 252181 | 253196 | 0 | 1229 |
| YJM1444 | VII | 1 |  | 1067392 | leftend_complete | YGL258W | 97.585 | 621 | 15 | 0 | 1 | 621 | 11659 | 12279 | 0 | 1064 |
| YJM1444 | VII | 2653 | 2662 | 1067392 | middle |  |  |  |  |  |  |  |  |  |  |  |
| YJM1444 | VII | 1066173 | 1066182 | 1067392 | middle |  |  |  |  |  |  |  |  |  |  |  |
| YJM1444 | VII | 1067383 | 1067392 | 1067392 | rightend | YGR287C | 99.209 | 1770 | 14 | 0 | 1 | 1770 | 1032097 | 1030328 | 0 | 3192 |
| YJM1444 | VIII | 1 | 10 | 536050 | leftend | YHL044W | 98.446 | 708 | 11 | 0 | 1 | 708 | 9052 | 9759 | 0 | 1247 |
| YJM1444 | VIII | 535074 | 535083 | 536050 | middle |  |  |  |  |  |  |  |  |  |  |  |
| YJM1444 | VIII | 536050 | 536050 | 536050 | rightend_complete | YHR211W | 98.031 | 2133 | 42 | 0 | 1096 | 3228 | 514185 | 516317 | 0 | 3707 |
| YJM1444 | IX | 1 |  | 428608 | leftend_complete | YIL169C | 96.401 | 2223 | 66 | 8 | 394 | 2612 | 17300 | 15088 | 0 | 3650 |
| YJM1444 | IX | 1168 | 1177 | 428608 | middle |  |  |  |  |  |  |  |  |  |  |  |
| YJM1444 | IX | 426912 | 426921 | 428608 | middle |  |  |  |  |  |  |  |  |  |  |  |
| YJM1444 | IX | 428599 | 428608 | 428608 | rightend | YIR039C | 98.39 | 1615 | 24 | 2 | 1 | 1614 | 420120 | 418507 | 0 | 2837 |
| YJM1444 | X | 1 |  | 725980 | leftend_complete | YJL213W | 89.859 | 996 | 101 | 0 | 1 | 996 | 34574 | 35569 | 0 | 1280 |
| YJM1444 | X | 1357 | 1366 | 725980 | middle |  |  |  |  |  |  |  |  |  |  |  |
| YJM1444 | X | 1566 | 1575 | 725980 | middle |  |  |  |  |  |  |  |  |  |  |  |
| YJM1444 | X | 2308 | 2317 | 725980 | middle |  |  |  |  |  |  |  |  |  |  |  |
| YJM1444 | X | 721696 | 721705 | 725980 | middle |  |  |  |  |  |  |  |  |  |  |  |
| YJM1444 | X | 725980 | 725980 | 725980 | rightend_complete | YJR152W | 99.571 | 1632 | 7 | 0 | 1 | 1632 | 705864 | 707495 | 0 | 2976 |
| YJM1444 | XI | 1 | 10 | 666628 | leftend | YKL222C | 99.528 | 2118 | 10 | 0 | 1 | 2118 | 4402 | 2285 | 0 | 3856 |
| YJM1444 | XI | 1026 | 1035 | 666628 | middle |  |  |  |  |  |  |  |  |  |  |  |
| YJM1444 | XI | 666619 | 666628 | 666628 | rightend | YKR101W | 99.135 | 1965 | 17 | 0 | 1 | 1965 | 638215 | 640179 | 0 | 3535 |
| YJM1444 | XII | 1 | 10 | 1657534 | leftend | YLL062C | 99.897 | 975 | 1 | 0 | 1 | 975 | 5807 | 4833 | 0 | 1796 |
| YJM1444 | XII | 1456 | 1465 | 1657534 | middle |  |  |  |  |  |  |  |  |  |  |  |
| YJM1444 | XII | 1657534 | 1657534 | 1657534 | rightend_complete | YLR460C | 99.823 | 1131 | 2 | 0 | 1 | 1131 | 1653061 | 1651931 | 0 | 2078 |
| YJM1444 | XIII | 1 | 10 | 900853 | leftend | YML131W | 99.454 | 1098 | 6 | 0 | 1 | 1098 | 4369 | 5466 | 0 | 1995 |
| YJM1444 | XIII | 522 | 531 | 900853 | middle |  |  |  |  |  |  |  |  |  |  |  |
| YJM1444 | XIII | 896181 | 896190 | 900853 | middle |  |  |  |  |  |  |  |  |  |  |  |
| YJM1444 | XIII | 900212 | 900221 | 900853 | middle |  |  |  |  |  |  |  |  |  |  |  |
| YJM1444 | XIII | 900844 | 900853 | 900853 | rightend | YMR321C | 99.361 | 313 | 2 | 0 | 6 | 318 | 888107 | 887795 | 2.48E-161 | 568 |
| YJM1444 | XIV | 1 |  | 768593 | leftend_complete | YNL332W | 99.022 | 1023 | 10 | 0 | 1 | 1023 | 2054 | 3076 | 0 | 1834 |
| YJM1444 | XIV | 1888 | 1897 | 768593 | middle |  |  |  |  |  |  |  |  |  |  |  |
| YJM1444 | XIV | 760940 | 760949 | 768593 | middle |  |  |  |  |  |  |  |  |  |  |  |
| YJM1444 | XIV | 766191 | 766200 | 768593 | middle |  |  |  |  |  |  |  |  |  |  |  |
| YJM1444 | XIV | 767131 | 767140 | 768593 | middle |  |  |  |  |  |  |  |  |  |  |  |
| YJM1444 | XIV | 767996 | 768005 | 768593 | middle |  |  |  |  |  |  |  |  |  |  |  |
| YJM1444 | XIV | 768593 | 768593 | 768593 | rightend_complete | YNR071C | 99.514 | 1029 | 5 | 0 | 1 | 1029 | 741737 | 740709 | 0 | 1873 |
| YJM1444 | XV | 1 | 10 | 1075835 | leftend | YOL154W | 99.6 | 750 | 3 | 0 | 1 | 750 | 48479 | 49228 | 0 | 1369 |
| YJM1444 | XV | 1075826 | 1075835 | 1075835 | rightend | YOR381W | 98.455 | 2136 | 33 | 0 | 1 | 2136 | 1044696 | 1046831 | 0 | 3762 |
| YJM1444 | XVI | 1 | 10 | 914131 | leftend | YPL274W | 98.866 | 1764 | 20 | 0 | 1 | 1764 | 22923 | 24686 | 0 | 3147 |
| YJM1444 | XVI | 1549 | 1558 | 914131 | middle |  |  |  |  |  |  |  |  |  |  |  |
| YJM1444 | XVI | 7918 | 7927 | 914131 | middle |  |  |  |  |  |  |  |  |  |  |  |
| YJM1444 | XVI | 912945 | 912954 | 914131 | middle |  |  |  |  |  |  |  |  |  |  |  |
| YJM1444 | XVI | 914122 | 914131 | 914131 | rightend | YPR201W | 99.588 | 1215 | 5 | 0 | 1 | 1215 | 909775 | 910989 | 0 | 2217 |
| YJM1447 | I | 1 | 10 | 198380 | leftend | YAL063C | 87.235 | 1559 | 196 | 1 | 1482 | 3037 | 2543 | 985 | 0 | 1773 |
| YJM1447 | I | 2544 | 2553 | 198380 | middle |  |  |  |  |  |  |  |  |  |  |  |
| YJM1447 | I | 193157 | 193166 | 198380 | middle |  |  |  |  |  |  |  |  |  |  |  |
| YJM1447 | I | 198371 | 198380 | 198380 | rightend | YAR050W | 90.696 | 1580 | 106 | 13 | 2716 | 4282 | 176748 | 178299 | 0 | 2065 |
| YJM1447 | II | 1 | 10 | 781068 | leftend | YNCB0001W | 100 | 84 | 0 | 0 | 1 | 84 | 1304 | 1387 | 4.16E-38 | 156 |
| YJM1447 | II | 781059 | 781068 | 781068 | rightend | YBR297W | 98.34 | 1205 | 20 | 0 | 1 | 1205 | 774836 | 776040 | 0 | 2115 |
| YJM1447 | III | 1 | 10 | 322864 | leftend | YCL073C | 99.08 | 1848 | 17 | 0 | 1 | 1848 | 6016 | 4169 | 0 | 3319 |
| YJM1447 | III | 322855 | 322864 | 322864 | rightend | YCR101C | 98.909 | 550 | 5 | 1 | 1 | 549 | 296682 | 296133 | 0 | 981 |
| YJM1447 | IV | 1 | 10 | 1463002 | leftend | YDL243C | 93.333 | 990 | 66 | 0 | 1 | 990 | 16998 | 16009 | 0 | 1463 |
| YJM1447 | IV | 1462993 | 1463002 | 1463002 | rightend | YDR541C | 99.807 | 1035 | 2 | 0 | 1 | 1035 | 1460202 | 1459168 | 0 | 1901 |
| YJM1447 | V | 1 | 10 | 554965 | leftend | YEL070W | 99.072 | 1509 | 14 | 0 | 1 | 1509 | 18761 | 20269 | 0 | 2710 |
| YJM1447 | V | 554956 | 554965 | 554965 | rightend | YER187W | 99.765 | 426 | 1 | 0 | 1 | 426 | 553532 | 553957 | 0 | 782 |
| YJM1447 | VI | 1 | 10 | 279663 | leftend | YFL051C | 98.894 | 452 | 5 | 0 | 1 | 452 | 41055 | 40604 | 0 | 808 |
| YJM1447 | VI | 279654 | 279663 | 279663 | rightend | YFR055W | 99.511 | 1022 | 5 | 0 | 1 | 1022 | 270073 | 271094 | 0 | 1860 |
| YJM1447 | VII | 1 | 10 | 1065894 | leftend | YGL258W | 98.39 | 621 | 10 | 0 | 1 | 621 | 18897 | 19517 | 0 | 1092 |
| YJM1447 | VII | 6401 | 6410 | 1065894 | middle |  |  |  |  |  |  |  |  |  |  |  |
| YJM1447 | VII | 1065885 | 1065894 | 1065894 | rightend | YGR287C | 97.232 | 1770 | 49 | 0 | 1 | 1770 | 1041879 | 1040110 | 0 | 2998 |
| YJM1447 | VIII | 1 | 10 | 521373 | leftend | YHL044W | 98.87 | 708 | 8 | 0 | 1 | 708 | 1521 | 2228 | 0 | 1264 |
| YJM1447 | VIII | 508926 | 508935 | 521373 | middle |  |  |  |  |  |  |  |  |  |  |  |
| YJM1447 | VIII | 521364 | 521373 | 521373 | rightend | YHR211W | 84.471 | 1275 | 187 | 7 | 1 | 1268 | 507655 | 508925 | 0 | 1247 |
| YJM1447 | IX | 1 | 10 | 416330 | leftend | YIL169C | 97.5 | 1400 | 29 | 4 | 520 | 1919 | 9616 | 8223 | 0 | 2386 |
| YJM1447 | IX | 999 | 1008 | 416330 | middle |  |  |  |  |  |  |  |  |  |  |  |
| YJM1447 | IX | 2337 | 2346 | 416330 | middle |  |  |  |  |  |  |  |  |  |  |  |
| YJM1447 | IX | 3115 | 3124 | 416330 | middle |  |  |  |  |  |  |  |  |  |  |  |
| YJM1447 | IX | 414488 | 414497 | 416330 | middle |  |  |  |  |  |  |  |  |  |  |  |
| YJM1447 | IX | 415431 | 415440 | 416330 | middle |  |  |  |  |  |  |  |  |  |  |  |
| YJM1447 | IX | 416321 | 416330 | 416330 | rightend | YIR039C | 98.327 | 1614 | 26 | 1 | 1 | 1614 | 408616 | 407004 | 0 | 2830 |
| YJM1447 | X | 1 | 10 | 701453 | leftend | YJL213W | 98.293 | 996 | 17 | 0 | 1 | 996 | 20949 | 21944 | 0 | 1746 |
| YJM1447 | X | 701444 | 701453 | 701453 | rightend | YJR152W | 99.265 | 1632 | 12 | 0 | 1 | 1632 | 691410 | 693041 | 0 | 2948 |
| YJM1447 | XI | 1 | 10 | 659649 | leftend | YKL222C | 99.717 | 2118 | 6 | 0 | 1 | 2118 | 3475 | 1358 | 0 | 3879 |
| YJM1447 | XI | 659640 | 659649 | 659649 | rightend | YKR101W | 99.135 | 1965 | 17 | 0 | 1 | 1965 | 639675 | 641639 | 0 | 3535 |
| YJM1447 | XII | 1 | 10 | 1562717 | leftend | YLL062C | 99.59 | 975 | 4 | 0 | 1 | 975 | 4013 | 3039 | 0 | 1779 |
| YJM1447 | XII | 1562708 | 1562717 | 1562717 | rightend | YLR460C | 94.876 | 1132 | 56 | 2 | 1 | 1131 | 1560944 | 1559814 | 0 | 1768 |
| YJM1447 | XIII | 1 | 10 | 901376 | leftend | YML131W | 98.816 | 1098 | 13 | 0 | 1 | 1098 | 2390 | 3487 | 0 | 1956 |
| YJM1447 | XIII | 897678 | 897687 | 901376 | middle |  |  |  |  |  |  |  |  |  |  |  |
| YJM1447 | XIII | 901367 | 901376 | 901376 | rightend | YMR321C | 98.722 | 313 | 4 | 0 | 6 | 318 | 889864 | 889552 | 5.37E-158 | 556 |
| YJM1447 | XIV | 1 | 10 | 765921 | leftend | YNL332W | 92.958 | 71 | 5 | 0 | 1 | 71 | 6995 | 7065 | 2.4E-21 | 104 |
| YJM1447 | XIV | 765912 | 765921 | 765921 | rightend | YNR071C | 99.417 | 1029 | 6 | 0 | 1 | 1029 | 746056 | 745028 | 0 | 1868 |
| YJM1447 | XV | 1 | 10 | 1060595 | leftend | YOL154W | 99.333 | 750 | 5 | 0 | 1 | 750 | 23057 | 23806 | 0 | 1358 |
| YJM1447 | XV | 1055528 | 1055537 | 1060595 | middle |  |  |  |  |  |  |  |  |  |  |  |
| YJM1447 | XV | 1060586 | 1060595 | 1060595 | rightend | YOR381W | 98.596 | 2136 | 30 | 0 | 1 | 2136 | 1028350 | 1030485 | 0 | 3779 |
| YJM1447 | XVI | 1 | 10 | 912634 | leftend | YPL274W | 97.562 | 1764 | 43 | 0 | 1 | 1764 | 13106 | 14869 | 0 | 3020 |
| YJM1447 | XVI | 261 | 270 | 912634 | middle |  |  |  |  |  |  |  |  |  |  |  |
| YJM1447 | XVI | 901110 | 901119 | 912634 | middle |  |  |  |  |  |  |  |  |  |  |  |
| YJM1447 | XVI | 912634 | 912634 | 912634 | rightend_complete | YPR198W | 99.326 | 1632 | 11 | 0 | 1 | 1632 | 895740 | 897371 | 0 | 2953 |
| YJM1450 | I | 1 |  | 194656 | leftend_complete | YAL063C | 85.355 | 1632 | 234 | 5 | 1 | 1630 | 16425 | 14797 | 0 | 1685 |
| YJM1450 | I | 1963 | 1972 | 194656 | middle |  |  |  |  |  |  |  |  |  |  |  |
| YJM1450 | I | 3547 | 3556 | 194656 | middle |  |  |  |  |  |  |  |  |  |  |  |
| YJM1450 | I | 191696 | 191705 | 194656 | middle |  |  |  |  |  |  |  |  |  |  |  |
| YJM1450 | I | 194647 | 194656 | 194656 | rightend | YAR050W | 95.08 | 813 | 40 | 0 | 1 | 813 | 190877 | 191689 | 0 | 1280 |
| YJM1450 | II | 1 | 10 | 791506 | leftend | YNCB0001W | 100 | 84 | 0 | 0 | 1 | 84 | 8036 | 8119 | 4.16E-38 | 156 |
| YJM1450 | II | 827 | 836 | 791506 | middle |  |  |  |  |  |  |  |  |  |  |  |
| YJM1450 | II | 3052 | 3061 | 791506 | middle |  |  |  |  |  |  |  |  |  |  |  |
| YJM1450 | II | 4225 | 4234 | 791506 | middle |  |  |  |  |  |  |  |  |  |  |  |
| YJM1450 | II | 4994 | 5003 | 791506 | middle |  |  |  |  |  |  |  |  |  |  |  |
| YJM1450 | II | 6285 | 6294 | 791506 | middle |  |  |  |  |  |  |  |  |  |  |  |
| YJM1450 | II | 790973 | 790982 | 791506 | middle |  |  |  |  |  |  |  |  |  |  |  |
| YJM1450 | II | 791497 | 791506 | 791506 | rightend | YBR297W | 91.825 | 1419 | 98 | 14 | 1 | 1407 | 780622 | 782034 | 0 | 1962 |
| YJM1450 | III | 1 | 10 | 310591 | leftend | YCL073C | 99.621 | 1848 | 7 | 0 | 1 | 1848 | 8481 | 6634 | 0 | 3374 |
| YJM1450 | III | 941 | 950 | 310591 | middle |  |  |  |  |  |  |  |  |  |  |  |
| YJM1450 | III | 310591 | 310591 | 310591 | rightend_complete | YCR101C | 98.182 | 550 | 9 | 1 | 1 | 549 | 297005 | 296456 | 0 | 959 |
| YJM1450 | IV | 1 | 10 | 1484524 | leftend | YDL243C | 93.939 | 990 | 60 | 0 | 1 | 990 | 27884 | 26895 | 0 | 1496 |
| YJM1450 | IV | 1473 | 1482 | 1484524 | middle |  |  |  |  |  |  |  |  |  |  |  |
| YJM1450 | IV | 1481958 | 1481967 | 1484524 | middle |  |  |  |  |  |  |  |  |  |  |  |
| YJM1450 | IV | 1483066 | 1483075 | 1484524 | middle |  |  |  |  |  |  |  |  |  |  |  |
| YJM1450 | IV | 1484286 | 1484295 | 1484524 | middle |  |  |  |  |  |  |  |  |  |  |  |
| YJM1450 | IV | 1484515 | 1484524 | 1484524 | rightend | YDR541C | 99.42 | 1035 | 6 | 0 | 1 | 1035 | 1479224 | 1478190 | 0 | 1879 |
| YJM1450 | V | 1 |  | 552442 | leftend_complete | YEL070W | 99.271 | 1509 | 11 | 0 | 1 | 1509 | 14641 | 16149 | 0 | 2726 |
| YJM1450 | V | 1764 | 1773 | 552442 | middle |  |  |  |  |  |  |  |  |  |  |  |
| YJM1450 | V | 552433 | 552442 | 552442 | rightend | YER187W | 99.296 | 426 | 3 | 0 | 1 | 426 | 549066 | 549491 | 0 | 771 |
| YJM1450 | VI | 1 | 10 | 236770 | leftend | YFL051C | 98.23 | 452 | 8 | 0 | 1 | 452 | 850 | 399 | 0 | 791 |
| YJM1450 | VI | 236761 | 236770 | 236770 | rightend | YFR055W | 99.609 | 1022 | 4 | 0 | 1 | 1022 | 227178 | 228199 | 0 | 1866 |
| YJM1450 | VII | 1 |  | 1050320 | leftend_complete | YGL258W | 97.585 | 621 | 15 | 0 | 1 | 621 | 11227 | 11847 | 0 | 1064 |
| YJM1450 | VII | 1048934 | 1048943 | 1050320 | middle |  |  |  |  |  |  |  |  |  |  |  |
| YJM1450 | VII | 1050311 | 1050320 | 1050320 | rightend | YGR287C | 98.757 | 1770 | 22 | 0 | 1 | 1770 | 1032518 | 1030749 | 0 | 3147 |
| YJM1450 | VIII | 1 | 10 | 545639 | leftend | YHL044W | 98.588 | 708 | 10 | 0 | 1 | 708 | 9023 | 9730 | 0 | 1253 |
| YJM1450 | VIII | 951 | 960 | 545639 | middle |  |  |  |  |  |  |  |  |  |  |  |
| YJM1450 | VIII | 520038 | 520047 | 545639 | middle |  |  |  |  |  |  |  |  |  |  |  |
| YJM1450 | VIII | 536807 | 536816 | 545639 | middle |  |  |  |  |  |  |  |  |  |  |  |
| YJM1450 | VIII | 540635 | 540644 | 545639 | middle |  |  |  |  |  |  |  |  |  |  |  |
| YJM1450 | VIII | 542255 | 542264 | 545639 | middle |  |  |  |  |  |  |  |  |  |  |  |
| YJM1450 | VIII | 543687 | 543696 | 545639 | middle |  |  |  |  |  |  |  |  |  |  |  |
| YJM1450 | VIII | 545639 | 545639 | 545639 | rightend_complete | YHR211W | 87.269 | 1516 | 172 | 5 | 826 | 2323 | 520688 | 522200 | 0 | 1711 |
| YJM1450 | IX | 1 | 10 | 421017 | leftend | YIL169C | 96.725 | 1374 | 39 | 2 | 478 | 1851 | 17615 | 16248 | 0 | 2283 |
| YJM1450 | IX | 1605 | 1614 | 421017 | middle |  |  |  |  |  |  |  |  |  |  |  |
| YJM1450 | IX | 420392 | 420401 | 421017 | middle |  |  |  |  |  |  |  |  |  |  |  |
| YJM1450 | IX | 421017 | 421017 | 421017 | rightend_complete | YIR039C | 99.009 | 1614 | 16 | 0 | 1 | 1614 | 418750 | 417137 | 0 | 2892 |
| YJM1450 | X | 1 |  | 809591 | leftend_complete | YJL213W | 99.699 | 996 | 3 | 0 | 1 | 996 | 33249 | 34244 | 0 | 1823 |
| YJM1450 | X | 1158 | 1167 | 809591 | middle |  |  |  |  |  |  |  |  |  |  |  |
| YJM1450 | X | 809591 | 809591 | 809591 | rightend_complete | YJR152W | 100 | 1632 | 0 | 0 | 1 | 1632 | 704366 | 705997 | 0 | 3014 |
| YJM1450 | XI | 1 | 10 | 656532 | leftend | YKL222C | 99.622 | 2118 | 8 | 0 | 1 | 2118 | 3122 | 1005 | 0 | 3868 |
| YJM1450 | XI | 656523 | 656532 | 656532 | rightend | YKR101W | 99.695 | 1965 | 6 | 0 | 1 | 1965 | 637583 | 639547 | 0 | 3596 |
| YJM1450 | XII | 1 |  | 2314440 | leftend_complete | YLL062C | 99.692 | 975 | 3 | 0 | 1 | 975 | 2606 | 1632 | 0 | 1784 |
| YJM1450 | XII | 2309568 | 2309577 | 2314440 | middle |  |  |  |  |  |  |  |  |  |  |  |
| YJM1450 | XII | 2310204 | 2310213 | 2314440 | middle |  |  |  |  |  |  |  |  |  |  |  |
| YJM1450 | XII | 2310945 | 2310954 | 2314440 | middle |  |  |  |  |  |  |  |  |  |  |  |
| YJM1450 | XII | 2314440 | 2314440 | 2314440 | rightend_complete | YLR460C | 94.346 | 1132 | 62 | 2 | 1 | 1131 | 2307944 | 2306814 | 0 | 1735 |
| YJM1450 | XIII | 1 | 10 | 902274 | leftend | YML131W | 100 | 1098 | 0 | 0 | 1 | 1098 | 2353 | 3450 | 0 | 2028 |
| YJM1450 | XIII | 899725 | 899734 | 902274 | middle |  |  |  |  |  |  |  |  |  |  |  |
| YJM1450 | XIII | 902265 | 902274 | 902274 | rightend | YMR321C | 99.361 | 313 | 2 | 0 | 6 | 318 | 885688 | 885376 | 2.48E-161 | 568 |
| YJM1450 | XIV | 1 | 10 | 761059 | leftend | YNL332W | 98.701 | 462 | 6 | 0 | 562 | 1023 | 11546 | 12007 | 0 | 821 |
| YJM1450 | XIV | 10431 | 10440 | 761059 | middle |  |  |  |  |  |  |  |  |  |  |  |
| YJM1450 | XIV | 11536 | 11545 | 761059 | middle |  |  |  |  |  |  |  |  |  |  |  |
| YJM1450 | XIV | 761050 | 761059 | 761059 | rightend | YNR071C | 99.806 | 1029 | 2 | 0 | 1 | 1029 | 750447 | 749419 | 0 | 1890 |
| YJM1450 | XV | 1 | 10 | 1049305 | leftend | YOL154W | 99.467 | 750 | 4 | 0 | 1 | 750 | 22871 | 23620 | 0 | 1363 |
| YJM1450 | XV | 1047866 | 1047875 | 1049305 | middle |  |  |  |  |  |  |  |  |  |  |  |
| YJM1450 | XV | 1048604 | 1048613 | 1049305 | middle |  |  |  |  |  |  |  |  |  |  |  |
| YJM1450 | XV | 1049296 | 1049305 | 1049305 | rightend | YOR381W | 98.408 | 2136 | 34 | 0 | 1 | 2136 | 1022358 | 1024493 | 0 | 3757 |
| YJM1450 | XVI | 1 |  | 920141 | leftend_complete | YPL274W | 99.943 | 1764 | 1 | 0 | 1 | 1764 | 29680 | 31443 | 0 | 3253 |
| YJM1450 | XVI | 920141 | 920141 | 920141 | rightend_complete | YPR201W | 100 | 1215 | 0 | 0 | 1 | 1215 | 916871 | 918085 | 0 | 2244 |
| YJM1460 | I | 1 | 10 | 195651 | leftend | YAL063C | 84.813 | 1633 | 242 | 6 | 1 | 1630 | 12251 | 10622 | 0 | 1637 |
| YJM1460 | I | 9485 | 9494 | 195651 | middle |  |  |  |  |  |  |  |  |  |  |  |
| YJM1460 | I | 9723 | 9732 | 195651 | middle |  |  |  |  |  |  |  |  |  |  |  |
| YJM1460 | I | 182242 | 182251 | 195651 | middle |  |  |  |  |  |  |  |  |  |  |  |
| YJM1460 | I | 195642 | 195651 | 195651 | rightend | YAR050W | 86.423 | 1694 | 181 | 15 | 2451 | 4100 | 11421 | 9733 | 0 | 1808 |
| YJM1460 | II | 1 | 10 | 787262 | leftend | YNCB0001W | 100 | 84 | 0 | 0 | 1 | 84 | 1313 | 1396 | 4.16E-38 | 156 |
| YJM1460 | II | 787253 | 787262 | 787262 | rightend | YBR297W | 97.925 | 1205 | 25 | 0 | 1 | 1205 | 774112 | 775316 | 0 | 2087 |
| YJM1460 | III | 1 | 10 | 300816 | leftend | YCL073C | 98.918 | 1848 | 20 | 0 | 1 | 1848 | 7533 | 5686 | 0 | 3302 |
| YJM1460 | III | 300807 | 300816 | 300816 | rightend | YCR101C | 98 | 550 | 10 | 1 | 1 | 549 | 297037 | 296488 | 0 | 953 |
| YJM1460 | IV | 1 | 10 | 1457464 | leftend | YDL243C | 93.838 | 990 | 61 | 0 | 1 | 990 | 11931 | 10942 | 0 | 1491 |
| YJM1460 | IV | 1456938 | 1456947 | 1457464 | middle |  |  |  |  |  |  |  |  |  |  |  |
| YJM1460 | IV | 1457455 | 1457464 | 1457464 | rightend | YDR541C | 99.614 | 1035 | 4 | 0 | 1 | 1035 | 1453989 | 1452955 | 0 | 1890 |
| YJM1460 | V | 1 | 10 | 551869 | leftend | YEL070W | 99.47 | 1509 | 8 | 0 | 1 | 1509 | 13334 | 14842 | 0 | 2743 |
| YJM1460 | V | 551860 | 551869 | 551869 | rightend | YER187W | 99.765 | 426 | 1 | 0 | 1 | 426 | 547120 | 547545 | 0 | 782 |
| YJM1460 | VI | 1 | 10 | 272513 | leftend | YFL051C | 100 | 483 | 0 | 0 | 1 | 483 | 33925 | 33443 | 0 | 893 |
| YJM1460 | VI | 272504 | 272513 | 272513 | rightend | YFR055W | 99.511 | 1022 | 5 | 0 | 1 | 1022 | 261971 | 262992 | 0 | 1860 |
| YJM1460 | VII | 1 | 10 | 1056068 | leftend | YGL258W | 99.356 | 621 | 4 | 0 | 1 | 621 | 4429 | 5049 | 0 | 1125 |
| YJM1460 | VII | 1039979 | 1039988 | 1056068 | middle |  |  |  |  |  |  |  |  |  |  |  |
| YJM1460 | VII | 1055503 | 1055512 | 1056068 | middle |  |  |  |  |  |  |  |  |  |  |  |
| YJM1460 | VII | 1056068 | 1056068 | 1056068 | rightend_complete | YGR287C | 97.966 | 1770 | 36 | 0 | 1 | 1770 | 1026787 | 1025018 | 0 | 3070 |
| YJM1460 | VIII | 1 | 10 | 527557 | leftend | YHL044W | 98.729 | 708 | 9 | 0 | 1 | 708 | 6958 | 7665 | 0 | 1258 |
| YJM1460 | VIII | 1445 | 1454 | 527557 | middle |  |  |  |  |  |  |  |  |  |  |  |
| YJM1460 | VIII | 515537 | 515546 | 527557 | middle |  |  |  |  |  |  |  |  |  |  |  |
| YJM1460 | VIII | 521570 | 521579 | 527557 | middle |  |  |  |  |  |  |  |  |  |  |  |
| YJM1460 | VIII | 525377 | 525386 | 527557 | middle |  |  |  |  |  |  |  |  |  |  |  |
| YJM1460 | VIII | 525596 | 525605 | 527557 | middle |  |  |  |  |  |  |  |  |  |  |  |
| YJM1460 | VIII | 527548 | 527557 | 527557 | rightend | YHR211W | 80.74 | 1594 | 281 | 17 | 1 | 1581 | 513954 | 515534 | 0 | 1219 |
| YJM1460 | IX | 1 | 10 | 409675 | leftend | YIL169C | 97.788 | 1989 | 38 | 2 | 478 | 2466 | 5285 | 3303 | 0 | 3424 |
| YJM1460 | IX | 408916 | 408925 | 409675 | middle |  |  |  |  |  |  |  |  |  |  |  |
| YJM1460 | IX | 409666 | 409675 | 409675 | rightend | YIR039C | 99.69 | 1614 | 5 | 0 | 1 | 1614 | 407384 | 405771 | 0 | 2953 |
| YJM1460 | X | 1 | 10 | 708426 | leftend | YJL213W | 99.9 | 996 | 1 | 0 | 1 | 996 | 10623 | 11618 | 0 | 1834 |
| YJM1460 | X | 706874 | 706883 | 708426 | middle |  |  |  |  |  |  |  |  |  |  |  |
| YJM1460 | X | 708209 | 708218 | 708426 | middle |  |  |  |  |  |  |  |  |  |  |  |
| YJM1460 | X | 708426 | 708426 | 708426 | rightend_complete | YJR152W | 99.326 | 1632 | 11 | 0 | 1 | 1632 | 681468 | 683099 | 0 | 2953 |
| YJM1460 | XI | 1 | 10 | 666135 | leftend | YKL222C | 99.906 | 2118 | 2 | 0 | 1 | 2118 | 3453 | 1336 | 0 | 3901 |
| YJM1460 | XI | 666126 | 666135 | 666135 | rightend | YKR101W | 99.695 | 1965 | 6 | 0 | 1 | 1965 | 638197 | 640161 | 0 | 3596 |
| YJM1460 | XII | 1 | 10 | 2131493 | leftend | YLL062C | 99.692 | 975 | 3 | 0 | 1 | 975 | 4171 | 3197 | 0 | 1784 |
| YJM1460 | XII | 2129739 | 2129748 | 2131493 | middle |  |  |  |  |  |  |  |  |  |  |  |
| YJM1460 | XII | 2131493 | 2131493 | 2131493 | rightend_complete | YLR460C | 99.912 | 1131 | 1 | 0 | 1 | 1131 | 2126960 | 2125830 | 0 | 2084 |
| YJM1460 | XIII | 1 | 10 | 895899 | leftend | YML131W | 98.816 | 1098 | 13 | 0 | 1 | 1098 | 4210 | 5307 | 0 | 1956 |
| YJM1460 | XIII | 1411 | 1420 | 895899 | middle |  |  |  |  |  |  |  |  |  |  |  |
| YJM1460 | XIII | 895899 | 895899 | 895899 | rightend_complete | YMR321C | 99.686 | 318 | 1 | 0 | 1 | 318 | 889443 | 889126 | 8.85E-166 | 582 |
| YJM1460 | XIV | 1 |  | 758643 | leftend_complete | YNL332W | 98.729 | 1023 | 13 | 0 | 1 | 1023 | 16113 | 17135 | 0 | 1818 |
| YJM1460 | XIV | 758634 | 758643 | 758643 | rightend | YNR071C | 99.515 | 1030 | 4 | 1 | 1 | 1029 | 755917 | 754888 | 0 | 1873 |
| YJM1460 | XV | 1 |  | 1044445 | leftend_complete | YOL154W | 99.733 | 750 | 2 | 0 | 1 | 750 | 25323 | 26072 | 0 | 1375 |
| YJM1460 | XV | 2594 | 2603 | 1044445 | middle |  |  |  |  |  |  |  |  |  |  |  |
| YJM1460 | XV | 3630 | 3639 | 1044445 | middle |  |  |  |  |  |  |  |  |  |  |  |
| YJM1460 | XV | 4097 | 4106 | 1044445 | middle |  |  |  |  |  |  |  |  |  |  |  |
| YJM1460 | XV | 5232 | 5241 | 1044445 | middle |  |  |  |  |  |  |  |  |  |  |  |
| YJM1460 | XV | 1043837 | 1043846 | 1044445 | middle |  |  |  |  |  |  |  |  |  |  |  |
| YJM1460 | XV | 1044436 | 1044445 | 1044445 | rightend | YOR381W | 98.596 | 2136 | 30 | 0 | 1 | 2136 | 1024209 | 1026344 | 0 | 3779 |
| YJM1460 | XVI | 1 | 10 | 897468 | leftend | YPL274W | 97.959 | 1764 | 36 | 0 | 1 | 1764 | 12574 | 14337 | 0 | 3059 |
| YJM1460 | XVI | 456 | 465 | 897468 | middle |  |  |  |  |  |  |  |  |  |  |  |
| YJM1460 | XVI | 674 | 683 | 897468 | middle |  |  |  |  |  |  |  |  |  |  |  |
| YJM1460 | XVI | 896886 | 896895 | 897468 | middle |  |  |  |  |  |  |  |  |  |  |  |
| YJM1460 | XVI | 897468 | 897468 | 897468 | rightend_complete | YPR201W | 99.588 | 1215 | 5 | 0 | 1 | 1215 | 894245 | 895459 | 0 | 2217 |
| YJM1463 | I | 1 | 10 | 184312 | leftend | YAL063C | 91.483 | 998 | 56 | 13 | 2966 | 3962 | 10444 | 9475 | 0 | 1345 |
| YJM1463 | I | 184303 | 184312 | 184312 | rightend | YAR050W | 92.687 | 670 | 28 | 11 | 3605 | 4253 | 10444 | 9775 | 0 | 946 |
| YJM1463 | II | 1 | 10 | 785342 | leftend | YNCB0001W | 100 | 84 | 0 | 0 | 1 | 84 | 3568 | 3651 | 4.16E-38 | 156 |
| YJM1463 | II | 500 | 509 | 785342 | middle |  |  |  |  |  |  |  |  |  |  |  |
| YJM1463 | II | 1240 | 1249 | 785342 | middle |  |  |  |  |  |  |  |  |  |  |  |
| YJM1463 | II | 784812 | 784821 | 785342 | middle |  |  |  |  |  |  |  |  |  |  |  |
| YJM1463 | II | 785333 | 785342 | 785342 | rightend | YBR297W | 99.431 | 1407 | 8 | 0 | 1 | 1407 | 776311 | 777717 | 0 | 2555 |
| YJM1463 | III | 1 | 10 | 310796 | leftend | YCL073C | 98.755 | 1848 | 23 | 0 | 1 | 1848 | 8365 | 6518 | 0 | 3286 |
| YJM1463 | III | 302914 | 302923 | 310796 | middle |  |  |  |  |  |  |  |  |  |  |  |
| YJM1463 | III | 310787 | 310796 | 310796 | rightend | YCR099C | 88.286 | 461 | 51 | 3 | 9 | 468 | 295333 | 294875 | 2.18E-156 | 549 |
| YJM1463 | IV | 1 |  | 1469848 | leftend_complete | YDL243C | 93.463 | 979 | 64 | 0 | 1 | 979 | 17020 | 16042 | 0 | 1454 |
| YJM1463 | IV | 1469180 | 1469189 | 1469848 | middle |  |  |  |  |  |  |  |  |  |  |  |
| YJM1463 | IV | 1469839 | 1469848 | 1469848 | rightend | YDR541C | 99.42 | 1035 | 6 | 0 | 1 | 1035 | 1466379 | 1465345 | 0 | 1879 |
| YJM1463 | V | 1 | 10 | 552447 | leftend | YEL070W | 99.337 | 1509 | 10 | 0 | 1 | 1509 | 15429 | 16937 | 0 | 2732 |
| YJM1463 | V | 707 | 716 | 552447 | middle |  |  |  |  |  |  |  |  |  |  |  |
| YJM1463 | V | 552438 | 552447 | 552447 | rightend | YER187W | 99.296 | 426 | 3 | 0 | 1 | 426 | 550372 | 550797 | 0 | 771 |
| YJM1463 | VI | 1 |  | 236499 | leftend_complete | YFL051C | 98.23 | 452 | 8 | 0 | 1 | 452 | 3091 | 2640 | 0 | 791 |
| YJM1463 | VI | 267 | 276 | 236499 | middle |  |  |  |  |  |  |  |  |  |  |  |
| YJM1463 | VI | 1003 | 1012 | 236499 | middle |  |  |  |  |  |  |  |  |  |  |  |
| YJM1463 | VI | 236499 | 236499 | 236499 | rightend_complete | YFR055W | 99.706 | 1022 | 3 | 0 | 1 | 1022 | 230527 | 231548 | 0 | 1871 |
| YJM1463 | VII | 1 |  | 1055615 | leftend_complete | YGL258W | 92.593 | 621 | 46 | 0 | 1 | 621 | 12732 | 13352 | 0 | 893 |
| YJM1463 | VII | 1721 | 1730 | 1055615 | middle |  |  |  |  |  |  |  |  |  |  |  |
| YJM1463 | VII | 8649 | 8658 | 1055615 | middle |  |  |  |  |  |  |  |  |  |  |  |
| YJM1463 | VII | 1052599 | 1052608 | 1055615 | middle |  |  |  |  |  |  |  |  |  |  |  |
| YJM1463 | VII | 1053240 | 1053249 | 1055615 | middle |  |  |  |  |  |  |  |  |  |  |  |
| YJM1463 | VII | 1055615 | 1055615 | 1055615 | rightend_complete | YGR287C | 98.927 | 1770 | 19 | 0 | 1 | 1770 | 1033747 | 1031978 | 0 | 3164 |
| YJM1463 | VIII | 1 | 10 | 536529 | leftend | YHL044W | 98.305 | 708 | 12 | 0 | 1 | 708 | 7519 | 8226 | 0 | 1242 |
| YJM1463 | VIII | 526394 | 526403 | 536529 | middle |  |  |  |  |  |  |  |  |  |  |  |
| YJM1463 | VIII | 534497 | 534506 | 536529 | middle |  |  |  |  |  |  |  |  |  |  |  |
| YJM1463 | VIII | 534720 | 534729 | 536529 | middle |  |  |  |  |  |  |  |  |  |  |  |
| YJM1463 | VIII | 535538 | 535547 | 536529 | middle |  |  |  |  |  |  |  |  |  |  |  |
| YJM1463 | VIII | 536520 | 536529 | 536529 | rightend | YHR211W | 88.194 | 1711 | 156 | 14 | 1231 | 2918 | 516087 | 517774 | 0 | 1999 |
| YJM1463 | IX | 1 | 10 | 402650 | leftend | YIL169C | 94.249 | 1252 | 70 | 2 | 562 | 1812 | 2396 | 1146 | 0 | 1912 |
| YJM1463 | IX | 402641 | 402650 | 402650 | rightend | YIR039C | 99.009 | 1614 | 16 | 0 | 1 | 1614 | 401198 | 399585 | 0 | 2892 |
| YJM1463 | X | 1 |  | 738954 | leftend_complete | YJL213W | 99.498 | 996 | 5 | 0 | 1 | 996 | 49155 | 50150 | 0 | 1812 |
| YJM1463 | X | 598 | 607 | 738954 | middle |  |  |  |  |  |  |  |  |  |  |  |
| YJM1463 | X | 2036 | 2045 | 738954 | middle |  |  |  |  |  |  |  |  |  |  |  |
| YJM1463 | X | 2900 | 2909 | 738954 | middle |  |  |  |  |  |  |  |  |  |  |  |
| YJM1463 | X | 3210 | 3219 | 738954 | middle |  |  |  |  |  |  |  |  |  |  |  |
| YJM1463 | X | 6183 | 6192 | 738954 | middle |  |  |  |  |  |  |  |  |  |  |  |
| YJM1463 | X | 7030 | 7039 | 738954 | middle |  |  |  |  |  |  |  |  |  |  |  |
| YJM1463 | X | 7910 | 7919 | 738954 | middle |  |  |  |  |  |  |  |  |  |  |  |
| YJM1463 | X | 8395 | 8404 | 738954 | middle |  |  |  |  |  |  |  |  |  |  |  |
| YJM1463 | X | 738954 | 738954 | 738954 | rightend_complete | YJR152W | 99.816 | 1632 | 3 | 0 | 1 | 1632 | 718541 | 720172 | 0 | 2998 |
| YJM1463 | XI | 1 |  | 647495 | leftend_complete | YKL222C | 99.248 | 1330 | 10 | 0 | 789 | 2118 | 4163 | 2834 | 0 | 2401 |
| YJM1463 | XI | 1876 | 1885 | 647495 | middle |  |  |  |  |  |  |  |  |  |  |  |
| YJM1463 | XI | 647486 | 647495 | 647495 | rightend | YKR101W | 99.746 | 1965 | 5 | 0 | 1 | 1965 | 639695 | 641659 | 0 | 3602 |
| YJM1463 | XII | 1 | 10 | 2795807 | leftend | YLL062C | 99.897 | 975 | 1 | 0 | 1 | 975 | 1932 | 958 | 0 | 1796 |
| YJM1463 | XII | 2792780 | 2792789 | 2795807 | middle |  |  |  |  |  |  |  |  |  |  |  |
| YJM1463 | XII | 2793015 | 2793024 | 2795807 | middle |  |  |  |  |  |  |  |  |  |  |  |
| YJM1463 | XII | 2794864 | 2794873 | 2795807 | middle |  |  |  |  |  |  |  |  |  |  |  |
| YJM1463 | XII | 2795798 | 2795807 | 2795807 | rightend | YLR460C | 94.336 | 1130 | 62 | 2 | 3 | 1131 | 2791110 | 2789982 | 0 | 1731 |
| YJM1463 | XIII | 1 | 10 | 908702 | leftend | YML131W | 99.818 | 1098 | 2 | 0 | 1 | 1098 | 2947 | 4044 | 0 | 2017 |
| YJM1463 | XIII | 918 | 927 | 908702 | middle |  |  |  |  |  |  |  |  |  |  |  |
| YJM1463 | XIII | 904492 | 904501 | 908702 | middle |  |  |  |  |  |  |  |  |  |  |  |
| YJM1463 | XIII | 908693 | 908702 | 908702 | rightend | YMR321C | 99.361 | 313 | 2 | 0 | 6 | 318 | 887833 | 887521 | 2.48E-161 | 568 |
| YJM1463 | XIV | 1 |  | 759993 | leftend_complete | YNL332W | 98.24 | 1023 | 18 | 0 | 1 | 1023 | 8547 | 9569 | 0 | 1790 |
| YJM1463 | XIV | 764 | 773 | 759993 | middle |  |  |  |  |  |  |  |  |  |  |  |
| YJM1463 | XIV | 3025 | 3034 | 759993 | middle |  |  |  |  |  |  |  |  |  |  |  |
| YJM1463 | XIV | 759984 | 759993 | 759993 | rightend | YNR071C | 98.834 | 1029 | 12 | 0 | 1 | 1029 | 747208 | 746180 | 0 | 1834 |
| YJM1463 | XV | 1 | 10 | 1041323 | leftend | YOL154W | 99.333 | 750 | 5 | 0 | 1 | 750 | 15879 | 16628 | 0 | 1358 |
| YJM1463 | XV | 1040636 | 1040645 | 1041323 | middle |  |  |  |  |  |  |  |  |  |  |  |
| YJM1463 | XV | 1041314 | 1041323 | 1041323 | rightend | YOR381W | 100 | 2136 | 0 | 0 | 1 | 2136 | 1013519 | 1015654 | 0 | 3945 |
| YJM1463 | XVI | 1 | 10 | 915577 | leftend | YPL274W | 99.263 | 1764 | 13 | 0 | 1 | 1764 | 15772 | 17535 | 0 | 3186 |
| YJM1463 | XVI | 2548 | 2557 | 915577 | middle |  |  |  |  |  |  |  |  |  |  |  |
| YJM1463 | XVI | 909676 | 909685 | 915577 | middle |  |  |  |  |  |  |  |  |  |  |  |
| YJM1463 | XVI | 910392 | 910401 | 915577 | middle |  |  |  |  |  |  |  |  |  |  |  |
| YJM1463 | XVI | 911058 | 911067 | 915577 | middle |  |  |  |  |  |  |  |  |  |  |  |
| YJM1463 | XVI | 911311 | 911320 | 915577 | middle |  |  |  |  |  |  |  |  |  |  |  |
| YJM1463 | XVI | 913472 | 913481 | 915577 | middle |  |  |  |  |  |  |  |  |  |  |  |
| YJM1463 | XVI | 915159 | 915168 | 915577 | middle |  |  |  |  |  |  |  |  |  |  |  |
| YJM1463 | XVI | 915577 | 915577 | 915577 | rightend_complete | YPR201W | 99.753 | 1215 | 3 | 0 | 1 | 1215 | 906996 | 908210 | 0 | 2228 |
| YJM1477 | I | 1 | 10 | 172259 | leftend | YAL063C | 88.408 | 1087 | 122 | 4 | 1 | 1085 | 1555 | 471 | 0 | 1306 |
| YJM1477 | I | 461 | 470 | 172259 | middle |  |  |  |  |  |  |  |  |  |  |  |
| YJM1477 | I | 172250 | 172259 | 172259 | rightend | YAR050W | 83.717 | 1087 | 173 | 4 | 1 | 1085 | 171165 | 172249 | 0 | 1024 |
| YJM1477 | II | 1 | 10 | 784115 | leftend | YNCB0001W | 100 | 84 | 0 | 0 | 1 | 84 | 1731 | 1814 | 4.16E-38 | 156 |
| YJM1477 | II | 784106 | 784115 | 784115 | rightend | YBR297W | 92.243 | 1418 | 94 | 12 | 1 | 1407 | 774136 | 775548 | 0 | 1995 |
| YJM1477 | III | 1 | 10 | 299780 | leftend | YCL073C | 99.892 | 1848 | 2 | 0 | 1 | 1848 | 7607 | 5760 | 0 | 3402 |
| YJM1477 | III | 299771 | 299780 | 299780 | rightend | YCR101C | 99.818 | 549 | 1 | 0 | 1 | 549 | 295672 | 295124 | 0 | 1009 |
| YJM1477 | IV | 1 | 10 | 1474380 | leftend | YDL243C | 95.051 | 990 | 49 | 0 | 1 | 990 | 17416 | 16427 | 0 | 1557 |
| YJM1477 | IV | 1474371 | 1474380 | 1474380 | rightend | YDR541C | 99.42 | 1035 | 6 | 0 | 1 | 1035 | 1464670 | 1463636 | 0 | 1879 |
| YJM1477 | V | 1 | 10 | 556928 | leftend | YEL070W | 98.873 | 1509 | 17 | 0 | 1 | 1509 | 13406 | 14914 | 0 | 2693 |
| YJM1477 | V | 554565 | 554574 | 556928 | middle |  |  |  |  |  |  |  |  |  |  |  |
| YJM1477 | V | 556928 | 556928 | 556928 | rightend_complete | YER187W | 91.745 | 424 | 33 | 2 | 4 | 426 | 548294 | 548716 | 1.6E-167 | 588 |
| YJM1477 | VI | 1 | 10 | 233731 | leftend | YFL051C | 98.451 | 452 | 7 | 0 | 1 | 452 | 1092 | 641 | 0 | 797 |
| YJM1477 | VI | 233731 | 233731 | 233731 | rightend_complete | YFR055W | 100 | 1023 | 0 | 0 | 1 | 1023 | 227760 | 228782 | 0 | 1890 |
| YJM1477 | VII | 1 |  | 1042257 | leftend_complete | YGL258W | 97.585 | 621 | 15 | 0 | 1 | 621 | 11216 | 11836 | 0 | 1064 |
| YJM1477 | VII | 1042248 | 1042257 | 1042257 | rightend | YGR287C | 99.139 | 1626 | 14 | 0 | 145 | 1770 | 1031152 | 1029527 | 0 | 2926 |
| YJM1477 | VIII | 1 | 10 | 545046 | leftend | YHL044W | 98.305 | 708 | 12 | 0 | 1 | 708 | 10301 | 11008 | 0 | 1242 |
| YJM1477 | VIII | 977 | 986 | 545046 | middle |  |  |  |  |  |  |  |  |  |  |  |
| YJM1477 | VIII | 518336 | 518345 | 545046 | middle |  |  |  |  |  |  |  |  |  |  |  |
| YJM1477 | VIII | 532749 | 532758 | 545046 | middle |  |  |  |  |  |  |  |  |  |  |  |
| YJM1477 | VIII | 536019 | 536028 | 545046 | middle |  |  |  |  |  |  |  |  |  |  |  |
| YJM1477 | VIII | 544224 | 544233 | 545046 | middle |  |  |  |  |  |  |  |  |  |  |  |
| YJM1477 | VIII | 545037 | 545046 | 545046 | rightend | YHR211W | 92.463 | 2136 | 137 | 2 | 1096 | 3228 | 516200 | 518314 | 0 | 3031 |
| YJM1477 | IX | 1 | 10 | 407403 | leftend | YIL169C | 93.45 | 1252 | 80 | 2 | 562 | 1812 | 6524 | 5274 | 0 | 1857 |
| YJM1477 | IX | 407394 | 407403 | 407403 | rightend | YIR039C | 99.009 | 1614 | 16 | 0 | 1 | 1614 | 405739 | 404126 | 0 | 2892 |
| YJM1477 | X | 1 | 10 | 714726 | leftend | YJL213W | 99.699 | 996 | 3 | 0 | 1 | 996 | 26315 | 27310 | 0 | 1823 |
| YJM1477 | X | 713025 | 713034 | 714726 | middle |  |  |  |  |  |  |  |  |  |  |  |
| YJM1477 | X | 714717 | 714726 | 714726 | rightend | YJR152W | 99.755 | 1632 | 3 | 1 | 1 | 1632 | 695337 | 696967 | 0 | 2990 |
| YJM1477 | XI | 1 | 10 | 663055 | leftend | YKL222C | 99.386 | 2118 | 13 | 0 | 1 | 2118 | 3991 | 1874 | 0 | 3840 |
| YJM1477 | XI | 726 | 735 | 663055 | middle |  |  |  |  |  |  |  |  |  |  |  |
| YJM1477 | XI | 662440 | 662449 | 663055 | middle |  |  |  |  |  |  |  |  |  |  |  |
| YJM1477 | XI | 663055 | 663055 | 663055 | rightend_complete | YKR101W | 99.847 | 1965 | 3 | 0 | 1 | 1965 | 637841 | 639805 | 0 | 3613 |
| YJM1477 | XII | 1 | 10 | 1626484 | leftend | YLL062C | 100 | 975 | 0 | 0 | 1 | 975 | 1968 | 994 | 0 | 1801 |
| YJM1477 | XII | 1626475 | 1626484 | 1626484 | rightend | YLR460C | 94.336 | 1130 | 62 | 2 | 3 | 1131 | 1624835 | 1623707 | 0 | 1731 |
| YJM1477 | XIII | 1 | 10 | 896420 | leftend | YML131W | 99.818 | 1098 | 2 | 0 | 1 | 1098 | 5258 | 6355 | 0 | 2017 |
| YJM1477 | XIII | 2218 | 2227 | 896420 | middle |  |  |  |  |  |  |  |  |  |  |  |
| YJM1477 | XIII | 2951 | 2960 | 896420 | middle |  |  |  |  |  |  |  |  |  |  |  |
| YJM1477 | XIII | 896411 | 896420 | 896420 | rightend | YMR321C | 99.042 | 313 | 3 | 0 | 6 | 318 | 890173 | 889861 | 1.15E-159 | 562 |
| YJM1477 | XIV | 1 | 10 | 760000 | leftend | YNL332W | 98.436 | 1023 | 16 | 0 | 1 | 1023 | 9169 | 10191 | 0 | 1801 |
| YJM1477 | XIV | 357 | 366 | 760000 | middle |  |  |  |  |  |  |  |  |  |  |  |
| YJM1477 | XIV | 1934 | 1943 | 760000 | middle |  |  |  |  |  |  |  |  |  |  |  |
| YJM1477 | XIV | 2799 | 2808 | 760000 | middle |  |  |  |  |  |  |  |  |  |  |  |
| YJM1477 | XIV | 3649 | 3658 | 760000 | middle |  |  |  |  |  |  |  |  |  |  |  |
| YJM1477 | XIV | 758926 | 758935 | 760000 | middle |  |  |  |  |  |  |  |  |  |  |  |
| YJM1477 | XIV | 759991 | 760000 | 760000 | rightend | YNR071C | 100 | 1029 | 0 | 0 | 1 | 1029 | 747556 | 746528 | 0 | 1901 |
| YJM1477 | XV | 1 |  | 1039353 | leftend_complete | YOL154W | 99.467 | 750 | 4 | 0 | 1 | 750 | 9437 | 10186 | 0 | 1363 |
| YJM1477 | XV | 1038045 | 1038054 | 1039353 | middle |  |  |  |  |  |  |  |  |  |  |  |
| YJM1477 | XV | 1039344 | 1039353 | 1039353 | rightend | YOR381W | 99.532 | 2136 | 2 | 1 | 1 | 2136 | 1006135 | 1008262 | 0 | 3882 |
| YJM1477 | XVI | 1 | 10 | 894038 | leftend | YPL274W | 99.887 | 1764 | 2 | 0 | 1 | 1764 | 4470 | 6233 | 0 | 3247 |
| YJM1477 | XVI | 1037 | 1046 | 894038 | middle |  |  |  |  |  |  |  |  |  |  |  |
| YJM1477 | XVI | 893573 | 893582 | 894038 | middle |  |  |  |  |  |  |  |  |  |  |  |
| YJM1477 | XVI | 894029 | 894038 | 894038 | rightend | YPR201W | 100 | 1215 | 0 | 0 | 1 | 1215 | 890562 | 891776 | 0 | 2244 |
| YJM1478 | I | 1 | 10 | 173954 | leftend | YAL063C | 88.445 | 952 | 106 | 4 | 1 | 950 | 960 | 11 | 0 | 1146 |
| YJM1478 | I | 173945 | 173954 | 173954 | rightend | YAR050W | 83.613 | 952 | 152 | 4 | 1 | 950 | 172995 | 173944 | 0 | 891 |
| YJM1478 | II | 1 | 10 | 788732 | leftend | YNCB0001W | 100 | 84 | 0 | 0 | 1 | 84 | 1309 | 1392 | 4.16E-38 | 156 |
| YJM1478 | II | 787189 | 787198 | 788732 | middle |  |  |  |  |  |  |  |  |  |  |  |
| YJM1478 | II | 788723 | 788732 | 788732 | rightend | YBR297W | 98.091 | 1205 | 23 | 0 | 1 | 1205 | 772221 | 773425 | 0 | 2098 |
| YJM1478 | III | 1 | 10 | 301275 | leftend | YCL073C | 99.839 | 1242 | 2 | 0 | 607 | 1848 | 4264 | 3023 | 0 | 2283 |
| YJM1478 | III | 300460 | 300469 | 301275 | middle |  |  |  |  |  |  |  |  |  |  |  |
| YJM1478 | III | 301275 | 301275 | 301275 | rightend_complete | YCR101C | 99.455 | 550 | 2 | 1 | 1 | 549 | 294210 | 293661 | 0 | 998 |
| YJM1478 | IV | 1 | 10 | 1463868 | leftend | YDL243C | 93.535 | 990 | 64 | 0 | 1 | 990 | 18645 | 17656 | 0 | 1474 |
| YJM1478 | IV | 1463859 | 1463868 | 1463868 | rightend | YDR541C | 99.517 | 1035 | 5 | 0 | 1 | 1035 | 1461095 | 1460061 | 0 | 1884 |
| YJM1478 | V | 1 | 10 | 560965 | leftend | YEL070W | 99.139 | 1509 | 13 | 0 | 1 | 1509 | 13289 | 14797 | 0 | 2715 |
| YJM1478 | V | 560041 | 560050 | 560965 | middle |  |  |  |  |  |  |  |  |  |  |  |
| YJM1478 | V | 560965 | 560965 | 560965 | rightend_complete | YER187W | 91.981 | 424 | 32 | 2 | 4 | 426 | 552926 | 553348 | 3.44E-169 | 593 |
| YJM1478 | VI | 1 |  | 247007 | leftend_complete | YFL051C | 98.451 | 452 | 7 | 0 | 1 | 452 | 11233 | 10782 | 0 | 797 |
| YJM1478 | VI | 246998 | 247007 | 247007 | rightend | YFR055W | 99.804 | 1022 | 2 | 0 | 1 | 1022 | 239785 | 240806 | 0 | 1877 |
| YJM1478 | VII | 1 |  | 1046949 | leftend_complete | YGL258W | 97.424 | 621 | 16 | 0 | 1 | 621 | 11715 | 12335 | 0 | 1059 |
| YJM1478 | VII | 581 | 590 | 1046949 | middle |  |  |  |  |  |  |  |  |  |  |  |
| YJM1478 | VII | 1046940 | 1046949 | 1046949 | rightend | YGR287C | 98.927 | 1770 | 19 | 0 | 1 | 1770 | 1031726 | 1029957 | 0 | 3164 |
| YJM1478 | VIII | 1 | 10 | 563344 | leftend | YHL044W | 98.446 | 708 | 11 | 0 | 1 | 708 | 1379 | 2086 | 0 | 1247 |
| YJM1478 | VIII | 563344 | 563344 | 563344 | rightend_complete | YHR211W | 82.535 | 939 | 150 | 10 | 1 | 932 | 510163 | 511094 | 0 | 813 |
| YJM1478 | IX | 1 | 10 | 437733 | leftend | YIL169C | 97.003 | 1568 | 47 | 0 | 352 | 1919 | 35629 | 34062 | 0 | 2636 |
| YJM1478 | IX | 437724 | 437733 | 437733 | rightend | YIR039C | 99.071 | 1614 | 15 | 0 | 1 | 1614 | 436068 | 434455 | 0 | 2898 |
| YJM1478 | X | 1 |  | 757957 | leftend_complete | YJL213W | 99.598 | 996 | 4 | 0 | 1 | 996 | 12849 | 13844 | 0 | 1818 |
| YJM1478 | X | 566 | 575 | 757957 | middle |  |  |  |  |  |  |  |  |  |  |  |
| YJM1478 | X | 757957 | 757957 | 757957 | rightend_complete | YJR152W | 99.939 | 1632 | 1 | 0 | 1 | 1632 | 682777 | 684408 | 0 | 3009 |
| YJM1478 | XI | 1 | 10 | 665584 | leftend | YKL222C | 99.433 | 2118 | 12 | 0 | 1 | 2118 | 4247 | 2130 | 0 | 3845 |
| YJM1478 | XI | 665575 | 665584 | 665584 | rightend | YKR101W | 99.796 | 1965 | 4 | 0 | 1 | 1965 | 656031 | 657995 | 0 | 3607 |
| YJM1478 | XII | 1 | 10 | 1959572 | leftend | YLL062C | 99.487 | 975 | 5 | 0 | 1 | 975 | 4029 | 3055 | 0 | 1773 |
| YJM1478 | XII | 1959563 | 1959572 | 1959572 | rightend | YLR460C | 94.071 | 1130 | 65 | 2 | 3 | 1131 | 1958500 | 1957372 | 0 | 1714 |
| YJM1478 | XIII | 1 | 10 | 914868 | leftend | YML131W | 99.545 | 1098 | 5 | 0 | 1 | 1098 | 13135 | 14232 | 0 | 2001 |
| YJM1478 | XIII | 914859 | 914868 | 914868 | rightend | YMR321C | 99.361 | 313 | 2 | 0 | 6 | 318 | 897332 | 897020 | 2.48E-161 | 568 |
| YJM1478 | XIV | 1 |  | 736159 | leftend_complete | YNL332W | 99.022 | 1023 | 10 | 0 | 1 | 1023 | 9873 | 10895 | 0 | 1834 |
| YJM1478 | XIV | 1657 | 1666 | 736159 | middle |  |  |  |  |  |  |  |  |  |  |  |
| YJM1478 | XIV | 734966 | 734975 | 736159 | middle |  |  |  |  |  |  |  |  |  |  |  |
| YJM1478 | XIV | 736150 | 736159 | 736159 | rightend | YNR065C | 95.172 | 3335 | 124 | 23 | 19 | 3351 | 731551 | 728252 | 0 | 5232 |
| YJM1478 | XV | 1 |  | 1041348 | leftend_complete | YOL154W | 99.867 | 750 | 1 | 0 | 1 | 750 | 24877 | 25626 | 0 | 1380 |
| YJM1478 | XV | 943 | 952 | 1041348 | middle |  |  |  |  |  |  |  |  |  |  |  |
| YJM1478 | XV | 1473 | 1482 | 1041348 | middle |  |  |  |  |  |  |  |  |  |  |  |
| YJM1478 | XV | 1041339 | 1041348 | 1041348 | rightend | YOR381W | 99.906 | 2136 | 2 | 0 | 1 | 2136 | 1022793 | 1024928 | 0 | 3934 |
| YJM1478 | XVI | 1 | 10 | 914824 | leftend | YPL274W | 99.66 | 1764 | 6 | 0 | 1 | 1764 | 24743 | 26506 | 0 | 3225 |
| YJM1478 | XVI | 359 | 368 | 914824 | middle |  |  |  |  |  |  |  |  |  |  |  |
| YJM1478 | XVI | 1075 | 1084 | 914824 | middle |  |  |  |  |  |  |  |  |  |  |  |
| YJM1478 | XVI | 914815 | 914824 | 914824 | rightend | YPR201W | 99.588 | 1215 | 5 | 0 | 1 | 1215 | 911569 | 912783 | 0 | 2217 |
| YJM1479 | I | 1 | 10 | 186938 | leftend | YAL063C | 81.929 | 2053 | 335 | 30 | 1 | 2035 | 3833 | 1799 | 0 | 1703 |
| YJM1479 | I | 879 | 888 | 186938 | middle |  |  |  |  |  |  |  |  |  |  |  |
| YJM1479 | I | 186929 | 186938 | 186938 | rightend | YAR050W | 86.084 | 1660 | 203 | 15 | 2037 | 3676 | 3012 | 1361 | 0 | 1760 |
| YJM1479 | II | 1 |  | 781718 | leftend_complete | YNCB0001W | 100 | 84 | 0 | 0 | 1 | 84 | 3487 | 3570 | 4.16E-38 | 156 |
| YJM1479 | II | 1957 | 1966 | 781718 | middle |  |  |  |  |  |  |  |  |  |  |  |
| YJM1479 | II | 781709 | 781718 | 781718 | rightend | YBR297W | 97.842 | 1205 | 26 | 0 | 1 | 1205 | 777518 | 778722 | 0 | 2082 |
| YJM1479 | III | 1 | 10 | 310875 | leftend | YCL073C | 98.972 | 1848 | 19 | 0 | 1 | 1848 | 8016 | 6169 | 0 | 3308 |
| YJM1479 | III | 302025 | 302034 | 310875 | middle |  |  |  |  |  |  |  |  |  |  |  |
| YJM1479 | III | 310875 | 310875 | 310875 | rightend_complete | YCR101C | 98.727 | 550 | 6 | 1 | 1 | 549 | 297616 | 297067 | 0 | 976 |
| YJM1479 | IV | 1 | 10 | 1456051 | leftend | YDL243C | 92.072 | 883 | 68 | 2 | 1 | 882 | 892 | 11 | 0 | 1242 |
| YJM1479 | IV | 1453291 | 1453300 | 1456051 | middle |  |  |  |  |  |  |  |  |  |  |  |
| YJM1479 | IV | 1454697 | 1454706 | 1456051 | middle |  |  |  |  |  |  |  |  |  |  |  |
| YJM1479 | IV | 1456051 | 1456051 | 1456051 | rightend_complete | YDR541C | 99.807 | 1035 | 2 | 0 | 1 | 1035 | 1448307 | 1447273 | 0 | 1901 |
| YJM1479 | V | 1 | 10 | 547335 | leftend | YEL070W | 99.27 | 1507 | 11 | 0 | 3 | 1509 | 10508 | 12014 | 0 | 2723 |
| YJM1479 | V | 1959 | 1968 | 547335 | middle |  |  |  |  |  |  |  |  |  |  |  |
| YJM1479 | V | 547326 | 547335 | 547335 | rightend | YER187W | 98.122 | 426 | 8 | 0 | 1 | 426 | 544000 | 544425 | 0 | 743 |
| YJM1479 | VI | 1 | 10 | 284110 | leftend | YFL051C | 98.23 | 452 | 8 | 0 | 1 | 452 | 39792 | 39341 | 0 | 791 |
| YJM1479 | VI | 284101 | 284110 | 284110 | rightend | YFR055W | 99.609 | 1022 | 4 | 0 | 1 | 1022 | 268390 | 269411 | 0 | 1866 |
| YJM1479 | VII | 1 | 10 | 1048946 | leftend | YGL258W | 93.72 | 621 | 39 | 0 | 1 | 621 | 10692 | 11312 | 0 | 931 |
| YJM1479 | VII | 1043606 | 1043615 | 1048946 | middle |  |  |  |  |  |  |  |  |  |  |  |
| YJM1479 | VII | 1045248 | 1045257 | 1048946 | middle |  |  |  |  |  |  |  |  |  |  |  |
| YJM1479 | VII | 1047855 | 1047864 | 1048946 | middle |  |  |  |  |  |  |  |  |  |  |  |
| YJM1479 | VII | 1048937 | 1048946 | 1048946 | rightend | YGR287C | 99.887 | 1770 | 2 | 0 | 1 | 1770 | 1033701 | 1031932 | 0 | 3258 |
| YJM1479 | VIII | 1 | 10 | 546462 | leftend | YHL044W | 99.011 | 708 | 7 | 0 | 1 | 708 | 7983 | 8690 | 0 | 1269 |
| YJM1479 | VIII | 516234 | 516243 | 546462 | middle |  |  |  |  |  |  |  |  |  |  |  |
| YJM1479 | VIII | 532262 | 532271 | 546462 | middle |  |  |  |  |  |  |  |  |  |  |  |
| YJM1479 | VIII | 532910 | 532919 | 546462 | middle |  |  |  |  |  |  |  |  |  |  |  |
| YJM1479 | VIII | 546453 | 546462 | 546462 | rightend | YHR211W | 84.587 | 1369 | 200 | 7 | 1 | 1362 | 514863 | 516227 | 0 | 1349 |
| YJM1479 | IX | 1 |  | 418850 | leftend_complete | YIL169C | 96.479 | 1278 | 36 | 5 | 612 | 1880 | 10312 | 9035 | 0 | 2102 |
| YJM1479 | IX | 1076 | 1085 | 418850 | middle |  |  |  |  |  |  |  |  |  |  |  |
| YJM1479 | IX | 2340 | 2349 | 418850 | middle |  |  |  |  |  |  |  |  |  |  |  |
| YJM1479 | IX | 3435 | 3444 | 418850 | middle |  |  |  |  |  |  |  |  |  |  |  |
| YJM1479 | IX | 418841 | 418850 | 418850 | rightend | YIR039C | 99.69 | 1614 | 5 | 0 | 1 | 1614 | 412117 | 410504 | 0 | 2953 |
| YJM1479 | X | 1 |  | 716917 | leftend_complete | YJL213W | 98.896 | 996 | 11 | 0 | 1 | 996 | 23817 | 24812 | 0 | 1779 |
| YJM1479 | X | 622 | 631 | 716917 | middle |  |  |  |  |  |  |  |  |  |  |  |
| YJM1479 | X | 1598 | 1607 | 716917 | middle |  |  |  |  |  |  |  |  |  |  |  |
| YJM1479 | X | 8341 | 8350 | 716917 | middle |  |  |  |  |  |  |  |  |  |  |  |
| YJM1479 | X | 715909 | 715918 | 716917 | middle |  |  |  |  |  |  |  |  |  |  |  |
| YJM1479 | X | 716908 | 716917 | 716917 | rightend | YJR152W | 99.571 | 1632 | 7 | 0 | 1 | 1632 | 695967 | 697598 | 0 | 2976 |
| YJM1479 | XI | 1 | 10 | 663666 | leftend | YKL222C | 99.669 | 2118 | 7 | 0 | 1 | 2118 | 3529 | 1412 | 0 | 3873 |
| YJM1479 | XI | 663657 | 663666 | 663666 | rightend | YKR101W | 99.389 | 1965 | 12 | 0 | 1 | 1965 | 638570 | 640534 | 0 | 3563 |
| YJM1479 | XII | 1 | 10 | 1724776 | leftend | YLL062C | 98.872 | 975 | 11 | 0 | 1 | 975 | 4352 | 3378 | 0 | 1740 |
| YJM1479 | XII | 1723534 | 1723543 | 1724776 | middle |  |  |  |  |  |  |  |  |  |  |  |
| YJM1479 | XII | 1723993 | 1724002 | 1724776 | middle |  |  |  |  |  |  |  |  |  |  |  |
| YJM1479 | XII | 1724767 | 1724776 | 1724776 | rightend | YLR460C | 98.851 | 1131 | 13 | 0 | 1 | 1131 | 1722540 | 1721410 | 0 | 2017 |
| YJM1479 | XIII | 1 | 10 | 905700 | leftend | YML131W | 99.454 | 1098 | 6 | 0 | 1 | 1098 | 3623 | 4720 | 0 | 1995 |
| YJM1479 | XIII | 899064 | 899073 | 905700 | middle |  |  |  |  |  |  |  |  |  |  |  |
| YJM1479 | XIII | 904824 | 904833 | 905700 | middle |  |  |  |  |  |  |  |  |  |  |  |
| YJM1479 | XIII | 905691 | 905700 | 905700 | rightend | YMR321C | 99.361 | 313 | 2 | 0 | 6 | 318 | 890986 | 890674 | 2.48E-161 | 568 |
| YJM1479 | XIV | 1 | 10 | 756640 | leftend | YNL332W | 99.316 | 1023 | 7 | 0 | 1 | 1023 | 6612 | 7634 | 0 | 1851 |
| YJM1479 | XIV | 1154 | 1163 | 756640 | middle |  |  |  |  |  |  |  |  |  |  |  |
| YJM1479 | XIV | 2753 | 2762 | 756640 | middle |  |  |  |  |  |  |  |  |  |  |  |
| YJM1479 | XIV | 756631 | 756640 | 756640 | rightend | YNR071C | 99.514 | 1029 | 5 | 0 | 1 | 1029 | 745969 | 744941 | 0 | 1873 |
| YJM1479 | XV | 1 | 10 | 1046087 | leftend | YOL154W | 99.467 | 750 | 4 | 0 | 1 | 750 | 17863 | 18612 | 0 | 1363 |
| YJM1479 | XV | 1114 | 1123 | 1046087 | middle |  |  |  |  |  |  |  |  |  |  |  |
| YJM1479 | XV | 13368 | 13377 | 1046087 | middle |  |  |  |  |  |  |  |  |  |  |  |
| YJM1479 | XV | 1046078 | 1046087 | 1046087 | rightend | YOR381W | 98.689 | 2136 | 28 | 0 | 1 | 2136 | 1018874 | 1021009 | 0 | 3790 |
| YJM1479 | XVI | 1 | 10 | 917031 | leftend | YPL274W | 98.186 | 1764 | 32 | 0 | 1 | 1764 | 22087 | 23850 | 0 | 3081 |
| YJM1479 | XVI | 917022 | 917031 | 917031 | rightend | YPR198W | 99.203 | 1632 | 13 | 0 | 1 | 1632 | 908516 | 910147 | 0 | 2942 |
| YJM1526 | I | 1 | 10 | 184187 | leftend | YAL063C | 95.02 | 1004 | 23 | 11 | 2966 | 3969 | 171835 | 172811 | 0 | 1552 |
| YJM1526 | I | 184178 | 184187 | 184187 | rightend | YAR050W | 81.865 | 1362 | 243 | 4 | 1 | 1360 | 169944 | 171303 | 0 | 1144 |
| YJM1526 | II | 1 | 10 | 788619 | leftend | YNCB0001W | 100 | 84 | 0 | 0 | 1 | 84 | 6545 | 6628 | 4.16E-38 | 156 |
| YJM1526 | II | 2389 | 2398 | 788619 | middle |  |  |  |  |  |  |  |  |  |  |  |
| YJM1526 | II | 3403 | 3412 | 788619 | middle |  |  |  |  |  |  |  |  |  |  |  |
| YJM1526 | II | 787956 | 787965 | 788619 | middle |  |  |  |  |  |  |  |  |  |  |  |
| YJM1526 | II | 788610 | 788619 | 788619 | rightend | YBR297W | 99.716 | 1407 | 4 | 0 | 1 | 1407 | 779267 | 780673 | 0 | 2577 |
| YJM1526 | III | 1 | 10 | 308393 | leftend | YCL073C | 99.729 | 1848 | 5 | 0 | 1 | 1848 | 7329 | 5482 | 0 | 3386 |
| YJM1526 | III | 300085 | 300094 | 308393 | middle |  |  |  |  |  |  |  |  |  |  |  |
| YJM1526 | III | 308384 | 308393 | 308393 | rightend | YCR101C | 99.818 | 549 | 1 | 0 | 1 | 549 | 295887 | 295339 | 0 | 1009 |
| YJM1526 | IV | 1 | 10 | 1475681 | leftend | YDL243C | 95.152 | 990 | 48 | 0 | 1 | 990 | 17430 | 16441 | 0 | 1563 |
| YJM1526 | IV | 511 | 520 | 1475681 | middle |  |  |  |  |  |  |  |  |  |  |  |
| YJM1526 | IV | 1471353 | 1471362 | 1475681 | middle |  |  |  |  |  |  |  |  |  |  |  |
| YJM1526 | IV | 1475672 | 1475681 | 1475681 | rightend | YDR541C | 99.42 | 1035 | 6 | 0 | 1 | 1035 | 1468503 | 1467469 | 0 | 1879 |
| YJM1526 | V | 1 | 10 | 571437 | leftend | YEL070W | 98.873 | 1509 | 17 | 0 | 1 | 1509 | 30673 | 32181 | 0 | 2693 |
| YJM1526 | V | 571428 | 571437 | 571437 | rightend | YER187W | 91.981 | 424 | 32 | 2 | 4 | 426 | 565157 | 565579 | 3.44E-169 | 593 |
| YJM1526 | VI | 1 | 10 | 234157 | leftend | YFL051C | 98.23 | 452 | 8 | 0 | 1 | 452 | 1227 | 776 | 0 | 791 |
| YJM1526 | VI | 234148 | 234157 | 234157 | rightend | YFR055W | 99.902 | 1023 | 1 | 0 | 1 | 1023 | 227908 | 228930 | 0 | 1884 |
| YJM1526 | VII | 1 | 10 | 1042876 | leftend | YGL258W | 100 | 621 | 0 | 0 | 1 | 621 | 4072 | 4692 | 0 | 1147 |
| YJM1526 | VII | 1042867 | 1042876 | 1042876 | rightend | YGR287C | 98.588 | 1770 | 25 | 0 | 1 | 1770 | 1025507 | 1023738 | 0 | 3131 |
| YJM1526 | VIII | 1 |  | 887513 | leftend_complete | YPL274W | 99.83 | 1764 | 3 | 0 | 1 | 1764 | 20658 | 22421 | 0 | 3241 |
| YJM1526 | VIII | 887033 | 887042 | 887513 | middle |  |  |  |  |  |  |  |  |  |  |  |
| YJM1526 | VIII | 887513 | 887513 | 887513 | rightend_complete | YHR211W | 92.322 | 2136 | 140 | 2 | 1096 | 3228 | 874262 | 876376 | 0 | 3014 |
| YJM1526 | IX | 1 | 10 | 415927 | leftend | YIL169C | 91.549 | 1420 | 118 | 2 | 394 | 1812 | 8504 | 7086 | 0 | 1956 |
| YJM1526 | IX | 415918 | 415927 | 415927 | rightend | YIR039C | 99.009 | 1614 | 16 | 0 | 1 | 1614 | 413857 | 412244 | 0 | 2892 |
| YJM1526 | X | 1 |  | 723867 | leftend_complete | YJL213W | 99.699 | 996 | 3 | 0 | 1 | 996 | 34393 | 35388 | 0 | 1823 |
| YJM1526 | X | 2302 | 2311 | 723867 | middle |  |  |  |  |  |  |  |  |  |  |  |
| YJM1526 | X | 2522 | 2531 | 723867 | middle |  |  |  |  |  |  |  |  |  |  |  |
| YJM1526 | X | 723867 | 723867 | 723867 | rightend_complete | YJR152W | 99.755 | 1632 | 3 | 1 | 1 | 1632 | 703454 | 705084 | 0 | 2990 |
| YJM1526 | XI | 1 | 10 | 659863 | leftend | YKL222C | 99.433 | 2118 | 12 | 0 | 1 | 2118 | 3203 | 1086 | 0 | 3845 |
| YJM1526 | XI | 659854 | 659863 | 659863 | rightend | YKR101W | 99.847 | 1965 | 3 | 0 | 1 | 1965 | 636788 | 638752 | 0 | 3613 |
| YJM1526 | XII | 1 |  | 1891382 | leftend_complete | YLL062C | 99.795 | 975 | 2 | 0 | 1 | 975 | 2483 | 1509 | 0 | 1790 |
| YJM1526 | XII | 202 | 211 | 1891382 | middle |  |  |  |  |  |  |  |  |  |  |  |
| YJM1526 | XII | 1891373 | 1891382 | 1891382 | rightend | YLR460C | 94.336 | 1130 | 62 | 2 | 3 | 1131 | 1889632 | 1888504 | 0 | 1731 |
| YJM1526 | XIII | 1 | 10 | 894526 | leftend | YML131W | 100 | 1098 | 0 | 0 | 1 | 1098 | 4154 | 5251 | 0 | 2028 |
| YJM1526 | XIII | 1847 | 1856 | 894526 | middle |  |  |  |  |  |  |  |  |  |  |  |
| YJM1526 | XIII | 894517 | 894526 | 894526 | rightend | YMR321C | 99.361 | 313 | 2 | 0 | 6 | 318 | 886333 | 886021 | 2.48E-161 | 568 |
| YJM1526 | XIV | 1 | 10 | 754375 | leftend | YNL332W | 98.631 | 1023 | 14 | 0 | 1 | 1023 | 4140 | 5162 | 0 | 1812 |
| YJM1526 | XIV | 799 | 808 | 754375 | middle |  |  |  |  |  |  |  |  |  |  |  |
| YJM1526 | XIV | 1326 | 1335 | 754375 | middle |  |  |  |  |  |  |  |  |  |  |  |
| YJM1526 | XIV | 2632 | 2641 | 754375 | middle |  |  |  |  |  |  |  |  |  |  |  |
| YJM1526 | XIV | 4066 | 4075 | 754375 | middle |  |  |  |  |  |  |  |  |  |  |  |
| YJM1526 | XIV | 752934 | 752943 | 754375 | middle |  |  |  |  |  |  |  |  |  |  |  |
| YJM1526 | XIV | 754375 | 754375 | 754375 | rightend_complete | YNR071C | 99.903 | 1029 | 1 | 0 | 1 | 1029 | 742227 | 741199 | 0 | 1895 |
| YJM1526 | XV | 1 | 10 | 1054164 | leftend | YOL154W | 99.6 | 750 | 3 | 0 | 1 | 750 | 23202 | 23951 | 0 | 1369 |
| YJM1526 | XV | 1054155 | 1054164 | 1054164 | rightend | YOR381W | 100 | 2136 | 0 | 0 | 1 | 2136 | 1021274 | 1023409 | 0 | 3945 |
| YJM1526 | XVI | 1 |  | 560369 | leftend_complete | YHL044W | 98.446 | 708 | 11 | 0 | 1 | 708 | 9004 | 9711 | 0 | 1247 |
| YJM1526 | XVI | 560360 | 560369 | 560369 | rightend | YPR201W | 100 | 1215 | 0 | 0 | 1 | 1215 | 557644 | 558858 | 0 | 2244 |
| YJM1527 | I | 1 | 10 | 198615 | leftend | YAL063C | 94.295 | 1262 | 46 | 7 | 2709 | 3969 | 194265 | 195501 | 0 | 1908 |
| YJM1527 | I | 221 | 230 | 198615 | middle |  |  |  |  |  |  |  |  |  |  |  |
| YJM1527 | I | 2856 | 2865 | 198615 | middle |  |  |  |  |  |  |  |  |  |  |  |
| YJM1527 | I | 13999 | 14008 | 198615 | middle |  |  |  |  |  |  |  |  |  |  |  |
| YJM1527 | I | 15356 | 15365 | 198615 | middle |  |  |  |  |  |  |  |  |  |  |  |
| YJM1527 | I | 198606 | 198615 | 198615 | rightend | YAR050W | 93.37 | 1448 | 84 | 8 | 2844 | 4282 | 193752 | 195196 | 0 | 2132 |
| YJM1527 | II | 1 |  | 780475 | leftend_complete | YNCB0001W | 100 | 84 | 0 | 0 | 1 | 84 | 2650 | 2733 | 4.16E-38 | 156 |
| YJM1527 | II | 1185 | 1194 | 780475 | middle |  |  |  |  |  |  |  |  |  |  |  |
| YJM1527 | II | 780466 | 780475 | 780475 | rightend | YBR297W | 98.257 | 1205 | 21 | 0 | 1 | 1205 | 775453 | 776657 | 0 | 2109 |
| YJM1527 | III | 1 |  | 313597 | leftend_complete | YCL073C | 100 | 1848 | 0 | 0 | 1 | 1848 | 10642 | 8795 | 0 | 3413 |
| YJM1527 | III | 2214 | 2223 | 313597 | middle |  |  |  |  |  |  |  |  |  |  |  |
| YJM1527 | III | 313597 | 313597 | 313597 | rightend_complete | YCR101C | 99.818 | 549 | 1 | 0 | 1 | 549 | 300003 | 299455 | 0 | 1009 |
| YJM1527 | IV | 1 |  | 1467869 | leftend_complete | YDL243C | 95.152 | 990 | 48 | 0 | 1 | 990 | 18450 | 17461 | 0 | 1563 |
| YJM1527 | IV | 1467860 | 1467869 | 1467869 | rightend | YDR541C | 99.42 | 1035 | 6 | 0 | 1 | 1035 | 1464284 | 1463250 | 0 | 1879 |
| YJM1527 | V | 1 | 10 | 552400 | leftend | YEL070W | 99.735 | 1509 | 4 | 0 | 1 | 1509 | 14621 | 16129 | 0 | 2765 |
| YJM1527 | V | 1042 | 1051 | 552400 | middle |  |  |  |  |  |  |  |  |  |  |  |
| YJM1527 | V | 552391 | 552400 | 552400 | rightend | YER187W | 91.981 | 424 | 32 | 2 | 4 | 426 | 548093 | 548515 | 3.44E-169 | 593 |
| YJM1527 | VI | 1 | 10 | 240669 | leftend | YFL051C | 98.23 | 452 | 8 | 0 | 1 | 452 | 850 | 399 | 0 | 791 |
| YJM1527 | VI | 240669 | 240669 | 240669 | rightend_complete | YFR055W | 99.511 | 1022 | 5 | 0 | 1 | 1022 | 229126 | 230147 | 0 | 1860 |
| YJM1527 | VII | 1 |  | 1065795 | leftend_complete | YGL258W | 97.585 | 621 | 15 | 0 | 1 | 621 | 11194 | 11814 | 0 | 1064 |
| YJM1527 | VII | 1065219 | 1065228 | 1065795 | middle |  |  |  |  |  |  |  |  |  |  |  |
| YJM1527 | VII | 1065786 | 1065795 | 1065795 | rightend | YGR287C | 99.831 | 1770 | 3 | 0 | 1 | 1770 | 1030799 | 1029030 | 0 | 3253 |
| YJM1527 | VIII | 1 | 10 | 543777 | leftend | YHL044W | 98.305 | 708 | 12 | 0 | 1 | 708 | 7317 | 8024 | 0 | 1242 |
| YJM1527 | VIII | 533786 | 533795 | 543777 | middle |  |  |  |  |  |  |  |  |  |  |  |
| YJM1527 | VIII | 534141 | 534150 | 543777 | middle |  |  |  |  |  |  |  |  |  |  |  |
| YJM1527 | VIII | 535712 | 535721 | 543777 | middle |  |  |  |  |  |  |  |  |  |  |  |
| YJM1527 | VIII | 542072 | 542081 | 543777 | middle |  |  |  |  |  |  |  |  |  |  |  |
| YJM1527 | VIII | 543768 | 543777 | 543777 | rightend | YHR211W | 89.935 | 1838 | 143 | 15 | 1096 | 2918 | 517936 | 519746 | 0 | 2331 |
| YJM1527 | IX | 1 |  | 425882 | leftend_complete | YIL169C | 84.633 | 2128 | 278 | 31 | 520 | 2618 | 18052 | 15945 | 0 | 2073 |
| YJM1527 | IX | 425085 | 425094 | 425882 | middle |  |  |  |  |  |  |  |  |  |  |  |
| YJM1527 | IX | 425882 | 425882 | 425882 | rightend_complete | YIR039C | 98.947 | 1614 | 17 | 0 | 1 | 1614 | 418761 | 417148 | 0 | 2887 |
| YJM1527 | X | 1 |  | 708161 | leftend_complete | YJL213W | 99.699 | 996 | 3 | 0 | 1 | 996 | 27111 | 28106 | 0 | 1823 |
| YJM1527 | X | 1509 | 1518 | 708161 | middle |  |  |  |  |  |  |  |  |  |  |  |
| YJM1527 | X | 706806 | 706815 | 708161 | middle |  |  |  |  |  |  |  |  |  |  |  |
| YJM1527 | X | 708161 | 708161 | 708161 | rightend_complete | YJR152W | 99.755 | 1632 | 3 | 1 | 1 | 1632 | 696632 | 698262 | 0 | 2990 |
| YJM1527 | XI | 1 | 10 | 667549 | leftend | YKL222C | 99.953 | 2118 | 1 | 0 | 1 | 2118 | 8777 | 6660 | 0 | 3906 |
| YJM1527 | XI | 667540 | 667549 | 667549 | rightend | YKR101W | 99.847 | 1965 | 3 | 0 | 1 | 1965 | 642935 | 644899 | 0 | 3613 |
| YJM1527 | XII | 1 |  | 2075171 | leftend_complete | YLL062C | 99.692 | 975 | 3 | 0 | 1 | 975 | 12261 | 11287 | 0 | 1784 |
| YJM1527 | XII | 574 | 583 | 2075171 | middle |  |  |  |  |  |  |  |  |  |  |  |
| YJM1527 | XII | 2075171 | 2075171 | 2075171 | rightend_complete | YLR460C | 98.585 | 1131 | 16 | 0 | 1 | 1131 | 2070413 | 2069283 | 0 | 2001 |
| YJM1527 | XIII | 1 | 10 | 897157 | leftend | YML131W | 99.545 | 1098 | 5 | 0 | 1 | 1098 | 3741 | 4838 | 0 | 2001 |
| YJM1527 | XIII | 936 | 945 | 897157 | middle |  |  |  |  |  |  |  |  |  |  |  |
| YJM1527 | XIII | 897148 | 897157 | 897157 | rightend | YMR321C | 99.361 | 313 | 2 | 0 | 6 | 318 | 888927 | 888615 | 2.48E-161 | 568 |
| YJM1527 | XIV | 1 |  | 764978 | leftend_complete | YNL332W | 98.338 | 1023 | 17 | 0 | 1 | 1023 | 6629 | 7651 | 0 | 1796 |
| YJM1527 | XIV | 997 | 1006 | 764978 | middle |  |  |  |  |  |  |  |  |  |  |  |
| YJM1527 | XIV | 1211 | 1220 | 764978 | middle |  |  |  |  |  |  |  |  |  |  |  |
| YJM1527 | XIV | 764969 | 764978 | 764978 | rightend | YNR071C | 99.514 | 1029 | 5 | 0 | 1 | 1029 | 744931 | 743903 | 0 | 1873 |
| YJM1527 | XV | 1 |  | 1034307 | leftend_complete | YOL154W | 99.467 | 750 | 4 | 0 | 1 | 750 | 9391 | 10140 | 0 | 1363 |
| YJM1527 | XV | 1033459 | 1033468 | 1034307 | middle |  |  |  |  |  |  |  |  |  |  |  |
| YJM1527 | XV | 1034298 | 1034307 | 1034307 | rightend | YOR381W | 99.953 | 2136 | 1 | 0 | 1 | 2136 | 1005629 | 1007764 | 0 | 3940 |
| YJM1527 | XVI | 1 |  | 908792 | leftend_complete | YPL274W | 99.943 | 1764 | 1 | 0 | 1 | 1764 | 16614 | 18377 | 0 | 3253 |
| YJM1527 | XVI | 905886 | 905895 | 908792 | middle |  |  |  |  |  |  |  |  |  |  |  |
| YJM1527 | XVI | 907271 | 907280 | 908792 | middle |  |  |  |  |  |  |  |  |  |  |  |
| YJM1527 | XVI | 908783 | 908792 | 908792 | rightend | YPR201W | 99.506 | 1215 | 6 | 0 | 1 | 1215 | 903152 | 904366 | 0 | 2211 |
| YJM1549 | I | 1 | 10 | 205180 | leftend | YAL063C | 91.984 | 998 | 51 | 13 | 2966 | 3962 | 16476 | 15507 | 0 | 1373 |
| YJM1549 | I | 17522 | 17531 | 205180 | middle |  |  |  |  |  |  |  |  |  |  |  |
| YJM1549 | I | 192765 | 192774 | 205180 | middle |  |  |  |  |  |  |  |  |  |  |  |
| YJM1549 | I | 196337 | 196346 | 205180 | middle |  |  |  |  |  |  |  |  |  |  |  |
| YJM1549 | I | 205171 | 205180 | 205180 | rightend | YAR050W | 86.85 | 1019 | 117 | 15 | 2665 | 3676 | 17509 | 16501 | 0 | 1123 |
| YJM1549 | II | 1 |  | 803802 | leftend_complete | YNCB0001W | 100 | 84 | 0 | 0 | 1 | 84 | 4897 | 4980 | 4.16E-38 | 156 |
| YJM1549 | II | 1040 | 1049 | 803802 | middle |  |  |  |  |  |  |  |  |  |  |  |
| YJM1549 | II | 1841 | 1850 | 803802 | middle |  |  |  |  |  |  |  |  |  |  |  |
| YJM1549 | II | 3533 | 3542 | 803802 | middle |  |  |  |  |  |  |  |  |  |  |  |
| YJM1549 | II | 802278 | 802287 | 803802 | middle |  |  |  |  |  |  |  |  |  |  |  |
| YJM1549 | II | 803793 | 803802 | 803802 | rightend | YBR297W | 99.076 | 1407 | 13 | 0 | 1 | 1407 | 793301 | 794707 | 0 | 2527 |
| YJM1549 | III | 1 |  | 310510 | leftend_complete | YCL073C | 99.567 | 1848 | 8 | 0 | 1 | 1848 | 8001 | 6154 | 0 | 3369 |
| YJM1549 | III | 301678 | 301687 | 310510 | middle |  |  |  |  |  |  |  |  |  |  |  |
| YJM1549 | III | 310510 | 310510 | 310510 | rightend_complete | YCR101C | 98.364 | 550 | 8 | 1 | 1 | 549 | 296753 | 296204 | 0 | 965 |
| YJM1549 | IV | 1 |  | 1466260 | leftend_complete | YDL243C | 93.769 | 979 | 61 | 0 | 1 | 979 | 9665 | 8687 | 0 | 1471 |
| YJM1549 | IV | 1390 | 1399 | 1466260 | middle |  |  |  |  |  |  |  |  |  |  |  |
| YJM1549 | IV | 1466251 | 1466260 | 1466260 | rightend | YDR541C | 99.614 | 1035 | 4 | 0 | 1 | 1035 | 1462179 | 1461145 | 0 | 1890 |
| YJM1549 | V | 1 |  | 558122 | leftend_complete | YEL070W | 99.072 | 1509 | 14 | 0 | 1 | 1509 | 20021 | 21529 | 0 | 2710 |
| YJM1549 | V | 558113 | 558122 | 558122 | rightend | YER187W | 91.981 | 424 | 32 | 2 | 4 | 426 | 555411 | 555833 | 3.44E-169 | 593 |
| YJM1549 | VI | 1 | 10 | 236413 | leftend | YFL051C | 98.451 | 452 | 7 | 0 | 1 | 452 | 2263 | 1812 | 0 | 797 |
| YJM1549 | VI | 236413 | 236413 | 236413 | rightend_complete | YFR055W | 100 | 1023 | 0 | 0 | 1 | 1023 | 230457 | 231479 | 0 | 1890 |
| YJM1549 | VII | 1 | 10 | 1035220 | leftend | YGL258W | 97.585 | 621 | 15 | 0 | 1 | 621 | 4562 | 5182 | 0 | 1064 |
| YJM1549 | VII | 1035211 | 1035220 | 1035220 | rightend | YGR287C | 99.379 | 1770 | 11 | 0 | 1 | 1770 | 1025310 | 1023541 | 0 | 3208 |
| YJM1549 | VIII | 1 | 10 | 889787 | leftend | YPL274W | 98.356 | 1764 | 29 | 0 | 1 | 1764 | 6472 | 8235 | 0 | 3097 |
| YJM1549 | VIII | 525 | 534 | 889787 | middle |  |  |  |  |  |  |  |  |  |  |  |
| YJM1549 | VIII | 889177 | 889186 | 889787 | middle |  |  |  |  |  |  |  |  |  |  |  |
| YJM1549 | VIII | 889787 | 889787 | 889787 | rightend_complete | YHR211W | 86.096 | 1496 | 197 | 7 | 1 | 1489 | 877031 | 878522 | 0 | 1600 |
| YJM1549 | IX | 1 |  | 407704 | leftend_complete | YIL169C | 92.964 | 1336 | 92 | 2 | 478 | 1812 | 2534 | 1200 | 0 | 1945 |
| YJM1549 | IX | 567 | 576 | 407704 | middle |  |  |  |  |  |  |  |  |  |  |  |
| YJM1549 | IX | 407695 | 407704 | 407704 | rightend | YIR039C | 99.009 | 1614 | 16 | 0 | 1 | 1614 | 406046 | 404433 | 0 | 2892 |
| YJM1549 | X | 1 | 10 | 712875 | leftend | YJL213W | 99.398 | 996 | 6 | 0 | 1 | 996 | 25994 | 26989 | 0 | 1807 |
| YJM1549 | X | 711126 | 711135 | 712875 | middle |  |  |  |  |  |  |  |  |  |  |  |
| YJM1549 | X | 712875 | 712875 | 712875 | rightend_complete | YJR152W | 99.755 | 1632 | 4 | 0 | 1 | 1632 | 696101 | 697732 | 0 | 2992 |
| YJM1549 | XI | 1 |  | 684120 | leftend_complete | YKL222C | 99.953 | 2118 | 1 | 0 | 1 | 2118 | 26799 | 24682 | 0 | 3906 |
| YJM1549 | XI | 684120 | 684120 | 684120 | rightend_complete | YKR101W | 99.847 | 1965 | 3 | 0 | 1 | 1965 | 660642 | 662606 | 0 | 3613 |
| YJM1549 | XII | 1 | 10 | 2549209 | leftend | YLL062C | 100 | 975 | 0 | 0 | 1 | 975 | 5167 | 4193 | 0 | 1801 |
| YJM1549 | XII | 1130 | 1139 | 2549209 | middle |  |  |  |  |  |  |  |  |  |  |  |
| YJM1549 | XII | 2547625 | 2547634 | 2549209 | middle |  |  |  |  |  |  |  |  |  |  |  |
| YJM1549 | XII | 2549209 | 2549209 | 2549209 | rightend_complete | YLR460C | 94.159 | 1130 | 64 | 2 | 3 | 1131 | 2546383 | 2545255 | 0 | 1720 |
| YJM1549 | XIII | 1 |  | 905641 | leftend_complete | YML131W | 100 | 1098 | 0 | 0 | 1 | 1098 | 15242 | 16339 | 0 | 2028 |
| YJM1549 | XIII | 640 | 649 | 905641 | middle |  |  |  |  |  |  |  |  |  |  |  |
| YJM1549 | XIII | 1802 | 1811 | 905641 | middle |  |  |  |  |  |  |  |  |  |  |  |
| YJM1549 | XIII | 904220 | 904229 | 905641 | middle |  |  |  |  |  |  |  |  |  |  |  |
| YJM1549 | XIII | 905641 | 905641 | 905641 | rightend_complete | YMR321C | 100 | 318 | 0 | 0 | 1 | 318 | 899546 | 899229 | 1.9E-167 | 588 |
| YJM1549 | XIV | 1 | 10 | 759332 | leftend | YNL332W | 98.631 | 1023 | 14 | 0 | 1 | 1023 | 8275 | 9297 | 0 | 1812 |
| YJM1549 | XIV | 2155 | 2164 | 759332 | middle |  |  |  |  |  |  |  |  |  |  |  |
| YJM1549 | XIV | 3174 | 3183 | 759332 | middle |  |  |  |  |  |  |  |  |  |  |  |
| YJM1549 | XIV | 758687 | 758696 | 759332 | middle |  |  |  |  |  |  |  |  |  |  |  |
| YJM1549 | XIV | 759332 | 759332 | 759332 | rightend_complete | YNR071C | 99.903 | 1029 | 1 | 0 | 1 | 1029 | 748470 | 747442 | 0 | 1895 |
| YJM1549 | XV | 1 | 10 | 1113104 | leftend | YOL154W | 99.867 | 750 | 1 | 0 | 1 | 750 | 23386 | 24135 | 0 | 1380 |
| YJM1549 | XV | 311 | 320 | 1113104 | middle |  |  |  |  |  |  |  |  |  |  |  |
| YJM1549 | XV | 1112228 | 1112237 | 1113104 | middle |  |  |  |  |  |  |  |  |  |  |  |
| YJM1549 | XV | 1113095 | 1113104 | 1113104 | rightend | YOR381W | 99.953 | 2136 | 1 | 0 | 1 | 2136 | 1020517 | 1022652 | 0 | 3940 |
| YJM1549 | XVI | 1 |  | 558153 | leftend_complete | YHL044W | 98.446 | 708 | 11 | 0 | 1 | 708 | 8948 | 9655 | 0 | 1247 |
| YJM1549 | XVI | 556583 | 556592 | 558153 | middle |  |  |  |  |  |  |  |  |  |  |  |
| YJM1549 | XVI | 558144 | 558153 | 558153 | rightend | YPR201W | 99.918 | 1215 | 1 | 0 | 1 | 1215 | 552079 | 553293 | 0 | 2239 |
| YJM1573 | I | 1 |  | 220843 | leftend_complete | YAL063C | 85.469 | 2037 | 292 | 4 | 1 | 2035 | 21371 | 19337 | 0 | 2119 |
| YJM1573 | I | 220843 | 220843 | 220843 | rightend_complete | YAR050W | 85.491 | 1923 | 245 | 17 | 1777 | 3676 | 195732 | 197643 | 0 | 1975 |
| YJM1573 | II | 1 |  | 780342 | leftend_complete | YNCB0001W | 100 | 84 | 0 | 0 | 1 | 84 | 2564 | 2647 | 4.16E-38 | 156 |
| YJM1573 | II | 1401 | 1410 | 780342 | middle |  |  |  |  |  |  |  |  |  |  |  |
| YJM1573 | II | 780333 | 780342 | 780342 | rightend | YBR297W | 98.257 | 1205 | 21 | 0 | 1 | 1205 | 775492 | 776696 | 0 | 2109 |
| YJM1573 | III | 1 | 10 | 302887 | leftend | YCL073C | 99.08 | 1848 | 17 | 0 | 1 | 1848 | 7361 | 5514 | 0 | 3319 |
| YJM1573 | III | 302878 | 302887 | 302887 | rightend | YCR101C | 97.636 | 550 | 12 | 1 | 1 | 549 | 296134 | 295585 | 0 | 942 |
| YJM1573 | IV | 1 |  | 1465733 | leftend_complete | YDL243C | 93.131 | 990 | 68 | 0 | 1 | 990 | 18117 | 17128 | 0 | 1452 |
| YJM1573 | IV | 1812 | 1821 | 1465733 | middle |  |  |  |  |  |  |  |  |  |  |  |
| YJM1573 | IV | 1463777 | 1463786 | 1465733 | middle |  |  |  |  |  |  |  |  |  |  |  |
| YJM1573 | IV | 1465733 | 1465733 | 1465733 | rightend_complete | YDR541C | 99.517 | 1035 | 5 | 0 | 1 | 1035 | 1461054 | 1460020 | 0 | 1884 |
| YJM1573 | V | 1 |  | 557232 | leftend_complete | YEL070W | 99.536 | 1509 | 7 | 0 | 1 | 1509 | 20207 | 21715 | 0 | 2748 |
| YJM1573 | V | 556068 | 556077 | 557232 | middle |  |  |  |  |  |  |  |  |  |  |  |
| YJM1573 | V | 557232 | 557232 | 557232 | rightend_complete | YER187W | 99.061 | 426 | 4 | 0 | 1 | 426 | 552385 | 552810 | 0 | 765 |
| YJM1573 | VI | 1 |  | 277048 | leftend_complete | YFL051C | 100 | 452 | 0 | 0 | 1 | 452 | 38775 | 38324 | 0 | 835 |
| YJM1573 | VI | 1154 | 1163 | 277048 | middle |  |  |  |  |  |  |  |  |  |  |  |
| YJM1573 | VI | 3169 | 3178 | 277048 | middle |  |  |  |  |  |  |  |  |  |  |  |
| YJM1573 | VI | 5419 | 5428 | 277048 | middle |  |  |  |  |  |  |  |  |  |  |  |
| YJM1573 | VI | 277039 | 277048 | 277048 | rightend | YFR055W | 99.609 | 1022 | 4 | 0 | 1 | 1022 | 267335 | 268356 | 0 | 1866 |
| YJM1573 | VII | 1 |  | 1076414 | leftend_complete | YGL258W | 99.678 | 621 | 2 | 0 | 1 | 621 | 11399 | 12019 | 0 | 1136 |
| YJM1573 | VII | 6580 | 6589 | 1076414 | middle |  |  |  |  |  |  |  |  |  |  |  |
| YJM1573 | VII | 1075560 | 1075569 | 1076414 | middle |  |  |  |  |  |  |  |  |  |  |  |
| YJM1573 | VII | 1076405 | 1076414 | 1076414 | rightend | YGR287C | 99.548 | 1770 | 8 | 0 | 1 | 1770 | 1039255 | 1037486 | 0 | 3225 |
| YJM1573 | VIII | 1 |  | 515612 | leftend_complete | YHL044W | 98.729 | 708 | 9 | 0 | 1 | 708 | 8792 | 9499 | 0 | 1258 |
| YJM1573 | VIII | 513826 | 513835 | 515612 | middle |  |  |  |  |  |  |  |  |  |  |  |
| YJM1573 | VIII | 515603 | 515612 | 515612 | rightend | YHR211W | 91.589 | 1403 | 94 | 2 | 1519 | 2918 | 513836 | 515217 | 0 | 1916 |
| YJM1573 | IX | 1 |  | 422341 | leftend_complete | YIL169C | 96.131 | 2223 | 75 | 7 | 394 | 2612 | 13926 | 11711 | 0 | 3618 |
| YJM1573 | IX | 633 | 642 | 422341 | middle |  |  |  |  |  |  |  |  |  |  |  |
| YJM1573 | IX | 422341 | 422341 | 422341 | rightend_complete | YIR039C | 99.69 | 1614 | 5 | 0 | 1 | 1614 | 414608 | 412995 | 0 | 2953 |
| YJM1573 | X | 1 | 10 | 718800 | leftend | YJL213W | 99.799 | 996 | 2 | 0 | 1 | 996 | 27493 | 28488 | 0 | 1829 |
| YJM1573 | X | 718800 | 718800 | 718800 | rightend_complete | YJR152W | 99.571 | 1632 | 7 | 0 | 1 | 1632 | 697953 | 699584 | 0 | 2976 |
| YJM1573 | XI | 1 | 10 | 656996 | leftend | YKL222C | 99.622 | 2118 | 8 | 0 | 1 | 2118 | 3942 | 1825 | 0 | 3868 |
| YJM1573 | XI | 656987 | 656996 | 656996 | rightend | YKR101W | 99.542 | 1965 | 9 | 0 | 1 | 1965 | 638221 | 640185 | 0 | 3579 |
| YJM1573 | XII | 1 |  | 1539518 | leftend_complete | YLL062C | 99.59 | 975 | 4 | 0 | 1 | 975 | 5897 | 4923 | 0 | 1779 |
| YJM1573 | XII | 1425 | 1434 | 1539518 | middle |  |  |  |  |  |  |  |  |  |  |  |
| YJM1573 | XII | 1538613 | 1538622 | 1539518 | middle |  |  |  |  |  |  |  |  |  |  |  |
| YJM1573 | XII | 1539509 | 1539518 | 1539518 | rightend | YLR460C | 99.646 | 1131 | 4 | 0 | 1 | 1131 | 1536560 | 1535430 | 0 | 2067 |
| YJM1573 | XIII | 1 |  | 909936 | leftend_complete | YML131W | 99.545 | 1098 | 5 | 0 | 1 | 1098 | 4659 | 5756 | 0 | 2001 |
| YJM1573 | XIII | 899149 | 899158 | 909936 | middle |  |  |  |  |  |  |  |  |  |  |  |
| YJM1573 | XIII | 905728 | 905737 | 909936 | middle |  |  |  |  |  |  |  |  |  |  |  |
| YJM1573 | XIII | 909927 | 909936 | 909936 | rightend | YMR321C | 99.042 | 313 | 3 | 0 | 6 | 318 | 891651 | 891339 | 1.15E-159 | 562 |
| YJM1573 | XIV | 1 |  | 771369 | leftend_complete | YNL332W | 98.925 | 1023 | 11 | 0 | 1 | 1023 | 7417 | 8439 | 0 | 1829 |
| YJM1573 | XIV | 745 | 754 | 771369 | middle |  |  |  |  |  |  |  |  |  |  |  |
| YJM1573 | XIV | 765551 | 765560 | 771369 | middle |  |  |  |  |  |  |  |  |  |  |  |
| YJM1573 | XIV | 771360 | 771369 | 771369 | rightend | YNR071C | 99.611 | 1029 | 4 | 0 | 1 | 1029 | 746147 | 745119 | 0 | 1879 |
| YJM1573 | XV | 1 | 10 | 1059853 | leftend | YOL154W | 99.333 | 750 | 5 | 0 | 1 | 750 | 35188 | 35937 | 0 | 1358 |
| YJM1573 | XV | 1059844 | 1059853 | 1059853 | rightend | YOR381W | 98.876 | 2136 | 24 | 0 | 1 | 2136 | 1032620 | 1034755 | 0 | 3812 |
| YJM1573 | XVI | 1 |  | 897414 | leftend_complete | YPL274W | 98.639 | 1764 | 24 | 0 | 1 | 1764 | 5961 | 7724 | 0 | 3125 |
| YJM1573 | XVI | 568 | 577 | 897414 | middle |  |  |  |  |  |  |  |  |  |  |  |
| YJM1573 | XVI | 897414 | 897414 | 897414 | rightend_complete | YPR201W | 99.588 | 1215 | 5 | 0 | 1 | 1215 | 894200 | 895414 | 0 | 2217 |
| YJM1574 | I | 1 | 10 | 200724 | leftend | YAL063C | 87.454 | 1092 | 133 | 4 | 1 | 1090 | 2166 | 1077 | 0 | 1254 |
| YJM1574 | I | 608 | 617 | 200724 | middle |  |  |  |  |  |  |  |  |  |  |  |
| YJM1574 | I | 173081 | 173090 | 200724 | middle |  |  |  |  |  |  |  |  |  |  |  |
| YJM1574 | I | 192225 | 192234 | 200724 | middle |  |  |  |  |  |  |  |  |  |  |  |
| YJM1574 | I | 200715 | 200724 | 200724 | rightend | YAR050W | 83.922 | 1132 | 178 | 4 | 1 | 1130 | 171951 | 173080 | 0 | 1079 |
| YJM1574 | II | 1 | 10 | 784187 | leftend | YNCB0001W | 100 | 84 | 0 | 0 | 1 | 84 | 1808 | 1891 | 4.16E-38 | 156 |
| YJM1574 | II | 562 | 571 | 784187 | middle |  |  |  |  |  |  |  |  |  |  |  |
| YJM1574 | II | 1368 | 1377 | 784187 | middle |  |  |  |  |  |  |  |  |  |  |  |
| YJM1574 | II | 784178 | 784187 | 784187 | rightend | YBR297W | 99.076 | 1407 | 13 | 0 | 1 | 1407 | 774623 | 776029 | 0 | 2527 |
| YJM1574 | III | 1 |  | 309125 | leftend_complete | YCL073C | 99.675 | 1848 | 6 | 0 | 1 | 1848 | 8209 | 6362 | 0 | 3380 |
| YJM1574 | III | 710 | 719 | 309125 | middle |  |  |  |  |  |  |  |  |  |  |  |
| YJM1574 | III | 300515 | 300524 | 309125 | middle |  |  |  |  |  |  |  |  |  |  |  |
| YJM1574 | III | 308635 | 308644 | 309125 | middle |  |  |  |  |  |  |  |  |  |  |  |
| YJM1574 | III | 309125 | 309125 | 309125 | rightend_complete | YCR101C | 99.818 | 549 | 1 | 0 | 1 | 549 | 296553 | 296005 | 0 | 1009 |
| YJM1574 | IV | 1 | 10 | 1466940 | leftend | YDL243C | 95.152 | 990 | 48 | 0 | 1 | 990 | 16832 | 15843 | 0 | 1563 |
| YJM1574 | IV | 1466931 | 1466940 | 1466940 | rightend | YDR541C | 99.517 | 1035 | 5 | 0 | 1 | 1035 | 1462892 | 1461858 | 0 | 1884 |
| YJM1574 | V | 1 | 10 | 552748 | leftend | YEL070W | 99.072 | 1509 | 14 | 0 | 1 | 1509 | 13095 | 14603 | 0 | 2710 |
| YJM1574 | V | 552283 | 552292 | 552748 | middle |  |  |  |  |  |  |  |  |  |  |  |
| YJM1574 | V | 552739 | 552748 | 552748 | rightend | YER187W | 91.981 | 424 | 32 | 2 | 4 | 426 | 548292 | 548714 | 3.44E-169 | 593 |
| YJM1574 | VI | 1 | 10 | 242554 | leftend | YFL051C | 98.451 | 452 | 7 | 0 | 1 | 452 | 9673 | 9222 | 0 | 797 |
| YJM1574 | VI | 955 | 964 | 242554 | middle |  |  |  |  |  |  |  |  |  |  |  |
| YJM1574 | VI | 242554 | 242554 | 242554 | rightend_complete | YFR055W | 100 | 1023 | 0 | 0 | 1 | 1023 | 236576 | 237598 | 0 | 1890 |
| YJM1574 | VII | 1 | 10 | 1039731 | leftend | YGL258W | 97.585 | 621 | 15 | 0 | 1 | 621 | 11399 | 12019 | 0 | 1064 |
| YJM1574 | VII | 6920 | 6929 | 1039731 | middle |  |  |  |  |  |  |  |  |  |  |  |
| YJM1574 | VII | 1039722 | 1039731 | 1039731 | rightend | YGR287C | 99.139 | 1626 | 14 | 0 | 145 | 1770 | 1031390 | 1029765 | 0 | 2926 |
| YJM1574 | VIII | 1 | 10 | 895609 | leftend | YPL274W | 99.887 | 1764 | 2 | 0 | 1 | 1764 | 7627 | 9390 | 0 | 3247 |
| YJM1574 | VIII | 469 | 478 | 895609 | middle |  |  |  |  |  |  |  |  |  |  |  |
| YJM1574 | VIII | 1505 | 1514 | 895609 | middle |  |  |  |  |  |  |  |  |  |  |  |
| YJM1574 | VIII | 880767 | 880776 | 895609 | middle |  |  |  |  |  |  |  |  |  |  |  |
| YJM1574 | VIII | 891055 | 891064 | 895609 | middle |  |  |  |  |  |  |  |  |  |  |  |
| YJM1574 | VIII | 894295 | 894304 | 895609 | middle |  |  |  |  |  |  |  |  |  |  |  |
| YJM1574 | VIII | 894983 | 894992 | 895609 | middle |  |  |  |  |  |  |  |  |  |  |  |
| YJM1574 | VIII | 895404 | 895413 | 895609 | middle |  |  |  |  |  |  |  |  |  |  |  |
| YJM1574 | VIII | 895609 | 895609 | 895609 | rightend_complete | YHR211W | 90.581 | 1826 | 148 | 2 | 1096 | 2918 | 878577 | 880381 | 0 | 2398 |
| YJM1574 | IX | 1 | 10 | 406014 | leftend | YIL169C | 93.199 | 1294 | 80 | 6 | 520 | 1812 | 5856 | 4570 | 0 | 1895 |
| YJM1574 | IX | 406005 | 406014 | 406014 | rightend | YIR039C | 99.009 | 1614 | 16 | 0 | 1 | 1614 | 403902 | 402289 | 0 | 2892 |
| YJM1574 | X | 1 | 10 | 715077 | leftend | YJL213W | 99.699 | 996 | 3 | 0 | 1 | 996 | 24434 | 25429 | 0 | 1823 |
| YJM1574 | X | 715077 | 715077 | 715077 | rightend_complete | YJR152W | 99.939 | 1632 | 1 | 0 | 1 | 1632 | 695043 | 696674 | 0 | 3009 |
| YJM1574 | XI | 1 | 10 | 655452 | leftend | YKL222C | 99.481 | 2118 | 11 | 0 | 1 | 2118 | 3135 | 1018 | 0 | 3851 |
| YJM1574 | XI | 654854 | 654863 | 655452 | middle |  |  |  |  |  |  |  |  |  |  |  |
| YJM1574 | XI | 655452 | 655452 | 655452 | rightend_complete | YKR101W | 99.746 | 1965 | 5 | 0 | 1 | 1965 | 636533 | 638497 | 0 | 3602 |
| YJM1574 | XII | 1 |  | 1810783 | leftend_complete | YLL062C | 100 | 975 | 0 | 0 | 1 | 975 | 2524 | 1550 | 0 | 1801 |
| YJM1574 | XII | 221 | 230 | 1810783 | middle |  |  |  |  |  |  |  |  |  |  |  |
| YJM1574 | XII | 813 | 822 | 1810783 | middle |  |  |  |  |  |  |  |  |  |  |  |
| YJM1574 | XII | 1810774 | 1810783 | 1810783 | rightend | YLR460C | 94.159 | 1130 | 64 | 2 | 3 | 1131 | 1808421 | 1807293 | 0 | 1720 |
| YJM1574 | XIII | 1 | 10 | 899489 | leftend | YML131W | 99.818 | 1098 | 2 | 0 | 1 | 1098 | 4421 | 5518 | 0 | 2017 |
| YJM1574 | XIII | 709 | 718 | 899489 | middle |  |  |  |  |  |  |  |  |  |  |  |
| YJM1574 | XIII | 2032 | 2041 | 899489 | middle |  |  |  |  |  |  |  |  |  |  |  |
| YJM1574 | XIII | 899480 | 899489 | 899489 | rightend | YMR321C | 99.361 | 313 | 2 | 0 | 6 | 318 | 890552 | 890240 | 2.48E-161 | 568 |
| YJM1574 | XIV | 1 | 10 | 768434 | leftend | YNL332W | 98.534 | 1023 | 15 | 0 | 1 | 1023 | 5635 | 6657 | 0 | 1807 |
| YJM1574 | XIV | 766476 | 766485 | 768434 | middle |  |  |  |  |  |  |  |  |  |  |  |
| YJM1574 | XIV | 768434 | 768434 | 768434 | rightend_complete | YNR071C | 99.903 | 1029 | 1 | 0 | 1 | 1029 | 761192 | 760164 | 0 | 1895 |
| YJM1574 | XV | 1 | 10 | 1047917 | leftend | YOL154W | 99.467 | 750 | 4 | 0 | 1 | 750 | 22250 | 22999 | 0 | 1363 |
| YJM1574 | XV | 1045538 | 1045547 | 1047917 | middle |  |  |  |  |  |  |  |  |  |  |  |
| YJM1574 | XV | 1047917 | 1047917 | 1047917 | rightend_complete | YOR381W | 99.953 | 2136 | 1 | 0 | 1 | 2136 | 1018972 | 1021107 | 0 | 3940 |
| YJM1574 | XVI | 1 | 10 | 557146 | leftend | YHL044W | 98.446 | 708 | 11 | 0 | 1 | 708 | 8431 | 9138 | 0 | 1247 |
| YJM1574 | XVI | 555437 | 555446 | 557146 | middle |  |  |  |  |  |  |  |  |  |  |  |
| YJM1574 | XVI | 555757 | 555766 | 557146 | middle |  |  |  |  |  |  |  |  |  |  |  |
| YJM1574 | XVI | 557137 | 557146 | 557146 | rightend | YPR201W | 100 | 1215 | 0 | 0 | 1 | 1215 | 551762 | 552976 | 0 | 2244 |
| YJM1592 | I | 1 | 10 | 192254 | leftend | YAL063C | 87.69 | 1779 | 156 | 34 | 1912 | 3660 | 190499 | 192244 | 0 | 2013 |
| YJM1592 | I | 3889 | 3898 | 192254 | middle |  |  |  |  |  |  |  |  |  |  |  |
| YJM1592 | I | 192245 | 192254 | 192254 | rightend | YAR050W | 91.534 | 1760 | 93 | 13 | 2574 | 4278 | 190486 | 192244 | 0 | 2374 |
| YJM1592 | II | 1 | 10 | 793231 | leftend | YNCB0001W | 100 | 84 | 0 | 0 | 1 | 84 | 1570 | 1653 | 4.16E-38 | 156 |
| YJM1592 | II | 793222 | 793231 | 793231 | rightend | YBR297W | 97.925 | 1205 | 25 | 0 | 1 | 1205 | 773382 | 774586 | 0 | 2087 |
| YJM1592 | III | 1 | 10 | 302308 | leftend | YCL073C | 98.755 | 1848 | 23 | 0 | 1 | 1848 | 6946 | 5099 | 0 | 3286 |
| YJM1592 | III | 298120 | 298129 | 302308 | middle |  |  |  |  |  |  |  |  |  |  |  |
| YJM1592 | III | 301717 | 301726 | 302308 | middle |  |  |  |  |  |  |  |  |  |  |  |
| YJM1592 | III | 302308 | 302308 | 302308 | rightend_complete | YCR101C | 98 | 550 | 10 | 1 | 1 | 549 | 295560 | 295011 | 0 | 953 |
| YJM1592 | IV | 1 | 10 | 1451639 | leftend | YDL243C | 93.838 | 990 | 61 | 0 | 1 | 990 | 2505 | 1516 | 0 | 1491 |
| YJM1592 | IV | 1451630 | 1451639 | 1451639 | rightend | YDR541C | 99.517 | 1035 | 5 | 0 | 1 | 1035 | 1448926 | 1447892 | 0 | 1884 |
| YJM1592 | V | 1 |  | 550177 | leftend_complete | YEL070W | 99.47 | 1509 | 8 | 0 | 1 | 1509 | 13371 | 14879 | 0 | 2743 |
| YJM1592 | V | 549655 | 549664 | 550177 | middle |  |  |  |  |  |  |  |  |  |  |  |
| YJM1592 | V | 550177 | 550177 | 550177 | rightend_complete | YER187W | 99.765 | 426 | 1 | 0 | 1 | 426 | 546046 | 546471 | 0 | 782 |
| YJM1592 | VI | 1 |  | 241423 | leftend_complete | YFL050C | 99.321 | 1472 | 10 | 0 | 1 | 1472 | 5821 | 4350 | 0 | 2663 |
| YJM1592 | VI | 1218 | 1227 | 241423 | middle |  |  |  |  |  |  |  |  |  |  |  |
| YJM1592 | VI | 238854 | 238863 | 241423 | middle |  |  |  |  |  |  |  |  |  |  |  |
| YJM1592 | VI | 239282 | 239291 | 241423 | middle |  |  |  |  |  |  |  |  |  |  |  |
| YJM1592 | VI | 241414 | 241423 | 241423 | rightend | YFR055W | 99.609 | 1022 | 4 | 0 | 1 | 1022 | 228514 | 229535 | 0 | 1866 |
| YJM1592 | VII | 1 | 10 | 1058650 | leftend | YGL258W | 99.195 | 621 | 5 | 0 | 1 | 621 | 4800 | 5420 | 0 | 1120 |
| YJM1592 | VII | 1055813 | 1055822 | 1058650 | middle |  |  |  |  |  |  |  |  |  |  |  |
| YJM1592 | VII | 1057114 | 1057123 | 1058650 | middle |  |  |  |  |  |  |  |  |  |  |  |
| YJM1592 | VII | 1058641 | 1058650 | 1058650 | rightend | YGR287C | 99.04 | 1770 | 17 | 0 | 1 | 1770 | 1033583 | 1031814 | 0 | 3175 |
| YJM1592 | VIII | 1 | 10 | 516466 | leftend | YHL044W | 98.588 | 708 | 10 | 0 | 1 | 708 | 8132 | 8839 | 0 | 1253 |
| YJM1592 | VIII | 516457 | 516466 | 516466 | rightend | YHR210C | 99.025 | 1026 | 8 | 1 | 1 | 1026 | 506889 | 505866 | 0 | 1838 |
| YJM1592 | IX | 1 | 10 | 409478 | leftend | YIL169C | 98.45 | 2322 | 30 | 2 | 478 | 2799 | 7058 | 4743 | 0 | 4084 |
| YJM1592 | IX | 408136 | 408145 | 409478 | middle |  |  |  |  |  |  |  |  |  |  |  |
| YJM1592 | IX | 409469 | 409478 | 409478 | rightend | YIR039C | 99.69 | 1614 | 5 | 0 | 1 | 1614 | 407066 | 405453 | 0 | 2953 |
| YJM1592 | X | 1 | 10 | 727970 | leftend | YJL213W | 99.9 | 996 | 1 | 0 | 1 | 996 | 28710 | 29705 | 0 | 1834 |
| YJM1592 | X | 263 | 272 | 727970 | middle |  |  |  |  |  |  |  |  |  |  |  |
| YJM1592 | X | 3457 | 3466 | 727970 | middle |  |  |  |  |  |  |  |  |  |  |  |
| YJM1592 | X | 6415 | 6424 | 727970 | middle |  |  |  |  |  |  |  |  |  |  |  |
| YJM1592 | X | 727961 | 727970 | 727970 | rightend | YJR152W | 99.449 | 1632 | 9 | 0 | 1 | 1632 | 704473 | 706104 | 0 | 2964 |
| YJM1592 | XI | 1 | 10 | 656922 | leftend | YKL222C | 99.858 | 2118 | 3 | 0 | 1 | 2118 | 3333 | 1216 | 0 | 3895 |
| YJM1592 | XI | 656913 | 656922 | 656922 | rightend | YKR101W | 99.644 | 1965 | 7 | 0 | 1 | 1965 | 637370 | 639334 | 0 | 3591 |
| YJM1592 | XII | 1 | 10 | 1779744 | leftend | YLL062C | 99.795 | 975 | 2 | 0 | 1 | 975 | 13556 | 12582 | 0 | 1790 |
| YJM1592 | XII | 1776256 | 1776265 | 1779744 | middle |  |  |  |  |  |  |  |  |  |  |  |
| YJM1592 | XII | 1777890 | 1777899 | 1779744 | middle |  |  |  |  |  |  |  |  |  |  |  |
| YJM1592 | XII | 1778608 | 1778617 | 1779744 | middle |  |  |  |  |  |  |  |  |  |  |  |
| YJM1592 | XII | 1779364 | 1779373 | 1779744 | middle |  |  |  |  |  |  |  |  |  |  |  |
| YJM1592 | XII | 1779735 | 1779744 | 1779744 | rightend | YLR460C | 99.823 | 1131 | 2 | 0 | 1 | 1131 | 1775233 | 1774103 | 0 | 2078 |
| YJM1592 | XIII | 1 |  | 890874 | leftend_complete | YML131W | 98.998 | 1098 | 11 | 0 | 1 | 1098 | 4679 | 5776 | 0 | 1967 |
| YJM1592 | XIII | 890865 | 890874 | 890874 | rightend | YMR321C | 99.371 | 318 | 2 | 0 | 1 | 318 | 889801 | 889484 | 4.12E-164 | 577 |
| YJM1592 | XIV | 1 | 10 | 766222 | leftend | YNL332W | 99.413 | 1023 | 6 | 0 | 1 | 1023 | 6782 | 7804 | 0 | 1857 |
| YJM1592 | XIV | 885 | 894 | 766222 | middle |  |  |  |  |  |  |  |  |  |  |  |
| YJM1592 | XIV | 766213 | 766222 | 766222 | rightend | YNR071C | 99.515 | 1030 | 4 | 1 | 1 | 1029 | 746931 | 745902 | 0 | 1873 |
| YJM1592 | XV | 1 |  | 1058281 | leftend_complete | YOL154W | 99.6 | 750 | 3 | 0 | 1 | 750 | 31264 | 32013 | 0 | 1369 |
| YJM1592 | XV | 1058272 | 1058281 | 1058281 | rightend | YOR381W | 98.642 | 2136 | 29 | 0 | 1 | 2136 | 1029407 | 1031542 | 0 | 3784 |
| YJM1592 | XVI | 1 | 10 | 904157 | leftend | YPL274W | 98.526 | 1764 | 26 | 0 | 1 | 1764 | 14007 | 15770 | 0 | 3114 |
| YJM1592 | XVI | 904157 | 904157 | 904157 | rightend_complete | YPR201W | 99.671 | 1215 | 4 | 0 | 1 | 1215 | 900857 | 902071 | 0 | 2222 |
| YJM1615 | I | 1 | 10 | 217876 | leftend | YAL063C | 89.254 | 2038 | 211 | 6 | 1 | 2035 | 13471 | 11439 | 0 | 2562 |
| YJM1615 | I | 10661 | 10670 | 217876 | middle |  |  |  |  |  |  |  |  |  |  |  |
| YJM1615 | I | 12178 | 12187 | 217876 | middle |  |  |  |  |  |  |  |  |  |  |  |
| YJM1615 | I | 190080 | 190089 | 217876 | middle |  |  |  |  |  |  |  |  |  |  |  |
| YJM1615 | I | 210085 | 210094 | 217876 | middle |  |  |  |  |  |  |  |  |  |  |  |
| YJM1615 | I | 217867 | 217876 | 217876 | rightend | YAR050W | 93.344 | 1818 | 76 | 10 | 2839 | 4614 | 190090 | 191904 | 0 | 2645 |
| YJM1615 | II | 1 |  | 783969 | leftend_complete | YNCB0001W | 100 | 84 | 0 | 0 | 1 | 84 | 4628 | 4711 | 4.16E-38 | 156 |
| YJM1615 | II | 1494 | 1503 | 783969 | middle |  |  |  |  |  |  |  |  |  |  |  |
| YJM1615 | II | 3044 | 3053 | 783969 | middle |  |  |  |  |  |  |  |  |  |  |  |
| YJM1615 | II | 783960 | 783969 | 783969 | rightend | YBR297W | 97.759 | 1205 | 27 | 0 | 1 | 1205 | 778441 | 779645 | 0 | 2076 |
| YJM1615 | III | 1 | 10 | 300711 | leftend | YCL073C | 99.892 | 1848 | 2 | 0 | 1 | 1848 | 8331 | 6484 | 0 | 3402 |
| YJM1615 | III | 300702 | 300711 | 300711 | rightend | YCR101C | 100 | 549 | 0 | 0 | 1 | 549 | 295956 | 295408 | 0 | 1014 |
| YJM1615 | IV | 1 |  | 1471855 | leftend_complete | YDL243C | 93.769 | 979 | 61 | 0 | 1 | 979 | 12580 | 11602 | 0 | 1471 |
| YJM1615 | IV | 3119 | 3128 | 1471855 | middle |  |  |  |  |  |  |  |  |  |  |  |
| YJM1615 | IV | 1467928 | 1467937 | 1471855 | middle |  |  |  |  |  |  |  |  |  |  |  |
| YJM1615 | IV | 1468120 | 1468129 | 1471855 | middle |  |  |  |  |  |  |  |  |  |  |  |
| YJM1615 | IV | 1470235 | 1470244 | 1471855 | middle |  |  |  |  |  |  |  |  |  |  |  |
| YJM1615 | IV | 1471855 | 1471855 | 1471855 | rightend_complete | YDR541C | 99.903 | 1035 | 1 | 0 | 1 | 1035 | 1465330 | 1464296 | 0 | 1906 |
| YJM1615 | V | 1 | 10 | 553423 | leftend | YEL070W | 100 | 1509 | 0 | 0 | 1 | 1509 | 15155 | 16663 | 0 | 2787 |
| YJM1615 | V | 1568 | 1577 | 553423 | middle |  |  |  |  |  |  |  |  |  |  |  |
| YJM1615 | V | 552399 | 552408 | 553423 | middle |  |  |  |  |  |  |  |  |  |  |  |
| YJM1615 | V | 553414 | 553423 | 553423 | rightend | YER187W | 99.531 | 426 | 2 | 0 | 1 | 426 | 548601 | 549026 | 0 | 776 |
| YJM1615 | VI | 1 | 10 | 250040 | leftend | YFL051C | 100 | 483 | 0 | 0 | 1 | 483 | 16875 | 16393 | 0 | 893 |
| YJM1615 | VI | 250040 | 250040 | 250040 | rightend_complete | YFR055W | 99.902 | 1023 | 1 | 0 | 1 | 1023 | 244102 | 245124 | 0 | 1884 |
| YJM1615 | VII | 1 |  | 1040875 | leftend_complete | YGL258W | 99.356 | 621 | 4 | 0 | 1 | 621 | 8468 | 9088 | 0 | 1125 |
| YJM1615 | VII | 6459 | 6468 | 1040875 | middle |  |  |  |  |  |  |  |  |  |  |  |
| YJM1615 | VII | 1040350 | 1040359 | 1040875 | middle |  |  |  |  |  |  |  |  |  |  |  |
| YJM1615 | VII | 1040875 | 1040875 | 1040875 | rightend_complete | YGR287C | 98.588 | 1770 | 25 | 0 | 1 | 1770 | 1031606 | 1029837 | 0 | 3131 |
| YJM1615 | VIII | 1 |  | 526580 | leftend_complete | YHL044W | 98.588 | 708 | 10 | 0 | 1 | 708 | 9624 | 10331 | 0 | 1253 |
| YJM1615 | VIII | 8228 | 8237 | 526580 | middle |  |  |  |  |  |  |  |  |  |  |  |
| YJM1615 | VIII | 516610 | 516619 | 526580 | middle |  |  |  |  |  |  |  |  |  |  |  |
| YJM1615 | VIII | 519003 | 519012 | 526580 | middle |  |  |  |  |  |  |  |  |  |  |  |
| YJM1615 | VIII | 521678 | 521687 | 526580 | middle |  |  |  |  |  |  |  |  |  |  |  |
| YJM1615 | VIII | 526580 | 526580 | 526580 | rightend_complete | YHR211W | 83.537 | 1148 | 171 | 14 | 1 | 1139 | 515453 | 516591 | 0 | 1057 |
| YJM1615 | IX | 1 |  | 425910 | leftend_complete | YIL169C | 96.725 | 1374 | 39 | 2 | 478 | 1851 | 19329 | 17962 | 0 | 2283 |
| YJM1615 | IX | 832 | 841 | 425910 | middle |  |  |  |  |  |  |  |  |  |  |  |
| YJM1615 | IX | 425901 | 425910 | 425910 | rightend | YIR039C | 99.814 | 1614 | 3 | 0 | 1 | 1614 | 420038 | 418425 | 0 | 2964 |
| YJM1615 | X | 1 |  | 703461 | leftend_complete | YJL213W | 99.699 | 996 | 3 | 0 | 1 | 996 | 13887 | 14882 | 0 | 1823 |
| YJM1615 | X | 2127 | 2136 | 703461 | middle |  |  |  |  |  |  |  |  |  |  |  |
| YJM1615 | X | 2821 | 2830 | 703461 | middle |  |  |  |  |  |  |  |  |  |  |  |
| YJM1615 | X | 703461 | 703461 | 703461 | rightend_complete | YJR152W | 99.939 | 1632 | 1 | 0 | 1 | 1632 | 683350 | 684981 | 0 | 3009 |
| YJM1615 | XI | 1 | 10 | 660069 | leftend | YKL222C | 99.669 | 2118 | 7 | 0 | 1 | 2118 | 3280 | 1163 | 0 | 3873 |
| YJM1615 | XI | 660060 | 660069 | 660069 | rightend | YKR101W | 99.796 | 1965 | 4 | 0 | 1 | 1965 | 637398 | 639362 | 0 | 3607 |
| YJM1615 | XII | 1 |  | 1919394 | leftend_complete | YLL062C | 99.282 | 975 | 7 | 0 | 1 | 975 | 6218 | 5244 | 0 | 1762 |
| YJM1615 | XII | 1997 | 2006 | 1919394 | middle |  |  |  |  |  |  |  |  |  |  |  |
| YJM1615 | XII | 1917705 | 1917714 | 1919394 | middle |  |  |  |  |  |  |  |  |  |  |  |
| YJM1615 | XII | 1919394 | 1919394 | 1919394 | rightend_complete | YLR460C | 98.055 | 1131 | 22 | 0 | 1 | 1131 | 1916119 | 1914989 | 0 | 1967 |
| YJM1615 | XIII | 1 | 10 | 906128 | leftend | YML131W | 99.545 | 1098 | 5 | 0 | 1 | 1098 | 15241 | 16338 | 0 | 2001 |
| YJM1615 | XIII | 759 | 768 | 906128 | middle |  |  |  |  |  |  |  |  |  |  |  |
| YJM1615 | XIII | 2707 | 2716 | 906128 | middle |  |  |  |  |  |  |  |  |  |  |  |
| YJM1615 | XIII | 904388 | 904397 | 906128 | middle |  |  |  |  |  |  |  |  |  |  |  |
| YJM1615 | XIII | 906128 | 906128 | 906128 | rightend_complete | YMR321C | 99.371 | 318 | 2 | 0 | 1 | 318 | 900366 | 900049 | 4.12E-164 | 577 |
| YJM1615 | XIV | 1 | 10 | 768287 | leftend | YNL332W | 99.902 | 1023 | 1 | 0 | 1 | 1023 | 7448 | 8470 | 0 | 1884 |
| YJM1615 | XIV | 768278 | 768287 | 768287 | rightend | YNR071C | 99.611 | 1029 | 4 | 0 | 1 | 1029 | 746941 | 745913 | 0 | 1879 |
| YJM1615 | XV | 1 | 10 | 1061915 | leftend | YOL154W | 99.467 | 750 | 4 | 0 | 1 | 750 | 34170 | 34919 | 0 | 1363 |
| YJM1615 | XV | 1055373 | 1055382 | 1061915 | middle |  |  |  |  |  |  |  |  |  |  |  |
| YJM1615 | XV | 1061915 | 1061915 | 1061915 | rightend_complete | YOR381W | 99.345 | 2136 | 14 | 0 | 1 | 2136 | 1034192 | 1036327 | 0 | 3868 |
| YJM1615 | XVI | 1 | 10 | 904765 | leftend | YPL274W | 98.129 | 1764 | 33 | 0 | 1 | 1764 | 13842 | 15605 | 0 | 3075 |
| YJM1615 | XVI | 809 | 818 | 904765 | middle |  |  |  |  |  |  |  |  |  |  |  |
| YJM1615 | XVI | 2599 | 2608 | 904765 | middle |  |  |  |  |  |  |  |  |  |  |  |
| YJM1615 | XVI | 3095 | 3104 | 904765 | middle |  |  |  |  |  |  |  |  |  |  |  |
| YJM1615 | XVI | 903261 | 903270 | 904765 | middle |  |  |  |  |  |  |  |  |  |  |  |
| YJM1615 | XVI | 904756 | 904765 | 904765 | rightend | YPR201W | 99.588 | 1215 | 5 | 0 | 1 | 1215 | 900301 | 901515 | 0 | 2217 |

Leftend and rightend indicate the chromosome sequence currently terminates short of the terminal telomere sequence. Leftend_complete and rightend_complete indicate the chromosome ends with the terminal telomere repeat.

Figure S1 *YFL051C*, *FLO12* is a FLO gene in some strains, and a truncated ORF in S288c and some other strains.

A) Alignment of *YFL051C* from S288c (top) and YJM1443 (bottom).

B) Smith-Waterman alignment of *FLO12* from YJM1443 with *FLO10* from S288c, the most similar flocculin gene to *FLO12* in S288c. Only the conserved regions near each end of the gene are shown. The central regions of the FLO genes share little sequence conservation and contain low complexity repetitive sequences.

A)

```
S288c    MSIPHSVFSALLVFVALATTTLASTEACLPTNKREDGMNINFYEYTIGDQTTYLEPEYMG
         ||||||||||||||||||||||||||||||||||||||||||||||||||||||||||||
YJM1443  MSIPHSVFSALLVFVALATTTLASTEACLPTNKREDGMNINFYEYTIGDQTTYLEPEYMG

S288c    YEYSNTKKLGSVSGQTNLSIYYSPPCESTPTCVTYAVLKRDEDGYDPCGPLYETKKRDTE
         ||||||||||||||||||||||||||||||||||||||||||||||||||||||||||||
YJM1443  YEYSNTKKLGSVSGQTNLSIYYSPPCESTPTCVTYAVLKRDEDGYDPCGPLYETKKRDTE

S288c    YCDPNTAYWSSDLFGFYTTPTNVTVEMTGYLIWSMGNRRR*
         ||||||||||||||||||||||||||||||
YJM1443  YCDPNTAYWSSDLFGFYTTPTNVTVEMTGYFLAPENGSYTFSFYTVDDSAILSIGGDNAF

YJM1443  ACCEQQQSSVTSTNFTIDAVGGNDGSSSEGSVYLYAGYYYPMKIVFSNAEESATLPVAVS
YJM1443  LPNGTSVMDDFSGYVYSFENNPAQDDCTITNPASHTIDALARTRTHAWTGTYTTTVVQTS
YJM1443  TYTGLNGLTTEETVFEIQTPTATSSSMITNTYSSASTPTAANITTSTSTSSEIVTSSSIS
YJM1443  ASTSSEIVTSSSTSASTSSEIVTSSSTSTSASSEMATSSSTSTLTSSYSTSSSQNEVISS
YJM1443  STSSSNLTTTSSISSSIPTSSVITSSSQSEETSASTTSIDSTSSSDIASSTQSVSITSTF
YJM1443  TSDLTTSTSASSTVPTSSVITSSNQNEVISSSTSSSNLTTSSSISSSIPTSSAITSSSQS
YJM1443  EETSASTTSIDSTSSSDIASSTQSVSITSTFTSDLTTSTSASSTVPTSSFITSSSESGET
YJM1443  SASTTSINSTSSSIISSSTESVSVTSTVSFKQTSSSSSSTYSTEKEVTSSLISSTSTSAI
YJM1443  TSSSAQSLTTSSFSSFLHSITPIYPSNETVVTSSSFSTTSGSSNPVITSTSSNRGSSTLD
YJM1443  ADARTASTSSGELSSSFSTTTGIDSIQSLIASTSSSVSASTTLDDSTSSASASRTYLTIS
YJM1443  SSSGSLNMPNASSSSTISSRHQETELSSTRSFTSSGSIKFSESSGSSAISSTASSTEVFS
YJM1443  SQTSIISSSDSNSKNYEFSSTPASSRSFKASSSLSISPSTTSSTTSSTFRNQTETISFTT
YJM1443  EIKSSMNESFTSISQSQTIDDKITSTASHASTTVPSLAPVATEKREQTTFVTITSCGSNI
YJM1443  CSKTVSPALISTSTTTVKGVATEYTTWCPISSSKSTRQTTLVTVTSCESGICSETASPAI
YJM1443  ISTATTTINDAVTVYTTWCSLTANDKGDTVEMNTSVGSTPAILEGSQTETVTKTNEKYSI
YJM1443  SESETHLPTATQNIIHSNGASSAPDTSKVEAIASTYLTTISQQPRNTYTNSVLTSATASV
YJM1443  EISSYIGVANNLLANSAISIFIASLLLAIV*
```

B)

Smith-Waterman score: 2331; 39.9% identity (68.4% similar) in 1217 aa overlap (8-1170:7-1171)

```
//
       120       130       140       150       160       170
FLO10  LCSDNTTLSSKTEKRENDDCDQGAAYWSSDLFGFYTTPTNVTVEMTGYFLPPKTGTYTFG
        :.       .:.::... :: ..:::::::::::::::::::::::::: :..:.:::.
FLO12_ PCGP----LYETKKRDTEYCDPNTAYWSSDLFGFYTTPTNVTVEMTGYFLAPENGSYTFS
        110           120       130       140       150       160

       180       190       200       210       220       230
FLO10  FATVDDSAILSVGGNVAFECCKQEQPPITSTDFTINGIKPWNADAPTDIKGSTYMYAGYY
       : :::::::::.::. :: ::.:.:  .:::.:::...   ...  .. .::.:.:::::
FLO12_ FYTVDDSAILSIGGDNAFACCEQQQSSVTSTNFTIDAV---GGNDGSSSEGSVYLYAGYY
            170       180       190       200          210

       240       250       260       270       280       290
FLO10  YPIKIVYSNAVSWGTLPVSVVLPDGTEVNDDFEGYVFSFDDNATQAHCSVPNPAEH----
       ::.:::.:::   .::::.: ::.:: : ::: :::.::..: .:  :.. ::: :
FLO12_ YPMKIVFSNAEESATLPVAVSLPNGTSVMDDFSGYVYSFENNPAQDDCTITNPASHTIDA
     220       230       240       250       260       270

//
       970       980       990      1000      1010      1020
FLO10  GITTEYTTWCPLSATELTTVSKLESEEKTTLITVTSCESGVCSETASPAIVSTATATVND
       :..::::::::.:...        : ..:::.::::::::.:::::::::.::::.:.::
FLO12_ GVATEYTTWCPISSSK--------STRQTTLVTVTSCESGICSETASPAIISTATTTIND
      980       990              1000      1010      1020      1030

      1030      1040      1050      1060      1070      1080
FLO10  VVTVYSTWSPQATNKLAVSSDIENSASKASFVSEAAETKSISRNNNFVPTSGTTSIETHT
       .::::.::   ..:  . . ....:....  . :...:......:.    : .   :::
FLO12_ AVTVYTTWCSLTANDKGDTVEMNTSVGSTPAILEGSQTETVTKTNEKYSISES---ETHL
             1040      1050      1060      1070      1080

       1090      1100      1110      1120      1130      1140
FLO10  -TTTSNASENSDNVSASEAVSSKSVTNPVLISVSQQPRGTPASSMIGSSTASLEMSSYLG
        :.:.:  ...   :: .. . .....  : ..:::::.: ..:.. :.:::.:.:::.:
FLO12_ PTATQNIIHSNGASSAPDTSKVEAIASTYLTTISQQPRNTYTNSVLTSATASVEISSYIG
      1090      1100      1110      1120      1130      1140

       1150      1160      1170
FLO10  IANHLLTNSGISIFIASLLLAIV*
       .::.::.::.::::::::::::::
FLO12_ VANNLLANSAISIFIASLLLAIV*
      1150      1160      1170
```

Figure S2 Smith-Waterman alignment of alleles of *YDR534C* (*FIT1*) from S288c and YJM248

Smith-Waterman score: 1282; 52.7% identity (66.8% similar) in 539 aa overlap (1-449:1-529)

```
               10        20        30        40        50        60
YJM248 MRFSSALALSAITVAALGESITTTITATKNGHVYTKTVTQDATFVWTGEGERAPAATSAV
       :..:::..::::::::::::::::::::::::::::::::::::::.:::    :.::::
S288c  MKLSSAFVLSAITVAALGESITTTITATKNGHVYTKTVTQDATFVWAGEGA---AVTSAV
               10        20        30        40        50

               70
YJM248 TSAVSSAANSAV------------------------------------------------
       : : . ::.::.
S288c  TEASTVAATSAAAETSVAAETSIVEPSTSAQGTSADEGSGSSITTTITATKNGHVYTKTV
        60        70        80        90       100       110

                                                  80        90
YJM248 ------------------------------SSAAEPS-------ADEGSGSSITTTITAT
                                     .:.::::       ::::::::::::::::
S288c  TQDATFVWTGEGERAPASTVATVETSVAAETSVAEPSTSAQGTSADEGSGSSITTTITAT
       120       130       140       150       160       170

         100       110       120        130         140       150
YJM248 KNGHVYTKTVTQDATFVWTGEGERAPAATSA-VTSAA--ITSAAEPSASAEGTTTDEGSG
       ::::::::::::::::::::::::::..: : : .::  .::.::::::     ::::::
S288c  KNGHVYTKTVTQDATFVWTGEGERAPVSTVATVETAASPVTSVAEPSAS-----TDEGSG
       180       190       200       210       220            230

            160       170       180       190
YJM248 SSITTTITATKNGHVYTKTVTQEATFVWTGEGERAPAST-----------------AVIS
       ::::::::::::::::::::::.::::::::::::::::                 ...
S288c  SSITTTITATKNGHVYTKTVTQDATFVWTGEGERAPASTVATSSISAIEIPSTTEASIVE
            240       250       260       270       280       290

         200            210        220              230       240
YJM248 GTSTLETSS-----SAPEISSA-ETSSAVQTSSTAEMSST-------ETSSAATISSALQ
       ..:..::::     :: : ::: ::::::.:::.:: ::.       :::::. ::::..
S288c  ASSAVETSSAAETSSAVETSSAVETSSAVETSSAAETSSAAETSSAVETSSAVEISSAVE
            300       310       320       330       340       350

            250                    260                          270
YJM248 SSASYTRASSAV-------------SKISSTSKASSAV-------------------SKI
       .::  : .::..             ...::. .::: .                   ..
S288c  TSAVETSSSSSTIETTSVKSLSPTQTSLSSSVQASSPIETSSAAKTSSVVPTFSSTTTEN
            360       370       380       390       400       410

              280       290       300       310       320       330
YJM248 SSASKASSAVEASTTTSSKSSITTVVTATKNGQARTKSVTQDAEFVHTGEGSNGGASGSF
       :: ::..::: :::::::.:: .:.:: :. ::: :.: ..::. :.: : ::. .:.:
S288c  SSNSKSTSAVVASTTTSSESS-ATIVTPTRIGQAYTESSSRDAQSVRTHE-SNNWSSSSS
            420       430        440       450       460        470

              340       350       360       370       380
YJM248 ASSKAISSIPKVPTTTVETLTNGKSSTTAQVVNYTGAADSIAAGTGLMGAALAAVIFL*
       ::.: .::: .: :::.  .:::::::: :.::::::::::::::::::::::::::::
S288c  ASTKMVSSITRVQTTTAGIFTNGKSSTTPQIVNYTGAADSIAAGTGLMGAALAAVIFL*
              480       490       500       510       520
```

Figure S3 Smith-Waterman alignment of alleles of *YBR188C* (*NTC20*) from strains S288c and YJM1400.

Smith-Waterman score: 293; 38.3% identity (70.9% similar) in 141 aa overlap (1-131:1-138)

```
                10        20        30        40        50        60
S288c   MPSLRDLSLERDQELNQLRARINQLGKTGKEEANDFVGLNISNEPVYDTVIQTGQSSNAT
        ::.:..:.:::.:..:.::.::....: .  . .     .: .. . . .. : :.:..
YJM1400 MPTLQNLALERNQKINELRGRIDKVNKRNDTKYSRRSYSEIPDKSLNNLAFGTVQGSGSL
                10        20        30        40        50        60

                70        80        90        100       110
S288c   NSFVQETIQKTKQKESGQPYIIPQKNEHQRY-IDKVCETSDLKAKLAPIMEVLEKKTNEK
         :. :: :.. .          :.::..:.   .:.:::::::::.::  :.:::.::.:
YJM1400 YSLEQEIIHEEE----------PKKNQYQQPGANKICETSDLKAKFAPYTEILEKETNKK
                70                  80        90       100       110

      120         130       140
S288c   IKGIIRKRVL--QEPDRDNDDSG*
        :: ::::..:    :. ...:
YJM1400 IKEIIRKKMLFLTYPNDSSSD*
               120       130
```

Figure S4 *ABM1*

*ABM1* and related duplicate genes *YIL014C-A* and *YIL102C* sharing some sequence similarity and appear to be related genes. *YIL102C* contains two stop codons in 94/94 strains, and in all strains in the 1011 strains (Peter et al. 2018), but has the full-length open reading frame in *S. paradoxus* strain UWOPS91. The related open reading frame found in some strains on chromosome I has a valine at the location where a start codon is conserved in other strains. This valine is conserved in all 94 strains in this gene at this location and in the 1011 strain dataset.

```
                                         ..:.: ::   *  :   * : .     :  :       .** **  .*
S. paradoxus UWOPS91 chrX ABM1      MSCSLGISTI LSPTDASMVA TWLNSTESNT YVHYSTCWCT SDYTLNISVI
YJM1418 chrX ABM1                   MSCSLGVSTI LSPSDASMVA AWLNSTESNT YIHYSTCWGT SDYTLNISVI
YIL102C                             MKYSCSITTI MSNGSASLLM NWEELTPGHC FTSYTTNPIA GDYGLNASAI
YIL014C-A                           MNYP.SITTL MSNESANLLI IWGNATPDIS YLSYTTNPML GDYVLNVSAI
S. paradoxus N44 chrIX              MKYPCSITTI MSNDSANLLM NWEELTPGNC YISYTTNPML GDYVLNVSAI
YJM1356 chrI                        VKYPGDFTTI MADETAKTLT KWLDSAADNC YFRYTTTPAL GDYNLNVLAI
S. paradoxus YPS138 chrI            MKYPGDFTTI MADETAKTLT KWLDSAADNC YFRYTTTPAL GDYNLNVLAI
YEL073C                                MVNLANV LTNATAATLS AWSNTVPLET YFHFDEASGF GDYYLNVSVI
S. paradoxus N44 chrXVI                MVNLANV LTNATAATLS AWSNTVPLET YFHFDEASGF GDYHLNVSVI

                                        * *  : :   :: *.    :         .    .
S. paradoxus UWOPS91 chrX ABM1      DATTEKLVDT RLFTTLENAT AWINSNSIDE DEDDMAHATD VADRLDGLSL
YJM1418 chrX ABM1                   DATTEKLVDT RLLTTLENAT AWINSNSIDE DEDDMPHATN VADRLDGLSL
YIL102C                             DGHTEELVAT HPAGTLENAT Q*VHSIRK*W HSLTFADENG TILPQSCSFF
YIL014C-A                           NGCTEELIAT HLVPTLENAT QWVYDAGEYW DNYSFTDES. TPLPGLSWPF
S. paradoxus N44 chrIX              NGCTEELVAT HLVPTLENAT QWVHEMGEYW DSLTFVDENG TSLPEYYYFF
YJM1356 chrI                        NADTETLVEA KSVPTLKNAK QWVNDMEVYW RQFDYDEKSD ISSPNPP...
S. paradoxus YPS138 chrI            NADTETLVEA KSVPTLKNAK QWVNDMEVYW RQFDYDEKSD ISSPNPP...
YEL073C                             WMN.ETLYET RIVPAIINVR EWLDHMEAND PSPSVTNPYE TSGYYAFSTV
S. paradoxus N44 chrXVI.            WMN.ETLYET RIVPAIINVR EWLDDMEAND PSPSITNPYE TSGYYAFSTV

S. paradoxus UWOPS91 chrX ABM1      SKRIYGISPC GF*
YJM1418 chrX ABM1                   SKRVYSICHY EF*
YIL102C                             TENEEH*
YIL014C-A                           NE*
S. paradoxus N44 chrIX.             QENEEQ*
YJM1356 chrI                        DSN*
S. paradoxus YPS138 chrI            DSN*
YEL073C                             VPVLMGNMK VA*
S. paradoxus N44 chrXVI.            VPVLMGNMK VA*
```

Figure S5 Gene Remnant REMYDR439C-A.

A) Gene order in the region of REMYDR439C-A. “X” indicates WGD paralogs lost in *S. cerevisiae.* The pre-WGD gene order in *A. gossypii* is shown to confirm the pre-WGD gene order. Red and blue arrows indicate relative orientation of gene on chromosome.

B) Alignment of a portion of REMYDR439C-A with S288c paralog *YML020W* and *S. paradoxus* (S.p) *YDR439C-A* shows sequence conservation. Stop codons shown as “*”.

A)

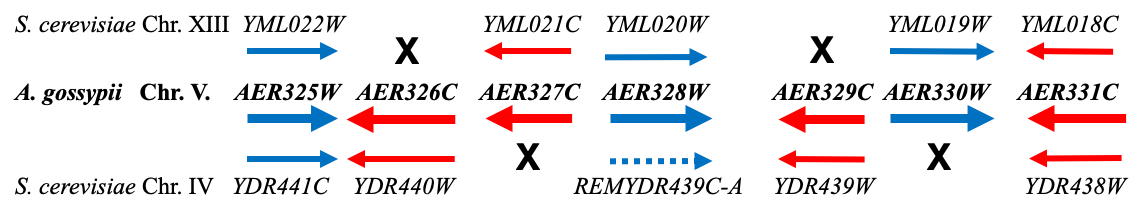

B)

```
S.c REMYDR439C-A  TVLNKLDVLVNRGVKISKIGSLNDQVMLFILHRGL*CNIQILFRALYVPGG*LQYPIFLLHVIKLSVN*LI*NTLTVPR*RN*LNFLWDQF*VLV-SIFYEDNKVY
                  + +  L  +++  VKI+ +GS+NDQ++       L  N   +FRA+++     Q P F+  ++K++   L                L         S  Y + +VY
S288c YML020W     SFIEGLRTIISNNVKITFVGSINDQLVPLYSSTCLFANHPNIFRAIFI-DRG-QTPAFITRIVKIAGSLLNLGYNDHGIIKEISGSLAGTLTGGGHSTIYNEKQVY
                   F+ GL  +++ +VKI+ +GS+NDQ+VPLYSS C+F  HPNIFRA+++ D+G Q P F+  +++I+  L+NLGY+DH +IKE+S    G +   GHS +Y + +VY
S.p YDR439C-A     QFLTGLNVLVNADVKISMIGSLNDQVVPLYSSLCIFLQHPNIFRALYVPDKGRQYPVFLVHVIEISVKLINLGYSDHAMIKELSQFFMGPILSDGHSIVYEDSKVY
```

Figure S6 Gene Remnant REMYFL041W-A.

A) Gene order in the region of REMYFL041W-A. “X” indicates WGD genes lost in *S. cerevisiae.* Red and Blue shows the relative orientation of the genes and gene remnant. *A. gossypii* is representative of the pre-WGD gene order.

B) Smith-Waterman alignment with the intact *S. paradoxus* strain CBS432 *YFL041W-A* gene on top, S288c REMYFL041W-A bottom. Bases underlined are the currently annotated dubious hypothetical gene *YFL041W-A*. Bold indicates the dubious hypothetical gene *610-snap_masked-2999-BGP_1*. F1, F2, F2 stand for reading frames 1,2,3.

C) Mafft Alignment of fragment of dubious gene *YFL041W-A* and the equivalent region from *YLR070C* from *S. cerevisiae* and *S. paradoxus* strain CBS432 gene *YFL041W-A*, and Dubious gene *YFL041W-A* appears to be a fragment of a lost gene, a duplicate copy of *YLR070C* that resulted from the WGD. Selected conserved residues shown in bold.

D) Mafft alignment of *YLR070C* orthologs from *S. cerevisiae* (SCER), *S. paradoxus* (SPAR), *S. uvarum* (SUVA), *S. eubayanus* (SEUB), and REMYLR070C from *S. kudriavzevii* (SKDI). These genes and the gene remnant are all at the same location and orientation relative to flanking genes. The *S. kudriavzevii* gene remnant contains nine frame shifts marked by “/” and six internal stop codons though still retains extensive protein sequence conservation.

E) Smith-Waterman alignment of *S. cerevisiae* (SCER) *YLR070C* with *S. mikitae* (SMIK) REMYLR070C. Most of this gene has been deleted in this species, with a small portion of N-terminal end homology left. This gene remnant is at the same location and orientation relative to flanking genes that *YLR070C* is in *S. cerevisiae.*

F) Smith-Waterman alignment of *S. cerevisiae* (SCER) *YLR070C* with *S. arboricola* (SARB) REMYLR070C. Most of this gene has been deleted in this species, with a small portion of N-terminal end homology left. This gene remnant is at the same location and orientation relative to flanking genes that *YLR070C* is in *S. cerevisiae.*

G) Partial alignment of intact genes *YFL041W-A* from *S. paradoxus* (SPAR) and *S. arboricola* (SARB) and the *S. cerevisiae (*SCER) gene remnant REMYFL041W-A.

A)

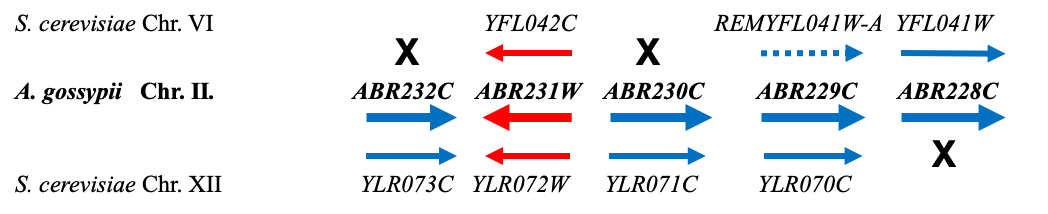

B)

```
Spara  ATGCAGAAATCTATCAGACTTATAGAGCCGAAGAAGATAGAAATTCAACAAGTTCCCAGG
F1     M  Q  K  S  I  R  L  I  E  P  K  K  I  E  I  Q  Q  V  P  R
       ::::::::: :::::: :  : : :: :: :: : :: ::: :::::  ::::: ::: :
S288c  ATGCAGAAAGCTATCAAATGTTTTGAACCAAACATGAAAGAGATTCAGAAAGTTGCCAAG
F1     M  Q  K  A  I  K  C  F  E  P  N  M  K  E  I  Q  K  V  A  K

Spara  CCCAAGGTCACGACGCCATATCAGGTTAAAGTGCAAGTGAAGGCCACTGGGATCTGCGGA
F1     P  K  V  T  T  P  Y  Q  V  K  V  Q  V  K  A  T  G  I  C  G
       :: :: ::   :: : ::   ::::: :::::  :::::::::: : ::: :::::::::
S288c  CC-AACGT---GATGTCA---CAGGTCAAAGTTTAAGTGAAGGCTATTGGAATCTGCGGA
F1     P  fs
F3        N  V     M  S     Q  V  K  V  *  V  K  A  I  G  I  C  G

Spara  TCAGACATTCACTACTACACAGATGGCCAGGTCGGTGGAAACCGACTTGAGAAGCCCATG
F1     S  D  I  H  Y  Y  T  D  G  Q  V  G  G  N  R  L  E  K  P  M
       :::::::::::::::::::: :::::::::::: : : :::   :: :: ::::::::::
S288c  TCAGACATTCACTACTACACGGATGGCCAGGTCAGCGTAAAGTAACCTGGGAAGCCCATG
F3     S  D  I  H  Y  Y  T  D  G  Q  V  S  V  K  *  P  G  K  P  M

Spara  ACTTTGGGCCACGAATCCAGCGGTGTTGTTATCGAGGTTGGTAATAAAGTGTCTCCAAAC
F1     T  L  G  H  E  S  S  G  V  V  I  E  V  G  N  K  V  S  P  N
       ::::::::::::::::::: ::::: :::: : ::  :::      :::: :::: ::::
S288c  ACTTTGGGCCACGAATCCAACGGTGCTGTTCTTGAAATTG------AAGTATCTC-AAAC
F2                                                          fs  N
F3     T  L  G  H  E  S  N  G  A  V  L  E  I  E        V  S  Q

Spara  TACTGTAAAGTGGGTGATGCAGTAAGCATAGAGCCCTCGTATCCTAATAGGTATACGAAA
F1     Y  C  K  V  G  D  A  V  S  I  E  P  S  Y  P  N  R  Y  T  K
       :::::::::: ::::::::::::::::::: ::::::: :: :: ::::::: ::: :::
S288c  TACTGTAAAGCGGGTGATGCAGTAAGCATATAGCCCTCATACCCAAATAGGTTTACCAAA
F2     Y  C  K  A  G  D  A  V  S  I  *  P  S  Y  P  N  R  F  T  K

Spara  GAGTTTCGGTCGGGCCACTATAATTTGGAGCCCAATATGCCATTTGCTGCGTCACCGCCC
F1     E  F  R  S  G  H  Y  N  L  E  P  N  M  P  F  A  A  S  P  P
       ::::::: ::  ::::: :::: :::: ::: ::: :: ::::::::::: :::: : ::
S288c  GAGTTTCTGTGAGGCCATTATAGTTTGCAGCTCAACATACCATTTGCTGCATCACTGACC
F2     E  F  L  *  G  H  Y  S  L  Q  L  N  I  P  F  A  A  S  L  T

Spara  TGTGACGGGACCCTTTGCGAAGAGTTTGTAGTTGACATGGACTACGTTTACAGATTACCA
F1     C  D  G  T  L  C  E  E  F  V  V  D  M  D  Y  V  Y  R  L  P
        ::::::: ::::::::  ::::::: :: ::: :: : :::: ::::::: :::::::
S288c  -GTGACGGCACCCTTTGTCAAGAGTTCGTGGTTAACGTCGACTGCGTTTACTGATTACCG
F1   fs   D  G  T  L  C  Q  E  F  V  V  N  V  D  C  V  Y  *  L  P

Spara  CCACATGTGAAATTTGAGCAAGGTGCTTTGATTGAGCCATTATCCGTCGCTGTTCATGCG
F1     P  H  V  K  F  E  Q  G  A  L  I  E  P  L  S  V  A  V  H  A
         :::    :  : :    :  :: :: ::     :::: :::  :::::: : ::::
S288c  TTACAGTGAATTTCTAGCAAGAGTCCTCTGTAGATGCCACTATTTGTCGCTCTCCATGTA
F1     L  Q  *  I  S  S  K  S  P  L  *  M  P  L  F  V  A  L  H  V

Spara  GTAAGGTTGTCCCAGGTGAAGTGTGGGGACCGCGCCGTAGTTATAGGTGTTGGAACTATT
F1     V  R  L  S  Q  V  K  C  G  D  R  A  V  V  I  G  V  G  T  I
       ::::::  :::: :: :::::::::::::    :: ::::::::::::: :::::: :::
S288c  GTAAGGCCGTCCTAGCTGAAGTGTGGGGAGTAAGCTGTAGTTATAGGTGCTGGAACAATT
F1     V  R  P  S  *  L  K  C  G  E  *  A  V  V  I  G  A  G  T  I

Spara  GGTCTACTGGTGGGTGCGGTGCTTAGAGAGTTCGGTGTCAACAAGTTGGTGTTCATGAGC
F1     G  L  L  V  G  A  V  L  R  E  F  G  V  N  K  L  V  F  M  S
       : ::: :::: ::::::::: :: :::: :::              :::::::::::::
S288c  GATCTGCTGGCGGGTGCGGTACTCAGAGGGTT--------------TGGTGTTCATGAGT
F1     D  L  L  A  G  A  V  L  R  G  F
F2                                           fs        V  F  M  S

Spara  ACACACGACTCCAAGAAAGAGCAAAGTGAGCAGTTTGGTGGTACCGGGTTTATCAACTCT
F1     T  H  D  S  K  K  E  Q  S  E  Q  F  G  G  T  G  F  I  N  S
       ::::: :: :::::::::::::::: ::  ::::::               :: ::: ::
S288c  ACACATGATTCCAAGAAAGAGCAAAATGGACAGTTT---------------ATAAACGCT
F2     T  H  D  S  K  K  E  Q  N  G  Q  F                 I  N  A

Spara  ACGAAGGATAGTTATGACCAGGAGAAGGAAGGGCTGGTGGATGTTGTATTCGAATGCTCA
F1     T  K  D  S  Y  D  Q  E  K  E  G  L  V  D  V  V  F  E  C  S
       : :::: :  : ::: : ::: : ::  :::    :  :  :: :: :::::: : ::::
S288c  ATGAAGAACTGCTATAATCAGAAAAA--AAG----GAAGCCTGATGGATTCGAGTCCTCA
F2     M  K  N  C  Y  N  Q  K  K    R      K  P  D  G  F  E  S  S

Spara  GGTGCTCAAAAGAGTGTTGAGAATGCACTGAAATGTGTCAAGAACGGCGGAACTGTTATG
F1     G  A  Q  K  S  V  E  N  A  L  K  C  V  K  N  G  G  T  V  M
       ::: :::::::::: :::::::::   :: ::::::::::::::::: ::::: :::: :
S288c  GGTACTCAAAAGAGCGTTGAGAATATGCTAAAATGTGTCAAGAACGGTGGAACAGTTAAG
F2     G  T  Q  K  S  V  E  N  M  L  K  C  V  K  N  G  G  T  V  K

Spara  CAGATTGGTATGCATCCCTCAAATGAAAATGTAGGCATTCCTCTTAACGAGATTTCCTCT
F1     Q  I  G  M  H  P  S  N  E  N  V  G  I  P  L  N  E  I  S  S
        :: ::: ::: :: ::::: ::::::::::: :: :  ::::  ::::::::: :::
S288c  TAGGTTGATATCCACCCCTCGAATGAAAATGTGGGGAC-CCTCCCAACGAGATT-CCTTC
F1                                        fs  P  P  N  E  I
F2     *  V  D  I  H  P  S  N  E  N  V  G  T
F3                                                        fs    F

Spara  AGAGAGATTACGTATAAAGGGTCTTTTAGGTATTACGAAGGTGACTACTCACTAGCCATT
F1     R  E  I  T  Y  K  G  S  F  R  Y  Y  E  G  D  Y  S  L  A  I
       ::::::::::: ::::::: :   :: : :::: :::: :::::::::::  : ::::::
S288c  AGAGAGATTACATATAAAGTGAAATTCAAGTATGACGATGGTGACTACTCCTTGGCCATT
F3     R  E  I  T  Y  K  V  K  F  K  Y  D  D  G  D  Y  S  L  A  I

Spara  GATTTAATGTCCAGAGGTTGTATCGATGTCGATGAGTTGATTACTGATAGGTTCCCCTTT
F1     D  L  M  S  R  G  C  I  D  V  D  E  L  I  T  D  R  F  P  F
       ::::: ::::: :: : ::::::  ::::: ::::::::::::: :::::: : :   ::
S288c  GATTTGATGTCTAGGGATTGTATTAATGTCTATGAGTTGATTACCGATAGGCTACTGGTT
F3     D  L  M  S  R  D  C  I  N  V  Y  E  L  I  T  D  R  L  L  V

Spara  GAGAAAGCAGCCGATGCGTTCGCATTTGCAGTAGAGAACTCT-GCTAAGGTAGGTAAGGT
F1     E  K  A  A  D  A  F  A  F  A  V  E  N  S   A  K  V  G  K  V
       ::            : : :::  :   :: :       :::  :::::: ::::::::::
S288c  GATT----------TCCTTTCAAAAAAGCTG-------CTCAAGCTAAGATAGGTAAGGT
F2             fs     P  F  K  K  A  A         Q  A  K  I  G  K  V
F3     D  F

Spara  CATTATAGAACATTAA
F1      I  I  E  H  *
         ::::::::  :
S288c  GTTTATAGAATTTATCGAACTGTATCTAAATATTTTTAATATTCTTGTTTTATTACTTTG
F2      F  I  E  F  I  E  L  Y  L  N  I  F  N  I  L  V  L  L  L  C

S288c  TGGCTTTTGTATTCAGTCTGCACCCTGTTTCGCCATCGATTCTTATAATACATACATTTT
F2      G  F  C  I  Q  S  A  P  C  F  A  I  D  S  Y  N  T  Y  I  F

S288c  TATATAA
F2      I  *
```

C)

```
S288c REMYFL041W-A            MKMWGPREITYKVKFKYDDGDYSLAIDLMSRDCINVYELITDRLLVD
S288c YLR070C                 ISIIPTKELTFQGCFRYCQGDYSDSIELVSSRKLSLKPFITHRYSFK
S. paradoxus CBS432 chrVI     LNEISSREITYKGSFRYYEGDYSLAIDLMSRGCIDVDELITDRFPFE
S. uvarum                     ISKIPAKEITFQGSFRYCQGDYSDSIELVSSGKLSLKPLITRRYQFK
A. gossypii ABR229            LAEATVKQLRILGSFRYAAGDYATAVELLASKRVDAARLVTHRFPFD
K. lactis                     IAMIGAKELRVLGSFRYYFNDYKIAVKLISEGKVNVKKMITHTFKFE
Z. rouxii                     ISAVGSKEITFHGCFRYKKGDFADSVALLSSGRINGKPLISHRFAFD
```

D)

```
SCER_YLR070C    -MTDLTTQEAIV----LERPGKITLTNVSIPKISDPNEVIIQIKATGICGSDIHYYTHGR
SPAR_YLR070C    -MTESITQEAVV----LEQPGKITLMDVSIPTISDPNDVIIQIKATGICGSDIHYYTHGK
SUVA_YLR070C    MLTELTTQEAVI----LEQPGKIVLTNVNIPKISDANDVIVQIKATGICGSDIHYYTHGR
SEUB_YLR070C    -MTELTTQQAVV----LEQPGKIVLTNVSIPKISDANEVIIQIKATGICGSDIHYYTHGR
SKDI_YLR070C    -MTE*TT*EAIIHQSDLE---KIVLTDVNISNISVPNRTIIQIGATRMYGSGIHHYTHGI
                 :*: ** :*::    **   **.* :*.*..** .* .*:** ** : **.**:****

SCER_YLR070C    IANYVVESPMVLGHESS GIVALIGENVKTLKVG DRVALEPGIPDRFSPEMKE-GRYN LDP
SPAR_YLR070C    IANYVVKSPMVLGHESS GIVALIGENVKTLKVG DRVALEPGLPDRFSPEMRE-GRYN LDP
SUVA_YLR070C    ISHFVVKSPMVLGHESS GVVVLIGDNVKTLKVG DRVALEPGFPNRFSSELKE-GRYN LDP
SEUB_YLR070C    ISHFVVKSPMVLGHESS GVVALIGDNVKTLKVG DRVALEPGFPNRFSSKLKE-GRYN LDP
SKDI_YLR070C    ISHFVVKLLTVLARNSC/GIAAFTCHHVKILGRG/ERVVLKPCYLTNFLPELKESGHFG/LDP
                *:::**:   **.::*. *:..:  .:** *  * :**.*:*    .* .:::* *::. ***

SCER_YLR070C    NLKFAATP---PFDGTLTKYYKTMKDFVYKLPDDVSFEEGALIEPLSVAIHANKLAKIKF
SPAR_YLR070C    NLKFAATP---PFDGTLTKYYKTAKDFVYKLPDHLSFEEGALMEPLSVAVHANKIAKMKF
SUVA_YLR070C    DLEFAATP---PYDGTLTKYYRAPKDFVYKIPDHVSFEEGALMEPLSVAVHANRLAKMRF
SEUB_YLR070C    DLKFAATP---PYDGTLTKYYRAPKDFVYKIPDHVSFEEGALMEPLSVAVHANKLAKMRF
SKDI_YLR070C    DSKFAGLLRLYPLGGILTKCYTAVKDLFYKLSDEVIFGKRALVEPLSVVVNASKTAKTIS
                : :**.     * .* *** * : **:.**:.*.: * : **:*****.::*.: **

SCER_YLR070C    GARCVVFGAGP IGLLAGKV ASVFGAADVVF VDLLENKLETARQFGATHIVNSGDLPHGVT
SPAR_YLR070C    GARCVVFGAGP IGLLAGKV ASVFGATDVVF VDLSETKLERASQFGATHVVNSGGLPHGVT
SUVA_YLR070C    GARCVVFGAGP VGLLTGKV ASIFGATDVVF VDLSETKLERAKQFGATHMVNSGDLPPGVS
SEUB_YLR070C    GARCVVFGAGP VGLLTGKV ASVFGATEVVF VDLSETKLERAKQFGATHTVNSGDLPPGVS
SKDI_YLR070C    GSHCVVFGTGF/VALLTRKV/ASEFGATEVVW/VDLSKTNLQGVEHFGSSTLIDH--VCSEVT
                *::*****:*  :.**: ** ** ***::**: *** :.:*: . :**::  ::   :   *:

SCER_YLR070C    VDSVIKKAIGKKGAD VVFECSGAEPCVRAGIEVCKAGGTIVQVGMGQEEIQF----PIS IIP
SPAR_YLR070C    VDSVIKKEIGKKGAD IIFECSGAQACIRTGIEVCKAGGTFIQVGMGPEEIQF----PIS MIP
SUVA_YLR070C    AETVIRKAIGKRGAD VVFECSGAQPCIRTGIEVCKAGGTYVQVGMGQEEIQF----PIS KIP
SEUB_YLR070C    VESVIRKAIGKKDAD VIFECSGAQPCIRTGIEVCKAGGTYVQVGMGQEEIQF----PIS KIS
SKDI_YLR070C    ADGAIRRAISKEGA-/IVFELSGAQPCVRTGIEVCKEGEMFVQVGMGKR*SGSWYQ*SLS/MI-
                .: .*:: *.*..*  ::** ***:.*:*:****** *   :***** .. ..    .:*  *

SCER_YLR070C    TKELTFQGCFRYCQGDYSDSIELVSSRKLSLKPFITHRYSF KDAV-EAFEETSHHPLNNIK
SPAR_YLR070C    TKELTFQGCLRYCQGDYSDSIELASSGKLSLKPLITHRYKF KDAI-DAFDENRDHPLNNVK
SUVA_YLR070C    TKEITFQGSFRYCQGDYSDSIELVSSGKLSLKPLITHRYQF KEAI-EAFDENRDHPLDNVK
SEUB_YLR070C    TKEITLQGSFRYCQGDYSDSIELVSTGKLSLKPLITHRYQF KEAI-EAFDENRDHPLDNVK
SKDI_YLR070C    --------SFRYCERDYSDPTNLVSGVGPSLTILITHHYQY/RSVQ*E*I-----HPLDNVK
                        .:***: ****. :*.*    **. :**::*.: :..  :       ***:*:*

SCER_YLR070C    TIIEGPE*
SPAR_YLR070C    TIIEGPA*
SUVA_YLR070C    TIIEGPE*
SEUB_YLR070C    TIIEGPE*
SKDI_YLR070C    TVIEGPEE
                *:****
```

E)

```
                   310       320       330       340       350
SCER_YLR070C CQGDYSDSIELVSSRKLSLKPFITHRYSFKDAVEAFEETSHHPLNNIKTIIEGPE*
                                           .:::::. .    :.: : ::: :
SMIK_YLR070C GYCFITA*NNRACSC*HPQDLRPNQISVRLEAVEAFNGNRD*SLGNTKIIIERP*WKGKS
```

F)

```
           290       300       310       320       330       340
SCER_YLR070C TKELTFQGCFRYCQGDYSDSIELVSSRKLSLKPFITHRYSFKDAVEAFEETS-HHPLN
                                 : :::  ::::::.:..::.:::: .::.... ..::.
SARB_YLR070C KKLYNIYINISINIKRHIHPIMLVSIGKLSLKPLIARRYKFKDAFKAFNDNKDYYPLD
```

G)

```
SPAR_YFL041W-A  VSIPTISDPNDVIIQIKATGICGSDIHYYTHGKIANYVVKSPMVLGHESSGIVALI
SARB_YFL041W-A  VLRPKITTPYQVKVQVKATGICGSDIHYYTDGQVHGNRLKKPMTLGHESSGVIIDI
SCER_YFL041W-A  ----------QVKV*VKAIGICGSDIHYYTDGQVSVK*PGKPMTLGHESNGAVLEI
                          :* : :** ***********.*::      .**.*****.* :  *
```

Figure S7 Gene Remnant REMYKL162C-A (REMPIR6)

A) Gene order in the region of REMYKL162C-A (REMPIR6). “[“ indicates translocation break point, where the translocation occurred after the WGD; YKL162C is not an ortholog of any gene in this region of the *A. gossypii* genome. “(..)” indicates an inverted region in the *A. gossypii* lineage. *AEL111C* is an ortholog of paralogs *YKL164C* *PIR1* and *YJL160C* *PIR5*, *AEL110W* is an ortholog of paralogs *YKL163W* *PIR2* and *YJL159W* *PIR2*, and *AEL103W* is an ortholog of paralogs REMYKL162C-A REMPIR6 and *YJL158C* *PIR4*. Due to translocation *YKL162C* is not an ortholog of *AEL104W*. Red and blue arrows indicate relative orientation of gene on chromosome.

B) Smith-Waterman alignment of *S. cerevisiae* gene remnant REMYKL162C-A (REMPIR6) gene from strain YJM1463 (bottom), with PIR6 from *S. mikatae* (S. mik) strain IFO1815 (top). Bold is area of out of frame overlap with dubious hypothetical gene *YKL162C-A*. F1, F2, and F3 refer to reading frames.

C) Alignment conserved region from 3' end of *S. cerevisiae* PIR genes. REMYKL162C-A (REMPIR6) from strain YJM1463.

A)

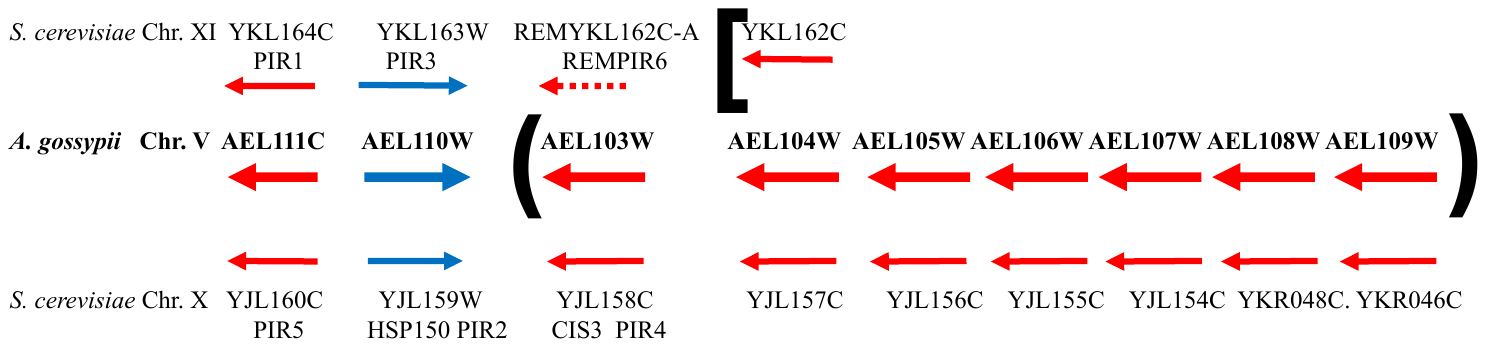

B)

```
S. mik ATGCAACTCAAAAAATTCTGGATAATTGCAGTCTCGCTAGCAGTTGTCATTGCCCAAGGG
F1     M  Q  L  K  K  F  W  I  I  A  V  S  L  A  V  V  I  A  Q  G
       ::: : : :::: :  :  :  : ::  :::: :: :::     : ::  :::::::  :
S288c  ATGTAGCCCAAATATATTGGTTTGATAACAGTTTCACTATACTGTTTC--TGCCCAACAG
F1     M  *  P  K  Y  I  G  L  I  T  V  S  L  Y  C  F
F2                                                   fs   A  Q  Q

S. mik TATACACCAGGTGAGCAATGGTCATCGTTAACTCCTTCAGCAACCTTTTCTTGTGCTGCT
F1     Y  T  P  G  E  Q  W  S  S  L  T  P  S  A  T  F  S  C  A  A
       ::::  ::: :::::: ::::::: : ::::::::::: :: ::    :: :: :  :::
S288c  TATATGCCAAGTGAGCCATGGTCAACCTTAACTCCTTCTGCTACTACCTCATGCGGCGCT
F2     Y  M  P  S  E  P  W  S  T  L  T  P  S  A  T  T  S  C  G  A

S. mik ACTGAGTTCACTTCTACTTTCGGTATCGCTGTTCAAACTATTTCAACCGATTGTATCAGT
F1     T  E  F  T  S  T  F  G  I  A  V  Q  T  I  S  T  D  C  I  S
       ::::: :: :::::    ::::::::: :::: ::::: ::: :::: :  : :: ::
S288c  ACTGAATTTACTTCCTTCTTCGGTATCACTGTCCAAACAATTACAACAGGATCTACCACG
F2     T  E  F  T  S  F  F  G  I  T  V  Q  T  I  T  T  G  S  T  T

S. mik GCTAATAAGAGAGATCTTATCTCTCAAATCGGTGATGGTCAAATTCAAGTCACTGCTGTC
F1     A  N  K  R  D  L  I  S  Q  I  G  D  G  Q  I  Q  V  T  A  V
       : :::   ::::::  :: : ::: : :::: ::: :: :::::: :: ::: :::: ::
S288c  GTTAA---GAGAGACGTTTTTTCTTAGATCGTTGACGGCCAAATTGAAATCATTGCTATC
F2     V  K     R  D  V  F  S  *  I  V  D  G  Q  I  E  I  I  A  I

S. mik CCTACTAAAGCCGG---ATTGGTTTTCGTTGGCGAAAG---TGTCGATATCACAACCACT
F1     P  T  K  A  G     L  V  F  V  G  E  S     V  D  I  T  T  T
       :: :: :::  : :   ::    :::  :: :  :::    :   :: ::::: ::::::
S288c  CCCACAAAAAACAGTGCATGTACTTTGATTAGTAAAAAAAATACTGA-ATCACGACCACT
F1                                                 fs  I  T  T  T
F2     P  T  K  N  S  A  C  T  L  I  S  K  K  N  T  E

S. mik TTAACTAGAACTTCTATTATTACAGTAAGAGAGAGAGCCACTACAACGTATGTTGGCTAC
F1     L  T  R  T  S  I  I  T  V  R  E  R  A  T  T  T  Y  V  G  Y
       ::::   :::: :  ::   :::  : : :::: :::::::  :::    ::::    ::
S288c  TTAATCGGAACCTACATAGATACGATTAAAGAGGGAGCCACAGCAATCACTGTTACTGAC
F1     L  I  G  T  Y  I  D  T  I  K  E  G  A  T  A  I  T  V  T  D

S. mik CAGGCTAGTTCTTTCAACAAAAACCCCGCATCTATCCTATCCCCATCACCGAGCCTTCCC
F1     Q  A  S  S  F  N  K  N  P  A  S  I  L  S  P  S  P  S  L  P
           ::: :: :: : : :::::   : : ::::   :  :   ::: :  ::  :: :
S288c  TTCTCTATTTATTCCGATAAAAAAAGCACCTCTACATTGCCGTTATCGCTTAGTATTTCT
F1     F  S  I  Y  S  D  K  K  S  T  S  T  L  P  L  S  L  S  I  S

S. mik TCTGTTATTTCCTTTATTTTTGCTTCA---AAATGCCAAACCACTGCCGTTGTTTCTTTA
F1     S  V  I  S  F  I  F  A  S     K  C  Q  T  T  A  V  V  S  L
       ::::  :: :: ::: : ::: :::     : ::: :::  :: ::: : :::::::  :
S288c  TCTGCCATCTCTTTTTTCTTTTCTTACGTTACATGTCAAGGCATTGCTGCTGTTTCTA-A
F1     S  A  I  S  F  F  F  S  Y  V  T  C  Q  G  I  A  A  V  S  K
F3                                                               fs

S. mik AAAAACGGTTCTTGTAGCAGTGATAGTACTTTGAAAGTAACTTTAATGGGCGGTATCTTG
F1     K  N  G  S  C  S  S  D  S  T  L  K  V  T  L  M  G  G  I  L
       :::::  : : : :::::::::: ::  :::::::::::::::: : : :::: :: :::
S288c  AAAAAAAGCTGTCGTAGCAGTGACAGCGCTTTGAAAGTAACTTTGAAGAGCGGCATTTTG
F3     K  K  S  C  R  S  S  D  S  A  L  K  V  T  L  K  S  G  I  L

S. mik ACTGACGGTAAGGGTAGAATCGGTTCTATTGTTGCCAACAGACAATTCCAATTCGATGGC
F1     T  D  G  K  G  R  I  G  S  I  V  A  N  R  Q  F  Q  F  D  G
       :: ::  : ::::: :: ::: :::::: ::::::::::::::::::::::::::::: :
S288c  ACAGATAGCAAGGGCAGCATCTGTTCTACTGTTGCCAACAGACAATTCCAATTCGATGAC
F3     T  D  S  K  G  S  I  C  S  T  V  A  N  R  Q  F  Q  F  D  D

S. mik CCTCCACCACAAGCTGGTGCCGTCTACGCGGCTGGC-TGGTCTATTACTCCAGAAGGTAA
F1     P  P  P  Q  A  G  A  V  Y  A  A  G   W  S  I  T  P  E  G  N
       ::::::::::::: :::::   :::: ::  ::::  : :::::: :: ::::::::  :
S288c  CCTCCACCACAAGTTGGTGTTATCTATGCTCCTGGGGTTGTCTATCACACCAGAAGGGGA
F1                                      fs  L  S  I  T  P  E  G  D
F3     P  P  P  Q  V  G  V  I  Y  A  P  G

S. mik CCTGGCCCTTGGTGATGAGGATGTCTTCTATCAATGCTTATCTGGCGATTTTTACAATTT
F1      L  A  L  G  D  E  D  V  F  Y  Q  C  L  S  G  D  F  Y  N  L
        ::: ::::: :::   : ::: :::: ::::::: ::  :   ::  :::::::::: :
S288c  TCTGCCCCTTAGTGGCCAAGATATCTTTTATCAATACTCGTTCAGCAGTTTTTACAATAT
F1      L  P  L  S  G  Q  D  I  F  Y  Q  Y  S  F  S  S  F  Y  N  I

S. mik GTACGACCAAAAAATTGGAGAGCAATGCAGCAGAATTCACTTACAAGCAGTTTCCTTGAT
F1      Y  D  Q  K  I  G  E  Q  C  S  R  I  H  L  Q  A  V  S  L  I
         ::::: :::  ::: :::: ::::: ::   : :  :: :::::::  :::::::: :
S288c  ACACGACGAAAGGATTAGAGAACAATGAAGTTAAGTCTACCTACAAGCTTTTTCCTTGGT
F1      H  D  E  R  I  R  E  Q  *  S  *  V  Y  L  Q  A  F  S  L  V

S. mik AAGTTGCTGA
        S  C  *
        ::::::: :
S288c  CAGTTGCTAA
F1      S  C  *
```

C)

```
YKL164C PIR1         LKGGILTDGK GRIGSIVANR QFQFDGPPPQ AGAIYAAGWS
YJL159W HSP150 PIR2  LKGGILTDGK GRIGSIVANR QFQFDGPPPQ AGAIYAAGWS
YKL163W PIR3         LSKGILTDRK GRIGSIVANR QFQFDGPPPQ AGAIYAAGWS
YJL158C CIS3 PIR4    LKDGVLTDAK GRIGSIVANR QFQFDGPPPQ AGAIYAAGWS
YJL160C PIR5         LQGGVLIDSS GRIGSIVANR QFQFDGPPPQ AGAIYAGGWS
REMYKL162C-A REMPIR6 LKSGILTDSK GSICSTVANR QFQFDDPPPQ VGVIYAPGVV
```

*. *:* * . * * * **** *****.**** .*.*** *

Figure S8 REMYCR099C (REMVTH4)

A) Alignment of *PEP1* (top) with REMYCR099C (REMVTH4) (bottom). Only the first 470 nucleotides of *PEP1*, out of 4740 nucleotides. There is week homology across the entire length of *PEP1*, except for from 2720 to 4535 in *PEP1* that is deleted from REMYCR099C (REMVTH4).

A)

```
PEP1   ATGATATTACTTCATTTTGTCTATTCTCTTTGGGCCTTACTTCTCATTCCTTTAACTAAT
F1     M  I  L  L  H  F  V  Y  S  L  W  A  L  L  L  I  P  L  T  N
       :::::::::::::::    : ::: :::::::::   :  : ::  ::::  ::   :::
S288c  ATGATATTACTTCATGCCATATATACTCTTTGGGTAATTATACTACTTCCGCTACTCAAT
F1     M  I  L  L  H  A  I  Y  T  L  W  V  I  I  L  L  P  L  L  N

PEP1   GCCGAAGAATTCACCCCCAAAGTGACAAAGACTATCGCGCAAGATTCATTTGATATATTA
F1     A  E  E  F  T  P  K  V  T  K  T  I  A  Q  D  S  F  D  I  L
       :: ::  ::::   ::: ::::: ::  :: ::       ::   :::::: :: :: :
S288c  GCAGAGAAATTTGTCCCAAAAGTAACGGAGGCTCCTATAGAAACATCATTTAATCTAGTG
F1     A  E  K  F  V  P  K  V  T  E  A  P  I  E  T  S  F  N  L  V

PEP1   AGCTTTGATGATTCCAACACTTTAATTAGAAAACAAGACACCTCCGTTACT---ATAAGC
F1     S  F  D  D  S  N  T  L  I  R  K  Q  D  T  S  V  T     I  S
       :: :::::::::::::::::::  :: :::  : : :        :::      :::::
S288c  AGTTTTGATGATTCCAACACTTCTATCAGATTAGATGGTTGGGGGGTTGTATGGATAAGT
F1     S  F  D  D  S  N  T  S  I  R  L  D  G  W  G  V  V  W  I  S

PEP1   TTTGACGATGGTGAAACATGGGAAAAAGTTGAAGGCATTGAAG-GCGAAATCACTTGGAT
F1     F  D  D  G  E  T  W  E  K  V  E  G  I  E  G   E  I  T  W  I
       :: :::: ::: ::::  :::::::  ::  :::  ::::::: ::: : :   :  :::
S288c  TTCGACGCTGGAGAAAATTGGGAAACGGTCAAAGAAATTGAAGAGCGCATT--TTCAGAT
F1     F  D  A  G  E  N  W  E  T  V  K  E  I  E  E
F2                                               fs  A  F    S  D
F3                                                       fs   Q  I

PEP1   ATACA-TCGACCCCTTCAATAGACATGACAGAGCCGTTGCAACGGCAA-TGAATGGGTCG
F1      Y  I   D  P  F  N  R  H  D  R  A  V  A  T  A  M   N  G  S
        :::  : :: :: ::: :: :::: :: ::::   : ::      :  :::::    :
S288c  TTACTGTTGATCCTTTCCATGGACAGGAAAGAGGTTTCGCTTTTATATGTGAATCACCCA
F1        fs   D  P  F  H  G  Q  E  R  G  F  A  F  I  C
F2                                                   fs   N  H  P
F3      Y  C

PEP1   TACCTTTATATAACCAATGATCAAGGTAAGTCATGGGAGCGTATAACGCTACCTGACTCC
F1     Y  L  Y  I  T  N  D  Q  G  K  S  W  E  R  I  T  L  P  D  S
        :  : :: :: ::: : :: :  ::  ::::::::  :  : ::::  ::::    ::
S288c  AAT-TCTACATTACCGACGACCGTGGGGAGTCATGGAGGGCTTTAACTATACCCTCATCA
F1           Y  I  T  D  D  R  G  E  S  W  R  A  L  T  I  P  S  S
F2     N

PEP1   GGAGAAAGTATTTCACCTCGTGAATGCTATATAGAAACCCATCCCTTGAAT--AAGAACT
F1     G  E  S  I  S  P  R  E  C  Y  I  E  T  H  P  L  N    K  N  Y
       : ::::  ::::: :  : : :: :: : ::::   :: ::::: : :::   :::::::
S288c  GAAGAA--TATTT-AGATGGCGACTGTTTTATAACTACTCATCC-TAGAAACAAAGAACT
F1     E  E           D  G  D  C  F  I  T  T  H  P
F2         fs   I  *  fs                                    K  N  F
F3                                               fs *  K  fs

PEP1   ATTTTCTTGCAAAGTGCAACTATTGTGAGAAAACAGAAGTAAACAATGACAATGAAGAGA
F1       F  L  A  K  C  N  Y  C  E  K  T  E  V  N  N  D  N  E  E  N
        : :: :::: :: :::   ::: :  : :  : ::: : : ::  :    ::::
S288c  -TCTTATTGCGAATTGC---TATAGCTATATGATAGACGCAGACGTTTTATATGAC----
         L  I  A  N  C     Y  S  Y  M  I  D  A  D  V  L  Y  D
```

Figure S9 REMYCL001W-A

A) Gene order in the region of REMYCL001W-A. “]” indicates translocation breakpoint. *S. cerevisiae* gene YNL002C does not have a homolog in this region. Red and blue arrows indicate relative orientation of gene on chromosome.

B) Smith-Waterman alignment of *YCL001W-A* from *S. paradoxus* strain CBS432 with S288c REMYCL001W-A. Underlined are the currently annotated *YCL001W-A* and *YCL001W-B.* Both of these dubious genes are fragments of the full length REMYCL001W-A. The final 13 amino acids of *YCL001W-B* is out of frame relative to the full length alleles due to a frame shift.

A)

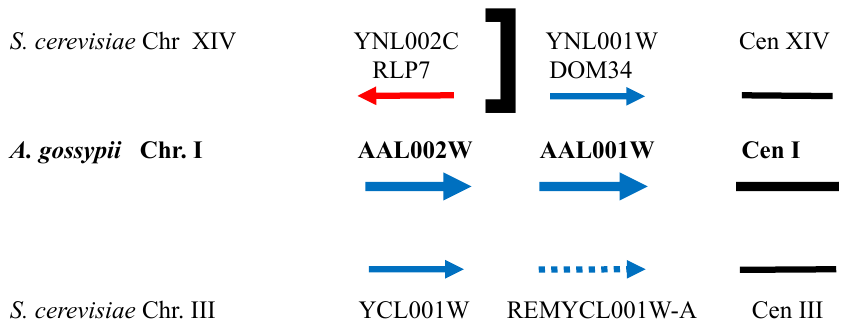

B)

```
Frame = +1
S. paradoxus MKIITVAEDSADNDNSIITLIPQNK---DDLFTIYQIIDKEDE----VIFKKLFTTKRDE
             MKII+VA+ +A      IT+   ++   +   T  Q I+   +     I +KLF TK++E
S288c        MKIISVAKTTAK-----ITVTQLSR*FVNLKMTFLQFINNNRQEGQGYISEKLFKTKKNE
                                            -----------YCL001W-A---------
S. paradoxus IIRKTITDYVAVRLRILSHEFDIKEEYLRYKGITVTDGTNISSANVPLGKFLSFNITYEH
             +IRKT+T+ VAVRL+ LSHEFD+ E YLRY          I+S             T EH
S288c        MIRKTVTNLVAVRLKNLSHEFDVIENYLRY----------IAS-------------TSEH
             ------------------------------------------------------------
S. paradoxus QFTIFKRHFNKYARKLLKEALDSEMKSDTAAVVLQEGISHICLLTTSSTILKQKIEVSIP
              FT  KRHFNK ARKLLKEA+DS+  S+TA VVLQEG S ICLL  SS ILK K++
S288c        LFTAIKRHFNKCARKLLKEAIDSKSNSETATVVLQEGFSGICLLKASSIILKLKLKFP--
             ------------------------------------------------------------
S. paradoxus KKKDTVDISKLDEKTEAFYRRTYDAVVKNFNFDELRVIILC-------SPGFYAKTLMEK
             KKKD  DISKL +K E   +              L + IL        +P  + +    K
S288c        KKKDRTDISKLCDKKERMTQW-------------LEISILMN*E**YYAPLDFMRRHC*K
             ----------------------------------------->
S. paradoxus ILKYAEKDQNNSVLQNQEMFVIAHCS--TGYIQGINEVLRDPEYSKILQNTKYLKEIHIM
                Y ++    +  +     V AHCS  T Y+QGI+EV+R+PEYS I+ NTK LKE  IM
S288c        NF*YMQRRSK*ASFKEPR*AVFAHCSSSTKYLQGIDEVVRNPEYSMIVHNTKKLKESRIM
                                                                        =
S. paradoxus DEFLEHLNKDDDRAWYGVEETERAAKLGAIETLLITDSVLKSDDVKKREKYLDLIENIEQ
             D+FLEHL+KDD++AWYG EETERAAKL AIETLLITDSVLK +DVKKREKYLDLIEN
S288c        DDFLEHLSKDDNKAWYGAEETERAAKLDAIETLLITDSVLKRNDVKKREKYLDLIENSGN
             ============YCL001W-B=======================================
S. paradoxus NNGEVFVFST-fs-
             NNG++FV ST
S288c        NNGKIFVLSTSKITVSNLTNQQI*
             =======================>

Frame = +3
S. paradoxus            LQSHGKELETLTGLACILKYPIHNLDEIFQ  384
                         ++H ++L+  T + C+LKY I  LDE+F+
S288c                   FKNHSEQLDKPTDIGCVLKYTIQYLDELFK  3992
```

Figure S10 REMYHR054

A) Gene order in the region of REMYCL001W-A. Duplication including part of YHR056C “()”. Duplication involving part of YHR052W “[]”. A duplication of CUP1 without duplicating part of either flanking gene or duplicating the entire flanking gene should in theory also be possible but were not observed.

B) Examples of structures of REMYHR054. Other lengths should also be possible.

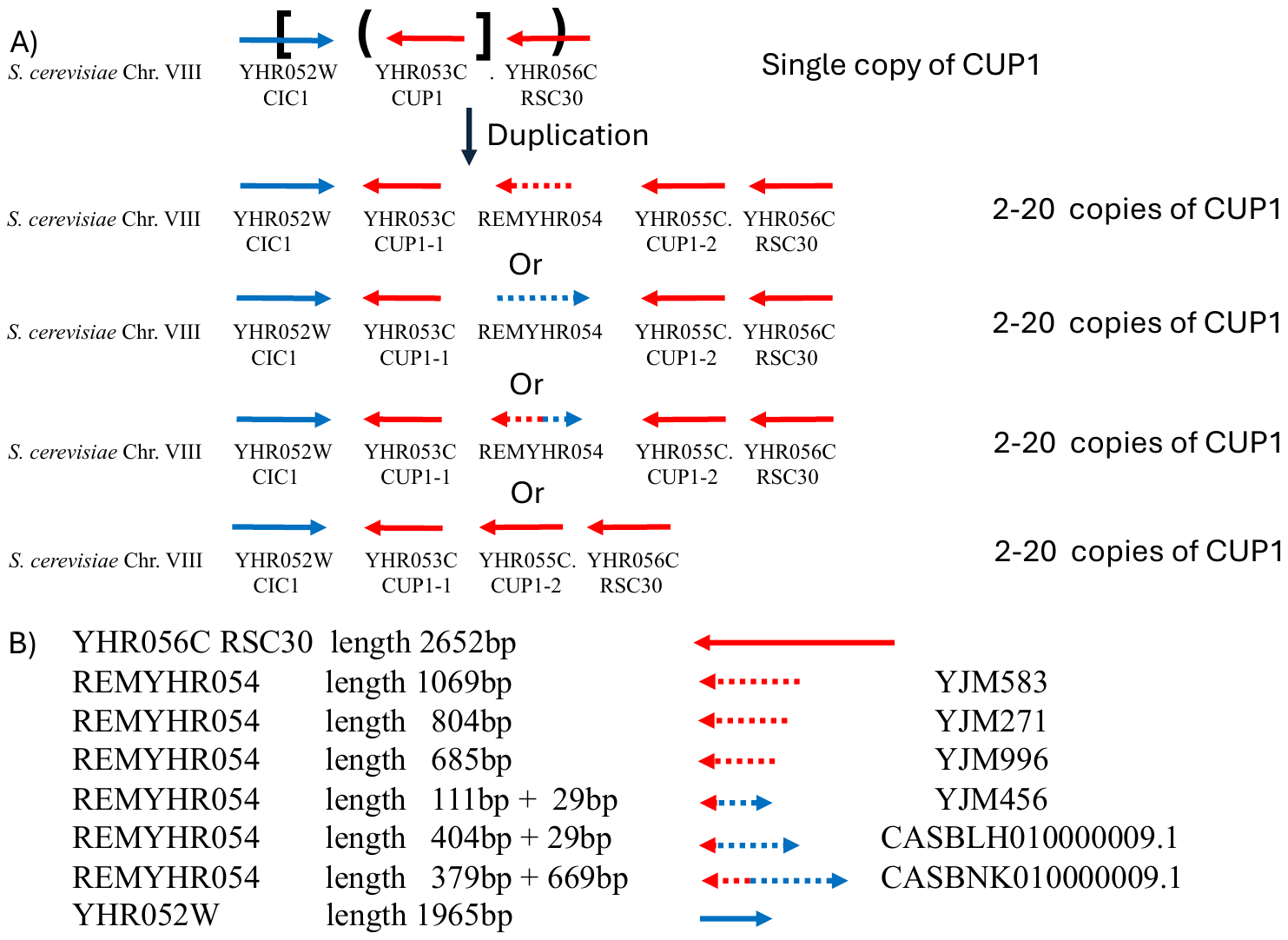

Figure S11 REMYDR508W-A

A) Gene order in the region of REMYDR508W-A. Duplication includes a portion of *YDR510W*.

B) Alignment of a portion of *YDR510W* and REMYDR510W from strain YJM1400. The “/” indicates site of a frame shift. Two in frame stop codons.

A)

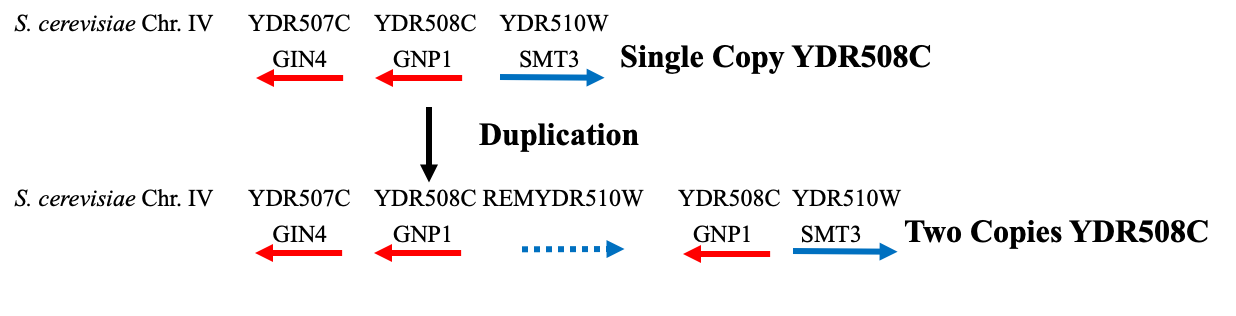

B)

```
YDR510W 15   KPEVKPETHINLKVSDGSSEIFFKIKKTTPLRRLMEAFAK RQGKEMDSLRFLYDGIRIQAD
REMYDR510W       QSRTHIN*KVSYVSSEIFSMIRKTTSLKIFMESFTK/RQGRKMDS*KFLYCG
                    **** ***  *****  * *** *   ** * * ***  ***  *** *
```

Figure S12 REMADL001C-A

A gene remnant found in *A. gossypii.* Gene order in the region of REMYCL001W-A

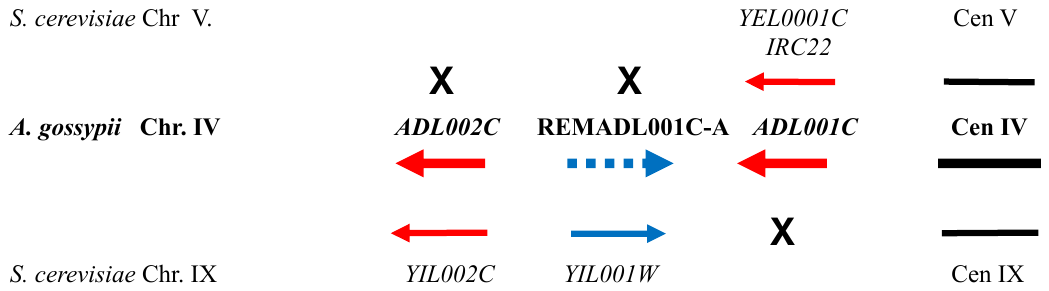

Figure S13

A) Alignment of 5' UTR region of *YBR089C*-A (*NHP6B*), dubious gene *YBR090C,* comparing *S. cerevisiae* S288c (top) and *S. paradoxus* strain N44 (bottom). Frameshifts (fs) both upstream and downstream of intron disrupt potential reading frames. Dashes under sequence indicate intron features: 5' splice site is the canonical GTATGT, branch site is the canonical TACTAAC, and 3' splice site is one of the two canonical sequences TAG). Equal signs underline start codon and beginning of *YBR089C-A* (*NHP6B*) open reading frame.

B) One of several RNA-seq reads from SRR488142 confirming splicing, and location of 3' splice site. Carrot indicates splice site.

A)

```
S288c  1       ATGGTACCCGCACCTGGGTCACGTGCATTCCCCTCACCCGtttttttaggtggtgtcttt  60
               ||| |||||||||||  |||||||||||||| ||||||| |||||||||||||||| |||
S. para 414700 ATGTTACCCGCACCTAAGTCACGTGCATTCCTCTCACCC-TTTTTTTAGGTGGTGT-TTT  414643
                                                      fs               fs
S288c  61      tttgtttttttttttAGGTGGCGCGGCAACTATAAAGTACAGCAAGTGAGGTTGAGGCAA  120
               ||| | |||||||||||||||||||||||||||||||||||||||||||||||||||||
S. para 414642 TTT-TGTTTTTTTTTAGGTGGCGCGGCAACTATAAAGTACAGCAAGTGAGGTTGAGGCAG  414584
                  fs
S288c  121     TACTGGGAGTTTACACTATGGGAGACAGCTCCTAACACCAAGCAGAAAAACGACTTTTTC  180
               || ||| |||||||||||||||||||||||||||||||||||||||||||||||||||||
S. para 414583 TAATGGAAGTTTACACTATGGGAGACAGCTCCTAACACCAAGCAGAAAAACGACTTTTTC  414524
                     5' splice stie
S288c  181     GCAAAGTATGTAAGGCGCTGGGTGAGCCCAGCGGACGAGGATGGGCTTAATAAGAACGTA  240
               ||||||||||||||||||||| |||||||||||||||| ||||||||| |||||||| ||
S. para 414523 GCAAAGTATGTAAGGCGCTGG-TGAGCCCAGCGGACGACGATGGGCTTGATAAGAACATA  414465
                    ------
S288c  241     CAGTTTAGCACAGCTAGAACAGGATACAGCTAAGGGCAACTCTGCTTTTCGGGAGAAGTT  300
               |||||||||||| |||||||||||||||||||||||||||||||||||||  |||| |||
S. para 414464 CAGTTTAGCACAACTAGAACAGGATACAGCTAAGGGCAACTCTGCTTTTCATGAGAGGTT  414405

S288c  301     AAAGAGGGGTAGACAATGATGGTAATCT-TATAAACCGGCTACAATGAAGGTTGTAGCAG  359
               ||||||||||| | |||||| || || | || |||  |||||  || |||||| ||||||
S. para 414404 AAAGAGGGGTA-ATAATGATAGTTATTTCTACAAATTGGCTATGATAAAGGTTATAGCAG  414346

S288c  360     CAAGGAAGATGATATTTTAATACGGTTCAGGTGAAATGAAATAGCCGCCCATAACGGCAT  419
               ||||| | ||| ||||  | |||  |||||  | || || |||||||||||||  |||||
S. para 414345 CAAGGGAAATGGTATTCCATTACAATTCAGTGGTAACGATATAGCCGCCCATAGTGGCAT  414286

S288c  420     GCTCAAGTTGTAAGTCAGGACTCTAGCTTTCTACTGTAGTATCCTCTAAAGGACT-GCTG  478
                ||| | |||||||| |||||| |||||||  | ||||  |||||||||  | || | ||
S. para 414285 ACTCGAATTGTAAGTTAGGACTTTAGCTTTACATTGTAACATCCTCTAACCGGCTAGTTG  414226
                                                             Branch site
S288c  479     TTCTGTGCAC-CCCCTTCCTTTGTTTATCATAGCGCACGACAAGAGTACTAACTAATTAA  537
               |||||  ||| ||||||| | |||||||| || ||||||||||||||||||||||||||
S. para 414225 TTCTG--CACGCCCCTTCTTCTGTTTATCGTAACGCACGACAAGAGTACTAACTAATTAG  414168
                 3' splice site                              -------
S288c  538     CTTAGAACATTAACATATATAAAACTAGCGCTATGGCCGCAACTAAAGAAGCAAAGCAAC  597
               |||||| | | |||||| ||  ||| |||||||||||||||||||| |||||||||||||
S. para 414167 CTTAGAGCTTAAACATACAT--AACGAGCGCTATGGCCGCAACTAAGGAAGCAAAGCAAC  414110
                 ---               fs          ==================NPH6B====>

B)
CTAACACCAAGCAGAAAAACGACTTTTTCGCAAAAACATTAACATATATAAAACTAGCGCTATGGCCGCAACTAAAGAAGCAAAGC
                                  ^
```

Figure S14

A) The *CUP1* protein sequence is identical in all 94 strains. However, there are four non-synonymous polymorphisms in the *YHR052C-B* protein, including a loss of start codon.

B) Comparing the *Saccharomyces* *sensu stricto* species *CUP1* protein sequences, there are eight conservative amino acid changes, and seven other changes.

C) Comparing the *Saccharomyces* *sensu stricto* species *YHR052C-B* protein sequences. There are four conservative amino acid changes, and thirty-one other changes, including loss of start codon.

A)

```
S288c YHR052C-B MKVMSANANVVAAKIMNNAKNHVAAQRGVTATTNAPAVTSLKKPRSHAALGNETNSL
YJM1083 chrVIII MKVMSANANVVAAKIMNNAKNHVAAQRGVTATTNAPAVTSLKKPRNHAALGNKTNSL
YJM1129 chrVIII MKVMSANANVVAAKIMNNAKNHVAAQRGVTATTNAPAVTSLKKPRSHAALGNKTNSL
YJM1248 chrVIII TKVMSANANVVAAKIMNNAKNHVAAQRGVTATTNAPAVTSLKKPRSHAALGNKTNSL
YJM1402 chrVIII MKVMSANANVVAAKIMNNAKNHVAAQRGVTATTNALAVTSLKKPRSHAALGNKTNSL
                 ********************************** *********.******:****
```

B)

```
S288c YHR053C  MFSELINFQNEGHECQCQCGSCKNNEQCQKSCSCPTGCNSDDKCPCGNKSEETKKSCCSGK
Spara CBS432    MFSELINFQNEGHECQCKCGSCKNNEQCQKSCSCPTGCDSDDKCPCGNKSEKTKKPCCSE
S. jurei        MFSKLVNFQNEGHECQCKCGSCKNNEQCQKSCSCPTGCNSDDKCPCGNKSEKTKKPCCSE
S. uvarum       MFSELINFQNEGHQCQCQCGSCKTNEQCQKSCNCPTGCNSDDKCPCGNKSEKTKKPCCSA
S. kudriavzevii   MFSELIDFQNEGHECQCQCGSCKNNEQCQKSCRCPIGCNSDDKCPCGAKSEKTKKPCCSE
S. eubayanus    MFSELINFQNEGHQCQCKCGSCKTNEQCQKSCNCPTGCDSDDKCPCGNKSEKTKKPCCSA
S. pastorianus  MFSELINFQNEGHQCQCKCGSCKTNEQCQKSCNCPTGCDSDDKCPCGNKSEKTKKPCCSA
S. mikatae      MFSELVNSQNEGHECQCKCGSCKNNEQCQKSCSCPTGCNSDEKCPCGNKSEKTKKPCCSE
                ***:*:: *****:***:*****.******** ** **:**:***** ***:***.***
```

C)

```
S288c YHR052C-B     MKVMSANANVVAAKIMNNAKNHVAAQRGVTATTNAPAVTSLKKPRSHAALGNETNSL
Spara CBS432.       MKVMNANANVAAAKIMNNAKNRAAAQRAATATTSALAATSLKKLRSHAALNKLMNNL
S. jurei        LTSKTKVTNANANVVAAKTMNNVRNRAAAQQVVTATTNALAVTSLKKLRSHAVLNN
S. uvarum       LISKTKATNVNANVVVAKPTNNVKNRATVQLAVIVMTNAPVVTSLKRPRSHAARHK
S. kudriavzevii        MNANASAVAARTMNNAKNRADAQSAVTAMINALVEQSLKKLRSHAALSKRRNYLQIHSCAH
S. eubayanus    SISKTKATNVNANVVVAKPMNSVKNRATVQLAVTVMTNAPVATSLKKPRSHAARLK
S. pastorianus  SISKTKATNVNANVVVAKPMNSVKNRATVQLAVTVMTNAPVATSLKKPRSHAARLK
S. mikatae             MNANANVVAAKTMNNVKNRAVAQQAVIATKNALAVTSLKKLRSHAVLNR
                        ..**....*:  *..:*:. .*  . .  .* .  ***: ****.  .
```

Figure S15

A) S288c sequence of *YKL104W-A*. Underneath are the nucleotide polymorphisms found in *S. paradoxus* strain CBS432. Sixteen of the polymorphisms are in position one of the codon, one is in position 2, and zero are in position 3. Position 1 of the *YKL104W-A* reading frame corresponds to position 3 of the protein coding gene found on the other strand, *YKL104C* (*GFA1*)

B) S288c sequence of *YLR379W-A*. Underneath are the nucleotide polymorphisms found in the 93 strains. Reading frame one: three. Reading frame two: seven. Reading frame three: seven.

A)

```
S288c     ATGAACGACCACAAATTGGTCTTCTGGGTCAGATCTTTGAGGGTGACAGTTAACTTGTTC
CBS432                   C     C           C
S288c     M  N  D  H  K  L  V  F  W  V  R  S  L  R  V  T  V  N  L  F

S288c     TGGTCGACCGTGAGTAGCCCATCTAGTATGCGCAATACCACAATGAGAGACAAAAGTAAC
CBS432        T                                  G
S288c     W  S  T  V  S  S  P  S  S  M  R  N  T  T  M  R  D  K  S  N

S288c     GTCTCTGTTCGGATTTTGCTTAGTAATCTCCTCTTTCAAAGCACTCACTTTACCGATTTG
CBS432    A  C  A  T  G                 T           G
S288c     V  S  V  R  I  L  L  S  N  L  L  F  Q  S  T  H  F  T  D  L

S288c    CTTATAGATGAAAGTAGAATCAGCTTCGTCACCATCGATAGCAATACCGGTGGAATCATA
CBS432         A  A     G                          G        A
S288c    L  I  D  E  S  R  I  S  F  V  T  I  D  S  N  T  G  G  I  I

S288c     GCCTCTATATTCTAA
CBS432
S288c     A  S  I  F  *
```

Figure S16

A) S288c sequence of *YLR379W-A*. Underneath are the nucleotide polymorphisms found in the 93 strains. Reading frame one: three. Reading frame two: seven. Reading frame three: seven.

A)

```
S288c     ATGCTTGTTGTTTGGCAAAAGTTGCCGAAAGAAAGAATGAGAGAGAACGACAAAGAGAAT
          G                                              T
S288c     M  L  V  V  W  Q  K  L  P  K  E  R  M  R  E  N  D  K  E  N

S288c     GAGAGAGAACACGTAAAGCGAAAAATCTGGAGCGCTGCCGCGCCGGGTAACAAGTGTGAA
                                             A                    G
S288c     E  R  E  H  V  K  R  K  I  W  S  A  A  A  P  G  N  K  C  E

S288c     GAGAGAGAGAGATGTAAGGAAGATGAATACATTCAATACATTCTAGCTTATAGCAAGCTT
           T  2-base_fs                T G      T   AA
S288c     E  R  E  R  C  K  E  D  E  Y  I  Q  Y  I  L  A  Y  S  K  L

S288c     TCAGCAGGCTCAATTGCTTATAGCGGTATAACTGCAACAAGAGTGAAAATTAATATGGTG
           T                     T       T                    G
S288c     S  A  G  S  I  A  Y  S  G  I  T  A  T  R  V  K  I  N  M  V

S288c     ACGAACGACGGATTCGGTAGTACAGTGTTATAA
           AA                  T
S288c     T  N  D  G  F  G  S  T  V  L  *
```

Figure S17 A case of plus one ribosomal frameshifting currently misannotated as an intron in S288c, *YJR112W-A*.

A) Potential intron splice and branch sites are underlined. None of the species below has consensus 5' splice sites, branch site, and 3' splice sites. The potential splice sites are at different positions and reading frames. The consensus +1 ribosomal frame shifting sites (CTTAGGC) are marked in bold with equal signs over them. *Kluyveromyces lactis* strain NRRL Y-1140 gene is all in the same reading frame and is missing the ribosomal frame shifting site and has no intron.

A)

```
YJR112W-A S288c                                                            =======
            ATGGTACAGCTGAGAAGAACGTATGTATATATGTTCGTGGTAAGCTCGTTGGTATGCACGTATCTTAGGCATCACTACCAACAAAGTGTTTCAAGCGATCACGTCGACTAACGACAA
            M  V  Q  L  R  R  T  Y  V  Y  M  F  V  V  S  S  L  V  C  T  Y  L  R  H  H  Y  Q  Q  S  V  S  S  D  H  V  D  *
                                                    *  A  R  W  Y  A  R  I  L  G  I  T  T  N  K  V  F  Q  A  I  T  S  T  N  D  K

S. eubayanus 450.1                                                         =======
            ATGGCGCAGGTAAGAAGGACATACGCGTGTATGTTTGTGGTGAGCTCGCTGGCGTGCGCGTACCTTAGGCATTACTACGAACAAATTATTCCAGGCGGTCACCTCGACTAACGACAA
            M  A  Q  V  R  R  T  Y  A  C  M  F  V  V  S  S  L  A  C  A  Y  L  R  H  Y  Y  E  Q  I  I  P  G  G  H  L  D  *
                                                    *  A  R  W  R  A  R  T  L  G  I  T  T  N  K  L  F  Q  A  V  T  S  T  N  D  K

S. mikatae IFO1815                                                         =======
            ATGGTGCTGTTGAGAAGAACGTATGTGTGCATGTTTGTGGTATGCTCGTTGGCATCCGCCTACCTTAGGCATCACTACGAACAAAATATTTCAAGCAATTACATCGACTAATGACAA
            M  V  L  L  R  R  T  Y  V  C  M  F  V  V  C  S  L  A  S  A  Y  L  R  H  H  Y  E  Q  N  I  S  S  N  Y  I  D  *
                      *  E  E  R  M  C  A  C  L  W  Y  A  R  W  H  P  P  T  L  G  I  T  T  N  K  I  F  Q  A  I  T  S  T  N  D  K

S. paradoxus N44                                                           =======
            ATGGTACAGCTAAGAAGAACGTATGCGTGCATGTTCGTGGTGAGCTCGTTGGTATGCGCGTACCTTAGGCATCACTACCAACAAATTATTTCAAGCGATCACATCGACTAACGACAA
            M  V  Q  L  R  R  T  Y  A  C  M  F  V  V  S  S  L  V  C  A  Y  L  R  H  H  Y  Q  Q  I  I  S  S  D  H  I  D  *
                                                    *  A  R  W  Y  A  R  T  L  G  I  T  T  N  K  L  F  Q  A  I  T  S  T  N  D  K

S. uvarum CBS7001                                                          =======
            ATGGTGCAGGTAAGAAGGACATATGCGTGCATGTTTGGGGTGAGCTCGCTGGCGTGCGTGTACCTTAGGCATTACTACGAACAAGTTATTCCAGGCGGTGACCTCGACTAACGACAA
            M  V  Q  V  R  R  T  Y  A  C  M  F  G  V  S  S  L  A  C  V  Y  L  R  H  Y  Y  E  Q  V  I  P  G  G  D  L  D  *
                                                    *  A  R  W  R  A  C  T  L  G  I  T  T  N  K  L  F  Q  A  V  T  S  T  N  D  K

K. lactis NRRL_Y-1140
                   ATGATAAGATATAAATACGCAGTAACATTTTTCTCGAGTGTATGTTTTGTTGTCTTCCTCGGATTAACTACTGATGCCAGGTTAAAGCTAATAACATCAACTCATGATAA
                   M  I  R  Y  K  Y  A  V  T  F  F  S  S  V  C  F  V  V  F  L  G  L  T  T  D  A  R  L  K  L  I  T  S  T  H  D  K

A. gossypii                                                                =======
ATGGAGCTATCAGGGCTTGGTATACGGTGGCGCTATTTCGGGTTACTTTGCGCGACAGGCTGCGTGGCGGTTTTTCTTAGGCATCACTACAGATAAGCGGTTCAACGCGATCACAGCGGA
M  E  L  S  G  L  G  I  R  W  R  Y  F  G  L  L  C  A  T  G  C  V  A  V  F  L  R  H  H  Y  R  *
 W  S  Y  Q  G  L  V  Y  G  G  A  I  S  G  Y  F  A  R  Q  A  A  W  R  F  F  L  G  I  T  T  D  K  R  F  N  A  I  T  A  E
```

Figure S18 Putative NAG1 open reading frame conserved in *S. cerevisiae*, *S. uvarum*, and *S. arboricola*, but not other related fungi.

```
NAG1 S. cerevisiae      MNSAGRVHRS RAGSRGHAAI SPLTMASFSV ARGIRSSNVY DDTDD---EL
NAG1 S. uvarum          MNNAGRVQRS CVGSRGHAAI SPFTMASLSV PSGMRSWNVY DDAPD---AP
NAG1 S. arboricola      MNSAGRVHRS CAGSRGHAAI SPLTIALFSV ARGMRSWNVY DDANDGLFVA

NAG1 S. mikatae         MNSAGRVHRS RAGSRGHAAI SPLTIASFRV VRGIRSWNVY DDVDD---GS
NAG1 S. paradoxus       MNSAGRVHRS RTGSRGQAAI SPLTIASFRV SRGIRSSNVY DDADD---GL
NAG1 S. bayanus         MNSAGRVQRS CVGSRGHAAI SPFTMASFSV PSGMRS*NVY DDAPN---AP
NAG1 S. kudriavzevii    MNSAGPVHRS WAGSKGHAAI SPLTIASFSV DRGIRSWNVY DDEGD---EP
NAG1 A. gossypii        RNSAGPVR*W VAGESSQPAM APRTTASLSV RRGMRGRRNL PEAGS-----

NAG1 S. cerevisiae      SILTFFSAVR RNRLTSSLPP ILSARCSSAC FSVRIVLPLS LTISISALMY
NAG1 S. uvarum          SALALFSVVS RNRLTSSLPP ILSARCSSAC FSVWMVLPLS LTISSSALMY
NAG1 S. arboricola      SDLTLFSVVS RKRRTSSLPP ILSARRSSAC FNVWIVFPLS LTISERALMY

NAG1 S. mikatae         LILILFNVVN KNRRISSLAP ILFAR*SSAC FSVWMYLPLS LTISSNALMY
NAG1 S. paradoxus       SILTLFNVVR RNRLTSSLPP ILSARRSSAC FSVWIVLPLS LTISISALMY
NAG1 S. bayanus         SVLTLFSVVS RNRLTSSLPP ILSARRSSAC FSVWMVLPLS FTISSSALMY
NAG1 S. kudriavzevii    SIVTLFRVVS RNRLTSSLPP ILPARYSSAC FSVWIVLPLS LTISISALIS
NAG1 A. gossypii        ----RCSMSE RRNRRTSGSD STAARARCAC AAE----SSG CAARRSRAPM

NAG1 S. cerevisiae      STNSALGRKL TGAFSIQTNI EQSCGFFRTS IMATLPPIEC PIIIGPPL-V
NAG1 S. uvarum          STNLALGRRL TGAFSIQTIM EHNCGFFRTS IMATLPPIEW PTIMGPPW-V
NAG1 S. arboricola      WTNSVLGCKL TGAFSIQTIM EHNCGFLRTS IMATLPPIEW PTMMGPPL-M

NAG1 S. mikatae         FTKSALGRKL TGAFSIQTIM EHNCGFLRTS IIATLPPIE* PTIIGPPL-M
NAG1 S. paradoxus       STNSALGRRL TGAFSIQTNI EHNCGFFKTS IIATLLPIE* PTMIGPPS-V
NAG1 S. bayanus         STNSILGRRL TGAFSIQTIM EHNCGFFRTS IMATFPPIE* PTIMGPSW-V
NAG1 S. kudriavzevii    -TNSALGRKL TGAFSIQTII EHNCGFFRTS IIATLPPIE* PTIIGPPW-A
NAG1 A. gossypii        RGNSGSGTVA TGAFSMEPRA EHRCGWRRAR PMATLGPIEC PTRTETARPV

NAG1 S. cerevisiae      FNSCFVIKCF TSSDMTS*
NAG1 S. uvarum          FSSCFVMKRF TSSDIAP*
NAG1 S. arboricola      LSSCFVMKCL TSSDITR*

NAG1 S. mikatae         FNSCFVIKCF TSSDIAA*
NAG1 S. paradoxus       FNSCFAMKCF TSSDIAA*
NAG1 S. bayanus         FSSCFVMKCF TSSDIAP*
NAG1 S. kudriavzevii    FNLCFAMKCL TSSDIAL*
NAG1 A. gossypii        FSRSQRATSA TMAG*SWR
```

Figure S19

Alignment of dubious gene *YNL269W* (*BSC4*) from multiple *S. cerevisiae* strains. In approximately half of strains examined, the open reading frame is intact, as in S288c. In the other half there are one or more frame shifts, marked here by “**/**” and in a few cases internal stop codons, marked here by “*****”. Most of the frame shifts are at homopolymer runs. Strains are from the 94 strains or the 1011 strains (Peter et al. 2018).

```
S288c           MSIVLRKSN-KKNKNCITSKFYTIHIIKISTPVFRAPIAIGESPYVEWSCLQVVFRKDMV
CNB_4-26425     MSIVLRKSN/KKNKNCITSKFYTIHIIKISTPVFRAPIAIGESPYVEWSCLQVVFRKDMV
yjm270_Chr14    MSIVLRKSN-KKNKNCITSKFYTIHIIKISTPVFRAPIAIGESPYVEWSCLQVVFRKDMV
AHQ_1-6338      MSIVLQKSN-KKNKNCITSKFYTIHIIKISTPVFRAPIAIGENPYV*WSCLQVVFRKDMV
ACI_7-6814      MSIVLRKSN-KKNKNCITSKFYTIHIIKISTPVFRAPIAIGENP*VEWSCLQVVFRKDMV
yjm1479_Chr14   MSIVLRKSN-KKNKNCITSKFYTIHIIKISTPVFRAPIAIGENPYVEWSCLQVVFR*DMV
yjm428_Chr14    MSIVLQKSN-KKNKNCITSKFYTIHIIKISTPVFRAPIAIGENPYVEWSCLQVVFRKDMV
yjm195_Chr14b   MSIVLQKSN/KKNKNCITSKFYTIHIIKISTPVFRPPIAIGENPYVEWSCLQVVFTKDMV
yjm1463_Chr14   MSIVLRKSN-KKNKNCITSKFYTIHIIKISTPVFRAPIAIGENPYVEWSCLQVVFRKDMV
yjm1355_Chr14   MSIVLRKSNKK/NKNCITSKFYTIHIIKISTPVSRAPIAIGENPYVEWSCLQVVFRKDMV
                *****:*** * ********************* *.******.* * ********  ***

S288c           TKKTTFAQLITRLNHFLCQALKRR-DSKTYILCRTAVFGAMTPFSPRKSH-INN-KLPMQ
CNB_4-26425     TKKTTFAQLITRLNHFLCQALKRR-DSKTYILCRTAVFGAMTPFSPRKSH-INN-KLPMQ
yjm270_Chr14    TKKTTFAQLITRLNHFLCQALKRR-DSKTYILCRTAVFGAMTPFSPRKSH-INN-KLPMQ
AHQ_1-6338      TKKTTFAQLITRLNHFLCQALKRR-DSKTYILCRTAVFGAMTPFSPRKSH-INN-KLPMQ
ACI_7-6814      TKKTTFAQLITRLNHFLCQALKRR-DSKTYILCRTAVFGAMTPFSPRKSH-INN-KLPMQ
yjm1479_Chr14   TKKTTFAQLITRLNHFLCQALKRR-DSKTYILCRTAVFGAMTPFSPRKSH-INN-KLPMQ
yjm428_Chr14    TKKTTFAQLITRLNHFLCQALKRR/DSKTYILCRTAVFGAMTPFSPRKSH-INN-KLPMQ
yjm195_Chr14b   TKKTTFAQLITRLNHFLCQALKRR-DSKTYILCRTAVFGAMTPFSPRKSH-INN-KLPMQ
yjm1463_Chr14   TKKTTFAQLITRLNHFLCQALKRR-DSKTYILCRTAVFGAMTPFSPRKSHLINN-/LPVQ
yjm1355_Chr14   TKKKTFAQLITRLNHFLCQALKRR/DSKTCILCRTSVLGPMAPF/PRKSHIINN/QLPVQ
                ***.******************** **** *****:*:*.*:** ***** ***  **:*

S288c           PRK--KK-IVIIYVVRFH*
CNB_4-26425     PRK--KK-IVIIYVVRFH*
yjm270_Chr14    PRK--KK/IVIIYVVRFH*
AHQ_1-6338      PRK--KK/IVIIYVVRFH*
ACI_7-6814      PRK--KK/IVIIYVVRFH*
yjm1479_Chr14   PRK--KK/IVIIYVVRFH*
yjm428_Chr14    PRK--KK/IVIIYVVRFH*
yjm195_Chr14b   PRK--KK/IVILYVVRFH*
yjm1463_Chr14   SRKT-KKRIVIIYAVRFR*
yjm1355_Chr14   SRKT/KKRIVIIYVVRFR*
                .**  ** ***:*.***:
```

Figure S20 Alignment of the *TAR1* region of *S. cerevisiae* and several related species. Entire alignment not shown.

A) The start codon ATG, also found in *S. paradoxus*, but is CTG in most related species.

B) The lower block is found approximately two thirds of the way through the gene. At the position marked *S. cerevisiae* has an AAG codon (lysine). Many of the species have a TAG codon (stop).

A)

```
               Start codon
T. pretoriensis         CTG CGAGATT CCCCCACCCA CAAGGAGCAG AGGGCGCAAA ACACCATGTC
S. uvarum               CTG CGAGATT CCCCCACCCA CAAGGAGCAG AGGGCGCAAA ACACCATGTC
S. bayanus              CTG CGAGATT CCCCCACCCA CAAGGAGCAG AGGGCGCAAA ACACCATGTC
K. africana             CTG CGAGATT CCCCTACCCA CAAGGAGCAG AGGGCGCAAA ACACCATGTC
A. gossypii             CTG CGG..TG ACCCCACCCA CAAGGAGCGA GGGCCGCAAA ACACCATGTC
T. globosa              CTG CGAGATT CCCCCACCCA CAAGGAGCAG AGGGCGCAAA ACACCATGTC
TAR1 S. cerevisiae      ATG CGAGATT CCCCTACCCA CAAGGAGCAG AGGGCACAAA ACACCATGTC
S. dairenensis           CTG CGAGAGT CCCCCACCCA CAAGGAGCAG AGGGCGCAAA ACACCATGTC
S. paradoxus            ATG CGAGATT CCCCTACCCA CAAGGAGCAG AGGGCACAAA ACACCATGTC
K. unispora             CTG CGAGATT CCCCCACCCA CAAGGAGCAG AGGGCGCAAA ACACCATGTC
K. exigua               CTG CGAGATT CCCCCACCCA CAAGGAGCGA GGGGCACAAA ACACCATGTC
                        M/L
```

B)

```
                                                                 ---
T. pretoriensis         ATTCCCAAAC AACTCGACTC TTCGAAAGCA CTTTACA AAG AACTGGGATC
S. uvarum               ATTCCCAAAC AACTCGACTC TTCGAAGGCA CTTTACA TAG AACCGCACTC
S. bayanus              ATTCCCAAAC AACTCGACTC TTCGAAGGCA CTTTACA TAG AACCGSACTC
K. africana             ATTCCCAAAC AACTCGACTC TTCGAGAGAA CTTTACA AAG AACCGCACTC
A. gossypii             ATTCCCAAAC AACTCGACTC GTCGAAAGAA CCTTAGA TGG CACTAGCACC
T. globosa              ATTCCCAAAC AACTCGACTC TTCGAAAGCA CTTTACA TAG GACTGGGATC
TAR1 S. cerevisiae      ATTCCCAAAC AACTCGACTC TTCGAAGGCA CTTTACA AAG AACCGCACTC
S. dairenensis           ATTCCCAAAC AACTCGACTC TTCGAAGGCA CTTTACA AAG GACTGGGTTC
S. paradoxus            ATTCCCAAAC AACTCGACTC TTCGAAGGCA CTTTACA TAG AACCGCACTC
K. unispora             ATTCCCAAAC AACTCGACTC TTCGAAGGCA CTTTACA TAG AACCGCACTC
K. exigua               ATTCCCAAAC AACTCGACTC TTCGAGAGCG CTTTACA CGG AACCGCACTC
                                                                K/R/W/stop
```

Figure S21 Start codon errors

A) Some start codon differences between strains appear to be real and the result of polymorphisms such as in the case of *YDL144C*.

B) *YDR512C* Third start codon appears to be the conserved start codon suggesting that divergent genes *YDR512C* and *YDR513W* do not overlap. Among the *sensu stricto* species the first 96 residues of the currently annotated *YDR512C*, X, contain multiple frame shifts and stop codons and are quite diverged, whereas the remainder of the protein is quite conserved.

C) *YFL021W* Second start codon appears to be the conserved start codon.

D) *YBR073W* Second start codon appears to be the conserved start codon.

E) *YBR245C* Fourth start codon appears to be the conserved start codon.

F) *YGL086W* Second start codon appears to be the conserved start codon.

G) *YGR079W* Second start codon appears to be the conserved start codon.

H) *YGR140W* (*CBF2*) Fourth start codon appears to be the conserved start codon.

I) *YGR174C* Second start codon appears to be the conserved start codon.

J) *YML126C* (*ERG13*) Second start codon appears to be the conserved start codon.

K) *YJR107W* Misannotated in S288c should use upstream ATG – current start codon is non-conserved Methionine.

L) *YJR126C* Third start codon appears to be the conserved start codon.

M) *YJR127C* (*RSF2*) Wrong start codon (WGD duplicated gene)

N) *YKL021C* Second start codon appears to be the conserved start codon.

O) *YKR056W* Second start codon appears to be the conserved start codon.

P) *YLR119W* Second start codon appears to be the conserved start codon.

Q) *YLR176C* Second start codon appears to be the conserved start codon.

R) *YLR218C* Second start codon appears to be the conserved start codon.

S) *YLR408C* Second start codon appears to be the conserved start codon.

T) *YMR019W* Second start codon appears to be the conserved start codon.

U) *YNR020C* Fourth start codon appears to be the conserved start codon.

V) *YNR056C* Second start codon appears to be the conserved start codon.

W) *YOR241W* Third start codon appears to be the conserved start codon.

X) *YHR175W* Second start codon appears to be the conserved start codon.

Y) *YIR017C* Second start codon appears to be the conserved start codon.

A)

```
S288c + 91 strains  YDL144C:   ATG---------TCTAGAATACTAGTTATCGGG
                               M           S  R  I  L  V  I  G
YJM1244 and YJM1250 YDL144C:   ATGTCGAAAATATCTAGAATACTAGTTATCGGG
                               M  S  K  I  S  R  I  L  V  I  G
```

B)

```
S288c YDR512C current annotation           XMSTK YPTTMSCREA FDQLTSCYSI GGQFRSYYRY GDFTSCDKQV
YJM1242 YDR512C                             MSTK YPTTMSCREA FDQLTSCYSI GGQFRSYYRY GDFTSCDKQV
S. paradoxus                                MSAK YPTTMSCREA FDQLTSCYSI GGQFRSYYRY GDFTSCDKQV
S. arboricola                               MSAK YPTTMSCREA FDQLTSCYSI GGQFRSYYRH GDFTSCEKQV
A. gossypii AFR707C             MPDSA PVEPQQPAEQ YPSQISCTEA FDRLTDCYSI GGLVRHYYRH GEFNGCSKEI
where X=MIITIKFESKEKLVSMKQYITVVGVSVNQEGTFSWRPGTFYMKIHEESAAGPFAKGAEISDLSNSSAIGCGISINQEGGDGSIRVKESSLRKKQER
```

C)

```
S288c  YFL021W           MHVFFPLLFR PSPVLFIACA YIYIDIYIHC TRCTVVNITM STNRVPNLDP
YJM627 YFL021W                                                     M STNRVPNLDP
S. paradoxus                                                       M STNRAPNPDP
A. gossypii AFR513C                                        METWSQDAR TAQRRLREAS
```

D)

```
S288c  YBR073W                   MAVISVKPRR REKILQEVKN SSVYQTVFDS GTTQMQIPKY ENKPFKPPRR
YJM693 YBR073W                                                        MQIPKY ENKPFKPPRR
S. paradoxus                                                         MQIPKY ENKPFRPPRR
S. mikatae                                                            MQLPKY ENKPFKPPRR
S. kudriavzevii                                                       MQIPKY QNKPFKPPRR
S. uvarum                                                             MQIPKF ENKPFKPPRR
A. gossypii AGL212W                                               MSNGLSLPKY ANMPFRPPRM
K. lactis                                                               MNHY VNKPFKRPRM
```

E)

```
S288c   YBR245C                               X MTASEEYLEN LKPFQVGLPP H.DPESNKKR YLLKDA.NGK
YJM1463 YBR245C                                 MTASEEYLEN LKPFQVGLPP H.DPESNKKR YLLKDA.SGK
S. paradoxus                                    MTASEEYLEN LKPFQVGLPP H.DPESNKKR YLLKDS.SGK
S. mikatae.                                     MTASEEYLEN LKPFQAGVPP H.DPESNKKR YLLKDS.NGK
S. uvarum                                       MTASEEYLEK LKPFQVGVPS R.DPESNKER YLLKDA.NNK
S. kudriavzevii                                 MTASEEYLEN LKPFQVGLPP H.DPETNKKR YLLKDA.NGK
A. gossypii AFL040W                              MENQEYAEK LRPFQEGLGH KPDAVETRAR YLVGGGQNRR
where   X=MAYMLAIANF HFFKFYTRMR KKHENNSCNE KDKDENLFKI ILAIFLQEKK KYDCISSGSI
```

F)

```
S288c   YGL086W              MDVRAALQCF FSALSGRFTG KKLGLEIYSI QYKMSNSGGS SPFLESPGGS PDVGSTNGQS
YJM1460 YGL086W                                                  MSNSGGS SPFLESPGGS PDVGSTNGQS
S. paradoxus                                                     MSNSGGS SPFLESPGGS PGAGSASGQS
A. gossypii AFL065C                                              MSQTGDS SSFVEEP.AS PEARQR..EL
```

G)

```
S288c    YGR079W                   XMNK SSKAFQVPNK VITKEDITPL SRSHTKKADT RGTADGKNTT
YJM1388  YGR079W                    MNK SSKAFQVPNK VITKEDITPL SRSHTKKADT RGTADGKNTT
S. paradoxus                        MNK SSRAFQAPNK VITKEDITPL SRSHTKKADA RGTADGNNTT
A. gossypii ABR106C                MLQQ GATASRRAGR RYTNEYVETV GSGMTAMRGS KRRTQQFQIT
YGR079W  X= MSSAANEGCVYLFIVVLRLSSFSCVNSFIHSFTRSRTRSSYSLDERSLVSYSIVYVA
```

H)

```
S288c YGR140W CBF2    MRSSILFLLK LMKIMDVQQQ QEAMSSEDRF QELVDSLKPR TAHQYKTYYT KYIQWCQLNQ
YJM1273 CBF2                                   MSSEDRF QELVDSLKPR TAHQYKTYYT KYIQWCQLNQ
S. mikatae CBF2                                MSSEDKF QELVDSLKPR TAHQYKTYYT KYIQWCQLNQ
S. uvarum CBF2                                 MSPKDKF QELVDSLKPR TAHQYKTYYT KYIQWCQLNQ
S. kudriavzevii                       MYSTI EEPMSSEDKF QELVDSLKPR TAHQYKTYYT KYIQWCQLNQ
A. gossypii AFR323W CBF2                         MELQL DRAFSRFSSR DVHRYKSYVK QYLEWCRERG
K. waltii CBF2                            M SPSPAPNPQI QALVDTLAPR TAHQYKLYHS KYIQWCRDNN
Z. rouxii CBF2                                    MEEI RKLVDTLSPR TAHQYRSYCK RYVEWLVQKG
```

I)

```
S288c   YGR174C                 MQCAITPREA VIAKQRQYKH YLGMERPLWV RWLKVYAIGG AIIGSGFLLF
YJM1252 YGR174C                 MQCAITPREA VIAK*RQYKH YLGMERPLWV RWLKVYAIGG AIIGSGFLLF
S. paradoxus                                             MERPLWV RWLKVYAIGG AIIGSGFLLF
S. mikatae                                               MERPLWV RWLKVYAIGG AIIGSGFLLF
S. kudriavzevii                                          MERPLWV RWLKVYAIGG AIIGSGFLLF
A. gossypii AAL142C                                      MERPVWF RWLRVYAIGF GIIGAGVLLF
```

J)

```
S288c YML126C               M KLSTKLCWCG QQLHNTNLQM TELKKQKTAE QKTRPQNVGI
S. cerevisiae BPN_2-22481   M KLSTKLCWCG IKGRLRPQKQ QQ*HNTNLQM TELKKQKTAE QKTRPQNVGI
S. uvarum                                                    M TELKKQKTTE QKARPQNVGI
S. mikatae                                                   M TELKKQKTAE QKARPQNVGI
S. kudriavzevii                                              M TELKKQKTTE QKARPQNVGI
A. gossypii ADL356C                                       MTDP NQAKKQKIEA QPPRPQNIGI
```

Where BPN_2-22481 is from (Peter et al. 2018)

K)

```
S288c          YJR107W                                                    M PVVHCSSNLP
Proposed S288c YJR107W  (10/93 strains)                   MMPARL FILYLFVVLM PVVHCSSNLP
YJM1190        YJR107W  (83/93 strains                    MMPARL FILYLFVVLT PVVHCSSNLP
S. paradoxus                                              MMSARL FILYLFVAFT PVVRCSSNLP
S. mikatae                                                MISTRI FILYLFITLA PFINCSPNLS
A. gossypii ABR159C  MHLMMILTLL CLFGRFGRGL APRLGGLVGI GGVADVGGEE LNSSALQTSE TLAAEASHVT
```

L)

```
S288c   YJR126C           MRMIQRERKR EKEEGQLKER TVVNMADPDD NEAEATGLQQ YS...GETTR
YJM1326 YJR126C                                     MADPDD NEAEATGLQQ YS...GETTR
S. paradoxus                                        MADPDD NEAEATGLQQ YS...GETTR
S. mikatae                                          MADPDD NEAEATGLQH YS...GETTR
S. kudriavzevii                                     MTDPDD NEAEATGLQQ YS...GETIL
S. uvarum                                           MVEPDD NEAEATGLQQ YS...GDSTR
A. gossypii AER157C                                 MATNEN LVDETNPLLP RPNTFAERQV
```

M)

```
S288c   YJR127C RSF2                                                 XMFVNG NQSNFAKPAG QGILPIPKKS
YJM1252 YJR127C RSF2                                                  MFVNG NQSNFAKPAG QGILPIPKKS
S. paradoxus    RSF2                                                  MFVNG NQSNFAKPAG QGILPIPKKS
S. mikatae.     RSF2                                                  MFVNG TQSNFAKPAG QGILPIPKKS
S. kudriavzevii RSF2                                                  MFVNG NQSNFAKPAG QGILPIPKKS
S. uvarum       RSF2                                                  MFANG NQSNFAKPTG HGILPIPKKS
A. gossypii AER159C   MELRSAESD GSGRAGATGR QKTTDSVRGR AGGKHSGVIV MRHTPGVSKE TGARGGGRED DGIVPIPKKS
S288c YML081W   TDA9                       MSSEEFKGLP IKRDISSTIY ADRPPALSAP PCVGATGNDK IQVLPIPKKS
S. paradoxus.   TDA9                       MSSEEFKGLP IKRDISSTIH TDRPPALSAP PCVGSTGNDK IQVLPIPKKS
S. mikatae      TDA9                       MSSEDFKELP IKRDIFSTIL VDRPPSLSAP PCVGSTGDDE TQVLPIPKKS
S. kudriavzevii TDA9                       MSSEEFKGLP FKRDIFSTGP ADRPPAFSAP PCVISTGNDE AQVLPIPKKS
S. uvarum       TDA9                       MSSEEFKGLP IKRDLSATDH ADRPPALSAP PCVISVAGDE LQVLPIPKKS
X= MEPFAFGRGAPALCILTAAARINLDNFVPCCWALFRLSFFFPLDPAYIRNENKETRTSWISIEFFFFVK
    HCLSQHTFFSKTLAPKRNFRAKKLKDIGDTRIDRADKDFLLVPEPS
```

N)

```
S288c   YKL021C        XMSAIGD KNQFRIIVGS YEHNILCLSL DIPNQKENDA AKTPHFMPIF
YJM1574 YKL021C         MSAIGD KNQFRIIVGS YEHNILCLSL DIPNQKENDA AKTPHFMPIF
S. paradoxus            MSGIGD KNQFRIIVGS YEHNILCLSL DIPTQKENDA AKTPHFIPIF
S. mikatae              MSDIGG KNQFRIIVGS YEHNILCLSL DIPTQKKDDV AKTPHFMPVF
S. kudriavzevii         MSDIEG KNHFRIIVGS YEHNILCLSL DIPVQKKDDA TKTPHFMPIF
S. uvarum               MSGIEG KNQFRIIVGS YEHNILCLSL DIPVQKKNDE AKTPHFMPIF
A. gossypii ACR166W     MAG     NQFRIVVGS YEHNLLCLSL        NLE LETPLFTPIF
X= MLLNTTYTVFIFFLLRIIKFKIKAENFLEHEKLKTHGTSSRKDPKGQVKNSCRK
```

O)

```
S288c   YKR056W        XMRC FWTVQKSIFK ARFFACRNFV KKHNYKLIST MTGSTEMVPP
YJM1463 YKR056W         MRC FWIVQKSIFK ARFFACRNFV KKHNYKLIST MTGSTEMVPP
S. paradoxus            MRC FWVIQKPIFR SRFFVCKNFV KKHNYTLTAT MTENTGTALP
S. mikatae              MRR FWIIRKPVFR SSFYTCNSFI KKYYYTLITT MTDNTTTTSL
S. kudriavzevii         MRQ FWLLQRPIFR SSLYFRHNLT KSHYCAPIFG MTENITTPAS
S. uvarum               MLR FWISKKPIFR PSYYLYRNIV KNYCCKPITK MTETTTKETS
A. gossypii ACR256C     MFI RRTVLKSIPP LQFHLLRRYT MATDPKLATF PAGSPANVGI
X= MYEQFEFSFFFFENSDNKVKYKAHLISSIKRWSIITC
```

P)

```
S288c  YLR119W     MKVKATKLRI KQRRKNKGLN ISRLDIIRAE MDVVPSPGLP EKVNEKSK     NIPLPEGI
YJM195 YLR119W                                      MDVVPSPGLP EKVNEKSK     NIPLPEGI
S. paradoxus                                        MDVIPSPSAQ KRVNEKNK     NIPLPEGI
S. mikatae                                          MNAASAPSVQ EGVNRRNT     NIPLPEGI
S. uvarum                                           MDMPPPPDVH ERTQKQNK     NIPLPEGI
S. kudriavzevii                                     MDVLPTLSIQ EKDNERNDKR NDSIPLPEAI
A. gossypii AGL189C                                 MGTPVLPPKP CAAAGGPR    AEVPLPRKA
```

Q)

```
S288c   YLR176C     MVIFKERKPT ENLFTRKIPA KYFIFSPSFL SVHYFEFYLP MSGDNNIEPT SRGSNDNSNG
YJM1419 YLR176C                                                 MSGDNNIEPT SRGSNDNSNG
S. paradoxus                                                    MSGDNNIEPT SRGSNDNSNG
S. mikatae                                                      MSDNNKIEPT RRGSNDNSNG
A. gossypii AAL010W              MNVDQASQ EPWASASLLN RSTPSTSTGT PLNTAQSSLP SAPTTSNSHG
```

R)

```
S288c   YLR218C         XMSETGETSEY YKQALEEYKE VQEDEDPDVW DTRISKTGCY
YJM1383 YLR218C          MSETGETSEY YKQALEEYKE VQEDEDPDAW DTRISKTGCY
S. paradoxus             MSETGETSEY YKQALEEYKE VQEDEDPDAW DTRISKTGCY
S. mikatae               MSETENNSEY YKQALEEYKE VQEDEDPDVW DTRISKTGCY
S. kudriavzevii          MSKTGKNSEY YKQALEEYKE VQEDEDPDVW DTRISKTGCY
S. uvarum                MSSVGENSEY YKQALEEYKE VQEDNDPDVW DSRISKTGCY
A. gossypii ADR089C      MAG...DTEY YKQAVEEYAA LKQDTDPEEW DRRIAQTGCY
X= MLLFSSFFSHNKCPKQKKGPQPLKVVKGTSKSCAKQGKIKGKEKLWQAKTGRLV
```

S)

```
S288C   YLR408C     MFLTFSMCVN WIIVKMPNRS EELDRLLDKI INSPHRTEAS KTLQEIENNQ
YJM1401 YLR408C                     MPNRS EELDRLLDKI INSPHRTEAS KTLQEIENNQ
S. paradoxus                        MPNRT EELDRLVDKI INSPHRTEAS KTLEEIENNQ
S. mikatae                  MT YMIVKMPNRT EELDRLLDRI INSPQGTEAS KTLEEIENNQ
S. kudriavzevii                     MPSRT EELDRLLEKI INSPHGTEAS KTLEEIENNQ
S. uvarum                           MSNRT EELDRLLDNI INSPHGTEAS KTLEEIENNQ
A. gossypii  AER079C                      MELDQDVNRI IKSKTATKVS AMQAEIEANH
```

T)

```
S288c   YMR019W                   MTAIG NTDDALDT.. .STAASKENG KGRLRVQKAC ELCKKRKVKC
YJM1383 YMR019W                   MTAIG NTDDALDT.. .STAASKENG KGRLRVQKAC ELCKKRKVKC
S. paradoxus                      MTAIR NTGDALET.. .STAASKENG RGRLRVQKAC ELCKKRKVKC
S. kudriavzevii    MTFMLTENR KRFREMTAIG NTDDALQN.. .VAAASKENS RGRLRVQKAC ELCKKRKVKC
S. uvarum                         MTAIG NIDDALPT.. .NTAASNENS KTRLRVQKAC ELCKKRKVKC
S. mikatae            MFSYER NRFREMTTIG NTTDALEINA AAAAVSKENG RGRLRVQKAC EICKRRKVKC
L. kluyveri                       MDPSK YNDPSLYG.. .....KDNGS KQRLRVRKAC EICKKRKIKC
X= MSINHEIYYILVFEHRSVAIKLIIVVIVLLQFFLARSRQIDRTWAHTNRKERFRE
```

U)

```
S288c  YNR020C                           XMNSSGDN AGFEWWRRTM QYKTGIGLTP EEKTRYEDDS KARELKKECL
YJM248 YNR020C                            MNSSGDN AGFEWWRRTM QYKTGIGLTP EEKTRYEDDS KARELKKECL
S. paradoxus                              MGTSGNN AGFEWWRRTM QYKTGIGLTP EEKTRYEEDS KARALKKECL
S. kudriavzevii                           MGTNESG TGFEWWRRTM QYKTGLGLTP EEKARYEEDS KVRALKKECL
S. uvarum                                 MGANENE TGFEWWRRTM QYKTGLGLTP EEKGRYEEDS KARALKKECL
S. mikatae                                MGTGGND TGFEWWRRTM QYKTGIGLTP EEKIRYEEDS KARALKLECL
A. gossypii AAR064W   MSKEP NFPPSEKLEK VEAPADLRSI AGFEWWRRTF EYKTGLGLTP EAKVQYEKDY QYVLQREQCK
X= MTMRTNKKNKSSNIKMIRRVRHNLYNQNQGKLIVPNFFFFWHD
```

V)

```
S288c   YNR056C       MNRVGAVFLF VYERNFFLSI VPDRHRTEIR MSSSERSEVK FDKHFNWWSL
YJM1479 YNR056C                                        MSSSERSEVK FDKHFNWWSL
S. paradoxus                                           MSSSERSEVK FDKHFNWWSL
S. mikatae                   MPM YIREKLHLSL IPDHHITGVK MSSSERSEVK FDKHFNWWSL
S. kudriavzevii                                        MSSSERPEVK FDKHFNWWSL
S. uvarum                                              MSSSETSEVK FDKHFNWWSL
K. lactis                                   MSSVSSCSLG TTSVSQKEGK LARNYSWWPL
```

W)

```
S288c   YOR241W        XMIIKNP LRMVGKTYRD AVTALNSLQS NYANIMAIRQ TGDRKNTMTL
YJM1418 YOR241W         MIIKNP LRMVGKTYRD AVTALNSLQS NYANIMAIRQ TGDRKNTMTL
S. paradoxus            MIIKNP LRMVGKTYRH AVTALNSLQS NYANIMAIRQ TGDRKNTMAL
S. mikatae              MIIRNP LRMVGKTYHD AVTALNSLQS NYANIMAIRQ TGDRKNIMTL
S. kudriavzevii         MIIRNP LRMIGKTYHD AVTALNSLQS NYANIMAIRQ TGDRKNIMTL
S. uvarum              YMLIRNP LRMIGKTYHD AVTALNSLQS NYANIMAIRQ TGDRKNIMTL
A. gossypii ACR134W          M TRMTRRGYQE AIGCLNGLQS NYANVMAVRA SGERRNMMNL
X=MHKGKKNYPNLITSFRMNLKKIILNHDRFSHPERWKTNALLRFTFVYIKFLFDL
Y=MNLKIKFDHDSFSHLERWKTKTFDRFAFVH
```

X)

```
S288c  YHR175W                  XMDSMNHKMEGNAGHDHSDMHMGDGDDTCSMNMLFSW
YJM248 YHR175W                   MDSMTHKMEGNADHDHSGMHMGDGDDTCSMNMLFSW
S. paradoxus                     MDSMTHKMEGNADHDHSGMHMGDGDDTCSMNMLFSW
S. mikatae                       MESSTHRMEVSAGHDHSGMHMGDEDDMCSMNMLFSW
S. kudriavzevii      MTTTSFVGDTKKVRSPTHMMEASTGHDHDGMHMGDGDDTCSMNMLFSW
S. uvarum                        MFKS   MEGSTDHEHGGMHMGDGDDTCSMNMLFSW
A. gossypii AER293C              MSD  TVHKHEAGH SGHEHV   AHSCAMNTSFTW
X=MDDKKTWSTVTLRTFNQLVTSSLIGYSKK
```

Y)

```
S288c  YIR017C         MSAKQGWEKK STNIDIASRK GMNVNNLSEH LQNLISSDS
YJM195 YIR017C                                MNVNNLSEH LQNLISSDS
S. paradoxus                                  MNVNNLSEH LQNLISSDS
S. mikatae                                    MNVNNLSEH LQNLISSDS
S. kudriavzevii                               MNVNNLSEH LQNLISSDS
S. uvarum                                     MNVNNLSEH LQSLISNDS
A. gossypii AAR010W                          YDE RQLSMQ LQDLIANNS
Y=MSSSVYERPAKQGAEDTEELAPLSPEAYRLKYKGIYSRLISDIWDDAPETAESLGMADSSSQPW
   SAGDVRNTDSPGAKARAATADVTDKAEGQGGGDDSFEKRPLSSVTLNLS
```

Figure S22 Evolutionarily conserved overlap of orthologs of *YPL360W* (*VPS38*) and *YPL361C* (*DCR2*).

A) Alignment of C terminal ends *YPL166W* (*ATG29*) orthologs from *S. cerevisiae, A. gossypii, Z. rouxii, K. lactis.*

B) Alignment of C terminal ends *YPL165C* (*SET6*) orthologs from *S. cerevisiae, A. gossypii, Z. rouxii, K. lactis.*

C) Forward and reverse translation across overlap region of *YPL166W* (*ATG29*) and *YPL165C* (*SET6*) for these four species.

D) Alignment of C terminal ends *YPL360W* (*VPS38*) orthologs for six species.

E) Alignment of C terminal ends *YPL361C* (*DCR2*) orthologs for six species.

F) Forward and reverse translation across overlap region of *YPL360W* (*VPS38*) and *YPL361C* (*DCR2*) for six species.

A)

```
S. cerevisiae     SSLSVSKSALEEALMDRLQF*
A. gossypii       SSLSVSKSALEEALMDRLQL*
Z. rouxii         SSLSVSNSALEEALMDRLKL*
K. lactis         SNLSVSKSALEDALMDCLQL*
```

B)

```
S. cerevisiae     EQICIDYSGVLDLPTVKRRAFLADSWFFDCACERCKSELQSVH*
A. gossypii       EQLCIDYKGILDLPVVERRNILHSNWFFDCACERCALELQSIH*
Z. rouxii         TQLCIAYNCDLTLPVSGRQQTMKDNWFFECMCDRCSSELQTVH*
K. lactis         EELCIDYKGILHLDVNRRRAILKDNWFFECKCSRCLSEL*
```

C)

```
S. cerevisiae S288c
                                ------------------------
          TAAATCTGCGTTGGAAGAAGCGCTAATGGACAGATTGCAATTCTGATTTACATCTTTCAC
          -- ATG29 ------------------------------------>
+ strand   K  S  A  L  E  E  A  L  M  D  R  L  Q  F  *  F  T  S  F  T
- strand  T  F  R  R  Q  F  F  R  *  H  V  S  Q  L  E  S  K  C  R  E
                                <-------- SET6 -----------------------
A. gossypii
                                ------------------------
          CAAGTCTGCATTGGAAGAGGCATTAATGGATCGACTGCAACTCTAGAGCGCAGCGCTCGC
          -- A. gossypii ATG29 ------------------------>
+ strand   K  S  A  L  E  E  A  L  M  D  R  L  Q  L  *  S  A  A  L  A
- strand  A  L  R  C  Q  F  L  C  *  H  I  S  Q  L  E  L  A  C  R  E
                                <-------- A. gossypii SET6 -----------
Z. rouxii
                                ------------------------
          TAACTCAGCTCTGGAAGAAGCACTAATGGACCGTTTGAAGCTCTGAACTGCATCTATCAC
          --Z. rouxii ATG29 --------------------------->
+ strand   N  S  A  L  E  E  A  L  M  D  R  L  K  L  *  T  A  S  I  T
- strand  T  V  *  S  Q  F  F  C  *  H  V  T  Q  L  E  S  S  C  R  D
                                <-------- Z. rouxii SET6 -------------

K. lactis
                                            ------------
          CAAGTCAGCACTTGAGGACGCATTGATGGATTGTTTACAACTCTGAGAGGCATCTACTAC
          -- K. lactis ATG29 -------------------------->
+ strand   K  S  A  L  E  D  A  L  M  D  C  L  Q  L  *  E  A  S  T  T
- strand  A  L  *  C  K  L  V  C  Q  H  I  T  *  L  E  S  L  C  R  S
                                            <-- K. lactis SET6 -------
```

D)

```
S. cerevisiae      DSQYPLYFTDQM--ISKHQAKLIKAIHYFNADILQFKQILENYRPT*
A. gossypii        DNKFPLYLPTSS--TQHHLDRFSEAVHLFNINIIQVRQFLNNLKDK*
E. cymbalariae     NNVLPLYLTASP--TQQHIESFSTAVHLFNMNLMQVKQYLYNHRG*
K. lactis          DGELPFYALTKA--NQQHNSKLKMATARFNINVNQLRQYMEHHR*
Z. rouxii          NGTTPLYLDN-----IKDIELLSQAIDCFNGNIIQVIQFLKHRRTQ*
L. kluyveri        KNQFPLYLTASP--TQQHLANFVTAVSYFNADILQIKQYLEHHR*
C. glabrata        GRKYRLSFPESR—STKSKEEFHIALRAFNENIMQFTQYLGHITQSFTI*
```

E)

```
S. cerevisiae      KIWLCYGGGGGEGGYAGYG-GTERRIRIYEINVNENNIHTWKRLNGSPKEIFDFQSMLDGNSPESV*
A. gossypii        KIWMCYGGAAGEGGYGGYG-GTERRIRIFEIDTREKRIETWKRLNSSPNDKFDAHLI----YP*
E. cymbalariae     SLWLCYGGAVGERGYAGYG-GTERRIRIFEFNAKKKVINTWKRLHGSPDRKFDFQ-IIHDGYTDTA*
K. lactis          KIWLCYGGAAGEGGYAGYG-GTERRIRVYEIDALKKDIYSWKRLNGSPENTFDHQKLISDGVPYTA*
Z. rouxii          KIWFCYGGAVGEGGYGDHNDGYERRVRIYHFETKDGNIYTWKRLNSSPINYFDYQLIASGGVSKTG*
L. kluyveri        KIWLCYGGGGGEGGYAGYG-GTERRIRIFEIDTQGNSIYTWKRLNGTPDEIFDHQTIVSDGVPNTA*
C. glabrata        NIWLCYGGGAGEGGYAGYG-GTERRIRTYKFDPQTGKITTWKRLNSKPSEIFDFQIVKD*
```

F)

```
S. cerevisiae S288c
          TATTTTAATGCTGATATATTGCAATTTAAACAGATTCTGGAGAATTACCGTCCAACATAGACTGAAAATCAA
          -------------------VPS38----------------------------------->
          Y  F  N  A  D  I  L  Q  F  K  Q  I  L  E  N  Y  R  P  T  *  T  E  N  Q
          N  N  *  H  Q  Y  I  A  I  *  V  S  E  P  S  N  G  D  L  M  S  Q  F  D
                                   <------------------DCR2------------------------

A. gossypii
          CAAGTTCGGCAATTTCTAAATAATCTCAAGGATAAATGAGGTGGGCATCAAATTTATCGTTTGGTGAACTGT
          -------------------VPS38-------------->
          Q  V  R  Q  F  L  N  N  L  K  D  K  *  G  G  H  Q  I  Y  R  L  V  N  C
          E  L  E  A  I  E  L  Y  D  *  P  Y  I  L  H  A  D  F  K  D  N  P  S  S
                                   <------------------DCR2------------------------

E. cymbalariae
          TTGTTCAATATGAATCTAATGCAAGTTAAGCAGTATCTGTATAACCATCGTGGATAATTTGAAAATCAAATT
          -------------------VPS38-------------------------------->
          L  F  N  M  N  L  M  Q  V  K  Q  Y  L  Y  N  H  R  G  *  F  E  N  Q  I
          S  T  *  Y  S  D  L  A  L  *  A  T  D  T  Y  G  D  H  I  I  Q  F  D  F
                                   <------------------DCR2------------------------

K. lactis
          AGGTTCAATATAAACGTCAATCAGCTCAGGCAGTATATGGAACACCATCGCTGATTAGTTTTTGATGATCGA
          -------------------VPS38----------------------------->
          R  F  N  I  N  V  N  Q  L  R  Q  Y  M  E  H  H  R  *  L  V  F  D  D  R
          L  T  *  Y  L  R  *  D  A  *  A  T  Y  P  V  G  D  S  I  L  K  Q  H  D
                                   <------------------DCR2------------------------

Z. rouxii
          TGTTTTAACGGTAATATAATACAAGTTATCCAGTTTTTGAAACACCGCCGGACGCAATGAGCTGGTAATCAA
          -------------------VPS38----------------------------------->
          C  F  N  G  N  I  I  Q  V  I  Q  F  L  K  H  R  R  T  Q  *  A  G  N  Q
          N  N  *  R  Y  Y  L  V  L  *  G  T  K  S  V  G  G  S  A  I  L  Q  Y  D
                                   <------------------DCR2------------------------

S. kluyveri
          TACTTCAACGCTGACATACTACAGATCAAGCAGTATTTGGAACACCATCGCTGACGATGGTTTGGTGGTCAA
          -------------------VPS38----------------------------->
          Y  F  N  A  D  I  L  Q  I  K  Q  Y  L  E  H  H  R  *  R  W  F  G  G  Q
          N  S  *  R  Q  C  V  V  S  *  A  T  N  P  V  G  D  S  V  I  T  Q  H  D
                                   <------------------DCR2------------------------

C. glabrata
          TTACACAGTATCTAGGTCATATTACTCAGTCCTTTACTATCTGAAAATCAAAAATCTCACTTGGTTTGCTAT
          -------------------VPS38------------------->
            T  Q  Y  L  G  H  I  T  Q  S  F  T  I  *  K  S  K  I  S  L  G  L  L  F
          N  V  C  Y  R  P  *  I  V  *  D  K  V  I  Q  F  D  F  I  E  S  P  K  S
                                   <------------------DCR2------------------------
```

Figure S23) Single stop codon in *YLR046C* of strains YJM1388 and YJM1592, length 270 amino acids, arises at amino acid 64 by inversion at 9 bp micro-homology.

A) DNA sequence and translation across sites of inversion in *YLR046C* in strains YJM1388 and YJM1592.

B) Protein alignment of strains S288c and YJM1388 of gene *YLR046C*.

A)

```
                   130       140       150       160       170       180
          ----:----|----:----|----:----|----:----|----:----|----:----|
          GTTTTAGGGATTTGTTTAGAGTTCGTGGGTTATGTGTTGAAAATAGTCTTTATCAATAGA
                                                            -------->
          V  L  G  I  C  L  E  F  V  G  Y  V  L  K  I  V  F  I  N  R
                   190       200       210       220       230       240
          ----:----|----:----|----:----|----:----|----:----|----:----|
          AGCACCGATTGAGCAATGAAAGTCGAAATCGAGGGGCTATTGATAGCACCTTCACTTTAC
                                              <--------
          S  T  D  *  A  M  K  V  E  I  E  G  L  L  I  A  P  S  L  Y
```

B)

```
Frame = -3 of YJM1388 chromosome XII

S288c   1       MATHTSKRRIHRWENNELSEENSTIIYFPARGLMWTHFPFVLGICLEFVGYVLKIVFINS  60
                MATHTSKRRIHRWENNELSEENSTIIYFPARGLMWTHFPFVLGICLEFVGYVLKIVFIN
YJM1388 220184  MATHTSKRRIHRWENNELSEENSTIIYFPARGLMWTHFPFVLGICLEFVGYVLKIVFINR  220005

S288c   61      PSISTFIAQSVLLLIAPSLYALSIFMLFSKMARLILMEAYMLIPAKFSTVSFVVADMIGR  120
                 +      +   LLIAPSLYALSIFMLFSKMARLILMEAYMLIPAKFSTVSFVVADMIGR
YJM1388 220004  STD*AMKVEIEGLLIAPSLYALSIFMLFSKMARLILMEAYMLIPAKFSTVSFVVADMIGR  219825

S288c   121     VLQAVGGGLLSSWNSRNTGRILIIVGLFIQIFCYTFLTFSQLFLHYKMKATPSKIVRDSN  180
                VLQAVGGGLLSSWNSRNTGRILIIVGLFIQIFCYTFLTFSQLFLHYKMKATPSKIVRDSN
YJM1388 219824  VLQAVGGGLLSSWNSRNTGRILIIVGLFIQIFCYTFLTFSQLFLHYKMKATPSKIVRDSN  219645

S288c   181     EWFQYNFILLAGILLVNGRTIVRVVQFLMGLQSYIGQHEWCLYVFDTVLMFLLPLIFLAT  240
                EWFQYNFILLAGILLVNGRTIVRVVQFLMGLQSYIGQHEWCLYVFDTVLMFLLPLIFLAT
YJM1388 219644  EWFQYNFILLAGILLVNGRTIVRVVQFLMGLQSYIGQHEWCLYVFDTVLMFLLPLIFLAT  219465

S288c   241     FRARNLFKLQDKSVNIQLNKLLDKESVSED*  271
                FRARNLFKLQDKSVNIQLNKLLDKESVSED*
YJM1388 219464  FRARNLFKLQDKSVNIQLNKLLDKESVSED*  219372

Frame = +1

S288c   58      INSPSISTFIAQSVLLLI  75
                INSPSISTFIAQSVLLLI
YJM1388 219961  INSPSISTFIAQSVLLLI  220014
```

Figure S24 DNA homopolymer runs in protein coding genes

A) *YPR157W* (*TDA6*), a non-essential protein coding gene, one of a duplicate pair of genes resulting from the whole genome duplication, with a run of three phenylalanine residues in *S. cerevisiae*. Phenylalanine is encoded by TTT and TTC. Note that the run of Ts at this location seems to be conserved across the *sensu stricto* species.

B) *YLR075W* (*RPL10*) an essential ribosomal protein with a run of three lysines. Note that none of the species listed has a run of more than five As at this location.

C) Coding frequencies of the two codons used for lysine (AAA and AAG) and phenylalanine (TTT and TTC) in essential genes. Essential genes obtained using the yeastmine tool at SGD.

D) Preference for AAA vs AAG in runs of two lysine residues in essential genes [measured number] (expected number). The expected number of the two-lysine coding sequence is bases on the codon frequencies from C). Underlines show sequences observed less frequently than expected based on the codon usage.

E) Preference for AAA vs AAG in runs of three lysine residues in essential genes [measured number] (expected number). The expected number of the three-lysine coding sequence is bases on the codon frequencies from C). Underlines show sequences observed less frequently than expected based on the codon usage.

F) Preference for TTT vs TTC in runs of two phenylalanine residues in essential genes [measured] (expected). The expected number of the two-lysine coding sequence is bases on the codon frequencies from C). Underlines show sequences observed less frequently than expected based on the codon usage.

G) Preference for TTT vs TTC in runs of three phenylalanine residues in essential genes [measured] (expected). The expected number of the two-lysine coding sequence is bases on the codon frequencies from C). Underlines show sequences observed less frequently than expected based on the codon usage.

H) Coding frequencies of the two codons used for lysine (AAA and AAG) and phenylalanine (TTT and TTC) in nonessential genes. Nonessential genes obtained using the yeastmine tool at SGD.

I) Preference for AAA vs AAG in runs of two lysine residues in nonessential genes [measured number] (expected number). The expected number of the two-lysine coding sequence is bases on the codon frequencies from H). Underlines show sequences observed less frequently than expected based on the codon usage.

J) Preference for AAA vs AAG in runs of three lysine residues in nonessential genes [measured number] (expected number). The expected number of the three-lysine coding sequence is bases on the codon frequencies from H). Underlines show sequences observed less frequently than expected based on the codon usage.

K) Preference for TTT vs TTC in runs of two phenylalanine residues in nonessential genes [measured] (expected). The expected number of the two-lysine coding sequence is bases on the codon frequencies from H). Underlines show sequences observed less frequently than expected based on the codon usage.

L) Preference for TTT vs TTC in runs of three phenylalanine residues in nonessential genes [measured] (expected). The expected number of the two-lysine coding sequence is bases on the codon frequencies from H). Underlines show sequences observed less frequently than expected based on the codon usage.

A)

```
S. cerevisiae                 TGGGCAGCATGCTCATGACATTCCT TTTTTTTTC ATGCCTTTGAGTGATT
S. paradoxus                  TGGGCAACATGCTCACGACATTCCC TTTTTTTTC ATGCCCCTGAGTGATT
S. kudriavzevii               GGGGCAACATGCTCATGATATTCCC TTTTTTTTT ATGCCATTGAGCGACT
S. mikatae                    CGGGCAACATGCTCACGATATTCCC TTTTTTTTT ATGCCCTTGAGTGATT
S. jurei                      GGGGCAACATGCTCACGATATTCCC TTTTTTTTT ATGCCCTTGAGTGATT
S. uvarum                     CGGACAACATGCCCACGATATTCCC TTTTTTTTC ATGCCCTTAAGCGATT
S. eubayanus                  CGGACAGCATGCACACGACATTCCC TTTTTTTTC ATGCCCTTGAGCGATT
Kazachstania naganishii       AGGGCAGCATTCACACGACGTTCCC TTTTTTTTC ATGCCGCTAAGTGACT
Holleya sinecaudum            GGGTCAACATGCCCATGATGTTCCC TTTTTTTTT AGTCCGTTAAGTGACT
A. gossypii                   TGGACAGCATGCACACGACGTGCCA TTTTTCTTT AGCCCGCTCAGTGACT
Ashbya aceri                  TGGACAACACGCGCACGACGTGCCA TTTTTCTTC AGCCCGCTGAGCGACT
Zygosaccharomyces parabailii  TGGGCAACACGCTCACGACGTGCCC TTTTTCTTC ATGCCCTTGAGTGATT
Zygosaccharomyces bailii      TGGGCAACACGCTCACGACGTGCCC TTTTTCTTC ATGCCCCTGAGCGATT
Tetrapisispora blattae        AGGTCAACATCCTCATGATGTTCCC TTTTTCTTT ATGCCATTGAGTGATT
Naumovozyma castellii         CGGGCAACATTCTCATGATGTGCCC TTTTTCTTT ATGCCCTTGAGTGATT
Z. rouxii                     GGGCCAACATGCTCATGATGTTCCA TTTTTCTTT ATGCCACTAAGTGATT
Vanderwaltozyma polyspora     AGGCCAACATCCTCATGATGTACCA TTCTTCTTT ATGCCGTTAAGTGATT
Naumovozyma dairenensis       AGGGCAACATGCACATGATGTACCG TTTTATTTC ATGCCCTTAAGTGACT
```

B)

```
S. cerevisiae                 ATCAGAATCTACGATTTGGGT AAGAAGAAG GCTACCGTCGATGAATTCCC
Z. bailii                     ATTAGAATTTATGACTTGGGT AAGAAGAAG GCCACTGTTGATGAGTTCCC
Z. rouxii                     ATCAGAATCTACGACTTGGGT AAGAAGAAG GCCACCGTCGATGAATTCCC
Candida tanzawaensis          ATCAGAATCTACGATTTGGGT AGAAAGAAG GCCCCTGTCGATGAATTCCC
S. paradoxus                  ATCAGAATTTACGATTTGGGT AAGAAGAAG GCTACCGTCGATGAATTCCC
S. jurei                      ATCAGAATTTACGATTTGGGT AAGAAGAAG GCTACCGTCGATGAATTCCC
S. kudriavzevii               ATCAGAATCTACGATTTGGGT AAGAAGAAG GCCACTGTCGATGAATTCCC
S. eubayanus                  ATCAGAATTTACGATCTAGGT AAGAAGAAG GCCACTGTCGATGAATTTCC
Torulaspora delbrueckii       ATTAGAATCTACGATTTGGGT AAGAAGAAG GCTACTGTCGATGAATTCCC

Kigelia africana              ATCAGAATCTACGATTTAGGT AAGAAGAAG GCTACCGTCGATGAATTCCC
Tetrapisispora phaffii        ATCAGAATCTATGATTTAGGT AAAAAGAAG GCCACAGTCGGAGAATTCCC
V. polyspora                  ATCAGAATTTACGATTTAGGT AAAAAGAAG GCTACTGTTGATGAGTTCCC
T. blattae                    ATCAGAATTTACGATTTAGGT AAAAAGAAG GCTACTGTCGATGAATTCCC
N. castellii                  ATCAGAATCTACGATTTAGGT AAGAAGAAG GCCACTGTCGATGAATTCCC
N. dairenensis                ATCAGAATCTACGATTTAGGT AAGAAGAAG GCTACTGTTGATGAATTCCC
Saccharomycopsis fibuligera   ATCAGAATCTACGATTTAGGT AAAAAGAAG GCCACTGTTGATGAATTCCC
K. lactis                     ATCAGAATTTATGATTTGGGT AAGAAGAAG GCCACTGTCGATGAATTCCC
Candida glabrata              ATCAGAATCTATGACTTGGGT AAGAAGAAG GCCACTGTCGATGAGTTCCC
H. sinecaudum                 ATCAGAATCTACGACTTGGGT AAGAAGAAG GCTACTGTTGATGAGTTCCC
E. cymbalariae                ATTAGAATCTATGACTTGGGT AAGAAGAAG GCCACTGTCGATGAATTTCC
A. gossypii                   ATCAGAATCTACGACTTGGGT AAGAAGAAG GCCACCGTTGATGAGTTCCC
A. aceri                      ATCAGAATCTACGACTTGGGT AAGAAGAAG GCCACTGTCGATGAGTTCCC
Lachancea thermotolerans      ATCAGAATCTACGATCTGGGT AAGAAGAAG GCTACTGTCGATGAGTTCCC
Kazachstania naganishii       ATCAGAATCTACGATTTGGGT AAGAAGAAG GCGACCGTCGATGAGTTCCC
Lachancea lanzarotensis       ATCAGAATCTACGATTTGGGT AAGAAGAAG GCCACCGTTGATGAGTTCCC
```

C) Essential genes:
AAA	K	57.4%

```
AAG    K     42.6%
TTT    F     58.7%
TTC    F     41.3%
```

D)

AAAAAA [1176](1539)

AAAAAG [992](1142)

AAGAAA [1412](1142)

AAGAAG [1094](848)

E)
AAAAAAAAA [27](97)

AAGAAAAAA [61](72)

AAAAAAAAG [35](72)

AAAAAGAAA [83](72)

AAGAAGAAA [67](53)

AAGAAAAAG [78](53)

AAAAAGAAG [68](53)

AAGAAGAAG [96](39)

F)
TTTTTT [334](392)

TTCTTT [202](276)

TTTTTC [389](276)

TTCTTC [214](194)

G)
TTTTTTTTT [3](7)

TTCTTTTTT [1](5)

TTCTTTTTC [2](3)

TTTTTCTTT [6](5)

TTTTTTTTC [8](5)

TTCTTCTTT [5](3)

TTTTTCTTC [10](3)

TTCTTCTTC [2](2)

H) Nonessential genes

```
AAA    K     58.1%
AAG    K     41.9%
TTT    F     59.4%
TTC    F     40.6%
```

I)
AAAAAA [4006](4714)

AAAAAG [2984](3399)

AAGAAA [3965](3399)

AAGAAG [3011](2451)

J)
AAAAAAAAA [136](270)

AAGAAAAAA [192](195)

AAAAAGAAA [191](195)

AAAAAAAAG [91](195)

AAGAAGAAA [206](140)

AAGAAAAAG [184](140)

AAAAAGAAG [154](140)

AAGAAGAAG [227](101)

K)
TTTTTT [1399](1641)

TTCTTT [967](1121)

TTTTTC [1436](1121)

TTCTTC [850](766)

L)

TTTTTTTTT [25](41)

TTCTTTTTT [17](28)

TTTTTCTTT [41](28)

TTTTTTTTC [22](28)

TTCTTCTTT [22](19)

TTCTTTTTC [27](19)

TTTTTCTTC [25](19)

TTCTTCTTC [18](13)

Figure S25 Alignment of a portion of YDR111C (ALT2) nonfunctional alanine transaminase and YLR089C (ALT1) alanine transaminase and the orthologs from other species. Highly conserved glycine residues are shown in bold. 93/93 refers to the 93 *S. cerevisiae* strains discussed in this paper. The single residue substitution creating a potential null allele is underlined.

```
S288c YLR089C              FDNVQLASLHSTSK G VS G EC G QR GG YMELTGFSHEMRQVILKLASISLCPVVTGQALVDL
S. mikatae                 FDNVQLASLHSTSK G VS G EC G QR GG YMELTGFSHEMRQVILKLASISLCPVVTGQALVDL
S. kudriavzevii            FDNVQLASLHSTSK G VS G EC G QR GG YMELTGFSHEMRQVILKLASISLCPVVTGQALVDL
S. uvarum                  FDNVQLASLHSTSK G VS G EC G QR GG YMELTGFSHEMRQVILKLASISLCPVVTGQALVDL
S288c YDR111C              FDNVQLASLHSISK G FM D EC G QR GG YMEIIGFSQEIRDALFKLMSISICSVVTGQAVVDL
93/93 YDR111C              FDNVQLASLHSISK G FM G EC G QR GG YMEIIGFSQEIRDALFKLMSISICSVVTGQAVVDL
S. mikatae                 FDNVQLASLHSISK G FM G EC G QR GG YMEVVGFSKDIRDALFKLMSISICSVVTGQAVVDL
S. kudriavzevii            FDNVQLASLHSISK G FM G EC G QR GG YMEIVGFSQDIRDALFKLMSISICSVVTGQAIVDL
S. uvarum                  FDNVQLASLHSISK G FM G EC G QR GG YMEIVGFSQDIRDALFKLMSISICSVATGQAIIDL
A. gossypii AGR085W        YKNVQLASLHSISK G LS G EC G QR GG YMELIGFREELRKVFVKLASISLCPVVTGQALVDL
Candida albicans           YKHVQLASLHSTSK G VS G EC G QR GG YMELVGFKPEVKDVVFKLASINLCSVVSGQALMEL
Schizosaccharomyces pombe  WDQVSLISVNSVSK G QF G EC G QR GG YLDVVNIPEPAKDQILKLATIDICPPVAGQLLVDM
Cryptococcus neoformans    RESVELVSFHSISK G VS G EC G RR GG YFECVNIDKEVMEQVYKMASVTLCPPVSGQIGVDL
Homo sapiens               SSNVELASFHSTSK G YM G EC G YR GG YMEVINLHPEIKGQLVKLLSVRLCPPVSGQAAMDI
Arabidopsis thaliana       EKDISLVSFQSVSK G YY G EC G KR GG YMEVTGFTSDVREQIYKMASVNLCSNISGQILASL
Escherichia coli           SKEVQLISFHTVSK G YW G EC G QR GG YFEMTNIPPETVDEIYKVASISLSPNVPAQIFMGL
```

Figure S26 Two of four serine GCU tRNA genes, *YNCD0024C* *tX(XXX)D* and *YNCL0039W* *tX(XXX)L* in stain S288c have sequence errors/annotation errors.

A) Alignment of four serine GCU anticodon tRNA genes from *S. cerevisiae* strain S288c. In all 93 strains from Strope et al these four genes are serine GCU anticodon tRNA genes. In S288c two copies as currently annotated are missing one base from the anti-codon loop, at a classic Applied Biosystems dye primer sequencing compression site GGC…GCC. This was the primary method used in the original yeast sequencing.

B) Alignment of sequence reads for S288c serine GCU anticodon tRNA gene region spanning the anti-codon site from S288c genomic sequence fastq file SRR24682795, submitted from KU Leuven. No sign of the “GCCT” sequence found in *tX(XXX)D* and *tX(XXX)L* was found in this fastq files or several other S288c and BY4741 genomic fastq files examined suggesting that “GCCT” is a sequence error. These sequence reads are long enough to span from the site of the sequence error to the unique sequence flanking these tRNA genes, showing that sequences were found for all four of the tRNA genes. The numbers at the end of each line indicate the number of identical reads found, and the sign is whether the reads were on the plus strand, relative to the tRNA gene, or the minus strand.

A)

```
tX(XXX)L GTCCCAGTGGCCGAGTGGTTAAGGCGAT GCC  TGCTATTTCCTCAGAAAAGCAATTAGGCATTGGGTTTTACCTGCGCAGGTTCGAATCCTGTCTGTGACG
tX(XXX)D GTCCCAGTGGCCGAGTGGTTAAGGCGAT GCC  TGCTATTTCCTCAGAAAAGCAATTAGGCATTGGGTTTTACCTGCGCAGGTTCGAATCCTGTCTGTGACG
tS(GCU)O GTCCCAGTGGCCGAGTGGTTAAGGCGAT GCCC TGCTATTTCCTCAGAAAAGCAATTAGGCATTGGGTTTTACCTGCGCAGGTTCGAATCCTGTCTGTGACG
tS(GCU)F GTCCCAGTGGCCGAGTGGTTAAGGCGAT GCCC TGCTATTTCCTCAGAAAAGCAATTAGGCATTGGGTTTTACCTGCGCAGGTTCGAATCCTGTCTGTGACG

```

B)

```
YNCL0039W tX(XXX)L
GTCCCAGTGGCCGAGTGGTTAAGGCGAT  GCCC  TGCTATTTCCTCAGAAAAGCAATTAGGCATTGGGTTTTACCTGCGCAGGTTCGAATCCTGTCTGTGACGC  TTTTTTTAATTTCTTTACTCCATGACAAAAGCGGATAAAAATTCCCGC -2
GTCCCAGTGGCCGAGTGGTTAAGGCGAT  GCCC  TGCTATTTCCTCAGAAAAGCAATTAGGCATTGGGTTTTACCTGCGCAGGTTCGAATCCTGTCTGTGACGC  TTTTTTTAATTTCTTTACTCCATGACAAAAGCGGATAAAAATTCCCGC +4
  CCCAGTGGCCGAGTGGTTAAGGCGAT  GCCC  TGCTATTTCCTCAGAAAAGCAATTAGGCATTGGGTTTTACCTGCGCAGGTTCGAATCCTGTCTGTGACGC  TTTTTTTAATTTCTTTACTCCATGACAAAAGCGGATAAAAATTCCCGCAT -1
   CCAGTGGCCGAGTGGTTAAGGCGAT  GCCC  TGCTATTTCCTCAGAAAAGCAATTAGGCATTGGGTTTTACCTGCGCAGGTTCGAATCCTGTCTGTGACGC  TTTTTTTAATTTCTTTACTCCATGACAAAAGCGGATAAAAATTCCCGCACT +1
     AGTGGCCGAGTGGTTAAGGCGAT  GCCC  TGCTATTTCCTCAGAAAAGCAATTAGGCATTGGGTTTTACCTGCGCAGGTTCGAATCCTGTCTGTGACGC  TTTTTTTAATTTCTTTACTCCATGACAAAAGCGGATAAAAATTCCCGCATTCG -1
      GTGGCCGAGTGGTTAAGGCGAT  GCCC  TGCTATTTCCTCAGAAAAGCAATTAGGCATTGGGTTTTACCTGCGCAGGTTCGAATCCTGTCTGTGACGC  TTTTTTTAATTTCTTTACTCCATGACAAAAGCGGATAAAAATTCCCGCATTCGG +1
YNCD0024C tX(XXX)D
GTCCCAGTGGCCGAGTGGTTAAGGCGAT  GCCC  TGCTATTTCCTCAGAAAAGCAATTAGGCATTGGGTTTTACCTGCGCAGGTTCGAATCCTGTCTGTGACGC  ACTTTATTTTTTCTACATCCACTATTTTAAAGCTTGATTTTTTTGATT -1
GTCCCAGTGGCCGAGTGGTTAAGGCGAT  GCCC  TGCTATTTCCTCAGAAAAGCAATTAGGCATTGGGTTTTACCTGCGCAGGTTCGAATCCTGTCTGTGACGC  ACTTTATTTTTTCTACATCCACTATTTTAAAGCTTGATTTTTTTGATT +3
  CCCAGTGGCCGAGTGGTTAAGGCGAT  GCCC  TGCTATTTCCTCAGAAAAGCAATTAGGCATTGGGTTTTACCTGCGCAGGTTCGAATCCTGTCTGTGACGC  ACTTTATTTTTTCTACATCCACTATTTTAAAGCTTGATTTTTTTGATTGT -1
  CCCAGTGGCCGAGTGGTTAAGGCGAT  GCCC  TGCTATTTCCTCAGAAAAGCAATTAGGCATTGGGTTTTACCTGCGCAGGTTCGAATCCTGTCTGTGACGC  ACTTTATTTTTTCTACATCCACTATTTTAAAGCTTGATTTTTTTGATTGT +2
   CCAGTGGCCGAGTGGTTAAGGCGAT  GCCC  TGCTATTTCCTCAGAAAAGCAATTAGGCATTGGGTTTTACCTGCGCAGGTTCGAATCCTGTCTGTGACGC  ACTTTATTTTTTCTACATCCACTATTTTAAAGCTTGATTTTTTTGATTGTG -2
    CAGTGGCCGAGTGGTTAAGGCGAT  GCCC  TGCTATTTCCTCAGAAAAGCAATTAGGCATTGGGTTTTACCTGCGCAGGTTCGAATCCTGTCTGTGACGC  ACTTTATTTTTTCTACATCCACTATTTTAAAGCTTGATTTTTTTGATTGTGA +1
      GTGGCCGAGTGGTTAAGGCGAT  GCCC  TGCTATTTCCTCAGAAAAGCAATTAGGCATTGGGTTTTACCTGCGCAGGTTCGAATCCTGTCTGTGACGC  ACTTTATTTTTTCTACATCCACTATTTTAAAGCTTGATTTTTTTGATTGTGAGC -2
      GTGGCCGAGTGGTTAAGGCGAT  GCCC  TGCTATTTCCTCAGAAAAGCAATTAGGCATTGGGTTTTACCTGCGCAGGTTCGAATCCTGTCTGTGACGC  ACTTTATTTTTTCTACATCCACTATTTTAAAGCTTGATTTTTTTGATTGTGAGC +1
YNCO0008W tS(GCU)O
GTCCCAGTGGCCGAGTGGTTAAGGCGAT  GCCC  TGCTATTTCCTCAGAAAAGCAATTAGGCATTGGGTTTTACCTGCGCAGGTTCGAATCCTGTCTGTGACGC  ATTTCTTTTTTGTGAATTCTTGCTTCTTATCCTTCTACAATCCTTTCG -3
GTCCCAGTGGCCGAGTGGTTAAGGCGAT  GCCC  TGCTATTTCCTCAGAAAAGCAATTAGGCATTGGGTTTTACCTGCGCAGGTTCGAATCCTGTCTGTGACGC  ATTTCTTTTTTGTGAATTCTTGCTTCTTATCCTTCTACAATCCTTTCG +3
 TCCCAGTGGCCGAGTGGTTAAGGCGAT  GCCC  TGCTATTTCCTCAGAAAAGCAATTAGGCATTGGGTTTTACCTGCGCAGGTTCGAATCCTGTCTGTGACGC  ATTTCTTTTTTGTGAATTCTTGCTTCTTATCCTTCTACAATCCTTTCGT -1
  CCCAGTGGCCGAGTGGTTAAGGCGAT  GCCC  TGCTATTTCCTCAGAAAAGCAATTAGGCATTGGGTTTTACCTGCGCAGGTTCGAATCCTGTCTGTGACGC  ATTTCTTTTTTGTGAATTCTTGCTTCTTATCCTTCTACAATCCTTTCGTG -3
  CCCAGTGGCCGAGTGGTTAAGGCGAT  GCCC  TGCTATTTCCTCAGAAAAGCAATTAGGCATTGGGTTTTACCTGCGCAGGTTCGAATCCTGTCTGTGACGC  ATTTCTTTTTTGTGAATTCTTGCTTCTTATCCTTCTACAATCCTTTCGTG +2
    CAGTGGCCGAGTGGTTAAGGCGAT  GCCC  TGCTATTTCCTCAGAAAAGCAATTAGGCATTGGGTTTTACCTGCGCAGGTTCGAATCCTGTCTGTGACGC  ATTTCTTTTTTGTGAATTCTTGCTTCTTATCCTTCTACAATCCTTTCGTGAC -1
     AGTGGCCGAGTGGTTAAGGCGAT  GCCC  TGCTATTTCCTCAGAAAAGCAATTAGGCATTGGGTTTTACCTGCGCAGGTTCGAATCCTGTCTGTGACGC  ATTTCTTTTTTGTGAATTCTTGCTTCTTATCCTTCTACAATCCTTTCGTGACA -1
     AGTGGCCGAGTGGTTAAGGCGAT  GCCC  TGCTATTTCCTCAGAAAAGCAATTAGGCATTGGGTTTTACCTGCGCAGGTTCGAATCCTGTCTGTGACGC  ATTTCTTTTTTGTGAATTCTTGCTTCTTATCCTTCTACAATCCTTTCGAGACA +1
      GTGGCCGAGTGGTTAAGGCGAT  GCCC  TGCTATTTCCTCAGAAAAGCAATTAGGCATTGGGTTTTACCTGCGCAGGTTCGAATCCTGTCTGTGACGC  ATTTCTTTTTTGTGAATTCTTGCTTCTTATCCTTCTACAATCCTTTCGTGACAT -1
      GTGGCCGAGTGGTTAAGGCGAT  GCCC  TGCTATTTCCTCAGAAAAGCAATTAGGCATTGGGTTTTACCTGCGCAGGTTCGAATCCTGTCTGTGACGC  ATTTCTTTTTTGTGAATTCTTGCTTCTTATCCTTCTACAATCCTTTCGTGACAT +1
YNCF0009C tS(GCU)F
GTCCCAGTGGCCGAGTGGTTAAGGCGAT  GCCC  TGCTATTTCCTCAGAAAAGCAATTAGGCATTGGGTTTTACCTGCGCAGGTTCGAATCCTGTCTGTGACGC  ATTTCTTTTTTAAATTTTTGCTTCTTATCCTTCTATAATAAACCTTTC -12
GTCCCAGTGGCCGAGTGGTTAAGGCGAT  GCCC  TGCTATTTCCTCAGAAAAGCAATTAGGCATTGGGTTTTACCTGCGCAGGTTCGAATCCTGTCTGTGACGC  ATTTCTTTTTTAAATTTTTGCTTCTTATCCTTCTATAATAAACCTTTC +1
 TCCCAGTGGCCGAGTGGTTAAGGCGAT  GCCC  TGCTATTTCCTCAGAAAAGCAATTAGGCATTGGGTTTTACCTGCGCAGGTTCGAATCCTGTCTGTGACGC  ATTTCTTTTTTAAATTTTTGCTTCTTATCCTTCTATAATAAACCTTTCG -6
  CCCAGTGGCCGAGTGGTTAAGGCGAT  GCCC  TGCTATTTCCTCAGAAAAGCAATTAGGCATTGGGTTTTACCTGCGCAGGTTCGAATCCTGTCTGTGACGC  ATTTCTTTTTTAAATTTTTGCTTCTTATCCTTCTATAATAAACCTTTCGT +2
   CCAGTGGCCGAGTGGTTAAGGCGAT  GCCC  TGCTATTTCCTCAGAAAAGCAATTAGGCATTGGGTTTTACCTGCGCAGGTTCGAATCCTGTCTGTGACGC  ATTTCTTTTTTAAATTTTTGCTTCTTATCCTTCTATAATAAACCTTTCGTA -2
   CCAGTGGCCGAGTGGTTAAGGCGAT  GCCC  TGCTATTTCCTCAGAAAAGCAATTAGGCATTGGGTTTTACCTGCGCAGGTTCGAATCCTGTCTGTGACGC  ATTTCTTTTTTAAATTTTTGCTTCTTATCCTTCTATAATAAACCTTTCGTA +2
    CAGTGGCCGAGTGGTTAAGGCGAT  GCCC  TGCTATTTCCTCAGAAAAGCAATTAGGCATTGGGTTTTACCTGCGCAGGTTCGAATCCTGTCTGTGACGC  ATTTCTTTTTTAAATTTTTGCTTCTTATCCTTCTATAATAAACCTTTCGTAA +1
     AGTGGCCGAGTGGTTAAGGCGAT  GCCC  TGCTATTTCCTCAGAAAAGCAATTAGGCATTGGGTTTTACCTGCGCAGGTTCGAATCCTGTCTGTGACGC  ATTTCTTTTTTAAATTTTTGCTTCTTATCCTTCTATAATAAACCTTTCGTAAC -4
     AGTGGCCGAGTGGTTAAGGCGAT  GCCC  TGCTATTTCCTCAGAAAAGCAATTAGGCATTGGGTTTTACCTGCGCAGGTTCGAATCCTGTCTGTGACGC  ATTTCTTTTTTAAATTTTTGCTTCTTATCCTTCTATAATAAACCTTTCGTAAC +2
      GTGGCCGAGTGGTTAAGGCGAT  GCCC  TGCTATTTCCTCAGAAAAGCAATTAGGCATTGGGTTTTACCTGCGCAGGTTCGAATCCTGTCTGTGACGC  ATTTCTTTTTTAAATTTTTGCTTCTTATCCTTCTATAATAAACCTTTCGTAACA -4
      GTGGCCGAGTGGTTAAGGCGAT  GCCC  TGCTATTTCCTCAGAAAAGCAATTAGGCATTGGGTTTTACCTGCGCAGGTTCGAATCCTGTCTGTGACGC  ATTTCTTTTTTAAATTTTTGCTTCTTATCCTTCTATAATAAACCTTTCGTAACA +1
                              Site             3' end of Serine GCU tRNA gene                               Unique sequence flanking the four Serine GCU tRNA genes
                              of
                              sequence
                              error
```

Figure S27

Alignment of the most conserved portion of the carboxy terminal end of *S. cerevisiae* gene *YMR008C-A* and portions of the putative gene remnants for this gene from *S. paradoxus,* and *S. mikatae,* and *S. jurei.*

```
S. cererivisiae     SAPFVPEIALRKSGAVGS*  156
S. paradoxus        SAPFVPEIALRKAAQRG  277205
S. mikatae           SPYRPETALRNSGAVG  289337
S. jurei                 PRIALRNSGAVG  278271
```

Figure S28

Alignment of *S. cerevisiae* *YFR035C* with *S. paradoxus* and *S. kudriavzevii*.

A) The sequence of *YFR035C* in *S. paradoxus* is missing the start codon, while on the opposite strand the leucine is conserved despite the single nucleotide G->A polymorphism. The start codon of *YFR035W-A* is much more conserved.

```
                 M         I
Forward strand  ATGAGCG   ATAAGCG where ATG/ATA  YFR035C start codon
Reverse strand    CTC       TTC
```

L	L

*S. cerevisiae	S. paradoxus*

B) The sequence of *YFR035W-A* is conserved across the whole open reading frame, whereas conservation of the *YFR035C* is more pronounced in the region overlapping *YFR035W-A* than outside the overlap

```
S. cerevisiae vs. S. paradoxus overlap

Scer YFR035C  MSASDKTKLCNKGMSRTSRTTTFVITPAFRERDDEGANSLCKAFLNTFSNLKSGMFKCLL
              +SASDKTKLCN+GMS TSR TTFVITP FRERDDEGA+SL  AFLNTFSNLKSGM KCLL
Spar YFR035C  ISASDKTKLCNRGMSNTSRITTFVITPNFRERDDEGASSLYNAFLNTFSNLKSGMLKCLL

              |overlap| | not overlapping       |
Scer YFR035C  GVGAVGTFI STFPQFFL--LPCLLCVRCVCVCL
              GVGAVG FI STFPQFFL  LP L   R VC+CL
Spar YFR035C  GVGAVGKFI STFPQFFLAALPSL---RMVCMCL
              <--------^ Start of YFR035W-A, on opposite strand

S. cerevisiae vs. S. kudriavzevii overlap

Scer YFR035C  MSASDKTKLCNKGMSRTSRTTTFVITPAFRERDDEGANSLCKAFLNTFSNLKSGMFKCLL
              MS SDKTKLC+ G+S TSRTTTFVIT    E DDEGA+SLC AFLNTFSNL+SG+F CLL
Skud YFR035C  MSTSDKTKLCSSGISNTSRTTTFVITAHLLETDDEGASSLCNAFLNTFSNLRSGIFICLL

              |overlap| | not overlapping     |
Scer YFR035C  GVGAVGTFI STFPQFFLLPCLLCVRCVCVCL
              GVGA GTFI ST PQFFLLPC L + C C+C+
Skud YFR035C  GVGANGTFI STVPQFFLLPC-LALPCPCMCV
              <--------^ Start of YFR035W-A, on opposite strand
```

Figure S29

A) Alignment of nanopore RNA-seq reads spanning *YNL155C-A* from SRA file SRR11550239 (Jenjaroenpun et al. 2021) displayed using jbrowse2. The sequence reads displayed are all on the coding strand relative to *YNL155C-A*, and terminate after the *YNL156C* stop codon (not shown).

B) Alignment of possible -1 frameshifted and un-frameshifted *YNL156C* proteins. Amino terminal end only is shown. Sarb *S. arboricola*; Scer *S. cerevisiae*; Spar *S. paradoxus*; Skdz *S. kudriavzevii*; Sjur *S. jurei*; Smik *S. mikatae*; Suva *S. uvarum*; Seub *S. eubayanus*; Lklu *L. kluyveri*; Nade *Nakaseomyces delphensis*; Cabr *Candida bracarensis*; _fs with putative frameshift combining *YNL155C-A and YNL156C*; _nofs without frameshift, in-frame start codon proposed to be used.

C) RNA sequence of putative frameshifting site (bold) and downstream sequence, showing inverted repeats.

A)

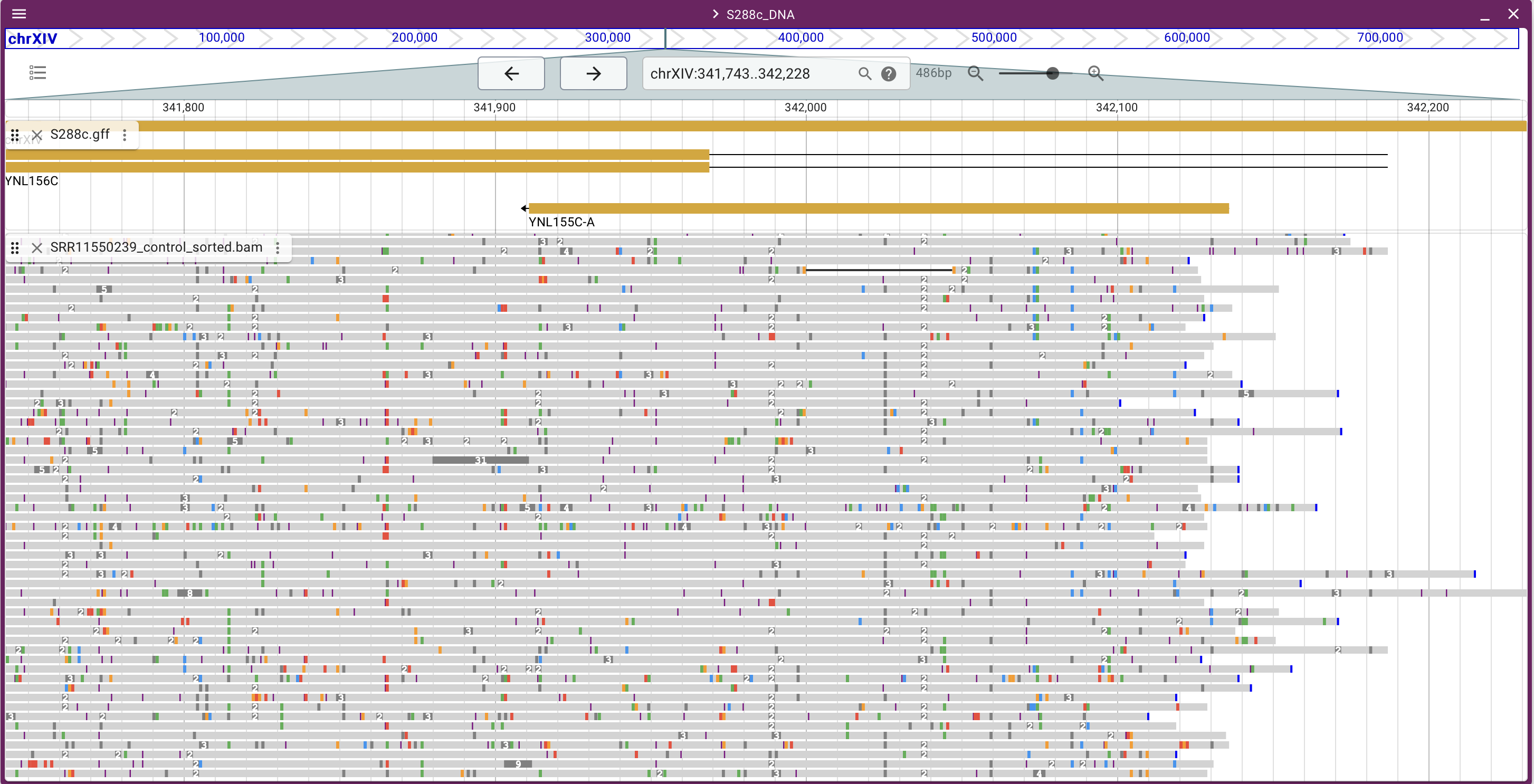

B)

```
Possible frameshift site from seven Saccharomyces species           [AG]GAGAAC
YNL155C-A frame                                                         E  N
YNL156C   frame                                                     [RG]  E  P
                                                                          fs
Sarb_fs   MIGLIVVPILFAIKGIVLGVQKVKKKFGKHSESKEQEGAGKDKRKR--QSISQKKQKTEWPIEEEPDPKKSTESISSLTKPQLYSLYDDDVVRSED
Scer_fs   MIGLIVVPILFAIKGIVVGVQKVKKKFGKHSNTKDQK-EDKDEDKR--QSISQRKQHTEWPIEEEPDPKKSTESICSLTKPQLYSLYDDDVVRSED
Spar_fs   MIGLIVVPILFAIKGIVVGVQKVKKKFGKNSESKDPI-EDKREDKR--QSIPQRKQHTEWPIEEEPDPKRSTESICSLTKPQLYSLYDDDVVRSED
Skdz_fs   MIGLIVVPILFAIKGIIVGVQKVKKKFGKHSEAKNQE-EGKSKNKR--QNSSQKKQETEWPIEEEPDPKRSTESISSLTKPQLYSLYDDDVVRSED
Sjur_fs   MLGLIVVPILFAIKGIIVGVQKAKKKFGKHSESKDQK-EGQSKDKR--QSISQKKQHTEWPIEEEPASEKSTESICSLTKPQLYSLYDDDVVRSED
Smik_fs   MIGLILVPILFAIKGIVVGVQKAKKKFGKHSESKNLK-EGQVKDKR--QDIFQKKQHTEWPIKEEPAPEKSTESICSLTKPQLYSLYDDDVVRSED
Suva_fs   MIGLIIVPIIFAIKGIVVGVQKVKQKFGKHSKTKHQEEEDKEKDKGQGQGISKTKQKTEWPIEEKPDPKKSTESISSLTKPQLYSLYDDDVVRSED
Seub_fs   MIGLIIVPIIFAIKGIVVGVQKVKQKFGKHSKSKHQEEEDRDKYKG--QGITKTKQKTEWPIEEEPDPKKSTESISSLTKPQLYSLYDDDVVRSED
Sarb_nofs --------------------------------------------------MANRR---------EPDPKKSTESISSLTKPQLYSLYDDDVVRSED
Scer_nofs --------------------------------------------------MANRG---------EPDPKKSTESICSLTKPQLYSLYDDDVVRSED
Spar_nofs --------------------------------------------------MANRR---------EPDPKRSTESICSLTKPQLYSLYDDDVVRSED
Skdz_nofs --------------------------------------------------MANRR---------EPDPKRSTESISSLTKPQLYSLYDDDVVRSED
Sjur_nofs --------------------------------------------------MANRR---------EPASEKSTESICSLTKPQLYSLYDDDVVRSED
Smik_nofs --------------------------------------------------MANKR---------EPAPEKSTESICSLTKPQLYSLYDDDVVRSED
Suva_nofs --------------------------------------------------MANRG---------KPDPKKSTESISSLTKPQLYSLYDDDVVRSED
Seub_nofs --------------------------------------------------MANRR---------EPDPKKSTESISSLTKPQLYSLYDDDVVRSED
Lklu      --------------------------------------------------MA-RKKSKA-----GNELSNSNESITTLTKPQLFGLYDEEIVKSED
Nade      --------------------------------------------------MA-KKLKKRGSP--SVASVGSTESISSLTTPQLYSIYDQDIVKSED
Cabr      --------------------------------------------------MA-KKLKKRGSP--PVKSVQSTESISSLTTPQLYSIYDQDIIKSED
                                                                                *.*** :**.***:.:**:::::***
```

C)

```
Possible                   downstream inverted repeats
fs site                    111111                              2222      2222       2222                     2222                    3333311111                            11111
Sarb                       >>>>>>                              >>>>      <<<<       <<<<                     <<<<                    >>>>><<<<<                            <<<<<
AGAGAACCAGAUCCAAAGAAGAGCACUGAAUCAAUCAGUUCAUUAACUAAGCCCCAGCUGUAUUCACUGUACGAUGAUGACGUAGUGAGAUCAGAAGAUAACGAAAUUUAUGAAGAAUUGAAACGAUCCGUGUCUAUAGAUUCUAGCAGGUAUAGUAGAGAUCAAACCAUUGAUUCG
Scer
GGAGAACCGGAUCCAAAGAAGAGCACCGAAUCAAUCUGCUCUCUGACUAAACCGCAGCUUUAUUCAUUAUAUGAUGAUGACGUCGUGAGAUCGGAAGAUAACGAAAUUUACGAAGAAUUGAAACGCUCUGUCUCUAUAGAUUCUACCAAAUAUAGUAGAGAUCAAACUAUUGAUUCG
Spar
AGAGAACCGGAUCCAAAAAGGAGCACCGAAUCGAUCUGCUCUCUGACUAAACCGCAGCUAUAUUCACUAUAUGAUGACGACGUAGUGAGAUCAGAAGAUAACGAAAUUUAUGAAGAAUUGAAACGGUCUGUAUCUAUAGAUUCUGGCAAAUAUAGUAGAGAUCAAACCAUUGAUUCA
Skdz
AGAGAACCGGAUCCAAAAAGGAGCACUGAAUCGAUCAGCUCUUUAACUAAGCCACAGUUAUACUCACUGUAUGAUGAUGACGUCGUGAGAUCCGAAGAUAACGAAAUAUACGAGGAAUUGAAACGAUCCGUCUCUAUAGAUUCCGGCAAGUAUAGUAGAGAUCAAACUAUUGAUGCG
Sjur
AGAGAACCAGCUUCAGAGAAGAGUACCGAAUCAAUCUGCUCUUUAACUAAACCGCAGCUAUAUUCAUUAUAUGAUGAUGACGUAGUGAGAUCAGAAGACAACGAAAUUUAUGAAGAGCUAAAACGCUCUGUCUCUGUAGACUCUAGCAAAUAUAGUAGAGACCAAACUAUCGAUUCA
Smki
AGAGAACCAGCUCCAGAGAAGAGUACCGAAUCAAUCUGCUCUUUAACUAAACCGCAGCUGUAUUCAUUAUAUGAUGAUGACGUAGUGAGAUCAGAAGAUAACGAAAUUUAUGAAGAGUUGAAACGCUCUGUCUCUAUAGACUCUGGCAAAUAUAGUAGAGAUCAAACUAUUGAUUCA
Suva
GGAAAACCGGAUCCAAAAAAGAGCACCGAAUCAAUUAGUUCUUUAACUAAGCCCCAGCUGUAUUCAUUAUAUGACGAUGACGUAGUGAGAUCGGAAGAUAACGACAUUUACGAGGAACUGAAACGAUCCGUCUCCAUAGACUCUGGCAGAUAUAGCAGAGACCAAACGAUUGAUUCA
Seub
AGAGAACCGGAUCCAAAAAAAAGCACCGAAUCAAUUAGUUCUUUAACUAAGCCCCAGCUUUAUUCAUUGUAUGACGAUGACGUAGUGAGAUCGGAAGAUAACGACAUUUAUGAGGAAUUGAAACGGUCUGUCUCUAUAGACUCUGGCAGAUAUAGCAGAGAUCAAACGAUUGAUUCU
Lklu
GCAGGUAACGAAUUAAGCAACAGCAAUGAAUCGAUUACCACACUAACUAAACCCCAAUUGUUUGGGCUAUACGAUGAAGAAAUCGUCAAAAGUGAAGACAACGAAAUUUACGAAGAGGCCAAAAAAUCUGCUGGCAAGCUAACUCCCGCAGGAGUAAUGGAAAGACGCAGUAUUUUC
```

Figure S30

*YJM106W-A* and *YJM106W-B*.

YMR106W-A: S288c *YMR106W-A*; YJM1479_106W-A: *S. cerevisiae* YJM1479 *YMR106W-A*; Spara_106W-A: *S. paradoxus* strain LL2012-028 *YMR106W-A*; YJM1479_106W-B: *S. cerevisiae* YJM1479 *YMR106W-B*; Spara_106W-B: *S. paradoxus* strain LL2012-028 *YMR106W-B*; S288c_106W-B: *S. cerevisiae* S288c *YMR106W-B* truncated allele.

```
YMR106W-A       -------------------MISMEAINNFIKTAPKHDYLTGGVHHSGNVDVLQLSGNKED
YJM1479_106W-A  -------------------MISLEAINNFIKTAPKHDYLTGGVHHSGNVDVLQLSGNKED
Spara_106W-A    -------------------MISMEAINNFIKAAPKHDYLTGGVHHSGNVDVLQLSGNKED
YJM1479_106W-B  MLPFSVLISYLFTFKLVVCAISIETVNDFINRAADHDYVTGGIFESNDVKVLYLGGYQDN
Spara_106W-B    MLPFTVLISYLFTFKLIACAISIETVNDYVNSAADHDYVTGGIFESSDVKVLYLGGYQDN
S288c_106W-B    MLPFSVLVSYLFTFKLVVCAITSKKVN---------------------------------
                ****:**:********:.* **:*::*:::: *..***:***:..*.:*.** *.* :::

YMR106W-A       GSLVWNHTFVDVDNNVVAKF--------------------EDALEKLESLHRR-SSSSTG
YJM1479_106W-A  GTLVWSHTFVDVDNNVVAKF--------------------EDALEKLESLHRR-SSSSTG
Spara_106W-A    GTLVWSHTFVDVDNNVVAKF--------------------KDALEKLEALHR---PSATD
YJM1479_106W-B  ATLVWTHQFIDVDSNVVAKINTGLNILADLASGNITEDIGSDALNKRWSCHQEFCRNAAD
Spara_106W-B    GTLVWTHQFTDVDSNVVAKINSGLNILADLASGNITESVASNGLNKRWSCHQEFCRNAAD
S288c_106W-B    ------------------------------------------------------------
                .:***.* * ***.*****:*:****************:..:.*:*  : *:    .::.

YMR106W-A       NEEHANV----------------
YJM1479_106W-A  NEEQANV----------------
Spara_106W-A    DEDQANV----------------
YJM1479_106W-B  CEKHSCSWCYKGLFRLPLTGICF
Spara_106W-B    CEKHSCSWCYKGLFRLPLTGICF
S288c_106W-B    -----------------------
```

*.:: ****************

**References Supplemental material**

Jenjaroenpun, P., T. Wongsurawat, T. D. Wadley, T. M. Wassenaar, J. Liu, Q. Dai, V. Wanchai, N. S. Akel, A. Jamshidi-Parsian, A. T. Franco, G. Boysen, M. L. Jennings, D. W. Ussery, C. He and I. Nookaew (2021). "Decoding the epitranscriptional landscape from native RNA sequences." Nucleic Acids Res **49**(2): e7.

Kessler, M. M., Q. Zeng, S. Hogan, R. Cook, A. J. Morales and G. Cottarel (2003). "Systematic discovery of new genes in the Saccharomyces cerevisiae genome." Genome Res **13**(2): 264-271.

Peter, J., M. De Chiara, A. Friedrich, J. X. Yue, D. Pflieger, A. Bergstrom, A. Sigwalt, B. Barre, K. Freel, A. Llored, C. Cruaud, K. Labadie, J. M. Aury, B. Istace, K. Lebrigand, P. Barbry, S. Engelen, A. Lemainque, P. Wincker, G. Liti and J. Schacherer (2018). "Genome evolution across 1,011 *Saccharomyces cerevisiae* isolates." Nature **556**(7701): 339-344.

Zhang, C. T. and J. Wang (2000). "Recognition of protein coding genes in the yeast genome at better than 95% accuracy based on the Z curve." Nucleic Acids Res **28**(14): 2804-2814.
